# CommDivMap: Modelling and mapping species richness at different spatial scales

**DOI:** 10.1101/2020.05.11.089029

**Authors:** Julia E Miller, Dirk Steinke

**Affiliations:** Centre for Biodiversity Genomics, University of Guelph, 50 Stone Road East, Guelph, Ontario, N1G 2W1, Canada; Department of Integrative Biology, University of Guelph, 50 Stone Road East, Guelph, Ontario, N1G 2W1, Canada

## Abstract

1. Modern ecosystem models have the potential to greatly enhance our capacity to predict community responses to change, but they demand comprehensive spatial distribution information, creating the need for new approaches to gather and synthesize biodiversity data.

2. Metabarcoding or metagenomics can generate comprehensive biodiversity data sets at species-level resolution but they are limited to point samples.

3. CommDivMap contains a number of functions that can be used to turn OTU tables resulting from metabarcoding runs of bulk samples into species richness maps. We tested the method on a series of arthropod bulk samples obtained from various experimental agricultural plots.

4. The script runs smoothly and is reasonably fast. We hope that our assemble first, predict later approach to statistical modelling of species richness will set the stage for the transition from data-rich but finite sets of point samples to spatially continuous biodiversity maps.

## Introduction

The observation and quantification of change in ecosystems are fundamental tools for assessing the response of species communities to environmental change (Bonanda et al. 2006; Moog et al. 2018). Past studies have typically monitored the response of a few indicator species through repeated surveys of sites to measure impacts on the entire community, e.g. quantified by shifts in abundance or by variation in alpha and beta diversity (Carignan & Villard 2002; Siddig et al 2016). Although such studies can deliver a basic understanding of biodiversity, they fall short of providing the observational data needed to manage and protect it at larger scales. New computationally demanding ecosystem models have the potential to greatly enhance our capacity to predict community responses to change, but they demand more comprehensive spatial distribution information, creating the need for new approaches to gather and synthesize biodiversity data (Bush et al. 2017).

High-throughput DNA sequencing methods used in metabarcoding or metagenomic studies are capable of generating the required comprehensive biodiversity data sets at species-level resolution (Beng et al 2016; Bell 2017; Braukmann et al. 2019; Yu et al. 2012), but they are still limited to point samples. In order to be able to describe or even forecast regional- to large-scale changes in patterns of biodiversity we need methods of statistical modelling that allow us to use such point-samples to build continuous community maps and to calculate metrics such as richness or dissimilarity (Bush et al 2017; Derocles et al. 2018; Ferrier & Guisan 2006).

One common type of modeling community distribution maps is to start with individual species distribution models (SDMs – Elith & Leathwick 2009) in a so-called *Predict first, assemble later approach.* Continuous community-level patterns are modeled by stacking individual SDMs (Ferrier & Guisan 2006). This strategy is based on the view that communities result from chance assembly of individual ecological responses of species (Amen et al. 2017). However, metabarcoding or metagenomic studies often yield hundreds of species per individual bulk sample which means that this approach is not scalable. These datasets could rather be used as *a priori* assemblies to predict community distribution or other cumulative community attributes. Such an *Assemble first, predict later* approach assumes that communities are combinations of a fixed set of co-occurring species. Distributions are modelled under the premise that the total species pool is stable over time and space (Ferrier & Guisan 2006).

Following the latter concept we developed a R script (R Core Team 2018), CommDivMap, which contains a set of functions that can turn OTU tables resulting from metabarcoding runs of bulk point samples into continuous species richness maps by using either linear or binomial regression models.

## The CommDivMap package

The package workflow is divided into three distinct stages that encompass data input and conversion, statistical analysis as well as mapping (Figure 1). All functions associated with those are briefly described in the balance of this section.

**Figure 1:**
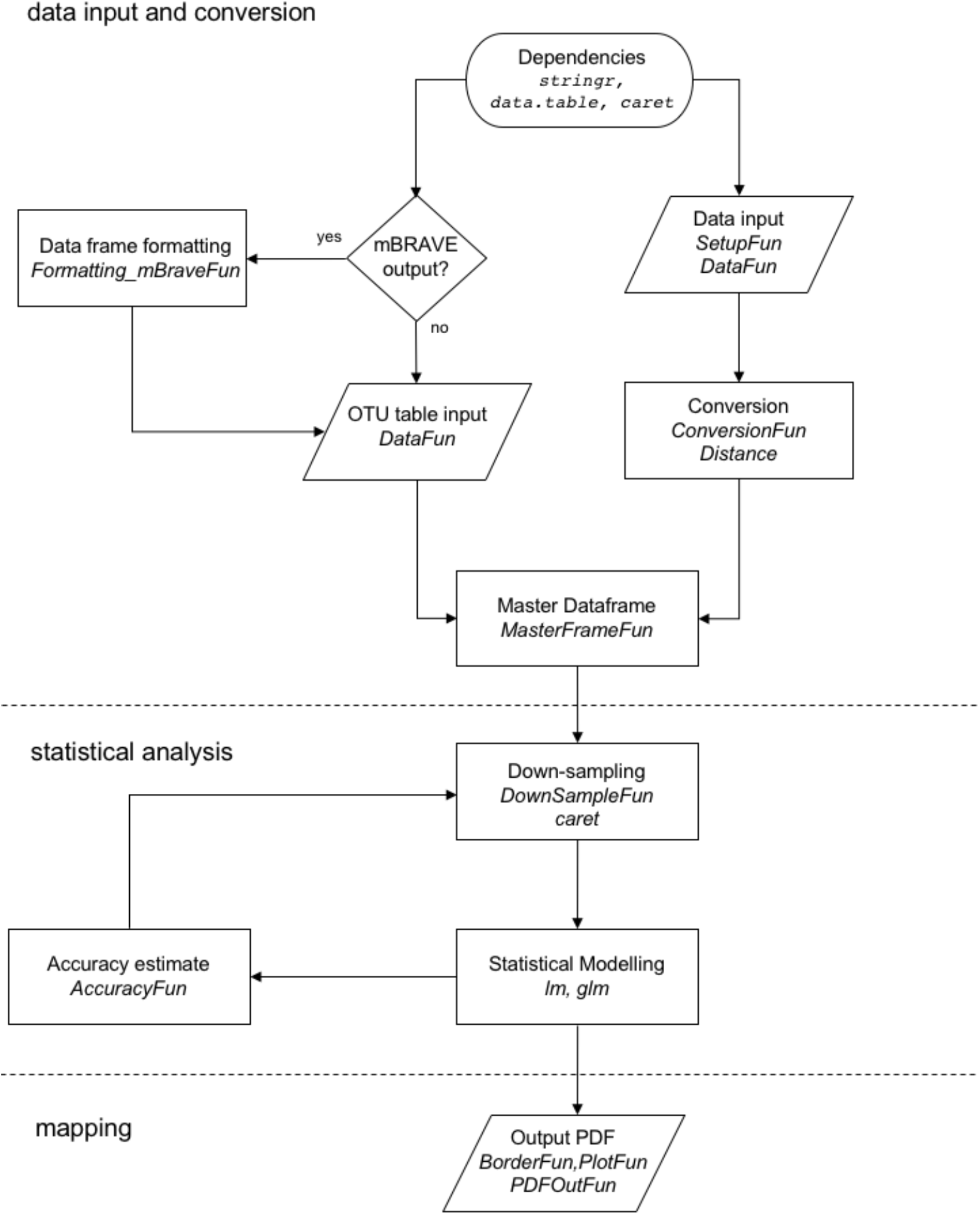
Flow diagram illustrating all functions and stages of the CommDivMap package.

### Data input and conversion

#### SetupFun

This function defines the number of temporal and spatial sampling points. It also uses the variables *Border_Coordinates* and *Trap_Coodinates* for input of either a matrix or a data frame outlining the decimal degree lat/lon coordinates of the area to be mapped and for the positions of the bulk sampling points.

#### DataFun

Function to input OTU lists (Supplementary Figure 1) for each bulk sample. Each column represents a single bulk sample. Two alternative input formats, both used as test cases for this study, are given in Figure 2.

**Figure 2:**
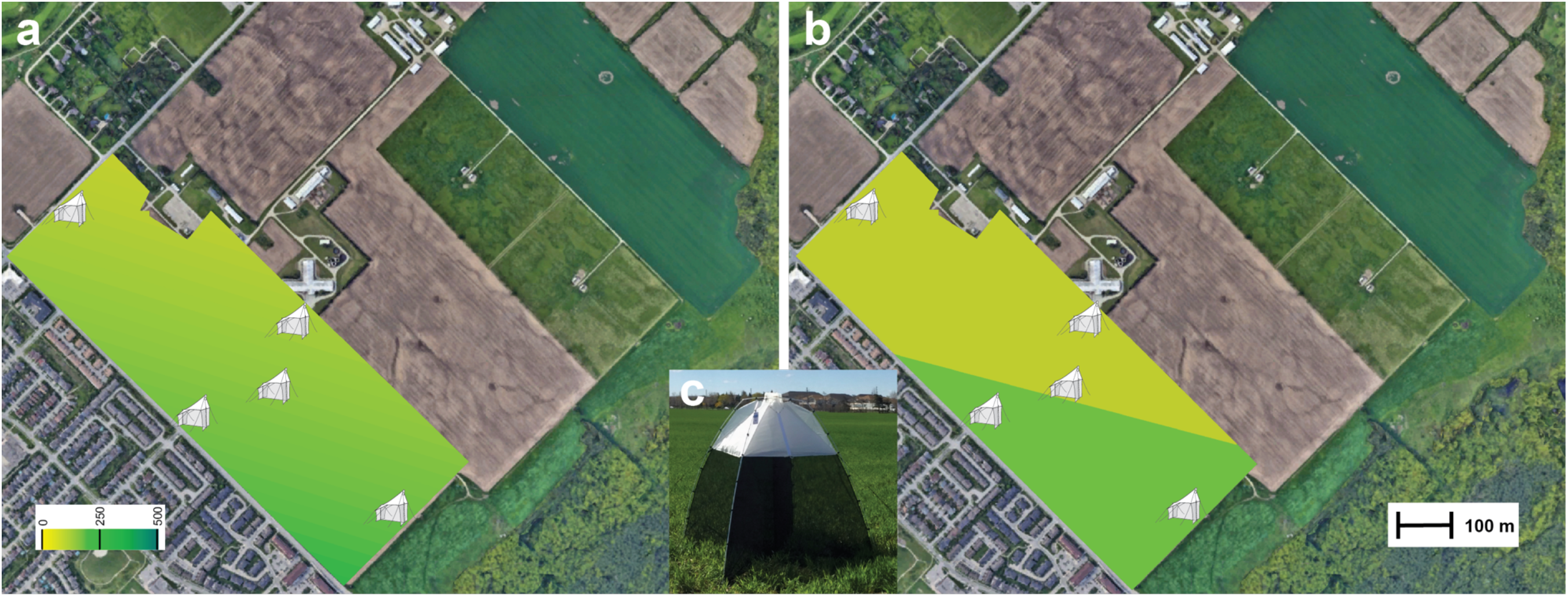
Map of the experimental wheat plot at Arkell Research Station (University of Guelph). CommDivMap heatmaps using a linear model (a) and a binomial model (b) are superimposed on a Google Earth Map (distance legend at bottom right). Locations for Malaisetraps (SLAM type as shown in c) are indicated by a trap icon. Heatmap colours represent species richness (legend at bottom left).

#### Formatting_mBraveFun

The function *Formatting_mBraveFun* can be used to convert OTU tables (tsv format) generated by the Multiplex Barcode Research And Visualization Environment (mBRAVE – Ratnasingham 2019) into the data frame required for *MasterFrameFun.*

#### ConversionFun

*ConversionFun* converts lat/lon coordinates into Cartesian coordinates. Two additional variables in *ConversionFun* are used to indicate if the scale of the map will be in kilometres (instead of default setting meters) and if the area being mapped is located in a polar region. Both variables are set to FALSE by default.

### Statistical analysis

Traditionally the first regression model chosen is the linear model which determines the equation for the line of best fit (Quinn & Keough 2002). It is considered the most appropriate choice for continuous data and has been implemented in our script. However, at the core of our modelling approach lies the prediction if a taxon is present or absent, consequently, the response variable is binary and results are assumed to be binomially distributed with n=1. It is more common to fit those as generalised linear model. Therefore, our script also contains a binomial regression approach which determines the probabilities of an OTU being present at a given range of geographic coordinates.

To find out how well a particular model performs we used a cross-validation approach by creating training and testing data through random sampling from the initial dataset without replacement (down-sampling). Different measures can be used to evaluate the quality of a prediction and those that evaluate presence-absence data are usually threshold-based (Liu et al. 2005). Predicted values above a chosen threshold indicate presence and values below indicate absence.

#### MasterFrameFun

This function links each OTU with spatio-temporal information. The resulting data frame contains OTUs, sample x-y coordinates, sampling dates, species (OTU) richness, count of samples that contained an OTU at a given sampling date, OTU presence/absence for each sample.

#### DownSampleFun

Down-sampling imbalanced data sets usually yields better generalised linear model performance. This function uses a 70/30 default ratio of training to test data. The object created by this function is a list including down-sampled training data, original training data and test data. The downsampled data can be inputted into any statistical modeling function in R. Our script uses the basic R functions *lm* (linear model) and *glm* (Generalized linear model). For the latter the argument *family* is set to binomial.

#### AccuracyFun

This function helps optimizing the model by applying it to the training data set. It returns a decimal showing the percentage of overlap between model predictions and observations.

### Mapping

#### BorderFun

*BorderFun* uses the coordinates that define the map boundaries to identify all the ordered pairs that will need to be predicted and mapped to generate a heatmap.

#### PlotFUN

This function generates a matrix of predicted species richness for each ordered pair.

#### PDFOutFun

The function converts the matrix into a heatmap that is saved as pdf file. The user can define colour, gradient and title of the heatmap.

## Application - Diversity at the farm scale

For development and test of our script we used a subset of data of an agricultural monitoring pilot project based on metabarcoding of weekly collected bulk Malaise trap samples (deWaard et al. 2017). From May to September 2017, a series of Malaise traps of the SLAM type (Figure 2c) was deployed across various agricultural plots at University of Guelph research stations in Southern Ontario to collect arthropod communities. SLAM traps contain four sampling bottles in separate compartments that were oriented along cardinal directions. Each plot hosted five traps at specific locations (Figure 2 a,b). DNA of each sample bottle (4 per week per trap) was isolated using a non-destructive membrane-based protocol (Steinke et al. in prep). Subsequently, PCR amplifications for two technical extract replicates per sample were performed using a standard DNA barcode primer set (AncientLepF3 (Prosser et al. 2016) and LepR1 (Hebert et al. 2004)). Pooled and purified amplicons were sequenced using an Ion torrent S5 with a 530 chip kit, following the manufacturer’s instructions (Thermo Fisher Scientific). The resulting high-throughput sequencing (HTS) datasets were analyzed using mBRAVE (mbrave.net – Ratnasingham 2019). All raw HTS datasets were deposited in SRA under Bioproject PRJNA631160. We used accession numbers SAMN14853989 to SAMN14853993. OTU output tables from mBRAVE are available as supplementary data (Supplementary file 1).

We used seasonal data for a wheat plot to generate species richness maps at m^2^-resolution for each set of bi-weekly samples. Multiple OTU tables representative of multiple collection events can be run simultaneously with our script. Examples for possible forms of input data are shown in Supplementary Figure 1. The runtime for the entire script was 22566.4 s on a MacBook Air 1.6 GHz Dual-Core Intel Core i5 for our application dataset (nine sampling events with four bulk samples each, OTU count varying from 80 to 350).

All resulting pdf files are available as supplementary data. One example is provided in Figure 2. Two heatmaps of local arthropod species diversity are provided – one using a linear model (Figure 2a) and one using a binomial model (Figure 2b). We superimposed heatmaps on a Google Earth layer for better visualization but outputs can also be used to generate standard shapefiles for use in GIS packages. We have started development of an R script to create shapefiles for use with QGIS (2020) and an experimental version is available at https://github.com/JuliaHarvie/Community-Diversity-Heatmap/blob/master/Rough-Code/Shapefile_Simplified.R

Figure 2 illustrates the differences between the two models used. For linear regression, the dependent variable is continuous and can have any one of an infinite number of possible values. For a binomial logistic regression, the dependent variable has only a limited number of possible values and produces a less continuous map. Nevertheless, both models show agreement in the prediction of higher species richness for the Southern tip of the Wheat plot chosen for illustration. This can be explained by pastures and woodlots at the South-East boundary of the plot which can serve as refugia for arthropod species. To further illustrate temporal variation throughout the season we used individual map plots to assemble animated GIFs (Supplementary Files 2 and 3) for each model type.

## Conclusions

We believe that the CommDivMap script can be very useful for both ecological study design and community analysis. The combination of species richness data obtained via high-throughput DNA sequencing and statistical modelling augments our capacity to create spatially continuous biodiversity maps that can aid in the prediction of community responses to change.

## Supporting information

Suppl File 1 - Input file example

Suppl File 2 - animated GIF linear model

Suppl File 3 - animated GIF binomial model

## Author’s contributions

J.E.H. and D.S. conceived the project. J.H. developed and tested the script. J.E.H. and D.S. wrote the publication.

## Acknowledgements

We like to thank Jarrett Phillips for helpful input on earlier versions of the script. This study was supported by funding through the Canada First Research Excellence Fund. The funders had no role in study design, data collection and analysis, decision to publish, or preparation of the manuscript. This work represents a contribution to the *Food From Thought* research programme

## Data accessibility

The CommDivMap package and documentation are hosted at https://github.com/JuliaHarvie/Community-Diversity-Heatmap

---

**Supplementary Figure 1:**
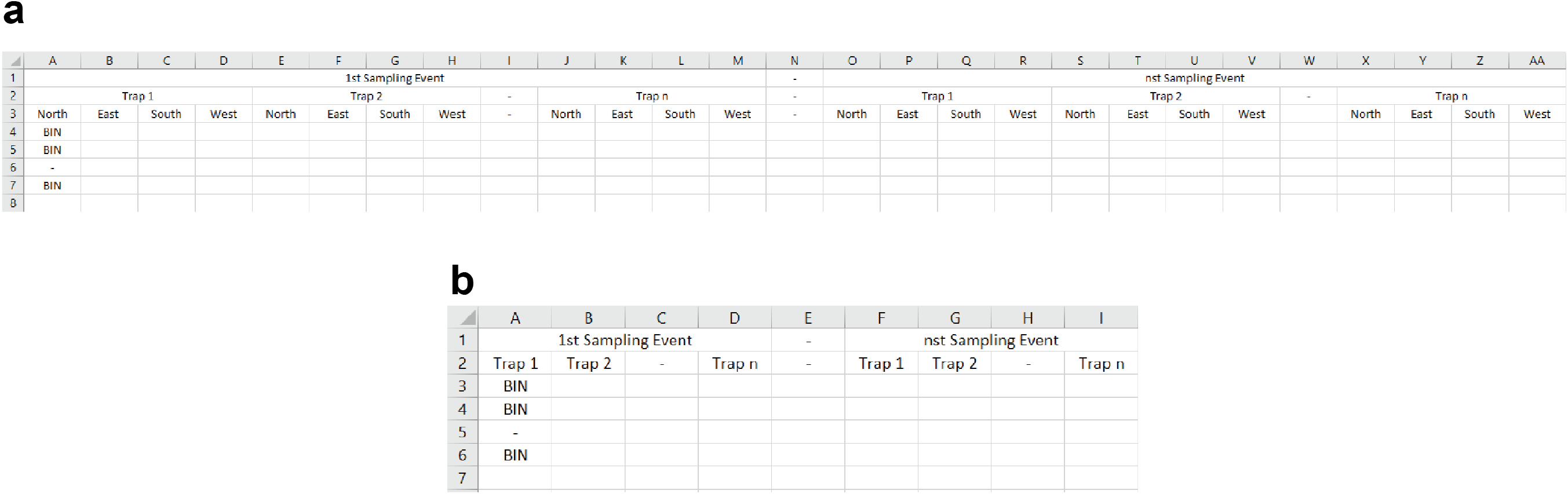
Alternative Input formats accepted by CommDivMap. (a) used for application presented in this study, designed for SLAM trap with directional subsamples in cardinal direction. Can be modified for other strategies involving subsamples; (b) more general input format for bulk sample results. Inputs can be BINs, OUT#s, species names etc.

## Notes

https://github.com/JuliaHarvie/Community-Diversity-Heatmap

### Competing Interest Statement

The authors have declared no competing interest.

