## Supplementary material for "CommDivMap: Modelling and mapping species richness at different spatial scales": Suppl File 1 - Input file example

"Site","Event","Library","BIN.URI","Phylum","Class","Order","Family","Subfamily","Genus","Species","Sequences","Dereplicates","PreClusters","Abundance.Rank","Dereplication.Reduction....","Preclustering.Reduction....","Mean.Similarity....","Mean.Overlap","Mean.ID.Match.Gaps","Length","GC.Composition","MID"
"AGAW1","02E","BAC","TAX:175768","Proteobacteria","Alphaproteobacteria","Rickettsiales","Rickettsiaceae",NA,"Wolbachia","Wolbachia pipientis","16","12","12","1","25","0","98.73","448.5","0","449.25","39.55","0"
"AGAW1","02E","BAC","TAX:376688","Proteobacteria","Alphaproteobacteria","Rickettsiales","Rickettsiaceae",NA,"Wolbachia","Wolbachia sp. elachBioLep01 BioLep373","10","7","7","2","30","0","97.49","449.9","0","450","41.11","0"
"AGAW1","02E","BAC","TAX:375763","Proteobacteria","Alphaproteobacteria","Rickettsiales","Rickettsiaceae",NA,"Wolbachia","Wolbachia sp. Pheidole nebulosa_ASLAM107-08","5","5","5","3","0","0","98.95","417.2","0","449.2","41.23","0"
"AGAW1","02E","BAC","TAX:91393","Proteobacteria","Alphaproteobacteria","Rickettsiales","Rickettsiaceae",NA,"Wolbachia","","3","3","3","4","0","0","99.15","391.67","0","439.33","39.53","0"
"AGAW1","02E","BAC","TAX:375762","Proteobacteria","Alphaproteobacteria","Rickettsiales","Rickettsiaceae",NA,"Wolbachia","Wolbachia sp. Myrmicinae_ACGAN556-09","2","2","2","5","0","0","97.33","449","0","450","41.11","0"
"AGAW1","02E","BAC","TAX:453459","Proteobacteria","Alphaproteobacteria","Rickettsiales","Rickettsiaceae",NA,"Wolbachia","Wolbachia endosymbiont of Ichneumonidae","1","1","1","6","0","0","97.56","450","0","450","41.11","0"
"AGAW1","02E","CHO","no match","no match","no match","no match","no match","no match","no match","no match","no match","no match","no match","no match","no match","no match","no match","no match","no match","no match","no match","no match"
"AGAW1","02E","INS","BOLD:AAG2511","Arthropoda","Insecta","Diptera","Anthomyiidae","Anthomyiinae","Delia","Delia platura","73246","47798","47798","1","34.74","0","99.46","443.38","0","445.56","33.18","0"
"AGAW1","02E","INS","BOLD:AAG1742","Arthropoda","Insecta","Diptera","Muscidae","Phaoniinae","Helina","Helina rufitibia","8886","5895","5895","2","33.66","0","99.22","445.15","0","446.86","30.32","0"
"AGAW1","02E","INS","BOLD:AAQ0582","Arthropoda","Insecta","Diptera","Anthomyiidae","Anthomyiinae","Lasiomma","","6924","4815","4815","3","30.46","0","99.37","445.18","0","446.25","31.72","0"
"AGAW1","02E","INS","BOLD:AAA3453","Arthropoda","Insecta","Diptera","Anthomyiidae","Anthomyiinae","Delia","","4402","3382","3382","4","23.17","0","99.27","444.1","0","445.86","34.7","0"
"AGAW1","02E","INS","BOLD:AAB2384","Arthropoda","Insecta","Diptera","Syrphidae","Syrphinae","Eupeodes","Eupeodes duseki, Eupeodes volucris, Eupeodes nielseni, Eupeodes americanus, Eupeodes flukei, Eupeodes vockerothi, Eupeodes lucasi, Eupeodes montivagus, Eupeodes rufipunctatus, Eupeodes rojasi, Eupeodes goeldlini, Eupeodes luniger, Eupeodes punctifer, Eupeodes bucculatus, Eupeodes perplexus, Eupeodes abiskoensis, Eupeodes corollae, Eupeodes pomus, Eupeodes lundbecki, Eupeodes nitens, Eupeodes sp. BOLDAAB2384, Eupeodes curtus, Eupeodes latifasciatus","3407","2924","2924","5","14.18","0","98.87","426.62","0","430.64","31.01","0"
"AGAW1","02E","INS","BOLD:AAD0853","Arthropoda","Insecta","Diptera","Scathophagidae","Scathophaginae","Scathophaga","Scathophaga lutaria, Scathophaga hiemalis, Scathophaga furcata, Scathophaga stercoraria, Scathophaga inquinata, Scathophaga suilla","2743","2267","2267","6","17.35","0","99.24","437.6","0","439.17","31.59","0"
"AGAW1","02E","INS","BOLD:AAN5355","Arthropoda","Insecta","Diptera","Chironomidae","Orthocladiinae","Smittia","Smittia cf. stercoraria, Smittia stercoraria","1803","1518","1518","7","15.81","0","99.14","415.67","0","444.81","36.31","0"
"AGAW1","02E","INS","BOLD:AAB1098","Arthropoda","Insecta","Diptera","Culicidae","Culicinae","Ochlerotatus, Aedes","Aedes excrucians, Aedes cantans, Aedes stimulans, Aedes euedes, Aedes riparius, Aedes annulipes, Ochlerotatus cantans, Aedes fitchii, Aedes sp., Ochlerotatus annulipes, Ochlerotatus excrucians","1663","1512","1512","8","9.08","0","98.6","431.34","0","432.65","35.2","0"
"AGAW1","02E","INS","BOLD:AAB2644","Arthropoda","Insecta","Diptera","Chironomidae","Orthocladiinae","Orthocladius","Orthocladius lapponicus, Orthocladius dorenus, Orthocladius mallochi","1454","1095","1095","9","24.69","0","99.45","446.42","0","448.07","32.3","0"
"AGAW1","02E","INS","BOLD:AAA9251","Arthropoda","Insecta","Lepidoptera","Gelechiidae","Gelechiinae","Scrobipalpa","Scrobipalpa atriplicella, Scrobipalpa nr. atriplicella sp. 1","407","347","347","10","14.74","0","99.43","442.48","0","444.24","34.48","0"
"AGAW1","02E","INS","BOLD:AAC4197","Arthropoda","Insecta","Diptera","Chironomidae","Orthocladiinae","Paraphaenocladius","Paraphaenocladius impensus","351","322","322","11","8.26","0","99.2","437.7","0","440.1","35.12","0"
"AGAW1","02E","INS","BOLD:AAN5358","Arthropoda","Insecta","Diptera","Chironomidae","Orthocladiinae","Smittia","Smittia stercoraria, Smittia sp. 22ES","274","241","241","12","12.04","0","99.27","442.21","0","443.97","37.32","0"
"AGAW1","02E","INS","BOLD:ABU5525","Arthropoda","Insecta","Diptera","Chironomidae","Orthocladiinae","Limnophyes","Limnophyes sp. 14ES","191","172","172","13","9.95","0","99.13","429.21","0","430.86","37.28","0"
"AGAW1","02E","INS","BOLD:AAM7726","Arthropoda","Insecta","Coleoptera","Curculionidae","Curculioninae","Orchestes","Orchestes steppensis, Orchestes alni","156","132","132","14","15.38","0","99.1","437.85","0","440.45","33.31","0"
"AGAW1","02E","INS","BOLD:ACV5349","Arthropoda","Insecta","Diptera","Chironomidae","Tanypodinae","","","154","120","120","15","22.08","0","98.99","388.71","0","443.6","36.49","0"
"AGAW1","02E","INS","BOLD:ACJ0208","Arthropoda","Insecta","Diptera","Chironomidae","Orthocladiinae","","","87","78","78","16","10.34","0","99.03","436.2","0","449.68","34.93","0"
"AGAW1","02E","INS","BOLD:ACN3917","Arthropoda","Insecta","Diptera","Anthomyiidae","Anthomyiinae","Strobilomyia","Strobilomyia anthracina","65","63","63","17","3.08","0","97.78","426.14","0","434.66","31.36","0"
"AGAW1","02E","INS","BOLD:AAN5267","Arthropoda","Insecta","Diptera","Cecidomyiidae","","","","64","53","53","18","17.19","0","99.36","448.72","0","449.47","30.65","0"
"AGAW1","02E","INS","BOLD:AAA5803","Arthropoda","Insecta","Hemiptera","Miridae","Mirinae","Lygus","Lygus lineolaris","61","49","49","19","19.67","0","99.45","445.97","0","447.26","34.13","0"
"AGAW1","02E","INS","BOLD:AAM9252","Arthropoda","Insecta","Diptera","Sciaridae","","Bradysia, Leptosciarella","Bradysia vagans, Leptosciarella zanti","45","41","41","20","8.89","0","99.6","448.56","0","449.58","39.22","0"
"AGAW1","02E","INS","BOLD:AAB9140","Arthropoda","Insecta","Diptera","Calliphoridae","Chrysomyinae","Phormia","Phormia regina","44","44","44","21","0","0","99.1","442.89","0","444.16","33","0"
"AGAW1","02E","INS","BOLD:AAG1000","Arthropoda","Insecta","Diptera","Chironomidae","Orthocladiinae","Orthocladius","Orthocladius carlatus","32","31","31","22","3.13","0","98.02","426.25","0","428.28","33.92","0"
"AGAW1","02E","INS","BOLD:ACX4619","Arthropoda","Insecta","Coleoptera","Scirtidae","Scirtinae","Cyphon","Cyphon laevipennis, Cyphon variabilis","32","30","30","22","6.25","0","99.21","425.84","0","427.56","39.68","0"
"AGAW1","02E","INS","BOLD:AAF9014","Arthropoda","Insecta","Diptera","Limoniidae","Chioneinae","Symplecta","Symplecta cana, Symplecta sp. BOLDAAF9014","31","31","31","24","0","0","98.29","414.39","0","418.19","33.89","0"
"AGAW1","02E","INS","BOLD:ADH4889","Arthropoda","Insecta","Diptera","","","","","28","26","26","25","7.14","0","98.83","440.36","0","441.93","35.36","0"
"AGAW1","02E","INS","BOLD:AAY6412","Arthropoda","Insecta","Diptera","Sciaridae","","Bradysia","Bradysia urticae","21","21","21","26","0","0","99.26","438.43","0","440.52","32.09","0"
"AGAW1","02E","INS","BOLD:AAY3269","Arthropoda","Insecta","Diptera","Chironomidae","Diamesinae","Potthastia","Potthastia gaedii","21","21","21","26","0","0","98.3","433.24","0","435.19","34.2","0"
"AGAW1","02E","INS","BOLD:AAP5343","Arthropoda","Insecta","Diptera","Cecidomyiidae","","","","19","17","17","28","10.53","0","99.33","448.37","0","449.37","29.46","0"
"AGAW1","02E","INS","BOLD:AAV1295","Arthropoda","Insecta","Diptera","Sciaridae","","Bradysia","Bradysia difformis, Bradysia impatiens","15","14","14","29","6.67","0","99.51","415.73","0","437.27","31.55","0"
"AGAW1","02E","INS","BOLD:AAB0868","Arthropoda","Insecta","Diptera","Calliphoridae","Calliphorinae","Cynomya","Cynomya cadaverina, Cynomya mortuorum","15","15","15","29","0","0","99","438","0","438.73","34.92","0"
"AGAW1","02E","INS","BOLD:ACF8552","Arthropoda","Insecta","Coleoptera","Anthicidae","Anthicinae","Malporus","Malporus formicarius","13","10","10","31","23.08","0","99.66","364.85","0","450","34.53","0"
"AGAW1","02E","INS","BOLD:ADH9867","Arthropoda","Insecta","Diptera","","","","","12","12","12","32","0","0","98.6","386.33","0","390.83","37.02","0"
"AGAW1","02E","INS","BOLD:ABU5520","Arthropoda","Insecta","Diptera","Sciaridae","","Corynoptera","","11","11","11","33","0","0","99.03","401.18","0","403.55","34.93","0"
"AGAW1","02E","INS","BOLD:ACL3317","Arthropoda","Insecta","Diptera","Anthomyiidae","","","","10","6","6","34","40","0","97.37","448.7","0","450","31.49","0"
"AGAW1","02E","INS","BOLD:ACR4394","Arthropoda","Insecta","Diptera","Anthomyiidae","Anthomyiinae","Delia","Delia florilega","8","8","8","35","0","0","98.87","433.13","0","435.75","32.91","0"
"AGAW1","02E","INS","BOLD:ACB1079","Arthropoda","Insecta","Diptera","Cecidomyiidae","Micromyinae","Campylomyza","Campylomyza dilatata, CecidInt44 sp. BOLD:ACB1079","6","6","6","36","0","0","98.14","439.17","0","440","28.06","0"
"AGAW1","02E","INS","BOLD:AAZ5599","Arthropoda","Insecta","Diptera","Chironomidae","Orthocladiinae","","","5","5","5","37","0","0","99.58","432.2","0","433.8","36.72","0"
"AGAW1","02E","INS","BOLD:AAD4355","Arthropoda","Insecta","Diptera","Culicidae","Culicinae","Aedes","Aedes provocans","4","4","4","38","0","0","98.42","433.75","0","436.25","32.03","0"
"AGAW1","02E","INS","BOLD:AAN5356","Arthropoda","Insecta","Diptera","Chironomidae","Orthocladiinae","Smittia","","4","4","4","38","0","0","99.5","448.5","0","450","35.11","0"
"AGAW1","02E","INS","BOLD:ACN5521","Arthropoda","Insecta","Diptera","Anthomyiidae","Anthomyiinae","Strobilomyia","Strobilomyia carbonaria","4","4","4","38","0","0","97.83","420.25","0","429.75","31.39","0"
"AGAW1","02E","INS","BOLD:ACL4768","Arthropoda","Insecta","Coleoptera","Chrysomelidae","Galerucinae","Chaetocnema","Chaetocnema confinis","3","3","3","41","0","0","100","348.33","0","450","33.48","0"
"AGAW1","02E","INS","BOLD:ACF6272","Arthropoda","Insecta","Diptera","Chironomidae","Orthocladiinae","","","3","3","3","41","0","0","97.35","377","0","450","36.96","0"
"AGAW1","02E","INS","BOLD:ACN6379","Arthropoda","Insecta","Diptera","Anthomyiidae","Anthomyiinae","Lasiomma","","3","3","3","41","0","0","98.29","448.67","0","450","31.78","0"
"AGAW1","02E","INS","BOLD:AAG6743","Arthropoda","Insecta","Diptera","Sarcophagidae","Sarcophaginae","Sarcophaga","Sarcophaga sp. 1 MG, Sarcophaga subvicina","3","3","3","41","0","0","98.61","435.67","0","437","34.3","0"
"AGAW1","02E","INS","BOLD:AAG6753","Arthropoda","Insecta","Diptera","Lauxaniidae","Lauxaniinae","Camptoprosopella","","3","3","3","41","0","0","97.82","424","0","450","36.37","0"
"AGAW1","02E","INS","BOLD:AAA4222","Arthropoda","Insecta","Lepidoptera","Erebidae","Hypeninae","Hypena","Hypena scabra","3","3","3","41","0","0","99.76","426.67","0","428","32.62","0"
"AGAW1","02E","INS","BOLD:AAK8831","Arthropoda","Insecta","Diptera","Limoniidae","Limoniinae","Metalimnobia, Limonia","","2","2","2","47","0","0","99.55","449","0","450","33.33","0"
"AGAW1","02E","INS","BOLD:AAL7534","Arthropoda","Insecta","Diptera","Anisopodidae","Anisopodinae","Sylvicola","Sylvicola stackelbergi, Sylvicola stackelbergii 1, Sylvicola punctatus","2","2","2","47","0","0","97.89","449.5","0","450","33.89","0"
"AGAW1","02E","INS","BOLD:ABU9891","Arthropoda","Insecta","Diptera","Muscidae","Phaoniinae","Phaonia","","2","2","2","47","0","0","98.99","445","0","450","32.22","0"
"AGAW1","02E","INS","BOLD:AAB5640","Arthropoda","Insecta","Coleoptera","Coccinellidae","Coccinellinae","Harmonia","Harmonia axyridis","2","2","2","47","0","0","99.63","417.5","0","418.5","32.1","0"
"AGAW1","02E","INS","BOLD:AAG1421","Arthropoda","Insecta","Hymenoptera","Braconidae","Aphidiinae","Diaeretiella","Diaeretiella rapae","2","2","2","47","0","0","99.72","358.5","0","442","27.71","0"
"AGAW1","02E","INS","BOLD:AAP9022","Arthropoda","Insecta","Diptera","Cecidomyiidae","","","","2","2","2","47","0","0","100","448","0","450","27.56","0"
"AGAW1","02E","INS","BOLD:ACC7611","Arthropoda","Insecta","Diptera","Cecidomyiidae","","","","1","1","1","53","0","0","97.76","446","0","450","28.22","0"
"AGAW1","02E","INS","BOLD:AAZ5973","Arthropoda","Insecta","Diptera","Sepsidae","Sepsinae","Themira","Themira annulipes","1","1","1","53","0","0","98.73","394","0","395","35.7","0"
"AGAW1","02E","INS","BOLD:AAG9055","Arthropoda","Insecta","Hymenoptera","Vespidae","Vespinae","Vespula","Vespula germanica","1","1","1","53","0","0","98.79","414","0","416","29.81","0"
"AGAW1","02E","INS","BOLD:AAL5267","Arthropoda","Insecta","Coleoptera","Chrysomelidae","Galerucinae","Phyllotreta","Phyllotreta striolata","1","1","1","53","0","0","98.67","450","0","450","39.11","0"
"AGAW1","02E","INS","BOLD:ACF6270","Arthropoda","Insecta","Diptera","Chironomidae","Orthocladiinae","Orthocladius","","1","1","1","53","0","0","99.3","427","0","430","30.7","0"
"AGAW1","02E","INS","BOLD:ADA3072","Arthropoda","Insecta","Diptera","Phoridae","Metopininae","Megaselia","Megaselia arcticae","1","1","1","53","0","0","97.11","450","0","450","32.22","0"
"AGAW1","02E","INS","BOLD:AAL2750","Arthropoda","Insecta","Diptera","Muscidae","Phaoniinae","Phaonia","Phaonia tuguriorum","1","1","1","53","0","0","98.44","450","0","450","32.67","0"
"AGAW1","02E","INS","BOLD:AAN1738","Arthropoda","Insecta","Coleoptera","Staphylinidae","Oxytelinae","Anotylus","Anotylus tetracarinatus","1","1","1","53","0","0","99.3","431","0","437","34.32","0"
"AGAW1","02E","INS","BOLD:AAA2144","Arthropoda","Insecta","Lepidoptera","Noctuidae","Noctuinae","Xestia","Xestia kollari, Xestia c-nigrum, Xestia dolosa","1","1","1","53","0","0","98.87","441","0","442","31.67","0"
"AGAW1","02E","INS","BOLD:AAJ1224","Arthropoda","Insecta","Hymenoptera","Braconidae","Aphidiinae","Diaeretiella","","1","1","1","53","0","0","97.33","450","0","450","26.89","0"
"AGAW1","02E","INS","BOLD:AAP2970","Arthropoda","Insecta","Diptera","Anthomyiidae","Anthomyiinae","Anthomyia","Anthomyia procellaris, Anthomyia pluvialis","1","1","1","53","0","0","97.32","448","0","450","33.33","0"
"AGAW1","02E","INS","BOLD:AAB4237","Arthropoda","Insecta","Lepidoptera","Gelechiidae","Gelechiinae","Telphusa","Telphusa longifasciella","1","1","1","53","0","0","98.65","444","0","445","33.03","0"
"AGAW1","02E","INS","BOLD:AAN5341","Arthropoda","Insecta","Diptera","Chironomidae","Orthocladiinae","Camptocladius","Camptocladius stercorarius","1","1","1","53","0","0","99.24","395","0","398","35.43","0"
"AGAW1","02E","INS","BOLD:ACX5521","Arthropoda","Insecta","Diptera","Chironomidae","Orthocladiinae","","","1","1","1","53","0","0","99.55","446","0","450","34.89","0"
"AGAW1","02E","INS","BOLD:AAB4093","Arthropoda","Insecta","Lepidoptera","Noctuidae","Noctuinae","Elaphria","Elaphria grata","1","1","1","53","0","0","97.99","447","0","450","30.22","0"
"AGAW1","02E","INS","BOLD:AAV6112","Arthropoda","Insecta","Diptera","Chloropidae","Oscinellinae","Oscinella","Oscinella sp. 2 AGT, Oscinella sp. TAW6","1","1","1","53","0","0","99.09","438","0","440","33.41","0"
"AGAW1","02E","INS","BOLD:AAA4121","Arthropoda","Insecta","Diptera","Simuliidae","Simuliinae","Simulium","Simulium vittatum complex, Simulium vittatum, Simulium sp. BOLDAAA4121, Simulium argus, Simulium vittatum s.str.","1","1","1","53","0","0","99.18","365","0","366","33.06","0"
"AGAW1","02E","INS","BOLD:AAA2955","Arthropoda","Insecta","Lepidoptera","Tortricidae","Tortricinae","Argyrotaenia","Argyrotaenia occultana, Argyrotaenia ljungiana, Argyrotaenia pinatubana, Argyrotaenia floridana, Argyrotaenia velutinana, Argyrotaenia tabulana, Argyrotaenia gogana","1","1","1","53","0","0","99.74","384","0","388","31.7","0"
"AGAW1","02E","INS","BOLD:AAA2482","Arthropoda","Insecta","Lepidoptera","Noctuidae","Noctuinae","Mythimna","Mythimna unipuncta","1","1","1","53","0","0","98.88","448","0","450","30.89","0"
"AGAW1","02E","INS","BOLD:AAG6745","Arthropoda","Insecta","Diptera","Calliphoridae","Calliphorinae","Pollenia","Pollenia pediculata","1","1","1","53","0","0","98.89","449","0","450","33.78","0"
"AGAW1","02E","INS","BOLD:ABA5313","Arthropoda","Insecta","Coleoptera","Staphylinidae","Aleocharinae","Amischa","Amischa analis, Amischa bifoveolata, Amischa nigrofusca","1","1","1","53","0","0","97.85","326","0","450","33.11","0"
"AGAW1","02E","NAR","no match","no match","no match","no match","no match","no match","no match","no match","no match","no match","no match","no match","no match","no match","no match","no match","no match","no match","no match","no match"
"AGAW1","02E","NIN","BOLD:AAB7668","Arthropoda","Arachnida","Araneae","Lycosidae",NA,"Pardosa","Pardosa milvina","205","186","186","1","9.27","0","99.24","433.2","0","435.19","30.99","0"
"AGAW1","02E","NIN","BOLD:AAN6707","Arthropoda","Arachnida","Mesostigmata","Ascidae",NA,"Arctoseius","","4","4","4","2","0","0","99.04","445.25","0","450","42.17","0"
"AGAW1","02N","BAC","TAX:91391","Proteobacteria","Alphaproteobacteria","Rickettsiales",NA,NA,NA,NA,"4","3","3","1","25","0","99.42","386","0","450","39.89","0"
"AGAW1","02N","CHO","no match","no match","no match","no match","no match","no match","no match","no match","no match","no match","no match","no match","no match","no match","no match","no match","no match","no match","no match","no match"
"AGAW1","02N","INS","BOLD:AAG2511","Arthropoda","Insecta","Diptera","Anthomyiidae","Anthomyiinae","Delia","Delia platura","48867","32492","32492","1","33.51","0","99.47","443.22","0","445.54","33.15","0"
"AGAW1","02N","INS","BOLD:AAA2482","Arthropoda","Insecta","Lepidoptera","Noctuidae","Noctuinae","Mythimna","Mythimna unipuncta","18887","14431","14431","2","23.59","0","99.47","430.46","0","432.93","30.42","0"
"AGAW1","02N","INS","BOLD:AAN5358","Arthropoda","Insecta","Diptera","Chironomidae","Orthocladiinae","Smittia","Smittia stercoraria, Smittia sp. 22ES","7181","5552","5552","3","22.68","0","99.27","443.36","0","445.33","37.28","0"
"AGAW1","02N","INS","BOLD:AAP9022","Arthropoda","Insecta","Diptera","Cecidomyiidae","","","","6397","4144","4144","4","35.22","0","99.69","446.85","0","447.91","27.66","0"
"AGAW1","02N","INS","BOLD:AAG1742","Arthropoda","Insecta","Diptera","Muscidae","Phaoniinae","Helina","Helina rufitibia","6251","4360","4360","5","30.25","0","99.09","443.97","0","445.88","30.13","0"
"AGAW1","02N","INS","BOLD:ACM5604","Arthropoda","Insecta","Diptera","Chironomidae","Orthocladiinae","Orthocladius","","2488","1954","1954","6","21.46","0","98.82","441.39","0","443.15","34.65","0"
"AGAW1","02N","INS","BOLD:AAN5355","Arthropoda","Insecta","Diptera","Chironomidae","Orthocladiinae","Smittia","Smittia cf. stercoraria, Smittia stercoraria","2450","2132","2132","7","12.98","0","98.72","442.15","0","445.12","36.18","0"
"AGAW1","02N","INS","BOLD:AAB2644","Arthropoda","Insecta","Diptera","Chironomidae","Orthocladiinae","Orthocladius","Orthocladius lapponicus, Orthocladius dorenus, Orthocladius mallochi","2044","1808","1808","8","11.55","0","98.84","438.77","0","440.93","32.76","0"
"AGAW1","02N","INS","BOLD:AAB1098","Arthropoda","Insecta","Diptera","Culicidae","Culicinae","Ochlerotatus, Aedes","Aedes excrucians, Aedes cantans, Aedes stimulans, Aedes euedes, Aedes riparius, Aedes annulipes, Ochlerotatus cantans, Aedes fitchii, Aedes sp., Ochlerotatus annulipes, Ochlerotatus excrucians","1191","1104","1104","9","7.3","0","98.67","429.58","0","430.98","35.26","0"
"AGAW1","02N","INS","BOLD:ACI6860","Arthropoda","Insecta","Diptera","Chironomidae","Orthocladiinae","","","1136","1071","1071","10","5.72","0","98.55","426.33","0","428.49","35.8","0"
"AGAW1","02N","INS","BOLD:AAC4197","Arthropoda","Insecta","Diptera","Chironomidae","Orthocladiinae","Paraphaenocladius","Paraphaenocladius impensus","555","494","494","11","10.99","0","98.9","438.18","0","440.59","34.65","0"
"AGAW1","02N","INS","BOLD:AAA3453","Arthropoda","Insecta","Diptera","Anthomyiidae","Anthomyiinae","Delia","","516","457","457","12","11.43","0","99.25","443.17","0","445.27","34.66","0"
"AGAW1","02N","INS","BOLD:ACJ0208","Arthropoda","Insecta","Diptera","Chironomidae","Orthocladiinae","","","440","335","335","13","23.86","0","99.1","436.04","0","449.51","34.92","0"
"AGAW1","02N","INS","BOLD:ACJ5126","Arthropoda","Insecta","Diptera","Chironomidae","Orthocladiinae","Metriocnemus","","162","158","158","14","2.47","0","99.12","439.04","0","441.01","34.85","0"
"AGAW1","02N","INS","BOLD:AAG2740","Arthropoda","Insecta","Diptera","Ephydridae","Parydrinae","Philygria","Philygria debilis","155","150","150","15","3.23","0","98.2","432.75","0","436.12","33.89","0"
"AGAW1","02N","INS","BOLD:ABW5541","Arthropoda","Insecta","Diptera","Sciaridae","","Lycoriella","Lycoriella agraria","102","74","74","16","27.45","0","99.67","447.92","0","448.73","36.93","0"
"AGAW1","02N","INS","BOLD:AAN6447","Arthropoda","Insecta","Diptera","Sciaridae","","Corynoptera","Corynoptera perpusilla","97","81","81","17","16.49","0","99.16","447.3","0","448.54","36.11","0"
"AGAW1","02N","INS","BOLD:ABA1215","Arthropoda","Insecta","Diptera","Sciaridae","","Lycoriella","Lycoriella sativae","95","73","73","18","23.16","0","99.57","442.86","0","443.99","35.76","0"
"AGAW1","02N","INS","BOLD:AAP6465","Arthropoda","Insecta","Diptera","Sciaridae","","Corynoptera","Corynoptera cursor","81","66","66","19","18.52","0","98.49","447.27","0","448.59","32.32","0"
"AGAW1","02N","INS","BOLD:AAM8996","Arthropoda","Insecta","Diptera","Mycetophilidae","Mycetophilinae","Exechia","","79","72","72","20","8.86","0","99.54","445.75","0","447.04","36.79","0"
"AGAW1","02N","INS","BOLD:AAL5267","Arthropoda","Insecta","Coleoptera","Chrysomelidae","Galerucinae","Phyllotreta","Phyllotreta striolata","67","63","63","21","5.97","0","99.37","437.48","0","441.34","39.23","0"
"AGAW1","02N","INS","BOLD:AAM9252","Arthropoda","Insecta","Diptera","Sciaridae","","Bradysia, Leptosciarella","Bradysia vagans, Leptosciarella zanti","67","61","61","21","8.96","0","99.47","448.4","0","449.19","39.08","0"
"AGAW1","02N","INS","BOLD:ACB1079","Arthropoda","Insecta","Diptera","Cecidomyiidae","Micromyinae","Campylomyza","Campylomyza dilatata, CecidInt44 sp. BOLD:ACB1079","63","51","51","23","19.05","0","98.34","441.98","0","446.79","28.33","0"
"AGAW1","02N","INS","BOLD:AAN8770","Arthropoda","Insecta","Diptera","Psychodidae","Psychodinae","Psychoda","Psychoda trinodulosa","55","55","55","24","0","0","99.19","439.87","0","440.93","33.55","0"
"AGAW1","02N","INS","BOLD:ABU5525","Arthropoda","Insecta","Diptera","Chironomidae","Orthocladiinae","Limnophyes","Limnophyes sp. 14ES","49","48","48","25","2.04","0","99.14","422.02","0","425.63","37.45","0"
"AGAW1","02N","INS","BOLD:AAN6433","Arthropoda","Insecta","Diptera","Sciaridae","","Corynoptera","Corynoptera deserta","45","43","43","26","4.44","0","98.89","442.16","0","443.93","30.62","0"
"AGAW1","02N","INS","BOLD:AAY6412","Arthropoda","Insecta","Diptera","Sciaridae","","Bradysia","Bradysia urticae","44","41","41","27","6.82","0","99.34","443.23","0","445.02","32.22","0"
"AGAW1","02N","INS","BOLD:AAN5267","Arthropoda","Insecta","Diptera","Cecidomyiidae","","","","42","36","36","28","14.29","0","99.5","447.07","0","450","30.66","0"
"AGAW1","02N","INS","BOLD:AAB9140","Arthropoda","Insecta","Diptera","Calliphoridae","Chrysomyinae","Phormia","Phormia regina","32","30","30","29","6.25","0","98.95","440.97","0","442.25","33.2","0"
"AGAW1","02N","INS","BOLD:AAG4944","Arthropoda","Insecta","Diptera","Mycetophilidae","Sciophilinae","Sciophila","Sciophila sp. CJB1","25","25","25","30","0","0","98.08","431.36","0","433.48","33.97","0"
"AGAW1","02N","INS","BOLD:ACO1622","Arthropoda","Insecta","Diptera","Cecidomyiidae","","","","24","24","24","31","0","0","98.96","434.13","0","438.67","31.55","0"
"AGAW1","02N","INS","BOLD:AAF9014","Arthropoda","Insecta","Diptera","Limoniidae","Chioneinae","Symplecta","Symplecta cana, Symplecta sp. BOLDAAF9014","22","20","20","32","9.09","0","98.69","436.09","0","443.27","34","0"
"AGAW1","02N","INS","BOLD:ABW5634","Arthropoda","Insecta","Thysanoptera","Thripidae","Thripinae","Thrips","","16","16","16","33","0","0","99.3","441.06","0","444.19","36.2","0"
"AGAW1","02N","INS","BOLD:ACF8552","Arthropoda","Insecta","Coleoptera","Anthicidae","Anthicinae","Malporus","Malporus formicarius","12","11","11","34","8.33","0","99.7","364.67","0","450","34.24","0"
"AGAW1","02N","INS","BOLD:AAB0868","Arthropoda","Insecta","Diptera","Calliphoridae","Calliphorinae","Cynomya","Cynomya cadaverina, Cynomya mortuorum","11","11","11","35","0","0","99.37","432.27","0","435.55","34.74","0"
"AGAW1","02N","INS","BOLD:AAD0853","Arthropoda","Insecta","Diptera","Scathophagidae","Scathophaginae","Scathophaga","Scathophaga lutaria, Scathophaga hiemalis, Scathophaga furcata, Scathophaga stercoraria, Scathophaga inquinata, Scathophaga suilla","10","10","10","36","0","0","98.85","437.7","0","440","31.76","0"
"AGAW1","02N","INS","BOLD:ADH9867","Arthropoda","Insecta","Diptera","","","","","9","9","9","37","0","0","99.16","447.22","0","450","37.63","0"
"AGAW1","02N","INS","BOLD:AAI1981","Arthropoda","Insecta","Diptera","Chironomidae","Orthocladiinae","Gymnometriocnemus","","9","9","9","37","0","0","97.77","449.11","0","450","36.3","0"
"AGAW1","02N","INS","BOLD:AAO8538","Arthropoda","Insecta","Diptera","Psychodidae","Psychodinae","Psychoda","","9","9","9","37","0","0","99.3","428.44","0","430.89","32.43","0"
"AGAW1","02N","INS","BOLD:AAB2384","Arthropoda","Insecta","Diptera","Syrphidae","Syrphinae","Eupeodes","Eupeodes duseki, Eupeodes volucris, Eupeodes nielseni, Eupeodes americanus, Eupeodes flukei, Eupeodes vockerothi, Eupeodes lucasi, Eupeodes montivagus, Eupeodes rufipunctatus, Eupeodes rojasi, Eupeodes goeldlini, Eupeodes luniger, Eupeodes punctifer, Eupeodes bucculatus, Eupeodes perplexus, Eupeodes abiskoensis, Eupeodes corollae, Eupeodes pomus, Eupeodes lundbecki, Eupeodes nitens, Eupeodes sp. BOLDAAB2384, Eupeodes curtus, Eupeodes latifasciatus","8","8","8","40","0","0","98.74","433.75","0","434.88","31.05","0"
"AGAW1","02N","INS","BOLD:AAD4355","Arthropoda","Insecta","Diptera","Culicidae","Culicinae","Aedes","Aedes provocans","5","5","5","41","0","0","98.64","434.8","0","436.2","32.13","0"
"AGAW1","02N","INS","BOLD:ACC7611","Arthropoda","Insecta","Diptera","Cecidomyiidae","","","","4","4","4","42","0","0","97.54","446.75","0","450","28.28","0"
"AGAW1","02N","INS","BOLD:AAA4222","Arthropoda","Insecta","Lepidoptera","Erebidae","Hypeninae","Hypena","Hypena scabra","4","4","4","42","0","0","99.38","428.25","0","431","32.76","0"
"AGAW1","02N","INS","BOLD:AAG2462","Arthropoda","Insecta","Diptera","Anthomyiidae","Anthomyiinae","Botanophila","Botanophila fugax","4","4","4","42","0","0","99.28","403.5","0","404.25","31.87","0"
"AGAW1","02N","INS","BOLD:ABU9891","Arthropoda","Insecta","Diptera","Muscidae","Phaoniinae","Phaonia","","4","4","4","42","0","0","98.87","443.25","0","449.25","32.67","0"
"AGAW1","02N","INS","BOLD:AAN5356","Arthropoda","Insecta","Diptera","Chironomidae","Orthocladiinae","Smittia","","3","3","3","46","0","0","98.96","448","0","450","35.33","0"
"AGAW1","02N","INS","BOLD:ABU5546","Arthropoda","Insecta","Diptera","Psychodidae","Psychodinae","Psychoda","","3","3","3","46","0","0","98.67","450.33","0","450","31.26","0"
"AGAW1","02N","INS","BOLD:AAN8583","Arthropoda","Insecta","Diptera","Mycetophilidae","Sciophilinae","Sciophila","Sciophila pseudoflexuosa, Sciophila lutea","2","2","2","48","0","0","97.53","445.5","0","450","33.56","0"
"AGAW1","02N","INS","BOLD:AAL2750","Arthropoda","Insecta","Diptera","Muscidae","Phaoniinae","Phaonia","Phaonia tuguriorum","2","2","2","48","0","0","98.1","448.5","0","450","33.56","0"
"AGAW1","02N","INS","BOLD:AAG6745","Arthropoda","Insecta","Diptera","Calliphoridae","Calliphorinae","Pollenia","Pollenia pediculata","2","2","2","48","0","0","99.55","418.5","0","420","33.81","0"
"AGAW1","02N","INS","BOLD:AAA8933","Arthropoda","Insecta","Coleoptera","Coccinellidae","Coccinellinae","Coccinella","Coccinella septempunctata","2","2","2","48","0","0","98.66","448","0","450","32.78","0"
"AGAW1","02N","INS","BOLD:ACV3021","Arthropoda","Insecta","Diptera","Cecidomyiidae","Cecidomyiinae","Contarinia","Contarinia verrucicola","2","2","2","48","0","0","99.78","449.5","0","450","29.56","0"
"AGAW1","02N","INS","BOLD:AAD0583","Arthropoda","Insecta","Hemiptera","Lygaeidae","Lygaeinae","Lygaeus","Lygaeus reclivatus, Lygaeus kalmii","1","1","1","53","0","0","97.54","447","0","450","32.22","0"
"AGAW1","02N","INS","BOLD:AAG9055","Arthropoda","Insecta","Hymenoptera","Vespidae","Vespinae","Vespula","Vespula germanica","1","1","1","53","0","0","99.11","450","0","450","31.11","0"
"AGAW1","02N","INS","BOLD:ACV3368","","","","","","","","1","1","1","53","0","0","99.76","422","0","422","32.94","0"
"AGAW1","02N","INS","BOLD:AAK8831","Arthropoda","Insecta","Diptera","Limoniidae","Limoniinae","Metalimnobia, Limonia","","1","1","1","53","0","0","98.67","375","0","376","33.24","0"
"AGAW1","02N","INS","BOLD:AAG1711","Arthropoda","Insecta","Diptera","Muscidae","Phaoniinae","Helina","Helina depuncta","1","1","1","53","0","0","99.74","387","0","387","31.01","0"
"AGAW1","02N","INS","BOLD:AAA8109","Arthropoda","Insecta","Lepidoptera","Gelechiidae","Dichomeridinae","Dichomeris","Dichomeris ligulella","1","1","1","53","0","0","99.11","450","0","450","32.89","0"
"AGAW1","02N","INS","BOLD:AAE4299","Arthropoda","Insecta","Diptera","Chironomidae","Orthocladiinae","Cricotopus","","1","1","1","53","0","0","97.98","445","0","450","33.78","0"
"AGAW1","02N","INS","BOLD:AAK5928","Arthropoda","Insecta","Coleoptera","Chrysomelidae","Criocerinae","Oulema","Oulema melanopus","1","1","1","53","0","0","99.33","446","0","450","36.44","0"
"AGAW1","02N","INS","BOLD:AAC9062","Arthropoda","Insecta","Diptera","Culicidae","Culicinae","Ochlerotatus, Aedes","Aedes fitchii, Ochlerotatus fitchii, Aedes grossbecki, Aedes ventrovittis","1","1","1","53","0","0","100","360","0","362","32.87","0"
"AGAW1","02N","INS","BOLD:AAA7067","Arthropoda","Insecta","Diptera","Culicidae","Culicinae","Aedes, Aedimorphus","Aedimorphus vexans, Aedes vexans","1","1","1","53","0","0","97.54","448","0","450","33.11","0"
"AGAW1","02N","INS","BOLD:ACA2954","Arthropoda","Insecta","Diptera","Chironomidae","Orthocladiinae","","","1","1","1","53","0","0","99.74","382","0","383","36.03","0"
"AGAW1","02N","INS","BOLD:AAB5640","Arthropoda","Insecta","Coleoptera","Coccinellidae","Coccinellinae","Harmonia","Harmonia axyridis","1","1","1","53","0","0","97.27","256","0","450","31.56","0"
"AGAW1","02N","INS","BOLD:ACF4289","Arthropoda","Insecta","Coleoptera","Staphylinidae","","","","1","1","1","53","0","0","98.89","449","0","450","34","0"
"AGAW1","02N","INS","BOLD:ADC6237","Arthropoda","Insecta","Diptera","Chironomidae","Chironominae","Micropsectra","","1","1","1","53","0","0","99.75","400","0","401","32.92","0"
"AGAW1","02N","INS","BOLD:AAO3939","Arthropoda","Insecta","Diptera","Limoniidae","Limoniinae","Dicranomyia","","1","1","1","53","0","0","99.78","450","0","450","36.44","0"
"AGAW1","02N","INS","BOLD:AAL7534","Arthropoda","Insecta","Diptera","Anisopodidae","Anisopodinae","Sylvicola","Sylvicola stackelbergi, Sylvicola stackelbergii 1, Sylvicola punctatus","1","1","1","53","0","0","99.33","450","0","450","34","0"
"AGAW1","02N","INS","BOLD:AAD2085","Arthropoda","Insecta","Lepidoptera","Bucculatricidae","","Bucculatrix","Bucculatrix angustata","1","1","1","53","0","0","99.33","450","0","450","33.11","0"
"AGAW1","02N","INS","BOLD:ACN9514","","","","","","","","1","1","1","53","0","0","97.84","417","0","423","33.1","0"
"AGAW1","02N","INS","BOLD:AAA1513","Arthropoda","Insecta","Lepidoptera","Plutellidae","Plutellinae","Plutella, Rhigognostis","Plutella xylostella, Rhigognostis sp.","1","1","1","53","0","0","99.55","449","0","450","32.67","0"
"AGAW1","02N","INS","BOLD:AAM7726","Arthropoda","Insecta","Coleoptera","Curculionidae","Curculioninae","Orchestes","Orchestes steppensis, Orchestes alni","1","1","1","53","0","0","99.11","450","0","450","33.78","0"
"AGAW1","02N","INS","BOLD:AAG7306","Arthropoda","Insecta","Diptera","Sphaeroceridae","Limosininae","Minilimosina","Minilimosina vitripennis","1","1","1","53","0","0","97.97","444","0","450","33.33","0"
"AGAW1","02N","INS","BOLD:ABA5313","Arthropoda","Insecta","Coleoptera","Staphylinidae","Aleocharinae","Amischa","Amischa analis, Amischa bifoveolata, Amischa nigrofusca","1","1","1","53","0","0","99.55","449","0","450","33.56","0"
"AGAW1","02N","NAR","no match","no match","no match","no match","no match","no match","no match","no match","no match","no match","no match","no match","no match","no match","no match","no match","no match","no match","no match","no match"
"AGAW1","02N","NIN","BOLD:AAB5726","Arthropoda","Arachnida","Araneae","Agelenidae",NA,"Agelenopsis","Agelenopsis pennsylvanica, Agelenopsis emertoni, Agelenopsis potteri, Agelenopsis actuosa, Agelenopsis kastoni","1","1","1","1","0","0","99.78","449","0","450","33.33","0"
"AGAW1","02S","BAC","TAX:175768","Proteobacteria","Alphaproteobacteria","Rickettsiales","Rickettsiaceae",NA,"Wolbachia","Wolbachia pipientis","17","15","15","1","11.76","0","98.68","447.47","0","448.41","39.54","0"
"AGAW1","02S","BAC","TAX:91391","Proteobacteria","Alphaproteobacteria","Rickettsiales","",NA,"","","6","3","3","2","50","0","98.87","397.83","0","442.83","40.76","0"
"AGAW1","02S","BAC","TAX:91393","Proteobacteria","Alphaproteobacteria","Rickettsiales","Rickettsiaceae",NA,"Wolbachia","","5","5","5","3","0","0","99.69","391.2","0","427","39.09","0"
"AGAW1","02S","BAC","TAX:91389","Proteobacteria","","","",NA,"","","1","1","1","4","0","0","98.9","362","0","372","38.98","0"
"AGAW1","02S","CHO","no match","no match","no match","no match","no match","no match","no match","no match","no match","no match","no match","no match","no match","no match","no match","no match","no match","no match","no match","no match"
"AGAW1","02S","INS","BOLD:AAB0868","Arthropoda","Insecta","Diptera","Calliphoridae","Calliphorinae","Cynomya","Cynomya cadaverina, Cynomya mortuorum","38774","24176","24176","1","37.65","0","99.52","436.93","0","438.7","34.97","0"
"AGAW1","02S","INS","BOLD:AAG2511","Arthropoda","Insecta","Diptera","Anthomyiidae","Anthomyiinae","Delia","Delia platura","34712","23924","23924","2","31.08","0","99.37","443.21","0","445.51","33.22","0"
"AGAW1","02S","INS","BOLD:AAA3453","Arthropoda","Insecta","Diptera","Anthomyiidae","Anthomyiinae","Delia","","9030","6514","6514","3","27.86","0","99.37","443.97","0","445.96","34.79","0"
"AGAW1","02S","INS","BOLD:AAN5355","Arthropoda","Insecta","Diptera","Chironomidae","Orthocladiinae","Smittia","Smittia cf. stercoraria, Smittia stercoraria","3590","3053","3053","4","14.96","0","99.18","415.13","0","445.88","36.33","0"
"AGAW1","02S","INS","BOLD:AAB2644","Arthropoda","Insecta","Diptera","Chironomidae","Orthocladiinae","Orthocladius","Orthocladius lapponicus, Orthocladius dorenus, Orthocladius mallochi","2432","1967","1967","5","19.12","0","99.29","442.05","0","443.87","32.44","0"
"AGAW1","02S","INS","BOLD:AAB4658","Arthropoda","Insecta","Diptera","Chironomidae","Chironominae","Chironomus","Chironomus dilutus","2269","2132","2132","6","6.04","0","98.47","443.56","0","446.25","37.86","0"
"AGAW1","02S","INS","BOLD:AAA9251","Arthropoda","Insecta","Lepidoptera","Gelechiidae","Gelechiinae","Scrobipalpa","Scrobipalpa atriplicella, Scrobipalpa nr. atriplicella sp. 1","1242","952","952","7","23.35","0","99.51","440.89","0","442.67","34.5","0"
"AGAW1","02S","INS","BOLD:ACF8552","Arthropoda","Insecta","Coleoptera","Anthicidae","Anthicinae","Malporus","Malporus formicarius","769","541","541","8","29.65","0","99.77","364.69","0","446.02","34.28","0"
"AGAW1","02S","INS","BOLD:AAN5358","Arthropoda","Insecta","Diptera","Chironomidae","Orthocladiinae","Smittia","Smittia stercoraria, Smittia sp. 22ES","556","481","481","9","13.49","0","99.22","443.28","0","445.27","37.32","0"
"AGAW1","02S","INS","BOLD:AAB1098","Arthropoda","Insecta","Diptera","Culicidae","Culicinae","Ochlerotatus, Aedes","Aedes excrucians, Aedes cantans, Aedes stimulans, Aedes euedes, Aedes riparius, Aedes annulipes, Ochlerotatus cantans, Aedes fitchii, Aedes sp., Ochlerotatus annulipes, Ochlerotatus excrucians","377","355","355","10","5.84","0","98.54","430.84","0","431.8","35.38","0"
"AGAW1","02S","INS","BOLD:AAC4197","Arthropoda","Insecta","Diptera","Chironomidae","Orthocladiinae","Paraphaenocladius","Paraphaenocladius impensus","303","279","279","11","7.92","0","98.98","435.33","0","440.86","34.75","0"
"AGAW1","02S","INS","BOLD:ACP4736","Arthropoda","Insecta","Diptera","Chironomidae","Orthocladiinae","Smittia","Smittia sp. 8ES","159","136","136","12","14.47","0","99.27","432.53","0","434.07","33.24","0"
"AGAW1","02S","INS","BOLD:AAN5267","Arthropoda","Insecta","Diptera","Cecidomyiidae","","","","148","117","117","13","20.95","0","99.23","447.11","0","448.25","30.64","0"
"AGAW1","02S","INS","BOLD:AAG9713","Arthropoda","Insecta","Diptera","Dolichopodidae","Sympycninae","Sympycnus","Sympycnus lineatus","136","111","111","14","18.38","0","98.9","447.07","0","448.51","35.49","0"
"AGAW1","02S","INS","BOLD:AAI4303","Arthropoda","Insecta","Diptera","Chironomidae","Chironominae","Chironomus","Chironomus melanescens","109","100","100","15","8.26","0","98.49","439.49","0","445.12","34.44","0"
"AGAW1","02S","INS","BOLD:ABA5313","Arthropoda","Insecta","Coleoptera","Staphylinidae","Aleocharinae","Amischa","Amischa analis, Amischa bifoveolata, Amischa nigrofusca","74","65","65","16","12.16","0","99.42","441.05","0","444.15","33.48","0"
"AGAW1","02S","INS","BOLD:AAA2482","Arthropoda","Insecta","Lepidoptera","Noctuidae","Noctuinae","Mythimna","Mythimna unipuncta","67","64","64","17","4.48","0","99.48","430.15","0","432.75","30.5","0"
"AGAW1","02S","INS","BOLD:AAG1000","Arthropoda","Insecta","Diptera","Chironomidae","Orthocladiinae","Orthocladius","Orthocladius carlatus","48","48","48","18","0","0","98","429.71","0","431.73","33.97","0"
"AGAW1","02S","INS","BOLD:ABA1215","Arthropoda","Insecta","Diptera","Sciaridae","","Lycoriella","Lycoriella sativae","46","40","40","19","13.04","0","99.25","440.17","0","441.65","35.78","0"
"AGAW1","02S","INS","BOLD:AAB9140","Arthropoda","Insecta","Diptera","Calliphoridae","Chrysomyinae","Phormia","Phormia regina","38","38","38","20","0","0","99.18","445.92","0","447.5","33.05","0"
"AGAW1","02S","INS","BOLD:AAO8538","Arthropoda","Insecta","Diptera","Psychodidae","Psychodinae","Psychoda","","30","29","29","21","3.33","0","99.12","445.2","0","446.63","33.1","0"
"AGAW1","02S","INS","BOLD:AAG4944","Arthropoda","Insecta","Diptera","Mycetophilidae","Sciophilinae","Sciophila","Sciophila sp. CJB1","30","30","30","21","0","0","97.87","420.87","0","423.43","33.9","0"
"AGAW1","02S","INS","BOLD:ACE3123","Arthropoda","Insecta","Diptera","Sciaridae","","Corynoptera","","27","23","23","23","14.81","0","99.37","441.48","0","442.26","35.33","0"
"AGAW1","02S","INS","BOLD:AAL7306","Arthropoda","Insecta","Diptera","Chironomidae","Chironominae","Micropsectra","","27","26","26","23","3.7","0","99.14","414.07","0","416.41","33.02","0"
"AGAW1","02S","INS","BOLD:ACL8053","Arthropoda","Insecta","Diptera","Chironomidae","Chironominae","Tanytarsus","","19","18","18","25","5.26","0","99.31","445.42","0","446.79","33.9","0"
"AGAW1","02S","INS","BOLD:ACF7678","Arthropoda","Insecta","Coleoptera","Staphylinidae","Aleocharinae","Dinaraea","Dinaraea angustula","12","12","12","26","0","0","99.14","378","0","429.17","36.15","0"
"AGAW1","02S","INS","BOLD:AAM7726","Arthropoda","Insecta","Coleoptera","Curculionidae","Curculioninae","Orchestes","Orchestes steppensis, Orchestes alni","12","12","12","26","0","0","99.51","443.25","0","444.67","32.95","0"
"AGAW1","02S","INS","BOLD:ABV1455","Arthropoda","Insecta","Diptera","Chironomidae","Chironominae","","","9","8","8","28","11.11","0","97.72","440.67","0","443.11","31.96","0"
"AGAW1","02S","INS","BOLD:AAY6412","Arthropoda","Insecta","Diptera","Sciaridae","","Bradysia","Bradysia urticae","8","8","8","29","0","0","98.8","445.75","0","447.25","32.17","0"
"AGAW1","02S","INS","BOLD:AAB2384","Arthropoda","Insecta","Diptera","Syrphidae","Syrphinae","Eupeodes","Eupeodes duseki, Eupeodes volucris, Eupeodes nielseni, Eupeodes americanus, Eupeodes flukei, Eupeodes vockerothi, Eupeodes lucasi, Eupeodes montivagus, Eupeodes rufipunctatus, Eupeodes rojasi, Eupeodes goeldlini, Eupeodes luniger, Eupeodes punctifer, Eupeodes bucculatus, Eupeodes perplexus, Eupeodes abiskoensis, Eupeodes corollae, Eupeodes pomus, Eupeodes lundbecki, Eupeodes nitens, Eupeodes sp. BOLDAAB2384, Eupeodes curtus, Eupeodes latifasciatus","7","7","7","30","0","0","98.82","447.57","0","449.57","31.2","0"
"AGAW1","02S","INS","BOLD:AAP6827","Arthropoda","Insecta","Diptera","Cecidomyiidae","","","","7","7","7","30","0","0","98.89","448.86","0","450","31.97","0"
"AGAW1","02S","INS","BOLD:AAG6433","Arthropoda","Insecta","Diptera","Ceratopogonidae","Forcipomyiinae","Forcipomyia","Forcipomyia sp. BOLD:AAG6433, Forcipomyia sp. 17","7","7","7","30","0","0","99.51","436.14","0","437.14","31.28","0"
"AGAW1","02S","INS","BOLD:AAC9062","Arthropoda","Insecta","Diptera","Culicidae","Culicinae","Ochlerotatus, Aedes","Aedes fitchii, Ochlerotatus fitchii, Aedes grossbecki, Aedes ventrovittis","4","4","4","33","0","0","98.77","448.75","0","450","33.44","0"
"AGAW1","02S","INS","BOLD:AAQ0582","Arthropoda","Insecta","Diptera","Anthomyiidae","Anthomyiinae","Lasiomma","","4","4","4","33","0","0","98.38","447.5","0","450","31.83","0"
"AGAW1","02S","INS","BOLD:AAD4355","Arthropoda","Insecta","Diptera","Culicidae","Culicinae","Aedes","Aedes provocans","3","3","3","35","0","0","98.96","449","0","450","32.37","0"
"AGAW1","02S","INS","BOLD:AAD0853","Arthropoda","Insecta","Diptera","Scathophagidae","Scathophaginae","Scathophaga","Scathophaga lutaria, Scathophaga hiemalis, Scathophaga furcata, Scathophaga stercoraria, Scathophaga inquinata, Scathophaga suilla","3","3","3","35","0","0","98.81","448","0","450","32.07","0"
"AGAW1","02S","INS","BOLD:AAK5928","Arthropoda","Insecta","Coleoptera","Chrysomelidae","Criocerinae","Oulema","Oulema melanopus","2","2","2","37","0","0","99.07","427.5","0","428.5","36.04","0"
"AGAW1","02S","INS","BOLD:ABU9891","Arthropoda","Insecta","Diptera","Muscidae","Phaoniinae","Phaonia","","2","2","2","37","0","0","99.09","394.5","0","398.5","31.51","0"
"AGAW1","02S","INS","BOLD:AAN6437","Arthropoda","Insecta","Diptera","Sciaridae","","Corynoptera","Corynoptera saccata","2","2","2","37","0","0","98.56","441","0","441.5","40.2","0"
"AGAW1","02S","INS","BOLD:AAA4222","Arthropoda","Insecta","Lepidoptera","Erebidae","Hypeninae","Hypena","Hypena scabra","2","2","2","37","0","0","98.35","423","0","425","32.56","0"
"AGAW1","02S","INS","BOLD:AAG6743","Arthropoda","Insecta","Diptera","Sarcophagidae","Sarcophaginae","Sarcophaga","Sarcophaga sp. 1 MG, Sarcophaga subvicina","2","2","2","37","0","0","100","447.5","0","450","35","0"
"AGAW1","02S","INS","BOLD:AAM9252","Arthropoda","Insecta","Diptera","Sciaridae","","Bradysia, Leptosciarella","Bradysia vagans, Leptosciarella zanti","2","2","2","37","0","0","98","448","0","450","38.44","0"
"AGAW1","02S","INS","BOLD:AAP9022","Arthropoda","Insecta","Diptera","Cecidomyiidae","","","","2","2","2","37","0","0","98.78","448.5","0","450","28","0"
"AGAW1","02S","INS","BOLD:AAK8831","Arthropoda","Insecta","Diptera","Limoniidae","Limoniinae","Metalimnobia, Limonia","","1","1","1","44","0","0","99.73","366","0","367","33.51","0"
"AGAW1","02S","INS","BOLD:AAL7534","Arthropoda","Insecta","Diptera","Anisopodidae","Anisopodinae","Sylvicola","Sylvicola stackelbergi, Sylvicola stackelbergii 1, Sylvicola punctatus","1","1","1","44","0","0","99.55","447","0","450","33.78","0"
"AGAW1","02S","INS","BOLD:AAN8583","Arthropoda","Insecta","Diptera","Mycetophilidae","Sciophilinae","Sciophila","Sciophila pseudoflexuosa, Sciophila lutea","1","1","1","44","0","0","97.01","402","0","406","33.25","0"
"AGAW1","02S","INS","BOLD:AAN5356","Arthropoda","Insecta","Diptera","Chironomidae","Orthocladiinae","Smittia","","1","1","1","44","0","0","97.32","447","0","450","35.11","0"
"AGAW1","02S","INS","BOLD:ACF6270","Arthropoda","Insecta","Diptera","Chironomidae","Orthocladiinae","Orthocladius","","1","1","1","44","0","0","99.55","445","0","450","30.89","0"
"AGAW1","02S","INS","BOLD:AAL2750","Arthropoda","Insecta","Diptera","Muscidae","Phaoniinae","Phaonia","Phaonia tuguriorum","1","1","1","44","0","0","98.45","451","0","450","32.89","0"
"AGAW1","02S","INS","BOLD:AAH0127","Arthropoda","Insecta","Coleoptera","Pyrochroidae","Pedilinae","Pedilus","Pedilus lugubris","1","1","1","44","0","0","99.25","399","0","450","33.11","0"
"AGAW1","02S","INS","BOLD:AAA2144","Arthropoda","Insecta","Lepidoptera","Noctuidae","Noctuinae","Xestia","Xestia kollari, Xestia c-nigrum, Xestia dolosa","1","1","1","44","0","0","98.15","432","0","439","31.21","0"
"AGAW1","02S","INS","BOLD:AAH1890","Arthropoda","Insecta","Hymenoptera","Ichneumonidae","Cryptinae","Stibeutes","Stibeutes brevicornis, Stibeutes heinemanni","1","1","1","44","0","0","100","450","0","450","29.78","0"
"AGAW1","02S","INS","BOLD:AAA4121","Arthropoda","Insecta","Diptera","Simuliidae","Simuliinae","Simulium","Simulium vittatum complex, Simulium vittatum, Simulium sp. BOLDAAA4121, Simulium argus, Simulium vittatum s.str.","1","1","1","44","0","0","99.73","369","0","371","33.42","0"
"AGAW1","02S","INS","BOLD:ABX9211","Arthropoda","Insecta","Diptera","Cecidomyiidae","","","","1","1","1","44","0","0","98.66","449","0","450","27.56","0"
"AGAW1","02S","INS","BOLD:AAG6745","Arthropoda","Insecta","Diptera","Calliphoridae","Calliphorinae","Pollenia","Pollenia pediculata","1","1","1","44","0","0","98.22","449","0","450","33.11","0"
"AGAW1","02S","INS","BOLD:AAA3080","Arthropoda","Insecta","Trichoptera","Limnephilidae","Limnephilinae","Limnephilus","Limnephilus submonilifer","1","1","1","44","0","0","98.21","448","0","450","34.67","0"
"AGAW1","02S","INS","BOLD:AAA4264","Arthropoda","Insecta","Diptera","Simuliidae","Simuliinae","Simulium","Simulium venustum/truncatum, Simulium venustum s.str., Simulium verecundum/verecundum complex, Simulium venustum complex, Simulium venustum, Simulium tormentor, Simulium truncatum, Simulium sp. BOLDAAA4264, Simulium venustum complex sp. JMR-2009","1","1","1","44","0","0","97.33","449","0","450","36.67","0"
"AGAW1","02S","INS","BOLD:ACR4394","Arthropoda","Insecta","Diptera","Anthomyiidae","Anthomyiinae","Delia","Delia florilega","1","1","1","44","0","0","98.17","328","0","350","32","0"
"AGAW1","02S","INS","BOLD:AAG1742","Arthropoda","Insecta","Diptera","Muscidae","Phaoniinae","Helina","Helina rufitibia","1","1","1","44","0","0","99.1","446","0","450","29.78","0"
"AGAW1","02S","INS","BOLD:ACN9514","","","","","","","","1","1","1","44","0","0","97.35","377","0","378","33.6","0"
"AGAW1","02S","INS","BOLD:ACA9275","Arthropoda","Insecta","Diptera","Chironomidae","Chironominae","Chironomus","","1","1","1","44","0","0","99.55","449","0","450","36.44","0"
"AGAW1","02S","INS","BOLD:AAA2955","Arthropoda","Insecta","Lepidoptera","Tortricidae","Tortricinae","Argyrotaenia","Argyrotaenia occultana, Argyrotaenia ljungiana, Argyrotaenia pinatubana, Argyrotaenia floridana, Argyrotaenia velutinana, Argyrotaenia tabulana, Argyrotaenia gogana","1","1","1","44","0","0","99.55","449","0","450","32.22","0"
"AGAW1","02S","INS","BOLD:ABW2407","Arthropoda","Insecta","Diptera","Anthomyiidae","Anthomyiinae","Hylemya","","1","1","1","44","0","0","99.76","422","0","427","32.08","0"
"AGAW1","02S","INS","BOLD:ACX5521","Arthropoda","Insecta","Diptera","Chironomidae","Orthocladiinae","","","1","1","1","44","0","0","98.89","449","0","450","34.89","0"
"AGAW1","02S","INS","BOLD:ACF4289","Arthropoda","Insecta","Coleoptera","Staphylinidae","","","","1","1","1","44","0","0","99.78","448","0","450","33.78","0"
"AGAW1","02S","INS","BOLD:AAG2462","Arthropoda","Insecta","Diptera","Anthomyiidae","Anthomyiinae","Botanophila","Botanophila fugax","1","1","1","44","0","0","97.77","449","0","450","32","0"
"AGAW1","02S","NAR","no match","no match","no match","no match","no match","no match","no match","no match","no match","no match","no match","no match","no match","no match","no match","no match","no match","no match","no match","no match"
"AGAW1","02S","NIN","BOLD:AAI4087","Arthropoda","Arachnida","Araneae","Clubionidae",NA,"Clubiona","Clubiona pallidula","1","1","1","1","0","0","99.32","440","0","450","32","0"
"AGAW1","02W","BAC","TAX:175768","Proteobacteria","Alphaproteobacteria","Rickettsiales","Rickettsiaceae",NA,"Wolbachia","Wolbachia pipientis","125","93","93","1","25.6","0","98.61","448.54","0","449.41","39.58","0"
"AGAW1","02W","BAC","TAX:91393","Proteobacteria","Alphaproteobacteria","Rickettsiales","Rickettsiaceae",NA,"Wolbachia","","42","42","42","2","0","0","99.48","391.17","0","424.74","39.26","0"
"AGAW1","02W","CHO","no match","no match","no match","no match","no match","no match","no match","no match","no match","no match","no match","no match","no match","no match","no match","no match","no match","no match","no match","no match"
"AGAW1","02W","INS","BOLD:AAA8933","Arthropoda","Insecta","Coleoptera","Coccinellidae","Coccinellinae","Coccinella","Coccinella septempunctata","31889","17244","17244","1","45.92","0","99.67","446.49","0","447.45","33.05","0"
"AGAW1","02W","INS","BOLD:AAA3453","Arthropoda","Insecta","Diptera","Anthomyiidae","Anthomyiinae","Delia","","30653","20627","20627","2","32.71","0","99.49","443.96","0","446.01","34.78","0"
"AGAW1","02W","INS","BOLD:AAG2511","Arthropoda","Insecta","Diptera","Anthomyiidae","Anthomyiinae","Delia","Delia platura","24582","17758","17758","3","27.76","0","99.31","443.24","0","445.48","33.2","0"
"AGAW1","02W","INS","BOLD:AAB2384","Arthropoda","Insecta","Diptera","Syrphidae","Syrphinae","Eupeodes","Eupeodes duseki, Eupeodes volucris, Eupeodes nielseni, Eupeodes americanus, Eupeodes flukei, Eupeodes vockerothi, Eupeodes lucasi, Eupeodes montivagus, Eupeodes rufipunctatus, Eupeodes rojasi, Eupeodes goeldlini, Eupeodes luniger, Eupeodes punctifer, Eupeodes bucculatus, Eupeodes perplexus, Eupeodes abiskoensis, Eupeodes corollae, Eupeodes pomus, Eupeodes lundbecki, Eupeodes nitens, Eupeodes sp. BOLDAAB2384, Eupeodes curtus, Eupeodes latifasciatus","13187","10198","10198","4","22.67","0","98.82","430.15","0","432.26","30.95","0"
"AGAW1","02W","INS","BOLD:AAB2644","Arthropoda","Insecta","Diptera","Chironomidae","Orthocladiinae","Orthocladius","Orthocladius lapponicus, Orthocladius dorenus, Orthocladius mallochi","13033","10595","10595","5","18.71","0","98.84","438.55","0","440.7","32.84","0"
"AGAW1","02W","INS","BOLD:AAB1098","Arthropoda","Insecta","Diptera","Culicidae","Culicinae","Ochlerotatus, Aedes","Aedes excrucians, Aedes cantans, Aedes stimulans, Aedes euedes, Aedes riparius, Aedes annulipes, Ochlerotatus cantans, Aedes fitchii, Aedes sp., Ochlerotatus annulipes, Ochlerotatus excrucians","7675","6859","6859","6","10.63","0","98.85","429.7","0","431.16","35.16","0"
"AGAW1","02W","INS","BOLD:AAA9251","Arthropoda","Insecta","Lepidoptera","Gelechiidae","Gelechiinae","Scrobipalpa","Scrobipalpa atriplicella, Scrobipalpa nr. atriplicella sp. 1","7408","4875","4875","7","34.19","0","99.36","446.02","0","447.54","34.73","0"
"AGAW1","02W","INS","BOLD:AAN5358","Arthropoda","Insecta","Diptera","Chironomidae","Orthocladiinae","Smittia","Smittia stercoraria, Smittia sp. 22ES","950","784","784","8","17.47","0","99.32","444.29","0","446.22","37.32","0"
"AGAW1","02W","INS","BOLD:AAN5355","Arthropoda","Insecta","Diptera","Chironomidae","Orthocladiinae","Smittia","Smittia cf. stercoraria, Smittia stercoraria","902","821","821","9","8.98","0","99.28","439.76","0","442.86","36.34","0"
"AGAW1","02W","INS","BOLD:ACJ0208","Arthropoda","Insecta","Diptera","Chironomidae","Orthocladiinae","","","404","322","322","10","20.3","0","99.1","435.36","0","448.66","34.91","0"
"AGAW1","02W","INS","BOLD:ACP4736","Arthropoda","Insecta","Diptera","Chironomidae","Orthocladiinae","Smittia","Smittia sp. 8ES","207","188","188","11","9.18","0","99.32","429.75","0","431.56","33.07","0"
"AGAW1","02W","INS","BOLD:ABW7690","Arthropoda","Insecta","Hemiptera","Aphalaridae","Aphalarinae","","","184","170","170","12","7.61","0","99.23","373.54","0","441.6","32.1","0"
"AGAW1","02W","INS","BOLD:AAQ0602","Arthropoda","Insecta","Diptera","Chironomidae","Chironominae","Micropsectra","","125","109","109","13","12.8","0","99.24","447.17","0","448.42","36.52","0"
"AGAW1","02W","INS","BOLD:AAN5267","Arthropoda","Insecta","Diptera","Cecidomyiidae","","","","97","77","77","14","20.62","0","99.5","446.8","0","447.94","30.64","0"
"AGAW1","02W","INS","BOLD:AAG1000","Arthropoda","Insecta","Diptera","Chironomidae","Orthocladiinae","Orthocladius","Orthocladius carlatus","30","30","30","15","0","0","98.09","428.97","0","430.87","33.89","0"
"AGAW1","02W","INS","BOLD:AAB9140","Arthropoda","Insecta","Diptera","Calliphoridae","Chrysomyinae","Phormia","Phormia regina","28","28","28","16","0","0","99.28","444.29","0","446.07","33.03","0"
"AGAW1","02W","INS","BOLD:AAY6412","Arthropoda","Insecta","Diptera","Sciaridae","","Bradysia","Bradysia urticae","21","18","18","17","14.29","0","99.5","448.38","0","449.95","32.28","0"
"AGAW1","02W","INS","BOLD:AAO8538","Arthropoda","Insecta","Diptera","Psychodidae","Psychodinae","Psychoda","","20","20","20","18","0","0","98.58","430.05","0","431.8","32.4","0"
"AGAW1","02W","INS","BOLD:AAM9252","Arthropoda","Insecta","Diptera","Sciaridae","","Bradysia, Leptosciarella","Bradysia vagans, Leptosciarella zanti","16","16","16","19","0","0","99.52","444.31","0","445.75","39.03","0"
"AGAW1","02W","INS","BOLD:AAB0868","Arthropoda","Insecta","Diptera","Calliphoridae","Calliphorinae","Cynomya","Cynomya cadaverina, Cynomya mortuorum","10","10","10","20","0","0","98.75","429.1","0","430.8","34.52","0"
"AGAW1","02W","INS","BOLD:AAD0853","Arthropoda","Insecta","Diptera","Scathophagidae","Scathophaginae","Scathophaga","Scathophaga lutaria, Scathophaga hiemalis, Scathophaga furcata, Scathophaga stercoraria, Scathophaga inquinata, Scathophaga suilla","9","9","9","21","0","0","98.81","448.44","0","449.56","32.13","0"
"AGAW1","02W","INS","BOLD:AAN6447","Arthropoda","Insecta","Diptera","Sciaridae","","Corynoptera","Corynoptera perpusilla","6","6","6","22","0","0","98.98","428.83","0","433.67","36","0"
"AGAW1","02W","INS","BOLD:AAP9022","Arthropoda","Insecta","Diptera","Cecidomyiidae","","","","6","6","6","22","0","0","98.37","441.17","0","442.83","27.8","0"
"AGAW1","02W","INS","BOLD:ACF2924","","","","","","","","5","5","5","24","0","0","99.29","449.8","0","450","25.69","0"
"AGAW1","02W","INS","BOLD:AAN5356","Arthropoda","Insecta","Diptera","Chironomidae","Orthocladiinae","Smittia","","4","4","4","25","0","0","98.66","443.75","0","445","35.27","0"
"AGAW1","02W","INS","BOLD:AAN8770","Arthropoda","Insecta","Diptera","Psychodidae","Psychodinae","Psychoda","Psychoda trinodulosa","3","3","3","26","0","0","99.85","449.67","0","450","33.93","0"
"AGAW1","02W","INS","BOLD:ABU9891","Arthropoda","Insecta","Diptera","Muscidae","Phaoniinae","Phaonia","","3","3","3","26","0","0","98.35","445","0","450","32.89","0"
"AGAW1","02W","INS","BOLD:AAA4222","Arthropoda","Insecta","Lepidoptera","Erebidae","Hypeninae","Hypena","Hypena scabra","3","3","3","26","0","0","98.63","443","0","445","32.66","0"
"AGAW1","02W","INS","BOLD:ABX3225","Arthropoda","Insecta","Coleoptera","Chrysomelidae","Galerucinae","Psylliodes","Psylliodes cucullatus","3","3","3","26","0","0","98.96","447.67","0","450","33.85","0"
"AGAW1","02W","INS","BOLD:AAG6743","Arthropoda","Insecta","Diptera","Sarcophagidae","Sarcophaginae","Sarcophaga","Sarcophaga sp. 1 MG, Sarcophaga subvicina","3","3","3","26","0","0","99.77","442.67","0","444.67","35.16","0"
"AGAW1","02W","INS","BOLD:AAG2462","Arthropoda","Insecta","Diptera","Anthomyiidae","Anthomyiinae","Botanophila","Botanophila fugax","3","3","3","26","0","0","98.88","447.33","0","450","31.56","0"
"AGAW1","02W","INS","BOLD:ACL0119","Arthropoda","Insecta","Diptera","Chironomidae","Orthocladiinae","Paraphaenocladius","","2","1","1","32","50","0","99.55","449","0","450","33.78","0"
"AGAW1","02W","INS","BOLD:AAD4355","Arthropoda","Insecta","Diptera","Culicidae","Culicinae","Aedes","Aedes provocans","2","2","2","32","0","0","99.76","420.5","0","424","31.81","0"
"AGAW1","02W","INS","BOLD:ABW7747","Arthropoda","Insecta","Coleoptera","Staphylinidae","Aleocharinae","Nehemitropia","Nehemitropia lividipennis","2","2","2","32","0","0","98.54","446","0","450","36.56","0"
"AGAW1","02W","INS","BOLD:AAA3836","Arthropoda","Insecta","Lepidoptera","Noctuidae","Plusiinae","Autographa","Autographa precationis","2","2","2","32","0","0","98.55","448.5","0","450","32.11","0"
"AGAW1","02W","INS","BOLD:AAG7276","Arthropoda","Insecta","Diptera","Sphaeroceridae","Limosininae","Leptocera","Leptocera erythrocera","2","2","2","32","0","0","98.43","434.5","0","436","32.67","0"
"AGAW1","02W","INS","BOLD:AAB4093","Arthropoda","Insecta","Lepidoptera","Noctuidae","Noctuinae","Elaphria","Elaphria grata","2","2","2","32","0","0","98.65","443.5","0","450","30.44","0"
"AGAW1","02W","INS","BOLD:AAB2076","Arthropoda","Insecta","Lepidoptera","Tortricidae","Olethreutinae","Ancylis","Ancylis metamelana, Ancylis sp.","2","2","2","32","0","0","99.77","427","0","428.5","31.15","0"
"AGAW1","02W","INS","BOLD:AAG1742","Arthropoda","Insecta","Diptera","Muscidae","Phaoniinae","Helina","Helina rufitibia","2","2","2","32","0","0","99","449","0","450","31","0"
"AGAW1","02W","INS","BOLD:AAQ0582","Arthropoda","Insecta","Diptera","Anthomyiidae","Anthomyiinae","Lasiomma","","2","2","2","32","0","0","99.89","449.5","0","450","31.89","0"
"AGAW1","02W","INS","BOLD:AAG9055","Arthropoda","Insecta","Hymenoptera","Vespidae","Vespinae","Vespula","Vespula germanica","1","1","1","41","0","0","99.56","450","0","450","31.33","0"
"AGAW1","02W","INS","BOLD:AAL7534","Arthropoda","Insecta","Diptera","Anisopodidae","Anisopodinae","Sylvicola","Sylvicola stackelbergi, Sylvicola stackelbergii 1, Sylvicola punctatus","1","1","1","41","0","0","99.55","449","0","450","34","0"
"AGAW1","02W","INS","BOLD:AAA8109","Arthropoda","Insecta","Lepidoptera","Gelechiidae","Dichomeridinae","Dichomeris","Dichomeris ligulella","1","1","1","41","0","0","97.78","450","0","450","32.22","0"
"AGAW1","02W","INS","BOLD:AAQ0558","Arthropoda","Insecta","Thysanoptera","Thripidae","Thripinae","Anaphothrips","Anaphothrips obscurus","1","1","1","41","0","0","99.56","450","0","450","31.56","0"
"AGAW1","02W","INS","BOLD:AAC9062","Arthropoda","Insecta","Diptera","Culicidae","Culicinae","Ochlerotatus, Aedes","Aedes fitchii, Ochlerotatus fitchii, Aedes grossbecki, Aedes ventrovittis","1","1","1","41","0","0","99.55","449","0","450","33.56","0"
"AGAW1","02W","INS","BOLD:AAG6745","Arthropoda","Insecta","Diptera","Calliphoridae","Calliphorinae","Pollenia","Pollenia pediculata","1","1","1","41","0","0","99.56","450","0","450","33.78","0"
"AGAW1","02W","INS","BOLD:ABA1215","Arthropoda","Insecta","Diptera","Sciaridae","","Lycoriella","Lycoriella sativae","1","1","1","41","0","0","99.78","448","0","450","35.78","0"
"AGAW1","02W","INS","BOLD:AAA7374","Arthropoda","Insecta","Diptera","Syrphidae","Syrphinae","Sphaerophoria","Sphaerophoria bankowskae, Sphaerophoria infuscata, Sphaerophoria nr. philanthus, Sphaerophoria batava, Sphaerophoria sp., Sphaerophoria rueppellii, Sphaerophoria interrupta-group, Sphaerophoria sp. MA-1, Sphaerophoria cleoae, Sphaerophoria asymmetrica, Sphaerophoria fatarum, Sphaerophoria bengalensis, Sphaerophoria bifurcata, Sphaerophoria taeniata, Sphaerophoria interrupta, Sphaerophoria contigua, Sphaerophoria kaa, Sphaerophoria laurae, Sphaerophoria chongjini, Sphaerophoria boreoalpina, Sphaerophoria sp. BOLDAAA7374, Sphaerophoria macrogaster, Sphaerophoria sulphuripes, Sphaerophoria brevipilosa, Sphaerophoria philanthus, Sphaerophoria pyrrhina, Sphaerophoria abbreviata, Sphaerophoria scripta, Sphaerophoria potentillae, Sphaerophoria longipilosa, Sphaerophoria virgata","1","1","1","41","0","0","99.11","449","0","450","31.56","0"
"AGAW1","02W","INS","BOLD:AAA4121","Arthropoda","Insecta","Diptera","Simuliidae","Simuliinae","Simulium","Simulium vittatum complex, Simulium vittatum, Simulium sp. BOLDAAA4121, Simulium argus, Simulium vittatum s.str.","1","1","1","41","0","0","99.75","405","0","407","32.43","0"
"AGAW1","02W","INS","BOLD:AAB5640","Arthropoda","Insecta","Coleoptera","Coccinellidae","Coccinellinae","Harmonia","Harmonia axyridis","1","1","1","41","0","0","100","450","0","450","32.44","0"
"AGAW1","02W","INS","BOLD:AAG1021","Arthropoda","Insecta","Diptera","Chironomidae","Orthocladiinae","Bryophaenocladius","Bryophaenocladius sp. 8ES","1","1","1","41","0","0","99.33","449","0","450","36.67","0"
"AGAW1","02W","INS","BOLD:AAB4658","Arthropoda","Insecta","Diptera","Chironomidae","Chironominae","Chironomus","Chironomus dilutus","1","1","1","41","0","0","97.54","447","0","450","37.11","0"
"AGAW1","02W","INS","BOLD:ABW2407","Arthropoda","Insecta","Diptera","Anthomyiidae","Anthomyiinae","Hylemya","","1","1","1","41","0","0","100","449","0","450","32.67","0"
"AGAW1","02W","INS","BOLD:ACK1651","Arthropoda","Insecta","Diptera","Cecidomyiidae","","","","1","1","1","41","0","0","99.55","449","0","450","26.89","0"
"AGAW1","02W","INS","BOLD:AAH3920","Arthropoda","Insecta","Diptera","Sciaridae","","Dichopygina, Scatopsciara","","1","1","1","41","0","0","97.49","399","0","450","35.33","0"
"AGAW1","02W","INS","BOLD:AAA3080","Arthropoda","Insecta","Trichoptera","Limnephilidae","Limnephilinae","Limnephilus","Limnephilus submonilifer","1","1","1","41","0","0","99.78","448","0","450","35.33","0"
"AGAW1","02W","INS","BOLD:ABU5525","Arthropoda","Insecta","Diptera","Chironomidae","Orthocladiinae","Limnophyes","Limnophyes sp. 14ES","1","1","1","41","0","0","99.78","449","0","450","37.78","0"
"AGAW1","02W","INS","BOLD:AAN7643","Arthropoda","Insecta","Hymenoptera","Tenthredinidae","Heterarthrinae","Fenusa","Fenusa dohrnii, Fenusa ulmi, Fenusa altenhoferi","1","1","1","41","0","0","99.55","449","0","450","30.89","0"
"AGAW1","02W","NAR","no match","no match","no match","no match","no match","no match","no match","no match","no match","no match","no match","no match","no match","no match","no match","no match","no match","no match","no match","no match"
"AGAW1","02W","NIN","BOLD:ACF8020","Arthropoda","Arachnida","Mesostigmata","Ascidae",NA,NA,NA,"2","2","2","1","0","0","99.56","449.5","0","450","36.78","0"
"AGAW1","05E","BAC","TAX:375853","Proteobacteria","Alphaproteobacteria","Rickettsiales","Rickettsiaceae",NA,"Wolbachia","Wolbachia sp. Hymenoptera_ASGLE646-10","12","11","11","1","8.33","0","97.64","448.83","0","449.33","40.95","0"
"AGAW1","05E","BAC","TAX:376688","Proteobacteria","Alphaproteobacteria","Rickettsiales","Rickettsiaceae",NA,"Wolbachia","Wolbachia sp. elachBioLep01 BioLep373","2","1","1","2","50","0","98","450","0","450","41.11","0"
"AGAW1","05E","BAC","TAX:392873","Proteobacteria","Alphaproteobacteria","Rickettsiales","Rickettsiaceae",NA,"Wolbachia","Wolbachia endosymbiont of Scrapter algoensis","2","2","2","2","0","0","97.69","412","0","448","41.18","0"
"AGAW1","05E","BAC","TAX:376694","Proteobacteria","Alphaproteobacteria","Rickettsiales","Rickettsiaceae",NA,"Wolbachia","Wolbachia sp. Anacrusis nephrodes","1","1","1","4","0","0","98.21","447","0","450","41.11","0"
"AGAW1","05E","BAC","TAX:375845","Proteobacteria","Alphaproteobacteria","Rickettsiales","Rickettsiaceae",NA,"Wolbachia","Wolbachia sp. Hymenoptera_ASGLE1440-10","1","1","1","4","0","0","98.42","442","0","446","41.03","0"
"AGAW1","05E","BAC","TAX:375772","Proteobacteria","Alphaproteobacteria","Rickettsiales","Rickettsiaceae",NA,"Wolbachia","Wolbachia sp. Leptogenys JCR25_ASANE483-10","1","1","1","4","0","0","97.54","447","0","450","40.89","0"
"AGAW1","05E","BAC","TAX:392914","Proteobacteria","Alphaproteobacteria","Rickettsiales","Rickettsiaceae",NA,"Wolbachia","Wolbachia endosymbiont of chalJanzen01 Janzen01","1","1","1","4","0","0","97.05","441","0","450","40.89","0"
"AGAW1","05E","BAC","TAX:392917","Proteobacteria","Alphaproteobacteria","Rickettsiales","Rickettsiaceae",NA,"Wolbachia","Wolbachia endosymbiont of Glyptapanteles Whitfield131","1","1","1","4","0","0","97.78","450","0","450","40.89","0"
"AGAW1","05E","CHO","no match","no match","no match","no match","no match","no match","no match","no match","no match","no match","no match","no match","no match","no match","no match","no match","no match","no match","no match","no match"
"AGAW1","05E","INS","BOLD:AAC2498","Arthropoda","Insecta","Diptera","Muscidae","Phaoniinae","Helina","Helina evecta, Helina aldrichi","22074","14953","14953","1","32.26","0","98.56","436.41","0","437.85","30.79","0"
"AGAW1","05E","INS","BOLD:AAG2511","Arthropoda","Insecta","Diptera","Anthomyiidae","Anthomyiinae","Delia","Delia platura","10955","9076","9076","2","17.15","0","99.05","444.5","0","447.04","33.18","0"
"AGAW1","05E","INS","BOLD:AAD4355","Arthropoda","Insecta","Diptera","Culicidae","Culicinae","Aedes","Aedes provocans","6972","6121","6121","3","12.21","0","98.89","431.53","0","433.62","32.06","0"
"AGAW1","05E","INS","BOLD:AAB7030","Arthropoda","Insecta","Diptera","Chironomidae","Chironominae","Chironomus","Chironomus sp. BOLDAAP2996, Chironomus quinnitukqut, Chironomus cf. decorus 2g, Chironomus decorus group sp. 2 IP2013, Chironomus cf. decorus, Chironomus decorus gr.","1734","1727","1727","4","0.4","0","98.3","405.62","0","409.85","34.32","0"
"AGAW1","05E","INS","BOLD:AAI4303","Arthropoda","Insecta","Diptera","Chironomidae","Chironominae","Chironomus","Chironomus melanescens","1727","1414","1414","5","18.12","0","98.65","433.31","0","436.62","34.55","0"
"AGAW1","05E","INS","BOLD:AAF9771","Arthropoda","Insecta","Diptera","Hybotidae","Tachydromiinae","Platypalpus","Platypalpus longiseta, Platypalpus trivialis, Platypalpus sp. 5, Platypalpus sp. 17, Platypalpus pallidiventris","1514","1145","1145","6","24.37","0","99.1","444.23","0","444.76","35.05","0"
"AGAW1","05E","INS","BOLD:AAO3939","Arthropoda","Insecta","Diptera","Limoniidae","Limoniinae","Dicranomyia","","1076","858","858","7","20.26","0","98.87","438.23","0","439.52","35.81","0"
"AGAW1","05E","INS","BOLD:AAV1261","Arthropoda","Insecta","Diptera","Sciaridae","","Bradysia","Bradysia splendida","902","801","801","8","11.2","0","99.2","434.56","0","435.73","34.11","0"
"AGAW1","05E","INS","BOLD:AAN5342","Arthropoda","Insecta","Diptera","Chironomidae","Orthocladiinae","Heleniella","","472","436","436","9","7.63","0","98.98","418.71","0","427.03","35.12","0"
"AGAW1","05E","INS","BOLD:ADB5997","Arthropoda","Insecta","Diptera","Muscidae","Phaoniinae","Helina","","399","393","393","10","1.5","0","97.38","445.17","0","448.04","30.81","0"
"AGAW1","05E","INS","BOLD:AAG4773","Arthropoda","Insecta","Diptera","Agromyzidae","","","","295","295","295","11","0","0","98.35","412.11","0","414.66","33.42","0"
"AGAW1","05E","INS","BOLD:ACA3730","","","","","","","","237","214","214","12","9.7","0","98.22","435.57","0","436.85","35.07","0"
"AGAW1","05E","INS","BOLD:AAF9014","Arthropoda","Insecta","Diptera","Limoniidae","Chioneinae","Symplecta","Symplecta cana, Symplecta sp. BOLDAAF9014","216","200","200","13","7.41","0","98.37","431.01","0","436.61","33.79","0"
"AGAW1","05E","INS","BOLD:ACP4736","Arthropoda","Insecta","Diptera","Chironomidae","Orthocladiinae","Smittia","Smittia sp. 8ES","208","200","200","14","3.85","0","99.18","417.83","0","419.44","33.06","0"
"AGAW1","05E","INS","BOLD:AAH3227","Arthropoda","Insecta","Psocodea","Lachesillidae","Lachesillinae","Lachesilla","","164","140","140","15","14.63","0","99.03","443.68","0","444.63","40.36","0"
"AGAW1","05E","INS","BOLD:AAI3059","Arthropoda","Insecta","Lepidoptera","Tineidae","Nemapogoninae","Nemapogon","Nemapogon variatella","143","117","117","16","18.18","0","99.61","441.61","0","442.36","30.38","0"
"AGAW1","05E","INS","BOLD:AAN5355","Arthropoda","Insecta","Diptera","Chironomidae","Orthocladiinae","Smittia","Smittia cf. stercoraria, Smittia stercoraria","143","142","142","16","0.7","0","98.85","431.83","0","434.97","36.21","0"
"AGAW1","05E","INS","BOLD:AAN6447","Arthropoda","Insecta","Diptera","Sciaridae","","Corynoptera","Corynoptera perpusilla","139","95","95","18","31.65","0","99.28","448.12","0","449","36.27","0"
"AGAW1","05E","INS","BOLD:ABU5521","Arthropoda","Insecta","Diptera","Sciaridae","","Corynoptera","Corynoptera furcata","108","101","101","19","6.48","0","98.62","414.23","0","424.51","36.02","0"
"AGAW1","05E","INS","BOLD:ABU5525","Arthropoda","Insecta","Diptera","Chironomidae","Orthocladiinae","Limnophyes","Limnophyes sp. 14ES","105","105","105","20","0","0","98.84","395.01","0","397.43","37.15","0"
"AGAW1","05E","INS","BOLD:AAG6441","Arthropoda","Insecta","Diptera","Ceratopogonidae","Ceratopogoninae","Culicoides","Culicoides stilobezzioides","93","73","73","21","21.51","0","98.15","435.14","0","435.78","35.48","0"
"AGAW1","05E","INS","BOLD:AAB1787","Arthropoda","Insecta","Hemiptera","Aphididae","Aphidinae","Rhopalosiphum, Acyrthosiphon","Acyrthosiphon pisum, Acyrthosiphon pisum 5A, Rhopalosiphum maidis","93","77","77","21","17.2","0","99.53","432.96","0","435.46","25.58","0"
"AGAW1","05E","INS","BOLD:AAM9252","Arthropoda","Insecta","Diptera","Sciaridae","","Bradysia, Leptosciarella","Bradysia vagans, Leptosciarella zanti","82","82","82","23","0","0","99.26","439.09","0","441.76","38.95","0"
"AGAW1","05E","INS","BOLD:ACK3431","Arthropoda","Insecta","Diptera","Cecidomyiidae","","","","81","76","76","24","6.17","0","99.34","427.22","0","428.72","32.34","0"
"AGAW1","05E","INS","BOLD:AAN6433","Arthropoda","Insecta","Diptera","Sciaridae","","Corynoptera","Corynoptera deserta","76","75","75","25","1.32","0","98.96","435.25","0","437.62","30.51","0"
"AGAW1","05E","INS","BOLD:AAC4197","Arthropoda","Insecta","Diptera","Chironomidae","Orthocladiinae","Paraphaenocladius","Paraphaenocladius impensus","73","62","62","26","15.07","0","99.35","441.44","0","445.01","35","0"
"AGAW1","05E","INS","BOLD:ADF6661","Arthropoda","Insecta","Diptera","Cecidomyiidae","","","","47","41","41","27","12.77","0","99.25","415.64","0","443.94","29.87","0"
"AGAW1","05E","INS","BOLD:AAA3453","Arthropoda","Insecta","Diptera","Anthomyiidae","Anthomyiinae","Delia","","43","43","43","28","0","0","99.05","431.86","0","434","34.48","0"
"AGAW1","05E","INS","BOLD:AAM9347","Arthropoda","Insecta","Diptera","Phoridae","Metopininae","Megaselia","","40","35","35","29","12.5","0","97.22","448.35","0","450","32.78","0"
"AGAW1","05E","INS","BOLD:ADH4889","Arthropoda","Insecta","Diptera","","","","","35","34","34","30","2.86","0","98.99","443.97","0","445.11","35.45","0"
"AGAW1","05E","INS","BOLD:AAF7013","","","","","","","","27","27","27","31","0","0","98.05","363.78","0","365.56","34.22","0"
"AGAW1","05E","INS","BOLD:AAV5575","Arthropoda","Insecta","Diptera","Cecidomyiidae","","","CecidInt30 sp. BOLD:AAV5575","27","24","24","31","11.11","0","99.49","439.93","0","441.48","25.94","0"
"AGAW1","05E","INS","BOLD:AAM7726","Arthropoda","Insecta","Coleoptera","Curculionidae","Curculioninae","Orchestes","Orchestes steppensis, Orchestes alni","22","20","20","33","9.09","0","99.1","444.77","0","445.95","33.35","0"
"AGAW1","05E","INS","BOLD:AAN5356","Arthropoda","Insecta","Diptera","Chironomidae","Orthocladiinae","Smittia","","20","18","18","34","10","0","98.79","434.25","0","435.65","35.2","0"
"AGAW1","05E","INS","BOLD:AAA4222","Arthropoda","Insecta","Lepidoptera","Erebidae","Hypeninae","Hypena","Hypena scabra","19","19","19","35","0","0","99.04","385.11","0","386.26","32.83","0"
"AGAW1","05E","INS","BOLD:ABU5520","Arthropoda","Insecta","Diptera","Sciaridae","","Corynoptera","","14","14","14","36","0","0","98.81","409.29","0","411","35.24","0"
"AGAW1","05E","INS","BOLD:AAA7374","Arthropoda","Insecta","Diptera","Syrphidae","Syrphinae","Sphaerophoria","Sphaerophoria bankowskae, Sphaerophoria infuscata, Sphaerophoria nr. philanthus, Sphaerophoria batava, Sphaerophoria sp., Sphaerophoria rueppellii, Sphaerophoria interrupta-group, Sphaerophoria sp. MA-1, Sphaerophoria cleoae, Sphaerophoria asymmetrica, Sphaerophoria fatarum, Sphaerophoria bengalensis, Sphaerophoria bifurcata, Sphaerophoria taeniata, Sphaerophoria interrupta, Sphaerophoria contigua, Sphaerophoria kaa, Sphaerophoria laurae, Sphaerophoria chongjini, Sphaerophoria boreoalpina, Sphaerophoria sp. BOLDAAA7374, Sphaerophoria macrogaster, Sphaerophoria sulphuripes, Sphaerophoria brevipilosa, Sphaerophoria philanthus, Sphaerophoria pyrrhina, Sphaerophoria abbreviata, Sphaerophoria scripta, Sphaerophoria potentillae, Sphaerophoria longipilosa, Sphaerophoria virgata","14","14","14","36","0","0","98.98","430.36","0","432.29","31.09","0"
"AGAW1","05E","INS","BOLD:ADH9867","Arthropoda","Insecta","Diptera","","","","","11","11","11","38","0","0","98.73","383.55","0","386.45","37.4","0"
"AGAW1","05E","INS","BOLD:AAG6743","Arthropoda","Insecta","Diptera","Sarcophagidae","Sarcophaginae","Sarcophaga","Sarcophaga sp. 1 MG, Sarcophaga subvicina","10","10","10","39","0","0","99.53","435.9","0","438.2","34.8","0"
"AGAW1","05E","INS","BOLD:AAB2644","Arthropoda","Insecta","Diptera","Chironomidae","Orthocladiinae","Orthocladius","Orthocladius lapponicus, Orthocladius dorenus, Orthocladius mallochi","8","8","8","40","0","0","98.48","430.5","0","432.63","32.79","0"
"AGAW1","05E","INS","BOLD:AAF9187","Arthropoda","Insecta","Coleoptera","Curculionidae","Entiminae","Phyllobius","Phyllobius oblongus","8","8","8","40","0","0","99.92","449.38","0","450","39.14","0"
"AGAW1","05E","INS","BOLD:AAB9140","Arthropoda","Insecta","Diptera","Calliphoridae","Chrysomyinae","Phormia","Phormia regina","7","7","7","42","0","0","99.23","429.86","0","431.57","33.05","0"
"AGAW1","05E","INS","BOLD:AAG2462","Arthropoda","Insecta","Diptera","Anthomyiidae","Anthomyiinae","Botanophila","Botanophila fugax","7","7","7","42","0","0","98.18","433.86","0","436.71","31.88","0"
"AGAW1","05E","INS","BOLD:AAA7661","Arthropoda","Insecta","Diptera","Culicidae","Culicinae","Culiseta, Culex","Culiseta melanura, Culex articularis, Culex restuans, Culex acharistus","5","5","5","44","0","0","98.83","412","0","412.6","32.72","0"
"AGAW1","05E","INS","BOLD:AAV5788","Arthropoda","Insecta","Diptera","Cecidomyiidae","Cecidomyiinae","Asphondylia","","4","4","4","45","0","0","98.98","442.75","0","444.25","24.13","0"
"AGAW1","05E","INS","BOLD:AAN5358","Arthropoda","Insecta","Diptera","Chironomidae","Orthocladiinae","Smittia","Smittia stercoraria, Smittia sp. 22ES","4","4","4","45","0","0","99.5","447.75","0","450","37.44","0"
"AGAW1","05E","INS","BOLD:AAU6630","Arthropoda","Insecta","Diptera","Fanniidae","Fanniinae","Fannia","Fannia armata","4","4","4","45","0","0","98.97","434.75","0","437","33.63","0"
"AGAW1","05E","INS","BOLD:ACU7906","Arthropoda","Insecta","Diptera","Chironomidae","Orthocladiinae","","","3","3","3","48","0","0","98.36","413.67","0","414.67","36.78","0"
"AGAW1","05E","INS","BOLD:AAB0373","Arthropoda","Insecta","Neuroptera","Chrysopidae","Chrysopinae","Chrysoperla","Chrysoperla sp. isolate 82, Chrysoperla zastrowi, Chrysoperla plorabunda, Chrysoperla sp. isolate 71, Chrysoperla agilis, Italochrysa italica, Chrysoperla sp. isolate 88, Chrysoperla lucasina, Chrysoperla carnea, Chrysoperla mediterranea, Chrysoperla pallida, Chrysoperla nipponensis, Chrysoperla sillemi","3","2","2","48","33.33","0","99.62","433","0","433","31.08","0"
"AGAW1","05E","INS","BOLD:AAN8770","Arthropoda","Insecta","Diptera","Psychodidae","Psychodinae","Psychoda","Psychoda trinodulosa","3","3","3","48","0","0","98.7","413","0","413.67","33.01","0"
"AGAW1","05E","INS","BOLD:ACV2677","Arthropoda","Insecta","Diptera","Cecidomyiidae","","","","3","3","3","48","0","0","97.55","449.67","0","450","30.07","0"
"AGAW1","05E","INS","BOLD:ABV1243","Arthropoda","Insecta","Diptera","Sarcophagidae","Paramacronychiinae, Miltogramminae","Miltogramminae_incertae_sedis, Brachicoma","Miltogramminae_incertae_sedis sp. 1 MG, Brachicoma sarcophagina","2","2","2","52","0","0","98.22","449","0","450","34.56","0"
"AGAW1","05E","INS","BOLD:AAG1711","Arthropoda","Insecta","Diptera","Muscidae","Phaoniinae","Helina","Helina depuncta","2","2","2","52","0","0","98.97","425","0","426","31.2","0"
"AGAW1","05E","INS","BOLD:ACF4734","Arthropoda","Insecta","Diptera","Syrphidae","Syrphinae","Platycheirus","Platycheirus hyperboreus","2","2","2","52","0","0","97.75","413","0","416","31.26","0"
"AGAW1","05E","INS","BOLD:ABU5546","Arthropoda","Insecta","Diptera","Psychodidae","Psychodinae","Psychoda","","2","2","2","52","0","0","99.33","448.5","0","450","31.78","0"
"AGAW1","05E","INS","BOLD:AAA4121","Arthropoda","Insecta","Diptera","Simuliidae","Simuliinae","Simulium","Simulium vittatum complex, Simulium vittatum, Simulium sp. BOLDAAA4121, Simulium argus, Simulium vittatum s.str.","2","2","2","52","0","0","98.2","445.5","0","450","33.44","0"
"AGAW1","05E","INS","BOLD:AAB4657","Arthropoda","Insecta","Diptera","Chironomidae","Chironominae","Chironomus","Chironomus maturus","2","2","2","52","0","0","99","449","0","450","37.67","0"
"AGAW1","05E","INS","BOLD:AAA3817","Arthropoda","Insecta","Lepidoptera","Geometridae","Larentiinae","Xanthorhoe","Xanthorhoe ferrugata","2","2","2","52","0","0","98.66","448.5","0","450","34","0"
"AGAW1","05E","INS","BOLD:AAB8817","Arthropoda","Insecta","Diptera","Muscidae","Azeliinae","Muscina","Muscina levida","2","2","2","52","0","0","98.89","449","0","450","33.67","0"
"AGAW1","05E","INS","BOLD:AAA2815","Arthropoda","Insecta","Lepidoptera","Noctuidae","Noctuinae","Mythimna","Mythimna yukonensis, Mythimna oxygala","2","2","2","52","0","0","98.74","420.5","0","422","33.96","0"
"AGAW1","05E","INS","BOLD:AAA9251","Arthropoda","Insecta","Lepidoptera","Gelechiidae","Gelechiinae","Scrobipalpa","Scrobipalpa atriplicella, Scrobipalpa nr. atriplicella sp. 1","2","2","2","52","0","0","98.12","426.5","0","430.5","33.8","0"
"AGAW1","05E","INS","BOLD:AAH0408","Arthropoda","Insecta","Coleoptera","Curculionidae","Entiminae","Sitona","Sitona cylindricollis","2","2","2","52","0","0","99.44","447.5","0","450","36","0"
"AGAW1","05E","INS","BOLD:AAG6745","Arthropoda","Insecta","Diptera","Calliphoridae","Calliphorinae","Pollenia","Pollenia pediculata","2","2","2","52","0","0","98.83","384.5","0","389","33.17","0"
"AGAW1","05E","INS","BOLD:AAQ3566","Arthropoda","Insecta","Diptera","Anthomyiidae","Pegomyinae","Pegomya","","2","2","2","52","0","0","98.75","422","0","423","31.77","0"
"AGAW1","05E","INS","BOLD:AAQ0582","Arthropoda","Insecta","Diptera","Anthomyiidae","Anthomyiinae","Lasiomma","","2","2","2","52","0","0","97.09","446.5","0","446.5","31.8","0"
"AGAW1","05E","INS","BOLD:AAH7158","Arthropoda","Insecta","Diptera","Sarcophagidae","Sarcophaginae","Ravinia","Ravinia stimulans","2","2","2","52","0","0","98.78","396.5","0","405","32.74","0"
"AGAW1","05E","INS","BOLD:AAF9041","Arthropoda","Insecta","Diptera","Tipulidae","Tipulinae","Tipula","Tipula oleracea","2","2","2","52","0","0","98.58","382","0","383.5","34.95","0"
"AGAW1","05E","INS","BOLD:AAB2384","Arthropoda","Insecta","Diptera","Syrphidae","Syrphinae","Eupeodes","Eupeodes duseki, Eupeodes volucris, Eupeodes nielseni, Eupeodes americanus, Eupeodes flukei, Eupeodes vockerothi, Eupeodes lucasi, Eupeodes montivagus, Eupeodes rufipunctatus, Eupeodes rojasi, Eupeodes goeldlini, Eupeodes luniger, Eupeodes punctifer, Eupeodes bucculatus, Eupeodes perplexus, Eupeodes abiskoensis, Eupeodes corollae, Eupeodes pomus, Eupeodes lundbecki, Eupeodes nitens, Eupeodes sp. BOLDAAB2384, Eupeodes curtus, Eupeodes latifasciatus","2","2","2","52","0","0","99.75","397.5","0","399","30.33","0"
"AGAW1","05E","INS","BOLD:ADH9310","Arthropoda","Insecta","Diptera","Anthomyiidae","Anthomyiinae","Anthomyia","Anthomyia procellaris, Anthomyia pluvialis","2","2","2","52","0","0","99.89","433.5","0","434.5","32.09","0"
"AGAW1","05E","INS","BOLD:AAA3654","Arthropoda","Insecta","Lepidoptera","Crambidae","Crambinae","Crambus","Crambus bigelovi, Crambus johnsoni, Crambus praefectellus","2","2","2","52","0","0","98.21","448.5","0","450","32.44","0"
"AGAW1","05E","INS","BOLD:AAG1701","Arthropoda","Insecta","Diptera","Muscidae","Coenosiinae","Macrorchis","Macrorchis ausoba","2","2","2","52","0","0","99.11","449.5","0","450","33.22","0"
"AGAW1","05E","INS","BOLD:AAP3004","Arthropoda","Insecta","Diptera","Chironomidae","Chironominae","Chironomus","Chironomus staegeri, Chironomus frommeri","2","2","2","52","0","0","98.22","449.5","0","450","34.78","0"
"AGAW1","05E","INS","BOLD:AAA8451","Arthropoda","Insecta","Lepidoptera","Hepialidae","","Korscheltellus","Korscheltellus lupulina","2","2","2","52","0","0","99.86","401.5","0","403","27.73","0"
"AGAW1","05E","INS","BOLD:ACE8735","Arthropoda","Insecta","Diptera","Cecidomyiidae","","","","2","2","2","52","0","0","99.06","423","0","423.5","25.97","0"
"AGAW1","05E","INS","BOLD:AAA3181","Arthropoda","Insecta","Diptera","Muscidae","Muscinae","Stomoxys","Stomoxys calcitrans","1","1","1","75","0","0","98.36","426","0","430","32.79","0"
"AGAW1","05E","INS","BOLD:AAA2590","Arthropoda","Insecta","Lepidoptera","Noctuidae","Noctuinae","Xestia","Xestia baja, Xestia smithii","1","1","1","75","0","0","99.05","419","0","419","31.98","0"
"AGAW1","05E","INS","BOLD:AAH0349","Arthropoda","Insecta","Coleoptera","Chrysomelidae","Cassidinae","Charidotella","Charidotella sexpunctata, Charidotella purpurata","1","1","1","75","0","0","97.31","446","0","450","32.89","0"
"AGAW1","05E","INS","BOLD:AAA3836","Arthropoda","Insecta","Lepidoptera","Noctuidae","Plusiinae","Autographa","Autographa precationis","1","1","1","75","0","0","99.78","448","0","450","31.78","0"
"AGAW1","05E","INS","BOLD:AAD1945","Arthropoda","Insecta","Diptera","Agromyzidae","Phytomyzinae","Cerodontha","Cerodontha dorsalis","1","1","1","75","0","0","99.78","449","0","450","32.22","0"
"AGAW1","05E","INS","BOLD:AAC9132","Arthropoda","Insecta","Diptera","Culicidae","Culicinae","Culiseta","Culiseta inornata","1","1","1","75","0","0","99.19","370","0","371","32.88","0"
"AGAW1","05E","INS","BOLD:AAN5233","Arthropoda","Insecta","Diptera","Cecidomyiidae","","","","1","1","1","75","0","0","99.33","451","0","450","26.89","0"
"AGAW1","05E","INS","BOLD:AAY3269","Arthropoda","Insecta","Diptera","Chironomidae","Diamesinae","Potthastia","Potthastia gaedii","1","1","1","75","0","0","98.25","399","0","401","34.16","0"
"AGAW1","05E","INS","BOLD:AAF9768","Arthropoda","Insecta","Diptera","Hybotidae","Tachydromiinae","Platypalpus","Platypalpus holosericus, Platypalpus candicans, Platypalpus sp. BOLD:AAF9768, Platypalpus sp. 22","1","1","1","75","0","0","99.33","450","0","450","32.22","0"
"AGAW1","05E","INS","BOLD:AAK7242","Arthropoda","Insecta","Coleoptera","Melandryidae","Melandryinae","Melandrya","Melandrya striata","1","1","1","75","0","0","98.99","395","0","450","33.11","0"
"AGAW1","05E","INS","BOLD:AAG0991","Arthropoda","Insecta","Diptera","Chironomidae","Orthocladiinae","Orthocladius","","1","1","1","75","0","0","98.37","369","0","371","32.88","0"
"AGAW1","05E","INS","BOLD:AAG8493","Arthropoda","Insecta","Diptera","Drosophilidae","Drosophilinae","Scaptomyza","Scaptomyza pallida","1","1","1","75","0","0","98.66","448","0","450","34","0"
"AGAW1","05E","INS","BOLD:AAD0583","Arthropoda","Insecta","Hemiptera","Lygaeidae","Lygaeinae","Lygaeus","Lygaeus reclivatus, Lygaeus kalmii","1","1","1","75","0","0","98.67","450","0","450","33.33","0"
"AGAW1","05E","INS","BOLD:AAG6753","Arthropoda","Insecta","Diptera","Lauxaniidae","Lauxaniinae","Camptoprosopella","","1","1","1","75","0","0","97.33","450","0","450","36.44","0"
"AGAW1","05E","INS","BOLD:AAC1312","Arthropoda","Insecta","Diptera","Syrphidae","Syrphinae","Toxomerus","Toxomerus geminatus, Toxomerus 75-5, Toxomerus rombicus, Toxomerus sp. 75-5","1","1","1","75","0","0","97.51","442","0","450","30","0"
"AGAW1","05E","INS","BOLD:AAB1098","Arthropoda","Insecta","Diptera","Culicidae","Culicinae","Ochlerotatus, Aedes","Aedes excrucians, Aedes cantans, Aedes stimulans, Aedes euedes, Aedes riparius, Aedes annulipes, Ochlerotatus cantans, Aedes fitchii, Aedes sp., Ochlerotatus annulipes, Ochlerotatus excrucians","1","1","1","75","0","0","98.56","417","0","419","34.84","0"
"AGAW1","05E","INS","BOLD:AAV1055","Arthropoda","Insecta","Diptera","Stratiomyidae","Beridinae","Allognosta","Allognosta fuscitarsis","1","1","1","75","0","0","99.2","375","0","376","35.9","0"
"AGAW1","05E","INS","BOLD:AAD0853","Arthropoda","Insecta","Diptera","Scathophagidae","Scathophaginae","Scathophaga","Scathophaga lutaria, Scathophaga hiemalis, Scathophaga furcata, Scathophaga stercoraria, Scathophaga inquinata, Scathophaga suilla","1","1","1","75","0","0","97.08","411","0","414","31.16","0"
"AGAW1","05E","INS","BOLD:AAG2128","Arthropoda","Insecta","Diptera","Tachinidae","Exoristinae","Cryptomeigenia","Cryptomeigenia sp. 3, Cryptomeigenia triangularis, Cryptomeigenia muscoides, Cryptomeigenia CR 82, Cryptomeigenia sp. GER5, Cryptomeigenia theutis, Cryptomeigenia sp. GER3","1","1","1","75","0","0","97.09","447","0","450","32","0"
"AGAW1","05E","INS","BOLD:ACF5939","Arthropoda","Insecta","Diptera","Tachinidae","Exoristinae","Winthemia","Winthemia sinuata, Winthemia illinoensis","1","1","1","75","0","0","100","450","0","450","29.78","0"
"AGAW1","05E","INS","BOLD:AAM6234","Arthropoda","Insecta","Diptera","Chironomidae","Tanypodinae","","","1","1","1","75","0","0","99.76","416","0","419","35.08","0"
"AGAW1","05E","INS","BOLD:ACK7296","Arthropoda","Insecta","Diptera","Cecidomyiidae","","","","1","1","1","75","0","0","97.29","442","0","443","25.06","0"
"AGAW1","05E","INS","BOLD:ACF7232","Arthropoda","Insecta","Diptera","Chironomidae","Orthocladiinae","Parakiefferiella","","1","1","1","75","0","0","98.43","383","0","398","37.19","0"
"AGAW1","05E","INS","BOLD:AAD8086","Arthropoda","Insecta","Lepidoptera","Elachistidae","Elachistinae","Elachista","Elachista griseicornis, Elachista acenteta","1","1","1","75","0","0","98.88","447","0","450","30.89","0"
"AGAW1","05E","INS","BOLD:AAE4298","Arthropoda","Insecta","Diptera","Chironomidae","Orthocladiinae","Cricotopus","Cricotopus tremulus","1","1","1","75","0","0","97.37","380","0","382","32.46","0"
"AGAW1","05E","INS","BOLD:AAG0161","Arthropoda","Insecta","Diptera","Milichiidae","Madizinae","Leptometopa","Leptometopa latipes","1","1","1","75","0","0","99.74","391","0","394","32.23","0"
"AGAW1","05E","INS","BOLD:ABW7897","Arthropoda","Insecta","Diptera","Cecidomyiidae","","","","1","1","1","75","0","0","98.21","448","0","450","30.44","0"
"AGAW1","05E","INS","BOLD:AAG4653","Arthropoda","Insecta","Diptera","Syrphidae","Syrphinae","Heringia","Heringia latitarsis, Heringia pubescens, Heringia jakutorum, Heringia rita, Heringia fulvimanus, Heringia sp. BOLDAAG4653, Heringia nr. pubescens, Heringia coxalis, Heringia larusi, Heringia vitripennis","1","1","1","75","0","0","99.78","450","0","450","31.56","0"
"AGAW1","05E","INS","BOLD:AAD0484","Arthropoda","Insecta","Diptera","Chironomidae","","","","1","1","1","75","0","0","97.77","448","0","450","34","0"
"AGAW1","05E","NAR","no match","no match","no match","no match","no match","no match","no match","no match","no match","no match","no match","no match","no match","no match","no match","no match","no match","no match","no match","no match"
"AGAW1","05E","NIN","BOLD:AAN6394","Arthropoda","Arachnida","Araneae","Anyphaenidae",NA,"Hibana","Hibana gracilis","2315","2241","2241","1","3.2","0","98.85","425.7","0","428.1","30.9","0"
"AGAW1","05E","NIN","BOLD:AAN6371","Arthropoda","Arachnida","Araneae","Theridiidae",NA,"Wamba","Wamba crispulus","229","179","179","2","21.83","0","99.22","439.35","0","445.38","30.14","0"
"AGAW1","05E","NIN","BOLD:AAA4125","Arthropoda","Arachnida","Araneae","Araneidae",NA,"Araneus","Araneus diadematus","126","118","118","3","6.35","0","99.32","431.44","0","432.6","34.13","0"
"AGAW1","05E","NIN","BOLD:AAI4346","Arthropoda","Arachnida","Opiliones","Phalangiidae",NA,"Paroligolophus, Phalangium","Paroligolophus agrestis, Phalangium opilio","64","64","64","4","0","0","98.9","422.06","0","430.95","35.65","0"
"AGAW1","05E","NIN","BOLD:ABW2639","Arthropoda","Arachnida","Trombidiformes","Anystidae",NA,"","","1","1","1","5","0","0","99.33","448","0","450","31.78","0"
"AGAW1","05E","NIN","BOLD:AAB7915","Arthropoda","Collembola","Symphypleona","Bourletiellidae",NA,"Deuterosminthurus","Deuterosminthurus sp1","1","1","1","5","0","0","99.78","449","0","450","35.11","0"
"AGAW1","05N","BAC","TAX:375853","Proteobacteria","Alphaproteobacteria","Rickettsiales","Rickettsiaceae",NA,"Wolbachia","Wolbachia sp. Hymenoptera_ASGLE646-10","230","120","120","1","47.83","0","97.67","449.53","0","449.89","40.9","0"
"AGAW1","05N","BAC","TAX:392873","Proteobacteria","Alphaproteobacteria","Rickettsiales","Rickettsiaceae",NA,"Wolbachia","Wolbachia endosymbiont of Scrapter algoensis","47","46","46","2","2.13","0","97.78","412.68","0","435.66","41.04","0"
"AGAW1","05N","BAC","TAX:91393","Proteobacteria","Alphaproteobacteria","Rickettsiales","Rickettsiaceae",NA,"Wolbachia","","24","24","24","3","0","0","99.36","386.5","0","417.71","40.32","0"
"AGAW1","05N","BAC","TAX:376688","Proteobacteria","Alphaproteobacteria","Rickettsiales","Rickettsiaceae",NA,"Wolbachia","Wolbachia sp. elachBioLep01 BioLep373","15","15","15","4","0","0","97.83","446.53","0","448.93","40.82","0"
"AGAW1","05N","BAC","TAX:375807","Proteobacteria","Alphaproteobacteria","Rickettsiales","Rickettsiaceae",NA,"Wolbachia","Wolbachia sp. Cerapachys lividus_ASANV557-09","13","13","13","5","0","0","99.48","399.54","0","442.54","40.81","0"
"AGAW1","05N","BAC","TAX:375772","Proteobacteria","Alphaproteobacteria","Rickettsiales","Rickettsiaceae",NA,"Wolbachia","Wolbachia sp. Leptogenys JCR25_ASANE483-10","12","12","12","6","0","0","97.44","446.67","0","450","40.81","0"
"AGAW1","05N","BAC","TAX:375782","Proteobacteria","Alphaproteobacteria","Rickettsiales","Rickettsiaceae",NA,"Wolbachia","Wolbachia sp. Cerapachys L_MG11_ASANH426-10","11","10","10","7","9.09","0","97.79","448.91","0","450","40.99","0"
"AGAW1","05N","BAC","TAX:375845","Proteobacteria","Alphaproteobacteria","Rickettsiales","Rickettsiaceae",NA,"Wolbachia","Wolbachia sp. Hymenoptera_ASGLE1440-10","6","6","6","8","0","0","98.19","431.67","0","435","41","0"
"AGAW1","05N","BAC","TAX:265077","Proteobacteria","Alphaproteobacteria","Rickettsiales","Rickettsiaceae",NA,"Wolbachia","Wolbachia sp. FormicidaeASPNA479-09","6","5","5","8","16.67","0","98.06","435.83","0","437.33","41.16","0"
"AGAW1","05N","BAC","TAX:392914","Proteobacteria","Alphaproteobacteria","Rickettsiales","Rickettsiaceae",NA,"Wolbachia","Wolbachia endosymbiont of chalJanzen01 Janzen01","4","4","4","10","0","0","97.54","446.75","0","450","40.89","0"
"AGAW1","05N","BAC","TAX:91391","Proteobacteria","Alphaproteobacteria","Rickettsiales","",NA,"","","4","4","4","10","0","0","98.32","406.75","0","438","40.26","0"
"AGAW1","05N","BAC","TAX:392874","Proteobacteria","Alphaproteobacteria","Rickettsiales","Rickettsiaceae",NA,"Wolbachia","Wolbachia endosymbiont of Hymenoptera","3","3","3","12","0","0","97.36","429","0","431","40.74","0"
"AGAW1","05N","BAC","TAX:375810","Proteobacteria","Alphaproteobacteria","Rickettsiales","Rickettsiaceae",NA,"Wolbachia","Wolbachia sp. Cerapachys L_MG10_ASANV571-09","3","3","3","12","0","0","97.26","450","0","450","40.89","0"
"AGAW1","05N","BAC","TAX:392881","Proteobacteria","Alphaproteobacteria","Rickettsiales","Rickettsiaceae",NA,"Wolbachia","Wolbachia endosymbiont of Blastophaga sp. 1-ERE","3","3","3","12","0","0","97.68","431.67","0","438.67","40.89","0"
"AGAW1","05N","BAC","TAX:392917","Proteobacteria","Alphaproteobacteria","Rickettsiales","Rickettsiaceae",NA,"Wolbachia","Wolbachia endosymbiont of Glyptapanteles Whitfield131","2","2","2","15","0","0","97.52","444","0","444.5","41.29","0"
"AGAW1","05N","BAC","TAX:453459","Proteobacteria","Alphaproteobacteria","Rickettsiales","Rickettsiaceae",NA,"Wolbachia","Wolbachia endosymbiont of Ichneumonidae","2","2","2","15","0","0","98","451","0","450","40.33","0"
"AGAW1","05N","BAC","TAX:375847","Proteobacteria","Alphaproteobacteria","Rickettsiales","Rickettsiaceae",NA,"Wolbachia","Wolbachia sp. Hymenoptera_ASGLE1705-10","1","1","1","17","0","0","98.33","420","0","424","39.86","0"
"AGAW1","05N","BAC","TAX:703648","Proteobacteria","Alphaproteobacteria","Rickettsiales","Rickettsiaceae",NA,"Wolbachia","Wolbachia endosymbiont","1","1","1","17","0","0","97.72","351","0","353","39.09","0"
"AGAW1","05N","BAC","TAX:376694","Proteobacteria","Alphaproteobacteria","Rickettsiales","Rickettsiaceae",NA,"Wolbachia","Wolbachia sp. Anacrusis nephrodes","1","1","1","17","0","0","98.43","446","0","450","40.89","0"
"AGAW1","05N","BAC","TAX:175768","Proteobacteria","Alphaproteobacteria","Rickettsiales","Rickettsiaceae",NA,"Wolbachia","Wolbachia pipientis","1","1","1","17","0","0","97.77","448","0","450","39.33","0"
"AGAW1","05N","BAC","TAX:392915","Proteobacteria","Alphaproteobacteria","Rickettsiales","Rickettsiaceae",NA,"Wolbachia","Wolbachia endosymbiont of Cubusvalidus","1","1","1","17","0","0","97.34","414","0","431","40.14","0"
"AGAW1","05N","BAC","TAX:375768","Proteobacteria","Alphaproteobacteria","Rickettsiales","Rickettsiaceae",NA,"Wolbachia","Wolbachia sp. Cerapachys L_MG09_ASAMX569-06","1","1","1","17","0","0","97.58","413","0","448","41.29","0"
"AGAW1","05N","CHO","no match","no match","no match","no match","no match","no match","no match","no match","no match","no match","no match","no match","no match","no match","no match","no match","no match","no match","no match","no match"
"AGAW1","05N","INS","BOLD:AAD0853","Arthropoda","Insecta","Diptera","Scathophagidae","Scathophaginae","Scathophaga","Scathophaga lutaria, Scathophaga hiemalis, Scathophaga furcata, Scathophaga stercoraria, Scathophaga inquinata, Scathophaga suilla","18855","14081","14081","1","25.32","0","99.45","436.38","0","438.02","31.91","0"
"AGAW1","05N","INS","BOLD:AAI4303","Arthropoda","Insecta","Diptera","Chironomidae","Chironominae","Chironomus","Chironomus melanescens","16913","14075","14075","2","16.78","0","98.49","431.24","0","435.42","34.5","0"
"AGAW1","05N","INS","BOLD:AAA7661","Arthropoda","Insecta","Diptera","Culicidae","Culicinae","Culiseta, Culex","Culiseta melanura, Culex articularis, Culex restuans, Culex acharistus","13032","11058","11058","3","15.15","0","99.24","432.82","0","434.65","32.73","0"
"AGAW1","05N","INS","BOLD:AAO3939","Arthropoda","Insecta","Diptera","Limoniidae","Limoniinae","Dicranomyia","","5423","4497","4497","4","17.08","0","98.85","439.42","0","441.44","36.83","0"
"AGAW1","05N","INS","BOLD:AAF9014","Arthropoda","Insecta","Diptera","Limoniidae","Chioneinae","Symplecta","Symplecta cana, Symplecta sp. BOLDAAF9014","3379","3052","3052","5","9.68","0","98.53","425.63","0","431.4","33.93","0"
"AGAW1","05N","INS","BOLD:AAB1098","Arthropoda","Insecta","Diptera","Culicidae","Culicinae","Ochlerotatus, Aedes","Aedes excrucians, Aedes cantans, Aedes stimulans, Aedes euedes, Aedes riparius, Aedes annulipes, Ochlerotatus cantans, Aedes fitchii, Aedes sp., Ochlerotatus annulipes, Ochlerotatus excrucians","2301","2091","2091","6","9.13","0","98.98","426.96","0","428.42","35.19","0"
"AGAW1","05N","INS","BOLD:AAA4121","Arthropoda","Insecta","Diptera","Simuliidae","Simuliinae","Simulium","Simulium vittatum complex, Simulium vittatum, Simulium sp. BOLDAAA4121, Simulium argus, Simulium vittatum s.str.","2089","1979","1979","7","5.27","0","98.3","409.67","0","412.29","33.99","0"
"AGAW1","05N","INS","BOLD:AAN5342","Arthropoda","Insecta","Diptera","Chironomidae","Orthocladiinae","Heleniella","","1050","1017","1017","8","3.14","0","99.08","401.56","0","412.3","34.88","0"
"AGAW1","05N","INS","BOLD:AAM7726","Arthropoda","Insecta","Coleoptera","Curculionidae","Curculioninae","Orchestes","Orchestes steppensis, Orchestes alni","911","659","659","9","27.66","0","99.44","442.77","0","444.11","32.97","0"
"AGAW1","05N","INS","BOLD:AAO1339","Arthropoda","Insecta","Coleoptera","Anthribidae","Anthribinae","Anthribus","Anthribus nebulosus","790","719","719","10","8.99","0","99.06","414.68","0","418.08","36.01","0"
"AGAW1","05N","INS","BOLD:ACW5117","Arthropoda","Insecta","Diptera","Chironomidae","Orthocladiinae","Smittia","Smittia leucopogon","777","720","720","11","7.34","0","99.28","424.61","0","425.91","31.48","0"
"AGAW1","05N","INS","BOLD:ABU5525","Arthropoda","Insecta","Diptera","Chironomidae","Orthocladiinae","Limnophyes","Limnophyes sp. 14ES","544","533","533","12","2.02","0","98.91","408.53","0","411.62","37.37","0"
"AGAW1","05N","INS","BOLD:AAG5423","Arthropoda","Insecta","Diptera","Chironomidae","Chironominae","Dicrotendipes","","524","515","515","13","1.72","0","98.46","400.36","0","404.37","37.31","0"
"AGAW1","05N","INS","BOLD:ACK3431","Arthropoda","Insecta","Diptera","Cecidomyiidae","","","","428","409","409","14","4.44","0","99.06","420.84","0","422.21","32.78","0"
"AGAW1","05N","INS","BOLD:ACP4736","Arthropoda","Insecta","Diptera","Chironomidae","Orthocladiinae","Smittia","Smittia sp. 8ES","403","388","388","15","3.72","0","99.13","417.13","0","418.85","32.98","0"
"AGAW1","05N","INS","BOLD:AAH3664","Arthropoda","Insecta","Diptera","Cecidomyiidae","","","CecidInt3 sp. BOLD:AAH3664","397","383","383","16","3.53","0","98.81","433.21","0","435.79","27.21","0"
"AGAW1","05N","INS","BOLD:ACX5521","Arthropoda","Insecta","Diptera","Chironomidae","Orthocladiinae","","","347","327","327","17","5.76","0","99.17","421.61","0","423.12","34.66","0"
"AGAW1","05N","INS","BOLD:AAP2998","Arthropoda","Insecta","Diptera","Chironomidae","Tanypodinae","","","306","306","306","18","0","0","98.79","340.3","0","385.51","36.7","0"
"AGAW1","05N","INS","BOLD:AAM7064","Arthropoda","Insecta","Diptera","Chironomidae","Orthocladiinae","Smittia","Smittia sp. 14ES","267","241","241","19","9.74","0","98.83","432.1","0","436.43","31.9","0"
"AGAW1","05N","INS","BOLD:ACB3186","Arthropoda","Insecta","Diptera","Chironomidae","Orthocladiinae","","","114","110","110","20","3.51","0","99.49","385.2","0","423.07","37.05","0"
"AGAW1","05N","INS","BOLD:AAG2868","Arthropoda","Insecta","Hemiptera","Cicadellidae","Typhlocybinae","Empoasca","Empoasca sp., Empoasca fabae","57","46","46","21","19.3","0","99.49","441.6","0","448.63","32.35","0"
"AGAW1","05N","INS","BOLD:ADH9867","Arthropoda","Insecta","Diptera","","","","","55","54","54","22","1.82","0","98.86","425.35","0","428.04","37.44","0"
"AGAW1","05N","INS","BOLD:AAN5304","Arthropoda","Insecta","Diptera","Cecidomyiidae","","","","53","49","49","23","7.55","0","99.39","442.15","0","443.3","30.47","0"
"AGAW1","05N","INS","BOLD:ABV1474","Arthropoda","Insecta","Diptera","Cecidomyiidae","","","","51","49","49","24","3.92","0","99.13","442.27","0","443.53","30.99","0"
"AGAW1","05N","INS","BOLD:AAN5356","Arthropoda","Insecta","Diptera","Chironomidae","Orthocladiinae","Smittia","","48","48","48","25","0","0","99.14","427.25","0","430.83","34.48","0"
"AGAW1","05N","INS","BOLD:AAN6154","Arthropoda","Insecta","Coleoptera","Latridiidae","Corticariinae","Melanophthalma","Melanophthalma inermis","45","41","41","26","8.89","0","98.75","439.76","0","442.84","31.34","0"
"AGAW1","05N","INS","BOLD:AAH3662","Arthropoda","Insecta","Diptera","Cecidomyiidae","","","CecidInt3 sp. BOLD:AAH3662","32","32","32","27","0","0","98.89","436.38","0","439.41","26.23","0"
"AGAW1","05N","INS","BOLD:ABA1215","Arthropoda","Insecta","Diptera","Sciaridae","","Lycoriella","Lycoriella sativae","23","22","22","28","4.35","0","99.51","439.52","0","440.22","35.88","0"
"AGAW1","05N","INS","BOLD:AAA3453","Arthropoda","Insecta","Diptera","Anthomyiidae","Anthomyiinae","Delia","","23","23","23","28","0","0","98.85","435.39","0","438.22","34.81","0"
"AGAW1","05N","INS","BOLD:ABV1375","Arthropoda","Insecta","Diptera","Cecidomyiidae","","","CecidInt3 sp. BOLD:ABV1375","22","21","21","30","4.55","0","99.2","436.95","0","437.77","26.59","0"
"AGAW1","05N","INS","BOLD:AAV6213","Arthropoda","Insecta","Diptera","Chironomidae","Chironominae","Paratanytarsus","","20","20","20","31","0","0","99","414.2","0","429.4","34.49","0"
"AGAW1","05N","INS","BOLD:AAG2511","Arthropoda","Insecta","Diptera","Anthomyiidae","Anthomyiinae","Delia","Delia platura","18","18","18","32","0","0","99.12","446.22","0","448.78","33.5","0"
"AGAW1","05N","INS","BOLD:AAB9140","Arthropoda","Insecta","Diptera","Calliphoridae","Chrysomyinae","Phormia","Phormia regina","14","14","14","33","0","0","98.65","431.21","0","432.86","33.03","0"
"AGAW1","05N","INS","BOLD:AAA4222","Arthropoda","Insecta","Lepidoptera","Erebidae","Hypeninae","Hypena","Hypena scabra","13","13","13","34","0","0","98.84","395.31","0","396.54","32.61","0"
"AGAW1","05N","INS","BOLD:AAM7855","Arthropoda","Insecta","Thysanoptera","Thripidae","Thripinae","Frankliniella","Frankliniella tritici","12","12","12","35","0","0","99.44","430.25","0","433.08","31.27","0"
"AGAW1","05N","INS","BOLD:AAG2462","Arthropoda","Insecta","Diptera","Anthomyiidae","Anthomyiinae","Botanophila","Botanophila fugax","10","10","10","36","0","0","98.22","435.3","0","437.5","31.78","0"
"AGAW1","05N","INS","BOLD:AAF9187","Arthropoda","Insecta","Coleoptera","Curculionidae","Entiminae","Phyllobius","Phyllobius oblongus","9","9","9","37","0","0","98.96","429.11","0","429.67","39.17","0"
"AGAW1","05N","INS","BOLD:AAG6743","Arthropoda","Insecta","Diptera","Sarcophagidae","Sarcophaginae","Sarcophaga","Sarcophaga sp. 1 MG, Sarcophaga subvicina","7","7","7","38","0","0","99.06","421.57","0","423","34.79","0"
"AGAW1","05N","INS","BOLD:AAY6412","Arthropoda","Insecta","Diptera","Sciaridae","","Bradysia","Bradysia urticae","7","7","7","38","0","0","99.38","443.43","0","445.43","32.3","0"
"AGAW1","05N","INS","BOLD:AAA7374","Arthropoda","Insecta","Diptera","Syrphidae","Syrphinae","Sphaerophoria","Sphaerophoria bankowskae, Sphaerophoria infuscata, Sphaerophoria nr. philanthus, Sphaerophoria batava, Sphaerophoria sp., Sphaerophoria rueppellii, Sphaerophoria interrupta-group, Sphaerophoria sp. MA-1, Sphaerophoria cleoae, Sphaerophoria asymmetrica, Sphaerophoria fatarum, Sphaerophoria bengalensis, Sphaerophoria bifurcata, Sphaerophoria taeniata, Sphaerophoria interrupta, Sphaerophoria contigua, Sphaerophoria kaa, Sphaerophoria laurae, Sphaerophoria chongjini, Sphaerophoria boreoalpina, Sphaerophoria sp. BOLDAAA7374, Sphaerophoria macrogaster, Sphaerophoria sulphuripes, Sphaerophoria brevipilosa, Sphaerophoria philanthus, Sphaerophoria pyrrhina, Sphaerophoria abbreviata, Sphaerophoria scripta, Sphaerophoria potentillae, Sphaerophoria longipilosa, Sphaerophoria virgata","7","7","7","38","0","0","99","412.57","0","414.43","31.09","0"
"AGAW1","05N","INS","BOLD:AAA9251","Arthropoda","Insecta","Lepidoptera","Gelechiidae","Gelechiinae","Scrobipalpa","Scrobipalpa atriplicella, Scrobipalpa nr. atriplicella sp. 1","5","5","5","41","0","0","99.25","390.4","0","393.8","34.08","0"
"AGAW1","05N","INS","BOLD:AAB2644","Arthropoda","Insecta","Diptera","Chironomidae","Orthocladiinae","Orthocladius","Orthocladius lapponicus, Orthocladius dorenus, Orthocladius mallochi","2","2","2","42","0","0","99.33","447.5","0","450","32.22","0"
"AGAW1","05N","INS","BOLD:AAA8933","Arthropoda","Insecta","Coleoptera","Coccinellidae","Coccinellinae","Coccinella","Coccinella septempunctata","2","2","2","42","0","0","98.77","448","0","450","33.11","0"
"AGAW1","05N","INS","BOLD:AAC9132","Arthropoda","Insecta","Diptera","Culicidae","Culicinae","Culiseta","Culiseta inornata","2","2","2","42","0","0","98.81","392","0","396","32.35","0"
"AGAW1","05N","INS","BOLD:AAH3914","Arthropoda","Insecta","Diptera","Sciaridae","","Sciara","Sciara hamatilis, Sciara humeralis","2","2","2","42","0","0","98.78","449.5","0","450","30.56","0"
"AGAW1","05N","INS","BOLD:AAD0484","Arthropoda","Insecta","Diptera","Chironomidae","","","","2","2","2","42","0","0","98.22","449","0","450","35.11","0"
"AGAW1","05N","INS","BOLD:ACF7004","Arthropoda","Insecta","Thysanoptera","Thripidae","Thripinae","Limothrips","Limothrips denticornis","2","2","2","42","0","0","97.74","376.5","0","450","35.89","0"
"AGAW1","05N","INS","BOLD:AAN8770","Arthropoda","Insecta","Diptera","Psychodidae","Psychodinae","Psychoda","Psychoda trinodulosa","2","2","2","42","0","0","99.89","449.5","0","450","34.11","0"
"AGAW1","05N","INS","BOLD:AAC9062","Arthropoda","Insecta","Diptera","Culicidae","Culicinae","Ochlerotatus, Aedes","Aedes fitchii, Ochlerotatus fitchii, Aedes grossbecki, Aedes ventrovittis","2","2","2","42","0","0","99","450","0","450","33.11","0"
"AGAW1","05N","INS","BOLD:AAU6630","Arthropoda","Insecta","Diptera","Fanniidae","Fanniinae","Fannia","Fannia armata","2","2","2","42","0","0","98.66","449","0","450","33.67","0"
"AGAW1","05N","INS","BOLD:AAG1701","Arthropoda","Insecta","Diptera","Muscidae","Coenosiinae","Macrorchis","Macrorchis ausoba","2","2","2","42","0","0","98.78","449.5","0","450","33.33","0"
"AGAW1","05N","INS","BOLD:AAA3181","Arthropoda","Insecta","Diptera","Muscidae","Muscinae","Stomoxys","Stomoxys calcitrans","1","1","1","52","0","0","99.11","448","0","450","33.56","0"
"AGAW1","05N","INS","BOLD:AAG9055","Arthropoda","Insecta","Hymenoptera","Vespidae","Vespinae","Vespula","Vespula germanica","1","1","1","52","0","0","100","449","0","450","30.67","0"
"AGAW1","05N","INS","BOLD:AAG1711","Arthropoda","Insecta","Diptera","Muscidae","Phaoniinae","Helina","Helina depuncta","1","1","1","52","0","0","100","448","0","450","31.78","0"
"AGAW1","05N","INS","BOLD:AAG1717","Arthropoda","Insecta","Diptera","Muscidae","Phaoniinae","Helina","Helina troene","1","1","1","52","0","0","99.33","450","0","450","31.11","0"
"AGAW1","05N","INS","BOLD:AAA2590","Arthropoda","Insecta","Lepidoptera","Noctuidae","Noctuinae","Xestia","Xestia baja, Xestia smithii","1","1","1","52","0","0","99.78","450","0","450","33.11","0"
"AGAW1","05N","INS","BOLD:AAA3933","Arthropoda","Insecta","Lepidoptera","Erebidae","Herminiinae","Palthis","Palthis angulalis","1","1","1","52","0","0","97.55","449","0","450","32.22","0"
"AGAW1","05N","INS","BOLD:AAH0349","Arthropoda","Insecta","Coleoptera","Chrysomelidae","Cassidinae","Charidotella","Charidotella sexpunctata, Charidotella purpurata","1","1","1","52","0","0","99.51","405","0","408","31.13","0"
"AGAW1","05N","INS","BOLD:ACT0257","","","","","","","","1","1","1","52","0","0","99.78","450","0","450","37.78","0"
"AGAW1","05N","INS","BOLD:AAA3836","Arthropoda","Insecta","Lepidoptera","Noctuidae","Plusiinae","Autographa","Autographa precationis","1","1","1","52","0","0","99.56","450","0","450","32","0"
"AGAW1","05N","INS","BOLD:AAG7276","Arthropoda","Insecta","Diptera","Sphaeroceridae","Limosininae","Leptocera","Leptocera erythrocera","1","1","1","52","0","0","98.89","449","0","450","32.44","0"
"AGAW1","05N","INS","BOLD:AAG5507","Arthropoda","Insecta","Diptera","Chironomidae","Chironominae","Chironomus","Chironomus atrella","1","1","1","52","0","0","99.02","408","0","414","34.78","0"
"AGAW1","05N","INS","BOLD:AAN5249","Arthropoda","Insecta","Diptera","Cecidomyiidae","","","","1","1","1","52","0","0","99.52","418","0","418","33.25","0"
"AGAW1","05N","INS","BOLD:AAH4135","Arthropoda","Insecta","Diptera","Chloropidae","Chloropinae","Thaumatomyia","Thaumatomyia glabra, Thaumatomyia grata, Thaumatomyia sp. 1 TAW","1","1","1","52","0","0","99.73","367","0","367","31.61","0"
"AGAW1","05N","INS","BOLD:AAD8749","Arthropoda","Insecta","Lepidoptera","Gelechiidae","Pexicopiinae","Platyedra","Platyedra subcinerea","1","1","1","52","0","0","97.69","389","0","391","31.46","0"
"AGAW1","05N","INS","BOLD:AAG5457","Arthropoda","Insecta","Diptera","Chironomidae","Chironominae","","","1","1","1","52","0","0","99.76","413","0","416","32.69","0"
"AGAW1","05N","INS","BOLD:AAA2815","Arthropoda","Insecta","Lepidoptera","Noctuidae","Noctuinae","Mythimna","Mythimna yukonensis, Mythimna oxygala","1","1","1","52","0","0","98.22","450","0","450","33.33","0"
"AGAW1","05N","INS","BOLD:AAF9771","Arthropoda","Insecta","Diptera","Hybotidae","Tachydromiinae","Platypalpus","Platypalpus longiseta, Platypalpus trivialis, Platypalpus sp. 5, Platypalpus sp. 17, Platypalpus pallidiventris","1","1","1","52","0","0","98.89","450","0","450","35.56","0"
"AGAW1","05N","INS","BOLD:AAZ5973","Arthropoda","Insecta","Diptera","Sepsidae","Sepsinae","Themira","Themira annulipes","1","1","1","52","0","0","98.89","450","0","450","36.44","0"
"AGAW1","05N","INS","BOLD:AAV5853","Arthropoda","Insecta","Diptera","Chironomidae","Tanypodinae","","","1","1","1","52","0","0","98.92","372","0","375","36","0"
"AGAW1","05N","INS","BOLD:AAB8817","Arthropoda","Insecta","Diptera","Muscidae","Azeliinae","Muscina","Muscina levida","1","1","1","52","0","0","98.44","450","0","450","32.44","0"
"AGAW1","05N","INS","BOLD:AAG0995","Arthropoda","Insecta","Diptera","Chironomidae","Orthocladiinae","Parakiefferiella","","1","1","1","52","0","0","98.22","449","0","450","39.11","0"
"AGAW1","05N","INS","BOLD:AAG0991","Arthropoda","Insecta","Diptera","Chironomidae","Orthocladiinae","Orthocladius","","1","1","1","52","0","0","99.73","366","0","373","32.17","0"
"AGAW1","05N","INS","BOLD:AAE0707","Arthropoda","Insecta","Diptera","Chironomidae","Chironominae","Microtendipes","Microtendipes pedellus","1","1","1","52","0","0","97.33","374","0","376","31.38","0"
"AGAW1","05N","INS","BOLD:AAA2482","Arthropoda","Insecta","Lepidoptera","Noctuidae","Noctuinae","Mythimna","Mythimna unipuncta","1","1","1","52","0","0","97.69","389","0","391","30.95","0"
"AGAW1","05N","INS","BOLD:AAG1743","Arthropoda","Insecta","Diptera","Muscidae","Phaoniinae","Phaonia","Phaonia fuscana","1","1","1","52","0","0","97.51","401","0","403","32.26","0"
"AGAW1","05N","INS","BOLD:AAD0583","Arthropoda","Insecta","Hemiptera","Lygaeidae","Lygaeinae","Lygaeus","Lygaeus reclivatus, Lygaeus kalmii","1","1","1","52","0","0","100","449","0","450","32.67","0"
"AGAW1","05N","INS","BOLD:AAN5358","Arthropoda","Insecta","Diptera","Chironomidae","Orthocladiinae","Smittia","Smittia stercoraria, Smittia sp. 22ES","1","1","1","52","0","0","97.78","450","0","450","37.56","0"
"AGAW1","05N","INS","BOLD:AAN5355","Arthropoda","Insecta","Diptera","Chironomidae","Orthocladiinae","Smittia","Smittia cf. stercoraria, Smittia stercoraria","1","1","1","52","0","0","98.88","448","0","450","36.22","0"
"AGAW1","05N","INS","BOLD:AAN5219","Arthropoda","Insecta","Diptera","Cecidomyiidae","","","","1","1","1","52","0","0","97.55","449","0","450","30.22","0"
"AGAW1","05N","INS","BOLD:AAG0161","Arthropoda","Insecta","Diptera","Milichiidae","Madizinae","Leptometopa","Leptometopa latipes","1","1","1","52","0","0","99.55","449","0","450","32.22","0"
"AGAW1","05N","INS","BOLD:AAU6970","Arthropoda","Insecta","Coleoptera","Cleridae","Clerinae","Enoclerus","Enoclerus nigripes","1","1","1","52","0","0","98.79","413","0","442","33.48","0"
"AGAW1","05N","INS","BOLD:AAB7030","Arthropoda","Insecta","Diptera","Chironomidae","Chironominae","Chironomus","Chironomus sp. BOLDAAP2996, Chironomus quinnitukqut, Chironomus cf. decorus 2g, Chironomus decorus group sp. 2 IP2013, Chironomus cf. decorus, Chironomus decorus gr.","1","1","1","52","0","0","98.61","361","0","365","33.42","0"
"AGAW1","05N","INS","BOLD:AAH3548","Arthropoda","Insecta","Diptera","Sphaeroceridae","Copromyzinae","Lotophila","Lotophila sp., Lotophila atra","1","1","1","52","0","0","98.44","450","0","450","32.44","0"
"AGAW1","05N","INS","BOLD:ACP6694","Arthropoda","Insecta","Diptera","Chironomidae","Orthocladiinae","","","1","1","1","52","0","0","98.22","450","0","450","34","0"
"AGAW1","05N","INS","BOLD:ACF5939","Arthropoda","Insecta","Diptera","Tachinidae","Exoristinae","Winthemia","Winthemia sinuata, Winthemia illinoensis","1","1","1","52","0","0","100","448","0","450","29.33","0"
"AGAW1","05N","INS","BOLD:ABA9949","Arthropoda","Insecta","Coleoptera","Chrysomelidae","Galerucinae","Chaetocnema","Chaetocnema concinna","1","1","1","52","0","0","98.44","450","0","450","36.67","0"
"AGAW1","05N","INS","BOLD:AAB1148","Arthropoda","Insecta","Diptera","Chloropidae","Oscinellinae","Oscinella","Oscinella sp. TAW3, Oscinella sp. 4, Oscinella sp. BOLD:AAB1148","1","1","1","52","0","0","97.1","449","0","450","32","0"
"AGAW1","05N","INS","BOLD:AAU6967","Arthropoda","Insecta","Coleoptera","Chrysomelidae","Galerucinae","Psylliodes","Psylliodes affinis","1","1","1","52","0","0","97.05","440","0","450","35.11","0"
"AGAW1","05N","INS","BOLD:AAG6588","Arthropoda","Insecta","Coleoptera","Chrysomelidae","Cassidinae","Deloyala","Deloyala guttata","1","1","1","52","0","0","99.46","367","0","369","31.17","0"
"AGAW1","05N","INS","BOLD:AAG6745","Arthropoda","Insecta","Diptera","Calliphoridae","Calliphorinae","Pollenia","Pollenia pediculata","1","1","1","52","0","0","99.55","449","0","450","33.78","0"
"AGAW1","05N","INS","BOLD:AAQ3566","Arthropoda","Insecta","Diptera","Anthomyiidae","Pegomyinae","Pegomya","","1","1","1","52","0","0","97.56","450","0","450","32","0"
"AGAW1","05N","INS","BOLD:AAG3236","Arthropoda","Insecta","Diptera","Phoridae","Phorinae","Conicera, Diplonevra","Diplonevra nitidula","1","1","1","52","0","0","98.22","450","0","450","32.67","0"
"AGAW1","05N","INS","BOLD:AAP9022","Arthropoda","Insecta","Diptera","Cecidomyiidae","","","","1","1","1","52","0","0","99.07","430","0","431","28.31","0"
"AGAW1","05N","INS","BOLD:AAH7158","Arthropoda","Insecta","Diptera","Sarcophagidae","Sarcophaginae","Ravinia","Ravinia stimulans","1","1","1","52","0","0","98.88","446","0","450","33.33","0"
"AGAW1","05N","INS","BOLD:AAD4355","Arthropoda","Insecta","Diptera","Culicidae","Culicinae","Aedes","Aedes provocans","1","1","1","52","0","0","99.27","411","0","416","31.49","0"
"AGAW1","05N","INS","BOLD:ABX6186","Arthropoda","Insecta","Diptera","Tipulidae","Tipulinae","Nephrotoma","Nephrotoma ferruginea","1","1","1","52","0","0","98.43","447","0","450","34.67","0"
"AGAW1","05N","INS","BOLD:AAB2384","Arthropoda","Insecta","Diptera","Syrphidae","Syrphinae","Eupeodes","Eupeodes duseki, Eupeodes volucris, Eupeodes nielseni, Eupeodes americanus, Eupeodes flukei, Eupeodes vockerothi, Eupeodes lucasi, Eupeodes montivagus, Eupeodes rufipunctatus, Eupeodes rojasi, Eupeodes goeldlini, Eupeodes luniger, Eupeodes punctifer, Eupeodes bucculatus, Eupeodes perplexus, Eupeodes abiskoensis, Eupeodes corollae, Eupeodes pomus, Eupeodes lundbecki, Eupeodes nitens, Eupeodes sp. BOLDAAB2384, Eupeodes curtus, Eupeodes latifasciatus","1","1","1","52","0","0","97.29","369","0","372","31.18","0"
"AGAW1","05N","INS","BOLD:ACK7296","Arthropoda","Insecta","Diptera","Cecidomyiidae","","","","1","1","1","52","0","0","99.33","448","0","450","24.22","0"
"AGAW1","05N","INS","BOLD:ABV1237","Arthropoda","Insecta","Diptera","Chironomidae","Chironominae","Chironomus","","1","1","1","52","0","0","98","450","0","450","34.22","0"
"AGAW1","05N","INS","BOLD:AAH3228","Arthropoda","Insecta","Psocodea","Caeciliusidae","Caeciliusinae","Valenzuela","Valenzuela flavidus","1","1","1","52","0","0","98.5","400","0","400","35.25","0"
"AGAW1","05N","INS","BOLD:AAE9431","Arthropoda","Insecta","Coleoptera","Cerambycidae","Lamiinae","Tetrops","Tetrops praeusta, Tetrops starkii","1","1","1","52","0","0","99.55","449","0","450","35.56","0"
"AGAW1","05N","INS","BOLD:ACB0984","Arthropoda","Insecta","Psocodea","Stenopsocidae","","Graphopsocus","Graphopsocus cruciatus","1","1","1","52","0","0","99.52","414","0","450","37.78","0"
"AGAW1","05N","INS","BOLD:AAE7016","Arthropoda","Insecta","Lepidoptera","Gelechiidae","Dichomeridinae","Helcystogramma","Helcystogramma hystricella","1","1","1","52","0","0","98.75","399","0","404","30.94","0"
"AGAW1","05N","INS","BOLD:AAA8451","Arthropoda","Insecta","Lepidoptera","Hepialidae","","Korscheltellus","Korscheltellus lupulina","1","1","1","52","0","0","100","449","0","450","28.22","0"
"AGAW1","05N","INS","BOLD:ADH9310","Arthropoda","Insecta","Diptera","Anthomyiidae","Anthomyiinae","Anthomyia","Anthomyia procellaris, Anthomyia pluvialis","1","1","1","52","0","0","99.43","352","0","352","32.67","0"
"AGAW1","05N","NAR","no match","no match","no match","no match","no match","no match","no match","no match","no match","no match","no match","no match","no match","no match","no match","no match","no match","no match","no match","no match"
"AGAW1","05N","NIN","BOLD:AAV0589","Arthropoda","Arachnida","Mesostigmata","Blattisociidae","Platyseiinae","Cheiroseius","","34","32","32","1","5.88","0","99.32","447.12","0","448.53","27.56","0"
"AGAW1","05N","NIN","BOLD:AAI4346","Arthropoda","Arachnida","Opiliones","Phalangiidae","","Paroligolophus, Phalangium","Paroligolophus agrestis, Phalangium opilio","2","2","2","2","0","0","98.51","436.5","0","449","36.75","0"
"AGAW1","05N","NIN","BOLD:ACX8969","Arthropoda","Arachnida","Trombidiformes","Eupodidae","","","","1","1","1","3","0","0","99.27","410","0","410","27.56","0"
"AGAW1","05N","NIN","BOLD:AAB5726","Arthropoda","Arachnida","Araneae","Agelenidae","","Agelenopsis","Agelenopsis pennsylvanica, Agelenopsis emertoni, Agelenopsis potteri, Agelenopsis actuosa, Agelenopsis kastoni","1","1","1","3","0","0","99.78","450","0","450","33.11","0"
"AGAW1","05S","BAC","TAX:91391","Proteobacteria","Alphaproteobacteria","Rickettsiales",NA,NA,NA,NA,"1","1","1","1","0","0","99.29","422","0","450","39.78","0"
"AGAW1","05S","CHO","no match","no match","no match","no match","no match","no match","no match","no match","no match","no match","no match","no match","no match","no match","no match","no match","no match","no match","no match","no match"
"AGAW1","05S","INS","BOLD:AAA4222","Arthropoda","Insecta","Lepidoptera","Erebidae","Hypeninae","Hypena","Hypena scabra","48336","39753","39753","1","17.76","0","99.25","398.05","0","399.67","32.45","0"
"AGAW1","05S","INS","BOLD:AAG2511","Arthropoda","Insecta","Diptera","Anthomyiidae","Anthomyiinae","Delia","Delia platura","7022","6010","6010","2","14.41","0","98.94","443.99","0","446.38","33.15","0"
"AGAW1","05S","INS","BOLD:AAG1717","Arthropoda","Insecta","Diptera","Muscidae","Phaoniinae","Helina","Helina troene","4968","4112","4112","3","17.23","0","99.23","435.66","0","436.68","30.89","0"
"AGAW1","05S","INS","BOLD:ABX6186","Arthropoda","Insecta","Diptera","Tipulidae","Tipulinae","Nephrotoma","Nephrotoma ferruginea","936","857","857","4","8.44","0","99.17","427.72","0","429.58","34.01","0"
"AGAW1","05S","INS","BOLD:AAG3236","Arthropoda","Insecta","Diptera","Phoridae","Phorinae","Conicera, Diplonevra","Diplonevra nitidula","727","584","584","5","19.67","0","99.42","446.32","0","447.79","32.09","0"
"AGAW1","05S","INS","BOLD:AAF9014","Arthropoda","Insecta","Diptera","Limoniidae","Chioneinae","Symplecta","Symplecta cana, Symplecta sp. BOLDAAF9014","693","665","665","6","4.04","0","98.06","421.51","0","426.59","34.27","0"
"AGAW1","05S","INS","BOLD:ACU7906","Arthropoda","Insecta","Diptera","Chironomidae","Orthocladiinae","","","585","583","583","7","0.34","0","98.8","405.99","0","408.38","36.11","0"
"AGAW1","05S","INS","BOLD:AAF9771","Arthropoda","Insecta","Diptera","Hybotidae","Tachydromiinae","Platypalpus","Platypalpus longiseta, Platypalpus trivialis, Platypalpus sp. 5, Platypalpus sp. 17, Platypalpus pallidiventris","532","427","427","8","19.74","0","99.39","443.7","0","444.34","35.15","0"
"AGAW1","05S","INS","BOLD:AAN5342","Arthropoda","Insecta","Diptera","Chironomidae","Orthocladiinae","Heleniella","","385","372","372","9","3.38","0","98.94","421.11","0","428.69","35.12","0"
"AGAW1","05S","INS","BOLD:AAG2868","Arthropoda","Insecta","Hemiptera","Cicadellidae","Typhlocybinae","Empoasca","Empoasca sp., Empoasca fabae","312","232","232","10","25.64","0","99.34","442.7","0","447.45","32.27","0"
"AGAW1","05S","INS","BOLD:AAN5355","Arthropoda","Insecta","Diptera","Chironomidae","Orthocladiinae","Smittia","Smittia cf. stercoraria, Smittia stercoraria","292","289","289","11","1.03","0","98.85","432.65","0","435.5","36.15","0"
"AGAW1","05S","INS","BOLD:ACP6694","Arthropoda","Insecta","Diptera","Chironomidae","Orthocladiinae","","","279","270","270","12","3.23","0","98.83","414.99","0","416.82","34.23","0"
"AGAW1","05S","INS","BOLD:ABA9949","Arthropoda","Insecta","Coleoptera","Chrysomelidae","Galerucinae","Chaetocnema","Chaetocnema concinna","256","203","203","13","20.7","0","99.35","438.9","0","439.84","36.3","0"
"AGAW1","05S","INS","BOLD:ACF8552","Arthropoda","Insecta","Coleoptera","Anthicidae","Anthicinae","Malporus","Malporus formicarius","187","154","154","14","17.65","0","99.71","364.58","0","433.59","34.26","0"
"AGAW1","05S","INS","BOLD:AAY3269","Arthropoda","Insecta","Diptera","Chironomidae","Diamesinae","Potthastia","Potthastia gaedii","183","168","168","15","8.2","0","98.05","432.1","0","433.2","34.47","0"
"AGAW1","05S","INS","BOLD:ACK3431","Arthropoda","Insecta","Diptera","Cecidomyiidae","","","","113","112","112","16","0.88","0","99.12","422.73","0","424.12","32.69","0"
"AGAW1","05S","INS","BOLD:AAN6447","Arthropoda","Insecta","Diptera","Sciaridae","","Corynoptera","Corynoptera perpusilla","76","65","65","17","14.47","0","99.36","448.39","0","449.59","36.39","0"
"AGAW1","05S","INS","BOLD:ACI4127","Arthropoda","Insecta","Diptera","Chironomidae","Chironominae","","","45","45","45","18","0","0","98.72","442.56","0","443.84","35.27","0"
"AGAW1","05S","INS","BOLD:AAF4817","Arthropoda","Insecta","Diptera","Chironomidae","Orthocladiinae","Smittia","Smittia edwardsi","40","36","36","19","10","0","98.67","375.55","0","443.3","33.08","0"
"AGAW1","05S","INS","BOLD:AAA3453","Arthropoda","Insecta","Diptera","Anthomyiidae","Anthomyiinae","Delia","","32","32","32","20","0","0","98.83","430","0","433.16","34.61","0"
"AGAW1","05S","INS","BOLD:ABZ0908","Arthropoda","Insecta","Diptera","Tipulidae","Tipulinae","Nephrotoma","","30","30","30","21","0","0","97.49","375.97","0","378.33","33.95","0"
"AGAW1","05S","INS","BOLD:ABA5798","Arthropoda","Insecta","Hemiptera","Cicadellidae","Typhlocybinae","Erythroneura","Erythroneura elegans","29","28","28","22","3.45","0","97.91","424.83","0","426.07","36.06","0"
"AGAW1","05S","INS","BOLD:AAM7064","Arthropoda","Insecta","Diptera","Chironomidae","Orthocladiinae","Smittia","Smittia sp. 14ES","26","26","26","23","0","0","98.81","432.81","0","438.92","31.79","0"
"AGAW1","05S","INS","BOLD:ABU5525","Arthropoda","Insecta","Diptera","Chironomidae","Orthocladiinae","Limnophyes","Limnophyes sp. 14ES","24","24","24","24","0","0","98.94","407.71","0","412.92","37.2","0"
"AGAW1","05S","INS","BOLD:AAL7819","Arthropoda","Insecta","Diptera","Psychodidae","Psychodinae","Psychoda","Psychoda sp., Psychoda sp. BOLD:AAL7819","20","20","20","25","0","0","99.27","396.15","0","397.85","32.74","0"
"AGAW1","05S","INS","BOLD:AAN5351","Arthropoda","Insecta","Diptera","Chironomidae","Tanypodinae","Conchapelopia","Conchapelopia telema","20","20","20","25","0","0","99.01","424.55","0","426.35","33.83","0"
"AGAW1","05S","INS","BOLD:ABU5521","Arthropoda","Insecta","Diptera","Sciaridae","","Corynoptera","Corynoptera furcata","17","17","17","27","0","0","98.65","395.94","0","398.29","36.05","0"
"AGAW1","05S","INS","BOLD:AAN5356","Arthropoda","Insecta","Diptera","Chironomidae","Orthocladiinae","Smittia","","14","14","14","28","0","0","98.67","425.07","0","426.86","34.87","0"
"AGAW1","05S","INS","BOLD:AAZ5601","Arthropoda","Insecta","Diptera","Chironomidae","Orthocladiinae","Smittia","","11","11","11","29","0","0","99.61","434.09","0","436","33.68","0"
"AGAW1","05S","INS","BOLD:AAF9187","Arthropoda","Insecta","Coleoptera","Curculionidae","Entiminae","Phyllobius","Phyllobius oblongus","10","10","10","30","0","0","99.32","442","0","442.4","39.06","0"
"AGAW1","05S","INS","BOLD:AAA7374","Arthropoda","Insecta","Diptera","Syrphidae","Syrphinae","Sphaerophoria","Sphaerophoria bankowskae, Sphaerophoria infuscata, Sphaerophoria nr. philanthus, Sphaerophoria batava, Sphaerophoria sp., Sphaerophoria rueppellii, Sphaerophoria interrupta-group, Sphaerophoria sp. MA-1, Sphaerophoria cleoae, Sphaerophoria asymmetrica, Sphaerophoria fatarum, Sphaerophoria bengalensis, Sphaerophoria bifurcata, Sphaerophoria taeniata, Sphaerophoria interrupta, Sphaerophoria contigua, Sphaerophoria kaa, Sphaerophoria laurae, Sphaerophoria chongjini, Sphaerophoria boreoalpina, Sphaerophoria sp. BOLDAAA7374, Sphaerophoria macrogaster, Sphaerophoria sulphuripes, Sphaerophoria brevipilosa, Sphaerophoria philanthus, Sphaerophoria pyrrhina, Sphaerophoria abbreviata, Sphaerophoria scripta, Sphaerophoria potentillae, Sphaerophoria longipilosa, Sphaerophoria virgata","8","8","8","31","0","0","99.04","424.38","0","426.63","31.1","0"
"AGAW1","05S","INS","BOLD:ADH4973","Arthropoda","Insecta","Diptera","Tipulidae","Tipulinae","Nephrotoma","","7","7","7","32","0","0","97.8","397.14","0","400.43","33.39","0"
"AGAW1","05S","INS","BOLD:AAV5325","Arthropoda","Insecta","Diptera","Cecidomyiidae","","","CecidInt12 sp. BOLD:AAV5325","7","7","7","32","0","0","99.42","431.43","0","431.14","28.19","0"
"AGAW1","05S","INS","BOLD:AAG6743","Arthropoda","Insecta","Diptera","Sarcophagidae","Sarcophaginae","Sarcophaga","Sarcophaga sp. 1 MG, Sarcophaga subvicina","6","6","6","34","0","0","99.1","444.67","0","445.17","34.95","0"
"AGAW1","05S","INS","BOLD:AAB9140","Arthropoda","Insecta","Diptera","Calliphoridae","Chrysomyinae","Phormia","Phormia regina","6","6","6","34","0","0","98.93","432","0","433.5","33.17","0"
"AGAW1","05S","INS","BOLD:AAB2644","Arthropoda","Insecta","Diptera","Chironomidae","Orthocladiinae","Orthocladius","Orthocladius lapponicus, Orthocladius dorenus, Orthocladius mallochi","5","5","5","36","0","0","98.4","448.6","0","450","32.98","0"
"AGAW1","05S","INS","BOLD:AAO3939","Arthropoda","Insecta","Diptera","Limoniidae","Limoniinae","Dicranomyia","","5","5","5","36","0","0","98.79","445.6","0","448","36.56","0"
"AGAW1","05S","INS","BOLD:AAG2462","Arthropoda","Insecta","Diptera","Anthomyiidae","Anthomyiinae","Botanophila","Botanophila fugax","5","5","5","36","0","0","98.57","448.2","0","449.6","32.12","0"
"AGAW1","05S","INS","BOLD:ABX6298","Arthropoda","Insecta","Diptera","Tipulidae","Tipulinae","Nephrotoma","Nephrotoma ferruginea","4","4","4","39","0","0","98.6","442.75","0","443.25","34.17","0"
"AGAW1","05S","INS","BOLD:AAA3654","Arthropoda","Insecta","Lepidoptera","Crambidae","Crambinae","Crambus","Crambus bigelovi, Crambus johnsoni, Crambus praefectellus","4","4","4","39","0","0","98.98","436.5","0","438","31.82","0"
"AGAW1","05S","INS","BOLD:ACR6129","","","","","","","","4","4","4","39","0","0","97.83","449.75","0","450","36.33","0"
"AGAW1","05S","INS","BOLD:AAH4135","Arthropoda","Insecta","Diptera","Chloropidae","Chloropinae","Thaumatomyia","Thaumatomyia glabra, Thaumatomyia grata, Thaumatomyia sp. 1 TAW","3","3","3","42","0","0","98.89","448.67","0","450","32.15","0"
"AGAW1","05S","INS","BOLD:ACR6132","","","","","","","","3","3","3","42","0","0","98.14","403.33","0","417","35.27","0"
"AGAW1","05S","INS","BOLD:AAH3664","Arthropoda","Insecta","Diptera","Cecidomyiidae","","","CecidInt3 sp. BOLD:AAH3664","3","3","3","42","0","0","98.59","449.67","0","450","27.63","0"
"AGAW1","05S","INS","BOLD:AAA9251","Arthropoda","Insecta","Lepidoptera","Gelechiidae","Gelechiinae","Scrobipalpa","Scrobipalpa atriplicella, Scrobipalpa nr. atriplicella sp. 1","3","3","3","42","0","0","99.07","415.67","0","418.33","34.24","0"
"AGAW1","05S","INS","BOLD:AAA9101","Arthropoda","Insecta","Lepidoptera","Pyralidae","Phycitinae","Moodna, Caudellia","Moodna ostrinella, Moodna bisinuella, Caudellia nigrella","3","2","2","42","33.33","0","99.33","450","0","450","34.44","0"
"AGAW1","05S","INS","BOLD:AAH7158","Arthropoda","Insecta","Diptera","Sarcophagidae","Sarcophaginae","Ravinia","Ravinia stimulans","3","3","3","42","0","0","98.25","392.33","0","397.67","33.54","0"
"AGAW1","05S","INS","BOLD:AAN6437","Arthropoda","Insecta","Diptera","Sciaridae","","Corynoptera","Corynoptera saccata","3","3","3","42","0","0","99.27","405","0","406.33","39.29","0"
"AGAW1","05S","INS","BOLD:AAG1701","Arthropoda","Insecta","Diptera","Muscidae","Coenosiinae","Macrorchis","Macrorchis ausoba","3","3","3","42","0","0","99.29","439.33","0","441.67","33.05","0"
"AGAW1","05S","INS","BOLD:AAG4653","Arthropoda","Insecta","Diptera","Syrphidae","Syrphinae","Heringia","Heringia latitarsis, Heringia pubescens, Heringia jakutorum, Heringia rita, Heringia fulvimanus, Heringia sp. BOLDAAG4653, Heringia nr. pubescens, Heringia coxalis, Heringia larusi, Heringia vitripennis","3","3","3","42","0","0","99.93","449.67","0","450","31.41","0"
"AGAW1","05S","INS","BOLD:AAP6465","Arthropoda","Insecta","Diptera","Sciaridae","","Corynoptera","Corynoptera cursor","3","2","2","42","33.33","0","99.33","450","0","450","32.67","0"
"AGAW1","05S","INS","BOLD:AAA3933","Arthropoda","Insecta","Lepidoptera","Erebidae","Herminiinae","Palthis","Palthis angulalis","2","2","2","52","0","0","98.97","437.5","0","438.5","32.61","0"
"AGAW1","05S","INS","BOLD:AAA4121","Arthropoda","Insecta","Diptera","Simuliidae","Simuliinae","Simulium","Simulium vittatum complex, Simulium vittatum, Simulium sp. BOLDAAA4121, Simulium argus, Simulium vittatum s.str.","2","2","2","52","0","0","97.96","402.5","0","404","33.09","0"
"AGAW1","05S","INS","BOLD:ABY9574","Arthropoda","Insecta","Lepidoptera","Noctuidae","Noctuinae","Hyppa","Hyppa xylinoides","2","2","2","52","0","0","98.86","425.5","0","426.5","31.74","0"
"AGAW1","05S","INS","BOLD:AAU6630","Arthropoda","Insecta","Diptera","Fanniidae","Fanniinae","Fannia","Fannia armata","2","2","2","52","0","0","99","448.5","0","450","33.11","0"
"AGAW1","05S","INS","BOLD:AAG5457","Arthropoda","Insecta","Diptera","Chironomidae","Chironominae","","","2","2","2","52","0","0","99.03","412.5","0","414.5","33.65","0"
"AGAW1","05S","INS","BOLD:ABW7995","Arthropoda","Insecta","Diptera","Dolichopodidae","Dolichopodinae","Gymnopternus","Gymnopternus nigribarbus","2","2","2","52","0","0","99.86","392","0","402","35.94","0"
"AGAW1","05S","INS","BOLD:ABZ7601","Arthropoda","Insecta","Lepidoptera","Noctuidae","Noctuinae","Anarta","Anarta trifolii","2","2","2","52","0","0","98.56","450","0","450","31.44","0"
"AGAW1","05S","INS","BOLD:AAH3914","Arthropoda","Insecta","Diptera","Sciaridae","","Sciara","Sciara hamatilis, Sciara humeralis","2","2","2","52","0","0","99.24","399.5","0","401","29.85","0"
"AGAW1","05S","INS","BOLD:ABY4763","Arthropoda","Insecta","Hemiptera","Aphididae","Aphidinae","Uroleucon","Uroleucon taraxaci","2","2","2","52","0","0","99.67","449.5","0","450","25.11","0"
"AGAW1","05S","INS","BOLD:AAB5609","Arthropoda","Insecta","Diptera","Muscidae","Coenosiinae","Coenosia","Coenosia sp.BOLD:AAB5609, Coenosia tigrina","2","2","2","52","0","0","99.01","403.5","0","403.5","32.22","0"
"AGAW1","05S","INS","BOLD:AAA2482","Arthropoda","Insecta","Lepidoptera","Noctuidae","Noctuinae","Mythimna","Mythimna unipuncta","2","2","2","52","0","0","98.96","427","0","431","30.73","0"
"AGAW1","05S","INS","BOLD:AAD2548","Arthropoda","Insecta","Hemiptera","Aphididae","Aphidinae","Acyrthosiphon","Acyrthosiphon caraganae, Acyrthosiphon sp. 1, Acyrthosiphon kondoi, Acyrthosiphon churchillense","2","2","2","52","0","0","99.89","450","0","450","26.11","0"
"AGAW1","05S","INS","BOLD:AAH2876","Arthropoda","Insecta","Diptera","Cecidomyiidae","","","CecidInt51 sp. BOLD:AAH2876","2","2","2","52","0","0","99.56","450","0","450","27","0"
"AGAW1","05S","INS","BOLD:AAB8817","Arthropoda","Insecta","Diptera","Muscidae","Azeliinae","Muscina","Muscina levida","2","2","2","52","0","0","99.33","442","0","444.5","33.18","0"
"AGAW1","05S","INS","BOLD:AAC9062","Arthropoda","Insecta","Diptera","Culicidae","Culicinae","Ochlerotatus, Aedes","Aedes fitchii, Ochlerotatus fitchii, Aedes grossbecki, Aedes ventrovittis","2","2","2","52","0","0","98.95","423","0","428.5","32.77","0"
"AGAW1","05S","INS","BOLD:ACB0984","Arthropoda","Insecta","Psocodea","Stenopsocidae","","Graphopsocus","Graphopsocus cruciatus","2","2","2","52","0","0","99.52","413","0","450","37.33","0"
"AGAW1","05S","INS","BOLD:ADH9310","Arthropoda","Insecta","Diptera","Anthomyiidae","Anthomyiinae","Anthomyia","Anthomyia procellaris, Anthomyia pluvialis","2","2","2","52","0","0","97.63","418.5","0","421.5","31.89","0"
"AGAW1","05S","INS","BOLD:AAP2998","Arthropoda","Insecta","Diptera","Chironomidae","Tanypodinae","","","1","1","1","69","0","0","100","313","0","351","36.47","0"
"AGAW1","05S","INS","BOLD:AAA2590","Arthropoda","Insecta","Lepidoptera","Noctuidae","Noctuinae","Xestia","Xestia baja, Xestia smithii","1","1","1","69","0","0","99.78","450","0","450","33.11","0"
"AGAW1","05S","INS","BOLD:ACT0257","","","","","","","","1","1","1","69","0","0","99.78","450","0","450","38.22","0"
"AGAW1","05S","INS","BOLD:AAA3836","Arthropoda","Insecta","Lepidoptera","Noctuidae","Plusiinae","Autographa","Autographa precationis","1","1","1","69","0","0","99.78","449","0","450","32","0"
"AGAW1","05S","INS","BOLD:ACE0699","Arthropoda","Insecta","Diptera","Chironomidae","","","","1","1","1","69","0","0","97.44","430","0","450","34.67","0"
"AGAW1","05S","INS","BOLD:AAG7276","Arthropoda","Insecta","Diptera","Sphaeroceridae","Limosininae","Leptocera","Leptocera erythrocera","1","1","1","69","0","0","97.7","434","0","435","33.79","0"
"AGAW1","05S","INS","BOLD:AAP7030","Arthropoda","Insecta","Coleoptera","Cryptophagidae","Atomariinae","Atomaria","Atomaria ephippiata","1","1","1","69","0","0","99.31","433","0","433","37.18","0"
"AGAW1","05S","INS","BOLD:AAN6433","Arthropoda","Insecta","Diptera","Sciaridae","","Corynoptera","Corynoptera deserta","1","1","1","69","0","0","97.32","447","0","450","30.67","0"
"AGAW1","05S","INS","BOLD:ACU3412","Arthropoda","Insecta","Diptera","Chironomidae","Orthocladiinae","","","1","1","1","69","0","0","97.77","448","0","450","34.67","0"
"AGAW1","05S","INS","BOLD:AAO8538","Arthropoda","Insecta","Diptera","Psychodidae","Psychodinae","Psychoda","","1","1","1","69","0","0","97.25","436","0","437","33.18","0"
"AGAW1","05S","INS","BOLD:AAB7490","Arthropoda","Insecta","Diptera","Chironomidae","Orthocladiinae","Tvetenia","Tvetenia paucunca","1","1","1","69","0","0","99.11","447","0","450","34.22","0"
"AGAW1","05S","INS","BOLD:AAN6406","Arthropoda","Insecta","Diptera","Sphaeroceridae","Limosininae","Opalimosina","Opalimosina mirabilis","1","1","1","69","0","0","97.32","448","0","450","33.33","0"
"AGAW1","05S","INS","BOLD:ADE1333","Arthropoda","Insecta","Hemiptera","Cicadellidae","Typhlocybinae","Erythroneura","","1","1","1","69","0","0","97.91","383","0","450","36","0"
"AGAW1","05S","INS","BOLD:ACA7493","Arthropoda","Insecta","Diptera","Chironomidae","Chironominae","","","1","1","1","69","0","0","97.04","372","0","372","40.59","0"
"AGAW1","05S","INS","BOLD:ABY2036","Arthropoda","Insecta","Diptera","Sciaridae","","","","1","1","1","69","0","0","99.55","448","0","450","29.33","0"
"AGAW1","05S","INS","BOLD:AAF9041","Arthropoda","Insecta","Diptera","Tipulidae","Tipulinae","Tipula","Tipula oleracea","1","1","1","69","0","0","99.15","351","0","352","35.23","0"
"AGAW1","05S","INS","BOLD:AAZ5973","Arthropoda","Insecta","Diptera","Sepsidae","Sepsinae","Themira","Themira annulipes","1","1","1","69","0","0","98.66","449","0","450","36.22","0"
"AGAW1","05S","INS","BOLD:AAG4827","Arthropoda","Insecta","Diptera","Anthomyzidae","Anthomyzinae","Mumetopia","Mumetopia occipitalis","1","1","1","69","0","0","97.1","448","0","450","36","0"
"AGAW1","05S","INS","BOLD:AAF9768","Arthropoda","Insecta","Diptera","Hybotidae","Tachydromiinae","Platypalpus","Platypalpus holosericus, Platypalpus candicans, Platypalpus sp. BOLD:AAF9768, Platypalpus sp. 22","1","1","1","69","0","0","97.66","385","0","387","31.78","0"
"AGAW1","05S","INS","BOLD:AAA5448","Arthropoda","Insecta","Lepidoptera","Geometridae","Ennominae","Eutrapela","Eutrapela clemataria","1","1","1","69","0","0","98.89","450","0","450","30.44","0"
"AGAW1","05S","INS","BOLD:ACU4327","Arthropoda","Insecta","Diptera","Chironomidae","Orthocladiinae","","","1","1","1","69","0","0","98.02","353","0","359","34.26","0"
"AGAW1","05S","INS","BOLD:AAG0991","Arthropoda","Insecta","Diptera","Chironomidae","Orthocladiinae","Orthocladius","","1","1","1","69","0","0","99.74","381","0","383","32.11","0"
"AGAW1","05S","INS","BOLD:ACG5862","Arthropoda","Insecta","Diptera","Agromyzidae","","","","1","1","1","69","0","0","98.65","446","0","450","33.78","0"
"AGAW1","05S","INS","BOLD:AAM6047","Arthropoda","Insecta","Diptera","Cecidomyiidae","","","","1","1","1","69","0","0","99.11","450","0","450","32.67","0"
"AGAW1","05S","INS","BOLD:AAN6506","Arthropoda","Insecta","Diptera","Stratiomyidae","Sarginae","Microchrysa","Microchrysa polita, Microchrysa flavicornis","1","1","1","69","0","0","99.11","448","0","450","39.33","0"
"AGAW1","05S","INS","BOLD:ABA6514","Arthropoda","Insecta","Diptera","Chironomidae","Chironominae","Rheotanytarsus","","1","1","1","69","0","0","97.99","448","0","450","35.78","0"
"AGAW1","05S","INS","BOLD:AAN5334","Arthropoda","Insecta","Diptera","Chironomidae","Chironominae","Polypedilum","","1","1","1","69","0","0","99.11","449","0","450","35.78","0"
"AGAW1","05S","INS","BOLD:AAN5358","Arthropoda","Insecta","Diptera","Chironomidae","Orthocladiinae","Smittia","Smittia stercoraria, Smittia sp. 22ES","1","1","1","69","0","0","99.55","449","0","450","37.56","0"
"AGAW1","05S","INS","BOLD:AAZ4669","Arthropoda","Insecta","Diptera","Tachinidae","Exoristinae","Oswaldia","Oswaldia assimilis","1","1","1","69","0","0","98.88","446","0","450","32.67","0"
"AGAW1","05S","INS","BOLD:AAA2815","Arthropoda","Insecta","Lepidoptera","Noctuidae","Noctuinae","Mythimna","Mythimna yukonensis, Mythimna oxygala","1","1","1","69","0","0","99.78","449","0","450","33.33","0"
"AGAW1","05S","INS","BOLD:AAJ7105","Arthropoda","Insecta","Diptera","Agromyzidae","Agromyzinae","Agromyza","Agromyza frontella","1","1","1","69","0","0","99.75","400","0","450","36.44","0"
"AGAW1","05S","INS","BOLD:AAB1098","Arthropoda","Insecta","Diptera","Culicidae","Culicinae","Ochlerotatus, Aedes","Aedes excrucians, Aedes cantans, Aedes stimulans, Aedes euedes, Aedes riparius, Aedes annulipes, Ochlerotatus cantans, Aedes fitchii, Aedes sp., Ochlerotatus annulipes, Ochlerotatus excrucians","1","1","1","69","0","0","99.33","449","0","450","35.33","0"
"AGAW1","05S","INS","BOLD:ACF5939","Arthropoda","Insecta","Diptera","Tachinidae","Exoristinae","Winthemia","Winthemia sinuata, Winthemia illinoensis","1","1","1","69","0","0","99.76","409","0","412","28.88","0"
"AGAW1","05S","INS","BOLD:AAD0853","Arthropoda","Insecta","Diptera","Scathophagidae","Scathophaginae","Scathophaga","Scathophaga lutaria, Scathophaga hiemalis, Scathophaga furcata, Scathophaga stercoraria, Scathophaga inquinata, Scathophaga suilla","1","1","1","69","0","0","98.65","446","0","450","31.78","0"
"AGAW1","05S","INS","BOLD:AAG6745","Arthropoda","Insecta","Diptera","Calliphoridae","Calliphorinae","Pollenia","Pollenia pediculata","1","1","1","69","0","0","100","428","0","428","33.88","0"
"AGAW1","05S","INS","BOLD:AAP9022","Arthropoda","Insecta","Diptera","Cecidomyiidae","","","","1","1","1","69","0","0","99.56","450","0","450","27.56","0"
"AGAW1","05S","INS","BOLD:AAD4355","Arthropoda","Insecta","Diptera","Culicidae","Culicinae","Aedes","Aedes provocans","1","1","1","69","0","0","97.11","450","0","450","32.44","0"
"AGAW1","05S","INS","BOLD:AAA4555","Arthropoda","Insecta","Orthoptera","Acrididae","Melanoplinae","Melanoplus","Melanoplus stupefactus, Melanoplus angustipennis, Melanoplus ponderosus, Melanoplus bruneri, Melanoplus frigidus, Melanoplus scapularis, Melanoplus cinereus, Melanoplus madeleineae, Melanoplus sanguinipes, Melanoplus nigrescens group, Melanoplus tunicae, Melanoplus walshi, Melanoplus femurrubrum, Melanoplus dawsoni, Melanoplus sp., Melanoplus fasciatus, Melanoplus bowditchi, Melanoplus stonei, Melanoplus bivittatus, Melanoplus huroni, Melanoplus thomasi, Melanoplus borealis, Melanoplus packardii","1","1","1","69","0","0","98.22","449","0","450","36.44","0"
"AGAW1","05S","INS","BOLD:AAN5183","Arthropoda","Insecta","Diptera","Cecidomyiidae","","","","1","1","1","69","0","0","99.07","432","0","432","28.01","0"
"AGAW1","05S","INS","BOLD:AAN5505","Arthropoda","Insecta","Diptera","Hybotidae","Tachydromiinae","Platypalpus","Platypalpus sp. BOLD:AAN5505, Platypalpus minutus, Platypalpus nr. minutus, Platypalpus australominutus","1","1","1","69","0","0","99.78","448","0","450","38","0"
"AGAW1","05S","INS","BOLD:ACA2933","Arthropoda","Insecta","Psocodea","Stenopsocidae","","Graphopsocus","Graphopsocus cruciatus","1","1","1","69","0","0","99.33","450","0","450","38.89","0"
"AGAW1","05S","INS","BOLD:ACP4736","Arthropoda","Insecta","Diptera","Chironomidae","Orthocladiinae","Smittia","Smittia sp. 8ES","1","1","1","69","0","0","98.21","447","0","450","32.44","0"
"AGAW1","05S","INS","BOLD:AAC4197","Arthropoda","Insecta","Diptera","Chironomidae","Orthocladiinae","Paraphaenocladius","Paraphaenocladius impensus","1","1","1","69","0","0","99.26","406","0","410","34.63","0"
"AGAW1","05S","INS","BOLD:ACX5521","Arthropoda","Insecta","Diptera","Chironomidae","Orthocladiinae","","","1","1","1","69","0","0","97.12","382","0","383","34.46","0"
"AGAW1","05S","INS","BOLD:ABV0255","Arthropoda","Insecta","Diptera","Chironomidae","Orthocladiinae","Limnophyes","","1","1","1","69","0","0","99.51","411","0","414","32.85","0"
"AGAW1","05S","INS","BOLD:AAY9557","Arthropoda","Insecta","Lepidoptera","Coleophoridae","Coleophorinae","Coleophora","Coleophora deauratella","1","1","1","69","0","0","97.33","450","0","450","32.44","0"
"AGAW1","05S","INS","BOLD:AAG0314","Arthropoda","Insecta","Diptera","Chironomidae","Tanypodinae","Psectrotanypus","Psectrotanypus sp. ES01","1","1","1","69","0","0","100","449","0","450","34.67","0"
"AGAW1","05S","NAR","no match","no match","no match","no match","no match","no match","no match","no match","no match","no match","no match","no match","no match","no match","no match","no match","no match","no match","no match","no match"
"AGAW1","05S","NIN","BOLD:AAB7667","Arthropoda","Arachnida","Araneae","Lycosidae","","Pardosa","Pardosa saxatilis","100","99","99","1","1","0","98.35","405.29","0","408.36","31.16","0"
"AGAW1","05S","NIN","BOLD:AAB7914","Arthropoda","Collembola","Symphypleona","Bourletiellidae","","Bourletiella","","5","5","5","2","0","0","98.94","422.8","0","424.8","37.46","0"
"AGAW1","05S","NIN","BOLD:AAV0589","Arthropoda","Arachnida","Mesostigmata","Blattisociidae","Platyseiinae","Cheiroseius","","3","3","3","3","0","0","99.92","447","0","450","27.41","0"
"AGAW1","05S","NIN","BOLD:AAI4346","Arthropoda","Arachnida","Opiliones","Phalangiidae","","Paroligolophus, Phalangium","Paroligolophus agrestis, Phalangium opilio","1","1","1","4","0","0","99.78","447","0","447","36.69","0"
"AGAW1","05S","NIN","BOLD:AAE6867","Arthropoda","Arachnida","Araneae","Salticidae","","Habronattus","Habronattus decorus","1","1","1","4","0","0","97.78","406","0","450","32.67","0"
"AGAW1","05S","NIN","BOLD:AAN6642","Arthropoda","Arachnida","Mesostigmata","Blattisociidae","Platyseiinae","Cheiroseius","","1","1","1","4","0","0","100","450","0","450","26.44","0"
"AGAW1","05S","NIN","BOLD:ABZ1645","Arthropoda","Arachnida","Araneae","Philodromidae","","Philodromus","Philodromus vulgaris","1","1","1","4","0","0","97.51","402","0","403","30.02","0"
"AGAW1","05W","BAC","TAX:375847","Proteobacteria","Alphaproteobacteria","Rickettsiales","Rickettsiaceae",NA,"Wolbachia","Wolbachia sp. Hymenoptera_ASGLE1705-10","1","1","1","1","0","0","99.09","440","0","442","40.05","0"
"AGAW1","05W","BAC","TAX:376694","Proteobacteria","Alphaproteobacteria","Rickettsiales","Rickettsiaceae",NA,"Wolbachia","Wolbachia sp. Anacrusis nephrodes","1","1","1","1","0","0","100","408","0","413","38.74","0"
"AGAW1","05W","BAC","TAX:376688","Proteobacteria","Alphaproteobacteria","Rickettsiales","Rickettsiaceae",NA,"Wolbachia","Wolbachia sp. elachBioLep01 BioLep373","1","1","1","1","0","0","99.53","427","0","427","39.58","0"
"AGAW1","05W","CHO","no match","no match","no match","no match","no match","no match","no match","no match","no match","no match","no match","no match","no match","no match","no match","no match","no match","no match","no match","no match"
"AGAW1","05W","INS","BOLD:AAO3939","Arthropoda","Insecta","Diptera","Limoniidae","Limoniinae","Dicranomyia","","27360","20079","20079","1","26.61","0","99.14","439.06","0","440.37","36.18","0"
"AGAW1","05W","INS","BOLD:AAA9101","Arthropoda","Insecta","Lepidoptera","Pyralidae","Phycitinae","Moodna, Caudellia","Moodna ostrinella, Moodna bisinuella, Caudellia nigrella","24597","15703","15703","2","36.16","0","99.59","441.66","0","442.98","34.38","0"
"AGAW1","05W","INS","BOLD:ACA2933","Arthropoda","Insecta","Psocodea","Stenopsocidae","","Graphopsocus","Graphopsocus cruciatus","11391","6869","6869","3","39.7","0","99.27","444.04","0","445.04","38.63","0"
"AGAW1","05W","INS","BOLD:AAN5342","Arthropoda","Insecta","Diptera","Chironomidae","Orthocladiinae","Heleniella","","3393","3042","3042","4","10.34","0","98.99","422.54","0","427.23","35.19","0"
"AGAW1","05W","INS","BOLD:AAB1098","Arthropoda","Insecta","Diptera","Culicidae","Culicinae","Ochlerotatus, Aedes","Aedes excrucians, Aedes cantans, Aedes stimulans, Aedes euedes, Aedes riparius, Aedes annulipes, Ochlerotatus cantans, Aedes fitchii, Aedes sp., Ochlerotatus annulipes, Ochlerotatus excrucians","3123","2982","2982","5","4.51","0","98.33","406.36","0","408.32","34.92","0"
"AGAW1","05W","INS","BOLD:ACE0699","Arthropoda","Insecta","Diptera","Chironomidae","","","","2847","2167","2167","6","23.88","0","98.28","423.79","0","438.91","35.57","0"
"AGAW1","05W","INS","BOLD:AAN6154","Arthropoda","Insecta","Coleoptera","Latridiidae","Corticariinae","Melanophthalma","Melanophthalma inermis","2311","1645","1645","7","28.82","0","98.42","437.59","0","439.6","31.32","0"
"AGAW1","05W","INS","BOLD:AAB2644","Arthropoda","Insecta","Diptera","Chironomidae","Orthocladiinae","Orthocladius","Orthocladius lapponicus, Orthocladius dorenus, Orthocladius mallochi","2189","2094","2094","8","4.34","0","98.5","423.88","0","426.54","32.89","0"
"AGAW1","05W","INS","BOLD:AAN5355","Arthropoda","Insecta","Diptera","Chironomidae","Orthocladiinae","Smittia","Smittia cf. stercoraria, Smittia stercoraria","1996","1693","1693","9","15.18","0","99.38","397.52","0","441.96","36.3","0"
"AGAW1","05W","INS","BOLD:ABX6186","Arthropoda","Insecta","Diptera","Tipulidae","Tipulinae","Nephrotoma","Nephrotoma ferruginea","1883","1591","1591","10","15.51","0","99.11","440.47","0","442.02","34.15","0"
"AGAW1","05W","INS","BOLD:AAF9014","Arthropoda","Insecta","Diptera","Limoniidae","Chioneinae","Symplecta","Symplecta cana, Symplecta sp. BOLDAAF9014","1069","1005","1005","11","5.99","0","98.12","420.23","0","428.13","33.6","0"
"AGAW1","05W","INS","BOLD:AAF9771","Arthropoda","Insecta","Diptera","Hybotidae","Tachydromiinae","Platypalpus","Platypalpus longiseta, Platypalpus trivialis, Platypalpus sp. 5, Platypalpus sp. 17, Platypalpus pallidiventris","858","627","627","12","26.92","0","98.87","444.84","0","445.26","35.05","0"
"AGAW1","05W","INS","BOLD:ACK4716","Arthropoda","Insecta","Diptera","Cecidomyiidae","","","","627","592","592","13","5.58","0","99.13","425.1","0","426.24","31.66","0"
"AGAW1","05W","INS","BOLD:AAD2085","Arthropoda","Insecta","Lepidoptera","Bucculatricidae","","Bucculatrix","Bucculatrix angustata","621","510","510","14","17.87","0","99.2","439.77","0","446.61","33","0"
"AGAW1","05W","INS","BOLD:ABW7897","Arthropoda","Insecta","Diptera","Cecidomyiidae","","","","516","399","399","15","22.67","0","99.6","441.68","0","443.28","29.72","0"
"AGAW1","05W","INS","BOLD:AAV1295","Arthropoda","Insecta","Diptera","Sciaridae","","Bradysia","Bradysia difformis, Bradysia impatiens","416","338","338","16","18.75","0","99.44","416.79","0","438.01","31.63","0"
"AGAW1","05W","INS","BOLD:ABU5546","Arthropoda","Insecta","Diptera","Psychodidae","Psychodinae","Psychoda","","374","357","357","17","4.55","0","98.61","423.33","0","428.26","31.18","0"
"AGAW1","05W","INS","BOLD:ACK7296","Arthropoda","Insecta","Diptera","Cecidomyiidae","","","","333","287","287","18","13.81","0","99.41","428.83","0","430.05","24.21","0"
"AGAW1","05W","INS","BOLD:ACX5521","Arthropoda","Insecta","Diptera","Chironomidae","Orthocladiinae","","","329","327","327","19","0.61","0","98.85","403.63","0","405.21","35.08","0"
"AGAW1","05W","INS","BOLD:AAG6934","Arthropoda","Insecta","Diptera","Hybotidae","","","","318","291","291","20","8.49","0","99.22","425.14","0","426.82","38.63","0"
"AGAW1","05W","INS","BOLD:ABZ0908","Arthropoda","Insecta","Diptera","Tipulidae","Tipulinae","Nephrotoma","","299","287","287","21","4.01","0","97.61","366.87","0","369.82","33.8","0"
"AGAW1","05W","INS","BOLD:AAU6513","Arthropoda","Insecta","Diptera","Sciaridae","","Corynoptera","Corynoptera bicuspidata","283","279","279","22","1.41","0","98.6","400.49","0","405.73","32.1","0"
"AGAW1","05W","INS","BOLD:AAV5788","Arthropoda","Insecta","Diptera","Cecidomyiidae","Cecidomyiinae","Asphondylia","","234","208","208","23","11.11","0","99.45","438.8","0","440.21","24.05","0"
"AGAW1","05W","INS","BOLD:AAN6437","Arthropoda","Insecta","Diptera","Sciaridae","","Corynoptera","Corynoptera saccata","216","216","216","24","0","0","99.27","403.24","0","404.38","39.67","0"
"AGAW1","05W","INS","BOLD:AAN6447","Arthropoda","Insecta","Diptera","Sciaridae","","Corynoptera","Corynoptera perpusilla","207","145","145","25","29.95","0","99.48","447.62","0","448.76","36.38","0"
"AGAW1","05W","INS","BOLD:AAN5249","Arthropoda","Insecta","Diptera","Cecidomyiidae","","","","178","176","176","26","1.12","0","98.73","421.7","0","423","33.37","0"
"AGAW1","05W","INS","BOLD:ADH4889","Arthropoda","Insecta","Diptera","","","","","103","91","91","27","11.65","0","98.92","445.11","0","446.36","35.52","0"
"AGAW1","05W","INS","BOLD:ABA9949","Arthropoda","Insecta","Coleoptera","Chrysomelidae","Galerucinae","Chaetocnema","Chaetocnema concinna","100","85","85","28","15","0","99.27","440.55","0","441.36","36.35","0"
"AGAW1","05W","INS","BOLD:ABA6409","Arthropoda","Insecta","Diptera","Sciaridae","","","","74","74","74","29","0","0","99.06","437.89","0","440.88","31.81","0"
"AGAW1","05W","INS","BOLD:AAG6460","Arthropoda","Insecta","Diptera","Cecidomyiidae","","","","71","71","71","30","0","0","99","439.51","0","441.42","34.84","0"
"AGAW1","05W","INS","BOLD:AAA8469","Arthropoda","Insecta","Hymenoptera","Braconidae","Euphorinae","Peristenus","Peristenus sp.","65","60","60","31","7.69","0","99.57","417.8","0","441.58","28.56","0"
"AGAW1","05W","INS","BOLD:AAC9199","Arthropoda","Insecta","Diptera","Chironomidae","Tanypodinae","Krenopelopia","","57","57","57","32","0","0","98.23","400.26","0","418.12","34.36","0"
"AGAW1","05W","INS","BOLD:ABU5520","Arthropoda","Insecta","Diptera","Sciaridae","","Corynoptera","","49","49","49","33","0","0","98.75","414.98","0","416.41","35.21","0"
"AGAW1","05W","INS","BOLD:AAH3664","Arthropoda","Insecta","Diptera","Cecidomyiidae","","","CecidInt3 sp. BOLD:AAH3664","32","32","32","34","0","0","99.22","422.31","0","427.66","26.96","0"
"AGAW1","05W","INS","BOLD:ABU5525","Arthropoda","Insecta","Diptera","Chironomidae","Orthocladiinae","Limnophyes","Limnophyes sp. 14ES","29","29","29","35","0","0","99.12","398.97","0","401.38","37.52","0"
"AGAW1","05W","INS","BOLD:ABX6298","Arthropoda","Insecta","Diptera","Tipulidae","Tipulinae","Nephrotoma","Nephrotoma ferruginea","24","24","24","36","0","0","99.29","422.21","0","424.13","33.03","0"
"AGAW1","05W","INS","BOLD:AAA3453","Arthropoda","Insecta","Diptera","Anthomyiidae","Anthomyiinae","Delia","","22","22","22","37","0","0","99","423.95","0","426.5","34.66","0"
"AGAW1","05W","INS","BOLD:ACF2457","Arthropoda","Insecta","Diptera","Chironomidae","Chironominae","Paratanytarsus","Paratanytarsus laccophilus","18","17","17","38","5.56","0","99.59","442.61","0","444.06","32.44","0"
"AGAW1","05W","INS","BOLD:AAA4222","Arthropoda","Insecta","Lepidoptera","Erebidae","Hypeninae","Hypena","Hypena scabra","15","15","15","39","0","0","99.09","389.33","0","390.4","32.62","0"
"AGAW1","05W","INS","BOLD:AAY6412","Arthropoda","Insecta","Diptera","Sciaridae","","Bradysia","Bradysia urticae","14","14","14","40","0","0","99.16","434.29","0","435.5","31.96","0"
"AGAW1","05W","INS","BOLD:AAN6433","Arthropoda","Insecta","Diptera","Sciaridae","","Corynoptera","Corynoptera deserta","12","12","12","41","0","0","99.09","420","0","422.92","30.34","0"
"AGAW1","05W","INS","BOLD:AAG2511","Arthropoda","Insecta","Diptera","Anthomyiidae","Anthomyiinae","Delia","Delia platura","10","10","10","42","0","0","98.63","440.2","0","442.5","33.39","0"
"AGAW1","05W","INS","BOLD:ADH4973","Arthropoda","Insecta","Diptera","Tipulidae","Tipulinae","Nephrotoma","","8","8","8","43","0","0","97.58","421","0","424.13","34.12","0"
"AGAW1","05W","INS","BOLD:AAA7374","Arthropoda","Insecta","Diptera","Syrphidae","Syrphinae","Sphaerophoria","Sphaerophoria bankowskae, Sphaerophoria infuscata, Sphaerophoria nr. philanthus, Sphaerophoria batava, Sphaerophoria sp., Sphaerophoria rueppellii, Sphaerophoria interrupta-group, Sphaerophoria sp. MA-1, Sphaerophoria cleoae, Sphaerophoria asymmetrica, Sphaerophoria fatarum, Sphaerophoria bengalensis, Sphaerophoria bifurcata, Sphaerophoria taeniata, Sphaerophoria interrupta, Sphaerophoria contigua, Sphaerophoria kaa, Sphaerophoria laurae, Sphaerophoria chongjini, Sphaerophoria boreoalpina, Sphaerophoria sp. BOLDAAA7374, Sphaerophoria macrogaster, Sphaerophoria sulphuripes, Sphaerophoria brevipilosa, Sphaerophoria philanthus, Sphaerophoria pyrrhina, Sphaerophoria abbreviata, Sphaerophoria scripta, Sphaerophoria potentillae, Sphaerophoria longipilosa, Sphaerophoria virgata","7","7","7","44","0","0","99.17","433.43","0","435.43","30.86","0"
"AGAW1","05W","INS","BOLD:AAU6595","Arthropoda","Insecta","Diptera","Sciaridae","","Corynoptera","Corynoptera perpusilla","6","6","6","45","0","0","98.76","393.33","0","396.67","35.84","0"
"AGAW1","05W","INS","BOLD:AAG2462","Arthropoda","Insecta","Diptera","Anthomyiidae","Anthomyiinae","Botanophila","Botanophila fugax","6","6","6","45","0","0","99.33","448.67","0","450","31.85","0"
"AGAW1","05W","INS","BOLD:AAF9187","Arthropoda","Insecta","Coleoptera","Curculionidae","Entiminae","Phyllobius","Phyllobius oblongus","5","5","5","47","0","0","99.51","449","0","449.8","38.95","0"
"AGAW1","05W","INS","BOLD:AAF9041","Arthropoda","Insecta","Diptera","Tipulidae","Tipulinae","Tipula","Tipula oleracea","5","5","5","47","0","0","99.15","398.8","0","400.2","34.98","0"
"AGAW1","05W","INS","BOLD:ABA6037","Arthropoda","Insecta","Hymenoptera","Pteromalidae","Pteromalinae","Dibrachys","Dibrachys cavus","5","5","5","47","0","0","99.43","422.6","0","426.4","26.36","0"
"AGAW1","05W","INS","BOLD:AAA4121","Arthropoda","Insecta","Diptera","Simuliidae","Simuliinae","Simulium","Simulium vittatum complex, Simulium vittatum, Simulium sp. BOLDAAA4121, Simulium argus, Simulium vittatum s.str.","3","3","3","50","0","0","99.14","431.33","0","433.33","33.59","0"
"AGAW1","05W","INS","BOLD:ACL2355","Arthropoda","Insecta","Diptera","Tipulidae","Tipulinae","Nephrotoma","","3","3","3","50","0","0","97.35","364.33","0","366.33","33.76","0"
"AGAW1","05W","INS","BOLD:AAN6440","Arthropoda","Insecta","Diptera","Sciaridae","","Corynoptera","","3","3","3","50","0","0","98.06","448","0","449","31.85","0"
"AGAW1","05W","INS","BOLD:AAU6630","Arthropoda","Insecta","Diptera","Fanniidae","Fanniinae","Fannia","Fannia armata","3","3","3","50","0","0","99.1","440.67","0","442.67","33.21","0"
"AGAW1","05W","INS","BOLD:AAI4303","Arthropoda","Insecta","Diptera","Chironomidae","Chironominae","Chironomus","Chironomus melanescens","2","2","2","54","0","0","97.64","444.5","0","450","34.11","0"
"AGAW1","05W","INS","BOLD:ACU7906","Arthropoda","Insecta","Diptera","Chironomidae","Orthocladiinae","","","2","2","2","54","0","0","99.86","407.5","0","421","37.21","0"
"AGAW1","05W","INS","BOLD:AAE0707","Arthropoda","Insecta","Diptera","Chironomidae","Chironominae","Microtendipes","Microtendipes pedellus","2","2","2","54","0","0","97.77","449","0","450","31.78","0"
"AGAW1","05W","INS","BOLD:ACF4734","Arthropoda","Insecta","Diptera","Syrphidae","Syrphinae","Platycheirus","Platycheirus hyperboreus","2","2","2","54","0","0","98.68","410","0","413.5","31.3","0"
"AGAW1","05W","INS","BOLD:AAD0484","Arthropoda","Insecta","Diptera","Chironomidae","","","","2","2","2","54","0","0","99.3","434","0","434","34.08","0"
"AGAW1","05W","INS","BOLD:AAN5219","Arthropoda","Insecta","Diptera","Cecidomyiidae","","","","2","2","2","54","0","0","98.56","449.5","0","450","29.44","0"
"AGAW1","05W","INS","BOLD:AAA2815","Arthropoda","Insecta","Lepidoptera","Noctuidae","Noctuinae","Mythimna","Mythimna yukonensis, Mythimna oxygala","2","2","2","54","0","0","98.73","402.5","0","405","33.09","0"
"AGAW1","05W","INS","BOLD:AAL5267","Arthropoda","Insecta","Coleoptera","Chrysomelidae","Galerucinae","Phyllotreta","Phyllotreta striolata","2","2","2","54","0","0","97.77","449","0","450","39.56","0"
"AGAW1","05W","INS","BOLD:AAG6743","Arthropoda","Insecta","Diptera","Sarcophagidae","Sarcophaginae","Sarcophaga","Sarcophaga sp. 1 MG, Sarcophaga subvicina","2","2","2","54","0","0","99.55","448.5","0","450","34.78","0"
"AGAW1","05W","INS","BOLD:ADH6353","Arthropoda","Insecta","Diptera","Tipulidae","Tipulinae","Nephrotoma","","2","2","2","54","0","0","97.55","449","0","450","34.33","0"
"AGAW1","05W","INS","BOLD:AAG4653","Arthropoda","Insecta","Diptera","Syrphidae","Syrphinae","Heringia","Heringia latitarsis, Heringia pubescens, Heringia jakutorum, Heringia rita, Heringia fulvimanus, Heringia sp. BOLDAAG4653, Heringia nr. pubescens, Heringia coxalis, Heringia larusi, Heringia vitripennis","2","2","2","54","0","0","100","450","0","450","31.33","0"
"AGAW1","05W","INS","BOLD:ABV1243","Arthropoda","Insecta","Diptera","Sarcophagidae","Paramacronychiinae, Miltogramminae","Miltogramminae_incertae_sedis, Brachicoma","Miltogramminae_incertae_sedis sp. 1 MG, Brachicoma sarcophagina","1","1","1","65","0","0","99.33","448","0","450","34.67","0"
"AGAW1","05W","INS","BOLD:ACF4733","Arthropoda","Insecta","Diptera","Syrphidae","Syrphinae","Platycheirus","Platycheirus europaeus, Platycheirus sp. BOLDACF4733, Platycheirus occultus, Platycheirus pennipes, Platycheirus angustatus2, Platycheirus angustatus3, Platycheirus angustatus","1","1","1","65","0","0","97.56","450","0","450","31.78","0"
"AGAW1","05W","INS","BOLD:AAG7279","Arthropoda","Insecta","Diptera","Sphaeroceridae","Limosininae","Spelobia","Spelobia ochripes","1","1","1","65","0","0","97.78","450","0","450","36","0"
"AGAW1","05W","INS","BOLD:AAD7145","Arthropoda","Insecta","Diptera","Muscidae","Mydaeinae","Myospila","Myospila meditabunda","1","1","1","65","0","0","99.33","449","0","450","31.56","0"
"AGAW1","05W","INS","BOLD:ABA1215","Arthropoda","Insecta","Diptera","Sciaridae","","Lycoriella","Lycoriella sativae","1","1","1","65","0","0","97.76","446","0","444","36.71","0"
"AGAW1","05W","INS","BOLD:AAY6739","Arthropoda","Insecta","Hemiptera","Cicadellidae","Deltocephalinae","Prescottia","Prescottia lobata","1","1","1","65","0","0","99.78","448","0","450","36.22","0"
"AGAW1","05W","INS","BOLD:AAA8933","Arthropoda","Insecta","Coleoptera","Coccinellidae","Coccinellinae","Coccinella","Coccinella septempunctata","1","1","1","65","0","0","98.44","448","0","450","33.33","0"
"AGAW1","05W","INS","BOLD:AAD4513","Arthropoda","Insecta","Coleoptera","Cerambycidae","Cerambycinae","Cyrtophorus","Cyrtophorus verrucosus","1","1","1","65","0","0","98.66","449","0","450","38.44","0"
"AGAW1","05W","INS","BOLD:AAG5457","Arthropoda","Insecta","Diptera","Chironomidae","Chironominae","","","1","1","1","65","0","0","99.77","426","0","430","33.49","0"
"AGAW1","05W","INS","BOLD:AAP2969","Arthropoda","Insecta","Diptera","Anthomyiidae","","","","1","1","1","65","0","0","99.11","450","0","450","33.78","0"
"AGAW1","05W","INS","BOLD:AAF9772","Arthropoda","Insecta","Diptera","Hybotidae","Tachydromiinae","Platypalpus","Platypalpus sp. 24, Platypalpus sp. BOLD:AAF9772, Platypalpus harpiger, Platypalpus major","1","1","1","65","0","0","98.67","376","0","379","34.83","0"
"AGAW1","05W","INS","BOLD:AAA3817","Arthropoda","Insecta","Lepidoptera","Geometridae","Larentiinae","Xanthorhoe","Xanthorhoe ferrugata","1","1","1","65","0","0","99.29","424","0","427","32.32","0"
"AGAW1","05W","INS","BOLD:AAB8817","Arthropoda","Insecta","Diptera","Muscidae","Azeliinae","Muscina","Muscina levida","1","1","1","65","0","0","98.19","442","0","446","32.74","0"
"AGAW1","05W","INS","BOLD:AAG0995","Arthropoda","Insecta","Diptera","Chironomidae","Orthocladiinae","Parakiefferiella","","1","1","1","65","0","0","98.66","449","0","450","40.22","0"
"AGAW1","05W","INS","BOLD:AAB9140","Arthropoda","Insecta","Diptera","Calliphoridae","Chrysomyinae","Phormia","Phormia regina","1","1","1","65","0","0","98.34","422","0","425","32.47","0"
"AGAW1","05W","INS","BOLD:AAC1312","Arthropoda","Insecta","Diptera","Syrphidae","Syrphinae","Toxomerus","Toxomerus geminatus, Toxomerus 75-5, Toxomerus rombicus, Toxomerus sp. 75-5","1","1","1","65","0","0","99.33","449","0","450","30.89","0"
"AGAW1","05W","INS","BOLD:AAU6970","Arthropoda","Insecta","Coleoptera","Cleridae","Clerinae","Enoclerus","Enoclerus nigripes","1","1","1","65","0","0","97.47","395","0","402","32.59","0"
"AGAW1","05W","INS","BOLD:AAN8770","Arthropoda","Insecta","Diptera","Psychodidae","Psychodinae","Psychoda","Psychoda trinodulosa","1","1","1","65","0","0","100","449","0","450","34","0"
"AGAW1","05W","INS","BOLD:AAA9251","Arthropoda","Insecta","Lepidoptera","Gelechiidae","Gelechiinae","Scrobipalpa","Scrobipalpa atriplicella, Scrobipalpa nr. atriplicella sp. 1","1","1","1","65","0","0","97.56","409","0","411","32.12","0"
"AGAW1","05W","INS","BOLD:AAC2498","Arthropoda","Insecta","Diptera","Muscidae","Phaoniinae","Helina","Helina evecta, Helina aldrichi","1","1","1","65","0","0","98.23","451","0","450","30.67","0"
"AGAW1","05W","INS","BOLD:ACF2821","Arthropoda","Insecta","Diptera","Mycetophilidae","Mycetophilinae","Mycetophila","Mycetophila fungorum","1","1","1","65","0","0","99.78","449","0","450","35.33","0"
"AGAW1","05W","INS","BOLD:AAD0853","Arthropoda","Insecta","Diptera","Scathophagidae","Scathophaginae","Scathophaga","Scathophaga lutaria, Scathophaga hiemalis, Scathophaga furcata, Scathophaga stercoraria, Scathophaga inquinata, Scathophaga suilla","1","1","1","65","0","0","99.55","446","0","450","32","0"
"AGAW1","05W","INS","BOLD:AAG6588","Arthropoda","Insecta","Coleoptera","Chrysomelidae","Cassidinae","Deloyala","Deloyala guttata","1","1","1","65","0","0","99.46","369","0","370","31.08","0"
"AGAW1","05W","INS","BOLD:AAQ3566","Arthropoda","Insecta","Diptera","Anthomyiidae","Pegomyinae","Pegomya","","1","1","1","65","0","0","99.25","400","0","403","31.27","0"
"AGAW1","05W","INS","BOLD:ACL6050","Arthropoda","Insecta","Trichoptera","Hydroptilidae","Orthotrichiinae","Orthotrichia","Orthotrichia cristata","1","1","1","65","0","0","97.99","349","0","350","27.43","0"
"AGAW1","05W","INS","BOLD:AAP9021","Arthropoda","Insecta","Diptera","Cecidomyiidae","","","","1","1","1","65","0","0","97.54","448","0","450","30.22","0"
"AGAW1","05W","INS","BOLD:AAN5505","Arthropoda","Insecta","Diptera","Hybotidae","Tachydromiinae","Platypalpus","Platypalpus sp. BOLD:AAN5505, Platypalpus minutus, Platypalpus nr. minutus, Platypalpus australominutus","1","1","1","65","0","0","98.22","449","0","450","37.56","0"
"AGAW1","05W","INS","BOLD:AAB2384","Arthropoda","Insecta","Diptera","Syrphidae","Syrphinae","Eupeodes","Eupeodes duseki, Eupeodes volucris, Eupeodes nielseni, Eupeodes americanus, Eupeodes flukei, Eupeodes vockerothi, Eupeodes lucasi, Eupeodes montivagus, Eupeodes rufipunctatus, Eupeodes rojasi, Eupeodes goeldlini, Eupeodes luniger, Eupeodes punctifer, Eupeodes bucculatus, Eupeodes perplexus, Eupeodes abiskoensis, Eupeodes corollae, Eupeodes pomus, Eupeodes lundbecki, Eupeodes nitens, Eupeodes sp. BOLDAAB2384, Eupeodes curtus, Eupeodes latifasciatus","1","1","1","65","0","0","98.4","375","0","378","30.95","0"
"AGAW1","05W","INS","BOLD:AAC9062","Arthropoda","Insecta","Diptera","Culicidae","Culicinae","Ochlerotatus, Aedes","Aedes fitchii, Ochlerotatus fitchii, Aedes grossbecki, Aedes ventrovittis","1","1","1","65","0","0","100","448","0","450","33.33","0"
"AGAW1","05W","INS","BOLD:AAI3059","Arthropoda","Insecta","Lepidoptera","Tineidae","Nemapogoninae","Nemapogon","Nemapogon variatella","1","1","1","65","0","0","100","449","0","450","30.22","0"
"AGAW1","05W","INS","BOLD:AAB5640","Arthropoda","Insecta","Coleoptera","Coccinellidae","Coccinellinae","Harmonia","Harmonia axyridis","1","1","1","65","0","0","100","450","0","450","32.44","0"
"AGAW1","05W","INS","BOLD:AAF4817","Arthropoda","Insecta","Diptera","Chironomidae","Orthocladiinae","Smittia","Smittia edwardsi","1","1","1","65","0","0","98.21","392","0","450","32.89","0"
"AGAW1","05W","INS","BOLD:AAC2438","Arthropoda","Insecta","Diptera","Syrphidae","Syrphinae","Paragus","Paragus haemorrhous, Paragus angustifrons","1","1","1","65","0","0","100","449","0","450","32.67","0"
"AGAW1","05W","INS","BOLD:AAE4298","Arthropoda","Insecta","Diptera","Chironomidae","Orthocladiinae","Cricotopus","Cricotopus tremulus","1","1","1","65","0","0","98.67","450","0","450","32.22","0"
"AGAW1","05W","INS","BOLD:AAA8451","Arthropoda","Insecta","Lepidoptera","Hepialidae","","Korscheltellus","Korscheltellus lupulina","1","1","1","65","0","0","98.44","450","0","450","28.67","0"
"AGAW1","05W","INS","BOLD:ADH9310","Arthropoda","Insecta","Diptera","Anthomyiidae","Anthomyiinae","Anthomyia","Anthomyia procellaris, Anthomyia pluvialis","1","1","1","65","0","0","99.78","448","0","450","32.67","0"
"AGAW1","05W","INS","BOLD:ABW1310","Arthropoda","Insecta","Diptera","Anthomyiidae","","","","1","1","1","65","0","0","98.88","447","0","450","36.67","0"
"AGAW1","05W","NAR","no match","no match","no match","no match","no match","no match","no match","no match","no match","no match","no match","no match","no match","no match","no match","no match","no match","no match","no match","no match"
"AGAW1","05W","NIN","BOLD:AAN6642","Arthropoda","Arachnida","Mesostigmata","Blattisociidae","Platyseiinae","Cheiroseius","","41","37","37","1","9.76","0","99.28","442.56","0","443.54","26.51","0"
"AGAW1","05W","NIN","BOLD:AAV0589","Arthropoda","Arachnida","Mesostigmata","Blattisociidae","Platyseiinae","Cheiroseius","","12","12","12","2","0","0","99.2","447.75","0","450","27.41","0"
"AGAW1","05W","NIN","BOLD:AAI4346","Arthropoda","Arachnida","Opiliones","Phalangiidae","","Paroligolophus, Phalangium","Paroligolophus agrestis, Phalangium opilio","2","2","2","3","0","0","98.77","448.5","0","450","36.56","0"
"AGAW1","05W","NIN","BOLD:AAA8399","Arthropoda","Arachnida","Araneae","Araneidae","","Araniella","Araniella displicata","1","1","1","4","0","0","98.27","404","0","450","37.33","0"
"AGAW1","05W","NIN","BOLD:AAN4906","Arthropoda","Collembola","Symphypleona","Bourletiellidae","","","","1","1","1","4","0","0","98.91","368","0","372","36.02","0"
"AGAW1","05W","NIN","BOLD:AAN6394","Arthropoda","Arachnida","Araneae","Anyphaenidae","","Hibana","Hibana gracilis","1","1","1","4","0","0","99.53","424","0","426","31.22","0"
"AGAW1","05W","NIN","BOLD:AAI6251","Arthropoda","Arachnida","Araneae","Dictynidae","","Emblyna","Emblyna hentzi","1","1","1","4","0","0","98.22","450","0","450","32.89","0"
"AGAW1","05W","NIN","BOLD:ACC8538","Arthropoda","Arachnida","Mesostigmata","Blattisociidae","","","","1","1","1","4","0","0","98.88","448","0","450","33.11","0"
"AGAW1","08E","BAC","TAX:91393","Proteobacteria","Alphaproteobacteria","Rickettsiales","Rickettsiaceae",NA,"Wolbachia","","38","37","37","1","2.63","0","98.97","390.74","0","422.39","39.13","0"
"AGAW1","08E","BAC","TAX:175768","Proteobacteria","Alphaproteobacteria","Rickettsiales","Rickettsiaceae",NA,"Wolbachia","Wolbachia pipientis","19","19","19","2","0","0","98.25","445.37","0","446.95","39.59","0"
"AGAW1","08E","BAC","TAX:91391","Proteobacteria","Alphaproteobacteria","Rickettsiales","",NA,"","","2","2","2","3","0","0","98.34","422","0","450","39.67","0"
"AGAW1","08E","BAC","TAX:91389","Proteobacteria","","","",NA,"","","2","2","2","3","0","0","99.17","362","0","414.5","38.17","0"
"AGAW1","08E","CHO","no match","no match","no match","no match","no match","no match","no match","no match","no match","no match","no match","no match","no match","no match","no match","no match","no match","no match","no match","no match"
"AGAW1","08E","INS","BOLD:AAA3453","Arthropoda","Insecta","Diptera","Anthomyiidae","Anthomyiinae","Delia","","14724","12405","12405","1","15.75","0","99.32","432.07","0","434.68","34.62","0"
"AGAW1","08E","INS","BOLD:AAA6292","Arthropoda","Insecta","Lepidoptera","Tortricidae","Tortricinae","Cnephasia","Cnephasia genitalana, Cnephasia stephensiana","13804","9105","9105","2","34.04","0","99.62","438.51","0","439.7","35.45","0"
"AGAW1","08E","INS","BOLD:AAP8739","Arthropoda","Insecta","Diptera","Platystomatidae","Platystomatinae","Rivellia","Rivellia steyskali","12704","8499","8499","3","33.1","0","99.25","442.12","0","444.03","37.94","0"
"AGAW1","08E","INS","BOLD:AAG2511","Arthropoda","Insecta","Diptera","Anthomyiidae","Anthomyiinae","Delia","Delia platura","12437","9912","9912","4","20.3","0","99.03","444.72","0","447.6","33.16","0"
"AGAW1","08E","INS","BOLD:AAB0373","Arthropoda","Insecta","Neuroptera","Chrysopidae","Chrysopinae","Chrysoperla","Chrysoperla sp. isolate 82, Chrysoperla zastrowi, Chrysoperla plorabunda, Chrysoperla sp. isolate 71, Chrysoperla agilis, Italochrysa italica, Chrysoperla sp. isolate 88, Chrysoperla lucasina, Chrysoperla carnea, Chrysoperla mediterranea, Chrysoperla pallida, Chrysoperla nipponensis, Chrysoperla sillemi","5059","3836","3836","5","24.17","0","99.21","434.31","0","435.24","31.63","0"
"AGAW1","08E","INS","BOLD:AAG6745","Arthropoda","Insecta","Diptera","Calliphoridae","Calliphorinae","Pollenia","Pollenia pediculata","4969","4382","4382","6","11.81","0","99.41","405.5","0","407.2","33.55","0"
"AGAW1","08E","INS","BOLD:AAD2041","Arthropoda","Insecta","Diptera","Dolichopodidae","Sciapodinae","Condylostylus","Condylostylus inermis, Condylostylus nigrofemoratus, Condylostylus connectans, Condylostylus caudatus, Condylostylus flavipes","3894","2764","2764","7","29.02","0","99.5","439.66","0","440.86","35.99","0"
"AGAW1","08E","INS","BOLD:AAA7374","Arthropoda","Insecta","Diptera","Syrphidae","Syrphinae","Sphaerophoria","Sphaerophoria bankowskae, Sphaerophoria infuscata, Sphaerophoria nr. philanthus, Sphaerophoria batava, Sphaerophoria sp., Sphaerophoria rueppellii, Sphaerophoria interrupta-group, Sphaerophoria sp. MA-1, Sphaerophoria cleoae, Sphaerophoria asymmetrica, Sphaerophoria fatarum, Sphaerophoria bengalensis, Sphaerophoria bifurcata, Sphaerophoria taeniata, Sphaerophoria interrupta, Sphaerophoria contigua, Sphaerophoria kaa, Sphaerophoria laurae, Sphaerophoria chongjini, Sphaerophoria boreoalpina, Sphaerophoria sp. BOLDAAA7374, Sphaerophoria macrogaster, Sphaerophoria sulphuripes, Sphaerophoria brevipilosa, Sphaerophoria philanthus, Sphaerophoria pyrrhina, Sphaerophoria abbreviata, Sphaerophoria scripta, Sphaerophoria potentillae, Sphaerophoria longipilosa, Sphaerophoria virgata","1941","1673","1673","8","13.81","0","99.32","428.51","0","431.68","30.73","0"
"AGAW1","08E","INS","BOLD:AAA2815","Arthropoda","Insecta","Lepidoptera","Noctuidae","Noctuinae","Mythimna","Mythimna yukonensis, Mythimna oxygala","1696","1549","1549","9","8.67","0","98.93","426.14","0","428.28","33.17","0"
"AGAW1","08E","INS","BOLD:AAG2123","Arthropoda","Insecta","Diptera","Tachinidae","Dexiinae","Dinera","Dinera grisescens","815","649","649","10","20.37","0","99.32","434.48","0","436.39","31.9","0"
"AGAW1","08E","INS","BOLD:AAG9626","Arthropoda","Insecta","Diptera","Dolichopodidae","","","","689","566","566","11","17.85","0","98.32","441.71","0","443.53","36.26","0"
"AGAW1","08E","INS","BOLD:AAA5299","Arthropoda","Insecta","Diptera","Chironomidae","Orthocladiinae","Cricotopus","Cricotopus laetus, Cricotopus relucens, Cricotopus trifasciatus, Cricotopus sp., Cricotopus sp. 3ES, Cricotopus sylvestris, Cricotopus glacialis","621","573","573","12","7.73","0","99.21","433.87","0","436.29","34.68","0"
"AGAW1","08E","INS","BOLD:AAG2868","Arthropoda","Insecta","Hemiptera","Cicadellidae","Typhlocybinae","Empoasca","Empoasca sp., Empoasca fabae","612","395","395","13","35.46","0","99.69","443.46","0","447.45","32.46","0"
"AGAW1","08E","INS","BOLD:AAH4171","Arthropoda","Insecta","Diptera","Chloropidae","Oscinellinae","Apallates","Apallates montanus, Apallates particeps","472","408","408","14","13.56","0","99.13","436.7","0","438.42","32.75","0"
"AGAW1","08E","INS","BOLD:AAN5199","Arthropoda","Insecta","Diptera","Cecidomyiidae","","","CecidInt20 sp. BOLD:AAN5199","413","315","315","15","23.73","0","99.58","435.01","0","446.22","26.72","0"
"AGAW1","08E","INS","BOLD:AAG2898","Arthropoda","Insecta","Hemiptera","Cicadellidae","Deltocephalinae","Paraphlepsius","Paraphlepsius apertus, Paraphlepsius irroratus","377","289","289","16","23.34","0","99.56","444.53","0","445.36","31.91","0"
"AGAW1","08E","INS","BOLD:AAG7276","Arthropoda","Insecta","Diptera","Sphaeroceridae","Limosininae","Leptocera","Leptocera erythrocera","337","305","305","17","9.5","0","99.2","432.89","0","435.97","32.89","0"
"AGAW1","08E","INS","BOLD:AAH4208","Arthropoda","Insecta","Diptera","Chloropidae","Oscinellinae","Elachiptera","Elachiptera erythropleura, Elachiptera costata","327","267","267","18","18.35","0","98.86","440.69","0","442.44","31.37","0"
"AGAW1","08E","INS","BOLD:AAF9771","Arthropoda","Insecta","Diptera","Hybotidae","Tachydromiinae","Platypalpus","Platypalpus longiseta, Platypalpus trivialis, Platypalpus sp. 5, Platypalpus sp. 17, Platypalpus pallidiventris","324","257","257","19","20.68","0","99.33","432.54","0","433.59","35","0"
"AGAW1","08E","INS","BOLD:AAN5342","Arthropoda","Insecta","Diptera","Chironomidae","Orthocladiinae","Heleniella","","215","210","210","20","2.33","0","99.01","415.48","0","423.64","35.03","0"
"AGAW1","08E","INS","BOLD:AAP5169","Arthropoda","Insecta","Diptera","Chloropidae","Oscinellinae","Elachiptera","Elachiptera nigriceps","211","169","169","21","19.91","0","99.5","429.45","0","430.49","31.43","0"
"AGAW1","08E","INS","BOLD:AAN5225","Arthropoda","Insecta","Diptera","Cecidomyiidae","","","","182","150","150","22","17.58","0","99.03","447.27","0","448.3","28.26","0"
"AGAW1","08E","INS","BOLD:AAA5803","Arthropoda","Insecta","Hemiptera","Miridae","Mirinae","Lygus","Lygus lineolaris","166","118","118","23","28.92","0","98.64","447.29","0","448.72","33.92","0"
"AGAW1","08E","INS","BOLD:AAG7279","Arthropoda","Insecta","Diptera","Sphaeroceridae","Limosininae","Spelobia","Spelobia ochripes","157","111","111","24","29.3","0","98.75","444.08","0","446.11","35.46","0"
"AGAW1","08E","INS","BOLD:AAG7891","Arthropoda","Insecta","Hymenoptera","Platygastridae","Scelioninae","Telenomus","Telenomus podisi","153","145","145","25","5.23","0","99.08","425.37","0","433.93","26.96","0"
"AGAW1","08E","INS","BOLD:ABW1379","Arthropoda","Insecta","Diptera","Chloropidae","Oscinellinae","Malloewia","Malloewia abdominalis","149","119","119","26","20.13","0","98.24","443.4","0","448.03","31.42","0"
"AGAW1","08E","INS","BOLD:AAA5300","Arthropoda","Insecta","Diptera","Chironomidae","Orthocladiinae","Cricotopus","Cricotopus tibialis","139","139","139","27","0","0","99.23","410.37","0","412.42","36.82","0"
"AGAW1","08E","INS","BOLD:AAA7041","Arthropoda","Insecta","Trichoptera","Limnephilidae","Limnephilinae","Limnephilus","Limnephilus sericeus, Limnephilus abbreviatus, Limnephilus fagus, Limnephilus sp., Limnephilus DERaa084","117","115","115","28","1.71","0","98.9","403.54","0","408.68","34.89","0"
"AGAW1","08E","INS","BOLD:AAH3661","Arthropoda","Insecta","Diptera","Cecidomyiidae","","","","117","110","110","28","5.98","0","99.19","431.82","0","433.35","25.42","0"
"AGAW1","08E","INS","BOLD:ACI4401","Arthropoda","Insecta","Diptera","Chironomidae","Orthocladiinae","Pseudorthocladius","","115","103","103","30","10.43","0","98.39","436.08","0","437.9","36.96","0"
"AGAW1","08E","INS","BOLD:AAN5660","Arthropoda","Insecta","Diptera","Chloropidae","Oscinellinae","Incertella","Incertella minor","113","91","91","31","19.47","0","99.6","447.87","0","449.06","32.59","0"
"AGAW1","08E","INS","BOLD:AAG9668","Arthropoda","Insecta","Diptera","Dolichopodidae","Diaphorinae","Chrysotus","Chrysotus sp., Chrysotus sp. nr. obliquus","106","93","93","32","12.26","0","98.96","445.74","0","448.2","32.86","0"
"AGAW1","08E","INS","BOLD:AAN6154","Arthropoda","Insecta","Coleoptera","Latridiidae","Corticariinae","Melanophthalma","Melanophthalma inermis","106","95","95","32","10.38","0","98.97","442","0","443.48","31.39","0"
"AGAW1","08E","INS","BOLD:AAA7661","Arthropoda","Insecta","Diptera","Culicidae","Culicinae","Culiseta, Culex","Culiseta melanura, Culex articularis, Culex restuans, Culex acharistus","103","94","94","34","8.74","0","99.34","433.5","0","435.38","32.97","0"
"AGAW1","08E","INS","BOLD:AAN5238","Arthropoda","Insecta","Diptera","Cecidomyiidae","","","","99","89","89","35","10.1","0","99.36","435.25","0","436.62","25.71","0"
"AGAW1","08E","INS","BOLD:ACU7906","Arthropoda","Insecta","Diptera","Chironomidae","Orthocladiinae","","","99","99","99","35","0","0","98.89","408.51","0","411.25","35.91","0"
"AGAW1","08E","INS","BOLD:ACH4728","Arthropoda","Insecta","Psocodea","Elipsocidae","Elipsocinae","Cuneopalpus","Cuneopalpus cyanops","97","85","85","37","12.37","0","99.47","420.58","0","422.16","38.82","0"
"AGAW1","08E","INS","BOLD:AAC4197","Arthropoda","Insecta","Diptera","Chironomidae","Orthocladiinae","Paraphaenocladius","Paraphaenocladius impensus","96","77","77","38","19.79","0","99.33","436.58","0","437.66","34.86","0"
"AGAW1","08E","INS","BOLD:ACE0554","Arthropoda","Insecta","Diptera","Phoridae","Metopininae","Megaselia","","82","72","72","39","12.2","0","99.68","439.65","0","441.59","31.96","0"
"AGAW1","08E","INS","BOLD:AAG3236","Arthropoda","Insecta","Diptera","Phoridae","Phorinae","Conicera, Diplonevra","Diplonevra nitidula","81","74","74","40","8.64","0","99.5","445.48","0","447.65","32.03","0"
"AGAW1","08E","INS","BOLD:AAN5267","Arthropoda","Insecta","Diptera","Cecidomyiidae","","","","81","63","63","40","22.22","0","99.49","448.09","0","449.68","30.62","0"
"AGAW1","08E","INS","BOLD:ACK7314","Arthropoda","Insecta","Diptera","Chironomidae","Chironominae","Polypedilum","","78","77","77","42","1.28","0","98.88","404.62","0","408.28","35.23","0"
"AGAW1","08E","INS","BOLD:ADF6661","Arthropoda","Insecta","Diptera","Cecidomyiidae","","","","73","71","71","43","2.74","0","98.98","415.41","0","441.14","29.31","0"
"AGAW1","08E","INS","BOLD:ABW7897","Arthropoda","Insecta","Diptera","Cecidomyiidae","","","","67","53","53","44","20.9","0","99.65","444.39","0","446.4","29.67","0"
"AGAW1","08E","INS","BOLD:ACG9026","Arthropoda","Insecta","Diptera","Chironomidae","Chironominae","Tanytarsus","","66","65","65","45","1.52","0","98.07","408.45","0","409.74","36.08","0"
"AGAW1","08E","INS","BOLD:AAH3949","Arthropoda","Insecta","Diptera","Sciaridae","","Pseudolycoriella","","63","54","54","46","14.29","0","99.64","435.52","0","440.6","35.91","0"
"AGAW1","08E","INS","BOLD:AAC4491","Arthropoda","Insecta","Diptera","Tipulidae","Tipulinae","Tipula","Tipula morrisoni, Tipula bicornis, Tipula johnsoniana, Tipula bicolor","57","56","56","47","1.75","0","98.67","433.07","0","433.4","34","0"
"AGAW1","08E","INS","BOLD:ABU5525","Arthropoda","Insecta","Diptera","Chironomidae","Orthocladiinae","Limnophyes","Limnophyes sp. 14ES","57","57","57","47","0","0","99.09","408","0","410.89","37.15","0"
"AGAW1","08E","INS","BOLD:ACV2677","Arthropoda","Insecta","Diptera","Cecidomyiidae","","","","56","33","33","49","41.07","0","97.74","449.23","0","449.57","29.72","0"
"AGAW1","08E","INS","BOLD:AAM6655","Arthropoda","Insecta","Diptera","Hybotidae","Tachydromiinae","Platypalpus","Platypalpus stabilis","53","42","42","50","20.75","0","99.43","447.77","0","448.43","35.02","0"
"AGAW1","08E","INS","BOLD:AAP6465","Arthropoda","Insecta","Diptera","Sciaridae","","Corynoptera","Corynoptera cursor","51","36","36","51","29.41","0","98.68","445.59","0","446.37","32.33","0"
"AGAW1","08E","INS","BOLD:ABA7871","Arthropoda","Insecta","Diptera","Cecidomyiidae","","","","47","45","45","52","4.26","0","99.03","432.89","0","434.17","26.55","0"
"AGAW1","08E","INS","BOLD:AAL4232","Arthropoda","Insecta","Diptera","Agromyzidae","Phytomyzinae","Phytoliriomyza","Phytoliriomyza arctica","46","43","43","53","6.52","0","99.48","442.7","0","443.72","32.13","0"
"AGAW1","08E","INS","BOLD:ACF6513","Arthropoda","Insecta","Diptera","Psychodidae","Psychodinae","Psychoda","Psychoda cinerea","35","33","33","54","5.71","0","99.52","426.4","0","427.89","30.66","0"
"AGAW1","08E","INS","BOLD:AAV5796","Arthropoda","Insecta","Diptera","Cecidomyiidae","","","","35","34","34","54","2.86","0","99.12","445.37","0","446.89","29.12","0"
"AGAW1","08E","INS","BOLD:AAU6592","Arthropoda","Insecta","Diptera","Ceratopogonidae","Dasyheleinae","Dasyhelea","Dasyhelea arenivaga","32","32","32","56","0","0","98.9","405.75","0","410.81","34.85","0"
"AGAW1","08E","INS","BOLD:ABA0863","Arthropoda","Insecta","Diptera","Cecidomyiidae","","","","30","25","25","57","16.67","0","99.44","443.97","0","445.5","29.87","0"
"AGAW1","08E","INS","BOLD:AAN5332","Arthropoda","Insecta","Diptera","Chironomidae","Chironominae","Polypedilum","","30","30","30","57","0","0","98.04","437.07","0","438.67","34.5","0"
"AGAW1","08E","INS","BOLD:ACA6930","Arthropoda","Insecta","Diptera","Cecidomyiidae","","","","28","28","28","59","0","0","98.44","418.57","0","432.32","27.6","0"
"AGAW1","08E","INS","BOLD:AAG2740","Arthropoda","Insecta","Diptera","Ephydridae","Parydrinae","Philygria","Philygria debilis","26","26","26","60","0","0","98.33","397.31","0","400.23","31.9","0"
"AGAW1","08E","INS","BOLD:ACK3402","Arthropoda","Insecta","Coleoptera","Mordellidae","Mordellinae","Mordellistena","","25","22","22","61","12","0","99.72","434.24","0","434.8","39.92","0"
"AGAW1","08E","INS","BOLD:AAP5921","Arthropoda","Insecta","Diptera","Chironomidae","Orthocladiinae","Cricotopus","","24","22","22","62","8.33","0","98.19","445.04","0","446.08","37.65","0"
"AGAW1","08E","INS","BOLD:AAG1711","Arthropoda","Insecta","Diptera","Muscidae","Phaoniinae","Helina","Helina depuncta","22","22","22","63","0","0","99.02","428.64","0","433.95","31.32","0"
"AGAW1","08E","INS","BOLD:AAG3154","Arthropoda","Insecta","Diptera","Psychodidae","Psychodinae","Psychoda","","22","22","22","63","0","0","97.84","435.82","0","445.23","32.77","0"
"AGAW1","08E","INS","BOLD:AAD1396","Arthropoda","Insecta","Diptera","Chironomidae","Chironominae","Polypedilum","Polypedilum convictum","22","21","21","63","4.55","0","98.91","449.18","0","450","33.79","0"
"AGAW1","08E","INS","BOLD:AAH0134","Arthropoda","Insecta","Coleoptera","Phalacridae","Phalacrinae","Stilbus","Stilbus apicialis","22","20","20","63","9.09","0","99.42","429.32","0","430.55","32.12","0"
"AGAW1","08E","INS","BOLD:ACS8354","Arthropoda","Insecta","Diptera","Milichiidae","","","","21","21","21","67","0","0","98.41","442.9","0","444","30.57","0"
"AGAW1","08E","INS","BOLD:ABA0811","Arthropoda","Insecta","Diptera","Cecidomyiidae","","","","21","19","19","67","9.52","0","99.29","438.24","0","439.57","26.44","0"
"AGAW1","08E","INS","BOLD:ACJ6984","Arthropoda","Insecta","Hymenoptera","Platygastridae","Scelioninae","Trissolcus","Trissolcus thyantae","21","19","19","67","9.52","0","99.43","426.9","0","434.81","26.98","0"
"AGAW1","08E","INS","BOLD:ABV1227","Arthropoda","Insecta","Diptera","Chironomidae","Orthocladiinae","","","21","21","21","67","0","0","98.25","389","0","391","34.28","0"
"AGAW1","08E","INS","BOLD:AAI1474","Arthropoda","Insecta","Diptera","Tachinidae","Dexiinae","Periscepsia","Periscepsia laevigata, Periscepsia helymus","19","19","19","71","0","0","99.54","443.58","0","445.84","31.75","0"
"AGAW1","08E","INS","BOLD:AAL7359","Arthropoda","Insecta","Diptera","Chironomidae","Chironominae","Tanytarsus","","17","16","16","72","5.88","0","99.42","441.59","0","443.24","36.2","0"
"AGAW1","08E","INS","BOLD:AAM9255","Arthropoda","Insecta","Diptera","Sciaridae","","Pseudolycoriella","","17","17","17","72","0","0","99.09","403.82","0","406.53","30.65","0"
"AGAW1","08E","INS","BOLD:AAG6934","Arthropoda","Insecta","Diptera","Hybotidae","","","","16","16","16","74","0","0","99.2","435.13","0","436.75","38.8","0"
"AGAW1","08E","INS","BOLD:AAN5183","Arthropoda","Insecta","Diptera","Cecidomyiidae","","","","16","16","16","74","0","0","98.52","445.31","0","446.44","27.85","0"
"AGAW1","08E","INS","BOLD:AAP6790","Arthropoda","Insecta","Diptera","Chloropidae","Oscinellinae","Incertella","Incertella incerta","15","15","15","76","0","0","99.24","440.47","0","441.6","31.48","0"
"AGAW1","08E","INS","BOLD:AAG9897","Arthropoda","Insecta","Dermaptera","Forficulidae","Forficulinae","Forficula","Forficula auricularia-A","13","13","13","77","0","0","98.92","384.08","0","390.92","45.81","0"
"AGAW1","08E","INS","BOLD:ACZ2431","Arthropoda","Insecta","Coleoptera","Staphylinidae","","","","12","9","9","78","25","0","97.63","443.58","0","443.92","33.27","0"
"AGAW1","08E","INS","BOLD:AAM6040","Arthropoda","Insecta","Diptera","Cecidomyiidae","","","","12","11","11","78","8.33","0","99.18","448.42","0","448.92","27.16","0"
"AGAW1","08E","INS","BOLD:AAC0903","Arthropoda","Insecta","Diptera","Chironomidae","Chironominae","Chironomus","Chironomus acidophilus","12","12","12","78","0","0","99.14","441.83","0","442.75","36.47","0"
"AGAW1","08E","INS","BOLD:AAG6460","Arthropoda","Insecta","Diptera","Cecidomyiidae","","","","12","12","12","78","0","0","99.1","434.25","0","435.92","34.87","0"
"AGAW1","08E","INS","BOLD:AAG8839","Arthropoda","Insecta","Hemiptera","Cicadellidae","Deltocephalinae","Errastunus, Latalus","Latalus ocellaris, Errastunus ocellaris","11","9","9","82","18.18","0","99.82","448.55","0","450","35.98","0"
"AGAW1","08E","INS","BOLD:AAV5573","Arthropoda","Insecta","Diptera","Cecidomyiidae","","","","11","11","11","82","0","0","98.83","428.09","0","429.18","27.2","0"
"AGAW1","08E","INS","BOLD:ACN2537","Arthropoda","Insecta","Diptera","Chironomidae","Orthocladiinae","Cricotopus","","10","10","10","84","0","0","97.91","396.7","0","444.3","37.84","0"
"AGAW1","08E","INS","BOLD:AAN5242","Arthropoda","Insecta","Diptera","Cecidomyiidae","","","","9","9","9","85","0","0","99.01","393.67","0","395.56","27.71","0"
"AGAW1","08E","INS","BOLD:AAV1261","Arthropoda","Insecta","Diptera","Sciaridae","","Bradysia","Bradysia splendida","9","9","9","85","0","0","99.42","433.56","0","435.11","34.07","0"
"AGAW1","08E","INS","BOLD:AAN5220","Arthropoda","Insecta","Diptera","Cecidomyiidae","","","","8","8","8","87","0","0","98.69","413.88","0","425.13","25.63","0"
"AGAW1","08E","INS","BOLD:AAF1013","Arthropoda","Insecta","Diptera","Chironomidae","Chironominae","Tanytarsus","Tanytarsus allicis","8","8","8","87","0","0","97.88","424.63","0","431.75","36.27","0"
"AGAW1","08E","INS","BOLD:AAD1485","Arthropoda","Insecta","Diptera","Chironomidae","Chironominae","Paratanytarsus","Paratanytarsus grimmii, Paratanytarsus sp.","8","8","8","87","0","0","99.56","445.5","0","446.63","33.83","0"
"AGAW1","08E","INS","BOLD:ADH9867","Arthropoda","Insecta","Diptera","","","","","8","8","8","87","0","0","98.98","394.25","0","397.63","37.16","0"
"AGAW1","08E","INS","BOLD:ACK9435","Arthropoda","Insecta","Hemiptera","Cicadellidae","Macropsinae","Macropsis","Macropsis fumipennis","6","6","6","91","0","0","99.03","448.17","0","450","38","0"
"AGAW1","08E","INS","BOLD:AAG8091","Arthropoda","Insecta","Hymenoptera","Ichneumonidae","Campopleginae","Campoletis","Campoletis sp. AB.2","6","6","6","91","0","0","99.44","420.33","0","423.17","24.5","0"
"AGAW1","08E","INS","BOLD:ABV1337","Arthropoda","Insecta","Diptera","Cecidomyiidae","","","","6","6","6","91","0","0","98.94","432.67","0","434.17","28.42","0"
"AGAW1","08E","INS","BOLD:AAM6184","Arthropoda","Insecta","Diptera","Ceratopogonidae","Ceratopogoninae","Culicoides","","6","6","6","91","0","0","97.7","411","0","413.33","37.28","0"
"AGAW1","08E","INS","BOLD:ACJ6954","Arthropoda","Insecta","Diptera","Cecidomyiidae","","","","5","5","5","95","0","0","99.82","338.2","0","429.8","26.5","0"
"AGAW1","08E","INS","BOLD:AAA2632","Arthropoda","Insecta","Lepidoptera","Noctuidae","Noctuinae","Noctua","Noctua atlantica, Noctua pronuba","5","5","5","95","0","0","99.15","448.6","0","450","31.42","0"
"AGAW1","08E","INS","BOLD:AAM7855","Arthropoda","Insecta","Thysanoptera","Thripidae","Thripinae","Frankliniella","Frankliniella tritici","4","4","4","97","0","0","99.61","448","0","450","31.5","0"
"AGAW1","08E","INS","BOLD:AAN8770","Arthropoda","Insecta","Diptera","Psychodidae","Psychodinae","Psychoda","Psychoda trinodulosa","4","4","4","97","0","0","98.72","449","0","450","34.28","0"
"AGAW1","08E","INS","BOLD:AAH3148","Arthropoda","Insecta","Hymenoptera","","","","","3","3","3","99","0","0","99.18","449.33","0","450","28.59","0"
"AGAW1","08E","INS","BOLD:AAN5255","Arthropoda","Insecta","Diptera","Cecidomyiidae","Cecidomyiinae","Dasineura","","3","3","3","99","0","0","98.51","448.67","0","450","26.89","0"
"AGAW1","08E","INS","BOLD:ACE7444","Arthropoda","Insecta","Hemiptera","Miridae","Mirinae, Phylinae","Adelphocoris, Megalocoleus","Adelphocoris lineolatus, Megalocoleus molliculus","3","3","3","99","0","0","99.63","450","0","450","34.89","0"
"AGAW1","08E","INS","BOLD:ACA7694","Arthropoda","Insecta","Diptera","Chironomidae","Chironominae","","","3","3","3","99","0","0","98.5","411.67","0","412.67","32.86","0"
"AGAW1","08E","INS","BOLD:AAB4321","Arthropoda","Insecta","Lepidoptera","Gelechiidae","Gelechiinae","Metzneria","Metzneria lappella","3","3","3","99","0","0","98.97","419.67","0","430.33","31.82","0"
"AGAW1","08E","INS","BOLD:AAM6037","Arthropoda","Insecta","Diptera","Cecidomyiidae","Cecidomyiinae","Contarinia, Dasineura","","3","3","3","99","0","0","98.62","406.33","0","409.67","25.76","0"
"AGAW1","08E","INS","BOLD:AAH3947","Arthropoda","Insecta","Diptera","Sciaridae","","Bradysia","Bradysia trivittata","3","3","3","99","0","0","97.45","444","0","446.33","35.1","0"
"AGAW1","08E","INS","BOLD:AAA9422","Arthropoda","Insecta","Hemiptera","Cicadellidae","Deltocephalinae","Macrosteles","Macrosteles quadrilineatus","3","3","3","99","0","0","98.96","416.33","0","447.33","36.74","0"
"AGAW1","08E","INS","BOLD:AAA3181","Arthropoda","Insecta","Diptera","Muscidae","Muscinae","Stomoxys","Stomoxys calcitrans","2","2","2","107","0","0","98.99","445.5","0","450","32.67","0"
"AGAW1","08E","INS","BOLD:AAH4135","Arthropoda","Insecta","Diptera","Chloropidae","Chloropinae","Thaumatomyia","Thaumatomyia glabra, Thaumatomyia grata, Thaumatomyia sp. 1 TAW","2","2","2","107","0","0","99.65","404","0","409","31.39","0"
"AGAW1","08E","INS","BOLD:AAA4277","Arthropoda","Insecta","Diptera","Syrphidae","Syrphinae","Toxomerus","Toxomerus marginatus, Toxomerus sp. CR-3, Toxomerus CR-3","2","2","2","107","0","0","99.44","447.5","0","450","31","0"
"AGAW1","08E","INS","BOLD:AAA3889","Arthropoda","Insecta","Lepidoptera","Crambidae","Crambinae","Parapediasia","Parapediasia teterrellus","2","2","2","107","0","0","98.99","398.5","0","401.5","31.94","0"
"AGAW1","08E","INS","BOLD:AAG2460","Arthropoda","Insecta","Diptera","Anthomyiidae","Anthomyiinae","Hydrophoria","Hydrophoria lancifer","2","2","2","107","0","0","97.65","447.5","0","450","33.78","0"
"AGAW1","08E","INS","BOLD:AAD0484","Arthropoda","Insecta","Diptera","Chironomidae","","","","2","2","2","107","0","0","98.73","433.5","0","434","33.97","0"
"AGAW1","08E","INS","BOLD:AAE0707","Arthropoda","Insecta","Diptera","Chironomidae","Chironominae","Microtendipes","Microtendipes pedellus","2","2","2","107","0","0","97.88","404","0","408","30.58","0"
"AGAW1","08E","INS","BOLD:AAF9187","Arthropoda","Insecta","Coleoptera","Curculionidae","Entiminae","Phyllobius","Phyllobius oblongus","2","2","2","107","0","0","99.78","425.5","0","428.5","38.82","0"
"AGAW1","08E","INS","BOLD:ACP6694","Arthropoda","Insecta","Diptera","Chironomidae","Orthocladiinae","","","2","2","2","107","0","0","98.26","418","0","420.5","33.53","0"
"AGAW1","08E","INS","BOLD:AAU7339","Arthropoda","Insecta","Coleoptera","Throscidae","Throscinae","Aulonothroscus","Aulonothroscus constrictor","2","2","2","107","0","0","98.27","435.5","0","435.5","36.5","0"
"AGAW1","08E","INS","BOLD:AAA3867","Arthropoda","Insecta","Lepidoptera","Noctuidae","Noctuinae","Apamea","Apamea amputatrix","2","2","2","107","0","0","98.99","447.5","0","450","33.78","0"
"AGAW1","08E","INS","BOLD:ACF7023","Arthropoda","Insecta","Diptera","Dolichopodidae","","","","2","2","2","107","0","0","97.36","227","0","450","32.78","0"
"AGAW1","08E","INS","BOLD:AAE0845","Arthropoda","Insecta","Hemiptera","Pentatomidae","Pentatominae","Euschistus","Euschistus servus","2","2","2","107","0","0","99.16","423.5","0","424.5","37.69","0"
"AGAW1","08E","INS","BOLD:AAB3656","Arthropoda","Insecta","Lepidoptera","Pterophoridae","Pterophorinae","Hellinsia","Hellinsia glenni, Hellinsia lacteodactylus, Hellinsia paleaceus","2","2","2","107","0","0","97.73","441.5","0","450","33.33","0"
"AGAW1","08E","INS","BOLD:ABV9293","Arthropoda","Insecta","Diptera","Chironomidae","Chironominae","Dicrotendipes","","2","2","2","107","0","0","99.86","356.5","0","438.5","35","0"
"AGAW1","08E","INS","BOLD:AAA5953","Arthropoda","Insecta","Lepidoptera","Gelechiidae","Gelechiinae","Coleotechnites","Coleotechnites sp., Coleotechnites piceaella, Coleotechnites laricis, Coleotechnites sp. 3, Coleotechnites carbonaria, Coleotechnites granti, Coleotechnites macleodi, Coleotechnites gibsonella, Coleotechnites coniferella, Coleotechnites martini","2","2","2","107","0","0","98.21","447.5","0","450","34","0"
"AGAW1","08E","INS","BOLD:AAG1717","Arthropoda","Insecta","Diptera","Muscidae","Phaoniinae","Helina","Helina troene","1","1","1","123","0","0","100","450","0","450","31.11","0"
"AGAW1","08E","INS","BOLD:AAN5497","Arthropoda","Insecta","Diptera","Anthomyiidae","Pegomyinae","Pegomya","Pegomya solennis, Pegomya bicolor","1","1","1","123","0","0","97.78","450","0","450","30","0"
"AGAW1","08E","INS","BOLD:AAH7146","Arthropoda","Insecta","Diptera","Sarcophagidae","Sarcophaginae","Boettcheria","Boettcheria latisterna","1","1","1","123","0","0","98.66","447","0","450","34.67","0"
"AGAW1","08E","INS","BOLD:AAM6229","Arthropoda","Insecta","Diptera","Chironomidae","Tanypodinae","","","1","1","1","123","0","0","98.31","414","0","415","32.53","0"
"AGAW1","08E","INS","BOLD:ABZ7164","Arthropoda","Insecta","Diptera","Cecidomyiidae","","","","1","1","1","123","0","0","97.96","392","0","392","25.51","0"
"AGAW1","08E","INS","BOLD:ACE5438","Arthropoda","Insecta","Diptera","Cecidomyiidae","","","","1","1","1","123","0","0","97.33","449","0","450","26","0"
"AGAW1","08E","INS","BOLD:ADF1156","Arthropoda","Insecta","Diptera","Chironomidae","Orthocladiinae","Cricotopus","","1","1","1","123","0","0","99.33","450","0","450","33.78","0"
"AGAW1","08E","INS","BOLD:AAB4657","Arthropoda","Insecta","Diptera","Chironomidae","Chironominae","Chironomus","Chironomus maturus","1","1","1","123","0","0","99.56","450","0","450","37.56","0"
"AGAW1","08E","INS","BOLD:ABA8103","Arthropoda","Insecta","Hemiptera","Cicadellidae","Typhlocybinae","Empoasca","","1","1","1","123","0","0","97.61","376","0","381","32.55","0"
"AGAW1","08E","INS","BOLD:AAG0734","Arthropoda","Insecta","Thysanoptera","Thripidae","Thripinae","Frankliniella","Frankliniella tritici","1","1","1","123","0","0","98.88","448","0","450","29.56","0"
"AGAW1","08E","INS","BOLD:AAU6681","Arthropoda","Insecta","Diptera","Tipulidae","Tipulinae","Nephrotoma","Nephrotoma cornicina","1","1","1","123","0","0","99.78","449","0","450","36.89","0"
"AGAW1","08E","INS","BOLD:AAQ0275","Arthropoda","Insecta","Diptera","Sciaridae","","Bradysia","Bradysia lembkei","1","1","1","123","0","0","99.53","423","0","428","32.71","0"
"AGAW1","08E","INS","BOLD:AAL7819","Arthropoda","Insecta","Diptera","Psychodidae","Psychodinae","Psychoda","Psychoda sp., Psychoda sp. BOLD:AAL7819","1","1","1","123","0","0","98.18","385","0","387","32.04","0"
"AGAW1","08E","INS","BOLD:AAV5788","Arthropoda","Insecta","Diptera","Cecidomyiidae","Cecidomyiinae","Asphondylia","","1","1","1","123","0","0","100","435","0","437","24.49","0"
"AGAW1","08E","INS","BOLD:AAA2482","Arthropoda","Insecta","Lepidoptera","Noctuidae","Noctuinae","Mythimna","Mythimna unipuncta","1","1","1","123","0","0","100","384","0","387","29.46","0"
"AGAW1","08E","INS","BOLD:AAA8796","Arthropoda","Insecta","Lepidoptera","Tortricidae","Tortricinae","Acleris","Acleris forsskaleana","1","1","1","123","0","0","99.73","375","0","376","32.45","0"
"AGAW1","08E","INS","BOLD:AAH3664","Arthropoda","Insecta","Diptera","Cecidomyiidae","","","CecidInt3 sp. BOLD:AAH3664","1","1","1","123","0","0","97.11","450","0","450","28.44","0"
"AGAW1","08E","INS","BOLD:AAI4303","Arthropoda","Insecta","Diptera","Chironomidae","Chironominae","Chironomus","Chironomus melanescens","1","1","1","123","0","0","98.2","445","0","450","33.11","0"
"AGAW1","08E","INS","BOLD:ABX6186","Arthropoda","Insecta","Diptera","Tipulidae","Tipulinae","Nephrotoma","Nephrotoma ferruginea","1","1","1","123","0","0","99.33","447","0","450","34","0"
"AGAW1","08E","INS","BOLD:AAA3654","Arthropoda","Insecta","Lepidoptera","Crambidae","Crambinae","Crambus","Crambus bigelovi, Crambus johnsoni, Crambus praefectellus","1","1","1","123","0","0","98.89","449","0","450","31.33","0"
"AGAW1","08E","INS","BOLD:AAZ3286","Arthropoda","Insecta","Hymenoptera","Platygastridae","","","","1","1","1","123","0","0","97.25","436","0","438","28.31","0"
"AGAW1","08E","INS","BOLD:ABX0893","Arthropoda","Insecta","Diptera","Cecidomyiidae","","","","1","1","1","123","0","0","97.3","444","0","450","26.44","0"
"AGAW1","08E","INS","BOLD:AAD8086","Arthropoda","Insecta","Lepidoptera","Elachistidae","Elachistinae","Elachista","Elachista griseicornis, Elachista acenteta","1","1","1","123","0","0","99.11","448","0","450","30.89","0"
"AGAW1","08E","INS","BOLD:ABX0657","Arthropoda","Insecta","Coleoptera","Anthicidae","Notoxinae","Notoxus","Notoxus desertus","1","1","1","123","0","0","98.89","450","0","450","36.44","0"
"AGAW1","08E","INS","BOLD:AAA1513","Arthropoda","Insecta","Lepidoptera","Plutellidae","Plutellinae","Plutella, Rhigognostis","Plutella xylostella, Rhigognostis sp.","1","1","1","123","0","0","97.11","450","0","450","31.78","0"
"AGAW1","08E","INS","BOLD:ACE4701","Arthropoda","Insecta","Lepidoptera","Crambidae","Crambinae","Crambus","Crambus multilinellus","1","1","1","123","0","0","100","450","0","450","31.78","0"
"AGAW1","08E","INS","BOLD:ABA0772","Arthropoda","Insecta","Diptera","Chironomidae","Chironominae","Polypedilum","","1","1","1","123","0","0","97.33","449","0","450","32.89","0"
"AGAW1","08E","INS","BOLD:AAF9035","Arthropoda","Insecta","Diptera","Tipulidae","Tipulinae","Nephrotoma","Nephrotoma alterna","1","1","1","123","0","0","99.55","449","0","450","33.78","0"
"AGAW1","08E","NAR","no match","no match","no match","no match","no match","no match","no match","no match","no match","no match","no match","no match","no match","no match","no match","no match","no match","no match","no match","no match"
"AGAW1","08E","NIN","BOLD:AAA7242","Arthropoda","Collembola","Entomobryomorpha","Entomobryidae","Seirinae, Entomobryinae","Entomobrya, Lepidosira","Entomobrya multifasciata, Lepidosira sp.","261","255","255","1","2.3","0","99.16","417.85","0","421.36","39.25","0"
"AGAW1","08E","NIN","BOLD:AAH6630","Arthropoda","Collembola","Entomobryomorpha","Entomobryidae","Entomobryinae","Entomobrya","Entomobrya unostrigata","31","29","29","2","6.45","0","99.61","443.19","0","444.16","38.8","0"
"AGAW1","08E","NIN","BOLD:ACV5610","","","","","","","","18","12","12","3","33.33","0","99.75","449.17","0","449.89","38.26","0"
"AGAW1","08E","NIN","BOLD:AAB7914","Arthropoda","Collembola","Symphypleona","Bourletiellidae","","Bourletiella","","6","6","6","4","0","0","99.85","449.67","0","450","38.33","0"
"AGAW1","08E","NIN","BOLD:ADH9851","Arthropoda","Collembola","Entomobryomorpha","","","","","4","4","4","5","0","0","98.33","449.25","0","450","39.56","0"
"AGAW1","08E","NIN","BOLD:ACU7748","Arthropoda","Arachnida","Trombidiformes","Pygmephoridae","","","","3","3","3","6","0","0","99.17","401","0","435.33","28.76","0"
"AGAW1","08E","NIN","BOLD:ACP6423","Arthropoda","Arachnida","Trombidiformes","Microtrombidiidae","","","","1","1","1","7","0","0","99.74","388","0","450","37.56","0"
"AGAW1","08E","NIN","BOLD:ACI9908","Arthropoda","Collembola","Symphypleona","Bourletiellidae","","","","1","1","1","7","0","0","98.64","367","0","396","33.59","0"
"AGAW1","08E","NIN","BOLD:AAB3836","Arthropoda","Arachnida","Araneae","Philodromidae","","Philodromus","Philodromus cespitum","1","1","1","7","0","0","97.22","395","0","403","30.02","0"
"AGAW1","08E","NIN","BOLD:AAA8924","Arthropoda","Collembola","Entomobryomorpha","Entomobryidae","Entomobryinae","Entomobrya","Entomobrya nivalis","1","1","1","7","0","0","98.88","446","0","450","38.89","0"
"AGAW1","08E","NIN","BOLD:AAA5267","Arthropoda","Collembola","Entomobryomorpha","Isotomidae","","","","1","1","1","7","0","0","99.55","446","0","450","38.67","0"
"AGAW1","08E","NIN","BOLD:ACX8969","Arthropoda","Arachnida","Trombidiformes","Eupodidae","","","","1","1","1","7","0","0","98.93","374","0","375","27.2","0"
"AGAW1","08N","BAC","TAX:91393","Proteobacteria","Alphaproteobacteria","Rickettsiales","Rickettsiaceae",NA,"Wolbachia","","32","32","32","1","0","0","98.98","390.75","0","422.56","39.13","0"
"AGAW1","08N","BAC","TAX:175768","Proteobacteria","Alphaproteobacteria","Rickettsiales","Rickettsiaceae",NA,"Wolbachia","Wolbachia pipientis","29","25","25","2","13.79","0","98.29","443.79","0","445","39.49","0"
"AGAW1","08N","BAC","TAX:91391","Proteobacteria","Alphaproteobacteria","Rickettsiales","",NA,"","","13","11","11","3","15.38","0","99","421.85","0","448.54","39.67","0"
"AGAW1","08N","CHO","no match","no match","no match","no match","no match","no match","no match","no match","no match","no match","no match","no match","no match","no match","no match","no match","no match","no match","no match","no match"
"AGAW1","08N","INS","BOLD:AAA2815","Arthropoda","Insecta","Lepidoptera","Noctuidae","Noctuinae","Mythimna","Mythimna yukonensis, Mythimna oxygala","25359","19340","19340","1","23.74","0","99.39","425.23","0","426.86","33.47","0"
"AGAW1","08N","INS","BOLD:AAG2511","Arthropoda","Insecta","Diptera","Anthomyiidae","Anthomyiinae","Delia","Delia platura","12169","9707","9707","2","20.23","0","99.12","444.11","0","447.04","33.09","0"
"AGAW1","08N","INS","BOLD:AAP8739","Arthropoda","Insecta","Diptera","Platystomatidae","Platystomatinae","Rivellia","Rivellia steyskali","8646","5412","5412","3","37.4","0","99.31","441.76","0","443.76","37.97","0"
"AGAW1","08N","INS","BOLD:AAB0373","Arthropoda","Insecta","Neuroptera","Chrysopidae","Chrysopinae","Chrysoperla","Chrysoperla sp. isolate 82, Chrysoperla zastrowi, Chrysoperla plorabunda, Chrysoperla sp. isolate 71, Chrysoperla agilis, Italochrysa italica, Chrysoperla sp. isolate 88, Chrysoperla lucasina, Chrysoperla carnea, Chrysoperla mediterranea, Chrysoperla pallida, Chrysoperla nipponensis, Chrysoperla sillemi","3338","2437","2437","4","26.99","0","99.53","436.84","0","437.76","31.33","0"
"AGAW1","08N","INS","BOLD:AAI4303","Arthropoda","Insecta","Diptera","Chironomidae","Chironominae","Chironomus","Chironomus melanescens","2496","2148","2148","5","13.94","0","97.88","432.5","0","437.14","33.41","0"
"AGAW1","08N","INS","BOLD:AAA7374","Arthropoda","Insecta","Diptera","Syrphidae","Syrphinae","Sphaerophoria","Sphaerophoria bankowskae, Sphaerophoria infuscata, Sphaerophoria nr. philanthus, Sphaerophoria batava, Sphaerophoria sp., Sphaerophoria rueppellii, Sphaerophoria interrupta-group, Sphaerophoria sp. MA-1, Sphaerophoria cleoae, Sphaerophoria asymmetrica, Sphaerophoria fatarum, Sphaerophoria bengalensis, Sphaerophoria bifurcata, Sphaerophoria taeniata, Sphaerophoria interrupta, Sphaerophoria contigua, Sphaerophoria kaa, Sphaerophoria laurae, Sphaerophoria chongjini, Sphaerophoria boreoalpina, Sphaerophoria sp. BOLDAAA7374, Sphaerophoria macrogaster, Sphaerophoria sulphuripes, Sphaerophoria brevipilosa, Sphaerophoria philanthus, Sphaerophoria pyrrhina, Sphaerophoria abbreviata, Sphaerophoria scripta, Sphaerophoria potentillae, Sphaerophoria longipilosa, Sphaerophoria virgata","1743","1517","1517","6","12.97","0","99.33","429.7","0","432.83","30.71","0"
"AGAW1","08N","INS","BOLD:ACR4394","Arthropoda","Insecta","Diptera","Anthomyiidae","Anthomyiinae","Delia","Delia florilega","1258","1165","1165","7","7.39","0","99.21","431.06","0","432.91","32.85","0"
"AGAW1","08N","INS","BOLD:AAD2041","Arthropoda","Insecta","Diptera","Dolichopodidae","Sciapodinae","Condylostylus","Condylostylus inermis, Condylostylus nigrofemoratus, Condylostylus connectans, Condylostylus caudatus, Condylostylus flavipes","1032","741","741","8","28.2","0","99.52","438.75","0","439.96","35.93","0"
"AGAW1","08N","INS","BOLD:AAB2384","Arthropoda","Insecta","Diptera","Syrphidae","Syrphinae","Eupeodes","Eupeodes duseki, Eupeodes volucris, Eupeodes nielseni, Eupeodes americanus, Eupeodes flukei, Eupeodes vockerothi, Eupeodes lucasi, Eupeodes montivagus, Eupeodes rufipunctatus, Eupeodes rojasi, Eupeodes goeldlini, Eupeodes luniger, Eupeodes punctifer, Eupeodes bucculatus, Eupeodes perplexus, Eupeodes abiskoensis, Eupeodes corollae, Eupeodes pomus, Eupeodes lundbecki, Eupeodes nitens, Eupeodes sp. BOLDAAB2384, Eupeodes curtus, Eupeodes latifasciatus","832","801","801","9","3.73","0","98.95","402.52","0","405.09","30.69","0"
"AGAW1","08N","INS","BOLD:AAF9771","Arthropoda","Insecta","Diptera","Hybotidae","Tachydromiinae","Platypalpus","Platypalpus longiseta, Platypalpus trivialis, Platypalpus sp. 5, Platypalpus sp. 17, Platypalpus pallidiventris","831","568","568","10","31.65","0","99.24","446.52","0","447.06","35.1","0"
"AGAW1","08N","INS","BOLD:AAG6745","Arthropoda","Insecta","Diptera","Calliphoridae","Calliphorinae","Pollenia","Pollenia pediculata","648","576","576","11","11.11","0","99.33","432.32","0","434.78","33.52","0"
"AGAW1","08N","INS","BOLD:AAG2868","Arthropoda","Insecta","Hemiptera","Cicadellidae","Typhlocybinae","Empoasca","Empoasca sp., Empoasca fabae","630","383","383","12","39.21","0","99.71","444.29","0","447.38","32.43","0"
"AGAW1","08N","INS","BOLD:AAH4135","Arthropoda","Insecta","Diptera","Chloropidae","Chloropinae","Thaumatomyia","Thaumatomyia glabra, Thaumatomyia grata, Thaumatomyia sp. 1 TAW","474","449","449","13","5.27","0","99.2","430.69","0","435.75","31.67","0"
"AGAW1","08N","INS","BOLD:AAO3939","Arthropoda","Insecta","Diptera","Limoniidae","Limoniinae","Dicranomyia","","354","279","279","14","21.19","0","99.26","440.37","0","441.7","36.24","0"
"AGAW1","08N","INS","BOLD:AAG7279","Arthropoda","Insecta","Diptera","Sphaeroceridae","Limosininae","Spelobia","Spelobia ochripes","331","265","265","15","19.94","0","99.34","440.83","0","442.11","35.09","0"
"AGAW1","08N","INS","BOLD:AAO7783","Arthropoda","Insecta","Diptera","Dolichopodidae","Diaphorinae","","","224","214","214","16","4.46","0","98.1","425.65","0","428","33.23","0"
"AGAW1","08N","INS","BOLD:ACK7314","Arthropoda","Insecta","Diptera","Chironomidae","Chironominae","Polypedilum","","220","219","219","17","0.45","0","98.91","392.94","0","397.13","35.16","0"
"AGAW1","08N","INS","BOLD:ACK3431","Arthropoda","Insecta","Diptera","Cecidomyiidae","","","","220","193","193","17","12.27","0","98.98","427.48","0","429.52","32.6","0"
"AGAW1","08N","INS","BOLD:AAG0999","Arthropoda","Insecta","Diptera","Chironomidae","Orthocladiinae","","","218","196","196","19","10.09","0","98.31","434.47","0","435.8","35.27","0"
"AGAW1","08N","INS","BOLD:AAH4205","Arthropoda","Insecta","Diptera","Chloropidae","Chloropinae","Meromyza","Meromyza nigriseta, Meromyza sp.","202","140","140","20","30.69","0","99.52","446.88","0","447.98","32.85","0"
"AGAW1","08N","INS","BOLD:AAP6790","Arthropoda","Insecta","Diptera","Chloropidae","Oscinellinae","Incertella","Incertella incerta","197","163","163","21","17.26","0","99.46","435.41","0","437.22","31.36","0"
"AGAW1","08N","INS","BOLD:AAA3453","Arthropoda","Insecta","Diptera","Anthomyiidae","Anthomyiinae","Delia","","195","192","192","22","1.54","0","99.19","433.16","0","436.19","34.55","0"
"AGAW1","08N","INS","BOLD:AAN5342","Arthropoda","Insecta","Diptera","Chironomidae","Orthocladiinae","Heleniella","","163","157","157","23","3.68","0","99.4","419.07","0","420.23","35.39","0"
"AGAW1","08N","INS","BOLD:AAA7661","Arthropoda","Insecta","Diptera","Culicidae","Culicinae","Culiseta, Culex","Culiseta melanura, Culex articularis, Culex restuans, Culex acharistus","156","153","153","24","1.92","0","99.24","432.03","0","434.21","32.59","0"
"AGAW1","08N","INS","BOLD:AAN5199","Arthropoda","Insecta","Diptera","Cecidomyiidae","","","CecidInt20 sp. BOLD:AAN5199","124","93","93","25","25","0","99.59","438.26","0","446.29","26.67","0"
"AGAW1","08N","INS","BOLD:ACP8516","Arthropoda","Insecta","Diptera","Chironomidae","Tanypodinae","Procladius","","115","95","95","26","17.39","0","99.3","417.05","0","440.9","34.73","0"
"AGAW1","08N","INS","BOLD:AAC4197","Arthropoda","Insecta","Diptera","Chironomidae","Orthocladiinae","Paraphaenocladius","Paraphaenocladius impensus","75","68","68","27","9.33","0","98.96","430.16","0","433.61","34.54","0"
"AGAW1","08N","INS","BOLD:AAG7891","Arthropoda","Insecta","Hymenoptera","Platygastridae","Scelioninae","Telenomus","Telenomus podisi","70","68","68","28","2.86","0","99.01","431.21","0","440.56","27.05","0"
"AGAW1","08N","INS","BOLD:AAB1097","Arthropoda","Insecta","Lepidoptera","Blastobasidae","Blastobasinae","Blastobasis","Blastobasis floridella","64","60","60","29","6.25","0","98.75","436.17","0","439.05","32.73","0"
"AGAW1","08N","INS","BOLD:AAA3080","Arthropoda","Insecta","Trichoptera","Limnephilidae","Limnephilinae","Limnephilus","Limnephilus submonilifer","55","53","53","30","3.64","0","99.22","402.35","0","404.4","33.97","0"
"AGAW1","08N","INS","BOLD:AAN5238","Arthropoda","Insecta","Diptera","Cecidomyiidae","","","","50","43","43","31","14","0","99.37","428.96","0","430.26","25.64","0"
"AGAW1","08N","INS","BOLD:ABA7871","Arthropoda","Insecta","Diptera","Cecidomyiidae","","","","49","43","43","32","12.24","0","99.04","439.9","0","440.67","26.79","0"
"AGAW1","08N","INS","BOLD:AAG3154","Arthropoda","Insecta","Diptera","Psychodidae","Psychodinae","Psychoda","","47","43","43","33","8.51","0","98.09","442.09","0","443.21","32.47","0"
"AGAW1","08N","INS","BOLD:AAA5299","Arthropoda","Insecta","Diptera","Chironomidae","Orthocladiinae","Cricotopus","Cricotopus laetus, Cricotopus relucens, Cricotopus trifasciatus, Cricotopus sp., Cricotopus sp. 3ES, Cricotopus sylvestris, Cricotopus glacialis","41","39","39","34","4.88","0","99.22","431.1","0","432.71","34.84","0"
"AGAW1","08N","INS","BOLD:AAG9897","Arthropoda","Insecta","Dermaptera","Forficulidae","Forficulinae","Forficula","Forficula auricularia-A","40","40","40","35","0","0","99.01","392.38","0","401.05","45.75","0"
"AGAW1","08N","INS","BOLD:ABY1179","Arthropoda","Insecta","Hymenoptera","Sierolomorphidae","","Sierolomorpha","","40","32","32","35","20","0","99.59","435.68","0","437.23","28.09","0"
"AGAW1","08N","INS","BOLD:AAH3968","Arthropoda","Insecta","Diptera","Sciaridae","","Cratyna","Cratyna cryptospina","31","30","30","37","3.23","0","99.18","421.84","0","433.65","27.63","0"
"AGAW1","08N","INS","BOLD:ABX5069","Arthropoda","Insecta","Trichoptera","Hydroptilidae","Hydroptilinae","Hydroptila","Hydroptila consimilis","27","16","16","38","40.74","0","99.58","448.37","0","449.89","33.3","0"
"AGAW1","08N","INS","BOLD:AAH3661","Arthropoda","Insecta","Diptera","Cecidomyiidae","","","","25","24","24","39","4","0","99.35","436.84","0","438.64","25.53","0"
"AGAW1","08N","INS","BOLD:ACF4455","Arthropoda","Insecta","Hemiptera","Aphididae","Lachninae","Eulachnus","Eulachnus agilis","25","25","25","39","0","0","99.48","433.68","0","434.56","26.27","0"
"AGAW1","08N","INS","BOLD:AAM7855","Arthropoda","Insecta","Thysanoptera","Thripidae","Thripinae","Frankliniella","Frankliniella tritici","23","23","23","41","0","0","99.27","439.43","0","442.74","31.18","0"
"AGAW1","08N","INS","BOLD:AAG5453","Arthropoda","Insecta","Diptera","Chironomidae","Chironominae","Chironomus","Chironomus bifurcatus","20","20","20","42","0","0","98.27","448.55","0","450","34.7","0"
"AGAW1","08N","INS","BOLD:ABU5525","Arthropoda","Insecta","Diptera","Chironomidae","Orthocladiinae","Limnophyes","Limnophyes sp. 14ES","18","18","18","43","0","0","99.26","376.83","0","380.78","37.59","0"
"AGAW1","08N","INS","BOLD:AAG1711","Arthropoda","Insecta","Diptera","Muscidae","Phaoniinae","Helina","Helina depuncta","17","17","17","44","0","0","99.44","439.76","0","441.71","31.35","0"
"AGAW1","08N","INS","BOLD:AAA3882","Arthropoda","Insecta","Lepidoptera","Pterophoridae","Pterophorinae","Emmelina","Emmelina monodactyla","14","14","14","45","0","0","98.52","411.93","0","431","32.09","0"
"AGAW1","08N","INS","BOLD:ABU5546","Arthropoda","Insecta","Diptera","Psychodidae","Psychodinae","Psychoda","","14","14","14","45","0","0","98.66","426.86","0","428.79","30.83","0"
"AGAW1","08N","INS","BOLD:AAH0152","Arthropoda","Insecta","Coleoptera","Cantharidae","Cantharinae","Cantharis","Cantharis rufa","14","13","13","45","7.14","0","98.31","438.14","0","442.64","35.11","0"
"AGAW1","08N","INS","BOLD:AAH0134","Arthropoda","Insecta","Coleoptera","Phalacridae","Phalacrinae","Stilbus","Stilbus apicialis","12","9","9","48","25","0","99.54","439","0","440.33","31.88","0"
"AGAW1","08N","INS","BOLD:ACJ5126","Arthropoda","Insecta","Diptera","Chironomidae","Orthocladiinae","Metriocnemus","","10","10","10","49","0","0","99.26","416.4","0","419.9","34.57","0"
"AGAW1","08N","INS","BOLD:ABV9277","","","","","","","","9","9","9","50","0","0","99.63","448.44","0","450","28.3","0"
"AGAW1","08N","INS","BOLD:AAS1281","Arthropoda","Insecta","Diptera","Chironomidae","Chironominae","Chironomus","Chironomus nr. sollicitus","8","8","8","51","0","0","97.25","409.13","0","412.13","32.64","0"
"AGAW1","08N","INS","BOLD:AAN6154","Arthropoda","Insecta","Coleoptera","Latridiidae","Corticariinae","Melanophthalma","Melanophthalma inermis","8","8","8","51","0","0","98.93","439.13","0","440.13","31.4","0"
"AGAW1","08N","INS","BOLD:AAC0903","Arthropoda","Insecta","Diptera","Chironomidae","Chironominae","Chironomus","Chironomus acidophilus","5","5","5","53","0","0","99.22","420.6","0","431.2","36.57","0"
"AGAW1","08N","INS","BOLD:ABW7897","Arthropoda","Insecta","Diptera","Cecidomyiidae","","","","5","5","5","53","0","0","99.29","448.4","0","450","29.87","0"
"AGAW1","08N","INS","BOLD:AAU6681","Arthropoda","Insecta","Diptera","Tipulidae","Tipulinae","Nephrotoma","Nephrotoma cornicina","4","4","4","55","0","0","99.2","436","0","437.5","36.2","0"
"AGAW1","08N","INS","BOLD:ACJ8771","Arthropoda","Insecta","Diptera","Chironomidae","Chironominae","Sublettea","","4","4","4","55","0","0","99.1","389","0","442.75","32.64","0"
"AGAW1","08N","INS","BOLD:AAV5573","Arthropoda","Insecta","Diptera","Cecidomyiidae","","","","4","4","4","55","0","0","99.67","449.5","0","450","27.67","0"
"AGAW1","08N","INS","BOLD:AAA4277","Arthropoda","Insecta","Diptera","Syrphidae","Syrphinae","Toxomerus","Toxomerus marginatus, Toxomerus sp. CR-3, Toxomerus CR-3","3","3","3","58","0","0","98.52","450","0","450","31.26","0"
"AGAW1","08N","INS","BOLD:ABY5318","Arthropoda","Insecta","Diptera","Chloropidae","Oscinellinae","Incertella, Rhopalopterum","Rhopalopterum umbrosum, Rhopalopterum luteiceps, Incertella bispina, Rhopalopterum painteri","3","3","3","58","0","0","99.17","443.33","0","444","32.13","0"
"AGAW1","08N","INS","BOLD:AAA2632","Arthropoda","Insecta","Lepidoptera","Noctuidae","Noctuinae","Noctua","Noctua atlantica, Noctua pronuba","3","3","3","58","0","0","99.04","449.33","0","450","31.19","0"
"AGAW1","08N","INS","BOLD:AAH3664","Arthropoda","Insecta","Diptera","Cecidomyiidae","","","CecidInt3 sp. BOLD:AAH3664","3","3","3","58","0","0","98.77","433","0","435.33","27.01","0"
"AGAW1","08N","INS","BOLD:AAG5507","Arthropoda","Insecta","Diptera","Chironomidae","Chironominae","Chironomus","Chironomus atrella","2","2","2","62","0","0","98.98","433","0","432.5","35.81","0"
"AGAW1","08N","INS","BOLD:ABA1215","Arthropoda","Insecta","Diptera","Sciaridae","","Lycoriella","Lycoriella sativae","2","1","1","62","50","0","99.78","450","0","450","36.22","0"
"AGAW1","08N","INS","BOLD:AAB7869","Arthropoda","Insecta","Lepidoptera","Tortricidae","Tortricinae","Phtheochroa","Phtheochroa baracana, Phtheochroa riscana, Phtheochroa birdana, Phtheochroa fulviplicana, Phtheochroa canariana","2","2","2","62","0","0","99.44","449","0","450","29.78","0"
"AGAW1","08N","INS","BOLD:AAA3889","Arthropoda","Insecta","Lepidoptera","Crambidae","Crambinae","Parapediasia","Parapediasia teterrellus","2","2","2","62","0","0","98.74","400","0","421.5","32.01","0"
"AGAW1","08N","INS","BOLD:ACJ6984","Arthropoda","Insecta","Hymenoptera","Platygastridae","Scelioninae","Trissolcus","Trissolcus thyantae","2","2","2","62","0","0","99.05","427.5","0","434.5","27.39","0"
"AGAW1","08N","INS","BOLD:AAB4894","Arthropoda","Insecta","Hemiptera","Aphididae","Aphidinae","Sitobion","Sitobion smilacifoliae, Sitobion avenae, Sitobion phyllanthi","2","2","2","62","0","0","97.67","450","0","450","25.67","0"
"AGAW1","08N","INS","BOLD:AAP6580","Arthropoda","Insecta","Diptera","Chironomidae","Chironominae","Paratanytarsus","Paratanytarsus sp. 5TE","2","2","2","62","0","0","98.65","408.5","0","409.5","32.57","0"
"AGAW1","08N","INS","BOLD:AAD2548","Arthropoda","Insecta","Hemiptera","Aphididae","Aphidinae","Acyrthosiphon","Acyrthosiphon caraganae, Acyrthosiphon sp. 1, Acyrthosiphon kondoi, Acyrthosiphon churchillense","2","2","2","62","0","0","99.89","450","0","450","26.11","0"
"AGAW1","08N","INS","BOLD:ABW1379","Arthropoda","Insecta","Diptera","Chloropidae","Oscinellinae","Malloewia","Malloewia abdominalis","2","2","2","62","0","0","97.89","449","0","450","31.89","0"
"AGAW1","08N","INS","BOLD:AAG2123","Arthropoda","Insecta","Diptera","Tachinidae","Dexiinae","Dinera","Dinera grisescens","2","2","2","62","0","0","99.38","387","0","388.5","30.76","0"
"AGAW1","08N","INS","BOLD:AAU6656","Arthropoda","Insecta","Diptera","Limoniidae","Chioneinae","Neolimnophila","Neolimnophila placida","2","2","2","62","0","0","97.64","403","0","406.5","35.07","0"
"AGAW1","08N","INS","BOLD:AAH1890","Arthropoda","Insecta","Hymenoptera","Ichneumonidae","Cryptinae","Stibeutes","Stibeutes brevicornis, Stibeutes heinemanni","2","2","2","62","0","0","99.44","450","0","450","29.56","0"
"AGAW1","08N","INS","BOLD:AAZ0271","Arthropoda","Insecta","Diptera","Ceratopogonidae","Ceratopogoninae","Culicoides","","2","2","2","62","0","0","98.1","446.5","0","450","41","0"
"AGAW1","08N","INS","BOLD:AAG2460","Arthropoda","Insecta","Diptera","Anthomyiidae","Anthomyiinae","Hydrophoria","Hydrophoria lancifer","2","2","2","62","0","0","98.6","434.5","0","450","34.22","0"
"AGAW1","08N","INS","BOLD:AAH3949","Arthropoda","Insecta","Diptera","Sciaridae","","Pseudolycoriella","","2","2","2","62","0","0","99.89","449","0","450","36.11","0"
"AGAW1","08N","INS","BOLD:AAA3181","Arthropoda","Insecta","Diptera","Muscidae","Muscinae","Stomoxys","Stomoxys calcitrans","1","1","1","77","0","0","98.5","400","0","405","32.35","0"
"AGAW1","08N","INS","BOLD:ABW3687","Arthropoda","Insecta","Diptera","Lauxaniidae","Homoneurinae","Homoneura","Homoneura bispina","1","1","1","77","0","0","98.52","406","0","406","33.74","0"
"AGAW1","08N","INS","BOLD:AAA6292","Arthropoda","Insecta","Lepidoptera","Tortricidae","Tortricinae","Cnephasia","Cnephasia genitalana, Cnephasia stephensiana","1","1","1","77","0","0","98.48","395","0","396","34.09","0"
"AGAW1","08N","INS","BOLD:AAA2590","Arthropoda","Insecta","Lepidoptera","Noctuidae","Noctuinae","Xestia","Xestia baja, Xestia smithii","1","1","1","77","0","0","97.74","442","0","443","33.41","0"
"AGAW1","08N","INS","BOLD:AAG7276","Arthropoda","Insecta","Diptera","Sphaeroceridae","Limosininae","Leptocera","Leptocera erythrocera","1","1","1","77","0","0","99.78","450","0","450","33.11","0"
"AGAW1","08N","INS","BOLD:AAA6560","Arthropoda","Insecta","Lepidoptera","Crambidae","Crambinae","Crambus","Crambus multilinellus, Crambus saltuellus","1","1","1","77","0","0","98.82","425","0","427","30.91","0"
"AGAW1","08N","INS","BOLD:AAP5169","Arthropoda","Insecta","Diptera","Chloropidae","Oscinellinae","Elachiptera","Elachiptera nigriceps","1","1","1","77","0","0","97.55","449","0","450","32.67","0"
"AGAW1","08N","INS","BOLD:ABV9319","Arthropoda","Insecta","Diptera","Chironomidae","Tanypodinae","Procladius","","1","1","1","77","0","0","99.46","370","0","374","34.76","0"
"AGAW1","08N","INS","BOLD:AAF9949","Arthropoda","Insecta","Hemiptera","Miridae","Mirinae","Trigonotylus","Trigonotylus pulchellus, Trigonotylus americanus, Trigonotylus psammaecolor, Trigonotylus caelestialium, Trigonotylus viridis, Trigonotylus ruficornis","1","1","1","77","0","0","99.78","450","0","450","34.44","0"
"AGAW1","08N","INS","BOLD:AAA3867","Arthropoda","Insecta","Lepidoptera","Noctuidae","Noctuinae","Apamea","Apamea amputatrix","1","1","1","77","0","0","100","447","0","450","33.56","0"
"AGAW1","08N","INS","BOLD:AAG5457","Arthropoda","Insecta","Diptera","Chironomidae","Chironominae","","","1","1","1","77","0","0","99.55","448","0","450","34.22","0"
"AGAW1","08N","INS","BOLD:AAA5803","Arthropoda","Insecta","Hemiptera","Miridae","Mirinae","Lygus","Lygus lineolaris","1","1","1","77","0","0","99.11","450","0","450","33.56","0"
"AGAW1","08N","INS","BOLD:ABY4763","Arthropoda","Insecta","Hemiptera","Aphididae","Aphidinae","Uroleucon","Uroleucon taraxaci","1","1","1","77","0","0","100","448","0","450","24.89","0"
"AGAW1","08N","INS","BOLD:AAH3914","Arthropoda","Insecta","Diptera","Sciaridae","","Sciara","Sciara hamatilis, Sciara humeralis","1","1","1","77","0","0","99.5","404","0","406","29.56","0"
"AGAW1","08N","INS","BOLD:AAD1395","Arthropoda","Insecta","Diptera","Chironomidae","Chironominae","Polypedilum","Polypedilum convictum, Polypedilum flavum","1","1","1","77","0","0","98.89","449","0","450","33.78","0"
"AGAW1","08N","INS","BOLD:AAZ5483","Arthropoda","Insecta","Diptera","Muscidae","Phaoniinae","Helina","Helina setiventris","1","1","1","77","0","0","99.11","449","0","450","29.78","0"
"AGAW1","08N","INS","BOLD:AAD1485","Arthropoda","Insecta","Diptera","Chironomidae","Chironominae","Paratanytarsus","Paratanytarsus grimmii, Paratanytarsus sp.","1","1","1","77","0","0","97.44","351","0","352","33.52","0"
"AGAW1","08N","INS","BOLD:AAH2293","Arthropoda","Insecta","Diptera","Asilidae","Stenopogoninae","Dioctria, Eudioctria","Dioctria hyalipennis, Dioctria baumhaueri, Eudioctria propinqua","1","1","1","77","0","0","99.23","392","0","396","30.56","0"
"AGAW1","08N","INS","BOLD:ADF6661","Arthropoda","Insecta","Diptera","Cecidomyiidae","","","","1","1","1","77","0","0","99.76","416","0","434","29.72","0"
"AGAW1","08N","INS","BOLD:AAG6753","Arthropoda","Insecta","Diptera","Lauxaniidae","Lauxaniinae","Camptoprosopella","","1","1","1","77","0","0","97.08","411","0","414","35.02","0"
"AGAW1","08N","INS","BOLD:AAF9187","Arthropoda","Insecta","Coleoptera","Curculionidae","Entiminae","Phyllobius","Phyllobius oblongus","1","1","1","77","0","0","100","438","0","450","38.44","0"
"AGAW1","08N","INS","BOLD:AAA4188","Arthropoda","Insecta","Hymenoptera","Braconidae","Aphidiinae","Aphidius","Aphidius microlophii, Aphidius ervi","1","1","1","77","0","0","98.67","450","0","450","27.11","0"
"AGAW1","08N","INS","BOLD:ACF2821","Arthropoda","Insecta","Diptera","Mycetophilidae","Mycetophilinae","Mycetophila","Mycetophila fungorum","1","1","1","77","0","0","99.77","443","0","450","34.89","0"
"AGAW1","08N","INS","BOLD:AAG6588","Arthropoda","Insecta","Coleoptera","Chrysomelidae","Cassidinae","Deloyala","Deloyala guttata","1","1","1","77","0","0","99.27","413","0","415","29.88","0"
"AGAW1","08N","INS","BOLD:AAA9246","Arthropoda","Insecta","Lepidoptera","Tortricidae","Tortricinae","Clepsis","Clepsis clemensiana","1","1","1","77","0","0","98.67","450","0","450","31.11","0"
"AGAW1","08N","INS","BOLD:AAH0256","Arthropoda","Insecta","Coleoptera","Latridiidae","Corticariinae","Corticaria, Corticarina","","1","1","1","77","0","0","99.54","438","0","450","31.78","0"
"AGAW1","08N","INS","BOLD:AAH2876","Arthropoda","Insecta","Diptera","Cecidomyiidae","","","CecidInt51 sp. BOLD:AAH2876","1","1","1","77","0","0","99.78","450","0","450","26.67","0"
"AGAW1","08N","INS","BOLD:AAF9306","Arthropoda","Insecta","Diptera","Psychodidae","Psychodinae","Psychoda","Psychoda alternata","1","1","1","77","0","0","98.44","450","0","450","31.56","0"
"AGAW1","08N","INS","BOLD:ACF7023","Arthropoda","Insecta","Diptera","Dolichopodidae","","","","1","1","1","77","0","0","98.24","227","0","450","33.78","0"
"AGAW1","08N","INS","BOLD:ABX2489","Arthropoda","Insecta","Coleoptera","Phalacridae","Phalacrinae","Stilbus","","1","1","1","77","0","0","97.47","396","0","397","31.74","0"
"AGAW1","08N","INS","BOLD:AAH0006","Arthropoda","Insecta","Diptera","Chironomidae","Chironominae","Sublettea","","1","1","1","77","0","0","97.32","448","0","450","32.44","0"
"AGAW1","08N","INS","BOLD:ABV1237","Arthropoda","Insecta","Diptera","Chironomidae","Chironominae","Chironomus","","1","1","1","77","0","0","97.56","450","0","450","34.67","0"
"AGAW1","08N","INS","BOLD:AAG4944","Arthropoda","Insecta","Diptera","Mycetophilidae","Sciophilinae","Sciophila","Sciophila sp. CJB1","1","1","1","77","0","0","98.66","447","0","450","33.56","0"
"AGAW1","08N","INS","BOLD:AAF1051","Arthropoda","Insecta","Diptera","Agromyzidae","Phytomyzinae","Cerodontha","Cerodontha muscina, Cerodontha calamagrostis","1","1","1","77","0","0","98.77","408","0","410","35.12","0"
"AGAW1","08N","INS","BOLD:AAG7036","Arthropoda","Insecta","Diptera","Agromyzidae","","","","1","1","1","77","0","0","99.06","426","0","450","33.11","0"
"AGAW1","08N","INS","BOLD:AAY6534","Arthropoda","Insecta","Coleoptera","Mordellidae","Mordellinae","Mordellina","Mordellina ancilla","1","1","1","77","0","0","100","445","0","445","38.65","0"
"AGAW1","08N","NAR","no match","no match","no match","no match","no match","no match","no match","no match","no match","no match","no match","no match","no match","no match","no match","no match","no match","no match","no match","no match"
"AGAW1","08N","NIN","BOLD:AAA7242","Arthropoda","Collembola","Entomobryomorpha","Entomobryidae","Seirinae, Entomobryinae","Entomobrya, Lepidosira","Entomobrya multifasciata, Lepidosira sp.","276","262","262","1","5.07","0","99.25","424.24","0","427.2","39.3","0"
"AGAW1","08N","NIN","BOLD:AAF1092","Arthropoda","Arachnida","Trombidiformes","Anystidae","","","","7","7","7","2","0","0","97.73","441.43","0","442.86","30.19","0"
"AGAW1","08N","NIN","BOLD:AAH6630","Arthropoda","Collembola","Entomobryomorpha","Entomobryidae","Entomobryinae","Entomobrya","Entomobrya unostrigata","3","3","3","3","0","0","99.6","438.67","0","440.67","38.42","0"
"AGAW1","08N","NIN","BOLD:AAZ2181","Arthropoda","Collembola","Symphypleona","Bourletiellidae","","","","2","2","2","4","0","0","97.44","449","0","450","39.56","0"
"AGAW1","08N","NIN","BOLD:ACP9512","Arthropoda","Collembola","Entomobryomorpha","Entomobryidae","","","","2","2","2","4","0","0","98.73","396","0","399.5","39.05","0"
"AGAW1","08N","NIN","BOLD:ABW2639","Arthropoda","Arachnida","Trombidiformes","Anystidae","","","","1","1","1","6","0","0","98.97","484","0","450","31.33","0"
"AGAW1","08N","NIN","BOLD:ACC8538","Arthropoda","Arachnida","Mesostigmata","Blattisociidae","","","","1","1","1","6","0","0","99.74","383","0","384","32.29","0"
"AGAW1","08N","NIN","BOLD:AAM3234","Arthropoda","Collembola","Entomobryomorpha","Entomobryidae","Seirinae","Lepidosira","","1","1","1","6","0","0","98.67","450","0","450","39.11","0"
"AGAW1","08N","NIN","BOLD:AAI4346","Arthropoda","Arachnida","Opiliones","Phalangiidae","","Paroligolophus, Phalangium","Paroligolophus agrestis, Phalangium opilio","1","1","1","6","0","0","99.11","447","0","450","36","0"
"AGAW1","08S","BAC","TAX:91393","Proteobacteria","Alphaproteobacteria","Rickettsiales","Rickettsiaceae",NA,"Wolbachia","","40","40","40","1","0","0","99","391.73","0","421.43","39.08","0"
"AGAW1","08S","BAC","TAX:175768","Proteobacteria","Alphaproteobacteria","Rickettsiales","Rickettsiaceae",NA,"Wolbachia","Wolbachia pipientis","35","30","30","2","14.29","0","98.36","446.06","0","447.26","39.53","0"
"AGAW1","08S","BAC","TAX:91391","Proteobacteria","Alphaproteobacteria","Rickettsiales","",NA,"","","15","14","14","3","6.67","0","98.93","412.6","0","442.73","39.52","0"
"AGAW1","08S","BAC","TAX:91389","Proteobacteria","","","",NA,"","","2","2","2","4","0","0","99.01","354","0","357","38.52","0"
"AGAW1","08S","BAC","TAX:392909","Proteobacteria","Alphaproteobacteria","Rickettsiales","Rickettsiaceae",NA,"Wolbachia","Wolbachia endosymbiont of Microgastrinae","1","1","1","5","0","0","99.56","450","0","450","40.44","0"
"AGAW1","08S","BAC","TAX:376099","Proteobacteria","Alphaproteobacteria","Rickettsiales","Rickettsiaceae",NA,"Wolbachia","Wolbachia sp. Hymenoptera","1","1","1","5","0","0","97.04","439","0","441","40.36","0"
"AGAW1","08S","CHO","no match","no match","no match","no match","no match","no match","no match","no match","no match","no match","no match","no match","no match","no match","no match","no match","no match","no match","no match","no match"
"AGAW1","08S","INS","BOLD:AAB0373","Arthropoda","Insecta","Neuroptera","Chrysopidae","Chrysopinae","Chrysoperla","Chrysoperla sp. isolate 82, Chrysoperla zastrowi, Chrysoperla plorabunda, Chrysoperla sp. isolate 71, Chrysoperla agilis, Italochrysa italica, Chrysoperla sp. isolate 88, Chrysoperla lucasina, Chrysoperla carnea, Chrysoperla mediterranea, Chrysoperla pallida, Chrysoperla nipponensis, Chrysoperla sillemi","14629","10391","10391","1","28.97","0","99.39","436.55","0","437.45","31.24","0"
"AGAW1","08S","INS","BOLD:AAG2511","Arthropoda","Insecta","Diptera","Anthomyiidae","Anthomyiinae","Delia","Delia platura","14527","11599","11599","2","20.16","0","99.04","444.77","0","447.64","33.15","0"
"AGAW1","08S","INS","BOLD:AAA3453","Arthropoda","Insecta","Diptera","Anthomyiidae","Anthomyiinae","Delia","","12222","10360","10360","3","15.23","0","99.29","432.34","0","434.9","34.59","0"
"AGAW1","08S","INS","BOLD:AAG2460","Arthropoda","Insecta","Diptera","Anthomyiidae","Anthomyiinae","Hydrophoria","Hydrophoria lancifer","5267","4292","4292","4","18.51","0","99.2","433.91","0","437.59","33.65","0"
"AGAW1","08S","INS","BOLD:AAH4205","Arthropoda","Insecta","Diptera","Chloropidae","Chloropinae","Meromyza","Meromyza nigriseta, Meromyza sp.","3330","1976","1976","5","40.66","0","99.6","447.39","0","448.25","32.89","0"
"AGAW1","08S","INS","BOLD:AAD2041","Arthropoda","Insecta","Diptera","Dolichopodidae","Sciapodinae","Condylostylus","Condylostylus inermis, Condylostylus nigrofemoratus, Condylostylus connectans, Condylostylus caudatus, Condylostylus flavipes","2543","1818","1818","6","28.51","0","99.26","440.06","0","441.2","36.02","0"
"AGAW1","08S","INS","BOLD:AAP8739","Arthropoda","Insecta","Diptera","Platystomatidae","Platystomatinae","Rivellia","Rivellia steyskali","2382","1674","1674","7","29.72","0","99.15","443.32","0","445.43","37.89","0"
"AGAW1","08S","INS","BOLD:AAA7368","Arthropoda","Insecta","Lepidoptera","Cosmopterigidae","Cosmopteriginae","Limnaecia","Limnaecia phragmitella","1939","1673","1673","8","13.72","0","99.38","427.89","0","432.81","30.99","0"
"AGAW1","08S","INS","BOLD:AAH4135","Arthropoda","Insecta","Diptera","Chloropidae","Chloropinae","Thaumatomyia","Thaumatomyia glabra, Thaumatomyia grata, Thaumatomyia sp. 1 TAW","1824","1681","1681","9","7.84","0","99.26","430.88","0","434.85","31.81","0"
"AGAW1","08S","INS","BOLD:AAB7030","Arthropoda","Insecta","Diptera","Chironomidae","Chironominae","Chironomus","Chironomus sp. BOLDAAP2996, Chironomus quinnitukqut, Chironomus cf. decorus 2g, Chironomus decorus group sp. 2 IP2013, Chironomus cf. decorus, Chironomus decorus gr.","1177","1134","1134","10","3.65","0","98.6","428.63","0","433.6","34.48","0"
"AGAW1","08S","INS","BOLD:AAA2815","Arthropoda","Insecta","Lepidoptera","Noctuidae","Noctuinae","Mythimna","Mythimna yukonensis, Mythimna oxygala","1042","923","923","11","11.42","0","99.24","425.92","0","427.47","33.47","0"
"AGAW1","08S","INS","BOLD:AAG9626","Arthropoda","Insecta","Diptera","Dolichopodidae","","","","722","582","582","12","19.39","0","97.44","433.72","0","435.13","36.65","0"
"AGAW1","08S","INS","BOLD:AAG2868","Arthropoda","Insecta","Hemiptera","Cicadellidae","Typhlocybinae","Empoasca","Empoasca sp., Empoasca fabae","674","462","462","13","31.45","0","99.68","444.49","0","447.91","32.43","0"
"AGAW1","08S","INS","BOLD:AAG2123","Arthropoda","Insecta","Diptera","Tachinidae","Dexiinae","Dinera","Dinera grisescens","638","486","486","14","23.82","0","99.31","435.82","0","437.94","31.87","0"
"AGAW1","08S","INS","BOLD:ACY6524","Arthropoda","Insecta","Diptera","Chironomidae","Orthocladiinae","Psectrocladius","Psectrocladius cf. limbatellus","620","587","587","15","5.32","0","98.64","419","0","427.09","36.01","0"
"AGAW1","08S","INS","BOLD:AAP5169","Arthropoda","Insecta","Diptera","Chloropidae","Oscinellinae","Elachiptera","Elachiptera nigriceps","617","519","519","16","15.88","0","99.04","424.58","0","425.84","31.41","0"
"AGAW1","08S","INS","BOLD:AAA4277","Arthropoda","Insecta","Diptera","Syrphidae","Syrphinae","Toxomerus","Toxomerus marginatus, Toxomerus sp. CR-3, Toxomerus CR-3","614","457","457","17","25.57","0","99.22","446.23","0","447.28","30.88","0"
"AGAW1","08S","INS","BOLD:AAD1395","Arthropoda","Insecta","Diptera","Chironomidae","Chironominae","Polypedilum","Polypedilum convictum, Polypedilum flavum","565","389","389","18","31.15","0","98.57","443.59","0","444.87","33.4","0"
"AGAW1","08S","INS","BOLD:AAD4936","Arthropoda","Insecta","Lepidoptera","Gracillariidae","Gracillariinae","Caloptilia","Caloptilia rhoifoliella, Caloptilia sassafrasella","510","454","454","19","10.98","0","98.34","417.64","0","428.98","29.35","0"
"AGAW1","08S","INS","BOLD:AAG9668","Arthropoda","Insecta","Diptera","Dolichopodidae","Diaphorinae","Chrysotus","Chrysotus sp., Chrysotus sp. nr. obliquus","364","322","322","20","11.54","0","98.81","446.42","0","448.92","32.77","0"
"AGAW1","08S","INS","BOLD:AAN6154","Arthropoda","Insecta","Coleoptera","Latridiidae","Corticariinae","Melanophthalma","Melanophthalma inermis","336","277","277","21","17.56","0","99.18","435.61","0","436.81","31.18","0"
"AGAW1","08S","INS","BOLD:AAL4232","Arthropoda","Insecta","Diptera","Agromyzidae","Phytomyzinae","Phytoliriomyza","Phytoliriomyza arctica","321","242","242","22","24.61","0","99.52","442.93","0","444.24","32.06","0"
"AGAW1","08S","INS","BOLD:AAG7279","Arthropoda","Insecta","Diptera","Sphaeroceridae","Limosininae","Spelobia","Spelobia ochripes","259","197","197","23","23.94","0","99.49","446.56","0","447.84","35.22","0"
"AGAW1","08S","INS","BOLD:AAG5507","Arthropoda","Insecta","Diptera","Chironomidae","Chironominae","Chironomus","Chironomus atrella","229","222","222","24","3.06","0","99.06","429.47","0","432.21","36.05","0"
"AGAW1","08S","INS","BOLD:AAN5199","Arthropoda","Insecta","Diptera","Cecidomyiidae","","","CecidInt20 sp. BOLD:AAN5199","208","165","165","25","20.67","0","99.53","432.08","0","445.71","26.82","0"
"AGAW1","08S","INS","BOLD:AAN5342","Arthropoda","Insecta","Diptera","Chironomidae","Orthocladiinae","Heleniella","","208","202","202","25","2.88","0","99","415.14","0","425.77","34.89","0"
"AGAW1","08S","INS","BOLD:AAB4658","Arthropoda","Insecta","Diptera","Chironomidae","Chironominae","Chironomus","Chironomus dilutus","207","201","201","27","2.9","0","98.41","438.17","0","440.3","37.84","0"
"AGAW1","08S","INS","BOLD:AAA3889","Arthropoda","Insecta","Lepidoptera","Crambidae","Crambinae","Parapediasia","Parapediasia teterrellus","196","186","186","28","5.1","0","99.23","428.42","0","429.7","32.17","0"
"AGAW1","08S","INS","BOLD:ACK7314","Arthropoda","Insecta","Diptera","Chironomidae","Chironominae","Polypedilum","","165","164","164","29","0.61","0","98.45","387.47","0","391.57","35.17","0"
"AGAW1","08S","INS","BOLD:AAG1000","Arthropoda","Insecta","Diptera","Chironomidae","Orthocladiinae","Orthocladius","Orthocladius carlatus","163","156","156","30","4.29","0","98.94","421","0","422.87","33.53","0"
"AGAW1","08S","INS","BOLD:ACX5521","Arthropoda","Insecta","Diptera","Chironomidae","Orthocladiinae","","","139","137","137","31","1.44","0","98.88","415.47","0","417.44","34.96","0"
"AGAW1","08S","INS","BOLD:AAF9771","Arthropoda","Insecta","Diptera","Hybotidae","Tachydromiinae","Platypalpus","Platypalpus longiseta, Platypalpus trivialis, Platypalpus sp. 5, Platypalpus sp. 17, Platypalpus pallidiventris","88","61","61","32","30.68","0","99.62","442.24","0","442.98","35.15","0"
"AGAW1","08S","INS","BOLD:AAC4197","Arthropoda","Insecta","Diptera","Chironomidae","Orthocladiinae","Paraphaenocladius","Paraphaenocladius impensus","88","71","71","32","19.32","0","99.57","432.45","0","437.97","35.05","0"
"AGAW1","08S","INS","BOLD:AAA4555","Arthropoda","Insecta","Orthoptera","Acrididae","Melanoplinae","Melanoplus","Melanoplus stupefactus, Melanoplus angustipennis, Melanoplus ponderosus, Melanoplus bruneri, Melanoplus frigidus, Melanoplus scapularis, Melanoplus cinereus, Melanoplus madeleineae, Melanoplus sanguinipes, Melanoplus nigrescens group, Melanoplus tunicae, Melanoplus walshi, Melanoplus femurrubrum, Melanoplus dawsoni, Melanoplus sp., Melanoplus fasciatus, Melanoplus bowditchi, Melanoplus stonei, Melanoplus bivittatus, Melanoplus huroni, Melanoplus thomasi, Melanoplus borealis, Melanoplus packardii","86","79","79","34","8.14","0","99.29","448.53","0","449.43","35.99","0"
"AGAW1","08S","INS","BOLD:AAG7276","Arthropoda","Insecta","Diptera","Sphaeroceridae","Limosininae","Leptocera","Leptocera erythrocera","82","75","75","35","8.54","0","99.19","437.07","0","438.87","32.86","0"
"AGAW1","08S","INS","BOLD:AAG7891","Arthropoda","Insecta","Hymenoptera","Platygastridae","Scelioninae","Telenomus","Telenomus podisi","66","60","60","36","9.09","0","99.19","431.92","0","441","27.18","0"
"AGAW1","08S","INS","BOLD:ABA7871","Arthropoda","Insecta","Diptera","Cecidomyiidae","","","","60","53","53","37","11.67","0","98.93","437.8","0","438.67","26.83","0"
"AGAW1","08S","INS","BOLD:AAZ5592","Arthropoda","Insecta","Diptera","Psychodidae","Psychodinae","","","58","56","56","38","3.45","0","99.14","425.19","0","427.64","32.4","0"
"AGAW1","08S","INS","BOLD:AAN5238","Arthropoda","Insecta","Diptera","Cecidomyiidae","","","","51","46","46","39","9.8","0","99.4","431.14","0","432.43","25.67","0"
"AGAW1","08S","INS","BOLD:AAN5225","Arthropoda","Insecta","Diptera","Cecidomyiidae","","","","50","38","38","40","24","0","99.38","446.76","0","447.3","28.53","0"
"AGAW1","08S","INS","BOLD:ABW1379","Arthropoda","Insecta","Diptera","Chloropidae","Oscinellinae","Malloewia","Malloewia abdominalis","47","36","36","41","23.4","0","98.29","439.47","0","446.74","31.62","0"
"AGAW1","08S","INS","BOLD:AAN6433","Arthropoda","Insecta","Diptera","Sciaridae","","Corynoptera","Corynoptera deserta","42","41","41","42","2.38","0","99.52","434.48","0","436.79","30.37","0"
"AGAW1","08S","INS","BOLD:ABA0822","Arthropoda","Insecta","Diptera","Cecidomyiidae","","","","41","37","37","43","9.76","0","98.65","438.34","0","439.71","26.66","0"
"AGAW1","08S","INS","BOLD:AAI2681","Arthropoda","Insecta","Diptera","Chironomidae","Orthocladiinae","Parakiefferiella","","40","34","34","44","15","0","99.48","444.28","0","444.93","34.6","0"
"AGAW1","08S","INS","BOLD:AAH3661","Arthropoda","Insecta","Diptera","Cecidomyiidae","","","","40","40","40","44","0","0","99.14","430.45","0","432.75","25.4","0"
"AGAW1","08S","INS","BOLD:ACE6783","","","","","","","","36","33","33","46","8.33","0","99.63","420.69","0","422.58","25.88","0"
"AGAW1","08S","INS","BOLD:AAB4657","Arthropoda","Insecta","Diptera","Chironomidae","Chironominae","Chironomus","Chironomus maturus","34","32","32","47","5.88","0","99.45","434.85","0","437.44","37.47","0"
"AGAW1","08S","INS","BOLD:ACU7906","Arthropoda","Insecta","Diptera","Chironomidae","Orthocladiinae","","","34","34","34","47","0","0","98.92","404.47","0","407.18","36.05","0"
"AGAW1","08S","INS","BOLD:AAM7855","Arthropoda","Insecta","Thysanoptera","Thripidae","Thripinae","Frankliniella","Frankliniella tritici","34","30","30","47","11.76","0","99.59","445.18","0","447.03","31.37","0"
"AGAW1","08S","INS","BOLD:ACA9441","Arthropoda","Insecta","Diptera","Chironomidae","Chironominae","Paratanytarsus","","33","33","33","50","0","0","99.3","413.64","0","415.61","34.37","0"
"AGAW1","08S","INS","BOLD:AAN5220","Arthropoda","Insecta","Diptera","Cecidomyiidae","","","","31","31","31","51","0","0","98.05","393.58","0","417.48","26.3","0"
"AGAW1","08S","INS","BOLD:AAV5575","Arthropoda","Insecta","Diptera","Cecidomyiidae","","","CecidInt30 sp. BOLD:AAV5575","25","20","20","52","20","0","99.72","446.32","0","447.24","25.93","0"
"AGAW1","08S","INS","BOLD:AAA3080","Arthropoda","Insecta","Trichoptera","Limnephilidae","Limnephilinae","Limnephilus","Limnephilus submonilifer","25","25","25","52","0","0","98.95","402.08","0","404.08","33.87","0"
"AGAW1","08S","INS","BOLD:ABA6514","Arthropoda","Insecta","Diptera","Chironomidae","Chironominae","Rheotanytarsus","","24","24","24","54","0","0","98.37","419.46","0","421.17","35.98","0"
"AGAW1","08S","INS","BOLD:AAN5660","Arthropoda","Insecta","Diptera","Chloropidae","Oscinellinae","Incertella","Incertella minor","23","19","19","55","17.39","0","99.49","447.35","0","448.09","32.65","0"
"AGAW1","08S","INS","BOLD:AAV5573","Arthropoda","Insecta","Diptera","Cecidomyiidae","","","","22","19","19","56","13.64","0","99.49","446.45","0","447.41","27.66","0"
"AGAW1","08S","INS","BOLD:AAG2740","Arthropoda","Insecta","Diptera","Ephydridae","Parydrinae","Philygria","Philygria debilis","21","21","21","57","0","0","98.27","402.81","0","407.95","32.5","0"
"AGAW1","08S","INS","BOLD:AAG1711","Arthropoda","Insecta","Diptera","Muscidae","Phaoniinae","Helina","Helina depuncta","19","19","19","58","0","0","99.02","438.11","0","440.47","31.39","0"
"AGAW1","08S","INS","BOLD:ACM7774","Arthropoda","Insecta","Trichoptera","Hydroptilidae","Orthotrichiinae","Orthotrichia","Orthotrichia cristata","19","19","19","58","0","0","99.14","415.21","0","416.95","29.2","0"
"AGAW1","08S","INS","BOLD:AAG9897","Arthropoda","Insecta","Dermaptera","Forficulidae","Forficulinae","Forficula","Forficula auricularia-A","18","18","18","60","0","0","99.29","390.83","0","400.89","45.68","0"
"AGAW1","08S","INS","BOLD:AAG6934","Arthropoda","Insecta","Diptera","Hybotidae","","","","18","17","17","60","5.56","0","99.04","436.72","0","438.44","38.69","0"
"AGAW1","08S","INS","BOLD:AAJ1250","Arthropoda","Insecta","Hymenoptera","Braconidae","Aphidiinae","Aphidius","Aphidius avenaphis, Aphidius uzbekistanicus","16","16","16","62","0","0","99.21","421.38","0","434.06","26.95","0"
"AGAW1","08S","INS","BOLD:AAA9422","Arthropoda","Insecta","Hemiptera","Cicadellidae","Deltocephalinae","Macrosteles","Macrosteles quadrilineatus","16","16","16","62","0","0","98.67","408.06","0","412","35.88","0"
"AGAW1","08S","INS","BOLD:AAN5267","Arthropoda","Insecta","Diptera","Cecidomyiidae","","","","15","15","15","64","0","0","99.11","449.27","0","450","30.67","0"
"AGAW1","08S","INS","BOLD:AAH0152","Arthropoda","Insecta","Coleoptera","Cantharidae","Cantharinae","Cantharis","Cantharis rufa","13","13","13","65","0","0","98.26","426.85","0","430.08","35.21","0"
"AGAW1","08S","INS","BOLD:AAP6790","Arthropoda","Insecta","Diptera","Chloropidae","Oscinellinae","Incertella","Incertella incerta","11","11","11","66","0","0","99.33","432.45","0","434.36","31.34","0"
"AGAW1","08S","INS","BOLD:AAN6447","Arthropoda","Insecta","Diptera","Sciaridae","","Corynoptera","Corynoptera perpusilla","10","8","8","67","20","0","99.42","447.8","0","450","36.31","0"
"AGAW1","08S","INS","BOLD:ACP4736","Arthropoda","Insecta","Diptera","Chironomidae","Orthocladiinae","Smittia","Smittia sp. 8ES","8","7","7","68","12.5","0","98.5","425.5","0","428.13","32.55","0"
"AGAW1","08S","INS","BOLD:ACJ6984","Arthropoda","Insecta","Hymenoptera","Platygastridae","Scelioninae","Trissolcus","Trissolcus thyantae","8","7","7","68","12.5","0","99.26","438.5","0","450","27.36","0"
"AGAW1","08S","INS","BOLD:ABU5525","Arthropoda","Insecta","Diptera","Chironomidae","Orthocladiinae","Limnophyes","Limnophyes sp. 14ES","8","8","8","68","0","0","98.9","410.13","0","412.13","37.51","0"
"AGAW1","08S","INS","BOLD:AAG6745","Arthropoda","Insecta","Diptera","Calliphoridae","Calliphorinae","Pollenia","Pollenia pediculata","8","8","8","68","0","0","99.6","415.88","0","417.25","33.61","0"
"AGAW1","08S","INS","BOLD:AAF5705","Arthropoda","Insecta","Hemiptera","Miridae","Orthotylinae","Blepharidopterus","Blepharidopterus chlorionis","8","7","7","68","12.5","0","99.65","361.13","0","449","42.9","0"
"AGAW1","08S","INS","BOLD:AAC0903","Arthropoda","Insecta","Diptera","Chironomidae","Chironominae","Chironomus","Chironomus acidophilus","7","7","7","73","0","0","99.29","436.43","0","436.71","36.49","0"
"AGAW1","08S","INS","BOLD:AAN5241","Arthropoda","Insecta","Diptera","Cecidomyiidae","Cecidomyiinae","Vitisiella","Vitisiella brevicauda","5","5","5","74","0","0","98.74","440.6","0","450","28.18","0"
"AGAW1","08S","INS","BOLD:AAH0134","Arthropoda","Insecta","Coleoptera","Phalacridae","Phalacrinae","Stilbus","Stilbus apicialis","5","5","5","74","0","0","99.31","422.4","0","423.6","31.36","0"
"AGAW1","08S","INS","BOLD:ABX4944","","","","","","","","5","5","5","74","0","0","97.7","391.2","0","395","34.09","0"
"AGAW1","08S","INS","BOLD:ACK1360","Arthropoda","Insecta","Diptera","Cecidomyiidae","Cecidomyiinae","Vitisiella","Vitisiella brevicauda","5","5","5","74","0","0","98.22","435.6","0","437.4","27.77","0"
"AGAW1","08S","INS","BOLD:AAN5183","Arthropoda","Insecta","Diptera","Cecidomyiidae","","","","5","5","5","74","0","0","98.39","448.2","0","449.4","27.86","0"
"AGAW1","08S","INS","BOLD:ABA2981","Arthropoda","Insecta","Thysanoptera","Aeolothripidae","Aeolothripinae","Aeolothrips","Aeolothrips ericae","4","4","4","79","0","0","98.9","412.75","0","413.5","36.57","0"
"AGAW1","08S","INS","BOLD:AAA5299","Arthropoda","Insecta","Diptera","Chironomidae","Orthocladiinae","Cricotopus","Cricotopus laetus, Cricotopus relucens, Cricotopus trifasciatus, Cricotopus sp., Cricotopus sp. 3ES, Cricotopus sylvestris, Cricotopus glacialis","4","4","4","79","0","0","98.88","445.75","0","448","34.66","0"
"AGAW1","08S","INS","BOLD:AAP5870","Arthropoda","Insecta","Diptera","Chironomidae","Chironominae","Tanytarsus","","4","4","4","79","0","0","97.87","409","0","411.25","36.37","0"
"AGAW1","08S","INS","BOLD:AAU6327","Arthropoda","Insecta","Diptera","Chironomidae","Orthocladiinae","Psectrocladius","Psectrocladius schlienzi","4","4","4","79","0","0","97.39","421.25","0","423","36.13","0"
"AGAW1","08S","INS","BOLD:AAB9140","Arthropoda","Insecta","Diptera","Calliphoridae","Chrysomyinae","Phormia","Phormia regina","3","3","3","83","0","0","99.78","449.33","0","450","33.33","0"
"AGAW1","08S","INS","BOLD:AAH3968","Arthropoda","Insecta","Diptera","Sciaridae","","Cratyna","Cratyna cryptospina","3","3","3","83","0","0","99.11","434.33","0","437.67","26.7","0"
"AGAW1","08S","INS","BOLD:AAG8683","Arthropoda","Insecta","Hemiptera","Cicadellidae","Eurymelinae, Typhlocybinae","Empoasca, Idiocerus","Empoasca luda, Idiocerus populi","3","3","3","83","0","0","97.28","428.67","0","429.33","30.35","0"
"AGAW1","08S","INS","BOLD:AAA6759","Arthropoda","Insecta","Lepidoptera","Hesperiidae","Hesperiinae","Thymelicus","Thymelicus lineola","2","2","2","86","0","0","99.56","450","0","450","30.33","0"
"AGAW1","08S","INS","BOLD:AAA8108","Arthropoda","Insecta","Lepidoptera","Noctuidae","Acontiinae","Ponometia","Ponometia erastrioides","2","2","2","86","0","0","98.1","448.5","0","450","32.78","0"
"AGAW1","08S","INS","BOLD:AAH7146","Arthropoda","Insecta","Diptera","Sarcophagidae","Sarcophaginae","Boettcheria","Boettcheria latisterna","2","2","2","86","0","0","98.92","431.5","0","433.5","34.6","0"
"AGAW1","08S","INS","BOLD:AAC9132","Arthropoda","Insecta","Diptera","Culicidae","Culicinae","Culiseta","Culiseta inornata","2","2","2","86","0","0","98.54","446.5","0","450","32.89","0"
"AGAW1","08S","INS","BOLD:AAL7819","Arthropoda","Insecta","Diptera","Psychodidae","Psychodinae","Psychoda","Psychoda sp., Psychoda sp. BOLD:AAL7819","2","2","2","86","0","0","99.48","387","0","390","32.3","0"
"AGAW1","08S","INS","BOLD:AAA4188","Arthropoda","Insecta","Hymenoptera","Braconidae","Aphidiinae","Aphidius","Aphidius microlophii, Aphidius ervi","2","2","2","86","0","0","99.11","449","0","450","26.67","0"
"AGAW1","08S","INS","BOLD:ACF2821","Arthropoda","Insecta","Diptera","Mycetophilidae","Mycetophilinae","Mycetophila","Mycetophila fungorum","2","2","2","86","0","0","99.31","435.5","0","437","35.92","0"
"AGAW1","08S","INS","BOLD:AAI4303","Arthropoda","Insecta","Diptera","Chironomidae","Chironominae","Chironomus","Chironomus melanescens","2","2","2","86","0","0","98.39","435.5","0","438.5","34.32","0"
"AGAW1","08S","INS","BOLD:AAH4208","Arthropoda","Insecta","Diptera","Chloropidae","Oscinellinae","Elachiptera","Elachiptera erythropleura, Elachiptera costata","2","2","2","86","0","0","98.56","450","0","450","32.33","0"
"AGAW1","08S","INS","BOLD:AAA6292","Arthropoda","Insecta","Lepidoptera","Tortricidae","Tortricinae","Cnephasia","Cnephasia genitalana, Cnephasia stephensiana","1","1","1","95","0","0","100","449","0","450","35.78","0"
"AGAW1","08S","INS","BOLD:AAG1714","Arthropoda","Insecta","Diptera","Muscidae","Azeliinae","Muscina","Muscina pascuorum","1","1","1","95","0","0","100","430","0","450","34.22","0"
"AGAW1","08S","INS","BOLD:ACE0699","Arthropoda","Insecta","Diptera","Chironomidae","","","","1","1","1","95","0","0","97.91","431","0","450","35.78","0"
"AGAW1","08S","INS","BOLD:ACV1953","Arthropoda","Insecta","Diptera","Cecidomyiidae","","","","1","1","1","95","0","0","99.33","448","0","450","29.56","0"
"AGAW1","08S","INS","BOLD:AAN5497","Arthropoda","Insecta","Diptera","Anthomyiidae","Pegomyinae","Pegomya","Pegomya solennis, Pegomya bicolor","1","1","1","95","0","0","100","449","0","450","29.33","0"
"AGAW1","08S","INS","BOLD:ABY9688","Arthropoda","Insecta","Diptera","Chloropidae","Chloropinae","Thaumatomyia","Thaumatomyia glabra","1","1","1","95","0","0","97.54","448","0","450","32.22","0"
"AGAW1","08S","INS","BOLD:ACR4394","Arthropoda","Insecta","Diptera","Anthomyiidae","Anthomyiinae","Delia","Delia florilega","1","1","1","95","0","0","100","404","0","406","32.02","0"
"AGAW1","08S","INS","BOLD:AAN5255","Arthropoda","Insecta","Diptera","Cecidomyiidae","Cecidomyiinae","Dasineura","","1","1","1","95","0","0","98.44","448","0","450","26.89","0"
"AGAW1","08S","INS","BOLD:ADF3809","Arthropoda","Insecta","Diptera","Dolichopodidae","","","","1","1","1","95","0","0","97.62","420","0","422","34.83","0"
"AGAW1","08S","INS","BOLD:ABX9225","Arthropoda","Insecta","Diptera","Limoniidae","Limoniinae","Rhipidia","","1","1","1","95","0","0","98.98","392","0","393","32.57","0"
"AGAW1","08S","INS","BOLD:AAN5230","Arthropoda","Insecta","Diptera","Cecidomyiidae","","","","1","1","1","95","0","0","98.44","450","0","450","26.89","0"
"AGAW1","08S","INS","BOLD:AAH3914","Arthropoda","Insecta","Diptera","Sciaridae","","Sciara","Sciara hamatilis, Sciara humeralis","1","1","1","95","0","0","99.3","426","0","427","30.68","0"
"AGAW1","08S","INS","BOLD:AAZ5973","Arthropoda","Insecta","Diptera","Sepsidae","Sepsinae","Themira","Themira annulipes","1","1","1","95","0","0","98.99","396","0","399","35.59","0"
"AGAW1","08S","INS","BOLD:AAZ5504","Arthropoda","Insecta","Diptera","Dolichopodidae","","","","1","1","1","95","0","0","97.11","450","0","450","34.44","0"
"AGAW1","08S","INS","BOLD:AAU6681","Arthropoda","Insecta","Diptera","Tipulidae","Tipulinae","Nephrotoma","Nephrotoma cornicina","1","1","1","95","0","0","99.32","442","0","442","36.2","0"
"AGAW1","08S","INS","BOLD:ACF6513","Arthropoda","Insecta","Diptera","Psychodidae","Psychodinae","Psychoda","Psychoda cinerea","1","1","1","95","0","0","99.56","450","0","450","30.89","0"
"AGAW1","08S","INS","BOLD:AAQ0275","Arthropoda","Insecta","Diptera","Sciaridae","","Bradysia","Bradysia lembkei","1","1","1","95","0","0","100","425","0","427","33.49","0"
"AGAW1","08S","INS","BOLD:AAA2632","Arthropoda","Insecta","Lepidoptera","Noctuidae","Noctuinae","Noctua","Noctua atlantica, Noctua pronuba","1","1","1","95","0","0","99.55","449","0","450","31.11","0"
"AGAW1","08S","INS","BOLD:ACE7444","Arthropoda","Insecta","Hemiptera","Miridae","Mirinae, Phylinae","Adelphocoris, Megalocoleus","Adelphocoris lineolatus, Megalocoleus molliculus","1","1","1","95","0","0","97.17","424","0","425","33.88","0"
"AGAW1","08S","INS","BOLD:ACJ6954","Arthropoda","Insecta","Diptera","Cecidomyiidae","","","","1","1","1","95","0","0","99.7","338","0","450","26.89","0"
"AGAW1","08S","INS","BOLD:ACY2979","","","","","","","","1","1","1","95","0","0","98.36","366","0","367","36.24","0"
"AGAW1","08S","INS","BOLD:ACV5892","Arthropoda","Insecta","Diptera","Cecidomyiidae","","","","1","1","1","95","0","0","99.01","403","0","407","23.83","0"
"AGAW1","08S","INS","BOLD:AAD5253","Arthropoda","Insecta","Diptera","Chironomidae","Orthocladiinae","Thienemanniella","Thienemanniella xena, Thienemanniella vittata","1","1","1","95","0","0","98.66","448","0","450","34.22","0"
"AGAW1","08S","INS","BOLD:ADF6661","Arthropoda","Insecta","Diptera","Cecidomyiidae","","","","1","1","1","95","0","0","97.14","420","0","450","28.89","0"
"AGAW1","08S","INS","BOLD:AAF9187","Arthropoda","Insecta","Coleoptera","Curculionidae","Entiminae","Phyllobius","Phyllobius oblongus","1","1","1","95","0","0","99.78","450","0","450","38.89","0"
"AGAW1","08S","INS","BOLD:AAH4180","Arthropoda","Insecta","Diptera","Chloropidae","Oscinellinae","Rhopalopterum","","1","1","1","95","0","0","98.38","431","0","450","31.78","0"
"AGAW1","08S","INS","BOLD:AAD4214","Arthropoda","Insecta","Hymenoptera","Ichneumonidae","Diplazontinae","Diplazon","Diplazon laetatorius","1","1","1","95","0","0","100","449","0","450","25.78","0"
"AGAW1","08S","INS","BOLD:AAB1148","Arthropoda","Insecta","Diptera","Chloropidae","Oscinellinae","Oscinella","Oscinella sp. TAW3, Oscinella sp. 4, Oscinella sp. BOLD:AAB1148","1","1","1","95","0","0","98.89","450","0","450","33.11","0"
"AGAW1","08S","INS","BOLD:AAB1098","Arthropoda","Insecta","Diptera","Culicidae","Culicinae","Ochlerotatus, Aedes","Aedes excrucians, Aedes cantans, Aedes stimulans, Aedes euedes, Aedes riparius, Aedes annulipes, Ochlerotatus cantans, Aedes fitchii, Aedes sp., Ochlerotatus annulipes, Ochlerotatus excrucians","1","1","1","95","0","0","99.11","447","0","450","35.33","0"
"AGAW1","08S","INS","BOLD:ABY5318","Arthropoda","Insecta","Diptera","Chloropidae","Oscinellinae","Incertella, Rhopalopterum","Rhopalopterum umbrosum, Rhopalopterum luteiceps, Incertella bispina, Rhopalopterum painteri","1","1","1","95","0","0","98.21","447","0","450","32.22","0"
"AGAW1","08S","INS","BOLD:ADH9867","Arthropoda","Insecta","Diptera","","","","","1","1","1","95","0","0","98.35","364","0","366","37.43","0"
"AGAW1","08S","INS","BOLD:AAA3654","Arthropoda","Insecta","Lepidoptera","Crambidae","Crambinae","Crambus","Crambus bigelovi, Crambus johnsoni, Crambus praefectellus","1","1","1","95","0","0","99.33","448","0","450","31.56","0"
"AGAW1","08S","INS","BOLD:ACC5677","Arthropoda","Insecta","Diptera","Cecidomyiidae","Cecidomyiinae","Asteromyia","","1","1","1","95","0","0","99.78","450","0","450","26.67","0"
"AGAW1","08S","INS","BOLD:AAD8086","Arthropoda","Insecta","Lepidoptera","Elachistidae","Elachistinae","Elachista","Elachista griseicornis, Elachista acenteta","1","1","1","95","0","0","99.52","419","0","420","29.76","0"
"AGAW1","08S","INS","BOLD:ACB0984","Arthropoda","Insecta","Psocodea","Stenopsocidae","","Graphopsocus","Graphopsocus cruciatus","1","1","1","95","0","0","99.76","413","0","450","37.78","0"
"AGAW1","08S","INS","BOLD:AAE7016","Arthropoda","Insecta","Lepidoptera","Gelechiidae","Dichomeridinae","Helcystogramma","Helcystogramma hystricella","1","1","1","95","0","0","97.1","449","0","450","31.11","0"
"AGAW1","08S","INS","BOLD:ACL3713","Arthropoda","Insecta","Diptera","Cecidomyiidae","","","CecidInt12 sp. BOLD:ACL3713","1","1","1","95","0","0","99.55","448","0","450","27.78","0"
"AGAW1","08S","INS","BOLD:AAA7661","Arthropoda","Insecta","Diptera","Culicidae","Culicinae","Culiseta, Culex","Culiseta melanura, Culex articularis, Culex restuans, Culex acharistus","1","1","1","95","0","0","100","365","0","367","33.24","0"
"AGAW1","08S","INS","BOLD:AAA5953","Arthropoda","Insecta","Lepidoptera","Gelechiidae","Gelechiinae","Coleotechnites","Coleotechnites sp., Coleotechnites piceaella, Coleotechnites laricis, Coleotechnites sp. 3, Coleotechnites carbonaria, Coleotechnites granti, Coleotechnites macleodi, Coleotechnites gibsonella, Coleotechnites coniferella, Coleotechnites martini","1","1","1","95","0","0","100","450","0","450","34.44","0"
"AGAW1","08S","INS","BOLD:AAA7374","Arthropoda","Insecta","Diptera","Syrphidae","Syrphinae","Sphaerophoria","Sphaerophoria bankowskae, Sphaerophoria infuscata, Sphaerophoria nr. philanthus, Sphaerophoria batava, Sphaerophoria sp., Sphaerophoria rueppellii, Sphaerophoria interrupta-group, Sphaerophoria sp. MA-1, Sphaerophoria cleoae, Sphaerophoria asymmetrica, Sphaerophoria fatarum, Sphaerophoria bengalensis, Sphaerophoria bifurcata, Sphaerophoria taeniata, Sphaerophoria interrupta, Sphaerophoria contigua, Sphaerophoria kaa, Sphaerophoria laurae, Sphaerophoria chongjini, Sphaerophoria boreoalpina, Sphaerophoria sp. BOLDAAA7374, Sphaerophoria macrogaster, Sphaerophoria sulphuripes, Sphaerophoria brevipilosa, Sphaerophoria philanthus, Sphaerophoria pyrrhina, Sphaerophoria abbreviata, Sphaerophoria scripta, Sphaerophoria potentillae, Sphaerophoria longipilosa, Sphaerophoria virgata","1","1","1","95","0","0","100","448","0","450","30.89","0"
"AGAW1","08S","INS","BOLD:ACF5830","Arthropoda","Insecta","Diptera","Chloropidae","Chloropinae","Thaumatomyia","","1","1","1","95","0","0","98.22","450","0","450","32","0"
"AGAW1","08S","NAR","no match","no match","no match","no match","no match","no match","no match","no match","no match","no match","no match","no match","no match","no match","no match","no match","no match","no match","no match","no match"
"AGAW1","08S","NIN","BOLD:AAA7242","Arthropoda","Collembola","Entomobryomorpha","Entomobryidae","Seirinae, Entomobryinae","Entomobrya, Lepidosira","Entomobrya multifasciata, Lepidosira sp.","300","276","276","1","8","0","99.33","428.96","0","431.92","39.33","0"
"AGAW1","08S","NIN","BOLD:AAB7914","Arthropoda","Collembola","Symphypleona","Bourletiellidae","","Bourletiella","","23","22","22","2","4.35","0","99.38","436.35","0","437.78","37.99","0"
"AGAW1","08S","NIN","BOLD:AAF9234","Arthropoda","Arachnida","Trombidiformes","Eupodidae","","","","3","3","3","3","0","0","99.63","447","0","450","29.93","0"
"AGAW1","08S","NIN","BOLD:AAB3836","Arthropoda","Arachnida","Araneae","Philodromidae","","Philodromus","Philodromus cespitum","2","2","2","4","0","0","97.46","394","0","426","30.25","0"
"AGAW1","08S","NIN","BOLD:ACX8969","Arthropoda","Arachnida","Trombidiformes","Eupodidae","","","","2","2","2","4","0","0","98.48","423","0","426","27.56","0"
"AGAW1","08S","NIN","BOLD:AAB4300","Arthropoda","Arachnida","Araneae","Thomisidae","","Xysticus","Xysticus emertoni","1","1","1","6","0","0","98.44","448","0","450","32","0"
"AGAW1","08S","NIN","BOLD:AAH0900","Arthropoda","Ostracoda","Podocopida","","","","","1","1","1","6","0","0","98.57","421","0","428","40.89","0"
"AGAW1","08S","NIN","BOLD:AAM3234","Arthropoda","Collembola","Entomobryomorpha","Entomobryidae","Seirinae","Lepidosira","","1","1","1","6","0","0","97.55","449","0","450","40.22","0"
"AGAW1","08W","BAC","TAX:91391","Proteobacteria","Alphaproteobacteria","Rickettsiales","",NA,"","","41","30","30","1","26.83","0","99.31","407.37","0","443.02","40.6","0"
"AGAW1","08W","BAC","TAX:91393","Proteobacteria","Alphaproteobacteria","Rickettsiales","Rickettsiaceae",NA,"Wolbachia","","1","1","1","2","0","0","99.74","392","0","441","39.68","0"
"AGAW1","08W","BAC","TAX:175768","Proteobacteria","Alphaproteobacteria","Rickettsiales","Rickettsiaceae",NA,"Wolbachia","Wolbachia pipientis","1","1","1","2","0","0","98.89","449","0","450","39.56","0"
"AGAW1","08W","CHO","no match","no match","no match","no match","no match","no match","no match","no match","no match","no match","no match","no match","no match","no match","no match","no match","no match","no match","no match","no match"
"AGAW1","08W","INS","BOLD:AAG1711","Arthropoda","Insecta","Diptera","Muscidae","Phaoniinae","Helina","Helina depuncta","24758","18681","18681","1","24.55","0","99.46","433.53","0","435.79","31.36","0"
"AGAW1","08W","INS","BOLD:AAA3654","Arthropoda","Insecta","Lepidoptera","Crambidae","Crambinae","Crambus","Crambus bigelovi, Crambus johnsoni, Crambus praefectellus","9313","7292","7292","2","21.7","0","99.05","437.65","0","439.27","31.53","0"
"AGAW1","08W","INS","BOLD:ACE7444","Arthropoda","Insecta","Hemiptera","Miridae","Mirinae, Phylinae","Adelphocoris, Megalocoleus","Adelphocoris lineolatus, Megalocoleus molliculus","8093","4799","4799","3","40.7","0","99.7","441.67","0","442.34","34.8","0"
"AGAW1","08W","INS","BOLD:AAA3453","Arthropoda","Insecta","Diptera","Anthomyiidae","Anthomyiinae","Delia","","7569","6610","6610","4","12.67","0","99.36","430.69","0","433.28","34.65","0"
"AGAW1","08W","INS","BOLD:AAG2868","Arthropoda","Insecta","Hemiptera","Cicadellidae","Typhlocybinae","Empoasca","Empoasca sp., Empoasca fabae","4937","2849","2849","5","42.29","0","99.47","443.51","0","447.69","32.24","0"
"AGAW1","08W","INS","BOLD:AAG9626","Arthropoda","Insecta","Diptera","Dolichopodidae","","","","4755","3172","3172","6","33.29","0","98.14","441.26","0","442.63","36.1","0"
"AGAW1","08W","INS","BOLD:AAG6745","Arthropoda","Insecta","Diptera","Calliphoridae","Calliphorinae","Pollenia","Pollenia pediculata","4690","4050","4050","7","13.65","0","99.27","433.4","0","435.49","33.48","0"
"AGAW1","08W","INS","BOLD:AAP8739","Arthropoda","Insecta","Diptera","Platystomatidae","Platystomatinae","Rivellia","Rivellia steyskali","3710","2521","2521","8","32.05","0","99.27","441.46","0","443.5","37.96","0"
"AGAW1","08W","INS","BOLD:AAO3939","Arthropoda","Insecta","Diptera","Limoniidae","Limoniinae","Dicranomyia","","3219","2395","2395","9","25.6","0","99.26","439.61","0","441","36.23","0"
"AGAW1","08W","INS","BOLD:ACB0984","Arthropoda","Insecta","Psocodea","Stenopsocidae","","Graphopsocus","Graphopsocus cruciatus","2967","1857","1857","10","37.41","0","99.59","412.44","0","448.8","37.47","0"
"AGAW1","08W","INS","BOLD:AAA8108","Arthropoda","Insecta","Lepidoptera","Noctuidae","Acontiinae","Ponometia","Ponometia erastrioides","2767","2240","2240","11","19.05","0","98.94","432.98","0","434.08","32.33","0"
"AGAW1","08W","INS","BOLD:AAG6435","Arthropoda","Insecta","Diptera","Ceratopogonidae","Forcipomyiinae","Forcipomyia","","2756","1787","1787","12","35.16","0","98.29","399.94","0","447.22","32.56","0"
"AGAW1","08W","INS","BOLD:AAG2511","Arthropoda","Insecta","Diptera","Anthomyiidae","Anthomyiinae","Delia","Delia platura","2693","2370","2370","13","11.99","0","98.96","444.19","0","447.02","33.15","0"
"AGAW1","08W","INS","BOLD:AAD4214","Arthropoda","Insecta","Hymenoptera","Ichneumonidae","Diplazontinae","Diplazon","Diplazon laetatorius","2645","1948","1948","14","26.35","0","99.48","445.87","0","447.33","25.82","0"
"AGAW1","08W","INS","BOLD:ABW1379","Arthropoda","Insecta","Diptera","Chloropidae","Oscinellinae","Malloewia","Malloewia abdominalis","2512","2008","2008","15","20.06","0","97.95","439.98","0","443.68","31.59","0"
"AGAW1","08W","INS","BOLD:AAH0131","Arthropoda","Insecta","Coleoptera","Nitidulidae","Cryptarchinae","Cryptarcha","Cryptarcha ampla","1769","1253","1253","16","29.17","0","99.5","402.79","0","446.46","38.53","0"
"AGAW1","08W","INS","BOLD:AAA4277","Arthropoda","Insecta","Diptera","Syrphidae","Syrphinae","Toxomerus","Toxomerus marginatus, Toxomerus sp. CR-3, Toxomerus CR-3","970","704","704","17","27.42","0","99.42","444.5","0","445.35","30.65","0"
"AGAW1","08W","INS","BOLD:AAN6154","Arthropoda","Insecta","Coleoptera","Latridiidae","Corticariinae","Melanophthalma","Melanophthalma inermis","807","673","673","18","16.6","0","98.82","437.28","0","438.41","31.29","0"
"AGAW1","08W","INS","BOLD:AAG0999","Arthropoda","Insecta","Diptera","Chironomidae","Orthocladiinae","","","539","399","399","19","25.97","0","97.79","442.21","0","443.29","35.51","0"
"AGAW1","08W","INS","BOLD:AAP6790","Arthropoda","Insecta","Diptera","Chloropidae","Oscinellinae","Incertella","Incertella incerta","463","405","405","20","12.53","0","99.23","437.78","0","439.44","31.39","0"
"AGAW1","08W","INS","BOLD:ABW8037","Arthropoda","Insecta","Diptera","Phoridae","Metopininae","Megaselia","","343","290","290","21","15.45","0","99.5","439.85","0","441.73","35.19","0"
"AGAW1","08W","INS","BOLD:ACX5521","Arthropoda","Insecta","Diptera","Chironomidae","Orthocladiinae","","","337","322","322","22","4.45","0","98.85","413.42","0","415.53","34.98","0"
"AGAW1","08W","INS","BOLD:ADE0510","","","","","","","","307","307","307","23","0","0","97.75","389.85","0","419.26","35.82","0"
"AGAW1","08W","INS","BOLD:AAM7657","","","","","","","","292","246","246","24","15.75","0","99.18","426.03","0","427.42","34.33","0"
"AGAW1","08W","INS","BOLD:ACU7906","Arthropoda","Insecta","Diptera","Chironomidae","Orthocladiinae","","","260","255","255","25","1.92","0","99.09","380.13","0","408.04","37.28","0"
"AGAW1","08W","INS","BOLD:AAC0706","Arthropoda","Insecta","Diptera","Chironomidae","Chironominae","Dicrotendipes","Dicrotendipes tritomus","242","242","242","26","0","0","97.68","396.2","0","398.94","35.84","0"
"AGAW1","08W","INS","BOLD:AAN5199","Arthropoda","Insecta","Diptera","Cecidomyiidae","","","CecidInt20 sp. BOLD:AAN5199","218","175","175","27","19.72","0","99.55","437.92","0","447.58","26.71","0"
"AGAW1","08W","INS","BOLD:ABA7871","Arthropoda","Insecta","Diptera","Cecidomyiidae","","","","177","159","159","28","10.17","0","99.08","440.35","0","441.21","26.74","0"
"AGAW1","08W","INS","BOLD:AAG9897","Arthropoda","Insecta","Dermaptera","Forficulidae","Forficulinae","Forficula","Forficula auricularia-A","117","117","117","29","0","0","99.09","397.64","0","409.15","45.82","0"
"AGAW1","08W","INS","BOLD:AAQ0296","Arthropoda","Insecta","Diptera","Cecidomyiidae","","","","105","94","94","30","10.48","0","99.3","436","0","437.83","27.68","0"
"AGAW1","08W","INS","BOLD:AAM6037","Arthropoda","Insecta","Diptera","Cecidomyiidae","Cecidomyiinae","Contarinia, Dasineura","","89","79","79","31","11.24","0","99.53","437.35","0","442.49","26.55","0"
"AGAW1","08W","INS","BOLD:AAN5183","Arthropoda","Insecta","Diptera","Cecidomyiidae","","","","75","63","63","32","16","0","98.58","445.51","0","446.31","27.71","0"
"AGAW1","08W","INS","BOLD:AAG6934","Arthropoda","Insecta","Diptera","Hybotidae","","","","61","61","61","33","0","0","99.11","415.84","0","417.67","38.41","0"
"AGAW1","08W","INS","BOLD:AAM7855","Arthropoda","Insecta","Thysanoptera","Thripidae","Thripinae","Frankliniella","Frankliniella tritici","51","45","45","34","11.76","0","99.36","436.53","0","438.88","31.2","0"
"AGAW1","08W","INS","BOLD:AAV5573","Arthropoda","Insecta","Diptera","Cecidomyiidae","","","","34","33","33","35","2.94","0","99.7","442.91","0","444.29","27.68","0"
"AGAW1","08W","INS","BOLD:AAN5225","Arthropoda","Insecta","Diptera","Cecidomyiidae","","","","31","22","22","36","29.03","0","99.3","449.13","0","450","28.41","0"
"AGAW1","08W","INS","BOLD:AAH3661","Arthropoda","Insecta","Diptera","Cecidomyiidae","","","","28","28","28","37","0","0","99.05","432","0","433.32","25.51","0"
"AGAW1","08W","INS","BOLD:ACC2901","Arthropoda","Insecta","Diptera","Chironomidae","Chironominae","","","24","24","24","38","0","0","97.99","424.67","0","436.75","36.73","0"
"AGAW1","08W","INS","BOLD:AAD5253","Arthropoda","Insecta","Diptera","Chironomidae","Orthocladiinae","Thienemanniella","Thienemanniella xena, Thienemanniella vittata","23","18","18","39","21.74","0","98.86","442.3","0","443.3","34.19","0"
"AGAW1","08W","INS","BOLD:AAC0903","Arthropoda","Insecta","Diptera","Chironomidae","Chironominae","Chironomus","Chironomus acidophilus","19","16","16","40","15.79","0","99.62","444.63","0","448.21","36.46","0"
"AGAW1","08W","INS","BOLD:ABV9322","Arthropoda","Insecta","Diptera","Cecidomyiidae","","","","18","18","18","41","0","0","98.97","440.39","0","441","27.12","0"
"AGAW1","08W","INS","BOLD:AAH3947","Arthropoda","Insecta","Diptera","Sciaridae","","Bradysia","Bradysia trivittata","18","18","18","41","0","0","99.02","426.83","0","428.22","34.34","0"
"AGAW1","08W","INS","BOLD:AAB4894","Arthropoda","Insecta","Hemiptera","Aphididae","Aphidinae","Sitobion","Sitobion smilacifoliae, Sitobion avenae, Sitobion phyllanthi","14","14","14","43","0","0","97.91","443.79","0","444.79","25.54","0"
"AGAW1","08W","INS","BOLD:ACF1257","","","","","","","","13","10","10","44","23.08","0","99.59","449.62","0","450","34.67","0"
"AGAW1","08W","INS","BOLD:AAH0135","Arthropoda","Insecta","Coleoptera","Phalacridae","Phalacrinae","Acylomus","Acylomus pugetanus, Acylomus ergoti","13","12","12","44","7.69","0","98.45","433.77","0","435.15","31.68","0"
"AGAW1","08W","INS","BOLD:AAB0373","Arthropoda","Insecta","Neuroptera","Chrysopidae","Chrysopinae","Chrysoperla","Chrysoperla sp. isolate 82, Chrysoperla zastrowi, Chrysoperla plorabunda, Chrysoperla sp. isolate 71, Chrysoperla agilis, Italochrysa italica, Chrysoperla sp. isolate 88, Chrysoperla lucasina, Chrysoperla carnea, Chrysoperla mediterranea, Chrysoperla pallida, Chrysoperla nipponensis, Chrysoperla sillemi","10","9","9","46","10","0","98.97","429.5","0","429.9","31.71","0"
"AGAW1","08W","INS","BOLD:AAG7891","Arthropoda","Insecta","Hymenoptera","Platygastridae","Scelioninae","Telenomus","Telenomus podisi","8","8","8","47","0","0","98.93","433.75","0","442.5","27.1","0"
"AGAW1","08W","INS","BOLD:AAU6681","Arthropoda","Insecta","Diptera","Tipulidae","Tipulinae","Nephrotoma","Nephrotoma cornicina","8","7","7","47","12.5","0","99.48","438.63","0","440.13","36.35","0"
"AGAW1","08W","INS","BOLD:AAF9035","Arthropoda","Insecta","Diptera","Tipulidae","Tipulinae","Nephrotoma","Nephrotoma alterna","7","7","7","49","0","0","98.87","430.71","0","433.29","33.56","0"
"AGAW1","08W","INS","BOLD:AAG2740","Arthropoda","Insecta","Diptera","Ephydridae","Parydrinae","Philygria","Philygria debilis","7","7","7","49","0","0","98.04","433.29","0","437.43","33.74","0"
"AGAW1","08W","INS","BOLD:ABA1215","Arthropoda","Insecta","Diptera","Sciaridae","","Lycoriella","Lycoriella sativae","6","3","3","51","50","0","99.91","440.17","0","440.33","35.82","0"
"AGAW1","08W","INS","BOLD:AAA5299","Arthropoda","Insecta","Diptera","Chironomidae","Orthocladiinae","Cricotopus","Cricotopus laetus, Cricotopus relucens, Cricotopus trifasciatus, Cricotopus sp., Cricotopus sp. 3ES, Cricotopus sylvestris, Cricotopus glacialis","5","5","5","52","0","0","99.05","434","0","435.6","35.13","0"
"AGAW1","08W","INS","BOLD:AAA2815","Arthropoda","Insecta","Lepidoptera","Noctuidae","Noctuinae","Mythimna","Mythimna yukonensis, Mythimna oxygala","5","5","5","52","0","0","99.37","414.2","0","415.8","33.59","0"
"AGAW1","08W","INS","BOLD:AAA6560","Arthropoda","Insecta","Lepidoptera","Crambidae","Crambinae","Crambus","Crambus multilinellus, Crambus saltuellus","4","4","4","54","0","0","98.89","435.25","0","436.75","30.89","0"
"AGAW1","08W","INS","BOLD:AAH3320","Arthropoda","Insecta","Coleoptera","Coccinellidae","Coccinellinae","Psyllobora","Psyllobora vigintimaculata","4","4","4","54","0","0","99.49","446.5","0","447.5","34.75","0"
"AGAW1","08W","INS","BOLD:AAE7186","Arthropoda","Insecta","Hymenoptera","Braconidae","Aphidiinae","Praon","Praon sp. 1 MM-2013, Praon volucre, Praon bicolor","4","4","4","54","0","0","99.77","438.25","0","439.25","25.54","0"
"AGAW1","08W","INS","BOLD:AAA3889","Arthropoda","Insecta","Lepidoptera","Crambidae","Crambinae","Parapediasia","Parapediasia teterrellus","4","4","4","54","0","0","99.83","449.5","0","450","32.5","0"
"AGAW1","08W","INS","BOLD:AAH0134","Arthropoda","Insecta","Coleoptera","Phalacridae","Phalacrinae","Stilbus","Stilbus apicialis","4","4","4","54","0","0","99.36","417.75","0","418.5","31.74","0"
"AGAW1","08W","INS","BOLD:ACF4925","Arthropoda","Insecta","Hemiptera","Miridae","Mirinae","Adelphocoris","Adelphocoris detritus","4","4","4","54","0","0","98.22","449.5","0","450","34.78","0"
"AGAW1","08W","INS","BOLD:AAG2460","Arthropoda","Insecta","Diptera","Anthomyiidae","Anthomyiinae","Hydrophoria","Hydrophoria lancifer","4","4","4","54","0","0","98.16","447.75","0","450","34.28","0"
"AGAW1","08W","INS","BOLD:AAG5507","Arthropoda","Insecta","Diptera","Chironomidae","Chironominae","Chironomus","Chironomus atrella","3","3","3","61","0","0","97.98","446.67","0","450","36.07","0"
"AGAW1","08W","INS","BOLD:AAA2632","Arthropoda","Insecta","Lepidoptera","Noctuidae","Noctuinae","Noctua","Noctua atlantica, Noctua pronuba","3","3","3","61","0","0","99.52","419.33","0","420","31.07","0"
"AGAW1","08W","INS","BOLD:AAD2041","Arthropoda","Insecta","Diptera","Dolichopodidae","Sciapodinae","Condylostylus","Condylostylus inermis, Condylostylus nigrofemoratus, Condylostylus connectans, Condylostylus caudatus, Condylostylus flavipes","3","3","3","61","0","0","98.86","431.67","0","433.67","35.71","0"
"AGAW1","08W","INS","BOLD:AAD0484","Arthropoda","Insecta","Diptera","Chironomidae","","","","3","3","3","61","0","0","98.66","448.33","0","450","34.96","0"
"AGAW1","08W","INS","BOLD:AAL2750","Arthropoda","Insecta","Diptera","Muscidae","Phaoniinae","Phaonia","Phaonia tuguriorum","3","3","3","61","0","0","99.24","401.33","0","403.33","30.99","0"
"AGAW1","08W","INS","BOLD:AAA3867","Arthropoda","Insecta","Lepidoptera","Noctuidae","Noctuinae","Apamea","Apamea amputatrix","3","3","3","61","0","0","99.63","447.67","0","450","33.7","0"
"AGAW1","08W","INS","BOLD:AAI4303","Arthropoda","Insecta","Diptera","Chironomidae","Chironominae","Chironomus","Chironomus melanescens","3","3","3","61","0","0","97.52","393.67","0","397","34.16","0"
"AGAW1","08W","INS","BOLD:AAA4555","Arthropoda","Insecta","Orthoptera","Acrididae","Melanoplinae","Melanoplus","Melanoplus stupefactus, Melanoplus angustipennis, Melanoplus ponderosus, Melanoplus bruneri, Melanoplus frigidus, Melanoplus scapularis, Melanoplus cinereus, Melanoplus madeleineae, Melanoplus sanguinipes, Melanoplus nigrescens group, Melanoplus tunicae, Melanoplus walshi, Melanoplus femurrubrum, Melanoplus dawsoni, Melanoplus sp., Melanoplus fasciatus, Melanoplus bowditchi, Melanoplus stonei, Melanoplus bivittatus, Melanoplus huroni, Melanoplus thomasi, Melanoplus borealis, Melanoplus packardii","3","3","3","61","0","0","98.89","449","0","450","36.74","0"
"AGAW1","08W","INS","BOLD:ABV1237","Arthropoda","Insecta","Diptera","Chironomidae","Chironominae","Chironomus","","3","3","3","61","0","0","99.78","449","0","450","34.81","0"
"AGAW1","08W","INS","BOLD:ACE0699","Arthropoda","Insecta","Diptera","Chironomidae","","","","2","2","2","70","0","0","98.84","430.5","0","450","36.67","0"
"AGAW1","08W","INS","BOLD:AAG7276","Arthropoda","Insecta","Diptera","Sphaeroceridae","Limosininae","Leptocera","Leptocera erythrocera","2","2","2","70","0","0","99","382","0","445.5","35.11","0"
"AGAW1","08W","INS","BOLD:ACL8669","Arthropoda","Insecta","Lepidoptera","Psychidae","Psychinae","Psyche","Psyche casta","2","2","2","70","0","0","98","382","0","385.5","36.85","0"
"AGAW1","08W","INS","BOLD:AAA8796","Arthropoda","Insecta","Lepidoptera","Tortricidae","Tortricinae","Acleris","Acleris forsskaleana","2","2","2","70","0","0","98.44","399","0","412","32.15","0"
"AGAW1","08W","INS","BOLD:AAN5342","Arthropoda","Insecta","Diptera","Chironomidae","Orthocladiinae","Heleniella","","2","2","2","70","0","0","99.42","426","0","426","35.05","0"
"AGAW1","08W","INS","BOLD:AAA9246","Arthropoda","Insecta","Lepidoptera","Tortricidae","Tortricinae","Clepsis","Clepsis clemensiana","2","2","2","70","0","0","98.21","398.5","0","399","29.47","0"
"AGAW1","08W","INS","BOLD:AAG2123","Arthropoda","Insecta","Diptera","Tachinidae","Dexiinae","Dinera","Dinera grisescens","2","2","2","70","0","0","99.78","449.5","0","450","32.22","0"
"AGAW1","08W","INS","BOLD:ABX6186","Arthropoda","Insecta","Diptera","Tipulidae","Tipulinae","Nephrotoma","Nephrotoma ferruginea","2","2","2","70","0","0","99.27","426.5","0","428.5","34.78","0"
"AGAW1","08W","INS","BOLD:AAU0577","Arthropoda","Insecta","Thysanoptera","Aeolothripidae","Aeolothripinae","Aeolothrips","Aeolothrips fasciatus","2","2","2","70","0","0","97.29","406","0","407.5","35.7","0"
"AGAW1","08W","INS","BOLD:AAE7016","Arthropoda","Insecta","Lepidoptera","Gelechiidae","Dichomeridinae","Helcystogramma","Helcystogramma hystricella","2","2","2","70","0","0","98.98","438.5","0","439.5","31.51","0"
"AGAW1","08W","INS","BOLD:ABA0863","Arthropoda","Insecta","Diptera","Cecidomyiidae","","","","1","1","1","80","0","0","99.56","450","0","450","30.22","0"
"AGAW1","08W","INS","BOLD:AAA2590","Arthropoda","Insecta","Lepidoptera","Noctuidae","Noctuinae","Xestia","Xestia baja, Xestia smithii","1","1","1","80","0","0","99.78","446","0","450","32.67","0"
"AGAW1","08W","INS","BOLD:ABA0856","Arthropoda","Insecta","Diptera","Cecidomyiidae","","","","1","1","1","80","0","0","100","450","0","450","29.33","0"
"AGAW1","08W","INS","BOLD:ABA6163","Arthropoda","Insecta","Psocodea","Caeciliusidae","Caeciliusinae","Valenzuela","Valenzuela piceus","1","1","1","80","0","0","100","446","0","450","36.89","0"
"AGAW1","08W","INS","BOLD:AAL4232","Arthropoda","Insecta","Diptera","Agromyzidae","Phytomyzinae","Phytoliriomyza","Phytoliriomyza arctica","1","1","1","80","0","0","99.78","450","0","450","32","0"
"AGAW1","08W","INS","BOLD:AAH4135","Arthropoda","Insecta","Diptera","Chloropidae","Chloropinae","Thaumatomyia","Thaumatomyia glabra, Thaumatomyia grata, Thaumatomyia sp. 1 TAW","1","1","1","80","0","0","97.09","447","0","450","33.11","0"
"AGAW1","08W","INS","BOLD:ACR4394","Arthropoda","Insecta","Diptera","Anthomyiidae","Anthomyiinae","Delia","Delia florilega","1","1","1","80","0","0","98.44","450","0","450","33.33","0"
"AGAW1","08W","INS","BOLD:AAM6062","Arthropoda","Insecta","Diptera","Cecidomyiidae","","","","1","1","1","80","0","0","100","447","0","450","26","0"
"AGAW1","08W","INS","BOLD:AAG5453","Arthropoda","Insecta","Diptera","Chironomidae","Chironominae","Chironomus","Chironomus bifurcatus","1","1","1","80","0","0","97.33","450","0","450","34.89","0"
"AGAW1","08W","INS","BOLD:AAH3920","Arthropoda","Insecta","Diptera","Sciaridae","","Dichopygina, Scatopsciara","","1","1","1","80","0","0","99.25","398","0","411","34.31","0"
"AGAW1","08W","INS","BOLD:AAV5594","Arthropoda","Insecta","Diptera","Cecidomyiidae","","","","1","1","1","80","0","0","99.72","356","0","357","25.49","0"
"AGAW1","08W","INS","BOLD:AAN5238","Arthropoda","Insecta","Diptera","Cecidomyiidae","","","","1","1","1","80","0","0","98.88","448","0","450","27.11","0"
"AGAW1","08W","INS","BOLD:AAF9771","Arthropoda","Insecta","Diptera","Hybotidae","Tachydromiinae","Platypalpus","Platypalpus longiseta, Platypalpus trivialis, Platypalpus sp. 5, Platypalpus sp. 17, Platypalpus pallidiventris","1","1","1","80","0","0","100","450","0","450","35.33","0"
"AGAW1","08W","INS","BOLD:AAH3914","Arthropoda","Insecta","Diptera","Sciaridae","","Sciara","Sciara hamatilis, Sciara humeralis","1","1","1","80","0","0","99.78","446","0","450","30.22","0"
"AGAW1","08W","INS","BOLD:AAG5443","Arthropoda","Insecta","Diptera","Chironomidae","Orthocladiinae","","","1","1","1","80","0","0","99.51","410","0","412","33.01","0"
"AGAW1","08W","INS","BOLD:AAG0996","Arthropoda","Insecta","Diptera","Chironomidae","Orthocladiinae","Cricotopus","Cricotopus sp. 18ES, Cricotopus bicinctus gr., Cricotopus bicinctus grp.","1","1","1","80","0","0","97.95","390","0","391","32.74","0"
"AGAW1","08W","INS","BOLD:AAL7359","Arthropoda","Insecta","Diptera","Chironomidae","Chironominae","Tanytarsus","","1","1","1","80","0","0","99.78","448","0","450","36.44","0"
"AGAW1","08W","INS","BOLD:AAN5205","Arthropoda","Insecta","Diptera","Cecidomyiidae","","","CecidInt42 sp. BOLD:AAN5205","1","1","1","80","0","0","98.88","448","0","450","31.11","0"
"AGAW1","08W","INS","BOLD:AAE0707","Arthropoda","Insecta","Diptera","Chironomidae","Chironominae","Microtendipes","Microtendipes pedellus","1","1","1","80","0","0","98.89","449","0","450","31.11","0"
"AGAW1","08W","INS","BOLD:ADH4973","Arthropoda","Insecta","Diptera","Tipulidae","Tipulinae","Nephrotoma","","1","1","1","80","0","0","98.44","448","0","450","33.78","0"
"AGAW1","08W","INS","BOLD:ACK7314","Arthropoda","Insecta","Diptera","Chironomidae","Chironominae","Polypedilum","","1","1","1","80","0","0","99.72","351","0","354","34.75","0"
"AGAW1","08W","INS","BOLD:AAD1485","Arthropoda","Insecta","Diptera","Chironomidae","Chironominae","Paratanytarsus","Paratanytarsus grimmii, Paratanytarsus sp.","1","1","1","80","0","0","100","450","0","450","34","0"
"AGAW1","08W","INS","BOLD:AAU6970","Arthropoda","Insecta","Coleoptera","Cleridae","Clerinae","Enoclerus","Enoclerus nigripes","1","1","1","80","0","0","97.47","396","0","398","33.42","0"
"AGAW1","08W","INS","BOLD:AAN8770","Arthropoda","Insecta","Diptera","Psychodidae","Psychodinae","Psychoda","Psychoda trinodulosa","1","1","1","80","0","0","97.78","450","0","450","34.22","0"
"AGAW1","08W","INS","BOLD:AAP5120","Arthropoda","Insecta","Diptera","Chironomidae","Chironominae","Polypedilum","Polypedilum halterale gr., Polypedilum scalaenum gr.","1","1","1","80","0","0","98.24","397","0","401","32.67","0"
"AGAW1","08W","INS","BOLD:AAF9187","Arthropoda","Insecta","Coleoptera","Curculionidae","Entiminae","Phyllobius","Phyllobius oblongus","1","1","1","80","0","0","100","450","0","450","38.89","0"
"AGAW1","08W","INS","BOLD:AAG2452","Arthropoda","Insecta","Diptera","Anthomyiidae","Anthomyiinae","","","1","1","1","80","0","0","97.56","450","0","450","32","0"
"AGAW1","08W","INS","BOLD:ACB9147","Arthropoda","Insecta","Psocodea","Psocidae","","","","1","1","1","80","0","0","97.1","448","0","450","37.78","0"
"AGAW1","08W","INS","BOLD:AAA4893","Arthropoda","Insecta","Lepidoptera","Crambidae","Crambinae","Microcrambus","Microcrambus elegans","1","1","1","80","0","0","99.29","421","0","450","32.67","0"
"AGAW1","08W","INS","BOLD:ABW2870","Arthropoda","Insecta","Coleoptera","Staphylinidae","Aleocharinae","Meronera","Meronera venustula","1","1","1","80","0","0","99.78","449","0","450","35.56","0"
"AGAW1","08W","INS","BOLD:ABV9293","Arthropoda","Insecta","Diptera","Chironomidae","Chironominae","Dicrotendipes","","1","1","1","80","0","0","98.03","355","0","386","33.68","0"
"AGAW1","08W","INS","BOLD:AAA3782","Arthropoda","Insecta","Hymenoptera","Halictidae","Halictinae","Lasioglossum","Lasioglossum weemsi, Lasioglossum leviense, Lasioglossum hitchensi","1","1","1","80","0","0","98.66","447","0","450","27.78","0"
"AGAW1","08W","INS","BOLD:ABY5257","Arthropoda","Insecta","Lepidoptera","Noctuidae","Noctuinae","Apamea","Apamea devastator","1","1","1","80","0","0","98.81","421","0","424","30.19","0"
"AGAW1","08W","INS","BOLD:ACP9679","Arthropoda","Insecta","Hymenoptera","Braconidae","Euphorinae","Microctonus","","1","1","1","80","0","0","98","449","0","450","31.33","0"
"AGAW1","08W","INS","BOLD:AAP9022","Arthropoda","Insecta","Diptera","Cecidomyiidae","","","","1","1","1","80","0","0","99.78","446","0","450","27.56","0"
"AGAW1","08W","INS","BOLD:AAG2898","Arthropoda","Insecta","Hemiptera","Cicadellidae","Deltocephalinae","Paraphlepsius","Paraphlepsius apertus, Paraphlepsius irroratus","1","1","1","80","0","0","100","449","0","450","32.22","0"
"AGAW1","08W","INS","BOLD:ADG1318","","","","","","","","1","1","1","80","0","0","100","448","0","450","25.11","0"
"AGAW1","08W","INS","BOLD:ABV1430","Arthropoda","Insecta","Diptera","Dolichopodidae","Medeterinae","Medetera","","1","1","1","80","0","0","100","342","0","352","37.5","0"
"AGAW1","08W","INS","BOLD:AAB2384","Arthropoda","Insecta","Diptera","Syrphidae","Syrphinae","Eupeodes","Eupeodes duseki, Eupeodes volucris, Eupeodes nielseni, Eupeodes americanus, Eupeodes flukei, Eupeodes vockerothi, Eupeodes lucasi, Eupeodes montivagus, Eupeodes rufipunctatus, Eupeodes rojasi, Eupeodes goeldlini, Eupeodes luniger, Eupeodes punctifer, Eupeodes bucculatus, Eupeodes perplexus, Eupeodes abiskoensis, Eupeodes corollae, Eupeodes pomus, Eupeodes lundbecki, Eupeodes nitens, Eupeodes sp. BOLDAAB2384, Eupeodes curtus, Eupeodes latifasciatus","1","1","1","80","0","0","99.75","397","0","399","30.33","0"
"AGAW1","08W","INS","BOLD:ADH9310","Arthropoda","Insecta","Diptera","Anthomyiidae","Anthomyiinae","Anthomyia","Anthomyia procellaris, Anthomyia pluvialis","1","1","1","80","0","0","98.89","450","0","450","32.67","0"
"AGAW1","08W","INS","BOLD:AAP8651","Arthropoda","Insecta","Diptera","Tachinidae","Exoristinae","Oswaldia","Oswaldia albifacies","1","1","1","80","0","0","99.18","366","0","366","32.79","0"
"AGAW1","08W","INS","BOLD:AAH4205","Arthropoda","Insecta","Diptera","Chloropidae","Chloropinae","Meromyza","Meromyza nigriseta, Meromyza sp.","1","1","1","80","0","0","99.78","450","0","450","33.11","0"
"AGAW1","08W","INS","BOLD:ACE7482","Arthropoda","Insecta","Hemiptera","Miridae","","","","1","1","1","80","0","0","98.51","404","0","405","34.32","0"
"AGAW1","08W","INS","BOLD:AAA7661","Arthropoda","Insecta","Diptera","Culicidae","Culicinae","Culiseta, Culex","Culiseta melanura, Culex articularis, Culex restuans, Culex acharistus","1","1","1","80","0","0","98.61","360","0","363","32.51","0"
"AGAW1","08W","INS","BOLD:ABW7897","Arthropoda","Insecta","Diptera","Cecidomyiidae","","","","1","1","1","80","0","0","99.55","449","0","450","29.33","0"
"AGAW1","08W","INS","BOLD:AAA5953","Arthropoda","Insecta","Lepidoptera","Gelechiidae","Gelechiinae","Coleotechnites","Coleotechnites sp., Coleotechnites piceaella, Coleotechnites laricis, Coleotechnites sp. 3, Coleotechnites carbonaria, Coleotechnites granti, Coleotechnites macleodi, Coleotechnites gibsonella, Coleotechnites coniferella, Coleotechnites martini","1","1","1","80","0","0","97.85","419","0","422","32.46","0"
"AGAW1","08W","INS","BOLD:ACE4701","Arthropoda","Insecta","Lepidoptera","Crambidae","Crambinae","Crambus","Crambus multilinellus","1","1","1","80","0","0","99.76","425","0","426","31.92","0"
"AGAW1","08W","INS","BOLD:AAA5803","Arthropoda","Insecta","Hemiptera","Miridae","Mirinae","Lygus","Lygus lineolaris","1","1","1","80","0","0","98.89","449","0","450","34.22","0"
"AGAW1","08W","INS","BOLD:ACK3589","Arthropoda","Insecta","Diptera","Cecidomyiidae","","","","1","1","1","80","0","0","99.33","449","0","450","32.89","0"
"AGAW1","08W","NAR","no match","no match","no match","no match","no match","no match","no match","no match","no match","no match","no match","no match","no match","no match","no match","no match","no match","no match","no match","no match"
"AGAW1","08W","NIN","BOLD:AAA7242","Arthropoda","Collembola","Entomobryomorpha","Entomobryidae","Seirinae, Entomobryinae","Entomobrya, Lepidosira","Entomobrya multifasciata, Lepidosira sp.","264","257","257","1","2.65","0","99.1","418.88","0","422.91","39.3","0"
"AGAW1","08W","NIN","BOLD:AAB7914","Arthropoda","Collembola","Symphypleona","Bourletiellidae","","Bourletiella","","45","36","36","2","20","0","99.76","446.8","0","448.02","38.15","0"
"AGAW1","08W","NIN","BOLD:AAM7961","Arthropoda","Arachnida","Trombidiformes","Anystidae","","","","28","25","25","3","10.71","0","99.3","438","0","434.46","31.04","0"
"AGAW1","08W","NIN","BOLD:AAH6630","Arthropoda","Collembola","Entomobryomorpha","Entomobryidae","Entomobryinae","Entomobrya","Entomobrya unostrigata","1","1","1","4","0","0","100","384","0","387","38.76","0"
"AGAW1","08W","NIN","BOLD:AAC9044","Arthropoda","Arachnida","Araneae","Salticidae","","Salticus","Salticus scenicus","1","1","1","4","0","0","99.3","427","0","430","33.26","0"
"AGAW1","08W","NIN","BOLD:ABW2639","Arthropoda","Arachnida","Trombidiformes","Anystidae","","","","1","1","1","4","0","0","100","448","0","450","31.56","0"
"AGAW1","08W","NIN","BOLD:AAA8820","Arthropoda","Collembola","Entomobryomorpha","Entomobryidae","Lepidocyrtinae","Lepidocyrtus","Lepidocyrtus cyaneus","1","1","1","4","0","0","99.11","449","0","450","43.56","0"
"AGAW1","08W","NIN","BOLD:AAI4346","Arthropoda","Arachnida","Opiliones","Phalangiidae","","Paroligolophus, Phalangium","Paroligolophus agrestis, Phalangium opilio","1","1","1","4","0","0","98.14","431","0","450","35.11","0"
"AGAW1","09E","BAC","TAX:175768","Proteobacteria","Alphaproteobacteria","Rickettsiales","Rickettsiaceae",NA,"Wolbachia","Wolbachia pipientis","111","87","87","1","21.62","0","98.06","448.13","0","449.59","39.45","0"
"AGAW1","09E","BAC","TAX:91393","Proteobacteria","Alphaproteobacteria","Rickettsiales","Rickettsiaceae",NA,"Wolbachia","","100","100","100","2","0","0","98.94","389.39","0","418.95","39.02","0"
"AGAW1","09E","BAC","TAX:91391","Proteobacteria","Alphaproteobacteria","Rickettsiales","",NA,"","","2","2","2","3","0","0","98.58","423","0","450","40","0"
"AGAW1","09E","BAC","TAX:91389","Proteobacteria","","","",NA,"","","1","1","1","4","0","0","98.33","359","0","364","37.91","0"
"AGAW1","09E","BAC","TAX:375763","Proteobacteria","Alphaproteobacteria","Rickettsiales","Rickettsiaceae",NA,"Wolbachia","Wolbachia sp. Pheidole nebulosa_ASLAM107-08","1","1","1","4","0","0","99.52","417","0","450","40.22","0"
"AGAW1","09E","CHO","no match","no match","no match","no match","no match","no match","no match","no match","no match","no match","no match","no match","no match","no match","no match","no match","no match","no match","no match","no match"
"AGAW1","09E","INS","BOLD:AAB0373","Arthropoda","Insecta","Neuroptera","Chrysopidae","Chrysopinae","Chrysoperla","Chrysoperla sp. isolate 82, Chrysoperla zastrowi, Chrysoperla plorabunda, Chrysoperla sp. isolate 71, Chrysoperla agilis, Italochrysa italica, Chrysoperla sp. isolate 88, Chrysoperla lucasina, Chrysoperla carnea, Chrysoperla mediterranea, Chrysoperla pallida, Chrysoperla nipponensis, Chrysoperla sillemi","19389","13402","13402","1","30.88","0","99.46","436.55","0","437.44","31.31","0"
"AGAW1","09E","INS","BOLD:AAP8739","Arthropoda","Insecta","Diptera","Platystomatidae","Platystomatinae","Rivellia","Rivellia steyskali","14240","8735","8735","2","38.66","0","99.33","443.4","0","444.86","38","0"
"AGAW1","09E","INS","BOLD:AAG1717","Arthropoda","Insecta","Diptera","Muscidae","Phaoniinae","Helina","Helina troene","12463","9511","9511","3","23.69","0","99.36","438.18","0","439.33","30.87","0"
"AGAW1","09E","INS","BOLD:AAA3453","Arthropoda","Insecta","Diptera","Anthomyiidae","Anthomyiinae","Delia","","6773","5816","5816","4","14.13","0","99.37","432.09","0","434.67","34.64","0"
"AGAW1","09E","INS","BOLD:AAH7142","Arthropoda","Insecta","Diptera","Sarcophagidae","Sarcophaginae","Oxysarcodexia, Sarcophaginae_incertae_sedis","Oxysarcodexia cingarus, Sarcophaginae_incertae_sedis sp. 1 AGT, Oxysarcodexia ventricosa","6299","3839","3839","5","39.05","0","99.03","445.99","0","447.02","34.09","0"
"AGAW1","09E","INS","BOLD:AAG2511","Arthropoda","Insecta","Diptera","Anthomyiidae","Anthomyiinae","Delia","Delia platura","5534","4635","4635","6","16.25","0","99.03","444.74","0","447.57","33.16","0"
"AGAW1","09E","INS","BOLD:AAC0903","Arthropoda","Insecta","Diptera","Chironomidae","Chironominae","Chironomus","Chironomus acidophilus","1631","1228","1228","7","24.71","0","99.5","438.44","0","440.86","36.31","0"
"AGAW1","09E","INS","BOLD:AAG9626","Arthropoda","Insecta","Diptera","Dolichopodidae","","","","1327","1033","1033","8","22.16","0","98.29","442.91","0","444.65","36.4","0"
"AGAW1","09E","INS","BOLD:AAA7661","Arthropoda","Insecta","Diptera","Culicidae","Culicinae","Culiseta, Culex","Culiseta melanura, Culex articularis, Culex restuans, Culex acharistus","1309","1191","1191","9","9.01","0","99.15","429.84","0","431.69","33.02","0"
"AGAW1","09E","INS","BOLD:ACL1796","Arthropoda","Insecta","Lepidoptera","Ypsolophidae","Ochsenheimeriinae","Ochsenheimeria","Ochsenheimeria vacculella","874","755","755","10","13.62","0","99.1","428.71","0","437.74","32.42","0"
"AGAW1","09E","INS","BOLD:AAD4936","Arthropoda","Insecta","Lepidoptera","Gracillariidae","Gracillariinae","Caloptilia","Caloptilia rhoifoliella, Caloptilia sassafrasella","862","832","832","11","3.48","0","98.92","425.14","0","427.34","29.45","0"
"AGAW1","09E","INS","BOLD:AAP8989","Arthropoda","Insecta","Diptera","Micropezidae","Calobatinae","Compsobata","Compsobata univitta, Compsobata sp.","830","609","609","12","26.63","0","98.74","440.24","0","441.1","36.58","0"
"AGAW1","09E","INS","BOLD:ABW7897","Arthropoda","Insecta","Diptera","Cecidomyiidae","","","","793","539","539","13","32.03","0","99.6","443.71","0","445.91","29.72","0"
"AGAW1","09E","INS","BOLD:AAF9414","Arthropoda","Insecta","Coleoptera","Meloidae","Meloinae","Epicauta","Epicauta murina","771","671","671","14","12.97","0","98.52","433.6","0","435.34","42","0"
"AGAW1","09E","INS","BOLD:AAU6681","Arthropoda","Insecta","Diptera","Tipulidae","Tipulinae","Nephrotoma","Nephrotoma cornicina","713","570","570","15","20.06","0","99.56","442.58","0","443.68","36.32","0"
"AGAW1","09E","INS","BOLD:ABW1379","Arthropoda","Insecta","Diptera","Chloropidae","Oscinellinae","Malloewia","Malloewia abdominalis","697","522","522","16","25.11","0","98.31","443.93","0","447.07","31.53","0"
"AGAW1","09E","INS","BOLD:AAD4214","Arthropoda","Insecta","Hymenoptera","Ichneumonidae","Diplazontinae","Diplazon","Diplazon laetatorius","651","492","492","17","24.42","0","99.52","446.72","0","447.75","25.8","0"
"AGAW1","09E","INS","BOLD:AAH3914","Arthropoda","Insecta","Diptera","Sciaridae","","Sciara","Sciara hamatilis, Sciara humeralis","649","471","471","18","27.43","0","99.17","442.54","0","443.5","30.57","0"
"AGAW1","09E","INS","BOLD:AAN6154","Arthropoda","Insecta","Coleoptera","Latridiidae","Corticariinae","Melanophthalma","Melanophthalma inermis","625","509","509","19","18.56","0","99.09","435.63","0","436.89","31.3","0"
"AGAW1","09E","INS","BOLD:AAN5342","Arthropoda","Insecta","Diptera","Chironomidae","Orthocladiinae","Heleniella","","588","552","552","20","6.12","0","99.05","423.06","0","427.14","35.11","0"
"AGAW1","09E","INS","BOLD:AAF9028","Arthropoda","Insecta","Diptera","Limoniidae","","","","511","412","412","21","19.37","0","99.35","436.82","0","438.96","38.61","0"
"AGAW1","09E","INS","BOLD:AAF9771","Arthropoda","Insecta","Diptera","Hybotidae","Tachydromiinae","Platypalpus","Platypalpus longiseta, Platypalpus trivialis, Platypalpus sp. 5, Platypalpus sp. 17, Platypalpus pallidiventris","473","362","362","22","23.47","0","99.47","444.72","0","445.39","35.12","0"
"AGAW1","09E","INS","BOLD:ACK7314","Arthropoda","Insecta","Diptera","Chironomidae","Chironominae","Polypedilum","","429","426","426","23","0.7","0","98.97","384.87","0","389.14","35.13","0"
"AGAW1","09E","INS","BOLD:AAD2041","Arthropoda","Insecta","Diptera","Dolichopodidae","Sciapodinae","Condylostylus","Condylostylus inermis, Condylostylus nigrofemoratus, Condylostylus connectans, Condylostylus caudatus, Condylostylus flavipes","385","288","288","24","25.19","0","99.53","441.97","0","443.11","35.94","0"
"AGAW1","09E","INS","BOLD:AAE0707","Arthropoda","Insecta","Diptera","Chironomidae","Chironominae","Microtendipes","Microtendipes pedellus","382","359","359","25","6.02","0","98.55","424.79","0","426.86","30.84","0"
"AGAW1","09E","INS","BOLD:AAG2898","Arthropoda","Insecta","Hemiptera","Cicadellidae","Deltocephalinae","Paraphlepsius","Paraphlepsius apertus, Paraphlepsius irroratus","359","281","281","26","21.73","0","99.63","442.3","0","443.06","31.88","0"
"AGAW1","09E","INS","BOLD:AAG5507","Arthropoda","Insecta","Diptera","Chironomidae","Chironominae","Chironomus","Chironomus atrella","358","356","356","27","0.56","0","98.98","424.22","0","430.99","35.89","0"
"AGAW1","09E","INS","BOLD:ACE7444","Arthropoda","Insecta","Hemiptera","Miridae","Mirinae, Phylinae","Adelphocoris, Megalocoleus","Adelphocoris lineolatus, Megalocoleus molliculus","349","233","233","28","33.24","0","99.67","442.81","0","443.4","34.83","0"
"AGAW1","09E","INS","BOLD:AAG4386","Arthropoda","Insecta","Hymenoptera","Pompilidae","","","","341","302","302","29","11.44","0","99.52","403","0","431.77","26.07","0"
"AGAW1","09E","INS","BOLD:ABU5525","Arthropoda","Insecta","Diptera","Chironomidae","Orthocladiinae","Limnophyes","Limnophyes sp. 14ES","316","300","300","30","5.06","0","99.02","402.57","0","405.89","37.17","0"
"AGAW1","09E","INS","BOLD:AAG5453","Arthropoda","Insecta","Diptera","Chironomidae","Chironominae","Chironomus","Chironomus bifurcatus","278","237","237","31","14.75","0","97.84","438.06","0","446.29","34.72","0"
"AGAW1","09E","INS","BOLD:ACK3589","Arthropoda","Insecta","Diptera","Cecidomyiidae","","","","277","226","226","32","18.41","0","99.34","447.81","0","448.68","32.25","0"
"AGAW1","09E","INS","BOLD:ACA9275","Arthropoda","Insecta","Diptera","Chironomidae","Chironominae","Chironomus","","274","233","233","33","14.96","0","99.46","433.17","0","434.85","35.48","0"
"AGAW1","09E","INS","BOLD:ACE0699","Arthropoda","Insecta","Diptera","Chironomidae","","","","244","182","182","34","25.41","0","98.8","423.6","0","439.46","36.31","0"
"AGAW1","09E","INS","BOLD:ACX5521","Arthropoda","Insecta","Diptera","Chironomidae","Orthocladiinae","","","234","230","230","35","1.71","0","99.04","417.94","0","419.92","34.85","0"
"AGAW1","09E","INS","BOLD:AAB7030","Arthropoda","Insecta","Diptera","Chironomidae","Chironominae","Chironomus","Chironomus sp. BOLDAAP2996, Chironomus quinnitukqut, Chironomus cf. decorus 2g, Chironomus decorus group sp. 2 IP2013, Chironomus cf. decorus, Chironomus decorus gr.","184","184","184","36","0","0","98.66","407.88","0","413.87","33.87","0"
"AGAW1","09E","INS","BOLD:AAH4135","Arthropoda","Insecta","Diptera","Chloropidae","Chloropinae","Thaumatomyia","Thaumatomyia glabra, Thaumatomyia grata, Thaumatomyia sp. 1 TAW","183","167","167","37","8.74","0","99.47","429.93","0","434.33","31.9","0"
"AGAW1","09E","INS","BOLD:ABX5095","Arthropoda","Insecta","Hemiptera","Miridae","Mirinae","Neolygus","Neolygus tiliae","181","137","137","38","24.31","0","99.58","441.64","0","442.29","33.34","0"
"AGAW1","09E","INS","BOLD:ACK3431","Arthropoda","Insecta","Diptera","Cecidomyiidae","","","","172","150","150","39","12.79","0","99.22","428.51","0","430.12","32.56","0"
"AGAW1","09E","INS","BOLD:AAN5199","Arthropoda","Insecta","Diptera","Cecidomyiidae","","","CecidInt20 sp. BOLD:AAN5199","167","142","142","40","14.97","0","99.53","433.15","0","444.86","26.7","0"
"AGAW1","09E","INS","BOLD:AAD1395","Arthropoda","Insecta","Diptera","Chironomidae","Chironominae","Polypedilum","Polypedilum convictum, Polypedilum flavum","161","122","122","41","24.22","0","98.76","443.84","0","445.66","33.56","0"
"AGAW1","09E","INS","BOLD:AAN5332","Arthropoda","Insecta","Diptera","Chironomidae","Chironominae","Polypedilum","","159","158","158","42","0.63","0","97.71","409.4","0","412.84","33.21","0"
"AGAW1","09E","INS","BOLD:AAH3661","Arthropoda","Insecta","Diptera","Cecidomyiidae","","","","159","152","152","42","4.4","0","99.32","432.4","0","433.76","25.42","0"
"AGAW1","09E","INS","BOLD:AAP5019","Arthropoda","Insecta","Diptera","Dolichopodidae","","","","158","122","122","44","22.78","0","99.22","447.77","0","448.78","32.4","0"
"AGAW1","09E","INS","BOLD:ACF2821","Arthropoda","Insecta","Diptera","Mycetophilidae","Mycetophilinae","Mycetophila","Mycetophila fungorum","139","120","120","45","13.67","0","99.18","426.28","0","429.32","35.12","0"
"AGAW1","09E","INS","BOLD:ACE9668","Arthropoda","Insecta","Hymenoptera","Ichneumonidae","Ophioninae","Enicospilus","Enicospilus purgatusDHJ02","138","117","117","46","15.22","0","99.33","444.89","0","445.43","25.99","0"
"AGAW1","09E","INS","BOLD:AAA4277","Arthropoda","Insecta","Diptera","Syrphidae","Syrphinae","Toxomerus","Toxomerus marginatus, Toxomerus sp. CR-3, Toxomerus CR-3","133","90","90","47","32.33","0","99.49","446.95","0","447.84","30.67","0"
"AGAW1","09E","INS","BOLD:AAG6460","Arthropoda","Insecta","Diptera","Cecidomyiidae","","","","131","129","129","48","1.53","0","98.39","436.34","0","438.15","34.3","0"
"AGAW1","09E","INS","BOLD:AAN5161","Arthropoda","Insecta","Diptera","Ceratopogonidae","Dasyheleinae","Dasyhelea","","125","121","121","49","3.2","0","99.09","401.97","0","404.16","37.47","0"
"AGAW1","09E","INS","BOLD:AAF1051","Arthropoda","Insecta","Diptera","Agromyzidae","Phytomyzinae","Cerodontha","Cerodontha muscina, Cerodontha calamagrostis","117","117","117","50","0","0","98.28","405.04","0","408.34","36.02","0"
"AGAW1","09E","INS","BOLD:AAN5267","Arthropoda","Insecta","Diptera","Cecidomyiidae","","","","115","91","91","51","20.87","0","99.39","447.52","0","448.1","30.58","0"
"AGAW1","09E","INS","BOLD:AAP2870","Arthropoda","Insecta","Diptera","Dolichopodidae","","","","111","93","93","52","16.22","0","99.19","438.49","0","439.73","38.36","0"
"AGAW1","09E","INS","BOLD:ACE0554","Arthropoda","Insecta","Diptera","Phoridae","Metopininae","Megaselia","","108","94","94","53","12.96","0","99.48","438.95","0","441.02","31.98","0"
"AGAW1","09E","INS","BOLD:AAN5238","Arthropoda","Insecta","Diptera","Cecidomyiidae","","","","98","85","85","54","13.27","0","99.39","428.41","0","429.43","25.59","0"
"AGAW1","09E","INS","BOLD:AAH3229","Arthropoda","Insecta","Psocodea","Dasydemellidae","Dasydemellinae","Teliapsocus","","98","93","93","54","5.1","0","99.34","418.96","0","420.32","35.13","0"
"AGAW1","09E","INS","BOLD:AAN8446","Arthropoda","Insecta","Psocodea","Peripsocidae","","Peripsocus","Peripsocus madidus","92","89","89","56","3.26","0","99.17","418.61","0","419.72","41.39","0"
"AGAW1","09E","INS","BOLD:AAN5183","Arthropoda","Insecta","Diptera","Cecidomyiidae","","","","87","75","75","57","13.79","0","98.29","445.26","0","446.51","27.8","0"
"AGAW1","09E","INS","BOLD:AAH2753","Arthropoda","Insecta","Coleoptera","Anthicidae","Anthicinae","Anthicus","Anthicus cervinus","86","81","81","58","5.81","0","98.59","414.59","0","426.42","33.39","0"
"AGAW1","09E","INS","BOLD:AAP5169","Arthropoda","Insecta","Diptera","Chloropidae","Oscinellinae","Elachiptera","Elachiptera nigriceps","85","67","67","59","21.18","0","99.47","433.58","0","434.53","31.5","0"
"AGAW1","09E","INS","BOLD:AAA1259","Arthropoda","Insecta","Diptera","Cecidomyiidae","","","","81","72","72","60","11.11","0","97.85","432.53","0","440.41","31.14","0"
"AGAW1","09E","INS","BOLD:AAM9347","Arthropoda","Insecta","Diptera","Phoridae","Metopininae","Megaselia","","80","80","80","61","0","0","97.82","394.98","0","397.64","33.37","0"
"AGAW1","09E","INS","BOLD:ABV1297","Arthropoda","Insecta","Diptera","Sciaridae","","","","79","77","77","62","2.53","0","99.03","435.33","0","437.8","30.3","0"
"AGAW1","09E","INS","BOLD:AAU7040","Arthropoda","Insecta","Coleoptera","Corylophidae","Corylophinae","Orthoperus","Orthoperus scutellaris","73","73","73","63","0","0","98.75","428.1","0","430.48","32.75","0"
"AGAW1","09E","INS","BOLD:AAG7891","Arthropoda","Insecta","Hymenoptera","Platygastridae","Scelioninae","Telenomus","Telenomus podisi","67","66","66","64","1.49","0","98.97","427.76","0","435.7","27.21","0"
"AGAW1","09E","INS","BOLD:AAG5443","Arthropoda","Insecta","Diptera","Chironomidae","Orthocladiinae","","","62","62","62","65","0","0","98.94","408.44","0","410.71","33.19","0"
"AGAW1","09E","INS","BOLD:AAM6248","Arthropoda","Insecta","Diptera","Chironomidae","Chironominae","Microtendipes","Microtendipes pedellus grp.","53","49","49","66","7.55","0","98.06","437.74","0","438.98","32.28","0"
"AGAW1","09E","INS","BOLD:AAM9255","Arthropoda","Insecta","Diptera","Sciaridae","","Pseudolycoriella","","50","49","49","67","2","0","99.03","413.68","0","416.02","30.94","0"
"AGAW1","09E","INS","BOLD:AAN6447","Arthropoda","Insecta","Diptera","Sciaridae","","Corynoptera","Corynoptera perpusilla","46","37","37","68","19.57","0","99.54","447.83","0","449.15","36.32","0"
"AGAW1","09E","INS","BOLD:ABV9018","Arthropoda","Insecta","Diptera","Mycetophilidae","Sciophilinae","Sciophila","","45","45","45","69","0","0","98.95","427.98","0","429.93","31.9","0"
"AGAW1","09E","INS","BOLD:AAH0135","Arthropoda","Insecta","Coleoptera","Phalacridae","Phalacrinae","Acylomus","Acylomus pugetanus, Acylomus ergoti","44","44","44","70","0","0","99.27","426.77","0","428.7","31.4","0"
"AGAW1","09E","INS","BOLD:AAL7819","Arthropoda","Insecta","Diptera","Psychodidae","Psychodinae","Psychoda","Psychoda sp., Psychoda sp. BOLD:AAL7819","43","43","43","71","0","0","99.24","413.65","0","415.67","32.61","0"
"AGAW1","09E","INS","BOLD:ACI3976","Arthropoda","Insecta","Diptera","Chironomidae","Orthocladiinae","","","43","43","43","71","0","0","99.16","385.53","0","400.05","37.48","0"
"AGAW1","09E","INS","BOLD:AAP6465","Arthropoda","Insecta","Diptera","Sciaridae","","Corynoptera","Corynoptera cursor","41","30","30","73","26.83","0","98.73","446.63","0","447.46","32.37","0"
"AGAW1","09E","INS","BOLD:AAH3300","Arthropoda","Insecta","Coleoptera","Coccinellidae","Coccinellinae","Cycloneda","Cycloneda munda","40","40","40","74","0","0","99.02","420.03","0","422.18","35.49","0"
"AGAW1","09E","INS","BOLD:ABA0863","Arthropoda","Insecta","Diptera","Cecidomyiidae","","","","37","36","36","75","2.7","0","99.31","439.43","0","440.7","29.76","0"
"AGAW1","09E","INS","BOLD:ACX8294","Arthropoda","Insecta","Diptera","Cecidomyiidae","","","","36","33","33","76","8.33","0","99.37","441.03","0","441.75","32.45","0"
"AGAW1","09E","INS","BOLD:ABY5267","Arthropoda","Insecta","Hymenoptera","Ichneumonidae","Campopleginae","Sinophorus","","36","35","35","76","2.78","0","98.4","423.5","0","425","25.95","0"
"AGAW1","09E","INS","BOLD:AAG6756","Arthropoda","Insecta","Diptera","Sarcophagidae","Sarcophaginae","Boettcheria","Boettcheria cimbicis","34","31","31","78","8.82","0","99.14","441.5","0","443.03","34.86","0"
"AGAW1","09E","INS","BOLD:AAI1549","Arthropoda","Insecta","Hymenoptera","Braconidae","Euphorinae","Meteorus","","33","30","30","79","9.09","0","98.89","440.88","0","441.79","31.06","0"
"AGAW1","09E","INS","BOLD:AAL4232","Arthropoda","Insecta","Diptera","Agromyzidae","Phytomyzinae","Phytoliriomyza","Phytoliriomyza arctica","32","30","30","80","6.25","0","99.37","440.56","0","441.88","31.93","0"
"AGAW1","09E","INS","BOLD:AAH3664","Arthropoda","Insecta","Diptera","Cecidomyiidae","","","CecidInt3 sp. BOLD:AAH3664","31","31","31","81","0","0","99.07","433.77","0","437","27.17","0"
"AGAW1","09E","INS","BOLD:AAG9897","Arthropoda","Insecta","Dermaptera","Forficulidae","Forficulinae","Forficula","Forficula auricularia-A","31","31","31","81","0","0","98.94","391.74","0","401.74","45.7","0"
"AGAW1","09E","INS","BOLD:AAJ4275","Arthropoda","Insecta","Diptera","Chironomidae","Chironominae","Chironomus","Chironomus harpi","29","26","26","83","10.34","0","98.39","429.17","0","444.41","38.43","0"
"AGAW1","09E","INS","BOLD:AAG1711","Arthropoda","Insecta","Diptera","Muscidae","Phaoniinae","Helina","Helina depuncta","28","28","28","84","0","0","98.68","435.79","0","438","31.16","0"
"AGAW1","09E","INS","BOLD:ABV9551","Arthropoda","Insecta","Hymenoptera","Pompilidae","Pompilinae","Episyron","Episyron biguttatus","28","28","28","84","0","0","98.18","418.21","0","422.75","31.72","0"
"AGAW1","09E","INS","BOLD:AAG2868","Arthropoda","Insecta","Hemiptera","Cicadellidae","Typhlocybinae","Empoasca","Empoasca sp., Empoasca fabae","24","19","19","86","20.83","0","99.71","449.29","0","450","32.35","0"
"AGAW1","09E","INS","BOLD:AAN5205","Arthropoda","Insecta","Diptera","Cecidomyiidae","","","CecidInt42 sp. BOLD:AAN5205","22","20","20","87","9.09","0","99.44","443.55","0","445.23","31.39","0"
"AGAW1","09E","INS","BOLD:AAG2740","Arthropoda","Insecta","Diptera","Ephydridae","Parydrinae","Philygria","Philygria debilis","20","20","20","88","0","0","98.51","388.05","0","391.6","32.37","0"
"AGAW1","09E","INS","BOLD:ABA1215","Arthropoda","Insecta","Diptera","Sciaridae","","Lycoriella","Lycoriella sativae","17","15","15","89","11.76","0","99.63","436.59","0","438","35.67","0"
"AGAW1","09E","INS","BOLD:ABV1455","Arthropoda","Insecta","Diptera","Chironomidae","Chironominae","","","17","17","17","89","0","0","97.83","441.47","0","443.94","31.62","0"
"AGAW1","09E","INS","BOLD:AAN5283","Arthropoda","Insecta","Diptera","Cecidomyiidae","","","","17","17","17","89","0","0","99.46","412.88","0","414.29","30.13","0"
"AGAW1","09E","INS","BOLD:ACM7977","Arthropoda","Insecta","Diptera","Cecidomyiidae","","","","17","16","16","89","5.88","0","99.61","448.71","0","450","24.84","0"
"AGAW1","09E","INS","BOLD:ACL6043","Arthropoda","Insecta","Diptera","Chironomidae","Chironominae","","","17","17","17","89","0","0","99.1","416","0","417.76","36.91","0"
"AGAW1","09E","INS","BOLD:ACU7906","Arthropoda","Insecta","Diptera","Chironomidae","Orthocladiinae","","","16","16","16","94","0","0","98.51","407.31","0","409.19","35.79","0"
"AGAW1","09E","INS","BOLD:ACA6930","Arthropoda","Insecta","Diptera","Cecidomyiidae","","","","16","16","16","94","0","0","98.52","430.88","0","450","28.03","0"
"AGAW1","09E","INS","BOLD:AAN5289","Arthropoda","Insecta","Diptera","Cecidomyiidae","","","","16","16","16","94","0","0","99.37","445.44","0","447.44","27.92","0"
"AGAW1","09E","INS","BOLD:AAH3947","Arthropoda","Insecta","Diptera","Sciaridae","","Bradysia","Bradysia trivittata","15","15","15","97","0","0","99.02","418.4","0","419.67","34.32","0"
"AGAW1","09E","INS","BOLD:AAH0134","Arthropoda","Insecta","Coleoptera","Phalacridae","Phalacrinae","Stilbus","Stilbus apicialis","14","13","13","98","7.14","0","99.08","433.71","0","434.71","32.1","0"
"AGAW1","09E","INS","BOLD:ABV1201","Arthropoda","Insecta","Diptera","Sciaridae","","Scatopsciara","Scatopsciara calamophila","14","13","13","98","7.14","0","99.24","445.14","0","445.43","36","0"
"AGAW1","09E","INS","BOLD:ACW5117","Arthropoda","Insecta","Diptera","Chironomidae","Orthocladiinae","Smittia","Smittia leucopogon","13","12","12","100","7.69","0","99.55","440","0","441.15","31.34","0"
"AGAW1","09E","INS","BOLD:AAF9768","Arthropoda","Insecta","Diptera","Hybotidae","Tachydromiinae","Platypalpus","Platypalpus holosericus, Platypalpus candicans, Platypalpus sp. BOLD:AAF9768, Platypalpus sp. 22","12","11","11","101","8.33","0","99.43","449.58","0","450","31.85","0"
"AGAW1","09E","INS","BOLD:ABY1179","Arthropoda","Insecta","Hymenoptera","Sierolomorphidae","","Sierolomorpha","","12","11","11","101","8.33","0","99.74","446.58","0","447.25","28.19","0"
"AGAW1","09E","INS","BOLD:AAM6037","Arthropoda","Insecta","Diptera","Cecidomyiidae","Cecidomyiinae","Contarinia, Dasineura","","11","11","11","103","0","0","98.25","439.36","0","441.82","26.41","0"
"AGAW1","09E","INS","BOLD:AAF1729","Arthropoda","Insecta","Psocodea","Lachesillidae","Lachesillinae","Lachesilla","Lachesilla pedicularia","11","11","11","103","0","0","99.28","444.73","0","446.09","37.27","0"
"AGAW1","09E","INS","BOLD:ABV1366","Arthropoda","Insecta","Diptera","Cecidomyiidae","Cecidomyiinae","Asteromyia","Asteromyia carbonifera","10","10","10","105","0","0","98.97","431.6","0","433.8","26.88","0"
"AGAW1","09E","INS","BOLD:ACZ1491","Arthropoda","Insecta","Diptera","Chironomidae","","","","10","10","10","105","0","0","97.66","448.8","0","450","34.53","0"
"AGAW1","09E","INS","BOLD:AAZ5506","Arthropoda","Insecta","Diptera","Dolichopodidae","Dolichopodinae","Dolichopus","Dolichopus sp. SEB39, Dolichopus gratus","10","10","10","105","0","0","98.83","401.6","0","401.4","41.06","0"
"AGAW1","09E","INS","BOLD:AAV5573","Arthropoda","Insecta","Diptera","Cecidomyiidae","","","","10","10","10","105","0","0","98.82","437.5","0","438.3","27.41","0"
"AGAW1","09E","INS","BOLD:ACX2014","Arthropoda","Insecta","Diptera","Chironomidae","Chironominae","Tanytarsus","Tanytarsus mendax","10","10","10","105","0","0","97.99","389.5","0","417.6","36.55","0"
"AGAW1","09E","INS","BOLD:AAD1485","Arthropoda","Insecta","Diptera","Chironomidae","Chironominae","Paratanytarsus","Paratanytarsus grimmii, Paratanytarsus sp.","9","9","9","110","0","0","99.38","449.22","0","450","34","0"
"AGAW1","09E","INS","BOLD:AAM7855","Arthropoda","Insecta","Thysanoptera","Thripidae","Thripinae","Frankliniella","Frankliniella tritici","9","8","8","110","11.11","0","99.6","448.22","0","450","31.33","0"
"AGAW1","09E","INS","BOLD:ADH9867","Arthropoda","Insecta","Diptera","","","","","8","8","8","112","0","0","98.71","388.63","0","391.63","37.05","0"
"AGAW1","09E","INS","BOLD:ABU5546","Arthropoda","Insecta","Diptera","Psychodidae","Psychodinae","Psychoda","","7","7","7","113","0","0","98.11","422","0","424.43","31.53","0"
"AGAW1","09E","INS","BOLD:AAE0766","Arthropoda","Insecta","Hemiptera","Miridae","Phylinae","Atractotomus","Atractotomus magnicornis","7","7","7","113","0","0","99.8","433.14","0","441","37.72","0"
"AGAW1","09E","INS","BOLD:AAY6412","Arthropoda","Insecta","Diptera","Sciaridae","","Bradysia","Bradysia urticae","7","7","7","113","0","0","99.16","440.29","0","443","31.91","0"
"AGAW1","09E","INS","BOLD:AAV5400","Arthropoda","Insecta","Diptera","Cecidomyiidae","","","","7","7","7","113","0","0","98.19","435.71","0","436.29","29.39","0"
"AGAW1","09E","INS","BOLD:ACV5892","Arthropoda","Insecta","Diptera","Cecidomyiidae","","","","7","7","7","113","0","0","98.66","428.14","0","430.29","24.54","0"
"AGAW1","09E","INS","BOLD:AAM6062","Arthropoda","Insecta","Diptera","Cecidomyiidae","","","","6","6","6","118","0","0","98.95","440.67","0","442.5","26.01","0"
"AGAW1","09E","INS","BOLD:AAN7616","Arthropoda","Insecta","Hymenoptera","Platygastridae","","","","6","6","6","118","0","0","97.23","439.67","0","450","25.59","0"
"AGAW1","09E","INS","BOLD:ACX6289","Arthropoda","Insecta","Diptera","Phoridae","Metopininae","Megaselia","","6","6","6","118","0","0","98.91","402.5","0","404.67","37.86","0"
"AGAW1","09E","INS","BOLD:AAZ0271","Arthropoda","Insecta","Diptera","Ceratopogonidae","Ceratopogoninae","Culicoides","","6","6","6","118","0","0","98.75","426.5","0","430","41.29","0"
"AGAW1","09E","INS","BOLD:ACF7633","Arthropoda","Insecta","Diptera","Cecidomyiidae","","","","5","5","5","122","0","0","99.51","449.2","0","450","30.67","0"
"AGAW1","09E","INS","BOLD:ABA0858","Arthropoda","Insecta","Diptera","Cecidomyiidae","","","","5","5","5","122","0","0","99.51","429.4","0","430","26.26","0"
"AGAW1","09E","INS","BOLD:AAO8538","Arthropoda","Insecta","Diptera","Psychodidae","Psychodinae","Psychoda","","5","5","5","122","0","0","99.31","417.6","0","420.8","31.9","0"
"AGAW1","09E","INS","BOLD:AAB4894","Arthropoda","Insecta","Hemiptera","Aphididae","Aphidinae","Sitobion","Sitobion smilacifoliae, Sitobion avenae, Sitobion phyllanthi","5","5","5","122","0","0","97.87","432.8","0","433.8","25.08","0"
"AGAW1","09E","INS","BOLD:ACX6055","Arthropoda","Insecta","Diptera","Phoridae","Metopininae","Megaselia","Megaselia sp. AK-2010-2, Megaselia sp. BOLD:ACX6055","5","5","5","122","0","0","98.75","448","0","450","33.29","0"
"AGAW1","09E","INS","BOLD:ACF0915","Arthropoda","Insecta","Diptera","Chironomidae","Orthocladiinae","","","4","4","4","127","0","0","98.66","406","0","407.5","33.72","0"
"AGAW1","09E","INS","BOLD:AAN5242","Arthropoda","Insecta","Diptera","Cecidomyiidae","","","","4","4","4","127","0","0","99.38","401","0","400.5","28.62","0"
"AGAW1","09E","INS","BOLD:AAN5255","Arthropoda","Insecta","Diptera","Cecidomyiidae","Cecidomyiinae","Dasineura","","4","4","4","127","0","0","97.36","436.75","0","450","26.89","0"
"AGAW1","09E","INS","BOLD:AAA5299","Arthropoda","Insecta","Diptera","Chironomidae","Orthocladiinae","Cricotopus","Cricotopus laetus, Cricotopus relucens, Cricotopus trifasciatus, Cricotopus sp., Cricotopus sp. 3ES, Cricotopus sylvestris, Cricotopus glacialis","4","4","4","127","0","0","99.01","425.75","0","427.25","34.5","0"
"AGAW1","09E","INS","BOLD:AAG6745","Arthropoda","Insecta","Diptera","Calliphoridae","Calliphorinae","Pollenia","Pollenia pediculata","4","4","4","127","0","0","99.15","390.25","0","392.25","33.47","0"
"AGAW1","09E","INS","BOLD:AAQ0558","Arthropoda","Insecta","Thysanoptera","Thripidae","Thripinae","Anaphothrips","Anaphothrips obscurus","3","3","3","132","0","0","98.91","432.67","0","432.33","31.34","0"
"AGAW1","09E","INS","BOLD:ACJ6954","Arthropoda","Insecta","Diptera","Cecidomyiidae","","","","3","3","3","132","0","0","99.8","337.67","0","431.33","26.87","0"
"AGAW1","09E","INS","BOLD:AAY6409","Arthropoda","Insecta","Diptera","Cecidomyiidae","","","","3","3","3","132","0","0","97.7","449.67","0","450","25.7","0"
"AGAW1","09E","INS","BOLD:AAH2334","Arthropoda","Insecta","Diptera","Asilidae","Asilinae","Neoitamus","Neoitamus flavofemoratus","3","3","3","132","0","0","98.03","429","0","430","39.51","0"
"AGAW1","09E","INS","BOLD:ACJ6984","Arthropoda","Insecta","Hymenoptera","Platygastridae","Scelioninae","Trissolcus","Trissolcus thyantae","3","3","3","132","0","0","98.99","415","0","419.33","26.61","0"
"AGAW1","09E","INS","BOLD:AAA2815","Arthropoda","Insecta","Lepidoptera","Noctuidae","Noctuinae","Mythimna","Mythimna yukonensis, Mythimna oxygala","3","3","3","132","0","0","98.51","448.67","0","450","33.63","0"
"AGAW1","09E","INS","BOLD:AAU6970","Arthropoda","Insecta","Coleoptera","Cleridae","Clerinae","Enoclerus","Enoclerus nigripes","3","3","3","132","0","0","97.25","449","0","450","33.19","0"
"AGAW1","09E","INS","BOLD:AAN8770","Arthropoda","Insecta","Diptera","Psychodidae","Psychodinae","Psychoda","Psychoda trinodulosa","3","3","3","132","0","0","98.7","423.33","0","424.33","33.21","0"
"AGAW1","09E","INS","BOLD:AAH3662","Arthropoda","Insecta","Diptera","Cecidomyiidae","","","CecidInt3 sp. BOLD:AAH3662","3","3","3","132","0","0","99.12","432.33","0","433.33","26.05","0"
"AGAW1","09E","INS","BOLD:AAI4303","Arthropoda","Insecta","Diptera","Chironomidae","Chironominae","Chironomus","Chironomus melanescens","3","3","3","132","0","0","97.68","431.33","0","433.67","34.11","0"
"AGAW1","09E","INS","BOLD:AAI5147","Arthropoda","Insecta","Hymenoptera","Ichneumonidae","Ophioninae","Enicospilus","Enicospilus sp. MS2189","2","2","2","142","0","0","98.89","449.5","0","450","26.78","0"
"AGAW1","09E","INS","BOLD:AAN5220","Arthropoda","Insecta","Diptera","Cecidomyiidae","","","","2","2","2","142","0","0","97.38","439.5","0","445.5","26.49","0"
"AGAW1","09E","INS","BOLD:AAF9314","Arthropoda","Insecta","Diptera","Psychodidae","Psychodinae","Psychoda","Psychoda sp., Psychoda minuta","2","2","2","142","0","0","98.44","450","0","450","34","0"
"AGAW1","09E","INS","BOLD:AAF9187","Arthropoda","Insecta","Coleoptera","Curculionidae","Entiminae","Phyllobius","Phyllobius oblongus","2","2","2","142","0","0","99.89","446.5","0","450","38.67","0"
"AGAW1","09E","INS","BOLD:AAN5178","Arthropoda","Insecta","Diptera","Cecidomyiidae","Cecidomyiinae","Rhopalomyia","Rhopalomyia sp. BOLD:AAN5178","2","2","2","142","0","0","98.47","422.5","0","431.5","26.73","0"
"AGAW1","09E","INS","BOLD:ACJ1142","Arthropoda","Insecta","Hymenoptera","Ichneumonidae","Campopleginae","Sinophorus","","2","2","2","142","0","0","97.44","449.5","0","450","25.67","0"
"AGAW1","09E","INS","BOLD:AAA5953","Arthropoda","Insecta","Lepidoptera","Gelechiidae","Gelechiinae","Coleotechnites","Coleotechnites sp., Coleotechnites piceaella, Coleotechnites laricis, Coleotechnites sp. 3, Coleotechnites carbonaria, Coleotechnites granti, Coleotechnites macleodi, Coleotechnites gibsonella, Coleotechnites coniferella, Coleotechnites martini","2","2","2","142","0","0","99.55","445.5","0","447","34.23","0"
"AGAW1","09E","INS","BOLD:AAA6292","Arthropoda","Insecta","Lepidoptera","Tortricidae","Tortricinae","Cnephasia","Cnephasia genitalana, Cnephasia stephensiana","1","1","1","149","0","0","98.44","449","0","450","35.56","0"
"AGAW1","09E","INS","BOLD:AAN8030","Arthropoda","Insecta","Hymenoptera","Platygastridae","Scelioninae","Telenomus","","1","1","1","149","0","0","99.03","414","0","450","26","0"
"AGAW1","09E","INS","BOLD:AAA9246","Arthropoda","Insecta","Lepidoptera","Tortricidae","Tortricinae","Clepsis","Clepsis clemensiana","1","1","1","149","0","0","100","450","0","450","30.44","0"
"AGAW1","09E","INS","BOLD:AAA6560","Arthropoda","Insecta","Lepidoptera","Crambidae","Crambinae","Crambus","Crambus multilinellus, Crambus saltuellus","1","1","1","149","0","0","99.46","367","0","370","30.81","0"
"AGAW1","09E","INS","BOLD:AAY6730","Arthropoda","Insecta","Hemiptera","Derbidae","Cedusinae","Cedusa","Cedusa incisa","1","1","1","149","0","0","99.55","449","0","450","35.56","0"
"AGAW1","09E","INS","BOLD:ABV0271","Arthropoda","Insecta","Diptera","Chloropidae","Oscinellinae","","","1","1","1","149","0","0","99.56","450","0","450","33.11","0"
"AGAW1","09E","INS","BOLD:AAB7869","Arthropoda","Insecta","Lepidoptera","Tortricidae","Tortricinae","Phtheochroa","Phtheochroa baracana, Phtheochroa riscana, Phtheochroa birdana, Phtheochroa fulviplicana, Phtheochroa canariana","1","1","1","149","0","0","99.78","450","0","450","29.56","0"
"AGAW1","09E","INS","BOLD:AAF9035","Arthropoda","Insecta","Diptera","Tipulidae","Tipulinae","Nephrotoma","Nephrotoma alterna","1","1","1","149","0","0","99.56","450","0","450","33.78","0"
"AGAW1","09E","INS","BOLD:ABW3192","Arthropoda","Insecta","Hymenoptera","Platygastridae","","","","1","1","1","149","0","0","99.55","446","0","450","26.67","0"
"AGAW1","09E","INS","BOLD:ACJ8771","Arthropoda","Insecta","Diptera","Chironomidae","Chironominae","Sublettea","","1","1","1","149","0","0","98.97","389","0","450","32.44","0"
"AGAW1","09E","INS","BOLD:AAA3889","Arthropoda","Insecta","Lepidoptera","Crambidae","Crambinae","Parapediasia","Parapediasia teterrellus","1","1","1","149","0","0","98.64","440","0","440","32.73","0"
"AGAW1","09E","INS","BOLD:ACF1257","","","","","","","","1","1","1","149","0","0","99.23","388","0","391","33.76","0"
"AGAW1","09E","INS","BOLD:AAA2632","Arthropoda","Insecta","Lepidoptera","Noctuidae","Noctuinae","Noctua","Noctua atlantica, Noctua pronuba","1","1","1","149","0","0","99.33","450","0","450","31.11","0"
"AGAW1","09E","INS","BOLD:ACV5891","Arthropoda","Insecta","Diptera","Cecidomyiidae","","","","1","1","1","149","0","0","97.11","346","0","350","23.14","0"
"AGAW1","09E","INS","BOLD:AAN5358","Arthropoda","Insecta","Diptera","Chironomidae","Orthocladiinae","Smittia","Smittia stercoraria, Smittia sp. 22ES","1","1","1","149","0","0","100","450","0","450","37.78","0"
"AGAW1","09E","INS","BOLD:ACF0890","Arthropoda","Insecta","Hymenoptera","Ichneumonidae","Ophioninae","Enicospilus","Enicospilus purgatusDHJ02","1","1","1","149","0","0","97.56","450","0","450","26.44","0"
"AGAW1","09E","INS","BOLD:AAB3870","Arthropoda","Insecta","Thysanoptera","Thripidae","Thripinae","Thrips","Thrips tabaci","1","1","1","149","0","0","98.44","448","0","450","34.67","0"
"AGAW1","09E","INS","BOLD:ABY5318","Arthropoda","Insecta","Diptera","Chloropidae","Oscinellinae","Incertella, Rhopalopterum","Rhopalopterum umbrosum, Rhopalopterum luteiceps, Incertella bispina, Rhopalopterum painteri","1","1","1","149","0","0","99.78","450","0","450","31.78","0"
"AGAW1","09E","INS","BOLD:ADG6553","Arthropoda","Insecta","Diptera","Chironomidae","","","","1","1","1","149","0","0","97.17","389","0","450","34","0"
"AGAW1","09E","INS","BOLD:AAG3236","Arthropoda","Insecta","Diptera","Phoridae","Phorinae","Conicera, Diplonevra","Diplonevra nitidula","1","1","1","149","0","0","98","449","0","450","32.89","0"
"AGAW1","09E","INS","BOLD:ACD8563","Arthropoda","Insecta","Diptera","Cecidomyiidae","","","","1","1","1","149","0","0","97.33","450","0","450","25.56","0"
"AGAW1","09E","INS","BOLD:AAC9062","Arthropoda","Insecta","Diptera","Culicidae","Culicinae","Ochlerotatus, Aedes","Aedes fitchii, Ochlerotatus fitchii, Aedes grossbecki, Aedes ventrovittis","1","1","1","149","0","0","98.22","450","0","450","33.11","0"
"AGAW1","09E","INS","BOLD:AAA3654","Arthropoda","Insecta","Lepidoptera","Crambidae","Crambinae","Crambus","Crambus bigelovi, Crambus johnsoni, Crambus praefectellus","1","1","1","149","0","0","98.44","449","0","450","31.56","0"
"AGAW1","09E","INS","BOLD:ABV2554","Arthropoda","Insecta","Hemiptera","Cicadellidae","Deltocephalinae","Sorhoanus, Stenometopiellus","Stenometopiellus sigillatus, Sorhoanus uhleri, Sorhoanus pascuellus","1","1","1","149","0","0","97.56","450","0","450","38.89","0"
"AGAW1","09E","INS","BOLD:AAA7374","Arthropoda","Insecta","Diptera","Syrphidae","Syrphinae","Sphaerophoria","Sphaerophoria bankowskae, Sphaerophoria infuscata, Sphaerophoria nr. philanthus, Sphaerophoria batava, Sphaerophoria sp., Sphaerophoria rueppellii, Sphaerophoria interrupta-group, Sphaerophoria sp. MA-1, Sphaerophoria cleoae, Sphaerophoria asymmetrica, Sphaerophoria fatarum, Sphaerophoria bengalensis, Sphaerophoria bifurcata, Sphaerophoria taeniata, Sphaerophoria interrupta, Sphaerophoria contigua, Sphaerophoria kaa, Sphaerophoria laurae, Sphaerophoria chongjini, Sphaerophoria boreoalpina, Sphaerophoria sp. BOLDAAA7374, Sphaerophoria macrogaster, Sphaerophoria sulphuripes, Sphaerophoria brevipilosa, Sphaerophoria philanthus, Sphaerophoria pyrrhina, Sphaerophoria abbreviata, Sphaerophoria scripta, Sphaerophoria potentillae, Sphaerophoria longipilosa, Sphaerophoria virgata","1","1","1","149","0","0","99.46","370","0","374","30.48","0"
"AGAW1","09E","INS","BOLD:ACF4925","Arthropoda","Insecta","Hemiptera","Miridae","Mirinae","Adelphocoris","Adelphocoris detritus","1","1","1","149","0","0","97.1","449","0","450","36.67","0"
"AGAW1","09E","INS","BOLD:AAA8796","Arthropoda","Insecta","Lepidoptera","Tortricidae","Tortricinae","Acleris","Acleris forsskaleana","1","1","1","149","0","0","97.17","424","0","450","32.44","0"
"AGAW1","09E","INS","BOLD:AAY6534","Arthropoda","Insecta","Coleoptera","Mordellidae","Mordellinae","Mordellina","Mordellina ancilla","1","1","1","149","0","0","98","449","0","450","38.89","0"
"AGAW1","09E","NAR","no match","no match","no match","no match","no match","no match","no match","no match","no match","no match","no match","no match","no match","no match","no match","no match","no match","no match","no match","no match"
"AGAW1","09E","NIN","BOLD:AAA7242","Arthropoda","Collembola","Entomobryomorpha","Entomobryidae","Seirinae, Entomobryinae","Entomobrya, Lepidosira","Entomobrya multifasciata, Lepidosira sp.","176","166","166","1","5.68","0","99.23","422.62","0","426.24","39.39","0"
"AGAW1","09E","NIN","BOLD:AAA6381","Arthropoda","Arachnida","Araneae","Tetragnathidae","","Tetragnatha","Tetragnatha laboriosa","118","116","116","2","1.69","0","99.31","429.1","0","431.12","39.45","0"
"AGAW1","09E","NIN","BOLD:AAH6630","Arthropoda","Collembola","Entomobryomorpha","Entomobryidae","Entomobryinae","Entomobrya","Entomobrya unostrigata","56","50","50","3","10.71","0","99.36","434.48","0","435.52","39.06","0"
"AGAW1","09E","NIN","BOLD:AAH8314","Arthropoda","Arachnida","Araneae","Linyphiidae","","Walckenaeria","Walckenaeria microspiralis","17","17","17","4","0","0","97.97","399.71","0","402.12","35.7","0"
"AGAW1","09E","NIN","BOLD:ABW2639","Arthropoda","Arachnida","Trombidiformes","Anystidae","","","","6","6","6","5","0","0","99.81","442.33","0","438.33","31.48","0"
"AGAW1","09E","NIN","BOLD:AAB7914","Arthropoda","Collembola","Symphypleona","Bourletiellidae","","Bourletiella","","4","4","4","6","0","0","99.94","448.75","0","450","38.22","0"
"AGAW1","09E","NIN","BOLD:ACX8969","Arthropoda","Arachnida","Trombidiformes","Eupodidae","","","","4","4","4","6","0","0","98.22","434.25","0","437.75","28.27","0"
"AGAW1","09E","NIN","BOLD:ACU7748","Arthropoda","Arachnida","Trombidiformes","Pygmephoridae","","","","2","2","2","8","0","0","99.12","395.5","0","450","29.56","0"
"AGAW1","09E","NIN","BOLD:ACN3178","Arthropoda","Collembola","Entomobryomorpha","Entomobryidae","","","","1","1","1","9","0","0","98.89","450","0","450","39.33","0"
"AGAW1","09E","NIN","BOLD:ACZ1295","Arthropoda","Arachnida","Trombidiformes","Anystidae","","","","1","1","1","9","0","0","100","382","0","384","31.25","0"
"AGAW1","09E","NIN","BOLD:AAC9044","Arthropoda","Arachnida","Araneae","Salticidae","","Salticus","Salticus scenicus","1","1","1","9","0","0","98.84","431","0","432","33.33","0"
"AGAW1","09E","NIN","BOLD:AAA4123","Arthropoda","Arachnida","Araneae","Araneidae","","Neoscona","Neoscona arabesca","1","1","1","9","0","0","98.43","383","0","398","30.9","0"
"AGAW1","09N","BAC","TAX:91393","Proteobacteria","Alphaproteobacteria","Rickettsiales","Rickettsiaceae",NA,"Wolbachia","","9","9","9","1","0","0","99.06","392","0","418.44","39.26","0"
"AGAW1","09N","BAC","TAX:175768","Proteobacteria","Alphaproteobacteria","Rickettsiales","Rickettsiaceae",NA,"Wolbachia","Wolbachia pipientis","6","6","6","2","0","0","98.33","448","0","450","39.48","0"
"AGAW1","09N","BAC","TAX:91391","Proteobacteria","Alphaproteobacteria","Rickettsiales","",NA,"","","1","1","1","3","0","0","100","384","0","450","42","0"
"AGAW1","09N","CHO","no match","no match","no match","no match","no match","no match","no match","no match","no match","no match","no match","no match","no match","no match","no match","no match","no match","no match","no match","no match"
"AGAW1","09N","INS","BOLD:AAA3453","Arthropoda","Insecta","Diptera","Anthomyiidae","Anthomyiinae","Delia","","22413","18505","18505","1","17.44","0","99.31","432.38","0","435.01","34.63","0"
"AGAW1","09N","INS","BOLD:AAP8739","Arthropoda","Insecta","Diptera","Platystomatidae","Platystomatinae","Rivellia","Rivellia steyskali","18042","11379","11379","2","36.93","0","99.36","441.24","0","443.21","37.97","0"
"AGAW1","09N","INS","BOLD:AAG2511","Arthropoda","Insecta","Diptera","Anthomyiidae","Anthomyiinae","Delia","Delia platura","11178","9109","9109","3","18.51","0","98.91","444.71","0","447.54","33.21","0"
"AGAW1","09N","INS","BOLD:AAG6743","Arthropoda","Insecta","Diptera","Sarcophagidae","Sarcophaginae","Sarcophaga","Sarcophaga sp. 1 MG, Sarcophaga subvicina","8810","6600","6600","4","25.09","0","99.37","436.75","0","438.33","34.83","0"
"AGAW1","09N","INS","BOLD:AAG6745","Arthropoda","Insecta","Diptera","Calliphoridae","Calliphorinae","Pollenia","Pollenia pediculata","8535","7451","7451","5","12.7","0","99.32","407.55","0","409.29","33.53","0"
"AGAW1","09N","INS","BOLD:ACK9316","Arthropoda","Insecta","Hemiptera","Cicadellidae","Macropsinae","Macropsis","Macropsis trivialis","2405","1512","1512","6","37.13","0","98.49","447.53","0","448.31","33.94","0"
"AGAW1","09N","INS","BOLD:AAH4135","Arthropoda","Insecta","Diptera","Chloropidae","Chloropinae","Thaumatomyia","Thaumatomyia glabra, Thaumatomyia grata, Thaumatomyia sp. 1 TAW","1550","1392","1392","7","10.19","0","98.92","429.45","0","432.99","31.76","0"
"AGAW1","09N","INS","BOLD:AAB7030","Arthropoda","Insecta","Diptera","Chironomidae","Chironominae","Chironomus","Chironomus sp. BOLDAAP2996, Chironomus quinnitukqut, Chironomus cf. decorus 2g, Chironomus decorus group sp. 2 IP2013, Chironomus cf. decorus, Chironomus decorus gr.","1024","1020","1020","8","0.39","0","97.9","404.48","0","414.11","33.74","0"
"AGAW1","09N","INS","BOLD:AAN6154","Arthropoda","Insecta","Coleoptera","Latridiidae","Corticariinae","Melanophthalma","Melanophthalma inermis","607","504","504","9","16.97","0","99.16","437.36","0","438.61","31.33","0"
"AGAW1","09N","INS","BOLD:AAM6227","Arthropoda","Insecta","Diptera","Chironomidae","Tanypodinae","Procladius","Procladius denticulatus, Procladius sp.","509","494","494","10","2.95","0","99.25","422.73","0","425.71","31.08","0"
"AGAW1","09N","INS","BOLD:AAD2041","Arthropoda","Insecta","Diptera","Dolichopodidae","Sciapodinae","Condylostylus","Condylostylus inermis, Condylostylus nigrofemoratus, Condylostylus connectans, Condylostylus caudatus, Condylostylus flavipes","410","326","326","11","20.49","0","99.44","436.1","0","437.58","36.05","0"
"AGAW1","09N","INS","BOLD:AAG5507","Arthropoda","Insecta","Diptera","Chironomidae","Chironominae","Chironomus","Chironomus atrella","388","383","383","12","1.29","0","99.1","425.31","0","432.11","36","0"
"AGAW1","09N","INS","BOLD:ACK7314","Arthropoda","Insecta","Diptera","Chironomidae","Chironominae","Polypedilum","","367","363","363","13","1.09","0","98.67","391.37","0","395.98","35.17","0"
"AGAW1","09N","INS","BOLD:AAC9062","Arthropoda","Insecta","Diptera","Culicidae","Culicinae","Ochlerotatus, Aedes","Aedes fitchii, Ochlerotatus fitchii, Aedes grossbecki, Aedes ventrovittis","353","311","311","14","11.9","0","99.36","432.55","0","434.15","33.13","0"
"AGAW1","09N","INS","BOLD:ACK3589","Arthropoda","Insecta","Diptera","Cecidomyiidae","","","","329","265","265","15","19.45","0","99.19","447.93","0","448.83","32.26","0"
"AGAW1","09N","INS","BOLD:AAU6681","Arthropoda","Insecta","Diptera","Tipulidae","Tipulinae","Nephrotoma","Nephrotoma cornicina","294","242","242","16","17.69","0","99.54","443.92","0","445.02","36.29","0"
"AGAW1","09N","INS","BOLD:AAA7661","Arthropoda","Insecta","Diptera","Culicidae","Culicinae","Culiseta, Culex","Culiseta melanura, Culex articularis, Culex restuans, Culex acharistus","281","270","270","17","3.91","0","99.04","436.23","0","438.08","32.86","0"
"AGAW1","09N","INS","BOLD:AAD0484","Arthropoda","Insecta","Diptera","Chironomidae","","","","280","225","225","18","19.64","0","98.67","446.04","0","447.39","35.1","0"
"AGAW1","09N","INS","BOLD:AAB0373","Arthropoda","Insecta","Neuroptera","Chrysopidae","Chrysopinae","Chrysoperla","Chrysoperla sp. isolate 82, Chrysoperla zastrowi, Chrysoperla plorabunda, Chrysoperla sp. isolate 71, Chrysoperla agilis, Italochrysa italica, Chrysoperla sp. isolate 88, Chrysoperla lucasina, Chrysoperla carnea, Chrysoperla mediterranea, Chrysoperla pallida, Chrysoperla nipponensis, Chrysoperla sillemi","262","219","219","19","16.41","0","99.47","437.01","0","438.02","31.36","0"
"AGAW1","09N","INS","BOLD:AAH3661","Arthropoda","Insecta","Diptera","Cecidomyiidae","","","","243","228","228","20","6.17","0","99.27","434.07","0","435.95","25.5","0"
"AGAW1","09N","INS","BOLD:ACX5521","Arthropoda","Insecta","Diptera","Chironomidae","Orthocladiinae","","","224","213","213","21","4.91","0","98.94","415.06","0","417.27","34.94","0"
"AGAW1","09N","INS","BOLD:AAG2868","Arthropoda","Insecta","Hemiptera","Cicadellidae","Typhlocybinae","Empoasca","Empoasca sp., Empoasca fabae","213","151","151","22","29.11","0","99.55","443.05","0","447.85","32.55","0"
"AGAW1","09N","INS","BOLD:AAN5342","Arthropoda","Insecta","Diptera","Chironomidae","Orthocladiinae","Heleniella","","213","209","209","22","1.88","0","98.93","422.18","0","429.36","35.26","0"
"AGAW1","09N","INS","BOLD:ABA0863","Arthropoda","Insecta","Diptera","Cecidomyiidae","","","","211","176","176","24","16.59","0","99.36","439.85","0","441.2","29.76","0"
"AGAW1","09N","INS","BOLD:AAK8751","Arthropoda","Insecta","Diptera","Agromyzidae","Phytomyzinae","Liriomyza","Liriomyza sp. SB, Liriomyza trifoliearum","203","174","174","25","14.29","0","98.75","437.5","0","438.72","32.48","0"
"AGAW1","09N","INS","BOLD:AAN5238","Arthropoda","Insecta","Diptera","Cecidomyiidae","","","","197","157","157","26","20.3","0","99.59","433.72","0","434.83","25.69","0"
"AGAW1","09N","INS","BOLD:ABU5525","Arthropoda","Insecta","Diptera","Chironomidae","Orthocladiinae","Limnophyes","Limnophyes sp. 14ES","179","177","177","27","1.12","0","99.03","401.57","0","406.28","37.08","0"
"AGAW1","09N","INS","BOLD:ACA3036","Arthropoda","Insecta","Diptera","Chironomidae","Chironominae","Cladotanytarsus","","172","160","160","28","6.98","0","99.16","424","0","426.52","39.25","0"
"AGAW1","09N","INS","BOLD:AAG9897","Arthropoda","Insecta","Dermaptera","Forficulidae","Forficulinae","Forficula","Forficula auricularia-A","160","159","159","29","0.63","0","98.97","398.78","0","409.71","45.7","0"
"AGAW1","09N","INS","BOLD:ABV1297","Arthropoda","Insecta","Diptera","Sciaridae","","","","152","150","150","30","1.32","0","98.97","430.56","0","433.3","30.22","0"
"AGAW1","09N","INS","BOLD:AAN5148","Arthropoda","Insecta","Diptera","Ceratopogonidae","Forcipomyiinae","Forcipomyia","Forcipomyia bipunctata, Forcipomyia sp. CK-2012","135","125","125","31","7.41","0","97.84","424.1","0","425.78","35.25","0"
"AGAW1","09N","INS","BOLD:ACE2676","Arthropoda","Insecta","Diptera","Chloropidae","Oscinellinae","Dicraeus","Dicraeus fennicus","132","120","120","32","9.09","0","98.36","431.99","0","433.9","34.06","0"
"AGAW1","09N","INS","BOLD:AAM9255","Arthropoda","Insecta","Diptera","Sciaridae","","Pseudolycoriella","","118","117","117","33","0.85","0","99.18","416.25","0","418.81","30.68","0"
"AGAW1","09N","INS","BOLD:AAI1549","Arthropoda","Insecta","Hymenoptera","Braconidae","Euphorinae","Meteorus","","115","95","95","34","17.39","0","98.89","444.42","0","445.54","31.25","0"
"AGAW1","09N","INS","BOLD:ABW1379","Arthropoda","Insecta","Diptera","Chloropidae","Oscinellinae","Malloewia","Malloewia abdominalis","105","82","82","35","21.9","0","97.79","438.05","0","447.36","32.19","0"
"AGAW1","09N","INS","BOLD:AAO3132","Arthropoda","Insecta","Coleoptera","Chrysomelidae","Galerucinae","Neocrepidodera","Neocrepidodera ferruginea","103","102","102","36","0.97","0","99.13","378.16","0","380.39","35.13","0"
"AGAW1","09N","INS","BOLD:ABA7871","Arthropoda","Insecta","Diptera","Cecidomyiidae","","","","102","87","87","37","14.71","0","98.87","441.21","0","442.01","26.85","0"
"AGAW1","09N","INS","BOLD:ABA0772","Arthropoda","Insecta","Diptera","Chironomidae","Chironominae","Polypedilum","","94","91","91","38","3.19","0","98.65","424.62","0","432.35","33.31","0"
"AGAW1","09N","INS","BOLD:AAG3236","Arthropoda","Insecta","Diptera","Phoridae","Phorinae","Conicera, Diplonevra","Diplonevra nitidula","90","77","77","39","14.44","0","99.46","448.58","0","450","32.08","0"
"AGAW1","09N","INS","BOLD:AAN5199","Arthropoda","Insecta","Diptera","Cecidomyiidae","","","CecidInt20 sp. BOLD:AAN5199","89","73","73","40","17.98","0","99.49","430.72","0","447.2","26.86","0"
"AGAW1","09N","INS","BOLD:ABV3225","Arthropoda","Insecta","Diptera","Sphaeroceridae","","","SphaerocerIntGen5 sp. BOLD:ABV3225","84","72","72","41","14.29","0","99.36","442.96","0","444.71","37.11","0"
"AGAW1","09N","INS","BOLD:ACK3431","Arthropoda","Insecta","Diptera","Cecidomyiidae","","","","76","73","73","42","3.95","0","99.17","426.21","0","428.38","32.76","0"
"AGAW1","09N","INS","BOLD:AAG7891","Arthropoda","Insecta","Hymenoptera","Platygastridae","Scelioninae","Telenomus","Telenomus podisi","74","63","63","43","14.86","0","99.15","425.85","0","434.3","26.92","0"
"AGAW1","09N","INS","BOLD:AAH0135","Arthropoda","Insecta","Coleoptera","Phalacridae","Phalacrinae","Acylomus","Acylomus pugetanus, Acylomus ergoti","74","72","72","43","2.7","0","99.22","431.43","0","433.62","31.43","0"
"AGAW1","09N","INS","BOLD:AAF9771","Arthropoda","Insecta","Diptera","Hybotidae","Tachydromiinae","Platypalpus","Platypalpus longiseta, Platypalpus trivialis, Platypalpus sp. 5, Platypalpus sp. 17, Platypalpus pallidiventris","70","60","60","45","14.29","0","98.95","442.9","0","443.8","34.93","0"
"AGAW1","09N","INS","BOLD:AAD5253","Arthropoda","Insecta","Diptera","Chironomidae","Orthocladiinae","Thienemanniella","Thienemanniella xena, Thienemanniella vittata","68","57","57","46","16.18","0","98.78","440.63","0","441.99","33.82","0"
"AGAW1","09N","INS","BOLD:AAG3154","Arthropoda","Insecta","Diptera","Psychodidae","Psychodinae","Psychoda","","68","55","55","46","19.12","0","98.45","444.69","0","448.96","32.48","0"
"AGAW1","09N","INS","BOLD:ADH5973","Arthropoda","Insecta","Diptera","Psychodidae","","","","66","62","62","48","6.06","0","99.38","388.83","0","433.98","34.56","0"
"AGAW1","09N","INS","BOLD:AAH3664","Arthropoda","Insecta","Diptera","Cecidomyiidae","","","CecidInt3 sp. BOLD:AAH3664","58","57","57","49","1.72","0","99.14","430.84","0","435.33","26.99","0"
"AGAW1","09N","INS","BOLD:ACE7444","Arthropoda","Insecta","Hemiptera","Miridae","Mirinae, Phylinae","Adelphocoris, Megalocoleus","Adelphocoris lineolatus, Megalocoleus molliculus","58","46","46","49","20.69","0","99.64","446.86","0","447.67","34.87","0"
"AGAW1","09N","INS","BOLD:AAG6934","Arthropoda","Insecta","Diptera","Hybotidae","","","","47","40","40","51","14.89","0","99.38","434.28","0","435.49","38.68","0"
"AGAW1","09N","INS","BOLD:ACD3384","Arthropoda","Insecta","Hemiptera","Miridae","Orthotylinae","Orthotylus","Orthotylus robiniae","41","39","39","52","4.88","0","99.6","431.17","0","432.39","41.45","0"
"AGAW1","09N","INS","BOLD:AAN6447","Arthropoda","Insecta","Diptera","Sciaridae","","Corynoptera","Corynoptera perpusilla","41","36","36","52","12.2","0","99.46","448.12","0","449.73","36.3","0"
"AGAW1","09N","INS","BOLD:AAL4232","Arthropoda","Insecta","Diptera","Agromyzidae","Phytomyzinae","Phytoliriomyza","Phytoliriomyza arctica","38","34","34","54","10.53","0","99.41","442.82","0","444.37","32.25","0"
"AGAW1","09N","INS","BOLD:ACX8294","Arthropoda","Insecta","Diptera","Cecidomyiidae","","","","37","35","35","55","5.41","0","99.32","444.7","0","445.86","32.45","0"
"AGAW1","09N","INS","BOLD:AAD4214","Arthropoda","Insecta","Hymenoptera","Ichneumonidae","Diplazontinae","Diplazon","Diplazon laetatorius","37","31","31","55","16.22","0","99.46","445.54","0","446.62","25.74","0"
"AGAW1","09N","INS","BOLD:AAL7819","Arthropoda","Insecta","Diptera","Psychodidae","Psychodinae","Psychoda","Psychoda sp., Psychoda sp. BOLD:AAL7819","34","32","32","57","5.88","0","99.14","415.94","0","421.09","32.8","0"
"AGAW1","09N","INS","BOLD:ABX0657","Arthropoda","Insecta","Coleoptera","Anthicidae","Notoxinae","Notoxus","Notoxus desertus","32","32","32","58","0","0","99.07","436.25","0","437.56","35.87","0"
"AGAW1","09N","INS","BOLD:AAN5225","Arthropoda","Insecta","Diptera","Cecidomyiidae","","","","31","28","28","59","9.68","0","99.21","447.94","0","448.94","28.41","0"
"AGAW1","09N","INS","BOLD:AAM7680","Arthropoda","Insecta","Coleoptera","Latridiidae","Corticariinae","Melanophthalma","Melanophthalma pumila","30","24","24","60","20","0","97.65","432.47","0","433.13","34.03","0"
"AGAW1","09N","INS","BOLD:AAG1711","Arthropoda","Insecta","Diptera","Muscidae","Phaoniinae","Helina","Helina depuncta","30","29","29","60","3.33","0","99.21","437.77","0","440.1","31.37","0"
"AGAW1","09N","INS","BOLD:ACC8303","Arthropoda","Insecta","Diptera","Cecidomyiidae","","","","27","27","27","62","0","0","99.24","431.07","0","433.78","26.27","0"
"AGAW1","09N","INS","BOLD:AAA5299","Arthropoda","Insecta","Diptera","Chironomidae","Orthocladiinae","Cricotopus","Cricotopus laetus, Cricotopus relucens, Cricotopus trifasciatus, Cricotopus sp., Cricotopus sp. 3ES, Cricotopus sylvestris, Cricotopus glacialis","26","26","26","63","0","0","98.97","431.58","0","433.27","34.86","0"
"AGAW1","09N","INS","BOLD:AAH0134","Arthropoda","Insecta","Coleoptera","Phalacridae","Phalacrinae","Stilbus","Stilbus apicialis","25","23","23","64","8","0","99.05","428.8","0","429.72","31.81","0"
"AGAW1","09N","INS","BOLD:AAM7726","Arthropoda","Insecta","Coleoptera","Curculionidae","Curculioninae","Orchestes","Orchestes steppensis, Orchestes alni","22","17","17","65","22.73","0","99.16","439.55","0","442.59","33.39","0"
"AGAW1","09N","INS","BOLD:AAN8770","Arthropoda","Insecta","Diptera","Psychodidae","Psychodinae","Psychoda","Psychoda trinodulosa","20","14","14","66","30","0","99.8","445.95","0","446.3","34.03","0"
"AGAW1","09N","INS","BOLD:AAG5443","Arthropoda","Insecta","Diptera","Chironomidae","Orthocladiinae","","","17","17","17","67","0","0","98.74","409.71","0","411.24","33.39","0"
"AGAW1","09N","INS","BOLD:ABV1455","Arthropoda","Insecta","Diptera","Chironomidae","Chironominae","","","17","17","17","67","0","0","98.12","439.65","0","442","31.81","0"
"AGAW1","09N","INS","BOLD:AAM9013","Arthropoda","Insecta","Diptera","Keroplatidae","Macrocerinae","Macrocera","Macrocera pusilla","17","17","17","67","0","0","99.08","429.35","0","431.18","30.04","0"
"AGAW1","09N","INS","BOLD:AAH3662","Arthropoda","Insecta","Diptera","Cecidomyiidae","","","CecidInt3 sp. BOLD:AAH3662","16","16","16","70","0","0","99.23","448.31","0","450","26.22","0"
"AGAW1","09N","INS","BOLD:ACM0828","Arthropoda","Insecta","Diptera","Chironomidae","Chironominae","Tanytarsus","","15","15","15","71","0","0","98.19","420.33","0","422.07","35.65","0"
"AGAW1","09N","INS","BOLD:AAP6384","Arthropoda","Insecta","Diptera","Psychodidae","","","","14","10","10","72","28.57","0","97.38","449.21","0","450","35.4","0"
"AGAW1","09N","INS","BOLD:ADH9867","Arthropoda","Insecta","Diptera","","","","","13","13","13","73","0","0","98.71","397.46","0","401.23","37.4","0"
"AGAW1","09N","INS","BOLD:ABV1475","Arthropoda","Insecta","Diptera","Cecidomyiidae","","","","12","11","11","74","8.33","0","99.06","445","0","446.08","26.43","0"
"AGAW1","09N","INS","BOLD:AAN5267","Arthropoda","Insecta","Diptera","Cecidomyiidae","","","","12","10","10","74","16.67","0","99.74","449.42","0","450","30.7","0"
"AGAW1","09N","INS","BOLD:AAG8493","Arthropoda","Insecta","Diptera","Drosophilidae","Drosophilinae","Scaptomyza","Scaptomyza pallida","10","10","10","76","0","0","98.87","380.8","0","382.3","32.91","0"
"AGAW1","09N","INS","BOLD:AAN8446","Arthropoda","Insecta","Psocodea","Peripsocidae","","Peripsocus","Peripsocus madidus","9","9","9","77","0","0","99.12","420","0","420.56","41.3","0"
"AGAW1","09N","INS","BOLD:ACJ5126","Arthropoda","Insecta","Diptera","Chironomidae","Orthocladiinae","Metriocnemus","","9","9","9","77","0","0","98.66","426.22","0","429.33","34.84","0"
"AGAW1","09N","INS","BOLD:AAC0903","Arthropoda","Insecta","Diptera","Chironomidae","Chironominae","Chironomus","Chironomus acidophilus","7","7","7","79","0","0","98.94","446.57","0","447.43","36.33","0"
"AGAW1","09N","INS","BOLD:ACK5742","Arthropoda","Insecta","Diptera","Cecidomyiidae","","","","6","6","6","80","0","0","99.25","448","0","448.83","29.07","0"
"AGAW1","09N","INS","BOLD:ABV9277","","","","","","","","6","6","6","80","0","0","98.76","434.33","0","435.67","27.95","0"
"AGAW1","09N","INS","BOLD:AAV5573","Arthropoda","Insecta","Diptera","Cecidomyiidae","","","","6","6","6","80","0","0","99.93","442.17","0","443.33","27.46","0"
"AGAW1","09N","INS","BOLD:AAN5205","Arthropoda","Insecta","Diptera","Cecidomyiidae","","","CecidInt42 sp. BOLD:AAN5205","5","5","5","83","0","0","98.93","447.2","0","450","31.56","0"
"AGAW1","09N","INS","BOLD:AAI4303","Arthropoda","Insecta","Diptera","Chironomidae","Chironominae","Chironomus","Chironomus melanescens","5","5","5","83","0","0","98.74","444.4","0","450","34.4","0"
"AGAW1","09N","INS","BOLD:ACF5830","Arthropoda","Insecta","Diptera","Chloropidae","Chloropinae","Thaumatomyia","","5","5","5","83","0","0","98.32","419","0","422.4","31.41","0"
"AGAW1","09N","INS","BOLD:AAM7855","Arthropoda","Insecta","Thysanoptera","Thripidae","Thripinae","Frankliniella","Frankliniella tritici","4","4","4","86","0","0","99.28","448.25","0","450","31.22","0"
"AGAW1","09N","INS","BOLD:AAF9187","Arthropoda","Insecta","Coleoptera","Curculionidae","Entiminae","Phyllobius","Phyllobius oblongus","4","4","4","86","0","0","99.2","390.5","0","438.5","38.93","0"
"AGAW1","09N","INS","BOLD:AAF4817","Arthropoda","Insecta","Diptera","Chironomidae","Orthocladiinae","Smittia","Smittia edwardsi","4","4","4","86","0","0","98.31","375.75","0","450","33.67","0"
"AGAW1","09N","INS","BOLD:ADC1124","Arthropoda","Insecta","Diptera","Chironomidae","Chironominae","Tanytarsus","","3","3","3","89","0","0","97.26","389","0","396.67","35.38","0"
"AGAW1","09N","INS","BOLD:ACM0158","Arthropoda","Insecta","Trichoptera","Hydroptilidae","Hydroptilinae","Hydroptila","","3","3","3","89","0","0","97.77","434.67","0","434.67","32.58","0"
"AGAW1","09N","INS","BOLD:AAA4277","Arthropoda","Insecta","Diptera","Syrphidae","Syrphinae","Toxomerus","Toxomerus marginatus, Toxomerus sp. CR-3, Toxomerus CR-3","3","3","3","89","0","0","98.96","450","0","450","30.81","0"
"AGAW1","09N","INS","BOLD:AAG6460","Arthropoda","Insecta","Diptera","Cecidomyiidae","","","","3","3","3","89","0","0","98.81","448","0","450","34.59","0"
"AGAW1","09N","INS","BOLD:AAN5178","Arthropoda","Insecta","Diptera","Cecidomyiidae","Cecidomyiinae","Rhopalomyia","Rhopalomyia sp. BOLD:AAN5178","3","3","3","89","0","0","98.2","410.33","0","415.67","26.54","0"
"AGAW1","09N","INS","BOLD:AAG2460","Arthropoda","Insecta","Diptera","Anthomyiidae","Anthomyiinae","Hydrophoria","Hydrophoria lancifer","3","2","2","89","33.33","0","98.06","433","0","433.33","33.9","0"
"AGAW1","09N","INS","BOLD:AAA3181","Arthropoda","Insecta","Diptera","Muscidae","Muscinae","Stomoxys","Stomoxys calcitrans","2","2","2","95","0","0","98.65","445.5","0","450","33.11","0"
"AGAW1","09N","INS","BOLD:AAA6292","Arthropoda","Insecta","Lepidoptera","Tortricidae","Tortricinae","Cnephasia","Cnephasia genitalana, Cnephasia stephensiana","2","2","2","95","0","0","99.43","441.5","0","444","35.59","0"
"AGAW1","09N","INS","BOLD:ABA0858","Arthropoda","Insecta","Diptera","Cecidomyiidae","","","","2","2","2","95","0","0","98.64","371","0","372.5","26.04","0"
"AGAW1","09N","INS","BOLD:AAG5453","Arthropoda","Insecta","Diptera","Chironomidae","Chironominae","Chironomus","Chironomus bifurcatus","2","2","2","95","0","0","97.44","449.5","0","450","34.89","0"
"AGAW1","09N","INS","BOLD:ABA1231","Arthropoda","Insecta","Coleoptera, Diptera","Sciaridae","","Lycoriella","","2","2","2","95","0","0","99.4","425.5","0","427.5","35.75","0"
"AGAW1","09N","INS","BOLD:AAA2632","Arthropoda","Insecta","Lepidoptera","Noctuidae","Noctuinae","Noctua","Noctua atlantica, Noctua pronuba","2","2","2","95","0","0","99.22","449.5","0","450","31.22","0"
"AGAW1","09N","INS","BOLD:ABW3236","Arthropoda","Insecta","Hymenoptera","Platygastridae","","","","2","2","2","95","0","0","98.88","444.5","0","450","27.89","0"
"AGAW1","09N","INS","BOLD:ACF7189","Arthropoda","Insecta","Hymenoptera","Platygastridae","","","","2","2","2","95","0","0","98.87","442","0","446.5","26.2","0"
"AGAW1","09N","INS","BOLD:AAA2815","Arthropoda","Insecta","Lepidoptera","Noctuidae","Noctuinae","Mythimna","Mythimna yukonensis, Mythimna oxygala","2","2","2","95","0","0","99.35","387.5","0","389","33.43","0"
"AGAW1","09N","INS","BOLD:AAV5400","Arthropoda","Insecta","Diptera","Cecidomyiidae","","","","2","2","2","95","0","0","98.34","419","0","421.5","29.04","0"
"AGAW1","09N","INS","BOLD:AAC1312","Arthropoda","Insecta","Diptera","Syrphidae","Syrphinae","Toxomerus","Toxomerus geminatus, Toxomerus 75-5, Toxomerus rombicus, Toxomerus sp. 75-5","2","2","2","95","0","0","97.91","432.5","0","433.5","30.19","0"
"AGAW1","09N","INS","BOLD:ABV9293","Arthropoda","Insecta","Diptera","Chironomidae","Chironominae","Dicrotendipes","","2","2","2","95","0","0","98.87","355","0","435","34.71","0"
"AGAW1","09N","INS","BOLD:AAH7142","Arthropoda","Insecta","Diptera","Sarcophagidae","Sarcophaginae","Oxysarcodexia, Sarcophaginae_incertae_sedis","Oxysarcodexia cingarus, Sarcophaginae_incertae_sedis sp. 1 AGT, Oxysarcodexia ventricosa","2","2","2","95","0","0","99.19","431.5","0","433","33.58","0"
"AGAW1","09N","INS","BOLD:AAZ0271","Arthropoda","Insecta","Diptera","Ceratopogonidae","Ceratopogoninae","Culicoides","","2","2","2","95","0","0","98.81","410","0","413.5","41.1","0"
"AGAW1","09N","INS","BOLD:ABX0893","Arthropoda","Insecta","Diptera","Cecidomyiidae","","","","2","2","2","95","0","0","98.11","424.5","0","425.5","25.72","0"
"AGAW1","09N","INS","BOLD:AAY6469","Arthropoda","Insecta","Diptera","Cecidomyiidae","","","","2","2","2","95","0","0","99.76","414.5","0","416","25.68","0"
"AGAW1","09N","INS","BOLD:ABW3687","Arthropoda","Insecta","Diptera","Lauxaniidae","Homoneurinae","Homoneura","Homoneura bispina","1","1","1","111","0","0","98.24","398","0","399","33.83","0"
"AGAW1","09N","INS","BOLD:AAG1714","Arthropoda","Insecta","Diptera","Muscidae","Azeliinae","Muscina","Muscina pascuorum","1","1","1","111","0","0","97.91","431","0","450","33.56","0"
"AGAW1","09N","INS","BOLD:ABV1415","Arthropoda","Insecta","Diptera","Chironomidae","Chironominae","","","1","1","1","111","0","0","99.78","449","0","450","32.67","0"
"AGAW1","09N","INS","BOLD:ACJ6954","Arthropoda","Insecta","Diptera","Cecidomyiidae","","","","1","1","1","111","0","0","99.12","340","0","438","26.48","0"
"AGAW1","09N","INS","BOLD:AAB4321","Arthropoda","Insecta","Lepidoptera","Gelechiidae","Gelechiinae","Metzneria","Metzneria lappella","1","1","1","111","0","0","98.33","359","0","360","31.39","0"
"AGAW1","09N","INS","BOLD:AAF9035","Arthropoda","Insecta","Diptera","Tipulidae","Tipulinae","Nephrotoma","Nephrotoma alterna","1","1","1","111","0","0","97.11","450","0","450","34","0"
"AGAW1","09N","INS","BOLD:ABY9688","Arthropoda","Insecta","Diptera","Chloropidae","Chloropinae","Thaumatomyia","Thaumatomyia glabra","1","1","1","111","0","0","97.51","402","0","404","31.19","0"
"AGAW1","09N","INS","BOLD:AAN6406","Arthropoda","Insecta","Diptera","Sphaeroceridae","Limosininae","Opalimosina","Opalimosina mirabilis","1","1","1","111","0","0","99.33","449","0","450","32.67","0"
"AGAW1","09N","INS","BOLD:AAC9132","Arthropoda","Insecta","Diptera","Culicidae","Culicinae","Culiseta","Culiseta inornata","1","1","1","111","0","0","97.09","446","0","450","32.89","0"
"AGAW1","09N","INS","BOLD:AAG5457","Arthropoda","Insecta","Diptera","Chironomidae","Chironominae","","","1","1","1","111","0","0","98.07","415","0","417","33.81","0"
"AGAW1","09N","INS","BOLD:AAN5255","Arthropoda","Insecta","Diptera","Cecidomyiidae","Cecidomyiinae","Dasineura","","1","1","1","111","0","0","98","449","0","450","27.11","0"
"AGAW1","09N","INS","BOLD:ACE8878","Arthropoda","Insecta","Orthoptera","Acrididae","Melanoplinae","Melanoplus","Melanoplus scudderi group","1","1","1","111","0","0","97.18","425","0","426","36.15","0"
"AGAW1","09N","INS","BOLD:AAY6412","Arthropoda","Insecta","Diptera","Sciaridae","","Bradysia","Bradysia urticae","1","1","1","111","0","0","97.99","447","0","450","31.56","0"
"AGAW1","09N","INS","BOLD:ACE1229","Arthropoda","Insecta","Diptera","Sciaridae","","","","1","1","1","111","0","0","99.04","416","0","433","30.25","0"
"AGAW1","09N","INS","BOLD:AAV5575","Arthropoda","Insecta","Diptera","Cecidomyiidae","","","CecidInt30 sp. BOLD:AAV5575","1","1","1","111","0","0","98.67","451","0","450","25.78","0"
"AGAW1","09N","INS","BOLD:AAA8796","Arthropoda","Insecta","Lepidoptera","Tortricidae","Tortricinae","Acleris","Acleris forsskaleana","1","1","1","111","0","0","99.15","352","0","355","31.83","0"
"AGAW1","09N","INS","BOLD:AAD1485","Arthropoda","Insecta","Diptera","Chironomidae","Chironominae","Paratanytarsus","Paratanytarsus grimmii, Paratanytarsus sp.","1","1","1","111","0","0","99.55","449","0","450","33.78","0"
"AGAW1","09N","INS","BOLD:AAD1945","Arthropoda","Insecta","Diptera","Agromyzidae","Phytomyzinae","Cerodontha","Cerodontha dorsalis","1","1","1","111","0","0","99.78","449","0","450","32.22","0"
"AGAW1","09N","INS","BOLD:AAA2314","Arthropoda","Insecta","Lepidoptera","Crambidae","Crambinae","Pediasia","Pediasia trisecta, Pediasia laciniellus, Pediasia sp.","1","1","1","111","0","0","98.66","448","0","450","33.56","0"
"AGAW1","09N","INS","BOLD:AAB5609","Arthropoda","Insecta","Diptera","Muscidae","Coenosiinae","Coenosia","Coenosia sp.BOLD:AAB5609, Coenosia tigrina","1","1","1","111","0","0","99.55","448","0","450","34","0"
"AGAW1","09N","INS","BOLD:AAG9626","Arthropoda","Insecta","Diptera","Dolichopodidae","","","","1","1","1","111","0","0","98.89","450","0","450","37.56","0"
"AGAW1","09N","INS","BOLD:AAL7370","Arthropoda","Insecta","Diptera","Chironomidae","Tanypodinae","Procladius","","1","1","1","111","0","0","98.22","449","0","450","32","0"
"AGAW1","09N","INS","BOLD:AAN5341","Arthropoda","Insecta","Diptera","Chironomidae","Orthocladiinae","Camptocladius","Camptocladius stercorarius","1","1","1","111","0","0","99.21","378","0","383","34.99","0"
"AGAW1","09N","INS","BOLD:AAA3867","Arthropoda","Insecta","Lepidoptera","Noctuidae","Noctuinae","Apamea","Apamea amputatrix","1","1","1","111","0","0","98.51","402","0","407","32.43","0"
"AGAW1","09N","INS","BOLD:ABY5318","Arthropoda","Insecta","Diptera","Chloropidae","Oscinellinae","Incertella, Rhopalopterum","Rhopalopterum umbrosum, Rhopalopterum luteiceps, Incertella bispina, Rhopalopterum painteri","1","1","1","111","0","0","97.77","448","0","449","31.85","0"
"AGAW1","09N","INS","BOLD:ACZ5284","Arthropoda","Insecta","Diptera","Sciomyzidae","Sciomyzinae","Dictya","Dictya stricta, Dictya expansa, Dictya floridensis, Dictya borealis, Dictya gaigei, Dictya sp. TAW1, Dictya umbroides, Dictya texensis, Dictya steyskali, Dictya pictipes","1","1","1","111","0","0","99.55","447","0","450","35.78","0"
"AGAW1","09N","INS","BOLD:AAB1098","Arthropoda","Insecta","Diptera","Culicidae","Culicinae","Ochlerotatus, Aedes","Aedes excrucians, Aedes cantans, Aedes stimulans, Aedes euedes, Aedes riparius, Aedes annulipes, Ochlerotatus cantans, Aedes fitchii, Aedes sp., Ochlerotatus annulipes, Ochlerotatus excrucians","1","1","1","111","0","0","99.33","447","0","450","35.33","0"
"AGAW1","09N","INS","BOLD:AAM6037","Arthropoda","Insecta","Diptera","Cecidomyiidae","Cecidomyiinae","Contarinia, Dasineura","","1","1","1","111","0","0","98.67","450","0","450","26.67","0"
"AGAW1","09N","INS","BOLD:AAA8279","Arthropoda","Insecta","Lepidoptera","Noctuidae","Noctuinae","Neoligia","Neoligia subjuncta","1","1","1","111","0","0","100","450","0","450","30.89","0"
"AGAW1","09N","INS","BOLD:ABV3224","Arthropoda","Insecta","Diptera","Sphaeroceridae","","","","1","1","1","111","0","0","99.33","450","0","450","32.67","0"
"AGAW1","09N","INS","BOLD:AAA3654","Arthropoda","Insecta","Lepidoptera","Crambidae","Crambinae","Crambus","Crambus bigelovi, Crambus johnsoni, Crambus praefectellus","1","1","1","111","0","0","98.22","449","0","450","31.78","0"
"AGAW1","09N","INS","BOLD:ABV0488","Arthropoda","Insecta","Diptera","Cecidomyiidae","","","","1","1","1","111","0","0","98","450","0","450","28.44","0"
"AGAW1","09N","INS","BOLD:AAC4197","Arthropoda","Insecta","Diptera","Chironomidae","Orthocladiinae","Paraphaenocladius","Paraphaenocladius impensus","1","1","1","111","0","0","97.99","447","0","450","34.89","0"
"AGAW1","09N","INS","BOLD:AAC2637","Arthropoda","Insecta","Lepidoptera","Pterophoridae","Pterophorinae","Geina","Geina sheppardi","1","1","1","111","0","0","98.92","369","0","369","30.08","0"
"AGAW1","09N","INS","BOLD:AAA1513","Arthropoda","Insecta","Lepidoptera","Plutellidae","Plutellinae","Plutella, Rhigognostis","Plutella xylostella, Rhigognostis sp.","1","1","1","111","0","0","99.78","450","0","450","32.22","0"
"AGAW1","09N","INS","BOLD:AAO3939","Arthropoda","Insecta","Diptera","Limoniidae","Limoniinae","Dicranomyia","","1","1","1","111","0","0","98.67","450","0","450","36.67","0"
"AGAW1","09N","INS","BOLD:ACE4701","Arthropoda","Insecta","Lepidoptera","Crambidae","Crambinae","Crambus","Crambus multilinellus","1","1","1","111","0","0","97.55","449","0","450","32.44","0"
"AGAW1","09N","NAR","no match","no match","no match","no match","no match","no match","no match","no match","no match","no match","no match","no match","no match","no match","no match","no match","no match","no match","no match","no match"
"AGAW1","09N","NIN","BOLD:AAA7242","Arthropoda","Collembola","Entomobryomorpha","Entomobryidae","Seirinae, Entomobryinae","Entomobrya, Lepidosira","Entomobrya multifasciata, Lepidosira sp.","1613","1518","1518","1","5.89","0","99.07","420.3","0","424.24","39.33","0"
"AGAW1","09N","NIN","BOLD:AAC0175","Arthropoda","Arachnida","Araneae","Theridiidae","","Parasteatoda","Parasteatoda tepidariorum, Parasteatoda simulans","67","63","63","2","5.97","0","99.26","428.1","0","430.3","32.59","0"
"AGAW1","09N","NIN","BOLD:AAB7914","Arthropoda","Collembola","Symphypleona","Bourletiellidae","","Bourletiella","","28","27","27","3","3.57","0","99.44","443.89","0","445","38.12","0"
"AGAW1","09N","NIN","BOLD:ACN3178","Arthropoda","Collembola","Entomobryomorpha","Entomobryidae","","","","7","7","7","4","0","0","97.86","433.29","0","434.86","39.9","0"
"AGAW1","09N","NIN","BOLD:ACC0359","Arthropoda","Collembola","Symphypleona","Bourletiellidae","","Bourletiella","","7","7","7","4","0","0","98.95","447","0","450","37.68","0"
"AGAW1","09N","NIN","BOLD:AAH6630","Arthropoda","Collembola","Entomobryomorpha","Entomobryidae","Entomobryinae","Entomobrya","Entomobrya unostrigata","3","3","3","6","0","0","99.31","421","0","421.67","39.02","0"
"AGAW1","09N","NIN","BOLD:AAI4346","Arthropoda","Arachnida","Opiliones","Phalangiidae","","Paroligolophus, Phalangium","Paroligolophus agrestis, Phalangium opilio","3","3","3","6","0","0","99.16","442.33","0","445.33","36.74","0"
"AGAW1","09N","NIN","BOLD:ACP9512","Arthropoda","Collembola","Entomobryomorpha","Entomobryidae","","","","3","3","3","6","0","0","98.08","416.67","0","429","39.16","0"
"AGAW1","09N","NIN","BOLD:ACI9908","Arthropoda","Collembola","Symphypleona","Bourletiellidae","","","","1","1","1","9","0","0","99.78","450","0","450","35.56","0"
"AGAW1","09N","NIN","BOLD:ACX8969","Arthropoda","Arachnida","Trombidiformes","Eupodidae","","","","1","1","1","9","0","0","98.6","430","0","433","28.18","0"
"AGAW1","09N","NIN","BOLD:AAM3234","Arthropoda","Collembola","Entomobryomorpha","Entomobryidae","Seirinae","Lepidosira","","1","1","1","9","0","0","98.66","449","0","450","39.11","0"
"AGAW1","09S","BAC","TAX:91391","Proteobacteria","Alphaproteobacteria","Rickettsiales","",NA,"","","51","38","38","1","25.49","0","98.55","412.02","0","438.47","40.08","0"
"AGAW1","09S","BAC","TAX:175768","Proteobacteria","Alphaproteobacteria","Rickettsiales","Rickettsiaceae",NA,"Wolbachia","Wolbachia pipientis","15","15","15","2","0","0","98.21","440.53","0","442.93","39.45","0"
"AGAW1","09S","BAC","TAX:91393","Proteobacteria","Alphaproteobacteria","Rickettsiales","Rickettsiaceae",NA,"Wolbachia","","4","4","4","3","0","0","99.11","391.5","0","413","39.21","0"
"AGAW1","09S","CHO","no match","no match","no match","no match","no match","no match","no match","no match","no match","no match","no match","no match","no match","no match","no match","no match","no match","no match","no match","no match"
"AGAW1","09S","INS","BOLD:AAP8739","Arthropoda","Insecta","Diptera","Platystomatidae","Platystomatinae","Rivellia","Rivellia steyskali","25162","15454","15454","1","38.58","0","99.3","441.66","0","443.4","37.96","0"
"AGAW1","09S","INS","BOLD:AAA3453","Arthropoda","Insecta","Diptera","Anthomyiidae","Anthomyiinae","Delia","","19284","15866","15866","2","17.72","0","99.39","432.62","0","435.23","34.66","0"
"AGAW1","09S","INS","BOLD:AAI4303","Arthropoda","Insecta","Diptera","Chironomidae","Chironominae","Chironomus","Chironomus melanescens","5163","4585","4585","3","11.2","0","98.41","429.75","0","434.1","33.95","0"
"AGAW1","09S","INS","BOLD:AAG2511","Arthropoda","Insecta","Diptera","Anthomyiidae","Anthomyiinae","Delia","Delia platura","4545","3866","3866","4","14.94","0","98.93","444.82","0","447.64","33.23","0"
"AGAW1","09S","INS","BOLD:AAG2123","Arthropoda","Insecta","Diptera","Tachinidae","Dexiinae","Dinera","Dinera grisescens","3464","2544","2544","5","26.56","0","99.3","434.63","0","436.42","31.89","0"
"AGAW1","09S","INS","BOLD:ABX6186","Arthropoda","Insecta","Diptera","Tipulidae","Tipulinae","Nephrotoma","Nephrotoma ferruginea","2477","2094","2094","6","15.46","0","99.37","430.51","0","432.65","34.25","0"
"AGAW1","09S","INS","BOLD:AAC0903","Arthropoda","Insecta","Diptera","Chironomidae","Chironominae","Chironomus","Chironomus acidophilus","2365","1834","1834","7","22.45","0","99.42","438.95","0","441.46","36.31","0"
"AGAW1","09S","INS","BOLD:AAA7374","Arthropoda","Insecta","Diptera","Syrphidae","Syrphinae","Sphaerophoria","Sphaerophoria bankowskae, Sphaerophoria infuscata, Sphaerophoria nr. philanthus, Sphaerophoria batava, Sphaerophoria sp., Sphaerophoria rueppellii, Sphaerophoria interrupta-group, Sphaerophoria sp. MA-1, Sphaerophoria cleoae, Sphaerophoria asymmetrica, Sphaerophoria fatarum, Sphaerophoria bengalensis, Sphaerophoria bifurcata, Sphaerophoria taeniata, Sphaerophoria interrupta, Sphaerophoria contigua, Sphaerophoria kaa, Sphaerophoria laurae, Sphaerophoria chongjini, Sphaerophoria boreoalpina, Sphaerophoria sp. BOLDAAA7374, Sphaerophoria macrogaster, Sphaerophoria sulphuripes, Sphaerophoria brevipilosa, Sphaerophoria philanthus, Sphaerophoria pyrrhina, Sphaerophoria abbreviata, Sphaerophoria scripta, Sphaerophoria potentillae, Sphaerophoria longipilosa, Sphaerophoria virgata","1824","1595","1595","8","12.55","0","99.33","428.75","0","432.13","30.72","0"
"AGAW1","09S","INS","BOLD:AAA5299","Arthropoda","Insecta","Diptera","Chironomidae","Orthocladiinae","Cricotopus","Cricotopus laetus, Cricotopus relucens, Cricotopus trifasciatus, Cricotopus sp., Cricotopus sp. 3ES, Cricotopus sylvestris, Cricotopus glacialis","1664","1491","1491","9","10.4","0","99.19","426.16","0","430.39","34.58","0"
"AGAW1","09S","INS","BOLD:AAG1711","Arthropoda","Insecta","Diptera","Muscidae","Phaoniinae","Helina","Helina depuncta","1515","1312","1312","10","13.4","0","99.29","433.17","0","435.39","31.38","0"
"AGAW1","09S","INS","BOLD:ACE7444","Arthropoda","Insecta","Hemiptera","Miridae","Mirinae, Phylinae","Adelphocoris, Megalocoleus","Adelphocoris lineolatus, Megalocoleus molliculus","1461","940","940","11","35.66","0","99.62","442.12","0","442.75","34.83","0"
"AGAW1","09S","INS","BOLD:AAN6154","Arthropoda","Insecta","Coleoptera","Latridiidae","Corticariinae","Melanophthalma","Melanophthalma inermis","1205","985","985","12","18.26","0","98.95","436.92","0","438.71","31.44","0"
"AGAW1","09S","INS","BOLD:AAG5499","Arthropoda","Insecta","Diptera","Chironomidae","Orthocladiinae","Psectrocladius","Psectrocladius obvius, Psectrocladius platypus","1009","904","904","13","10.41","0","97.86","424.21","0","427.21","36.41","0"
"AGAW1","09S","INS","BOLD:AAG2868","Arthropoda","Insecta","Hemiptera","Cicadellidae","Typhlocybinae","Empoasca","Empoasca sp., Empoasca fabae","993","638","638","14","35.75","0","99.62","441.59","0","448.12","32.42","0"
"AGAW1","09S","INS","BOLD:ACO8630","","","","","","","","679","476","476","15","29.9","0","98.23","443","0","443.99","34.09","0"
"AGAW1","09S","INS","BOLD:AAU6681","Arthropoda","Insecta","Diptera","Tipulidae","Tipulinae","Nephrotoma","Nephrotoma cornicina","645","538","538","16","16.59","0","99.51","443.02","0","444.18","36.32","0"
"AGAW1","09S","INS","BOLD:AAB0373","Arthropoda","Insecta","Neuroptera","Chrysopidae","Chrysopinae","Chrysoperla","Chrysoperla sp. isolate 82, Chrysoperla zastrowi, Chrysoperla plorabunda, Chrysoperla sp. isolate 71, Chrysoperla agilis, Italochrysa italica, Chrysoperla sp. isolate 88, Chrysoperla lucasina, Chrysoperla carnea, Chrysoperla mediterranea, Chrysoperla pallida, Chrysoperla nipponensis, Chrysoperla sillemi","630","503","503","17","20.16","0","99.42","436","0","436.86","31.37","0"
"AGAW1","09S","INS","BOLD:AAD0484","Arthropoda","Insecta","Diptera","Chironomidae","","","","556","473","473","18","14.93","0","99.05","444.54","0","446.13","34.66","0"
"AGAW1","09S","INS","BOLD:ACK3469","Arthropoda","Insecta","Coleoptera","Chrysomelidae","Eumolpinae","Brachypnoea","Brachypnoea puncticollis","544","390","390","19","28.31","0","99.46","442.96","0","444.22","36.34","0"
"AGAW1","09S","INS","BOLD:AAA5803","Arthropoda","Insecta","Hemiptera","Miridae","Mirinae","Lygus","Lygus lineolaris","415","318","318","20","23.37","0","98.58","446.63","0","448.31","33.93","0"
"AGAW1","09S","INS","BOLD:AAH4135","Arthropoda","Insecta","Diptera","Chloropidae","Chloropinae","Thaumatomyia","Thaumatomyia glabra, Thaumatomyia grata, Thaumatomyia sp. 1 TAW","379","352","352","21","7.12","0","98.56","435.09","0","439.05","32.19","0"
"AGAW1","09S","INS","BOLD:ACD3384","Arthropoda","Insecta","Hemiptera","Miridae","Orthotylinae","Orthotylus","Orthotylus robiniae","337","262","262","22","22.26","0","99.59","434.28","0","435.2","41.43","0"
"AGAW1","09S","INS","BOLD:AAH3664","Arthropoda","Insecta","Diptera","Cecidomyiidae","","","CecidInt3 sp. BOLD:AAH3664","313","303","303","23","3.19","0","99.1","432.95","0","435.66","27.18","0"
"AGAW1","09S","INS","BOLD:AAG9626","Arthropoda","Insecta","Diptera","Dolichopodidae","","","","283","237","237","24","16.25","0","97.89","431.71","0","433.33","36.24","0"
"AGAW1","09S","INS","BOLD:ACK7314","Arthropoda","Insecta","Diptera","Chironomidae","Chironominae","Polypedilum","","278","277","277","25","0.36","0","98.57","385.26","0","390","35.08","0"
"AGAW1","09S","INS","BOLD:AAP5169","Arthropoda","Insecta","Diptera","Chloropidae","Oscinellinae","Elachiptera","Elachiptera nigriceps","270","221","221","26","18.15","0","99.16","435.84","0","436.95","31.78","0"
"AGAW1","09S","INS","BOLD:AAN5342","Arthropoda","Insecta","Diptera","Chironomidae","Orthocladiinae","Heleniella","","269","260","260","27","3.35","0","99.06","421.66","0","428.41","35.11","0"
"AGAW1","09S","INS","BOLD:AAH3661","Arthropoda","Insecta","Diptera","Cecidomyiidae","","","","256","229","229","28","10.55","0","99.28","432.77","0","434.57","25.44","0"
"AGAW1","09S","INS","BOLD:ACK3431","Arthropoda","Insecta","Diptera","Cecidomyiidae","","","","249","224","224","29","10.04","0","99.22","428.39","0","430.25","32.44","0"
"AGAW1","09S","INS","BOLD:AAN5199","Arthropoda","Insecta","Diptera","Cecidomyiidae","","","CecidInt20 sp. BOLD:AAN5199","231","194","194","30","16.02","0","99.55","434.33","0","445.37","26.82","0"
"AGAW1","09S","INS","BOLD:AAP6790","Arthropoda","Insecta","Diptera","Chloropidae","Oscinellinae","Incertella","Incertella incerta","222","197","197","31","11.26","0","99.29","440.23","0","441.98","31.47","0"
"AGAW1","09S","INS","BOLD:AAJ9681","Arthropoda","Insecta","Diptera","Agromyzidae","Phytomyzinae","Aulagromyza","Aulagromyza luteoscutellata","219","198","198","32","9.59","0","99.37","441.45","0","443.72","34.06","0"
"AGAW1","09S","INS","BOLD:AAG5443","Arthropoda","Insecta","Diptera","Chironomidae","Orthocladiinae","","","207","206","206","33","0.48","0","98.78","412.12","0","414.21","33.43","0"
"AGAW1","09S","INS","BOLD:AAG1517","Arthropoda","Insecta","Diptera","Chloropidae","Oscinellinae","Gaurax","Gaurax pallidipes","206","203","203","34","1.46","0","98.91","418.73","0","420.76","31.88","0"
"AGAW1","09S","INS","BOLD:AAG5507","Arthropoda","Insecta","Diptera","Chironomidae","Chironominae","Chironomus","Chironomus atrella","204","202","202","35","0.98","0","98.71","422.81","0","429.88","35.68","0"
"AGAW1","09S","INS","BOLD:AAC4197","Arthropoda","Insecta","Diptera","Chironomidae","Orthocladiinae","Paraphaenocladius","Paraphaenocladius impensus","204","175","175","35","14.22","0","99.09","432.95","0","437.83","34.82","0"
"AGAW1","09S","INS","BOLD:AAP5087","Arthropoda","Insecta","Diptera","Ceratopogonidae","Dasyheleinae","Dasyhelea","","173","148","148","37","14.45","0","98.84","435.16","0","436.66","34.83","0"
"AGAW1","09S","INS","BOLD:AAB3380","Arthropoda","Insecta","Hemiptera","Miridae","Phylinae","Plagiognathus","Plagiognathus fuscosus, Plagiognathus politus, Plagiognathus obscurus, Plagiognathus blatchleyi","155","126","126","38","18.71","0","99.32","434.08","0","435.14","37.89","0"
"AGAW1","09S","INS","BOLD:ABV9018","Arthropoda","Insecta","Diptera","Mycetophilidae","Sciophilinae","Sciophila","","141","136","136","39","3.55","0","98.65","423.3","0","425.48","31.86","0"
"AGAW1","09S","INS","BOLD:ABY5318","Arthropoda","Insecta","Diptera","Chloropidae","Oscinellinae","Incertella, Rhopalopterum","Rhopalopterum umbrosum, Rhopalopterum luteiceps, Incertella bispina, Rhopalopterum painteri","119","93","93","40","21.85","0","99.64","447.25","0","448.27","32.09","0"
"AGAW1","09S","INS","BOLD:ABU5525","Arthropoda","Insecta","Diptera","Chironomidae","Orthocladiinae","Limnophyes","Limnophyes sp. 14ES","115","115","115","41","0","0","99.09","401.09","0","404.38","37.18","0"
"AGAW1","09S","INS","BOLD:AAA7661","Arthropoda","Insecta","Diptera","Culicidae","Culicinae","Culiseta, Culex","Culiseta melanura, Culex articularis, Culex restuans, Culex acharistus","110","106","106","42","3.64","0","99.35","438.01","0","440.16","32.79","0"
"AGAW1","09S","INS","BOLD:AAD2041","Arthropoda","Insecta","Diptera","Dolichopodidae","Sciapodinae","Condylostylus","Condylostylus inermis, Condylostylus nigrofemoratus, Condylostylus connectans, Condylostylus caudatus, Condylostylus flavipes","100","79","79","43","21","0","99.52","433.16","0","434.22","35.94","0"
"AGAW1","09S","INS","BOLD:ABZ0908","Arthropoda","Insecta","Diptera","Tipulidae","Tipulinae","Nephrotoma","","97","97","97","44","0","0","97.46","367.41","0","370.93","34.05","0"
"AGAW1","09S","INS","BOLD:ADF6661","Arthropoda","Insecta","Diptera","Cecidomyiidae","","","","82","67","67","45","18.29","0","98.91","415.26","0","440.88","29.85","0"
"AGAW1","09S","INS","BOLD:AAG9897","Arthropoda","Insecta","Dermaptera","Forficulidae","Forficulinae","Forficula","Forficula auricularia-A","79","79","79","46","0","0","98.82","394.38","0","404.16","45.65","0"
"AGAW1","09S","INS","BOLD:AAM9255","Arthropoda","Insecta","Diptera","Sciaridae","","Pseudolycoriella","","79","79","79","46","0","0","98.87","414.53","0","417.15","30.87","0"
"AGAW1","09S","INS","BOLD:AAN5220","Arthropoda","Insecta","Diptera","Cecidomyiidae","","","","72","72","72","48","0","0","97.99","418.76","0","427.36","26.04","0"
"AGAW1","09S","INS","BOLD:ABA0863","Arthropoda","Insecta","Diptera","Cecidomyiidae","","","","71","62","62","49","12.68","0","99.4","441.28","0","442.44","29.73","0"
"AGAW1","09S","INS","BOLD:AAN5267","Arthropoda","Insecta","Diptera","Cecidomyiidae","","","","66","50","50","50","24.24","0","99.57","447.03","0","447.74","30.59","0"
"AGAW1","09S","INS","BOLD:ACY6524","Arthropoda","Insecta","Diptera","Chironomidae","Orthocladiinae","Psectrocladius","Psectrocladius cf. limbatellus","64","61","61","51","4.69","0","99.11","434.72","0","444.78","35.9","0"
"AGAW1","09S","INS","BOLD:AAP6703","Arthropoda","Insecta","Diptera","Ceratopogonidae","Dasyheleinae","Dasyhelea","","60","56","56","52","6.67","0","99.35","439.92","0","442.3","40.4","0"
"AGAW1","09S","INS","BOLD:AAH4171","Arthropoda","Insecta","Diptera","Chloropidae","Oscinellinae","Apallates","Apallates montanus, Apallates particeps","56","51","51","53","8.93","0","99","438.29","0","440.11","32.7","0"
"AGAW1","09S","INS","BOLD:AAJ9671","Arthropoda","Insecta","Diptera","Agromyzidae","Phytomyzinae","Aulagromyza","Aulagromyza tridentata","51","49","49","54","3.92","0","99.12","440.94","0","443.76","32.99","0"
"AGAW1","09S","INS","BOLD:AAG7891","Arthropoda","Insecta","Hymenoptera","Platygastridae","Scelioninae","Telenomus","Telenomus podisi","50","47","47","55","6","0","98.85","427.06","0","435.66","27.06","0"
"AGAW1","09S","INS","BOLD:AAB7072","Arthropoda","Insecta","Lepidoptera","Coleophoridae","Coleophorinae","Coleophora","Coleophora coxi, Coleophora texanella","49","47","47","56","4.08","0","99.09","430.94","0","432.33","33.03","0"
"AGAW1","09S","INS","BOLD:AAN5205","Arthropoda","Insecta","Diptera","Cecidomyiidae","","","CecidInt42 sp. BOLD:AAN5205","48","42","42","57","12.5","0","99.59","448.19","0","449.35","31.4","0"
"AGAW1","09S","INS","BOLD:AAG1004","Arthropoda","Insecta","Diptera","Chironomidae","Orthocladiinae","Cricotopus","Cricotopus trifascia, Cricotopus trifascia grp.","48","41","41","57","14.58","0","99.24","436.5","0","438.46","37.04","0"
"AGAW1","09S","INS","BOLD:ACC7718","Arthropoda","Insecta","Diptera","Chironomidae","Orthocladiinae","Psectrocladius","","47","42","42","59","10.64","0","99.19","415.38","0","428.4","37.06","0"
"AGAW1","09S","INS","BOLD:AAN5230","Arthropoda","Insecta","Diptera","Cecidomyiidae","","","","45","45","45","60","0","0","99.15","425.44","0","427.11","25.96","0"
"AGAW1","09S","INS","BOLD:ABA7871","Arthropoda","Insecta","Diptera","Cecidomyiidae","","","","44","40","40","61","9.09","0","99.11","445.2","0","446.36","26.56","0"
"AGAW1","09S","INS","BOLD:AAD1485","Arthropoda","Insecta","Diptera","Chironomidae","Chironominae","Paratanytarsus","Paratanytarsus grimmii, Paratanytarsus sp.","42","38","38","62","9.52","0","99.52","439.57","0","441.31","33.87","0"
"AGAW1","09S","INS","BOLD:AAN5225","Arthropoda","Insecta","Diptera","Cecidomyiidae","","","","37","32","32","63","13.51","0","99.1","443.54","0","444.86","28.31","0"
"AGAW1","09S","INS","BOLD:AAG0996","Arthropoda","Insecta","Diptera","Chironomidae","Orthocladiinae","Cricotopus","Cricotopus sp. 18ES, Cricotopus bicinctus gr., Cricotopus bicinctus grp.","37","37","37","63","0","0","98.57","408.86","0","411.68","32.53","0"
"AGAW1","09S","INS","BOLD:AAN5238","Arthropoda","Insecta","Diptera","Cecidomyiidae","","","","33","26","26","65","21.21","0","99.81","442.06","0","443.15","25.78","0"
"AGAW1","09S","INS","BOLD:AAG2898","Arthropoda","Insecta","Hemiptera","Cicadellidae","Deltocephalinae","Paraphlepsius","Paraphlepsius apertus, Paraphlepsius irroratus","33","30","30","65","9.09","0","99.42","448.76","0","449.39","32.26","0"
"AGAW1","09S","INS","BOLD:AAB4657","Arthropoda","Insecta","Diptera","Chironomidae","Chironominae","Chironomus","Chironomus maturus","29","28","28","67","3.45","0","99.4","441.17","0","444.34","37.49","0"
"AGAW1","09S","INS","BOLD:AAC4403","Arthropoda","Insecta","Diptera","Chironomidae","Chironominae","Micropsectra","Micropsectra polita","27","27","27","68","0","0","98.44","436.26","0","438.07","34.69","0"
"AGAW1","09S","INS","BOLD:ABA0856","Arthropoda","Insecta","Diptera","Cecidomyiidae","","","","26","21","21","69","19.23","0","98.9","449.69","0","450","29.09","0"
"AGAW1","09S","INS","BOLD:AAH0134","Arthropoda","Insecta","Coleoptera","Phalacridae","Phalacrinae","Stilbus","Stilbus apicialis","26","23","23","69","11.54","0","99.38","431.62","0","432.38","32.01","0"
"AGAW1","09S","INS","BOLD:ABV0255","Arthropoda","Insecta","Diptera","Chironomidae","Orthocladiinae","Limnophyes","","26","24","24","69","7.69","0","99.27","435.96","0","436.92","34.7","0"
"AGAW1","09S","INS","BOLD:ABV9319","Arthropoda","Insecta","Diptera","Chironomidae","Tanypodinae","Procladius","","22","22","22","72","0","0","99.01","432.18","0","433.64","34.4","0"
"AGAW1","09S","INS","BOLD:AAH0135","Arthropoda","Insecta","Coleoptera","Phalacridae","Phalacrinae","Acylomus","Acylomus pugetanus, Acylomus ergoti","21","21","21","73","0","0","98.94","433.52","0","435.67","31.66","0"
"AGAW1","09S","INS","BOLD:AAG6934","Arthropoda","Insecta","Diptera","Hybotidae","","","","18","18","18","74","0","0","99.14","408.89","0","411.28","38.26","0"
"AGAW1","09S","INS","BOLD:AAN5242","Arthropoda","Insecta","Diptera","Cecidomyiidae","","","","17","17","17","75","0","0","98.74","426.06","0","427.76","28.56","0"
"AGAW1","09S","INS","BOLD:AAM6037","Arthropoda","Insecta","Diptera","Cecidomyiidae","Cecidomyiinae","Contarinia, Dasineura","","17","15","15","75","11.76","0","98.72","440.94","0","442.82","26.36","0"
"AGAW1","09S","INS","BOLD:ACF5830","Arthropoda","Insecta","Diptera","Chloropidae","Chloropinae","Thaumatomyia","","17","17","17","75","0","0","98.12","394.18","0","397.35","31.81","0"
"AGAW1","09S","INS","BOLD:AAB4894","Arthropoda","Insecta","Hemiptera","Aphididae","Aphidinae","Sitobion","Sitobion smilacifoliae, Sitobion avenae, Sitobion phyllanthi","16","16","16","78","0","0","97.75","447.44","0","447.81","25.31","0"
"AGAW1","09S","INS","BOLD:AAN5260","Arthropoda","Insecta","Diptera","Cecidomyiidae","","","","16","16","16","78","0","0","99.19","414.94","0","416.19","25.92","0"
"AGAW1","09S","INS","BOLD:AAH2876","Arthropoda","Insecta","Diptera","Cecidomyiidae","","","CecidInt51 sp. BOLD:AAH2876","16","15","15","78","6.25","0","99.06","437.94","0","439.44","26.79","0"
"AGAW1","09S","INS","BOLD:ADH4973","Arthropoda","Insecta","Diptera","Tipulidae","Tipulinae","Nephrotoma","","15","15","15","81","0","0","97.74","379","0","382.4","33.85","0"
"AGAW1","09S","INS","BOLD:ABV1297","Arthropoda","Insecta","Diptera","Sciaridae","","","","15","15","15","81","0","0","99.38","433.2","0","435.53","30.44","0"
"AGAW1","09S","INS","BOLD:ABA0772","Arthropoda","Insecta","Diptera","Chironomidae","Chironominae","Polypedilum","","15","15","15","81","0","0","98.99","429.67","0","431.2","33.36","0"
"AGAW1","09S","INS","BOLD:AAU8441","Arthropoda","Insecta","Hymenoptera","Ichneumonidae","Campopleginae","Meloboris","","14","9","9","84","35.71","0","99.89","449.29","0","450","25.59","0"
"AGAW1","09S","INS","BOLD:ABV1354","Arthropoda","Insecta","Diptera","Ceratopogonidae","Dasyheleinae","Dasyhelea","","14","11","11","84","21.43","0","98.91","439.5","0","440.79","35.89","0"
"AGAW1","09S","INS","BOLD:ABV1452","Arthropoda","Insecta","Diptera","Chironomidae","Chironominae","","","13","10","10","86","23.08","0","98.05","448.77","0","450","35.69","0"
"AGAW1","09S","INS","BOLD:AAF9771","Arthropoda","Insecta","Diptera","Hybotidae","Tachydromiinae","Platypalpus","Platypalpus longiseta, Platypalpus trivialis, Platypalpus sp. 5, Platypalpus sp. 17, Platypalpus pallidiventris","11","11","11","87","0","0","99.33","445.82","0","446","35.3","0"
"AGAW1","09S","INS","BOLD:AAV5400","Arthropoda","Insecta","Diptera","Cecidomyiidae","","","","10","10","10","88","0","0","97.85","436.4","0","437.5","29.32","0"
"AGAW1","09S","INS","BOLD:AAN5249","Arthropoda","Insecta","Diptera","Cecidomyiidae","","","","9","9","9","89","0","0","99.07","413.67","0","415.44","33.01","0"
"AGAW1","09S","INS","BOLD:AAM6257","Arthropoda","Insecta","Diptera","Chironomidae","Orthocladiinae","","","7","7","7","90","0","0","98.48","387.86","0","394.86","35.13","0"
"AGAW1","09S","INS","BOLD:ABV0451","Arthropoda","Insecta","Diptera","Cecidomyiidae","","","","7","5","5","90","28.57","0","97.68","449.71","0","450","28.67","0"
"AGAW1","09S","INS","BOLD:AAP8816","Arthropoda","Insecta","Diptera","Cecidomyiidae","Cecidomyiinae","Sitodiplosis","Sitodiplosis sp. BOLD:AAP8816","7","7","7","90","0","0","99.48","443.14","0","445.71","26.38","0"
"AGAW1","09S","INS","BOLD:AAG6745","Arthropoda","Insecta","Diptera","Calliphoridae","Calliphorinae","Pollenia","Pollenia pediculata","7","7","7","90","0","0","99.53","428.14","0","429.57","33.3","0"
"AGAW1","09S","INS","BOLD:AAB8911","Arthropoda","Insecta","Coleoptera","Curculionidae","Entiminae","Polydrusus","Polydrusus formosus","7","6","6","90","14.29","0","97.3","449.43","0","450","34.63","0"
"AGAW1","09S","INS","BOLD:ACG4376","Arthropoda","Insecta","Diptera","Chironomidae","Chironominae","Tanytarsus","","7","5","5","90","28.57","0","97.17","438.57","0","439.71","37.02","0"
"AGAW1","09S","INS","BOLD:ADH9867","Arthropoda","Insecta","Diptera","","","","","6","6","6","96","0","0","98.76","418","0","421.5","37.45","0"
"AGAW1","09S","INS","BOLD:AAZ0271","Arthropoda","Insecta","Diptera","Ceratopogonidae","Ceratopogoninae","Culicoides","","6","6","6","96","0","0","98.36","441.33","0","444","41.43","0"
"AGAW1","09S","INS","BOLD:AAN5255","Arthropoda","Insecta","Diptera","Cecidomyiidae","Cecidomyiinae","Dasineura","","5","5","5","98","0","0","98.74","439.6","0","450","26.98","0"
"AGAW1","09S","INS","BOLD:AAM7855","Arthropoda","Insecta","Thysanoptera","Thripidae","Thripinae","Frankliniella","Frankliniella tritici","5","4","4","98","20","0","99.91","438","0","441.2","31.18","0"
"AGAW1","09S","INS","BOLD:AAN5183","Arthropoda","Insecta","Diptera","Cecidomyiidae","","","","5","5","5","98","0","0","98.43","447.4","0","449.2","28.36","0"
"AGAW1","09S","INS","BOLD:AAA4781","Arthropoda","Insecta","Hymenoptera","Braconidae","Microgastrinae","Glyptapanteles","Glyptapanteles carlossarmientoi","5","5","5","98","0","0","99.62","428.2","0","429.8","23.75","0"
"AGAW1","09S","INS","BOLD:ABY9688","Arthropoda","Insecta","Diptera","Chloropidae","Chloropinae","Thaumatomyia","Thaumatomyia glabra","4","4","4","102","0","0","97.57","427.75","0","431.25","32.14","0"
"AGAW1","09S","INS","BOLD:AAA3889","Arthropoda","Insecta","Lepidoptera","Crambidae","Crambinae","Parapediasia","Parapediasia teterrellus","4","4","4","102","0","0","99.48","435","0","435.75","32.29","0"
"AGAW1","09S","INS","BOLD:AAN5200","Arthropoda","Insecta","Diptera","Cecidomyiidae","","","","4","4","4","102","0","0","98.3","424","0","426.25","26.97","0"
"AGAW1","09S","INS","BOLD:AAF9187","Arthropoda","Insecta","Coleoptera","Curculionidae","Entiminae","Phyllobius","Phyllobius oblongus","4","4","4","102","0","0","99.89","449","0","450","38.89","0"
"AGAW1","09S","INS","BOLD:AAH3947","Arthropoda","Insecta","Diptera","Sciaridae","","Bradysia","Bradysia trivittata","4","4","4","102","0","0","99.16","389.25","0","390","33.54","0"
"AGAW1","09S","INS","BOLD:ACV2677","Arthropoda","Insecta","Diptera","Cecidomyiidae","","","","4","4","4","102","0","0","97.44","448.5","0","450","30.22","0"
"AGAW1","09S","INS","BOLD:AAH0152","Arthropoda","Insecta","Coleoptera","Cantharidae","Cantharinae","Cantharis","Cantharis rufa","4","4","4","102","0","0","97.97","433","0","438.25","35.29","0"
"AGAW1","09S","INS","BOLD:AAG2740","Arthropoda","Insecta","Diptera","Ephydridae","Parydrinae","Philygria","Philygria debilis","4","4","4","102","0","0","97.94","393.25","0","394.5","32.59","0"
"AGAW1","09S","INS","BOLD:AAA6292","Arthropoda","Insecta","Lepidoptera","Tortricidae","Tortricinae","Cnephasia","Cnephasia genitalana, Cnephasia stephensiana","3","3","3","110","0","0","100","446.67","0","447.67","35.59","0"
"AGAW1","09S","INS","BOLD:AAY6412","Arthropoda","Insecta","Diptera","Sciaridae","","Bradysia","Bradysia urticae","3","3","3","110","0","0","99.47","441.67","0","446.33","32.26","0"
"AGAW1","09S","INS","BOLD:ABX4079","Arthropoda","Insecta","Diptera","Cecidomyiidae","","","","3","3","3","110","0","0","97.92","449","0","450","28.44","0"
"AGAW1","09S","INS","BOLD:ACR3349","Arthropoda","Insecta","Diptera","Agromyzidae","Phytomyzinae","Aulagromyza","Aulagromyza sp. BOLD:ACR3349","3","3","3","110","0","0","97.15","445.33","0","447.33","33.68","0"
"AGAW1","09S","INS","BOLD:ACF1257","","","","","","","","3","2","2","110","33.33","0","99.41","450","0","450","34.67","0"
"AGAW1","09S","INS","BOLD:AAA3867","Arthropoda","Insecta","Lepidoptera","Noctuidae","Noctuinae","Apamea","Apamea amputatrix","3","3","3","110","0","0","98.21","446","0","450","33.33","0"
"AGAW1","09S","INS","BOLD:AAA3654","Arthropoda","Insecta","Lepidoptera","Crambidae","Crambinae","Crambus","Crambus bigelovi, Crambus johnsoni, Crambus praefectellus","3","3","3","110","0","0","99.26","448.33","0","450","31.63","0"
"AGAW1","09S","INS","BOLD:AAG7276","Arthropoda","Insecta","Diptera","Sphaeroceridae","Limosininae","Leptocera","Leptocera erythrocera","2","2","2","117","0","0","99.51","413.5","0","415","33.11","0"
"AGAW1","09S","INS","BOLD:AAA6560","Arthropoda","Insecta","Lepidoptera","Crambidae","Crambinae","Crambus","Crambus multilinellus, Crambus saltuellus","2","2","2","117","0","0","99.54","424.5","0","425","30.54","0"
"AGAW1","09S","INS","BOLD:AAE7186","Arthropoda","Insecta","Hymenoptera","Braconidae","Aphidiinae","Praon","Praon sp. 1 MM-2013, Praon volucre, Praon bicolor","2","2","2","117","0","0","99.87","418","0","420","25.45","0"
"AGAW1","09S","INS","BOLD:AAA4277","Arthropoda","Insecta","Diptera","Syrphidae","Syrphinae","Toxomerus","Toxomerus marginatus, Toxomerus sp. CR-3, Toxomerus CR-3","2","2","2","117","0","0","99.33","449.5","0","450","31.11","0"
"AGAW1","09S","INS","BOLD:AAH3914","Arthropoda","Insecta","Diptera","Sciaridae","","Sciara","Sciara hamatilis, Sciara humeralis","2","2","2","117","0","0","98.09","445","0","450","30.67","0"
"AGAW1","09S","INS","BOLD:AAL7819","Arthropoda","Insecta","Diptera","Psychodidae","Psychodinae","Psychoda","Psychoda sp., Psychoda sp. BOLD:AAL7819","2","2","2","117","0","0","99.33","405","0","406.5","32.56","0"
"AGAW1","09S","INS","BOLD:AAF1013","Arthropoda","Insecta","Diptera","Chironomidae","Chironominae","Tanytarsus","Tanytarsus allicis","2","2","2","117","0","0","97.2","392","0","393.5","36.38","0"
"AGAW1","09S","INS","BOLD:ACL2355","Arthropoda","Insecta","Diptera","Tipulidae","Tipulinae","Nephrotoma","","2","2","2","117","0","0","97.57","375.5","0","376","33.41","0"
"AGAW1","09S","INS","BOLD:ACI3271","Arthropoda","Insecta","Diptera","Cecidomyiidae","","","","2","2","2","117","0","0","98.76","364","0","391.5","26.71","0"
"AGAW1","09S","INS","BOLD:AAY8918","Arthropoda","Insecta","Hemiptera","Cicadellidae","Deltocephalinae","Deltocephalus","Deltocephalus pulicaris","2","2","2","117","0","0","98.45","425","0","430.5","36.31","0"
"AGAW1","09S","INS","BOLD:ACF4925","Arthropoda","Insecta","Hemiptera","Miridae","Mirinae","Adelphocoris","Adelphocoris detritus","2","2","2","117","0","0","97.55","449","0","450","36.44","0"
"AGAW1","09S","INS","BOLD:ACU6925","Arthropoda","Insecta","Diptera","Cecidomyiidae","","","","1","1","1","128","0","0","99.78","450","0","450","28.44","0"
"AGAW1","09S","INS","BOLD:ACK5742","Arthropoda","Insecta","Diptera","Cecidomyiidae","","","","1","1","1","128","0","0","100","450","0","450","29.33","0"
"AGAW1","09S","INS","BOLD:ABW3687","Arthropoda","Insecta","Diptera","Lauxaniidae","Homoneurinae","Homoneura","Homoneura bispina","1","1","1","128","0","0","98","450","0","450","36","0"
"AGAW1","09S","INS","BOLD:AAF8971","Arthropoda","Insecta","Diptera","Limoniidae","","","","1","1","1","128","0","0","98.65","444","0","450","35.56","0"
"AGAW1","09S","INS","BOLD:ABZ1405","Arthropoda","Insecta","Hemiptera","Miridae","Mirinae","Lygus","Lygus abroniae, Lygus unctuosus, Lygus plagiatus, Lygus ceanothi, Lygus potentillae, Lygus humeralis, Lygus keltoni, Lygus rolfsi, Lygus rufidorsus, Lygus hesperus, Lygus rugulipennis, Lygus solidaginis, Lygus atriflavus, Lygus lupini, Lygus atritibialis, Lygus scudderi, Lygus aeratus, Lygus elisus, Lygus rubroclarus, Lygus punctatus, Lygus bradleyi, Lygus borealis, Lygus rubrosignatus, Lygus oregonae, Lygus convexicollis, Lygus robustus, Lygus shulli, Lygus striatus","1","1","1","128","0","0","97.69","390","0","392","33.42","0"
"AGAW1","09S","INS","BOLD:AAB9404","Arthropoda","Insecta","Lepidoptera","Tortricidae","Tortricinae","Archips","Archips rosana, Archips grisea","1","1","1","128","0","0","98.44","448","0","450","31.11","0"
"AGAW1","09S","INS","BOLD:AAY6730","Arthropoda","Insecta","Hemiptera","Derbidae","Cedusinae","Cedusa","Cedusa incisa","1","1","1","128","0","0","98.43","381","0","382","34.82","0"
"AGAW1","09S","INS","BOLD:ACF0916","Arthropoda","Insecta","Diptera","Chironomidae","","","","1","1","1","128","0","0","97.27","403","0","406","32.76","0"
"AGAW1","09S","INS","BOLD:ACF0915","Arthropoda","Insecta","Diptera","Chironomidae","Orthocladiinae","","","1","1","1","128","0","0","97.1","449","0","450","34.67","0"
"AGAW1","09S","INS","BOLD:AAN5497","Arthropoda","Insecta","Diptera","Anthomyiidae","Pegomyinae","Pegomya","Pegomya solennis, Pegomya bicolor","1","1","1","128","0","0","97.33","449","0","450","30.22","0"
"AGAW1","09S","INS","BOLD:AAC9132","Arthropoda","Insecta","Diptera","Culicidae","Culicinae","Culiseta","Culiseta inornata","1","1","1","128","0","0","98.85","349","0","351","32.48","0"
"AGAW1","09S","INS","BOLD:AAG5453","Arthropoda","Insecta","Diptera","Chironomidae","Chironominae","Chironomus","Chironomus bifurcatus","1","1","1","128","0","0","97.32","448","0","450","34.44","0"
"AGAW1","09S","INS","BOLD:ACC8511","Arthropoda","Insecta","Diptera","Cecidomyiidae","Cecidomyiinae","Sitodiplosis","Sitodiplosis mosellana","1","1","1","128","0","0","100","449","0","450","26.67","0"
"AGAW1","09S","INS","BOLD:AAC4491","Arthropoda","Insecta","Diptera","Tipulidae","Tipulinae","Tipula","Tipula morrisoni, Tipula bicornis, Tipula johnsoniana, Tipula bicolor","1","1","1","128","0","0","98.36","426","0","428","34.58","0"
"AGAW1","09S","INS","BOLD:ACM6342","Arthropoda","Insecta","Diptera","Cecidomyiidae","","","","1","1","1","128","0","0","97.22","431","0","433","25.87","0"
"AGAW1","09S","INS","BOLD:AAD1945","Arthropoda","Insecta","Diptera","Agromyzidae","Phytomyzinae","Cerodontha","Cerodontha dorsalis","1","1","1","128","0","0","98.66","448","0","450","32","0"
"AGAW1","09S","INS","BOLD:AAA2815","Arthropoda","Insecta","Lepidoptera","Noctuidae","Noctuinae","Mythimna","Mythimna yukonensis, Mythimna oxygala","1","1","1","128","0","0","99.33","449","0","450","33.33","0"
"AGAW1","09S","INS","BOLD:AAB7030","Arthropoda","Insecta","Diptera","Chironomidae","Chironominae","Chironomus","Chironomus sp. BOLDAAP2996, Chironomus quinnitukqut, Chironomus cf. decorus 2g, Chironomus decorus group sp. 2 IP2013, Chironomus cf. decorus, Chironomus decorus gr.","1","1","1","128","0","0","97.54","366","0","370","33.51","0"
"AGAW1","09S","INS","BOLD:ACC8079","Arthropoda","Insecta","Diptera","Cecidomyiidae","","","","1","1","1","128","0","0","98.89","449","0","450","29.56","0"
"AGAW1","09S","INS","BOLD:AAH3662","Arthropoda","Insecta","Diptera","Cecidomyiidae","","","CecidInt3 sp. BOLD:AAH3662","1","1","1","128","0","0","100","448","0","450","26","0"
"AGAW1","09S","INS","BOLD:ACX8610","Arthropoda","Insecta","Diptera","Cecidomyiidae","","","","1","1","1","128","0","0","99.56","450","0","450","30.67","0"
"AGAW1","09S","INS","BOLD:ACV0176","Arthropoda","Insecta","Diptera","Cecidomyiidae","Cecidomyiinae","Sitodiplosis","","1","1","1","128","0","0","99.11","450","0","450","27.56","0"
"AGAW1","09S","INS","BOLD:AAN5341","Arthropoda","Insecta","Diptera","Chironomidae","Orthocladiinae","Camptocladius","Camptocladius stercorarius","1","1","1","128","0","0","97.62","378","0","380","36.05","0"
"AGAW1","09S","INS","BOLD:ABV9293","Arthropoda","Insecta","Diptera","Chironomidae","Chironominae","Dicrotendipes","","1","1","1","128","0","0","99.15","355","0","401","34.41","0"
"AGAW1","09S","INS","BOLD:AAM6227","Arthropoda","Insecta","Diptera","Chironomidae","Tanypodinae","Procladius","Procladius denticulatus, Procladius sp.","1","1","1","128","0","0","100","431","0","432","31.48","0"
"AGAW1","09S","INS","BOLD:AAA5953","Arthropoda","Insecta","Lepidoptera","Gelechiidae","Gelechiinae","Coleotechnites","Coleotechnites sp., Coleotechnites piceaella, Coleotechnites laricis, Coleotechnites sp. 3, Coleotechnites carbonaria, Coleotechnites granti, Coleotechnites macleodi, Coleotechnites gibsonella, Coleotechnites coniferella, Coleotechnites martini","1","1","1","128","0","0","99.33","449","0","450","34.44","0"
"AGAW1","09S","INS","BOLD:AAB3656","Arthropoda","Insecta","Lepidoptera","Pterophoridae","Pterophorinae","Hellinsia","Hellinsia glenni, Hellinsia lacteodactylus, Hellinsia paleaceus","1","1","1","128","0","0","98.69","383","0","387","31.52","0"
"AGAW1","09S","INS","BOLD:AAB2384","Arthropoda","Insecta","Diptera","Syrphidae","Syrphinae","Eupeodes","Eupeodes duseki, Eupeodes volucris, Eupeodes nielseni, Eupeodes americanus, Eupeodes flukei, Eupeodes vockerothi, Eupeodes lucasi, Eupeodes montivagus, Eupeodes rufipunctatus, Eupeodes rojasi, Eupeodes goeldlini, Eupeodes luniger, Eupeodes punctifer, Eupeodes bucculatus, Eupeodes perplexus, Eupeodes abiskoensis, Eupeodes corollae, Eupeodes pomus, Eupeodes lundbecki, Eupeodes nitens, Eupeodes sp. BOLDAAB2384, Eupeodes curtus, Eupeodes latifasciatus","1","1","1","128","0","0","98.42","380","0","386","30.31","0"
"AGAW1","09S","INS","BOLD:AAC9062","Arthropoda","Insecta","Diptera","Culicidae","Culicinae","Ochlerotatus, Aedes","Aedes fitchii, Ochlerotatus fitchii, Aedes grossbecki, Aedes ventrovittis","1","1","1","128","0","0","97.85","419","0","426","32.39","0"
"AGAW1","09S","INS","BOLD:AAV1136","Arthropoda","Insecta","Diptera","Scatopsidae","","","","1","1","1","128","0","0","99.75","398","0","400","35.5","0"
"AGAW1","09S","INS","BOLD:AAO7694","Arthropoda","Insecta","Diptera","Cecidomyiidae","","","","1","1","1","128","0","0","99.5","399","0","400","25.25","0"
"AGAW1","09S","INS","BOLD:ACE4701","Arthropoda","Insecta","Lepidoptera","Crambidae","Crambinae","Crambus","Crambus multilinellus","1","1","1","128","0","0","100","450","0","450","31.78","0"
"AGAW1","09S","INS","BOLD:AAG2460","Arthropoda","Insecta","Diptera","Anthomyiidae","Anthomyiinae","Hydrophoria","Hydrophoria lancifer","1","1","1","128","0","0","98.64","441","0","444","34.23","0"
"AGAW1","09S","NAR","no match","no match","no match","no match","no match","no match","no match","no match","no match","no match","no match","no match","no match","no match","no match","no match","no match","no match","no match","no match"
"AGAW1","09S","NIN","BOLD:AAA7242","Arthropoda","Collembola","Entomobryomorpha","Entomobryidae","Seirinae, Entomobryinae","Entomobrya, Lepidosira","Entomobrya multifasciata, Lepidosira sp.","414","386","386","1","6.76","0","99.24","424.24","0","427.27","39.36","0"
"AGAW1","09S","NIN","BOLD:AAB7914","Arthropoda","Collembola","Symphypleona","Bourletiellidae","","Bourletiella","","11","11","11","2","0","0","99.39","444","0","448.36","38.14","0"
"AGAW1","09S","NIN","BOLD:ACP2815","Arthropoda","Arachnida","Mesostigmata","Ascidae","","","","6","6","6","3","0","0","99.23","387.83","0","442.33","34.54","0"
"AGAW1","09S","NIN","BOLD:AAH6630","Arthropoda","Collembola","Entomobryomorpha","Entomobryidae","Entomobryinae","Entomobrya","Entomobrya unostrigata","5","5","5","4","0","0","99.33","447.6","0","450","38.8","0"
"AGAW1","09S","NIN","BOLD:ACN3178","Arthropoda","Collembola","Entomobryomorpha","Entomobryidae","","","","1","1","1","5","0","0","98.88","447","0","450","39.56","0"
"AGAW1","09S","NIN","BOLD:ACR0932","Arthropoda","Collembola","Symphypleona","Bourletiellidae","","","","1","1","1","5","0","0","99.47","379","0","441","40.36","0"
"AGAW1","09W","BAC","TAX:175768","Proteobacteria","Alphaproteobacteria","Rickettsiales","Rickettsiaceae",NA,"Wolbachia","Wolbachia pipientis","82","71","71","1","13.41","0","98.44","446.4","0","447.74","39.72","0"
"AGAW1","09W","BAC","TAX:91391","Proteobacteria","Alphaproteobacteria","Rickettsiales","",NA,"","","40","34","34","2","15","0","98.66","409.23","0","445.65","40.23","0"
"AGAW1","09W","BAC","TAX:91393","Proteobacteria","Alphaproteobacteria","Rickettsiales","Rickettsiaceae",NA,"Wolbachia","","28","28","28","3","0","0","99.11","391.61","0","421.39","39.38","0"
"AGAW1","09W","BAC","TAX:703648","Proteobacteria","Alphaproteobacteria","Rickettsiales","Rickettsiaceae",NA,"Wolbachia","Wolbachia endosymbiont","2","2","2","4","0","0","97.47","395","0","450","40","0"
"AGAW1","09W","CHO","no match","no match","no match","no match","no match","no match","no match","no match","no match","no match","no match","no match","no match","no match","no match","no match","no match","no match","no match","no match"
"AGAW1","09W","INS","BOLD:AAG2511","Arthropoda","Insecta","Diptera","Anthomyiidae","Anthomyiinae","Delia","Delia platura","12197","9746","9746","1","20.1","0","99.1","444.56","0","447.44","33.14","0"
"AGAW1","09W","INS","BOLD:AAA3453","Arthropoda","Insecta","Diptera","Anthomyiidae","Anthomyiinae","Delia","","7974","6923","6923","2","13.18","0","99.32","432.21","0","434.89","34.63","0"
"AGAW1","09W","INS","BOLD:ACE0699","Arthropoda","Insecta","Diptera","Chironomidae","","","","7695","5484","5484","3","28.73","0","98.36","425.09","0","441.2","36.03","0"
"AGAW1","09W","INS","BOLD:AAG9626","Arthropoda","Insecta","Diptera","Dolichopodidae","","","","5980","4674","4674","4","21.84","0","98.47","436.04","0","438.06","36.97","0"
"AGAW1","09W","INS","BOLD:ACE7444","Arthropoda","Insecta","Hemiptera","Miridae","Mirinae, Phylinae","Adelphocoris, Megalocoleus","Adelphocoris lineolatus, Megalocoleus molliculus","5560","3325","3325","5","40.2","0","99.69","442.51","0","443.18","34.82","0"
"AGAW1","09W","INS","BOLD:AAP8739","Arthropoda","Insecta","Diptera","Platystomatidae","Platystomatinae","Rivellia","Rivellia steyskali","5150","3386","3386","6","34.25","0","99.3","441.8","0","443.67","37.98","0"
"AGAW1","09W","INS","BOLD:ABV1415","Arthropoda","Insecta","Diptera","Chironomidae","Chironominae","","","3049","2830","2830","7","7.18","0","98.91","421.8","0","423.56","32.74","0"
"AGAW1","09W","INS","BOLD:AAH4205","Arthropoda","Insecta","Diptera","Chloropidae","Chloropinae","Meromyza","Meromyza nigriseta, Meromyza sp.","2889","1786","1786","8","38.18","0","99.57","447.04","0","447.92","32.9","0"
"AGAW1","09W","INS","BOLD:AAA4555","Arthropoda","Insecta","Orthoptera","Acrididae","Melanoplinae","Melanoplus","Melanoplus stupefactus, Melanoplus angustipennis, Melanoplus ponderosus, Melanoplus bruneri, Melanoplus frigidus, Melanoplus scapularis, Melanoplus cinereus, Melanoplus madeleineae, Melanoplus sanguinipes, Melanoplus nigrescens group, Melanoplus tunicae, Melanoplus walshi, Melanoplus femurrubrum, Melanoplus dawsoni, Melanoplus sp., Melanoplus fasciatus, Melanoplus bowditchi, Melanoplus stonei, Melanoplus bivittatus, Melanoplus huroni, Melanoplus thomasi, Melanoplus borealis, Melanoplus packardii","2829","2321","2321","9","17.96","0","98.91","443.4","0","444.89","36.6","0"
"AGAW1","09W","INS","BOLD:AAH4135","Arthropoda","Insecta","Diptera","Chloropidae","Chloropinae","Thaumatomyia","Thaumatomyia glabra, Thaumatomyia grata, Thaumatomyia sp. 1 TAW","1564","1454","1454","10","7.03","0","99.2","431.21","0","435.7","31.72","0"
"AGAW1","09W","INS","BOLD:AAN6154","Arthropoda","Insecta","Coleoptera","Latridiidae","Corticariinae","Melanophthalma","Melanophthalma inermis","1388","1070","1070","11","22.91","0","99.01","438.5","0","440.31","31.45","0"
"AGAW1","09W","INS","BOLD:AAH0134","Arthropoda","Insecta","Coleoptera","Phalacridae","Phalacrinae","Stilbus","Stilbus apicialis","1379","1154","1154","12","16.32","0","99.32","428.93","0","430.12","31.99","0"
"AGAW1","09W","INS","BOLD:ACK3431","Arthropoda","Insecta","Diptera","Cecidomyiidae","","","","1203","997","997","13","17.12","0","99.14","433.64","0","435.51","32.65","0"
"AGAW1","09W","INS","BOLD:AAA7661","Arthropoda","Insecta","Diptera","Culicidae","Culicinae","Culiseta, Culex","Culiseta melanura, Culex articularis, Culex restuans, Culex acharistus","1176","1058","1058","14","10.03","0","99.16","434.15","0","436.17","32.66","0"
"AGAW1","09W","INS","BOLD:AAU6681","Arthropoda","Insecta","Diptera","Tipulidae","Tipulinae","Nephrotoma","Nephrotoma cornicina","982","776","776","15","20.98","0","99.54","443.64","0","444.77","36.32","0"
"AGAW1","09W","INS","BOLD:AAG5507","Arthropoda","Insecta","Diptera","Chironomidae","Chironominae","Chironomus","Chironomus atrella","913","902","902","16","1.2","0","98.85","416.1","0","420.65","36.06","0"
"AGAW1","09W","INS","BOLD:AAF9771","Arthropoda","Insecta","Diptera","Hybotidae","Tachydromiinae","Platypalpus","Platypalpus longiseta, Platypalpus trivialis, Platypalpus sp. 5, Platypalpus sp. 17, Platypalpus pallidiventris","780","576","576","17","26.15","0","99.47","444.52","0","445.44","35.24","0"
"AGAW1","09W","INS","BOLD:AAN5199","Arthropoda","Insecta","Diptera","Cecidomyiidae","","","CecidInt20 sp. BOLD:AAN5199","718","537","537","18","25.21","0","99.58","438.9","0","446.99","26.7","0"
"AGAW1","09W","INS","BOLD:AAG5453","Arthropoda","Insecta","Diptera","Chironomidae","Chironominae","Chironomus","Chironomus bifurcatus","711","542","542","19","23.77","0","97.83","442.01","0","445.69","35.27","0"
"AGAW1","09W","INS","BOLD:AAB0373","Arthropoda","Insecta","Neuroptera","Chrysopidae","Chrysopinae","Chrysoperla","Chrysoperla sp. isolate 82, Chrysoperla zastrowi, Chrysoperla plorabunda, Chrysoperla sp. isolate 71, Chrysoperla agilis, Italochrysa italica, Chrysoperla sp. isolate 88, Chrysoperla lucasina, Chrysoperla carnea, Chrysoperla mediterranea, Chrysoperla pallida, Chrysoperla nipponensis, Chrysoperla sillemi","587","475","475","20","19.08","0","99.45","436.62","0","437.5","31.36","0"
"AGAW1","09W","INS","BOLD:AAH0135","Arthropoda","Insecta","Coleoptera","Phalacridae","Phalacrinae","Acylomus","Acylomus pugetanus, Acylomus ergoti","560","494","494","21","11.79","0","98.99","436.3","0","438.25","31.68","0"
"AGAW1","09W","INS","BOLD:AAH3661","Arthropoda","Insecta","Diptera","Cecidomyiidae","","","","480","427","427","22","11.04","0","99.32","431.11","0","433.03","25.42","0"
"AGAW1","09W","INS","BOLD:ABX6186","Arthropoda","Insecta","Diptera","Tipulidae","Tipulinae","Nephrotoma","Nephrotoma ferruginea","430","395","395","23","8.14","0","99.29","429.11","0","431.35","34.25","0"
"AGAW1","09W","INS","BOLD:AAH3664","Arthropoda","Insecta","Diptera","Cecidomyiidae","","","CecidInt3 sp. BOLD:AAH3664","383","362","362","24","5.48","0","99.15","429.42","0","432.65","27.01","0"
"AGAW1","09W","INS","BOLD:AAN5225","Arthropoda","Insecta","Diptera","Cecidomyiidae","","","","377","303","303","25","19.63","0","99.3","447.19","0","448.14","28.41","0"
"AGAW1","09W","INS","BOLD:AAG5443","Arthropoda","Insecta","Diptera","Chironomidae","Orthocladiinae","","","365","362","362","26","0.82","0","98.87","411.39","0","413.56","33.54","0"
"AGAW1","09W","INS","BOLD:AAD5253","Arthropoda","Insecta","Diptera","Chironomidae","Orthocladiinae","Thienemanniella","Thienemanniella xena, Thienemanniella vittata","361","287","287","27","20.5","0","98.92","440.49","0","442.04","34.27","0"
"AGAW1","09W","INS","BOLD:AAE0707","Arthropoda","Insecta","Diptera","Chironomidae","Chironominae","Microtendipes","Microtendipes pedellus","356","335","335","28","5.9","0","99.09","433.81","0","436.09","30.71","0"
"AGAW1","09W","INS","BOLD:ABV1297","Arthropoda","Insecta","Diptera","Sciaridae","","","","350","345","345","29","1.43","0","98.98","431.02","0","433.62","30.34","0"
"AGAW1","09W","INS","BOLD:AAL4232","Arthropoda","Insecta","Diptera","Agromyzidae","Phytomyzinae","Phytoliriomyza","Phytoliriomyza arctica","349","283","283","30","18.91","0","99.56","438.95","0","440.28","32.04","0"
"AGAW1","09W","INS","BOLD:AAN6407","Arthropoda","Insecta","Diptera","Sphaeroceridae","Limosininae","Coproica","Coproica ferruginata","335","288","288","31","14.03","0","99.19","444.68","0","446.73","33.37","0"
"AGAW1","09W","INS","BOLD:AAG2898","Arthropoda","Insecta","Hemiptera","Cicadellidae","Deltocephalinae","Paraphlepsius","Paraphlepsius apertus, Paraphlepsius irroratus","317","248","248","32","21.77","0","99.57","441","0","441.84","31.87","0"
"AGAW1","09W","INS","BOLD:AAC0903","Arthropoda","Insecta","Diptera","Chironomidae","Chironominae","Chironomus","Chironomus acidophilus","287","233","233","33","18.82","0","99.41","440.09","0","442.89","36.26","0"
"AGAW1","09W","INS","BOLD:AAP6948","Arthropoda","Insecta","Coleoptera","Carabidae","Trechinae","Bembidion","Bembidion mimus","251","216","216","34","13.94","0","99.47","427.57","0","428.84","34.94","0"
"AGAW1","09W","INS","BOLD:AAG9897","Arthropoda","Insecta","Dermaptera","Forficulidae","Forficulinae","Forficula","Forficula auricularia-A","231","229","229","35","0.87","0","99.01","392.92","0","403.01","45.73","0"
"AGAW1","09W","INS","BOLD:AAZ5592","Arthropoda","Insecta","Diptera","Psychodidae","Psychodinae","","","215","195","195","36","9.3","0","99.22","435.62","0","438","32.58","0"
"AGAW1","09W","INS","BOLD:AAA5803","Arthropoda","Insecta","Hemiptera","Miridae","Mirinae","Lygus","Lygus lineolaris","199","151","151","37","24.12","0","98.59","447.08","0","448.74","33.91","0"
"AGAW1","09W","INS","BOLD:AAN5238","Arthropoda","Insecta","Diptera","Cecidomyiidae","","","","169","137","137","38","18.93","0","99.18","439.76","0","440.95","26.3","0"
"AGAW1","09W","INS","BOLD:ABA0863","Arthropoda","Insecta","Diptera","Cecidomyiidae","","","","167","126","126","39","24.55","0","99.58","442.46","0","443.7","29.78","0"
"AGAW1","09W","INS","BOLD:ABA9112","Arthropoda","Insecta","Diptera","Chironomidae","Chironominae","Chironomus","","165","164","164","40","0.61","0","99","400.79","0","402.47","38.34","0"
"AGAW1","09W","INS","BOLD:AAV5561","Arthropoda","Insecta","Diptera","Cecidomyiidae","","","","161","152","152","41","5.59","0","99.11","443.42","0","445.32","27.17","0"
"AGAW1","09W","INS","BOLD:AAM6249","","","","","","","","156","131","131","42","16.03","0","98.65","444.62","0","446.37","32.49","0"
"AGAW1","09W","INS","BOLD:ACF6513","Arthropoda","Insecta","Diptera","Psychodidae","Psychodinae","Psychoda","Psychoda cinerea","149","137","137","43","8.05","0","99.31","429.03","0","431.74","30.65","0"
"AGAW1","09W","INS","BOLD:ACC8303","Arthropoda","Insecta","Diptera","Cecidomyiidae","","","","146","145","145","44","0.68","0","99.11","426.86","0","429.86","26.19","0"
"AGAW1","09W","INS","BOLD:ACD3384","Arthropoda","Insecta","Hemiptera","Miridae","Orthotylinae","Orthotylus","Orthotylus robiniae","142","112","112","45","21.13","0","99.58","437.37","0","438.43","41.53","0"
"AGAW1","09W","INS","BOLD:AAN6447","Arthropoda","Insecta","Diptera","Sciaridae","","Corynoptera","Corynoptera perpusilla","124","96","96","46","22.58","0","99.36","446.56","0","449.92","36.35","0"
"AGAW1","09W","INS","BOLD:AAG2868","Arthropoda","Insecta","Hemiptera","Cicadellidae","Typhlocybinae","Empoasca","Empoasca sp., Empoasca fabae","99","76","76","47","23.23","0","99.49","446.64","0","447.42","32.26","0"
"AGAW1","09W","INS","BOLD:ABX4079","Arthropoda","Insecta","Diptera","Cecidomyiidae","","","","97","85","85","48","12.37","0","98.49","444.52","0","445.67","28.25","0"
"AGAW1","09W","INS","BOLD:AAE9936","Arthropoda","Insecta","Coleoptera","Lampyridae","Lampyrinae","Lucidota","Lucidota atra","82","69","69","49","15.85","0","98.9","431.22","0","432.65","32.46","0"
"AGAW1","09W","INS","BOLD:AAC4197","Arthropoda","Insecta","Diptera","Chironomidae","Orthocladiinae","Paraphaenocladius","Paraphaenocladius impensus","74","62","62","50","16.22","0","99.48","437.38","0","438.78","35.02","0"
"AGAW1","09W","INS","BOLD:AAZ0271","Arthropoda","Insecta","Diptera","Ceratopogonidae","Ceratopogoninae","Culicoides","","65","65","65","51","0","0","98.46","430.6","0","435.88","41.4","0"
"AGAW1","09W","INS","BOLD:AAG7891","Arthropoda","Insecta","Hymenoptera","Platygastridae","Scelioninae","Telenomus","Telenomus podisi","61","50","50","52","18.03","0","98.84","425.05","0","433.69","27.13","0"
"AGAW1","09W","INS","BOLD:AAF1013","Arthropoda","Insecta","Diptera","Chironomidae","Chironominae","Tanytarsus","Tanytarsus allicis","57","51","51","53","10.53","0","98.04","435.3","0","442.93","36.17","0"
"AGAW1","09W","INS","BOLD:AAN5267","Arthropoda","Insecta","Diptera","Cecidomyiidae","","","","55","38","38","54","30.91","0","99.65","447.02","0","447.67","30.63","0"
"AGAW1","09W","INS","BOLD:AAM9255","Arthropoda","Insecta","Diptera","Sciaridae","","Pseudolycoriella","","52","52","52","55","0","0","98.86","407.38","0","409.96","30.76","0"
"AGAW1","09W","INS","BOLD:ABU5525","Arthropoda","Insecta","Diptera","Chironomidae","Orthocladiinae","Limnophyes","Limnophyes sp. 14ES","51","51","51","56","0","0","99.03","403.43","0","405.65","37.23","0"
"AGAW1","09W","INS","BOLD:ACP4736","Arthropoda","Insecta","Diptera","Chironomidae","Orthocladiinae","Smittia","Smittia sp. 8ES","45","45","45","57","0","0","99.08","423.89","0","425.8","32.88","0"
"AGAW1","09W","INS","BOLD:AAL7819","Arthropoda","Insecta","Diptera","Psychodidae","Psychodinae","Psychoda","Psychoda sp., Psychoda sp. BOLD:AAL7819","42","39","39","58","7.14","0","99.39","416.57","0","418.19","32.69","0"
"AGAW1","09W","INS","BOLD:AAN5183","Arthropoda","Insecta","Diptera","Cecidomyiidae","","","","39","31","31","59","20.51","0","98.37","445.46","0","446.28","27.92","0"
"AGAW1","09W","INS","BOLD:AAN5205","Arthropoda","Insecta","Diptera","Cecidomyiidae","","","CecidInt42 sp. BOLD:AAN5205","38","34","34","60","10.53","0","99.11","448.18","0","449.63","31.52","0"
"AGAW1","09W","INS","BOLD:AAM6655","Arthropoda","Insecta","Diptera","Hybotidae","Tachydromiinae","Platypalpus","Platypalpus stabilis","37","32","32","61","13.51","0","99.54","446.86","0","447.46","35.02","0"
"AGAW1","09W","INS","BOLD:ACM0303","Arthropoda","Insecta","Coleoptera","Anthicidae","Anthicinae","Sapintus","Sapintus lemniscatus","29","27","27","62","6.9","0","99.26","424.48","0","426.21","33.9","0"
"AGAW1","09W","INS","BOLD:ABV1327","Arthropoda","Insecta","Diptera","Cecidomyiidae","","","","28","23","23","63","17.86","0","99.38","436.96","0","437.82","29.18","0"
"AGAW1","09W","INS","BOLD:AAN8770","Arthropoda","Insecta","Diptera","Psychodidae","Psychodinae","Psychoda","Psychoda trinodulosa","27","24","24","64","11.11","0","98.98","433.93","0","434.37","33.37","0"
"AGAW1","09W","INS","BOLD:ACJ6975","","","","","","","","26","26","26","65","0","0","99.44","381.54","0","419.69","36.35","0"
"AGAW1","09W","INS","BOLD:ACV5892","Arthropoda","Insecta","Diptera","Cecidomyiidae","","","","25","25","25","66","0","0","98.42","440.56","0","443.24","25.08","0"
"AGAW1","09W","INS","BOLD:ACY3929","","","","","","","","25","25","25","66","0","0","98.05","448.68","0","450","37.08","0"
"AGAW1","09W","INS","BOLD:ADB5463","Arthropoda","Insecta","Diptera","Chironomidae","","","","23","23","23","68","0","0","99.32","400.22","0","420.87","35.86","0"
"AGAW1","09W","INS","BOLD:AAG1711","Arthropoda","Insecta","Diptera","Muscidae","Phaoniinae","Helina","Helina depuncta","21","20","20","69","4.76","0","98.94","426.52","0","428.67","31.42","0"
"AGAW1","09W","INS","BOLD:ABZ0908","Arthropoda","Insecta","Diptera","Tipulidae","Tipulinae","Nephrotoma","","17","17","17","70","0","0","97.56","364.24","0","367.65","34.17","0"
"AGAW1","09W","INS","BOLD:AAZ2075","Arthropoda","Insecta","Hemiptera","Anthocoridae","Anthocorinae","Orius","Orius insidiosus","17","15","15","70","11.76","0","99.61","444.35","0","445.71","35.15","0"
"AGAW1","09W","INS","BOLD:AAN5342","Arthropoda","Insecta","Diptera","Chironomidae","Orthocladiinae","Heleniella","","15","15","15","72","0","0","99.08","430.27","0","434.8","35.14","0"
"AGAW1","09W","INS","BOLD:AAA7631","Arthropoda","Insecta","Hemiptera","Miridae","Mirinae","Adelphocoris","Adelphocoris lineolatus","13","10","10","73","23.08","0","99.07","449.31","0","450","35.49","0"
"AGAW1","09W","INS","BOLD:AAM9254","Arthropoda","Insecta","Diptera","Sciaridae","","Bradysia","Bradysia pallipes","13","13","13","73","0","0","99.36","430.69","0","431.15","33.19","0"
"AGAW1","09W","INS","BOLD:AAM7855","Arthropoda","Insecta","Thysanoptera","Thripidae","Thripinae","Frankliniella","Frankliniella tritici","12","12","12","75","0","0","99.42","441.25","0","444.17","31.24","0"
"AGAW1","09W","INS","BOLD:AAV5400","Arthropoda","Insecta","Diptera","Cecidomyiidae","","","","12","12","12","75","0","0","97.92","421.75","0","423.5","29.33","0"
"AGAW1","09W","INS","BOLD:AAY6412","Arthropoda","Insecta","Diptera","Sciaridae","","Bradysia","Bradysia urticae","11","10","10","77","9.09","0","98.95","445.73","0","446.73","31.98","0"
"AGAW1","09W","INS","BOLD:AAG8091","Arthropoda","Insecta","Hymenoptera","Ichneumonidae","Campopleginae","Campoletis","Campoletis sp. AB.2","11","11","11","77","0","0","99.02","443.09","0","445.18","24.65","0"
"AGAW1","09W","INS","BOLD:ACJ6984","Arthropoda","Insecta","Hymenoptera","Platygastridae","Scelioninae","Trissolcus","Trissolcus thyantae","10","10","10","79","0","0","99.41","420.2","0","425.8","27.05","0"
"AGAW1","09W","INS","BOLD:AAM6037","Arthropoda","Insecta","Diptera","Cecidomyiidae","Cecidomyiinae","Contarinia, Dasineura","","9","9","9","80","0","0","98.14","439.67","0","442.11","26.07","0"
"AGAW1","09W","INS","BOLD:ACE5981","","","","","","","","8","8","8","81","0","0","97.31","437.38","0","438.75","36.82","0"
"AGAW1","09W","INS","BOLD:ABX2489","Arthropoda","Insecta","Coleoptera","Phalacridae","Phalacrinae","Stilbus","","7","7","7","82","0","0","97.5","434.86","0","436.29","32.35","0"
"AGAW1","09W","INS","BOLD:AAV1136","Arthropoda","Insecta","Diptera","Scatopsidae","","","","6","6","6","83","0","0","99.19","427.33","0","429.17","36.37","0"
"AGAW1","09W","INS","BOLD:ACP6694","Arthropoda","Insecta","Diptera","Chironomidae","Orthocladiinae","","","6","6","6","83","0","0","99.18","402.17","0","404","33.69","0"
"AGAW1","09W","INS","BOLD:AAG6745","Arthropoda","Insecta","Diptera","Calliphoridae","Calliphorinae","Pollenia","Pollenia pediculata","6","6","6","83","0","0","99.28","437.17","0","438.5","33.7","0"
"AGAW1","09W","INS","BOLD:ACP1407","Arthropoda","Insecta","Orthoptera","Acrididae","Melanoplinae","Melanoplus","Melanoplus walshi","5","5","5","86","0","0","97.3","444.8","0","447.8","36.67","0"
"AGAW1","09W","INS","BOLD:AAN5242","Arthropoda","Insecta","Diptera","Cecidomyiidae","","","","4","4","4","87","0","0","99.49","404","0","405.75","28.46","0"
"AGAW1","09W","INS","BOLD:AAV5796","Arthropoda","Insecta","Diptera","Cecidomyiidae","","","","4","4","4","87","0","0","99.72","430.75","0","431.75","29.1","0"
"AGAW1","09W","INS","BOLD:ACJ0111","Arthropoda","Insecta","Hymenoptera","Platygastridae","Scelioninae","Telenomus","","4","4","4","87","0","0","99.15","407.25","0","426.5","28.69","0"
"AGAW1","09W","INS","BOLD:AAH3662","Arthropoda","Insecta","Diptera","Cecidomyiidae","","","CecidInt3 sp. BOLD:AAH3662","4","4","4","87","0","0","99.45","428.5","0","430","26.04","0"
"AGAW1","09W","INS","BOLD:ADH9867","Arthropoda","Insecta","Diptera","","","","","4","4","4","87","0","0","98.76","383","0","386.75","37.38","0"
"AGAW1","09W","INS","BOLD:ACK6193","Arthropoda","Insecta","Diptera","Cecidomyiidae","","","","3","3","3","92","0","0","98.95","443","0","444.33","26.78","0"
"AGAW1","09W","INS","BOLD:AAA6560","Arthropoda","Insecta","Lepidoptera","Crambidae","Crambinae","Crambus","Crambus multilinellus, Crambus saltuellus","3","3","3","92","0","0","98.68","435.67","0","439","31.29","0"
"AGAW1","09W","INS","BOLD:ADB4826","Arthropoda","Insecta","Diptera","Chironomidae","","","","3","3","3","92","0","0","98.46","389","0","450","34.3","0"
"AGAW1","09W","INS","BOLD:AAA2632","Arthropoda","Insecta","Lepidoptera","Noctuidae","Noctuinae","Noctua","Noctua atlantica, Noctua pronuba","3","3","3","92","0","0","99.11","449.33","0","450","31.56","0"
"AGAW1","09W","INS","BOLD:AAF9187","Arthropoda","Insecta","Coleoptera","Curculionidae","Entiminae","Phyllobius","Phyllobius oblongus","3","3","3","92","0","0","99.93","448","0","450","38.89","0"
"AGAW1","09W","INS","BOLD:ACF4925","Arthropoda","Insecta","Hemiptera","Miridae","Mirinae","Adelphocoris","Adelphocoris detritus","3","2","2","92","33.33","0","97.85","448.67","0","450","36.07","0"
"AGAW1","09W","INS","BOLD:AAF9014","Arthropoda","Insecta","Diptera","Limoniidae","Chioneinae","Symplecta","Symplecta cana, Symplecta sp. BOLDAAF9014","3","3","3","92","0","0","99.45","417.33","0","427.67","33.55","0"
"AGAW1","09W","INS","BOLD:AAG2740","Arthropoda","Insecta","Diptera","Ephydridae","Parydrinae","Philygria","Philygria debilis","3","3","3","92","0","0","97.89","397.67","0","401","32.99","0"
"AGAW1","09W","INS","BOLD:AAG2460","Arthropoda","Insecta","Diptera","Anthomyiidae","Anthomyiinae","Hydrophoria","Hydrophoria lancifer","3","3","3","92","0","0","98.89","435","0","436.33","33.65","0"
"AGAW1","09W","INS","BOLD:ACU3040","Arthropoda","Insecta","Diptera","Sciaridae","","","","3","3","3","92","0","0","98.77","416.33","0","417.67","31.91","0"
"AGAW1","09W","INS","BOLD:AAN5249","Arthropoda","Insecta","Diptera","Cecidomyiidae","","","","2","2","2","102","0","0","98.31","407","0","407.5","33.38","0"
"AGAW1","09W","INS","BOLD:AAB4657","Arthropoda","Insecta","Diptera","Chironomidae","Chironominae","Chironomus","Chironomus maturus","2","2","2","102","0","0","99.33","447.5","0","450","37.33","0"
"AGAW1","09W","INS","BOLD:AAA3889","Arthropoda","Insecta","Lepidoptera","Crambidae","Crambinae","Parapediasia","Parapediasia teterrellus","2","2","2","102","0","0","97.04","439.5","0","440","32.61","0"
"AGAW1","09W","INS","BOLD:AAB9140","Arthropoda","Insecta","Diptera","Calliphoridae","Chrysomyinae","Phormia","Phormia regina","2","2","2","102","0","0","98.77","448","0","450","33","0"
"AGAW1","09W","INS","BOLD:AAN5334","Arthropoda","Insecta","Diptera","Chironomidae","Chironominae","Polypedilum","","2","2","2","102","0","0","97.71","370","0","372","34.55","0"
"AGAW1","09W","INS","BOLD:AAI4303","Arthropoda","Insecta","Diptera","Chironomidae","Chironominae","Chironomus","Chironomus melanescens","2","2","2","102","0","0","99.18","434","0","439.5","34.34","0"
"AGAW1","09W","INS","BOLD:AAA3867","Arthropoda","Insecta","Lepidoptera","Noctuidae","Noctuinae","Apamea","Apamea amputatrix","2","2","2","102","0","0","98.45","359.5","0","434.5","34.8","0"
"AGAW1","09W","INS","BOLD:ABY5318","Arthropoda","Insecta","Diptera","Chloropidae","Oscinellinae","Incertella, Rhopalopterum","Rhopalopterum umbrosum, Rhopalopterum luteiceps, Incertella bispina, Rhopalopterum painteri","2","2","2","102","0","0","99.78","450","0","450","32.33","0"
"AGAW1","09W","INS","BOLD:AAD3251","Arthropoda","Insecta","Orthoptera","Acrididae","Oedipodinae","Dissosteira","Dissosteira carolina","2","2","2","102","0","0","99.89","449","0","450","33.78","0"
"AGAW1","09W","INS","BOLD:AAD8086","Arthropoda","Insecta","Lepidoptera","Elachistidae","Elachistinae","Elachista","Elachista griseicornis, Elachista acenteta","2","2","2","102","0","0","97.31","405","0","410","30.09","0"
"AGAW1","09W","INS","BOLD:AAA3181","Arthropoda","Insecta","Diptera","Muscidae","Muscinae","Stomoxys","Stomoxys calcitrans","1","1","1","112","0","0","99.11","448","0","450","33.33","0"
"AGAW1","09W","INS","BOLD:AAB2644","Arthropoda","Insecta","Diptera","Chironomidae","Orthocladiinae","Orthocladius","Orthocladius lapponicus, Orthocladius dorenus, Orthocladius mallochi","1","1","1","112","0","0","100","447","0","450","32","0"
"AGAW1","09W","INS","BOLD:ACG4376","Arthropoda","Insecta","Diptera","Chironomidae","Chironominae","Tanytarsus","","1","1","1","112","0","0","97.1","449","0","450","36.44","0"
"AGAW1","09W","INS","BOLD:ACE4100","Arthropoda","Insecta","Orthoptera","Acrididae","Melanoplinae","Melanoplus","Melanoplus pinicola","1","1","1","112","0","0","97.09","447","0","450","36.22","0"
"AGAW1","09W","INS","BOLD:AAA2518","Arthropoda","Insecta","Lepidoptera","Crambidae","Crambinae","Chrysoteuchia","Chrysoteuchia topiaria, Chrysoteuchia topiarius","1","1","1","112","0","0","97.98","446","0","450","36","0"
"AGAW1","09W","INS","BOLD:AAN5497","Arthropoda","Insecta","Diptera","Anthomyiidae","Pegomyinae","Pegomya","Pegomya solennis, Pegomya bicolor","1","1","1","112","0","0","97.54","448","0","450","29.56","0"
"AGAW1","09W","INS","BOLD:AAA3759","Arthropoda","Insecta","Hemiptera","Aphididae","Calaphidinae","Therioaphis","Therioaphis trifolii","1","1","1","112","0","0","99.78","449","0","450","28.22","0"
"AGAW1","09W","INS","BOLD:AAA8933","Arthropoda","Insecta","Coleoptera","Coccinellidae","Coccinellinae","Coccinella","Coccinella septempunctata","1","1","1","112","0","0","100","450","0","450","33.11","0"
"AGAW1","09W","INS","BOLD:ACA9275","Arthropoda","Insecta","Diptera","Chironomidae","Chironominae","Chironomus","","1","1","1","112","0","0","100","447","0","450","35.78","0"
"AGAW1","09W","INS","BOLD:AAG4477","Arthropoda","Insecta","Coleoptera","Chrysomelidae","Chrysomelinae","Plagiodera","Plagiodera versicolora","1","1","1","112","0","0","99.56","451","0","450","34.22","0"
"AGAW1","09W","INS","BOLD:AAG5457","Arthropoda","Insecta","Diptera","Chironomidae","Chironominae","","","1","1","1","112","0","0","98.19","443","0","450","34.44","0"
"AGAW1","09W","INS","BOLD:AAH3914","Arthropoda","Insecta","Diptera","Sciaridae","","Sciara","Sciara hamatilis, Sciara humeralis","1","1","1","112","0","0","99.75","405","0","406","29.31","0"
"AGAW1","09W","INS","BOLD:AAA5299","Arthropoda","Insecta","Diptera","Chironomidae","Orthocladiinae","Cricotopus","Cricotopus laetus, Cricotopus relucens, Cricotopus trifasciatus, Cricotopus sp., Cricotopus sp. 3ES, Cricotopus sylvestris, Cricotopus glacialis","1","1","1","112","0","0","98.44","448","0","450","35.11","0"
"AGAW1","09W","INS","BOLD:ACE0554","Arthropoda","Insecta","Diptera","Phoridae","Metopininae","Megaselia","","1","1","1","112","0","0","100","450","0","450","32.22","0"
"AGAW1","09W","INS","BOLD:AAV5575","Arthropoda","Insecta","Diptera","Cecidomyiidae","","","CecidInt30 sp. BOLD:AAV5575","1","1","1","112","0","0","100","450","0","450","25.78","0"
"AGAW1","09W","INS","BOLD:AAD0484","Arthropoda","Insecta","Diptera","Chironomidae","","","","1","1","1","112","0","0","98.86","440","0","444","35.14","0"
"AGAW1","09W","INS","BOLD:ACD4236","","","","","","","","1","1","1","112","0","0","97.41","348","0","350","33.14","0"
"AGAW1","09W","INS","BOLD:AAB4894","Arthropoda","Insecta","Hemiptera","Aphididae","Aphidinae","Sitobion","Sitobion smilacifoliae, Sitobion avenae, Sitobion phyllanthi","1","1","1","112","0","0","97.35","452","0","450","24.67","0"
"AGAW1","09W","INS","BOLD:AAM6295","Arthropoda","Insecta","Diptera","Chaoboridae","","","","1","1","1","112","0","0","100","450","0","450","37.11","0"
"AGAW1","09W","INS","BOLD:AAB7030","Arthropoda","Insecta","Diptera","Chironomidae","Chironominae","Chironomus","Chironomus sp. BOLDAAP2996, Chironomus quinnitukqut, Chironomus cf. decorus 2g, Chironomus decorus group sp. 2 IP2013, Chironomus cf. decorus, Chironomus decorus gr.","1","1","1","112","0","0","98.44","448","0","450","34.89","0"
"AGAW1","09W","INS","BOLD:AAD5601","Arthropoda","Insecta","Lepidoptera","Noctuidae","Noctuinae","Apamea","Apamea impulsa","1","1","1","112","0","0","97.99","447","0","450","31.56","0"
"AGAW1","09W","INS","BOLD:ACX6065","Arthropoda","Insecta","Diptera","Cecidomyiidae","","","","1","1","1","112","0","0","99.77","438","0","443","30.02","0"
"AGAW1","09W","INS","BOLD:AAA1525","Arthropoda","Insecta","Trichoptera","Leptoceridae","Leptocerinae","Oecetis","Oecetis inconspicua","1","1","1","112","0","0","98.22","449","0","450","31.78","0"
"AGAW1","09W","INS","BOLD:AAD4214","Arthropoda","Insecta","Hymenoptera","Ichneumonidae","Diplazontinae","Diplazon","Diplazon laetatorius","1","1","1","112","0","0","99.11","448","0","450","26","0"
"AGAW1","09W","INS","BOLD:ACK7314","Arthropoda","Insecta","Diptera","Chironomidae","Chironominae","Polypedilum","","1","1","1","112","0","0","98.91","366","0","370","35.95","0"
"AGAW1","09W","INS","BOLD:AAG2123","Arthropoda","Insecta","Diptera","Tachinidae","Dexiinae","Dinera","Dinera grisescens","1","1","1","112","0","0","98.79","412","0","412","30.58","0"
"AGAW1","09W","INS","BOLD:ACN0444","Arthropoda","Insecta","Hymenoptera","Megaspilidae","","","","1","1","1","112","0","0","99.26","405","0","406","32.27","0"
"AGAW1","09W","INS","BOLD:AAE0845","Arthropoda","Insecta","Hemiptera","Pentatomidae","Pentatominae","Euschistus","Euschistus servus","1","1","1","112","0","0","99.11","450","0","450","38","0"
"AGAW1","09W","INS","BOLD:ACG3646","Arthropoda","Insecta","Coleoptera","Corylophidae","Corylophinae","Orthoperus","","1","1","1","112","0","0","97.1","449","0","450","34.44","0"
"AGAW1","09W","INS","BOLD:AAN5505","Arthropoda","Insecta","Diptera","Hybotidae","Tachydromiinae","Platypalpus","Platypalpus sp. BOLD:AAN5505, Platypalpus minutus, Platypalpus nr. minutus, Platypalpus australominutus","1","1","1","112","0","0","97.54","366","0","369","38.48","0"
"AGAW1","09W","INS","BOLD:ACF0127","Arthropoda","Insecta","Orthoptera","Acrididae","Melanoplinae","Melanoplus","Melanoplus nigrescens group","1","1","1","112","0","0","98","450","0","450","36","0"
"AGAW1","09W","INS","BOLD:ACD8563","Arthropoda","Insecta","Diptera","Cecidomyiidae","","","","1","1","1","112","0","0","97.11","450","0","450","25.78","0"
"AGAW1","09W","INS","BOLD:ABZ4994","Arthropoda","Insecta","Orthoptera","Acrididae","Melanoplinae","Melanoplus","Melanoplus gladstoni, Melanoplus sanguinipes","1","1","1","112","0","0","97.32","447","0","450","36.44","0"
"AGAW1","09W","INS","BOLD:AAB2384","Arthropoda","Insecta","Diptera","Syrphidae","Syrphinae","Eupeodes","Eupeodes duseki, Eupeodes volucris, Eupeodes nielseni, Eupeodes americanus, Eupeodes flukei, Eupeodes vockerothi, Eupeodes lucasi, Eupeodes montivagus, Eupeodes rufipunctatus, Eupeodes rojasi, Eupeodes goeldlini, Eupeodes luniger, Eupeodes punctifer, Eupeodes bucculatus, Eupeodes perplexus, Eupeodes abiskoensis, Eupeodes corollae, Eupeodes pomus, Eupeodes lundbecki, Eupeodes nitens, Eupeodes sp. BOLDAAB2384, Eupeodes curtus, Eupeodes latifasciatus","1","1","1","112","0","0","99.2","374","0","376","30.59","0"
"AGAW1","09W","INS","BOLD:AAB5640","Arthropoda","Insecta","Coleoptera","Coccinellidae","Coccinellinae","Harmonia","Harmonia axyridis","1","1","1","112","0","0","99.1","445","0","450","32","0"
"AGAW1","09W","INS","BOLD:AAC0706","Arthropoda","Insecta","Diptera","Chironomidae","Chironominae","Dicrotendipes","Dicrotendipes tritomus","1","1","1","112","0","0","97.53","445","0","450","36","0"
"AGAW1","09W","INS","BOLD:AAK8756","Arthropoda","Insecta","Diptera","Agromyzidae","Phytomyzinae","Liriomyza","Liriomyza sp. 2 AGT, Liriomyza fricki","1","1","1","112","0","0","98.94","376","0","376","31.65","0"
"AGAW1","09W","INS","BOLD:AAA5953","Arthropoda","Insecta","Lepidoptera","Gelechiidae","Gelechiinae","Coleotechnites","Coleotechnites sp., Coleotechnites piceaella, Coleotechnites laricis, Coleotechnites sp. 3, Coleotechnites carbonaria, Coleotechnites granti, Coleotechnites macleodi, Coleotechnites gibsonella, Coleotechnites coniferella, Coleotechnites martini","1","1","1","112","0","0","98.22","450","0","450","33.33","0"
"AGAW1","09W","INS","BOLD:AAA4277","Arthropoda","Insecta","Diptera","Syrphidae","Syrphinae","Toxomerus","Toxomerus marginatus, Toxomerus sp. CR-3, Toxomerus CR-3","1","1","1","112","0","0","99.78","449","0","450","30.89","0"
"AGAW1","09W","NAR","no match","no match","no match","no match","no match","no match","no match","no match","no match","no match","no match","no match","no match","no match","no match","no match","no match","no match","no match","no match"
"AGAW1","09W","NIN","BOLD:AAI4346","Arthropoda","Arachnida","Opiliones","Phalangiidae","","Paroligolophus, Phalangium","Paroligolophus agrestis, Phalangium opilio","615","591","591","1","3.9","0","98.91","412.31","0","420.17","35.3","0"
"AGAW1","09W","NIN","BOLD:AAA7242","Arthropoda","Collembola","Entomobryomorpha","Entomobryidae","Seirinae, Entomobryinae","Entomobrya, Lepidosira","Entomobrya multifasciata, Lepidosira sp.","429","421","421","2","1.86","0","99.13","416.29","0","420.39","39.29","0"
"AGAW1","09W","NIN","BOLD:AAM7944","Arthropoda","Diplopoda","Julida","Julidae","","Brachyiulus","Brachyiulus pusillus","58","55","55","3","5.17","0","99.44","426.52","0","428.14","40.69","0"
"AGAW1","09W","NIN","BOLD:AAH6630","Arthropoda","Collembola","Entomobryomorpha","Entomobryidae","Entomobryinae","Entomobrya","Entomobrya unostrigata","57","55","55","4","3.51","0","99.42","438.96","0","440.02","39.04","0"
"AGAW1","09W","NIN","BOLD:ABW2639","Arthropoda","Arachnida","Trombidiformes","Anystidae","","","","53","49","49","5","7.55","0","99.54","448.11","0","446.7","31.49","0"
"AGAW1","09W","NIN","BOLD:AAN6371","Arthropoda","Arachnida","Araneae","Theridiidae","","Wamba","Wamba crispulus","46","41","41","6","10.87","0","98.95","440.13","0","441.78","29.41","0"
"AGAW1","09W","NIN","BOLD:AAN6702","Arthropoda","Arachnida","Trombidiformes","Pionidae","","","","10","10","10","7","0","0","98.64","418.3","0","420.4","35.12","0"
"AGAW1","09W","NIN","BOLD:AAB7914","Arthropoda","Collembola","Symphypleona","Bourletiellidae","","Bourletiella","","4","4","4","8","0","0","98.33","449.5","0","450","37.78","0"
"AGAW1","09W","NIN","BOLD:ACZ1295","Arthropoda","Arachnida","Trombidiformes","Anystidae","","","","3","3","3","9","0","0","99.82","374.33","0","379.67","31.35","0"
"AGAW1","09W","NIN","BOLD:ACP9512","Arthropoda","Collembola","Entomobryomorpha","Entomobryidae","","","","3","3","3","9","0","0","98.66","422.33","0","440","39.32","0"
"AGAW1","09W","NIN","BOLD:AAC9044","Arthropoda","Arachnida","Araneae","Salticidae","","Salticus","Salticus scenicus","1","1","1","11","0","0","97.1","449","0","450","33.56","0"
"AGAW1","09W","NIN","BOLD:AAB5726","Arthropoda","Arachnida","Araneae","Agelenidae","","Agelenopsis","Agelenopsis pennsylvanica, Agelenopsis emertoni, Agelenopsis potteri, Agelenopsis actuosa, Agelenopsis kastoni","1","1","1","11","0","0","97.74","442","0","450","33.78","0"
"AGAW1","09W","NIN","BOLD:ADE2039","Arthropoda","Arachnida","Trombidiformes","Tarsonemidae","","","","1","1","1","11","0","0","99.02","408","0","410","27.07","0"
"AGAW1","09W","NIN","BOLD:AAB3836","Arthropoda","Arachnida","Araneae","Philodromidae","","Philodromus","Philodromus cespitum","1","1","1","11","0","0","100","392","0","427","31.62","0"
"AGAW1","09W","NIN","BOLD:AAM3234","Arthropoda","Collembola","Entomobryomorpha","Entomobryidae","Seirinae","Lepidosira","","1","1","1","11","0","0","97.33","450","0","450","40.89","0"
"AGAW1","09W","NIN","BOLD:AAF9234","Arthropoda","Arachnida","Trombidiformes","Eupodidae","","","","1","1","1","11","0","0","99.1","442","0","450","29.33","0"
"AGAW1","10E","BAC","TAX:175768","Proteobacteria","Alphaproteobacteria","Rickettsiales","Rickettsiaceae",NA,"Wolbachia","Wolbachia pipientis","46","27","27","1","41.3","0","97.58","447.35","0","448.43","40.03","0"
"AGAW1","10E","BAC","TAX:91391","Proteobacteria","Alphaproteobacteria","Rickettsiales","",NA,"","","46","43","43","1","6.52","0","98.44","392.33","0","425.3","39.74","0"
"AGAW1","10E","BAC","TAX:91393","Proteobacteria","Alphaproteobacteria","Rickettsiales","Rickettsiaceae",NA,"Wolbachia","","3","3","3","3","0","0","99.58","392.33","0","431.67","39.07","0"
"AGAW1","10E","CHO","no match","no match","no match","no match","no match","no match","no match","no match","no match","no match","no match","no match","no match","no match","no match","no match","no match","no match","no match","no match"
"AGAW1","10E","INS","BOLD:AAP8739","Arthropoda","Insecta","Diptera","Platystomatidae","Platystomatinae","Rivellia","Rivellia steyskali","15540","9689","9689","1","37.65","0","99.32","441.82","0","443.53","37.97","0"
"AGAW1","10E","INS","BOLD:AAA3654","Arthropoda","Insecta","Lepidoptera","Crambidae","Crambinae","Crambus","Crambus bigelovi, Crambus johnsoni, Crambus praefectellus","14538","11456","11456","2","21.2","0","98.77","438.47","0","440.13","31.22","0"
"AGAW1","10E","INS","BOLD:AAA3453","Arthropoda","Insecta","Diptera","Anthomyiidae","Anthomyiinae","Delia","","8675","7447","7447","3","14.16","0","99.34","430.89","0","433.62","34.64","0"
"AGAW1","10E","INS","BOLD:AAB7869","Arthropoda","Insecta","Lepidoptera","Tortricidae","Tortricinae","Phtheochroa","Phtheochroa baracana, Phtheochroa riscana, Phtheochroa birdana, Phtheochroa fulviplicana, Phtheochroa canariana","8094","4904","4904","4","39.41","0","99.33","447.32","0","448.43","29.54","0"
"AGAW1","10E","INS","BOLD:AAU6681","Arthropoda","Insecta","Diptera","Tipulidae","Tipulinae","Nephrotoma","Nephrotoma cornicina","5517","4105","4105","5","25.59","0","99.56","443.17","0","444.35","36.3","0"
"AGAW1","10E","INS","BOLD:AAB4321","Arthropoda","Insecta","Lepidoptera","Gelechiidae","Gelechiinae","Metzneria","Metzneria lappella","3546","3436","3436","6","3.1","0","98.79","410.06","0","427.66","31.84","0"
"AGAW1","10E","INS","BOLD:AAA2087","Arthropoda","Insecta","Lepidoptera","Erebidae","Herminiinae","Idia","Idia sp., Idia americalis","3118","2531","2531","7","18.83","0","99.02","430.5","0","433.82","30.29","0"
"AGAW1","10E","INS","BOLD:AAG2511","Arthropoda","Insecta","Diptera","Anthomyiidae","Anthomyiinae","Delia","Delia platura","2617","2315","2315","8","11.54","0","98.95","443.97","0","446.82","33.14","0"
"AGAW1","10E","INS","BOLD:AAI4303","Arthropoda","Insecta","Diptera","Chironomidae","Chironominae","Chironomus","Chironomus melanescens","2180","1810","1810","9","16.97","0","98.13","431.54","0","437.98","34.3","0"
"AGAW1","10E","INS","BOLD:AAN6154","Arthropoda","Insecta","Coleoptera","Latridiidae","Corticariinae","Melanophthalma","Melanophthalma inermis","1728","1433","1433","10","17.07","0","98.94","434.46","0","437.04","31.23","0"
"AGAW1","10E","INS","BOLD:AAG2868","Arthropoda","Insecta","Hemiptera","Cicadellidae","Typhlocybinae","Empoasca","Empoasca sp., Empoasca fabae","1279","824","824","11","35.57","0","99.56","443.53","0","448.18","32.36","0"
"AGAW1","10E","INS","BOLD:ACE0699","Arthropoda","Insecta","Diptera","Chironomidae","","","","1045","746","746","12","28.61","0","98.44","424.92","0","441.49","36.38","0"
"AGAW1","10E","INS","BOLD:AAA4555","Arthropoda","Insecta","Orthoptera","Acrididae","Melanoplinae","Melanoplus","Melanoplus stupefactus, Melanoplus angustipennis, Melanoplus ponderosus, Melanoplus bruneri, Melanoplus frigidus, Melanoplus scapularis, Melanoplus cinereus, Melanoplus madeleineae, Melanoplus sanguinipes, Melanoplus nigrescens group, Melanoplus tunicae, Melanoplus walshi, Melanoplus femurrubrum, Melanoplus dawsoni, Melanoplus sp., Melanoplus fasciatus, Melanoplus bowditchi, Melanoplus stonei, Melanoplus bivittatus, Melanoplus huroni, Melanoplus thomasi, Melanoplus borealis, Melanoplus packardii","994","844","844","13","15.09","0","99.32","447","0","448.61","36.22","0"
"AGAW1","10E","INS","BOLD:AAH3661","Arthropoda","Insecta","Diptera","Cecidomyiidae","","","","821","716","716","14","12.79","0","99.27","434.1","0","435.79","25.49","0"
"AGAW1","10E","INS","BOLD:ABW7897","Arthropoda","Insecta","Diptera","Cecidomyiidae","","","","775","550","550","15","29.03","0","99.57","443.29","0","445.49","29.66","0"
"AGAW1","10E","INS","BOLD:ACE0626","Arthropoda","Insecta","Coleoptera","Tetratomidae","Eustrophinae","Eustrophopsis","Eustrophopsis bicolor","749","642","642","16","14.29","0","99.31","422.48","0","426.43","32.83","0"
"AGAW1","10E","INS","BOLD:AAA7661","Arthropoda","Insecta","Diptera","Culicidae","Culicinae","Culiseta, Culex","Culiseta melanura, Culex articularis, Culex restuans, Culex acharistus","733","676","676","17","7.78","0","99.11","433.05","0","435.11","32.97","0"
"AGAW1","10E","INS","BOLD:ABW1379","Arthropoda","Insecta","Diptera","Chloropidae","Oscinellinae","Malloewia","Malloewia abdominalis","667","531","531","18","20.39","0","98.34","443.37","0","446.91","31.61","0"
"AGAW1","10E","INS","BOLD:AAH4171","Arthropoda","Insecta","Diptera","Chloropidae","Oscinellinae","Apallates","Apallates montanus, Apallates particeps","597","512","512","19","14.24","0","99.14","434.75","0","436.51","32.7","0"
"AGAW1","10E","INS","BOLD:ACK7314","Arthropoda","Insecta","Diptera","Chironomidae","Chironominae","Polypedilum","","538","534","534","20","0.74","0","98.73","388.88","0","392.72","35.15","0"
"AGAW1","10E","INS","BOLD:AAB7030","Arthropoda","Insecta","Diptera","Chironomidae","Chironominae","Chironomus","Chironomus sp. BOLDAAP2996, Chironomus quinnitukqut, Chironomus cf. decorus 2g, Chironomus decorus group sp. 2 IP2013, Chironomus cf. decorus, Chironomus decorus gr.","517","517","517","21","0","0","98.62","411.43","0","416.2","33.69","0"
"AGAW1","10E","INS","BOLD:ACL2677","Arthropoda","Insecta","Psocodea","Mesopsocidae","","Mesopsocus","Mesopsocus laticeps","454","383","383","22","15.64","0","99.52","435.38","0","436.85","40.98","0"
"AGAW1","10E","INS","BOLD:AAU6966","Arthropoda","Insecta","Coleoptera","Throscidae","Throscinae","Trixagus","Trixagus chevrolati","425","280","280","23","34.12","0","99.78","411.95","0","448.41","39.72","0"
"AGAW1","10E","INS","BOLD:ACF1257","","","","","","","","356","214","214","24","39.89","0","99.72","446.39","0","446.87","34.66","0"
"AGAW1","10E","INS","BOLD:AAG1748","Arthropoda","Insecta","Diptera","Muscidae","Coenosiinae","Coenosia","","350","339","339","25","3.14","0","98.77","426.09","0","428.47","34.28","0"
"AGAW1","10E","INS","BOLD:AAM6781","Arthropoda","Insecta","Diptera","Dolichopodidae","Neurigoninae","Neurigona","Neurigona tenuis","349","324","324","26","7.16","0","99.19","421.36","0","423.49","35.04","0"
"AGAW1","10E","INS","BOLD:AAG7276","Arthropoda","Insecta","Diptera","Sphaeroceridae","Limosininae","Leptocera","Leptocera erythrocera","301","274","274","27","8.97","0","99.16","432.32","0","435.63","32.82","0"
"AGAW1","10E","INS","BOLD:AAG9705","Arthropoda","Insecta","Diptera","Dolichopodidae","","","","243","202","202","28","16.87","0","99.26","443.99","0","445.27","32.09","0"
"AGAW1","10E","INS","BOLD:AAN5199","Arthropoda","Insecta","Diptera","Cecidomyiidae","","","CecidInt20 sp. BOLD:AAN5199","218","180","180","29","17.43","0","99.43","437.71","0","445.57","26.6","0"
"AGAW1","10E","INS","BOLD:ACK3431","Arthropoda","Insecta","Diptera","Cecidomyiidae","","","","212","196","196","30","7.55","0","99.09","427.99","0","429.53","32.68","0"
"AGAW1","10E","INS","BOLD:ABX9581","Arthropoda","Insecta","Hymenoptera","Chrysididae","Chrysidinae","Caenochrysis","","185","143","143","31","22.7","0","98.2","436.35","0","440.36","29.25","0"
"AGAW1","10E","INS","BOLD:AAM9013","Arthropoda","Insecta","Diptera","Keroplatidae","Macrocerinae","Macrocera","Macrocera pusilla","176","167","167","32","5.11","0","99.37","433.86","0","435.35","30.34","0"
"AGAW1","10E","INS","BOLD:AAB8143","Arthropoda","Insecta","Hemiptera","Nabidae","Nabinae","Nabis","Nabis inscriptus, Nabis americoferus, Nabis alternatus, Nabis sp.","176","153","153","32","13.07","0","99.28","437.17","0","438.8","37.24","0"
"AGAW1","10E","INS","BOLD:ABA0863","Arthropoda","Insecta","Diptera","Cecidomyiidae","","","","168","148","148","34","11.9","0","99.18","441.06","0","442.15","29.85","0"
"AGAW1","10E","INS","BOLD:AAP9022","Arthropoda","Insecta","Diptera","Cecidomyiidae","","","","166","138","138","35","16.87","0","99.18","439.59","0","441.05","27.57","0"
"AGAW1","10E","INS","BOLD:AAL4232","Arthropoda","Insecta","Diptera","Agromyzidae","Phytomyzinae","Phytoliriomyza","Phytoliriomyza arctica","149","122","122","36","18.12","0","99.56","438.64","0","439.85","32.02","0"
"AGAW1","10E","INS","BOLD:AAB4657","Arthropoda","Insecta","Diptera","Chironomidae","Chironominae","Chironomus","Chironomus maturus","147","119","119","37","19.05","0","99.3","436.87","0","439.27","37.39","0"
"AGAW1","10E","INS","BOLD:ABV1297","Arthropoda","Insecta","Diptera","Sciaridae","","","","142","140","140","38","1.41","0","99.05","435.89","0","438.59","30.15","0"
"AGAW1","10E","INS","BOLD:AAN5342","Arthropoda","Insecta","Diptera","Chironomidae","Orthocladiinae","Heleniella","","135","128","128","39","5.19","0","99.17","422.21","0","427.3","35.16","0"
"AGAW1","10E","INS","BOLD:AAG6431","Arthropoda","Insecta","Diptera","Ceratopogonidae","Forcipomyiinae","Atrichopogon","","134","116","116","40","13.43","0","98.33","437.8","0","439.4","35.98","0"
"AGAW1","10E","INS","BOLD:ADH5210","Arthropoda","Insecta","Diptera","Cecidomyiidae","","","","133","110","110","41","17.29","0","99.51","445.74","0","446.84","26.37","0"
"AGAW1","10E","INS","BOLD:AAD2041","Arthropoda","Insecta","Diptera","Dolichopodidae","Sciapodinae","Condylostylus","Condylostylus inermis, Condylostylus nigrofemoratus, Condylostylus connectans, Condylostylus caudatus, Condylostylus flavipes","125","95","95","42","24","0","99.39","440.17","0","441.12","35.95","0"
"AGAW1","10E","INS","BOLD:ABU5525","Arthropoda","Insecta","Diptera","Chironomidae","Orthocladiinae","Limnophyes","Limnophyes sp. 14ES","122","119","119","43","2.46","0","98.86","406.02","0","409.17","37.19","0"
"AGAW1","10E","INS","BOLD:AAG9897","Arthropoda","Insecta","Dermaptera","Forficulidae","Forficulinae","Forficula","Forficula auricularia-A","108","108","108","44","0","0","98.96","389.56","0","398.94","45.57","0"
"AGAW1","10E","INS","BOLD:AAN5161","Arthropoda","Insecta","Diptera","Ceratopogonidae","Dasyheleinae","Dasyhelea","","104","102","102","45","1.92","0","98.98","397.47","0","399.86","37.43","0"
"AGAW1","10E","INS","BOLD:AAP5169","Arthropoda","Insecta","Diptera","Chloropidae","Oscinellinae","Elachiptera","Elachiptera nigriceps","103","89","89","46","13.59","0","99.15","434.09","0","435.28","31.24","0"
"AGAW1","10E","INS","BOLD:AAG2898","Arthropoda","Insecta","Hemiptera","Cicadellidae","Deltocephalinae","Paraphlepsius","Paraphlepsius apertus, Paraphlepsius irroratus","98","81","81","47","17.35","0","99.51","445.2","0","445.88","31.9","0"
"AGAW1","10E","INS","BOLD:ADF6661","Arthropoda","Insecta","Diptera","Cecidomyiidae","","","","94","85","85","48","9.57","0","99.27","416.15","0","441.41","29.38","0"
"AGAW1","10E","INS","BOLD:AAA3782","Arthropoda","Insecta","Hymenoptera","Halictidae","Halictinae","Lasioglossum","Lasioglossum weemsi, Lasioglossum leviense, Lasioglossum hitchensi","91","82","82","49","9.89","0","98.95","439.8","0","441.16","28.16","0"
"AGAW1","10E","INS","BOLD:ACY6524","Arthropoda","Insecta","Diptera","Chironomidae","Orthocladiinae","Psectrocladius","Psectrocladius cf. limbatellus","86","81","81","50","5.81","0","99.24","432.33","0","443.19","35.93","0"
"AGAW1","10E","INS","BOLD:AAZ9744","Arthropoda","Insecta","Hymenoptera","Braconidae","Brachistinae","Blacus","Blacus stelfoxi","82","59","59","51","28.05","0","98.44","447.01","0","448.06","32.31","0"
"AGAW1","10E","INS","BOLD:ADC9698","Arthropoda","Insecta","Lepidoptera","","","","","80","72","72","52","10","0","99.27","438.01","0","439.94","33.08","0"
"AGAW1","10E","INS","BOLD:AAN5225","Arthropoda","Insecta","Diptera","Cecidomyiidae","","","","79","66","66","53","16.46","0","99.41","446.38","0","447.34","28.38","0"
"AGAW1","10E","INS","BOLD:AAF9771","Arthropoda","Insecta","Diptera","Hybotidae","Tachydromiinae","Platypalpus","Platypalpus longiseta, Platypalpus trivialis, Platypalpus sp. 5, Platypalpus sp. 17, Platypalpus pallidiventris","75","60","60","54","20","0","99.03","437.36","0","438.01","34.77","0"
"AGAW1","10E","INS","BOLD:AAH0256","Arthropoda","Insecta","Coleoptera","Latridiidae","Corticariinae","Corticaria, Corticarina","","71","62","62","55","12.68","0","99.05","425.1","0","433.39","31.78","0"
"AGAW1","10E","INS","BOLD:ABY5318","Arthropoda","Insecta","Diptera","Chloropidae","Oscinellinae","Incertella, Rhopalopterum","Rhopalopterum umbrosum, Rhopalopterum luteiceps, Incertella bispina, Rhopalopterum painteri","69","50","50","56","27.54","0","99.14","442.64","0","443.67","32.25","0"
"AGAW1","10E","INS","BOLD:ABA0845","Arthropoda","Insecta","Diptera","Cecidomyiidae","","","","68","55","55","57","19.12","0","99.12","447.69","0","449.25","30.96","0"
"AGAW1","10E","INS","BOLD:AAV5938","Arthropoda","Insecta","Diptera","Chironomidae","Chironominae","","","64","51","51","58","20.31","0","99.06","443.94","0","445.09","36.48","0"
"AGAW1","10E","INS","BOLD:AAN5260","Arthropoda","Insecta","Diptera","Cecidomyiidae","","","","62","62","62","59","0","0","99.12","409.76","0","411.42","26","0"
"AGAW1","10E","INS","BOLD:AAN5242","Arthropoda","Insecta","Diptera","Cecidomyiidae","","","","61","61","61","60","0","0","98.9","401.26","0","403.38","28.18","0"
"AGAW1","10E","INS","BOLD:AAM6248","Arthropoda","Insecta","Diptera","Chironomidae","Chironominae","Microtendipes","Microtendipes pedellus grp.","60","43","43","61","28.33","0","97.46","445.03","0","446.07","32.57","0"
"AGAW1","10E","INS","BOLD:AAH0135","Arthropoda","Insecta","Coleoptera","Phalacridae","Phalacrinae","Acylomus","Acylomus pugetanus, Acylomus ergoti","58","57","57","62","1.72","0","98.95","434.41","0","435.9","31.6","0"
"AGAW1","10E","INS","BOLD:AAN5205","Arthropoda","Insecta","Diptera","Cecidomyiidae","","","CecidInt42 sp. BOLD:AAN5205","55","49","49","63","10.91","0","99.28","435.8","0","442.04","31.33","0"
"AGAW1","10E","INS","BOLD:AAE0707","Arthropoda","Insecta","Diptera","Chironomidae","Chironominae","Microtendipes","Microtendipes pedellus","54","50","50","64","7.41","0","99.04","437.24","0","439.02","30.78","0"
"AGAW1","10E","INS","BOLD:AAM9255","Arthropoda","Insecta","Diptera","Sciaridae","","Pseudolycoriella","","54","51","51","64","5.56","0","99.17","424.78","0","426.61","31.12","0"
"AGAW1","10E","INS","BOLD:AAB1098","Arthropoda","Insecta","Diptera","Culicidae","Culicinae","Ochlerotatus, Aedes","Aedes excrucians, Aedes cantans, Aedes stimulans, Aedes euedes, Aedes riparius, Aedes annulipes, Ochlerotatus cantans, Aedes fitchii, Aedes sp., Ochlerotatus annulipes, Ochlerotatus excrucians","53","49","49","66","7.55","0","98.81","436.06","0","437.79","35.35","0"
"AGAW1","10E","INS","BOLD:AAN8446","Arthropoda","Insecta","Psocodea","Peripsocidae","","Peripsocus","Peripsocus madidus","52","52","52","67","0","0","99.16","414.06","0","415.23","41.23","0"
"AGAW1","10E","INS","BOLD:AAC3517","Arthropoda","Insecta","Hemiptera","Delphacidae","Delphacinae","Kosswigianella, Muirodelphax, Delphacodes, Nothodelphax, Elachodelphax","Muirodelphax arvensis, Delphacodes nitens, Delphacodes lutulenta, Muirodelphax nigrinota, Kosswigianella analis, Delphacodes campestris, Muirodelphax anufrievi, Delphacodes indistincta, Muirodelphax unda, Muirodelphax nr. arvensis, Delphacodes stramineosa, Nothodelphax consimilis, Muirodelphax atrolabis, Muirodelphax atralabis, Kosswigianella lutulenta","51","46","46","68","9.8","0","99.43","446.59","0","447.51","37.08","0"
"AGAW1","10E","INS","BOLD:AAH4276","Arthropoda","Insecta","Lepidoptera","Gelechiidae","Gelechiinae","Bryotropha","Bryotropha hodgesi, Bryotropha altitudophila","50","46","46","69","8","0","99.37","439.78","0","441.16","33.67","0"
"AGAW1","10E","INS","BOLD:AAD1485","Arthropoda","Insecta","Diptera","Chironomidae","Chironominae","Paratanytarsus","Paratanytarsus grimmii, Paratanytarsus sp.","49","41","41","70","16.33","0","99.48","440.02","0","441.14","33.78","0"
"AGAW1","10E","INS","BOLD:AAN5238","Arthropoda","Insecta","Diptera","Cecidomyiidae","","","","46","43","43","71","6.52","0","99.31","429.76","0","430.67","25.67","0"
"AGAW1","10E","INS","BOLD:AAN5341","Arthropoda","Insecta","Diptera","Chironomidae","Orthocladiinae","Camptocladius","Camptocladius stercorarius","45","45","45","72","0","0","99.04","400.11","0","405.18","35.39","0"
"AGAW1","10E","INS","BOLD:ABA0822","Arthropoda","Insecta","Diptera","Cecidomyiidae","","","","44","42","42","73","4.55","0","99.01","435.86","0","437.23","26.01","0"
"AGAW1","10E","INS","BOLD:AAN5183","Arthropoda","Insecta","Diptera","Cecidomyiidae","","","","44","40","40","73","9.09","0","98.45","444.77","0","445.86","27.81","0"
"AGAW1","10E","INS","BOLD:AAY6730","Arthropoda","Insecta","Hemiptera","Derbidae","Cedusinae","Cedusa","Cedusa incisa","41","40","40","75","2.44","0","99.05","417.02","0","418.27","35.36","0"
"AGAW1","10E","INS","BOLD:AAV6213","Arthropoda","Insecta","Diptera","Chironomidae","Chironominae","Paratanytarsus","","40","40","40","76","0","0","99.21","420.2","0","427.63","34.22","0"
"AGAW1","10E","INS","BOLD:AAH3664","Arthropoda","Insecta","Diptera","Cecidomyiidae","","","CecidInt3 sp. BOLD:AAH3664","40","38","38","76","5","0","99.22","428.28","0","433.7","26.94","0"
"AGAW1","10E","INS","BOLD:AAN5220","Arthropoda","Insecta","Diptera","Cecidomyiidae","","","","39","38","38","78","2.56","0","98.1","421.38","0","433.74","26.27","0"
"AGAW1","10E","INS","BOLD:AAP2870","Arthropoda","Insecta","Diptera","Dolichopodidae","","","","32","32","32","79","0","0","98.13","416.31","0","418.22","38.25","0"
"AGAW1","10E","INS","BOLD:AAM6062","Arthropoda","Insecta","Diptera","Cecidomyiidae","","","","31","25","25","80","19.35","0","99.56","438.26","0","438.55","26.14","0"
"AGAW1","10E","INS","BOLD:AAZ5592","Arthropoda","Insecta","Diptera","Psychodidae","Psychodinae","","","30","29","29","81","3.33","0","99.19","432.03","0","434.07","32.47","0"
"AGAW1","10E","INS","BOLD:ACC8079","Arthropoda","Insecta","Diptera","Cecidomyiidae","","","","27","25","25","82","7.41","0","99.49","442.15","0","448.33","29.16","0"
"AGAW1","10E","INS","BOLD:ADB4457","Arthropoda","Insecta","Hymenoptera","Braconidae","Euphorinae","","","25","24","24","83","4","0","99.6","379.36","0","447.6","34.62","0"
"AGAW1","10E","INS","BOLD:AAH0134","Arthropoda","Insecta","Coleoptera","Phalacridae","Phalacrinae","Stilbus","Stilbus apicialis","24","23","23","84","4.17","0","99.17","423.79","0","424.71","31.86","0"
"AGAW1","10E","INS","BOLD:AAN5249","Arthropoda","Insecta","Diptera","Cecidomyiidae","","","","22","22","22","85","0","0","98.86","428.05","0","429.82","33","0"
"AGAW1","10E","INS","BOLD:ABV1279","Arthropoda","Insecta","Diptera","Cecidomyiidae","","","","22","19","19","85","13.64","0","99.4","444.91","0","446.05","29.13","0"
"AGAW1","10E","INS","BOLD:ACP8819","Arthropoda","Insecta","Hemiptera","Cicadellidae","Typhlocybinae","","","22","21","21","85","4.55","0","99.89","388.77","0","442.36","31.03","0"
"AGAW1","10E","INS","BOLD:ADG8758","Arthropoda","Insecta","Hymenoptera","Braconidae","Brachistinae","Blacus","Blacus stelfoxi","21","21","21","88","0","0","99.04","440.62","0","441.9","31.96","0"
"AGAW1","10E","INS","BOLD:AAN5250","Arthropoda","Insecta","Diptera","Cecidomyiidae","","","CecidInt50 sp. BOLD:AAN5250","21","18","18","88","14.29","0","99.26","443.24","0","444.29","28.68","0"
"AGAW1","10E","INS","BOLD:AAG1711","Arthropoda","Insecta","Diptera","Muscidae","Phaoniinae","Helina","Helina depuncta","20","20","20","90","0","0","98.87","431.9","0","434.3","31.32","0"
"AGAW1","10E","INS","BOLD:AAP6465","Arthropoda","Insecta","Diptera","Sciaridae","","Corynoptera","Corynoptera cursor","19","10","10","91","47.37","0","98.72","449.42","0","450","32.41","0"
"AGAW1","10E","INS","BOLD:ACF6513","Arthropoda","Insecta","Diptera","Psychodidae","Psychodinae","Psychoda","Psychoda cinerea","17","17","17","92","0","0","99.18","428.12","0","429.82","30.76","0"
"AGAW1","10E","INS","BOLD:ACV2677","Arthropoda","Insecta","Diptera","Cecidomyiidae","","","","15","15","15","93","0","0","97.71","449.07","0","449.93","29.68","0"
"AGAW1","10E","INS","BOLD:AAO8538","Arthropoda","Insecta","Diptera","Psychodidae","Psychodinae","Psychoda","","13","13","13","94","0","0","99.24","426.92","0","429.31","32.22","0"
"AGAW1","10E","INS","BOLD:ABA7871","Arthropoda","Insecta","Diptera","Cecidomyiidae","","","","13","13","13","94","0","0","98.6","443.15","0","443.38","27.02","0"
"AGAW1","10E","INS","BOLD:ACV5892","Arthropoda","Insecta","Diptera","Cecidomyiidae","","","","11","11","11","96","0","0","97.89","447.09","0","448.36","25.46","0"
"AGAW1","10E","INS","BOLD:ADH9867","Arthropoda","Insecta","Diptera","","","","","11","11","11","96","0","0","98.56","411.64","0","416.09","37.51","0"
"AGAW1","10E","INS","BOLD:ABZ3447","Arthropoda","Insecta","Lepidoptera","Tortricidae","Tortricinae","Phtheochroa","Phtheochroa aegrana, Phtheochroa sp.","10","10","10","98","0","0","97.58","437.3","0","441.7","29.4","0"
"AGAW1","10E","INS","BOLD:AAV5689","Arthropoda","Insecta","Diptera","Cecidomyiidae","","","","10","9","9","98","10","0","98.87","434.5","0","436.7","25.58","0"
"AGAW1","10E","INS","BOLD:AAL7356","Arthropoda","Insecta","Diptera","Chironomidae","Chironominae","Tanytarsus","","10","10","10","98","0","0","99.27","442.4","0","443.8","37.79","0"
"AGAW1","10E","INS","BOLD:ACE7444","Arthropoda","Insecta","Hemiptera","Miridae","Mirinae, Phylinae","Adelphocoris, Megalocoleus","Adelphocoris lineolatus, Megalocoleus molliculus","10","10","10","98","0","0","99.78","438.7","0","439.5","34.96","0"
"AGAW1","10E","INS","BOLD:ABV9308","Arthropoda","Insecta","Diptera","Chironomidae","Chironominae","Tanytarsus","","9","9","9","102","0","0","97.27","430.67","0","431.44","37.14","0"
"AGAW1","10E","INS","BOLD:AAP8816","Arthropoda","Insecta","Diptera","Cecidomyiidae","Cecidomyiinae","Sitodiplosis","Sitodiplosis sp. BOLD:AAP8816","9","9","9","102","0","0","98.7","436.56","0","438","26.45","0"
"AGAW1","10E","INS","BOLD:AAG7891","Arthropoda","Insecta","Hymenoptera","Platygastridae","Scelioninae","Telenomus","Telenomus podisi","9","9","9","102","0","0","99.27","422.78","0","430.56","27.12","0"
"AGAW1","10E","INS","BOLD:AAG3154","Arthropoda","Insecta","Diptera","Psychodidae","Psychodinae","Psychoda","","9","9","9","102","0","0","97.97","449.11","0","450","32.1","0"
"AGAW1","10E","INS","BOLD:AAY6469","Arthropoda","Insecta","Diptera","Cecidomyiidae","","","","9","9","9","102","0","0","99.1","429.89","0","431.22","26.23","0"
"AGAW1","10E","INS","BOLD:AAG8683","Arthropoda","Insecta","Hemiptera","Cicadellidae","Eurymelinae, Typhlocybinae","Empoasca, Idiocerus","Empoasca luda, Idiocerus populi","9","9","9","102","0","0","97.72","400.22","0","402.44","31.01","0"
"AGAW1","10E","INS","BOLD:AAN5221","Arthropoda","Insecta","Diptera","Cecidomyiidae","","","","9","9","9","102","0","0","99.13","431.44","0","433.44","25.53","0"
"AGAW1","10E","INS","BOLD:AAN5255","Arthropoda","Insecta","Diptera","Cecidomyiidae","Cecidomyiinae","Dasineura","","8","8","8","109","0","0","98.55","434.63","0","440","26.83","0"
"AGAW1","10E","INS","BOLD:AAA3882","Arthropoda","Insecta","Lepidoptera","Pterophoridae","Pterophorinae","Emmelina","Emmelina monodactyla","8","8","8","109","0","0","98.36","405.75","0","413.5","31.33","0"
"AGAW1","10E","INS","BOLD:ABX7402","Arthropoda","Insecta","Diptera","Cecidomyiidae","","","","7","6","6","111","14.29","0","99.97","419.43","0","436","27.71","0"
"AGAW1","10E","INS","BOLD:AAN5230","Arthropoda","Insecta","Diptera","Cecidomyiidae","","","","7","7","7","111","0","0","98.21","411.57","0","413.86","25.6","0"
"AGAW1","10E","INS","BOLD:ADF3937","","","","","","","","7","7","7","111","0","0","99.2","409.57","0","420.86","31.24","0"
"AGAW1","10E","INS","BOLD:AAV8368","Arthropoda","Insecta","Lepidoptera","Nepticulidae","Nepticulinae","Stigmella","Stigmella saginella","6","6","6","114","0","0","98.66","409.83","0","412.67","32.35","0"
"AGAW1","10E","INS","BOLD:AAB0079","Arthropoda","Insecta","Diptera","Chironomidae","Orthocladiinae","Corynoneura","Corynoneura arctica","6","6","6","114","0","0","98.29","431","0","431","34.8","0"
"AGAW1","10E","INS","BOLD:AAG6460","Arthropoda","Insecta","Diptera","Cecidomyiidae","","","","6","6","6","114","0","0","98.7","447.67","0","450","34.41","0"
"AGAW1","10E","INS","BOLD:AAN6447","Arthropoda","Insecta","Diptera","Sciaridae","","Corynoptera","Corynoptera perpusilla","6","6","6","114","0","0","99.4","448.33","0","450","36.26","0"
"AGAW1","10E","INS","BOLD:AAG9668","Arthropoda","Insecta","Diptera","Dolichopodidae","Diaphorinae","Chrysotus","Chrysotus sp., Chrysotus sp. nr. obliquus","5","5","5","118","0","0","99.33","446.6","0","448","32.95","0"
"AGAW1","10E","INS","BOLD:AAC0903","Arthropoda","Insecta","Diptera","Chironomidae","Chironominae","Chironomus","Chironomus acidophilus","5","5","5","118","0","0","99.09","441.8","0","442.8","36.58","0"
"AGAW1","10E","INS","BOLD:AAM6037","Arthropoda","Insecta","Diptera","Cecidomyiidae","Cecidomyiinae","Contarinia, Dasineura","","5","5","5","118","0","0","98.29","437","0","439","26.3","0"
"AGAW1","10E","INS","BOLD:AAA6292","Arthropoda","Insecta","Lepidoptera","Tortricidae","Tortricinae","Cnephasia","Cnephasia genitalana, Cnephasia stephensiana","4","4","4","121","0","0","99.78","445.75","0","446.5","35.67","0"
"AGAW1","10E","INS","BOLD:AAF1729","Arthropoda","Insecta","Psocodea","Lachesillidae","Lachesillinae","Lachesilla","Lachesilla pedicularia","4","4","4","121","0","0","99.16","444","0","446.25","36.87","0"
"AGAW1","10E","INS","BOLD:ABA1215","Arthropoda","Insecta","Diptera","Sciaridae","","Lycoriella","Lycoriella sativae","4","4","4","121","0","0","99.49","421.5","0","424.25","35.9","0"
"AGAW1","10E","INS","BOLD:ABA0858","Arthropoda","Insecta","Diptera","Cecidomyiidae","","","","4","4","4","121","0","0","98.73","403.5","0","405","25.7","0"
"AGAW1","10E","INS","BOLD:AAZ0293","Arthropoda","Insecta","Diptera","Phoridae","Metopininae","Puliciphora","","4","4","4","121","0","0","98.15","422.75","0","423.75","35.15","0"
"AGAW1","10E","INS","BOLD:AAA4277","Arthropoda","Insecta","Diptera","Syrphidae","Syrphinae","Toxomerus","Toxomerus marginatus, Toxomerus sp. CR-3, Toxomerus CR-3","4","4","4","121","0","0","99.77","440.25","0","441","30.47","0"
"AGAW1","10E","INS","BOLD:AAG3236","Arthropoda","Insecta","Diptera","Phoridae","Phorinae","Conicera, Diplonevra","Diplonevra nitidula","4","4","4","121","0","0","99.55","447.25","0","450","31.83","0"
"AGAW1","10E","INS","BOLD:ABX4079","Arthropoda","Insecta","Diptera","Cecidomyiidae","","","","4","4","4","121","0","0","98.5","449.25","0","450","28.33","0"
"AGAW1","10E","INS","BOLD:AAL7819","Arthropoda","Insecta","Diptera","Psychodidae","Psychodinae","Psychoda","Psychoda sp., Psychoda sp. BOLD:AAL7819","4","4","4","121","0","0","99.13","430.5","0","431.5","32.78","0"
"AGAW1","10E","INS","BOLD:AAB0373","Arthropoda","Insecta","Neuroptera","Chrysopidae","Chrysopinae","Chrysoperla","Chrysoperla sp. isolate 82, Chrysoperla zastrowi, Chrysoperla plorabunda, Chrysoperla sp. isolate 71, Chrysoperla agilis, Italochrysa italica, Chrysoperla sp. isolate 88, Chrysoperla lucasina, Chrysoperla carnea, Chrysoperla mediterranea, Chrysoperla pallida, Chrysoperla nipponensis, Chrysoperla sillemi","4","4","4","121","0","0","99.78","407.75","0","408","31.6","0"
"AGAW1","10E","INS","BOLD:AAD1395","Arthropoda","Insecta","Diptera","Chironomidae","Chironominae","Polypedilum","Polypedilum convictum, Polypedilum flavum","4","2","2","121","50","0","98.94","450","0","450","33.94","0"
"AGAW1","10E","INS","BOLD:AAG6745","Arthropoda","Insecta","Diptera","Calliphoridae","Calliphorinae","Pollenia","Pollenia pediculata","4","4","4","121","0","0","99.42","410","0","411","33.96","0"
"AGAW1","10E","INS","BOLD:AAU8379","Arthropoda","Insecta","Hymenoptera","Braconidae","Alysiinae","Dinotrema","Dinotrema JFT01","4","4","4","121","0","0","99.55","448.75","0","450","24.39","0"
"AGAW1","10E","INS","BOLD:ACC5649","Arthropoda","Insecta","Diptera","Cecidomyiidae","","","CecidInt23 sp. BOLD:ACC5649","3","3","3","134","0","0","99.12","416","0","418.33","25.14","0"
"AGAW1","10E","INS","BOLD:ACJ6954","Arthropoda","Insecta","Diptera","Cecidomyiidae","","","","3","3","3","134","0","0","98.53","339.33","0","396","26.73","0"
"AGAW1","10E","INS","BOLD:AAA3867","Arthropoda","Insecta","Lepidoptera","Noctuidae","Noctuinae","Apamea","Apamea amputatrix","3","3","3","134","0","0","98.8","433","0","436.67","33.49","0"
"AGAW1","10E","INS","BOLD:AAA3889","Arthropoda","Insecta","Lepidoptera","Crambidae","Crambinae","Parapediasia","Parapediasia teterrellus","3","3","3","134","0","0","99.16","428.33","0","430","32.07","0"
"AGAW1","10E","INS","BOLD:AAV1136","Arthropoda","Insecta","Diptera","Scatopsidae","","","","3","3","3","134","0","0","98.89","449","0","449.67","36.92","0"
"AGAW1","10E","INS","BOLD:AAN5215","Arthropoda","Insecta","Diptera","Cecidomyiidae","","","","3","3","3","134","0","0","98.25","415","0","418.33","29.02","0"
"AGAW1","10E","INS","BOLD:AAH3662","Arthropoda","Insecta","Diptera","Cecidomyiidae","","","CecidInt3 sp. BOLD:AAH3662","3","3","3","134","0","0","99.26","448.33","0","450","26.3","0"
"AGAW1","10E","INS","BOLD:ABV1337","Arthropoda","Insecta","Diptera","Cecidomyiidae","","","","3","3","3","134","0","0","98.95","445.67","0","450","28.15","0"
"AGAW1","10E","INS","BOLD:AAA4781","Arthropoda","Insecta","Hymenoptera","Braconidae","Microgastrinae","Glyptapanteles","Glyptapanteles carlossarmientoi","3","3","3","134","0","0","99.47","424.33","0","426.67","24.18","0"
"AGAW1","10E","INS","BOLD:AAV1261","Arthropoda","Insecta","Diptera","Sciaridae","","Bradysia","Bradysia splendida","3","3","3","134","0","0","99.18","441.67","0","443","34","0"
"AGAW1","10E","INS","BOLD:AAN6149","Arthropoda","Insecta","Coleoptera","Coccinellidae","Coccinellinae","Stethorus","Stethorus punctillum","3","3","3","134","0","0","98.18","374.33","0","388.67","36.19","0"
"AGAW1","10E","INS","BOLD:AAH4205","Arthropoda","Insecta","Diptera","Chloropidae","Chloropinae","Meromyza","Meromyza nigriseta, Meromyza sp.","3","2","2","134","33.33","0","98","450","0","450","33.26","0"
"AGAW1","10E","INS","BOLD:AAA1513","Arthropoda","Insecta","Lepidoptera","Plutellidae","Plutellinae","Plutella, Rhigognostis","Plutella xylostella, Rhigognostis sp.","3","3","3","134","0","0","99.19","407.67","0","408.67","30.67","0"
"AGAW1","10E","INS","BOLD:ABZ3448","Arthropoda","Insecta","Lepidoptera","Tortricidae","Tortricinae","Phtheochroa","Phtheochroa inopiana","2","2","2","147","0","0","97.4","442.5","0","450","29.33","0"
"AGAW1","10E","INS","BOLD:ACK5742","Arthropoda","Insecta","Diptera","Cecidomyiidae","","","","2","2","2","147","0","0","98.89","449","0","450","29.22","0"
"AGAW1","10E","INS","BOLD:AAA6560","Arthropoda","Insecta","Lepidoptera","Crambidae","Crambinae","Crambus","Crambus multilinellus, Crambus saltuellus","2","2","2","147","0","0","99.2","442","0","443.5","31.12","0"
"AGAW1","10E","INS","BOLD:AAM7398","Arthropoda","Insecta","Hymenoptera","Ichneumonidae","Campopleginae","Hyposoter","","2","2","2","147","0","0","97.48","397","0","405.5","26.1","0"
"AGAW1","10E","INS","BOLD:AAV5594","Arthropoda","Insecta","Diptera","Cecidomyiidae","","","","2","2","2","147","0","0","99.33","448","0","450","26","0"
"AGAW1","10E","INS","BOLD:AAA5299","Arthropoda","Insecta","Diptera","Chironomidae","Orthocladiinae","Cricotopus","Cricotopus laetus, Cricotopus relucens, Cricotopus trifasciatus, Cricotopus sp., Cricotopus sp. 3ES, Cricotopus sylvestris, Cricotopus glacialis","2","2","2","147","0","0","99.66","447","0","450","35","0"
"AGAW1","10E","INS","BOLD:AAU7040","Arthropoda","Insecta","Coleoptera","Corylophidae","Corylophinae","Orthoperus","Orthoperus scutellaris","2","2","2","147","0","0","97.76","423","0","424","32.29","0"
"AGAW1","10E","INS","BOLD:AAV5400","Arthropoda","Insecta","Diptera","Cecidomyiidae","","","","2","2","2","147","0","0","98.18","413","0","414.5","30.24","0"
"AGAW1","10E","INS","BOLD:ACK1456","Arthropoda","Insecta","Diptera","Cecidomyiidae","","","","2","2","2","147","0","0","100","414","0","424","24.95","0"
"AGAW1","10E","INS","BOLD:AAN5497","Arthropoda","Insecta","Diptera","Anthomyiidae","Pegomyinae","Pegomya","Pegomya solennis, Pegomya bicolor","2","2","2","147","0","0","99.22","448","0","450","29.56","0"
"AGAW1","10E","INS","BOLD:AAD4214","Arthropoda","Insecta","Hymenoptera","Ichneumonidae","Diplazontinae","Diplazon","Diplazon laetatorius","2","2","2","147","0","0","98.22","449.5","0","450","27","0"
"AGAW1","10E","INS","BOLD:ABX6186","Arthropoda","Insecta","Diptera","Tipulidae","Tipulinae","Nephrotoma","Nephrotoma ferruginea","2","2","2","147","0","0","99.43","439.5","0","441","34.58","0"
"AGAW1","10E","INS","BOLD:AAA2815","Arthropoda","Insecta","Lepidoptera","Noctuidae","Noctuinae","Mythimna","Mythimna yukonensis, Mythimna oxygala","2","2","2","147","0","0","98.15","432","0","434","33.51","0"
"AGAW1","10E","INS","BOLD:AAA6759","Arthropoda","Insecta","Lepidoptera","Hesperiidae","Hesperiinae","Thymelicus","Thymelicus lineola","1","1","1","160","0","0","97.97","444","0","450","29.56","0"
"AGAW1","10E","INS","BOLD:AAG1714","Arthropoda","Insecta","Diptera","Muscidae","Azeliinae","Muscina","Muscina pascuorum","1","1","1","160","0","0","98.6","429","0","450","33.78","0"
"AGAW1","10E","INS","BOLD:AAG1717","Arthropoda","Insecta","Diptera","Muscidae","Phaoniinae","Helina","Helina troene","1","1","1","160","0","0","99.73","374","0","377","30.77","0"
"AGAW1","10E","INS","BOLD:ACE8806","Arthropoda","Insecta","Diptera","Cecidomyiidae","","","","1","1","1","160","0","0","98.13","427","0","450","26.89","0"
"AGAW1","10E","INS","BOLD:ABA0811","Arthropoda","Insecta","Diptera","Cecidomyiidae","","","","1","1","1","160","0","0","99.78","449","0","450","26.44","0"
"AGAW1","10E","INS","BOLD:AAA4201","Arthropoda","Insecta","Lepidoptera","Noctuidae","Noctuinae","Apamea","Apamea sp., Apamea sordens, Apamea sp. nr. sordens, Apamea digitula","1","1","1","160","0","0","98.89","449","0","450","32","0"
"AGAW1","10E","INS","BOLD:ABV9293","Arthropoda","Insecta","Diptera","Chironomidae","Chironominae","Dicrotendipes","","1","1","1","160","0","0","99.16","356","0","415","34.22","0"
"AGAW1","10E","INS","BOLD:ACB2629","Arthropoda","Insecta","Diptera","Cecidomyiidae","","","","1","1","1","160","0","0","99.78","449","0","450","30.22","0"
"AGAW1","10E","INS","BOLD:ACI4417","Arthropoda","Insecta","Diptera","Cecidomyiidae","","","","1","1","1","160","0","0","98.4","376","0","408","29.17","0"
"AGAW1","10E","INS","BOLD:AAG5453","Arthropoda","Insecta","Diptera","Chironomidae","Chironominae","Chironomus","Chironomus bifurcatus","1","1","1","160","0","0","97.77","449","0","450","35.11","0"
"AGAW1","10E","INS","BOLD:AAC5210","Arthropoda","Insecta","Diptera","Culicidae","Culicinae","Aedes, Aedimorphus","Aedes japonicus, Aedimorphus caecus","1","1","1","160","0","0","97.88","377","0","392","34.44","0"
"AGAW1","10E","INS","BOLD:AAF9393","Arthropoda","Insecta","Coleoptera","Meloidae","Meloinae","Epicauta","Epicauta fabricii","1","1","1","160","0","0","98.88","448","0","450","42.44","0"
"AGAW1","10E","INS","BOLD:AAE7186","Arthropoda","Insecta","Hymenoptera","Braconidae","Aphidiinae","Praon","Praon sp. 1 MM-2013, Praon volucre, Praon bicolor","1","1","1","160","0","0","99.11","448","0","450","26","0"
"AGAW1","10E","INS","BOLD:ACA3113","Arthropoda","Insecta","Psocodea","Caeciliusidae","","","","1","1","1","160","0","0","98.55","413","0","417","36.69","0"
"AGAW1","10E","INS","BOLD:AAL7442","Arthropoda","Insecta","Diptera","Cecidomyiidae","","","","1","1","1","160","0","0","97.56","450","0","450","26.22","0"
"AGAW1","10E","INS","BOLD:AAD5253","Arthropoda","Insecta","Diptera","Chironomidae","Orthocladiinae","Thienemanniella","Thienemanniella xena, Thienemanniella vittata","1","1","1","160","0","0","99.11","449","0","450","34.22","0"
"AGAW1","10E","INS","BOLD:AAV5788","Arthropoda","Insecta","Diptera","Cecidomyiidae","Cecidomyiinae","Asphondylia","","1","1","1","160","0","0","100","448","0","450","24","0"
"AGAW1","10E","INS","BOLD:AAA2632","Arthropoda","Insecta","Lepidoptera","Noctuidae","Noctuinae","Noctua","Noctua atlantica, Noctua pronuba","1","1","1","160","0","0","99.09","438","0","450","30.67","0"
"AGAW1","10E","INS","BOLD:ACJ0111","Arthropoda","Insecta","Hymenoptera","Platygastridae","Scelioninae","Telenomus","","1","1","1","160","0","0","98.63","439","0","450","28.89","0"
"AGAW1","10E","INS","BOLD:AAB9140","Arthropoda","Insecta","Diptera","Calliphoridae","Chrysomyinae","Phormia","Phormia regina","1","1","1","160","0","0","100","450","0","450","33.11","0"
"AGAW1","10E","INS","BOLD:ACI0042","Arthropoda","Insecta","Psocodea","Liposcelidae","Liposcelidinae","Liposcelis","Liposcelis decolor","1","1","1","160","0","0","97.1","449","0","450","36.44","0"
"AGAW1","10E","INS","BOLD:ADF9260","Arthropoda","Insecta","Coleoptera","Phalacridae","Phalacrinae","Acylomus","","1","1","1","160","0","0","98.21","447","0","450","31.11","0"
"AGAW1","10E","INS","BOLD:AAB9332","Arthropoda","Insecta","Hemiptera","Aphididae","Aphidinae","Nasonovia","Nasonovia ribisnigri","1","1","1","160","0","0","100","450","0","450","25.11","0"
"AGAW1","10E","INS","BOLD:ACL2601","Arthropoda","Insecta","Hemiptera","Cicadellidae","","","","1","1","1","160","0","0","98.89","450","0","450","36.44","0"
"AGAW1","10E","INS","BOLD:ACA6596","Arthropoda","Insecta","Diptera","Cecidomyiidae","","","","1","1","1","160","0","0","99.3","431","0","450","27.33","0"
"AGAW1","10E","INS","BOLD:ACX6065","Arthropoda","Insecta","Diptera","Cecidomyiidae","","","","1","1","1","160","0","0","99.55","448","0","450","29.56","0"
"AGAW1","10E","INS","BOLD:AAV1055","Arthropoda","Insecta","Diptera","Stratiomyidae","Beridinae","Allognosta","Allognosta fuscitarsis","1","1","1","160","0","0","100","431","0","432","36.57","0"
"AGAW1","10E","INS","BOLD:AAN3785","Arthropoda","Insecta","Hemiptera","Aphididae","Aphidinae","Aphis","Aphis urticata","1","1","1","160","0","0","99.78","445","0","450","26.89","0"
"AGAW1","10E","INS","BOLD:AAG9626","Arthropoda","Insecta","Diptera","Dolichopodidae","","","","1","1","1","160","0","0","98.66","447","0","450","36.89","0"
"AGAW1","10E","INS","BOLD:AAG2123","Arthropoda","Insecta","Diptera","Tachinidae","Dexiinae","Dinera","Dinera grisescens","1","1","1","160","0","0","99.78","448","0","450","32","0"
"AGAW1","10E","INS","BOLD:AAD9720","Arthropoda","Insecta","Lepidoptera","Tortricidae","Tortricinae","Phtheochroa","Phtheochroa vulneratana","1","1","1","160","0","0","97.93","435","0","438","29.45","0"
"AGAW1","10E","INS","BOLD:AAH4135","Arthropoda","Insecta","Diptera","Chloropidae","Chloropinae","Thaumatomyia","Thaumatomyia glabra, Thaumatomyia grata, Thaumatomyia sp. 1 TAW","1","1","1","160","0","0","99.49","393","0","394","31.22","0"
"AGAW1","10E","INS","BOLD:ABZ0621","Arthropoda","Insecta","Diptera","Psychodidae","Psychodinae","","","1","1","1","160","0","0","99.78","450","0","450","33.33","0"
"AGAW1","10E","INS","BOLD:ABV1439","Arthropoda","Insecta","Diptera","Cecidomyiidae","","","","1","1","1","160","0","0","98.43","447","0","450","28.67","0"
"AGAW1","10E","INS","BOLD:AAI8935","Arthropoda","Insecta","Coleoptera","Latridiidae","Corticariinae","Cortinicara","Cortinicara gibbosa","1","1","1","160","0","0","98.38","371","0","372","33.06","0"
"AGAW1","10E","INS","BOLD:ABZ4994","Arthropoda","Insecta","Orthoptera","Acrididae","Melanoplinae","Melanoplus","Melanoplus gladstoni, Melanoplus sanguinipes","1","1","1","160","0","0","97.11","450","0","450","36.67","0"
"AGAW1","10E","INS","BOLD:AAU0577","Arthropoda","Insecta","Thysanoptera","Aeolothripidae","Aeolothripinae","Aeolothrips","Aeolothrips fasciatus","1","1","1","160","0","0","98.21","390","0","391","36.06","0"
"AGAW1","10E","INS","BOLD:ABZ2233","Arthropoda","Insecta","Lepidoptera","Noctuidae","Noctuinae","Oligia","Oligia modica","1","1","1","160","0","0","98.21","390","0","394","29.44","0"
"AGAW1","10E","INS","BOLD:ACK6937","Arthropoda","Insecta","Diptera","Cecidomyiidae","","","","1","1","1","160","0","0","100","449","0","450","26.67","0"
"AGAW1","10E","INS","BOLD:AAG9744","Arthropoda","Insecta","Diptera","Dolichopodidae","","","","1","1","1","160","0","0","99.33","448","0","450","38","0"
"AGAW1","10E","INS","BOLD:AAO7694","Arthropoda","Insecta","Diptera","Cecidomyiidae","","","","1","1","1","160","0","0","97.98","446","0","450","27.56","0"
"AGAW1","10E","INS","BOLD:AAG2460","Arthropoda","Insecta","Diptera","Anthomyiidae","Anthomyiinae","Hydrophoria","Hydrophoria lancifer","1","1","1","160","0","0","97.37","419","0","450","33.33","0"
"AGAW1","10E","NAR","no match","no match","no match","no match","no match","no match","no match","no match","no match","no match","no match","no match","no match","no match","no match","no match","no match","no match","no match","no match"
"AGAW1","10E","NIN","BOLD:AAU6252","Arthropoda","Arachnida","Trombidiformes","Bdellidae","","Cyta","","717","573","573","1","20.08","0","98.43","437.55","0","438.78","31.53","0"
"AGAW1","10E","NIN","BOLD:AAA7242","Arthropoda","Collembola","Entomobryomorpha","Entomobryidae","Seirinae, Entomobryinae","Entomobrya, Lepidosira","Entomobrya multifasciata, Lepidosira sp.","245","241","241","2","1.63","0","99.05","417.93","0","422.01","39.3","0"
"AGAW1","10E","NIN","BOLD:ACB9565","Arthropoda","Arachnida","Araneae","Clubionidae","","Clubiona","Clubiona pygmaea","222","210","210","3","5.41","0","99.25","431.59","0","434.13","30.74","0"
"AGAW1","10E","NIN","BOLD:AAB4300","Arthropoda","Arachnida","Araneae","Thomisidae","","Xysticus","Xysticus emertoni","199","159","159","4","20.1","0","98.87","403.19","0","447.46","32.34","0"
"AGAW1","10E","NIN","BOLD:ACY2550","Arthropoda","Arachnida","Trombidiformes","Bdellidae","","Cyta","","124","124","124","5","0","0","98.86","377.56","0","399.65","31.12","0"
"AGAW1","10E","NIN","BOLD:ABW2639","Arthropoda","Arachnida","Trombidiformes","Anystidae","","","","82","71","71","6","13.41","0","99.47","449.05","0","447.3","31.49","0"
"AGAW1","10E","NIN","BOLD:AAA6381","Arthropoda","Arachnida","Araneae","Tetragnathidae","","Tetragnatha","Tetragnatha laboriosa","18","18","18","7","0","0","99.04","439.06","0","442.28","39.43","0"
"AGAW1","10E","NIN","BOLD:ACX8969","Arthropoda","Arachnida","Trombidiformes","Eupodidae","","","","14","14","14","8","0","0","98.42","424.14","0","427.5","27.77","0"
"AGAW1","10E","NIN","BOLD:AAA9476","Arthropoda","Arachnida","Araneae","Linyphiidae","","Grammonota","Grammonota gentilis","11","11","11","9","0","0","98.91","449","0","450","30.87","0"
"AGAW1","10E","NIN","BOLD:AAB7914","Arthropoda","Collembola","Symphypleona","Bourletiellidae","","Bourletiella","","11","11","11","9","0","0","99.21","448.18","0","450","38.14","0"
"AGAW1","10E","NIN","BOLD:ACZ1295","Arthropoda","Arachnida","Trombidiformes","Anystidae","","","","6","6","6","11","0","0","99.69","372.5","0","380.5","30.97","0"
"AGAW1","10E","NIN","BOLD:AAM7961","Arthropoda","Arachnida","Trombidiformes","Anystidae","","","","6","6","6","11","0","0","99.48","449.17","0","450","31.44","0"
"AGAW1","10E","NIN","BOLD:AAH6630","Arthropoda","Collembola","Entomobryomorpha","Entomobryidae","Entomobryinae","Entomobrya","Entomobrya unostrigata","5","5","5","13","0","0","99.56","449.2","0","450","38.89","0"
"AGAW1","10E","NIN","BOLD:AAF9234","Arthropoda","Arachnida","Trombidiformes","Eupodidae","","","","3","3","3","14","0","0","99.78","447","0","450","29.93","0"
"AGAW1","10E","NIN","BOLD:AAA5267","Arthropoda","Collembola","Entomobryomorpha","Isotomidae","","","","2","2","2","15","0","0","99.02","407.5","0","413","37.41","0"
"AGAW1","10E","NIN","BOLD:AAG5658","Arthropoda","Arachnida","Araneae","Mimetidae","","Mimetus","Mimetus epeiroides","1","1","1","16","0","0","97.32","448","0","450","34.89","0"
"AGAW1","10N","BAC","TAX:91391","Proteobacteria","Alphaproteobacteria","Rickettsiales","",NA,"","","13","9","9","1","30.77","0","98.68","418.92","0","450","40.55","0"
"AGAW1","10N","BAC","TAX:175768","Proteobacteria","Alphaproteobacteria","Rickettsiales","Rickettsiaceae",NA,"Wolbachia","Wolbachia pipientis","6","6","6","2","0","0","98.33","441.67","0","443.17","39.63","0"
"AGAW1","10N","BAC","TAX:703648","Proteobacteria","Alphaproteobacteria","Rickettsiales","Rickettsiaceae",NA,"Wolbachia","Wolbachia endosymbiont","5","5","5","3","0","0","97.52","395.4","0","447.2","39.85","0"
"AGAW1","10N","BAC","TAX:91393","Proteobacteria","Alphaproteobacteria","Rickettsiales","Rickettsiaceae",NA,"Wolbachia","","4","4","4","4","0","0","99.11","391.5","0","418.25","39.49","0"
"AGAW1","10N","CHO","no match","no match","no match","no match","no match","no match","no match","no match","no match","no match","no match","no match","no match","no match","no match","no match","no match","no match","no match","no match"
"AGAW1","10N","INS","BOLD:AAP8739","Arthropoda","Insecta","Diptera","Platystomatidae","Platystomatinae","Rivellia","Rivellia steyskali","16463","11025","11025","1","33.03","0","99.25","441.61","0","443.35","38.06","0"
"AGAW1","10N","INS","BOLD:AAU6681","Arthropoda","Insecta","Diptera","Tipulidae","Tipulinae","Nephrotoma","Nephrotoma cornicina","12979","9358","9358","2","27.9","0","99.61","442.97","0","444.14","36.33","0"
"AGAW1","10N","INS","BOLD:AAA3453","Arthropoda","Insecta","Diptera","Anthomyiidae","Anthomyiinae","Delia","","11304","9655","9655","3","14.59","0","99.4","431.65","0","434.24","34.65","0"
"AGAW1","10N","INS","BOLD:AAC0903","Arthropoda","Insecta","Diptera","Chironomidae","Chironominae","Chironomus","Chironomus acidophilus","7851","5644","5644","4","28.11","0","99.54","439.3","0","441.54","36.32","0"
"AGAW1","10N","INS","BOLD:AAF9035","Arthropoda","Insecta","Diptera","Tipulidae","Tipulinae","Nephrotoma","Nephrotoma alterna","4984","4381","4381","5","12.1","0","98.81","426.7","0","428.98","33.57","0"
"AGAW1","10N","INS","BOLD:AAG2511","Arthropoda","Insecta","Diptera","Anthomyiidae","Anthomyiinae","Delia","Delia platura","2922","2520","2520","6","13.76","0","99.01","444.5","0","447.39","33.17","0"
"AGAW1","10N","INS","BOLD:ACE0699","Arthropoda","Insecta","Diptera","Chironomidae","","","","2689","1998","1998","7","25.7","0","98.47","425.06","0","441.14","36.13","0"
"AGAW1","10N","INS","BOLD:AAN6154","Arthropoda","Insecta","Coleoptera","Latridiidae","Corticariinae","Melanophthalma","Melanophthalma inermis","2333","1905","1905","8","18.35","0","99.19","436.06","0","437.65","31.21","0"
"AGAW1","10N","INS","BOLD:AAH3661","Arthropoda","Insecta","Diptera","Cecidomyiidae","","","","2278","1923","1923","9","15.58","0","99.27","433.35","0","435.14","25.44","0"
"AGAW1","10N","INS","BOLD:ACE7444","Arthropoda","Insecta","Hemiptera","Miridae","Mirinae, Phylinae","Adelphocoris, Megalocoleus","Adelphocoris lineolatus, Megalocoleus molliculus","1600","1032","1032","10","35.5","0","99.71","443.15","0","443.84","34.84","0"
"AGAW1","10N","INS","BOLD:AAB4657","Arthropoda","Insecta","Diptera","Chironomidae","Chironominae","Chironomus","Chironomus maturus","1580","1246","1246","11","21.14","0","99.16","438.65","0","441.17","37.24","0"
"AGAW1","10N","INS","BOLD:AAA7661","Arthropoda","Insecta","Diptera","Culicidae","Culicinae","Culiseta, Culex","Culiseta melanura, Culex articularis, Culex restuans, Culex acharistus","1195","1092","1092","12","8.62","0","99.33","434.93","0","437.01","32.84","0"
"AGAW1","10N","INS","BOLD:AAL4232","Arthropoda","Insecta","Diptera","Agromyzidae","Phytomyzinae","Phytoliriomyza","Phytoliriomyza arctica","959","742","742","13","22.63","0","99.53","441.29","0","442.6","32.09","0"
"AGAW1","10N","INS","BOLD:AAN5199","Arthropoda","Insecta","Diptera","Cecidomyiidae","","","CecidInt20 sp. BOLD:AAN5199","881","677","677","14","23.16","0","99.45","439.94","0","445.83","26.6","0"
"AGAW1","10N","INS","BOLD:ABW7897","Arthropoda","Insecta","Diptera","Cecidomyiidae","","","","848","616","616","15","27.36","0","99.57","444.61","0","446.78","29.73","0"
"AGAW1","10N","INS","BOLD:AAG2898","Arthropoda","Insecta","Hemiptera","Cicadellidae","Deltocephalinae","Paraphlepsius","Paraphlepsius apertus, Paraphlepsius irroratus","778","609","609","16","21.72","0","99.61","442.78","0","443.65","31.91","0"
"AGAW1","10N","INS","BOLD:ABW1379","Arthropoda","Insecta","Diptera","Chloropidae","Oscinellinae","Malloewia","Malloewia abdominalis","776","580","580","17","25.26","0","98.36","442.89","0","446.95","31.53","0"
"AGAW1","10N","INS","BOLD:ABA0863","Arthropoda","Insecta","Diptera","Cecidomyiidae","","","","659","528","528","18","19.88","0","99.32","441.36","0","442.59","29.87","0"
"AGAW1","10N","INS","BOLD:AAG2123","Arthropoda","Insecta","Diptera","Tachinidae","Dexiinae","Dinera","Dinera grisescens","631","496","496","19","21.39","0","99.33","435.15","0","436.72","31.92","0"
"AGAW1","10N","INS","BOLD:AAG7276","Arthropoda","Insecta","Diptera","Sphaeroceridae","Limosininae","Leptocera","Leptocera erythrocera","604","561","561","20","7.12","0","99.01","427.18","0","430.6","32.96","0"
"AGAW1","10N","INS","BOLD:AAC4197","Arthropoda","Insecta","Diptera","Chironomidae","Orthocladiinae","Paraphaenocladius","Paraphaenocladius impensus","564","514","514","21","8.87","0","99.2","428.03","0","431.32","34.83","0"
"AGAW1","10N","INS","BOLD:ABA0822","Arthropoda","Insecta","Diptera","Cecidomyiidae","","","","550","514","514","22","6.55","0","99.08","426.16","0","427.36","25.91","0"
"AGAW1","10N","INS","BOLD:AAG9897","Arthropoda","Insecta","Dermaptera","Forficulidae","Forficulinae","Forficula","Forficula auricularia-A","509","501","501","23","1.57","0","99.08","393.24","0","403.09","45.7","0"
"AGAW1","10N","INS","BOLD:AAN5219","Arthropoda","Insecta","Diptera","Cecidomyiidae","","","","431","356","356","24","17.4","0","99.5","438.32","0","439.52","29.03","0"
"AGAW1","10N","INS","BOLD:AAH2753","Arthropoda","Insecta","Coleoptera","Anthicidae","Anthicinae","Anthicus","Anthicus cervinus","413","335","335","25","18.89","0","98.12","424.73","0","438.91","33.9","0"
"AGAW1","10N","INS","BOLD:AAH0134","Arthropoda","Insecta","Coleoptera","Phalacridae","Phalacrinae","Stilbus","Stilbus apicialis","396","352","352","26","11.11","0","99.28","429.11","0","430.3","31.97","0"
"AGAW1","10N","INS","BOLD:AAG5507","Arthropoda","Insecta","Diptera","Chironomidae","Chironominae","Chironomus","Chironomus atrella","338","330","330","27","2.37","0","98.83","431.26","0","434.85","36.26","0"
"AGAW1","10N","INS","BOLD:AAP9022","Arthropoda","Insecta","Diptera","Cecidomyiidae","","","","337","279","279","28","17.21","0","99.53","440.27","0","441.58","27.55","0"
"AGAW1","10N","INS","BOLD:AAB1098","Arthropoda","Insecta","Diptera","Culicidae","Culicinae","Ochlerotatus, Aedes","Aedes excrucians, Aedes cantans, Aedes stimulans, Aedes euedes, Aedes riparius, Aedes annulipes, Ochlerotatus cantans, Aedes fitchii, Aedes sp., Ochlerotatus annulipes, Ochlerotatus excrucians","322","286","286","29","11.18","0","98.73","432.77","0","434.17","35.33","0"
"AGAW1","10N","INS","BOLD:AAH4171","Arthropoda","Insecta","Diptera","Chloropidae","Oscinellinae","Apallates","Apallates montanus, Apallates particeps","313","276","276","30","11.82","0","99.12","436.58","0","438.23","32.79","0"
"AGAW1","10N","INS","BOLD:AAN5242","Arthropoda","Insecta","Diptera","Cecidomyiidae","","","","259","258","258","31","0.39","0","98.57","412.37","0","414.87","28.25","0"
"AGAW1","10N","INS","BOLD:AAA4893","Arthropoda","Insecta","Lepidoptera","Crambidae","Crambinae","Microcrambus","Microcrambus elegans","223","193","193","32","13.45","0","99.08","413.61","0","438.13","32.75","0"
"AGAW1","10N","INS","BOLD:AAU8713","Arthropoda","Insecta","Hymenoptera","Dryinidae","Anteoninae","Anteon","Anteon tripartitum, Anteon gaullei, Anteon cameroni, Anteon pubicorne","220","218","218","33","0.91","0","99","400.04","0","402.91","27.78","0"
"AGAW1","10N","INS","BOLD:ACF8552","Arthropoda","Insecta","Coleoptera","Anthicidae","Anthicinae","Malporus","Malporus formicarius","190","172","172","34","9.47","0","99.57","364.81","0","430.24","34.58","0"
"AGAW1","10N","INS","BOLD:AAM6248","Arthropoda","Insecta","Diptera","Chironomidae","Chironominae","Microtendipes","Microtendipes pedellus grp.","181","161","161","35","11.05","0","98.72","428.83","0","430.44","31.38","0"
"AGAW1","10N","INS","BOLD:AAL7819","Arthropoda","Insecta","Diptera","Psychodidae","Psychodinae","Psychoda","Psychoda sp., Psychoda sp. BOLD:AAL7819","175","166","166","36","5.14","0","99.44","420.3","0","421.87","32.66","0"
"AGAW1","10N","INS","BOLD:ABV9018","Arthropoda","Insecta","Diptera","Mycetophilidae","Sciophilinae","Sciophila","","171","166","166","37","2.92","0","99.15","435.06","0","437.08","32.04","0"
"AGAW1","10N","INS","BOLD:AAN5205","Arthropoda","Insecta","Diptera","Cecidomyiidae","","","CecidInt42 sp. BOLD:AAN5205","171","149","149","37","12.87","0","99.4","443.56","0","445.08","31.37","0"
"AGAW1","10N","INS","BOLD:AAZ5592","Arthropoda","Insecta","Diptera","Psychodidae","Psychodinae","","","170","162","162","39","4.71","0","99.17","432.44","0","435.11","32.49","0"
"AGAW1","10N","INS","BOLD:AAF7193","Arthropoda","Insecta","Diptera","Cecidomyiidae","Cecidomyiinae","Aphidoletes, Dasineura","Aphidoletes aphidimyza, Dasineura leguminicola, Dasineura sp. BOLD:AAF7193","168","147","147","40","12.5","0","99.48","440.57","0","442.21","26.66","0"
"AGAW1","10N","INS","BOLD:AAH3664","Arthropoda","Insecta","Diptera","Cecidomyiidae","","","CecidInt3 sp. BOLD:AAH3664","155","150","150","41","3.23","0","99.14","431.37","0","435.18","26.98","0"
"AGAW1","10N","INS","BOLD:ACU6062","Arthropoda","Insecta","Diptera","Cecidomyiidae","","","","144","121","121","42","15.97","0","98.97","442.91","0","444.5","26.66","0"
"AGAW1","10N","INS","BOLD:AAH0135","Arthropoda","Insecta","Coleoptera","Phalacridae","Phalacrinae","Acylomus","Acylomus pugetanus, Acylomus ergoti","140","130","130","43","7.14","0","99.34","435.29","0","437.1","31.61","0"
"AGAW1","10N","INS","BOLD:AAM9013","Arthropoda","Insecta","Diptera","Keroplatidae","Macrocerinae","Macrocera","Macrocera pusilla","137","133","133","44","2.92","0","99.3","434.07","0","435.65","30.33","0"
"AGAW1","10N","INS","BOLD:ACK3431","Arthropoda","Insecta","Diptera","Cecidomyiidae","","","","123","113","113","45","8.13","0","99.33","427.24","0","429.34","32.52","0"
"AGAW1","10N","INS","BOLD:AAN5148","Arthropoda","Insecta","Diptera","Ceratopogonidae","Forcipomyiinae","Forcipomyia","Forcipomyia bipunctata, Forcipomyia sp. CK-2012","121","105","105","46","13.22","0","97.52","436.02","0","437.62","35.42","0"
"AGAW1","10N","INS","BOLD:AAA1513","Arthropoda","Insecta","Lepidoptera","Plutellidae","Plutellinae","Plutella, Rhigognostis","Plutella xylostella, Rhigognostis sp.","113","102","102","47","9.73","0","99.02","443.34","0","446.1","32.38","0"
"AGAW1","10N","INS","BOLD:AAG2868","Arthropoda","Insecta","Hemiptera","Cicadellidae","Typhlocybinae","Empoasca","Empoasca sp., Empoasca fabae","109","83","83","48","23.85","0","99.64","445.77","0","448.22","32.39","0"
"AGAW1","10N","INS","BOLD:ABV1297","Arthropoda","Insecta","Diptera","Sciaridae","","","","108","108","108","49","0","0","98.92","433.43","0","436.12","30.25","0"
"AGAW1","10N","INS","BOLD:ACJ6954","Arthropoda","Insecta","Diptera","Cecidomyiidae","","","","102","91","91","50","10.78","0","99.6","337.96","0","426.23","26.56","0"
"AGAW1","10N","INS","BOLD:AAN5255","Arthropoda","Insecta","Diptera","Cecidomyiidae","Cecidomyiinae","Dasineura","","88","74","74","51","15.91","0","98.39","442.66","0","446.18","26.84","0"
"AGAW1","10N","INS","BOLD:ABZ0621","Arthropoda","Insecta","Diptera","Psychodidae","Psychodinae","","","80","79","79","52","1.25","0","99.11","413.61","0","417.01","32.54","0"
"AGAW1","10N","INS","BOLD:ACE5778","Arthropoda","Insecta","Diptera","Dolichopodidae","","","","75","74","74","53","1.33","0","98.9","432.41","0","433.92","38.35","0"
"AGAW1","10N","INS","BOLD:AAN5220","Arthropoda","Insecta","Diptera","Cecidomyiidae","","","","69","69","69","54","0","0","98.06","419.57","0","430.62","26.22","0"
"AGAW1","10N","INS","BOLD:AAD1485","Arthropoda","Insecta","Diptera","Chironomidae","Chironominae","Paratanytarsus","Paratanytarsus grimmii, Paratanytarsus sp.","67","60","60","55","10.45","0","99.64","443.61","0","445.12","33.83","0"
"AGAW1","10N","INS","BOLD:AAI2606","Arthropoda","Insecta","Diptera","Chironomidae","Chironominae","Paratanytarsus","","65","59","59","56","9.23","0","99.27","435.95","0","437.37","33.92","0"
"AGAW1","10N","INS","BOLD:AAF9771","Arthropoda","Insecta","Diptera","Hybotidae","Tachydromiinae","Platypalpus","Platypalpus longiseta, Platypalpus trivialis, Platypalpus sp. 5, Platypalpus sp. 17, Platypalpus pallidiventris","65","57","57","56","12.31","0","98.93","444.52","0","445.22","35.07","0"
"AGAW1","10N","INS","BOLD:ADF6661","Arthropoda","Insecta","Diptera","Cecidomyiidae","","","","64","55","55","58","14.06","0","98.94","416.53","0","444.06","29.82","0"
"AGAW1","10N","INS","BOLD:AAN5183","Arthropoda","Insecta","Diptera","Cecidomyiidae","","","","64","57","57","58","10.94","0","98.35","443.86","0","445.14","27.7","0"
"AGAW1","10N","INS","BOLD:AAG6468","Arthropoda","Insecta","Diptera","Ceratopogonidae","Ceratopogoninae","Culicoides","Culicoides biguttatus","60","59","59","60","1.67","0","98.98","447.03","0","448.58","37.28","0"
"AGAW1","10N","INS","BOLD:AAN5221","Arthropoda","Insecta","Diptera","Cecidomyiidae","","","","60","60","60","60","0","0","98.7","416.72","0","419.85","25.31","0"
"AGAW1","10N","INS","BOLD:AAG4477","Arthropoda","Insecta","Coleoptera","Chrysomelidae","Chrysomelinae","Plagiodera","Plagiodera versicolora","58","57","57","62","1.72","0","99.24","427.29","0","429.07","34.57","0"
"AGAW1","10N","INS","BOLD:ADI0927","","","","","","","","54","45","45","63","16.67","0","97.67","422.11","0","423.35","34.18","0"
"AGAW1","10N","INS","BOLD:AAG8320","Arthropoda","Insecta","Hymenoptera","Crabronidae","Astatinae","Astata","Astata leuthstromi","53","48","48","64","9.43","0","99.33","427.21","0","444.38","28.18","0"
"AGAW1","10N","INS","BOLD:AAG3154","Arthropoda","Insecta","Diptera","Psychodidae","Psychodinae","Psychoda","","53","46","46","64","13.21","0","98.02","445.58","0","447.21","32.7","0"
"AGAW1","10N","INS","BOLD:ACV5892","Arthropoda","Insecta","Diptera","Cecidomyiidae","","","","51","51","51","66","0","0","98.3","435.71","0","437.37","24.73","0"
"AGAW1","10N","INS","BOLD:ACK7465","Arthropoda","Insecta","Psocodea","Liposcelidae","","","","49","49","49","67","0","0","98.54","386.31","0","399.67","36.57","0"
"AGAW1","10N","INS","BOLD:AAN5238","Arthropoda","Insecta","Diptera","Cecidomyiidae","","","","45","44","44","68","2.22","0","99.45","434.27","0","436.11","25.62","0"
"AGAW1","10N","INS","BOLD:AAG3236","Arthropoda","Insecta","Diptera","Phoridae","Phorinae","Conicera, Diplonevra","Diplonevra nitidula","40","39","39","69","2.5","0","99.44","442.4","0","444.03","32.12","0"
"AGAW1","10N","INS","BOLD:AAY6730","Arthropoda","Insecta","Hemiptera","Derbidae","Cedusinae","Cedusa","Cedusa incisa","36","34","34","70","5.56","0","98.86","427.78","0","429.08","35.44","0"
"AGAW1","10N","INS","BOLD:AAM9255","Arthropoda","Insecta","Diptera","Sciaridae","","Pseudolycoriella","","28","28","28","71","0","0","99.34","407.79","0","410.04","30.88","0"
"AGAW1","10N","INS","BOLD:ACF0915","Arthropoda","Insecta","Diptera","Chironomidae","Orthocladiinae","","","24","24","24","72","0","0","98.91","399.58","0","409.54","33.36","0"
"AGAW1","10N","INS","BOLD:AAM6037","Arthropoda","Insecta","Diptera","Cecidomyiidae","Cecidomyiinae","Contarinia, Dasineura","","24","22","22","72","8.33","0","98.88","438.42","0","439.88","26.1","0"
"AGAW1","10N","INS","BOLD:ACA3113","Arthropoda","Insecta","Psocodea","Caeciliusidae","","","","23","23","23","74","0","0","98.72","386.09","0","390.65","37.14","0"
"AGAW1","10N","INS","BOLD:AAN5260","Arthropoda","Insecta","Diptera","Cecidomyiidae","","","","23","23","23","74","0","0","99.26","413.65","0","415.13","26.17","0"
"AGAW1","10N","INS","BOLD:ABX0893","Arthropoda","Insecta","Diptera","Cecidomyiidae","","","","23","22","22","74","4.35","0","97.39","445.22","0","446.57","26.54","0"
"AGAW1","10N","INS","BOLD:ACX6055","Arthropoda","Insecta","Diptera","Phoridae","Metopininae","Megaselia","Megaselia sp. AK-2010-2, Megaselia sp. BOLD:ACX6055","19","16","16","77","15.79","0","99.33","446.37","0","447.95","33.28","0"
"AGAW1","10N","INS","BOLD:AAM6040","Arthropoda","Insecta","Diptera","Cecidomyiidae","","","","18","15","15","78","16.67","0","99.21","443.89","0","445","27.1","0"
"AGAW1","10N","INS","BOLD:ABW2870","Arthropoda","Insecta","Coleoptera","Staphylinidae","Aleocharinae","Meronera","Meronera venustula","18","18","18","78","0","0","99.26","434.83","0","437.39","35.17","0"
"AGAW1","10N","INS","BOLD:ACU7906","Arthropoda","Insecta","Diptera","Chironomidae","Orthocladiinae","","","17","17","17","80","0","0","99","392.12","0","396.12","36.04","0"
"AGAW1","10N","INS","BOLD:ABA1215","Arthropoda","Insecta","Diptera","Sciaridae","","Lycoriella","Lycoriella sativae","15","15","15","81","0","0","99.38","423.93","0","426.07","35.35","0"
"AGAW1","10N","INS","BOLD:AAC6157","Arthropoda","Insecta","Lepidoptera","Pyralidae","Phycitinae","Ephestia","Ephestia elutella","15","14","14","81","6.67","0","99.66","431.07","0","432.2","32.8","0"
"AGAW1","10N","INS","BOLD:ABU5525","Arthropoda","Insecta","Diptera","Chironomidae","Orthocladiinae","Limnophyes","Limnophyes sp. 14ES","15","15","15","81","0","0","99","402.47","0","404.93","37.05","0"
"AGAW1","10N","INS","BOLD:AAZ0271","Arthropoda","Insecta","Diptera","Ceratopogonidae","Ceratopogoninae","Culicoides","","15","15","15","81","0","0","98.64","436.67","0","439.93","41.58","0"
"AGAW1","10N","INS","BOLD:AAN5264","Arthropoda","Insecta","Diptera","Cecidomyiidae","","","","13","13","13","85","0","0","99.11","429.08","0","430.31","27.38","0"
"AGAW1","10N","INS","BOLD:AAG1711","Arthropoda","Insecta","Diptera","Muscidae","Phaoniinae","Helina","Helina depuncta","12","12","12","86","0","0","99.4","436.83","0","438.83","31.26","0"
"AGAW1","10N","INS","BOLD:ACN7725","Arthropoda","Insecta","Hymenoptera","Platygastridae","","","","12","12","12","86","0","0","99.35","392.83","0","429","26.83","0"
"AGAW1","10N","INS","BOLD:AAG8091","Arthropoda","Insecta","Hymenoptera","Ichneumonidae","Campopleginae","Campoletis","Campoletis sp. AB.2","12","12","12","86","0","0","99.27","436.08","0","438","24.59","0"
"AGAW1","10N","INS","BOLD:AAF1729","Arthropoda","Insecta","Psocodea","Lachesillidae","Lachesillinae","Lachesilla","Lachesilla pedicularia","12","8","8","86","33.33","0","99.13","433.92","0","435","36.87","0"
"AGAW1","10N","INS","BOLD:AAN5249","Arthropoda","Insecta","Diptera","Cecidomyiidae","","","","11","11","11","90","0","0","99.43","439.64","0","440.64","33.19","0"
"AGAW1","10N","INS","BOLD:AAH0256","Arthropoda","Insecta","Coleoptera","Latridiidae","Corticariinae","Corticaria, Corticarina","","11","9","9","90","18.18","0","99.26","442","0","443","31.6","0"
"AGAW1","10N","INS","BOLD:AAG7891","Arthropoda","Insecta","Hymenoptera","Platygastridae","Scelioninae","Telenomus","Telenomus podisi","10","10","10","92","0","0","99.16","439.9","0","448.1","27.32","0"
"AGAW1","10N","INS","BOLD:AAB1850","Arthropoda","Insecta","Hemiptera","Aphrophoridae","Aphrophorinae","Philaenus","Philaenus tesselatus, Philaenus spumarius, Philaenus maghresignus","10","10","10","92","0","0","99.39","441.2","0","442.2","33.28","0"
"AGAW1","10N","INS","BOLD:AAA5803","Arthropoda","Insecta","Hemiptera","Miridae","Mirinae","Lygus","Lygus lineolaris","10","9","9","92","10","0","98.54","447.2","0","449.4","33.98","0"
"AGAW1","10N","INS","BOLD:ABZ5071","Arthropoda","Insecta","Diptera","Chironomidae","Chironominae","Micropsectra","Micropsectra nigripila","9","9","9","95","0","0","98.08","417.33","0","431.78","36.76","0"
"AGAW1","10N","INS","BOLD:AAH3662","Arthropoda","Insecta","Diptera","Cecidomyiidae","","","CecidInt3 sp. BOLD:AAH3662","9","9","9","95","0","0","99.25","439.67","0","440.89","26.26","0"
"AGAW1","10N","INS","BOLD:ACC7779","Arthropoda","Insecta","Coleoptera","Elateridae","Cardiophorinae","Cardiophorus","Cardiophorus convexus","8","8","8","97","0","0","99.59","435","0","435.63","39.78","0"
"AGAW1","10N","INS","BOLD:AAU7170","Arthropoda","Insecta","Coleoptera","Cryptophagidae","Atomariinae","Atomaria","Atomaria wollastoni","7","7","7","98","0","0","98.78","434.71","0","436.14","39.29","0"
"AGAW1","10N","INS","BOLD:ACV5891","Arthropoda","Insecta","Diptera","Cecidomyiidae","","","","7","7","7","98","0","0","99.01","361.57","0","420.86","24.44","0"
"AGAW1","10N","INS","BOLD:AAN5342","Arthropoda","Insecta","Diptera","Chironomidae","Orthocladiinae","Heleniella","","7","7","7","98","0","0","98.01","400.57","0","416.71","34.85","0"
"AGAW1","10N","INS","BOLD:AAN5267","Arthropoda","Insecta","Diptera","Cecidomyiidae","","","","7","5","5","98","28.57","0","99.3","449.43","0","450","30.63","0"
"AGAW1","10N","INS","BOLD:AAF4817","Arthropoda","Insecta","Diptera","Chironomidae","Orthocladiinae","Smittia","Smittia edwardsi","7","6","6","98","14.29","0","98.14","391","0","450","33.08","0"
"AGAW1","10N","INS","BOLD:AAA3867","Arthropoda","Insecta","Lepidoptera","Noctuidae","Noctuinae","Apamea","Apamea amputatrix","6","6","6","103","0","0","98.82","433","0","434.83","33.78","0"
"AGAW1","10N","INS","BOLD:AAF9789","Arthropoda","Insecta","Diptera","Hybotidae","","","","6","6","6","103","0","0","99.08","403.67","0","404.5","33","0"
"AGAW1","10N","INS","BOLD:ADB2738","Arthropoda","Insecta","Psocodea","Liposcelidae","","","","6","6","6","103","0","0","97.24","446.83","0","450","36.33","0"
"AGAW1","10N","INS","BOLD:ABV1337","Arthropoda","Insecta","Diptera","Cecidomyiidae","","","","6","5","5","103","16.67","0","99","449","0","450","28.59","0"
"AGAW1","10N","INS","BOLD:ACA1917","Arthropoda","Insecta","Diptera","Cecidomyiidae","","","","5","5","5","107","0","0","98.66","448.8","0","450","26.18","0"
"AGAW1","10N","INS","BOLD:AAB0373","Arthropoda","Insecta","Neuroptera","Chrysopidae","Chrysopinae","Chrysoperla","Chrysoperla sp. isolate 82, Chrysoperla zastrowi, Chrysoperla plorabunda, Chrysoperla sp. isolate 71, Chrysoperla agilis, Italochrysa italica, Chrysoperla sp. isolate 88, Chrysoperla lucasina, Chrysoperla carnea, Chrysoperla mediterranea, Chrysoperla pallida, Chrysoperla nipponensis, Chrysoperla sillemi","5","5","5","107","0","0","99.36","433.2","0","434","31.04","0"
"AGAW1","10N","INS","BOLD:AAG6745","Arthropoda","Insecta","Diptera","Calliphoridae","Calliphorinae","Pollenia","Pollenia pediculata","5","5","5","107","0","0","99.31","426.2","0","427.2","33.49","0"
"AGAW1","10N","INS","BOLD:ABV1289","Arthropoda","Insecta","Diptera","Cecidomyiidae","","","","4","4","4","110","0","0","99","448.75","0","450","25.61","0"
"AGAW1","10N","INS","BOLD:ABA7871","Arthropoda","Insecta","Diptera","Cecidomyiidae","","","","4","4","4","110","0","0","98.99","432.5","0","433.75","26.78","0"
"AGAW1","10N","INS","BOLD:AAV5400","Arthropoda","Insecta","Diptera","Cecidomyiidae","","","","4","4","4","110","0","0","97.88","413","0","415","29.62","0"
"AGAW1","10N","INS","BOLD:AAF9187","Arthropoda","Insecta","Coleoptera","Curculionidae","Entiminae","Phyllobius","Phyllobius oblongus","4","4","4","110","0","0","99.27","449","0","450","39","0"
"AGAW1","10N","INS","BOLD:ACV5271","Arthropoda","Insecta","Diptera","Cecidomyiidae","","","","4","4","4","110","0","0","99.78","448.5","0","450","28.33","0"
"AGAW1","10N","INS","BOLD:ABY5318","Arthropoda","Insecta","Diptera","Chloropidae","Oscinellinae","Incertella, Rhopalopterum","Rhopalopterum umbrosum, Rhopalopterum luteiceps, Incertella bispina, Rhopalopterum painteri","4","4","4","110","0","0","99.11","450","0","450","32.44","0"
"AGAW1","10N","INS","BOLD:ADF3937","","","","","","","","4","4","4","110","0","0","98.85","411.75","0","429","32.12","0"
"AGAW1","10N","INS","BOLD:ABV1422","Arthropoda","Insecta","Diptera","Cecidomyiidae","","","","4","4","4","110","0","0","99.12","427.5","0","428","26.84","0"
"AGAW1","10N","INS","BOLD:AAG2740","Arthropoda","Insecta","Diptera","Ephydridae","Parydrinae","Philygria","Philygria debilis","4","4","4","110","0","0","97.68","415","0","417.75","32.96","0"
"AGAW1","10N","INS","BOLD:ACV1014","Arthropoda","Insecta","Diptera","Cecidomyiidae","","","","3","3","3","119","0","0","97.66","396","0","399.67","25.21","0"
"AGAW1","10N","INS","BOLD:ABA8103","Arthropoda","Insecta","Hemiptera","Cicadellidae","Typhlocybinae","Empoasca","","3","3","3","119","0","0","97.78","393.33","0","396.67","32.17","0"
"AGAW1","10N","INS","BOLD:AAN8408","Arthropoda","Insecta","Hemiptera","Aphididae","Anoeciinae","Anoecia","","3","3","3","119","0","0","98.41","441","0","443","29.05","0"
"AGAW1","10N","INS","BOLD:AAI4303","Arthropoda","Insecta","Diptera","Chironomidae","Chironominae","Chironomus","Chironomus melanescens","3","3","3","119","0","0","97.93","436.33","0","439","34.3","0"
"AGAW1","10N","INS","BOLD:AAN6447","Arthropoda","Insecta","Diptera","Sciaridae","","Corynoptera","Corynoptera perpusilla","3","3","3","119","0","0","99.63","447.33","0","450","36.22","0"
"AGAW1","10N","INS","BOLD:AAA7631","Arthropoda","Insecta","Hemiptera","Miridae","Mirinae","Adelphocoris","Adelphocoris lineolatus","3","3","3","119","0","0","99.4","440.33","0","441.67","35.39","0"
"AGAW1","10N","INS","BOLD:ABW8023","Arthropoda","Insecta","Diptera","Cecidomyiidae","","","","3","3","3","119","0","0","98.07","448.33","0","449","27.69","0"
"AGAW1","10N","INS","BOLD:ABX2489","Arthropoda","Insecta","Coleoptera","Phalacridae","Phalacrinae","Stilbus","","3","3","3","119","0","0","97.65","412","0","413","31.31","0"
"AGAW1","10N","INS","BOLD:AAA9246","Arthropoda","Insecta","Lepidoptera","Tortricidae","Tortricinae","Clepsis","Clepsis clemensiana","2","2","2","127","0","0","99.64","433","0","433","29.89","0"
"AGAW1","10N","INS","BOLD:AAG9668","Arthropoda","Insecta","Diptera","Dolichopodidae","Diaphorinae","Chrysotus","Chrysotus sp., Chrysotus sp. nr. obliquus","2","2","2","127","0","0","98.3","443.5","0","444","34.14","0"
"AGAW1","10N","INS","BOLD:ACJ8771","Arthropoda","Insecta","Diptera","Chironomidae","Chironominae","Sublettea","","2","2","2","127","0","0","99.61","389","0","445","32.92","0"
"AGAW1","10N","INS","BOLD:AAY6412","Arthropoda","Insecta","Diptera","Sciaridae","","Bradysia","Bradysia urticae","2","2","2","127","0","0","98.33","446.5","0","450","31.89","0"
"AGAW1","10N","INS","BOLD:AAA5299","Arthropoda","Insecta","Diptera","Chironomidae","Orthocladiinae","Cricotopus","Cricotopus laetus, Cricotopus relucens, Cricotopus trifasciatus, Cricotopus sp., Cricotopus sp. 3ES, Cricotopus sylvestris, Cricotopus glacialis","2","2","2","127","0","0","98.83","436.5","0","439","34.51","0"
"AGAW1","10N","INS","BOLD:AAD2041","Arthropoda","Insecta","Diptera","Dolichopodidae","Sciapodinae","Condylostylus","Condylostylus inermis, Condylostylus nigrofemoratus, Condylostylus connectans, Condylostylus caudatus, Condylostylus flavipes","2","2","2","127","0","0","99.2","413.5","0","415","35.32","0"
"AGAW1","10N","INS","BOLD:AAA2815","Arthropoda","Insecta","Lepidoptera","Noctuidae","Noctuinae","Mythimna","Mythimna yukonensis, Mythimna oxygala","2","2","2","127","0","0","98.89","428.5","0","429.5","33.16","0"
"AGAW1","10N","INS","BOLD:AAA4277","Arthropoda","Insecta","Diptera","Syrphidae","Syrphinae","Toxomerus","Toxomerus marginatus, Toxomerus sp. CR-3, Toxomerus CR-3","2","2","2","127","0","0","99.44","449","0","450","30.78","0"
"AGAW1","10N","INS","BOLD:ACJ0111","Arthropoda","Insecta","Hymenoptera","Platygastridae","Scelioninae","Telenomus","","2","2","2","127","0","0","98.52","439.5","0","450","28.78","0"
"AGAW1","10N","INS","BOLD:ADH9867","Arthropoda","Insecta","Diptera","","","","","2","2","2","127","0","0","98.45","383.5","0","388","37.36","0"
"AGAW1","10N","INS","BOLD:AAE0845","Arthropoda","Insecta","Hemiptera","Pentatomidae","Pentatominae","Euschistus","Euschistus servus","2","2","2","127","0","0","98.66","449","0","450","37.78","0"
"AGAW1","10N","INS","BOLD:ACX6289","Arthropoda","Insecta","Diptera","Phoridae","Metopininae","Megaselia","","2","2","2","127","0","0","99.03","430.5","0","430.5","38.08","0"
"AGAW1","10N","INS","BOLD:AAE0456","Arthropoda","Insecta","Dermaptera","Forficulidae","Forficulinae","Forficula","Forficula auricularia-B, Forficula auricularia","2","2","2","127","0","0","98.99","398","0","406","44.96","0"
"AGAW1","10N","INS","BOLD:AAE7016","Arthropoda","Insecta","Lepidoptera","Gelechiidae","Dichomeridinae","Helcystogramma","Helcystogramma hystricella","2","2","2","127","0","0","99","401.5","0","402.5","30.88","0"
"AGAW1","10N","INS","BOLD:ACD3588","Arthropoda","Insecta","Diptera","Sciaridae","","Lycoriella","Lycoriella sativae","2","2","2","127","0","0","99.41","418.5","0","445","37.08","0"
"AGAW1","10N","INS","BOLD:AAA3181","Arthropoda","Insecta","Diptera","Muscidae","Muscinae","Stomoxys","Stomoxys calcitrans","1","1","1","142","0","0","97.54","447","0","450","32.67","0"
"AGAW1","10N","INS","BOLD:AAC0022","Arthropoda","Insecta","Diptera","Chironomidae","Chironominae","Micropsectra","Micropsectra nigripila","1","1","1","142","0","0","97.33","449","0","450","36.89","0"
"AGAW1","10N","INS","BOLD:AAA9899","Arthropoda","Insecta","Hemiptera","Aphididae","Aphidinae","Rhopalosiphum","Rhopalosiphum padi","1","1","1","142","0","0","99.33","448","0","450","25.78","0"
"AGAW1","10N","INS","BOLD:AAG9661","Arthropoda","Insecta","Diptera","Dolichopodidae","","","","1","1","1","142","0","0","97.56","328","0","354","33.62","0"
"AGAW1","10N","INS","BOLD:AAN5497","Arthropoda","Insecta","Diptera","Anthomyiidae","Pegomyinae","Pegomya","Pegomya solennis, Pegomya bicolor","1","1","1","142","0","0","100","403","0","406","28.82","0"
"AGAW1","10N","INS","BOLD:ACA9275","Arthropoda","Insecta","Diptera","Chironomidae","Chironominae","Chironomus","","1","1","1","142","0","0","100","450","0","450","35.78","0"
"AGAW1","10N","INS","BOLD:AAC0635","Arthropoda","Insecta","Hemiptera","Miridae","Mirinae","Stenotus","Stenotus binotatus","1","1","1","142","0","0","97.33","450","0","450","34.44","0"
"AGAW1","10N","INS","BOLD:ACV2677","Arthropoda","Insecta","Diptera","Cecidomyiidae","","","","1","1","1","142","0","0","97.55","449","0","450","30","0"
"AGAW1","10N","INS","BOLD:AAA8108","Arthropoda","Insecta","Lepidoptera","Noctuidae","Acontiinae","Ponometia","Ponometia erastrioides","1","1","1","142","0","0","97.54","448","0","450","32.44","0"
"AGAW1","10N","INS","BOLD:ACI4417","Arthropoda","Insecta","Diptera","Cecidomyiidae","","","","1","1","1","142","0","0","97.62","378","0","450","30.67","0"
"AGAW1","10N","INS","BOLD:AAV1136","Arthropoda","Insecta","Diptera","Scatopsidae","","","","1","1","1","142","0","0","97.78","450","0","450","36.44","0"
"AGAW1","10N","INS","BOLD:AAM6286","Arthropoda","Insecta","Diptera","Chironomidae","Chironominae","","","1","1","1","142","0","0","99.78","447","0","450","35.11","0"
"AGAW1","10N","INS","BOLD:AAB5609","Arthropoda","Insecta","Diptera","Muscidae","Coenosiinae","Coenosia","Coenosia sp.BOLD:AAB5609, Coenosia tigrina","1","1","1","142","0","0","98.8","415","0","419","31.98","0"
"AGAW1","10N","INS","BOLD:ACF1257","","","","","","","","1","1","1","142","0","0","99.78","449","0","450","34.89","0"
"AGAW1","10N","INS","BOLD:AAH0131","Arthropoda","Insecta","Coleoptera","Nitidulidae","Cryptarchinae","Cryptarcha","Cryptarcha ampla","1","1","1","142","0","0","99.26","403","0","450","39.11","0"
"AGAW1","10N","INS","BOLD:AAV0259","Arthropoda","Insecta","Hemiptera","Cicadellidae","Deltocephalinae","Endria","Endria inimica","1","1","1","142","0","0","97.76","402","0","404","35.4","0"
"AGAW1","10N","INS","BOLD:AAD0484","Arthropoda","Insecta","Diptera","Chironomidae","","","","1","1","1","142","0","0","97.84","416","0","419","32.46","0"
"AGAW1","10N","INS","BOLD:ABU5546","Arthropoda","Insecta","Diptera","Psychodidae","Psychodinae","Psychoda","","1","1","1","142","0","0","97.55","449","0","450","31.78","0"
"AGAW1","10N","INS","BOLD:ACL7225","Arthropoda","Insecta","Diptera","Cecidomyiidae","","","","1","1","1","142","0","0","100","450","0","450","27.56","0"
"AGAW1","10N","INS","BOLD:AAM6295","Arthropoda","Insecta","Diptera","Chaoboridae","","","","1","1","1","142","0","0","100","450","0","450","37.33","0"
"AGAW1","10N","INS","BOLD:AAC0706","Arthropoda","Insecta","Diptera","Chironomidae","Chironominae","Dicrotendipes","Dicrotendipes tritomus","1","1","1","142","0","0","97.97","443","0","448","35.94","0"
"AGAW1","10N","INS","BOLD:AAG6753","Arthropoda","Insecta","Diptera","Lauxaniidae","Lauxaniinae","Camptoprosopella","","1","1","1","142","0","0","98.31","413","0","450","35.78","0"
"AGAW1","10N","INS","BOLD:AAG9626","Arthropoda","Insecta","Diptera","Dolichopodidae","","","","1","1","1","142","0","0","98.89","449","0","450","36.89","0"
"AGAW1","10N","INS","BOLD:AAP6882","Arthropoda","Insecta","Diptera","Chironomidae","Chironominae","Dicrotendipes","","1","1","1","142","0","0","97.78","450","0","450","36.22","0"
"AGAW1","10N","INS","BOLD:AAA4555","Arthropoda","Insecta","Orthoptera","Acrididae","Melanoplinae","Melanoplus","Melanoplus stupefactus, Melanoplus angustipennis, Melanoplus ponderosus, Melanoplus bruneri, Melanoplus frigidus, Melanoplus scapularis, Melanoplus cinereus, Melanoplus madeleineae, Melanoplus sanguinipes, Melanoplus nigrescens group, Melanoplus tunicae, Melanoplus walshi, Melanoplus femurrubrum, Melanoplus dawsoni, Melanoplus sp., Melanoplus fasciatus, Melanoplus bowditchi, Melanoplus stonei, Melanoplus bivittatus, Melanoplus huroni, Melanoplus thomasi, Melanoplus borealis, Melanoplus packardii","1","1","1","142","0","0","99.11","449","0","450","36.44","0"
"AGAW1","10N","INS","BOLD:ACE1335","Arthropoda","Insecta","Diptera","Hybotidae","Tachydromiinae","Platypalpus","Platypalpus sp. BOLD:ACE1335, Platypalpus calccutus, Platypalpus calceatus","1","1","1","142","0","0","99.33","449","0","450","35.56","0"
"AGAW1","10N","INS","BOLD:AAA7374","Arthropoda","Insecta","Diptera","Syrphidae","Syrphinae","Sphaerophoria","Sphaerophoria bankowskae, Sphaerophoria infuscata, Sphaerophoria nr. philanthus, Sphaerophoria batava, Sphaerophoria sp., Sphaerophoria rueppellii, Sphaerophoria interrupta-group, Sphaerophoria sp. MA-1, Sphaerophoria cleoae, Sphaerophoria asymmetrica, Sphaerophoria fatarum, Sphaerophoria bengalensis, Sphaerophoria bifurcata, Sphaerophoria taeniata, Sphaerophoria interrupta, Sphaerophoria contigua, Sphaerophoria kaa, Sphaerophoria laurae, Sphaerophoria chongjini, Sphaerophoria boreoalpina, Sphaerophoria sp. BOLDAAA7374, Sphaerophoria macrogaster, Sphaerophoria sulphuripes, Sphaerophoria brevipilosa, Sphaerophoria philanthus, Sphaerophoria pyrrhina, Sphaerophoria abbreviata, Sphaerophoria scripta, Sphaerophoria potentillae, Sphaerophoria longipilosa, Sphaerophoria virgata","1","1","1","142","0","0","99.74","384","0","387","29.97","0"
"AGAW1","10N","INS","BOLD:AAH4205","Arthropoda","Insecta","Diptera","Chloropidae","Chloropinae","Meromyza","Meromyza nigriseta, Meromyza sp.","1","1","1","142","0","0","99.55","448","0","450","32.67","0"
"AGAW1","10N","INS","BOLD:AAO3939","Arthropoda","Insecta","Diptera","Limoniidae","Limoniinae","Dicranomyia","","1","1","1","142","0","0","98.66","449","0","449","36.08","0"
"AGAW1","10N","INS","BOLD:AAA5953","Arthropoda","Insecta","Lepidoptera","Gelechiidae","Gelechiinae","Coleotechnites","Coleotechnites sp., Coleotechnites piceaella, Coleotechnites laricis, Coleotechnites sp. 3, Coleotechnites carbonaria, Coleotechnites granti, Coleotechnites macleodi, Coleotechnites gibsonella, Coleotechnites coniferella, Coleotechnites martini","1","1","1","142","0","0","98.67","450","0","450","34.44","0"
"AGAW1","10N","INS","BOLD:ACI7380","Arthropoda","Insecta","Diptera","Cecidomyiidae","","","","1","1","1","142","0","0","97.12","451","0","450","24.67","0"
"AGAW1","10N","NAR","no match","no match","no match","no match","no match","no match","no match","no match","no match","no match","no match","no match","no match","no match","no match","no match","no match","no match","no match","no match"
"AGAW1","10N","NIN","BOLD:AAU6252","Arthropoda","Arachnida","Trombidiformes","Bdellidae","","Cyta","","269","222","222","1","17.47","0","98.28","440.61","0","441.67","31.59","0"
"AGAW1","10N","NIN","BOLD:AAA7242","Arthropoda","Collembola","Entomobryomorpha","Entomobryidae","Seirinae, Entomobryinae","Entomobrya, Lepidosira","Entomobrya multifasciata, Lepidosira sp.","246","241","241","2","2.03","0","99.18","421.97","0","425.07","39.3","0"
"AGAW1","10N","NIN","BOLD:AAA6381","Arthropoda","Arachnida","Araneae","Tetragnathidae","","Tetragnatha","Tetragnatha laboriosa","186","180","180","3","3.23","0","99.26","427.23","0","429.75","39.34","0"
"AGAW1","10N","NIN","BOLD:AAC6924","Arthropoda","Arachnida","Araneae","Anyphaenidae","","Wulfila","Wulfila saltabundus","128","128","128","4","0","0","98.85","394.47","0","397.2","30.34","0"
"AGAW1","10N","NIN","BOLD:ABW2639","Arthropoda","Arachnida","Trombidiformes","Anystidae","","","","46","43","43","5","6.52","0","99.34","449.22","0","447.61","31.49","0"
"AGAW1","10N","NIN","BOLD:AAH6630","Arthropoda","Collembola","Entomobryomorpha","Entomobryidae","Entomobryinae","Entomobrya","Entomobrya unostrigata","38","36","36","6","5.26","0","99.03","433.58","0","434.55","39.13","0"
"AGAW1","10N","NIN","BOLD:ACY2550","Arthropoda","Arachnida","Trombidiformes","Bdellidae","","Cyta","","31","31","31","7","0","0","98.64","375.84","0","391.97","31.04","0"
"AGAW1","10N","NIN","BOLD:AAA6383","","","","","","","","20","20","20","8","0","0","98.26","430.35","0","433.4","37.02","0"
"AGAW1","10N","NIN","BOLD:AAG1792","Arthropoda","Arachnida","Araneae","Tetragnathidae","","Glenognatha","Glenognatha foxi","17","17","17","9","0","0","98.34","436.47","0","438.06","31.29","0"
"AGAW1","10N","NIN","BOLD:AAB7914","Arthropoda","Collembola","Symphypleona","Bourletiellidae","","Bourletiella","","11","11","11","10","0","0","99.26","442.64","0","443.91","38.12","0"
"AGAW1","10N","NIN","BOLD:ACZ1295","Arthropoda","Arachnida","Trombidiformes","Anystidae","","","","9","9","9","11","0","0","99.2","376.44","0","387.11","31.34","0"
"AGAW1","10N","NIN","BOLD:AAO8147","Arthropoda","Arachnida","Trombidiformes","Eupodidae","","Eupodes","","9","9","9","11","0","0","98.91","420.22","0","422.44","27.87","0"
"AGAW1","10N","NIN","BOLD:AAF9285","Arthropoda","Arachnida","Mesostigmata","Parasitidae","","","","8","8","8","13","0","0","98.83","428.5","0","430","35.23","0"
"AGAW1","10N","NIN","BOLD:AAF9234","Arthropoda","Arachnida","Trombidiformes","Eupodidae","","","","6","6","6","14","0","0","99.55","439.67","0","443.17","29.83","0"
"AGAW1","10N","NIN","BOLD:AAM7961","Arthropoda","Arachnida","Trombidiformes","Anystidae","","","","3","3","3","15","0","0","100","448.33","0","450","31.19","0"
"AGAW1","10N","NIN","BOLD:ACX8969","Arthropoda","Arachnida","Trombidiformes","Eupodidae","","","","2","2","2","16","0","0","98.75","435.5","0","439","27.45","0"
"AGAW1","10N","NIN","BOLD:AAI4346","Arthropoda","Arachnida","Opiliones","Phalangiidae","","Paroligolophus, Phalangium","Paroligolophus agrestis, Phalangium opilio","1","1","1","17","0","0","99.21","381","0","386","33.68","0"
"AGAW1","10S","BAC","TAX:91391","Proteobacteria","Alphaproteobacteria","Rickettsiales","",NA,"","","13","10","10","1","23.08","0","99.19","419.31","0","440.54","39.62","0"
"AGAW1","10S","BAC","TAX:175768","Proteobacteria","Alphaproteobacteria","Rickettsiales","Rickettsiaceae",NA,"Wolbachia","Wolbachia pipientis","3","3","3","2","0","0","98.36","447.67","0","448.67","39.75","0"
"AGAW1","10S","CHO","no match","no match","no match","no match","no match","no match","no match","no match","no match","no match","no match","no match","no match","no match","no match","no match","no match","no match","no match","no match"
"AGAW1","10S","INS","BOLD:AAA3453","Arthropoda","Insecta","Diptera","Anthomyiidae","Anthomyiinae","Delia","","22302","18142","18142","1","18.65","0","99.45","430.92","0","433.56","34.65","0"
"AGAW1","10S","INS","BOLD:AAA6560","Arthropoda","Insecta","Lepidoptera","Crambidae","Crambinae","Crambus","Crambus multilinellus, Crambus saltuellus","21191","14373","14373","2","32.17","0","99.57","439.71","0","440.98","30.82","0"
"AGAW1","10S","INS","BOLD:AAU6681","Arthropoda","Insecta","Diptera","Tipulidae","Tipulinae","Nephrotoma","Nephrotoma cornicina","10821","7723","7723","3","28.63","0","99.62","443.21","0","444.39","36.31","0"
"AGAW1","10S","INS","BOLD:AAP8739","Arthropoda","Insecta","Diptera","Platystomatidae","Platystomatinae","Rivellia","Rivellia steyskali","7951","4980","4980","4","37.37","0","99.31","441.77","0","443.55","37.98","0"
"AGAW1","10S","INS","BOLD:AAI4303","Arthropoda","Insecta","Diptera","Chironomidae","Chironominae","Chironomus","Chironomus melanescens","3559","3304","3304","5","7.16","0","98.37","427.99","0","432.22","34.12","0"
"AGAW1","10S","INS","BOLD:ACE7444","Arthropoda","Insecta","Hemiptera","Miridae","Mirinae, Phylinae","Adelphocoris, Megalocoleus","Adelphocoris lineolatus, Megalocoleus molliculus","2443","1590","1590","6","34.92","0","99.62","441.88","0","442.56","34.8","0"
"AGAW1","10S","INS","BOLD:AAA3889","Arthropoda","Insecta","Lepidoptera","Crambidae","Crambinae","Parapediasia","Parapediasia teterrellus","2430","2185","2185","7","10.08","0","99.23","428.54","0","430.36","32.12","0"
"AGAW1","10S","INS","BOLD:AAC9132","Arthropoda","Insecta","Diptera","Culicidae","Culicinae","Culiseta","Culiseta inornata","2011","1981","1981","8","1.49","0","98.56","406.93","0","409.8","32.44","0"
"AGAW1","10S","INS","BOLD:AAN6154","Arthropoda","Insecta","Coleoptera","Latridiidae","Corticariinae","Melanophthalma","Melanophthalma inermis","1851","1523","1523","9","17.72","0","98.96","437.7","0","439.34","31.36","0"
"AGAW1","10S","INS","BOLD:AAA7631","Arthropoda","Insecta","Hemiptera","Miridae","Mirinae","Adelphocoris","Adelphocoris lineolatus","1022","695","695","10","32","0","99.15","446.02","0","447.58","35.44","0"
"AGAW1","10S","INS","BOLD:AAC0903","Arthropoda","Insecta","Diptera","Chironomidae","Chironominae","Chironomus","Chironomus acidophilus","929","752","752","11","19.05","0","99.36","439.53","0","442.09","36.15","0"
"AGAW1","10S","INS","BOLD:AAG2868","Arthropoda","Insecta","Hemiptera","Cicadellidae","Typhlocybinae","Empoasca","Empoasca sp., Empoasca fabae","851","516","516","12","39.37","0","99.68","445.41","0","448.2","32.43","0"
"AGAW1","10S","INS","BOLD:AAG5453","Arthropoda","Insecta","Diptera","Chironomidae","Chironominae","Chironomus","Chironomus bifurcatus","639","515","515","13","19.41","0","98.29","442.54","0","446.42","34.93","0"
"AGAW1","10S","INS","BOLD:AAG6745","Arthropoda","Insecta","Diptera","Calliphoridae","Calliphorinae","Pollenia","Pollenia pediculata","563","511","511","14","9.24","0","99.31","432.44","0","434.37","33.52","0"
"AGAW1","10S","INS","BOLD:ACE0699","Arthropoda","Insecta","Diptera","Chironomidae","","","","477","346","346","15","27.46","0","98.57","425.89","0","442.71","36.57","0"
"AGAW1","10S","INS","BOLD:AAH3661","Arthropoda","Insecta","Diptera","Cecidomyiidae","","","","455","401","401","16","11.87","0","99.28","434.95","0","436.54","25.49","0"
"AGAW1","10S","INS","BOLD:AAA4555","Arthropoda","Insecta","Orthoptera","Acrididae","Melanoplinae","Melanoplus","Melanoplus stupefactus, Melanoplus angustipennis, Melanoplus ponderosus, Melanoplus bruneri, Melanoplus frigidus, Melanoplus scapularis, Melanoplus cinereus, Melanoplus madeleineae, Melanoplus sanguinipes, Melanoplus nigrescens group, Melanoplus tunicae, Melanoplus walshi, Melanoplus femurrubrum, Melanoplus dawsoni, Melanoplus sp., Melanoplus fasciatus, Melanoplus bowditchi, Melanoplus stonei, Melanoplus bivittatus, Melanoplus huroni, Melanoplus thomasi, Melanoplus borealis, Melanoplus packardii","404","353","353","17","12.62","0","99.33","446.99","0","448.37","36.23","0"
"AGAW1","10S","INS","BOLD:ABA8103","Arthropoda","Insecta","Hemiptera","Cicadellidae","Typhlocybinae","Empoasca","","403","329","329","18","18.36","0","97.78","437.02","0","438.22","32.87","0"
"AGAW1","10S","INS","BOLD:AAG2898","Arthropoda","Insecta","Hemiptera","Cicadellidae","Deltocephalinae","Paraphlepsius","Paraphlepsius apertus, Paraphlepsius irroratus","386","316","316","19","18.13","0","99.38","442.94","0","443.79","32.09","0"
"AGAW1","10S","INS","BOLD:AAH0256","Arthropoda","Insecta","Coleoptera","Latridiidae","Corticariinae","Corticaria, Corticarina","","299","221","221","20","26.09","0","98.91","439.97","0","444.1","31.74","0"
"AGAW1","10S","INS","BOLD:AAG5507","Arthropoda","Insecta","Diptera","Chironomidae","Chironominae","Chironomus","Chironomus atrella","286","284","284","21","0.7","0","98.68","413.58","0","416.83","35.61","0"
"AGAW1","10S","INS","BOLD:ACY6524","Arthropoda","Insecta","Diptera","Chironomidae","Orthocladiinae","Psectrocladius","Psectrocladius cf. limbatellus","274","249","249","22","9.12","0","99.15","433.76","0","444.75","35.89","0"
"AGAW1","10S","INS","BOLD:AAG9897","Arthropoda","Insecta","Dermaptera","Forficulidae","Forficulinae","Forficula","Forficula auricularia-A","220","219","219","23","0.45","0","99.05","399.21","0","410.54","45.66","0"
"AGAW1","10S","INS","BOLD:AAD0484","Arthropoda","Insecta","Diptera","Chironomidae","","","","179","149","149","24","16.76","0","99.12","444.92","0","446.28","34.87","0"
"AGAW1","10S","INS","BOLD:ABV1297","Arthropoda","Insecta","Diptera","Sciaridae","","","","178","174","174","25","2.25","0","98.98","435.56","0","438.31","30.24","0"
"AGAW1","10S","INS","BOLD:ACK3431","Arthropoda","Insecta","Diptera","Cecidomyiidae","","","","167","157","157","26","5.99","0","99.17","429.5","0","431.55","32.59","0"
"AGAW1","10S","INS","BOLD:AAV5853","Arthropoda","Insecta","Diptera","Chironomidae","Tanypodinae","","","159","157","157","27","1.26","0","98.52","416.53","0","418.23","34.65","0"
"AGAW1","10S","INS","BOLD:AAA7661","Arthropoda","Insecta","Diptera","Culicidae","Culicinae","Culiseta, Culex","Culiseta melanura, Culex articularis, Culex restuans, Culex acharistus","156","151","151","28","3.21","0","99.2","433.68","0","435.68","32.96","0"
"AGAW1","10S","INS","BOLD:ABW7897","Arthropoda","Insecta","Diptera","Cecidomyiidae","","","","153","120","120","29","21.57","0","99.6","442.35","0","444.12","29.66","0"
"AGAW1","10S","INS","BOLD:ABW1379","Arthropoda","Insecta","Diptera","Chloropidae","Oscinellinae","Malloewia","Malloewia abdominalis","149","120","120","30","19.46","0","98.13","445.39","0","447.81","31.71","0"
"AGAW1","10S","INS","BOLD:AAA5803","Arthropoda","Insecta","Hemiptera","Miridae","Mirinae","Lygus","Lygus lineolaris","144","104","104","31","27.78","0","98.76","447.29","0","448.78","33.96","0"
"AGAW1","10S","INS","BOLD:AAN5199","Arthropoda","Insecta","Diptera","Cecidomyiidae","","","CecidInt20 sp. BOLD:AAN5199","126","105","105","32","16.67","0","99.51","435.78","0","445.29","26.63","0"
"AGAW1","10S","INS","BOLD:AAF9789","Arthropoda","Insecta","Diptera","Hybotidae","","","","115","113","113","33","1.74","0","99.2","422.56","0","424.38","32.99","0"
"AGAW1","10S","INS","BOLD:ACU7906","Arthropoda","Insecta","Diptera","Chironomidae","Orthocladiinae","","","108","108","108","34","0","0","98.72","411.4","0","414.09","36.1","0"
"AGAW1","10S","INS","BOLD:AAI7960","Arthropoda","Insecta","Diptera","Agromyzidae","Agromyzinae","Japanagromyza","Japanagromyza viridula","99","99","99","35","0","0","98.82","411.45","0","416.09","31.75","0"
"AGAW1","10S","INS","BOLD:AAF9014","Arthropoda","Insecta","Diptera","Limoniidae","Chioneinae","Symplecta","Symplecta cana, Symplecta sp. BOLDAAF9014","77","73","73","36","5.19","0","99.04","413.92","0","425.04","34","0"
"AGAW1","10S","INS","BOLD:AAH0135","Arthropoda","Insecta","Coleoptera","Phalacridae","Phalacrinae","Acylomus","Acylomus pugetanus, Acylomus ergoti","75","71","71","37","5.33","0","99.01","438.99","0","441.11","31.73","0"
"AGAW1","10S","INS","BOLD:ACB6899","Arthropoda","Insecta","Diptera","Cecidomyiidae","","","","75","64","64","37","14.67","0","99.61","415.79","0","444.11","26.3","0"
"AGAW1","10S","INS","BOLD:AAN5242","Arthropoda","Insecta","Diptera","Cecidomyiidae","","","","74","73","73","39","1.35","0","98.95","400.5","0","402.31","28.03","0"
"AGAW1","10S","INS","BOLD:AAM9013","Arthropoda","Insecta","Diptera","Keroplatidae","Macrocerinae","Macrocera","Macrocera pusilla","74","73","73","39","1.35","0","99.19","435.7","0","437.38","30.42","0"
"AGAW1","10S","INS","BOLD:ABU5525","Arthropoda","Insecta","Diptera","Chironomidae","Orthocladiinae","Limnophyes","Limnophyes sp. 14ES","67","67","67","41","0","0","98.92","400.55","0","403.67","37.21","0"
"AGAW1","10S","INS","BOLD:ABA0863","Arthropoda","Insecta","Diptera","Cecidomyiidae","","","","65","56","56","42","13.85","0","99.26","442.45","0","443.72","29.74","0"
"AGAW1","10S","INS","BOLD:AAC4525","Arthropoda","Insecta","Diptera","Chironomidae","Chironominae","Tanytarsus","Tanytarsus guerlus","63","50","50","43","20.63","0","97.98","445.17","0","446.16","33.93","0"
"AGAW1","10S","INS","BOLD:ACL8053","Arthropoda","Insecta","Diptera","Chironomidae","Chironominae","Tanytarsus","","55","51","51","44","7.27","0","99.41","447.38","0","449.05","33.7","0"
"AGAW1","10S","INS","BOLD:AAN5205","Arthropoda","Insecta","Diptera","Cecidomyiidae","","","CecidInt42 sp. BOLD:AAN5205","53","50","50","45","5.66","0","99.47","445.85","0","447.58","31.36","0"
"AGAW1","10S","INS","BOLD:AAD1945","Arthropoda","Insecta","Diptera","Agromyzidae","Phytomyzinae","Cerodontha","Cerodontha dorsalis","49","44","44","46","10.2","0","99.28","442.14","0","443.94","32.06","0"
"AGAW1","10S","INS","BOLD:AAF9949","Arthropoda","Insecta","Hemiptera","Miridae","Mirinae","Trigonotylus","Trigonotylus pulchellus, Trigonotylus americanus, Trigonotylus psammaecolor, Trigonotylus caelestialium, Trigonotylus viridis, Trigonotylus ruficornis","48","35","35","47","27.08","0","99.47","449.5","0","449.79","34.63","0"
"AGAW1","10S","INS","BOLD:AAG6929","Arthropoda","Insecta","Diptera","Dolichopodidae","Medeterinae","Medetera","Medetera vittata","48","44","44","47","8.33","0","98.54","433.52","0","437.94","41.1","0"
"AGAW1","10S","INS","BOLD:ACC8079","Arthropoda","Insecta","Diptera","Cecidomyiidae","","","","48","36","36","47","25","0","99.47","445.52","0","447.13","29.1","0"
"AGAW1","10S","INS","BOLD:AAA7368","Arthropoda","Insecta","Lepidoptera","Cosmopterigidae","Cosmopteriginae","Limnaecia","Limnaecia phragmitella","48","44","44","47","8.33","0","99.43","425.85","0","432.69","30.95","0"
"AGAW1","10S","INS","BOLD:ACM0749","Arthropoda","Insecta","Coleoptera","Anthicidae","Notoxinae","Notoxus","Notoxus anchora","48","48","48","47","0","0","99.15","422.65","0","424.29","35.34","0"
"AGAW1","10S","INS","BOLD:ADF3937","","","","","","","","46","46","46","52","0","0","98.86","413.17","0","423.48","31.4","0"
"AGAW1","10S","INS","BOLD:AAH3664","Arthropoda","Insecta","Diptera","Cecidomyiidae","","","CecidInt3 sp. BOLD:AAH3664","42","42","42","53","0","0","99.27","425.02","0","428.02","26.8","0"
"AGAW1","10S","INS","BOLD:ACF2457","Arthropoda","Insecta","Diptera","Chironomidae","Chironominae","Paratanytarsus","Paratanytarsus laccophilus","42","42","42","53","0","0","98.83","438.1","0","440.31","33.1","0"
"AGAW1","10S","INS","BOLD:ABA7871","Arthropoda","Insecta","Diptera","Cecidomyiidae","","","","36","35","35","55","2.78","0","99.05","437.78","0","439","26.76","0"
"AGAW1","10S","INS","BOLD:ADF6661","Arthropoda","Insecta","Diptera","Cecidomyiidae","","","","35","34","34","56","2.86","0","99.18","415.34","0","440.57","29.52","0"
"AGAW1","10S","INS","BOLD:AAD9052","Arthropoda","Insecta","Lepidoptera","Elachistidae","Elachistinae","Elachista","Elachista herbigrada, Elachista rianella, Elachista illectella, Elachista sp., Elachista tuorella","34","30","30","57","11.76","0","99.45","440.65","0","443.44","31.47","0"
"AGAW1","10S","INS","BOLD:ADG8758","Arthropoda","Insecta","Hymenoptera","Braconidae","Brachistinae","Blacus","Blacus stelfoxi","34","33","33","57","2.94","0","99.06","436.44","0","438.71","32.09","0"
"AGAW1","10S","INS","BOLD:AAH0134","Arthropoda","Insecta","Coleoptera","Phalacridae","Phalacrinae","Stilbus","Stilbus apicialis","34","34","34","57","0","0","99.1","427.97","0","428.97","31.86","0"
"AGAW1","10S","INS","BOLD:AAN5341","Arthropoda","Insecta","Diptera","Chironomidae","Orthocladiinae","Camptocladius","Camptocladius stercorarius","34","34","34","57","0","0","98.75","394.29","0","400.06","35.48","0"
"AGAW1","10S","INS","BOLD:ABV9298","Arthropoda","Insecta","Diptera","Chironomidae","Chironominae","Chironomus","","32","32","32","61","0","0","98.98","420.47","0","423.59","34.64","0"
"AGAW1","10S","INS","BOLD:ABA0822","Arthropoda","Insecta","Diptera","Cecidomyiidae","","","","32","30","30","61","6.25","0","98.56","442.72","0","444.09","26.75","0"
"AGAW1","10S","INS","BOLD:AAZ9744","Arthropoda","Insecta","Hymenoptera","Braconidae","Brachistinae","Blacus","Blacus stelfoxi","31","19","19","63","38.71","0","98.58","449.23","0","450","32.41","0"
"AGAW1","10S","INS","BOLD:AAN5238","Arthropoda","Insecta","Diptera","Cecidomyiidae","","","","30","29","29","64","3.33","0","99.47","428.77","0","430.1","25.75","0"
"AGAW1","10S","INS","BOLD:ABA5313","Arthropoda","Insecta","Coleoptera","Staphylinidae","Aleocharinae","Amischa","Amischa analis, Amischa bifoveolata, Amischa nigrofusca","28","26","26","65","7.14","0","99.6","438.5","0","440.14","33.35","0"
"AGAW1","10S","INS","BOLD:AAH4171","Arthropoda","Insecta","Diptera","Chloropidae","Oscinellinae","Apallates","Apallates montanus, Apallates particeps","27","27","27","66","0","0","98.51","391.11","0","394.04","31.87","0"
"AGAW1","10S","INS","BOLD:ACE4701","Arthropoda","Insecta","Lepidoptera","Crambidae","Crambinae","Crambus","Crambus multilinellus","27","27","27","66","0","0","98.1","422.19","0","423.85","30.64","0"
"AGAW1","10S","INS","BOLD:AAD5253","Arthropoda","Insecta","Diptera","Chironomidae","Orthocladiinae","Thienemanniella","Thienemanniella xena, Thienemanniella vittata","26","24","24","68","7.69","0","98.34","433.54","0","434.92","34.31","0"
"AGAW1","10S","INS","BOLD:AAH4276","Arthropoda","Insecta","Lepidoptera","Gelechiidae","Gelechiinae","Bryotropha","Bryotropha hodgesi, Bryotropha altitudophila","26","24","24","68","7.69","0","99.36","441.96","0","443.58","33.6","0"
"AGAW1","10S","INS","BOLD:AAN5183","Arthropoda","Insecta","Diptera","Cecidomyiidae","","","","24","23","23","70","4.17","0","98.46","446.29","0","447.17","27.83","0"
"AGAW1","10S","INS","BOLD:AAN5255","Arthropoda","Insecta","Diptera","Cecidomyiidae","Cecidomyiinae","Dasineura","","22","22","22","71","0","0","98.45","424.82","0","440.82","26.67","0"
"AGAW1","10S","INS","BOLD:AAG5443","Arthropoda","Insecta","Diptera","Chironomidae","Orthocladiinae","","","22","22","22","71","0","0","98.98","412.23","0","414.05","33.44","0"
"AGAW1","10S","INS","BOLD:ABA0856","Arthropoda","Insecta","Diptera","Cecidomyiidae","","","","21","15","15","73","28.57","0","99.53","449.33","0","450","29.16","0"
"AGAW1","10S","INS","BOLD:ABV1337","Arthropoda","Insecta","Diptera","Cecidomyiidae","","","","21","21","21","73","0","0","99.11","434.52","0","435.81","28.22","0"
"AGAW1","10S","INS","BOLD:ACV2677","Arthropoda","Insecta","Diptera","Cecidomyiidae","","","","20","18","18","75","10","0","97.6","449.3","0","450","29.77","0"
"AGAW1","10S","INS","BOLD:AAG2511","Arthropoda","Insecta","Diptera","Anthomyiidae","Anthomyiinae","Delia","Delia platura","18","18","18","76","0","0","98.54","446.44","0","448.72","33.62","0"
"AGAW1","10S","INS","BOLD:AAV6213","Arthropoda","Insecta","Diptera","Chironomidae","Chironominae","Paratanytarsus","","17","17","17","77","0","0","99.25","427.82","0","430.24","34.25","0"
"AGAW1","10S","INS","BOLD:AAD1485","Arthropoda","Insecta","Diptera","Chironomidae","Chironominae","Paratanytarsus","Paratanytarsus grimmii, Paratanytarsus sp.","17","13","13","77","23.53","0","99.52","447.47","0","448.35","33.86","0"
"AGAW1","10S","INS","BOLD:AAM6037","Arthropoda","Insecta","Diptera","Cecidomyiidae","Cecidomyiinae","Contarinia, Dasineura","","17","16","16","77","5.88","0","98.48","436.71","0","438.53","25.66","0"
"AGAW1","10S","INS","BOLD:ACF7633","Arthropoda","Insecta","Diptera","Cecidomyiidae","","","","16","15","15","80","6.25","0","99.5","446.69","0","447.94","30.6","0"
"AGAW1","10S","INS","BOLD:AAA4781","Arthropoda","Insecta","Hymenoptera","Braconidae","Microgastrinae","Glyptapanteles","Glyptapanteles carlossarmientoi","16","16","16","80","0","0","99.17","437.88","0","440.5","24.29","0"
"AGAW1","10S","INS","BOLD:AAG1711","Arthropoda","Insecta","Diptera","Muscidae","Phaoniinae","Helina","Helina depuncta","15","15","15","82","0","0","99.03","426.33","0","429.13","31.13","0"
"AGAW1","10S","INS","BOLD:AAN5225","Arthropoda","Insecta","Diptera","Cecidomyiidae","","","","15","14","14","82","6.67","0","98.97","447.47","0","448.6","28.22","0"
"AGAW1","10S","INS","BOLD:AAP5087","Arthropoda","Insecta","Diptera","Ceratopogonidae","Dasyheleinae","Dasyhelea","","14","12","12","84","14.29","0","99.26","435.5","0","436.79","34.78","0"
"AGAW1","10S","INS","BOLD:AAG7891","Arthropoda","Insecta","Hymenoptera","Platygastridae","Scelioninae","Telenomus","Telenomus podisi","13","12","12","85","7.69","0","99.15","430.54","0","438.69","26.89","0"
"AGAW1","10S","INS","BOLD:ACA6930","Arthropoda","Insecta","Diptera","Cecidomyiidae","","","","13","11","11","85","15.38","0","98.36","427.54","0","445.92","27.95","0"
"AGAW1","10S","INS","BOLD:AAG9668","Arthropoda","Insecta","Diptera","Dolichopodidae","Diaphorinae","Chrysotus","Chrysotus sp., Chrysotus sp. nr. obliquus","12","12","12","87","0","0","98.88","447","0","449.08","33","0"
"AGAW1","10S","INS","BOLD:ABA9076","Arthropoda","Insecta","Coleoptera","Latridiidae","Latridiinae","Cartodere","Cartodere constricta","10","10","10","88","0","0","99.08","415.1","0","417.2","35.31","0"
"AGAW1","10S","INS","BOLD:AAZ5601","Arthropoda","Insecta","Diptera","Chironomidae","Orthocladiinae","Smittia","","10","9","9","88","10","0","99.17","427.6","0","428.5","33.65","0"
"AGAW1","10S","INS","BOLD:AAO4092","Arthropoda","Insecta","Psocodea","Caeciliusidae","Caeciliusinae","Valenzuela","Valenzuela burmeisteri","10","10","10","88","0","0","98.66","433.1","0","435.5","35.05","0"
"AGAW1","10S","INS","BOLD:AAL7819","Arthropoda","Insecta","Diptera","Psychodidae","Psychodinae","Psychoda","Psychoda sp., Psychoda sp. BOLD:AAL7819","9","9","9","91","0","0","99.12","421.89","0","423.33","32.56","0"
"AGAW1","10S","INS","BOLD:AAN5220","Arthropoda","Insecta","Diptera","Cecidomyiidae","","","","9","9","9","91","0","0","98.22","414","0","427.56","25.87","0"
"AGAW1","10S","INS","BOLD:ABV1475","Arthropoda","Insecta","Diptera","Cecidomyiidae","","","","8","7","7","93","12.5","0","99.32","440.75","0","441.38","26.26","0"
"AGAW1","10S","INS","BOLD:AAA5299","Arthropoda","Insecta","Diptera","Chironomidae","Orthocladiinae","Cricotopus","Cricotopus laetus, Cricotopus relucens, Cricotopus trifasciatus, Cricotopus sp., Cricotopus sp. 3ES, Cricotopus sylvestris, Cricotopus glacialis","8","8","8","93","0","0","99.03","431.38","0","432.5","34.78","0"
"AGAW1","10S","INS","BOLD:AAN5538","Arthropoda","Insecta","Diptera","Dolichopodidae","","","","8","8","8","93","0","0","98.29","436.63","0","436.88","39.16","0"
"AGAW1","10S","INS","BOLD:AAF1013","Arthropoda","Insecta","Diptera","Chironomidae","Chironominae","Tanytarsus","Tanytarsus allicis","7","7","7","96","0","0","97.67","434.71","0","438.57","36.53","0"
"AGAW1","10S","INS","BOLD:ADH9867","Arthropoda","Insecta","Diptera","","","","","7","7","7","96","0","0","98.92","368.43","0","371.29","37.48","0"
"AGAW1","10S","INS","BOLD:ACC8303","Arthropoda","Insecta","Diptera","Cecidomyiidae","","","","7","7","7","96","0","0","98.61","430.86","0","433.86","26.36","0"
"AGAW1","10S","INS","BOLD:AAG8839","Arthropoda","Insecta","Hemiptera","Cicadellidae","Deltocephalinae","Errastunus, Latalus","Latalus ocellaris, Errastunus ocellaris","6","6","6","99","0","0","99.74","449.67","0","450","36.15","0"
"AGAW1","10S","INS","BOLD:AAM6040","Arthropoda","Insecta","Diptera","Cecidomyiidae","","","","6","5","5","99","16.67","0","99.29","437.17","0","438","26.81","0"
"AGAW1","10S","INS","BOLD:ACF0915","Arthropoda","Insecta","Diptera","Chironomidae","Orthocladiinae","","","5","5","5","101","0","0","98.55","399.2","0","402.2","33.15","0"
"AGAW1","10S","INS","BOLD:AAD7604","Arthropoda","Insecta","Coleoptera","Coccinellidae","Coccinellinae","Coleomegilla","Coleomegilla maculata","5","5","5","101","0","0","98.63","426.2","0","428.6","31.66","0"
"AGAW1","10S","INS","BOLD:AAL7442","Arthropoda","Insecta","Diptera","Cecidomyiidae","","","","5","5","5","101","0","0","99.86","436.6","0","438.2","26.54","0"
"AGAW1","10S","INS","BOLD:ACF1257","","","","","","","","5","5","5","101","0","0","99.38","449","0","450","34.84","0"
"AGAW1","10S","INS","BOLD:AAN5332","Arthropoda","Insecta","Diptera","Chironomidae","Chironominae","Polypedilum","","5","5","5","101","0","0","98.12","418.8","0","422.2","33.26","0"
"AGAW1","10S","INS","BOLD:AAH3662","Arthropoda","Insecta","Diptera","Cecidomyiidae","","","CecidInt3 sp. BOLD:AAH3662","5","5","5","101","0","0","99.32","441","0","443.2","26.12","0"
"AGAW1","10S","INS","BOLD:AAF9306","Arthropoda","Insecta","Diptera","Psychodidae","Psychodinae","Psychoda","Psychoda alternata","5","5","5","101","0","0","98.37","442.4","0","443.6","31.42","0"
"AGAW1","10S","INS","BOLD:ABA4532","Arthropoda","Insecta","Coleoptera","Chrysomelidae","Galerucinae","Psylliodes","Psylliodes napi","4","4","4","108","0","0","99.43","418","0","418.5","35.82","0"
"AGAW1","10S","INS","BOLD:AAN5230","Arthropoda","Insecta","Diptera","Cecidomyiidae","","","","4","4","4","108","0","0","99.36","421","0","422","26.15","0"
"AGAW1","10S","INS","BOLD:AAG4905","Arthropoda","Insecta","Diptera","Mycetophilidae","Mycetophilinae","Mycetophila","","4","4","4","108","0","0","99.05","448.75","0","450","35.22","0"
"AGAW1","10S","INS","BOLD:ACX3809","Arthropoda","Insecta","Thysanoptera","Phlaeothripidae","Phlaeothripinae","Haplothrips","","4","4","4","108","0","0","99.94","442.25","0","450","27.89","0"
"AGAW1","10S","INS","BOLD:AAB0373","Arthropoda","Insecta","Neuroptera","Chrysopidae","Chrysopinae","Chrysoperla","Chrysoperla sp. isolate 82, Chrysoperla zastrowi, Chrysoperla plorabunda, Chrysoperla sp. isolate 71, Chrysoperla agilis, Italochrysa italica, Chrysoperla sp. isolate 88, Chrysoperla lucasina, Chrysoperla carnea, Chrysoperla mediterranea, Chrysoperla pallida, Chrysoperla nipponensis, Chrysoperla sillemi","4","4","4","108","0","0","99.42","442.5","0","442.75","31.16","0"
"AGAW1","10S","INS","BOLD:ACV5892","Arthropoda","Insecta","Diptera","Cecidomyiidae","","","","4","4","4","108","0","0","97.83","449","0","450","25.44","0"
"AGAW1","10S","INS","BOLD:ACA3036","Arthropoda","Insecta","Diptera","Chironomidae","Chironominae","Cladotanytarsus","","3","3","3","114","0","0","99.38","427.67","0","428.67","39.03","0"
"AGAW1","10S","INS","BOLD:AAY6469","Arthropoda","Insecta","Diptera","Cecidomyiidae","","","","3","3","3","114","0","0","99.23","403.33","0","404","25.95","0"
"AGAW1","10S","INS","BOLD:ACI3976","Arthropoda","Insecta","Diptera","Chironomidae","Orthocladiinae","","","3","3","3","114","0","0","99.49","392","0","410","37.32","0"
"AGAW1","10S","INS","BOLD:AAN6447","Arthropoda","Insecta","Diptera","Sciaridae","","Corynoptera","Corynoptera perpusilla","3","3","3","114","0","0","99.63","448.33","0","450","36.15","0"
"AGAW1","10S","INS","BOLD:AAP5169","Arthropoda","Insecta","Diptera","Chloropidae","Oscinellinae","Elachiptera","Elachiptera nigriceps","3","3","3","114","0","0","99.78","448.33","0","450","31.48","0"
"AGAW1","10S","INS","BOLD:ABY5318","Arthropoda","Insecta","Diptera","Chloropidae","Oscinellinae","Incertella, Rhopalopterum","Rhopalopterum umbrosum, Rhopalopterum luteiceps, Incertella bispina, Rhopalopterum painteri","3","3","3","114","0","0","100","449.67","0","450","32.15","0"
"AGAW1","10S","INS","BOLD:AAN5267","Arthropoda","Insecta","Diptera","Cecidomyiidae","","","","3","3","3","114","0","0","98.74","450","0","450","30.67","0"
"AGAW1","10S","INS","BOLD:ACF4925","Arthropoda","Insecta","Hemiptera","Miridae","Mirinae","Adelphocoris","Adelphocoris detritus","3","3","3","114","0","0","97.26","437.67","0","440","36.35","0"
"AGAW1","10S","INS","BOLD:AAG4944","Arthropoda","Insecta","Diptera","Mycetophilidae","Sciophilinae","Sciophila","Sciophila sp. CJB1","3","3","3","114","0","0","98.05","409.67","0","413","33","0"
"AGAW1","10S","INS","BOLD:AAA4277","Arthropoda","Insecta","Diptera","Syrphidae","Syrphinae","Toxomerus","Toxomerus marginatus, Toxomerus sp. CR-3, Toxomerus CR-3","3","3","3","114","0","0","99.41","449","0","450","30.81","0"
"AGAW1","10S","INS","BOLD:ABX4079","Arthropoda","Insecta","Diptera","Cecidomyiidae","","","","2","2","2","124","0","0","98.32","446.5","0","449.5","28.25","0"
"AGAW1","10S","INS","BOLD:AAV5796","Arthropoda","Insecta","Diptera","Cecidomyiidae","","","","2","2","2","124","0","0","99.55","447","0","450","29.11","0"
"AGAW1","10S","INS","BOLD:AAF9771","Arthropoda","Insecta","Diptera","Hybotidae","Tachydromiinae","Platypalpus","Platypalpus longiseta, Platypalpus trivialis, Platypalpus sp. 5, Platypalpus sp. 17, Platypalpus pallidiventris","2","2","2","124","0","0","98.55","449.5","0","450","34.89","0"
"AGAW1","10S","INS","BOLD:AAL7359","Arthropoda","Insecta","Diptera","Chironomidae","Chironominae","Tanytarsus","","2","2","2","124","0","0","99.78","447.5","0","450","36.44","0"
"AGAW1","10S","INS","BOLD:AAN5215","Arthropoda","Insecta","Diptera","Cecidomyiidae","","","","2","2","2","124","0","0","98.6","438.5","0","442","29.29","0"
"AGAW1","10S","INS","BOLD:AAG2123","Arthropoda","Insecta","Diptera","Tachinidae","Dexiinae","Dinera","Dinera grisescens","2","2","2","124","0","0","98.55","449","0","450","31.89","0"
"AGAW1","10S","INS","BOLD:AAG1528","Arthropoda","Insecta","Hemiptera","Aphididae","Chaitophorinae","Sipha","Sipha elegans","2","2","2","124","0","0","99.22","450","0","450","28.44","0"
"AGAW1","10S","INS","BOLD:AAO7694","Arthropoda","Insecta","Diptera","Cecidomyiidae","","","","2","2","2","124","0","0","99.43","442.5","0","444.5","26.66","0"
"AGAW1","10S","INS","BOLD:AAA3181","Arthropoda","Insecta","Diptera","Muscidae","Muscinae","Stomoxys","Stomoxys calcitrans","1","1","1","132","0","0","98.74","398","0","404","31.68","0"
"AGAW1","10S","INS","BOLD:ACK5742","Arthropoda","Insecta","Diptera","Cecidomyiidae","","","","1","1","1","132","0","0","100","447","0","450","29.11","0"
"AGAW1","10S","INS","BOLD:AAN5160","Arthropoda","Insecta","Diptera","Ceratopogonidae","Forcipomyiinae","Atrichopogon","","1","1","1","132","0","0","98.86","350","0","353","35.69","0"
"AGAW1","10S","INS","BOLD:AAA6292","Arthropoda","Insecta","Lepidoptera","Tortricidae","Tortricinae","Cnephasia","Cnephasia genitalana, Cnephasia stephensiana","1","1","1","132","0","0","100","449","0","450","35.56","0"
"AGAW1","10S","INS","BOLD:AAM6655","Arthropoda","Insecta","Diptera","Hybotidae","Tachydromiinae","Platypalpus","Platypalpus stabilis","1","1","1","132","0","0","99.56","450","0","450","35.11","0"
"AGAW1","10S","INS","BOLD:AAG5467","Arthropoda","Insecta","Diptera","Chironomidae","Chironominae","Tanytarsus","","1","1","1","132","0","0","99.03","413","0","414","30.43","0"
"AGAW1","10S","INS","BOLD:AAA2518","Arthropoda","Insecta","Lepidoptera","Crambidae","Crambinae","Chrysoteuchia","Chrysoteuchia topiaria, Chrysoteuchia topiarius","1","1","1","132","0","0","97.1","448","0","450","34.67","0"
"AGAW1","10S","INS","BOLD:ACL2677","Arthropoda","Insecta","Psocodea","Mesopsocidae","","Mesopsocus","Mesopsocus laticeps","1","1","1","132","0","0","99.78","450","0","450","40.67","0"
"AGAW1","10S","INS","BOLD:AAB4657","Arthropoda","Insecta","Diptera","Chironomidae","Chironominae","Chironomus","Chironomus maturus","1","1","1","132","0","0","98.22","450","0","450","37.11","0"
"AGAW1","10S","INS","BOLD:ACL1796","Arthropoda","Insecta","Lepidoptera","Ypsolophidae","Ochsenheimeriinae","Ochsenheimeria","Ochsenheimeria vacculella","1","1","1","132","0","0","99.55","448","0","450","32.67","0"
"AGAW1","10S","INS","BOLD:AAA4213","Arthropoda","Insecta","Lepidoptera","Geometridae","Sterrhinae","Idaea","Idaea dimidiata","1","1","1","132","0","0","97.32","447","0","450","30","0"
"AGAW1","10S","INS","BOLD:ABX9225","Arthropoda","Insecta","Diptera","Limoniidae","Limoniinae","Rhipidia","","1","1","1","132","0","0","97.51","442","0","444","33.33","0"
"AGAW1","10S","INS","BOLD:ACE7482","Arthropoda","Insecta","Hemiptera","Miridae","","","","1","1","1","132","0","0","98.08","416","0","416","35.1","0"
"AGAW1","10S","INS","BOLD:AAV5575","Arthropoda","Insecta","Diptera","Cecidomyiidae","","","CecidInt30 sp. BOLD:AAV5575","1","1","1","132","0","0","98.45","451","0","450","26","0"
"AGAW1","10S","INS","BOLD:AAA2632","Arthropoda","Insecta","Lepidoptera","Noctuidae","Noctuinae","Noctua","Noctua atlantica, Noctua pronuba","1","1","1","132","0","0","98","449","0","450","31.33","0"
"AGAW1","10S","INS","BOLD:ACK7314","Arthropoda","Insecta","Diptera","Chironomidae","Chironominae","Polypedilum","","1","1","1","132","0","0","97.22","360","0","364","35.44","0"
"AGAW1","10S","INS","BOLD:AAB9652","Arthropoda","Insecta","Trichoptera","Leptoceridae","Leptocerinae","Triaenodes","Triaenodes nox","1","1","1","132","0","0","97.99","448","0","450","32","0"
"AGAW1","10S","INS","BOLD:AAM9117","Arthropoda","Insecta","Hymenoptera","Ichneumonidae","Cryptinae","","","1","1","1","132","0","0","99.77","428","0","450","28.67","0"
"AGAW1","10S","INS","BOLD:ACJ6954","Arthropoda","Insecta","Diptera","Cecidomyiidae","","","","1","1","1","132","0","0","100","339","0","389","25.45","0"
"AGAW1","10S","INS","BOLD:ACV5891","Arthropoda","Insecta","Diptera","Cecidomyiidae","","","","1","1","1","132","0","0","98.61","360","0","450","25.78","0"
"AGAW1","10S","INS","BOLD:AAA8796","Arthropoda","Insecta","Lepidoptera","Tortricidae","Tortricinae","Acleris","Acleris forsskaleana","1","1","1","132","0","0","98.66","372","0","372","31.99","0"
"AGAW1","10S","INS","BOLD:ABV1453","Arthropoda","Insecta","Diptera","Cecidomyiidae","","","","1","1","1","132","0","0","98.65","371","0","376","27.13","0"
"AGAW1","10S","INS","BOLD:AAI1549","Arthropoda","Insecta","Hymenoptera","Braconidae","Euphorinae","Meteorus","","1","1","1","132","0","0","97.99","447","0","450","31.11","0"
"AGAW1","10S","INS","BOLD:AAB7030","Arthropoda","Insecta","Diptera","Chironomidae","Chironominae","Chironomus","Chironomus sp. BOLDAAP2996, Chironomus quinnitukqut, Chironomus cf. decorus 2g, Chironomus decorus group sp. 2 IP2013, Chironomus cf. decorus, Chironomus decorus gr.","1","1","1","132","0","0","97.34","413","0","415","33.73","0"
"AGAW1","10S","INS","BOLD:AAM6286","Arthropoda","Insecta","Diptera","Chironomidae","Chironominae","","","1","1","1","132","0","0","97.87","376","0","378","32.28","0"
"AGAW1","10S","INS","BOLD:AAA3877","Arthropoda","Insecta","Trichoptera","Hydroptilidae","Hydroptilinae","Agraylea","Agraylea multipunctata","1","1","1","132","0","0","98.89","449","0","450","39.56","0"
"AGAW1","10S","INS","BOLD:AAF9187","Arthropoda","Insecta","Coleoptera","Curculionidae","Entiminae","Phyllobius","Phyllobius oblongus","1","1","1","132","0","0","99.29","425","0","426","39.2","0"
"AGAW1","10S","INS","BOLD:AAN5260","Arthropoda","Insecta","Diptera","Cecidomyiidae","","","","1","1","1","132","0","0","97.6","416","0","419","25.54","0"
"AGAW1","10S","INS","BOLD:AAN5342","Arthropoda","Insecta","Diptera","Chironomidae","Orthocladiinae","Heleniella","","1","1","1","132","0","0","98.21","447","0","450","35.11","0"
"AGAW1","10S","INS","BOLD:AAU6966","Arthropoda","Insecta","Coleoptera","Throscidae","Throscinae","Trixagus","Trixagus chevrolati","1","1","1","132","0","0","99.51","412","0","450","39.56","0"
"AGAW1","10S","INS","BOLD:AAC0635","Arthropoda","Insecta","Hemiptera","Miridae","Mirinae","Stenotus","Stenotus binotatus","1","1","1","132","0","0","97.78","450","0","450","34.67","0"
"AGAW1","10S","INS","BOLD:AAG6743","Arthropoda","Insecta","Diptera","Sarcophagidae","Sarcophaginae","Sarcophaga","Sarcophaga sp. 1 MG, Sarcophaga subvicina","1","1","1","132","0","0","99.55","447","0","450","35.33","0"
"AGAW1","10S","INS","BOLD:AAG6468","Arthropoda","Insecta","Diptera","Ceratopogonidae","Ceratopogoninae","Culicoides","Culicoides biguttatus","1","1","1","132","0","0","98","450","0","450","36.89","0"
"AGAW1","10S","INS","BOLD:ACZ5284","Arthropoda","Insecta","Diptera","Sciomyzidae","Sciomyzinae","Dictya","Dictya stricta, Dictya expansa, Dictya floridensis, Dictya borealis, Dictya gaigei, Dictya sp. TAW1, Dictya umbroides, Dictya texensis, Dictya steyskali, Dictya pictipes","1","1","1","132","0","0","99.72","360","0","361","35.73","0"
"AGAW1","10S","INS","BOLD:AAD4214","Arthropoda","Insecta","Hymenoptera","Ichneumonidae","Diplazontinae","Diplazon","Diplazon laetatorius","1","1","1","132","0","0","99.78","450","0","450","25.56","0"
"AGAW1","10S","INS","BOLD:AAU6327","Arthropoda","Insecta","Diptera","Chironomidae","Orthocladiinae","Psectrocladius","Psectrocladius schlienzi","1","1","1","132","0","0","97.75","311","0","450","38","0"
"AGAW1","10S","INS","BOLD:AAN8583","Arthropoda","Insecta","Diptera","Mycetophilidae","Sciophilinae","Sciophila","Sciophila pseudoflexuosa, Sciophila lutea","1","1","1","132","0","0","97.76","447","0","450","33.11","0"
"AGAW1","10S","INS","BOLD:ABX6186","Arthropoda","Insecta","Diptera","Tipulidae","Tipulinae","Nephrotoma","Nephrotoma ferruginea","1","1","1","132","0","0","99.11","450","0","450","35.56","0"
"AGAW1","10S","INS","BOLD:AAB2384","Arthropoda","Insecta","Diptera","Syrphidae","Syrphinae","Eupeodes","Eupeodes duseki, Eupeodes volucris, Eupeodes nielseni, Eupeodes americanus, Eupeodes flukei, Eupeodes vockerothi, Eupeodes lucasi, Eupeodes montivagus, Eupeodes rufipunctatus, Eupeodes rojasi, Eupeodes goeldlini, Eupeodes luniger, Eupeodes punctifer, Eupeodes bucculatus, Eupeodes perplexus, Eupeodes abiskoensis, Eupeodes corollae, Eupeodes pomus, Eupeodes lundbecki, Eupeodes nitens, Eupeodes sp. BOLDAAB2384, Eupeodes curtus, Eupeodes latifasciatus","1","1","1","132","0","0","99.5","404","0","407","29.98","0"
"AGAW1","10S","INS","BOLD:ACD8563","Arthropoda","Insecta","Diptera","Cecidomyiidae","","","","1","1","1","132","0","0","97.11","450","0","450","25.78","0"
"AGAW1","10S","INS","BOLD:AAA3654","Arthropoda","Insecta","Lepidoptera","Crambidae","Crambinae","Crambus","Crambus bigelovi, Crambus johnsoni, Crambus praefectellus","1","1","1","132","0","0","97.49","399","0","401","29.93","0"
"AGAW1","10S","INS","BOLD:ACP4736","Arthropoda","Insecta","Diptera","Chironomidae","Orthocladiinae","Smittia","Smittia sp. 8ES","1","1","1","132","0","0","99.78","445","0","450","33.33","0"
"AGAW1","10S","INS","BOLD:AAZ0271","Arthropoda","Insecta","Diptera","Ceratopogonidae","Ceratopogoninae","Culicoides","","1","1","1","132","0","0","98.44","448","0","450","42","0"
"AGAW1","10S","INS","BOLD:AAH4205","Arthropoda","Insecta","Diptera","Chloropidae","Chloropinae","Meromyza","Meromyza nigriseta, Meromyza sp.","1","1","1","132","0","0","99.33","447","0","450","32.67","0"
"AGAW1","10S","INS","BOLD:AAU8628","Arthropoda","Insecta","Hymenoptera","Bethylidae","","","","1","1","1","132","0","0","99.78","449","0","450","27.11","0"
"AGAW1","10S","INS","BOLD:ACF4940","Arthropoda","Insecta","Hymenoptera","Ichneumonidae","Campopleginae","Campoletis","Campoletis sp. AB.2","1","1","1","132","0","0","99.11","450","0","450","25.33","0"
"AGAW1","10S","INS","BOLD:ABV1422","Arthropoda","Insecta","Diptera","Cecidomyiidae","","","","1","1","1","132","0","0","99.11","449","0","450","27.78","0"
"AGAW1","10S","INS","BOLD:AAA5953","Arthropoda","Insecta","Lepidoptera","Gelechiidae","Gelechiinae","Coleotechnites","Coleotechnites sp., Coleotechnites piceaella, Coleotechnites laricis, Coleotechnites sp. 3, Coleotechnites carbonaria, Coleotechnites granti, Coleotechnites macleodi, Coleotechnites gibsonella, Coleotechnites coniferella, Coleotechnites martini","1","1","1","132","0","0","98","449","0","450","34.22","0"
"AGAW1","10S","INS","BOLD:AAA7374","Arthropoda","Insecta","Diptera","Syrphidae","Syrphinae","Sphaerophoria","Sphaerophoria bankowskae, Sphaerophoria infuscata, Sphaerophoria nr. philanthus, Sphaerophoria batava, Sphaerophoria sp., Sphaerophoria rueppellii, Sphaerophoria interrupta-group, Sphaerophoria sp. MA-1, Sphaerophoria cleoae, Sphaerophoria asymmetrica, Sphaerophoria fatarum, Sphaerophoria bengalensis, Sphaerophoria bifurcata, Sphaerophoria taeniata, Sphaerophoria interrupta, Sphaerophoria contigua, Sphaerophoria kaa, Sphaerophoria laurae, Sphaerophoria chongjini, Sphaerophoria boreoalpina, Sphaerophoria sp. BOLDAAA7374, Sphaerophoria macrogaster, Sphaerophoria sulphuripes, Sphaerophoria brevipilosa, Sphaerophoria philanthus, Sphaerophoria pyrrhina, Sphaerophoria abbreviata, Sphaerophoria scripta, Sphaerophoria potentillae, Sphaerophoria longipilosa, Sphaerophoria virgata","1","1","1","132","0","0","98.34","421","0","424","30.19","0"
"AGAW1","10S","NAR","no match","no match","no match","no match","no match","no match","no match","no match","no match","no match","no match","no match","no match","no match","no match","no match","no match","no match","no match","no match"
"AGAW1","10S","NIN","BOLD:AAA7242","Arthropoda","Collembola","Entomobryomorpha","Entomobryidae","Seirinae, Entomobryinae","Entomobrya, Lepidosira","Entomobrya multifasciata, Lepidosira sp.","344","326","326","1","5.23","0","99.23","427.04","0","429.79","39.4","0"
"AGAW1","10S","NIN","BOLD:AAU6252","Arthropoda","Arachnida","Trombidiformes","Bdellidae","","Cyta","","130","111","111","2","14.62","0","98.29","438.1","0","439.59","31.5","0"
"AGAW1","10S","NIN","BOLD:AAB7914","Arthropoda","Collembola","Symphypleona","Bourletiellidae","","Bourletiella","","22","22","22","3","0","0","99.37","435.18","0","436.91","37.98","0"
"AGAW1","10S","NIN","BOLD:ACL6413","Arthropoda","Arachnida","Trombidiformes","Erythraeidae","","Balaustium","","20","17","17","4","15","0","99.37","439.2","0","440.2","35.85","0"
"AGAW1","10S","NIN","BOLD:ACY2550","Arthropoda","Arachnida","Trombidiformes","Bdellidae","","Cyta","","16","14","14","5","12.5","0","98.82","378.75","0","399.5","31.23","0"
"AGAW1","10S","NIN","BOLD:ABW2639","Arthropoda","Arachnida","Trombidiformes","Anystidae","","","","10","10","10","6","0","0","99.01","443.5","0","441.6","31.4","0"
"AGAW1","10S","NIN","BOLD:ACZ1295","Arthropoda","Arachnida","Trombidiformes","Anystidae","","","","3","3","3","7","0","0","99.3","375.67","0","390.67","31.18","0"
"AGAW1","10S","NIN","BOLD:AAH6630","Arthropoda","Collembola","Entomobryomorpha","Entomobryidae","Entomobryinae","Entomobrya","Entomobrya unostrigata","2","2","2","8","0","0","99.73","411","0","412","38.61","0"
"AGAW1","10W","BAC","TAX:175768","Proteobacteria","Alphaproteobacteria","Rickettsiales","Rickettsiaceae",NA,"Wolbachia","Wolbachia pipientis","37","32","32","1","13.51","0","98.37","448.46","0","449.7","39.7","0"
"AGAW1","10W","BAC","TAX:91393","Proteobacteria","Alphaproteobacteria","Rickettsiales","Rickettsiaceae",NA,"Wolbachia","","20","20","20","2","0","0","99.42","389.75","0","413.85","39.16","0"
"AGAW1","10W","BAC","TAX:91391","Proteobacteria","Alphaproteobacteria","Rickettsiales","",NA,"","","8","8","8","3","0","0","98.59","412.75","0","441.63","40.1","0"
"AGAW1","10W","BAC","TAX:703648","Proteobacteria","Alphaproteobacteria","Rickettsiales","Rickettsiaceae",NA,"Wolbachia","Wolbachia endosymbiont","1","1","1","4","0","0","97.72","395","0","450","39.78","0"
"AGAW1","10W","BAC","TAX:375853","Proteobacteria","Alphaproteobacteria","Rickettsiales","Rickettsiaceae",NA,"Wolbachia","Wolbachia sp. Hymenoptera_ASGLE646-10","1","1","1","4","0","0","98.44","450","0","450","40.22","0"
"AGAW1","10W","CHO","no match","no match","no match","no match","no match","no match","no match","no match","no match","no match","no match","no match","no match","no match","no match","no match","no match","no match","no match","no match"
"AGAW1","10W","INS","BOLD:AAG1711","Arthropoda","Insecta","Diptera","Muscidae","Phaoniinae","Helina","Helina depuncta","15893","12174","12174","1","23.4","0","99.43","433.51","0","435.83","31.37","0"
"AGAW1","10W","INS","BOLD:AAU6681","Arthropoda","Insecta","Diptera","Tipulidae","Tipulinae","Nephrotoma","Nephrotoma cornicina","15178","10611","10611","2","30.09","0","99.62","443.05","0","444.22","36.32","0"
"AGAW1","10W","INS","BOLD:AAA3453","Arthropoda","Insecta","Diptera","Anthomyiidae","Anthomyiinae","Delia","","7068","6156","6156","3","12.9","0","99.39","431.37","0","433.99","34.65","0"
"AGAW1","10W","INS","BOLD:ACD9117","Arthropoda","Insecta","Hemiptera","Cicadellidae","Eurymelinae","Idiocerus","Idiocerus levis","5809","4864","4864","4","16.27","0","99.49","349.63","0","422.94","32.44","0"
"AGAW1","10W","INS","BOLD:ACE0699","Arthropoda","Insecta","Diptera","Chironomidae","","","","3773","2551","2551","5","32.39","0","98.59","425.31","0","442.08","36.6","0"
"AGAW1","10W","INS","BOLD:AAP8739","Arthropoda","Insecta","Diptera","Platystomatidae","Platystomatinae","Rivellia","Rivellia steyskali","3374","2460","2460","6","27.09","0","99.36","441.33","0","443.21","37.95","0"
"AGAW1","10W","INS","BOLD:AAG1748","Arthropoda","Insecta","Diptera","Muscidae","Coenosiinae","Coenosia","","2932","2325","2325","7","20.7","0","99.25","440.47","0","441.86","34.36","0"
"AGAW1","10W","INS","BOLD:AAN6154","Arthropoda","Insecta","Coleoptera","Latridiidae","Corticariinae","Melanophthalma","Melanophthalma inermis","2521","2032","2032","8","19.4","0","98.81","437.32","0","438.75","31.37","0"
"AGAW1","10W","INS","BOLD:AAH3661","Arthropoda","Insecta","Diptera","Cecidomyiidae","","","","2219","1941","1941","9","12.53","0","99.28","432.32","0","434.22","25.4","0"
"AGAW1","10W","INS","BOLD:ABA0822","Arthropoda","Insecta","Diptera","Cecidomyiidae","","","","1603","1380","1380","10","13.91","0","98.97","426.73","0","427.78","25.79","0"
"AGAW1","10W","INS","BOLD:ACE7444","Arthropoda","Insecta","Hemiptera","Miridae","Mirinae, Phylinae","Adelphocoris, Megalocoleus","Adelphocoris lineolatus, Megalocoleus molliculus","1518","997","997","11","34.32","0","99.68","440.95","0","441.66","34.79","0"
"AGAW1","10W","INS","BOLD:AAI4303","Arthropoda","Insecta","Diptera","Chironomidae","Chironominae","Chironomus","Chironomus melanescens","1498","1327","1327","12","11.42","0","98.02","430.47","0","436.33","33.65","0"
"AGAW1","10W","INS","BOLD:AAG5507","Arthropoda","Insecta","Diptera","Chironomidae","Chironominae","Chironomus","Chironomus atrella","1495","1456","1456","13","2.61","0","99.05","425.78","0","432.48","35.94","0"
"AGAW1","10W","INS","BOLD:AAG2511","Arthropoda","Insecta","Diptera","Anthomyiidae","Anthomyiinae","Delia","Delia platura","1493","1333","1333","14","10.72","0","98.98","445.04","0","447.84","33.16","0"
"AGAW1","10W","INS","BOLD:AAG2868","Arthropoda","Insecta","Hemiptera","Cicadellidae","Typhlocybinae","Empoasca","Empoasca sp., Empoasca fabae","1359","847","847","15","37.67","0","99.68","443.85","0","447.69","32.44","0"
"AGAW1","10W","INS","BOLD:ABX0657","Arthropoda","Insecta","Coleoptera","Anthicidae","Notoxinae","Notoxus","Notoxus desertus","1238","1158","1158","16","6.46","0","98.84","433.94","0","435.18","35.97","0"
"AGAW1","10W","INS","BOLD:AAP5169","Arthropoda","Insecta","Diptera","Chloropidae","Oscinellinae","Elachiptera","Elachiptera nigriceps","1178","920","920","17","21.9","0","99.23","429.47","0","430.56","31.66","0"
"AGAW1","10W","INS","BOLD:AAM6286","Arthropoda","Insecta","Diptera","Chironomidae","Chironominae","","","780","747","747","18","4.23","0","99.11","407.64","0","410.17","34.02","0"
"AGAW1","10W","INS","BOLD:ABW7897","Arthropoda","Insecta","Diptera","Cecidomyiidae","","","","681","493","493","19","27.61","0","99.58","443.08","0","445.36","29.67","0"
"AGAW1","10W","INS","BOLD:AAA2326","Arthropoda","Insecta","Hymenoptera","Apidae","Apinae","Apis","Apis mellifera","610","430","430","20","29.51","0","99.6","447.8","0","448.66","26.43","0"
"AGAW1","10W","INS","BOLD:AAH0135","Arthropoda","Insecta","Coleoptera","Phalacridae","Phalacrinae","Acylomus","Acylomus pugetanus, Acylomus ergoti","603","526","526","21","12.77","0","99.3","436.28","0","438.23","31.66","0"
"AGAW1","10W","INS","BOLD:AAA7661","Arthropoda","Insecta","Diptera","Culicidae","Culicinae","Culiseta, Culex","Culiseta melanura, Culex articularis, Culex restuans, Culex acharistus","599","550","550","22","8.18","0","99.17","434.17","0","436.2","32.93","0"
"AGAW1","10W","INS","BOLD:AAC3517","Arthropoda","Insecta","Hemiptera","Delphacidae","Delphacinae","Kosswigianella, Muirodelphax, Delphacodes, Nothodelphax, Elachodelphax","Muirodelphax arvensis, Delphacodes nitens, Delphacodes lutulenta, Muirodelphax nigrinota, Kosswigianella analis, Delphacodes campestris, Muirodelphax anufrievi, Delphacodes indistincta, Muirodelphax unda, Muirodelphax nr. arvensis, Delphacodes stramineosa, Nothodelphax consimilis, Muirodelphax atrolabis, Muirodelphax atralabis, Kosswigianella lutulenta","458","367","367","23","19.87","0","99.47","442.46","0","443.41","36.98","0"
"AGAW1","10W","INS","BOLD:AAG9656","Arthropoda","Insecta","Diptera","Ephydridae","Notiphilinae","Hydrellia","Hydrellia notata, Hydrellia sp. 2 AGT","446","406","406","24","8.97","0","98.49","435.46","0","439.84","34.76","0"
"AGAW1","10W","INS","BOLD:ACF2821","Arthropoda","Insecta","Diptera","Mycetophilidae","Mycetophilinae","Mycetophila","Mycetophila fungorum","436","382","382","25","12.39","0","99.21","430.2","0","433","35.25","0"
"AGAW1","10W","INS","BOLD:AAN5199","Arthropoda","Insecta","Diptera","Cecidomyiidae","","","CecidInt20 sp. BOLD:AAN5199","430","340","340","26","20.93","0","99.41","435.9","0","444.38","26.66","0"
"AGAW1","10W","INS","BOLD:ABA0863","Arthropoda","Insecta","Diptera","Cecidomyiidae","","","","389","312","312","27","19.79","0","99.36","440.96","0","442.48","29.68","0"
"AGAW1","10W","INS","BOLD:ACK3431","Arthropoda","Insecta","Diptera","Cecidomyiidae","","","","322","300","300","28","6.83","0","99.28","424.05","0","425.71","32.61","0"
"AGAW1","10W","INS","BOLD:AAN5238","Arthropoda","Insecta","Diptera","Cecidomyiidae","","","","234","192","192","29","17.95","0","99.22","429.28","0","430.69","25.66","0"
"AGAW1","10W","INS","BOLD:AAN5183","Arthropoda","Insecta","Diptera","Cecidomyiidae","","","","232","196","196","30","15.52","0","98.46","443.06","0","444.06","27.85","0"
"AGAW1","10W","INS","BOLD:AAH3664","Arthropoda","Insecta","Diptera","Cecidomyiidae","","","CecidInt3 sp. BOLD:AAH3664","199","186","186","31","6.53","0","99.16","433.16","0","437","27.03","0"
"AGAW1","10W","INS","BOLD:AAP5113","Arthropoda","Insecta","Diptera","Chironomidae","Tanypodinae","","","181","167","167","32","7.73","0","97.55","426.92","0","428.99","37.23","0"
"AGAW1","10W","INS","BOLD:ABV1297","Arthropoda","Insecta","Diptera","Sciaridae","","","","145","141","141","33","2.76","0","99.08","431.42","0","433.88","30.19","0"
"AGAW1","10W","INS","BOLD:ACK8384","Arthropoda","Insecta","Hemiptera","Miridae","Phylinae","Phoenicocoris","Phoenicocoris rostratus","141","127","127","34","9.93","0","99.19","439.88","0","441.05","38.41","0"
"AGAW1","10W","INS","BOLD:AAD2041","Arthropoda","Insecta","Diptera","Dolichopodidae","Sciapodinae","Condylostylus","Condylostylus inermis, Condylostylus nigrofemoratus, Condylostylus connectans, Condylostylus caudatus, Condylostylus flavipes","136","104","104","35","23.53","0","99.5","440.26","0","441.36","35.94","0"
"AGAW1","10W","INS","BOLD:AAG7276","Arthropoda","Insecta","Diptera","Sphaeroceridae","Limosininae","Leptocera","Leptocera erythrocera","125","115","115","36","8","0","99.28","432.74","0","435.66","32.78","0"
"AGAW1","10W","INS","BOLD:AAN5260","Arthropoda","Insecta","Diptera","Cecidomyiidae","","","","123","117","117","37","4.88","0","99.09","411.23","0","412.81","25.98","0"
"AGAW1","10W","INS","BOLD:ADF6661","Arthropoda","Insecta","Diptera","Cecidomyiidae","","","","116","98","98","38","15.52","0","99.23","416.67","0","444.82","29.74","0"
"AGAW1","10W","INS","BOLD:AAN5242","Arthropoda","Insecta","Diptera","Cecidomyiidae","","","","106","105","105","39","0.94","0","98.98","406.92","0","408.69","28.25","0"
"AGAW1","10W","INS","BOLD:ABA9076","Arthropoda","Insecta","Coleoptera","Latridiidae","Latridiinae","Cartodere","Cartodere constricta","101","89","89","40","11.88","0","99.16","434.09","0","435.57","35.25","0"
"AGAW1","10W","INS","BOLD:AAG5443","Arthropoda","Insecta","Diptera","Chironomidae","Orthocladiinae","","","96","96","96","41","0","0","98.87","409.6","0","411.78","33.3","0"
"AGAW1","10W","INS","BOLD:AAN5205","Arthropoda","Insecta","Diptera","Cecidomyiidae","","","CecidInt42 sp. BOLD:AAN5205","91","83","83","42","8.79","0","99.23","440.44","0","441.8","31.4","0"
"AGAW1","10W","INS","BOLD:AAH0134","Arthropoda","Insecta","Coleoptera","Phalacridae","Phalacrinae","Stilbus","Stilbus apicialis","90","88","88","43","2.22","0","99.22","425.14","0","426.4","31.81","0"
"AGAW1","10W","INS","BOLD:ABV9018","Arthropoda","Insecta","Diptera","Mycetophilidae","Sciophilinae","Sciophila","","89","84","84","44","5.62","0","99.15","439.67","0","441.94","31.96","0"
"AGAW1","10W","INS","BOLD:AAN5230","Arthropoda","Insecta","Diptera","Cecidomyiidae","","","","89","85","85","44","4.49","0","98.99","423.82","0","425.47","25.84","0"
"AGAW1","10W","INS","BOLD:AAH4171","Arthropoda","Insecta","Diptera","Chloropidae","Oscinellinae","Apallates","Apallates montanus, Apallates particeps","81","74","74","46","8.64","0","98.93","436.73","0","438.42","32.79","0"
"AGAW1","10W","INS","BOLD:AAN5220","Arthropoda","Insecta","Diptera","Cecidomyiidae","","","","74","74","74","47","0","0","98.14","399.03","0","422.54","26.29","0"
"AGAW1","10W","INS","BOLD:AAF9014","Arthropoda","Insecta","Diptera","Limoniidae","Chioneinae","Symplecta","Symplecta cana, Symplecta sp. BOLDAAF9014","66","61","61","48","7.58","0","98.65","425.67","0","431.05","33.76","0"
"AGAW1","10W","INS","BOLD:ABW1379","Arthropoda","Insecta","Diptera","Chloropidae","Oscinellinae","Malloewia","Malloewia abdominalis","65","54","54","49","16.92","0","98.22","442.57","0","448.38","31.72","0"
"AGAW1","10W","INS","BOLD:AAN6447","Arthropoda","Insecta","Diptera","Sciaridae","","Corynoptera","Corynoptera perpusilla","59","51","51","50","13.56","0","99.4","448.24","0","449.66","36.49","0"
"AGAW1","10W","INS","BOLD:AAG4905","Arthropoda","Insecta","Diptera","Mycetophilidae","Mycetophilinae","Mycetophila","","54","50","50","51","7.41","0","99.35","418.02","0","419.37","35.41","0"
"AGAW1","10W","INS","BOLD:ABA0845","Arthropoda","Insecta","Diptera","Cecidomyiidae","","","","52","47","47","52","9.62","0","99.16","446.17","0","447.79","30.94","0"
"AGAW1","10W","INS","BOLD:AAD1485","Arthropoda","Insecta","Diptera","Chironomidae","Chironominae","Paratanytarsus","Paratanytarsus grimmii, Paratanytarsus sp.","49","45","45","53","8.16","0","99.36","441.94","0","443.88","33.79","0"
"AGAW1","10W","INS","BOLD:AAH4276","Arthropoda","Insecta","Lepidoptera","Gelechiidae","Gelechiinae","Bryotropha","Bryotropha hodgesi, Bryotropha altitudophila","41","38","38","54","7.32","0","99.46","435.85","0","437.73","33.53","0"
"AGAW1","10W","INS","BOLD:AAN5255","Arthropoda","Insecta","Diptera","Cecidomyiidae","Cecidomyiinae","Dasineura","","38","35","35","55","7.89","0","98.49","421.55","0","439.66","26.67","0"
"AGAW1","10W","INS","BOLD:ACK3469","Arthropoda","Insecta","Coleoptera","Chrysomelidae","Eumolpinae","Brachypnoea","Brachypnoea puncticollis","37","30","30","56","18.92","0","99.26","441.68","0","442.86","36.35","0"
"AGAW1","10W","INS","BOLD:ACJ8120","Arthropoda","Insecta","Diptera","Cecidomyiidae","","","","35","30","30","57","14.29","0","99.66","391.69","0","442.6","25.86","0"
"AGAW1","10W","INS","BOLD:AAF1729","Arthropoda","Insecta","Psocodea","Lachesillidae","Lachesillinae","Lachesilla","Lachesilla pedicularia","35","31","31","57","11.43","0","99.42","443.14","0","444.63","36.92","0"
"AGAW1","10W","INS","BOLD:AAL7819","Arthropoda","Insecta","Diptera","Psychodidae","Psychodinae","Psychoda","Psychoda sp., Psychoda sp. BOLD:AAL7819","32","32","32","59","0","0","99.35","413.22","0","414.91","32.76","0"
"AGAW1","10W","INS","BOLD:AAY6730","Arthropoda","Insecta","Hemiptera","Derbidae","Cedusinae","Cedusa","Cedusa incisa","27","27","27","60","0","0","99.02","425.85","0","427.59","35.37","0"
"AGAW1","10W","INS","BOLD:ABU5525","Arthropoda","Insecta","Diptera","Chironomidae","Orthocladiinae","Limnophyes","Limnophyes sp. 14ES","27","27","27","60","0","0","98.88","397.41","0","399.89","37.03","0"
"AGAW1","10W","INS","BOLD:ACC8079","Arthropoda","Insecta","Diptera","Cecidomyiidae","","","","26","23","23","62","11.54","0","99.33","444.73","0","446.15","29.18","0"
"AGAW1","10W","INS","BOLD:AAM7398","Arthropoda","Insecta","Hymenoptera","Ichneumonidae","Campopleginae","Hyposoter","","24","23","23","63","4.17","0","99.33","427","0","441.58","26.63","0"
"AGAW1","10W","INS","BOLD:ABA7871","Arthropoda","Insecta","Diptera","Cecidomyiidae","","","","23","23","23","64","0","0","98.88","439.22","0","440.26","26.75","0"
"AGAW1","10W","INS","BOLD:ABA1215","Arthropoda","Insecta","Diptera","Sciaridae","","Lycoriella","Lycoriella sativae","22","19","19","65","13.64","0","99.38","440.68","0","441.95","35.81","0"
"AGAW1","10W","INS","BOLD:AAG7891","Arthropoda","Insecta","Hymenoptera","Platygastridae","Scelioninae","Telenomus","Telenomus podisi","21","21","21","66","0","0","98.97","426.14","0","433.24","27.04","0"
"AGAW1","10W","INS","BOLD:ABX3229","Arthropoda","Insecta","Coleoptera","Chrysomelidae","Eumolpinae","Colaspis","Colaspis brunnea","21","20","20","66","4.76","0","98.74","446.52","0","447.9","36.02","0"
"AGAW1","10W","INS","BOLD:ACV5892","Arthropoda","Insecta","Diptera","Cecidomyiidae","","","","19","19","19","68","0","0","98.07","439.26","0","440.42","25.13","0"
"AGAW1","10W","INS","BOLD:ACF2457","Arthropoda","Insecta","Diptera","Chironomidae","Chironominae","Paratanytarsus","Paratanytarsus laccophilus","17","17","17","69","0","0","99.16","441.59","0","442.59","32.61","0"
"AGAW1","10W","INS","BOLD:AAN5221","Arthropoda","Insecta","Diptera","Cecidomyiidae","","","","16","16","16","70","0","0","98.68","423.81","0","429.25","25.72","0"
"AGAW1","10W","INS","BOLD:AAN5267","Arthropoda","Insecta","Diptera","Cecidomyiidae","","","","15","12","12","71","20","0","99.7","449.33","0","450","30.56","0"
"AGAW1","10W","INS","BOLD:AAV1261","Arthropoda","Insecta","Diptera","Sciaridae","","Bradysia","Bradysia splendida","14","13","13","72","7.14","0","99.5","439.64","0","440.71","34.12","0"
"AGAW1","10W","INS","BOLD:AAY6469","Arthropoda","Insecta","Diptera","Cecidomyiidae","","","","14","14","14","72","0","0","99.05","428.5","0","429.5","26.06","0"
"AGAW1","10W","INS","BOLD:ABX4079","Arthropoda","Insecta","Diptera","Cecidomyiidae","","","","13","12","12","74","7.69","0","98.6","449.08","0","450","28.31","0"
"AGAW1","10W","INS","BOLD:ABA6407","Arthropoda","Insecta","Diptera","Sciaridae","","Pseudolycoriella","","12","12","12","75","0","0","99.37","433.08","0","435.75","34.78","0"
"AGAW1","10W","INS","BOLD:AAI1549","Arthropoda","Insecta","Hymenoptera","Braconidae","Euphorinae","Meteorus","","11","9","9","76","18.18","0","98.77","427.18","0","431.09","30.91","0"
"AGAW1","10W","INS","BOLD:ABX7480","Arthropoda","Insecta","Diptera","Cecidomyiidae","","","","10","10","10","77","0","0","98.08","410.7","0","433.6","27.26","0"
"AGAW1","10W","INS","BOLD:ACI9539","Arthropoda","Insecta","Diptera","Cecidomyiidae","Cecidomyiinae","Sitodiplosis","Sitodiplosis sp. BOLD:ACI9539","10","10","10","77","0","0","99.16","412.3","0","414.7","26.29","0"
"AGAW1","10W","INS","BOLD:ADG5727","Arthropoda","Insecta","Diptera","Cecidomyiidae","","","","10","10","10","77","0","0","98.89","397.4","0","429.9","30.1","0"
"AGAW1","10W","INS","BOLD:AAM6097","Arthropoda","Insecta","Diptera","Cecidomyiidae","","","","9","9","9","80","0","0","99.22","419.33","0","420.89","34.13","0"
"AGAW1","10W","INS","BOLD:ACV1014","Arthropoda","Insecta","Diptera","Cecidomyiidae","","","","9","9","9","80","0","0","97.85","401.89","0","404.33","25.23","0"
"AGAW1","10W","INS","BOLD:AAB0373","Arthropoda","Insecta","Neuroptera","Chrysopidae","Chrysopinae","Chrysoperla","Chrysoperla sp. isolate 82, Chrysoperla zastrowi, Chrysoperla plorabunda, Chrysoperla sp. isolate 71, Chrysoperla agilis, Italochrysa italica, Chrysoperla sp. isolate 88, Chrysoperla lucasina, Chrysoperla carnea, Chrysoperla mediterranea, Chrysoperla pallida, Chrysoperla nipponensis, Chrysoperla sillemi","9","9","9","80","0","0","99","435","0","436.44","31.43","0"
"AGAW1","10W","INS","BOLD:ACK5742","Arthropoda","Insecta","Diptera","Cecidomyiidae","","","","8","8","8","83","0","0","99.52","445.25","0","446.75","29.06","0"
"AGAW1","10W","INS","BOLD:AAN5334","Arthropoda","Insecta","Diptera","Chironomidae","Chironominae","Polypedilum","","7","7","7","84","0","0","98.06","435.71","0","439.57","34.69","0"
"AGAW1","10W","INS","BOLD:AAZ5707","Arthropoda","Insecta","Hymenoptera","Dryinidae","","","","7","7","7","84","0","0","99.23","371.29","0","374","24.63","0"
"AGAW1","10W","INS","BOLD:ADI0927","","","","","","","","7","5","5","84","28.57","0","97.62","444.71","0","445.14","34.33","0"
"AGAW1","10W","INS","BOLD:ACX6289","Arthropoda","Insecta","Diptera","Phoridae","Metopininae","Megaselia","","7","7","7","84","0","0","99.18","425.57","0","426.71","37.99","0"
"AGAW1","10W","INS","BOLD:AAM9255","Arthropoda","Insecta","Diptera","Sciaridae","","Pseudolycoriella","","7","7","7","84","0","0","99.28","417.43","0","419.71","30.72","0"
"AGAW1","10W","INS","BOLD:AAF5980","Arthropoda","Insecta","Hemiptera","Cicadellidae","Typhlocybinae","Typhlocyba","Typhlocyba pomaria","6","6","6","89","0","0","97.93","431.17","0","431.67","37.12","0"
"AGAW1","10W","INS","BOLD:AAF9306","Arthropoda","Insecta","Diptera","Psychodidae","Psychodinae","Psychoda","Psychoda alternata","6","6","6","89","0","0","97.54","447.67","0","448.83","31.56","0"
"AGAW1","10W","INS","BOLD:ACV5891","Arthropoda","Insecta","Diptera","Cecidomyiidae","","","","6","6","6","89","0","0","99.08","361.17","0","418.33","24.99","0"
"AGAW1","10W","INS","BOLD:ABV1422","Arthropoda","Insecta","Diptera","Cecidomyiidae","","","","6","6","6","89","0","0","98.52","413.5","0","414.67","26.59","0"
"AGAW1","10W","INS","BOLD:AAH3662","Arthropoda","Insecta","Diptera","Cecidomyiidae","","","CecidInt3 sp. BOLD:AAH3662","5","5","5","93","0","0","98.57","449.2","0","450","26.71","0"
"AGAW1","10W","INS","BOLD:ACJ9529","Arthropoda","Insecta","Psocodea","Liposcelidae","Liposcelidinae","Liposcelis","Liposcelis paeta","5","5","5","93","0","0","99.38","447.8","0","449","39.91","0"
"AGAW1","10W","INS","BOLD:AAP6827","Arthropoda","Insecta","Diptera","Cecidomyiidae","","","","4","4","4","95","0","0","98.7","444.5","0","448","31.53","0"
"AGAW1","10W","INS","BOLD:AAA5299","Arthropoda","Insecta","Diptera","Chironomidae","Orthocladiinae","Cricotopus","Cricotopus laetus, Cricotopus relucens, Cricotopus trifasciatus, Cricotopus sp., Cricotopus sp. 3ES, Cricotopus sylvestris, Cricotopus glacialis","4","4","4","95","0","0","99.5","449","0","450","34.72","0"
"AGAW1","10W","INS","BOLD:ACU6062","Arthropoda","Insecta","Diptera","Cecidomyiidae","","","","4","4","4","95","0","0","98.72","448.75","0","450","26.61","0"
"AGAW1","10W","INS","BOLD:ACF1257","","","","","","","","4","4","4","95","0","0","99.27","448.75","0","450","34.94","0"
"AGAW1","10W","INS","BOLD:ADH9867","Arthropoda","Insecta","Diptera","","","","","4","4","4","95","0","0","99.11","364.25","0","367.5","37.96","0"
"AGAW1","10W","INS","BOLD:AAM6257","Arthropoda","Insecta","Diptera","Chironomidae","Orthocladiinae","","","3","3","3","100","0","0","99.11","448","0","448.33","36.36","0"
"AGAW1","10W","INS","BOLD:ACV2677","Arthropoda","Insecta","Diptera","Cecidomyiidae","","","","3","3","3","100","0","0","97.48","450","0","450","30.22","0"
"AGAW1","10W","INS","BOLD:AAH0256","Arthropoda","Insecta","Coleoptera","Latridiidae","Corticariinae","Corticaria, Corticarina","","3","3","3","100","0","0","99.3","431.33","0","441","31.89","0"
"AGAW1","10W","INS","BOLD:AAG6745","Arthropoda","Insecta","Diptera","Calliphoridae","Calliphorinae","Pollenia","Pollenia pediculata","3","3","3","100","0","0","99.03","447.67","0","449.33","33.53","0"
"AGAW1","10W","INS","BOLD:AAA5953","Arthropoda","Insecta","Lepidoptera","Gelechiidae","Gelechiinae","Coleotechnites","Coleotechnites sp., Coleotechnites piceaella, Coleotechnites laricis, Coleotechnites sp. 3, Coleotechnites carbonaria, Coleotechnites granti, Coleotechnites macleodi, Coleotechnites gibsonella, Coleotechnites coniferella, Coleotechnites martini","3","3","3","100","0","0","99.46","416.33","0","417","33.91","0"
"AGAW1","10W","INS","BOLD:AAA6292","Arthropoda","Insecta","Lepidoptera","Tortricidae","Tortricinae","Cnephasia","Cnephasia genitalana, Cnephasia stephensiana","2","2","2","105","0","0","99.11","449.5","0","450","35.33","0"
"AGAW1","10W","INS","BOLD:AAO8538","Arthropoda","Insecta","Diptera","Psychodidae","Psychodinae","Psychoda","","2","2","2","105","0","0","99.5","390","0","391.5","32.07","0"
"AGAW1","10W","INS","BOLD:AAA2632","Arthropoda","Insecta","Lepidoptera","Noctuidae","Noctuinae","Noctua","Noctua atlantica, Noctua pronuba","2","2","2","105","0","0","97.88","449","0","450","31","0"
"AGAW1","10W","INS","BOLD:AAD0484","Arthropoda","Insecta","Diptera","Chironomidae","","","","2","2","2","105","0","0","99.11","449","0","450","34.78","0"
"AGAW1","10W","INS","BOLD:AAF9771","Arthropoda","Insecta","Diptera","Hybotidae","Tachydromiinae","Platypalpus","Platypalpus longiseta, Platypalpus trivialis, Platypalpus sp. 5, Platypalpus sp. 17, Platypalpus pallidiventris","2","2","2","105","0","0","99.44","448","0","450","35.33","0"
"AGAW1","10W","INS","BOLD:ABX2489","Arthropoda","Insecta","Coleoptera","Phalacridae","Phalacrinae","Stilbus","","2","2","2","105","0","0","97.69","434.5","0","435.5","32.59","0"
"AGAW1","10W","INS","BOLD:AAC0903","Arthropoda","Insecta","Diptera","Chironomidae","Chironominae","Chironomus","Chironomus acidophilus","2","2","2","105","0","0","99.23","416","0","416.5","36.74","0"
"AGAW1","10W","INS","BOLD:AAP6790","Arthropoda","Insecta","Diptera","Chloropidae","Oscinellinae","Incertella","Incertella incerta","1","1","1","112","0","0","99.55","440","0","442","31.67","0"
"AGAW1","10W","INS","BOLD:AAA3181","Arthropoda","Insecta","Diptera","Muscidae","Muscinae","Stomoxys","Stomoxys calcitrans","1","1","1","112","0","0","99.11","447","0","450","33.56","0"
"AGAW1","10W","INS","BOLD:AAA9246","Arthropoda","Insecta","Lepidoptera","Tortricidae","Tortricinae","Clepsis","Clepsis clemensiana","1","1","1","112","0","0","99.78","449","0","450","30.67","0"
"AGAW1","10W","INS","BOLD:AAA3075","Arthropoda","Insecta","Lepidoptera","Geometridae","Larentiinae","Eupithecia, Pasiphila","Pasiphila rectangulata","1","1","1","112","0","0","98.56","416","0","419","29.59","0"
"AGAW1","10W","INS","BOLD:AAN5497","Arthropoda","Insecta","Diptera","Anthomyiidae","Pegomyinae","Pegomya","Pegomya solennis, Pegomya bicolor","1","1","1","112","0","0","98.22","450","0","450","29.78","0"
"AGAW1","10W","INS","BOLD:AAA4201","Arthropoda","Insecta","Lepidoptera","Noctuidae","Noctuinae","Apamea","Apamea sp., Apamea sordens, Apamea sp. nr. sordens, Apamea digitula","1","1","1","112","0","0","100","450","0","450","32","0"
"AGAW1","10W","INS","BOLD:AAC9132","Arthropoda","Insecta","Diptera","Culicidae","Culicinae","Culiseta","Culiseta inornata","1","1","1","112","0","0","97.04","406","0","407","32.43","0"
"AGAW1","10W","INS","BOLD:AAG5192","Arthropoda","Insecta","Coleoptera","Curculionidae","Entiminae","Barypeithes","Barypeithes pellucidus","1","1","1","112","0","0","98.22","450","0","450","38.67","0"
"AGAW1","10W","INS","BOLD:AAC0635","Arthropoda","Insecta","Hemiptera","Miridae","Mirinae","Stenotus","Stenotus binotatus","1","1","1","112","0","0","98","449","0","450","34.89","0"
"AGAW1","10W","INS","BOLD:AAU6592","Arthropoda","Insecta","Diptera","Ceratopogonidae","Dasyheleinae","Dasyhelea","Dasyhelea arenivaga","1","1","1","112","0","0","98.88","448","0","450","35.78","0"
"AGAW1","10W","INS","BOLD:AAG6431","Arthropoda","Insecta","Diptera","Ceratopogonidae","Forcipomyiinae","Atrichopogon","","1","1","1","112","0","0","97.33","412","0","413","35.11","0"
"AGAW1","10W","INS","BOLD:AAH3914","Arthropoda","Insecta","Diptera","Sciaridae","","Sciara","Sciara hamatilis, Sciara humeralis","1","1","1","112","0","0","98.12","426","0","427","30.68","0"
"AGAW1","10W","INS","BOLD:AAU6518","Arthropoda","Insecta","Diptera","Cecidomyiidae","Cecidomyiinae","Asteromyia","","1","1","1","112","0","0","97.11","450","0","450","26.89","0"
"AGAW1","10W","INS","BOLD:AAB9140","Arthropoda","Insecta","Diptera","Calliphoridae","Chrysomyinae","Phormia","Phormia regina","1","1","1","112","0","0","98.66","448","0","450","32.89","0"
"AGAW1","10W","INS","BOLD:ACK3142","Arthropoda","Insecta","Diptera","Cecidomyiidae","","","","1","1","1","112","0","0","97.99","447","0","450","28.44","0"
"AGAW1","10W","INS","BOLD:AAG5457","Arthropoda","Insecta","Diptera","Chironomidae","Chironominae","","","1","1","1","112","0","0","98.08","416","0","420","33.1","0"
"AGAW1","10W","INS","BOLD:AAA2815","Arthropoda","Insecta","Lepidoptera","Noctuidae","Noctuinae","Mythimna","Mythimna yukonensis, Mythimna oxygala","1","1","1","112","0","0","99.55","448","0","450","33.78","0"
"AGAW1","10W","INS","BOLD:AAU6970","Arthropoda","Insecta","Coleoptera","Cleridae","Clerinae","Enoclerus","Enoclerus nigripes","1","1","1","112","0","0","98","450","0","450","33.78","0"
"AGAW1","10W","INS","BOLD:AAF9316","Arthropoda","Insecta","Diptera","Psychodidae","Psychodinae","Psychoda","","1","1","1","112","0","0","98.67","450","0","450","31.78","0"
"AGAW1","10W","INS","BOLD:AAN5342","Arthropoda","Insecta","Diptera","Chironomidae","Orthocladiinae","Heleniella","","1","1","1","112","0","0","100","371","0","371","36.12","0"
"AGAW1","10W","INS","BOLD:AAN5341","Arthropoda","Insecta","Diptera","Chironomidae","Orthocladiinae","Camptocladius","Camptocladius stercorarius","1","1","1","112","0","0","99.19","371","0","373","36.19","0"
"AGAW1","10W","INS","BOLD:AAG1741","Arthropoda","Insecta","Diptera","Muscidae","Coenosiinae","Coenosia","","1","1","1","112","0","0","99.33","450","0","450","33.78","0"
"AGAW1","10W","INS","BOLD:ACU3718","Arthropoda","Insecta","Diptera","Agromyzidae","Phytomyzinae","Chromatomyia","","1","1","1","112","0","0","99.11","447","0","450","36","0"
"AGAW1","10W","INS","BOLD:AAB4321","Arthropoda","Insecta","Lepidoptera","Gelechiidae","Gelechiinae","Metzneria","Metzneria lappella","1","1","1","112","0","0","98.59","425","0","450","32.22","0"
"AGAW1","10W","INS","BOLD:AAG2123","Arthropoda","Insecta","Diptera","Tachinidae","Dexiinae","Dinera","Dinera grisescens","1","1","1","112","0","0","99.11","450","0","450","32.22","0"
"AGAW1","10W","INS","BOLD:AAA4555","Arthropoda","Insecta","Orthoptera","Acrididae","Melanoplinae","Melanoplus","Melanoplus stupefactus, Melanoplus angustipennis, Melanoplus ponderosus, Melanoplus bruneri, Melanoplus frigidus, Melanoplus scapularis, Melanoplus cinereus, Melanoplus madeleineae, Melanoplus sanguinipes, Melanoplus nigrescens group, Melanoplus tunicae, Melanoplus walshi, Melanoplus femurrubrum, Melanoplus dawsoni, Melanoplus sp., Melanoplus fasciatus, Melanoplus bowditchi, Melanoplus stonei, Melanoplus bivittatus, Melanoplus huroni, Melanoplus thomasi, Melanoplus borealis, Melanoplus packardii","1","1","1","112","0","0","99.78","450","0","450","35.78","0"
"AGAW1","10W","INS","BOLD:ACF7023","Arthropoda","Insecta","Diptera","Dolichopodidae","","","","1","1","1","112","0","0","98.22","225","0","414","30.19","0"
"AGAW1","10W","INS","BOLD:AAE0845","Arthropoda","Insecta","Hemiptera","Pentatomidae","Pentatominae","Euschistus","Euschistus servus","1","1","1","112","0","0","97.54","448","0","450","37.78","0"
"AGAW1","10W","INS","BOLD:ABX6186","Arthropoda","Insecta","Diptera","Tipulidae","Tipulinae","Nephrotoma","Nephrotoma ferruginea","1","1","1","112","0","0","99.11","447","0","450","35.11","0"
"AGAW1","10W","INS","BOLD:AAM6037","Arthropoda","Insecta","Diptera","Cecidomyiidae","Cecidomyiinae","Contarinia, Dasineura","","1","1","1","112","0","0","97.78","450","0","450","26.67","0"
"AGAW1","10W","INS","BOLD:AAM6230","Arthropoda","Insecta","Diptera","Chironomidae","Chironominae","Chironomus","","1","1","1","112","0","0","99.55","443","0","450","34.44","0"
"AGAW1","10W","INS","BOLD:ADH9310","Arthropoda","Insecta","Diptera","Anthomyiidae","Anthomyiinae","Anthomyia","Anthomyia procellaris, Anthomyia pluvialis","1","1","1","112","0","0","100","387","0","391","31.71","0"
"AGAW1","10W","INS","BOLD:AAA3654","Arthropoda","Insecta","Lepidoptera","Crambidae","Crambinae","Crambus","Crambus bigelovi, Crambus johnsoni, Crambus praefectellus","1","1","1","112","0","0","98.67","450","0","450","31.56","0"
"AGAW1","10W","INS","BOLD:AAA7374","Arthropoda","Insecta","Diptera","Syrphidae","Syrphinae","Sphaerophoria","Sphaerophoria bankowskae, Sphaerophoria infuscata, Sphaerophoria nr. philanthus, Sphaerophoria batava, Sphaerophoria sp., Sphaerophoria rueppellii, Sphaerophoria interrupta-group, Sphaerophoria sp. MA-1, Sphaerophoria cleoae, Sphaerophoria asymmetrica, Sphaerophoria fatarum, Sphaerophoria bengalensis, Sphaerophoria bifurcata, Sphaerophoria taeniata, Sphaerophoria interrupta, Sphaerophoria contigua, Sphaerophoria kaa, Sphaerophoria laurae, Sphaerophoria chongjini, Sphaerophoria boreoalpina, Sphaerophoria sp. BOLDAAA7374, Sphaerophoria macrogaster, Sphaerophoria sulphuripes, Sphaerophoria brevipilosa, Sphaerophoria philanthus, Sphaerophoria pyrrhina, Sphaerophoria abbreviata, Sphaerophoria scripta, Sphaerophoria potentillae, Sphaerophoria longipilosa, Sphaerophoria virgata","1","1","1","112","0","0","98.88","446","0","450","30.89","0"
"AGAW1","10W","INS","BOLD:AAH4205","Arthropoda","Insecta","Diptera","Chloropidae","Chloropinae","Meromyza","Meromyza nigriseta, Meromyza sp.","1","1","1","112","0","0","100","450","0","450","32.89","0"
"AGAW1","10W","INS","BOLD:AAC4197","Arthropoda","Insecta","Diptera","Chironomidae","Orthocladiinae","Paraphaenocladius","Paraphaenocladius impensus","1","1","1","112","0","0","99.11","449","0","450","35.33","0"
"AGAW1","10W","INS","BOLD:AAG4781","Arthropoda","Insecta","Diptera","Agromyzidae","","","","1","1","1","112","0","0","97.78","450","0","450","31.78","0"
"AGAW1","10W","INS","BOLD:ACE7482","Arthropoda","Insecta","Hemiptera","Miridae","","","","1","1","1","112","0","0","98.31","413","0","414","34.3","0"
"AGAW1","10W","INS","BOLD:AAD8086","Arthropoda","Insecta","Lepidoptera","Elachistidae","Elachistinae","Elachista","Elachista griseicornis, Elachista acenteta","1","1","1","112","0","0","99.3","429","0","431","30.86","0"
"AGAW1","10W","INS","BOLD:ACE4701","Arthropoda","Insecta","Lepidoptera","Crambidae","Crambinae","Crambus","Crambus multilinellus","1","1","1","112","0","0","98.66","449","0","450","31.11","0"
"AGAW1","10W","INS","BOLD:AAA4277","Arthropoda","Insecta","Diptera","Syrphidae","Syrphinae","Toxomerus","Toxomerus marginatus, Toxomerus sp. CR-3, Toxomerus CR-3","1","1","1","112","0","0","99.56","450","0","450","30.89","0"
"AGAW1","10W","NAR","no match","no match","no match","no match","no match","no match","no match","no match","no match","no match","no match","no match","no match","no match","no match","no match","no match","no match","no match","no match"
"AGAW1","10W","NIN","BOLD:AAI4346","Arthropoda","Arachnida","Opiliones","Phalangiidae","","Paroligolophus, Phalangium","Paroligolophus agrestis, Phalangium opilio","2052","1800","1800","1","12.28","0","99.31","438.26","0","439.9","36.4","0"
"AGAW1","10W","NIN","BOLD:AAA6381","Arthropoda","Arachnida","Araneae","Tetragnathidae","","Tetragnatha","Tetragnatha laboriosa","1576","1512","1512","2","4.06","0","99.04","428.61","0","430.82","39.69","0"
"AGAW1","10W","NIN","BOLD:AAA7242","Arthropoda","Collembola","Entomobryomorpha","Entomobryidae","Seirinae, Entomobryinae","Entomobrya, Lepidosira","Entomobrya multifasciata, Lepidosira sp.","737","707","707","3","4.07","0","99.09","424.06","0","427.95","39.44","0"
"AGAW1","10W","NIN","BOLD:AAH0003","Arthropoda","Arachnida","Araneae","Linyphiidae","","Agyneta","Agyneta unimaculata","86","86","86","4","0","0","99.21","403.17","0","404.74","32","0"
"AGAW1","10W","NIN","BOLD:AAH0001","Arthropoda","Arachnida","Araneae","Linyphiidae","","Erigone","Erigone autumnalis","25","24","24","5","4","0","98.61","429.84","0","438.84","32.64","0"
"AGAW1","10W","NIN","BOLD:ABW2639","Arthropoda","Arachnida","Trombidiformes","Anystidae","","","","18","18","18","6","0","0","99.6","446.78","0","442.44","31.39","0"
"AGAW1","10W","NIN","BOLD:AAH6630","Arthropoda","Collembola","Entomobryomorpha","Entomobryidae","Entomobryinae","Entomobrya","Entomobrya unostrigata","13","13","13","7","0","0","99.8","442.08","0","443.15","38.79","0"
"AGAW1","10W","NIN","BOLD:ACI9974","Arthropoda","Arachnida","Trombidiformes","Tydeidae","","","","10","10","10","8","0","0","99.69","424.1","0","450","41.09","0"
"AGAW1","10W","NIN","BOLD:AAU6252","Arthropoda","Arachnida","Trombidiformes","Bdellidae","","Cyta","","8","8","8","9","0","0","99.01","419.88","0","421.25","31.45","0"
"AGAW1","10W","NIN","BOLD:ACX8969","Arthropoda","Arachnida","Trombidiformes","Eupodidae","","","","4","4","4","10","0","0","98.6","426.5","0","428.25","28","0"
"AGAW1","10W","NIN","BOLD:ADA1360","Arthropoda","Arachnida","Mesostigmata","Digamasellidae","","Cornodendrolaelaps","Cornodendrolaelaps longiscutatus, Cornodendrolaelaps presepum","3","2","2","11","33.33","0","99.78","450","0","450","30.67","0"
"AGAW1","10W","NIN","BOLD:ACZ1295","Arthropoda","Arachnida","Trombidiformes","Anystidae","","","","2","2","2","12","0","0","99.87","372.5","0","373","31.5","0"
"AGAW1","10W","NIN","BOLD:AAE2695","Arthropoda","Arachnida","Sarcoptiformes","Oribatulidae","","Zygoribatula","","2","2","2","12","0","0","98.44","448","0","449","39.09","0"
"AGAW1","10W","NIN","BOLD:AAB7914","Arthropoda","Collembola","Symphypleona","Bourletiellidae","","Bourletiella","","1","1","1","14","0","0","97.09","447","0","450","39.11","0"
"AGAW1","10W","NIN","BOLD:AAM7961","Arthropoda","Arachnida","Trombidiformes","Anystidae","","","","1","1","1","14","0","0","99.78","450","0","450","31.56","0"
"AGAW1","10W","NIN","BOLD:ACR6863","","","","","","","","1","1","1","14","0","0","97.16","422","0","426","38.73","0"
"AGAW1","10W","NIN","BOLD:ACI9908","Arthropoda","Collembola","Symphypleona","Bourletiellidae","","","","1","1","1","14","0","0","97.78","450","0","450","36","0"
"AGAW1","10W","NIN","BOLD:AAM3234","Arthropoda","Collembola","Entomobryomorpha","Entomobryidae","Seirinae","Lepidosira","","1","1","1","14","0","0","97.77","449","0","450","40.22","0"
"AGAW1","10W","NIN","BOLD:AAF9234","Arthropoda","Arachnida","Trombidiformes","Eupodidae","","","","1","1","1","14","0","0","99.78","450","0","450","30","0"
"AGAW1","11E","BAC","TAX:91391","Proteobacteria","Alphaproteobacteria","Rickettsiales","",NA,"","","20","18","18","1","10","0","99.11","401.55","0","448","39.7","0"
"AGAW1","11E","BAC","TAX:175768","Proteobacteria","Alphaproteobacteria","Rickettsiales","Rickettsiaceae",NA,"Wolbachia","Wolbachia pipientis","17","17","17","2","0","0","98.31","448.29","0","449.47","39.67","0"
"AGAW1","11E","BAC","TAX:91393","Proteobacteria","Alphaproteobacteria","Rickettsiales","Rickettsiaceae",NA,"Wolbachia","","12","12","12","3","0","0","99.13","383.5","0","405.58","38.94","0"
"AGAW1","11E","BAC","TAX:91389","Proteobacteria","","","",NA,"","","1","1","1","4","0","0","99.72","358","0","360","38.61","0"
"AGAW1","11E","CHO","no match","no match","no match","no match","no match","no match","no match","no match","no match","no match","no match","no match","no match","no match","no match","no match","no match","no match","no match","no match"
"AGAW1","11E","INS","BOLD:AAP8739","Arthropoda","Insecta","Diptera","Platystomatidae","Platystomatinae","Rivellia","Rivellia steyskali","50056","30960","30960","1","38.15","0","99.32","440.73","0","442.22","37.97","0"
"AGAW1","11E","INS","BOLD:AAA3453","Arthropoda","Insecta","Diptera","Anthomyiidae","Anthomyiinae","Delia","","8817","7729","7729","2","12.34","0","99.37","428.57","0","431.25","34.65","0"
"AGAW1","11E","INS","BOLD:AAG6745","Arthropoda","Insecta","Diptera","Calliphoridae","Calliphorinae","Pollenia","Pollenia pediculata","7778","6422","6422","3","17.43","0","99.39","433.27","0","435.94","33.55","0"
"AGAW1","11E","INS","BOLD:AAU6681","Arthropoda","Insecta","Diptera","Tipulidae","Tipulinae","Nephrotoma","Nephrotoma cornicina","2649","2074","2074","4","21.71","0","99.61","441.96","0","443.16","36.32","0"
"AGAW1","11E","INS","BOLD:AAG1711","Arthropoda","Insecta","Diptera","Muscidae","Phaoniinae","Helina","Helina depuncta","2604","2200","2200","5","15.51","0","99.36","434.03","0","436.32","31.37","0"
"AGAW1","11E","INS","BOLD:AAB7030","Arthropoda","Insecta","Diptera","Chironomidae","Chironominae","Chironomus","Chironomus sp. BOLDAAP2996, Chironomus quinnitukqut, Chironomus cf. decorus 2g, Chironomus decorus group sp. 2 IP2013, Chironomus cf. decorus, Chironomus decorus gr.","2331","2307","2307","6","1.03","0","98.63","415.9","0","421.85","34.54","0"
"AGAW1","11E","INS","BOLD:AAA6337","Arthropoda","Insecta","Lepidoptera","Noctuidae","Plusiinae","Anagrapha","Anagrapha falcifera","1159","909","909","7","21.57","0","99.45","439.77","0","441.93","32.28","0"
"AGAW1","11E","INS","BOLD:AAG5453","Arthropoda","Insecta","Diptera","Chironomidae","Chironominae","Chironomus","Chironomus bifurcatus","1043","861","861","8","17.45","0","98.65","443.5","0","446.79","34.87","0"
"AGAW1","11E","INS","BOLD:AAG2511","Arthropoda","Insecta","Diptera","Anthomyiidae","Anthomyiinae","Delia","Delia platura","959","864","864","9","9.91","0","99.02","442.41","0","445.4","33.11","0"
"AGAW1","11E","INS","BOLD:AAA7374","Arthropoda","Insecta","Diptera","Syrphidae","Syrphinae","Sphaerophoria","Sphaerophoria bankowskae, Sphaerophoria infuscata, Sphaerophoria nr. philanthus, Sphaerophoria batava, Sphaerophoria sp., Sphaerophoria rueppellii, Sphaerophoria interrupta-group, Sphaerophoria sp. MA-1, Sphaerophoria cleoae, Sphaerophoria asymmetrica, Sphaerophoria fatarum, Sphaerophoria bengalensis, Sphaerophoria bifurcata, Sphaerophoria taeniata, Sphaerophoria interrupta, Sphaerophoria contigua, Sphaerophoria kaa, Sphaerophoria laurae, Sphaerophoria chongjini, Sphaerophoria boreoalpina, Sphaerophoria sp. BOLDAAA7374, Sphaerophoria macrogaster, Sphaerophoria sulphuripes, Sphaerophoria brevipilosa, Sphaerophoria philanthus, Sphaerophoria pyrrhina, Sphaerophoria abbreviata, Sphaerophoria scripta, Sphaerophoria potentillae, Sphaerophoria longipilosa, Sphaerophoria virgata","920","710","710","10","22.83","0","98.68","440.21","0","441.8","31.24","0"
"AGAW1","11E","INS","BOLD:AAC0903","Arthropoda","Insecta","Diptera","Chironomidae","Chironominae","Chironomus","Chironomus acidophilus","906","744","744","11","17.88","0","99.5","436.25","0","438.75","36.32","0"
"AGAW1","11E","INS","BOLD:AAN5162","Arthropoda","Insecta","Diptera","Ceratopogonidae","Dasyheleinae","Dasyhelea","","867","791","791","12","8.77","0","99.31","429.38","0","431.3","33.57","0"
"AGAW1","11E","INS","BOLD:ABW5431","Arthropoda","Insecta","Diptera","Agromyzidae","","","","780","633","633","13","18.85","0","99.24","431.14","0","432.25","34.11","0"
"AGAW1","11E","INS","BOLD:AAH2373","Arthropoda","Insecta","Coleoptera","Elateridae","Dendrometrinae","Hemicrepidius","Hemicrepidius memnonius","775","722","722","14","6.84","0","99.02","421.97","0","423.47","37.2","0"
"AGAW1","11E","INS","BOLD:ABX6186","Arthropoda","Insecta","Diptera","Tipulidae","Tipulinae","Nephrotoma","Nephrotoma ferruginea","669","614","614","15","8.22","0","98.96","427.66","0","429.68","34.7","0"
"AGAW1","11E","INS","BOLD:ACK9434","Arthropoda","Insecta","Hemiptera","Cicadellidae","Macropsinae","Macropsis","Macropsis ocellata","629","495","495","16","21.3","0","99.22","443.91","0","444.93","33.96","0"
"AGAW1","11E","INS","BOLD:AAG5055","Arthropoda","Insecta","Diptera","Xylomyidae","","Solva","","597","457","457","17","23.45","0","99.29","446.97","0","448.12","35.26","0"
"AGAW1","11E","INS","BOLD:AAN6154","Arthropoda","Insecta","Coleoptera","Latridiidae","Corticariinae","Melanophthalma","Melanophthalma inermis","582","504","504","18","13.4","0","98.93","435.19","0","436.78","31.39","0"
"AGAW1","11E","INS","BOLD:AAH3661","Arthropoda","Insecta","Diptera","Cecidomyiidae","","","","575","522","522","19","9.22","0","99.28","431.34","0","433.03","25.41","0"
"AGAW1","11E","INS","BOLD:AAU6592","Arthropoda","Insecta","Diptera","Ceratopogonidae","Dasyheleinae","Dasyhelea","Dasyhelea arenivaga","574","565","565","20","1.57","0","99.15","396.05","0","398.23","34.55","0"
"AGAW1","11E","INS","BOLD:AAK5928","Arthropoda","Insecta","Coleoptera","Chrysomelidae","Criocerinae","Oulema","Oulema melanopus","567","472","472","21","16.75","0","99.49","438.75","0","440.26","36.55","0"
"AGAW1","11E","INS","BOLD:AAG9897","Arthropoda","Insecta","Dermaptera","Forficulidae","Forficulinae","Forficula","Forficula auricularia-A","430","424","424","22","1.4","0","99.16","394.67","0","404.31","45.67","0"
"AGAW1","11E","INS","BOLD:ABW1379","Arthropoda","Insecta","Diptera","Chloropidae","Oscinellinae","Malloewia","Malloewia abdominalis","429","348","348","23","18.88","0","98.44","444.09","0","446.75","31.45","0"
"AGAW1","11E","INS","BOLD:AAG2123","Arthropoda","Insecta","Diptera","Tachinidae","Dexiinae","Dinera","Dinera grisescens","404","322","322","24","20.3","0","99.37","432.37","0","434.1","31.86","0"
"AGAW1","11E","INS","BOLD:AAH2306","Arthropoda","Insecta","Diptera","Asilidae","Asilinae","Efferia","Efferia aestuans","395","353","353","25","10.63","0","99.27","434.39","0","435.75","37.96","0"
"AGAW1","11E","INS","BOLD:ABW7897","Arthropoda","Insecta","Diptera","Cecidomyiidae","","","","365","264","264","26","27.67","0","99.65","442.87","0","445.06","29.69","0"
"AGAW1","11E","INS","BOLD:AAP5169","Arthropoda","Insecta","Diptera","Chloropidae","Oscinellinae","Elachiptera","Elachiptera nigriceps","363","294","294","27","19.01","0","99.34","430.32","0","431.4","31.66","0"
"AGAW1","11E","INS","BOLD:AAE5173","Arthropoda","Insecta","Diptera","Psychodidae","","PschodididGC","","363","360","360","27","0.83","0","98.96","405.04","0","407.51","34.64","0"
"AGAW1","11E","INS","BOLD:AAL4232","Arthropoda","Insecta","Diptera","Agromyzidae","Phytomyzinae","Phytoliriomyza","Phytoliriomyza arctica","346","268","268","29","22.54","0","99.58","440.9","0","442.32","32.05","0"
"AGAW1","11E","INS","BOLD:AAN5199","Arthropoda","Insecta","Diptera","Cecidomyiidae","","","CecidInt20 sp. BOLD:AAN5199","338","269","269","30","20.41","0","99.59","440.6","0","445.62","26.6","0"
"AGAW1","11E","INS","BOLD:AAG6471","Arthropoda","Insecta","Diptera","Ceratopogonidae","Dasyheleinae","Dasyhelea","","307","291","291","31","5.21","0","99","419.58","0","421.95","34.06","0"
"AGAW1","11E","INS","BOLD:ABY5318","Arthropoda","Insecta","Diptera","Chloropidae","Oscinellinae","Incertella, Rhopalopterum","Rhopalopterum umbrosum, Rhopalopterum luteiceps, Incertella bispina, Rhopalopterum painteri","302","234","234","32","22.52","0","99.51","443.74","0","444.92","31.97","0"
"AGAW1","11E","INS","BOLD:ACV4278","Arthropoda","Insecta","Diptera","Limoniidae","Chioneinae","Erioptera","Erioptera chlorophylla","277","263","263","33","5.05","0","99.33","428.4","0","431.04","32.93","0"
"AGAW1","11E","INS","BOLD:AAH2295","Arthropoda","Insecta","Diptera","Asilidae","Leptogastrinae","Leptogaster","Leptogaster flavipes","237","200","200","34","15.61","0","98.97","407.6","0","440.66","34.89","0"
"AGAW1","11E","INS","BOLD:AAA5803","Arthropoda","Insecta","Hemiptera","Miridae","Mirinae","Lygus","Lygus lineolaris","193","151","151","35","21.76","0","99.22","446.12","0","447.82","34.12","0"
"AGAW1","11E","INS","BOLD:ABA0822","Arthropoda","Insecta","Diptera","Cecidomyiidae","","","","188","166","166","36","11.7","0","98.88","427.35","0","428.54","26.07","0"
"AGAW1","11E","INS","BOLD:AAP6790","Arthropoda","Insecta","Diptera","Chloropidae","Oscinellinae","Incertella","Incertella incerta","181","162","162","37","10.5","0","99.05","435.25","0","437.24","31.15","0"
"AGAW1","11E","INS","BOLD:AAN5242","Arthropoda","Insecta","Diptera","Cecidomyiidae","","","","177","177","177","38","0","0","98.76","402.68","0","405.35","28.11","0"
"AGAW1","11E","INS","BOLD:AAG2868","Arthropoda","Insecta","Hemiptera","Cicadellidae","Typhlocybinae","Empoasca","Empoasca sp., Empoasca fabae","172","130","130","39","24.42","0","99.62","444.6","0","447.13","32.32","0"
"AGAW1","11E","INS","BOLD:AAB0373","Arthropoda","Insecta","Neuroptera","Chrysopidae","Chrysopinae","Chrysoperla","Chrysoperla sp. isolate 82, Chrysoperla zastrowi, Chrysoperla plorabunda, Chrysoperla sp. isolate 71, Chrysoperla agilis, Italochrysa italica, Chrysoperla sp. isolate 88, Chrysoperla lucasina, Chrysoperla carnea, Chrysoperla mediterranea, Chrysoperla pallida, Chrysoperla nipponensis, Chrysoperla sillemi","169","135","135","40","20.12","0","99.41","433.04","0","434.05","31.08","0"
"AGAW1","11E","INS","BOLD:AAP9022","Arthropoda","Insecta","Diptera","Cecidomyiidae","","","","167","152","152","41","8.98","0","99.58","439.38","0","440.81","27.52","0"
"AGAW1","11E","INS","BOLD:AAZ9943","Arthropoda","Insecta","Hymenoptera","Chrysididae","Chrysidinae","Hedychridium","Hedychridium dimidiatum","167","139","139","41","16.77","0","99.51","431.58","0","433.1","30.62","0"
"AGAW1","11E","INS","BOLD:AAZ9744","Arthropoda","Insecta","Hymenoptera","Braconidae","Brachistinae","Blacus","Blacus stelfoxi","163","116","116","43","28.83","0","98.41","445.17","0","446.39","32.33","0"
"AGAW1","11E","INS","BOLD:AAB8271","Arthropoda","Insecta","Lepidoptera","Autostichidae","Oegoconiinae","Oegoconia","Oegoconia quadripuncta, Oegoconia deauratella","160","160","160","44","0","0","99.01","398.61","0","400.54","34.28","0"
"AGAW1","11E","INS","BOLD:ACK3431","Arthropoda","Insecta","Diptera","Cecidomyiidae","","","","156","146","146","45","6.41","0","99.13","426.29","0","428.11","32.52","0"
"AGAW1","11E","INS","BOLD:ACE7444","Arthropoda","Insecta","Hemiptera","Miridae","Mirinae, Phylinae","Adelphocoris, Megalocoleus","Adelphocoris lineolatus, Megalocoleus molliculus","151","113","113","46","25.17","0","99.63","442.75","0","443.36","34.91","0"
"AGAW1","11E","INS","BOLD:AAN5183","Arthropoda","Insecta","Diptera","Cecidomyiidae","","","","148","131","131","47","11.49","0","98.43","442.88","0","444.19","27.8","0"
"AGAW1","11E","INS","BOLD:ABA0863","Arthropoda","Insecta","Diptera","Cecidomyiidae","","","","137","124","124","48","9.49","0","99.32","438.82","0","440.04","29.87","0"
"AGAW1","11E","INS","BOLD:AAA4277","Arthropoda","Insecta","Diptera","Syrphidae","Syrphinae","Toxomerus","Toxomerus marginatus, Toxomerus sp. CR-3, Toxomerus CR-3","133","108","108","49","18.8","0","99.44","441.01","0","441.98","30.62","0"
"AGAW1","11E","INS","BOLD:AAG4333","Arthropoda","Insecta","Coleoptera","Staphylinidae","Staphylininae","Xantholinus","Xantholinus linearis","133","100","100","49","24.81","0","99.48","444.3","0","446.73","33.59","0"
"AGAW1","11E","INS","BOLD:AAC4403","Arthropoda","Insecta","Diptera","Chironomidae","Chironominae","Micropsectra","Micropsectra polita","122","121","121","51","0.82","0","98.91","412.34","0","414.56","34.58","0"
"AGAW1","11E","INS","BOLD:AAA1259","Arthropoda","Insecta","Diptera","Cecidomyiidae","","","","121","114","114","52","5.79","0","99.17","422.91","0","424.69","30.17","0"
"AGAW1","11E","INS","BOLD:AAA3782","Arthropoda","Insecta","Hymenoptera","Halictidae","Halictinae","Lasioglossum","Lasioglossum weemsi, Lasioglossum leviense, Lasioglossum hitchensi","119","110","110","53","7.56","0","98.94","430.79","0","432.06","28.28","0"
"AGAW1","11E","INS","BOLD:AAN5660","Arthropoda","Insecta","Diptera","Chloropidae","Oscinellinae","Incertella","Incertella minor","116","87","87","54","25","0","99.41","447.82","0","448.7","32.89","0"
"AGAW1","11E","INS","BOLD:AAP5087","Arthropoda","Insecta","Diptera","Ceratopogonidae","Dasyheleinae","Dasyhelea","","115","98","98","55","14.78","0","99.12","440","0","441.51","34.79","0"
"AGAW1","11E","INS","BOLD:AAO7783","Arthropoda","Insecta","Diptera","Dolichopodidae","Diaphorinae","","","110","107","107","56","2.73","0","98.05","443.25","0","445.53","33.09","0"
"AGAW1","11E","INS","BOLD:ADF6661","Arthropoda","Insecta","Diptera","Cecidomyiidae","","","","109","93","93","57","14.68","0","99.26","414.67","0","441.19","29.74","0"
"AGAW1","11E","INS","BOLD:AAG9626","Arthropoda","Insecta","Diptera","Dolichopodidae","","","","101","98","98","58","2.97","0","97.8","424.76","0","426.69","36.27","0"
"AGAW1","11E","INS","BOLD:AAH0256","Arthropoda","Insecta","Coleoptera","Latridiidae","Corticariinae","Corticaria, Corticarina","","94","83","83","59","11.7","0","98.8","423.27","0","430.4","31.22","0"
"AGAW1","11E","INS","BOLD:AAA7661","Arthropoda","Insecta","Diptera","Culicidae","Culicinae","Culiseta, Culex","Culiseta melanura, Culex articularis, Culex restuans, Culex acharistus","91","88","88","60","3.3","0","99.15","427.47","0","429.51","32.92","0"
"AGAW1","11E","INS","BOLD:ABU5525","Arthropoda","Insecta","Diptera","Chironomidae","Orthocladiinae","Limnophyes","Limnophyes sp. 14ES","90","87","87","61","3.33","0","99.12","398.63","0","403.62","37.08","0"
"AGAW1","11E","INS","BOLD:AAG8493","Arthropoda","Insecta","Diptera","Drosophilidae","Drosophilinae","Scaptomyza","Scaptomyza pallida","89","89","89","62","0","0","98.87","393.65","0","395.73","33.02","0"
"AGAW1","11E","INS","BOLD:AAN5161","Arthropoda","Insecta","Diptera","Ceratopogonidae","Dasyheleinae","Dasyhelea","","84","82","82","63","2.38","0","98.99","405.63","0","407.67","37.41","0"
"AGAW1","11E","INS","BOLD:ADG8758","Arthropoda","Insecta","Hymenoptera","Braconidae","Brachistinae","Blacus","Blacus stelfoxi","81","70","70","64","13.58","0","99.23","433.25","0","435.32","32","0"
"AGAW1","11E","INS","BOLD:AAH0134","Arthropoda","Insecta","Coleoptera","Phalacridae","Phalacrinae","Stilbus","Stilbus apicialis","81","74","74","64","8.64","0","99.42","423.94","0","425.31","31.89","0"
"AGAW1","11E","INS","BOLD:AAA3882","Arthropoda","Insecta","Lepidoptera","Pterophoridae","Pterophorinae","Emmelina","Emmelina monodactyla","80","80","80","66","0","0","98.64","397.35","0","403.83","31.67","0"
"AGAW1","11E","INS","BOLD:AAY3940","Arthropoda","Insecta","Diptera","Agromyzidae","Phytomyzinae","Phytomyza","Phytomyza spinaciae","69","67","67","67","2.9","0","99.31","436.16","0","438.8","32.91","0"
"AGAW1","11E","INS","BOLD:AAD1485","Arthropoda","Insecta","Diptera","Chironomidae","Chironominae","Paratanytarsus","Paratanytarsus grimmii, Paratanytarsus sp.","67","62","62","68","7.46","0","99.49","441.46","0","442.97","33.76","0"
"AGAW1","11E","INS","BOLD:ACE0554","Arthropoda","Insecta","Diptera","Phoridae","Metopininae","Megaselia","","65","62","62","69","4.62","0","99.64","442.63","0","444.63","32.08","0"
"AGAW1","11E","INS","BOLD:AAN6447","Arthropoda","Insecta","Diptera","Sciaridae","","Corynoptera","Corynoptera perpusilla","63","51","51","70","19.05","0","99.15","445.6","0","446.86","36.3","0"
"AGAW1","11E","INS","BOLD:AAD2041","Arthropoda","Insecta","Diptera","Dolichopodidae","Sciapodinae","Condylostylus","Condylostylus inermis, Condylostylus nigrofemoratus, Condylostylus connectans, Condylostylus caudatus, Condylostylus flavipes","62","54","54","71","12.9","0","99.58","433.02","0","434.19","35.88","0"
"AGAW1","11E","INS","BOLD:AAC4197","Arthropoda","Insecta","Diptera","Chironomidae","Orthocladiinae","Paraphaenocladius","Paraphaenocladius impensus","62","51","51","71","17.74","0","99.36","431.52","0","435.56","34.92","0"
"AGAW1","11E","INS","BOLD:AAH0135","Arthropoda","Insecta","Coleoptera","Phalacridae","Phalacrinae","Acylomus","Acylomus pugetanus, Acylomus ergoti","59","58","58","73","1.69","0","99.23","432","0","434.19","31.55","0"
"AGAW1","11E","INS","BOLD:AAF9771","Arthropoda","Insecta","Diptera","Hybotidae","Tachydromiinae","Platypalpus","Platypalpus longiseta, Platypalpus trivialis, Platypalpus sp. 5, Platypalpus sp. 17, Platypalpus pallidiventris","58","45","45","74","22.41","0","99.69","443.84","0","444.71","35.22","0"
"AGAW1","11E","INS","BOLD:AAG6433","Arthropoda","Insecta","Diptera","Ceratopogonidae","Forcipomyiinae","Forcipomyia","Forcipomyia sp. BOLD:AAG6433, Forcipomyia sp. 17","57","50","50","75","12.28","0","98.51","436.26","0","437.7","32.71","0"
"AGAW1","11E","INS","BOLD:AAE4793","Arthropoda","Insecta","Diptera","Pipunculidae","Pipunculinae","Pipunculus","Pipunculus hertzogi, Pipunculus sp. MB1","56","42","42","76","25","0","99.34","443.34","0","444.46","37.92","0"
"AGAW1","11E","INS","BOLD:ACU7906","Arthropoda","Insecta","Diptera","Chironomidae","Orthocladiinae","","","50","50","50","77","0","0","98.95","411.68","0","413.98","36","0"
"AGAW1","11E","INS","BOLD:AAF1729","Arthropoda","Insecta","Psocodea","Lachesillidae","Lachesillinae","Lachesilla","Lachesilla pedicularia","47","37","37","78","21.28","0","99.54","446.7","0","447.91","37.14","0"
"AGAW1","11E","INS","BOLD:ACC8079","Arthropoda","Insecta","Diptera","Cecidomyiidae","","","","42","39","39","79","7.14","0","99.32","446.4","0","447.86","29.17","0"
"AGAW1","11E","INS","BOLD:AAM9013","Arthropoda","Insecta","Diptera","Keroplatidae","Macrocerinae","Macrocera","Macrocera pusilla","41","41","41","80","0","0","99.45","434.07","0","435.34","30.31","0"
"AGAW1","11E","INS","BOLD:AAF0738","Arthropoda","Insecta","Trichoptera","Hydroptilidae","Orthotrichiinae","Orthotrichia","Orthotrichia aegerfasciella","40","40","40","81","0","0","98.82","425.63","0","427.73","31.12","0"
"AGAW1","11E","INS","BOLD:AAN5238","Arthropoda","Insecta","Diptera","Cecidomyiidae","","","","39","38","38","82","2.56","0","99.24","431.51","0","432.72","25.82","0"
"AGAW1","11E","INS","BOLD:AAN5260","Arthropoda","Insecta","Diptera","Cecidomyiidae","","","","39","39","39","82","0","0","99.24","408.67","0","410.1","25.99","0"
"AGAW1","11E","INS","BOLD:ABV1279","Arthropoda","Insecta","Diptera","Cecidomyiidae","","","","35","33","33","84","5.71","0","99.43","440.6","0","442.54","28.85","0"
"AGAW1","11E","INS","BOLD:AAN5255","Arthropoda","Insecta","Diptera","Cecidomyiidae","Cecidomyiinae","Dasineura","","35","33","33","84","5.71","0","98.62","432.11","0","445.2","26.65","0"
"AGAW1","11E","INS","BOLD:ACR3866","Arthropoda","Insecta","Psocodea","Caeciliusidae","Caeciliusinae","Valenzuela","","33","29","29","86","12.12","0","99.57","439.3","0","440.3","35.55","0"
"AGAW1","11E","INS","BOLD:AAC4523","Arthropoda","Insecta","Diptera","Chironomidae","Chironominae","Tanytarsus","Tanytarsus guerlus","31","28","28","87","9.68","0","99.11","431.26","0","436.81","32.46","0"
"AGAW1","11E","INS","BOLD:ABA7871","Arthropoda","Insecta","Diptera","Cecidomyiidae","","","","29","23","23","88","20.69","0","99.52","437.21","0","438.34","26.49","0"
"AGAW1","11E","INS","BOLD:AAM6062","Arthropoda","Insecta","Diptera","Cecidomyiidae","","","","26","24","24","89","7.69","0","99.45","435.08","0","436.12","25.99","0"
"AGAW1","11E","INS","BOLD:AAM9255","Arthropoda","Insecta","Diptera","Sciaridae","","Pseudolycoriella","","26","26","26","89","0","0","99.16","414.42","0","416.81","30.75","0"
"AGAW1","11E","INS","BOLD:AAN5220","Arthropoda","Insecta","Diptera","Cecidomyiidae","","","","23","23","23","91","0","0","98.13","391.35","0","410.83","25.58","0"
"AGAW1","11E","INS","BOLD:AAG4477","Arthropoda","Insecta","Coleoptera","Chrysomelidae","Chrysomelinae","Plagiodera","Plagiodera versicolora","22","22","22","92","0","0","99.54","429.41","0","430.32","34.67","0"
"AGAW1","11E","INS","BOLD:ADF3937","","","","","","","","22","22","22","92","0","0","98.64","401.73","0","406.05","30.76","0"
"AGAW1","11E","INS","BOLD:ABV1337","Arthropoda","Insecta","Diptera","Cecidomyiidae","","","","21","19","19","94","9.52","0","99.19","432.86","0","438.81","28.34","0"
"AGAW1","11E","INS","BOLD:AAV5689","Arthropoda","Insecta","Diptera","Cecidomyiidae","","","","20","20","20","95","0","0","98.74","437.1","0","438.8","26.13","0"
"AGAW1","11E","INS","BOLD:ACB0061","Arthropoda","Insecta","Diptera","Cecidomyiidae","","","","19","19","19","96","0","0","98.67","437.74","0","439.37","27.56","0"
"AGAW1","11E","INS","BOLD:AAM6037","Arthropoda","Insecta","Diptera","Cecidomyiidae","Cecidomyiinae","Contarinia, Dasineura","","19","18","18","96","5.26","0","99.17","440.21","0","442.16","26.38","0"
"AGAW1","11E","INS","BOLD:AAG5507","Arthropoda","Insecta","Diptera","Chironomidae","Chironominae","Chironomus","Chironomus atrella","18","18","18","98","0","0","99.02","415.11","0","430.78","35.76","0"
"AGAW1","11E","INS","BOLD:ABV9018","Arthropoda","Insecta","Diptera","Mycetophilidae","Sciophilinae","Sciophila","","18","18","18","98","0","0","98.85","437.94","0","440.61","31.82","0"
"AGAW1","11E","INS","BOLD:AAC2157","Arthropoda","Insecta","Hymenoptera","Megachilidae","Megachilinae","Heriades","Heriades carinatus","18","18","18","98","0","0","98.9","386.83","0","389.11","23.5","0"
"AGAW1","11E","INS","BOLD:AAJ7123","Arthropoda","Insecta","Diptera","Culicidae","Culicinae","Culiseta","Culiseta minnesotae","18","18","18","98","0","0","99.11","433.67","0","434.83","34.68","0"
"AGAW1","11E","INS","BOLD:ACV5892","Arthropoda","Insecta","Diptera","Cecidomyiidae","","","","17","17","17","102","0","0","98.54","438.65","0","440.35","24.63","0"
"AGAW1","11E","INS","BOLD:AAN5332","Arthropoda","Insecta","Diptera","Chironomidae","Chironominae","Polypedilum","","17","17","17","102","0","0","97.63","408.12","0","410.76","33.37","0"
"AGAW1","11E","INS","BOLD:ACK5742","Arthropoda","Insecta","Diptera","Cecidomyiidae","","","","15","15","15","104","0","0","99.2","429.2","0","430.93","28.72","0"
"AGAW1","11E","INS","BOLD:AAH3760","Arthropoda","Insecta","Diptera","Cecidomyiidae","","","","15","12","12","104","20","0","98.86","439.67","0","441.33","25.88","0"
"AGAW1","11E","INS","BOLD:AAH3662","Arthropoda","Insecta","Diptera","Cecidomyiidae","","","CecidInt3 sp. BOLD:AAH3662","14","14","14","106","0","0","98.96","441.43","0","442.71","26.09","0"
"AGAW1","11E","INS","BOLD:AAG8091","Arthropoda","Insecta","Hymenoptera","Ichneumonidae","Campopleginae","Campoletis","Campoletis sp. AB.2","14","14","14","106","0","0","99.22","429.14","0","431.71","24.49","0"
"AGAW1","11E","INS","BOLD:AAG3236","Arthropoda","Insecta","Diptera","Phoridae","Phorinae","Conicera, Diplonevra","Diplonevra nitidula","14","14","14","106","0","0","99.75","448.93","0","450","32.03","0"
"AGAW1","11E","INS","BOLD:ABW8055","Arthropoda","Insecta","Diptera","Cecidomyiidae","","","","14","13","13","106","7.14","0","99.08","437.36","0","439.57","25.11","0"
"AGAW1","11E","INS","BOLD:AAN5230","Arthropoda","Insecta","Diptera","Cecidomyiidae","","","","12","12","12","110","0","0","99.4","431.67","0","433.25","26.28","0"
"AGAW1","11E","INS","BOLD:AAN5201","Arthropoda","Insecta","Diptera","Cecidomyiidae","","","","12","11","11","110","8.33","0","99.46","449.42","0","450","27.94","0"
"AGAW1","11E","INS","BOLD:AAV1261","Arthropoda","Insecta","Diptera","Sciaridae","","Bradysia","Bradysia splendida","12","12","12","110","0","0","99.34","425.42","0","427.92","33.97","0"
"AGAW1","11E","INS","BOLD:ACX6289","Arthropoda","Insecta","Diptera","Phoridae","Metopininae","Megaselia","","12","12","12","110","0","0","98.98","414.5","0","416.42","38.24","0"
"AGAW1","11E","INS","BOLD:AAO8538","Arthropoda","Insecta","Diptera","Psychodidae","Psychodinae","Psychoda","","11","11","11","114","0","0","98.57","421","0","423.55","32.28","0"
"AGAW1","11E","INS","BOLD:ABY3788","Arthropoda","Insecta","Thysanoptera","Phlaeothripidae","Idolothripinae","Bolothrips","","11","11","11","114","0","0","98.25","417.27","0","422.55","30.19","0"
"AGAW1","11E","INS","BOLD:AAA4222","Arthropoda","Insecta","Lepidoptera","Erebidae","Hypeninae","Hypena","Hypena scabra","11","11","11","114","0","0","98.94","395.36","0","397.91","32.24","0"
"AGAW1","11E","INS","BOLD:ABX4079","Arthropoda","Insecta","Diptera","Cecidomyiidae","","","","10","10","10","117","0","0","98.41","441","0","442.6","27.96","0"
"AGAW1","11E","INS","BOLD:ACA1917","Arthropoda","Insecta","Diptera","Cecidomyiidae","","","","10","10","10","117","0","0","98.59","413.6","0","414.9","24.85","0"
"AGAW1","11E","INS","BOLD:ABV1487","Arthropoda","Insecta","Diptera","Cecidomyiidae","","","","10","10","10","117","0","0","99.58","429","0","431.6","32.56","0"
"AGAW1","11E","INS","BOLD:ACK3142","Arthropoda","Insecta","Diptera","Cecidomyiidae","","","","10","10","10","117","0","0","99.46","447.5","0","449.1","28.9","0"
"AGAW1","11E","INS","BOLD:AAY6469","Arthropoda","Insecta","Diptera","Cecidomyiidae","","","","10","10","10","117","0","0","99.28","432","0","432.8","25.99","0"
"AGAW1","11E","INS","BOLD:ACX6055","Arthropoda","Insecta","Diptera","Phoridae","Metopininae","Megaselia","Megaselia sp. AK-2010-2, Megaselia sp. BOLD:ACX6055","9","9","9","122","0","0","99.13","409.22","0","411.11","33.08","0"
"AGAW1","11E","INS","BOLD:ADH7839","Arthropoda","Insecta","Diptera","","","","","9","9","9","122","0","0","98.83","412.89","0","416.33","25.56","0"
"AGAW1","11E","INS","BOLD:ABA0811","Arthropoda","Insecta","Diptera","Cecidomyiidae","","","","8","8","8","124","0","0","99.52","441","0","442.38","26.52","0"
"AGAW1","11E","INS","BOLD:AAF7193","Arthropoda","Insecta","Diptera","Cecidomyiidae","Cecidomyiinae","Aphidoletes, Dasineura","Aphidoletes aphidimyza, Dasineura leguminicola, Dasineura sp. BOLD:AAF7193","8","8","8","124","0","0","99.33","429.75","0","431.38","26.23","0"
"AGAW1","11E","INS","BOLD:AAV5609","Arthropoda","Insecta","Diptera","Cecidomyiidae","","","","7","7","7","126","0","0","99.61","442.57","0","443.29","27.41","0"
"AGAW1","11E","INS","BOLD:AAP4636","Arthropoda","Insecta","Psocodea","Caeciliusidae","","","","7","7","7","126","0","0","99.8","363.86","0","413.57","36.47","0"
"AGAW1","11E","INS","BOLD:AAN5205","Arthropoda","Insecta","Diptera","Cecidomyiidae","","","CecidInt42 sp. BOLD:AAN5205","7","7","7","126","0","0","98.74","422","0","423.86","30.92","0"
"AGAW1","11E","INS","BOLD:ACE5372","Arthropoda","Insecta","Hymenoptera","Halictidae","Halictinae","Lasioglossum","Lasioglossum simplex","7","5","5","126","28.57","0","97.47","447","0","448.43","28.36","0"
"AGAW1","11E","INS","BOLD:AAP6465","Arthropoda","Insecta","Diptera","Sciaridae","","Corynoptera","Corynoptera cursor","7","7","7","126","0","0","98.59","445.43","0","446.29","32.36","0"
"AGAW1","11E","INS","BOLD:AAL7819","Arthropoda","Insecta","Diptera","Psychodidae","Psychodinae","Psychoda","Psychoda sp., Psychoda sp. BOLD:AAL7819","6","6","6","131","0","0","99.17","422.67","0","424.5","32.8","0"
"AGAW1","11E","INS","BOLD:AAA4555","Arthropoda","Insecta","Orthoptera","Acrididae","Melanoplinae","Melanoplus","Melanoplus stupefactus, Melanoplus angustipennis, Melanoplus ponderosus, Melanoplus bruneri, Melanoplus frigidus, Melanoplus scapularis, Melanoplus cinereus, Melanoplus madeleineae, Melanoplus sanguinipes, Melanoplus nigrescens group, Melanoplus tunicae, Melanoplus walshi, Melanoplus femurrubrum, Melanoplus dawsoni, Melanoplus sp., Melanoplus fasciatus, Melanoplus bowditchi, Melanoplus stonei, Melanoplus bivittatus, Melanoplus huroni, Melanoplus thomasi, Melanoplus borealis, Melanoplus packardii","6","6","6","131","0","0","99.15","449.83","0","450","36.41","0"
"AGAW1","11E","INS","BOLD:AAN5221","Arthropoda","Insecta","Diptera","Cecidomyiidae","","","","6","6","6","131","0","0","98.77","421.83","0","422.83","25.45","0"
"AGAW1","11E","INS","BOLD:AAH3655","Arthropoda","Insecta","Diptera","Cecidomyiidae","","","","6","6","6","131","0","0","99.1","413.5","0","415","26.95","0"
"AGAW1","11E","INS","BOLD:AAU0577","Arthropoda","Insecta","Thysanoptera","Aeolothripidae","Aeolothripinae","Aeolothrips","Aeolothrips fasciatus","5","5","5","135","0","0","98.02","406","0","428.2","36.55","0"
"AGAW1","11E","INS","BOLD:ACL1796","Arthropoda","Insecta","Lepidoptera","Ypsolophidae","Ochsenheimeriinae","Ochsenheimeria","Ochsenheimeria vacculella","5","5","5","135","0","0","98.66","397","0","410","32.63","0"
"AGAW1","11E","INS","BOLD:ABZ0908","Arthropoda","Insecta","Diptera","Tipulidae","Tipulinae","Nephrotoma","","5","5","5","135","0","0","97.25","370.6","0","372.6","35.05","0"
"AGAW1","11E","INS","BOLD:ACO3101","Arthropoda","Insecta","Diptera","Psychodidae","Psychodinae","","","5","5","5","135","0","0","99.07","388","0","412","34.57","0"
"AGAW1","11E","INS","BOLD:ACV0785","Arthropoda","Insecta","Hymenoptera","Ichneumonidae","Mesochorinae","","","5","5","5","135","0","0","99.91","449.4","0","450","27.6","0"
"AGAW1","11E","INS","BOLD:ACV1014","Arthropoda","Insecta","Diptera","Cecidomyiidae","","","","4","4","4","140","0","0","98.06","400.5","0","405.75","25.39","0"
"AGAW1","11E","INS","BOLD:AAN5383","Arthropoda","Insecta","Diptera","Chironomidae","Chironominae","Dicrotendipes","","4","4","4","140","0","0","97.83","379.25","0","380","36.66","0"
"AGAW1","11E","INS","BOLD:ACJ6312","Arthropoda","Insecta","Diptera","Psychodidae","","","","4","4","4","140","0","0","98.16","447.5","0","450","35.72","0"
"AGAW1","11E","INS","BOLD:ACU6062","Arthropoda","Insecta","Diptera","Cecidomyiidae","","","","4","4","4","140","0","0","98.64","420.25","0","423.75","26.37","0"
"AGAW1","11E","INS","BOLD:AAF9306","Arthropoda","Insecta","Diptera","Psychodidae","Psychodinae","Psychoda","Psychoda alternata","4","4","4","140","0","0","98.17","430.5","0","431","31.64","0"
"AGAW1","11E","INS","BOLD:ADH9867","Arthropoda","Insecta","Diptera","","","","","4","4","4","140","0","0","98.95","384.5","0","388","37.71","0"
"AGAW1","11E","INS","BOLD:ABX2489","Arthropoda","Insecta","Coleoptera","Phalacridae","Phalacrinae","Stilbus","","4","4","4","140","0","0","97.22","424","0","422.75","31.74","0"
"AGAW1","11E","INS","BOLD:AAV1295","Arthropoda","Insecta","Diptera","Sciaridae","","Bradysia","Bradysia difformis, Bradysia impatiens","4","4","4","140","0","0","99.13","400.75","0","427.5","32.14","0"
"AGAW1","11E","INS","BOLD:AAZ2075","Arthropoda","Insecta","Hemiptera","Anthocoridae","Anthocorinae","Orius","Orius insidiosus","4","3","3","140","25","0","99.54","435","0","436","35.15","0"
"AGAW1","11E","INS","BOLD:AAG7276","Arthropoda","Insecta","Diptera","Sphaeroceridae","Limosininae","Leptocera","Leptocera erythrocera","3","3","3","149","0","0","99.56","449.67","0","450","33.04","0"
"AGAW1","11E","INS","BOLD:ABA1215","Arthropoda","Insecta","Diptera","Sciaridae","","Lycoriella","Lycoriella sativae","3","3","3","149","0","0","99.85","424.67","0","427","35.46","0"
"AGAW1","11E","INS","BOLD:AAA8933","Arthropoda","Insecta","Coleoptera","Coccinellidae","Coccinellinae","Coccinella","Coccinella septempunctata","3","3","3","149","0","0","99.77","443","0","445.33","33.01","0"
"AGAW1","11E","INS","BOLD:AAA3867","Arthropoda","Insecta","Lepidoptera","Noctuidae","Noctuinae","Apamea","Apamea amputatrix","3","3","3","149","0","0","99.05","409.33","0","411.67","33.61","0"
"AGAW1","11E","INS","BOLD:AAA5299","Arthropoda","Insecta","Diptera","Chironomidae","Orthocladiinae","Cricotopus","Cricotopus laetus, Cricotopus relucens, Cricotopus trifasciatus, Cricotopus sp., Cricotopus sp. 3ES, Cricotopus sylvestris, Cricotopus glacialis","3","3","3","149","0","0","98.55","419.67","0","422","34.64","0"
"AGAW1","11E","INS","BOLD:ACA5755","Arthropoda","Insecta","Diptera","Cecidomyiidae","","","","3","3","3","149","0","0","98.75","422.33","0","424","27.11","0"
"AGAW1","11E","INS","BOLD:ABA1661","Arthropoda","Insecta","Coleoptera","Chrysomelidae","Galerucinae","Longitarsus","Longitarsus pellucidus","3","3","3","149","0","0","99.34","400.67","0","404.33","35.45","0"
"AGAW1","11E","INS","BOLD:AAA2632","Arthropoda","Insecta","Lepidoptera","Noctuidae","Noctuinae","Noctua","Noctua atlantica, Noctua pronuba","3","3","3","149","0","0","98.96","450","0","450","31.04","0"
"AGAW1","11E","INS","BOLD:AAG5499","Arthropoda","Insecta","Diptera","Chironomidae","Orthocladiinae","Psectrocladius","Psectrocladius obvius, Psectrocladius platypus","3","3","3","149","0","0","99.13","422","0","424.33","35.66","0"
"AGAW1","11E","INS","BOLD:AAH2293","Arthropoda","Insecta","Diptera","Asilidae","Stenopogoninae","Dioctria, Eudioctria","Dioctria hyalipennis, Dioctria baumhaueri, Eudioctria propinqua","3","3","3","149","0","0","99.75","376.33","0","377.67","30.34","0"
"AGAW1","11E","INS","BOLD:ADB4698","Arthropoda","Insecta","Diptera","Cecidomyiidae","","","","3","3","3","149","0","0","98.81","405.67","0","408","31.47","0"
"AGAW1","11E","INS","BOLD:ACV0619","Arthropoda","Insecta","Diptera","Cecidomyiidae","","","","3","3","3","149","0","0","99.2","408","0","409.67","25.82","0"
"AGAW1","11E","INS","BOLD:AAB1352","Arthropoda","Insecta","Lepidoptera","Erebidae","Hypeninae","Hypena","Hypena sp.","3","3","3","149","0","0","99.04","449.67","0","450","33.11","0"
"AGAW1","11E","INS","BOLD:AAP8816","Arthropoda","Insecta","Diptera","Cecidomyiidae","Cecidomyiinae","Sitodiplosis","Sitodiplosis sp. BOLD:AAP8816","3","3","3","149","0","0","99.93","431.33","0","434","26.17","0"
"AGAW1","11E","INS","BOLD:AAY6412","Arthropoda","Insecta","Diptera","Sciaridae","","Bradysia","Bradysia urticae","3","3","3","149","0","0","99.53","418.33","0","420.33","32.21","0"
"AGAW1","11E","INS","BOLD:AAG2740","Arthropoda","Insecta","Diptera","Ephydridae","Parydrinae","Philygria","Philygria debilis","3","3","3","149","0","0","98.47","419.33","0","423.33","32.56","0"
"AGAW1","11E","INS","BOLD:AAA3933","Arthropoda","Insecta","Lepidoptera","Erebidae","Herminiinae","Palthis","Palthis angulalis","2","2","2","165","0","0","98.1","448","0","450","31.56","0"
"AGAW1","11E","INS","BOLD:AAF3338","Arthropoda","Insecta","Lepidoptera","Noctuidae","Noctuinae","Striacosta","Striacosta albicosta","2","2","2","165","0","0","98.16","381","0","382","33.64","0"
"AGAW1","11E","INS","BOLD:ACM8528","Arthropoda","Insecta","Diptera","Limoniidae","Chioneinae","Erioptera","Erioptera chlorophylla","2","1","1","165","50","0","98.67","450","0","450","33.78","0"
"AGAW1","11E","INS","BOLD:AAL1297","Arthropoda","Insecta","Coleoptera","Cantharidae","Cantharinae","Rhagonycha","Rhagonycha fulva","2","2","2","165","0","0","99.77","400.5","0","405","33.78","0"
"AGAW1","11E","INS","BOLD:ADF2969","Arthropoda","Insecta","Diptera","Sarcophagidae","Sarcophaginae","Blaesoxipha","Blaesoxipha falciformis","2","2","2","165","0","0","99.2","431.5","0","433","37.64","0"
"AGAW1","11E","INS","BOLD:ACV2677","Arthropoda","Insecta","Diptera","Cecidomyiidae","","","","2","2","2","165","0","0","97.22","449.5","0","450","30.11","0"
"AGAW1","11E","INS","BOLD:AAL7442","Arthropoda","Insecta","Diptera","Cecidomyiidae","","","","2","2","2","165","0","0","98.1","448.5","0","450","25.44","0"
"AGAW1","11E","INS","BOLD:AAU8592","Arthropoda","Insecta","Hymenoptera","Braconidae","Alysiinae","","","2","1","1","165","50","0","99.33","449","0","450","25.33","0"
"AGAW1","11E","INS","BOLD:AAV5575","Arthropoda","Insecta","Diptera","Cecidomyiidae","","","CecidInt30 sp. BOLD:AAV5575","2","2","2","165","0","0","99.65","429.5","0","430.5","26.13","0"
"AGAW1","11E","INS","BOLD:AAH3664","Arthropoda","Insecta","Diptera","Cecidomyiidae","","","CecidInt3 sp. BOLD:AAH3664","2","2","2","165","0","0","98.26","425","0","450","27.78","0"
"AGAW1","11E","INS","BOLD:ADH4973","Arthropoda","Insecta","Diptera","Tipulidae","Tipulinae","Nephrotoma","","2","2","2","165","0","0","97.33","393.5","0","397","33.75","0"
"AGAW1","11E","INS","BOLD:AAY6476","Arthropoda","Insecta","Diptera","Cecidomyiidae","","","","2","2","2","165","0","0","99.33","450","0","450","25.22","0"
"AGAW1","11E","INS","BOLD:AAB0651","Arthropoda","Insecta","Lepidoptera","Gelechiidae","Dichomeridinae","Dichomeris","Dichomeris leuconotella, Dichomeris gnoma","2","2","2","165","0","0","98.56","417.5","0","418","33.12","0"
"AGAW1","11E","INS","BOLD:AAY6450","Arthropoda","Insecta","Diptera","Cecidomyiidae","","","","2","2","2","165","0","0","99.89","449","0","450","27.89","0"
"AGAW1","11E","INS","BOLD:AAI4303","Arthropoda","Insecta","Diptera","Chironomidae","Chironominae","Chironomus","Chironomus melanescens","2","2","2","165","0","0","98.73","414.5","0","418","33.99","0"
"AGAW1","11E","INS","BOLD:ABA5839","Arthropoda","Insecta","Hemiptera","Cicadellidae","Deltocephalinae","Euscelis","Euscelis aemulans, Euscelis confinis","2","2","2","165","0","0","97.91","385.5","0","386.5","36.1","0"
"AGAW1","11E","INS","BOLD:AAA1513","Arthropoda","Insecta","Lepidoptera","Plutellidae","Plutellinae","Plutella, Rhigognostis","Plutella xylostella, Rhigognostis sp.","2","2","2","165","0","0","98.85","421.5","0","422.5","31.93","0"
"AGAW1","11E","INS","BOLD:AAA3181","Arthropoda","Insecta","Diptera","Muscidae","Muscinae","Stomoxys","Stomoxys calcitrans","1","1","1","182","0","0","99.11","448","0","450","33.56","0"
"AGAW1","11E","INS","BOLD:AAA6759","Arthropoda","Insecta","Lepidoptera","Hesperiidae","Hesperiinae","Thymelicus","Thymelicus lineola","1","1","1","182","0","0","99.78","448","0","450","30.22","0"
"AGAW1","11E","INS","BOLD:AAH4701","Arthropoda","Insecta","Lepidoptera","Pterophoridae","Pterophorinae","Sphenarches","Sphenarches ontario","1","1","1","182","0","0","97.15","421","0","422","30.33","0"
"AGAW1","11E","INS","BOLD:AAA6822","Arthropoda","Insecta","Orthoptera","Acrididae","Gomphocerinae","Chorthippus","Chorthippus curtipennis","1","1","1","182","0","0","99.19","371","0","372","35.48","0"
"AGAW1","11E","INS","BOLD:ACJ6954","Arthropoda","Insecta","Diptera","Cecidomyiidae","","","","1","1","1","182","0","0","100","337","0","392","25.77","0"
"AGAW1","11E","INS","BOLD:AAF9028","Arthropoda","Insecta","Diptera","Limoniidae","","","","1","1","1","182","0","0","97.19","427","0","427","37.94","0"
"AGAW1","11E","INS","BOLD:AAN5249","Arthropoda","Insecta","Diptera","Cecidomyiidae","","","","1","1","1","182","0","0","100","448","0","450","33.11","0"
"AGAW1","11E","INS","BOLD:AAA2518","Arthropoda","Insecta","Lepidoptera","Crambidae","Crambinae","Chrysoteuchia","Chrysoteuchia topiaria, Chrysoteuchia topiarius","1","1","1","182","0","0","99.78","450","0","450","35.78","0"
"AGAW1","11E","INS","BOLD:ACF0915","Arthropoda","Insecta","Diptera","Chironomidae","Orthocladiinae","","","1","1","1","182","0","0","97.15","351","0","356","32.3","0"
"AGAW1","11E","INS","BOLD:AAP4633","Arthropoda","Insecta","Psocodea","Psocidae","Psocinae","Loensia","","1","1","1","182","0","0","99.56","450","0","450","36.89","0"
"AGAW1","11E","INS","BOLD:ACU7271","Arthropoda","Insecta","Diptera","Cecidomyiidae","","","","1","1","1","182","0","0","99.44","360","0","361","31.86","0"
"AGAW1","11E","INS","BOLD:AAN5225","Arthropoda","Insecta","Diptera","Cecidomyiidae","","","","1","1","1","182","0","0","97.04","439","0","439","29.38","0"
"AGAW1","11E","INS","BOLD:ACA3626","Arthropoda","Insecta","Diptera","Ceratopogonidae","Forcipomyiinae","Atrichopogon","Atrichopogon sp. 2, Atrichopogon sp. BOLD:ACA3626","1","1","1","182","0","0","99.55","447","0","450","34.44","0"
"AGAW1","11E","INS","BOLD:AAD0952","Arthropoda","Insecta","Lepidoptera","Pyralidae","Chrysauginae","Galasa","Galasa nigrinodis, Galasa nigripunctalis","1","1","1","182","0","0","99.77","427","0","450","33.11","0"
"AGAW1","11E","INS","BOLD:AAD7604","Arthropoda","Insecta","Coleoptera","Coccinellidae","Coccinellinae","Coleomegilla","Coleomegilla maculata","1","1","1","182","0","0","99.33","449","0","450","31.78","0"
"AGAW1","11E","INS","BOLD:AAB4658","Arthropoda","Insecta","Diptera","Chironomidae","Chironominae","Chironomus","Chironomus dilutus","1","1","1","182","0","0","98.88","445","0","450","38","0"
"AGAW1","11E","INS","BOLD:AAB8143","Arthropoda","Insecta","Hemiptera","Nabidae","Nabinae","Nabis","Nabis inscriptus, Nabis americoferus, Nabis alternatus, Nabis sp.","1","1","1","182","0","0","99.78","450","0","450","36.89","0"
"AGAW1","11E","INS","BOLD:AAG5457","Arthropoda","Insecta","Diptera","Chironomidae","Chironominae","","","1","1","1","182","0","0","97.92","432","0","438","33.79","0"
"AGAW1","11E","INS","BOLD:AAG3273","Arthropoda","Insecta","Diptera","Phoridae","Metopininae","Megaselia","Megaselia albicaudata","1","1","1","182","0","0","97.56","450","0","450","32.67","0"
"AGAW1","11E","INS","BOLD:AAG6929","Arthropoda","Insecta","Diptera","Dolichopodidae","Medeterinae","Medetera","Medetera vittata","1","1","1","182","0","0","99.11","448","0","450","41.11","0"
"AGAW1","11E","INS","BOLD:ABX5883","Arthropoda","Insecta","Lepidoptera","Tortricidae","Tortricinae","Choristoneura","Choristoneura retiniana, Choristoneura lambertiana, Choristoneura biennis, Choristoneura orae, Choristoneura fumiferana, Choristoneura occidentalis, Choristoneura freemani, Choristoneura pinus, Choristoneura occidentalis group, Choristoneura sp., Choristoneura carnana","1","1","1","182","0","0","99.11","450","0","450","31.56","0"
"AGAW1","11E","INS","BOLD:ABZ3705","Arthropoda","Insecta","Lepidoptera","Crambidae","Pyraustinae","Sitochroa","Sitochroa chortalis","1","1","1","182","0","0","100","445","0","445","29.89","0"
"AGAW1","11E","INS","BOLD:ACX5521","Arthropoda","Insecta","Diptera","Chironomidae","Orthocladiinae","","","1","1","1","182","0","0","99.2","374","0","378","35.71","0"
"AGAW1","11E","INS","BOLD:AAG5639","Arthropoda","Insecta","Diptera","Sepsidae","Sepsinae","Sepsis","Sepsis punctum","1","1","1","182","0","0","98.39","434","0","436","34.86","0"
"AGAW1","11E","INS","BOLD:AAH3914","Arthropoda","Insecta","Diptera","Sciaridae","","Sciara","Sciara hamatilis, Sciara humeralis","1","1","1","182","0","0","99.33","447","0","450","30.89","0"
"AGAW1","11E","INS","BOLD:AAA8640","Arthropoda","Insecta","Lepidoptera","Tortricidae","Olethreutinae","Eucosma, Sonia, Pelochrista","Pelochrista suadana, Sonia new sp. 1, Sonia new sp. 2, Sonia new sp. 3, Sonia paraplesiana, Sonia canadana","1","1","1","182","0","0","99.78","449","0","450","32.89","0"
"AGAW1","11E","INS","BOLD:AAG8882","Arthropoda","Insecta","Hemiptera","Pentatomidae","Pentatominae","Thyanta","Thyanta pallidovirens, Thyanta custator, Thyanta accerra","1","1","1","182","0","0","98.88","445","0","446","36.55","0"
"AGAW1","11E","INS","BOLD:AAV1136","Arthropoda","Insecta","Diptera","Scatopsidae","","","","1","1","1","182","0","0","98.67","450","0","450","37.11","0"
"AGAW1","11E","INS","BOLD:ACN6040","Arthropoda","Insecta","Diptera","Chironomidae","Chironominae","Tanytarsus","Tanytarsus glabrescens","1","1","1","182","0","0","97.1","449","0","450","36.89","0"
"AGAW1","11E","INS","BOLD:ACA9284","Arthropoda","Insecta","Hymenoptera","Platygastridae","","","","1","1","1","182","0","0","98.41","440","0","450","28","0"
"AGAW1","11E","INS","BOLD:AAM6286","Arthropoda","Insecta","Diptera","Chironomidae","Chironominae","","","1","1","1","182","0","0","97.45","392","0","394","34.77","0"
"AGAW1","11E","INS","BOLD:AAZ5483","Arthropoda","Insecta","Diptera","Muscidae","Phaoniinae","Helina","Helina setiventris","1","1","1","182","0","0","99.03","413","0","417","28.06","0"
"AGAW1","11E","INS","BOLD:AAN5342","Arthropoda","Insecta","Diptera","Chironomidae","Orthocladiinae","Heleniella","","1","1","1","182","0","0","98.28","408","0","410","34.39","0"
"AGAW1","11E","INS","BOLD:AAA2482","Arthropoda","Insecta","Lepidoptera","Noctuidae","Noctuinae","Mythimna","Mythimna unipuncta","1","1","1","182","0","0","97.18","426","0","433","30.02","0"
"AGAW1","11E","INS","BOLD:AAG6431","Arthropoda","Insecta","Diptera","Ceratopogonidae","Forcipomyiinae","Atrichopogon","","1","1","1","182","0","0","97.32","448","0","450","36.89","0"
"AGAW1","11E","INS","BOLD:ACV5891","Arthropoda","Insecta","Diptera","Cecidomyiidae","","","","1","1","1","182","0","0","98.62","362","0","384","23.96","0"
"AGAW1","11E","INS","BOLD:AAY6468","Arthropoda","Insecta","Diptera","Cecidomyiidae","","","","1","1","1","182","0","0","100","450","0","450","29.11","0"
"AGAW1","11E","INS","BOLD:ABV1203","Arthropoda","Insecta","Diptera","Chironomidae","Orthocladiinae","","","1","1","1","182","0","0","99.04","418","0","418","33.49","0"
"AGAW1","11E","INS","BOLD:AAH4171","Arthropoda","Insecta","Diptera","Chloropidae","Oscinellinae","Apallates","Apallates montanus, Apallates particeps","1","1","1","182","0","0","97.33","450","0","450","33.11","0"
"AGAW1","11E","INS","BOLD:AAA5206","Arthropoda","Insecta","Lepidoptera","Erebidae","Herminiinae","Zanclognatha","Zanclognatha cruralis, Zanclognatha sp. 1, Zanclognatha jacchusalis, Zanclognatha minoralis, Zanclognatha minoralis sp. 1, Zanclognatha sp., Zanclognatha minoralis sp. 3, Zanclognatha gypsalis, Zanclognatha dentata, Zanclognatha theralis, Zanclognatha protumnusalis, Zanclognatha martha, Zanclognatha atrilineella, Zanclognatha obscuripennis, Zanclognatha minoralis sp. 2, Zanclognatha marcidilinea","1","1","1","182","0","0","97.62","378","0","381","31.23","0"
"AGAW1","11E","INS","BOLD:AAB9332","Arthropoda","Insecta","Hemiptera","Aphididae","Aphidinae","Nasonovia","Nasonovia ribisnigri","1","1","1","182","0","0","99.78","445","0","450","25.33","0"
"AGAW1","11E","INS","BOLD:ADB1122","Arthropoda","Insecta","Hemiptera","Cicadellidae","Deltocephalinae","Macustus","Macustus grisescens","1","1","1","182","0","0","97.61","376","0","377","33.69","0"
"AGAW1","11E","INS","BOLD:AAB1850","Arthropoda","Insecta","Hemiptera","Aphrophoridae","Aphrophorinae","Philaenus","Philaenus tesselatus, Philaenus spumarius, Philaenus maghresignus","1","1","1","182","0","0","98","449","0","450","32.67","0"
"AGAW1","11E","INS","BOLD:ACL6109","Arthropoda","Insecta","Psocodea","Psocidae","","","","1","1","1","182","0","0","98.09","366","0","370","35.95","0"
"AGAW1","11E","INS","BOLD:AAA6560","Arthropoda","Insecta","Lepidoptera","Crambidae","Crambinae","Crambus","Crambus multilinellus, Crambus saltuellus","1","1","1","182","0","0","99.51","405","0","410","29.51","0"
"AGAW1","11E","INS","BOLD:AAB0269","Arthropoda","Insecta","Lepidoptera","Tortricidae","Tortricinae","Acleris","Acleris albicomana","1","1","1","182","0","0","98.67","450","0","450","33.11","0"
"AGAW1","11E","INS","BOLD:ABW7905","Arthropoda","Insecta","Diptera","Cecidomyiidae","","","","1","1","1","182","0","0","97.9","429","0","450","26.22","0"
"AGAW1","11E","INS","BOLD:ACA7531","Arthropoda","Insecta","Hymenoptera","Megachilidae","Megachilinae","Heriades","","1","1","1","182","0","0","97.76","447","0","450","23.78","0"
"AGAW1","11E","INS","BOLD:AAB1098","Arthropoda","Insecta","Diptera","Culicidae","Culicinae","Ochlerotatus, Aedes","Aedes excrucians, Aedes cantans, Aedes stimulans, Aedes euedes, Aedes riparius, Aedes annulipes, Ochlerotatus cantans, Aedes fitchii, Aedes sp., Ochlerotatus annulipes, Ochlerotatus excrucians","1","1","1","182","0","0","97.75","445","0","450","34.89","0"
"AGAW1","11E","INS","BOLD:AAL2750","Arthropoda","Insecta","Diptera","Muscidae","Phaoniinae","Phaonia","Phaonia tuguriorum","1","1","1","182","0","0","98.25","400","0","403","31.02","0"
"AGAW1","11E","INS","BOLD:ABX6298","Arthropoda","Insecta","Diptera","Tipulidae","Tipulinae","Nephrotoma","Nephrotoma ferruginea","1","1","1","182","0","0","99.78","450","0","450","33.78","0"
"AGAW1","11E","INS","BOLD:ACL7238","Arthropoda","Insecta","Diptera","Xylomyidae","","Solva","","1","1","1","182","0","0","97.75","444","0","450","35.33","0"
"AGAW1","11E","INS","BOLD:AAV6112","Arthropoda","Insecta","Diptera","Chloropidae","Oscinellinae","Oscinella","Oscinella sp. 2 AGT, Oscinella sp. TAW6","1","1","1","182","0","0","100","450","0","450","33.33","0"
"AGAW1","11E","INS","BOLD:ABA6407","Arthropoda","Insecta","Diptera","Sciaridae","","Pseudolycoriella","","1","1","1","182","0","0","99.78","450","0","450","35.11","0"
"AGAW1","11E","INS","BOLD:AAG7042","Arthropoda","Insecta","Diptera","Agromyzidae","","","","1","1","1","182","0","0","98.2","444","0","450","32.44","0"
"AGAW1","11E","INS","BOLD:ACE9865","Arthropoda","Insecta","Lepidoptera","Noctuidae","Noctuinae","Feltia","Feltia subgothica, Feltia herilis, Feltia tricosa","1","1","1","182","0","0","100","450","0","450","30.89","0"
"AGAW1","11E","INS","BOLD:ACA2933","Arthropoda","Insecta","Psocodea","Stenopsocidae","","Graphopsocus","Graphopsocus cruciatus","1","1","1","182","0","0","99.73","370","0","372","38.71","0"
"AGAW1","11E","INS","BOLD:AAV5356","Arthropoda","Insecta","Diptera","Cecidomyiidae","","","","1","1","1","182","0","0","98.65","446","0","450","31.11","0"
"AGAW1","11E","INS","BOLD:AAD1169","Arthropoda","Insecta","Lepidoptera","Cosmopterigidae","Cosmopteriginae","Cosmopterix","Cosmopterix scirpicola, Cosmopterix fernaldella","1","1","1","182","0","0","99.78","449","0","450","32.89","0"
"AGAW1","11E","INS","BOLD:AAP6882","Arthropoda","Insecta","Diptera","Chironomidae","Chironominae","Dicrotendipes","","1","1","1","182","0","0","97.09","447","0","450","36.67","0"
"AGAW1","11E","INS","BOLD:ABX0130","Arthropoda","Insecta","Diptera","Cecidomyiidae","","","CecidInt43 sp. BOLD:ABX0130","1","1","1","182","0","0","99.76","421","0","422","29.86","0"
"AGAW1","11E","INS","BOLD:AAM6230","Arthropoda","Insecta","Diptera","Chironomidae","Chironominae","Chironomus","","1","1","1","182","0","0","97.28","441","0","450","34.67","0"
"AGAW1","11E","INS","BOLD:ADH9310","Arthropoda","Insecta","Diptera","Anthomyiidae","Anthomyiinae","Anthomyia","Anthomyia procellaris, Anthomyia pluvialis","1","1","1","182","0","0","99.03","412","0","416","31.25","0"
"AGAW1","11E","INS","BOLD:AAN6433","Arthropoda","Insecta","Diptera","Sciaridae","","Corynoptera","Corynoptera deserta","1","1","1","182","0","0","99.33","449","0","450","30.44","0"
"AGAW1","11E","INS","BOLD:AAN8310","Arthropoda","Insecta","Hemiptera","Cicadellidae","Deltocephalinae","Opsius","Opsius stactogalus","1","1","1","182","0","0","100","449","0","450","37.78","0"
"AGAW1","11E","INS","BOLD:AAO7890","Arthropoda","Insecta","Diptera","Agromyzidae","","","","1","1","1","182","0","0","99.5","398","0","450","34.22","0"
"AGAW1","11E","INS","BOLD:ACB0984","Arthropoda","Insecta","Psocodea","Stenopsocidae","","Graphopsocus","Graphopsocus cruciatus","1","1","1","182","0","0","98.79","413","0","450","37.11","0"
"AGAW1","11E","INS","BOLD:AAC0661","Arthropoda","Insecta","Coleoptera","Carabidae","Harpalinae","Pterostichus","Pterostichus melanarius","1","1","1","182","0","0","98.71","387","0","390","31.79","0"
"AGAW1","11E","INS","BOLD:AAA9422","Arthropoda","Insecta","Hemiptera","Cicadellidae","Deltocephalinae","Macrosteles","Macrosteles quadrilineatus","1","1","1","182","0","0","99.52","416","0","450","37.11","0"
"AGAW1","11E","INS","BOLD:AAB6095","Arthropoda","Insecta","Lepidoptera","Noctuidae","Noctuinae","Neoligia","Neoligia exhausta","1","1","1","182","0","0","100","450","0","450","32.44","0"
"AGAW1","11E","INS","BOLD:ABA0772","Arthropoda","Insecta","Diptera","Chironomidae","Chironominae","Polypedilum","","1","1","1","182","0","0","97.8","364","0","365","32.88","0"
"AGAW1","11E","INS","BOLD:AAY6534","Arthropoda","Insecta","Coleoptera","Mordellidae","Mordellinae","Mordellina","Mordellina ancilla","1","1","1","182","0","0","100","450","0","450","38.89","0"
"AGAW1","11E","NAR","BOLD:AAI9663","Mollusca","Gastropoda","Stylommatophora","Agriolimacidae",NA,"Deroceras","Deroceras reticulatum","2","2","2","1","0","0","99.75","404.5","0","417.5","35.44","0"
"AGAW1","11E","NIN","BOLD:AAU6252","Arthropoda","Arachnida","Trombidiformes","Bdellidae","","Cyta","","197","174","174","1","11.68","0","98.63","430.72","0","432.15","31.41","0"
"AGAW1","11E","NIN","BOLD:AAA7242","Arthropoda","Collembola","Entomobryomorpha","Entomobryidae","Seirinae, Entomobryinae","Entomobrya, Lepidosira","Entomobrya multifasciata, Lepidosira sp.","123","121","121","2","1.63","0","99.12","421.13","0","424.29","39.3","0"
"AGAW1","11E","NIN","BOLD:ABW2639","Arthropoda","Arachnida","Trombidiformes","Anystidae","","","","73","61","61","3","16.44","0","99.63","447.51","0","446","31.44","0"
"AGAW1","11E","NIN","BOLD:AAH6630","Arthropoda","Collembola","Entomobryomorpha","Entomobryidae","Entomobryinae","Entomobrya","Entomobrya unostrigata","53","48","48","4","9.43","0","99.67","434.55","0","435.58","38.91","0"
"AGAW1","11E","NIN","BOLD:ACY2550","Arthropoda","Arachnida","Trombidiformes","Bdellidae","","Cyta","","27","26","26","5","3.7","0","98.82","375.48","0","391.48","31.31","0"
"AGAW1","11E","NIN","BOLD:ACL6413","Arthropoda","Arachnida","Trombidiformes","Erythraeidae","","Balaustium","","19","19","19","6","0","0","99.44","428.74","0","429.68","35.87","0"
"AGAW1","11E","NIN","BOLD:ACN8290","Arthropoda","Arachnida","Araneae","Trachelidae","","Trachelas","Trachelas tranquillus","15","15","15","7","0","0","99.45","432.87","0","435.13","34.19","0"
"AGAW1","11E","NIN","BOLD:ACZ1295","Arthropoda","Arachnida","Trombidiformes","Anystidae","","","","14","14","14","8","0","0","99.39","377.64","0","385.43","31.3","0"
"AGAW1","11E","NIN","BOLD:AAF9285","Arthropoda","Arachnida","Mesostigmata","Parasitidae","","","","6","6","6","9","0","0","98.92","431.33","0","432.83","35.64","0"
"AGAW1","11E","NIN","BOLD:AAN6706","Arthropoda","Arachnida","Mesostigmata","Ascidae","","","","6","6","6","9","0","0","98.83","441.33","0","442","34.64","0"
"AGAW1","11E","NIN","BOLD:AAM7961","Arthropoda","Arachnida","Trombidiformes","Anystidae","","","","3","3","3","11","0","0","99.7","436.33","0","426","31.46","0"
"AGAW1","11E","NIN","BOLD:AAF9234","Arthropoda","Arachnida","Trombidiformes","Eupodidae","","","","3","3","3","11","0","0","99.11","447.33","0","450","30.07","0"
"AGAW1","11E","NIN","BOLD:AAB7914","Arthropoda","Collembola","Symphypleona","Bourletiellidae","","Bourletiella","","3","3","3","11","0","0","99.7","449","0","450","38.52","0"
"AGAW1","11E","NIN","BOLD:ACX8969","Arthropoda","Arachnida","Trombidiformes","Eupodidae","","","","3","3","3","11","0","0","98.8","421","0","423.33","28.33","0"
"AGAW1","11E","NIN","BOLD:AAM3234","Arthropoda","Collembola","Entomobryomorpha","Entomobryidae","Seirinae","Lepidosira","","2","2","2","15","0","0","98.55","449.5","0","450","39.22","0"
"AGAW1","11E","NIN","BOLD:AAE2695","Arthropoda","Arachnida","Sarcoptiformes","Oribatulidae","","Zygoribatula","","1","1","1","16","0","0","97.88","378","0","378","39.15","0"
"AGAW1","11E","NIN","BOLD:AAA8820","Arthropoda","Collembola","Entomobryomorpha","Entomobryidae","Lepidocyrtinae","Lepidocyrtus","Lepidocyrtus cyaneus","1","1","1","16","0","0","98.92","369","0","371","43.94","0"
"AGAW1","11E","NIN","BOLD:ACN3178","Arthropoda","Collembola","Entomobryomorpha","Entomobryidae","","","","1","1","1","16","0","0","98.67","450","0","450","39.33","0"
"AGAW1","11E","NIN","BOLD:AAB7601","Arthropoda","Arachnida","Araneae","Eutichuridae","","Cheiracanthium","Cheiracanthium mildei","1","1","1","16","0","0","98.22","449","0","450","30.22","0"
"AGAW1","11E","NIN","BOLD:ADE3666","Arthropoda","Arachnida","Mesostigmata","Phytoseiidae","","","","1","1","1","16","0","0","97.87","376","0","405","33.33","0"
"AGAW1","11E","NIN","BOLD:AAI4346","Arthropoda","Arachnida","Opiliones","Phalangiidae","","Paroligolophus, Phalangium","Paroligolophus agrestis, Phalangium opilio","1","1","1","16","0","0","97.47","396","0","396","35.1","0"
"AGAW1","11E","NIN","BOLD:AAI6247","Arthropoda","Arachnida","Araneae","Dictynidae","","Dictyna","Dictyna foliacea","1","1","1","16","0","0","99.21","380","0","382","32.2","0"
"AGAW1","11E","NIN","BOLD:ACP9512","Arthropoda","Collembola","Entomobryomorpha","Entomobryidae","","","","1","1","1","16","0","0","98.35","424","0","450","39.56","0"
"AGAW1","11N","BAC","TAX:91393","Proteobacteria","Alphaproteobacteria","Rickettsiales","Rickettsiaceae",NA,"Wolbachia","","39","39","39","1","0","0","99.44","391.18","0","420.15","39.11","0"
"AGAW1","11N","BAC","TAX:175768","Proteobacteria","Alphaproteobacteria","Rickettsiales","Rickettsiaceae",NA,"Wolbachia","Wolbachia pipientis","38","27","27","2","28.95","0","98.68","447.89","0","448.76","39.6","0"
"AGAW1","11N","BAC","TAX:91391","Proteobacteria","Alphaproteobacteria","Rickettsiales","",NA,"","","4","4","4","3","0","0","99.27","336.75","0","417.25","38.21","0"
"AGAW1","11N","BAC","TAX:375762","Proteobacteria","Alphaproteobacteria","Rickettsiales","Rickettsiaceae",NA,"Wolbachia","Wolbachia sp. Myrmicinae_ACGAN556-09","1","1","1","4","0","0","97.33","450","0","450","39.56","0"
"AGAW1","11N","BAC","TAX:376688","Proteobacteria","Alphaproteobacteria","Rickettsiales","Rickettsiaceae",NA,"Wolbachia","Wolbachia sp. elachBioLep01 BioLep373","1","1","1","4","0","0","97.11","450","0","450","39.33","0"
"AGAW1","11N","CHO","no match","no match","no match","no match","no match","no match","no match","no match","no match","no match","no match","no match","no match","no match","no match","no match","no match","no match","no match","no match"
"AGAW1","11N","INS","BOLD:AAP8739","Arthropoda","Insecta","Diptera","Platystomatidae","Platystomatinae","Rivellia","Rivellia steyskali","10282","6741","6741","1","34.44","0","99.34","440.94","0","442.59","38","0"
"AGAW1","11N","INS","BOLD:AAB4321","Arthropoda","Insecta","Lepidoptera","Gelechiidae","Gelechiinae","Metzneria","Metzneria lappella","6784","6479","6479","2","4.5","0","99.03","408.64","0","425.53","31.63","0"
"AGAW1","11N","INS","BOLD:AAF3338","Arthropoda","Insecta","Lepidoptera","Noctuidae","Noctuinae","Striacosta","Striacosta albicosta","5776","5352","5352","3","7.34","0","98.61","417.46","0","419.77","33.86","0"
"AGAW1","11N","INS","BOLD:AAA3453","Arthropoda","Insecta","Diptera","Anthomyiidae","Anthomyiinae","Delia","","5676","5079","5079","4","10.52","0","99.35","430.18","0","432.98","34.61","0"
"AGAW1","11N","INS","BOLD:AAB0269","Arthropoda","Insecta","Lepidoptera","Tortricidae","Tortricinae","Acleris","Acleris albicomana","4904","3990","3990","5","18.64","0","99.38","426.11","0","427.32","32.39","0"
"AGAW1","11N","INS","BOLD:AAU6681","Arthropoda","Insecta","Diptera","Tipulidae","Tipulinae","Nephrotoma","Nephrotoma cornicina","4717","3573","3573","6","24.25","0","99.6","442.53","0","443.77","36.31","0"
"AGAW1","11N","INS","BOLD:AAJ7123","Arthropoda","Insecta","Diptera","Culicidae","Culicinae","Culiseta","Culiseta minnesotae","4376","3418","3418","7","21.89","0","98.35","434.43","0","435.78","34.37","0"
"AGAW1","11N","INS","BOLD:AAG2511","Arthropoda","Insecta","Diptera","Anthomyiidae","Anthomyiinae","Delia","Delia platura","4122","3527","3527","8","14.43","0","99.07","443.71","0","446.76","33.13","0"
"AGAW1","11N","INS","BOLD:AAA7368","Arthropoda","Insecta","Lepidoptera","Cosmopterigidae","Cosmopteriginae","Limnaecia","Limnaecia phragmitella","1464","1265","1265","9","13.59","0","99.41","425.55","0","429.3","31","0"
"AGAW1","11N","INS","BOLD:ABX6298","Arthropoda","Insecta","Diptera","Tipulidae","Tipulinae","Nephrotoma","Nephrotoma ferruginea","1332","1192","1192","10","10.51","0","99.3","428.13","0","430.33","33.24","0"
"AGAW1","11N","INS","BOLD:AAA7661","Arthropoda","Insecta","Diptera","Culicidae","Culicinae","Culiseta, Culex","Culiseta melanura, Culex articularis, Culex restuans, Culex acharistus","1307","1209","1209","11","7.5","0","99.2","430.03","0","432.01","32.99","0"
"AGAW1","11N","INS","BOLD:ACA2933","Arthropoda","Insecta","Psocodea","Stenopsocidae","","Graphopsocus","Graphopsocus cruciatus","1020","725","725","12","28.92","0","99.66","442.19","0","443.36","38.55","0"
"AGAW1","11N","INS","BOLD:AAG2123","Arthropoda","Insecta","Diptera","Tachinidae","Dexiinae","Dinera","Dinera grisescens","947","744","744","13","21.44","0","99.37","432.3","0","434.26","31.87","0"
"AGAW1","11N","INS","BOLD:AAI4303","Arthropoda","Insecta","Diptera","Chironomidae","Chironominae","Chironomus","Chironomus melanescens","946","880","880","14","6.98","0","98.44","427.25","0","430.99","33.83","0"
"AGAW1","11N","INS","BOLD:ABW7897","Arthropoda","Insecta","Diptera","Cecidomyiidae","","","","771","543","543","15","29.57","0","99.63","443.66","0","445.65","29.7","0"
"AGAW1","11N","INS","BOLD:AAN5225","Arthropoda","Insecta","Diptera","Cecidomyiidae","","","","700","505","505","16","27.86","0","99.16","445.51","0","446.47","28.15","0"
"AGAW1","11N","INS","BOLD:AAN6154","Arthropoda","Insecta","Coleoptera","Latridiidae","Corticariinae","Melanophthalma","Melanophthalma inermis","661","525","525","17","20.57","0","98.9","434.89","0","437.01","31.44","0"
"AGAW1","11N","INS","BOLD:AAG5055","Arthropoda","Insecta","Diptera","Xylomyidae","","Solva","","651","495","495","18","23.96","0","99.44","446.48","0","447.76","35.03","0"
"AGAW1","11N","INS","BOLD:AAN5162","Arthropoda","Insecta","Diptera","Ceratopogonidae","Dasyheleinae","Dasyhelea","","625","584","584","19","6.56","0","99.22","431.61","0","433.6","33.64","0"
"AGAW1","11N","INS","BOLD:AAZ9744","Arthropoda","Insecta","Hymenoptera","Braconidae","Brachistinae","Blacus","Blacus stelfoxi","576","407","407","20","29.34","0","98.41","445.12","0","446.39","32.31","0"
"AGAW1","11N","INS","BOLD:AAU6592","Arthropoda","Insecta","Diptera","Ceratopogonidae","Dasyheleinae","Dasyhelea","Dasyhelea arenivaga","551","532","532","21","3.45","0","99.21","396.74","0","398.99","34.55","0"
"AGAW1","11N","INS","BOLD:AAA4751","Arthropoda","Insecta","Diptera","Culicidae","Culicinae","Culex","Culex pipiens sl, Culex nr. pipiens, Culex pipiens, Culex quinquefasciatus, Culex sp.","516","487","487","22","5.62","0","99.19","425.58","0","428.13","33.07","0"
"AGAW1","11N","INS","BOLD:AAG2868","Arthropoda","Insecta","Hemiptera","Cicadellidae","Typhlocybinae","Empoasca","Empoasca sp., Empoasca fabae","480","325","325","23","32.29","0","99.63","443.5","0","446.56","32.3","0"
"AGAW1","11N","INS","BOLD:ABA4732","Arthropoda","Insecta","Lepidoptera","Cosmopterigidae","Chrysopeleiinae","Perimede","Perimede falcata","473","463","463","24","2.11","0","99.13","423.51","0","425.68","32.64","0"
"AGAW1","11N","INS","BOLD:AAN5199","Arthropoda","Insecta","Diptera","Cecidomyiidae","","","CecidInt20 sp. BOLD:AAN5199","417","326","326","25","21.82","0","99.45","436.2","0","445.09","26.76","0"
"AGAW1","11N","INS","BOLD:ADF6661","Arthropoda","Insecta","Diptera","Cecidomyiidae","","","","364","281","281","26","22.8","0","99.07","416.65","0","445.09","29.9","0"
"AGAW1","11N","INS","BOLD:AAG9897","Arthropoda","Insecta","Dermaptera","Forficulidae","Forficulinae","Forficula","Forficula auricularia-A","340","337","337","27","0.88","0","99.15","396.79","0","407.43","45.76","0"
"AGAW1","11N","INS","BOLD:AAG9626","Arthropoda","Insecta","Diptera","Dolichopodidae","","","","340","293","293","27","13.82","0","98.6","431.45","0","434.37","36.63","0"
"AGAW1","11N","INS","BOLD:AAH3661","Arthropoda","Insecta","Diptera","Cecidomyiidae","","","","277","261","261","29","5.78","0","99.19","428.43","0","430.65","25.32","0"
"AGAW1","11N","INS","BOLD:ACR3866","Arthropoda","Insecta","Psocodea","Caeciliusidae","Caeciliusinae","Valenzuela","","235","216","216","30","8.09","0","99.43","437","0","438.67","35.38","0"
"AGAW1","11N","INS","BOLD:AAV5098","Arthropoda","Insecta","Diptera","Ceratopogonidae","Dasyheleinae","Dasyhelea","Dasyhelea turficola, Dasyhelea sp. 5ES","227","199","199","31","12.33","0","99.19","424.49","0","426.28","36.41","0"
"AGAW1","11N","INS","BOLD:AAC4197","Arthropoda","Insecta","Diptera","Chironomidae","Orthocladiinae","Paraphaenocladius","Paraphaenocladius impensus","193","168","168","32","12.95","0","98.87","431.93","0","434.46","34.61","0"
"AGAW1","11N","INS","BOLD:ACX5521","Arthropoda","Insecta","Diptera","Chironomidae","Orthocladiinae","","","190","176","176","33","7.37","0","99.19","420.87","0","422.68","34.58","0"
"AGAW1","11N","INS","BOLD:ABY7666","Arthropoda","Insecta","Diptera","Culicidae","Culicinae","Culex","Culex territans","187","163","163","34","12.83","0","98.28","442.59","0","444.03","33.81","0"
"AGAW1","11N","INS","BOLD:AAO4092","Arthropoda","Insecta","Psocodea","Caeciliusidae","Caeciliusinae","Valenzuela","Valenzuela burmeisteri","163","163","163","35","0","0","98.92","424.1","0","426.6","34.99","0"
"AGAW1","11N","INS","BOLD:AAN5238","Arthropoda","Insecta","Diptera","Cecidomyiidae","","","","155","132","132","36","14.84","0","99.53","424.8","0","426.85","25.51","0"
"AGAW1","11N","INS","BOLD:ABV0488","Arthropoda","Insecta","Diptera","Cecidomyiidae","","","","143","138","138","37","3.5","0","99","415.32","0","417.42","27.44","0"
"AGAW1","11N","INS","BOLD:ACI3976","Arthropoda","Insecta","Diptera","Chironomidae","Orthocladiinae","","","136","135","135","38","0.74","0","98.72","386.41","0","404.4","37.04","0"
"AGAW1","11N","INS","BOLD:AAN5538","Arthropoda","Insecta","Diptera","Dolichopodidae","","","","128","109","109","39","14.84","0","98.5","438.63","0","439.71","38.85","0"
"AGAW1","11N","INS","BOLD:AAM9013","Arthropoda","Insecta","Diptera","Keroplatidae","Macrocerinae","Macrocera","Macrocera pusilla","128","126","126","39","1.56","0","99.36","431.9","0","433.55","30.33","0"
"AGAW1","11N","INS","BOLD:AAN5161","Arthropoda","Insecta","Diptera","Ceratopogonidae","Dasyheleinae","Dasyhelea","","107","106","106","41","0.93","0","99.1","405.83","0","407.96","37.44","0"
"AGAW1","11N","INS","BOLD:ACE0554","Arthropoda","Insecta","Diptera","Phoridae","Metopininae","Megaselia","","102","92","92","42","9.8","0","99.61","439.25","0","441.8","31.95","0"
"AGAW1","11N","INS","BOLD:AAN5341","Arthropoda","Insecta","Diptera","Chironomidae","Orthocladiinae","Camptocladius","Camptocladius stercorarius","101","100","100","43","0.99","0","98.55","409.29","0","412.74","35.46","0"
"AGAW1","11N","INS","BOLD:AAB5609","Arthropoda","Insecta","Diptera","Muscidae","Coenosiinae","Coenosia","Coenosia sp.BOLD:AAB5609, Coenosia tigrina","94","89","89","44","5.32","0","99.11","408.87","0","414.84","32.85","0"
"AGAW1","11N","INS","BOLD:ABA6514","Arthropoda","Insecta","Diptera","Chironomidae","Chironominae","Rheotanytarsus","","92","86","86","45","6.52","0","98.94","423.62","0","425.42","36.1","0"
"AGAW1","11N","INS","BOLD:AAN5242","Arthropoda","Insecta","Diptera","Cecidomyiidae","","","","88","88","88","46","0","0","98.71","407.06","0","410.7","28.15","0"
"AGAW1","11N","INS","BOLD:ACF4289","Arthropoda","Insecta","Coleoptera","Staphylinidae","","","","87","79","79","47","9.2","0","99.15","435.18","0","437.44","33.72","0"
"AGAW1","11N","INS","BOLD:ADG8758","Arthropoda","Insecta","Hymenoptera","Braconidae","Brachistinae","Blacus","Blacus stelfoxi","80","70","70","48","12.5","0","99.49","434.35","0","436.09","32.05","0"
"AGAW1","11N","INS","BOLD:ABU5525","Arthropoda","Insecta","Diptera","Chironomidae","Orthocladiinae","Limnophyes","Limnophyes sp. 14ES","79","79","79","49","0","0","98.97","400.84","0","403.56","37.11","0"
"AGAW1","11N","INS","BOLD:ACE3123","Arthropoda","Insecta","Diptera","Sciaridae","","Corynoptera","","79","73","73","49","7.59","0","99.38","431.11","0","432.48","34.98","0"
"AGAW1","11N","INS","BOLD:ABV1297","Arthropoda","Insecta","Diptera","Sciaridae","","","","77","72","72","51","6.49","0","99.05","427.75","0","430.39","30.27","0"
"AGAW1","11N","INS","BOLD:ACV2677","Arthropoda","Insecta","Diptera","Cecidomyiidae","","","","76","57","57","52","25","0","97.62","448","0","448.58","29.95","0"
"AGAW1","11N","INS","BOLD:AAP1118","Arthropoda","Insecta","Diptera","Mycetophilidae","Sciophilinae","Sciophila","","75","63","63","53","16","0","98.13","446.96","0","448.17","33.12","0"
"AGAW1","11N","INS","BOLD:ABA0863","Arthropoda","Insecta","Diptera","Cecidomyiidae","","","","74","71","71","54","4.05","0","99.36","440.47","0","442.34","29.77","0"
"AGAW1","11N","INS","BOLD:AAN5205","Arthropoda","Insecta","Diptera","Cecidomyiidae","","","CecidInt42 sp. BOLD:AAN5205","74","58","58","54","21.62","0","99.74","445.81","0","446.97","31.38","0"
"AGAW1","11N","INS","BOLD:AAC2499","Arthropoda","Insecta","Diptera","Drosophilidae","Drosophilinae","Drosophila","Drosophila subpulchrella, Drosophila suzukii","70","66","66","56","5.71","0","98.67","437.36","0","440.06","34.33","0"
"AGAW1","11N","INS","BOLD:AAN6433","Arthropoda","Insecta","Diptera","Sciaridae","","Corynoptera","Corynoptera deserta","70","69","69","56","1.43","0","99.12","436.86","0","439.64","30.47","0"
"AGAW1","11N","INS","BOLD:AAN5183","Arthropoda","Insecta","Diptera","Cecidomyiidae","","","","69","65","65","58","5.8","0","98.19","440.16","0","441.28","27.73","0"
"AGAW1","11N","INS","BOLD:ACL2677","Arthropoda","Insecta","Psocodea","Mesopsocidae","","Mesopsocus","Mesopsocus laticeps","66","60","60","59","9.09","0","99.46","426.83","0","428.44","41.01","0"
"AGAW1","11N","INS","BOLD:AAN6447","Arthropoda","Insecta","Diptera","Sciaridae","","Corynoptera","Corynoptera perpusilla","63","52","52","60","17.46","0","99.43","443.7","0","448.19","36.26","0"
"AGAW1","11N","INS","BOLD:AAH3664","Arthropoda","Insecta","Diptera","Cecidomyiidae","","","CecidInt3 sp. BOLD:AAH3664","61","58","58","61","4.92","0","98.92","427.97","0","430.26","27.15","0"
"AGAW1","11N","INS","BOLD:ABA0822","Arthropoda","Insecta","Diptera","Cecidomyiidae","","","","60","60","60","62","0","0","99.22","416.78","0","418.22","26.02","0"
"AGAW1","11N","INS","BOLD:AAD1485","Arthropoda","Insecta","Diptera","Chironomidae","Chironominae","Paratanytarsus","Paratanytarsus grimmii, Paratanytarsus sp.","53","51","51","63","3.77","0","99.44","435.96","0","437.66","33.77","0"
"AGAW1","11N","INS","BOLD:AAV5575","Arthropoda","Insecta","Diptera","Cecidomyiidae","","","CecidInt30 sp. BOLD:AAV5575","48","44","44","64","8.33","0","99.51","438.25","0","440.67","25.87","0"
"AGAW1","11N","INS","BOLD:ACZ1491","Arthropoda","Insecta","Diptera","Chironomidae","","","","48","41","41","64","14.58","0","97.82","447.56","0","449.29","34.33","0"
"AGAW1","11N","INS","BOLD:AAG2898","Arthropoda","Insecta","Hemiptera","Cicadellidae","Deltocephalinae","Paraphlepsius","Paraphlepsius apertus, Paraphlepsius irroratus","47","44","44","66","6.38","0","99.58","433.06","0","433.98","31.74","0"
"AGAW1","11N","INS","BOLD:ABV1337","Arthropoda","Insecta","Diptera","Cecidomyiidae","","","","42","40","40","67","4.76","0","98.91","429.71","0","434.14","28.3","0"
"AGAW1","11N","INS","BOLD:AAN5332","Arthropoda","Insecta","Diptera","Chironomidae","Chironominae","Polypedilum","","42","42","42","67","0","0","98.66","405.5","0","408.36","33.78","0"
"AGAW1","11N","INS","BOLD:ABU5546","Arthropoda","Insecta","Diptera","Psychodidae","Psychodinae","Psychoda","","36","35","35","69","2.78","0","98.13","425.61","0","427.22","31.32","0"
"AGAW1","11N","INS","BOLD:ABA6407","Arthropoda","Insecta","Diptera","Sciaridae","","Pseudolycoriella","","36","36","36","69","0","0","98.9","430.56","0","433","35.05","0"
"AGAW1","11N","INS","BOLD:ACC8079","Arthropoda","Insecta","Diptera","Cecidomyiidae","","","","35","33","33","71","5.71","0","99.16","447.54","0","449.17","29.42","0"
"AGAW1","11N","INS","BOLD:AAN5260","Arthropoda","Insecta","Diptera","Cecidomyiidae","","","","34","34","34","72","0","0","99.19","408.44","0","409.94","26.03","0"
"AGAW1","11N","INS","BOLD:ABY5318","Arthropoda","Insecta","Diptera","Chloropidae","Oscinellinae","Incertella, Rhopalopterum","Rhopalopterum umbrosum, Rhopalopterum luteiceps, Incertella bispina, Rhopalopterum painteri","34","27","27","72","20.59","0","99.34","442.62","0","443.85","32.18","0"
"AGAW1","11N","INS","BOLD:AAG1000","Arthropoda","Insecta","Diptera","Chironomidae","Orthocladiinae","Orthocladius","Orthocladius carlatus","32","32","32","74","0","0","98.17","417.25","0","419.38","33.57","0"
"AGAW1","11N","INS","TAX:487439","Arthropoda","Insecta","Coleoptera","Chrysomelidae","Eumolpinae","Paria","Paria thoracica","31","31","31","75","0","0","99.28","403.19","0","405.13","38.86","0"
"AGAW1","11N","INS","BOLD:AAG8091","Arthropoda","Insecta","Hymenoptera","Ichneumonidae","Campopleginae","Campoletis","Campoletis sp. AB.2","30","30","30","76","0","0","98.84","434.17","0","436.63","24.8","0"
"AGAW1","11N","INS","BOLD:AAG3236","Arthropoda","Insecta","Diptera","Phoridae","Phorinae","Conicera, Diplonevra","Diplonevra nitidula","29","25","25","77","13.79","0","99.3","443.9","0","446.14","31.9","0"
"AGAW1","11N","INS","BOLD:AAH0135","Arthropoda","Insecta","Coleoptera","Phalacridae","Phalacrinae","Acylomus","Acylomus pugetanus, Acylomus ergoti","29","27","27","77","6.9","0","99.04","438.62","0","440.83","31.63","0"
"AGAW1","11N","INS","BOLD:AAH0134","Arthropoda","Insecta","Coleoptera","Phalacridae","Phalacrinae","Stilbus","Stilbus apicialis","28","27","27","79","3.57","0","99.1","425.86","0","427.39","31.75","0"
"AGAW1","11N","INS","BOLD:AAN5255","Arthropoda","Insecta","Diptera","Cecidomyiidae","Cecidomyiinae","Dasineura","","26","26","26","80","0","0","98.33","427.04","0","444.77","26.76","0"
"AGAW1","11N","INS","BOLD:AAG8493","Arthropoda","Insecta","Diptera","Drosophilidae","Drosophilinae","Scaptomyza","Scaptomyza pallida","26","26","26","80","0","0","99.12","400.73","0","402.5","33.11","0"
"AGAW1","11N","INS","BOLD:AAF9306","Arthropoda","Insecta","Diptera","Psychodidae","Psychodinae","Psychoda","Psychoda alternata","24","24","24","82","0","0","97.97","438.13","0","439.5","31.34","0"
"AGAW1","11N","INS","BOLD:ADG4616","Arthropoda","Insecta","Diptera","Cecidomyiidae","","","","23","23","23","83","0","0","99.61","385.09","0","440.61","26.9","0"
"AGAW1","11N","INS","BOLD:ACX6289","Arthropoda","Insecta","Diptera","Phoridae","Metopininae","Megaselia","","23","23","23","83","0","0","99.4","390.78","0","393.3","37.77","0"
"AGAW1","11N","INS","BOLD:AAH3228","Arthropoda","Insecta","Psocodea","Caeciliusidae","Caeciliusinae","Valenzuela","Valenzuela flavidus","23","22","22","83","4.35","0","98.97","431.65","0","433.22","35.69","0"
"AGAW1","11N","INS","BOLD:ABA7871","Arthropoda","Insecta","Diptera","Cecidomyiidae","","","","22","22","22","86","0","0","99.17","445.5","0","446.41","26.53","0"
"AGAW1","11N","INS","BOLD:AAG1711","Arthropoda","Insecta","Diptera","Muscidae","Phaoniinae","Helina","Helina depuncta","20","20","20","87","0","0","98.85","436.8","0","440.25","31.37","0"
"AGAW1","11N","INS","BOLD:AAU8483","","","","","","","","18","18","18","88","0","0","98.69","427.22","0","429.22","28.01","0"
"AGAW1","11N","INS","BOLD:ABZ0621","Arthropoda","Insecta","Diptera","Psychodidae","Psychodinae","","","18","18","18","88","0","0","99.17","427.89","0","431.33","33.01","0"
"AGAW1","11N","INS","BOLD:AAA4222","Arthropoda","Insecta","Lepidoptera","Erebidae","Hypeninae","Hypena","Hypena scabra","17","17","17","90","0","0","98.99","400.12","0","403","32.41","0"
"AGAW1","11N","INS","BOLD:ACU3040","Arthropoda","Insecta","Diptera","Sciaridae","","","","17","17","17","90","0","0","98.44","431.65","0","433.65","32.11","0"
"AGAW1","11N","INS","BOLD:AAM6040","Arthropoda","Insecta","Diptera","Cecidomyiidae","","","","16","14","14","92","12.5","0","99.79","436.25","0","437.44","26.69","0"
"AGAW1","11N","INS","BOLD:AAG6745","Arthropoda","Insecta","Diptera","Calliphoridae","Calliphorinae","Pollenia","Pollenia pediculata","16","16","16","92","0","0","99.11","435.19","0","437.06","33.71","0"
"AGAW1","11N","INS","BOLD:AAM9255","Arthropoda","Insecta","Diptera","Sciaridae","","Pseudolycoriella","","16","16","16","92","0","0","99.23","398.63","0","401.19","30.63","0"
"AGAW1","11N","INS","BOLD:AAN8337","Arthropoda","Insecta","Hemiptera","Cicadellidae","Typhlocybinae","Empoasca","","15","15","15","95","0","0","98.55","406.53","0","408.4","29.87","0"
"AGAW1","11N","INS","BOLD:AAD1185","Arthropoda","Insecta","Lepidoptera","Gracillariidae","Lithocolletinae","Phyllonorycter","Phyllonorycter blancardella","14","14","14","96","0","0","99.7","448","0","450","34.37","0"
"AGAW1","11N","INS","BOLD:ADC9915","Arthropoda","Insecta","Diptera","Cecidomyiidae","","","","14","13","13","96","7.14","0","99.71","445.21","0","446.64","27.63","0"
"AGAW1","11N","INS","BOLD:AAU6513","Arthropoda","Insecta","Diptera","Sciaridae","","Corynoptera","Corynoptera bicuspidata","14","14","14","96","0","0","98.94","399.21","0","401.5","32.01","0"
"AGAW1","11N","INS","BOLD:ABA9949","Arthropoda","Insecta","Coleoptera","Chrysomelidae","Galerucinae","Chaetocnema","Chaetocnema concinna","14","11","11","96","21.43","0","99.5","439","0","440.43","36.24","0"
"AGAW1","11N","INS","BOLD:AAG6471","Arthropoda","Insecta","Diptera","Ceratopogonidae","Dasyheleinae","Dasyhelea","","14","14","14","96","0","0","99.15","409.29","0","411.5","33.91","0"
"AGAW1","11N","INS","BOLD:AAC0903","Arthropoda","Insecta","Diptera","Chironomidae","Chironominae","Chironomus","Chironomus acidophilus","13","13","13","101","0","0","99.18","441.77","0","443.46","36.37","0"
"AGAW1","11N","INS","BOLD:AAF1729","Arthropoda","Insecta","Psocodea","Lachesillidae","Lachesillinae","Lachesilla","Lachesilla pedicularia","12","10","10","102","16.67","0","99.65","449.42","0","450","37.3","0"
"AGAW1","11N","INS","BOLD:ACU7906","Arthropoda","Insecta","Diptera","Chironomidae","Orthocladiinae","","","10","10","10","103","0","0","98.58","399.5","0","403","36.14","0"
"AGAW1","11N","INS","BOLD:AAI8935","Arthropoda","Insecta","Coleoptera","Latridiidae","Corticariinae","Cortinicara","Cortinicara gibbosa","10","10","10","103","0","0","98.96","436.8","0","439.2","33.73","0"
"AGAW1","11N","INS","BOLD:AAA3181","Arthropoda","Insecta","Diptera","Muscidae","Muscinae","Stomoxys","Stomoxys calcitrans","9","9","9","105","0","0","98.78","418.33","0","422.11","32.78","0"
"AGAW1","11N","INS","BOLD:ABW2929","Arthropoda","Insecta","Hemiptera","Cicadellidae","Typhlocybinae","Empoasca","","9","9","9","105","0","0","99.54","416.89","0","418.67","30.88","0"
"AGAW1","11N","INS","BOLD:AAK5928","Arthropoda","Insecta","Coleoptera","Chrysomelidae","Criocerinae","Oulema","Oulema melanopus","9","9","9","105","0","0","99.49","429.33","0","431","36.25","0"
"AGAW1","11N","INS","BOLD:AAN5221","Arthropoda","Insecta","Diptera","Cecidomyiidae","","","","9","9","9","105","0","0","98.66","402.44","0","408.11","25.17","0"
"AGAW1","11N","INS","BOLD:AAN5220","Arthropoda","Insecta","Diptera","Cecidomyiidae","","","","9","9","9","105","0","0","99.05","435.11","0","437.67","25.54","0"
"AGAW1","11N","INS","BOLD:AAN5201","Arthropoda","Insecta","Diptera","Cecidomyiidae","","","","8","8","8","110","0","0","99.75","449.5","0","450","27.92","0"
"AGAW1","11N","INS","BOLD:AAG1405","Arthropoda","Insecta","Hymenoptera","Braconidae","Euphorinae","Meteorus","","7","7","7","111","0","0","98.26","434.14","0","436.86","29.18","0"
"AGAW1","11N","INS","BOLD:ABA6848","Arthropoda","Insecta","Diptera","Mycetophilidae","Mycetophilinae","Exechia","","7","7","7","111","0","0","98.9","430","0","432.71","36.08","0"
"AGAW1","11N","INS","BOLD:AAV5400","Arthropoda","Insecta","Diptera","Cecidomyiidae","","","","7","7","7","111","0","0","98.4","433.57","0","434.43","29.5","0"
"AGAW1","11N","INS","BOLD:AAA4555","Arthropoda","Insecta","Orthoptera","Acrididae","Melanoplinae","Melanoplus","Melanoplus stupefactus, Melanoplus angustipennis, Melanoplus ponderosus, Melanoplus bruneri, Melanoplus frigidus, Melanoplus scapularis, Melanoplus cinereus, Melanoplus madeleineae, Melanoplus sanguinipes, Melanoplus nigrescens group, Melanoplus tunicae, Melanoplus walshi, Melanoplus femurrubrum, Melanoplus dawsoni, Melanoplus sp., Melanoplus fasciatus, Melanoplus bowditchi, Melanoplus stonei, Melanoplus bivittatus, Melanoplus huroni, Melanoplus thomasi, Melanoplus borealis, Melanoplus packardii","7","7","7","111","0","0","98.95","448.29","0","450","36.7","0"
"AGAW1","11N","INS","BOLD:AAA2632","Arthropoda","Insecta","Lepidoptera","Noctuidae","Noctuinae","Noctua","Noctua atlantica, Noctua pronuba","6","6","6","115","0","0","98.06","441.33","0","442.83","31.46","0"
"AGAW1","11N","INS","BOLD:AAM9355","Arthropoda","Insecta","Diptera","Phoridae","Metopininae","Megaselia","Megaselia sp. BOLD:AAM9355","6","6","6","115","0","0","99.06","422.33","0","424.17","34.78","0"
"AGAW1","11N","INS","BOLD:AAH3227","Arthropoda","Insecta","Psocodea","Lachesillidae","Lachesillinae","Lachesilla","","6","5","5","115","16.67","0","99.52","449.83","0","450","40.7","0"
"AGAW1","11N","INS","BOLD:AAO7616","Arthropoda","Insecta","Diptera","Chironomidae","Chironominae","Polypedilum","","6","6","6","115","0","0","98.9","408.33","0","411.17","34.45","0"
"AGAW1","11N","INS","BOLD:AAL7442","Arthropoda","Insecta","Diptera","Cecidomyiidae","","","","5","5","5","119","0","0","98.89","448.8","0","450","26.49","0"
"AGAW1","11N","INS","BOLD:AAA2518","Arthropoda","Insecta","Lepidoptera","Crambidae","Crambinae","Chrysoteuchia","Chrysoteuchia topiaria, Chrysoteuchia topiarius","4","4","4","120","0","0","99.24","431.75","0","433.75","35.42","0"
"AGAW1","11N","INS","BOLD:ACJ8771","Arthropoda","Insecta","Diptera","Chironomidae","Chironominae","Sublettea","","4","4","4","120","0","0","99.09","373","0","430.5","32.15","0"
"AGAW1","11N","INS","BOLD:ACU6062","Arthropoda","Insecta","Diptera","Cecidomyiidae","","","","4","4","4","120","0","0","99.05","448.25","0","450","26.67","0"
"AGAW1","11N","INS","BOLD:ACB4103","Arthropoda","Insecta","Diptera","Cecidomyiidae","","","","4","4","4","120","0","0","98.92","426.25","0","425.5","26.04","0"
"AGAW1","11N","INS","BOLD:AAH2293","Arthropoda","Insecta","Diptera","Asilidae","Stenopogoninae","Dioctria, Eudioctria","Dioctria hyalipennis, Dioctria baumhaueri, Eudioctria propinqua","4","4","4","120","0","0","98.81","426","0","427.5","30.81","0"
"AGAW1","11N","INS","BOLD:ACE9865","Arthropoda","Insecta","Lepidoptera","Noctuidae","Noctuinae","Feltia","Feltia subgothica, Feltia herilis, Feltia tricosa","4","4","4","120","0","0","100","423.5","0","425","30.61","0"
"AGAW1","11N","INS","BOLD:ADH9867","Arthropoda","Insecta","Diptera","","","","","4","4","4","120","0","0","98.55","389.5","0","393","37.32","0"
"AGAW1","11N","INS","BOLD:AAV1136","Arthropoda","Insecta","Diptera","Scatopsidae","","","","4","4","4","120","0","0","98.6","408.75","0","409.75","36.67","0"
"AGAW1","11N","INS","BOLD:AAG2740","Arthropoda","Insecta","Diptera","Ephydridae","Parydrinae","Philygria","Philygria debilis","4","4","4","120","0","0","98.28","393.25","0","395.5","32.88","0"
"AGAW1","11N","INS","BOLD:AAG6743","Arthropoda","Insecta","Diptera","Sarcophagidae","Sarcophaginae","Sarcophaga","Sarcophaga sp. 1 MG, Sarcophaga subvicina","3","3","3","129","0","0","99.7","446.33","0","447.33","35.39","0"
"AGAW1","11N","INS","BOLD:ABY5257","Arthropoda","Insecta","Lepidoptera","Noctuidae","Noctuinae","Apamea","Apamea devastator","3","3","3","129","0","0","99.47","440","0","440","30.37","0"
"AGAW1","11N","INS","BOLD:AAH2306","Arthropoda","Insecta","Diptera","Asilidae","Asilinae","Efferia","Efferia aestuans","3","3","3","129","0","0","99.26","449.67","0","450","37.93","0"
"AGAW1","11N","INS","BOLD:ACF8174","Arthropoda","Insecta","Diptera","Chironomidae","Chironominae","","","3","3","3","129","0","0","98.21","446.33","0","450","32.89","0"
"AGAW1","11N","INS","BOLD:ABX6186","Arthropoda","Insecta","Diptera","Tipulidae","Tipulinae","Nephrotoma","Nephrotoma ferruginea","3","3","3","129","0","0","99.55","448.33","0","450","34.44","0"
"AGAW1","11N","INS","BOLD:AAP6465","Arthropoda","Insecta","Diptera","Sciaridae","","Corynoptera","Corynoptera cursor","3","3","3","129","0","0","98.81","449.33","0","450","32.52","0"
"AGAW1","11N","INS","BOLD:ABV1237","Arthropoda","Insecta","Diptera","Chironomidae","Chironominae","Chironomus","","2","2","2","135","0","0","98.43","448","0","450","35.11","0"
"AGAW1","11N","INS","BOLD:AAN8030","Arthropoda","Insecta","Hymenoptera","Platygastridae","Scelioninae","Telenomus","","2","2","2","135","0","0","99.21","393.5","0","409.5","25.69","0"
"AGAW1","11N","INS","BOLD:AAA3836","Arthropoda","Insecta","Lepidoptera","Noctuidae","Plusiinae","Autographa","Autographa precationis","2","2","2","135","0","0","98.66","449","0","450","31.78","0"
"AGAW1","11N","INS","BOLD:ADH2911","Arthropoda","Insecta","Hymenoptera","Braconidae","","","","2","2","2","135","0","0","98.32","447.5","0","450","32","0"
"AGAW1","11N","INS","BOLD:AAB4657","Arthropoda","Insecta","Diptera","Chironomidae","Chironominae","Chironomus","Chironomus maturus","2","2","2","135","0","0","99.77","439.5","0","441","37.98","0"
"AGAW1","11N","INS","BOLD:AAG5453","Arthropoda","Insecta","Diptera","Chironomidae","Chironominae","Chironomus","Chironomus bifurcatus","2","2","2","135","0","0","97.66","449.5","0","450","35.33","0"
"AGAW1","11N","INS","BOLD:ACA9170","Arthropoda","Insecta","Hymenoptera","Braconidae","Alysiinae","","","2","2","2","135","0","0","99.89","436.5","0","439","26.18","0"
"AGAW1","11N","INS","BOLD:AAL7819","Arthropoda","Insecta","Diptera","Psychodidae","Psychodinae","Psychoda","Psychoda sp., Psychoda sp. BOLD:AAL7819","2","2","2","135","0","0","98.62","420.5","0","421","33.55","0"
"AGAW1","11N","INS","BOLD:AAB0373","Arthropoda","Insecta","Neuroptera","Chrysopidae","Chrysopinae","Chrysoperla","Chrysoperla sp. isolate 82, Chrysoperla zastrowi, Chrysoperla plorabunda, Chrysoperla sp. isolate 71, Chrysoperla agilis, Italochrysa italica, Chrysoperla sp. isolate 88, Chrysoperla lucasina, Chrysoperla carnea, Chrysoperla mediterranea, Chrysoperla pallida, Chrysoperla nipponensis, Chrysoperla sillemi","2","2","2","135","0","0","99.08","414","0","416.5","31.09","0"
"AGAW1","11N","INS","BOLD:AAH2295","Arthropoda","Insecta","Diptera","Asilidae","Leptogastrinae","Leptogaster","Leptogaster flavipes","2","2","2","135","0","0","99.15","413","0","450","35.11","0"
"AGAW1","11N","INS","BOLD:AAM6295","Arthropoda","Insecta","Diptera","Chaoboridae","","","","2","2","2","135","0","0","98.55","449.5","0","450","36.67","0"
"AGAW1","11N","INS","BOLD:AAA4277","Arthropoda","Insecta","Diptera","Syrphidae","Syrphinae","Toxomerus","Toxomerus marginatus, Toxomerus sp. CR-3, Toxomerus CR-3","2","2","2","135","0","0","97.55","449.5","0","450","31.22","0"
"AGAW1","11N","INS","BOLD:AAA4188","Arthropoda","Insecta","Hymenoptera","Braconidae","Aphidiinae","Aphidius","Aphidius microlophii, Aphidius ervi","2","2","2","135","0","0","99.55","439.5","0","443","26.3","0"
"AGAW1","11N","INS","BOLD:AAC9062","Arthropoda","Insecta","Diptera","Culicidae","Culicinae","Ochlerotatus, Aedes","Aedes fitchii, Ochlerotatus fitchii, Aedes grossbecki, Aedes ventrovittis","2","2","2","135","0","0","99.88","405","0","406.5","32.85","0"
"AGAW1","11N","INS","BOLD:AAA1517","Arthropoda","Insecta","Lepidoptera","Tortricidae","Tortricinae","Choristoneura","Choristoneura parallela, Choristoneura rosaceana, Choristoneura zapulata, Choristoneura obsoletana","2","2","2","135","0","0","98.88","448.5","0","450","30.56","0"
"AGAW1","11N","INS","BOLD:AAA6759","Arthropoda","Insecta","Lepidoptera","Hesperiidae","Hesperiinae","Thymelicus","Thymelicus lineola","1","1","1","150","0","0","99.73","372","0","374","29.14","0"
"AGAW1","11N","INS","BOLD:AAP7026","Arthropoda","Insecta","Coleoptera","Latridiidae","Corticariinae","Melanophthalma","Melanophthalma villosa","1","1","1","150","0","0","100","409","0","450","32","0"
"AGAW1","11N","INS","BOLD:AAG1713","Arthropoda","Insecta","Diptera","Muscidae","Mydaeinae","Gymnodia","Gymnodia humilis","1","1","1","150","0","0","97.78","450","0","450","31.33","0"
"AGAW1","11N","INS","BOLD:AAG7279","Arthropoda","Insecta","Diptera","Sphaeroceridae","Limosininae","Spelobia","Spelobia ochripes","1","1","1","150","0","0","98.99","398","0","403","33.75","0"
"AGAW1","11N","INS","BOLD:AAF9028","Arthropoda","Insecta","Diptera","Limoniidae","","","","1","1","1","150","0","0","99.33","446","0","450","38.44","0"
"AGAW1","11N","INS","BOLD:AAG7276","Arthropoda","Insecta","Diptera","Sphaeroceridae","Limosininae","Leptocera","Leptocera erythrocera","1","1","1","150","0","0","97.76","446","0","448","32.81","0"
"AGAW1","11N","INS","BOLD:AAA6560","Arthropoda","Insecta","Lepidoptera","Crambidae","Crambinae","Crambus","Crambus multilinellus, Crambus saltuellus","1","1","1","150","0","0","100","450","0","450","30.89","0"
"AGAW1","11N","INS","BOLD:AAY6730","Arthropoda","Insecta","Hemiptera","Derbidae","Cedusinae","Cedusa","Cedusa incisa","1","1","1","150","0","0","99.49","392","0","394","35.03","0"
"AGAW1","11N","INS","BOLD:AAI4367","Arthropoda","Insecta","Lepidoptera","Heliozelidae","","Antispila","Antispila freemani, Antispila oinophylla, Antispila ampelopsifoliella","1","1","1","150","0","0","99.78","449","0","450","30.89","0"
"AGAW1","11N","INS","BOLD:AAL4232","Arthropoda","Insecta","Diptera","Agromyzidae","Phytomyzinae","Phytoliriomyza","Phytoliriomyza arctica","1","1","1","150","0","0","99.3","426","0","426","31.69","0"
"AGAW1","11N","INS","BOLD:AAA8933","Arthropoda","Insecta","Coleoptera","Coccinellidae","Coccinellinae","Coccinella","Coccinella septempunctata","1","1","1","150","0","0","98.17","438","0","442","33.03","0"
"AGAW1","11N","INS","BOLD:AAB7869","Arthropoda","Insecta","Lepidoptera","Tortricidae","Tortricinae","Phtheochroa","Phtheochroa baracana, Phtheochroa riscana, Phtheochroa birdana, Phtheochroa fulviplicana, Phtheochroa canariana","1","1","1","150","0","0","100","450","0","450","29.78","0"
"AGAW1","11N","INS","BOLD:AAM6781","Arthropoda","Insecta","Diptera","Dolichopodidae","Neurigoninae","Neurigona","Neurigona tenuis","1","1","1","150","0","0","97.86","373","0","378","34.92","0"
"AGAW1","11N","INS","BOLD:AAA3867","Arthropoda","Insecta","Lepidoptera","Noctuidae","Noctuinae","Apamea","Apamea amputatrix","1","1","1","150","0","0","99.56","450","0","450","34","0"
"AGAW1","11N","INS","BOLD:ABA0755","Arthropoda","Insecta","Diptera","Drosophilidae","","","","1","1","1","150","0","0","99.11","448","0","450","31.56","0"
"AGAW1","11N","INS","BOLD:AAC9132","Arthropoda","Insecta","Diptera","Culicidae","Culicinae","Culiseta","Culiseta inornata","1","1","1","150","0","0","97.57","371","0","375","34.4","0"
"AGAW1","11N","INS","BOLD:AAI1549","Arthropoda","Insecta","Hymenoptera","Braconidae","Euphorinae","Meteorus","","1","1","1","150","0","0","98.66","372","0","373","31.37","0"
"AGAW1","11N","INS","BOLD:AAG3273","Arthropoda","Insecta","Diptera","Phoridae","Metopininae","Megaselia","Megaselia albicaudata","1","1","1","150","0","0","99.78","450","0","450","32.89","0"
"AGAW1","11N","INS","BOLD:ADF2969","Arthropoda","Insecta","Diptera","Sarcophagidae","Sarcophaginae","Blaesoxipha","Blaesoxipha falciformis","1","1","1","150","0","0","99.32","439","0","440","38.41","0"
"AGAW1","11N","INS","BOLD:AAB5962","Arthropoda","Insecta","Lepidoptera","Tortricidae","Tortricinae","Phalonidia","Phalonidia lepidana","1","1","1","150","0","0","97.27","439","0","441","33.56","0"
"AGAW1","11N","INS","BOLD:AAG6929","Arthropoda","Insecta","Diptera","Dolichopodidae","Medeterinae","Medetera","Medetera vittata","1","1","1","150","0","0","98.67","450","0","450","41.78","0"
"AGAW1","11N","INS","BOLD:ABX5883","Arthropoda","Insecta","Lepidoptera","Tortricidae","Tortricinae","Choristoneura","Choristoneura retiniana, Choristoneura lambertiana, Choristoneura biennis, Choristoneura orae, Choristoneura fumiferana, Choristoneura occidentalis, Choristoneura freemani, Choristoneura pinus, Choristoneura occidentalis group, Choristoneura sp., Choristoneura carnana","1","1","1","150","0","0","98.22","449","0","450","31.78","0"
"AGAW1","11N","INS","BOLD:AAG7891","Arthropoda","Insecta","Hymenoptera","Platygastridae","Scelioninae","Telenomus","Telenomus podisi","1","1","1","150","0","0","99.55","440","0","450","27.33","0"
"AGAW1","11N","INS","BOLD:AAG6433","Arthropoda","Insecta","Diptera","Ceratopogonidae","Forcipomyiinae","Forcipomyia","Forcipomyia sp. BOLD:AAG6433, Forcipomyia sp. 17","1","1","1","150","0","0","98","450","0","450","32.44","0"
"AGAW1","11N","INS","BOLD:AAM9105","Arthropoda","Insecta","Diptera","Muscidae","Coenosiinae","Coenosia","","1","1","1","150","0","0","99.11","449","0","450","36.89","0"
"AGAW1","11N","INS","BOLD:ABA6362","Arthropoda","Insecta","Coleoptera","Staphylinidae","Aleocharinae","Amischa","Amischa decipiens","1","1","1","150","0","0","99.78","447","0","449","34.97","0"
"AGAW1","11N","INS","BOLD:ACB1322","Arthropoda","Insecta","Coleoptera","Mordellidae","Mordellinae","Mordellistena","Mordellistena trifasciata","1","1","1","150","0","0","99.5","404","0","450","41.56","0"
"AGAW1","11N","INS","BOLD:AAH3225","Arthropoda","Insecta","Psocodea","Psocidae","Psocinae","Metylophorus","Metylophorus novaescotiae","1","1","1","150","0","0","98.66","449","0","450","35.11","0"
"AGAW1","11N","INS","BOLD:ACE6286","Arthropoda","Insecta","Diptera","Culicidae","Culicinae","Ochlerotatus, Aedes","Aedes dorsalis, Ochlerotatus dorsalis","1","1","1","150","0","0","97.78","450","0","450","32.44","0"
"AGAW1","11N","INS","BOLD:AAZ5483","Arthropoda","Insecta","Diptera","Muscidae","Phaoniinae","Helina","Helina setiventris","1","1","1","150","0","0","99.55","448","0","450","29.33","0"
"AGAW1","11N","INS","BOLD:AAA6822","Arthropoda","Insecta","Orthoptera","Acrididae","Gomphocerinae","Chorthippus","Chorthippus curtipennis","1","1","1","150","0","0","99.78","448","0","450","35.11","0"
"AGAW1","11N","INS","BOLD:ACG5862","Arthropoda","Insecta","Diptera","Agromyzidae","","","","1","1","1","150","0","0","99.11","450","0","450","34.67","0"
"AGAW1","11N","INS","BOLD:ACE7444","Arthropoda","Insecta","Hemiptera","Miridae","Mirinae, Phylinae","Adelphocoris, Megalocoleus","Adelphocoris lineolatus, Megalocoleus molliculus","1","1","1","150","0","0","99.56","450","0","450","35.33","0"
"AGAW1","11N","INS","BOLD:AAF5980","Arthropoda","Insecta","Hemiptera","Cicadellidae","Typhlocybinae","Typhlocyba","Typhlocyba pomaria","1","1","1","150","0","0","98","449","0","450","37.56","0"
"AGAW1","11N","INS","BOLD:AAA5206","Arthropoda","Insecta","Lepidoptera","Erebidae","Herminiinae","Zanclognatha","Zanclognatha cruralis, Zanclognatha sp. 1, Zanclognatha jacchusalis, Zanclognatha minoralis, Zanclognatha minoralis sp. 1, Zanclognatha sp., Zanclognatha minoralis sp. 3, Zanclognatha gypsalis, Zanclognatha dentata, Zanclognatha theralis, Zanclognatha protumnusalis, Zanclognatha martha, Zanclognatha atrilineella, Zanclognatha obscuripennis, Zanclognatha minoralis sp. 2, Zanclognatha marcidilinea","1","1","1","150","0","0","98.37","368","0","369","31.71","0"
"AGAW1","11N","INS","BOLD:AAM6291","Arthropoda","Insecta","Diptera","Chironomidae","Chironominae","Chironomus","","1","1","1","150","0","0","98.43","447","0","450","36.44","0"
"AGAW1","11N","INS","BOLD:AAA8144","Arthropoda","Insecta","Lepidoptera","Pterophoridae","Pterophorinae","Dejongia","Dejongia lobidactylus","1","1","1","150","0","0","98.36","366","0","373","29.49","0"
"AGAW1","11N","INS","BOLD:AAC0706","Arthropoda","Insecta","Diptera","Chironomidae","Chironominae","Dicrotendipes","Dicrotendipes tritomus","1","1","1","150","0","0","98.39","372","0","376","37.23","0"
"AGAW1","11N","INS","BOLD:AAP5087","Arthropoda","Insecta","Diptera","Ceratopogonidae","Dasyheleinae","Dasyhelea","","1","1","1","150","0","0","98.89","449","0","450","34.89","0"
"AGAW1","11N","INS","BOLD:AAL2750","Arthropoda","Insecta","Diptera","Muscidae","Phaoniinae","Phaonia","Phaonia tuguriorum","1","1","1","150","0","0","98.67","452","0","450","33.11","0"
"AGAW1","11N","INS","BOLD:ADH4973","Arthropoda","Insecta","Diptera","Tipulidae","Tipulinae","Nephrotoma","","1","1","1","150","0","0","97.78","450","0","450","34","0"
"AGAW1","11N","INS","BOLD:ACB0412","Arthropoda","Insecta","Diptera","Agromyzidae","Phytomyzinae","Liriomyza","","1","1","1","150","0","0","97.32","447","0","450","34.22","0"
"AGAW1","11N","INS","BOLD:AAF9187","Arthropoda","Insecta","Coleoptera","Curculionidae","Entiminae","Phyllobius","Phyllobius oblongus","1","1","1","150","0","0","99.78","449","0","450","38.67","0"
"AGAW1","11N","INS","BOLD:AAA4893","Arthropoda","Insecta","Lepidoptera","Crambidae","Crambinae","Microcrambus","Microcrambus elegans","1","1","1","150","0","0","97.63","422","0","450","33.33","0"
"AGAW1","11N","INS","BOLD:AAN5342","Arthropoda","Insecta","Diptera","Chironomidae","Orthocladiinae","Heleniella","","1","1","1","150","0","0","98.79","413","0","416","34.13","0"
"AGAW1","11N","INS","BOLD:AAF8096","Arthropoda","Insecta","Hymenoptera","Braconidae","Euphorinae","Meteorus","","1","1","1","150","0","0","97.31","409","0","440","29.77","0"
"AGAW1","11N","INS","BOLD:AAV6112","Arthropoda","Insecta","Diptera","Chloropidae","Oscinellinae","Oscinella","Oscinella sp. 2 AGT, Oscinella sp. TAW6","1","1","1","150","0","0","97.54","448","0","450","33.11","0"
"AGAW1","11N","INS","BOLD:ACX6055","Arthropoda","Insecta","Diptera","Phoridae","Metopininae","Megaselia","Megaselia sp. AK-2010-2, Megaselia sp. BOLD:ACX6055","1","1","1","150","0","0","100","450","0","450","33.33","0"
"AGAW1","11N","INS","BOLD:ACE4897","Arthropoda","Insecta","Diptera","Tachinidae","Tachininae","Microphthalma","","1","1","1","150","0","0","100","424","0","427","28.81","0"
"AGAW1","11N","INS","BOLD:ADG6553","Arthropoda","Insecta","Diptera","Chironomidae","","","","1","1","1","150","0","0","97.17","389","0","450","34.89","0"
"AGAW1","11N","INS","BOLD:AAG5499","Arthropoda","Insecta","Diptera","Chironomidae","Orthocladiinae","Psectrocladius","Psectrocladius obvius, Psectrocladius platypus","1","1","1","150","0","0","99.55","449","0","450","36","0"
"AGAW1","11N","INS","BOLD:AAG6460","Arthropoda","Insecta","Diptera","Cecidomyiidae","","","","1","1","1","150","0","0","98.17","437","0","450","35.56","0"
"AGAW1","11N","INS","BOLD:AAE0845","Arthropoda","Insecta","Hemiptera","Pentatomidae","Pentatominae","Euschistus","Euschistus servus","1","1","1","150","0","0","98.89","452","0","450","38","0"
"AGAW1","11N","INS","BOLD:ABX2489","Arthropoda","Insecta","Coleoptera","Phalacridae","Phalacrinae","Stilbus","","1","1","1","150","0","0","97.84","416","0","417","31.18","0"
"AGAW1","11N","INS","BOLD:ABV1430","Arthropoda","Insecta","Diptera","Dolichopodidae","Medeterinae","Medetera","","1","1","1","150","0","0","100","387","0","391","38.62","0"
"AGAW1","11N","INS","BOLD:AAL1297","Arthropoda","Insecta","Coleoptera","Cantharidae","Cantharinae","Rhagonycha","Rhagonycha fulva","1","1","1","150","0","0","99.77","442","0","450","33.78","0"
"AGAW1","11N","INS","BOLD:AAM6037","Arthropoda","Insecta","Diptera","Cecidomyiidae","Cecidomyiinae","Contarinia, Dasineura","","1","1","1","150","0","0","98.43","445","0","450","25.78","0"
"AGAW1","11N","INS","BOLD:AAU0577","Arthropoda","Insecta","Thysanoptera","Aeolothripidae","Aeolothripinae","Aeolothrips","Aeolothrips fasciatus","1","1","1","150","0","0","97","400","0","399","35.59","0"
"AGAW1","11N","INS","BOLD:AAE0456","Arthropoda","Insecta","Dermaptera","Forficulidae","Forficulinae","Forficula","Forficula auricularia-B, Forficula auricularia","1","1","1","150","0","0","99.11","448","0","450","45.78","0"
"AGAW1","11N","INS","BOLD:AAU8379","Arthropoda","Insecta","Hymenoptera","Braconidae","Alysiinae","Dinotrema","Dinotrema JFT01","1","1","1","150","0","0","98.43","447","0","450","24","0"
"AGAW1","11N","INS","BOLD:ACL7238","Arthropoda","Insecta","Diptera","Xylomyidae","","Solva","","1","1","1","150","0","0","97.99","448","0","450","35.56","0"
"AGAW1","11N","INS","BOLD:ABV1422","Arthropoda","Insecta","Diptera","Cecidomyiidae","","","","1","1","1","150","0","0","97.33","450","0","450","27.78","0"
"AGAW1","11N","INS","BOLD:AAB6095","Arthropoda","Insecta","Lepidoptera","Noctuidae","Noctuinae","Neoligia","Neoligia exhausta","1","1","1","150","0","0","99.17","362","0","365","32.33","0"
"AGAW1","11N","INS","BOLD:AAY6534","Arthropoda","Insecta","Coleoptera","Mordellidae","Mordellinae","Mordellina","Mordellina ancilla","1","1","1","150","0","0","100","450","0","450","38.89","0"
"AGAW1","11N","NAR","BOLD:AAL1512","Mollusca","Gastropoda","Stylommatophora","Agriolimacidae",NA,"Deroceras","Deroceras laeve","1","1","1","1","0","0","98.24","398","0","403","32.01","0"
"AGAW1","11N","NIN","BOLD:AAU6252","Arthropoda","Arachnida","Trombidiformes","Bdellidae","","Cyta","","425","360","360","1","15.29","0","98.65","434.58","0","435.85","31.49","0"
"AGAW1","11N","NIN","BOLD:AAA7242","Arthropoda","Collembola","Entomobryomorpha","Entomobryidae","Seirinae, Entomobryinae","Entomobrya, Lepidosira","Entomobrya multifasciata, Lepidosira sp.","125","125","125","2","0","0","98.97","418.1","0","421.54","39.57","0"
"AGAW1","11N","NIN","BOLD:ABW2639","Arthropoda","Arachnida","Trombidiformes","Anystidae","","","","96","87","87","3","9.38","0","99.5","446.41","0","445.85","31.41","0"
"AGAW1","11N","NIN","BOLD:AAA6383","","","","","","","","89","78","78","4","12.36","0","98.4","435","0","436.56","38.02","0"
"AGAW1","11N","NIN","BOLD:ACY2550","Arthropoda","Arachnida","Trombidiformes","Bdellidae","","Cyta","","56","56","56","5","0","0","98.67","374.21","0","383.71","31.12","0"
"AGAW1","11N","NIN","BOLD:AAH6630","Arthropoda","Collembola","Entomobryomorpha","Entomobryidae","Entomobryinae","Entomobrya","Entomobrya unostrigata","25","25","25","6","0","0","99.07","434.92","0","435.6","39.38","0"
"AGAW1","11N","NIN","BOLD:ACZ1295","Arthropoda","Arachnida","Trombidiformes","Anystidae","","","","16","16","16","7","0","0","99.52","374.88","0","382.44","31.28","0"
"AGAW1","11N","NIN","BOLD:AAB7914","Arthropoda","Collembola","Symphypleona","Bourletiellidae","","Bourletiella","","7","7","7","8","0","0","99.49","448.57","0","450","38.32","0"
"AGAW1","11N","NIN","BOLD:AAM7961","Arthropoda","Arachnida","Trombidiformes","Anystidae","","","","6","6","6","9","0","0","99.78","455.33","0","450","31.19","0"
"AGAW1","11N","NIN","BOLD:AAF9234","Arthropoda","Arachnida","Trombidiformes","Eupodidae","","","","3","3","3","10","0","0","98.88","448.67","0","450","29.85","0"
"AGAW1","11N","NIN","BOLD:ACV5609","","","","","","","","2","2","2","11","0","0","99.18","422.5","0","423","38.87","0"
"AGAW1","11N","NIN","BOLD:AAI4346","Arthropoda","Arachnida","Opiliones","Phalangiidae","","Paroligolophus, Phalangium","Paroligolophus agrestis, Phalangium opilio","2","2","2","11","0","0","98.49","383.5","0","386.5","34.29","0"
"AGAW1","11N","NIN","BOLD:AAA8820","Arthropoda","Collembola","Entomobryomorpha","Entomobryidae","Lepidocyrtinae","Lepidocyrtus","Lepidocyrtus cyaneus","2","2","2","11","0","0","99.15","355","0","357.5","42.94","0"
"AGAW1","11N","NIN","BOLD:AAA6381","Arthropoda","Arachnida","Araneae","Tetragnathidae","","Tetragnatha","Tetragnatha laboriosa","1","1","1","14","0","0","97.77","403","0","407","37.59","0"
"AGAW1","11N","NIN","BOLD:ACX8969","Arthropoda","Arachnida","Trombidiformes","Eupodidae","","","","1","1","1","14","0","0","99.77","426","0","427","27.63","0"
"AGAW1","11N","NIN","BOLD:AAB0726","Arthropoda","Arachnida","Araneae","Lycosidae","","Trochosa","Trochosa ruricola","1","1","1","14","0","0","100","409","0","412","30.1","0"
"AGAW1","11N","NIN","BOLD:AAF9237","Arthropoda","Arachnida","Trombidiformes","Erythraeidae","","Balaustium","","1","1","1","14","0","0","100","448","0","450","37.33","0"
"AGAW1","11N","NIN","BOLD:AAC6891","Arthropoda","Arachnida","Araneae","Salticidae","","Phidippus","Phidippus audax","1","1","1","14","0","0","97.01","401","0","438","31.74","0"
"AGAW1","11S","BAC","TAX:175768","Proteobacteria","Alphaproteobacteria","Rickettsiales","Rickettsiaceae",NA,"Wolbachia","Wolbachia pipientis","26","25","25","1","3.85","0","98.5","445.54","0","446.35","39.59","0"
"AGAW1","11S","BAC","TAX:91393","Proteobacteria","Alphaproteobacteria","Rickettsiales","Rickettsiaceae",NA,"Wolbachia","","25","25","25","2","0","0","99.02","393.76","0","419.48","39.22","0"
"AGAW1","11S","BAC","TAX:91389","Proteobacteria","","","",NA,"","","2","2","2","3","0","0","99.3","358.5","0","363.5","38.51","0"
"AGAW1","11S","BAC","TAX:91391","Proteobacteria","Alphaproteobacteria","Rickettsiales","",NA,"","","1","1","1","4","0","0","99.74","386","0","450","41.33","0"
"AGAW1","11S","CHO","no match","no match","no match","no match","no match","no match","no match","no match","no match","no match","no match","no match","no match","no match","no match","no match","no match","no match","no match","no match"
"AGAW1","11S","INS","BOLD:AAP8739","Arthropoda","Insecta","Diptera","Platystomatidae","Platystomatinae","Rivellia","Rivellia steyskali","33293","21263","21263","1","36.13","0","99.38","441.39","0","442.89","38.03","0"
"AGAW1","11S","INS","BOLD:AAG2017","Arthropoda","Insecta","Neuroptera","Chrysopidae","Chrysopinae","Chrysopa","Chrysopa oculata","6876","5278","5278","2","23.24","0","99.38","433.12","0","434.48","31.52","0"
"AGAW1","11S","INS","BOLD:AAG1711","Arthropoda","Insecta","Diptera","Muscidae","Phaoniinae","Helina","Helina depuncta","5602","4609","4609","3","17.73","0","99.4","433.03","0","435.41","31.34","0"
"AGAW1","11S","INS","BOLD:AAU6681","Arthropoda","Insecta","Diptera","Tipulidae","Tipulinae","Nephrotoma","Nephrotoma cornicina","4613","3433","3433","4","25.58","0","99.63","442.12","0","443.37","36.31","0"
"AGAW1","11S","INS","BOLD:AAG2123","Arthropoda","Insecta","Diptera","Tachinidae","Dexiinae","Dinera","Dinera grisescens","2518","1928","1928","5","23.43","0","99.36","432.28","0","433.95","31.84","0"
"AGAW1","11S","INS","BOLD:ADH9310","Arthropoda","Insecta","Diptera","Anthomyiidae","Anthomyiinae","Anthomyia","Anthomyia procellaris, Anthomyia pluvialis","1735","1641","1641","6","5.42","0","99.2","413.91","0","416.13","32.18","0"
"AGAW1","11S","INS","BOLD:AAM6291","Arthropoda","Insecta","Diptera","Chironomidae","Chironominae","Chironomus","","1236","981","981","7","20.63","0","99.33","439.04","0","442.03","36.44","0"
"AGAW1","11S","INS","BOLD:AAG2511","Arthropoda","Insecta","Diptera","Anthomyiidae","Anthomyiinae","Delia","Delia platura","1059","927","927","8","12.46","0","99.08","444.05","0","447.05","33.1","0"
"AGAW1","11S","INS","BOLD:AAA4555","Arthropoda","Insecta","Orthoptera","Acrididae","Melanoplinae","Melanoplus","Melanoplus stupefactus, Melanoplus angustipennis, Melanoplus ponderosus, Melanoplus bruneri, Melanoplus frigidus, Melanoplus scapularis, Melanoplus cinereus, Melanoplus madeleineae, Melanoplus sanguinipes, Melanoplus nigrescens group, Melanoplus tunicae, Melanoplus walshi, Melanoplus femurrubrum, Melanoplus dawsoni, Melanoplus sp., Melanoplus fasciatus, Melanoplus bowditchi, Melanoplus stonei, Melanoplus bivittatus, Melanoplus huroni, Melanoplus thomasi, Melanoplus borealis, Melanoplus packardii","999","779","779","9","22.02","0","99.45","447.41","0","448.57","36.33","0"
"AGAW1","11S","INS","BOLD:AAG6745","Arthropoda","Insecta","Diptera","Calliphoridae","Calliphorinae","Pollenia","Pollenia pediculata","969","925","925","10","4.54","0","99.34","403.31","0","405.13","33.53","0"
"AGAW1","11S","INS","BOLD:AAA3430","Arthropoda","Insecta","Lepidoptera","Geometridae","Larentiinae","Orthonama","Orthonama obstipata","860","744","744","11","13.49","0","98.93","441.91","0","444.81","32.78","0"
"AGAW1","11S","INS","BOLD:AAI3064","","","","","","","","802","565","565","12","29.55","0","98.79","443.64","0","444.59","33.47","0"
"AGAW1","11S","INS","BOLD:ACB0984","Arthropoda","Insecta","Psocodea","Stenopsocidae","","Graphopsocus","Graphopsocus cruciatus","791","578","578","13","26.93","0","99.16","412.18","0","447.51","37.23","0"
"AGAW1","11S","INS","BOLD:AAG5507","Arthropoda","Insecta","Diptera","Chironomidae","Chironominae","Chironomus","Chironomus atrella","735","666","666","14","9.39","0","98.91","429.98","0","436.03","36.03","0"
"AGAW1","11S","INS","BOLD:AAA5299","Arthropoda","Insecta","Diptera","Chironomidae","Orthocladiinae","Cricotopus","Cricotopus laetus, Cricotopus relucens, Cricotopus trifasciatus, Cricotopus sp., Cricotopus sp. 3ES, Cricotopus sylvestris, Cricotopus glacialis","455","420","420","15","7.69","0","99.09","428","0","431.02","34.72","0"
"AGAW1","11S","INS","BOLD:AAK8751","Arthropoda","Insecta","Diptera","Agromyzidae","Phytomyzinae","Liriomyza","Liriomyza sp. SB, Liriomyza trifoliearum","356","287","287","16","19.38","0","99.43","440.63","0","441.75","33.18","0"
"AGAW1","11S","INS","BOLD:AAH3661","Arthropoda","Insecta","Diptera","Cecidomyiidae","","","","354","329","329","17","7.06","0","99.26","427.45","0","429.42","25.36","0"
"AGAW1","11S","INS","BOLD:AAN6154","Arthropoda","Insecta","Coleoptera","Latridiidae","Corticariinae","Melanophthalma","Melanophthalma inermis","339","285","285","18","15.93","0","99.03","432.04","0","433.93","31.18","0"
"AGAW1","11S","INS","BOLD:ACE7444","Arthropoda","Insecta","Hemiptera","Miridae","Mirinae, Phylinae","Adelphocoris, Megalocoleus","Adelphocoris lineolatus, Megalocoleus molliculus","284","186","186","19","34.51","0","99.74","443.5","0","444.14","34.82","0"
"AGAW1","11S","INS","BOLD:AAC3517","Arthropoda","Insecta","Hemiptera","Delphacidae","Delphacinae","Kosswigianella, Muirodelphax, Delphacodes, Nothodelphax, Elachodelphax","Muirodelphax arvensis, Delphacodes nitens, Delphacodes lutulenta, Muirodelphax nigrinota, Kosswigianella analis, Delphacodes campestris, Muirodelphax anufrievi, Delphacodes indistincta, Muirodelphax unda, Muirodelphax nr. arvensis, Delphacodes stramineosa, Nothodelphax consimilis, Muirodelphax atrolabis, Muirodelphax atralabis, Kosswigianella lutulenta","228","189","189","20","17.11","0","99.49","440.08","0","441.18","36.94","0"
"AGAW1","11S","INS","BOLD:AAU6592","Arthropoda","Insecta","Diptera","Ceratopogonidae","Dasyheleinae","Dasyhelea","Dasyhelea arenivaga","226","221","221","21","2.21","0","99.16","396.49","0","398.94","34.52","0"
"AGAW1","11S","INS","BOLD:AAN5161","Arthropoda","Insecta","Diptera","Ceratopogonidae","Dasyheleinae","Dasyhelea","","224","220","220","22","1.79","0","99.11","402.16","0","404.4","37.38","0"
"AGAW1","11S","INS","BOLD:ABA5203","Arthropoda","Insecta","Diptera","Limoniidae","Limoniinae","Dicranomyia","Dicranomyia frontalis","215","169","169","23","21.4","0","99.49","444.83","0","445.8","32.33","0"
"AGAW1","11S","INS","BOLD:ACX5521","Arthropoda","Insecta","Diptera","Chironomidae","Orthocladiinae","","","190","178","178","24","6.32","0","99.24","417.82","0","419.89","34.62","0"
"AGAW1","11S","INS","BOLD:AAP5169","Arthropoda","Insecta","Diptera","Chloropidae","Oscinellinae","Elachiptera","Elachiptera nigriceps","179","158","158","25","11.73","0","99.46","429.46","0","430.79","31.42","0"
"AGAW1","11S","INS","BOLD:AAA7661","Arthropoda","Insecta","Diptera","Culicidae","Culicinae","Culiseta, Culex","Culiseta melanura, Culex articularis, Culex restuans, Culex acharistus","178","174","174","26","2.25","0","99.17","430.99","0","433.3","32.94","0"
"AGAW1","11S","INS","BOLD:AAN5199","Arthropoda","Insecta","Diptera","Cecidomyiidae","","","CecidInt20 sp. BOLD:AAN5199","165","138","138","27","16.36","0","99.47","435.65","0","446.6","26.74","0"
"AGAW1","11S","INS","BOLD:AAD1485","Arthropoda","Insecta","Diptera","Chironomidae","Chironominae","Paratanytarsus","Paratanytarsus grimmii, Paratanytarsus sp.","164","147","147","28","10.37","0","99.54","442.12","0","443.49","33.77","0"
"AGAW1","11S","INS","BOLD:ACV4278","Arthropoda","Insecta","Diptera","Limoniidae","Chioneinae","Erioptera","Erioptera chlorophylla","160","158","158","29","1.25","0","99.23","433.09","0","436.38","32.86","0"
"AGAW1","11S","INS","BOLD:AAA8933","Arthropoda","Insecta","Coleoptera","Coccinellidae","Coccinellinae","Coccinella","Coccinella septempunctata","152","140","140","30","7.89","0","99.34","442.16","0","443.88","32.83","0"
"AGAW1","11S","INS","BOLD:AAH1932","Arthropoda","Insecta","Hymenoptera","Ichneumonidae","Cryptinae","Dichrogaster","Dichrogaster perlae, Dichrogaster modesta","152","148","148","30","2.63","0","99.16","412","0","413.43","25.82","0"
"AGAW1","11S","INS","BOLD:AAA9667","Arthropoda","Insecta","Lepidoptera","Oecophoridae","Oecophorinae","Epicallima","Epicallima argenticinctella","146","145","145","32","0.68","0","98.74","422.16","0","424.49","32.39","0"
"AGAW1","11S","INS","BOLD:AAN5242","Arthropoda","Insecta","Diptera","Cecidomyiidae","","","","135","135","135","33","0","0","98.74","404.3","0","406.83","28.12","0"
"AGAW1","11S","INS","BOLD:AAF9028","Arthropoda","Insecta","Diptera","Limoniidae","","","","134","113","113","34","15.67","0","99.35","432.9","0","435.4","38.58","0"
"AGAW1","11S","INS","BOLD:AAJ7123","Arthropoda","Insecta","Diptera","Culicidae","Culicinae","Culiseta","Culiseta minnesotae","131","112","112","35","14.5","0","98.83","440.21","0","441.56","34.75","0"
"AGAW1","11S","INS","BOLD:ABY5318","Arthropoda","Insecta","Diptera","Chloropidae","Oscinellinae","Incertella, Rhopalopterum","Rhopalopterum umbrosum, Rhopalopterum luteiceps, Incertella bispina, Rhopalopterum painteri","129","107","107","36","17.05","0","99.52","440.35","0","441.49","32.22","0"
"AGAW1","11S","INS","BOLD:AAM9013","Arthropoda","Insecta","Diptera","Keroplatidae","Macrocerinae","Macrocera","Macrocera pusilla","122","118","118","37","3.28","0","99.29","435.66","0","437.38","30.42","0"
"AGAW1","11S","INS","BOLD:AAG6471","Arthropoda","Insecta","Diptera","Ceratopogonidae","Dasyheleinae","Dasyhelea","","121","118","118","38","2.48","0","98.99","423.06","0","425.53","34.04","0"
"AGAW1","11S","INS","BOLD:ABW7897","Arthropoda","Insecta","Diptera","Cecidomyiidae","","","","107","81","81","39","24.3","0","99.64","444.94","0","446.46","29.74","0"
"AGAW1","11S","INS","BOLD:ABY7666","Arthropoda","Insecta","Diptera","Culicidae","Culicinae","Culex","Culex territans","104","80","80","40","23.08","0","97.82","443.73","0","444.82","33.32","0"
"AGAW1","11S","INS","BOLD:ABW1379","Arthropoda","Insecta","Diptera","Chloropidae","Oscinellinae","Malloewia","Malloewia abdominalis","95","82","82","41","13.68","0","98.35","444","0","445.77","31.07","0"
"AGAW1","11S","INS","BOLD:AAN5260","Arthropoda","Insecta","Diptera","Cecidomyiidae","","","","86","81","81","42","5.81","0","99.21","408.56","0","410.09","25.93","0"
"AGAW1","11S","INS","BOLD:AAN5238","Arthropoda","Insecta","Diptera","Cecidomyiidae","","","","74","67","67","43","9.46","0","99.39","422.91","0","425.92","25.47","0"
"AGAW1","11S","INS","BOLD:AAP5087","Arthropoda","Insecta","Diptera","Ceratopogonidae","Dasyheleinae","Dasyhelea","","73","63","63","44","13.7","0","99.02","438.63","0","440.26","34.43","0"
"AGAW1","11S","INS","BOLD:ACK5742","Arthropoda","Insecta","Diptera","Cecidomyiidae","","","","70","61","61","45","12.86","0","99.51","442.67","0","443.7","29.04","0"
"AGAW1","11S","INS","BOLD:AAG2868","Arthropoda","Insecta","Hemiptera","Cicadellidae","Typhlocybinae","Empoasca","Empoasca sp., Empoasca fabae","70","53","53","45","24.29","0","99.67","443.5","0","447","32.43","0"
"AGAW1","11S","INS","BOLD:ACU7906","Arthropoda","Insecta","Diptera","Chironomidae","Orthocladiinae","","","69","69","69","47","0","0","99.08","394.58","0","409.16","36.59","0"
"AGAW1","11S","INS","BOLD:ACC8079","Arthropoda","Insecta","Diptera","Cecidomyiidae","","","","68","55","55","48","19.12","0","99.43","446.79","0","448.49","29.22","0"
"AGAW1","11S","INS","BOLD:AAO4092","Arthropoda","Insecta","Psocodea","Caeciliusidae","Caeciliusinae","Valenzuela","Valenzuela burmeisteri","68","68","68","48","0","0","99.16","416.9","0","418.96","35.09","0"
"AGAW1","11S","INS","BOLD:AAV5098","Arthropoda","Insecta","Diptera","Ceratopogonidae","Dasyheleinae","Dasyhelea","Dasyhelea turficola, Dasyhelea sp. 5ES","66","63","63","50","4.55","0","99.13","419.32","0","421.71","36.34","0"
"AGAW1","11S","INS","BOLD:AAN5205","Arthropoda","Insecta","Diptera","Cecidomyiidae","","","CecidInt42 sp. BOLD:AAN5205","66","56","56","50","15.15","0","99.53","445.15","0","446.45","31.43","0"
"AGAW1","11S","INS","BOLD:ABA0863","Arthropoda","Insecta","Diptera","Cecidomyiidae","","","","64","61","61","52","4.69","0","99.43","430.25","0","432.03","29.53","0"
"AGAW1","11S","INS","BOLD:AAN5162","Arthropoda","Insecta","Diptera","Ceratopogonidae","Dasyheleinae","Dasyhelea","","62","62","62","53","0","0","99.19","432.94","0","435.16","33.7","0"
"AGAW1","11S","INS","BOLD:ADF6661","Arthropoda","Insecta","Diptera","Cecidomyiidae","","","","61","42","42","54","31.15","0","99.33","415.8","0","443.16","30","0"
"AGAW1","11S","INS","BOLD:AAC4197","Arthropoda","Insecta","Diptera","Chironomidae","Orthocladiinae","Paraphaenocladius","Paraphaenocladius impensus","60","53","53","55","11.67","0","98.93","436.13","0","437.7","34.65","0"
"AGAW1","11S","INS","BOLD:AAN5183","Arthropoda","Insecta","Diptera","Cecidomyiidae","","","","57","53","53","56","7.02","0","98.37","434.42","0","435.91","27.63","0"
"AGAW1","11S","INS","BOLD:AAG9897","Arthropoda","Insecta","Dermaptera","Forficulidae","Forficulinae","Forficula","Forficula auricularia-A","52","52","52","57","0","0","99.37","402.46","0","415.77","45.86","0"
"AGAW1","11S","INS","BOLD:AAF9771","Arthropoda","Insecta","Diptera","Hybotidae","Tachydromiinae","Platypalpus","Platypalpus longiseta, Platypalpus trivialis, Platypalpus sp. 5, Platypalpus sp. 17, Platypalpus pallidiventris","48","39","39","58","18.75","0","99.53","444.63","0","445.33","35.27","0"
"AGAW1","11S","INS","BOLD:AAN5255","Arthropoda","Insecta","Diptera","Cecidomyiidae","Cecidomyiinae","Dasineura","","47","46","46","59","2.13","0","98.61","420.62","0","435.7","26.55","0"
"AGAW1","11S","INS","BOLD:AAM6241","Arthropoda","Insecta","Diptera","Chironomidae","","","","44","41","41","60","6.82","0","99.33","434.41","0","436.93","35.14","0"
"AGAW1","11S","INS","BOLD:AAH0134","Arthropoda","Insecta","Coleoptera","Phalacridae","Phalacrinae","Stilbus","Stilbus apicialis","43","42","42","61","2.33","0","99.14","412.88","0","414.28","31.34","0"
"AGAW1","11S","INS","BOLD:AAF1729","Arthropoda","Insecta","Psocodea","Lachesillidae","Lachesillinae","Lachesilla","Lachesilla pedicularia","41","37","37","62","9.76","0","99.63","446.61","0","448.27","37.05","0"
"AGAW1","11S","INS","BOLD:AAV5594","Arthropoda","Insecta","Diptera","Cecidomyiidae","","","","40","38","38","63","5","0","99.14","439.98","0","442.1","26.15","0"
"AGAW1","11S","INS","BOLD:AAE0707","Arthropoda","Insecta","Diptera","Chironomidae","Chironominae","Microtendipes","Microtendipes pedellus","37","32","32","64","13.51","0","97.62","436.43","0","437.86","30.98","0"
"AGAW1","11S","INS","BOLD:AAV1261","Arthropoda","Insecta","Diptera","Sciaridae","","Bradysia","Bradysia splendida","35","35","35","65","0","0","99.32","436.89","0","439.4","34.09","0"
"AGAW1","11S","INS","BOLD:AAZ9744","Arthropoda","Insecta","Hymenoptera","Braconidae","Brachistinae","Blacus","Blacus stelfoxi","35","29","29","65","17.14","0","98.38","445.43","0","446.94","32.32","0"
"AGAW1","11S","INS","BOLD:AAD2041","Arthropoda","Insecta","Diptera","Dolichopodidae","Sciapodinae","Condylostylus","Condylostylus inermis, Condylostylus nigrofemoratus, Condylostylus connectans, Condylostylus caudatus, Condylostylus flavipes","34","25","25","67","26.47","0","99.63","432.09","0","433.41","35.89","0"
"AGAW1","11S","INS","BOLD:AAH3227","Arthropoda","Insecta","Psocodea","Lachesillidae","Lachesillinae","Lachesilla","","33","32","32","68","3.03","0","99.32","438.48","0","439.73","40.24","0"
"AGAW1","11S","INS","BOLD:ACJ0616","","","","","","","","31","31","31","69","0","0","97.67","445.48","0","446.29","33.75","0"
"AGAW1","11S","INS","BOLD:AAH0135","Arthropoda","Insecta","Coleoptera","Phalacridae","Phalacrinae","Acylomus","Acylomus pugetanus, Acylomus ergoti","30","30","30","70","0","0","98.92","433.1","0","435.73","31.72","0"
"AGAW1","11S","INS","BOLD:AAM9255","Arthropoda","Insecta","Diptera","Sciaridae","","Pseudolycoriella","","30","30","30","70","0","0","99.16","406.23","0","408.97","30.74","0"
"AGAW1","11S","INS","BOLD:ACZ1491","Arthropoda","Insecta","Diptera","Chironomidae","","","","28","28","28","72","0","0","97.86","447.93","0","450","34.35","0"
"AGAW1","11S","INS","BOLD:AAV1295","Arthropoda","Insecta","Diptera","Sciaridae","","Bradysia","Bradysia difformis, Bradysia impatiens","28","25","25","72","10.71","0","99.3","418.75","0","446.68","31.9","0"
"AGAW1","11S","INS","BOLD:AAN6447","Arthropoda","Insecta","Diptera","Sciaridae","","Corynoptera","Corynoptera perpusilla","26","24","24","74","7.69","0","99.46","448.69","0","450","36.44","0"
"AGAW1","11S","INS","BOLD:AAH3910","Arthropoda","Insecta","Diptera","Sciaridae","","Bradysia","Bradysia fungicola","25","25","25","75","0","0","98.78","421.04","0","423.76","29.82","0"
"AGAW1","11S","INS","BOLD:ACL8053","Arthropoda","Insecta","Diptera","Chironomidae","Chironominae","Tanytarsus","","25","25","25","75","0","0","99.26","443.44","0","444.56","33.68","0"
"AGAW1","11S","INS","TAX:487439","Arthropoda","Insecta","Coleoptera","Chrysomelidae","Eumolpinae","Paria","Paria thoracica","25","24","24","75","4","0","99.32","408.72","0","410.24","38.81","0"
"AGAW1","11S","INS","BOLD:AAD7604","Arthropoda","Insecta","Coleoptera","Coccinellidae","Coccinellinae","Coleomegilla","Coleomegilla maculata","23","22","22","78","4.35","0","99.24","432.91","0","435.48","31.73","0"
"AGAW1","11S","INS","BOLD:ACN7725","Arthropoda","Insecta","Hymenoptera","Platygastridae","","","","22","22","22","79","0","0","99.43","388.95","0","434","26.76","0"
"AGAW1","11S","INS","BOLD:ACE6783","","","","","","","","20","20","20","80","0","0","99.25","417.65","0","419.15","25.97","0"
"AGAW1","11S","INS","BOLD:ABX4079","Arthropoda","Insecta","Diptera","Cecidomyiidae","","","","18","11","11","81","38.89","0","98.72","443.56","0","444.78","28.3","0"
"AGAW1","11S","INS","BOLD:AAN5220","Arthropoda","Insecta","Diptera","Cecidomyiidae","","","","17","17","17","82","0","0","98.49","421.88","0","427.24","25.96","0"
"AGAW1","11S","INS","BOLD:AAF9014","Arthropoda","Insecta","Diptera","Limoniidae","Chioneinae","Symplecta","Symplecta cana, Symplecta sp. BOLDAAF9014","16","16","16","83","0","0","99.36","428.63","0","435.25","33.8","0"
"AGAW1","11S","INS","BOLD:ACF2457","Arthropoda","Insecta","Diptera","Chironomidae","Chironominae","Paratanytarsus","Paratanytarsus laccophilus","14","13","13","84","7.14","0","99.14","447.43","0","450","32.71","0"
"AGAW1","11S","INS","BOLD:AAP2970","Arthropoda","Insecta","Diptera","Anthomyiidae","Anthomyiinae","Anthomyia","Anthomyia procellaris, Anthomyia pluvialis","13","13","13","85","0","0","99.19","313.92","0","385.54","32.68","0"
"AGAW1","11S","INS","BOLD:AAG2740","Arthropoda","Insecta","Diptera","Ephydridae","Parydrinae","Philygria","Philygria debilis","13","13","13","85","0","0","98.01","421.38","0","423.62","33.42","0"
"AGAW1","11S","INS","BOLD:ABA1215","Arthropoda","Insecta","Diptera","Sciaridae","","Lycoriella","Lycoriella sativae","12","12","12","87","0","0","99.55","447.75","0","449","35.97","0"
"AGAW1","11S","INS","BOLD:ABA6407","Arthropoda","Insecta","Diptera","Sciaridae","","Pseudolycoriella","","12","12","12","87","0","0","98.98","426","0","428.58","34.94","0"
"AGAW1","11S","INS","BOLD:AAY6450","Arthropoda","Insecta","Diptera","Cecidomyiidae","","","","12","9","9","87","25","0","99.35","438.5","0","442.17","27.48","0"
"AGAW1","11S","INS","BOLD:ADG8758","Arthropoda","Insecta","Hymenoptera","Braconidae","Brachistinae","Blacus","Blacus stelfoxi","11","11","11","90","0","0","99.28","419.64","0","421.27","31.9","0"
"AGAW1","11S","INS","BOLD:AAC0903","Arthropoda","Insecta","Diptera","Chironomidae","Chironominae","Chironomus","Chironomus acidophilus","11","11","11","90","0","0","99.04","446.91","0","448.18","36.39","0"
"AGAW1","11S","INS","BOLD:AAP4636","Arthropoda","Insecta","Psocodea","Caeciliusidae","","","","10","10","10","92","0","0","99.59","363.3","0","408.5","36.17","0"
"AGAW1","11S","INS","BOLD:AAG3154","Arthropoda","Insecta","Diptera","Psychodidae","Psychodinae","Psychoda","","10","10","10","92","0","0","97.74","439.1","0","440.7","32.43","0"
"AGAW1","11S","INS","BOLD:ABU5525","Arthropoda","Insecta","Diptera","Chironomidae","Orthocladiinae","Limnophyes","Limnophyes sp. 14ES","10","10","10","92","0","0","98.71","404.7","0","406.5","37.13","0"
"AGAW1","11S","INS","BOLD:AAA3453","Arthropoda","Insecta","Diptera","Anthomyiidae","Anthomyiinae","Delia","","9","9","9","95","0","0","99.48","428.56","0","434.89","34.61","0"
"AGAW1","11S","INS","BOLD:ACL7600","Arthropoda","Insecta","Diptera","Cecidomyiidae","Cecidomyiinae","","","8","8","8","96","0","0","99.63","439.88","0","442.38","27.06","0"
"AGAW1","11S","INS","BOLD:AAA4222","Arthropoda","Insecta","Lepidoptera","Erebidae","Hypeninae","Hypena","Hypena scabra","8","8","8","96","0","0","98.93","415","0","417.5","32.41","0"
"AGAW1","11S","INS","BOLD:ABV9081","Arthropoda","Insecta","Diptera","Cecidomyiidae","","","","6","6","6","98","0","0","99.26","412.33","0","415.33","30.96","0"
"AGAW1","11S","INS","BOLD:AAG7891","Arthropoda","Insecta","Hymenoptera","Platygastridae","Scelioninae","Telenomus","Telenomus podisi","6","6","6","98","0","0","98.31","418","0","425.67","26.21","0"
"AGAW1","11S","INS","BOLD:AAB6943","Arthropoda","Insecta","Diptera","Culicidae","Culicinae","Culex","Culex territans","6","6","6","98","0","0","97.22","415.33","0","417.83","33.14","0"
"AGAW1","11S","INS","BOLD:AAG8091","Arthropoda","Insecta","Hymenoptera","Ichneumonidae","Campopleginae","Campoletis","Campoletis sp. AB.2","6","6","6","98","0","0","99.18","446.5","0","448","24.93","0"
"AGAW1","11S","INS","BOLD:AAG6743","Arthropoda","Insecta","Diptera","Sarcophagidae","Sarcophaginae","Sarcophaga","Sarcophaga sp. 1 MG, Sarcophaga subvicina","6","6","6","98","0","0","98.87","432.17","0","435.5","34.89","0"
"AGAW1","11S","INS","BOLD:AAV5796","Arthropoda","Insecta","Diptera","Cecidomyiidae","","","","6","6","6","98","0","0","99.78","434","0","435.17","29.12","0"
"AGAW1","11S","INS","BOLD:AAM6037","Arthropoda","Insecta","Diptera","Cecidomyiidae","Cecidomyiinae","Contarinia, Dasineura","","6","6","6","98","0","0","98.66","427","0","429.5","25.57","0"
"AGAW1","11S","INS","BOLD:ABV1375","Arthropoda","Insecta","Diptera","Cecidomyiidae","","","CecidInt3 sp. BOLD:ABV1375","5","5","5","105","0","0","98.69","424.4","0","425.6","26.33","0"
"AGAW1","11S","INS","BOLD:ACK3431","Arthropoda","Insecta","Diptera","Cecidomyiidae","","","","5","5","5","105","0","0","99.76","428.2","0","430.8","32.77","0"
"AGAW1","11S","INS","BOLD:ACV2677","Arthropoda","Insecta","Diptera","Cecidomyiidae","","","","5","5","5","105","0","0","97.51","449.4","0","450","30.31","0"
"AGAW1","11S","INS","BOLD:AAG8882","Arthropoda","Insecta","Hemiptera","Pentatomidae","Pentatominae","Thyanta","Thyanta pallidovirens, Thyanta custator, Thyanta accerra","5","5","5","105","0","0","99.33","437.4","0","437.8","35.98","0"
"AGAW1","11S","INS","BOLD:AAH3664","Arthropoda","Insecta","Diptera","Cecidomyiidae","","","CecidInt3 sp. BOLD:AAH3664","5","5","5","105","0","0","98.61","446.2","0","447.8","27.24","0"
"AGAW1","11S","INS","BOLD:ABA1661","Arthropoda","Insecta","Coleoptera","Chrysomelidae","Galerucinae","Longitarsus","Longitarsus pellucidus","5","5","5","105","0","0","99.55","440.8","0","441.8","35.65","0"
"AGAW1","11S","INS","BOLD:AAH0256","Arthropoda","Insecta","Coleoptera","Latridiidae","Corticariinae","Corticaria, Corticarina","","5","5","5","105","0","0","98.6","426.8","0","435","31.86","0"
"AGAW1","11S","INS","BOLD:ABW8023","Arthropoda","Insecta","Diptera","Cecidomyiidae","","","","5","5","5","105","0","0","97.64","449.2","0","450","27.42","0"
"AGAW1","11S","INS","BOLD:AAZ0276","Arthropoda","Insecta","Diptera","Cecidomyiidae","","","","5","5","5","105","0","0","99.24","448.6","0","450","26.67","0"
"AGAW1","11S","INS","BOLD:AAN5244","Arthropoda","Insecta","Diptera","Cecidomyiidae","","","","4","4","4","114","0","0","99.09","440.25","0","441.75","25","0"
"AGAW1","11S","INS","BOLD:AAN5230","Arthropoda","Insecta","Diptera","Cecidomyiidae","","","","4","4","4","114","0","0","99.11","448.25","0","450","26.72","0"
"AGAW1","11S","INS","BOLD:AAN5334","Arthropoda","Insecta","Diptera","Chironomidae","Chironominae","Polypedilum","","4","4","4","114","0","0","98.06","425","0","427.75","34.11","0"
"AGAW1","11S","INS","BOLD:AAK5928","Arthropoda","Insecta","Coleoptera","Chrysomelidae","Criocerinae","Oulema","Oulema melanopus","4","4","4","114","0","0","99.44","448.5","0","450","36.83","0"
"AGAW1","11S","INS","BOLD:ACM1709","Arthropoda","Insecta","Diptera","Chironomidae","Tanypodinae","","","4","4","4","114","0","0","98.67","436.5","0","438.75","34.49","0"
"AGAW1","11S","INS","BOLD:AAM6184","Arthropoda","Insecta","Diptera","Ceratopogonidae","Ceratopogoninae","Culicoides","","4","4","4","114","0","0","97.61","408.75","0","410.5","37.27","0"
"AGAW1","11S","INS","BOLD:AAP4633","Arthropoda","Insecta","Psocodea","Psocidae","Psocinae","Loensia","","3","3","3","120","0","0","99.19","450","0","450","36.89","0"
"AGAW1","11S","INS","BOLD:AAY6396","Arthropoda","Insecta","Diptera","Cecidomyiidae","","","","3","3","3","120","0","0","97.54","422","0","422.33","24.68","0"
"AGAW1","11S","INS","BOLD:AAM6040","Arthropoda","Insecta","Diptera","Cecidomyiidae","","","","3","2","2","120","33.33","0","99.25","448","0","448","27.16","0"
"AGAW1","11S","INS","BOLD:AAV5561","Arthropoda","Insecta","Diptera","Cecidomyiidae","","","","3","3","3","120","0","0","99.25","446.33","0","450","27.26","0"
"AGAW1","11S","INS","BOLD:AAF9306","Arthropoda","Insecta","Diptera","Psychodidae","Psychodinae","Psychoda","Psychoda alternata","3","3","3","120","0","0","98.29","449","0","450","31.41","0"
"AGAW1","11S","INS","BOLD:ABV9336","Arthropoda","Insecta","Diptera","Cecidomyiidae","","","","3","3","3","120","0","0","98.94","418.67","0","422.33","25.4","0"
"AGAW1","11S","INS","BOLD:AAA3181","Arthropoda","Insecta","Diptera","Muscidae","Muscinae","Stomoxys","Stomoxys calcitrans","2","2","2","126","0","0","99.11","447","0","450","33.22","0"
"AGAW1","11S","INS","BOLD:ACE7452","Arthropoda","Insecta","Diptera","Chironomidae","Chironominae","Paratanytarsus","Paratanytarsus sp.","2","2","2","126","0","0","98.22","449","0","450","35","0"
"AGAW1","11S","INS","BOLD:AAF3338","Arthropoda","Insecta","Lepidoptera","Noctuidae","Noctuinae","Striacosta","Striacosta albicosta","2","2","2","126","0","0","98.66","447","0","450","34","0"
"AGAW1","11S","INS","BOLD:ACN1248","Arthropoda","Insecta","Diptera","Agromyzidae","Phytomyzinae","Nemorimyza","","2","1","1","126","50","0","97.11","450","0","450","34.22","0"
"AGAW1","11S","INS","BOLD:AAL4232","Arthropoda","Insecta","Diptera","Agromyzidae","Phytomyzinae","Phytoliriomyza","Phytoliriomyza arctica","2","2","2","126","0","0","98.5","408.5","0","409","31.67","0"
"AGAW1","11S","INS","BOLD:AAB4657","Arthropoda","Insecta","Diptera","Chironomidae","Chironominae","Chironomus","Chironomus maturus","2","2","2","126","0","0","99.07","436","0","438","36.63","0"
"AGAW1","11S","INS","BOLD:ACE8878","Arthropoda","Insecta","Orthoptera","Acrididae","Melanoplinae","Melanoplus","Melanoplus scudderi group","2","2","2","126","0","0","97.23","451","0","450","36.67","0"
"AGAW1","11S","INS","BOLD:ABA7871","Arthropoda","Insecta","Diptera","Cecidomyiidae","","","","2","2","2","126","0","0","99.78","448","0","450","26.33","0"
"AGAW1","11S","INS","BOLD:ABA5202","Arthropoda","Insecta","Diptera","Limoniidae","Limnophilinae","Pseudolimnophila","Pseudolimnophila luteipennis","2","2","2","126","0","0","98.07","443","0","443.5","33.94","0"
"AGAW1","11S","INS","BOLD:ACA6930","Arthropoda","Insecta","Diptera","Cecidomyiidae","","","","2","2","2","126","0","0","98.61","431","0","450","28","0"
"AGAW1","11S","INS","BOLD:AAL7819","Arthropoda","Insecta","Diptera","Psychodidae","Psychodinae","Psychoda","Psychoda sp., Psychoda sp. BOLD:AAL7819","2","2","2","126","0","0","99.65","415.5","0","416.5","32.85","0"
"AGAW1","11S","INS","BOLD:AAZ5483","Arthropoda","Insecta","Diptera","Muscidae","Phaoniinae","Helina","Helina setiventris","2","2","2","126","0","0","99.33","448.5","0","450","30","0"
"AGAW1","11S","INS","BOLD:ACX1842","Arthropoda","Insecta","Diptera","Agromyzidae","Phytomyzinae","Cerodontha","Cerodontha sp. BOLD:ACX1842","2","2","2","126","0","0","97.26","365","0","450","33.22","0"
"AGAW1","11S","INS","BOLD:AAA2482","Arthropoda","Insecta","Lepidoptera","Noctuidae","Noctuinae","Mythimna","Mythimna unipuncta","2","2","2","126","0","0","99.66","446","0","450","30.67","0"
"AGAW1","11S","INS","BOLD:AAG1743","Arthropoda","Insecta","Diptera","Muscidae","Phaoniinae","Phaonia","Phaonia fuscana","2","2","2","126","0","0","97.35","395.5","0","397.5","32.59","0"
"AGAW1","11S","INS","BOLD:AAH2293","Arthropoda","Insecta","Diptera","Asilidae","Stenopogoninae","Dioctria, Eudioctria","Dioctria hyalipennis, Dioctria baumhaueri, Eudioctria propinqua","2","2","2","126","0","0","99.44","445.5","0","447","30.54","0"
"AGAW1","11S","INS","BOLD:AAA5206","Arthropoda","Insecta","Lepidoptera","Erebidae","Herminiinae","Zanclognatha","Zanclognatha cruralis, Zanclognatha sp. 1, Zanclognatha jacchusalis, Zanclognatha minoralis, Zanclognatha minoralis sp. 1, Zanclognatha sp., Zanclognatha minoralis sp. 3, Zanclognatha gypsalis, Zanclognatha dentata, Zanclognatha theralis, Zanclognatha protumnusalis, Zanclognatha martha, Zanclognatha atrilineella, Zanclognatha obscuripennis, Zanclognatha minoralis sp. 2, Zanclognatha marcidilinea","2","2","2","126","0","0","99.18","424","0","424.5","31.18","0"
"AGAW1","11S","INS","BOLD:AAV5400","Arthropoda","Insecta","Diptera","Cecidomyiidae","","","","2","2","2","126","0","0","98.32","387","0","391","28.53","0"
"AGAW1","11S","INS","BOLD:AAL2750","Arthropoda","Insecta","Diptera","Muscidae","Phaoniinae","Phaonia","Phaonia tuguriorum","2","2","2","126","0","0","99.36","415","0","417","32.05","0"
"AGAW1","11S","INS","BOLD:AAI4303","Arthropoda","Insecta","Diptera","Chironomidae","Chironominae","Chironomus","Chironomus melanescens","2","2","2","126","0","0","98.19","415","0","415","33.61","0"
"AGAW1","11S","INS","BOLD:AAB0651","Arthropoda","Insecta","Lepidoptera","Gelechiidae","Dichomeridinae","Dichomeris","Dichomeris leuconotella, Dichomeris gnoma","2","2","2","126","0","0","99.89","448.5","0","450","33.22","0"
"AGAW1","11S","INS","BOLD:ABY5257","Arthropoda","Insecta","Lepidoptera","Noctuidae","Noctuinae","Apamea","Apamea devastator","2","2","2","126","0","0","98.99","446.5","0","450","31.22","0"
"AGAW1","11S","INS","BOLD:AAN6433","Arthropoda","Insecta","Diptera","Sciaridae","","Corynoptera","Corynoptera deserta","2","2","2","126","0","0","98.96","440","0","444.5","30.71","0"
"AGAW1","11S","INS","BOLD:AAB5640","Arthropoda","Insecta","Coleoptera","Coccinellidae","Coccinellinae","Harmonia","Harmonia axyridis","2","2","2","126","0","0","98.88","446.5","0","450","32.67","0"
"AGAW1","11S","INS","BOLD:AAY6376","Arthropoda","Insecta","Diptera","Cecidomyiidae","","","","2","2","2","126","0","0","97.49","358","0","360","25.42","0"
"AGAW1","11S","INS","BOLD:AAH4135","Arthropoda","Insecta","Diptera","Chloropidae","Chloropinae","Thaumatomyia","Thaumatomyia glabra, Thaumatomyia grata, Thaumatomyia sp. 1 TAW","1","1","1","151","0","0","98.18","384","0","386","32.38","0"
"AGAW1","11S","INS","BOLD:AAM6655","Arthropoda","Insecta","Diptera","Hybotidae","Tachydromiinae","Platypalpus","Platypalpus stabilis","1","1","1","151","0","0","100","450","0","450","35.11","0"
"AGAW1","11S","INS","BOLD:AAD0952","Arthropoda","Insecta","Lepidoptera","Pyralidae","Chrysauginae","Galasa","Galasa nigrinodis, Galasa nigripunctalis","1","1","1","151","0","0","98.49","397","0","397","32.24","0"
"AGAW1","11S","INS","BOLD:ACE4100","Arthropoda","Insecta","Orthoptera","Acrididae","Melanoplinae","Melanoplus","Melanoplus pinicola","1","1","1","151","0","0","98.65","445","0","450","35.11","0"
"AGAW1","11S","INS","BOLD:AAA6560","Arthropoda","Insecta","Lepidoptera","Crambidae","Crambinae","Crambus","Crambus multilinellus, Crambus saltuellus","1","1","1","151","0","0","98.96","383","0","385","30.13","0"
"AGAW1","11S","INS","BOLD:AAH2306","Arthropoda","Insecta","Diptera","Asilidae","Asilinae","Efferia","Efferia aestuans","1","1","1","151","0","0","99.52","413","0","416","36.78","0"
"AGAW1","11S","INS","BOLD:ABV1279","Arthropoda","Insecta","Diptera","Cecidomyiidae","","","","1","1","1","151","0","0","98.02","354","0","354","27.4","0"
"AGAW1","11S","INS","BOLD:ABA0858","Arthropoda","Insecta","Diptera","Cecidomyiidae","","","","1","1","1","151","0","0","97.11","450","0","450","26.22","0"
"AGAW1","11S","INS","BOLD:ADG3385","Arthropoda","Insecta","Diptera","Cecidomyiidae","","","","1","1","1","151","0","0","99.23","392","0","421","23.75","0"
"AGAW1","11S","INS","BOLD:AAM6781","Arthropoda","Insecta","Diptera","Dolichopodidae","Neurigoninae","Neurigona","Neurigona tenuis","1","1","1","151","0","0","98.1","422","0","424","34.91","0"
"AGAW1","11S","INS","BOLD:ADB5456","Arthropoda","Insecta","Hymenoptera","Eulophidae","Entedoninae","Neochrysocharis","","1","1","1","151","0","0","98.31","413","0","450","25.78","0"
"AGAW1","11S","INS","BOLD:ACA9275","Arthropoda","Insecta","Diptera","Chironomidae","Chironominae","Chironomus","","1","1","1","151","0","0","99.11","450","0","450","36","0"
"AGAW1","11S","INS","BOLD:ACL2677","Arthropoda","Insecta","Psocodea","Mesopsocidae","","Mesopsocus","Mesopsocus laticeps","1","1","1","151","0","0","97.33","450","0","450","40.22","0"
"AGAW1","11S","INS","BOLD:AAC1042","Arthropoda","Insecta","Lepidoptera","Crambidae","Spilomelinae","Lygropia","Lygropia rivulalis","1","1","1","151","0","0","97.44","351","0","352","28.41","0"
"AGAW1","11S","INS","BOLD:AAC9132","Arthropoda","Insecta","Diptera","Culicidae","Culicinae","Culiseta","Culiseta inornata","1","1","1","151","0","0","99.77","430","0","431","32.48","0"
"AGAW1","11S","INS","BOLD:AAG5453","Arthropoda","Insecta","Diptera","Chironomidae","Chironominae","Chironomus","Chironomus bifurcatus","1","1","1","151","0","0","98.22","449","0","450","34.89","0"
"AGAW1","11S","INS","BOLD:AAC5210","Arthropoda","Insecta","Diptera","Culicidae","Culicinae","Aedes, Aedimorphus","Aedes japonicus, Aedimorphus caecus","1","1","1","151","0","0","98","450","0","450","35.33","0"
"AGAW1","11S","INS","BOLD:AAB1352","Arthropoda","Insecta","Lepidoptera","Erebidae","Hypeninae","Hypena","Hypena sp.","1","1","1","151","0","0","99.33","447","0","450","32.44","0"
"AGAW1","11S","INS","BOLD:AAY6412","Arthropoda","Insecta","Diptera","Sciaridae","","Bradysia","Bradysia urticae","1","1","1","151","0","0","100","449","0","450","32.22","0"
"AGAW1","11S","INS","BOLD:AAH3914","Arthropoda","Insecta","Diptera","Sciaridae","","Sciara","Sciara hamatilis, Sciara humeralis","1","1","1","151","0","0","98.6","356","0","356","29.78","0"
"AGAW1","11S","INS","BOLD:AAG6432","Arthropoda","Insecta","Diptera","Ceratopogonidae","Forcipomyiinae","Atrichopogon","","1","1","1","151","0","0","97.32","448","0","450","34.89","0"
"AGAW1","11S","INS","BOLD:AAH3225","Arthropoda","Insecta","Psocodea","Psocidae","Psocinae","Metylophorus","Metylophorus novaescotiae","1","1","1","151","0","0","98.24","398","0","398","34.17","0"
"AGAW1","11S","INS","BOLD:ACF4905","Arthropoda","Insecta","Diptera","Cecidomyiidae","","","","1","1","1","151","0","0","98.8","418","0","418","26.08","0"
"AGAW1","11S","INS","BOLD:AAG7279","Arthropoda","Insecta","Diptera","Sphaeroceridae","Limosininae","Spelobia","Spelobia ochripes","1","1","1","151","0","0","99.33","449","0","450","35.11","0"
"AGAW1","11S","INS","BOLD:AAV5575","Arthropoda","Insecta","Diptera","Cecidomyiidae","","","CecidInt30 sp. BOLD:AAV5575","1","1","1","151","0","0","100","449","0","450","25.78","0"
"AGAW1","11S","INS","BOLD:AAB0373","Arthropoda","Insecta","Neuroptera","Chrysopidae","Chrysopinae","Chrysoperla","Chrysoperla sp. isolate 82, Chrysoperla zastrowi, Chrysoperla plorabunda, Chrysoperla sp. isolate 71, Chrysoperla agilis, Italochrysa italica, Chrysoperla sp. isolate 88, Chrysoperla lucasina, Chrysoperla carnea, Chrysoperla mediterranea, Chrysoperla pallida, Chrysoperla nipponensis, Chrysoperla sillemi","1","1","1","151","0","0","99.78","450","0","450","31.56","0"
"AGAW1","11S","INS","BOLD:AAA2632","Arthropoda","Insecta","Lepidoptera","Noctuidae","Noctuinae","Noctua","Noctua atlantica, Noctua pronuba","1","1","1","151","0","0","98.87","353","0","355","30.14","0"
"AGAW1","11S","INS","BOLD:ACG5862","Arthropoda","Insecta","Diptera","Agromyzidae","","","","1","1","1","151","0","0","98.21","447","0","450","34.22","0"
"AGAW1","11S","INS","BOLD:ACK3142","Arthropoda","Insecta","Diptera","Cecidomyiidae","","","","1","1","1","151","0","0","98.89","450","0","450","28.22","0"
"AGAW1","11S","INS","BOLD:AAB7030","Arthropoda","Insecta","Diptera","Chironomidae","Chironominae","Chironomus","Chironomus sp. BOLDAAP2996, Chironomus quinnitukqut, Chironomus cf. decorus 2g, Chironomus decorus group sp. 2 IP2013, Chironomus cf. decorus, Chironomus decorus gr.","1","1","1","151","0","0","97.1","448","0","450","35.56","0"
"AGAW1","11S","INS","BOLD:AAA9251","Arthropoda","Insecta","Lepidoptera","Gelechiidae","Gelechiinae","Scrobipalpa","Scrobipalpa atriplicella, Scrobipalpa nr. atriplicella sp. 1","1","1","1","151","0","0","97.39","422","0","427","33.02","0"
"AGAW1","11S","INS","BOLD:ABV1422","Arthropoda","Insecta","Diptera","Cecidomyiidae","","","","1","1","1","151","0","0","98.89","449","0","450","26.89","0"
"AGAW1","11S","INS","BOLD:ABA0838","Arthropoda","Insecta","Diptera","Cecidomyiidae","","","","1","1","1","151","0","0","99.78","445","0","450","26","0"
"AGAW1","11S","INS","BOLD:ACK3717","Arthropoda","Insecta","Diptera","Cecidomyiidae","","","","1","1","1","151","0","0","99.78","449","0","450","26.44","0"
"AGAW1","11S","INS","BOLD:AAD8027","Arthropoda","Insecta","Diptera","Culicidae","Culicinae","Aedes","Aedes fitchii","1","1","1","151","0","0","100","450","0","450","34.67","0"
"AGAW1","11S","INS","BOLD:AAC2657","Arthropoda","Insecta","Lepidoptera","Coleophoridae","Coleophorinae","Coleophora","Coleophora salinoidella, Coleophora JFL205, Coleophora littorella, Coleophora lineapulvella","1","1","1","151","0","0","99.78","450","0","450","32.67","0"
"AGAW1","11S","INS","BOLD:AAA3867","Arthropoda","Insecta","Lepidoptera","Noctuidae","Noctuinae","Apamea","Apamea amputatrix","1","1","1","151","0","0","99.74","384","0","386","33.68","0"
"AGAW1","11S","INS","BOLD:ABV9293","Arthropoda","Insecta","Diptera","Chironomidae","Chironominae","Dicrotendipes","","1","1","1","151","0","0","100","355","0","450","34.89","0"
"AGAW1","11S","INS","BOLD:ADG6553","Arthropoda","Insecta","Diptera","Chironomidae","","","","1","1","1","151","0","0","99.23","388","0","450","32.44","0"
"AGAW1","11S","INS","BOLD:ABA0822","Arthropoda","Insecta","Diptera","Cecidomyiidae","","","","1","1","1","151","0","0","98.5","401","0","403","25.06","0"
"AGAW1","11S","INS","BOLD:AAG2898","Arthropoda","Insecta","Hemiptera","Cicadellidae","Deltocephalinae","Paraphlepsius","Paraphlepsius apertus, Paraphlepsius irroratus","1","1","1","151","0","0","99.78","450","0","450","32.22","0"
"AGAW1","11S","INS","BOLD:ADH9867","Arthropoda","Insecta","Diptera","","","","","1","1","1","151","0","0","98.08","364","0","367","38.15","0"
"AGAW1","11S","INS","BOLD:ABZ4994","Arthropoda","Insecta","Orthoptera","Acrididae","Melanoplinae","Melanoplus","Melanoplus gladstoni, Melanoplus sanguinipes","1","1","1","151","0","0","97.11","450","0","450","36.44","0"
"AGAW1","11S","INS","BOLD:ABV1237","Arthropoda","Insecta","Diptera","Chironomidae","Chironominae","Chironomus","","1","1","1","151","0","0","98.44","449","0","450","34.44","0"
"AGAW1","11S","INS","BOLD:AAA7368","Arthropoda","Insecta","Lepidoptera","Cosmopterigidae","Cosmopteriginae","Limnaecia","Limnaecia phragmitella","1","1","1","151","0","0","98","449","0","450","31.11","0"
"AGAW1","11S","INS","BOLD:ABZ8007","Arthropoda","Insecta","Lepidoptera","Noctuidae","Cuculliinae","Cucullia","Cucullia similaris, Cucullia postera, Cucullia sp., Cucullia florea","1","1","1","151","0","0","99.56","450","0","450","31.78","0"
"AGAW1","11S","INS","BOLD:AAA1962","Arthropoda","Insecta","Diptera","Sarcophagidae","Sarcophaginae","Helicobia, sarcJanzen01, Sarcophaga","sarcJanzen01 Janzen14, Helicobia rapax, Sarcophaga rapax","1","1","1","151","0","0","100","448","0","450","34.67","0"
"AGAW1","11S","INS","BOLD:AAV0162","Arthropoda","Insecta","Hemiptera","Cicadellidae","Deltocephalinae","Eutettix","Eutettix variabilis","1","1","1","151","0","0","97.81","365","0","450","36.89","0"
"AGAW1","11S","INS","BOLD:AAZ0271","Arthropoda","Insecta","Diptera","Ceratopogonidae","Ceratopogoninae","Culicoides","","1","1","1","151","0","0","100","400","0","403","40.69","0"
"AGAW1","11S","INS","BOLD:ACR3866","Arthropoda","Insecta","Psocodea","Caeciliusidae","Caeciliusinae","Valenzuela","","1","1","1","151","0","0","97.67","429","0","434","35.48","0"
"AGAW1","11S","INS","BOLD:ABA0803","Arthropoda","Insecta","Diptera","Ceratopogonidae","Ceratopogoninae","Culicoides","Culicoides stellifer","1","1","1","151","0","0","98","449","0","450","36.89","0"
"AGAW1","11S","NAR","no match","no match","no match","no match","no match","no match","no match","no match","no match","no match","no match","no match","no match","no match","no match","no match","no match","no match","no match","no match"
"AGAW1","11S","NIN","BOLD:AAB3836","Arthropoda","Arachnida","Araneae","Philodromidae","","Philodromus","Philodromus cespitum","553","512","512","1","7.41","0","98.89","389.37","0","426.26","30.08","0"
"AGAW1","11S","NIN","BOLD:AAA6381","Arthropoda","Arachnida","Araneae","Tetragnathidae","","Tetragnatha","Tetragnatha laboriosa","383","366","366","2","4.44","0","99.2","425.5","0","427.7","39.4","0"
"AGAW1","11S","NIN","BOLD:AAU6252","Arthropoda","Arachnida","Trombidiformes","Bdellidae","","Cyta","","318","257","257","3","19.18","0","98.55","432.25","0","433.69","31.46","0"
"AGAW1","11S","NIN","BOLD:AAG5631","Arthropoda","Arachnida","Araneae","Linyphiidae","","Tennesseellum","Tennesseellum formica","241","181","181","4","24.9","0","97.95","445.55","0","446.81","31.65","0"
"AGAW1","11S","NIN","BOLD:ACY2550","Arthropoda","Arachnida","Trombidiformes","Bdellidae","","Cyta","","47","46","46","5","2.13","0","98.69","376.23","0","398.68","31.16","0"
"AGAW1","11S","NIN","BOLD:AAB7914","Arthropoda","Collembola","Symphypleona","Bourletiellidae","","Bourletiella","","27","24","24","6","11.11","0","99.6","445.26","0","446.26","38.17","0"
"AGAW1","11S","NIN","BOLD:AAB7601","Arthropoda","Arachnida","Araneae","Eutichuridae","","Cheiracanthium","Cheiracanthium mildei","24","24","24","7","0","0","98.5","438","0","439.17","28.59","0"
"AGAW1","11S","NIN","BOLD:ABW2639","Arthropoda","Arachnida","Trombidiformes","Anystidae","","","","16","15","15","8","6.25","0","99.44","448.94","0","450","31.51","0"
"AGAW1","11S","NIN","BOLD:AAM7961","Arthropoda","Arachnida","Trombidiformes","Anystidae","","","","11","11","11","9","0","0","99.23","446.45","0","444.73","31.3","0"
"AGAW1","11S","NIN","BOLD:AAA7242","Arthropoda","Collembola","Entomobryomorpha","Entomobryidae","Seirinae, Entomobryinae","Entomobrya, Lepidosira","Entomobrya multifasciata, Lepidosira sp.","7","7","7","10","0","0","99.56","419.57","0","422","39.31","0"
"AGAW1","11S","NIN","BOLD:ACZ1295","Arthropoda","Arachnida","Trombidiformes","Anystidae","","","","6","6","6","11","0","0","99.01","382.17","0","396","31.78","0"
"AGAW1","11S","NIN","BOLD:AAH6630","Arthropoda","Collembola","Entomobryomorpha","Entomobryidae","Entomobryinae","Entomobrya","Entomobrya unostrigata","4","4","4","12","0","0","98.98","430","0","431.25","39.3","0"
"AGAW1","11S","NIN","BOLD:AAB0726","Arthropoda","Arachnida","Araneae","Lycosidae","","Trochosa","Trochosa ruricola","3","3","3","13","0","0","99.11","446.33","0","450","31.26","0"
"AGAW1","11S","NIN","BOLD:ACL6413","Arthropoda","Arachnida","Trombidiformes","Erythraeidae","","Balaustium","","2","2","2","14","0","0","98.44","449","0","450","36.56","0"
"AGAW1","11S","NIN","BOLD:AAA7246","Arthropoda","Collembola","Entomobryomorpha","Entomobryidae","Entomobryinae","Entomobrya","Entomobrya nicoleti, Entomobrya multifasciata, Entomobrya intermedia","1","1","1","15","0","0","98.44","449","0","450","43.11","0"
"AGAW1","11S","NIN","BOLD:ADG1586","Arthropoda","Collembola","Entomobryomorpha","Entomobryidae","","","","1","1","1","15","0","0","99.75","401","0","417","40.05","0"
"AGAW1","11S","NIN","BOLD:ACP2815","Arthropoda","Arachnida","Mesostigmata","Ascidae","","","","1","1","1","15","0","0","99.49","389","0","428","33.18","0"
"AGAW1","11S","NIN","BOLD:ADE0402","","","","","","","","1","1","1","15","0","0","97.76","401","0","403","28.54","0"
"AGAW1","11W","BAC","TAX:175768","Proteobacteria","Alphaproteobacteria","Rickettsiales","Rickettsiaceae",NA,"Wolbachia","Wolbachia pipientis","44","41","41","1","6.82","0","98.38","446.16","0","447.39","39.46","0"
"AGAW1","11W","BAC","TAX:91393","Proteobacteria","Alphaproteobacteria","Rickettsiales","Rickettsiaceae",NA,"Wolbachia","","25","25","25","2","0","0","99.36","391.2","0","424.56","39.15","0"
"AGAW1","11W","BAC","TAX:91391","Proteobacteria","Alphaproteobacteria","Rickettsiales","",NA,"","","20","17","17","3","15","0","98.83","420","0","445.75","39.86","0"
"AGAW1","11W","BAC","TAX:91389","Proteobacteria","","","",NA,"","","3","3","3","4","0","0","99.44","357.33","0","363.33","38.35","0"
"AGAW1","11W","BAC","TAX:376688","Proteobacteria","Alphaproteobacteria","Rickettsiales","Rickettsiaceae",NA,"Wolbachia","Wolbachia sp. elachBioLep01 BioLep373","1","1","1","5","0","0","99.33","450","0","450","40.22","0"
"AGAW1","11W","CHO","no match","no match","no match","no match","no match","no match","no match","no match","no match","no match","no match","no match","no match","no match","no match","no match","no match","no match","no match","no match"
"AGAW1","11W","INS","BOLD:AAP8739","Arthropoda","Insecta","Diptera","Platystomatidae","Platystomatinae","Rivellia","Rivellia steyskali","43723","28097","28097","1","35.74","0","99.14","438.2","0","439.62","38.04","0"
"AGAW1","11W","INS","BOLD:AAG2123","Arthropoda","Insecta","Diptera","Tachinidae","Dexiinae","Dinera","Dinera grisescens","10516","7414","7414","2","29.5","0","99.37","431.69","0","433.26","31.84","0"
"AGAW1","11W","INS","BOLD:AAU6681","Arthropoda","Insecta","Diptera","Tipulidae","Tipulinae","Nephrotoma","Nephrotoma cornicina","10022","7320","7320","3","26.96","0","99.61","442.11","0","443.35","36.32","0"
"AGAW1","11W","INS","BOLD:ABW1379","Arthropoda","Insecta","Diptera","Chloropidae","Oscinellinae","Malloewia","Malloewia abdominalis","3899","2831","2831","4","27.39","0","98.27","444.06","0","446.48","31.51","0"
"AGAW1","11W","INS","BOLD:AAJ7123","Arthropoda","Insecta","Diptera","Culicidae","Culicinae","Culiseta","Culiseta minnesotae","3028","2506","2506","5","17.24","0","99.09","432.93","0","434.24","34.74","0"
"AGAW1","11W","INS","BOLD:AAA7661","Arthropoda","Insecta","Diptera","Culicidae","Culicinae","Culiseta, Culex","Culiseta melanura, Culex articularis, Culex restuans, Culex acharistus","2802","2508","2508","6","10.49","0","99.28","432.14","0","434.1","32.76","0"
"AGAW1","11W","INS","BOLD:AAG2868","Arthropoda","Insecta","Hemiptera","Cicadellidae","Typhlocybinae","Empoasca","Empoasca sp., Empoasca fabae","2628","1642","1642","7","37.52","0","99.63","445.1","0","447.58","32.33","0"
"AGAW1","11W","INS","BOLD:ACF8174","Arthropoda","Insecta","Diptera","Chironomidae","Chironominae","","","1859","1572","1572","8","15.44","0","98.4","436.75","0","438.12","33.31","0"
"AGAW1","11W","INS","BOLD:AAK8751","Arthropoda","Insecta","Diptera","Agromyzidae","Phytomyzinae","Liriomyza","Liriomyza sp. SB, Liriomyza trifoliearum","1586","1196","1196","9","24.59","0","99.49","427.66","0","441.24","33.6","0"
"AGAW1","11W","INS","BOLD:AAG5507","Arthropoda","Insecta","Diptera","Chironomidae","Chironominae","Chironomus","Chironomus atrella","1328","1283","1283","10","3.39","0","99.23","423.01","0","429.3","36","0"
"AGAW1","11W","INS","BOLD:AAF9028","Arthropoda","Insecta","Diptera","Limoniidae","","","","1033","1021","1021","11","1.16","0","98.87","402.44","0","405.71","38.23","0"
"AGAW1","11W","INS","BOLD:AAG9897","Arthropoda","Insecta","Dermaptera","Forficulidae","Forficulinae","Forficula","Forficula auricularia-A","904","892","892","12","1.33","0","99.14","396.31","0","406.51","45.74","0"
"AGAW1","11W","INS","BOLD:AAN5162","Arthropoda","Insecta","Diptera","Ceratopogonidae","Dasyheleinae","Dasyhelea","","803","681","681","13","15.19","0","99.4","430.52","0","432.5","33.75","0"
"AGAW1","11W","INS","BOLD:AAK5928","Arthropoda","Insecta","Coleoptera","Chrysomelidae","Criocerinae","Oulema","Oulema melanopus","731","603","603","14","17.51","0","99.6","439.35","0","440.63","36.48","0"
"AGAW1","11W","INS","BOLD:AAN5430","Arthropoda","Insecta","Diptera","Agromyzidae","","","","689","603","603","15","12.48","0","98.85","415.5","0","435.59","31.97","0"
"AGAW1","11W","INS","BOLD:AAB8271","Arthropoda","Insecta","Lepidoptera","Autostichidae","Oegoconiinae","Oegoconia","Oegoconia quadripuncta, Oegoconia deauratella","542","534","534","16","1.48","0","99.11","398.49","0","400.43","34.34","0"
"AGAW1","11W","INS","BOLD:AAB4657","Arthropoda","Insecta","Diptera","Chironomidae","Chironominae","Chironomus","Chironomus maturus","539","454","454","17","15.77","0","99.27","432.5","0","434.66","37.39","0"
"AGAW1","11W","INS","BOLD:AAD2041","Arthropoda","Insecta","Diptera","Dolichopodidae","Sciapodinae","Condylostylus","Condylostylus inermis, Condylostylus nigrofemoratus, Condylostylus connectans, Condylostylus caudatus, Condylostylus flavipes","520","406","406","18","21.92","0","99.04","436.64","0","438.47","35.9","0"
"AGAW1","11W","INS","BOLD:ACV4278","Arthropoda","Insecta","Diptera","Limoniidae","Chioneinae","Erioptera","Erioptera chlorophylla","517","497","497","19","3.87","0","99.22","427.64","0","430.61","32.91","0"
"AGAW1","11W","INS","BOLD:AAF3704","Arthropoda","Insecta","Hemiptera","Artheneidae","Artheneinae","Chilacis","Chilacis typhae","484","382","382","20","21.07","0","99.58","436.04","0","437.46","34.14","0"
"AGAW1","11W","INS","BOLD:AAG4333","Arthropoda","Insecta","Coleoptera","Staphylinidae","Staphylininae","Xantholinus","Xantholinus linearis","482","346","346","21","28.22","0","99.63","444.96","0","446.42","33.45","0"
"AGAW1","11W","INS","BOLD:ABV1430","Arthropoda","Insecta","Diptera","Dolichopodidae","Medeterinae","Medetera","","465","443","443","22","4.73","0","99.3","415.82","0","421.05","39","0"
"AGAW1","11W","INS","BOLD:AAO7783","Arthropoda","Insecta","Diptera","Dolichopodidae","Diaphorinae","","","452","377","377","23","16.59","0","99.14","438.24","0","440.13","33.44","0"
"AGAW1","11W","INS","BOLD:ABA0863","Arthropoda","Insecta","Diptera","Cecidomyiidae","","","","446","386","386","24","13.45","0","99.29","437.68","0","439","29.87","0"
"AGAW1","11W","INS","BOLD:ADG6553","Arthropoda","Insecta","Diptera","Chironomidae","","","","442","405","405","25","8.37","0","99.43","387.75","0","444.48","33.27","0"
"AGAW1","11W","INS","BOLD:ABW7897","Arthropoda","Insecta","Diptera","Cecidomyiidae","","","","439","327","327","26","25.51","0","99.65","443.32","0","445.51","29.71","0"
"AGAW1","11W","INS","BOLD:ABY5318","Arthropoda","Insecta","Diptera","Chloropidae","Oscinellinae","Incertella, Rhopalopterum","Rhopalopterum umbrosum, Rhopalopterum luteiceps, Incertella bispina, Rhopalopterum painteri","401","284","284","27","29.18","0","99.29","445.19","0","446.14","32.41","0"
"AGAW1","11W","INS","BOLD:AAN5199","Arthropoda","Insecta","Diptera","Cecidomyiidae","","","CecidInt20 sp. BOLD:AAN5199","393","307","307","28","21.88","0","99.59","436.45","0","444.59","26.59","0"
"AGAW1","11W","INS","BOLD:AAC4197","Arthropoda","Insecta","Diptera","Chironomidae","Orthocladiinae","Paraphaenocladius","Paraphaenocladius impensus","381","323","323","29","15.22","0","99.21","432.6","0","437.21","34.87","0"
"AGAW1","11W","INS","BOLD:AAL4232","Arthropoda","Insecta","Diptera","Agromyzidae","Phytomyzinae","Phytoliriomyza","Phytoliriomyza arctica","321","261","261","30","18.69","0","99.57","439.52","0","440.85","32.03","0"
"AGAW1","11W","INS","BOLD:ACF2457","Arthropoda","Insecta","Diptera","Chironomidae","Chironominae","Paratanytarsus","Paratanytarsus laccophilus","306","293","293","31","4.25","0","98.95","422.84","0","425.2","32.71","0"
"AGAW1","11W","INS","BOLD:AAH3661","Arthropoda","Insecta","Diptera","Cecidomyiidae","","","","288","267","267","32","7.29","0","99.21","429.22","0","431.07","25.28","0"
"AGAW1","11W","INS","BOLD:AAM9013","Arthropoda","Insecta","Diptera","Keroplatidae","Macrocerinae","Macrocera","Macrocera pusilla","282","259","259","33","8.16","0","99.36","428.75","0","430.46","30.24","0"
"AGAW1","11W","INS","BOLD:ACK3589","Arthropoda","Insecta","Diptera","Cecidomyiidae","","","","270","220","220","34","18.52","0","99.28","447.46","0","448.41","32.25","0"
"AGAW1","11W","INS","BOLD:AAN5242","Arthropoda","Insecta","Diptera","Cecidomyiidae","","","","256","254","254","35","0.78","0","98.35","409.38","0","411.64","28.21","0"
"AGAW1","11W","INS","BOLD:AAY6730","Arthropoda","Insecta","Hemiptera","Derbidae","Cedusinae","Cedusa","Cedusa incisa","230","214","214","36","6.96","0","99.17","422.73","0","424.47","35.35","0"
"AGAW1","11W","INS","BOLD:AAN5238","Arthropoda","Insecta","Diptera","Cecidomyiidae","","","","228","188","188","37","17.54","0","99.45","427.49","0","428.74","25.57","0"
"AGAW1","11W","INS","BOLD:AAZ5592","Arthropoda","Insecta","Diptera","Psychodidae","Psychodinae","","","211","196","196","38","7.11","0","99.15","430.11","0","432.55","32.44","0"
"AGAW1","11W","INS","BOLD:AAN5183","Arthropoda","Insecta","Diptera","Cecidomyiidae","","","","146","118","118","39","19.18","0","98.58","443.32","0","444.44","27.82","0"
"AGAW1","11W","INS","BOLD:AAU7126","Arthropoda","Insecta","Coleoptera","Cleridae","Hydnocerinae","Isohydnocera","Isohydnocera curtipennis","144","120","120","40","16.67","0","99.43","440.8","0","442.35","33.02","0"
"AGAW1","11W","INS","BOLD:ACR3866","Arthropoda","Insecta","Psocodea","Caeciliusidae","Caeciliusinae","Valenzuela","","138","125","125","41","9.42","0","99.46","437.41","0","438.62","35.39","0"
"AGAW1","11W","INS","BOLD:ACK4716","Arthropoda","Insecta","Diptera","Cecidomyiidae","","","","133","122","122","42","8.27","0","99.28","431.36","0","432.73","31.66","0"
"AGAW1","11W","INS","BOLD:AAF9949","Arthropoda","Insecta","Hemiptera","Miridae","Mirinae","Trigonotylus","Trigonotylus pulchellus, Trigonotylus americanus, Trigonotylus psammaecolor, Trigonotylus caelestialium, Trigonotylus viridis, Trigonotylus ruficornis","122","87","87","43","28.69","0","99.69","440.91","0","445.04","34.53","0"
"AGAW1","11W","INS","BOLD:ACB0984","Arthropoda","Insecta","Psocodea","Stenopsocidae","","Graphopsocus","Graphopsocus cruciatus","109","85","85","44","22.02","0","99.24","411.75","0","446.62","37.48","0"
"AGAW1","11W","INS","BOLD:AAF9771","Arthropoda","Insecta","Diptera","Hybotidae","Tachydromiinae","Platypalpus","Platypalpus longiseta, Platypalpus trivialis, Platypalpus sp. 5, Platypalpus sp. 17, Platypalpus pallidiventris","105","74","74","45","29.52","0","99.36","443.5","0","444.39","34.83","0"
"AGAW1","11W","INS","BOLD:ADF6661","Arthropoda","Insecta","Diptera","Cecidomyiidae","","","","102","91","91","46","10.78","0","98.87","416.96","0","443.64","29.96","0"
"AGAW1","11W","INS","BOLD:ACC8079","Arthropoda","Insecta","Diptera","Cecidomyiidae","","","","95","74","74","47","22.11","0","99.29","445.08","0","446.58","28.94","0"
"AGAW1","11W","INS","BOLD:AAH3741","Arthropoda","Insecta","Diptera","Cecidomyiidae","","","","91","91","91","48","0","0","97.78","428.43","0","431.8","26.94","0"
"AGAW1","11W","INS","BOLD:ABA9076","Arthropoda","Insecta","Coleoptera","Latridiidae","Latridiinae","Cartodere","Cartodere constricta","74","66","66","49","10.81","0","99.24","427.47","0","428.85","35.16","0"
"AGAW1","11W","INS","BOLD:ADD3648","Arthropoda","Insecta","Diptera","Cecidomyiidae","","","","69","65","65","50","5.8","0","99.21","434.65","0","436.48","26.24","0"
"AGAW1","11W","INS","BOLD:ABA0822","Arthropoda","Insecta","Diptera","Cecidomyiidae","","","","60","59","59","51","1.67","0","98.96","433.92","0","435.63","26.56","0"
"AGAW1","11W","INS","BOLD:ACU7906","Arthropoda","Insecta","Diptera","Chironomidae","Orthocladiinae","","","55","55","55","52","0","0","98.8","403.75","0","406.35","36.2","0"
"AGAW1","11W","INS","BOLD:AAM6040","Arthropoda","Insecta","Diptera","Cecidomyiidae","","","","52","50","50","53","3.85","0","99.27","429.4","0","430.63","26.81","0"
"AGAW1","11W","INS","BOLD:AAP4636","Arthropoda","Insecta","Psocodea","Caeciliusidae","","","","51","51","51","54","0","0","99.53","363.61","0","402.02","36.31","0"
"AGAW1","11W","INS","BOLD:AAN5148","Arthropoda","Insecta","Diptera","Ceratopogonidae","Forcipomyiinae","Forcipomyia","Forcipomyia bipunctata, Forcipomyia sp. CK-2012","47","44","44","55","6.38","0","99.32","436.13","0","438.43","35.9","0"
"AGAW1","11W","INS","BOLD:AAN6447","Arthropoda","Insecta","Diptera","Sciaridae","","Corynoptera","Corynoptera perpusilla","47","40","40","55","14.89","0","99.44","448.66","0","450","36.44","0"
"AGAW1","11W","INS","BOLD:AAM6037","Arthropoda","Insecta","Diptera","Cecidomyiidae","Cecidomyiinae","Contarinia, Dasineura","","44","39","39","57","11.36","0","99.24","440.5","0","441.91","26.47","0"
"AGAW1","11W","INS","BOLD:AAB0079","Arthropoda","Insecta","Diptera","Chironomidae","Orthocladiinae","Corynoneura","Corynoneura arctica","43","40","40","58","6.98","0","99.15","431.4","0","433.42","34.77","0"
"AGAW1","11W","INS","BOLD:AAG1711","Arthropoda","Insecta","Diptera","Muscidae","Phaoniinae","Helina","Helina depuncta","39","39","39","59","0","0","99.33","434.95","0","437.54","31.36","0"
"AGAW1","11W","INS","BOLD:AAD1485","Arthropoda","Insecta","Diptera","Chironomidae","Chironominae","Paratanytarsus","Paratanytarsus grimmii, Paratanytarsus sp.","38","37","37","60","2.63","0","99.34","439.66","0","441.45","33.83","0"
"AGAW1","11W","INS","BOLD:ACM0918","Arthropoda","Insecta","Diptera","Cecidomyiidae","","","","37","21","21","61","43.24","0","97.28","446.11","0","446.43","27.94","0"
"AGAW1","11W","INS","BOLD:AAG0892","Arthropoda","Insecta","Neuroptera","Hemerobiidae","Hemerobiinae","Hemerobius","Hemerobius simulans, Hemerobius perelegans, Hemerobius humulinus","36","35","35","62","2.78","0","99.29","418.28","0","420.06","29.87","0"
"AGAW1","11W","INS","BOLD:AAH0135","Arthropoda","Insecta","Coleoptera","Phalacridae","Phalacrinae","Acylomus","Acylomus pugetanus, Acylomus ergoti","36","34","34","62","5.56","0","99.02","436.58","0","438.67","31.81","0"
"AGAW1","11W","INS","BOLD:AAO4092","Arthropoda","Insecta","Psocodea","Caeciliusidae","Caeciliusinae","Valenzuela","Valenzuela burmeisteri","36","36","36","62","0","0","98.82","420.28","0","422.64","34.96","0"
"AGAW1","11W","INS","BOLD:ABA6407","Arthropoda","Insecta","Diptera","Sciaridae","","Pseudolycoriella","","33","33","33","65","0","0","99.27","429.79","0","431.76","34.95","0"
"AGAW1","11W","INS","BOLD:AAG6745","Arthropoda","Insecta","Diptera","Calliphoridae","Calliphorinae","Pollenia","Pollenia pediculata","28","28","28","66","0","0","99.23","418.93","0","421.68","33.51","0"
"AGAW1","11W","INS","BOLD:AAB8143","Arthropoda","Insecta","Hemiptera","Nabidae","Nabinae","Nabis","Nabis inscriptus, Nabis americoferus, Nabis alternatus, Nabis sp.","27","18","18","67","33.33","0","99.49","439.26","0","440.3","36.6","0"
"AGAW1","11W","INS","BOLD:AAN6154","Arthropoda","Insecta","Coleoptera","Latridiidae","Corticariinae","Melanophthalma","Melanophthalma inermis","25","25","25","68","0","0","98.57","431.04","0","432.28","31.49","0"
"AGAW1","11W","INS","BOLD:ACX5521","Arthropoda","Insecta","Diptera","Chironomidae","Orthocladiinae","","","24","24","24","69","0","0","98.7","426.46","0","428","34.93","0"
"AGAW1","11W","INS","BOLD:AAV5609","Arthropoda","Insecta","Diptera","Cecidomyiidae","","","","23","23","23","70","0","0","97.84","438.52","0","439.74","27.67","0"
"AGAW1","11W","INS","BOLD:AAN8770","Arthropoda","Insecta","Diptera","Psychodidae","Psychodinae","Psychoda","Psychoda trinodulosa","22","21","21","71","4.55","0","98.95","433.23","0","434.32","33.39","0"
"AGAW1","11W","INS","BOLD:ACK5742","Arthropoda","Insecta","Diptera","Cecidomyiidae","","","","20","19","19","72","5","0","99.46","430.8","0","431.45","28.9","0"
"AGAW1","11W","INS","BOLD:AAN5255","Arthropoda","Insecta","Diptera","Cecidomyiidae","Cecidomyiinae","Dasineura","","20","20","20","72","0","0","98.5","427.7","0","436.55","26.79","0"
"AGAW1","11W","INS","BOLD:AAC0903","Arthropoda","Insecta","Diptera","Chironomidae","Chironominae","Chironomus","Chironomus acidophilus","20","20","20","72","0","0","99.48","438.2","0","441.4","36.35","0"
"AGAW1","11W","INS","BOLD:ABU5546","Arthropoda","Insecta","Diptera","Psychodidae","Psychodinae","Psychoda","","20","19","19","72","5","0","98.56","441.35","0","443.15","31.43","0"
"AGAW1","11W","INS","BOLD:AAY6469","Arthropoda","Insecta","Diptera","Cecidomyiidae","","","","19","19","19","76","0","0","99.47","422.89","0","424.53","26.02","0"
"AGAW1","11W","INS","BOLD:AAN5220","Arthropoda","Insecta","Diptera","Cecidomyiidae","","","","16","16","16","77","0","0","98.16","422.38","0","426.25","25.54","0"
"AGAW1","11W","INS","BOLD:ADI1135","Arthropoda","Insecta","Diptera","Chironomidae","Chironominae","Microtendipes","","15","15","15","78","0","0","97.28","399.8","0","402.27","32.95","0"
"AGAW1","11W","INS","BOLD:ABA7871","Arthropoda","Insecta","Diptera","Cecidomyiidae","","","","14","14","14","79","0","0","99","447.21","0","447.93","26.93","0"
"AGAW1","11W","INS","BOLD:AAV0214","Arthropoda","Insecta","Hemiptera","Cicadellidae","Eurymelinae","Idiocerus","Idiocerus delongi, Idiocerus pallidus","13","13","13","80","0","0","98.58","417.62","0","425.77","32.73","0"
"AGAW1","11W","INS","BOLD:AAZ5483","Arthropoda","Insecta","Diptera","Muscidae","Phaoniinae","Helina","Helina setiventris","12","12","12","81","0","0","99.17","439.58","0","441.92","29.48","0"
"AGAW1","11W","INS","BOLD:AAA3453","Arthropoda","Insecta","Diptera","Anthomyiidae","Anthomyiinae","Delia","","11","11","11","82","0","0","99.62","425.18","0","430.27","34.72","0"
"AGAW1","11W","INS","BOLD:AAU8483","","","","","","","","11","11","11","82","0","0","98.69","432.55","0","434.73","28.94","0"
"AGAW1","11W","INS","BOLD:AAA3181","Arthropoda","Insecta","Diptera","Muscidae","Muscinae","Stomoxys","Stomoxys calcitrans","10","10","10","84","0","0","99.01","425.2","0","429.3","32.92","0"
"AGAW1","11W","INS","BOLD:ABV9336","Arthropoda","Insecta","Diptera","Cecidomyiidae","","","","10","10","10","84","0","0","98.91","447.1","0","448.6","26.28","0"
"AGAW1","11W","INS","BOLD:ABX7402","Arthropoda","Insecta","Diptera","Cecidomyiidae","","","","9","9","9","86","0","0","99.43","420.89","0","440.78","27.75","0"
"AGAW1","11W","INS","BOLD:ACC7907","Arthropoda","Insecta","Diptera","Cecidomyiidae","","","","9","9","9","86","0","0","98.98","394.56","0","396.56","25.42","0"
"AGAW1","11W","INS","BOLD:AAA4222","Arthropoda","Insecta","Lepidoptera","Erebidae","Hypeninae","Hypena","Hypena scabra","9","9","9","86","0","0","99.09","395.33","0","397.22","32.41","0"
"AGAW1","11W","INS","BOLD:AAP5870","Arthropoda","Insecta","Diptera","Chironomidae","Chironominae","Tanytarsus","","9","9","9","86","0","0","98.13","414","0","416.11","36.22","0"
"AGAW1","11W","INS","BOLD:AAH3662","Arthropoda","Insecta","Diptera","Cecidomyiidae","","","CecidInt3 sp. BOLD:AAH3662","9","9","9","86","0","0","99.21","448.33","0","450","26.02","0"
"AGAW1","11W","INS","BOLD:AAC4523","Arthropoda","Insecta","Diptera","Chironomidae","Chironominae","Tanytarsus","Tanytarsus guerlus","8","7","7","91","12.5","0","99.25","420.5","0","424.88","32.55","0"
"AGAW1","11W","INS","BOLD:ACL4792","Arthropoda","Insecta","Diptera","Cecidomyiidae","","","","8","8","8","91","0","0","98","417.63","0","419.88","27.32","0"
"AGAW1","11W","INS","BOLD:ABX2572","Arthropoda","Insecta","Thysanoptera","Phlaeothripidae","Phlaeothripinae","","","7","7","7","93","0","0","98.9","442.57","0","450","27.43","0"
"AGAW1","11W","INS","BOLD:AAV1136","Arthropoda","Insecta","Diptera","Scatopsidae","","","","7","6","6","93","14.29","0","99.29","418.14","0","420.29","36.54","0"
"AGAW1","11W","INS","BOLD:ABU5525","Arthropoda","Insecta","Diptera","Chironomidae","Orthocladiinae","Limnophyes","Limnophyes sp. 14ES","7","7","7","93","0","0","99.45","396.71","0","398.29","36.95","0"
"AGAW1","11W","INS","BOLD:AAV5561","Arthropoda","Insecta","Diptera","Cecidomyiidae","","","","7","7","7","93","0","0","99.27","448.71","0","450","27.27","0"
"AGAW1","11W","INS","BOLD:AAN6041","Arthropoda","Insecta","Coleoptera","Latridiidae","Corticariinae","Corticaria","Corticaria elongata","7","6","6","93","14.29","0","99.8","438.57","0","439.57","29.93","0"
"AGAW1","11W","INS","BOLD:ABY5257","Arthropoda","Insecta","Lepidoptera","Noctuidae","Noctuinae","Apamea","Apamea devastator","7","7","7","93","0","0","99.04","445","0","446.86","30.66","0"
"AGAW1","11W","INS","BOLD:AAM9255","Arthropoda","Insecta","Diptera","Sciaridae","","Pseudolycoriella","","7","7","7","93","0","0","99.06","399.29","0","402.43","30.61","0"
"AGAW1","11W","INS","BOLD:AAL2750","Arthropoda","Insecta","Diptera","Muscidae","Phaoniinae","Phaonia","Phaonia tuguriorum","6","6","6","100","0","0","99.58","440.17","0","441.17","32.77","0"
"AGAW1","11W","INS","BOLD:AAG2511","Arthropoda","Insecta","Diptera","Anthomyiidae","Anthomyiinae","Delia","Delia platura","6","6","6","100","0","0","98.82","442.5","0","445.33","33.27","0"
"AGAW1","11W","INS","BOLD:AAG2740","Arthropoda","Insecta","Diptera","Ephydridae","Parydrinae","Philygria","Philygria debilis","6","6","6","100","0","0","98.06","406.67","0","410.83","33.32","0"
"AGAW1","11W","INS","BOLD:AAU6595","Arthropoda","Insecta","Diptera","Sciaridae","","Corynoptera","Corynoptera perpusilla","5","5","5","103","0","0","99.52","412","0","414.4","35.84","0"
"AGAW1","11W","INS","BOLD:ACP8190","Arthropoda","Insecta","Hymenoptera","Platygastridae","Scelioninae","","","5","5","5","103","0","0","99.43","349.8","0","397.6","27.27","0"
"AGAW1","11W","INS","BOLD:AAG7891","Arthropoda","Insecta","Hymenoptera","Platygastridae","Scelioninae","Telenomus","Telenomus podisi","5","5","5","103","0","0","99.27","437.6","0","450","27.2","0"
"AGAW1","11W","INS","BOLD:AAI4303","Arthropoda","Insecta","Diptera","Chironomidae","Chironominae","Chironomus","Chironomus melanescens","5","5","5","103","0","0","97.99","436.6","0","439.8","34.1","0"
"AGAW1","11W","INS","BOLD:AAN5178","Arthropoda","Insecta","Diptera","Cecidomyiidae","Cecidomyiinae","Rhopalomyia","Rhopalomyia sp. BOLD:AAN5178","5","5","5","103","0","0","98.98","422","0","427","26.43","0"
"AGAW1","11W","INS","BOLD:AAA8933","Arthropoda","Insecta","Coleoptera","Coccinellidae","Coccinellinae","Coccinella","Coccinella septempunctata","4","4","4","108","0","0","99.72","448.25","0","450","33.06","0"
"AGAW1","11W","INS","BOLD:AAQ2138","Arthropoda","Insecta","Diptera","Sciaridae","","","","4","4","4","108","0","0","99.25","426.25","0","428.5","35.68","0"
"AGAW1","11W","INS","BOLD:AAA4277","Arthropoda","Insecta","Diptera","Syrphidae","Syrphinae","Toxomerus","Toxomerus marginatus, Toxomerus sp. CR-3, Toxomerus CR-3","4","3","3","108","25","0","99.61","449.5","0","450","30.44","0"
"AGAW1","11W","INS","BOLD:AAA4555","Arthropoda","Insecta","Orthoptera","Acrididae","Melanoplinae","Melanoplus","Melanoplus stupefactus, Melanoplus angustipennis, Melanoplus ponderosus, Melanoplus bruneri, Melanoplus frigidus, Melanoplus scapularis, Melanoplus cinereus, Melanoplus madeleineae, Melanoplus sanguinipes, Melanoplus nigrescens group, Melanoplus tunicae, Melanoplus walshi, Melanoplus femurrubrum, Melanoplus dawsoni, Melanoplus sp., Melanoplus fasciatus, Melanoplus bowditchi, Melanoplus stonei, Melanoplus bivittatus, Melanoplus huroni, Melanoplus thomasi, Melanoplus borealis, Melanoplus packardii","4","4","4","108","0","0","99","449","0","450","36.39","0"
"AGAW1","11W","INS","BOLD:AAF9306","Arthropoda","Insecta","Diptera","Psychodidae","Psychodinae","Psychoda","Psychoda alternata","4","4","4","108","0","0","98.21","446.75","0","447","31.43","0"
"AGAW1","11W","INS","BOLD:AAI8935","Arthropoda","Insecta","Coleoptera","Latridiidae","Corticariinae","Cortinicara","Cortinicara gibbosa","4","4","4","108","0","0","99.16","446.5","0","450","33.83","0"
"AGAW1","11W","INS","BOLD:AAP8816","Arthropoda","Insecta","Diptera","Cecidomyiidae","Cecidomyiinae","Sitodiplosis","Sitodiplosis sp. BOLD:AAP8816","4","4","4","108","0","0","99.67","441","0","441.5","26.38","0"
"AGAW1","11W","INS","BOLD:ABV1237","Arthropoda","Insecta","Diptera","Chironomidae","Chironominae","Chironomus","","4","4","4","108","0","0","99.44","449.5","0","450","35.28","0"
"AGAW1","11W","INS","BOLD:ABV1765","Arthropoda","Insecta","Diptera","Limoniidae","Chioneinae","Erioptera","Erioptera viridula, Erioptera villosa","3","3","3","116","0","0","98.66","426.67","0","429.67","33.83","0"
"AGAW1","11W","INS","BOLD:ACM8528","Arthropoda","Insecta","Diptera","Limoniidae","Chioneinae","Erioptera","Erioptera chlorophylla","3","3","3","116","0","0","97.98","446.67","0","448.67","32.69","0"
"AGAW1","11W","INS","BOLD:AAH2306","Arthropoda","Insecta","Diptera","Asilidae","Asilinae","Efferia","Efferia aestuans","3","3","3","116","0","0","99.29","426.67","0","427.33","37.96","0"
"AGAW1","11W","INS","BOLD:AAG5453","Arthropoda","Insecta","Diptera","Chironomidae","Chironominae","Chironomus","Chironomus bifurcatus","3","3","3","116","0","0","98.12","370","0","433.33","35.88","0"
"AGAW1","11W","INS","BOLD:AAA7631","Arthropoda","Insecta","Hemiptera","Miridae","Mirinae","Adelphocoris","Adelphocoris lineolatus","3","3","3","116","0","0","99.4","448.33","0","450","35.7","0"
"AGAW1","11W","INS","BOLD:AAV5573","Arthropoda","Insecta","Diptera","Cecidomyiidae","","","","3","3","3","116","0","0","98.92","432.67","0","432.67","27.47","0"
"AGAW1","11W","INS","BOLD:AAU6966","Arthropoda","Insecta","Coleoptera","Throscidae","Throscinae","Trixagus","Trixagus chevrolati","3","3","3","116","0","0","99.75","397.67","0","422.67","39.7","0"
"AGAW1","11W","INS","BOLD:AAA3867","Arthropoda","Insecta","Lepidoptera","Noctuidae","Noctuinae","Apamea","Apamea amputatrix","3","3","3","116","0","0","98.96","446.33","0","450","33.85","0"
"AGAW1","11W","INS","BOLD:AAP2527","Arthropoda","Insecta","Diptera","Mycetophilidae","Mycetophilinae","Brevicornu","Brevicornu fissicauda","3","2","2","116","33.33","0","99.33","450","0","450","37.26","0"
"AGAW1","11W","INS","BOLD:AAG6743","Arthropoda","Insecta","Diptera","Sarcophagidae","Sarcophaginae","Sarcophaga","Sarcophaga sp. 1 MG, Sarcophaga subvicina","3","3","3","116","0","0","99.53","418","0","420","34.63","0"
"AGAW1","11W","INS","BOLD:AAB0651","Arthropoda","Insecta","Lepidoptera","Gelechiidae","Dichomeridinae","Dichomeris","Dichomeris leuconotella, Dichomeris gnoma","3","3","3","116","0","0","98.74","448","0","450","33.33","0"
"AGAW1","11W","INS","BOLD:AAN6433","Arthropoda","Insecta","Diptera","Sciaridae","","Corynoptera","Corynoptera deserta","3","3","3","116","0","0","99.48","447.33","0","450","30.59","0"
"AGAW1","11W","INS","BOLD:ACZ1491","Arthropoda","Insecta","Diptera","Chironomidae","","","","3","3","3","116","0","0","97.1","448","0","450","33.19","0"
"AGAW1","11W","INS","BOLD:AAB5640","Arthropoda","Insecta","Coleoptera","Coccinellidae","Coccinellinae","Harmonia","Harmonia axyridis","3","3","3","116","0","0","99.26","427.67","0","429.33","31.86","0"
"AGAW1","11W","INS","BOLD:AAA6759","Arthropoda","Insecta","Lepidoptera","Hesperiidae","Hesperiinae","Thymelicus","Thymelicus lineola","2","2","2","130","0","0","99.66","445.5","0","450","29.22","0"
"AGAW1","11W","INS","BOLD:AAH4135","Arthropoda","Insecta","Diptera","Chloropidae","Chloropinae","Thaumatomyia","Thaumatomyia glabra, Thaumatomyia grata, Thaumatomyia sp. 1 TAW","2","2","2","130","0","0","99.22","448.5","0","450","32.56","0"
"AGAW1","11W","INS","BOLD:ACF4734","Arthropoda","Insecta","Diptera","Syrphidae","Syrphinae","Platycheirus","Platycheirus hyperboreus","2","2","2","130","0","0","98.78","449","0","450","31.67","0"
"AGAW1","11W","INS","BOLD:AAA7067","Arthropoda","Insecta","Diptera","Culicidae","Culicinae","Aedes, Aedimorphus","Aedimorphus vexans, Aedes vexans","2","2","2","130","0","0","99.47","380.5","0","381.5","32.13","0"
"AGAW1","11W","INS","BOLD:AAA2518","Arthropoda","Insecta","Lepidoptera","Crambidae","Crambinae","Chrysoteuchia","Chrysoteuchia topiaria, Chrysoteuchia topiarius","2","2","2","130","0","0","98.66","448","0","450","35.78","0"
"AGAW1","11W","INS","BOLD:AAH2295","Arthropoda","Insecta","Diptera","Asilidae","Leptogastrinae","Leptogaster","Leptogaster flavipes","2","2","2","130","0","0","99.27","413","0","450","35.11","0"
"AGAW1","11W","INS","BOLD:AAL1297","Arthropoda","Insecta","Coleoptera","Cantharidae","Cantharinae","Rhagonycha","Rhagonycha fulva","2","2","2","130","0","0","98.04","415.5","0","420","33.32","0"
"AGAW1","11W","INS","BOLD:ACV2677","Arthropoda","Insecta","Diptera","Cecidomyiidae","","","","2","2","2","130","0","0","97.55","449.5","0","450","30.11","0"
"AGAW1","11W","INS","BOLD:AAH3914","Arthropoda","Insecta","Diptera","Sciaridae","","Sciara","Sciara hamatilis, Sciara humeralis","2","2","2","130","0","0","99.2","388","0","389","29.44","0"
"AGAW1","11W","INS","BOLD:AAA2632","Arthropoda","Insecta","Lepidoptera","Noctuidae","Noctuinae","Noctua","Noctua atlantica, Noctua pronuba","2","2","2","130","0","0","99.33","448","0","450","30.89","0"
"AGAW1","11W","INS","BOLD:AAA6822","Arthropoda","Insecta","Orthoptera","Acrididae","Gomphocerinae","Chorthippus","Chorthippus curtipennis","2","2","2","130","0","0","99.33","449","0","450","34.78","0"
"AGAW1","11W","INS","BOLD:ACE7444","Arthropoda","Insecta","Hemiptera","Miridae","Mirinae, Phylinae","Adelphocoris, Megalocoleus","Adelphocoris lineolatus, Megalocoleus molliculus","2","2","2","130","0","0","99.22","439","0","441","35.05","0"
"AGAW1","11W","INS","BOLD:ABY7666","Arthropoda","Insecta","Diptera","Culicidae","Culicinae","Culex","Culex territans","2","2","2","130","0","0","98.78","449.5","0","450","33.56","0"
"AGAW1","11W","INS","BOLD:AAA6337","Arthropoda","Insecta","Lepidoptera","Noctuidae","Plusiinae","Anagrapha","Anagrapha falcifera","2","2","2","130","0","0","99.89","448.5","0","450","32.67","0"
"AGAW1","11W","INS","BOLD:AAH0256","Arthropoda","Insecta","Coleoptera","Latridiidae","Corticariinae","Corticaria, Corticarina","","2","2","2","130","0","0","98.29","399.5","0","401.5","31.27","0"
"AGAW1","11W","INS","BOLD:AAA7374","Arthropoda","Insecta","Diptera","Syrphidae","Syrphinae","Sphaerophoria","Sphaerophoria bankowskae, Sphaerophoria infuscata, Sphaerophoria nr. philanthus, Sphaerophoria batava, Sphaerophoria sp., Sphaerophoria rueppellii, Sphaerophoria interrupta-group, Sphaerophoria sp. MA-1, Sphaerophoria cleoae, Sphaerophoria asymmetrica, Sphaerophoria fatarum, Sphaerophoria bengalensis, Sphaerophoria bifurcata, Sphaerophoria taeniata, Sphaerophoria interrupta, Sphaerophoria contigua, Sphaerophoria kaa, Sphaerophoria laurae, Sphaerophoria chongjini, Sphaerophoria boreoalpina, Sphaerophoria sp. BOLDAAA7374, Sphaerophoria macrogaster, Sphaerophoria sulphuripes, Sphaerophoria brevipilosa, Sphaerophoria philanthus, Sphaerophoria pyrrhina, Sphaerophoria abbreviata, Sphaerophoria scripta, Sphaerophoria potentillae, Sphaerophoria longipilosa, Sphaerophoria virgata","2","2","2","130","0","0","98.77","447","0","450","31.22","0"
"AGAW1","11W","INS","BOLD:ABW3687","Arthropoda","Insecta","Diptera","Lauxaniidae","Homoneurinae","Homoneura","Homoneura bispina","1","1","1","146","0","0","98.44","449","0","450","35.11","0"
"AGAW1","11W","INS","BOLD:AAM6655","Arthropoda","Insecta","Diptera","Hybotidae","Tachydromiinae","Platypalpus","Platypalpus stabilis","1","1","1","146","0","0","99.78","450","0","450","35.33","0"
"AGAW1","11W","INS","BOLD:ABX4079","Arthropoda","Insecta","Diptera","Cecidomyiidae","","","","1","1","1","146","0","0","99.78","450","0","450","28.67","0"
"AGAW1","11W","INS","BOLD:AAA3836","Arthropoda","Insecta","Lepidoptera","Noctuidae","Plusiinae","Autographa","Autographa precationis","1","1","1","146","0","0","99.78","450","0","450","32.44","0"
"AGAW1","11W","INS","BOLD:AAF3338","Arthropoda","Insecta","Lepidoptera","Noctuidae","Noctuinae","Striacosta","Striacosta albicosta","1","1","1","146","0","0","98.67","450","0","450","34","0"
"AGAW1","11W","INS","BOLD:ABA1215","Arthropoda","Insecta","Diptera","Sciaridae","","Lycoriella","Lycoriella sativae","1","1","1","146","0","0","100","450","0","450","36","0"
"AGAW1","11W","INS","BOLD:ACV1559","Arthropoda","Insecta","Diptera","Tipulidae","Tipulinae","Tipula","Tipula mallochi, Tipula sp., Tipula submaculata","1","1","1","146","0","0","100","447","0","450","33.78","0"
"AGAW1","11W","INS","BOLD:AAY6795","Arthropoda","Insecta","Hymenoptera","Ichneumonidae","Mesochorinae","Mesochorus","Mesochorus americanus","1","1","1","146","0","0","99.1","444","0","450","27.56","0"
"AGAW1","11W","INS","BOLD:AAG2717","Arthropoda","Insecta","Orthoptera","Tettigoniidae","Phaneropterinae","Scudderia","Scudderia curvicauda, Scudderia pistillata","1","1","1","146","0","0","99.33","445","0","450","39.11","0"
"AGAW1","11W","INS","BOLD:AAM6781","Arthropoda","Insecta","Diptera","Dolichopodidae","Neurigoninae","Neurigona","Neurigona tenuis","1","1","1","146","0","0","99.03","414","0","417","35.01","0"
"AGAW1","11W","INS","BOLD:AAD0952","Arthropoda","Insecta","Lepidoptera","Pyralidae","Chrysauginae","Galasa","Galasa nigrinodis, Galasa nigripunctalis","1","1","1","146","0","0","99.03","414","0","415","31.33","0"
"AGAW1","11W","INS","BOLD:AAB2076","Arthropoda","Insecta","Lepidoptera","Tortricidae","Olethreutinae","Ancylis","Ancylis metamelana, Ancylis sp.","1","1","1","146","0","0","98.35","364","0","366","30.6","0"
"AGAW1","11W","INS","BOLD:AAE7309","Arthropoda","Insecta","Hemiptera","Aphididae","Aphidinae","Aphis","Aphis sambuci, Aphis horii","1","1","1","146","0","0","100","450","0","450","28","0"
"AGAW1","11W","INS","BOLD:ADF7493","Arthropoda","Insecta","Diptera","Phoridae","Metopininae","Megaselia","","1","1","1","146","0","0","99.74","390","0","422","31.52","0"
"AGAW1","11W","INS","BOLD:AAN5230","Arthropoda","Insecta","Diptera","Cecidomyiidae","","","","1","1","1","146","0","0","99.54","435","0","435","26.67","0"
"AGAW1","11W","INS","BOLD:AAU6592","Arthropoda","Insecta","Diptera","Ceratopogonidae","Dasyheleinae","Dasyhelea","Dasyhelea arenivaga","1","1","1","146","0","0","99.01","405","0","414","33.82","0"
"AGAW1","11W","INS","BOLD:AAG3236","Arthropoda","Insecta","Diptera","Phoridae","Phorinae","Conicera, Diplonevra","Diplonevra nitidula","1","1","1","146","0","0","100","448","0","450","31.78","0"
"AGAW1","11W","INS","BOLD:AAH4276","Arthropoda","Insecta","Lepidoptera","Gelechiidae","Gelechiinae","Bryotropha","Bryotropha hodgesi, Bryotropha altitudophila","1","1","1","146","0","0","98.7","384","0","385","32.99","0"
"AGAW1","11W","INS","BOLD:ACE5981","","","","","","","","1","1","1","146","0","0","98.67","450","0","450","36.67","0"
"AGAW1","11W","INS","BOLD:AAO0694","Arthropoda","Insecta","Coleoptera","Chrysomelidae","Criocerinae","Oulema","Oulema duftschmidi, Oulema cfr. melanopus","1","1","1","146","0","0","97.41","425","0","426","36.38","0"
"AGAW1","11W","INS","BOLD:AAG3154","Arthropoda","Insecta","Diptera","Psychodidae","Psychodinae","Psychoda","","1","1","1","146","0","0","97.08","445","0","450","32","0"
"AGAW1","11W","INS","BOLD:AAG8882","Arthropoda","Insecta","Hemiptera","Pentatomidae","Pentatominae","Thyanta","Thyanta pallidovirens, Thyanta custator, Thyanta accerra","1","1","1","146","0","0","98.67","450","0","450","36.44","0"
"AGAW1","11W","INS","BOLD:ABA5202","Arthropoda","Insecta","Diptera","Limoniidae","Limnophilinae","Pseudolimnophila","Pseudolimnophila luteipennis","1","1","1","146","0","0","98.44","450","0","450","34","0"
"AGAW1","11W","INS","BOLD:AAN5205","Arthropoda","Insecta","Diptera","Cecidomyiidae","","","CecidInt42 sp. BOLD:AAN5205","1","1","1","146","0","0","99.33","446","0","450","30.89","0"
"AGAW1","11W","INS","BOLD:AAG2355","Arthropoda","Insecta","Diptera","Tachinidae","Tachininae","Strongygaster","Strongygaster triangulifera, Strongygaster sp.","1","1","1","146","0","0","98.51","403","0","403","30.77","0"
"AGAW1","11W","INS","BOLD:ACE0554","Arthropoda","Insecta","Diptera","Phoridae","Metopininae","Megaselia","","1","1","1","146","0","0","99.52","413","0","415","30.6","0"
"AGAW1","11W","INS","BOLD:AAB5609","Arthropoda","Insecta","Diptera","Muscidae","Coenosiinae","Coenosia","Coenosia sp.BOLD:AAB5609, Coenosia tigrina","1","1","1","146","0","0","97.9","428","0","433","32.79","0"
"AGAW1","11W","INS","BOLD:AAA2482","Arthropoda","Insecta","Lepidoptera","Noctuidae","Noctuinae","Mythimna","Mythimna unipuncta","1","1","1","146","0","0","99.75","394","0","398","29.9","0"
"AGAW1","11W","INS","BOLD:ACG5862","Arthropoda","Insecta","Diptera","Agromyzidae","","","","1","1","1","146","0","0","98.89","449","0","450","34","0"
"AGAW1","11W","INS","BOLD:AAN8202","Arthropoda","Insecta","Hymenoptera","Braconidae","Alysiinae","","","1","1","1","146","0","0","100","450","0","450","26.44","0"
"AGAW1","11W","INS","BOLD:AAY6374","Arthropoda","Insecta","Diptera","Cecidomyiidae","","","","1","1","1","146","0","0","99.55","448","0","450","28.67","0"
"AGAW1","11W","INS","BOLD:AAH2293","Arthropoda","Insecta","Diptera","Asilidae","Stenopogoninae","Dioctria, Eudioctria","Dioctria hyalipennis, Dioctria baumhaueri, Eudioctria propinqua","1","1","1","146","0","0","97.99","448","0","450","30","0"
"AGAW1","11W","INS","BOLD:AAG6756","Arthropoda","Insecta","Diptera","Sarcophagidae","Sarcophaginae","Boettcheria","Boettcheria cimbicis","1","1","1","146","0","0","98.77","407","0","409","34.47","0"
"AGAW1","11W","INS","BOLD:AAA9251","Arthropoda","Insecta","Lepidoptera","Gelechiidae","Gelechiinae","Scrobipalpa","Scrobipalpa atriplicella, Scrobipalpa nr. atriplicella sp. 1","1","1","1","146","0","0","99.29","423","0","428","33.64","0"
"AGAW1","11W","INS","BOLD:AAN5260","Arthropoda","Insecta","Diptera","Cecidomyiidae","","","","1","1","1","146","0","0","99.01","403","0","406","25.62","0"
"AGAW1","11W","INS","BOLD:ACI1304","Arthropoda","Insecta","Diptera","Muscidae","Muscinae","Stomoxys","Stomoxys indicus","1","1","1","146","0","0","97.75","445","0","450","32.44","0"
"AGAW1","11W","INS","BOLD:AAA4893","Arthropoda","Insecta","Lepidoptera","Crambidae","Crambinae","Microcrambus","Microcrambus elegans","1","1","1","146","0","0","98.74","398","0","398","31.91","0"
"AGAW1","11W","INS","BOLD:ACX6055","Arthropoda","Insecta","Diptera","Phoridae","Metopininae","Megaselia","Megaselia sp. AK-2010-2, Megaselia sp. BOLD:ACX6055","1","1","1","146","0","0","98.51","404","0","408","32.35","0"
"AGAW1","11W","INS","BOLD:AAU6533","Arthropoda","Insecta","Diptera","Phoridae","Metopininae","Megaselia","","1","1","1","146","0","0","97.47","356","0","450","32.89","0"
"AGAW1","11W","INS","BOLD:AAG7284","Arthropoda","Insecta","Diptera","Sphaeroceridae","Limosininae","Pullimosina","Pullimosina pullula","1","1","1","146","0","0","99.02","408","0","410","32.68","0"
"AGAW1","11W","INS","BOLD:ACE9865","Arthropoda","Insecta","Lepidoptera","Noctuidae","Noctuinae","Feltia","Feltia subgothica, Feltia herilis, Feltia tricosa","1","1","1","146","0","0","100","448","0","450","30.67","0"
"AGAW1","11W","INS","BOLD:AAI6258","Arthropoda","Insecta","Lepidoptera","Gelechiidae","Dichomeridinae","Dichomeris","Dichomeris mercatrix","1","1","1","146","0","0","98.22","450","0","450","33.56","0"
"AGAW1","11W","INS","BOLD:ACF6498","Arthropoda","Insecta","Hymenoptera","Platygastridae","","","","1","1","1","146","0","0","99.77","436","0","450","25.78","0"
"AGAW1","11W","INS","BOLD:AAF1995","Arthropoda","Insecta","Hemiptera","Miridae","Phylinae","Plagiognathus, Parapsallus","Plagiognathus vitellinus, Parapsallus vitellinus","1","1","1","146","0","0","99.78","451","0","450","41.78","0"
"AGAW1","11W","INS","BOLD:AAE0844","Arthropoda","Insecta","Hemiptera","Pentatomidae","Pentatominae","Euschistus","Euschistus variolarius","1","1","1","146","0","0","99.48","387","0","388","36.08","0"
"AGAW1","11W","INS","BOLD:ACI5617","Arthropoda","Insecta","Diptera","Cecidomyiidae","","","","1","1","1","146","0","0","97.17","424","0","426","26.76","0"
"AGAW1","11W","INS","BOLD:AAG8822","Arthropoda","Insecta","Hemiptera","Cicadellidae","Eurymelinae","Idiocerus","Idiocerus snowi, Idiocerus raphus","1","1","1","146","0","0","97.88","425","0","428","34.81","0"
"AGAW1","11W","INS","BOLD:AAA3797","Arthropoda","Insecta","Hymenoptera","Megachilidae","Megachilinae","Megachile","Megachile sp. aff rotundata, Megachile cf. rotundata, Megachile rotundata","1","1","1","146","0","0","100","449","0","450","24","0"
"AGAW1","11W","INS","BOLD:AAH4171","Arthropoda","Insecta","Diptera","Chloropidae","Oscinellinae","Apallates","Apallates montanus, Apallates particeps","1","1","1","146","0","0","99.74","391","0","394","32.74","0"
"AGAW1","11W","INS","BOLD:AAF1729","Arthropoda","Insecta","Psocodea","Lachesillidae","Lachesillinae","Lachesilla","Lachesilla pedicularia","1","1","1","146","0","0","97.99","448","0","450","36.22","0"
"AGAW1","11W","INS","BOLD:ACE5778","Arthropoda","Insecta","Diptera","Dolichopodidae","","","","1","1","1","146","0","0","98.89","450","0","450","39.11","0"
"AGAW1","11W","INS","BOLD:AAN5342","Arthropoda","Insecta","Diptera","Chironomidae","Orthocladiinae","Heleniella","","1","1","1","146","0","0","99.55","449","0","450","35.33","0"
"AGAW1","11W","INS","BOLD:ACD8563","Arthropoda","Insecta","Diptera","Cecidomyiidae","","","","1","1","1","146","0","0","97.11","450","0","450","25.78","0"
"AGAW1","11W","INS","BOLD:AAA9246","Arthropoda","Insecta","Lepidoptera","Tortricidae","Tortricinae","Clepsis","Clepsis clemensiana","1","1","1","146","0","0","98.59","354","0","356","30.06","0"
"AGAW1","11W","INS","BOLD:AAD8027","Arthropoda","Insecta","Diptera","Culicidae","Culicinae","Aedes","Aedes fitchii","1","1","1","146","0","0","99.25","402","0","407","34.15","0"
"AGAW1","11W","INS","BOLD:AAN5186","Arthropoda","Insecta","Diptera","Cecidomyiidae","","","","1","1","1","146","0","0","98.47","326","0","382","24.87","0"
"AGAW1","11W","INS","BOLD:AAD0484","Arthropoda","Insecta","Diptera","Chironomidae","","","","1","1","1","146","0","0","98.66","449","0","450","34.67","0"
"AGAW1","11W","INS","BOLD:AAA5803","Arthropoda","Insecta","Hemiptera","Miridae","Mirinae","Lygus","Lygus lineolaris","1","1","1","146","0","0","97.32","447","0","450","34.89","0"
"AGAW1","11W","NAR","BOLD:AAL1512","Mollusca","Gastropoda","Stylommatophora","Agriolimacidae",NA,"Deroceras","Deroceras laeve","1","1","1","1","0","0","97.57","370","0","370","33.51","0"
"AGAW1","11W","NIN","BOLD:AAA7242","Arthropoda","Collembola","Entomobryomorpha","Entomobryidae","Seirinae, Entomobryinae","Entomobrya, Lepidosira","Entomobrya multifasciata, Lepidosira sp.","415","407","407","1","1.93","0","99.09","416.17","0","419.57","39.29","0"
"AGAW1","11W","NIN","BOLD:AAU6252","Arthropoda","Arachnida","Trombidiformes","Bdellidae","","Cyta","","305","253","253","2","17.05","0","98.47","435.55","0","436.83","31.5","0"
"AGAW1","11W","NIN","BOLD:AAH6630","Arthropoda","Collembola","Entomobryomorpha","Entomobryidae","Entomobryinae","Entomobrya","Entomobrya unostrigata","170","155","155","3","8.82","0","99.42","432.61","0","433.83","38.98","0"
"AGAW1","11W","NIN","BOLD:ABW2639","Arthropoda","Arachnida","Trombidiformes","Anystidae","","","","142","123","123","4","13.38","0","99.54","447.2","0","446","31.48","0"
"AGAW1","11W","NIN","BOLD:ACV5609","","","","","","","","114","92","92","5","19.3","0","99.03","440.83","0","442.08","38.67","0"
"AGAW1","11W","NIN","BOLD:AAA6381","Arthropoda","Arachnida","Araneae","Tetragnathidae","","Tetragnatha","Tetragnatha laboriosa","50","50","50","6","0","0","99.2","426.76","0","429.04","39.37","0"
"AGAW1","11W","NIN","BOLD:ACY2550","Arthropoda","Arachnida","Trombidiformes","Bdellidae","","Cyta","","43","43","43","7","0","0","98.56","373.49","0","388.58","31.18","0"
"AGAW1","11W","NIN","BOLD:AAB7914","Arthropoda","Collembola","Symphypleona","Bourletiellidae","","Bourletiella","","25","25","25","8","0","0","98.75","439.4","0","441.4","38.48","0"
"AGAW1","11W","NIN","BOLD:ACZ1295","Arthropoda","Arachnida","Trombidiformes","Anystidae","","","","23","23","23","9","0","0","99.44","373.09","0","380.96","31.24","0"
"AGAW1","11W","NIN","BOLD:AAL8179","Arthropoda","Arachnida","Trombidiformes","Tetranychidae","","","","7","7","7","10","0","0","97.69","438","0","438.71","23.47","0"
"AGAW1","11W","NIN","BOLD:AAM7961","Arthropoda","Arachnida","Trombidiformes","Anystidae","","","","5","5","5","11","0","0","99.41","437.8","0","440.6","31.14","0"
"AGAW1","11W","NIN","BOLD:ACX8969","Arthropoda","Arachnida","Trombidiformes","Eupodidae","","","","2","2","2","12","0","0","98.18","435","0","439","28.01","0"
"AGAW1","11W","NIN","BOLD:AAE3860","Arthropoda","Arachnida","Araneae","Linyphiidae","","Agyneta","Agyneta fabra, Agyneta unimaculata","2","2","2","12","0","0","98.66","447.5","0","450","32.56","0"
"AGAW1","11W","NIN","BOLD:AAB5726","Arthropoda","Arachnida","Araneae","Agelenidae","","Agelenopsis","Agelenopsis pennsylvanica, Agelenopsis emertoni, Agelenopsis potteri, Agelenopsis actuosa, Agelenopsis kastoni","2","2","2","12","0","0","99.67","449.5","0","450","33.11","0"
"AGAW1","11W","NIN","BOLD:AAI4346","Arthropoda","Arachnida","Opiliones","Phalangiidae","","Paroligolophus, Phalangium","Paroligolophus agrestis, Phalangium opilio","1","1","1","15","0","0","99.78","448","0","450","36.22","0"
"AGAW1","11W","NIN","BOLD:AAA8820","Arthropoda","Collembola","Entomobryomorpha","Entomobryidae","Lepidocyrtinae","Lepidocyrtus","Lepidocyrtus cyaneus","1","1","1","15","0","0","98.39","373","0","375","43.73","0"
"AGAW1","11W","NIN","BOLD:AAF9237","Arthropoda","Arachnida","Trombidiformes","Erythraeidae","","Balaustium","","1","1","1","15","0","0","99.78","446","0","450","37.33","0"
"AGAW1","11W","NIN","BOLD:ACC0536","Arthropoda","Arachnida","Mesostigmata","Blattisociidae","","","","1","1","1","15","0","0","99.28","416","0","417","35.73","0"
"AGAW2","02E","BAC","TAX:91391","Proteobacteria","Alphaproteobacteria","Rickettsiales","",NA,"","","9","8","8","1","11.11","0","97.58","266","0","450","37.75","0"
"AGAW2","02E","BAC","TAX:175768","Proteobacteria","Alphaproteobacteria","Rickettsiales","Rickettsiaceae",NA,"Wolbachia","Wolbachia pipientis","3","3","3","2","0","0","98.74","448.67","0","450","39.48","0"
"AGAW2","02E","BAC","TAX:91393","Proteobacteria","Alphaproteobacteria","Rickettsiales","Rickettsiaceae",NA,"Wolbachia","","1","1","1","3","0","0","99.74","391","0","417","38.61","0"
"AGAW2","02E","BAC","TAX:376688","Proteobacteria","Alphaproteobacteria","Rickettsiales","Rickettsiaceae",NA,"Wolbachia","Wolbachia sp. elachBioLep01 BioLep373","1","1","1","3","0","0","99.33","450","0","450","40.22","0"
"AGAW2","02E","BAC","TAX:375762","Proteobacteria","Alphaproteobacteria","Rickettsiales","Rickettsiaceae",NA,"Wolbachia","Wolbachia sp. Myrmicinae_ACGAN556-09","1","1","1","3","0","0","99.11","449","0","450","40","0"
"AGAW2","02E","CHO","no match","no match","no match","no match","no match","no match","no match","no match","no match","no match","no match","no match","no match","no match","no match","no match","no match","no match","no match","no match"
"AGAW2","02E","INS","BOLD:AAG2511","Arthropoda","Insecta","Diptera","Anthomyiidae","Anthomyiinae","Delia","Delia platura","66417","43729","43729","1","34.16","0","99.45","443.27","0","445.5","33.19","0"
"AGAW2","02E","INS","BOLD:AAB2384","Arthropoda","Insecta","Diptera","Syrphidae","Syrphinae","Eupeodes","Eupeodes duseki, Eupeodes volucris, Eupeodes nielseni, Eupeodes americanus, Eupeodes flukei, Eupeodes vockerothi, Eupeodes lucasi, Eupeodes montivagus, Eupeodes rufipunctatus, Eupeodes rojasi, Eupeodes goeldlini, Eupeodes luniger, Eupeodes punctifer, Eupeodes bucculatus, Eupeodes perplexus, Eupeodes abiskoensis, Eupeodes corollae, Eupeodes pomus, Eupeodes lundbecki, Eupeodes nitens, Eupeodes sp. BOLDAAB2384, Eupeodes curtus, Eupeodes latifasciatus","11519","8970","8970","2","22.13","0","98.79","430.65","0","432.71","30.96","0"
"AGAW2","02E","INS","BOLD:AAB1024","Arthropoda","Insecta","Lepidoptera","Noctuidae","Noctuinae","Pseudohermonassa","Pseudohermonassa bicarnea","8341","6964","6964","3","16.51","0","99.33","435.37","0","436.9","33.63","0"
"AGAW2","02E","INS","BOLD:AAD4355","Arthropoda","Insecta","Diptera","Culicidae","Culicinae","Aedes","Aedes provocans","6526","5485","5485","4","15.95","0","99.02","434.77","0","437.13","31.74","0"
"AGAW2","02E","INS","BOLD:AAA3453","Arthropoda","Insecta","Diptera","Anthomyiidae","Anthomyiinae","Delia","","6451","4912","4912","5","23.86","0","99.33","443.64","0","445.47","34.72","0"
"AGAW2","02E","INS","BOLD:AAB2644","Arthropoda","Insecta","Diptera","Chironomidae","Orthocladiinae","Orthocladius","Orthocladius lapponicus, Orthocladius dorenus, Orthocladius mallochi","4003","3154","3154","6","21.21","0","99.21","443.12","0","445.04","32.5","0"
"AGAW2","02E","INS","BOLD:AAE0844","Arthropoda","Insecta","Hemiptera","Pentatomidae","Pentatominae","Euschistus","Euschistus variolarius","2439","1819","1819","7","25.42","0","99.03","439.04","0","439.07","36.27","0"
"AGAW2","02E","INS","BOLD:AAN5355","Arthropoda","Insecta","Diptera","Chironomidae","Orthocladiinae","Smittia","Smittia cf. stercoraria, Smittia stercoraria","1876","1700","1700","8","9.38","0","99.04","427.94","0","443.56","36.24","0"
"AGAW2","02E","INS","BOLD:AAD0853","Arthropoda","Insecta","Diptera","Scathophagidae","Scathophaginae","Scathophaga","Scathophaga lutaria, Scathophaga hiemalis, Scathophaga furcata, Scathophaga stercoraria, Scathophaga inquinata, Scathophaga suilla","727","598","598","9","17.74","0","99.12","443.7","0","445.34","31.79","0"
"AGAW2","02E","INS","BOLD:AAN5358","Arthropoda","Insecta","Diptera","Chironomidae","Orthocladiinae","Smittia","Smittia stercoraria, Smittia sp. 22ES","571","508","508","10","11.03","0","99.25","442.56","0","444.46","37.27","0"
"AGAW2","02E","INS","BOLD:AAB1098","Arthropoda","Insecta","Diptera","Culicidae","Culicinae","Ochlerotatus, Aedes","Aedes excrucians, Aedes cantans, Aedes stimulans, Aedes euedes, Aedes riparius, Aedes annulipes, Ochlerotatus cantans, Aedes fitchii, Aedes sp., Ochlerotatus annulipes, Ochlerotatus excrucians","546","506","506","11","7.33","0","98.64","431.52","0","432.95","35.28","0"
"AGAW2","02E","INS","BOLD:AAI2661","Arthropoda","Insecta","Diptera","Chironomidae","Orthocladiinae","Paraphaenocladius","Paraphaenocladius impensus","489","402","402","12","17.79","0","98.76","423.59","0","442.78","35.6","0"
"AGAW2","02E","INS","BOLD:AAC4197","Arthropoda","Insecta","Diptera","Chironomidae","Orthocladiinae","Paraphaenocladius","Paraphaenocladius impensus","237","221","221","13","6.75","0","99.09","436.14","0","439.96","34.99","0"
"AGAW2","02E","INS","BOLD:AAV1295","Arthropoda","Insecta","Diptera","Sciaridae","","Bradysia","Bradysia difformis, Bradysia impatiens","178","150","150","14","15.73","0","99.42","416.91","0","440.3","31.55","0"
"AGAW2","02E","INS","BOLD:AAD1185","Arthropoda","Insecta","Lepidoptera","Gracillariidae","Lithocolletinae","Phyllonorycter","Phyllonorycter blancardella","159","137","137","15","13.84","0","99.32","439.42","0","440.6","34.12","0"
"AGAW2","02E","INS","BOLD:AAA4335","Arthropoda","Insecta","Lepidoptera","Depressariidae","Depressariinae","Semioscopis","Semioscopis inornata","154","131","131","16","14.94","0","99.49","437.21","0","444.34","30.84","0"
"AGAW2","02E","INS","BOLD:AAK5928","Arthropoda","Insecta","Coleoptera","Chrysomelidae","Criocerinae","Oulema","Oulema melanopus","137","112","112","17","18.25","0","99.28","447.17","0","448.09","36.3","0"
"AGAW2","02E","INS","BOLD:ACB1079","Arthropoda","Insecta","Diptera","Cecidomyiidae","Micromyinae","Campylomyza","Campylomyza dilatata, CecidInt44 sp. BOLD:ACB1079","130","115","115","18","11.54","0","98.29","443.11","0","446.18","28.35","0"
"AGAW2","02E","INS","BOLD:AAN5267","Arthropoda","Insecta","Diptera","Cecidomyiidae","","","","82","69","69","19","15.85","0","99.44","447.52","0","448.51","30.63","0"
"AGAW2","02E","INS","BOLD:ACK1651","Arthropoda","Insecta","Diptera","Cecidomyiidae","","","","41","39","39","20","4.88","0","98.92","439.71","0","441.27","26.99","0"
"AGAW2","02E","INS","BOLD:ACW4261","Arthropoda","Insecta","Diptera","Chironomidae","Orthocladiinae","","","40","40","40","21","0","0","98.3","404.18","0","417.5","31.62","0"
"AGAW2","02E","INS","BOLD:AAO8538","Arthropoda","Insecta","Diptera","Psychodidae","Psychodinae","Psychoda","","38","37","37","22","2.63","0","99.47","446","0","447.58","32.88","0"
"AGAW2","02E","INS","BOLD:AAB9140","Arthropoda","Insecta","Diptera","Calliphoridae","Chrysomyinae","Phormia","Phormia regina","36","31","31","23","13.89","0","99.11","439.44","0","444.33","33.01","0"
"AGAW2","02E","INS","BOLD:AAG1000","Arthropoda","Insecta","Diptera","Chironomidae","Orthocladiinae","Orthocladius","Orthocladius carlatus","21","21","21","24","0","0","97.98","428.33","0","430.14","33.95","0"
"AGAW2","02E","INS","BOLD:ACG4648","Arthropoda","Insecta","Diptera","Cecidomyiidae","","","","20","20","20","25","0","0","99.48","384.95","0","426.4","34.38","0"
"AGAW2","02E","INS","BOLD:ADH4889","Arthropoda","Insecta","Diptera","","","","","15","15","15","26","0","0","98.94","447.27","0","448.27","35.35","0"
"AGAW2","02E","INS","BOLD:AAV5688","Arthropoda","Insecta","Diptera","Cecidomyiidae","","","","14","14","14","27","0","0","99.33","428.71","0","430","34.14","0"
"AGAW2","02E","INS","BOLD:ABU5520","Arthropoda","Insecta","Diptera","Sciaridae","","Corynoptera","","13","13","13","28","0","0","98.77","426.54","0","429.54","35.16","0"
"AGAW2","02E","INS","BOLD:AAY6412","Arthropoda","Insecta","Diptera","Sciaridae","","Bradysia","Bradysia urticae","11","11","11","29","0","0","99.39","448.27","0","450","32.32","0"
"AGAW2","02E","INS","BOLD:ACC7611","Arthropoda","Insecta","Diptera","Cecidomyiidae","","","","10","10","10","30","0","0","97.49","446.5","0","448.9","28.31","0"
"AGAW2","02E","INS","BOLD:AAM7855","Arthropoda","Insecta","Thysanoptera","Thripidae","Thripinae","Frankliniella","Frankliniella tritici","9","9","9","31","0","0","99.15","443.89","0","446.33","31.36","0"
"AGAW2","02E","INS","BOLD:AAB0868","Arthropoda","Insecta","Diptera","Calliphoridae","Calliphorinae","Cynomya","Cynomya cadaverina, Cynomya mortuorum","8","8","8","32","0","0","99.08","432","0","433","34.65","0"
"AGAW2","02E","INS","BOLD:AAN6447","Arthropoda","Insecta","Diptera","Sciaridae","","Corynoptera","Corynoptera perpusilla","6","4","4","33","33.33","0","99.7","449.83","0","450","36.48","0"
"AGAW2","02E","INS","BOLD:AAP5146","Arthropoda","Insecta","Diptera","Chironomidae","Orthocladiinae","Orthocladius","","5","5","5","34","0","0","97.88","431","0","431.4","33.34","0"
"AGAW2","02E","INS","BOLD:AAN8770","Arthropoda","Insecta","Diptera","Psychodidae","Psychodinae","Psychoda","Psychoda trinodulosa","4","4","4","35","0","0","98.94","448.25","0","450","33.22","0"
"AGAW2","02E","INS","BOLD:AAP6827","Arthropoda","Insecta","Diptera","Cecidomyiidae","","","","4","4","4","35","0","0","98.83","448.75","0","450","31.72","0"
"AGAW2","02E","INS","BOLD:AAH3920","Arthropoda","Insecta","Diptera","Sciaridae","","Dichopygina, Scatopsciara","","4","4","4","35","0","0","98.72","404.25","0","442.75","35.39","0"
"AGAW2","02E","INS","BOLD:AAM9252","Arthropoda","Insecta","Diptera","Sciaridae","","Bradysia, Leptosciarella","Bradysia vagans, Leptosciarella zanti","4","3","3","35","25","0","99.94","450","0","450","39.22","0"
"AGAW2","02E","INS","BOLD:ABU9891","Arthropoda","Insecta","Diptera","Muscidae","Phaoniinae","Phaonia","","3","3","3","39","0","0","99.44","431","0","435.33","32.6","0"
"AGAW2","02E","INS","BOLD:AAC9062","Arthropoda","Insecta","Diptera","Culicidae","Culicinae","Ochlerotatus, Aedes","Aedes fitchii, Ochlerotatus fitchii, Aedes grossbecki, Aedes ventrovittis","3","3","3","39","0","0","98.89","436","0","436.67","32.96","0"
"AGAW2","02E","INS","BOLD:AAA4222","Arthropoda","Insecta","Lepidoptera","Erebidae","Hypeninae","Hypena","Hypena scabra","3","3","3","39","0","0","98.63","383.33","0","387","32.35","0"
"AGAW2","02E","INS","BOLD:AAU8441","Arthropoda","Insecta","Hymenoptera","Ichneumonidae","Campopleginae","Meloboris","","2","2","2","42","0","0","99.78","449","0","450","25.67","0"
"AGAW2","02E","INS","BOLD:AAN5356","Arthropoda","Insecta","Diptera","Chironomidae","Orthocladiinae","Smittia","","2","2","2","42","0","0","98.22","450","0","450","34.89","0"
"AGAW2","02E","INS","BOLD:AAA9251","Arthropoda","Insecta","Lepidoptera","Gelechiidae","Gelechiinae","Scrobipalpa","Scrobipalpa atriplicella, Scrobipalpa nr. atriplicella sp. 1","2","2","2","42","0","0","98","449","0","450","33.78","0"
"AGAW2","02E","INS","BOLD:ACF2821","Arthropoda","Insecta","Diptera","Mycetophilidae","Mycetophilinae","Mycetophila","Mycetophila fungorum","2","2","2","42","0","0","98.77","448.5","0","450","35.22","0"
"AGAW2","02E","INS","BOLD:AAQ0582","Arthropoda","Insecta","Diptera","Anthomyiidae","Anthomyiinae","Lasiomma","","2","2","2","42","0","0","97.44","449.5","0","450","31.67","0"
"AGAW2","02E","INS","BOLD:AAN1738","Arthropoda","Insecta","Coleoptera","Staphylinidae","Oxytelinae","Anotylus","Anotylus tetracarinatus","1","1","1","47","0","0","99.78","450","0","450","34","0"
"AGAW2","02E","INS","BOLD:AAB2645","","","","","","","","1","1","1","47","0","0","98.31","415","0","420","32.14","0"
"AGAW2","02E","INS","BOLD:AAL2750","Arthropoda","Insecta","Diptera","Muscidae","Phaoniinae","Phaonia","Phaonia tuguriorum","1","1","1","47","0","0","98.08","416","0","416","31.49","0"
"AGAW2","02E","INS","BOLD:AAG9055","Arthropoda","Insecta","Hymenoptera","Vespidae","Vespinae","Vespula","Vespula germanica","1","1","1","47","0","0","97.73","396","0","399","30.83","0"
"AGAW2","02E","INS","BOLD:ACI2687","Arthropoda","Insecta","Diptera","Cecidomyiidae","","","","1","1","1","47","0","0","97.32","448","0","450","35.33","0"
"AGAW2","02E","INS","BOLD:AAG7276","Arthropoda","Insecta","Diptera","Sphaeroceridae","Limosininae","Leptocera","Leptocera erythrocera","1","1","1","47","0","0","98.22","449","0","450","33.33","0"
"AGAW2","02E","INS","BOLD:AAP2970","Arthropoda","Insecta","Diptera","Anthomyiidae","Anthomyiinae","Anthomyia","Anthomyia procellaris, Anthomyia pluvialis","1","1","1","47","0","0","99.78","449","0","450","33.33","0"
"AGAW2","02E","INS","BOLD:ABY4763","Arthropoda","Insecta","Hemiptera","Aphididae","Aphidinae","Uroleucon","Uroleucon taraxaci","1","1","1","47","0","0","99.55","446","0","450","25.78","0"
"AGAW2","02E","INS","BOLD:AAI4303","Arthropoda","Insecta","Diptera","Chironomidae","Chironominae","Chironomus","Chironomus melanescens","1","1","1","47","0","0","97.75","444","0","447","34.9","0"
"AGAW2","02E","INS","BOLD:AAH0256","Arthropoda","Insecta","Coleoptera","Latridiidae","Corticariinae","Corticaria, Corticarina","","1","1","1","47","0","0","99.31","437","0","450","31.78","0"
"AGAW2","02E","INS","BOLD:ABV9277","","","","","","","","1","1","1","47","0","0","98.89","450","0","450","28.67","0"
"AGAW2","02E","INS","BOLD:AAA3080","Arthropoda","Insecta","Trichoptera","Limnephilidae","Limnephilinae","Limnephilus","Limnephilus submonilifer","1","1","1","47","0","0","98.86","438","0","441","34.24","0"
"AGAW2","02E","INS","BOLD:ABW2407","Arthropoda","Insecta","Diptera","Anthomyiidae","Anthomyiinae","Hylemya","","1","1","1","47","0","0","97.78","450","0","450","32.89","0"
"AGAW2","02E","INS","BOLD:ACX6055","Arthropoda","Insecta","Diptera","Phoridae","Metopininae","Megaselia","Megaselia sp. AK-2010-2, Megaselia sp. BOLD:ACX6055","1","1","1","47","0","0","97.54","448","0","450","32.67","0"
"AGAW2","02E","INS","BOLD:AAM9013","Arthropoda","Insecta","Diptera","Keroplatidae","Macrocerinae","Macrocera","Macrocera pusilla","1","1","1","47","0","0","99.33","449","0","450","30.67","0"
"AGAW2","02E","INS","BOLD:AAA2955","Arthropoda","Insecta","Lepidoptera","Tortricidae","Tortricinae","Argyrotaenia","Argyrotaenia occultana, Argyrotaenia ljungiana, Argyrotaenia pinatubana, Argyrotaenia floridana, Argyrotaenia velutinana, Argyrotaenia tabulana, Argyrotaenia gogana","1","1","1","47","0","0","99.56","450","0","450","32.44","0"
"AGAW2","02E","INS","BOLD:AAG1742","Arthropoda","Insecta","Diptera","Muscidae","Phaoniinae","Helina","Helina rufitibia","1","1","1","47","0","0","99.77","436","0","436","30.96","0"
"AGAW2","02E","INS","BOLD:AAB4658","Arthropoda","Insecta","Diptera","Chironomidae","Chironominae","Chironomus","Chironomus dilutus","1","1","1","47","0","0","97.33","450","0","450","37.56","0"
"AGAW2","02E","INS","BOLD:AAG6743","Arthropoda","Insecta","Diptera","Sarcophagidae","Sarcophaginae","Sarcophaga","Sarcophaga sp. 1 MG, Sarcophaga subvicina","1","1","1","47","0","0","99.78","450","0","450","35.33","0"
"AGAW2","02E","INS","BOLD:ACR4394","Arthropoda","Insecta","Diptera","Anthomyiidae","Anthomyiinae","Delia","Delia florilega","1","1","1","47","0","0","98.19","443","0","450","33.56","0"
"AGAW2","02E","INS","BOLD:AAG2462","Arthropoda","Insecta","Diptera","Anthomyiidae","Anthomyiinae","Botanophila","Botanophila fugax","1","1","1","47","0","0","97.1","449","0","450","32","0"
"AGAW2","02E","INS","BOLD:AAG7306","Arthropoda","Insecta","Diptera","Sphaeroceridae","Limosininae","Minilimosina","Minilimosina vitripennis","1","1","1","47","0","0","97.85","418","0","423","31.68","0"
"AGAW2","02E","INS","BOLD:AAP9022","Arthropoda","Insecta","Diptera","Cecidomyiidae","","","","1","1","1","47","0","0","99.33","450","0","450","28.44","0"
"AGAW2","02E","INS","BOLD:ACJ0208","Arthropoda","Insecta","Diptera","Chironomidae","Orthocladiinae","","","1","1","1","47","0","0","99.77","435","0","450","35.11","0"
"AGAW2","02E","INS","BOLD:AAA5803","Arthropoda","Insecta","Hemiptera","Miridae","Mirinae","Lygus","Lygus lineolaris","1","1","1","47","0","0","98.22","449","0","450","34.22","0"
"AGAW2","02E","INS","BOLD:AAH2370","Arthropoda","Insecta","Coleoptera","Elateridae","Dendrometrinae","Sylvanelater","Sylvanelater cylindriformis","1","1","1","47","0","0","98.8","418","0","450","39.33","0"
"AGAW2","02E","NAR","no match","no match","no match","no match","no match","no match","no match","no match","no match","no match","no match","no match","no match","no match","no match","no match","no match","no match","no match","no match"
"AGAW2","02E","NIN","BOLD:AAB7915","Arthropoda","Collembola","Symphypleona","Bourletiellidae",NA,"Deuterosminthurus","Deuterosminthurus sp1","19","18","18","1","5.26","0","99.31","434.63","0","436","35.23","0"
"AGAW2","02E","NIN","BOLD:ABX0027","Arthropoda","Collembola","Symphypleona","Bourletiellidae",NA,"Bourletiella","","18","17","17","2","5.56","0","99.57","418.17","0","448.56","35.74","0"
"AGAW2","02N","BAC","TAX:175768","Proteobacteria","Alphaproteobacteria","Rickettsiales","Rickettsiaceae",NA,"Wolbachia","Wolbachia pipientis","13","12","12","1","7.69","0","98.74","446.46","0","447.54","39.41","0"
"AGAW2","02N","BAC","TAX:91391","Proteobacteria","Alphaproteobacteria","Rickettsiales","",NA,"","","10","10","10","2","0","0","97.67","265.6","0","450","37.71","0"
"AGAW2","02N","BAC","TAX:91393","Proteobacteria","Alphaproteobacteria","Rickettsiales","Rickettsiaceae",NA,"Wolbachia","","3","3","3","3","0","0","98.87","388.67","0","395.67","38.93","0"
"AGAW2","02N","BAC","TAX:376688","Proteobacteria","Alphaproteobacteria","Rickettsiales","Rickettsiaceae",NA,"Wolbachia","Wolbachia sp. elachBioLep01 BioLep373","1","1","1","4","0","0","98.66","449","0","450","40.22","0"
"AGAW2","02N","BAC","TAX:392874","Proteobacteria","Alphaproteobacteria","Rickettsiales","Rickettsiaceae",NA,"Wolbachia","Wolbachia endosymbiont of Hymenoptera","1","1","1","4","0","0","98.67","450","0","450","40.67","0"
"AGAW2","02N","BAC","TAX:375772","Proteobacteria","Alphaproteobacteria","Rickettsiales","Rickettsiaceae",NA,"Wolbachia","Wolbachia sp. Leptogenys JCR25_ASANE483-10","1","1","1","4","0","0","97.99","447","0","450","40.22","0"
"AGAW2","02N","CHO","no match","no match","no match","no match","no match","no match","no match","no match","no match","no match","no match","no match","no match","no match","no match","no match","no match","no match","no match","no match"
"AGAW2","02N","INS","BOLD:AAB9140","Arthropoda","Insecta","Diptera","Calliphoridae","Chrysomyinae","Phormia","Phormia regina","53918","26909","26909","1","50.09","0","99.67","445.64","0","446.67","33.07","0"
"AGAW2","02N","INS","BOLD:AAG2511","Arthropoda","Insecta","Diptera","Anthomyiidae","Anthomyiinae","Delia","Delia platura","13806","10378","10378","2","24.83","0","99.31","443.02","0","445.23","33.17","0"
"AGAW2","02N","INS","BOLD:AAD4355","Arthropoda","Insecta","Diptera","Culicidae","Culicinae","Aedes","Aedes provocans","10499","8656","8656","3","17.55","0","99.05","435.05","0","437.14","31.93","0"
"AGAW2","02N","INS","BOLD:AAA3453","Arthropoda","Insecta","Diptera","Anthomyiidae","Anthomyiinae","Delia","","7451","5484","5484","4","26.4","0","99.39","443.81","0","445.65","34.78","0"
"AGAW2","02N","INS","BOLD:AAB2644","Arthropoda","Insecta","Diptera","Chironomidae","Orthocladiinae","Orthocladius","Orthocladius lapponicus, Orthocladius dorenus, Orthocladius mallochi","6027","5095","5095","5","15.46","0","98.82","438.93","0","440.95","32.81","0"
"AGAW2","02N","INS","BOLD:AAB1098","Arthropoda","Insecta","Diptera","Culicidae","Culicinae","Ochlerotatus, Aedes","Aedes excrucians, Aedes cantans, Aedes stimulans, Aedes euedes, Aedes riparius, Aedes annulipes, Ochlerotatus cantans, Aedes fitchii, Aedes sp., Ochlerotatus annulipes, Ochlerotatus excrucians","5226","4748","4748","6","9.15","0","98.62","429.92","0","431.26","35.11","0"
"AGAW2","02N","INS","BOLD:AAN5355","Arthropoda","Insecta","Diptera","Chironomidae","Orthocladiinae","Smittia","Smittia cf. stercoraria, Smittia stercoraria","5014","4280","4280","7","14.64","0","99.2","420.35","0","445.32","36.27","0"
"AGAW2","02N","INS","BOLD:AAD0853","Arthropoda","Insecta","Diptera","Scathophagidae","Scathophaginae","Scathophaga","Scathophaga lutaria, Scathophaga hiemalis, Scathophaga furcata, Scathophaga stercoraria, Scathophaga inquinata, Scathophaga suilla","3288","2345","2345","8","28.68","0","99","444.47","0","445.77","32.17","0"
"AGAW2","02N","INS","BOLD:AAN5358","Arthropoda","Insecta","Diptera","Chironomidae","Orthocladiinae","Smittia","Smittia stercoraria, Smittia sp. 22ES","2304","1919","1919","9","16.71","0","99.18","442.34","0","444.28","37.25","0"
"AGAW2","02N","INS","BOLD:ABW1509","Arthropoda","Insecta","Diptera","Chironomidae","Orthocladiinae","","","1047","857","857","10","18.15","0","99.36","438.96","0","440.64","35.39","0"
"AGAW2","02N","INS","BOLD:AAP9022","Arthropoda","Insecta","Diptera","Cecidomyiidae","","","","975","669","669","11","31.38","0","99.67","446.62","0","447.72","27.65","0"
"AGAW2","02N","INS","BOLD:ACF2821","Arthropoda","Insecta","Diptera","Mycetophilidae","Mycetophilinae","Mycetophila","Mycetophila fungorum","956","805","805","12","15.79","0","99.12","437.43","0","439.83","35.42","0"
"AGAW2","02N","INS","BOLD:AAN5267","Arthropoda","Insecta","Diptera","Cecidomyiidae","","","","649","475","475","13","26.81","0","99.18","447.88","0","448.89","30.58","0"
"AGAW2","02N","INS","BOLD:ACP4736","Arthropoda","Insecta","Diptera","Chironomidae","Orthocladiinae","Smittia","Smittia sp. 8ES","549","455","455","14","17.12","0","99.18","432.79","0","434.33","33.11","0"
"AGAW2","02N","INS","BOLD:ACJ0208","Arthropoda","Insecta","Diptera","Chironomidae","Orthocladiinae","","","503","393","393","15","21.87","0","99.11","435.93","0","449.35","34.74","0"
"AGAW2","02N","INS","BOLD:AAG1000","Arthropoda","Insecta","Diptera","Chironomidae","Orthocladiinae","Orthocladius","Orthocladius carlatus","317","312","312","16","1.58","0","98.07","427.56","0","429.66","33.9","0"
"AGAW2","02N","INS","BOLD:AAG7312","Arthropoda","Insecta","Diptera","Sphaeroceridae","Limosininae","Spelobia","Spelobia clunipes, Spelobia sp. B TAW","307","247","247","17","19.54","0","99.1","447.28","0","448.68","33.54","0"
"AGAW2","02N","INS","BOLD:AAK5928","Arthropoda","Insecta","Coleoptera","Chrysomelidae","Criocerinae","Oulema","Oulema melanopus","251","206","206","18","17.93","0","99.59","446.89","0","448.03","36.49","0"
"AGAW2","02N","INS","BOLD:AAN6445","Arthropoda","Insecta","Diptera","Sciaridae","","","","201","177","177","19","11.94","0","98.57","438.49","0","441.76","34.44","0"
"AGAW2","02N","INS","BOLD:ACZ4169","Arthropoda","Insecta","Diptera","Agromyzidae","Phytomyzinae","Cerodontha","Cerodontha incisa","187","177","177","20","5.35","0","99.3","434.37","0","436.31","31.1","0"
"AGAW2","02N","INS","BOLD:AAA1513","Arthropoda","Insecta","Lepidoptera","Plutellidae","Plutellinae","Plutella, Rhigognostis","Plutella xylostella, Rhigognostis sp.","96","80","80","21","16.67","0","99.34","445","0","446.17","32.25","0"
"AGAW2","02N","INS","BOLD:AAM9254","Arthropoda","Insecta","Diptera","Sciaridae","","Bradysia","Bradysia pallipes","91","89","89","22","2.2","0","98.87","432.59","0","432.87","33.39","0"
"AGAW2","02N","INS","BOLD:AAF9014","Arthropoda","Insecta","Diptera","Limoniidae","Chioneinae","Symplecta","Symplecta cana, Symplecta sp. BOLDAAF9014","74","69","69","23","6.76","0","98.51","432.92","0","440.62","33.49","0"
"AGAW2","02N","INS","BOLD:AAH3669","Arthropoda","Insecta","Diptera","Cecidomyiidae","","","","65","45","45","24","30.77","0","99.05","446.82","0","447.4","30.13","0"
"AGAW2","02N","INS","BOLD:ACB1079","Arthropoda","Insecta","Diptera","Cecidomyiidae","Micromyinae","Campylomyza","Campylomyza dilatata, CecidInt44 sp. BOLD:ACB1079","63","52","52","25","17.46","0","98.39","443.37","0","446.35","28.28","0"
"AGAW2","02N","INS","BOLD:AAC4197","Arthropoda","Insecta","Diptera","Chironomidae","Orthocladiinae","Paraphaenocladius","Paraphaenocladius impensus","61","56","56","26","8.2","0","99.31","439.39","0","443.28","35.11","0"
"AGAW2","02N","INS","BOLD:AAY6412","Arthropoda","Insecta","Diptera","Sciaridae","","Bradysia","Bradysia urticae","39","39","39","27","0","0","99.44","434.74","0","437.85","32.05","0"
"AGAW2","02N","INS","BOLD:ABA6440","Arthropoda","Insecta","Diptera","Chironomidae","Chironominae","Polypedilum","","24","20","20","28","16.67","0","97.95","433.96","0","434.29","33.32","0"
"AGAW2","02N","INS","BOLD:AAN6435","Arthropoda","Insecta","Diptera","Sciaridae","","Bradysia","Bradysia angustipennis","23","21","21","29","8.7","0","99.52","396","0","436.48","34.38","0"
"AGAW2","02N","INS","BOLD:AAM9252","Arthropoda","Insecta","Diptera","Sciaridae","","Bradysia, Leptosciarella","Bradysia vagans, Leptosciarella zanti","20","20","20","30","0","0","99.61","441.8","0","448.45","38.92","0"
"AGAW2","02N","INS","BOLD:ACX9624","Arthropoda","Insecta","Diptera","Cecidomyiidae","","","","19","15","15","31","21.05","0","97.33","448.16","0","450","30.05","0"
"AGAW2","02N","INS","BOLD:AAH2753","Arthropoda","Insecta","Coleoptera","Anthicidae","Anthicinae","Anthicus","Anthicus cervinus","19","19","19","31","0","0","98.69","419.95","0","435","34.19","0"
"AGAW2","02N","INS","BOLD:AAO8538","Arthropoda","Insecta","Diptera","Psychodidae","Psychodinae","Psychoda","","15","15","15","33","0","0","98.61","441.13","0","442.67","33.01","0"
"AGAW2","02N","INS","BOLD:AAB0868","Arthropoda","Insecta","Diptera","Calliphoridae","Calliphorinae","Cynomya","Cynomya cadaverina, Cynomya mortuorum","14","13","13","34","7.14","0","99.05","449.29","0","450","34.86","0"
"AGAW2","02N","INS","BOLD:AAN8770","Arthropoda","Insecta","Diptera","Psychodidae","Psychodinae","Psychoda","Psychoda trinodulosa","12","12","12","35","0","0","99","440.42","0","441.5","33.59","0"
"AGAW2","02N","INS","BOLD:AAB2384","Arthropoda","Insecta","Diptera","Syrphidae","Syrphinae","Eupeodes","Eupeodes duseki, Eupeodes volucris, Eupeodes nielseni, Eupeodes americanus, Eupeodes flukei, Eupeodes vockerothi, Eupeodes lucasi, Eupeodes montivagus, Eupeodes rufipunctatus, Eupeodes rojasi, Eupeodes goeldlini, Eupeodes luniger, Eupeodes punctifer, Eupeodes bucculatus, Eupeodes perplexus, Eupeodes abiskoensis, Eupeodes corollae, Eupeodes pomus, Eupeodes lundbecki, Eupeodes nitens, Eupeodes sp. BOLDAAB2384, Eupeodes curtus, Eupeodes latifasciatus","12","12","12","35","0","0","98.81","419.83","0","422","30.78","0"
"AGAW2","02N","INS","BOLD:ACX6055","Arthropoda","Insecta","Diptera","Phoridae","Metopininae","Megaselia","Megaselia sp. AK-2010-2, Megaselia sp. BOLD:ACX6055","10","10","10","37","0","0","99.75","430.3","0","433.4","32.85","0"
"AGAW2","02N","INS","BOLD:AAU8379","Arthropoda","Insecta","Hymenoptera","Braconidae","Alysiinae","Dinotrema","Dinotrema JFT01","8","8","8","38","0","0","99.8","429.75","0","430.63","24.19","0"
"AGAW2","02N","INS","BOLD:AAP5146","Arthropoda","Insecta","Diptera","Chironomidae","Orthocladiinae","Orthocladius","","8","8","8","38","0","0","98.53","434.38","0","435.63","34.17","0"
"AGAW2","02N","INS","BOLD:AAC1928","Arthropoda","Insecta","Coleoptera","Carabidae","Harpalinae","Harpalus","Harpalus sp. 2, Harpalus herbivagus","8","8","8","38","0","0","98.74","430.88","0","433.13","31.38","0"
"AGAW2","02N","INS","BOLD:AAG1742","Arthropoda","Insecta","Diptera","Muscidae","Phaoniinae","Helina","Helina rufitibia","7","5","5","41","28.57","0","99","445.86","0","446.29","30.38","0"
"AGAW2","02N","INS","BOLD:AAM7855","Arthropoda","Insecta","Thysanoptera","Thripidae","Thripinae","Frankliniella","Frankliniella tritici","6","6","6","42","0","0","98.99","446","0","449","31.33","0"
"AGAW2","02N","INS","BOLD:AAA4222","Arthropoda","Insecta","Lepidoptera","Erebidae","Hypeninae","Hypena","Hypena scabra","6","6","6","42","0","0","99.57","404.67","0","406.17","32.69","0"
"AGAW2","02N","INS","BOLD:AAG3316","Arthropoda","Insecta","Diptera","Phoridae","Metopininae","Megaselia","","5","5","5","44","0","0","98.88","448.4","0","450","33.91","0"
"AGAW2","02N","INS","BOLD:AAC6643","Arthropoda","Insecta","Diptera","Chironomidae","Orthocladiinae","Metriocnemus","Metriocnemus albolineatus","3","3","3","45","0","0","97.86","412","0","413.33","33.99","0"
"AGAW2","02N","INS","BOLD:AAG6743","Arthropoda","Insecta","Diptera","Sarcophagidae","Sarcophaginae","Sarcophaga","Sarcophaga sp. 1 MG, Sarcophaga subvicina","3","3","3","45","0","0","97.95","419.33","0","420.33","34.95","0"
"AGAW2","02N","INS","BOLD:AAG2462","Arthropoda","Insecta","Diptera","Anthomyiidae","Anthomyiinae","Botanophila","Botanophila fugax","3","3","3","45","0","0","99.4","448","0","450","31.7","0"
"AGAW2","02N","INS","BOLD:AAA8109","Arthropoda","Insecta","Lepidoptera","Gelechiidae","Dichomeridinae","Dichomeris","Dichomeris ligulella","2","2","2","48","0","0","99","449.5","0","450","32.44","0"
"AGAW2","02N","INS","BOLD:AAP6827","Arthropoda","Insecta","Diptera","Cecidomyiidae","","","","2","2","2","48","0","0","98.88","447.5","0","450","31.89","0"
"AGAW2","02N","INS","BOLD:AAC9062","Arthropoda","Insecta","Diptera","Culicidae","Culicinae","Ochlerotatus, Aedes","Aedes fitchii, Ochlerotatus fitchii, Aedes grossbecki, Aedes ventrovittis","2","2","2","48","0","0","98.43","417.5","0","418.5","33.07","0"
"AGAW2","02N","INS","BOLD:AAA2482","Arthropoda","Insecta","Lepidoptera","Noctuidae","Noctuinae","Mythimna","Mythimna unipuncta","2","2","2","48","0","0","99.89","448.5","0","450","30.89","0"
"AGAW2","02N","INS","BOLD:AAG6745","Arthropoda","Insecta","Diptera","Calliphoridae","Calliphorinae","Pollenia","Pollenia pediculata","2","2","2","48","0","0","99.42","428.5","0","428.5","33.36","0"
"AGAW2","02N","INS","BOLD:AAC9132","Arthropoda","Insecta","Diptera","Culicidae","Culicinae","Culiseta","Culiseta inornata","2","2","2","48","0","0","99.02","401.5","0","404","32.67","0"
"AGAW2","02N","INS","BOLD:AAH3914","Arthropoda","Insecta","Diptera","Sciaridae","","Sciara","Sciara hamatilis, Sciara humeralis","2","2","2","48","0","0","99.44","448","0","450","30","0"
"AGAW2","02N","INS","BOLD:ACC7611","Arthropoda","Insecta","Diptera","Cecidomyiidae","","","","1","1","1","55","0","0","97.77","448","0","450","28.22","0"
"AGAW2","02N","INS","BOLD:AAG9055","Arthropoda","Insecta","Hymenoptera","Vespidae","Vespinae","Vespula","Vespula germanica","1","1","1","55","0","0","97.2","428","0","429","32.4","0"
"AGAW2","02N","INS","BOLD:ACV3368","","","","","","","","1","1","1","55","0","0","99.78","446","0","449","33.41","0"
"AGAW2","02N","INS","BOLD:AAK8831","Arthropoda","Insecta","Diptera","Limoniidae","Limoniinae","Metalimnobia, Limonia","","1","1","1","55","0","0","98.22","449","0","450","32.67","0"
"AGAW2","02N","INS","BOLD:AAM6047","Arthropoda","Insecta","Diptera","Cecidomyiidae","","","","1","1","1","55","0","0","98.93","373","0","376","31.65","0"
"AGAW2","02N","INS","BOLD:AAL7534","Arthropoda","Insecta","Diptera","Anisopodidae","Anisopodinae","Sylvicola","Sylvicola stackelbergi, Sylvicola stackelbergii 1, Sylvicola punctatus","1","1","1","55","0","0","98.44","448","0","450","34.44","0"
"AGAW2","02N","INS","BOLD:AAE0844","Arthropoda","Insecta","Hemiptera","Pentatomidae","Pentatominae","Euschistus","Euschistus variolarius","1","1","1","55","0","0","98.22","450","0","450","36.44","0"
"AGAW2","02N","INS","BOLD:AAG7276","Arthropoda","Insecta","Diptera","Sphaeroceridae","Limosininae","Leptocera","Leptocera erythrocera","1","1","1","55","0","0","99.78","449","0","450","32.89","0"
"AGAW2","02N","INS","BOLD:AAU6970","Arthropoda","Insecta","Coleoptera","Cleridae","Clerinae","Enoclerus","Enoclerus nigripes","1","1","1","55","0","0","98.44","450","0","450","33.11","0"
"AGAW2","02N","INS","BOLD:ABX1202","Arthropoda","Insecta","Diptera","Chironomidae","Chironominae","Cladotanytarsus","","1","1","1","55","0","0","97.77","449","0","450","34.44","0"
"AGAW2","02N","INS","BOLD:AAB5640","Arthropoda","Insecta","Coleoptera","Coccinellidae","Coccinellinae","Harmonia","Harmonia axyridis","1","1","1","55","0","0","99.78","450","0","450","32.67","0"
"AGAW2","02N","INS","BOLD:AAL2750","Arthropoda","Insecta","Diptera","Muscidae","Phaoniinae","Phaonia","Phaonia tuguriorum","1","1","1","55","0","0","98","449","0","450","32.89","0"
"AGAW2","02N","INS","BOLD:ACB9399","Arthropoda","Insecta","Diptera","Chironomidae","Chironominae","Parachironomus","","1","1","1","55","0","0","97.32","447","0","450","35.78","0"
"AGAW2","02N","INS","BOLD:AAA4555","Arthropoda","Insecta","Orthoptera","Acrididae","Melanoplinae","Melanoplus","Melanoplus stupefactus, Melanoplus angustipennis, Melanoplus ponderosus, Melanoplus bruneri, Melanoplus frigidus, Melanoplus scapularis, Melanoplus cinereus, Melanoplus madeleineae, Melanoplus sanguinipes, Melanoplus nigrescens group, Melanoplus tunicae, Melanoplus walshi, Melanoplus femurrubrum, Melanoplus dawsoni, Melanoplus sp., Melanoplus fasciatus, Melanoplus bowditchi, Melanoplus stonei, Melanoplus bivittatus, Melanoplus huroni, Melanoplus thomasi, Melanoplus borealis, Melanoplus packardii","1","1","1","55","0","0","99.52","419","0","422","36.02","0"
"AGAW2","02N","INS","BOLD:ABU9891","Arthropoda","Insecta","Diptera","Muscidae","Phaoniinae","Phaonia","","1","1","1","55","0","0","97.78","450","0","450","32.89","0"
"AGAW2","02N","INS","BOLD:ACL6160","Arthropoda","Insecta","Coleoptera","Chrysomelidae","Galerucinae","Longitarsus","","1","1","1","55","0","0","98.44","450","0","450","32.89","0"
"AGAW2","02N","INS","BOLD:AAN1738","Arthropoda","Insecta","Coleoptera","Staphylinidae","Oxytelinae","Anotylus","Anotylus tetracarinatus","1","1","1","55","0","0","100","449","0","450","33.56","0"
"AGAW2","02N","INS","BOLD:AAB7513","Arthropoda","Insecta","Diptera","Simuliidae","Simuliinae","Stegopterna","Stegopterna mutata/diplomutata","1","1","1","55","0","0","98.42","380","0","383","37.08","0"
"AGAW2","02N","INS","BOLD:ABA1215","Arthropoda","Insecta","Diptera","Sciaridae","","Lycoriella","Lycoriella sativae","1","1","1","55","0","0","98","450","0","450","35.78","0"
"AGAW2","02N","INS","BOLD:AAA8933","Arthropoda","Insecta","Coleoptera","Coccinellidae","Coccinellinae","Coccinella","Coccinella septempunctata","1","1","1","55","0","0","99.78","449","0","450","32.89","0"
"AGAW2","02N","INS","BOLD:AAB4093","Arthropoda","Insecta","Lepidoptera","Noctuidae","Noctuinae","Elaphria","Elaphria grata","1","1","1","55","0","0","99.78","446","0","450","30.22","0"
"AGAW2","02N","INS","BOLD:ACR4394","Arthropoda","Insecta","Diptera","Anthomyiidae","Anthomyiinae","Delia","Delia florilega","1","1","1","55","0","0","97.54","448","0","450","34","0"
"AGAW2","02N","INS","BOLD:ABA8091","Arthropoda","Insecta","Hymenoptera","Braconidae","Aphidiinae","Trioxys","Trioxys pallidus","1","1","1","55","0","0","97.6","416","0","450","32.67","0"
"AGAW2","02N","INS","BOLD:AAP5142","Arthropoda","Insecta","Diptera","Chironomidae","Chironominae","","","1","1","1","55","0","0","99.33","448","0","450","32.44","0"
"AGAW2","02N","INS","BOLD:AAH0134","Arthropoda","Insecta","Coleoptera","Phalacridae","Phalacrinae","Stilbus","Stilbus apicialis","1","1","1","55","0","0","97.04","439","0","440","32.27","0"
"AGAW2","02N","INS","BOLD:AAM6241","Arthropoda","Insecta","Diptera","Chironomidae","","","","1","1","1","55","0","0","100","448","0","450","35.56","0"
"AGAW2","02N","NAR","no match","no match","no match","no match","no match","no match","no match","no match","no match","no match","no match","no match","no match","no match","no match","no match","no match","no match","no match","no match"
"AGAW2","02N","NIN","BOLD:AAB7667","Arthropoda","Arachnida","Araneae","Lycosidae","","Pardosa","Pardosa saxatilis","74","74","74","1","0","0","98.57","416.35","0","420.74","31.66","0"
"AGAW2","02N","NIN","BOLD:AAV0589","Arthropoda","Arachnida","Mesostigmata","Blattisociidae","Platyseiinae","Cheiroseius","","7","7","7","2","0","0","99.52","447.71","0","450","27.52","0"
"AGAW2","02N","NIN","BOLD:ABX0027","Arthropoda","Collembola","Symphypleona","Bourletiellidae","","Bourletiella","","3","3","3","3","0","0","99.6","418","0","450","35.48","0"
"AGAW2","02N","NIN","BOLD:AAB5726","Arthropoda","Arachnida","Araneae","Agelenidae","","Agelenopsis","Agelenopsis pennsylvanica, Agelenopsis emertoni, Agelenopsis potteri, Agelenopsis actuosa, Agelenopsis kastoni","2","2","2","4","0","0","98.09","444.5","0","445.5","33.56","0"
"AGAW2","02N","NIN","BOLD:AAN6642","Arthropoda","Arachnida","Mesostigmata","Blattisociidae","Platyseiinae","Cheiroseius","","2","2","2","4","0","0","99.21","447.5","0","450","26.56","0"
"AGAW2","02N","NIN","BOLD:ABY2066","Arthropoda","Arachnida","Mesostigmata","Macrochelidae","","Macrocheles","Macrocheles muscaedomesticae","1","1","1","6","0","0","99.48","386","0","398","31.91","0"
"AGAW2","02N","NIN","BOLD:AAA4664","Arthropoda","Collembola","Entomobryomorpha","Entomobryidae","Lepidocyrtinae","Willowsia","Willowsia buski","1","1","1","6","0","0","100","398","0","399","41.6","0"
"AGAW2","02S","BAC","no match","no match","no match","no match","no match","no match","no match","no match","no match","no match","no match","no match","no match","no match","no match","no match","no match","no match","no match","no match"
"AGAW2","02S","CHO","no match","no match","no match","no match","no match","no match","no match","no match","no match","no match","no match","no match","no match","no match","no match","no match","no match","no match","no match","no match"
"AGAW2","02S","INS","BOLD:AAB9140","Arthropoda","Insecta","Diptera","Calliphoridae","Chrysomyinae","Phormia","Phormia regina","37572","21112","21112","1","43.81","0","99.45","443.34","0","444.33","33.28","0"
"AGAW2","02S","INS","BOLD:AAG2511","Arthropoda","Insecta","Diptera","Anthomyiidae","Anthomyiinae","Delia","Delia platura","18341","13127","13127","2","28.43","0","99.39","443.18","0","445.45","33.16","0"
"AGAW2","02S","INS","BOLD:AAB2384","Arthropoda","Insecta","Diptera","Syrphidae","Syrphinae","Eupeodes","Eupeodes duseki, Eupeodes volucris, Eupeodes nielseni, Eupeodes americanus, Eupeodes flukei, Eupeodes vockerothi, Eupeodes lucasi, Eupeodes montivagus, Eupeodes rufipunctatus, Eupeodes rojasi, Eupeodes goeldlini, Eupeodes luniger, Eupeodes punctifer, Eupeodes bucculatus, Eupeodes perplexus, Eupeodes abiskoensis, Eupeodes corollae, Eupeodes pomus, Eupeodes lundbecki, Eupeodes nitens, Eupeodes sp. BOLDAAB2384, Eupeodes curtus, Eupeodes latifasciatus","13046","10103","10103","3","22.56","0","98.81","430.53","0","432.54","30.96","0"
"AGAW2","02S","INS","BOLD:AAD4355","Arthropoda","Insecta","Diptera","Culicidae","Culicinae","Aedes","Aedes provocans","9415","7867","7867","4","16.44","0","99.17","434.18","0","436.54","31.89","0"
"AGAW2","02S","INS","BOLD:AAB2644","Arthropoda","Insecta","Diptera","Chironomidae","Orthocladiinae","Orthocladius","Orthocladius lapponicus, Orthocladius dorenus, Orthocladius mallochi","7753","5807","5807","5","25.1","0","99.26","443.5","0","445.37","32.45","0"
"AGAW2","02S","INS","BOLD:AAA3453","Arthropoda","Insecta","Diptera","Anthomyiidae","Anthomyiinae","Delia","","5609","4191","4191","6","25.28","0","99.41","443.83","0","445.72","34.77","0"
"AGAW2","02S","INS","BOLD:ACL0865","Arthropoda","Insecta","Diptera","Sciaridae","","Leptosciarella","","359","352","352","7","1.95","0","98.26","421.52","0","428.09","32.94","0"
"AGAW2","02S","INS","BOLD:AAN6407","Arthropoda","Insecta","Diptera","Sphaeroceridae","Limosininae","Coproica","Coproica ferruginata","307","255","255","8","16.94","0","99.29","445.51","0","447.24","33.38","0"
"AGAW2","02S","INS","BOLD:ACJ0208","Arthropoda","Insecta","Diptera","Chironomidae","Orthocladiinae","","","132","108","108","9","18.18","0","99.16","435.67","0","449","34.95","0"
"AGAW2","02S","INS","BOLD:AAN5267","Arthropoda","Insecta","Diptera","Cecidomyiidae","","","","121","93","93","10","23.14","0","99.52","448.11","0","449.17","30.72","0"
"AGAW2","02S","INS","BOLD:ACF8552","Arthropoda","Insecta","Coleoptera","Anthicidae","Anthicinae","Malporus","Malporus formicarius","95","77","77","11","18.95","0","99.04","364.74","0","443.72","34.81","0"
"AGAW2","02S","INS","BOLD:ABA1215","Arthropoda","Insecta","Diptera","Sciaridae","","Lycoriella","Lycoriella sativae","92","73","73","12","20.65","0","99.55","444.01","0","445.36","35.86","0"
"AGAW2","02S","INS","BOLD:AAH2753","Arthropoda","Insecta","Coleoptera","Anthicidae","Anthicinae","Anthicus","Anthicus cervinus","86","72","72","13","16.28","0","98.05","428.42","0","446.7","34.56","0"
"AGAW2","02S","INS","BOLD:ADH4889","Arthropoda","Insecta","Diptera","","","","","67","56","56","14","16.42","0","99.14","446.75","0","447.45","35.72","0"
"AGAW2","02S","INS","BOLD:AAM7726","Arthropoda","Insecta","Coleoptera","Curculionidae","Curculioninae","Orchestes","Orchestes steppensis, Orchestes alni","65","61","61","15","6.15","0","99.18","438.25","0","440.72","33.13","0"
"AGAW2","02S","INS","BOLD:ABU5520","Arthropoda","Insecta","Diptera","Sciaridae","","Corynoptera","","58","57","57","16","1.72","0","99.36","434.86","0","435.83","35.94","0"
"AGAW2","02S","INS","BOLD:AAN5355","Arthropoda","Insecta","Diptera","Chironomidae","Orthocladiinae","Smittia","Smittia cf. stercoraria, Smittia stercoraria","56","53","53","17","5.36","0","98.73","437.55","0","444.38","36.1","0"
"AGAW2","02S","INS","BOLD:AAC4197","Arthropoda","Insecta","Diptera","Chironomidae","Orthocladiinae","Paraphaenocladius","Paraphaenocladius impensus","55","52","52","18","5.45","0","99.28","435.84","0","440.75","35.13","0"
"AGAW2","02S","INS","BOLD:AAN5358","Arthropoda","Insecta","Diptera","Chironomidae","Orthocladiinae","Smittia","Smittia stercoraria, Smittia sp. 22ES","53","51","51","19","3.77","0","99.21","439.58","0","441.3","37.58","0"
"AGAW2","02S","INS","BOLD:ACB1079","Arthropoda","Insecta","Diptera","Cecidomyiidae","Micromyinae","Campylomyza","Campylomyza dilatata, CecidInt44 sp. BOLD:ACB1079","49","42","42","20","14.29","0","98.19","442","0","443.94","28.37","0"
"AGAW2","02S","INS","BOLD:AAN8770","Arthropoda","Insecta","Diptera","Psychodidae","Psychodinae","Psychoda","Psychoda trinodulosa","25","25","25","21","0","0","99.24","442.76","0","443.64","33.84","0"
"AGAW2","02S","INS","BOLD:AAG1000","Arthropoda","Insecta","Diptera","Chironomidae","Orthocladiinae","Orthocladius","Orthocladius carlatus","17","17","17","22","0","0","97.94","429.24","0","431.59","33.83","0"
"AGAW2","02S","INS","BOLD:AAD0853","Arthropoda","Insecta","Diptera","Scathophagidae","Scathophaginae","Scathophaga","Scathophaga lutaria, Scathophaga hiemalis, Scathophaga furcata, Scathophaga stercoraria, Scathophaga inquinata, Scathophaga suilla","12","12","12","23","0","0","98.67","439.75","0","442.33","31.69","0"
"AGAW2","02S","INS","BOLD:ACC7611","Arthropoda","Insecta","Diptera","Cecidomyiidae","","","","9","9","9","24","0","0","97.32","447","0","450","28.37","0"
"AGAW2","02S","INS","BOLD:AAY6412","Arthropoda","Insecta","Diptera","Sciaridae","","Bradysia","Bradysia urticae","8","8","8","25","0","0","99.53","448.88","0","450","32.36","0"
"AGAW2","02S","INS","BOLD:AAN5356","Arthropoda","Insecta","Diptera","Chironomidae","Orthocladiinae","Smittia","","7","7","7","26","0","0","98.61","426.29","0","429.43","34.9","0"
"AGAW2","02S","INS","BOLD:AAB0868","Arthropoda","Insecta","Diptera","Calliphoridae","Calliphorinae","Cynomya","Cynomya cadaverina, Cynomya mortuorum","5","3","3","27","40","0","99.82","449.6","0","450","35.38","0"
"AGAW2","02S","INS","BOLD:AAP6827","Arthropoda","Insecta","Diptera","Cecidomyiidae","","","","4","4","4","28","0","0","98.67","431.75","0","434.5","31.4","0"
"AGAW2","02S","INS","BOLD:ACL9997","Arthropoda","Insecta","Coleoptera","Trogossitidae","Peltinae","Grynocharis","Grynocharis quadrilineata","4","4","4","28","0","0","97.43","447","0","448.5","40.86","0"
"AGAW2","02S","INS","BOLD:AAA4222","Arthropoda","Insecta","Lepidoptera","Erebidae","Hypeninae","Hypena","Hypena scabra","4","4","4","28","0","0","99.16","440.75","0","441","32.42","0"
"AGAW2","02S","INS","BOLD:AAP9022","Arthropoda","Insecta","Diptera","Cecidomyiidae","","","","3","3","3","31","0","0","99.7","450","0","450","27.85","0"
"AGAW2","02S","INS","BOLD:AAG9055","Arthropoda","Insecta","Hymenoptera","Vespidae","Vespinae","Vespula","Vespula germanica","2","2","2","32","0","0","99.55","435","0","437","30.75","0"
"AGAW2","02S","INS","BOLD:ACP4925","Arthropoda","Insecta","Diptera","Sciaridae","","","","2","2","2","32","0","0","97.69","389.5","0","450","36.33","0"
"AGAW2","02S","INS","BOLD:ACO2805","Arthropoda","Insecta","Diptera","Chironomidae","Chironominae","","","2","2","2","32","0","0","97.1","448","0","450","35.89","0"
"AGAW2","02S","INS","BOLD:AAA2482","Arthropoda","Insecta","Lepidoptera","Noctuidae","Noctuinae","Mythimna","Mythimna unipuncta","2","2","2","32","0","0","99.13","424","0","427.5","30.15","0"
"AGAW2","02S","INS","BOLD:AAG6743","Arthropoda","Insecta","Diptera","Sarcophagidae","Sarcophaginae","Sarcophaga","Sarcophaga sp. 1 MG, Sarcophaga subvicina","2","2","2","32","0","0","99.63","417","0","418.5","34.61","0"
"AGAW2","02S","INS","BOLD:AAA2955","Arthropoda","Insecta","Lepidoptera","Tortricidae","Tortricinae","Argyrotaenia","Argyrotaenia occultana, Argyrotaenia ljungiana, Argyrotaenia pinatubana, Argyrotaenia floridana, Argyrotaenia velutinana, Argyrotaenia tabulana, Argyrotaenia gogana","2","2","2","32","0","0","98.28","417.5","0","420","31.77","0"
"AGAW2","02S","INS","BOLD:ACK1651","Arthropoda","Insecta","Diptera","Cecidomyiidae","","","","2","2","2","32","0","0","98.1","448.5","0","450","27.56","0"
"AGAW2","02S","INS","BOLD:AAH0134","Arthropoda","Insecta","Coleoptera","Phalacridae","Phalacrinae","Stilbus","Stilbus apicialis","2","2","2","32","0","0","99.17","410.5","0","411","31.81","0"
"AGAW2","02S","INS","BOLD:AAQ0582","Arthropoda","Insecta","Diptera","Anthomyiidae","Anthomyiinae","Lasiomma","","2","2","2","32","0","0","98.78","449.5","0","450","31.67","0"
"AGAW2","02S","INS","BOLD:AAG2462","Arthropoda","Insecta","Diptera","Anthomyiidae","Anthomyiinae","Botanophila","Botanophila fugax","2","2","2","32","0","0","98.77","446.5","0","450","31.67","0"
"AGAW2","02S","INS","BOLD:ACD3588","Arthropoda","Insecta","Diptera","Sciaridae","","Lycoriella","Lycoriella sativae","2","2","2","32","0","0","99.3","427.5","0","430.5","36.23","0"
"AGAW2","02S","INS","BOLD:AAU6970","Arthropoda","Insecta","Coleoptera","Cleridae","Clerinae","Enoclerus","Enoclerus nigripes","1","1","1","43","0","0","97.94","437","0","439","33.49","0"
"AGAW2","02S","INS","BOLD:AAD0583","Arthropoda","Insecta","Hemiptera","Lygaeidae","Lygaeinae","Lygaeus","Lygaeus reclivatus, Lygaeus kalmii","1","1","1","43","0","0","100","449","0","450","32.22","0"
"AGAW2","02S","INS","BOLD:ACE5721","Arthropoda","Insecta","Lepidoptera","Noctuidae","Noctuinae","Pseudohermonassa","Pseudohermonassa tenuicula","1","1","1","43","0","0","97.1","449","0","450","32.89","0"
"AGAW2","02S","INS","BOLD:ACF7745","Arthropoda","Insecta","Diptera","Trichoceridae","Trichocerinae","Trichocera","Trichocera sp. BOLD:ACF7745","1","1","1","43","0","0","100","362","0","450","38.44","0"
"AGAW2","02S","INS","BOLD:AAA8109","Arthropoda","Insecta","Lepidoptera","Gelechiidae","Dichomeridinae","Dichomeris","Dichomeris ligulella","1","1","1","43","0","0","98.66","449","0","450","32.44","0"
"AGAW2","02S","INS","BOLD:ABU9891","Arthropoda","Insecta","Diptera","Muscidae","Phaoniinae","Phaonia","","1","1","1","43","0","0","98.4","437","0","440","33.86","0"
"AGAW2","02S","INS","BOLD:AAN1738","Arthropoda","Insecta","Coleoptera","Staphylinidae","Oxytelinae","Anotylus","Anotylus tetracarinatus","1","1","1","43","0","0","99.78","450","0","450","33.56","0"
"AGAW2","02S","INS","BOLD:ACJ5126","Arthropoda","Insecta","Diptera","Chironomidae","Orthocladiinae","Metriocnemus","","1","1","1","43","0","0","99.77","444","0","450","34.89","0"
"AGAW2","02S","INS","BOLD:ACA4749","Arthropoda","Insecta","Diptera","Chironomidae","Orthocladiinae","","","1","1","1","43","0","0","98.6","428","0","447","35.12","0"
"AGAW2","02S","INS","BOLD:AAB1098","Arthropoda","Insecta","Diptera","Culicidae","Culicinae","Ochlerotatus, Aedes","Aedes excrucians, Aedes cantans, Aedes stimulans, Aedes euedes, Aedes riparius, Aedes annulipes, Ochlerotatus cantans, Aedes fitchii, Aedes sp., Ochlerotatus annulipes, Ochlerotatus excrucians","1","1","1","43","0","0","98.65","446","0","447","35.35","0"
"AGAW2","02S","INS","BOLD:AAB5640","Arthropoda","Insecta","Coleoptera","Coccinellidae","Coccinellinae","Harmonia","Harmonia axyridis","1","1","1","43","0","0","98.83","428","0","429","31.7","0"
"AGAW2","02S","INS","BOLD:ACX3480","Arthropoda","Insecta","Diptera","Cecidomyiidae","","","","1","1","1","43","0","0","97.33","450","0","450","29.11","0"
"AGAW2","02S","INS","BOLD:AAF1051","Arthropoda","Insecta","Diptera","Agromyzidae","Phytomyzinae","Cerodontha","Cerodontha muscina, Cerodontha calamagrostis","1","1","1","43","0","0","98","449","0","450","36.44","0"
"AGAW2","02S","INS","BOLD:AAV2165","Arthropoda","Insecta","Hymenoptera","Braconidae","Microgastrinae","Cotesia","Cotesia vanessae","1","1","1","43","0","0","99.11","447","0","450","25.11","0"
"AGAW2","02S","INS","BOLD:ACF2821","Arthropoda","Insecta","Diptera","Mycetophilidae","Mycetophilinae","Mycetophila","Mycetophila fungorum","1","1","1","43","0","0","98.41","378","0","381","34.91","0"
"AGAW2","02S","INS","BOLD:AAB4093","Arthropoda","Insecta","Lepidoptera","Noctuidae","Noctuinae","Elaphria","Elaphria grata","1","1","1","43","0","0","97.91","431","0","432","30.32","0"
"AGAW2","02S","INS","BOLD:AAL7534","Arthropoda","Insecta","Diptera","Anisopodidae","Anisopodinae","Sylvicola","Sylvicola stackelbergi, Sylvicola stackelbergii 1, Sylvicola punctatus","1","1","1","43","0","0","97.32","448","0","450","34.22","0"
"AGAW2","02S","INS","BOLD:AAB1024","Arthropoda","Insecta","Lepidoptera","Noctuidae","Noctuinae","Pseudohermonassa","Pseudohermonassa bicarnea","1","1","1","43","0","0","97.31","446","0","449","33.63","0"
"AGAW2","02S","INS","BOLD:ACO2757","Arthropoda","Insecta","Diptera","Chironomidae","Chironominae","","","1","1","1","43","0","0","97.02","369","0","372","34.68","0"
"AGAW2","02S","INS","BOLD:AAF9014","Arthropoda","Insecta","Diptera","Limoniidae","Chioneinae","Symplecta","Symplecta cana, Symplecta sp. BOLDAAF9014","1","1","1","43","0","0","97.05","441","0","450","34","0"
"AGAW2","02S","INS","BOLD:AAG2123","Arthropoda","Insecta","Diptera","Tachinidae","Dexiinae","Dinera","Dinera grisescens","1","1","1","43","0","0","99.78","450","0","450","32.22","0"
"AGAW2","02S","INS","BOLD:AAU8503","Arthropoda","Insecta","Hymenoptera","Tenthredinidae","Nematinae","Pristiphora","Pristiphora subbifida","1","1","1","43","0","0","99.78","449","0","450","30.67","0"
"AGAW2","02S","INS","BOLD:AAL0908","Arthropoda","Insecta","Coleoptera","Chrysomelidae","Galerucinae","Dibolia, Psylliodes","Dibolia borealis","1","1","1","43","0","0","99.49","392","0","397","34.26","0"
"AGAW2","02S","NAR","no match","no match","no match","no match","no match","no match","no match","no match","no match","no match","no match","no match","no match","no match","no match","no match","no match","no match","no match","no match"
"AGAW2","02S","NIN","BOLD:AAH0055","Arthropoda","Arachnida","Araneae","Lycosidae",NA,"Schizocosa","Schizocosa mccooki","535","486","486","1","9.16","0","98.78","435.66","0","437.83","33.02","0"
"AGAW2","02S","NIN","BOLD:AAC6891","Arthropoda","Arachnida","Araneae","Salticidae",NA,"Phidippus","Phidippus audax","1","1","1","2","0","0","99.75","404","0","450","32","0"
"AGAW2","02W","BAC","TAX:91391","Proteobacteria","Alphaproteobacteria","Rickettsiales","",NA,"","","76","52","52","1","31.58","0","99.08","372.91","0","449.38","39.57","0"
"AGAW2","02W","BAC","TAX:175768","Proteobacteria","Alphaproteobacteria","Rickettsiales","Rickettsiaceae",NA,"Wolbachia","Wolbachia pipientis","10","10","10","2","0","0","98.33","444.6","0","446.4","39.78","0"
"AGAW2","02W","BAC","TAX:91393","Proteobacteria","Alphaproteobacteria","Rickettsiales","Rickettsiaceae",NA,"Wolbachia","","3","3","3","3","0","0","98.72","390.33","0","426.33","39.73","0"
"AGAW2","02W","CHO","no match","no match","no match","no match","no match","no match","no match","no match","no match","no match","no match","no match","no match","no match","no match","no match","no match","no match","no match","no match"
"AGAW2","02W","INS","BOLD:AAG2511","Arthropoda","Insecta","Diptera","Anthomyiidae","Anthomyiinae","Delia","Delia platura","46238","30777","30777","1","33.44","0","99.48","443.37","0","445.61","33.17","0"
"AGAW2","02W","INS","BOLD:AAB2644","Arthropoda","Insecta","Diptera","Chironomidae","Orthocladiinae","Orthocladius","Orthocladius lapponicus, Orthocladius dorenus, Orthocladius mallochi","17440","14210","14210","2","18.52","0","98.85","438.03","0","440.16","32.84","0"
"AGAW2","02W","INS","BOLD:AAB1098","Arthropoda","Insecta","Diptera","Culicidae","Culicinae","Ochlerotatus, Aedes","Aedes excrucians, Aedes cantans, Aedes stimulans, Aedes euedes, Aedes riparius, Aedes annulipes, Ochlerotatus cantans, Aedes fitchii, Aedes sp., Ochlerotatus annulipes, Ochlerotatus excrucians","6824","6058","6058","3","11.23","0","98.6","429.25","0","430.68","35.34","0"
"AGAW2","02W","INS","BOLD:AAN5355","Arthropoda","Insecta","Diptera","Chironomidae","Orthocladiinae","Smittia","Smittia cf. stercoraria, Smittia stercoraria","6647","5301","5301","4","20.25","0","99.04","423.49","0","446.11","36.23","0"
"AGAW2","02W","INS","BOLD:AAO3939","Arthropoda","Insecta","Diptera","Limoniidae","Limoniinae","Dicranomyia","","4771","3369","3369","5","29.39","0","99.31","442.82","0","443.88","36.3","0"
"AGAW2","02W","INS","BOLD:AAP9022","Arthropoda","Insecta","Diptera","Cecidomyiidae","","","","4733","3218","3218","6","32.01","0","99.64","446.81","0","447.86","27.65","0"
"AGAW2","02W","INS","BOLD:AAN5358","Arthropoda","Insecta","Diptera","Chironomidae","Orthocladiinae","Smittia","Smittia stercoraria, Smittia sp. 22ES","2172","1779","1779","7","18.09","0","99.24","443.53","0","445.46","37.28","0"
"AGAW2","02W","INS","BOLD:ACJ0208","Arthropoda","Insecta","Diptera","Chironomidae","Orthocladiinae","","","2074","1529","1529","8","26.28","0","99.11","435.74","0","449.19","34.95","0"
"AGAW2","02W","INS","BOLD:AAM7726","Arthropoda","Insecta","Coleoptera","Curculionidae","Curculioninae","Orchestes","Orchestes steppensis, Orchestes alni","1726","1347","1347","9","21.96","0","99.43","438.83","0","440.39","32.87","0"
"AGAW2","02W","INS","BOLD:AAA3453","Arthropoda","Insecta","Diptera","Anthomyiidae","Anthomyiinae","Delia","","1552","1268","1268","10","18.3","0","99.36","444.28","0","446.09","34.73","0"
"AGAW2","02W","INS","BOLD:ABV1228","Arthropoda","Insecta","Diptera","Chironomidae","Chironominae","Micropsectra","","1487","1038","1038","11","30.2","0","99.61","446.48","0","448.16","34.62","0"
"AGAW2","02W","INS","BOLD:AAN5267","Arthropoda","Insecta","Diptera","Cecidomyiidae","","","","837","598","598","12","28.55","0","99.39","447.75","0","448.82","30.67","0"
"AGAW2","02W","INS","BOLD:ACV3368","","","","","","","","654","481","481","13","26.45","0","99.14","444.59","0","445.6","33.39","0"
"AGAW2","02W","INS","BOLD:AAD5253","Arthropoda","Insecta","Diptera","Chironomidae","Orthocladiinae","Thienemanniella","Thienemanniella xena, Thienemanniella vittata","618","447","447","14","27.67","0","98.26","440.73","0","442.81","34.75","0"
"AGAW2","02W","INS","BOLD:AAH0134","Arthropoda","Insecta","Coleoptera","Phalacridae","Phalacrinae","Stilbus","Stilbus apicialis","458","416","416","15","9.17","0","99.25","432.3","0","433.68","32.1","0"
"AGAW2","02W","INS","BOLD:AAG1010","Arthropoda","Insecta","Diptera","Chironomidae","Orthocladiinae","Paraphaenocladius","","301","284","284","16","5.65","0","98.95","428.29","0","430.45","33.1","0"
"AGAW2","02W","INS","BOLD:AAO8538","Arthropoda","Insecta","Diptera","Psychodidae","Psychodinae","Psychoda","","184","176","176","17","4.35","0","99.08","436.55","0","437.99","32.85","0"
"AGAW2","02W","INS","BOLD:ABA1215","Arthropoda","Insecta","Diptera","Sciaridae","","Lycoriella","Lycoriella sativae","146","127","127","18","13.01","0","99.39","443.21","0","444.35","35.84","0"
"AGAW2","02W","INS","BOLD:ACB1079","Arthropoda","Insecta","Diptera","Cecidomyiidae","Micromyinae","Campylomyza","Campylomyza dilatata, CecidInt44 sp. BOLD:ACB1079","125","98","98","19","21.6","0","98.1","444.01","0","447.3","28.59","0"
"AGAW2","02W","INS","BOLD:AAG9713","Arthropoda","Insecta","Diptera","Dolichopodidae","Sympycninae","Sympycnus","Sympycnus lineatus","107","96","96","20","10.28","0","99.14","444.1","0","445.38","35.52","0"
"AGAW2","02W","INS","BOLD:ACK1651","Arthropoda","Insecta","Diptera","Cecidomyiidae","","","","103","102","102","21","0.97","0","98.78","428.38","0","430.11","26.57","0"
"AGAW2","02W","INS","BOLD:AAN8770","Arthropoda","Insecta","Diptera","Psychodidae","Psychodinae","Psychoda","Psychoda trinodulosa","91","78","78","22","14.29","0","99.04","438.4","0","439.2","33.38","0"
"AGAW2","02W","INS","BOLD:AAZ5599","Arthropoda","Insecta","Diptera","Chironomidae","Orthocladiinae","","","89","78","78","23","12.36","0","99.43","426.88","0","428.31","36.59","0"
"AGAW2","02W","INS","BOLD:AAC4197","Arthropoda","Insecta","Diptera","Chironomidae","Orthocladiinae","Paraphaenocladius","Paraphaenocladius impensus","65","60","60","24","7.69","0","99.19","440.45","0","441.92","34.89","0"
"AGAW2","02W","INS","BOLD:AAG1000","Arthropoda","Insecta","Diptera","Chironomidae","Orthocladiinae","Orthocladius","Orthocladius carlatus","57","57","57","25","0","0","98.02","428","0","430.53","33.84","0"
"AGAW2","02W","INS","BOLD:ABA1231","Arthropoda","Insecta","Coleoptera, Diptera","Sciaridae","","Lycoriella","","54","54","54","26","0","0","99.12","430.31","0","432.61","36.39","0"
"AGAW2","02W","INS","BOLD:AAZ5590","Arthropoda","Insecta","Diptera","Cecidomyiidae","","","","44","44","44","27","0","0","99.13","432.7","0","433.98","26.01","0"
"AGAW2","02W","INS","BOLD:AAY6412","Arthropoda","Insecta","Diptera","Sciaridae","","Bradysia","Bradysia urticae","44","36","36","27","18.18","0","99.44","442.45","0","443.89","32.18","0"
"AGAW2","02W","INS","BOLD:AAU8379","Arthropoda","Insecta","Hymenoptera","Braconidae","Alysiinae","Dinotrema","Dinotrema JFT01","34","32","32","29","5.88","0","99.68","436.47","0","437.56","24.31","0"
"AGAW2","02W","INS","BOLD:AAB9140","Arthropoda","Insecta","Diptera","Calliphoridae","Chrysomyinae","Phormia","Phormia regina","33","33","33","30","0","0","98.88","446.82","0","447.67","33.07","0"
"AGAW2","02W","INS","BOLD:AAN5356","Arthropoda","Insecta","Diptera","Chironomidae","Orthocladiinae","Smittia","","32","30","30","31","6.25","0","98.87","449.19","0","450","34.89","0"
"AGAW2","02W","INS","BOLD:AAL7819","Arthropoda","Insecta","Diptera","Psychodidae","Psychodinae","Psychoda","Psychoda sp., Psychoda sp. BOLD:AAL7819","31","29","29","32","6.45","0","99.27","432.84","0","433.9","32.83","0"
"AGAW2","02W","INS","BOLD:ABU5520","Arthropoda","Insecta","Diptera","Sciaridae","","Corynoptera","","26","25","25","33","3.85","0","99.32","429.08","0","430.08","35.97","0"
"AGAW2","02W","INS","BOLD:ACC7611","Arthropoda","Insecta","Diptera","Cecidomyiidae","","","","22","22","22","34","0","0","97.4","444.55","0","447.32","28.5","0"
"AGAW2","02W","INS","BOLD:AAM9243","Arthropoda","Insecta","Diptera","Sciaridae","","Bradysia","Bradysia nitidicollis","17","17","17","35","0","0","99.45","435.76","0","436.65","36.09","0"
"AGAW2","02W","INS","BOLD:ACF8829","Arthropoda","Insecta","Diptera","Chironomidae","Chironominae","Tanytarsus","","16","16","16","36","0","0","99.09","444.75","0","446.69","36.31","0"
"AGAW2","02W","INS","BOLD:AAN6435","Arthropoda","Insecta","Diptera","Sciaridae","","Bradysia","Bradysia angustipennis","13","12","12","37","7.69","0","99.59","397.69","0","443.46","34.51","0"
"AGAW2","02W","INS","BOLD:AAB2384","Arthropoda","Insecta","Diptera","Syrphidae","Syrphinae","Eupeodes","Eupeodes duseki, Eupeodes volucris, Eupeodes nielseni, Eupeodes americanus, Eupeodes flukei, Eupeodes vockerothi, Eupeodes lucasi, Eupeodes montivagus, Eupeodes rufipunctatus, Eupeodes rojasi, Eupeodes goeldlini, Eupeodes luniger, Eupeodes punctifer, Eupeodes bucculatus, Eupeodes perplexus, Eupeodes abiskoensis, Eupeodes corollae, Eupeodes pomus, Eupeodes lundbecki, Eupeodes nitens, Eupeodes sp. BOLDAAB2384, Eupeodes curtus, Eupeodes latifasciatus","9","9","9","38","0","0","98.36","426","0","427.56","31.13","0"
"AGAW2","02W","INS","BOLD:AAB0868","Arthropoda","Insecta","Diptera","Calliphoridae","Calliphorinae","Cynomya","Cynomya cadaverina, Cynomya mortuorum","9","9","9","38","0","0","99.33","422.33","0","425.22","34.58","0"
"AGAW2","02W","INS","BOLD:ABA6465","Arthropoda","Insecta","Diptera","Chironomidae","Orthocladiinae","","","7","7","7","40","0","0","97.84","448.86","0","450","34.19","0"
"AGAW2","02W","INS","BOLD:AAD0853","Arthropoda","Insecta","Diptera","Scathophagidae","Scathophaginae","Scathophaga","Scathophaga lutaria, Scathophaga hiemalis, Scathophaga furcata, Scathophaga stercoraria, Scathophaga inquinata, Scathophaga suilla","7","7","7","40","0","0","98.21","447.86","0","450","31.94","0"
"AGAW2","02W","INS","BOLD:ABX2489","Arthropoda","Insecta","Coleoptera","Phalacridae","Phalacrinae","Stilbus","","4","4","4","42","0","0","97.67","440.25","0","439.25","32.25","0"
"AGAW2","02W","INS","BOLD:AAD4355","Arthropoda","Insecta","Diptera","Culicidae","Culicinae","Aedes","Aedes provocans","4","4","4","42","0","0","98.53","440","0","444.5","32","0"
"AGAW2","02W","INS","BOLD:ACF2821","Arthropoda","Insecta","Diptera","Mycetophilidae","Mycetophilinae","Mycetophila","Mycetophila fungorum","3","3","3","44","0","0","98.04","408.33","0","409.67","35.22","0"
"AGAW2","02W","INS","BOLD:ABU5546","Arthropoda","Insecta","Diptera","Psychodidae","Psychodinae","Psychoda","","3","3","3","44","0","0","98.54","435.33","0","437","31.39","0"
"AGAW2","02W","INS","BOLD:AAG1742","Arthropoda","Insecta","Diptera","Muscidae","Phaoniinae","Helina","Helina rufitibia","3","3","3","44","0","0","98.73","408.33","0","450","30.67","0"
"AGAW2","02W","INS","BOLD:AAB7513","Arthropoda","Insecta","Diptera","Simuliidae","Simuliinae","Stegopterna","Stegopterna mutata/diplomutata","2","2","2","47","0","0","100","448.5","0","450","37.78","0"
"AGAW2","02W","INS","BOLD:AAK8831","Arthropoda","Insecta","Diptera","Limoniidae","Limoniinae","Metalimnobia, Limonia","","2","2","2","47","0","0","98.77","403","0","404.5","33.13","0"
"AGAW2","02W","INS","BOLD:AAG6743","Arthropoda","Insecta","Diptera","Sarcophagidae","Sarcophaginae","Sarcophaga","Sarcophaga sp. 1 MG, Sarcophaga subvicina","2","2","2","47","0","0","99.33","449","0","450","35.11","0"
"AGAW2","02W","INS","BOLD:ACF8552","Arthropoda","Insecta","Coleoptera","Anthicidae","Anthicinae","Malporus","Malporus formicarius","1","1","1","50","0","0","99.45","365","0","401","34.41","0"
"AGAW2","02W","INS","BOLD:AAL7534","Arthropoda","Insecta","Diptera","Anisopodidae","Anisopodinae","Sylvicola","Sylvicola stackelbergi, Sylvicola stackelbergii 1, Sylvicola punctatus","1","1","1","50","0","0","99.78","450","0","450","34.22","0"
"AGAW2","02W","INS","BOLD:ACM5604","Arthropoda","Insecta","Diptera","Chironomidae","Orthocladiinae","Orthocladius","","1","1","1","50","0","0","99.33","450","0","450","34.89","0"
"AGAW2","02W","INS","BOLD:AAL5267","Arthropoda","Insecta","Coleoptera","Chrysomelidae","Galerucinae","Phyllotreta","Phyllotreta striolata","1","1","1","50","0","0","99.78","450","0","450","39.11","0"
"AGAW2","02W","INS","BOLD:AAB1024","Arthropoda","Insecta","Lepidoptera","Noctuidae","Noctuinae","Pseudohermonassa","Pseudohermonassa bicarnea","1","1","1","50","0","0","100","447","0","450","33.56","0"
"AGAW2","02W","INS","BOLD:ABA1237","Arthropoda","Insecta","Diptera","Chironomidae","Orthocladiinae","","","1","1","1","50","0","0","98.89","449","0","450","34.67","0"
"AGAW2","02W","INS","BOLD:AAG9055","Arthropoda","Insecta","Hymenoptera","Vespidae","Vespinae","Vespula","Vespula germanica","1","1","1","50","0","0","98.89","450","0","450","31.56","0"
"AGAW2","02W","INS","BOLD:AAB5640","Arthropoda","Insecta","Coleoptera","Coccinellidae","Coccinellinae","Harmonia","Harmonia axyridis","1","1","1","50","0","0","100","450","0","450","32.44","0"
"AGAW2","02W","INS","BOLD:ABV0271","Arthropoda","Insecta","Diptera","Chloropidae","Oscinellinae","","","1","1","1","50","0","0","99.78","447","0","450","33.33","0"
"AGAW2","02W","INS","BOLD:AAN5178","Arthropoda","Insecta","Diptera","Cecidomyiidae","Cecidomyiinae","Rhopalomyia","Rhopalomyia sp. BOLD:AAN5178","1","1","1","50","0","0","99.54","433","0","442","27.6","0"
"AGAW2","02W","INS","BOLD:AAN5342","Arthropoda","Insecta","Diptera","Chironomidae","Orthocladiinae","Heleniella","","1","1","1","50","0","0","98.89","450","0","450","35.78","0"
"AGAW2","02W","INS","BOLD:ACF7714","Arthropoda","Insecta","Diptera","Sphaeroceridae","Limosininae","Coproica","Coproica hirtula","1","1","1","50","0","0","97.33","449","0","450","35.56","0"
"AGAW2","02W","INS","BOLD:AAM9252","Arthropoda","Insecta","Diptera","Sciaridae","","Bradysia, Leptosciarella","Bradysia vagans, Leptosciarella zanti","1","1","1","50","0","0","100","449","0","450","39.11","0"
"AGAW2","02W","INS","BOLD:AAM9376","Arthropoda","Insecta","Diptera","Phoridae","Phorinae","Phora","Phora sp. BOLD:AAM9376","1","1","1","50","0","0","100","433","0","450","40.44","0"
"AGAW2","02W","INS","BOLD:AAG2462","Arthropoda","Insecta","Diptera","Anthomyiidae","Anthomyiinae","Botanophila","Botanophila fugax","1","1","1","50","0","0","97.29","443","0","450","31.33","0"
"AGAW2","02W","NAR","no match","no match","no match","no match","no match","no match","no match","no match","no match","no match","no match","no match","no match","no match","no match","no match","no match","no match","no match","no match"
"AGAW2","02W","NIN","BOLD:AAB7667","Arthropoda","Arachnida","Araneae","Lycosidae",NA,"Pardosa","Pardosa saxatilis","417","396","396","1","5.04","0","99.39","422.74","0","428.5","31.37","0"
"AGAW2","02W","NIN","BOLD:ABX0027","Arthropoda","Collembola","Symphypleona","Bourletiellidae",NA,"Bourletiella","","152","129","129","2","15.13","0","99.59","415.08","0","442.38","35.6","0"
"AGAW2","02W","NIN","BOLD:AAN6707","Arthropoda","Arachnida","Mesostigmata","Ascidae",NA,"Arctoseius","","51","50","50","3","1.96","0","99.32","444.75","0","448.61","42.35","0"
"AGAW2","02W","NIN","BOLD:AAB7915","Arthropoda","Collembola","Symphypleona","Bourletiellidae",NA,"Deuterosminthurus","Deuterosminthurus sp1","12","12","12","4","0","0","99.29","443.5","0","444.58","35.12","0"
"AGAW2","02W","NIN","BOLD:ACU6668","Arthropoda","Collembola","Symphypleona","Bourletiellidae",NA,"","","2","2","2","5","0","0","99.77","439.5","0","442.5","34.58","0"
"AGAW2","02W","NIN","BOLD:AAN6711","Arthropoda","Arachnida","Mesostigmata","Parasitidae",NA,"","","1","1","1","6","0","0","98.67","450","0","450","39.33","0"
"AGAW2","02W","NIN","BOLD:ACW2271","Arthropoda","Collembola","Symphypleona","Bourletiellidae",NA,"Bourletiella","","1","1","1","6","0","0","97.11","450","0","450","35.78","0"
"AGAW2","02W","NIN","BOLD:ACF8021","Arthropoda","Arachnida","Mesostigmata","Ascidae",NA,"Arctoseius","Arctoseius cetratus","1","1","1","6","0","0","97.77","448","0","450","37.78","0"
"AGAW2","05E","BAC","TAX:175768","Proteobacteria","Alphaproteobacteria","Rickettsiales","Rickettsiaceae",NA,"Wolbachia","Wolbachia pipientis","4","4","4","1","0","0","99.28","449.25","0","449.75","39.74","0"
"AGAW2","05E","CHO","no match","no match","no match","no match","no match","no match","no match","no match","no match","no match","no match","no match","no match","no match","no match","no match","no match","no match","no match","no match"
"AGAW2","05E","INS","BOLD:AAA2815","Arthropoda","Insecta","Lepidoptera","Noctuidae","Noctuinae","Mythimna","Mythimna yukonensis, Mythimna oxygala","15746","13197","13197","1","16.19","0","99.15","421.34","0","422.89","33.37","0"
"AGAW2","05E","INS","BOLD:AAG2511","Arthropoda","Insecta","Diptera","Anthomyiidae","Anthomyiinae","Delia","Delia platura","15420","12569","12569","2","18.49","0","99.02","444.58","0","447.09","33.2","0"
"AGAW2","05E","INS","BOLD:AAA3453","Arthropoda","Insecta","Diptera","Anthomyiidae","Anthomyiinae","Delia","","8175","7200","7200","3","11.93","0","99.22","431.52","0","433.86","34.65","0"
"AGAW2","05E","INS","BOLD:AAA8933","Arthropoda","Insecta","Coleoptera","Coccinellidae","Coccinellinae","Coccinella","Coccinella septempunctata","4951","3785","3785","4","23.55","0","99.35","442.8","0","444.65","32.93","0"
"AGAW2","05E","INS","BOLD:AAB2384","Arthropoda","Insecta","Diptera","Syrphidae","Syrphinae","Eupeodes","Eupeodes duseki, Eupeodes volucris, Eupeodes nielseni, Eupeodes americanus, Eupeodes flukei, Eupeodes vockerothi, Eupeodes lucasi, Eupeodes montivagus, Eupeodes rufipunctatus, Eupeodes rojasi, Eupeodes goeldlini, Eupeodes luniger, Eupeodes punctifer, Eupeodes bucculatus, Eupeodes perplexus, Eupeodes abiskoensis, Eupeodes corollae, Eupeodes pomus, Eupeodes lundbecki, Eupeodes nitens, Eupeodes sp. BOLDAAB2384, Eupeodes curtus, Eupeodes latifasciatus","3190","3086","3086","5","3.26","0","98.75","387.17","0","395.25","30.91","0"
"AGAW2","05E","INS","BOLD:AAF9187","Arthropoda","Insecta","Coleoptera","Curculionidae","Entiminae","Phyllobius","Phyllobius oblongus","1399","1143","1143","6","18.3","0","99.44","436.72","0","440.12","38.86","0"
"AGAW2","05E","INS","BOLD:AAA4121","Arthropoda","Insecta","Diptera","Simuliidae","Simuliinae","Simulium","Simulium vittatum complex, Simulium vittatum, Simulium sp. BOLDAAA4121, Simulium argus, Simulium vittatum s.str.","1340","1038","1038","7","22.54","0","98.32","436.95","0","438.84","33.83","0"
"AGAW2","05E","INS","BOLD:ACT0257","","","","","","","","829","738","738","8","10.98","0","98.68","430.95","0","432.31","37.68","0"
"AGAW2","05E","INS","BOLD:AAO3939","Arthropoda","Insecta","Diptera","Limoniidae","Limoniinae","Dicranomyia","","721","648","648","9","10.12","0","98.98","437.41","0","439.29","36.27","0"
"AGAW2","05E","INS","BOLD:AAB1098","Arthropoda","Insecta","Diptera","Culicidae","Culicinae","Ochlerotatus, Aedes","Aedes excrucians, Aedes cantans, Aedes stimulans, Aedes euedes, Aedes riparius, Aedes annulipes, Ochlerotatus cantans, Aedes fitchii, Aedes sp., Ochlerotatus annulipes, Ochlerotatus excrucians","661","610","610","10","7.72","0","98.7","430.41","0","431.63","35.32","0"
"AGAW2","05E","INS","BOLD:AAP6369","Arthropoda","Insecta","Coleoptera","Elateridae","Agrypninae","Aeolus","Aeolus mellillus","603","455","455","11","24.54","0","99.7","440.61","0","441.71","38.97","0"
"AGAW2","05E","INS","BOLD:AAB4657","Arthropoda","Insecta","Diptera","Chironomidae","Chironominae","Chironomus","Chironomus maturus","534","467","467","12","12.55","0","99.31","441.08","0","444.02","37.48","0"
"AGAW2","05E","INS","BOLD:AAF9014","Arthropoda","Insecta","Diptera","Limoniidae","Chioneinae","Symplecta","Symplecta cana, Symplecta sp. BOLDAAF9014","371","354","354","13","4.58","0","97.98","430.31","0","438.24","33.78","0"
"AGAW2","05E","INS","BOLD:AAF9771","Arthropoda","Insecta","Diptera","Hybotidae","Tachydromiinae","Platypalpus","Platypalpus longiseta, Platypalpus trivialis, Platypalpus sp. 5, Platypalpus sp. 17, Platypalpus pallidiventris","297","221","221","14","25.59","0","98.95","445.63","0","446.19","34.93","0"
"AGAW2","05E","INS","BOLD:AAI4303","Arthropoda","Insecta","Diptera","Chironomidae","Chironominae","Chironomus","Chironomus melanescens","242","235","235","15","2.89","0","98.37","428.69","0","433.06","33.8","0"
"AGAW2","05E","INS","BOLD:AAB2644","Arthropoda","Insecta","Diptera","Chironomidae","Orthocladiinae","Orthocladius","Orthocladius lapponicus, Orthocladius dorenus, Orthocladius mallochi","236","210","210","16","11.02","0","99.25","442.94","0","444.84","32.23","0"
"AGAW2","05E","INS","BOLD:ACF2924","","","","","","","","226","180","180","17","20.35","0","99.61","444.9","0","445.84","25.59","0"
"AGAW2","05E","INS","BOLD:AAD0484","Arthropoda","Insecta","Diptera","Chironomidae","","","","163","146","146","18","10.43","0","98.75","446.87","0","447.83","34.49","0"
"AGAW2","05E","INS","BOLD:AAH3664","Arthropoda","Insecta","Diptera","Cecidomyiidae","","","CecidInt3 sp. BOLD:AAH3664","147","143","143","19","2.72","0","98.64","435.14","0","436.5","27.23","0"
"AGAW2","05E","INS","BOLD:AAN5342","Arthropoda","Insecta","Diptera","Chironomidae","Orthocladiinae","Heleniella","","145","139","139","20","4.14","0","98.9","420.81","0","428.43","35.12","0"
"AGAW2","05E","INS","BOLD:ACX5521","Arthropoda","Insecta","Diptera","Chironomidae","Orthocladiinae","","","145","145","145","20","0","0","98.52","397.14","0","399.18","34.54","0"
"AGAW2","05E","INS","BOLD:AAQ0265","Arthropoda","Insecta","Diptera","Hybotidae","Tachydromiinae","Platypalpus","Platypalpus pulicarius","143","138","138","22","3.5","0","99.34","408.36","0","410.24","34.68","0"
"AGAW2","05E","INS","BOLD:AAN6447","Arthropoda","Insecta","Diptera","Sciaridae","","Corynoptera","Corynoptera perpusilla","140","96","96","23","31.43","0","99.49","448.76","0","449.63","36.38","0"
"AGAW2","05E","INS","BOLD:AAV1261","Arthropoda","Insecta","Diptera","Sciaridae","","Bradysia","Bradysia splendida","84","83","83","24","1.19","0","99.25","430.56","0","431.93","33.97","0"
"AGAW2","05E","INS","BOLD:ACK7296","Arthropoda","Insecta","Diptera","Cecidomyiidae","","","","78","74","74","25","5.13","0","99.4","429.71","0","430.81","24.24","0"
"AGAW2","05E","INS","BOLD:ACU7906","Arthropoda","Insecta","Diptera","Chironomidae","Orthocladiinae","","","67","67","67","26","0","0","98.81","407.94","0","411.91","36.1","0"
"AGAW2","05E","INS","BOLD:AAN8770","Arthropoda","Insecta","Diptera","Psychodidae","Psychodinae","Psychoda","Psychoda trinodulosa","56","48","48","27","14.29","0","99.36","442.75","0","443.98","33.94","0"
"AGAW2","05E","INS","BOLD:AAF9768","Arthropoda","Insecta","Diptera","Hybotidae","Tachydromiinae","Platypalpus","Platypalpus holosericus, Platypalpus candicans, Platypalpus sp. BOLD:AAF9768, Platypalpus sp. 22","41","35","35","28","14.63","0","99.34","445.05","0","445.54","31.86","0"
"AGAW2","05E","INS","BOLD:AAN5205","Arthropoda","Insecta","Diptera","Cecidomyiidae","","","CecidInt42 sp. BOLD:AAN5205","35","29","29","29","17.14","0","99.52","448.97","0","450","31.47","0"
"AGAW2","05E","INS","BOLD:ABA1215","Arthropoda","Insecta","Diptera","Sciaridae","","Lycoriella","Lycoriella sativae","21","20","20","30","4.76","0","99.28","439.81","0","441.43","35.85","0"
"AGAW2","05E","INS","BOLD:AAG1000","Arthropoda","Insecta","Diptera","Chironomidae","Orthocladiinae","Orthocladius","Orthocladius carlatus","21","21","21","30","0","0","98.12","421.48","0","423.19","33.65","0"
"AGAW2","05E","INS","BOLD:AAN5356","Arthropoda","Insecta","Diptera","Chironomidae","Orthocladiinae","Smittia","","18","15","15","32","16.67","0","98.9","438.5","0","439.89","35.02","0"
"AGAW2","05E","INS","BOLD:AAG2868","Arthropoda","Insecta","Hemiptera","Cicadellidae","Typhlocybinae","Empoasca","Empoasca sp., Empoasca fabae","15","15","15","33","0","0","99.84","449.4","0","450","32.52","0"
"AGAW2","05E","INS","BOLD:AAY3269","Arthropoda","Insecta","Diptera","Chironomidae","Diamesinae","Potthastia","Potthastia gaedii","14","14","14","34","0","0","98.17","444.79","0","446.43","34.46","0"
"AGAW2","05E","INS","BOLD:AAA4222","Arthropoda","Insecta","Lepidoptera","Erebidae","Hypeninae","Hypena","Hypena scabra","13","13","13","35","0","0","99.05","394.92","0","397.15","32.35","0"
"AGAW2","05E","INS","BOLD:AAH0134","Arthropoda","Insecta","Coleoptera","Phalacridae","Phalacrinae","Stilbus","Stilbus apicialis","12","12","12","36","0","0","98.74","427.08","0","427.67","32.02","0"
"AGAW2","05E","INS","BOLD:AAU8733","Arthropoda","Insecta","Hymenoptera","Ichneumonidae","Tersilochinae","","","11","9","9","37","18.18","0","99.61","427.09","0","428.18","29.28","0"
"AGAW2","05E","INS","BOLD:AAE4298","Arthropoda","Insecta","Diptera","Chironomidae","Orthocladiinae","Cricotopus","Cricotopus tremulus","11","11","11","37","0","0","98.06","409.82","0","425.27","32.52","0"
"AGAW2","05E","INS","BOLD:AAU6967","Arthropoda","Insecta","Coleoptera","Chrysomelidae","Galerucinae","Psylliodes","Psylliodes affinis","10","10","10","39","0","0","99.41","438.7","0","450","35.04","0"
"AGAW2","05E","INS","BOLD:AAA9899","Arthropoda","Insecta","Hemiptera","Aphididae","Aphidinae","Rhopalosiphum","Rhopalosiphum padi","9","9","9","40","0","0","99.58","425.56","0","426.89","25.48","0"
"AGAW2","05E","INS","BOLD:AAV1136","Arthropoda","Insecta","Diptera","Scatopsidae","","","","8","8","8","41","0","0","98.15","430.25","0","430.38","35.71","0"
"AGAW2","05E","INS","BOLD:AAG2462","Arthropoda","Insecta","Diptera","Anthomyiidae","Anthomyiinae","Botanophila","Botanophila fugax","7","7","7","42","0","0","99.84","435.43","0","436.71","31.7","0"
"AGAW2","05E","INS","BOLD:AAA7374","Arthropoda","Insecta","Diptera","Syrphidae","Syrphinae","Sphaerophoria","Sphaerophoria bankowskae, Sphaerophoria infuscata, Sphaerophoria nr. philanthus, Sphaerophoria batava, Sphaerophoria sp., Sphaerophoria rueppellii, Sphaerophoria interrupta-group, Sphaerophoria sp. MA-1, Sphaerophoria cleoae, Sphaerophoria asymmetrica, Sphaerophoria fatarum, Sphaerophoria bengalensis, Sphaerophoria bifurcata, Sphaerophoria taeniata, Sphaerophoria interrupta, Sphaerophoria contigua, Sphaerophoria kaa, Sphaerophoria laurae, Sphaerophoria chongjini, Sphaerophoria boreoalpina, Sphaerophoria sp. BOLDAAA7374, Sphaerophoria macrogaster, Sphaerophoria sulphuripes, Sphaerophoria brevipilosa, Sphaerophoria philanthus, Sphaerophoria pyrrhina, Sphaerophoria abbreviata, Sphaerophoria scripta, Sphaerophoria potentillae, Sphaerophoria longipilosa, Sphaerophoria virgata","7","7","7","42","0","0","99.47","418.57","0","421","30.82","0"
"AGAW2","05E","INS","BOLD:AAM9252","Arthropoda","Insecta","Diptera","Sciaridae","","Bradysia, Leptosciarella","Bradysia vagans, Leptosciarella zanti","7","7","7","42","0","0","98.47","441.71","0","441.86","39.05","0"
"AGAW2","05E","INS","BOLD:ACK3431","Arthropoda","Insecta","Diptera","Cecidomyiidae","","","","5","5","5","45","0","0","99.3","425.6","0","427.6","32.13","0"
"AGAW2","05E","INS","BOLD:AAA9251","Arthropoda","Insecta","Lepidoptera","Gelechiidae","Gelechiinae","Scrobipalpa","Scrobipalpa atriplicella, Scrobipalpa nr. atriplicella sp. 1","5","5","5","45","0","0","98.83","413.2","0","414.6","34.31","0"
"AGAW2","05E","INS","BOLD:ACF5939","Arthropoda","Insecta","Diptera","Tachinidae","Exoristinae","Winthemia","Winthemia sinuata, Winthemia illinoensis","5","5","5","45","0","0","99.31","429","0","430.2","29.64","0"
"AGAW2","05E","INS","BOLD:ABX6186","Arthropoda","Insecta","Diptera","Tipulidae","Tipulinae","Nephrotoma","Nephrotoma ferruginea","4","4","4","48","0","0","98.66","447.75","0","450","34.28","0"
"AGAW2","05E","INS","BOLD:ABA0863","Arthropoda","Insecta","Diptera","Cecidomyiidae","","","","3","3","3","49","0","0","99.48","447.33","0","450","30","0"
"AGAW2","05E","INS","BOLD:AAU6592","Arthropoda","Insecta","Diptera","Ceratopogonidae","Dasyheleinae","Dasyhelea","Dasyhelea arenivaga","3","3","3","49","0","0","98.9","394.67","0","397.33","34.31","0"
"AGAW2","05E","INS","BOLD:AAH3299","Arthropoda","Insecta","Coleoptera","Coccinellidae","Coccinellinae","Hippodamia","Hippodamia variegata","3","3","3","49","0","0","99.7","450","0","450","32.81","0"
"AGAW2","05E","INS","BOLD:ADH4889","Arthropoda","Insecta","Diptera","","","","","3","3","3","49","0","0","98.81","449","0","450","35.48","0"
"AGAW2","05E","INS","BOLD:AAB9140","Arthropoda","Insecta","Diptera","Calliphoridae","Chrysomyinae","Phormia","Phormia regina","3","3","3","49","0","0","98.96","448","0","450","33.33","0"
"AGAW2","05E","INS","BOLD:AAD0853","Arthropoda","Insecta","Diptera","Scathophagidae","Scathophaginae","Scathophaga","Scathophaga lutaria, Scathophaga hiemalis, Scathophaga furcata, Scathophaga stercoraria, Scathophaga inquinata, Scathophaga suilla","3","3","3","49","0","0","98.12","445.33","0","447.67","31.94","0"
"AGAW2","05E","INS","BOLD:ADH6290","","","","","","","","3","3","3","49","0","0","99.31","440.33","0","440.67","38.8","0"
"AGAW2","05E","INS","BOLD:AAF9041","Arthropoda","Insecta","Diptera","Tipulidae","Tipulinae","Tipula","Tipula oleracea","3","3","3","49","0","0","99","401.67","0","402.67","35.51","0"
"AGAW2","05E","INS","BOLD:AAC9062","Arthropoda","Insecta","Diptera","Culicidae","Culicinae","Ochlerotatus, Aedes","Aedes fitchii, Ochlerotatus fitchii, Aedes grossbecki, Aedes ventrovittis","3","3","3","49","0","0","99","431","0","431.67","33.18","0"
"AGAW2","05E","INS","BOLD:AAH4135","Arthropoda","Insecta","Diptera","Chloropidae","Chloropinae","Thaumatomyia","Thaumatomyia glabra, Thaumatomyia grata, Thaumatomyia sp. 1 TAW","2","2","2","58","0","0","98.11","449","0","450","32.44","0"
"AGAW2","05E","INS","BOLD:AAN5355","Arthropoda","Insecta","Diptera","Chironomidae","Orthocladiinae","Smittia","Smittia cf. stercoraria, Smittia stercoraria","2","2","2","58","0","0","99.11","391.5","0","425.5","35.93","0"
"AGAW2","05E","INS","BOLD:AAG6743","Arthropoda","Insecta","Diptera","Sarcophagidae","Sarcophaginae","Sarcophaga","Sarcophaga sp. 1 MG, Sarcophaga subvicina","2","2","2","58","0","0","98.88","448","0","450","34.89","0"
"AGAW2","05E","INS","BOLD:AAB7490","Arthropoda","Insecta","Diptera","Chironomidae","Orthocladiinae","Tvetenia","Tvetenia paucunca","2","2","2","58","0","0","98.55","449","0","450","34.22","0"
"AGAW2","05E","INS","BOLD:ACK3142","Arthropoda","Insecta","Diptera","Cecidomyiidae","","","","2","2","2","58","0","0","98.59","422.5","0","425","27.71","0"
"AGAW2","05E","INS","BOLD:ABU5520","Arthropoda","Insecta","Diptera","Sciaridae","","Corynoptera","","2","2","2","58","0","0","98.16","415.5","0","419.5","35.39","0"
"AGAW2","05E","INS","BOLD:AAB7030","Arthropoda","Insecta","Diptera","Chironomidae","Chironominae","Chironomus","Chironomus sp. BOLDAAP2996, Chironomus quinnitukqut, Chironomus cf. decorus 2g, Chironomus decorus group sp. 2 IP2013, Chironomus cf. decorus, Chironomus decorus gr.","2","2","2","58","0","0","98.97","438","0","450","35.22","0"
"AGAW2","05E","INS","BOLD:AAN5260","Arthropoda","Insecta","Diptera","Cecidomyiidae","","","","2","2","2","58","0","0","99.88","395.5","0","396.5","25.86","0"
"AGAW2","05E","INS","BOLD:AAV5796","Arthropoda","Insecta","Diptera","Cecidomyiidae","","","","2","2","2","58","0","0","98.89","447.5","0","450","29","0"
"AGAW2","05E","INS","BOLD:AAD4355","Arthropoda","Insecta","Diptera","Culicidae","Culicinae","Aedes","Aedes provocans","2","2","2","58","0","0","98.55","448","0","450","32.56","0"
"AGAW2","05E","INS","BOLD:AAM7726","Arthropoda","Insecta","Coleoptera","Curculionidae","Curculioninae","Orchestes","Orchestes steppensis, Orchestes alni","2","2","2","58","0","0","99.22","449.5","0","450","33.56","0"
"AGAW2","05E","INS","BOLD:AAG1701","Arthropoda","Insecta","Diptera","Muscidae","Coenosiinae","Macrorchis","Macrorchis ausoba","2","2","2","58","0","0","99.08","434.5","0","437.5","32.46","0"
"AGAW2","05E","INS","BOLD:AAG4653","Arthropoda","Insecta","Diptera","Syrphidae","Syrphinae","Heringia","Heringia latitarsis, Heringia pubescens, Heringia jakutorum, Heringia rita, Heringia fulvimanus, Heringia sp. BOLDAAG4653, Heringia nr. pubescens, Heringia coxalis, Heringia larusi, Heringia vitripennis","2","2","2","58","0","0","99.44","450","0","450","31.56","0"
"AGAW2","05E","INS","BOLD:ABV1229","Arthropoda","Insecta","Diptera","Chironomidae","Orthocladiinae","Eukiefferiella","","2","2","2","58","0","0","98.88","448.5","0","450","35.78","0"
"AGAW2","05E","INS","BOLD:ADH9310","Arthropoda","Insecta","Diptera","Anthomyiidae","Anthomyiinae","Anthomyia","Anthomyia procellaris, Anthomyia pluvialis","2","2","2","58","0","0","98.5","417.5","0","420.5","32.19","0"
"AGAW2","05E","INS","BOLD:AAH3548","Arthropoda","Insecta","Diptera","Sphaeroceridae","Copromyzinae","Lotophila","Lotophila sp., Lotophila atra","1","1","1","73","0","0","99.56","450","0","450","32.44","0"
"AGAW2","05E","INS","BOLD:AAG1717","Arthropoda","Insecta","Diptera","Muscidae","Phaoniinae","Helina","Helina troene","1","1","1","73","0","0","98","449","0","450","31.33","0"
"AGAW2","05E","INS","BOLD:AAA2590","Arthropoda","Insecta","Lepidoptera","Noctuidae","Noctuinae","Xestia","Xestia baja, Xestia smithii","1","1","1","73","0","0","100","410","0","410","32.44","0"
"AGAW2","05E","INS","BOLD:AAD7145","Arthropoda","Insecta","Diptera","Muscidae","Mydaeinae","Myospila","Myospila meditabunda","1","1","1","73","0","0","99.5","403","0","405","30.37","0"
"AGAW2","05E","INS","BOLD:ACE0699","Arthropoda","Insecta","Diptera","Chironomidae","","","","1","1","1","73","0","0","98.84","430","0","450","36.67","0"
"AGAW2","05E","INS","BOLD:ABV1375","Arthropoda","Insecta","Diptera","Cecidomyiidae","","","CecidInt3 sp. BOLD:ABV1375","1","1","1","73","0","0","97.36","416","0","418","25.84","0"
"AGAW2","05E","INS","BOLD:AAN5219","Arthropoda","Insecta","Diptera","Cecidomyiidae","","","","1","1","1","73","0","0","99","401","0","403","28.29","0"
"AGAW2","05E","INS","BOLD:ACU4508","Arthropoda","Insecta","Diptera","Agromyzidae","","","","1","1","1","73","0","0","97.27","366","0","375","32.27","0"
"AGAW2","05E","INS","BOLD:AAC9132","Arthropoda","Insecta","Diptera","Culicidae","Culicinae","Culiseta","Culiseta inornata","1","1","1","73","0","0","98.2","388","0","390","33.33","0"
"AGAW2","05E","INS","BOLD:AAC4403","Arthropoda","Insecta","Diptera","Chironomidae","Chironominae","Micropsectra","Micropsectra polita","1","1","1","73","0","0","98.1","369","0","371","34.5","0"
"AGAW2","05E","INS","BOLD:AAU8613","Arthropoda","Insecta","Hymenoptera","Ichneumonidae","Diplazontinae","Sussaba","Sussaba dorsalis","1","1","1","73","0","0","99.55","449","0","450","30.22","0"
"AGAW2","05E","INS","BOLD:AAA3817","Arthropoda","Insecta","Lepidoptera","Geometridae","Larentiinae","Xanthorhoe","Xanthorhoe ferrugata","1","1","1","73","0","0","98.09","418","0","418","32.06","0"
"AGAW2","05E","INS","BOLD:ABZ7601","Arthropoda","Insecta","Lepidoptera","Noctuidae","Noctuinae","Anarta","Anarta trifolii","1","1","1","73","0","0","99.56","450","0","450","30.67","0"
"AGAW2","05E","INS","BOLD:AAH0127","Arthropoda","Insecta","Coleoptera","Pyrochroidae","Pedilinae","Pedilus","Pedilus lugubris","1","1","1","73","0","0","100","398","0","437","32.72","0"
"AGAW2","05E","INS","BOLD:ACL7849","","","","","","","","1","1","1","73","0","0","99.73","377","0","415","32.05","0"
"AGAW2","05E","INS","BOLD:AAA5299","Arthropoda","Insecta","Diptera","Chironomidae","Orthocladiinae","Cricotopus","Cricotopus laetus, Cricotopus relucens, Cricotopus trifasciatus, Cricotopus sp., Cricotopus sp. 3ES, Cricotopus sylvestris, Cricotopus glacialis","1","1","1","73","0","0","99.55","449","0","450","35.56","0"
"AGAW2","05E","INS","BOLD:AAG0991","Arthropoda","Insecta","Diptera","Chironomidae","Orthocladiinae","Orthocladius","","1","1","1","73","0","0","99.18","367","0","372","32.53","0"
"AGAW2","05E","INS","BOLD:AAB5609","Arthropoda","Insecta","Diptera","Muscidae","Coenosiinae","Coenosia","Coenosia sp.BOLD:AAB5609, Coenosia tigrina","1","1","1","73","0","0","100","374","0","375","32.8","0"
"AGAW2","05E","INS","BOLD:AAA2482","Arthropoda","Insecta","Lepidoptera","Noctuidae","Noctuinae","Mythimna","Mythimna unipuncta","1","1","1","73","0","0","98.44","450","0","450","31.56","0"
"AGAW2","05E","INS","BOLD:AAG8493","Arthropoda","Insecta","Diptera","Drosophilidae","Drosophilinae","Scaptomyza","Scaptomyza pallida","1","1","1","73","0","0","98","450","0","450","34","0"
"AGAW2","05E","INS","BOLD:AAC0903","Arthropoda","Insecta","Diptera","Chironomidae","Chironominae","Chironomus","Chironomus acidophilus","1","1","1","73","0","0","99.56","450","0","450","36.44","0"
"AGAW2","05E","INS","BOLD:AAU6630","Arthropoda","Insecta","Diptera","Fanniidae","Fanniinae","Fannia","Fannia armata","1","1","1","73","0","0","99.54","437","0","438","34.25","0"
"AGAW2","05E","INS","BOLD:AAM7064","Arthropoda","Insecta","Diptera","Chironomidae","Orthocladiinae","Smittia","Smittia sp. 14ES","1","1","1","73","0","0","98.64","442","0","442","32.35","0"
"AGAW2","05E","INS","BOLD:AAU6970","Arthropoda","Insecta","Coleoptera","Cleridae","Clerinae","Enoclerus","Enoclerus nigripes","1","1","1","73","0","0","98","449","0","450","34.22","0"
"AGAW2","05E","INS","BOLD:AAA9101","Arthropoda","Insecta","Lepidoptera","Pyralidae","Phycitinae","Moodna, Caudellia","Moodna ostrinella, Moodna bisinuella, Caudellia nigrella","1","1","1","73","0","0","99.56","450","0","450","34.22","0"
"AGAW2","05E","INS","BOLD:AAU8900","Arthropoda","Insecta","Hymenoptera","Braconidae","Euphorinae","Pygostolus","Pygostolus falcatus","1","1","1","73","0","0","98.17","382","0","386","26.17","0"
"AGAW2","05E","INS","BOLD:AAG8091","Arthropoda","Insecta","Hymenoptera","Ichneumonidae","Campopleginae","Campoletis","Campoletis sp. AB.2","1","1","1","73","0","0","97.11","450","0","450","24.67","0"
"AGAW2","05E","INS","BOLD:AAH0145","Arthropoda","Insecta","Coleoptera","Carabidae","Harpalinae","Anisodactylus","Anisodactylus pitychrous, Anisodactylus sanctaecrucis, Anisodactylus alternans, Anisodactylus discoideus","1","1","1","73","0","0","97.11","450","0","450","33.11","0"
"AGAW2","05E","INS","BOLD:ABA3960","Arthropoda","Insecta","Coleoptera","Chrysomelidae","Eumolpinae","Paria","Paria fragariae","1","1","1","73","0","0","100","444","0","450","36.67","0"
"AGAW2","05E","INS","BOLD:ACF2821","Arthropoda","Insecta","Diptera","Mycetophilidae","Mycetophilinae","Mycetophila","Mycetophila fungorum","1","1","1","73","0","0","98.65","444","0","450","36","0"
"AGAW2","05E","INS","BOLD:AAH2876","Arthropoda","Insecta","Diptera","Cecidomyiidae","","","CecidInt51 sp. BOLD:AAH2876","1","1","1","73","0","0","99.78","450","0","450","26.67","0"
"AGAW2","05E","INS","BOLD:AAQ0582","Arthropoda","Insecta","Diptera","Anthomyiidae","Anthomyiinae","Lasiomma","","1","1","1","73","0","0","99.42","344","0","351","30.77","0"
"AGAW2","05E","INS","BOLD:ACJ7008","Arthropoda","Insecta","Diptera","Sciaridae","","Leptosciarella","Leptosciarella belingi","1","1","1","73","0","0","98.57","350","0","392","32.14","0"
"AGAW2","05E","INS","BOLD:AAF1049","Arthropoda","Insecta","Diptera","Agromyzidae","Phytomyzinae","Cerodontha","Cerodontha fasciata","1","1","1","73","0","0","100","402","0","450","31.56","0"
"AGAW2","05E","INS","BOLD:AAA3654","Arthropoda","Insecta","Lepidoptera","Crambidae","Crambinae","Crambus","Crambus bigelovi, Crambus johnsoni, Crambus praefectellus","1","1","1","73","0","0","99.11","449","0","450","32","0"
"AGAW2","05E","INS","BOLD:AAC4197","Arthropoda","Insecta","Diptera","Chironomidae","Orthocladiinae","Paraphaenocladius","Paraphaenocladius impensus","1","1","1","73","0","0","100","412","0","413","34.14","0"
"AGAW2","05E","INS","BOLD:ACF7232","Arthropoda","Insecta","Diptera","Chironomidae","Orthocladiinae","Parakiefferiella","","1","1","1","73","0","0","99.48","383","0","450","37.56","0"
"AGAW2","05E","INS","BOLD:AAF4817","Arthropoda","Insecta","Diptera","Chironomidae","Orthocladiinae","Smittia","Smittia edwardsi","1","1","1","73","0","0","97.18","390","0","450","33.11","0"
"AGAW2","05E","INS","BOLD:ACB0984","Arthropoda","Insecta","Psocodea","Stenopsocidae","","Graphopsocus","Graphopsocus cruciatus","1","1","1","73","0","0","99.03","413","0","450","37.56","0"
"AGAW2","05E","INS","BOLD:ABV0255","Arthropoda","Insecta","Diptera","Chironomidae","Orthocladiinae","Limnophyes","","1","1","1","73","0","0","98.89","449","0","450","34.89","0"
"AGAW2","05E","INS","BOLD:AAA5803","Arthropoda","Insecta","Hemiptera","Miridae","Mirinae","Lygus","Lygus lineolaris","1","1","1","73","0","0","99.33","449","0","450","34.22","0"
"AGAW2","05E","INS","BOLD:ABW1310","Arthropoda","Insecta","Diptera","Anthomyiidae","","","","1","1","1","73","0","0","99.78","448","0","450","36.67","0"
"AGAW2","05E","INS","BOLD:ACD3588","Arthropoda","Insecta","Diptera","Sciaridae","","Lycoriella","Lycoriella sativae","1","1","1","73","0","0","99.33","450","0","450","36.89","0"
"AGAW2","05E","NAR","no match","no match","no match","no match","no match","no match","no match","no match","no match","no match","no match","no match","no match","no match","no match","no match","no match","no match","no match","no match"
"AGAW2","05E","NIN","BOLD:ABX0027","Arthropoda","Collembola","Symphypleona","Bourletiellidae","","Bourletiella","","80","66","66","1","17.5","0","99.6","417.15","0","445.06","35.62","0"
"AGAW2","05E","NIN","BOLD:AAB7915","Arthropoda","Collembola","Symphypleona","Bourletiellidae","","Deuterosminthurus","Deuterosminthurus sp1","14","14","14","2","0","0","99.21","441.43","0","442.79","35.27","0"
"AGAW2","05E","NIN","BOLD:AAV0589","Arthropoda","Arachnida","Mesostigmata","Blattisociidae","Platyseiinae","Cheiroseius","","8","8","8","3","0","0","99.36","447.5","0","448","27.68","0"
"AGAW2","05E","NIN","BOLD:AAN6711","Arthropoda","Arachnida","Mesostigmata","Parasitidae","","","","7","6","6","4","14.29","0","98.92","449.71","0","450","38.76","0"
"AGAW2","05E","NIN","BOLD:AAH6630","Arthropoda","Collembola","Entomobryomorpha","Entomobryidae","Entomobryinae","Entomobrya","Entomobrya unostrigata","5","5","5","5","0","0","98.96","416.6","0","417.8","39.24","0"
"AGAW2","05E","NIN","BOLD:ACZ1295","Arthropoda","Arachnida","Trombidiformes","Anystidae","","","","1","1","1","6","0","0","100","389","0","450","32.22","0"
"AGAW2","05E","NIN","BOLD:ABW2639","Arthropoda","Arachnida","Trombidiformes","Anystidae","","","","1","1","1","6","0","0","99.59","485","0","450","31.78","0"
"AGAW2","05E","NIN","BOLD:ACC0595","Arthropoda","Collembola","Entomobryomorpha","Entomobryidae","Entomobryinae","Entomobrya","","1","1","1","6","0","0","98","449","0","450","38.22","0"
"AGAW2","05E","NIN","BOLD:AAB7914","Arthropoda","Collembola","Symphypleona","Bourletiellidae","","Bourletiella","","1","1","1","6","0","0","97.54","447","0","450","38","0"
"AGAW2","05N","BAC","TAX:375853","Proteobacteria","Alphaproteobacteria","Rickettsiales","Rickettsiaceae",NA,"Wolbachia","Wolbachia sp. Hymenoptera_ASGLE646-10","8","6","6","1","25","0","97.69","449.75","0","450","40.83","0"
"AGAW2","05N","BAC","TAX:375807","Proteobacteria","Alphaproteobacteria","Rickettsiales","Rickettsiaceae",NA,"Wolbachia","Wolbachia sp. Cerapachys lividus_ASANV557-09","2","2","2","2","0","0","99.75","401","0","446","41.37","0"
"AGAW2","05N","BAC","TAX:392867","Proteobacteria","Alphaproteobacteria","Rickettsiales","Rickettsiaceae",NA,"Wolbachia","Wolbachia endosymbiont of Lasioglossum sp.","1","1","1","3","0","0","97.55","449","0","450","41.11","0"
"AGAW2","05N","BAC","TAX:392873","Proteobacteria","Alphaproteobacteria","Rickettsiales","Rickettsiaceae",NA,"Wolbachia","Wolbachia endosymbiont of Scrapter algoensis","1","1","1","3","0","0","97.58","413","0","450","41.33","0"
"AGAW2","05N","BAC","TAX:375772","Proteobacteria","Alphaproteobacteria","Rickettsiales","Rickettsiaceae",NA,"Wolbachia","Wolbachia sp. Leptogenys JCR25_ASANE483-10","1","1","1","3","0","0","97.54","447","0","450","40.89","0"
"AGAW2","05N","BAC","TAX:91393","Proteobacteria","Alphaproteobacteria","Rickettsiales","Rickettsiaceae",NA,"Wolbachia","","1","1","1","3","0","0","99.49","389","0","405","40","0"
"AGAW2","05N","BAC","TAX:392917","Proteobacteria","Alphaproteobacteria","Rickettsiales","Rickettsiaceae",NA,"Wolbachia","Wolbachia endosymbiont of Glyptapanteles Whitfield131","1","1","1","3","0","0","97.56","450","0","450","40.67","0"
"AGAW2","05N","CHO","no match","no match","no match","no match","no match","no match","no match","no match","no match","no match","no match","no match","no match","no match","no match","no match","no match","no match","no match","no match"
"AGAW2","05N","INS","BOLD:AAG2511","Arthropoda","Insecta","Diptera","Anthomyiidae","Anthomyiinae","Delia","Delia platura","24468","19139","19139","1","21.78","0","99.08","444.5","0","447.09","33.15","0"
"AGAW2","05N","INS","BOLD:AAF9187","Arthropoda","Insecta","Coleoptera","Curculionidae","Entiminae","Phyllobius","Phyllobius oblongus","9869","7430","7430","2","24.71","0","99.41","437.22","0","438.9","38.87","0"
"AGAW2","05N","INS","BOLD:AAA4121","Arthropoda","Insecta","Diptera","Simuliidae","Simuliinae","Simulium","Simulium vittatum complex, Simulium vittatum, Simulium sp. BOLDAAA4121, Simulium argus, Simulium vittatum s.str.","2934","2765","2765","3","5.76","0","98.66","412","0","415.04","33.44","0"
"AGAW2","05N","INS","BOLD:AAH4135","Arthropoda","Insecta","Diptera","Chloropidae","Chloropinae","Thaumatomyia","Thaumatomyia glabra, Thaumatomyia grata, Thaumatomyia sp. 1 TAW","2120","1929","1929","4","9.01","0","99.39","429.39","0","433.15","31.86","0"
"AGAW2","05N","INS","BOLD:AAB4657","Arthropoda","Insecta","Diptera","Chironomidae","Chironominae","Chironomus","Chironomus maturus","1787","1521","1521","5","14.89","0","99.19","433.37","0","436.32","37.78","0"
"AGAW2","05N","INS","BOLD:AAB1098","Arthropoda","Insecta","Diptera","Culicidae","Culicinae","Ochlerotatus, Aedes","Aedes excrucians, Aedes cantans, Aedes stimulans, Aedes euedes, Aedes riparius, Aedes annulipes, Ochlerotatus cantans, Aedes fitchii, Aedes sp., Ochlerotatus annulipes, Ochlerotatus excrucians","1599","1416","1416","6","11.44","0","98.63","434","0","435.11","35.51","0"
"AGAW2","05N","INS","BOLD:AAO3939","Arthropoda","Insecta","Diptera","Limoniidae","Limoniinae","Dicranomyia","","1374","1182","1182","7","13.97","0","99","440.62","0","442.05","36.18","0"
"AGAW2","05N","INS","BOLD:AAA7661","Arthropoda","Insecta","Diptera","Culicidae","Culicinae","Culiseta, Culex","Culiseta melanura, Culex articularis, Culex restuans, Culex acharistus","1282","1198","1198","8","6.55","0","99.19","433.85","0","435.61","32.94","0"
"AGAW2","05N","INS","BOLD:ACX5521","Arthropoda","Insecta","Diptera","Chironomidae","Orthocladiinae","","","1128","1084","1084","9","3.9","0","98.84","407.37","0","409.02","35.04","0"
"AGAW2","05N","INS","BOLD:AAA3453","Arthropoda","Insecta","Diptera","Anthomyiidae","Anthomyiinae","Delia","","1039","965","965","10","7.12","0","99.11","431.7","0","433.77","34.58","0"
"AGAW2","05N","INS","BOLD:AAH3664","Arthropoda","Insecta","Diptera","Cecidomyiidae","","","CecidInt3 sp. BOLD:AAH3664","797","755","755","11","5.27","0","99.1","428.34","0","430.94","26.98","0"
"AGAW2","05N","INS","BOLD:AAF9771","Arthropoda","Insecta","Diptera","Hybotidae","Tachydromiinae","Platypalpus","Platypalpus longiseta, Platypalpus trivialis, Platypalpus sp. 5, Platypalpus sp. 17, Platypalpus pallidiventris","602","476","476","12","20.93","0","98.99","444.86","0","445.38","35.01","0"
"AGAW2","05N","INS","BOLD:AAG2868","Arthropoda","Insecta","Hemiptera","Cicadellidae","Typhlocybinae","Empoasca","Empoasca sp., Empoasca fabae","568","382","382","13","32.75","0","99.69","442.39","0","446.83","32.45","0"
"AGAW2","05N","INS","BOLD:ABA0755","Arthropoda","Insecta","Diptera","Drosophilidae","","","","532","422","422","14","20.68","0","98.34","439.99","0","441.04","31.43","0"
"AGAW2","05N","INS","BOLD:ACU7906","Arthropoda","Insecta","Diptera","Chironomidae","Orthocladiinae","","","518","514","514","15","0.77","0","98.84","407.32","0","410.14","36.1","0"
"AGAW2","05N","INS","BOLD:AAH2372","Arthropoda","Insecta","Coleoptera","Elateridae","Elaterinae","Dalopius","Dalopius pallidus","463","349","349","16","24.62","0","99.36","438.27","0","439.15","36.46","0"
"AGAW2","05N","INS","BOLD:AAB2644","Arthropoda","Insecta","Diptera","Chironomidae","Orthocladiinae","Orthocladius","Orthocladius lapponicus, Orthocladius dorenus, Orthocladius mallochi","441","382","382","17","13.38","0","99.32","442.78","0","444.68","32.22","0"
"AGAW2","05N","INS","BOLD:AAG5507","Arthropoda","Insecta","Diptera","Chironomidae","Chironominae","Chironomus","Chironomus atrella","382","380","380","18","0.52","0","98.98","423.16","0","426.29","35.87","0"
"AGAW2","05N","INS","BOLD:ACF2924","","","","","","","","372","287","287","19","22.85","0","99.62","445.82","0","446.63","25.65","0"
"AGAW2","05N","INS","BOLD:AAD1945","Arthropoda","Insecta","Diptera","Agromyzidae","Phytomyzinae","Cerodontha","Cerodontha dorsalis","284","251","251","20","11.62","0","99.3","441.51","0","443.02","31.91","0"
"AGAW2","05N","INS","BOLD:AAC4197","Arthropoda","Insecta","Diptera","Chironomidae","Orthocladiinae","Paraphaenocladius","Paraphaenocladius impensus","220","166","166","21","24.55","0","99.4","439.56","0","442.02","35.46","0"
"AGAW2","05N","INS","BOLD:AAQ2557","Arthropoda","Insecta","Diptera","Sciaridae","","Peyerimhoffia, Cratyna","Cratyna vagabunda, Peyerimhoffia vagabunda","172","169","169","22","1.74","0","99.1","416.69","0","421.32","35.35","0"
"AGAW2","05N","INS","BOLD:AAM9347","Arthropoda","Insecta","Diptera","Phoridae","Metopininae","Megaselia","","160","157","157","23","1.88","0","98.05","392.44","0","394.26","33.17","0"
"AGAW2","05N","INS","BOLD:AAF9014","Arthropoda","Insecta","Diptera","Limoniidae","Chioneinae","Symplecta","Symplecta cana, Symplecta sp. BOLDAAF9014","129","128","128","24","0.78","0","98.69","420.46","0","426.16","33.71","0"
"AGAW2","05N","INS","BOLD:AAN5355","Arthropoda","Insecta","Diptera","Chironomidae","Orthocladiinae","Smittia","Smittia cf. stercoraria, Smittia stercoraria","122","118","118","25","3.28","0","98.61","439.11","0","442.56","36.02","0"
"AGAW2","05N","INS","BOLD:AAB1787","Arthropoda","Insecta","Hemiptera","Aphididae","Aphidinae","Rhopalosiphum, Acyrthosiphon","Acyrthosiphon pisum, Acyrthosiphon pisum 5A, Rhopalosiphum maidis","118","96","96","26","18.64","0","99.56","436.01","0","438.08","25.63","0"
"AGAW2","05N","INS","BOLD:AAD1485","Arthropoda","Insecta","Diptera","Chironomidae","Chironominae","Paratanytarsus","Paratanytarsus grimmii, Paratanytarsus sp.","90","87","87","27","3.33","0","99.24","437.72","0","439.27","33.79","0"
"AGAW2","05N","INS","BOLD:AAN5342","Arthropoda","Insecta","Diptera","Chironomidae","Orthocladiinae","Heleniella","","77","75","75","28","2.6","0","98.57","422.81","0","432.65","35.12","0"
"AGAW2","05N","INS","BOLD:AAA9899","Arthropoda","Insecta","Hemiptera","Aphididae","Aphidinae","Rhopalosiphum","Rhopalosiphum padi","76","76","76","29","0","0","99.32","437.22","0","438.79","25.77","0"
"AGAW2","05N","INS","BOLD:AAN5356","Arthropoda","Insecta","Diptera","Chironomidae","Orthocladiinae","Smittia","","57","50","50","30","12.28","0","99.25","445.04","0","446.4","35.17","0"
"AGAW2","05N","INS","BOLD:AAN1581","Arthropoda","Insecta","Coleoptera","Staphylinidae","Aleocharinae","Oxypoda","Oxypoda opaca","48","46","46","31","4.17","0","99.77","389.81","0","440.88","33.64","0"
"AGAW2","05N","INS","BOLD:AAM9252","Arthropoda","Insecta","Diptera","Sciaridae","","Bradysia, Leptosciarella","Bradysia vagans, Leptosciarella zanti","45","44","44","32","2.22","0","99.29","442.93","0","443.98","38.99","0"
"AGAW2","05N","INS","BOLD:AAD7604","Arthropoda","Insecta","Coleoptera","Coccinellidae","Coccinellinae","Coleomegilla","Coleomegilla maculata","43","42","42","33","2.33","0","99","432.77","0","435.21","31.77","0"
"AGAW2","05N","INS","BOLD:AAA5803","Arthropoda","Insecta","Hemiptera","Miridae","Mirinae","Lygus","Lygus lineolaris","35","30","30","34","14.29","0","99.26","446.31","0","447.89","34.1","0"
"AGAW2","05N","INS","BOLD:AAN5267","Arthropoda","Insecta","Diptera","Cecidomyiidae","","","","32","27","27","35","15.63","0","99.19","447.19","0","447.75","30.65","0"
"AGAW2","05N","INS","BOLD:AAN5205","Arthropoda","Insecta","Diptera","Cecidomyiidae","","","CecidInt42 sp. BOLD:AAN5205","31","29","29","36","6.45","0","98.71","444.9","0","446.39","31.89","0"
"AGAW2","05N","INS","BOLD:ABA0863","Arthropoda","Insecta","Diptera","Cecidomyiidae","","","","29","26","26","37","10.34","0","99.22","438.62","0","439.72","29.83","0"
"AGAW2","05N","INS","BOLD:ACO1622","Arthropoda","Insecta","Diptera","Cecidomyiidae","","","","26","26","26","38","0","0","98.5","418.85","0","420.35","31.52","0"
"AGAW2","05N","INS","BOLD:AAM7726","Arthropoda","Insecta","Coleoptera","Curculionidae","Curculioninae","Orchestes","Orchestes steppensis, Orchestes alni","24","21","21","39","12.5","0","98.94","435","0","442.83","33.5","0"
"AGAW2","05N","INS","BOLD:AAA4783","Arthropoda","Insecta","Hymenoptera","Braconidae","Microgastrinae","Glyptapanteles","","23","22","22","40","4.35","0","99.26","422.91","0","439.78","25.02","0"
"AGAW2","05N","INS","BOLD:AAB7490","Arthropoda","Insecta","Diptera","Chironomidae","Orthocladiinae","Tvetenia","Tvetenia paucunca","20","20","20","41","0","0","97.82","436.35","0","439.5","34","0"
"AGAW2","05N","INS","BOLD:AAU8585","Arthropoda","Insecta","Hymenoptera","Braconidae","Aphidiinae","Trioxys, Binodoxys","","17","17","17","42","0","0","99.33","435.94","0","437.12","27.56","0"
"AGAW2","05N","INS","BOLD:AAN6447","Arthropoda","Insecta","Diptera","Sciaridae","","Corynoptera","Corynoptera perpusilla","15","14","14","43","6.67","0","99.36","448.33","0","450","36.44","0"
"AGAW2","05N","INS","BOLD:AAV6213","Arthropoda","Insecta","Diptera","Chironomidae","Chironominae","Paratanytarsus","","11","11","11","44","0","0","99.18","416.64","0","418.64","34.44","0"
"AGAW2","05N","INS","BOLD:AAF9311","Arthropoda","Insecta","Diptera","Psychodidae","Psychodinae","Psychoda","Psychoda cf. satchelli, Psychoda satchelli","11","11","11","44","0","0","99.13","441.36","0","443","34.83","0"
"AGAW2","05N","INS","BOLD:AAA4222","Arthropoda","Insecta","Lepidoptera","Erebidae","Hypeninae","Hypena","Hypena scabra","10","10","10","46","0","0","98.89","405.1","0","406.9","32.42","0"
"AGAW2","05N","INS","BOLD:AAM6037","Arthropoda","Insecta","Diptera","Cecidomyiidae","Cecidomyiinae","Contarinia, Dasineura","","10","10","10","46","0","0","98.32","446.2","0","446","26.69","0"
"AGAW2","05N","INS","BOLD:ACH0608","Arthropoda","Insecta","Hymenoptera","Ichneumonidae","","","","9","9","9","48","0","0","99.4","433.33","0","435.89","29.01","0"
"AGAW2","05N","INS","BOLD:AAK5928","Arthropoda","Insecta","Coleoptera","Chrysomelidae","Criocerinae","Oulema","Oulema melanopus","9","9","9","48","0","0","99.72","436.89","0","438.44","36.54","0"
"AGAW2","05N","INS","BOLD:AAG2462","Arthropoda","Insecta","Diptera","Anthomyiidae","Anthomyiinae","Botanophila","Botanophila fugax","8","8","8","50","0","0","98.33","441.75","0","442.88","31.53","0"
"AGAW2","05N","INS","BOLD:AAG6743","Arthropoda","Insecta","Diptera","Sarcophagidae","Sarcophaginae","Sarcophaga","Sarcophaga sp. 1 MG, Sarcophaga subvicina","7","7","7","51","0","0","98.87","433.71","0","436","34.51","0"
"AGAW2","05N","INS","BOLD:AAG1421","Arthropoda","Insecta","Hymenoptera","Braconidae","Aphidiinae","Diaeretiella","Diaeretiella rapae","7","7","7","51","0","0","99.08","358.57","0","431.14","27.04","0"
"AGAW2","05N","INS","BOLD:ABY9541","Arthropoda","Insecta","Hymenoptera","Braconidae","Microgastrinae","Glyptapanteles","Glyptapanteles fulvipes, Glyptapanteles jft04, Glyptapanteles jft09","6","5","5","53","16.67","0","98.14","449.17","0","450","25.48","0"
"AGAW2","05N","INS","BOLD:AAA7374","Arthropoda","Insecta","Diptera","Syrphidae","Syrphinae","Sphaerophoria","Sphaerophoria bankowskae, Sphaerophoria infuscata, Sphaerophoria nr. philanthus, Sphaerophoria batava, Sphaerophoria sp., Sphaerophoria rueppellii, Sphaerophoria interrupta-group, Sphaerophoria sp. MA-1, Sphaerophoria cleoae, Sphaerophoria asymmetrica, Sphaerophoria fatarum, Sphaerophoria bengalensis, Sphaerophoria bifurcata, Sphaerophoria taeniata, Sphaerophoria interrupta, Sphaerophoria contigua, Sphaerophoria kaa, Sphaerophoria laurae, Sphaerophoria chongjini, Sphaerophoria boreoalpina, Sphaerophoria sp. BOLDAAA7374, Sphaerophoria macrogaster, Sphaerophoria sulphuripes, Sphaerophoria brevipilosa, Sphaerophoria philanthus, Sphaerophoria pyrrhina, Sphaerophoria abbreviata, Sphaerophoria scripta, Sphaerophoria potentillae, Sphaerophoria longipilosa, Sphaerophoria virgata","5","5","5","54","0","0","99.46","448","0","450","31.16","0"
"AGAW2","05N","INS","BOLD:ACX6357","Arthropoda","Insecta","Diptera","Chironomidae","","","","4","4","4","55","0","0","99.19","433","0","434.5","33.72","0"
"AGAW2","05N","INS","BOLD:AAP5146","Arthropoda","Insecta","Diptera","Chironomidae","Orthocladiinae","Orthocladius","","4","4","4","55","0","0","98.89","399.75","0","401.25","33.16","0"
"AGAW2","05N","INS","BOLD:AAB9140","Arthropoda","Insecta","Diptera","Calliphoridae","Chrysomyinae","Phormia","Phormia regina","3","3","3","57","0","0","98.96","424.67","0","426.67","32.56","0"
"AGAW2","05N","INS","BOLD:AAN5358","Arthropoda","Insecta","Diptera","Chironomidae","Orthocladiinae","Smittia","Smittia stercoraria, Smittia sp. 22ES","3","3","3","57","0","0","99.18","447.33","0","450","37.19","0"
"AGAW2","05N","INS","BOLD:AAA2815","Arthropoda","Insecta","Lepidoptera","Noctuidae","Noctuinae","Mythimna","Mythimna yukonensis, Mythimna oxygala","3","3","3","57","0","0","99.17","445","0","446.33","33.61","0"
"AGAW2","05N","INS","BOLD:AAA9101","Arthropoda","Insecta","Lepidoptera","Pyralidae","Phycitinae","Moodna, Caudellia","Moodna ostrinella, Moodna bisinuella, Caudellia nigrella","3","3","3","57","0","0","99.93","449.33","0","450","34.52","0"
"AGAW2","05N","INS","BOLD:AAC9062","Arthropoda","Insecta","Diptera","Culicidae","Culicinae","Ochlerotatus, Aedes","Aedes fitchii, Ochlerotatus fitchii, Aedes grossbecki, Aedes ventrovittis","3","3","3","57","0","0","99.55","449","0","450","33.26","0"
"AGAW2","05N","INS","BOLD:AAG4653","Arthropoda","Insecta","Diptera","Syrphidae","Syrphinae","Heringia","Heringia latitarsis, Heringia pubescens, Heringia jakutorum, Heringia rita, Heringia fulvimanus, Heringia sp. BOLDAAG4653, Heringia nr. pubescens, Heringia coxalis, Heringia larusi, Heringia vitripennis","3","3","3","57","0","0","99.33","421","0","423.67","31.39","0"
"AGAW2","05N","INS","BOLD:AAA3933","Arthropoda","Insecta","Lepidoptera","Erebidae","Herminiinae","Palthis","Palthis angulalis","2","2","2","63","0","0","98.95","429.5","0","434","32.25","0"
"AGAW2","05N","INS","BOLD:AAG6433","Arthropoda","Insecta","Diptera","Ceratopogonidae","Forcipomyiinae","Forcipomyia","Forcipomyia sp. BOLD:AAG6433, Forcipomyia sp. 17","2","2","2","63","0","0","97.87","447","0","450","32.67","0"
"AGAW2","05N","INS","BOLD:AAL7520","Arthropoda","Insecta","Diptera","Anthomyiidae","Anthomyiinae","Lasiomma","","2","2","2","63","0","0","99.55","447.5","0","450","32.89","0"
"AGAW2","05N","INS","BOLD:ABU5521","Arthropoda","Insecta","Diptera","Sciaridae","","Corynoptera","Corynoptera furcata","2","2","2","63","0","0","98.01","432.5","0","434.5","35.89","0"
"AGAW2","05N","INS","BOLD:AAU6630","Arthropoda","Insecta","Diptera","Fanniidae","Fanniinae","Fannia","Fannia armata","2","2","2","63","0","0","98.66","448","0","450","33.33","0"
"AGAW2","05N","INS","BOLD:AAA9251","Arthropoda","Insecta","Lepidoptera","Gelechiidae","Gelechiinae","Scrobipalpa","Scrobipalpa atriplicella, Scrobipalpa nr. atriplicella sp. 1","2","2","2","63","0","0","99.88","426.5","0","431.5","34.18","0"
"AGAW2","05N","INS","BOLD:ACX6065","Arthropoda","Insecta","Diptera","Cecidomyiidae","","","","2","2","2","63","0","0","99.33","450","0","450","29.33","0"
"AGAW2","05N","INS","BOLD:AAC2498","Arthropoda","Insecta","Diptera","Muscidae","Phaoniinae","Helina","Helina evecta, Helina aldrichi","2","2","2","63","0","0","99.16","358","0","359.5","30.46","0"
"AGAW2","05N","INS","BOLD:ACF2821","Arthropoda","Insecta","Diptera","Mycetophilidae","Mycetophilinae","Mycetophila","Mycetophila fungorum","2","2","2","63","0","0","98.49","421.5","0","422.5","35.23","0"
"AGAW2","05N","INS","BOLD:AAD0853","Arthropoda","Insecta","Diptera","Scathophagidae","Scathophaginae","Scathophaga","Scathophaga lutaria, Scathophaga hiemalis, Scathophaga furcata, Scathophaga stercoraria, Scathophaga inquinata, Scathophaga suilla","2","2","2","63","0","0","98.39","432","0","434","31.58","0"
"AGAW2","05N","INS","BOLD:ADH6290","","","","","","","","2","2","2","63","0","0","98.58","405","0","405","38.94","0"
"AGAW2","05N","INS","BOLD:ABA2925","Arthropoda","Insecta","Hymenoptera","Braconidae","Microgastrinae","Glyptapanteles","Glyptapanteles fulvipes","2","2","2","63","0","0","97.33","449","0","450","25.78","0"
"AGAW2","05N","INS","BOLD:ACI3511","Arthropoda","Insecta","Lepidoptera","Gelechiidae","Gelechiinae","Coleotechnites","Coleotechnites occidentis","1","1","1","75","0","0","97.1","449","0","450","33.56","0"
"AGAW2","05N","INS","BOLD:ACE5443","Arthropoda","Insecta","Coleoptera","Carabidae","Harpalinae","Anisodactylus","Anisodactylus kirbyi","1","1","1","75","0","0","99.09","439","0","448","33.04","0"
"AGAW2","05N","INS","BOLD:AAQ2892","Arthropoda","Insecta","Hymenoptera","Braconidae","Rhyssalinae","Proacrisis","","1","1","1","75","0","0","100","450","0","450","28","0"
"AGAW2","05N","INS","BOLD:AAA8933","Arthropoda","Insecta","Coleoptera","Coccinellidae","Coccinellinae","Coccinella","Coccinella septempunctata","1","1","1","75","0","0","99.33","448","0","450","32.89","0"
"AGAW2","05N","INS","BOLD:AAH1187","Arthropoda","Insecta","Hymenoptera","Braconidae","Microgastrinae","Glyptapanteles","Glyptapanteles jft10","1","1","1","75","0","0","97.53","445","0","450","24.89","0"
"AGAW2","05N","INS","BOLD:AAC9132","Arthropoda","Insecta","Diptera","Culicidae","Culicinae","Culiseta","Culiseta inornata","1","1","1","75","0","0","99.75","396","0","396","32.32","0"
"AGAW2","05N","INS","BOLD:AAF9041","Arthropoda","Insecta","Diptera","Tipulidae","Tipulinae","Tipula","Tipula oleracea","1","1","1","75","0","0","99.75","395","0","397","35.26","0"
"AGAW2","05N","INS","BOLD:ABZ7601","Arthropoda","Insecta","Lepidoptera","Noctuidae","Noctuinae","Anarta","Anarta trifolii","1","1","1","75","0","0","98.44","448","0","450","31.11","0"
"AGAW2","05N","INS","BOLD:AAG5443","Arthropoda","Insecta","Diptera","Chironomidae","Orthocladiinae","","","1","1","1","75","0","0","99.75","400","0","402","32.34","0"
"AGAW2","05N","INS","BOLD:AAG7312","Arthropoda","Insecta","Diptera","Sphaeroceridae","Limosininae","Spelobia","Spelobia clunipes, Spelobia sp. B TAW","1","1","1","75","0","0","99.33","450","0","450","33.56","0"
"AGAW2","05N","INS","BOLD:AAK7242","Arthropoda","Insecta","Coleoptera","Melandryidae","Melandryinae","Melandrya","Melandrya striata","1","1","1","75","0","0","97.99","398","0","450","33.33","0"
"AGAW2","05N","INS","BOLD:AAA5299","Arthropoda","Insecta","Diptera","Chironomidae","Orthocladiinae","Cricotopus","Cricotopus laetus, Cricotopus relucens, Cricotopus trifasciatus, Cricotopus sp., Cricotopus sp. 3ES, Cricotopus sylvestris, Cricotopus glacialis","1","1","1","75","0","0","98.24","398","0","399","33.33","0"
"AGAW2","05N","INS","BOLD:AAG0991","Arthropoda","Insecta","Diptera","Chironomidae","Orthocladiinae","Orthocladius","","1","1","1","75","0","0","97.99","447","0","450","32.67","0"
"AGAW2","05N","INS","BOLD:AAB5609","Arthropoda","Insecta","Diptera","Muscidae","Coenosiinae","Coenosia","Coenosia sp.BOLD:AAB5609, Coenosia tigrina","1","1","1","75","0","0","100","375","0","377","33.16","0"
"AGAW2","05N","INS","BOLD:AAV5788","Arthropoda","Insecta","Diptera","Cecidomyiidae","Cecidomyiinae","Asphondylia","","1","1","1","75","0","0","98.34","421","0","422","23.7","0"
"AGAW2","05N","INS","BOLD:AAC2438","Arthropoda","Insecta","Diptera","Syrphidae","Syrphinae","Paragus","Paragus haemorrhous, Paragus angustifrons","1","1","1","75","0","0","98.22","450","0","450","32.44","0"
"AGAW2","05N","INS","BOLD:ACJ5130","Arthropoda","Insecta","Diptera","Chironomidae","Chironominae","","","1","1","1","75","0","0","99.02","410","0","413","31.48","0"
"AGAW2","05N","INS","BOLD:AAG6768","Arthropoda","Insecta","Diptera","Scathophagidae","Scathophaginae","Cordilura","Cordilura pleuritica","1","1","1","75","0","0","99.11","450","0","450","34.22","0"
"AGAW2","05N","INS","BOLD:AAZ4669","Arthropoda","Insecta","Diptera","Tachinidae","Exoristinae","Oswaldia","Oswaldia assimilis","1","1","1","75","0","0","98.21","447","0","450","33.11","0"
"AGAW2","05N","INS","BOLD:AAJ7105","Arthropoda","Insecta","Diptera","Agromyzidae","Agromyzinae","Agromyza","Agromyza frontella","1","1","1","75","0","0","100","400","0","450","36.67","0"
"AGAW2","05N","INS","BOLD:AAB7030","Arthropoda","Insecta","Diptera","Chironomidae","Chironominae","Chironomus","Chironomus sp. BOLDAAP2996, Chironomus quinnitukqut, Chironomus cf. decorus 2g, Chironomus decorus group sp. 2 IP2013, Chironomus cf. decorus, Chironomus decorus gr.","1","1","1","75","0","0","97.8","363","0","365","33.97","0"
"AGAW2","05N","INS","BOLD:AAD5253","Arthropoda","Insecta","Diptera","Chironomidae","Orthocladiinae","Thienemanniella","Thienemanniella xena, Thienemanniella vittata","1","1","1","75","0","0","98","450","0","450","33.56","0"
"AGAW2","05N","INS","BOLD:ADF6661","Arthropoda","Insecta","Diptera","Cecidomyiidae","","","","1","1","1","75","0","0","99.04","418","0","450","30","0"
"AGAW2","05N","INS","BOLD:AAH0145","Arthropoda","Insecta","Coleoptera","Carabidae","Harpalinae","Anisodactylus","Anisodactylus pitychrous, Anisodactylus sanctaecrucis, Anisodactylus alternans, Anisodactylus discoideus","1","1","1","75","0","0","98.81","419","0","419","31.74","0"
"AGAW2","05N","INS","BOLD:AAF4976","Arthropoda","Insecta","Coleoptera","Lampyridae","Lampyrinae","Ellychnia","Ellychnia aff. corrusca KSH611, Ellychnia sp., Ellychnia facula, Ellychnia sp. KSH1376, Ellychnia lacustris, Ellychnia corrusca, Ellychnia hatchi, Ellychnia californica","1","1","1","75","0","0","99.33","450","0","450","36.67","0"
"AGAW2","05N","INS","BOLD:AAU6966","Arthropoda","Insecta","Coleoptera","Throscidae","Throscinae","Trixagus","Trixagus chevrolati","1","1","1","75","0","0","99.76","413","0","450","39.78","0"
"AGAW2","05N","INS","BOLD:AAG6745","Arthropoda","Insecta","Diptera","Calliphoridae","Calliphorinae","Pollenia","Pollenia pediculata","1","1","1","75","0","0","97.58","414","0","417","33.81","0"
"AGAW2","05N","INS","BOLD:ABW2407","Arthropoda","Insecta","Diptera","Anthomyiidae","Anthomyiinae","Hylemya","","1","1","1","75","0","0","98.73","393","0","396","32.07","0"
"AGAW2","05N","INS","BOLD:ACF5939","Arthropoda","Insecta","Diptera","Tachinidae","Exoristinae","Winthemia","Winthemia sinuata, Winthemia illinoensis","1","1","1","75","0","0","97.64","424","0","430","29.3","0"
"AGAW2","05N","INS","BOLD:ACK7296","Arthropoda","Insecta","Diptera","Cecidomyiidae","","","","1","1","1","75","0","0","100","450","0","450","24.44","0"
"AGAW2","05N","INS","BOLD:AAA3654","Arthropoda","Insecta","Lepidoptera","Crambidae","Crambinae","Crambus","Crambus bigelovi, Crambus johnsoni, Crambus praefectellus","1","1","1","75","0","0","99.55","449","0","450","31.33","0"
"AGAW2","05N","INS","BOLD:AAH1890","Arthropoda","Insecta","Hymenoptera","Ichneumonidae","Cryptinae","Stibeutes","Stibeutes brevicornis, Stibeutes heinemanni","1","1","1","75","0","0","99.53","427","0","450","29.33","0"
"AGAW2","05N","INS","BOLD:AAP5142","Arthropoda","Insecta","Diptera","Chironomidae","Chironominae","","","1","1","1","75","0","0","98.44","450","0","450","32.44","0"
"AGAW2","05N","INS","BOLD:AAG6647","Arthropoda","Insecta","Diptera","Bibionidae","Bibioninae","Bibio","","1","1","1","75","0","0","98.22","449","0","450","32","0"
"AGAW2","05N","INS","BOLD:AAE4298","Arthropoda","Insecta","Diptera","Chironomidae","Orthocladiinae","Cricotopus","Cricotopus tremulus","1","1","1","75","0","0","97.76","447","0","450","32","0"
"AGAW2","05N","INS","BOLD:AAG1701","Arthropoda","Insecta","Diptera","Muscidae","Coenosiinae","Macrorchis","Macrorchis ausoba","1","1","1","75","0","0","99.33","450","0","450","33.11","0"
"AGAW2","05N","INS","BOLD:AAE7016","Arthropoda","Insecta","Lepidoptera","Gelechiidae","Dichomeridinae","Helcystogramma","Helcystogramma hystricella","1","1","1","75","0","0","97.53","446","0","450","32","0"
"AGAW2","05N","INS","BOLD:ADH9310","Arthropoda","Insecta","Diptera","Anthomyiidae","Anthomyiinae","Anthomyia","Anthomyia procellaris, Anthomyia pluvialis","1","1","1","75","0","0","98.44","450","0","450","32.89","0"
"AGAW2","05N","INS","BOLD:AAG2465","Arthropoda","Insecta","Diptera","Anthomyiidae","Anthomyiinae","Delia","Delia radicum","1","1","1","75","0","0","98.67","450","0","450","34.22","0"
"AGAW2","05N","NAR","no match","no match","no match","no match","no match","no match","no match","no match","no match","no match","no match","no match","no match","no match","no match","no match","no match","no match","no match","no match"
"AGAW2","05N","NIN","BOLD:ABX0027","Arthropoda","Collembola","Symphypleona","Bourletiellidae","","Bourletiella","","113","90","90","1","20.35","0","99.56","415.55","0","443.03","35.59","0"
"AGAW2","05N","NIN","BOLD:AAG1792","Arthropoda","Arachnida","Araneae","Tetragnathidae","","Glenognatha","Glenognatha foxi","99","84","84","2","15.15","0","99.54","445.78","0","446.89","31.52","0"
"AGAW2","05N","NIN","BOLD:AAV0589","Arthropoda","Arachnida","Mesostigmata","Blattisociidae","Platyseiinae","Cheiroseius","","72","67","67","3","6.94","0","99.24","447.85","0","449","27.67","0"
"AGAW2","05N","NIN","BOLD:AAB8452","Arthropoda","Collembola","Entomobryomorpha","Entomobryidae","Lepidocyrtinae","Lepidocyrtus","Lepidocyrtus paradoxus","54","47","47","4","12.96","0","99.7","445.7","0","444.72","42.01","0"
"AGAW2","05N","NIN","BOLD:AAB7915","Arthropoda","Collembola","Symphypleona","Bourletiellidae","","Deuterosminthurus","Deuterosminthurus sp1","39","39","39","5","0","0","98.88","438.49","0","440.13","35.01","0"
"AGAW2","05N","NIN","BOLD:AAA5267","Arthropoda","Collembola","Entomobryomorpha","Isotomidae","","","","37","37","37","6","0","0","98.75","419.08","0","420.89","38.25","0"
"AGAW2","05N","NIN","BOLD:AAI4706","Arthropoda","Collembola","Symphypleona","Katiannidae","","","","13","13","13","7","0","0","98.95","417.92","0","419.54","41.49","0"
"AGAW2","05N","NIN","BOLD:AAH6630","Arthropoda","Collembola","Entomobryomorpha","Entomobryidae","Entomobryinae","Entomobrya","Entomobrya unostrigata","7","7","7","8","0","0","98.67","426","0","426.71","39.15","0"
"AGAW2","05N","NIN","BOLD:ACF8021","Arthropoda","Arachnida","Mesostigmata","Ascidae","","Arctoseius","Arctoseius cetratus","4","4","4","9","0","0","97.83","370.75","0","372.75","38.57","0"
"AGAW2","05N","NIN","BOLD:AAN6642","Arthropoda","Arachnida","Mesostigmata","Blattisociidae","Platyseiinae","Cheiroseius","","3","3","3","10","0","0","98.82","449.33","0","450","26.81","0"
"AGAW2","05N","NIN","BOLD:AAN4906","Arthropoda","Collembola","Symphypleona","Bourletiellidae","","","","2","2","2","11","0","0","99.27","426.5","0","427.5","35.74","0"
"AGAW2","05N","NIN","BOLD:ABW2639","Arthropoda","Arachnida","Trombidiformes","Anystidae","","","","1","1","1","12","0","0","99.55","449","0","450","31.56","0"
"AGAW2","05S","BAC","no match","no match","no match","no match","no match","no match","no match","no match","no match","no match","no match","no match","no match","no match","no match","no match","no match","no match","no match","no match"
"AGAW2","05S","CHO","no match","no match","no match","no match","no match","no match","no match","no match","no match","no match","no match","no match","no match","no match","no match","no match","no match","no match","no match","no match"
"AGAW2","05S","INS","BOLD:AAA3453","Arthropoda","Insecta","Diptera","Anthomyiidae","Anthomyiinae","Delia","","24529","20439","20439","1","16.67","0","99.29","431.08","0","433.28","34.67","0"
"AGAW2","05S","INS","BOLD:AAG2511","Arthropoda","Insecta","Diptera","Anthomyiidae","Anthomyiinae","Delia","Delia platura","12231","10277","10277","2","15.98","0","98.89","444.71","0","447.11","33.29","0"
"AGAW2","05S","INS","BOLD:AAO3939","Arthropoda","Insecta","Diptera","Limoniidae","Limoniinae","Dicranomyia","","11899","9900","9900","3","16.8","0","99.17","439.16","0","441.15","36.49","0"
"AGAW2","05S","INS","BOLD:AAC9132","Arthropoda","Insecta","Diptera","Culicidae","Culicinae","Culiseta","Culiseta inornata","4340","4290","4290","4","1.15","0","98.8","406.16","0","408.72","32.39","0"
"AGAW2","05S","INS","BOLD:AAA4555","Arthropoda","Insecta","Orthoptera","Acrididae","Melanoplinae","Melanoplus","Melanoplus stupefactus, Melanoplus angustipennis, Melanoplus ponderosus, Melanoplus bruneri, Melanoplus frigidus, Melanoplus scapularis, Melanoplus cinereus, Melanoplus madeleineae, Melanoplus sanguinipes, Melanoplus nigrescens group, Melanoplus tunicae, Melanoplus walshi, Melanoplus femurrubrum, Melanoplus dawsoni, Melanoplus sp., Melanoplus fasciatus, Melanoplus bowditchi, Melanoplus stonei, Melanoplus bivittatus, Melanoplus huroni, Melanoplus thomasi, Melanoplus borealis, Melanoplus packardii","1009","895","895","5","11.3","0","98.86","441.95","0","443.48","36.64","0"
"AGAW2","05S","INS","BOLD:AAN5358","Arthropoda","Insecta","Diptera","Chironomidae","Orthocladiinae","Smittia","Smittia stercoraria, Smittia sp. 22ES","924","798","798","6","13.64","0","99.33","441.19","0","442.73","37.3","0"
"AGAW2","05S","INS","BOLD:ACK3431","Arthropoda","Insecta","Diptera","Cecidomyiidae","","","","917","870","870","7","5.13","0","99.07","421.79","0","423.24","32.73","0"
"AGAW2","05S","INS","BOLD:ACA7493","Arthropoda","Insecta","Diptera","Chironomidae","Chironominae","","","878","862","862","8","1.82","0","97.43","383.39","0","391.74","39.51","0"
"AGAW2","05S","INS","BOLD:AAP9022","Arthropoda","Insecta","Diptera","Cecidomyiidae","","","","854","721","721","9","15.57","0","99.16","438.34","0","439.63","27.54","0"
"AGAW2","05S","INS","BOLD:AAI3268","Arthropoda","Insecta","Diptera","Chaoboridae","Chaoborinae","Mochlonyx","","551","436","436","10","20.87","0","99.03","443.45","0","444.55","34.2","0"
"AGAW2","05S","INS","BOLD:AAF9771","Arthropoda","Insecta","Diptera","Hybotidae","Tachydromiinae","Platypalpus","Platypalpus longiseta, Platypalpus trivialis, Platypalpus sp. 5, Platypalpus sp. 17, Platypalpus pallidiventris","543","402","402","11","25.97","0","99.47","444.98","0","445.61","35.19","0"
"AGAW2","05S","INS","BOLD:ACU7906","Arthropoda","Insecta","Diptera","Chironomidae","Orthocladiinae","","","538","537","537","12","0.19","0","98.89","403.64","0","406.76","36.09","0"
"AGAW2","05S","INS","BOLD:AAF8995","Arthropoda","Insecta","Diptera","Limoniidae","Limoniinae","Antocha","Antocha saxicola","466","404","404","13","13.3","0","98.37","429.24","0","431.04","37.46","0"
"AGAW2","05S","INS","BOLD:AAD1945","Arthropoda","Insecta","Diptera","Agromyzidae","Phytomyzinae","Cerodontha","Cerodontha dorsalis","404","358","358","14","11.39","0","99.34","440.74","0","442.29","31.92","0"
"AGAW2","05S","INS","BOLD:AAF9014","Arthropoda","Insecta","Diptera","Limoniidae","Chioneinae","Symplecta","Symplecta cana, Symplecta sp. BOLDAAF9014","268","262","262","15","2.24","0","98.53","422.29","0","427.83","33.83","0"
"AGAW2","05S","INS","BOLD:ACF8552","Arthropoda","Insecta","Coleoptera","Anthicidae","Anthicinae","Malporus","Malporus formicarius","204","184","184","16","9.8","0","99.03","364.33","0","425.75","34.32","0"
"AGAW2","05S","INS","BOLD:AAA5299","Arthropoda","Insecta","Diptera","Chironomidae","Orthocladiinae","Cricotopus","Cricotopus laetus, Cricotopus relucens, Cricotopus trifasciatus, Cricotopus sp., Cricotopus sp. 3ES, Cricotopus sylvestris, Cricotopus glacialis","182","166","166","17","8.79","0","99.09","431.06","0","432.13","34.62","0"
"AGAW2","05S","INS","BOLD:AAG2868","Arthropoda","Insecta","Hemiptera","Cicadellidae","Typhlocybinae","Empoasca","Empoasca sp., Empoasca fabae","158","104","104","18","34.18","0","99.65","441.55","0","446.62","32.51","0"
"AGAW2","05S","INS","BOLD:AAH3664","Arthropoda","Insecta","Diptera","Cecidomyiidae","","","CecidInt3 sp. BOLD:AAH3664","137","135","135","19","1.46","0","99.08","426.34","0","429.11","26.94","0"
"AGAW2","05S","INS","BOLD:ACF2924","","","","","","","","116","96","96","20","17.24","0","99.65","446.62","0","447.6","25.67","0"
"AGAW2","05S","INS","BOLD:ACI4729","Arthropoda","Insecta","Diptera","Chironomidae","Chironominae","Cladotanytarsus","Cladotanytarsus nigrovittatus","81","81","81","21","0","0","99.31","420.51","0","421.58","41.19","0"
"AGAW2","05S","INS","BOLD:ACK4716","Arthropoda","Insecta","Diptera","Cecidomyiidae","","","","72","69","69","22","4.17","0","98.95","428.03","0","429.31","31.53","0"
"AGAW2","05S","INS","BOLD:AAN5342","Arthropoda","Insecta","Diptera","Chironomidae","Orthocladiinae","Heleniella","","62","62","62","23","0","0","99.09","396.11","0","410.35","34.83","0"
"AGAW2","05S","INS","BOLD:ABA1215","Arthropoda","Insecta","Diptera","Sciaridae","","Lycoriella","Lycoriella sativae","60","49","49","24","18.33","0","99.31","437.17","0","438.77","35.96","0"
"AGAW2","05S","INS","BOLD:AAB1787","Arthropoda","Insecta","Hemiptera","Aphididae","Aphidinae","Rhopalosiphum, Acyrthosiphon","Acyrthosiphon pisum, Acyrthosiphon pisum 5A, Rhopalosiphum maidis","53","47","47","25","11.32","0","99.39","429.98","0","430.66","25.4","0"
"AGAW2","05S","INS","BOLD:AAV1295","Arthropoda","Insecta","Diptera","Sciaridae","","Bradysia","Bradysia difformis, Bradysia impatiens","46","46","46","26","0","0","99.21","415.74","0","437.54","31.61","0"
"AGAW2","05S","INS","BOLD:ABU5521","Arthropoda","Insecta","Diptera","Sciaridae","","Corynoptera","Corynoptera furcata","39","39","39","27","0","0","98.63","407.56","0","409.97","36.76","0"
"AGAW2","05S","INS","BOLD:AAQ0265","Arthropoda","Insecta","Diptera","Hybotidae","Tachydromiinae","Platypalpus","Platypalpus pulicarius","28","27","27","28","3.57","0","98.48","447.18","0","448.21","35.56","0"
"AGAW2","05S","INS","BOLD:AAN5356","Arthropoda","Insecta","Diptera","Chironomidae","Orthocladiinae","Smittia","","22","16","16","29","27.27","0","99.23","449.23","0","450","35.25","0"
"AGAW2","05S","INS","BOLD:AAA4222","Arthropoda","Insecta","Lepidoptera","Erebidae","Hypeninae","Hypena","Hypena scabra","20","20","20","30","0","0","98.92","398.65","0","400.95","32.53","0"
"AGAW2","05S","INS","BOLD:ADF6661","Arthropoda","Insecta","Diptera","Cecidomyiidae","","","","16","16","16","31","0","0","98.86","416.06","0","444.5","30.13","0"
"AGAW2","05S","INS","BOLD:AAG6756","Arthropoda","Insecta","Diptera","Sarcophagidae","Sarcophaginae","Boettcheria","Boettcheria cimbicis","14","14","14","32","0","0","99.45","437.43","0","438.57","34.81","0"
"AGAW2","05S","INS","BOLD:AAF9187","Arthropoda","Insecta","Coleoptera","Curculionidae","Entiminae","Phyllobius","Phyllobius oblongus","11","11","11","33","0","0","99.34","442.55","0","443.64","39.2","0"
"AGAW2","05S","INS","BOLD:AAB2644","Arthropoda","Insecta","Diptera","Chironomidae","Orthocladiinae","Orthocladius","Orthocladius lapponicus, Orthocladius dorenus, Orthocladius mallochi","8","8","8","34","0","0","98.76","420.25","0","422.88","32.44","0"
"AGAW2","05S","INS","BOLD:AAA7374","Arthropoda","Insecta","Diptera","Syrphidae","Syrphinae","Sphaerophoria","Sphaerophoria bankowskae, Sphaerophoria infuscata, Sphaerophoria nr. philanthus, Sphaerophoria batava, Sphaerophoria sp., Sphaerophoria rueppellii, Sphaerophoria interrupta-group, Sphaerophoria sp. MA-1, Sphaerophoria cleoae, Sphaerophoria asymmetrica, Sphaerophoria fatarum, Sphaerophoria bengalensis, Sphaerophoria bifurcata, Sphaerophoria taeniata, Sphaerophoria interrupta, Sphaerophoria contigua, Sphaerophoria kaa, Sphaerophoria laurae, Sphaerophoria chongjini, Sphaerophoria boreoalpina, Sphaerophoria sp. BOLDAAA7374, Sphaerophoria macrogaster, Sphaerophoria sulphuripes, Sphaerophoria brevipilosa, Sphaerophoria philanthus, Sphaerophoria pyrrhina, Sphaerophoria abbreviata, Sphaerophoria scripta, Sphaerophoria potentillae, Sphaerophoria longipilosa, Sphaerophoria virgata","8","8","8","34","0","0","99.22","442.38","0","443.13","31.17","0"
"AGAW2","05S","INS","BOLD:AAG2462","Arthropoda","Insecta","Diptera","Anthomyiidae","Anthomyiinae","Botanophila","Botanophila fugax","7","7","7","36","0","0","99.74","432.29","0","433.71","31.71","0"
"AGAW2","05S","INS","BOLD:AAG6743","Arthropoda","Insecta","Diptera","Sarcophagidae","Sarcophaginae","Sarcophaga","Sarcophaga sp. 1 MG, Sarcophaga subvicina","6","6","6","37","0","0","99.48","427.5","0","428.67","34.9","0"
"AGAW2","05S","INS","BOLD:AAB9140","Arthropoda","Insecta","Diptera","Calliphoridae","Chrysomyinae","Phormia","Phormia regina","6","5","5","37","16.67","0","98.89","428","0","429","32.7","0"
"AGAW2","05S","INS","BOLD:AAN5355","Arthropoda","Insecta","Diptera","Chironomidae","Orthocladiinae","Smittia","Smittia cf. stercoraria, Smittia stercoraria","6","6","6","37","0","0","98.33","414.5","0","422.5","35.8","0"
"AGAW2","05S","INS","BOLD:AAN8770","Arthropoda","Insecta","Diptera","Psychodidae","Psychodinae","Psychoda","Psychoda trinodulosa","6","6","6","37","0","0","99.28","441.5","0","442.17","33.42","0"
"AGAW2","05S","INS","BOLD:AAG6460","Arthropoda","Insecta","Diptera","Cecidomyiidae","","","","6","6","6","37","0","0","98.71","433.5","0","434.5","35","0"
"AGAW2","05S","INS","BOLD:ADF3937","","","","","","","","6","6","6","37","0","0","99.83","394.67","0","399.67","30.97","0"
"AGAW2","05S","INS","BOLD:AAA9899","Arthropoda","Insecta","Hemiptera","Aphididae","Aphidinae","Rhopalosiphum","Rhopalosiphum padi","5","5","5","43","0","0","99.22","436.4","0","437.8","25.69","0"
"AGAW2","05S","INS","BOLD:AAG4462","Arthropoda","Insecta","Coleoptera","Chrysomelidae","Galerucinae","Crepidodera","Crepidodera solita, Crepidodera decora, Crepidodera bella, Crepidodera heikertingeri","5","5","5","43","0","0","98.64","440.8","0","439.8","35.58","0"
"AGAW2","05S","INS","BOLD:AAI4303","Arthropoda","Insecta","Diptera","Chironomidae","Chironominae","Chironomus","Chironomus melanescens","5","5","5","43","0","0","98.41","422","0","425.2","34.17","0"
"AGAW2","05S","INS","BOLD:AAH0256","Arthropoda","Insecta","Coleoptera","Latridiidae","Corticariinae","Corticaria, Corticarina","","5","5","5","43","0","0","99.46","437.2","0","438.8","31.3","0"
"AGAW2","05S","INS","BOLD:AAL7819","Arthropoda","Insecta","Diptera","Psychodidae","Psychodinae","Psychoda","Psychoda sp., Psychoda sp. BOLD:AAL7819","5","5","5","43","0","0","99.4","399.2","0","401.2","32.8","0"
"AGAW2","05S","INS","BOLD:ACQ8463","Arthropoda","Insecta","Diptera","Limoniidae","Limoniinae","Antocha","","4","4","4","48","0","0","98.6","446.5","0","450","37.72","0"
"AGAW2","05S","INS","BOLD:AAB8817","Arthropoda","Insecta","Diptera","Muscidae","Azeliinae","Muscina","Muscina levida","4","4","4","48","0","0","99.61","448.25","0","450","33.44","0"
"AGAW2","05S","INS","BOLD:AAI2681","Arthropoda","Insecta","Diptera","Chironomidae","Orthocladiinae","Parakiefferiella","","4","4","4","48","0","0","97.94","449.25","0","450","34.78","0"
"AGAW2","05S","INS","BOLD:AAN5249","Arthropoda","Insecta","Diptera","Cecidomyiidae","","","","4","4","4","48","0","0","98.09","437.5","0","438.5","33.35","0"
"AGAW2","05S","INS","BOLD:AAC9062","Arthropoda","Insecta","Diptera","Culicidae","Culicinae","Ochlerotatus, Aedes","Aedes fitchii, Ochlerotatus fitchii, Aedes grossbecki, Aedes ventrovittis","4","4","4","48","0","0","99.23","410.75","0","413","33.52","0"
"AGAW2","05S","INS","BOLD:ADH9310","Arthropoda","Insecta","Diptera","Anthomyiidae","Anthomyiinae","Anthomyia","Anthomyia procellaris, Anthomyia pluvialis","4","4","4","48","0","0","98.98","417.75","0","421","32.06","0"
"AGAW2","05S","INS","BOLD:AAA3181","Arthropoda","Insecta","Diptera","Muscidae","Muscinae","Stomoxys","Stomoxys calcitrans","2","2","2","54","0","0","99.3","430.5","0","433.5","32.74","0"
"AGAW2","05S","INS","BOLD:AAA4121","Arthropoda","Insecta","Diptera","Simuliidae","Simuliinae","Simulium","Simulium vittatum complex, Simulium vittatum, Simulium sp. BOLDAAA4121, Simulium argus, Simulium vittatum s.str.","2","2","2","54","0","0","97.84","425","0","427","33.23","0"
"AGAW2","05S","INS","BOLD:AAL2750","Arthropoda","Insecta","Diptera","Muscidae","Phaoniinae","Phaonia","Phaonia tuguriorum","2","2","2","54","0","0","97.99","398.5","0","401.5","31.14","0"
"AGAW2","05S","INS","BOLD:AAB1098","Arthropoda","Insecta","Diptera","Culicidae","Culicinae","Ochlerotatus, Aedes","Aedes excrucians, Aedes cantans, Aedes stimulans, Aedes euedes, Aedes riparius, Aedes annulipes, Ochlerotatus cantans, Aedes fitchii, Aedes sp., Ochlerotatus annulipes, Ochlerotatus excrucians","2","2","2","54","0","0","97.53","446","0","449","35.41","0"
"AGAW2","05S","INS","BOLD:AAH7158","Arthropoda","Insecta","Diptera","Sarcophagidae","Sarcophaginae","Ravinia","Ravinia stimulans","2","2","2","54","0","0","99.07","418","0","421.5","33.22","0"
"AGAW2","05S","INS","BOLD:AAB2384","Arthropoda","Insecta","Diptera","Syrphidae","Syrphinae","Eupeodes","Eupeodes duseki, Eupeodes volucris, Eupeodes nielseni, Eupeodes americanus, Eupeodes flukei, Eupeodes vockerothi, Eupeodes lucasi, Eupeodes montivagus, Eupeodes rufipunctatus, Eupeodes rojasi, Eupeodes goeldlini, Eupeodes luniger, Eupeodes punctifer, Eupeodes bucculatus, Eupeodes perplexus, Eupeodes abiskoensis, Eupeodes corollae, Eupeodes pomus, Eupeodes lundbecki, Eupeodes nitens, Eupeodes sp. BOLDAAB2384, Eupeodes curtus, Eupeodes latifasciatus","2","2","2","54","0","0","99.07","427","0","430","30.43","0"
"AGAW2","05S","INS","BOLD:AAN6437","Arthropoda","Insecta","Diptera","Sciaridae","","Corynoptera","Corynoptera saccata","2","2","2","54","0","0","98.62","436.5","0","436.5","39.87","0"
"AGAW2","05S","INS","BOLD:ABV1237","Arthropoda","Insecta","Diptera","Chironomidae","Chironominae","Chironomus","","2","2","2","54","0","0","98.09","429.5","0","429.5","34.55","0"
"AGAW2","05S","INS","BOLD:AAD8086","Arthropoda","Insecta","Lepidoptera","Elachistidae","Elachistinae","Elachista","Elachista griseicornis, Elachista acenteta","2","2","2","54","0","0","98.43","422","0","423","30.01","0"
"AGAW2","05S","INS","BOLD:AAE4298","Arthropoda","Insecta","Diptera","Chironomidae","Orthocladiinae","Cricotopus","Cricotopus tremulus","2","2","2","54","0","0","98.34","386.5","0","387","32.06","0"
"AGAW2","05S","INS","BOLD:AAG4653","Arthropoda","Insecta","Diptera","Syrphidae","Syrphinae","Heringia","Heringia latitarsis, Heringia pubescens, Heringia jakutorum, Heringia rita, Heringia fulvimanus, Heringia sp. BOLDAAG4653, Heringia nr. pubescens, Heringia coxalis, Heringia larusi, Heringia vitripennis","2","2","2","54","0","0","99.88","435","0","435","31.02","0"
"AGAW2","05S","INS","BOLD:AAA8451","Arthropoda","Insecta","Lepidoptera","Hepialidae","","Korscheltellus","Korscheltellus lupulina","2","2","2","54","0","0","99.11","449.5","0","450","28.44","0"
"AGAW2","05S","INS","BOLD:AAH4135","Arthropoda","Insecta","Diptera","Chloropidae","Chloropinae","Thaumatomyia","Thaumatomyia glabra, Thaumatomyia grata, Thaumatomyia sp. 1 TAW","1","1","1","66","0","0","99.21","380","0","382","31.15","0"
"AGAW2","05S","INS","BOLD:AAA3933","Arthropoda","Insecta","Lepidoptera","Erebidae","Herminiinae","Palthis","Palthis angulalis","1","1","1","66","0","0","98.44","450","0","450","32","0"
"AGAW2","05S","INS","BOLD:ACF4734","Arthropoda","Insecta","Diptera","Syrphidae","Syrphinae","Platycheirus","Platycheirus hyperboreus","1","1","1","66","0","0","97.99","448","0","450","30.89","0"
"AGAW2","05S","INS","BOLD:AAU6630","Arthropoda","Insecta","Diptera","Fanniidae","Fanniinae","Fannia","Fannia armata","1","1","1","66","0","0","98.89","449","0","450","33.78","0"
"AGAW2","05S","INS","BOLD:ABV0271","Arthropoda","Insecta","Diptera","Chloropidae","Oscinellinae","","","1","1","1","66","0","0","99.47","380","0","381","32.55","0"
"AGAW2","05S","INS","BOLD:AAN6447","Arthropoda","Insecta","Diptera","Sciaridae","","Corynoptera","Corynoptera perpusilla","1","1","1","66","0","0","97.54","447","0","450","36.44","0"
"AGAW2","05S","INS","BOLD:AAN5250","Arthropoda","Insecta","Diptera","Cecidomyiidae","","","CecidInt50 sp. BOLD:AAN5250","1","1","1","66","0","0","99.56","450","0","450","29.11","0"
"AGAW2","05S","INS","BOLD:AAG2465","Arthropoda","Insecta","Diptera","Anthomyiidae","Anthomyiinae","Delia","Delia radicum","1","1","1","66","0","0","98.3","411","0","415","32.77","0"
"AGAW2","05S","INS","BOLD:ABY2036","Arthropoda","Insecta","Diptera","Sciaridae","","","","1","1","1","66","0","0","99.33","450","0","450","30.22","0"
"AGAW2","05S","INS","BOLD:AAU6592","Arthropoda","Insecta","Diptera","Ceratopogonidae","Dasyheleinae","Dasyhelea","Dasyhelea arenivaga","1","1","1","66","0","0","99.44","358","0","362","33.7","0"
"AGAW2","05S","INS","BOLD:AAP6843","Arthropoda","Insecta","Diptera","Cecidomyiidae","","","CecidInt3 sp. BOLD:AAP6843","1","1","1","66","0","0","98.97","389","0","450","29.11","0"
"AGAW2","05S","INS","BOLD:AAG5443","Arthropoda","Insecta","Diptera","Chironomidae","Orthocladiinae","","","1","1","1","66","0","0","100","385","0","386","33.16","0"
"AGAW2","05S","INS","BOLD:AAA5565","Arthropoda","Insecta","Hemiptera","Aphididae","Aphidinae","Aphis","Aphis fabae, Aphis newtoni, Aphis solanella, Aphis ilicis, Aphis hederae, Aphis viburni, Aphis lambersi, Aphis fukii","1","1","1","66","0","0","97.56","410","0","450","27.78","0"
"AGAW2","05S","INS","BOLD:ABY4763","Arthropoda","Insecta","Hemiptera","Aphididae","Aphidinae","Uroleucon","Uroleucon taraxaci","1","1","1","66","0","0","99.33","446","0","450","25.11","0"
"AGAW2","05S","INS","BOLD:AAB7105","Arthropoda","Insecta","Hymenoptera","Vespidae","Polistinae","Polistes","Polistes dominula","1","1","1","66","0","0","98.22","449","0","450","27.33","0"
"AGAW2","05S","INS","BOLD:ABZ4179","Arthropoda","Insecta","Hymenoptera","Braconidae","Agathidinae","Bassus","Bassus cinctus","1","1","1","66","0","0","100","381","0","383","22.98","0"
"AGAW2","05S","INS","BOLD:ABU5525","Arthropoda","Insecta","Diptera","Chironomidae","Orthocladiinae","Limnophyes","Limnophyes sp. 14ES","1","1","1","66","0","0","99.78","448","0","450","37.33","0"
"AGAW2","05S","INS","BOLD:ABV0456","Arthropoda","Insecta","Diptera","Cecidomyiidae","","","","1","1","1","66","0","0","98.83","429","0","431","27.38","0"
"AGAW2","05S","INS","BOLD:AAM7064","Arthropoda","Insecta","Diptera","Chironomidae","Orthocladiinae","Smittia","Smittia sp. 14ES","1","1","1","66","0","0","97.73","441","0","444","31.76","0"
"AGAW2","05S","INS","BOLD:AAA2815","Arthropoda","Insecta","Lepidoptera","Noctuidae","Noctuinae","Mythimna","Mythimna yukonensis, Mythimna oxygala","1","1","1","66","0","0","99.33","450","0","450","33.56","0"
"AGAW2","05S","INS","BOLD:AAB7030","Arthropoda","Insecta","Diptera","Chironomidae","Chironominae","Chironomus","Chironomus sp. BOLDAAP2996, Chironomus quinnitukqut, Chironomus cf. decorus 2g, Chironomus decorus group sp. 2 IP2013, Chironomus cf. decorus, Chironomus decorus gr.","1","1","1","66","0","0","97.77","448","0","450","34.22","0"
"AGAW2","05S","INS","BOLD:AAA9101","Arthropoda","Insecta","Lepidoptera","Pyralidae","Phycitinae","Moodna, Caudellia","Moodna ostrinella, Moodna bisinuella, Caudellia nigrella","1","1","1","66","0","0","99.78","450","0","450","34.22","0"
"AGAW2","05S","INS","BOLD:AAV5325","Arthropoda","Insecta","Diptera","Cecidomyiidae","","","CecidInt12 sp. BOLD:AAV5325","1","1","1","66","0","0","97.54","447","0","450","27.78","0"
"AGAW2","05S","INS","BOLD:AAG6745","Arthropoda","Insecta","Diptera","Calliphoridae","Calliphorinae","Pollenia","Pollenia pediculata","1","1","1","66","0","0","98.98","392","0","394","33.25","0"
"AGAW2","05S","INS","BOLD:AAG2128","Arthropoda","Insecta","Diptera","Tachinidae","Exoristinae","Cryptomeigenia","Cryptomeigenia sp. 3, Cryptomeigenia triangularis, Cryptomeigenia muscoides, Cryptomeigenia CR 82, Cryptomeigenia sp. GER5, Cryptomeigenia theutis, Cryptomeigenia sp. GER3","1","1","1","66","0","0","97.1","448","0","450","32","0"
"AGAW2","05S","INS","BOLD:ABW2407","Arthropoda","Insecta","Diptera","Anthomyiidae","Anthomyiinae","Hylemya","","1","1","1","66","0","0","97.1","449","0","450","32.67","0"
"AGAW2","05S","INS","BOLD:AAQ3566","Arthropoda","Insecta","Diptera","Anthomyiidae","Pegomyinae","Pegomya","","1","1","1","66","0","0","98.31","415","0","415","31.57","0"
"AGAW2","05S","INS","BOLD:ACV2677","Arthropoda","Insecta","Diptera","Cecidomyiidae","","","","1","1","1","66","0","0","97.56","450","0","450","30.67","0"
"AGAW2","05S","INS","BOLD:AAD4355","Arthropoda","Insecta","Diptera","Culicidae","Culicinae","Aedes","Aedes provocans","1","1","1","66","0","0","97.16","422","0","422","32.46","0"
"AGAW2","05S","INS","BOLD:AAN5264","Arthropoda","Insecta","Diptera","Cecidomyiidae","","","","1","1","1","66","0","0","97.09","447","0","450","27.11","0"
"AGAW2","05S","INS","BOLD:AAA3654","Arthropoda","Insecta","Lepidoptera","Crambidae","Crambinae","Crambus","Crambus bigelovi, Crambus johnsoni, Crambus praefectellus","1","1","1","66","0","0","98.89","449","0","450","31.78","0"
"AGAW2","05S","INS","BOLD:ACF7232","Arthropoda","Insecta","Diptera","Chironomidae","Orthocladiinae","Parakiefferiella","","1","1","1","66","0","0","98.43","383","0","406","36.7","0"
"AGAW2","05S","INS","BOLD:AAF4817","Arthropoda","Insecta","Diptera","Chironomidae","Orthocladiinae","Smittia","Smittia edwardsi","1","1","1","66","0","0","98.72","391","0","450","33.11","0"
"AGAW2","05S","INS","BOLD:AAG1701","Arthropoda","Insecta","Diptera","Muscidae","Coenosiinae","Macrorchis","Macrorchis ausoba","1","1","1","66","0","0","97.56","450","0","450","33.11","0"
"AGAW2","05S","INS","BOLD:AAG0161","Arthropoda","Insecta","Diptera","Milichiidae","Madizinae","Leptometopa","Leptometopa latipes","1","1","1","66","0","0","99.78","450","0","450","32.44","0"
"AGAW2","05S","INS","BOLD:AAA1513","Arthropoda","Insecta","Lepidoptera","Plutellidae","Plutellinae","Plutella, Rhigognostis","Plutella xylostella, Rhigognostis sp.","1","1","1","66","0","0","99.56","450","0","450","32.67","0"
"AGAW2","05S","INS","BOLD:AAA4782","Arthropoda","Insecta","Hymenoptera","Braconidae","Microgastrinae","Protapanteles, Glyptapanteles","","1","1","1","66","0","0","97.26","365","0","367","24.25","0"
"AGAW2","05S","INS","BOLD:AAA3080","Arthropoda","Insecta","Trichoptera","Limnephilidae","Limnephilinae","Limnephilus","Limnephilus submonilifer","1","1","1","66","0","0","97.88","424","0","428","34.11","0"
"AGAW2","05S","INS","BOLD:AAN6154","Arthropoda","Insecta","Coleoptera","Latridiidae","Corticariinae","Melanophthalma","Melanophthalma inermis","1","1","1","66","0","0","98.89","450","0","450","31.33","0"
"AGAW2","05S","INS","BOLD:AAA5803","Arthropoda","Insecta","Hemiptera","Miridae","Mirinae","Lygus","Lygus lineolaris","1","1","1","66","0","0","99.33","449","0","450","34.67","0"
"AGAW2","05S","INS","BOLD:AAD0853","Arthropoda","Insecta","Diptera","Scathophagidae","Scathophaginae","Scathophaga","Scathophaga lutaria, Scathophaga hiemalis, Scathophaga furcata, Scathophaga stercoraria, Scathophaga inquinata, Scathophaga suilla","1","1","1","66","0","0","98.19","441","0","441","31.97","0"
"AGAW2","05S","NAR","no match","no match","no match","no match","no match","no match","no match","no match","no match","no match","no match","no match","no match","no match","no match","no match","no match","no match","no match","no match"
"AGAW2","05S","NIN","BOLD:AAH6630","Arthropoda","Collembola","Entomobryomorpha","Entomobryidae","Entomobryinae","Entomobrya","Entomobrya unostrigata","67","64","64","1","4.48","0","99.25","434.49","0","435.97","38.89","0"
"AGAW2","05S","NIN","BOLD:AAV0589","Arthropoda","Arachnida","Mesostigmata","Blattisociidae","Platyseiinae","Cheiroseius","","51","50","50","2","1.96","0","99.1","440.57","0","444.57","27.52","0"
"AGAW2","05S","NIN","BOLD:ABW2639","Arthropoda","Arachnida","Trombidiformes","Anystidae","","","","3","3","3","3","0","0","99.4","446.67","0","448.33","31.6","0"
"AGAW2","05S","NIN","BOLD:AAC7802","Arthropoda","Arachnida","Araneae","Lycosidae","","Pardosa","Pardosa distincta","1","1","1","4","0","0","99.78","449","0","450","29.33","0"
"AGAW2","05W","BAC","TAX:91391","Proteobacteria","Alphaproteobacteria","Rickettsiales","",NA,"","","84","73","73","1","13.1","0","97.32","408.36","0","425.56","39.61","0"
"AGAW2","05W","BAC","TAX:392851","Proteobacteria","Alphaproteobacteria","Rickettsiales","Rickettsiaceae",NA,"Wolbachia","Wolbachia endosymbiont of Formicidae","2","1","1","2","50","0","98.87","443","0","450","41.78","0"
"AGAW2","05W","BAC","TAX:375853","Proteobacteria","Alphaproteobacteria","Rickettsiales","Rickettsiaceae",NA,"Wolbachia","Wolbachia sp. Hymenoptera_ASGLE646-10","1","1","1","3","0","0","97.77","449","0","450","40.67","0"
"AGAW2","05W","BAC","TAX:375847","Proteobacteria","Alphaproteobacteria","Rickettsiales","Rickettsiaceae",NA,"Wolbachia","Wolbachia sp. Hymenoptera_ASGLE1705-10","1","1","1","3","0","0","99.01","405","0","406","39.66","0"
"AGAW2","05W","BAC","TAX:392874","Proteobacteria","Alphaproteobacteria","Rickettsiales","Rickettsiaceae",NA,"Wolbachia","Wolbachia endosymbiont of Hymenoptera","1","1","1","3","0","0","98.88","445","0","450","40.44","0"
"AGAW2","05W","BAC","TAX:376688","Proteobacteria","Alphaproteobacteria","Rickettsiales","Rickettsiaceae",NA,"Wolbachia","Wolbachia sp. elachBioLep01 BioLep373","1","1","1","3","0","0","99.11","450","0","450","40.44","0"
"AGAW2","05W","CHO","no match","no match","no match","no match","no match","no match","no match","no match","no match","no match","no match","no match","no match","no match","no match","no match","no match","no match","no match","no match"
"AGAW2","05W","INS","BOLD:AAF9187","Arthropoda","Insecta","Coleoptera","Curculionidae","Entiminae","Phyllobius","Phyllobius oblongus","29335","20390","20390","1","30.49","0","99.49","436.44","0","438.92","38.88","0"
"AGAW2","05W","INS","BOLD:AAA8451","Arthropoda","Insecta","Lepidoptera","Hepialidae","","Korscheltellus","Korscheltellus lupulina","18335","11699","11699","2","36.19","0","99.62","445.36","0","446.43","28.18","0"
"AGAW2","05W","INS","BOLD:AAO3939","Arthropoda","Insecta","Diptera","Limoniidae","Limoniinae","Dicranomyia","","13541","10614","10614","3","21.62","0","99.12","438.71","0","440.06","36.24","0"
"AGAW2","05W","INS","BOLD:AAG2511","Arthropoda","Insecta","Diptera","Anthomyiidae","Anthomyiinae","Delia","Delia platura","5012","4279","4279","4","14.62","0","99.03","444.27","0","446.77","33.17","0"
"AGAW2","05W","INS","BOLD:AAA3453","Arthropoda","Insecta","Diptera","Anthomyiidae","Anthomyiinae","Delia","","2583","2358","2358","5","8.71","0","99.22","431.77","0","434.42","34.67","0"
"AGAW2","05W","INS","BOLD:ABW7897","Arthropoda","Insecta","Diptera","Cecidomyiidae","","","","1840","1305","1305","6","29.08","0","99.6","442.03","0","443.98","29.73","0"
"AGAW2","05W","INS","BOLD:ACX5521","Arthropoda","Insecta","Diptera","Chironomidae","Orthocladiinae","","","1838","1754","1754","7","4.57","0","98.83","407.66","0","409.4","35.08","0"
"AGAW2","05W","INS","BOLD:AAA7661","Arthropoda","Insecta","Diptera","Culicidae","Culicinae","Culiseta, Culex","Culiseta melanura, Culex articularis, Culex restuans, Culex acharistus","1827","1677","1677","8","8.21","0","99.07","432.63","0","434.66","33.02","0"
"AGAW2","05W","INS","BOLD:AAN5358","Arthropoda","Insecta","Diptera","Chironomidae","Orthocladiinae","Smittia","Smittia stercoraria, Smittia sp. 22ES","1672","1419","1419","9","15.13","0","99.31","440.98","0","442.59","37.31","0"
"AGAW2","05W","INS","BOLD:AAH3664","Arthropoda","Insecta","Diptera","Cecidomyiidae","","","CecidInt3 sp. BOLD:AAH3664","1579","1484","1484","10","6.02","0","99.14","427.67","0","431.17","27.02","0"
"AGAW2","05W","INS","BOLD:AAG0314","Arthropoda","Insecta","Diptera","Chironomidae","Tanypodinae","Psectrotanypus","Psectrotanypus sp. ES01","1499","1371","1371","11","8.54","0","99.09","440.06","0","441.99","34.37","0"
"AGAW2","05W","INS","BOLD:AAK5928","Arthropoda","Insecta","Coleoptera","Chrysomelidae","Criocerinae","Oulema","Oulema melanopus","377","329","329","12","12.73","0","99.52","436.94","0","438.16","36.42","0"
"AGAW2","05W","INS","BOLD:ACW5117","Arthropoda","Insecta","Diptera","Chironomidae","Orthocladiinae","Smittia","Smittia leucopogon","309","289","289","13","6.47","0","99.23","424.59","0","425.88","31.43","0"
"AGAW2","05W","INS","BOLD:AAU6544","Arthropoda","Insecta","Diptera","Limoniidae","Chioneinae","Ormosia","Ormosia affinis","220","215","215","14","2.27","0","98.85","428.43","0","430.3","38.75","0"
"AGAW2","05W","INS","BOLD:AAB2644","Arthropoda","Insecta","Diptera","Chironomidae","Orthocladiinae","Orthocladius","Orthocladius lapponicus, Orthocladius dorenus, Orthocladius mallochi","197","168","168","15","14.72","0","99.28","443.52","0","445.17","32.17","0"
"AGAW2","05W","INS","BOLD:AAD2548","Arthropoda","Insecta","Hemiptera","Aphididae","Aphidinae","Acyrthosiphon","Acyrthosiphon caraganae, Acyrthosiphon sp. 1, Acyrthosiphon kondoi, Acyrthosiphon churchillense","118","93","93","16","21.19","0","99.57","446.92","0","447.78","25.98","0"
"AGAW2","05W","INS","BOLD:ACO1622","Arthropoda","Insecta","Diptera","Cecidomyiidae","","","","103","103","103","17","0","0","98.8","417.83","0","419.43","31.62","0"
"AGAW2","05W","INS","BOLD:AAV1261","Arthropoda","Insecta","Diptera","Sciaridae","","Bradysia","Bradysia splendida","75","74","74","18","1.33","0","99.16","435.19","0","436.44","34.02","0"
"AGAW2","05W","INS","BOLD:ACF2924","","","","","","","","72","57","57","19","20.83","0","99.7","447.86","0","448.78","25.72","0"
"AGAW2","05W","INS","BOLD:ADF6661","Arthropoda","Insecta","Diptera","Cecidomyiidae","","","","63","55","55","20","12.7","0","99.24","416.78","0","445.56","30.06","0"
"AGAW2","05W","INS","BOLD:AAH0255","Arthropoda","Insecta","Coleoptera","Chrysomelidae","Galerucinae","Phyllotreta","Phyllotreta cruciferae, Phyllotreta albionica","39","38","38","21","2.56","0","99.29","432.28","0","433.95","32.72","0"
"AGAW2","05W","INS","BOLD:ACJ1357","Arthropoda","Insecta","Diptera","Cecidomyiidae","","","","38","37","37","22","2.63","0","99.3","405.53","0","442.42","29.1","0"
"AGAW2","05W","INS","BOLD:AAF9014","Arthropoda","Insecta","Diptera","Limoniidae","Chioneinae","Symplecta","Symplecta cana, Symplecta sp. BOLDAAF9014","35","35","35","23","0","0","97.93","403.91","0","408.06","33.31","0"
"AGAW2","05W","INS","BOLD:AAA9899","Arthropoda","Insecta","Hemiptera","Aphididae","Aphidinae","Rhopalosiphum","Rhopalosiphum padi","30","30","30","24","0","0","99.5","440.73","0","441.8","25.92","0"
"AGAW2","05W","INS","BOLD:AAM9242","Arthropoda","Insecta","Diptera","Sciaridae","","Corynoptera","Corynoptera melanochaeta","19","19","19","25","0","0","99.37","425.84","0","427.47","34.39","0"
"AGAW2","05W","INS","BOLD:ADH6290","","","","","","","","14","14","14","26","0","0","99.03","445.36","0","446.14","39.21","0"
"AGAW2","05W","INS","BOLD:AAN8770","Arthropoda","Insecta","Diptera","Psychodidae","Psychodinae","Psychoda","Psychoda trinodulosa","13","13","13","27","0","0","99.55","442.85","0","443.77","33.92","0"
"AGAW2","05W","INS","BOLD:AAN5267","Arthropoda","Insecta","Diptera","Cecidomyiidae","","","","9","9","9","28","0","0","99.55","438.44","0","439.67","30.5","0"
"AGAW2","05W","INS","BOLD:AAA7374","Arthropoda","Insecta","Diptera","Syrphidae","Syrphinae","Sphaerophoria","Sphaerophoria bankowskae, Sphaerophoria infuscata, Sphaerophoria nr. philanthus, Sphaerophoria batava, Sphaerophoria sp., Sphaerophoria rueppellii, Sphaerophoria interrupta-group, Sphaerophoria sp. MA-1, Sphaerophoria cleoae, Sphaerophoria asymmetrica, Sphaerophoria fatarum, Sphaerophoria bengalensis, Sphaerophoria bifurcata, Sphaerophoria taeniata, Sphaerophoria interrupta, Sphaerophoria contigua, Sphaerophoria kaa, Sphaerophoria laurae, Sphaerophoria chongjini, Sphaerophoria boreoalpina, Sphaerophoria sp. BOLDAAA7374, Sphaerophoria macrogaster, Sphaerophoria sulphuripes, Sphaerophoria brevipilosa, Sphaerophoria philanthus, Sphaerophoria pyrrhina, Sphaerophoria abbreviata, Sphaerophoria scripta, Sphaerophoria potentillae, Sphaerophoria longipilosa, Sphaerophoria virgata","9","9","9","28","0","0","99.46","432","0","434.11","30.8","0"
"AGAW2","05W","INS","BOLD:AAA4222","Arthropoda","Insecta","Lepidoptera","Erebidae","Hypeninae","Hypena","Hypena scabra","8","8","8","30","0","0","99.02","386.75","0","390","32.31","0"
"AGAW2","05W","INS","BOLD:AAL9076","Arthropoda","Insecta","Diptera","Phoridae","Metopininae","Megaselia","","8","8","8","30","0","0","98.87","404","0","406.25","32.36","0"
"AGAW2","05W","INS","BOLD:AAC9199","Arthropoda","Insecta","Diptera","Chironomidae","Tanypodinae","Krenopelopia","","8","8","8","30","0","0","98.22","382.38","0","393.38","34.67","0"
"AGAW2","05W","INS","BOLD:AAH7428","Arthropoda","Insecta","Hymenoptera","Braconidae","Aphidiinae","","","7","7","7","33","0","0","98.85","404.43","0","408.14","27.28","0"
"AGAW2","05W","INS","BOLD:AAG2868","Arthropoda","Insecta","Hemiptera","Cicadellidae","Typhlocybinae","Empoasca","Empoasca sp., Empoasca fabae","6","6","6","34","0","0","98.74","448.83","0","450","32.59","0"
"AGAW2","05W","INS","BOLD:AAH2753","Arthropoda","Insecta","Coleoptera","Anthicidae","Anthicinae","Anthicus","Anthicus cervinus","6","6","6","34","0","0","97.69","380.33","0","382.67","33.07","0"
"AGAW2","05W","INS","BOLD:AAB9140","Arthropoda","Insecta","Diptera","Calliphoridae","Chrysomyinae","Phormia","Phormia regina","5","5","5","36","0","0","98.89","449","0","450","33.07","0"
"AGAW2","05W","INS","BOLD:AAD8855","Arthropoda","Insecta","Diptera","Chironomidae","Chironominae","Tanytarsus","Tanytarsus glabrescens","5","5","5","36","0","0","98.07","392.8","0","394.4","35.79","0"
"AGAW2","05W","INS","BOLD:AAG6743","Arthropoda","Insecta","Diptera","Sarcophagidae","Sarcophaginae","Sarcophaga","Sarcophaga sp. 1 MG, Sarcophaga subvicina","5","5","5","36","0","0","99.82","449.4","0","450","35.11","0"
"AGAW2","05W","INS","BOLD:AAF9771","Arthropoda","Insecta","Diptera","Hybotidae","Tachydromiinae","Platypalpus","Platypalpus longiseta, Platypalpus trivialis, Platypalpus sp. 5, Platypalpus sp. 17, Platypalpus pallidiventris","4","3","3","39","25","0","98.45","450.25","0","450","34.94","0"
"AGAW2","05W","INS","BOLD:AAA9251","Arthropoda","Insecta","Lepidoptera","Gelechiidae","Gelechiinae","Scrobipalpa","Scrobipalpa atriplicella, Scrobipalpa nr. atriplicella sp. 1","4","4","4","39","0","0","98.6","388.25","0","391.25","33.58","0"
"AGAW2","05W","INS","BOLD:AAH3759","Arthropoda","Insecta","Diptera","Cecidomyiidae","","","","4","4","4","39","0","0","98.2","425.25","0","431.25","29.1","0"
"AGAW2","05W","INS","BOLD:ACV2677","Arthropoda","Insecta","Diptera","Cecidomyiidae","","","","4","4","4","39","0","0","97.44","449.5","0","450","30.22","0"
"AGAW2","05W","INS","BOLD:AAG4653","Arthropoda","Insecta","Diptera","Syrphidae","Syrphinae","Heringia","Heringia latitarsis, Heringia pubescens, Heringia jakutorum, Heringia rita, Heringia fulvimanus, Heringia sp. BOLDAAG4653, Heringia nr. pubescens, Heringia coxalis, Heringia larusi, Heringia vitripennis","4","4","4","39","0","0","98.77","447.75","0","450","31.61","0"
"AGAW2","05W","INS","BOLD:AAA3836","Arthropoda","Insecta","Lepidoptera","Noctuidae","Plusiinae","Autographa","Autographa precationis","3","3","3","44","0","0","98.81","449","0","450","32.07","0"
"AGAW2","05W","INS","BOLD:AAN5355","Arthropoda","Insecta","Diptera","Chironomidae","Orthocladiinae","Smittia","Smittia cf. stercoraria, Smittia stercoraria","3","3","3","44","0","0","98.65","408.67","0","449","36.53","0"
"AGAW2","05W","INS","BOLD:ACK3431","Arthropoda","Insecta","Diptera","Cecidomyiidae","","","","3","3","3","44","0","0","98.84","434.33","0","436.67","32.97","0"
"AGAW2","05W","INS","BOLD:AAU6592","Arthropoda","Insecta","Diptera","Ceratopogonidae","Dasyheleinae","Dasyhelea","Dasyhelea arenivaga","3","3","3","44","0","0","98.78","434.33","0","436.33","35.34","0"
"AGAW2","05W","INS","BOLD:AAG2462","Arthropoda","Insecta","Diptera","Anthomyiidae","Anthomyiinae","Botanophila","Botanophila fugax","3","3","3","44","0","0","99.55","448.67","0","450","31.7","0"
"AGAW2","05W","INS","BOLD:ABV1243","Arthropoda","Insecta","Diptera","Sarcophagidae","Paramacronychiinae, Miltogramminae","Miltogramminae_incertae_sedis, Brachicoma","Miltogramminae_incertae_sedis sp. 1 MG, Brachicoma sarcophagina","2","2","2","49","0","0","98.33","448.5","0","450","34.78","0"
"AGAW2","05W","INS","BOLD:ABV0261","Arthropoda","Insecta","Diptera","Chironomidae","Orthocladiinae","","","2","2","2","49","0","0","100","427.5","0","429","32.97","0"
"AGAW2","05W","INS","BOLD:AAG5457","Arthropoda","Insecta","Diptera","Chironomidae","Chironominae","","","2","2","2","49","0","0","98.78","450.5","0","450","34.22","0"
"AGAW2","05W","INS","BOLD:AAU8455","Arthropoda","Insecta","Hymenoptera","Platygastridae","Platygastrinae","Aceroteta","","2","2","2","49","0","0","97.13","400","0","402.5","25.2","0"
"AGAW2","05W","INS","BOLD:AAU6630","Arthropoda","Insecta","Diptera","Fanniidae","Fanniinae","Fannia","Fannia armata","2","2","2","49","0","0","97.77","448","0","450","33.22","0"
"AGAW2","05W","INS","BOLD:AAA2815","Arthropoda","Insecta","Lepidoptera","Noctuidae","Noctuinae","Mythimna","Mythimna yukonensis, Mythimna oxygala","2","2","2","49","0","0","99.78","448","0","450","33.11","0"
"AGAW2","05W","INS","BOLD:AAB1098","Arthropoda","Insecta","Diptera","Culicidae","Culicinae","Ochlerotatus, Aedes","Aedes excrucians, Aedes cantans, Aedes stimulans, Aedes euedes, Aedes riparius, Aedes annulipes, Ochlerotatus cantans, Aedes fitchii, Aedes sp., Ochlerotatus annulipes, Ochlerotatus excrucians","2","2","2","49","0","0","99.31","430.5","0","432","34.94","0"
"AGAW2","05W","INS","BOLD:AAN5342","Arthropoda","Insecta","Diptera","Chironomidae","Orthocladiinae","Heleniella","","2","2","2","49","0","0","99.01","402.5","0","404","34.57","0"
"AGAW2","05W","INS","BOLD:AAD0853","Arthropoda","Insecta","Diptera","Scathophagidae","Scathophaginae","Scathophaga","Scathophaga lutaria, Scathophaga hiemalis, Scathophaga furcata, Scathophaga stercoraria, Scathophaga inquinata, Scathophaga suilla","2","2","2","49","0","0","98.91","403","0","404","31.5","0"
"AGAW2","05W","INS","BOLD:AAG6745","Arthropoda","Insecta","Diptera","Calliphoridae","Calliphorinae","Pollenia","Pollenia pediculata","2","2","2","49","0","0","99.11","409.5","0","411.5","33.69","0"
"AGAW2","05W","INS","BOLD:AAG3236","Arthropoda","Insecta","Diptera","Phoridae","Phorinae","Conicera, Diplonevra","Diplonevra nitidula","2","2","2","49","0","0","99.21","413","0","414","32.4","0"
"AGAW2","05W","INS","BOLD:AAB2384","Arthropoda","Insecta","Diptera","Syrphidae","Syrphinae","Eupeodes","Eupeodes duseki, Eupeodes volucris, Eupeodes nielseni, Eupeodes americanus, Eupeodes flukei, Eupeodes vockerothi, Eupeodes lucasi, Eupeodes montivagus, Eupeodes rufipunctatus, Eupeodes rojasi, Eupeodes goeldlini, Eupeodes luniger, Eupeodes punctifer, Eupeodes bucculatus, Eupeodes perplexus, Eupeodes abiskoensis, Eupeodes corollae, Eupeodes pomus, Eupeodes lundbecki, Eupeodes nitens, Eupeodes sp. BOLDAAB2384, Eupeodes curtus, Eupeodes latifasciatus","2","2","2","49","0","0","98.51","438.5","0","441","31.29","0"
"AGAW2","05W","INS","BOLD:AAC9062","Arthropoda","Insecta","Diptera","Culicidae","Culicinae","Ochlerotatus, Aedes","Aedes fitchii, Ochlerotatus fitchii, Aedes grossbecki, Aedes ventrovittis","2","2","2","49","0","0","99.89","448.5","0","450","33.56","0"
"AGAW2","05W","INS","BOLD:ACP4736","Arthropoda","Insecta","Diptera","Chironomidae","Orthocladiinae","Smittia","Smittia sp. 8ES","2","2","2","49","0","0","98.78","449.5","0","450","33","0"
"AGAW2","05W","INS","BOLD:ACF7232","Arthropoda","Insecta","Diptera","Chironomidae","Orthocladiinae","Parakiefferiella","","2","2","2","49","0","0","97.76","378.5","0","383.5","36.91","0"
"AGAW2","05W","INS","BOLD:ADH9310","Arthropoda","Insecta","Diptera","Anthomyiidae","Anthomyiinae","Anthomyia","Anthomyia procellaris, Anthomyia pluvialis","2","2","2","49","0","0","98.63","419","0","420.5","32.35","0"
"AGAW2","05W","INS","BOLD:AAH3548","Arthropoda","Insecta","Diptera","Sphaeroceridae","Copromyzinae","Lotophila","Lotophila sp., Lotophila atra","1","1","1","65","0","0","99.33","449","0","450","32.22","0"
"AGAW2","05W","INS","BOLD:AAG9055","Arthropoda","Insecta","Hymenoptera","Vespidae","Vespinae","Vespula","Vespula germanica","1","1","1","65","0","0","99.78","449","0","450","31.11","0"
"AGAW2","05W","INS","BOLD:AAH4135","Arthropoda","Insecta","Diptera","Chloropidae","Chloropinae","Thaumatomyia","Thaumatomyia glabra, Thaumatomyia grata, Thaumatomyia sp. 1 TAW","1","1","1","65","0","0","98.67","450","0","450","32.67","0"
"AGAW2","05W","INS","BOLD:AAG5507","Arthropoda","Insecta","Diptera","Chironomidae","Chironominae","Chironomus","Chironomus atrella","1","1","1","65","0","0","98.66","448","0","450","36.22","0"
"AGAW2","05W","INS","BOLD:AAG9668","Arthropoda","Insecta","Diptera","Dolichopodidae","Diaphorinae","Chrysotus","Chrysotus sp., Chrysotus sp. nr. obliquus","1","1","1","65","0","0","98.89","450","0","450","32.22","0"
"AGAW2","05W","INS","BOLD:ACI2531","","","","","","","","1","1","1","65","0","0","98.44","448","0","450","26.67","0"
"AGAW2","05W","INS","BOLD:AAP6369","Arthropoda","Insecta","Coleoptera","Elateridae","Agrypninae","Aeolus","Aeolus mellillus","1","1","1","65","0","0","99.78","450","0","450","38.89","0"
"AGAW2","05W","INS","BOLD:AAI4303","Arthropoda","Insecta","Diptera","Chironomidae","Chironominae","Chironomus","Chironomus melanescens","1","1","1","65","0","0","98.14","376","0","377","33.95","0"
"AGAW2","05W","INS","BOLD:AAA8933","Arthropoda","Insecta","Coleoptera","Coccinellidae","Coccinellinae","Coccinella","Coccinella septempunctata","1","1","1","65","0","0","99.55","448","0","450","33.11","0"
"AGAW2","05W","INS","BOLD:ABA0755","Arthropoda","Insecta","Diptera","Drosophilidae","","","","1","1","1","65","0","0","98.67","376","0","377","31.3","0"
"AGAW2","05W","INS","BOLD:AAU6970","Arthropoda","Insecta","Coleoptera","Cleridae","Clerinae","Enoclerus","Enoclerus nigripes","1","1","1","65","0","0","98.22","450","0","450","33.33","0"
"AGAW2","05W","INS","BOLD:AAQ0265","Arthropoda","Insecta","Diptera","Hybotidae","Tachydromiinae","Platypalpus","Platypalpus pulicarius","1","1","1","65","0","0","98.44","448","0","450","35.11","0"
"AGAW2","05W","INS","BOLD:AAB8817","Arthropoda","Insecta","Diptera","Muscidae","Azeliinae","Muscina","Muscina levida","1","1","1","65","0","0","98","450","0","450","33.78","0"
"AGAW2","05W","INS","BOLD:ACU7906","Arthropoda","Insecta","Diptera","Chironomidae","Orthocladiinae","","","1","1","1","65","0","0","98.4","437","0","439","36.67","0"
"AGAW2","05W","INS","BOLD:AAA5448","Arthropoda","Insecta","Lepidoptera","Geometridae","Ennominae","Eutrapela","Eutrapela clemataria","1","1","1","65","0","0","99.56","450","0","450","30","0"
"AGAW2","05W","INS","BOLD:AAG0995","Arthropoda","Insecta","Diptera","Chironomidae","Orthocladiinae","Parakiefferiella","","1","1","1","65","0","0","97.49","359","0","360","38.61","0"
"AGAW2","05W","INS","BOLD:AAA5299","Arthropoda","Insecta","Diptera","Chironomidae","Orthocladiinae","Cricotopus","Cricotopus laetus, Cricotopus relucens, Cricotopus trifasciatus, Cricotopus sp., Cricotopus sp. 3ES, Cricotopus sylvestris, Cricotopus glacialis","1","1","1","65","0","0","99.28","417","0","422","33.89","0"
"AGAW2","05W","INS","BOLD:AAG0991","Arthropoda","Insecta","Diptera","Chironomidae","Orthocladiinae","Orthocladius","","1","1","1","65","0","0","98.2","389","0","389","31.62","0"
"AGAW2","05W","INS","BOLD:AAB5609","Arthropoda","Insecta","Diptera","Muscidae","Coenosiinae","Coenosia","Coenosia sp.BOLD:AAB5609, Coenosia tigrina","1","1","1","65","0","0","97.78","361","0","366","32.24","0"
"AGAW2","05W","INS","BOLD:AAE0707","Arthropoda","Insecta","Diptera","Chironomidae","Chironominae","Microtendipes","Microtendipes pedellus","1","1","1","65","0","0","99.19","370","0","372","31.18","0"
"AGAW2","05W","INS","BOLD:AAG1743","Arthropoda","Insecta","Diptera","Muscidae","Phaoniinae","Phaonia","Phaonia fuscana","1","1","1","65","0","0","97.21","394","0","395","32.91","0"
"AGAW2","05W","INS","BOLD:ADH4889","Arthropoda","Insecta","Diptera","","","","","1","1","1","65","0","0","98","449","0","450","35.56","0"
"AGAW2","05W","INS","BOLD:AAH0134","Arthropoda","Insecta","Coleoptera","Phalacridae","Phalacrinae","Stilbus","Stilbus apicialis","1","1","1","65","0","0","99.55","449","0","450","32.44","0"
"AGAW2","05W","INS","BOLD:AAG8493","Arthropoda","Insecta","Diptera","Drosophilidae","Drosophilinae","Scaptomyza","Scaptomyza pallida","1","1","1","65","0","0","98.08","417","0","419","32.7","0"
"AGAW2","05W","INS","BOLD:AAY9557","Arthropoda","Insecta","Lepidoptera","Coleophoridae","Coleophorinae","Coleophora","Coleophora deauratella","1","1","1","65","0","0","100","444","0","450","32","0"
"AGAW2","05W","INS","BOLD:ADH8687","Arthropoda","Insecta","Coleoptera","","","","","1","1","1","65","0","0","100","435","0","450","34.44","0"
"AGAW2","05W","INS","BOLD:AAN5332","Arthropoda","Insecta","Diptera","Chironomidae","Chironominae","Polypedilum","","1","1","1","65","0","0","97.54","447","0","450","34","0"
"AGAW2","05W","INS","BOLD:AAN5356","Arthropoda","Insecta","Diptera","Chironomidae","Orthocladiinae","Smittia","","1","1","1","65","0","0","99.55","449","0","450","35.33","0"
"AGAW2","05W","INS","BOLD:AAJ7105","Arthropoda","Insecta","Diptera","Agromyzidae","Agromyzinae","Agromyza","Agromyza frontella","1","1","1","65","0","0","99.25","399","0","450","37.56","0"
"AGAW2","05W","INS","BOLD:AAB7030","Arthropoda","Insecta","Diptera","Chironomidae","Chironominae","Chironomus","Chironomus sp. BOLDAAP2996, Chironomus quinnitukqut, Chironomus cf. decorus 2g, Chironomus decorus group sp. 2 IP2013, Chironomus cf. decorus, Chironomus decorus gr.","1","1","1","65","0","0","97.54","447","0","450","34.22","0"
"AGAW2","05W","INS","BOLD:AAA9101","Arthropoda","Insecta","Lepidoptera","Pyralidae","Phycitinae","Moodna, Caudellia","Moodna ostrinella, Moodna bisinuella, Caudellia nigrella","1","1","1","65","0","0","97.53","446","0","450","34","0"
"AGAW2","05W","INS","BOLD:AAH0145","Arthropoda","Insecta","Coleoptera","Carabidae","Harpalinae","Anisodactylus","Anisodactylus pitychrous, Anisodactylus sanctaecrucis, Anisodactylus alternans, Anisodactylus discoideus","1","1","1","65","0","0","97.99","448","0","450","32.44","0"
"AGAW2","05W","INS","BOLD:ACF5939","Arthropoda","Insecta","Diptera","Tachinidae","Exoristinae","Winthemia","Winthemia sinuata, Winthemia illinoensis","1","1","1","65","0","0","98","449","0","450","29.56","0"
"AGAW2","05W","INS","BOLD:AAN5341","Arthropoda","Insecta","Diptera","Chironomidae","Orthocladiinae","Camptocladius","Camptocladius stercorarius","1","1","1","65","0","0","97.34","376","0","378","35.71","0"
"AGAW2","05W","INS","BOLD:AAM9013","Arthropoda","Insecta","Diptera","Keroplatidae","Macrocerinae","Macrocera","Macrocera pusilla","1","1","1","65","0","0","98.44","450","0","450","30.22","0"
"AGAW2","05W","INS","BOLD:ACF0198","Arthropoda","Insecta","Diptera","Tachinidae","Exoristinae","Winthemia","Winthemia quadripustulata, Winthemia venusta, Winthemia sinuata, Winthemia occidentis, Winthemia erythura, Winthemia illinoensis","1","1","1","65","0","0","99.24","394","0","394","29.44","0"
"AGAW2","05W","INS","BOLD:AAB1787","Arthropoda","Insecta","Hemiptera","Aphididae","Aphidinae","Rhopalosiphum, Acyrthosiphon","Acyrthosiphon pisum, Acyrthosiphon pisum 5A, Rhopalosiphum maidis","1","1","1","65","0","0","97.78","450","0","450","26.22","0"
"AGAW2","05W","INS","BOLD:AAH7158","Arthropoda","Insecta","Diptera","Sarcophagidae","Sarcophaginae","Ravinia","Ravinia stimulans","1","1","1","65","0","0","99.55","449","0","450","33.78","0"
"AGAW2","05W","INS","BOLD:ABX6186","Arthropoda","Insecta","Diptera","Tipulidae","Tipulinae","Nephrotoma","Nephrotoma ferruginea","1","1","1","65","0","0","99.56","450","0","450","34","0"
"AGAW2","05W","INS","BOLD:ABX6298","Arthropoda","Insecta","Diptera","Tipulidae","Tipulinae","Nephrotoma","Nephrotoma ferruginea","1","1","1","65","0","0","98.03","406","0","406","32.02","0"
"AGAW2","05W","INS","BOLD:ACZ1491","Arthropoda","Insecta","Diptera","Chironomidae","","","","1","1","1","65","0","0","97.77","449","0","450","34.67","0"
"AGAW2","05W","INS","BOLD:AAG6971","Arthropoda","Insecta","Diptera","Sciomyzidae","Sciomyzinae","Anticheta","Anticheta sp., Anticheta melanosoma","1","1","1","65","0","0","97.45","431","0","431","39.68","0"
"AGAW2","05W","INS","BOLD:AAD8086","Arthropoda","Insecta","Lepidoptera","Elachistidae","Elachistinae","Elachista","Elachista griseicornis, Elachista acenteta","1","1","1","65","0","0","98.41","440","0","444","31.53","0"
"AGAW2","05W","INS","BOLD:AAD1485","Arthropoda","Insecta","Diptera","Chironomidae","Chironominae","Paratanytarsus","Paratanytarsus grimmii, Paratanytarsus sp.","1","1","1","65","0","0","98.82","425","0","428","32.71","0"
"AGAW2","05W","INS","BOLD:AAM7726","Arthropoda","Insecta","Coleoptera","Curculionidae","Curculioninae","Orchestes","Orchestes steppensis, Orchestes alni","1","1","1","65","0","0","99.56","450","0","450","33.33","0"
"AGAW2","05W","INS","BOLD:ABU5525","Arthropoda","Insecta","Diptera","Chironomidae","Orthocladiinae","Limnophyes","Limnophyes sp. 14ES","1","1","1","65","0","0","97.81","365","0","367","38.15","0"
"AGAW2","05W","INS","BOLD:ABV1229","Arthropoda","Insecta","Diptera","Chironomidae","Orthocladiinae","Eukiefferiella","","1","1","1","65","0","0","97.04","371","0","374","35.56","0"
"AGAW2","05W","INS","BOLD:AAN6154","Arthropoda","Insecta","Coleoptera","Latridiidae","Corticariinae","Melanophthalma","Melanophthalma inermis","1","1","1","65","0","0","98.66","449","0","450","31.56","0"
"AGAW2","05W","NAR","no match","no match","no match","no match","no match","no match","no match","no match","no match","no match","no match","no match","no match","no match","no match","no match","no match","no match","no match","no match"
"AGAW2","05W","NIN","BOLD:AAI4346","Arthropoda","Arachnida","Opiliones","Phalangiidae","","Paroligolophus, Phalangium","Paroligolophus agrestis, Phalangium opilio","4563","4176","4176","1","8.48","0","99.16","435.71","0","437.9","36.32","0"
"AGAW2","05W","NIN","BOLD:AAB7667","Arthropoda","Arachnida","Araneae","Lycosidae","","Pardosa","Pardosa saxatilis","1128","1086","1086","2","3.72","0","99.19","402.41","0","406.55","31","0"
"AGAW2","05W","NIN","BOLD:ABX0027","Arthropoda","Collembola","Symphypleona","Bourletiellidae","","Bourletiella","","143","114","114","3","20.28","0","99.63","415.39","0","442.67","35.62","0"
"AGAW2","05W","NIN","BOLD:AAH6630","Arthropoda","Collembola","Entomobryomorpha","Entomobryidae","Entomobryinae","Entomobrya","Entomobrya unostrigata","111","107","107","4","3.6","0","99.01","439.95","0","441.08","39.27","0"
"AGAW2","05W","NIN","BOLD:AAB7915","Arthropoda","Collembola","Symphypleona","Bourletiellidae","","Deuterosminthurus","Deuterosminthurus sp1","36","33","33","5","8.33","0","98.84","444.22","0","444.94","35.09","0"
"AGAW2","05W","NIN","BOLD:ABW2639","Arthropoda","Arachnida","Trombidiformes","Anystidae","","","","26","24","24","6","7.69","0","99.59","452.19","0","442.54","31.54","0"
"AGAW2","05W","NIN","BOLD:AAB8452","Arthropoda","Collembola","Entomobryomorpha","Entomobryidae","Lepidocyrtinae","Lepidocyrtus","Lepidocyrtus paradoxus","21","19","19","7","9.52","0","99.44","448.52","0","449.95","42.19","0"
"AGAW2","05W","NIN","BOLD:ACJ7405","Arthropoda","Arachnida","Mesostigmata","Ameroseiidae","","Ameroseius","Ameroseius corbiculus","15","15","15","8","0","0","98.85","388.53","0","446.87","27.82","0"
"AGAW2","05W","NIN","BOLD:ACI9974","Arthropoda","Arachnida","Trombidiformes","Tydeidae","","","","3","3","3","9","0","0","97.56","424","0","450","40.3","0"
"AGAW2","05W","NIN","BOLD:ACZ1295","Arthropoda","Arachnida","Trombidiformes","Anystidae","","","","2","2","2","10","0","0","100","370","0","403","31","0"
"AGAW2","05W","NIN","BOLD:AAB5726","Arthropoda","Arachnida","Araneae","Agelenidae","","Agelenopsis","Agelenopsis pennsylvanica, Agelenopsis emertoni, Agelenopsis potteri, Agelenopsis actuosa, Agelenopsis kastoni","1","1","1","11","0","0","100","449","0","450","33.56","0"
"AGAW2","05W","NIN","BOLD:ACC0359","Arthropoda","Collembola","Symphypleona","Bourletiellidae","","Bourletiella","","1","1","1","11","0","0","99.78","448","0","450","37.56","0"
"AGAW2","08E","BAC","TAX:91393","Proteobacteria","Alphaproteobacteria","Rickettsiales","Rickettsiaceae",NA,"Wolbachia","","83","83","83","1","0","0","98.96","390.13","0","421.36","39.02","0"
"AGAW2","08E","BAC","TAX:175768","Proteobacteria","Alphaproteobacteria","Rickettsiales","Rickettsiaceae",NA,"Wolbachia","Wolbachia pipientis","54","49","49","2","9.26","0","98.19","446.98","0","448.65","39.47","0"
"AGAW2","08E","BAC","TAX:91391","Proteobacteria","Alphaproteobacteria","Rickettsiales","",NA,"","","1","1","1","3","0","0","98.02","405","0","405","39.51","0"
"AGAW2","08E","BAC","TAX:91389","Proteobacteria","","","",NA,"","","1","1","1","3","0","0","98.34","362","0","381","39.37","0"
"AGAW2","08E","CHO","no match","no match","no match","no match","no match","no match","no match","no match","no match","no match","no match","no match","no match","no match","no match","no match","no match","no match","no match","no match"
"AGAW2","08E","INS","BOLD:AAA3453","Arthropoda","Insecta","Diptera","Anthomyiidae","Anthomyiinae","Delia","","22932","18828","18828","1","17.9","0","99.37","432.16","0","434.64","34.63","0"
"AGAW2","08E","INS","BOLD:AAG1711","Arthropoda","Insecta","Diptera","Muscidae","Phaoniinae","Helina","Helina depuncta","14507","11175","11175","2","22.97","0","99.4","434.88","0","437.14","31.37","0"
"AGAW2","08E","INS","BOLD:AAB0373","Arthropoda","Insecta","Neuroptera","Chrysopidae","Chrysopinae","Chrysoperla","Chrysoperla sp. isolate 82, Chrysoperla zastrowi, Chrysoperla plorabunda, Chrysoperla sp. isolate 71, Chrysoperla agilis, Italochrysa italica, Chrysoperla sp. isolate 88, Chrysoperla lucasina, Chrysoperla carnea, Chrysoperla mediterranea, Chrysoperla pallida, Chrysoperla nipponensis, Chrysoperla sillemi","9872","7095","7095","3","28.13","0","99.42","436.52","0","437.44","31.45","0"
"AGAW2","08E","INS","BOLD:AAG2511","Arthropoda","Insecta","Diptera","Anthomyiidae","Anthomyiinae","Delia","Delia platura","5388","4519","4519","4","16.13","0","98.92","444.36","0","447.13","33.19","0"
"AGAW2","08E","INS","BOLD:AAG2123","Arthropoda","Insecta","Diptera","Tachinidae","Dexiinae","Dinera","Dinera grisescens","5167","3726","3726","5","27.89","0","99.34","434.71","0","436.38","31.87","0"
"AGAW2","08E","INS","BOLD:AAI4303","Arthropoda","Insecta","Diptera","Chironomidae","Chironominae","Chironomus","Chironomus melanescens","4656","4091","4091","6","12.13","0","98.33","430.43","0","435.71","33.77","0"
"AGAW2","08E","INS","BOLD:AAG6745","Arthropoda","Insecta","Diptera","Calliphoridae","Calliphorinae","Pollenia","Pollenia pediculata","2377","2055","2055","7","13.55","0","99.27","434.49","0","436.54","33.49","0"
"AGAW2","08E","INS","BOLD:AAC0903","Arthropoda","Insecta","Diptera","Chironomidae","Chironominae","Chironomus","Chironomus acidophilus","2320","1791","1791","8","22.8","0","99.46","438.12","0","440.8","36.26","0"
"AGAW2","08E","INS","BOLD:AAA3836","Arthropoda","Insecta","Lepidoptera","Noctuidae","Plusiinae","Autographa","Autographa precationis","1762","1433","1433","9","18.67","0","99.17","443.09","0","445.74","32.2","0"
"AGAW2","08E","INS","BOLD:AAP8739","Arthropoda","Insecta","Diptera","Platystomatidae","Platystomatinae","Rivellia","Rivellia steyskali","1604","1130","1130","10","29.55","0","99.28","440.99","0","443.31","37.93","0"
"AGAW2","08E","INS","BOLD:AAA4277","Arthropoda","Insecta","Diptera","Syrphidae","Syrphinae","Toxomerus","Toxomerus marginatus, Toxomerus sp. CR-3, Toxomerus CR-3","1500","1069","1069","11","28.73","0","99.34","443.86","0","444.87","30.44","0"
"AGAW2","08E","INS","BOLD:AAF9771","Arthropoda","Insecta","Diptera","Hybotidae","Tachydromiinae","Platypalpus","Platypalpus longiseta, Platypalpus trivialis, Platypalpus sp. 5, Platypalpus sp. 17, Platypalpus pallidiventris","1323","939","939","12","29.02","0","99.5","444.31","0","445.11","35.13","0"
"AGAW2","08E","INS","BOLD:ACR4394","Arthropoda","Insecta","Diptera","Anthomyiidae","Anthomyiinae","Delia","Delia florilega","1254","1168","1168","13","6.86","0","99.11","430.6","0","432.31","32.91","0"
"AGAW2","08E","INS","BOLD:ACU7906","Arthropoda","Insecta","Diptera","Chironomidae","Orthocladiinae","","","861","853","853","14","0.93","0","98.83","407.48","0","412.66","36.14","0"
"AGAW2","08E","INS","BOLD:AAD0484","Arthropoda","Insecta","Diptera","Chironomidae","","","","764","648","648","15","15.18","0","98.94","445.28","0","446.86","34.57","0"
"AGAW2","08E","INS","BOLD:AAD2041","Arthropoda","Insecta","Diptera","Dolichopodidae","Sciapodinae","Condylostylus","Condylostylus inermis, Condylostylus nigrofemoratus, Condylostylus connectans, Condylostylus caudatus, Condylostylus flavipes","679","507","507","16","25.33","0","99.5","439.7","0","440.87","35.92","0"
"AGAW2","08E","INS","BOLD:ABW7897","Arthropoda","Insecta","Diptera","Cecidomyiidae","","","","659","462","462","17","29.89","0","99.6","444.36","0","446.46","29.7","0"
"AGAW2","08E","INS","BOLD:AAA7661","Arthropoda","Insecta","Diptera","Culicidae","Culicinae","Culiseta, Culex","Culiseta melanura, Culex articularis, Culex restuans, Culex acharistus","609","542","542","18","11","0","99.14","434.92","0","436.89","33","0"
"AGAW2","08E","INS","BOLD:AAN5199","Arthropoda","Insecta","Diptera","Cecidomyiidae","","","CecidInt20 sp. BOLD:AAN5199","599","457","457","19","23.71","0","99.53","434.57","0","446.86","26.75","0"
"AGAW2","08E","INS","BOLD:AAA3889","Arthropoda","Insecta","Lepidoptera","Crambidae","Crambinae","Parapediasia","Parapediasia teterrellus","559","524","524","20","6.26","0","99.25","430.76","0","432.26","32.15","0"
"AGAW2","08E","INS","BOLD:ACX5521","Arthropoda","Insecta","Diptera","Chironomidae","Orthocladiinae","","","471","457","457","21","2.97","0","98.86","415.76","0","418.06","34.95","0"
"AGAW2","08E","INS","BOLD:AAG7279","Arthropoda","Insecta","Diptera","Sphaeroceridae","Limosininae","Spelobia","Spelobia ochripes","389","305","305","22","21.59","0","99.32","440.26","0","441.87","35.08","0"
"AGAW2","08E","INS","BOLD:AAC4197","Arthropoda","Insecta","Diptera","Chironomidae","Orthocladiinae","Paraphaenocladius","Paraphaenocladius impensus","336","277","277","23","17.56","0","99.46","435.93","0","439.25","35.09","0"
"AGAW2","08E","INS","BOLD:ACE0554","Arthropoda","Insecta","Diptera","Phoridae","Metopininae","Megaselia","","299","260","260","24","13.04","0","99.5","441.42","0","443.42","31.97","0"
"AGAW2","08E","INS","BOLD:AAA5299","Arthropoda","Insecta","Diptera","Chironomidae","Orthocladiinae","Cricotopus","Cricotopus laetus, Cricotopus relucens, Cricotopus trifasciatus, Cricotopus sp., Cricotopus sp. 3ES, Cricotopus sylvestris, Cricotopus glacialis","242","228","228","25","5.79","0","99.16","432.2","0","434.77","34.57","0"
"AGAW2","08E","INS","BOLD:AAC1312","Arthropoda","Insecta","Diptera","Syrphidae","Syrphinae","Toxomerus","Toxomerus geminatus, Toxomerus 75-5, Toxomerus rombicus, Toxomerus sp. 75-5","240","207","207","26","13.75","0","99.39","437.13","0","439.16","30.03","0"
"AGAW2","08E","INS","BOLD:AAA4781","Arthropoda","Insecta","Hymenoptera","Braconidae","Microgastrinae","Glyptapanteles","Glyptapanteles carlossarmientoi","190","179","179","27","5.79","0","99.26","428.98","0","432.04","24.14","0"
"AGAW2","08E","INS","BOLD:AAG8091","Arthropoda","Insecta","Hymenoptera","Ichneumonidae","Campopleginae","Campoletis","Campoletis sp. AB.2","175","170","170","28","2.86","0","99.32","434.85","0","437.22","24.69","0"
"AGAW2","08E","INS","BOLD:AAO7783","Arthropoda","Insecta","Diptera","Dolichopodidae","Diaphorinae","","","163","151","151","29","7.36","0","98.27","435.87","0","438.12","32.79","0"
"AGAW2","08E","INS","BOLD:ACK3431","Arthropoda","Insecta","Diptera","Cecidomyiidae","","","","160","146","146","30","8.75","0","99.09","431.17","0","433.19","32.68","0"
"AGAW2","08E","INS","BOLD:ABV0271","Arthropoda","Insecta","Diptera","Chloropidae","Oscinellinae","","","143","125","125","31","12.59","0","99.2","438.72","0","440.07","32.96","0"
"AGAW2","08E","INS","BOLD:AAH3661","Arthropoda","Insecta","Diptera","Cecidomyiidae","","","","142","132","132","32","7.04","0","99.25","431.47","0","434.06","25.36","0"
"AGAW2","08E","INS","BOLD:AAP5087","Arthropoda","Insecta","Diptera","Ceratopogonidae","Dasyheleinae","Dasyhelea","","132","102","102","33","22.73","0","99.16","439.08","0","440.48","34.65","0"
"AGAW2","08E","INS","BOLD:AAN5238","Arthropoda","Insecta","Diptera","Cecidomyiidae","","","","130","114","114","34","12.31","0","99.34","428.23","0","429.38","25.55","0"
"AGAW2","08E","INS","BOLD:AAD8820","Arthropoda","Insecta","Trichoptera","Hydroptilidae","Hydroptilinae","Hydroptila","Hydroptila jackmanni","119","115","115","35","3.36","0","98.7","368.31","0","414.03","33.25","0"
"AGAW2","08E","INS","BOLD:AAN5183","Arthropoda","Insecta","Diptera","Cecidomyiidae","","","","105","95","95","36","9.52","0","98.28","442.96","0","444.05","27.77","0"
"AGAW2","08E","INS","BOLD:AAM9013","Arthropoda","Insecta","Diptera","Keroplatidae","Macrocerinae","Macrocera","Macrocera pusilla","98","94","94","37","4.08","0","99.32","433.43","0","435.33","30.28","0"
"AGAW2","08E","INS","BOLD:AAL9079","Arthropoda","Insecta","Diptera","Phoridae","Metopininae","Megaselia","Megaselia giraudii, Megaselia aequalis","95","71","71","38","25.26","0","99.29","445.35","0","446.42","34.17","0"
"AGAW2","08E","INS","BOLD:AAD1485","Arthropoda","Insecta","Diptera","Chironomidae","Chironominae","Paratanytarsus","Paratanytarsus grimmii, Paratanytarsus sp.","90","81","81","39","10","0","99.43","441.51","0","442.71","33.78","0"
"AGAW2","08E","INS","BOLD:ACK7314","Arthropoda","Insecta","Diptera","Chironomidae","Chironominae","Polypedilum","","85","85","85","40","0","0","98.49","383.87","0","388.28","35.06","0"
"AGAW2","08E","INS","BOLD:ABA0863","Arthropoda","Insecta","Diptera","Cecidomyiidae","","","","81","62","62","41","23.46","0","99.41","444.11","0","445.32","29.8","0"
"AGAW2","08E","INS","BOLD:ACL8053","Arthropoda","Insecta","Diptera","Chironomidae","Chironominae","Tanytarsus","","76","69","69","42","9.21","0","99.32","445.07","0","446.37","33.67","0"
"AGAW2","08E","INS","BOLD:AAH0135","Arthropoda","Insecta","Coleoptera","Phalacridae","Phalacrinae","Acylomus","Acylomus pugetanus, Acylomus ergoti","66","62","62","43","6.06","0","98.99","432.29","0","434.33","31.6","0"
"AGAW2","08E","INS","BOLD:ABU5525","Arthropoda","Insecta","Diptera","Chironomidae","Orthocladiinae","Limnophyes","Limnophyes sp. 14ES","50","50","50","44","0","0","99.13","394.96","0","398.88","37.25","0"
"AGAW2","08E","INS","BOLD:AAN6154","Arthropoda","Insecta","Coleoptera","Latridiidae","Corticariinae","Melanophthalma","Melanophthalma inermis","42","35","35","45","16.67","0","98.82","435.74","0","437.02","31.47","0"
"AGAW2","08E","INS","BOLD:AAG4827","Arthropoda","Insecta","Diptera","Anthomyzidae","Anthomyzinae","Mumetopia","Mumetopia occipitalis","39","39","39","46","0","0","98.6","437.38","0","439.9","36.64","0"
"AGAW2","08E","INS","BOLD:AAG7891","Arthropoda","Insecta","Hymenoptera","Platygastridae","Scelioninae","Telenomus","Telenomus podisi","38","36","36","47","5.26","0","99.33","429.16","0","437.47","27.05","0"
"AGAW2","08E","INS","BOLD:ADF5285","Arthropoda","Insecta","Hymenoptera","Braconidae","Opiinae","Opius","","37","31","31","48","16.22","0","98.86","387.14","0","442.92","25.77","0"
"AGAW2","08E","INS","BOLD:AAL7359","Arthropoda","Insecta","Diptera","Chironomidae","Chironominae","Tanytarsus","","31","29","29","49","6.45","0","99.25","445.65","0","448.16","36.17","0"
"AGAW2","08E","INS","BOLD:ABV1194","Arthropoda","Insecta","Diptera","Chironomidae","Chironominae","Micropsectra","","31","26","26","49","16.13","0","99.55","438.1","0","439.39","36.15","0"
"AGAW2","08E","INS","BOLD:AAG2740","Arthropoda","Insecta","Diptera","Ephydridae","Parydrinae","Philygria","Philygria debilis","30","30","30","51","0","0","98.19","407","0","409.57","32.85","0"
"AGAW2","08E","INS","BOLD:AAN5267","Arthropoda","Insecta","Diptera","Cecidomyiidae","","","","28","21","21","52","25","0","99.59","448.29","0","449.18","30.55","0"
"AGAW2","08E","INS","BOLD:AAL4232","Arthropoda","Insecta","Diptera","Agromyzidae","Phytomyzinae","Phytoliriomyza","Phytoliriomyza arctica","25","23","23","53","8","0","99.41","431.16","0","432.32","31.93","0"
"AGAW2","08E","INS","BOLD:AAH0256","Arthropoda","Insecta","Coleoptera","Latridiidae","Corticariinae","Corticaria, Corticarina","","25","24","24","53","4","0","98.62","418.64","0","425.6","31.27","0"
"AGAW2","08E","INS","BOLD:AAN6433","Arthropoda","Insecta","Diptera","Sciaridae","","Corynoptera","Corynoptera deserta","20","20","20","55","0","0","99.19","445.75","0","448.25","30.52","0"
"AGAW2","08E","INS","BOLD:ABA1215","Arthropoda","Insecta","Diptera","Sciaridae","","Lycoriella","Lycoriella sativae","19","14","14","56","26.32","0","99.56","443.89","0","445.47","35.78","0"
"AGAW2","08E","INS","BOLD:AAM6184","Arthropoda","Insecta","Diptera","Ceratopogonidae","Ceratopogoninae","Culicoides","","16","16","16","57","0","0","98.02","407.25","0","409.56","36.53","0"
"AGAW2","08E","INS","BOLD:AAB9332","Arthropoda","Insecta","Hemiptera","Aphididae","Aphidinae","Nasonovia","Nasonovia ribisnigri","15","13","13","58","13.33","0","99.3","435.6","0","436.47","24.64","0"
"AGAW2","08E","INS","BOLD:AAG8687","Arthropoda","Insecta","Hemiptera","Cicadellidae","Typhlocybinae","Empoasca","","15","15","15","58","0","0","98.62","400.13","0","402","31.98","0"
"AGAW2","08E","INS","BOLD:AAA3181","Arthropoda","Insecta","Diptera","Muscidae","Muscinae","Stomoxys","Stomoxys calcitrans","14","13","13","60","7.14","0","98.9","435.29","0","439.36","32.97","0"
"AGAW2","08E","INS","BOLD:ACK5742","Arthropoda","Insecta","Diptera","Cecidomyiidae","","","","13","10","10","61","23.08","0","99.42","442.62","0","443.08","29.08","0"
"AGAW2","08E","INS","BOLD:AAG2868","Arthropoda","Insecta","Hemiptera","Cicadellidae","Typhlocybinae","Empoasca","Empoasca sp., Empoasca fabae","13","11","11","61","15.38","0","99.61","435.62","0","450","32.48","0"
"AGAW2","08E","INS","BOLD:ABX6186","Arthropoda","Insecta","Diptera","Tipulidae","Tipulinae","Nephrotoma","Nephrotoma ferruginea","13","13","13","61","0","0","99.3","434.15","0","435.77","34.25","0"
"AGAW2","08E","INS","BOLD:ACL7225","Arthropoda","Insecta","Diptera","Cecidomyiidae","","","","12","10","10","64","16.67","0","99.53","446.17","0","447.42","27.65","0"
"AGAW2","08E","INS","BOLD:AAH3664","Arthropoda","Insecta","Diptera","Cecidomyiidae","","","CecidInt3 sp. BOLD:AAH3664","12","12","12","64","0","0","99.27","443","0","443.92","27","0"
"AGAW2","08E","INS","BOLD:ADF6661","Arthropoda","Insecta","Diptera","Cecidomyiidae","","","","12","11","11","64","8.33","0","99.4","418.75","0","447.17","30.02","0"
"AGAW2","08E","INS","BOLD:AAJ1250","Arthropoda","Insecta","Hymenoptera","Braconidae","Aphidiinae","Aphidius","Aphidius avenaphis, Aphidius uzbekistanicus","11","11","11","67","0","0","99.15","430.09","0","435.09","26.77","0"
"AGAW2","08E","INS","BOLD:AAG6934","Arthropoda","Insecta","Diptera","Hybotidae","","","","11","11","11","67","0","0","99.34","427.27","0","430.36","38.81","0"
"AGAW2","08E","INS","BOLD:AAN5260","Arthropoda","Insecta","Diptera","Cecidomyiidae","","","","11","11","11","67","0","0","99.13","403.09","0","405","25.77","0"
"AGAW2","08E","INS","BOLD:ACK1360","Arthropoda","Insecta","Diptera","Cecidomyiidae","Cecidomyiinae","Vitisiella","Vitisiella brevicauda","10","10","10","70","0","0","99.51","449","0","450","28.22","0"
"AGAW2","08E","INS","BOLD:ACE9668","Arthropoda","Insecta","Hymenoptera","Ichneumonidae","Ophioninae","Enicospilus","Enicospilus purgatusDHJ02","9","9","9","71","0","0","99.28","444.44","0","445.89","25.7","0"
"AGAW2","08E","INS","BOLD:AAV1136","Arthropoda","Insecta","Diptera","Scatopsidae","","","","8","8","8","72","0","0","98.72","441","0","441.5","36.6","0"
"AGAW2","08E","INS","BOLD:ADH9867","Arthropoda","Insecta","Diptera","","","","","8","8","8","72","0","0","98.55","397.88","0","401.88","37.32","0"
"AGAW2","08E","INS","BOLD:AAN5230","Arthropoda","Insecta","Diptera","Cecidomyiidae","","","","7","7","7","74","0","0","98.85","425.29","0","429","25.8","0"
"AGAW2","08E","INS","BOLD:ABA7871","Arthropoda","Insecta","Diptera","Cecidomyiidae","","","","7","7","7","74","0","0","98.72","442.43","0","442.86","26.76","0"
"AGAW2","08E","INS","BOLD:AAH3662","Arthropoda","Insecta","Diptera","Cecidomyiidae","","","CecidInt3 sp. BOLD:AAH3662","6","6","6","76","0","0","99.11","441.33","0","442.83","26.18","0"
"AGAW2","08E","INS","BOLD:ACX6289","Arthropoda","Insecta","Diptera","Phoridae","Metopininae","Megaselia","","6","6","6","76","0","0","99.55","376","0","401.67","37.2","0"
"AGAW2","08E","INS","BOLD:AAA2632","Arthropoda","Insecta","Lepidoptera","Noctuidae","Noctuinae","Noctua","Noctua atlantica, Noctua pronuba","4","4","4","78","0","0","99.22","448","0","450","31.28","0"
"AGAW2","08E","INS","BOLD:ABA0929","Arthropoda","Insecta","Diptera","Sciaridae","","Bradysia","Bradysia scabricornis","4","4","4","78","0","0","98.71","443.25","0","444.5","30.26","0"
"AGAW2","08E","INS","BOLD:ABA6407","Arthropoda","Insecta","Diptera","Sciaridae","","Pseudolycoriella","","4","4","4","78","0","0","99.31","434.25","0","436.25","34.71","0"
"AGAW2","08E","INS","BOLD:ACL6050","Arthropoda","Insecta","Trichoptera","Hydroptilidae","Orthotrichiinae","Orthotrichia","Orthotrichia cristata","4","4","4","78","0","0","98.86","384.5","0","387.25","29.1","0"
"AGAW2","08E","INS","BOLD:AAU8379","Arthropoda","Insecta","Hymenoptera","Braconidae","Alysiinae","Dinotrema","Dinotrema JFT01","4","4","4","78","0","0","100","432.25","0","433.5","24.2","0"
"AGAW2","08E","INS","BOLD:AAH4135","Arthropoda","Insecta","Diptera","Chloropidae","Chloropinae","Thaumatomyia","Thaumatomyia glabra, Thaumatomyia grata, Thaumatomyia sp. 1 TAW","3","3","3","83","0","0","99.84","427.33","0","434.33","31.76","0"
"AGAW2","08E","INS","BOLD:AAY6412","Arthropoda","Insecta","Diptera","Sciaridae","","Bradysia","Bradysia urticae","3","3","3","83","0","0","98.53","434","0","436.67","31.88","0"
"AGAW2","08E","INS","BOLD:AAM7855","Arthropoda","Insecta","Thysanoptera","Thripidae","Thripinae","Frankliniella","Frankliniella tritici","3","3","3","83","0","0","99.4","445.67","0","450","31.26","0"
"AGAW2","08E","INS","BOLD:ACE7444","Arthropoda","Insecta","Hemiptera","Miridae","Mirinae, Phylinae","Adelphocoris, Megalocoleus","Adelphocoris lineolatus, Megalocoleus molliculus","3","3","3","83","0","0","99.03","447.33","0","450","35.33","0"
"AGAW2","08E","INS","BOLD:AAH4171","Arthropoda","Insecta","Diptera","Chloropidae","Oscinellinae","Apallates","Apallates montanus, Apallates particeps","3","3","3","83","0","0","98.83","427","0","430","32.6","0"
"AGAW2","08E","INS","BOLD:AAN5219","Arthropoda","Insecta","Diptera","Cecidomyiidae","","","","3","3","3","83","0","0","98.94","445.33","0","447","29.61","0"
"AGAW2","08E","INS","BOLD:ACC8079","Arthropoda","Insecta","Diptera","Cecidomyiidae","","","","3","3","3","83","0","0","99.48","448.33","0","450","29.26","0"
"AGAW2","08E","INS","BOLD:AAN5220","Arthropoda","Insecta","Diptera","Cecidomyiidae","","","","3","3","3","83","0","0","97.97","392.67","0","414.67","26.17","0"
"AGAW2","08E","INS","BOLD:ACC0911","Arthropoda","Insecta","Hymenoptera","Braconidae","Aphidiinae","Praon","","3","3","3","83","0","0","99.48","449.67","0","450","26.3","0"
"AGAW2","08E","INS","BOLD:AAG2898","Arthropoda","Insecta","Hemiptera","Cicadellidae","Deltocephalinae","Paraphlepsius","Paraphlepsius apertus, Paraphlepsius irroratus","3","3","3","83","0","0","98.22","430.33","0","432.33","32.11","0"
"AGAW2","08E","INS","BOLD:AAE0845","Arthropoda","Insecta","Hemiptera","Pentatomidae","Pentatominae","Euschistus","Euschistus servus","3","3","3","83","0","0","98.49","443","0","445","37.91","0"
"AGAW2","08E","INS","BOLD:AAA5803","Arthropoda","Insecta","Hemiptera","Miridae","Mirinae","Lygus","Lygus lineolaris","3","3","3","83","0","0","98.66","448.67","0","450","33.85","0"
"AGAW2","08E","INS","BOLD:AAG2460","Arthropoda","Insecta","Diptera","Anthomyiidae","Anthomyiinae","Hydrophoria","Hydrophoria lancifer","3","3","3","83","0","0","99.1","431","0","432.67","33.56","0"
"AGAW2","08E","INS","BOLD:AAN5161","Arthropoda","Insecta","Diptera","Ceratopogonidae","Dasyheleinae","Dasyhelea","","2","2","2","96","0","0","98.08","418","0","420","37.34","0"
"AGAW2","08E","INS","BOLD:AAH2306","Arthropoda","Insecta","Diptera","Asilidae","Asilinae","Efferia","Efferia aestuans","2","2","2","96","0","0","99.11","448","0","450","38.33","0"
"AGAW2","08E","INS","BOLD:AAF9949","Arthropoda","Insecta","Hemiptera","Miridae","Mirinae","Trigonotylus","Trigonotylus pulchellus, Trigonotylus americanus, Trigonotylus psammaecolor, Trigonotylus caelestialium, Trigonotylus viridis, Trigonotylus ruficornis","2","2","2","96","0","0","99.32","445.5","0","447","35.02","0"
"AGAW2","08E","INS","BOLD:AAV5594","Arthropoda","Insecta","Diptera","Cecidomyiidae","","","","2","2","2","96","0","0","99.78","449","0","450","26.11","0"
"AGAW2","08E","INS","BOLD:ACJ0111","Arthropoda","Insecta","Hymenoptera","Platygastridae","Scelioninae","Telenomus","","2","2","2","96","0","0","99","402","0","408","27.97","0"
"AGAW2","08E","INS","BOLD:ACV5892","Arthropoda","Insecta","Diptera","Cecidomyiidae","","","","2","2","2","96","0","0","98","449","0","450","25.44","0"
"AGAW2","08E","INS","BOLD:AAN5342","Arthropoda","Insecta","Diptera","Chironomidae","Orthocladiinae","Heleniella","","2","2","2","96","0","0","98.44","389.5","0","402.5","34.53","0"
"AGAW2","08E","INS","BOLD:AAS1281","Arthropoda","Insecta","Diptera","Chironomidae","Chironominae","Chironomus","Chironomus nr. sollicitus","2","2","2","96","0","0","97.1","413.5","0","415.5","32.85","0"
"AGAW2","08E","INS","BOLD:AAB2384","Arthropoda","Insecta","Diptera","Syrphidae","Syrphinae","Eupeodes","Eupeodes duseki, Eupeodes volucris, Eupeodes nielseni, Eupeodes americanus, Eupeodes flukei, Eupeodes vockerothi, Eupeodes lucasi, Eupeodes montivagus, Eupeodes rufipunctatus, Eupeodes rojasi, Eupeodes goeldlini, Eupeodes luniger, Eupeodes punctifer, Eupeodes bucculatus, Eupeodes perplexus, Eupeodes abiskoensis, Eupeodes corollae, Eupeodes pomus, Eupeodes lundbecki, Eupeodes nitens, Eupeodes sp. BOLDAAB2384, Eupeodes curtus, Eupeodes latifasciatus","2","2","2","96","0","0","97.87","418","0","420","31.32","0"
"AGAW2","08E","INS","BOLD:AAN8010","Arthropoda","Insecta","Hymenoptera","Platygastridae","Scelioninae","Telenomus","","2","2","2","96","0","0","98.86","440","0","450","28.33","0"
"AGAW2","08E","INS","BOLD:AAH4205","Arthropoda","Insecta","Diptera","Chloropidae","Chloropinae","Meromyza","Meromyza nigriseta, Meromyza sp.","2","2","2","96","0","0","99.33","449.5","0","450","33","0"
"AGAW2","08E","INS","BOLD:ABY1179","Arthropoda","Insecta","Hymenoptera","Sierolomorphidae","","Sierolomorpha","","2","2","2","96","0","0","99.89","449","0","450","28.11","0"
"AGAW2","08E","INS","BOLD:ACI3514","Arthropoda","Insecta","Diptera","Chironomidae","Chironominae","Tanytarsus","","1","1","1","108","0","0","99.32","443","0","450","36.89","0"
"AGAW2","08E","INS","BOLD:AAA6292","Arthropoda","Insecta","Lepidoptera","Tortricidae","Tortricinae","Cnephasia","Cnephasia genitalana, Cnephasia stephensiana","1","1","1","108","0","0","100","450","0","450","35.78","0"
"AGAW2","08E","INS","BOLD:AAG1714","Arthropoda","Insecta","Diptera","Muscidae","Azeliinae","Muscina","Muscina pascuorum","1","1","1","108","0","0","98.38","431","0","450","33.56","0"
"AGAW2","08E","INS","BOLD:AAG5507","Arthropoda","Insecta","Diptera","Chironomidae","Chironominae","Chironomus","Chironomus atrella","1","1","1","108","0","0","99.33","450","0","450","36.89","0"
"AGAW2","08E","INS","BOLD:AAA6560","Arthropoda","Insecta","Lepidoptera","Crambidae","Crambinae","Crambus","Crambus multilinellus, Crambus saltuellus","1","1","1","108","0","0","99.78","450","0","450","31.11","0"
"AGAW2","08E","INS","BOLD:AAF9890","Arthropoda","Insecta","Orthoptera","Gryllidae","Oecanthinae","Oecanthus","Oecanthus niveus","1","1","1","108","0","0","100","450","0","450","36","0"
"AGAW2","08E","INS","BOLD:AAG9668","Arthropoda","Insecta","Diptera","Dolichopodidae","Diaphorinae","Chrysotus","Chrysotus sp., Chrysotus sp. nr. obliquus","1","1","1","108","0","0","99.56","450","0","450","33.11","0"
"AGAW2","08E","INS","BOLD:ACF4786","Arthropoda","Insecta","Hymenoptera","Braconidae","Aphidiinae","Aphidius","Aphidius urticae, Aphidius matricariae","1","1","1","108","0","0","97.34","414","0","420","24.52","0"
"AGAW2","08E","INS","BOLD:ACA9275","Arthropoda","Insecta","Diptera","Chironomidae","Chironominae","Chironomus","","1","1","1","108","0","0","99.56","450","0","450","35.33","0"
"AGAW2","08E","INS","BOLD:AAG5453","Arthropoda","Insecta","Diptera","Chironomidae","Chironominae","Chironomus","Chironomus bifurcatus","1","1","1","108","0","0","97.54","447","0","450","35.11","0"
"AGAW2","08E","INS","BOLD:ACV2677","Arthropoda","Insecta","Diptera","Cecidomyiidae","","","","1","1","1","108","0","0","97.56","450","0","450","30.22","0"
"AGAW2","08E","INS","BOLD:AAH3914","Arthropoda","Insecta","Diptera","Sciaridae","","Sciara","Sciara hamatilis, Sciara humeralis","1","1","1","108","0","0","97.42","426","0","430","30.93","0"
"AGAW2","08E","INS","BOLD:AAB9404","Arthropoda","Insecta","Lepidoptera","Tortricidae","Tortricinae","Archips","Archips rosana, Archips grisea","1","1","1","108","0","0","100","448","0","450","31.33","0"
"AGAW2","08E","INS","BOLD:AAV5853","Arthropoda","Insecta","Diptera","Chironomidae","Tanypodinae","","","1","1","1","108","0","0","98.96","385","0","388","34.79","0"
"AGAW2","08E","INS","BOLD:AAD7604","Arthropoda","Insecta","Coleoptera","Coccinellidae","Coccinellinae","Coleomegilla","Coleomegilla maculata","1","1","1","108","0","0","97.19","427","0","428","32.01","0"
"AGAW2","08E","INS","BOLD:ABY4763","Arthropoda","Insecta","Hemiptera","Aphididae","Aphidinae","Uroleucon","Uroleucon taraxaci","1","1","1","108","0","0","100","450","0","450","25.11","0"
"AGAW2","08E","INS","BOLD:AAH0237","Arthropoda","Insecta","Coleoptera","Cleridae","Clerinae","Enoclerus","Enoclerus nigripes, Enoclerus rosmarus","1","1","1","108","0","0","97.56","450","0","450","32","0"
"AGAW2","08E","INS","BOLD:ACU6062","Arthropoda","Insecta","Diptera","Cecidomyiidae","","","","1","1","1","108","0","0","99.11","450","0","450","26.89","0"
"AGAW2","08E","INS","BOLD:AAG9897","Arthropoda","Insecta","Dermaptera","Forficulidae","Forficulinae","Forficula","Forficula auricularia-A","1","1","1","108","0","0","99.76","414","0","417","45.32","0"
"AGAW2","08E","INS","BOLD:AAB9652","Arthropoda","Insecta","Trichoptera","Leptoceridae","Leptocerinae","Triaenodes","Triaenodes nox","1","1","1","108","0","0","98.11","424","0","428","31.31","0"
"AGAW2","08E","INS","BOLD:AAH2293","Arthropoda","Insecta","Diptera","Asilidae","Stenopogoninae","Dioctria, Eudioctria","Dioctria hyalipennis, Dioctria baumhaueri, Eudioctria propinqua","1","1","1","108","0","0","98.01","403","0","401","29.93","0"
"AGAW2","08E","INS","BOLD:AAA2815","Arthropoda","Insecta","Lepidoptera","Noctuidae","Noctuinae","Mythimna","Mythimna yukonensis, Mythimna oxygala","1","1","1","108","0","0","98.66","448","0","450","34","0"
"AGAW2","08E","INS","BOLD:AAD4936","Arthropoda","Insecta","Lepidoptera","Gracillariidae","Gracillariinae","Caloptilia","Caloptilia rhoifoliella, Caloptilia sassafrasella","1","1","1","108","0","0","99.51","411","0","411","28.47","0"
"AGAW2","08E","INS","BOLD:AAF9187","Arthropoda","Insecta","Coleoptera","Curculionidae","Entiminae","Phyllobius","Phyllobius oblongus","1","1","1","108","0","0","97.66","427","0","431","39.21","0"
"AGAW2","08E","INS","BOLD:AAG9626","Arthropoda","Insecta","Diptera","Dolichopodidae","","","","1","1","1","108","0","0","98.66","448","0","450","36","0"
"AGAW2","08E","INS","BOLD:ACS4193","Arthropoda","Insecta","Hemiptera","Cicadellidae","Typhlocybinae","Empoasca","","1","1","1","108","0","0","97.54","406","0","408","31.62","0"
"AGAW2","08E","INS","BOLD:AAA3867","Arthropoda","Insecta","Lepidoptera","Noctuidae","Noctuinae","Apamea","Apamea amputatrix","1","1","1","108","0","0","99.78","450","0","450","33.78","0"
"AGAW2","08E","INS","BOLD:ACF7023","Arthropoda","Insecta","Diptera","Dolichopodidae","","","","1","1","1","108","0","0","98.24","227","0","450","33.11","0"
"AGAW2","08E","INS","BOLD:AAG1739","Arthropoda","Insecta","Diptera","Muscidae","Coenosiinae","Neodexiopsis","Neodexiopsis rufitibia","1","1","1","108","0","0","100","447","0","450","34","0"
"AGAW2","08E","INS","BOLD:AAA3654","Arthropoda","Insecta","Lepidoptera","Crambidae","Crambinae","Crambus","Crambus bigelovi, Crambus johnsoni, Crambus praefectellus","1","1","1","108","0","0","98.65","443","0","450","31.56","0"
"AGAW2","08E","INS","BOLD:AAH7142","Arthropoda","Insecta","Diptera","Sarcophagidae","Sarcophaginae","Oxysarcodexia, Sarcophaginae_incertae_sedis","Oxysarcodexia cingarus, Sarcophaginae_incertae_sedis sp. 1 AGT, Oxysarcodexia ventricosa","1","1","1","108","0","0","98.89","450","0","450","34.67","0"
"AGAW2","08E","INS","BOLD:AAH7146","Arthropoda","Insecta","Diptera","Sarcophagidae","Sarcophaginae","Boettcheria","Boettcheria latisterna","1","1","1","108","0","0","100","437","0","438","35.16","0"
"AGAW2","08E","INS","BOLD:ABX6298","Arthropoda","Insecta","Diptera","Tipulidae","Tipulinae","Nephrotoma","Nephrotoma ferruginea","1","1","1","108","0","0","97.77","449","0","450","33.33","0"
"AGAW2","08E","INS","BOLD:ADF3937","","","","","","","","1","1","1","108","0","0","98.44","384","0","386","31.35","0"
"AGAW2","08E","INS","BOLD:AAD8086","Arthropoda","Insecta","Lepidoptera","Elachistidae","Elachistinae","Elachista","Elachista griseicornis, Elachista acenteta","1","1","1","108","0","0","99.2","376","0","382","30.37","0"
"AGAW2","08E","INS","BOLD:AAC0706","Arthropoda","Insecta","Diptera","Chironomidae","Chironominae","Dicrotendipes","Dicrotendipes tritomus","1","1","1","108","0","0","97.32","410","0","411","35.04","0"
"AGAW2","08E","INS","BOLD:AAA9422","Arthropoda","Insecta","Hemiptera","Cicadellidae","Deltocephalinae","Macrosteles","Macrosteles quadrilineatus","1","1","1","108","0","0","98.56","418","0","444","36.94","0"
"AGAW2","08E","INS","BOLD:AAC0635","Arthropoda","Insecta","Hemiptera","Miridae","Mirinae","Stenotus","Stenotus binotatus","1","1","1","108","0","0","98.29","410","0","413","33.41","0"
"AGAW2","08E","NAR","no match","no match","no match","no match","no match","no match","no match","no match","no match","no match","no match","no match","no match","no match","no match","no match","no match","no match","no match","no match"
"AGAW2","08E","NIN","BOLD:AAH6630","Arthropoda","Collembola","Entomobryomorpha","Entomobryidae","Entomobryinae","Entomobrya","Entomobrya unostrigata","218","192","192","1","11.93","0","99.41","440.34","0","441.53","38.94","0"
"AGAW2","08E","NIN","BOLD:AAI4346","Arthropoda","Arachnida","Opiliones","Phalangiidae","","Paroligolophus, Phalangium","Paroligolophus agrestis, Phalangium opilio","208","203","203","2","2.4","0","99.09","408.79","0","417.94","35.21","0"
"AGAW2","08E","NIN","BOLD:AAA6383","","","","","","","","98","92","92","3","6.12","0","98.35","434.47","0","435.71","37.76","0"
"AGAW2","08E","NIN","BOLD:ABW2639","Arthropoda","Arachnida","Trombidiformes","Anystidae","","","","68","59","59","4","13.24","0","99.45","449.49","0","447.4","31.45","0"
"AGAW2","08E","NIN","BOLD:AAA6381","Arthropoda","Arachnida","Araneae","Tetragnathidae","","Tetragnatha","Tetragnatha laboriosa","35","35","35","5","0","0","98.74","419.51","0","422","39.27","0"
"AGAW2","08E","NIN","BOLD:AAB7914","Arthropoda","Collembola","Symphypleona","Bourletiellidae","","Bourletiella","","20","20","20","6","0","0","99.37","442.85","0","444.4","38.09","0"
"AGAW2","08E","NIN","BOLD:ACZ1295","Arthropoda","Arachnida","Trombidiformes","Anystidae","","","","12","12","12","7","0","0","99.24","376.75","0","397.17","31.28","0"
"AGAW2","08E","NIN","BOLD:AAA5267","Arthropoda","Collembola","Entomobryomorpha","Isotomidae","","","","6","6","6","8","0","0","99.35","418.67","0","421.33","37.77","0"
"AGAW2","08E","NIN","BOLD:AAB3836","Arthropoda","Arachnida","Araneae","Philodromidae","","Philodromus","Philodromus cespitum","4","4","4","9","0","0","98.17","385.25","0","413.25","30.48","0"
"AGAW2","08E","NIN","BOLD:AAH8314","Arthropoda","Arachnida","Araneae","Linyphiidae","","Walckenaeria","Walckenaeria microspiralis","2","2","2","10","0","0","98.2","445","0","450","36.78","0"
"AGAW2","08E","NIN","BOLD:AAA7242","Arthropoda","Collembola","Entomobryomorpha","Entomobryidae","Seirinae, Entomobryinae","Entomobrya, Lepidosira","Entomobrya multifasciata, Lepidosira sp.","2","2","2","10","0","0","99.11","402.5","0","406","39.06","0"
"AGAW2","08E","NIN","BOLD:ACR6860","","","","","","","","1","1","1","12","0","0","98.97","387","0","415","38.8","0"
"AGAW2","08E","NIN","BOLD:AAB5726","Arthropoda","Arachnida","Araneae","Agelenidae","","Agelenopsis","Agelenopsis pennsylvanica, Agelenopsis emertoni, Agelenopsis potteri, Agelenopsis actuosa, Agelenopsis kastoni","1","1","1","12","0","0","99.07","430","0","433","33.03","0"
"AGAW2","08E","NIN","BOLD:AAG5658","Arthropoda","Arachnida","Araneae","Mimetidae","","Mimetus","Mimetus epeiroides","1","1","1","12","0","0","99.33","447","0","450","34.22","0"
"AGAW2","08N","BAC","TAX:175768","Proteobacteria","Alphaproteobacteria","Rickettsiales","Rickettsiaceae",NA,"Wolbachia","Wolbachia pipientis","47","41","41","1","12.77","0","98.18","447.6","0","449.49","39.52","0"
"AGAW2","08N","BAC","TAX:91393","Proteobacteria","Alphaproteobacteria","Rickettsiales","Rickettsiaceae",NA,"Wolbachia","","43","43","43","2","0","0","99.02","388.93","0","419.91","39.01","0"
"AGAW2","08N","BAC","TAX:91391","Proteobacteria","Alphaproteobacteria","Rickettsiales","",NA,"","","9","9","9","3","0","0","98","421.67","0","450","39.98","0"
"AGAW2","08N","BAC","TAX:703648","Proteobacteria","Alphaproteobacteria","Rickettsiales","Rickettsiaceae",NA,"Wolbachia","Wolbachia endosymbiont","3","3","3","4","0","0","97.55","395","0","450","39.63","0"
"AGAW2","08N","BAC","TAX:91389","Proteobacteria","","","",NA,"","","2","2","2","5","0","0","98.19","359","0","363.5","38.92","0"
"AGAW2","08N","BAC","TAX:375762","Proteobacteria","Alphaproteobacteria","Rickettsiales","Rickettsiaceae",NA,"Wolbachia","Wolbachia sp. Myrmicinae_ACGAN556-09","2","2","2","5","0","0","98.33","449","0","450","39.89","0"
"AGAW2","08N","BAC","TAX:376688","Proteobacteria","Alphaproteobacteria","Rickettsiales","Rickettsiaceae",NA,"Wolbachia","Wolbachia sp. elachBioLep01 BioLep373","1","1","1","7","0","0","98.44","450","0","450","39.78","0"
"AGAW2","08N","CHO","no match","no match","no match","no match","no match","no match","no match","no match","no match","no match","no match","no match","no match","no match","no match","no match","no match","no match","no match","no match"
"AGAW2","08N","INS","BOLD:AAG1711","Arthropoda","Insecta","Diptera","Muscidae","Phaoniinae","Helina","Helina depuncta","15425","12165","12165","1","21.13","0","99.38","433.4","0","435.73","31.35","0"
"AGAW2","08N","INS","BOLD:AAA4277","Arthropoda","Insecta","Diptera","Syrphidae","Syrphinae","Toxomerus","Toxomerus marginatus, Toxomerus sp. CR-3, Toxomerus CR-3","13589","8487","8487","2","37.55","0","99.35","445.28","0","446.17","30.65","0"
"AGAW2","08N","INS","BOLD:AAA3181","Arthropoda","Insecta","Diptera","Muscidae","Muscinae","Stomoxys","Stomoxys calcitrans","9865","7440","7440","3","24.58","0","98.84","433.3","0","437.01","33.1","0"
"AGAW2","08N","INS","BOLD:ABX6186","Arthropoda","Insecta","Diptera","Tipulidae","Tipulinae","Nephrotoma","Nephrotoma ferruginea","9524","6620","6620","4","30.49","0","99.13","443.98","0","445.57","34.31","0"
"AGAW2","08N","INS","BOLD:AAB0373","Arthropoda","Insecta","Neuroptera","Chrysopidae","Chrysopinae","Chrysoperla","Chrysoperla sp. isolate 82, Chrysoperla zastrowi, Chrysoperla plorabunda, Chrysoperla sp. isolate 71, Chrysoperla agilis, Italochrysa italica, Chrysoperla sp. isolate 88, Chrysoperla lucasina, Chrysoperla carnea, Chrysoperla mediterranea, Chrysoperla pallida, Chrysoperla nipponensis, Chrysoperla sillemi","9412","6521","6521","5","30.72","0","99.08","436.59","0","437.52","31.34","0"
"AGAW2","08N","INS","BOLD:AAA3453","Arthropoda","Insecta","Diptera","Anthomyiidae","Anthomyiinae","Delia","","7522","6520","6520","6","13.32","0","99.37","431.39","0","433.99","34.64","0"
"AGAW2","08N","INS","BOLD:AAG6745","Arthropoda","Insecta","Diptera","Calliphoridae","Calliphorinae","Pollenia","Pollenia pediculata","6140","5396","5396","7","12.12","0","99.32","427.16","0","429.31","33.48","0"
"AGAW2","08N","INS","BOLD:AAA5803","Arthropoda","Insecta","Hemiptera","Miridae","Mirinae","Lygus","Lygus lineolaris","2432","1576","1576","8","35.2","0","98.66","446.71","0","448.39","33.9","0"
"AGAW2","08N","INS","BOLD:AAH4135","Arthropoda","Insecta","Diptera","Chloropidae","Chloropinae","Thaumatomyia","Thaumatomyia glabra, Thaumatomyia grata, Thaumatomyia sp. 1 TAW","2220","1962","1962","9","11.62","0","99.42","430.6","0","435.45","31.83","0"
"AGAW2","08N","INS","BOLD:AAH4171","Arthropoda","Insecta","Diptera","Chloropidae","Oscinellinae","Apallates","Apallates montanus, Apallates particeps","1540","1251","1251","10","18.77","0","99.02","436.8","0","438.58","32.72","0"
"AGAW2","08N","INS","BOLD:ADH4973","Arthropoda","Insecta","Diptera","Tipulidae","Tipulinae","Nephrotoma","","1371","1238","1238","11","9.7","0","98.3","422.53","0","424.55","34.06","0"
"AGAW2","08N","INS","BOLD:AAN5219","Arthropoda","Insecta","Diptera","Cecidomyiidae","","","","1156","892","892","12","22.84","0","99.5","440.06","0","441.31","29.03","0"
"AGAW2","08N","INS","BOLD:ACU7906","Arthropoda","Insecta","Diptera","Chironomidae","Orthocladiinae","","","1070","1066","1066","13","0.37","0","98.89","403.5","0","411.19","36.18","0"
"AGAW2","08N","INS","BOLD:AAN6154","Arthropoda","Insecta","Coleoptera","Latridiidae","Corticariinae","Melanophthalma","Melanophthalma inermis","1049","840","840","14","19.92","0","98.52","437.74","0","439.98","31.36","0"
"AGAW2","08N","INS","BOLD:AAF9771","Arthropoda","Insecta","Diptera","Hybotidae","Tachydromiinae","Platypalpus","Platypalpus longiseta, Platypalpus trivialis, Platypalpus sp. 5, Platypalpus sp. 17, Platypalpus pallidiventris","996","723","723","15","27.41","0","99.43","444.04","0","444.96","35.14","0"
"AGAW2","08N","INS","BOLD:AAP8739","Arthropoda","Insecta","Diptera","Platystomatidae","Platystomatinae","Rivellia","Rivellia steyskali","848","655","655","16","22.76","0","99.4","441.36","0","443.23","37.79","0"
"AGAW2","08N","INS","BOLD:AAA5299","Arthropoda","Insecta","Diptera","Chironomidae","Orthocladiinae","Cricotopus","Cricotopus laetus, Cricotopus relucens, Cricotopus trifasciatus, Cricotopus sp., Cricotopus sp. 3ES, Cricotopus sylvestris, Cricotopus glacialis","846","728","728","17","13.95","0","99.4","433.63","0","436.4","34.56","0"
"AGAW2","08N","INS","BOLD:AAF9949","Arthropoda","Insecta","Hemiptera","Miridae","Mirinae","Trigonotylus","Trigonotylus pulchellus, Trigonotylus americanus, Trigonotylus psammaecolor, Trigonotylus caelestialium, Trigonotylus viridis, Trigonotylus ruficornis","720","511","511","18","29.03","0","99.52","444.63","0","445.48","34.58","0"
"AGAW2","08N","INS","BOLD:AAN5199","Arthropoda","Insecta","Diptera","Cecidomyiidae","","","CecidInt20 sp. BOLD:AAN5199","680","519","519","19","23.68","0","99.54","435.07","0","446.51","26.76","0"
"AGAW2","08N","INS","BOLD:ACZ1491","Arthropoda","Insecta","Diptera","Chironomidae","","","","607","412","412","20","32.13","0","97.86","446.91","0","448.39","34.35","0"
"AGAW2","08N","INS","BOLD:AAG2511","Arthropoda","Insecta","Diptera","Anthomyiidae","Anthomyiinae","Delia","Delia platura","589","537","537","21","8.83","0","98.94","444.43","0","447.15","33.15","0"
"AGAW2","08N","INS","BOLD:AAY3269","Arthropoda","Insecta","Diptera","Chironomidae","Diamesinae","Potthastia","Potthastia gaedii","383","322","322","22","15.93","0","98.1","438.33","0","439.73","34.5","0"
"AGAW2","08N","INS","BOLD:AAG2740","Arthropoda","Insecta","Diptera","Ephydridae","Parydrinae","Philygria","Philygria debilis","361","359","359","23","0.55","0","98.3","408.33","0","411.75","32.85","0"
"AGAW2","08N","INS","BOLD:AAA7661","Arthropoda","Insecta","Diptera","Culicidae","Culicinae","Culiseta, Culex","Culiseta melanura, Culex articularis, Culex restuans, Culex acharistus","309","295","295","24","4.53","0","99.15","436.17","0","438.23","32.67","0"
"AGAW2","08N","INS","BOLD:AAN5161","Arthropoda","Insecta","Diptera","Ceratopogonidae","Dasyheleinae","Dasyhelea","","306","300","300","25","1.96","0","98.92","398.68","0","400.93","37.29","0"
"AGAW2","08N","INS","BOLD:AAN5183","Arthropoda","Insecta","Diptera","Cecidomyiidae","","","","219","183","183","26","16.44","0","98.46","443.81","0","444.98","27.85","0"
"AGAW2","08N","INS","BOLD:AAH0134","Arthropoda","Insecta","Coleoptera","Phalacridae","Phalacrinae","Stilbus","Stilbus apicialis","199","177","177","27","11.06","0","99.17","429.32","0","430.51","32.03","0"
"AGAW2","08N","INS","BOLD:AAL4232","Arthropoda","Insecta","Diptera","Agromyzidae","Phytomyzinae","Phytoliriomyza","Phytoliriomyza arctica","189","166","166","28","12.17","0","99.08","443.44","0","444.92","32.02","0"
"AGAW2","08N","INS","BOLD:ACX6055","Arthropoda","Insecta","Diptera","Phoridae","Metopininae","Megaselia","Megaselia sp. AK-2010-2, Megaselia sp. BOLD:ACX6055","188","156","156","29","17.02","0","99.45","438.98","0","440.98","33.13","0"
"AGAW2","08N","INS","BOLD:AAC4197","Arthropoda","Insecta","Diptera","Chironomidae","Orthocladiinae","Paraphaenocladius","Paraphaenocladius impensus","170","134","134","30","21.18","0","99.56","437.42","0","440.84","35.14","0"
"AGAW2","08N","INS","BOLD:AAH3661","Arthropoda","Insecta","Diptera","Cecidomyiidae","","","","152","144","144","31","5.26","0","99.31","428.14","0","429.6","25.32","0"
"AGAW2","08N","INS","BOLD:ABZ0908","Arthropoda","Insecta","Diptera","Tipulidae","Tipulinae","Nephrotoma","","152","152","152","31","0","0","97.69","378.14","0","381.2","33.99","0"
"AGAW2","08N","INS","BOLD:ABW7897","Arthropoda","Insecta","Diptera","Cecidomyiidae","","","","98","79","79","33","19.39","0","99.53","442.35","0","445.79","29.67","0"
"AGAW2","08N","INS","BOLD:AAU8441","Arthropoda","Insecta","Hymenoptera","Ichneumonidae","Campopleginae","Meloboris","","92","76","76","34","17.39","0","99.34","448.64","0","449.73","25.41","0"
"AGAW2","08N","INS","BOLD:AAL7352","Arthropoda","Insecta","Diptera","Chironomidae","Chironominae","Polypedilum","","69","69","69","35","0","0","98.7","420.48","0","423.43","33.99","0"
"AGAW2","08N","INS","BOLD:ACI1304","Arthropoda","Insecta","Diptera","Muscidae","Muscinae","Stomoxys","Stomoxys indicus","66","58","58","36","12.12","0","97.36","426.98","0","429.52","32.4","0"
"AGAW2","08N","INS","BOLD:AAH1890","Arthropoda","Insecta","Hymenoptera","Ichneumonidae","Cryptinae","Stibeutes","Stibeutes brevicornis, Stibeutes heinemanni","45","35","35","37","22.22","0","99.66","443.22","0","444.78","29.62","0"
"AGAW2","08N","INS","BOLD:ACP9679","Arthropoda","Insecta","Hymenoptera","Braconidae","Euphorinae","Microctonus","","43","43","43","38","0","0","98.53","433.6","0","435.84","31.3","0"
"AGAW2","08N","INS","BOLD:AAV1136","Arthropoda","Insecta","Diptera","Scatopsidae","","","","41","41","41","39","0","0","99.07","420.56","0","422.27","36.38","0"
"AGAW2","08N","INS","BOLD:AAN5220","Arthropoda","Insecta","Diptera","Cecidomyiidae","","","","40","40","40","40","0","0","98.18","410.9","0","426.58","26.32","0"
"AGAW2","08N","INS","BOLD:ACK4901","Arthropoda","Insecta","Diptera","Chironomidae","Chironominae","","","37","36","36","41","2.7","0","99.55","415.97","0","418.41","36.09","0"
"AGAW2","08N","INS","BOLD:ABV1455","Arthropoda","Insecta","Diptera","Chironomidae","Chironominae","","","37","34","34","41","8.11","0","98","435.78","0","438.41","31.79","0"
"AGAW2","08N","INS","BOLD:ABA7871","Arthropoda","Insecta","Diptera","Cecidomyiidae","","","","33","30","30","43","9.09","0","98.95","446.03","0","447.06","26.44","0"
"AGAW2","08N","INS","BOLD:AAB4894","Arthropoda","Insecta","Hemiptera","Aphididae","Aphidinae","Sitobion","Sitobion smilacifoliae, Sitobion avenae, Sitobion phyllanthi","28","28","28","44","0","0","97.8","433.14","0","434.82","25.13","0"
"AGAW2","08N","INS","BOLD:AAE4612","Arthropoda","Insecta","Hemiptera","Miridae","Phylinae","Plagiognathus","Plagiognathus suffuscipennis, Plagiognathus tumidifrons","27","20","20","45","25.93","0","99.68","447.41","0","448.07","38.24","0"
"AGAW2","08N","INS","BOLD:AAU7332","Arthropoda","Insecta","Coleoptera","Chrysomelidae","Galerucinae","Epitrix","Epitrix tuberis","25","25","25","46","0","0","99.15","393.12","0","394.76","36.7","0"
"AGAW2","08N","INS","BOLD:AAN5230","Arthropoda","Insecta","Diptera","Cecidomyiidae","","","","21","21","21","47","0","0","98.91","418.19","0","419.71","25.69","0"
"AGAW2","08N","INS","BOLD:AAG3154","Arthropoda","Insecta","Diptera","Psychodidae","Psychodinae","Psychoda","","21","21","21","47","0","0","98.28","447.95","0","450","31.99","0"
"AGAW2","08N","INS","BOLD:ADG6553","Arthropoda","Insecta","Diptera","Chironomidae","","","","21","20","20","47","4.76","0","97.2","388.9","0","448.86","34.07","0"
"AGAW2","08N","INS","BOLD:AAB1188","Arthropoda","Insecta","Hymenoptera","Ichneumonidae","Orthocentrinae","Orthocentrus","Orthocentrus corrugatus, Orthocentrus sannio, Orthocentrus attenuatus, Orthocentrus jft10, Orthocentrus jft09, Orthocentrus frontator","18","16","16","50","11.11","0","99.76","449.11","0","450","27.31","0"
"AGAW2","08N","INS","BOLD:AAC0903","Arthropoda","Insecta","Diptera","Chironomidae","Chironominae","Chironomus","Chironomus acidophilus","13","13","13","51","0","0","98.69","436.69","0","439.15","36.22","0"
"AGAW2","08N","INS","BOLD:AAC1374","Arthropoda","Insecta","Hemiptera","Aphididae","Aphidinae","Aphis","Aphis nasturtii, Aphis davletshinae, Aphis sp. rostella like, Aphis cf. rostella, Aphis althaeae, Aphis umbrella","13","13","13","51","0","0","99.33","438.85","0","440.31","26.08","0"
"AGAW2","08N","INS","BOLD:AAL7819","Arthropoda","Insecta","Diptera","Psychodidae","Psychodinae","Psychoda","Psychoda sp., Psychoda sp. BOLD:AAL7819","9","9","9","53","0","0","99.52","401.33","0","403.89","32.35","0"
"AGAW2","08N","INS","BOLD:AAN5267","Arthropoda","Insecta","Diptera","Cecidomyiidae","","","","9","8","8","53","11.11","0","99.07","441.11","0","450","30.42","0"
"AGAW2","08N","INS","BOLD:ACJ0111","Arthropoda","Insecta","Hymenoptera","Platygastridae","Scelioninae","Telenomus","","7","7","7","55","0","0","98.82","436.86","0","446.14","29.02","0"
"AGAW2","08N","INS","BOLD:AAG0734","Arthropoda","Insecta","Thysanoptera","Thripidae","Thripinae","Frankliniella","Frankliniella tritici","6","6","6","56","0","0","99.24","423.17","0","425","29.26","0"
"AGAW2","08N","INS","BOLD:AAM7855","Arthropoda","Insecta","Thysanoptera","Thripidae","Thripinae","Frankliniella","Frankliniella tritici","6","6","6","56","0","0","98.45","401.83","0","405.5","30.89","0"
"AGAW2","08N","INS","BOLD:ABU5525","Arthropoda","Insecta","Diptera","Chironomidae","Orthocladiinae","Limnophyes","Limnophyes sp. 14ES","6","6","6","56","0","0","99.11","370.17","0","374.83","37.44","0"
"AGAW2","08N","INS","BOLD:AAI4303","Arthropoda","Insecta","Diptera","Chironomidae","Chironominae","Chironomus","Chironomus melanescens","6","6","6","56","0","0","98.41","411.5","0","414","33.37","0"
"AGAW2","08N","INS","BOLD:ABX6298","Arthropoda","Insecta","Diptera","Tipulidae","Tipulinae","Nephrotoma","Nephrotoma ferruginea","5","5","5","60","0","0","97.83","436.6","0","437.4","34.32","0"
"AGAW2","08N","INS","BOLD:AAU6681","Arthropoda","Insecta","Diptera","Tipulidae","Tipulinae","Nephrotoma","Nephrotoma cornicina","5","5","5","60","0","0","99.33","449.2","0","450","36.53","0"
"AGAW2","08N","INS","BOLD:ABX2489","Arthropoda","Insecta","Coleoptera","Phalacridae","Phalacrinae","Stilbus","","5","5","5","60","0","0","97.56","416.4","0","417.6","32.81","0"
"AGAW2","08N","INS","BOLD:AAG5467","Arthropoda","Insecta","Diptera","Chironomidae","Chironominae","Tanytarsus","","4","4","4","63","0","0","98.3","415","0","417","31.98","0"
"AGAW2","08N","INS","BOLD:AAP8816","Arthropoda","Insecta","Diptera","Cecidomyiidae","Cecidomyiinae","Sitodiplosis","Sitodiplosis sp. BOLD:AAP8816","4","4","4","63","0","0","99.65","434.5","0","438","26.07","0"
"AGAW2","08N","INS","BOLD:AAP7843","Arthropoda","Insecta","Coleoptera","Cantharidae","Malthininae","Malthodes","Malthodes pumilus","4","4","4","63","0","0","97.81","411.5","0","414.5","38.1","0"
"AGAW2","08N","INS","BOLD:AAD2041","Arthropoda","Insecta","Diptera","Dolichopodidae","Sciapodinae","Condylostylus","Condylostylus inermis, Condylostylus nigrofemoratus, Condylostylus connectans, Condylostylus caudatus, Condylostylus flavipes","4","4","4","63","0","0","97.88","448.25","0","450","36.11","0"
"AGAW2","08N","INS","BOLD:ACK1456","Arthropoda","Insecta","Diptera","Cecidomyiidae","","","","4","4","4","63","0","0","99.02","431","0","443.25","25.83","0"
"AGAW2","08N","INS","BOLD:AAM6184","Arthropoda","Insecta","Diptera","Ceratopogonidae","Ceratopogoninae","Culicoides","","4","4","4","63","0","0","97.93","399.25","0","400","37","0"
"AGAW2","08N","INS","BOLD:AAN5235","Arthropoda","Insecta","Diptera","Cecidomyiidae","","","","3","3","3","69","0","0","99.7","449.67","0","450","26.96","0"
"AGAW2","08N","INS","BOLD:AAA3889","Arthropoda","Insecta","Lepidoptera","Crambidae","Crambinae","Parapediasia","Parapediasia teterrellus","3","3","3","69","0","0","98.5","445","0","447.33","32.41","0"
"AGAW2","08N","INS","BOLD:AAA2815","Arthropoda","Insecta","Lepidoptera","Noctuidae","Noctuinae","Mythimna","Mythimna yukonensis, Mythimna oxygala","3","3","3","69","0","0","98.72","421.33","0","422.33","33.94","0"
"AGAW2","08N","INS","BOLD:AAU6968","Arthropoda","Insecta","Coleoptera","Staphylinidae","Aleocharinae","Strigota","Strigota obscurata","3","3","3","69","0","0","99.25","395","0","401.33","37.45","0"
"AGAW2","08N","INS","BOLD:AAB2384","Arthropoda","Insecta","Diptera","Syrphidae","Syrphinae","Eupeodes","Eupeodes duseki, Eupeodes volucris, Eupeodes nielseni, Eupeodes americanus, Eupeodes flukei, Eupeodes vockerothi, Eupeodes lucasi, Eupeodes montivagus, Eupeodes rufipunctatus, Eupeodes rojasi, Eupeodes goeldlini, Eupeodes luniger, Eupeodes punctifer, Eupeodes bucculatus, Eupeodes perplexus, Eupeodes abiskoensis, Eupeodes corollae, Eupeodes pomus, Eupeodes lundbecki, Eupeodes nitens, Eupeodes sp. BOLDAAB2384, Eupeodes curtus, Eupeodes latifasciatus","3","3","3","69","0","0","98.64","403.67","0","405.33","30.92","0"
"AGAW2","08N","INS","BOLD:ABZ1405","Arthropoda","Insecta","Hemiptera","Miridae","Mirinae","Lygus","Lygus abroniae, Lygus unctuosus, Lygus plagiatus, Lygus ceanothi, Lygus potentillae, Lygus humeralis, Lygus keltoni, Lygus rolfsi, Lygus rufidorsus, Lygus hesperus, Lygus rugulipennis, Lygus solidaginis, Lygus atriflavus, Lygus lupini, Lygus atritibialis, Lygus scudderi, Lygus aeratus, Lygus elisus, Lygus rubroclarus, Lygus punctatus, Lygus bradleyi, Lygus borealis, Lygus rubrosignatus, Lygus oregonae, Lygus convexicollis, Lygus robustus, Lygus shulli, Lygus striatus","2","2","2","74","0","0","97.28","405.5","0","409","33.44","0"
"AGAW2","08N","INS","BOLD:AAA6292","Arthropoda","Insecta","Lepidoptera","Tortricidae","Tortricinae","Cnephasia","Cnephasia genitalana, Cnephasia stephensiana","2","2","2","74","0","0","99.22","440","0","442","35.52","0"
"AGAW2","08N","INS","BOLD:ACD9348","Arthropoda","Insecta","Hymenoptera","Platygastridae","","","","2","2","2","74","0","0","98.77","445.5","0","450","23.67","0"
"AGAW2","08N","INS","BOLD:AAC9132","Arthropoda","Insecta","Diptera","Culicidae","Culicinae","Culiseta","Culiseta inornata","2","2","2","74","0","0","99.35","390.5","0","393","31.98","0"
"AGAW2","08N","INS","BOLD:AAD5253","Arthropoda","Insecta","Diptera","Chironomidae","Orthocladiinae","Thienemanniella","Thienemanniella xena, Thienemanniella vittata","2","2","2","74","0","0","99.43","439","0","441.5","34.54","0"
"AGAW2","08N","INS","BOLD:AAH2293","Arthropoda","Insecta","Diptera","Asilidae","Stenopogoninae","Dioctria, Eudioctria","Dioctria hyalipennis, Dioctria baumhaueri, Eudioctria propinqua","2","2","2","74","0","0","98.81","396","0","398.5","30.52","0"
"AGAW2","08N","INS","BOLD:AAA4555","Arthropoda","Insecta","Orthoptera","Acrididae","Melanoplinae","Melanoplus","Melanoplus stupefactus, Melanoplus angustipennis, Melanoplus ponderosus, Melanoplus bruneri, Melanoplus frigidus, Melanoplus scapularis, Melanoplus cinereus, Melanoplus madeleineae, Melanoplus sanguinipes, Melanoplus nigrescens group, Melanoplus tunicae, Melanoplus walshi, Melanoplus femurrubrum, Melanoplus dawsoni, Melanoplus sp., Melanoplus fasciatus, Melanoplus bowditchi, Melanoplus stonei, Melanoplus bivittatus, Melanoplus huroni, Melanoplus thomasi, Melanoplus borealis, Melanoplus packardii","2","2","2","74","0","0","99.67","448","0","450","36","0"
"AGAW2","08N","INS","BOLD:AAG1739","Arthropoda","Insecta","Diptera","Muscidae","Coenosiinae","Neodexiopsis","Neodexiopsis rufitibia","2","2","2","74","0","0","99.89","447","0","450","34","0"
"AGAW2","08N","INS","BOLD:AAA3654","Arthropoda","Insecta","Lepidoptera","Crambidae","Crambinae","Crambus","Crambus bigelovi, Crambus johnsoni, Crambus praefectellus","2","2","2","74","0","0","98.11","450","0","450","31.67","0"
"AGAW2","08N","INS","BOLD:AAA7374","Arthropoda","Insecta","Diptera","Syrphidae","Syrphinae","Sphaerophoria","Sphaerophoria bankowskae, Sphaerophoria infuscata, Sphaerophoria nr. philanthus, Sphaerophoria batava, Sphaerophoria sp., Sphaerophoria rueppellii, Sphaerophoria interrupta-group, Sphaerophoria sp. MA-1, Sphaerophoria cleoae, Sphaerophoria asymmetrica, Sphaerophoria fatarum, Sphaerophoria bengalensis, Sphaerophoria bifurcata, Sphaerophoria taeniata, Sphaerophoria interrupta, Sphaerophoria contigua, Sphaerophoria kaa, Sphaerophoria laurae, Sphaerophoria chongjini, Sphaerophoria boreoalpina, Sphaerophoria sp. BOLDAAA7374, Sphaerophoria macrogaster, Sphaerophoria sulphuripes, Sphaerophoria brevipilosa, Sphaerophoria philanthus, Sphaerophoria pyrrhina, Sphaerophoria abbreviata, Sphaerophoria scripta, Sphaerophoria potentillae, Sphaerophoria longipilosa, Sphaerophoria virgata","2","2","2","74","0","0","99.35","417.5","0","420.5","30.41","0"
"AGAW2","08N","INS","BOLD:AAA6759","Arthropoda","Insecta","Lepidoptera","Hesperiidae","Hesperiinae","Thymelicus","Thymelicus lineola","1","1","1","84","0","0","99.54","437","0","438","30.37","0"
"AGAW2","08N","INS","BOLD:AAB0079","Arthropoda","Insecta","Diptera","Chironomidae","Orthocladiinae","Corynoneura","Corynoneura arctica","1","1","1","84","0","0","99.33","450","0","450","35.33","0"
"AGAW2","08N","INS","BOLD:AAG5507","Arthropoda","Insecta","Diptera","Chironomidae","Chironominae","Chironomus","Chironomus atrella","1","1","1","84","0","0","98.75","400","0","402","35.07","0"
"AGAW2","08N","INS","BOLD:AAG9668","Arthropoda","Insecta","Diptera","Dolichopodidae","Diaphorinae","Chrysotus","Chrysotus sp., Chrysotus sp. nr. obliquus","1","1","1","84","0","0","99.78","449","0","450","32.89","0"
"AGAW2","08N","INS","BOLD:AAP5169","Arthropoda","Insecta","Diptera","Chloropidae","Oscinellinae","Elachiptera","Elachiptera nigriceps","1","1","1","84","0","0","99.33","448","0","450","31.78","0"
"AGAW2","08N","INS","BOLD:AAA4751","Arthropoda","Insecta","Diptera","Culicidae","Culicinae","Culex","Culex pipiens sl, Culex nr. pipiens, Culex pipiens, Culex quinquefasciatus, Culex sp.","1","1","1","84","0","0","99.75","400","0","402","32.84","0"
"AGAW2","08N","INS","BOLD:ABV9018","Arthropoda","Insecta","Diptera","Mycetophilidae","Sciophilinae","Sciophila","","1","1","1","84","0","0","97.53","445","0","450","32.22","0"
"AGAW2","08N","INS","BOLD:ABY9688","Arthropoda","Insecta","Diptera","Chloropidae","Chloropinae","Thaumatomyia","Thaumatomyia glabra","1","1","1","84","0","0","97.03","370","0","375","32.53","0"
"AGAW2","08N","INS","BOLD:ACA9275","Arthropoda","Insecta","Diptera","Chironomidae","Chironominae","Chironomus","","1","1","1","84","0","0","99.78","449","0","450","36","0"
"AGAW2","08N","INS","BOLD:AAF9393","Arthropoda","Insecta","Coleoptera","Meloidae","Meloinae","Epicauta","Epicauta fabricii","1","1","1","84","0","0","98.67","450","0","450","40.67","0"
"AGAW2","08N","INS","BOLD:AAU6592","Arthropoda","Insecta","Diptera","Ceratopogonidae","Dasyheleinae","Dasyhelea","Dasyhelea arenivaga","1","1","1","84","0","0","99.11","449","0","450","35.56","0"
"AGAW2","08N","INS","BOLD:AAH3914","Arthropoda","Insecta","Diptera","Sciaridae","","Sciara","Sciara hamatilis, Sciara humeralis","1","1","1","84","0","0","99.78","450","0","450","30.44","0"
"AGAW2","08N","INS","BOLD:ACC8511","Arthropoda","Insecta","Diptera","Cecidomyiidae","Cecidomyiinae","Sitodiplosis","Sitodiplosis mosellana","1","1","1","84","0","0","98.32","358","0","360","25.28","0"
"AGAW2","08N","INS","BOLD:ACE7444","Arthropoda","Insecta","Hemiptera","Miridae","Mirinae, Phylinae","Adelphocoris, Megalocoleus","Adelphocoris lineolatus, Megalocoleus molliculus","1","1","1","84","0","0","99.78","450","0","450","34.67","0"
"AGAW2","08N","INS","BOLD:AAZ5967","Arthropoda","Insecta","Diptera","Limoniidae","Chioneinae","Molophilus","Molophilus forcipulus, Molophilus pubipennis","1","1","1","84","0","0","99.33","450","0","450","38.89","0"
"AGAW2","08N","INS","BOLD:AAB9652","Arthropoda","Insecta","Trichoptera","Leptoceridae","Leptocerinae","Triaenodes","Triaenodes nox","1","1","1","84","0","0","98.1","422","0","422","31.28","0"
"AGAW2","08N","INS","BOLD:AAO8324","Arthropoda","Insecta","Hemiptera","Cicadellidae","Typhlocybinae","Dikraneura","","1","1","1","84","0","0","97.31","409","0","412","35.68","0"
"AGAW2","08N","INS","BOLD:AAB9140","Arthropoda","Insecta","Diptera","Calliphoridae","Chrysomyinae","Phormia","Phormia regina","1","1","1","84","0","0","100","449","0","450","33.11","0"
"AGAW2","08N","INS","BOLD:AAD1485","Arthropoda","Insecta","Diptera","Chironomidae","Chironominae","Paratanytarsus","Paratanytarsus grimmii, Paratanytarsus sp.","1","1","1","84","0","0","98.89","450","0","450","34.89","0"
"AGAW2","08N","INS","BOLD:AAB7030","Arthropoda","Insecta","Diptera","Chironomidae","Chironominae","Chironomus","Chironomus sp. BOLDAAP2996, Chironomus quinnitukqut, Chironomus cf. decorus 2g, Chironomus decorus group sp. 2 IP2013, Chironomus cf. decorus, Chironomus decorus gr.","1","1","1","84","0","0","97.77","449","0","450","35.78","0"
"AGAW2","08N","INS","BOLD:AAF9187","Arthropoda","Insecta","Coleoptera","Curculionidae","Entiminae","Phyllobius","Phyllobius oblongus","1","1","1","84","0","0","99.18","368","0","368","39.13","0"
"AGAW2","08N","INS","BOLD:AAB1098","Arthropoda","Insecta","Diptera","Culicidae","Culicinae","Ochlerotatus, Aedes","Aedes excrucians, Aedes cantans, Aedes stimulans, Aedes euedes, Aedes riparius, Aedes annulipes, Ochlerotatus cantans, Aedes fitchii, Aedes sp., Ochlerotatus annulipes, Ochlerotatus excrucians","1","1","1","84","0","0","97.76","446","0","449","35.63","0"
"AGAW2","08N","INS","BOLD:AAN5342","Arthropoda","Insecta","Diptera","Chironomidae","Orthocladiinae","Heleniella","","1","1","1","84","0","0","98.88","447","0","450","35.56","0"
"AGAW2","08N","INS","BOLD:AAN5341","Arthropoda","Insecta","Diptera","Chironomidae","Orthocladiinae","Camptocladius","Camptocladius stercorarius","1","1","1","84","0","0","98.68","379","0","382","35.34","0"
"AGAW2","08N","INS","BOLD:ABV9293","Arthropoda","Insecta","Diptera","Chironomidae","Chironominae","Dicrotendipes","","1","1","1","84","0","0","100","356","0","450","35.11","0"
"AGAW2","08N","INS","BOLD:AAA3867","Arthropoda","Insecta","Lepidoptera","Noctuidae","Noctuinae","Apamea","Apamea amputatrix","1","1","1","84","0","0","98.87","444","0","450","32.89","0"
"AGAW2","08N","INS","BOLD:AAG6460","Arthropoda","Insecta","Diptera","Cecidomyiidae","","","","1","1","1","84","0","0","98","449","0","450","33.78","0"
"AGAW2","08N","INS","BOLD:AAH1888","Arthropoda","Insecta","Hymenoptera","Ichneumonidae","Ctenopelmatinae","","","1","1","1","84","0","0","99.78","450","0","450","29.56","0"
"AGAW2","08N","INS","BOLD:AAD5425","Arthropoda","Insecta","Ephemeroptera","Baetidae","Baetinae","Callibaetis","Callibaetis fluctuans","1","1","1","84","0","0","98.22","450","0","450","38.67","0"
"AGAW2","08N","INS","BOLD:AAB3656","Arthropoda","Insecta","Lepidoptera","Pterophoridae","Pterophorinae","Hellinsia","Hellinsia glenni, Hellinsia lacteodactylus, Hellinsia paleaceus","1","1","1","84","0","0","98.41","378","0","379","31.66","0"
"AGAW2","08N","INS","BOLD:AAC9062","Arthropoda","Insecta","Diptera","Culicidae","Culicinae","Ochlerotatus, Aedes","Aedes fitchii, Ochlerotatus fitchii, Aedes grossbecki, Aedes ventrovittis","1","1","1","84","0","0","99.5","398","0","398","32.66","0"
"AGAW2","08N","INS","BOLD:AAH7146","Arthropoda","Insecta","Diptera","Sarcophagidae","Sarcophaginae","Boettcheria","Boettcheria latisterna","1","1","1","84","0","0","99.55","449","0","450","34.67","0"
"AGAW2","08N","INS","BOLD:ABV1237","Arthropoda","Insecta","Diptera","Chironomidae","Chironominae","Chironomus","","1","1","1","84","0","0","100","450","0","450","35.33","0"
"AGAW2","08N","INS","BOLD:AAC2637","Arthropoda","Insecta","Lepidoptera","Pterophoridae","Pterophorinae","Geina","Geina sheppardi","1","1","1","84","0","0","99.78","446","0","450","30.89","0"
"AGAW2","08N","INS","BOLD:AAA4222","Arthropoda","Insecta","Lepidoptera","Erebidae","Hypeninae","Hypena","Hypena scabra","1","1","1","84","0","0","97.69","389","0","391","32.23","0"
"AGAW2","08N","INS","BOLD:AAY6534","Arthropoda","Insecta","Coleoptera","Mordellidae","Mordellinae","Mordellina","Mordellina ancilla","1","1","1","84","0","0","99.78","449","0","450","38.44","0"
"AGAW2","08N","NAR","no match","no match","no match","no match","no match","no match","no match","no match","no match","no match","no match","no match","no match","no match","no match","no match","no match","no match","no match","no match"
"AGAW2","08N","NIN","BOLD:AAH6630","Arthropoda","Collembola","Entomobryomorpha","Entomobryidae","Entomobryinae","Entomobrya","Entomobrya unostrigata","2483","1985","1985","1","20.06","0","99.39","439.15","0","440.36","39.03","0"
"AGAW2","08N","NIN","BOLD:AAI4346","Arthropoda","Arachnida","Opiliones","Phalangiidae","","Paroligolophus, Phalangium","Paroligolophus agrestis, Phalangium opilio","918","863","863","2","5.99","0","99","414.37","0","423.84","35.43","0"
"AGAW2","08N","NIN","BOLD:AAA5267","Arthropoda","Collembola","Entomobryomorpha","Isotomidae","","","","130","130","130","3","0","0","98.89","421.65","0","424.37","38.04","0"
"AGAW2","08N","NIN","BOLD:AAA4125","Arthropoda","Arachnida","Araneae","Araneidae","","Araneus","Araneus diadematus","110","101","101","4","8.18","0","99.31","437.91","0","439.22","34.03","0"
"AGAW2","08N","NIN","BOLD:ABW2639","Arthropoda","Arachnida","Trombidiformes","Anystidae","","","","76","66","66","5","13.16","0","99.6","447.82","0","446.87","31.46","0"
"AGAW2","08N","NIN","BOLD:ACR6860","","","","","","","","50","46","46","6","8","0","99.23","385.78","0","443.66","40.32","0"
"AGAW2","08N","NIN","BOLD:AAB7914","Arthropoda","Collembola","Symphypleona","Bourletiellidae","","Bourletiella","","41","38","38","7","7.32","0","99.48","443.44","0","444.59","38.04","0"
"AGAW2","08N","NIN","BOLD:AAC2734","Arthropoda","Arachnida","Araneae","Theridiidae","","Theridion","Theridion frondeum","27","27","27","8","0","0","98.01","414.26","0","419.93","28.06","0"
"AGAW2","08N","NIN","BOLD:AAB7915","Arthropoda","Collembola","Symphypleona","Bourletiellidae","","Deuterosminthurus","Deuterosminthurus sp1","23","20","20","9","13.04","0","99.32","445.74","0","446.39","35.22","0"
"AGAW2","08N","NIN","BOLD:AAM7961","Arthropoda","Arachnida","Trombidiformes","Anystidae","","","","20","20","20","10","0","0","99.66","448.65","0","446.85","31.32","0"
"AGAW2","08N","NIN","BOLD:ACZ1295","Arthropoda","Arachnida","Trombidiformes","Anystidae","","","","13","13","13","11","0","0","99.14","379.08","0","396.92","31.41","0"
"AGAW2","08N","NIN","BOLD:ACC0359","Arthropoda","Collembola","Symphypleona","Bourletiellidae","","Bourletiella","","9","9","9","12","0","0","99.38","444.44","0","447.67","37.75","0"
"AGAW2","08N","NIN","BOLD:AAB8452","Arthropoda","Collembola","Entomobryomorpha","Entomobryidae","Lepidocyrtinae","Lepidocyrtus","Lepidocyrtus paradoxus","6","6","6","13","0","0","99.78","455","0","450","42.26","0"
"AGAW2","08N","NIN","BOLD:AAO8147","Arthropoda","Arachnida","Trombidiformes","Eupodidae","","Eupodes","","5","5","5","14","0","0","98.49","409.8","0","411.2","28.19","0"
"AGAW2","08N","NIN","BOLD:ACI9974","Arthropoda","Arachnida","Trombidiformes","Tydeidae","","","","4","4","4","15","0","0","99.65","425","0","450","41","0"
"AGAW2","08N","NIN","BOLD:ADD2059","Arthropoda","Arachnida","Mesostigmata","Phytoseiidae","","","","4","3","3","15","25","0","99.82","428","0","450","33.39","0"
"AGAW2","08N","NIN","BOLD:ACM6751","Arthropoda","Arachnida","Trombidiformes","Anystidae","","","","2","2","2","17","0","0","97.02","403","0","450","30","0"
"AGAW2","08N","NIN","BOLD:AAA8820","Arthropoda","Collembola","Entomobryomorpha","Entomobryidae","Lepidocyrtinae","Lepidocyrtus","Lepidocyrtus cyaneus","2","2","2","17","0","0","99.01","411.5","0","415.5","42.96","0"
"AGAW2","08S","BAC","no match","no match","no match","no match","no match","no match","no match","no match","no match","no match","no match","no match","no match","no match","no match","no match","no match","no match","no match","no match"
"AGAW2","08S","CHO","no match","no match","no match","no match","no match","no match","no match","no match","no match","no match","no match","no match","no match","no match","no match","no match","no match","no match","no match","no match"
"AGAW2","08S","INS","BOLD:AAA3453","Arthropoda","Insecta","Diptera","Anthomyiidae","Anthomyiinae","Delia","","24139","19822","19822","1","17.88","0","99.31","432.07","0","434.51","34.61","0"
"AGAW2","08S","INS","BOLD:AAG1711","Arthropoda","Insecta","Diptera","Muscidae","Phaoniinae","Helina","Helina depuncta","22111","16683","16683","2","24.55","0","99.39","433.69","0","435.91","31.36","0"
"AGAW2","08S","INS","BOLD:AAG6745","Arthropoda","Insecta","Diptera","Calliphoridae","Calliphorinae","Pollenia","Pollenia pediculata","14902","12023","12023","3","19.32","0","99.29","433.57","0","435.58","33.49","0"
"AGAW2","08S","INS","BOLD:AAG2511","Arthropoda","Insecta","Diptera","Anthomyiidae","Anthomyiinae","Delia","Delia platura","12945","10539","10539","4","18.59","0","98.89","444.57","0","447.36","33.2","0"
"AGAW2","08S","INS","BOLD:AAH4135","Arthropoda","Insecta","Diptera","Chloropidae","Chloropinae","Thaumatomyia","Thaumatomyia glabra, Thaumatomyia grata, Thaumatomyia sp. 1 TAW","938","887","887","5","5.44","0","99.28","428.31","0","434.19","31.77","0"
"AGAW2","08S","INS","BOLD:AAA4277","Arthropoda","Insecta","Diptera","Syrphidae","Syrphinae","Toxomerus","Toxomerus marginatus, Toxomerus sp. CR-3, Toxomerus CR-3","849","616","616","6","27.44","0","99.2","444.86","0","445.67","30.77","0"
"AGAW2","08S","INS","BOLD:AAA7661","Arthropoda","Insecta","Diptera","Culicidae","Culicinae","Culiseta, Culex","Culiseta melanura, Culex articularis, Culex restuans, Culex acharistus","616","573","573","7","6.98","0","99.11","433.43","0","435.29","32.96","0"
"AGAW2","08S","INS","BOLD:AAG5507","Arthropoda","Insecta","Diptera","Chironomidae","Chironominae","Chironomus","Chironomus atrella","600","579","579","8","3.5","0","99.11","424.62","0","430.09","35.91","0"
"AGAW2","08S","INS","BOLD:ACR4394","Arthropoda","Insecta","Diptera","Anthomyiidae","Anthomyiinae","Delia","Delia florilega","534","510","510","9","4.49","0","99.05","431.31","0","433.23","32.96","0"
"AGAW2","08S","INS","BOLD:ACU7906","Arthropoda","Insecta","Diptera","Chironomidae","Orthocladiinae","","","345","344","344","10","0.29","0","98.87","398.41","0","409.85","36.37","0"
"AGAW2","08S","INS","BOLD:AAN5341","Arthropoda","Insecta","Diptera","Chironomidae","Orthocladiinae","Camptocladius","Camptocladius stercorarius","345","336","336","10","2.61","0","98.97","409.91","0","414.4","35.52","0"
"AGAW2","08S","INS","BOLD:AAL7382","Arthropoda","Insecta","Diptera","Chironomidae","Orthocladiinae","Psectrocladius","Psectrocladius sordidellus","257","211","211","12","17.9","0","99.48","438.52","0","439.77","36.51","0"
"AGAW2","08S","INS","BOLD:AAN5199","Arthropoda","Insecta","Diptera","Cecidomyiidae","","","CecidInt20 sp. BOLD:AAN5199","220","176","176","13","20","0","99.65","431.24","0","446.79","26.81","0"
"AGAW2","08S","INS","BOLD:ACK6562","Arthropoda","Insecta","Hemiptera","Miridae","Phylinae","Plagiognathus","Plagiognathus davisi, Plagiognathus delicatus","205","157","157","14","23.41","0","99.46","426.84","0","427.98","40.29","0"
"AGAW2","08S","INS","BOLD:AAG5457","Arthropoda","Insecta","Diptera","Chironomidae","Chironominae","","","188","175","175","15","6.91","0","98.27","429.48","0","432.1","33.92","0"
"AGAW2","08S","INS","BOLD:AAC0022","Arthropoda","Insecta","Diptera","Chironomidae","Chironominae","Micropsectra","Micropsectra nigripila","158","156","156","16","1.27","0","98.61","418.34","0","420.55","36.48","0"
"AGAW2","08S","INS","BOLD:AAD0484","Arthropoda","Insecta","Diptera","Chironomidae","","","","148","137","137","17","7.43","0","98.88","445.09","0","446.59","34.43","0"
"AGAW2","08S","INS","BOLD:AAA5299","Arthropoda","Insecta","Diptera","Chironomidae","Orthocladiinae","Cricotopus","Cricotopus laetus, Cricotopus relucens, Cricotopus trifasciatus, Cricotopus sp., Cricotopus sp. 3ES, Cricotopus sylvestris, Cricotopus glacialis","117","101","101","18","13.68","0","99.24","433.56","0","434.85","34.79","0"
"AGAW2","08S","INS","BOLD:AAF9771","Arthropoda","Insecta","Diptera","Hybotidae","Tachydromiinae","Platypalpus","Platypalpus longiseta, Platypalpus trivialis, Platypalpus sp. 5, Platypalpus sp. 17, Platypalpus pallidiventris","103","83","83","19","19.42","0","99.43","443.86","0","444.88","35.08","0"
"AGAW2","08S","INS","BOLD:AAN6154","Arthropoda","Insecta","Coleoptera","Latridiidae","Corticariinae","Melanophthalma","Melanophthalma inermis","99","91","91","20","8.08","0","99.14","435","0","437.33","31.29","0"
"AGAW2","08S","INS","BOLD:AAG3154","Arthropoda","Insecta","Diptera","Psychodidae","Psychodinae","Psychoda","","92","79","79","21","14.13","0","98.76","433.57","0","434.58","31.86","0"
"AGAW2","08S","INS","BOLD:ACE0554","Arthropoda","Insecta","Diptera","Phoridae","Metopininae","Megaselia","","88","75","75","22","14.77","0","99.5","442.6","0","444.39","32.07","0"
"AGAW2","08S","INS","BOLD:ABW7897","Arthropoda","Insecta","Diptera","Cecidomyiidae","","","","87","66","66","23","24.14","0","99.56","438.25","0","440.86","29.55","0"
"AGAW2","08S","INS","BOLD:AAC4197","Arthropoda","Insecta","Diptera","Chironomidae","Orthocladiinae","Paraphaenocladius","Paraphaenocladius impensus","70","65","65","24","7.14","0","99.07","427.23","0","432.51","34.61","0"
"AGAW2","08S","INS","BOLD:ABW1379","Arthropoda","Insecta","Diptera","Chloropidae","Oscinellinae","Malloewia","Malloewia abdominalis","70","63","63","24","10","0","97.83","431.99","0","442.73","31.58","0"
"AGAW2","08S","INS","BOLD:AAN6447","Arthropoda","Insecta","Diptera","Sciaridae","","Corynoptera","Corynoptera perpusilla","57","46","46","26","19.3","0","99.36","446.37","0","447.7","36.27","0"
"AGAW2","08S","INS","BOLD:AAD1945","Arthropoda","Insecta","Diptera","Agromyzidae","Phytomyzinae","Cerodontha","Cerodontha dorsalis","53","46","46","27","13.21","0","99.56","442.17","0","444.15","31.86","0"
"AGAW2","08S","INS","BOLD:AAG2868","Arthropoda","Insecta","Hemiptera","Cicadellidae","Typhlocybinae","Empoasca","Empoasca sp., Empoasca fabae","48","41","41","28","14.58","0","99.18","442.73","0","447.13","32.34","0"
"AGAW2","08S","INS","BOLD:ACZ1491","Arthropoda","Insecta","Diptera","Chironomidae","","","","43","38","38","29","11.63","0","97.76","445.81","0","447.88","34.26","0"
"AGAW2","08S","INS","BOLD:ACK3431","Arthropoda","Insecta","Diptera","Cecidomyiidae","","","","41","34","34","30","17.07","0","98.82","440.02","0","441.49","32.51","0"
"AGAW2","08S","INS","BOLD:AAG7036","Arthropoda","Insecta","Diptera","Agromyzidae","","","","29","29","29","31","0","0","99.02","399.31","0","407.31","32.62","0"
"AGAW2","08S","INS","BOLD:AAD4214","Arthropoda","Insecta","Hymenoptera","Ichneumonidae","Diplazontinae","Diplazon","Diplazon laetatorius","26","23","23","32","11.54","0","99.41","448.19","0","449.04","25.83","0"
"AGAW2","08S","INS","BOLD:AAV5688","Arthropoda","Insecta","Diptera","Cecidomyiidae","","","","23","20","20","33","13.04","0","99.16","425.96","0","427.26","34.55","0"
"AGAW2","08S","INS","BOLD:AAH3661","Arthropoda","Insecta","Diptera","Cecidomyiidae","","","","22","21","21","34","4.55","0","99.05","434.27","0","435.82","25.53","0"
"AGAW2","08S","INS","BOLD:ABU5525","Arthropoda","Insecta","Diptera","Chironomidae","Orthocladiinae","Limnophyes","Limnophyes sp. 14ES","21","21","21","35","0","0","98.92","400.29","0","407.86","37.3","0"
"AGAW2","08S","INS","BOLD:AAN5342","Arthropoda","Insecta","Diptera","Chironomidae","Orthocladiinae","Heleniella","","19","17","17","36","10.53","0","98.46","424.84","0","431.42","35.28","0"
"AGAW2","08S","INS","BOLD:AAH0135","Arthropoda","Insecta","Coleoptera","Phalacridae","Phalacrinae","Acylomus","Acylomus pugetanus, Acylomus ergoti","18","18","18","37","0","0","99.02","440.11","0","441.61","31.72","0"
"AGAW2","08S","INS","BOLD:AAA4555","Arthropoda","Insecta","Orthoptera","Acrididae","Melanoplinae","Melanoplus","Melanoplus stupefactus, Melanoplus angustipennis, Melanoplus ponderosus, Melanoplus bruneri, Melanoplus frigidus, Melanoplus scapularis, Melanoplus cinereus, Melanoplus madeleineae, Melanoplus sanguinipes, Melanoplus nigrescens group, Melanoplus tunicae, Melanoplus walshi, Melanoplus femurrubrum, Melanoplus dawsoni, Melanoplus sp., Melanoplus fasciatus, Melanoplus bowditchi, Melanoplus stonei, Melanoplus bivittatus, Melanoplus huroni, Melanoplus thomasi, Melanoplus borealis, Melanoplus packardii","13","13","13","38","0","0","99.26","444","0","445.92","36.09","0"
"AGAW2","08S","INS","BOLD:AAH0134","Arthropoda","Insecta","Coleoptera","Phalacridae","Phalacrinae","Stilbus","Stilbus apicialis","10","10","10","39","0","0","98.92","431.6","0","433.3","31.83","0"
"AGAW2","08S","INS","BOLD:AAP8739","Arthropoda","Insecta","Diptera","Platystomatidae","Platystomatinae","Rivellia","Rivellia steyskali","9","9","9","40","0","0","98.86","442","0","443.89","37.73","0"
"AGAW2","08S","INS","BOLD:AAC0903","Arthropoda","Insecta","Diptera","Chironomidae","Chironominae","Chironomus","Chironomus acidophilus","8","8","8","41","0","0","99.32","416","0","418","36.3","0"
"AGAW2","08S","INS","BOLD:AAN5183","Arthropoda","Insecta","Diptera","Cecidomyiidae","","","","8","7","7","41","12.5","0","98.61","449","0","450","27.94","0"
"AGAW2","08S","INS","BOLD:AAU6681","Arthropoda","Insecta","Diptera","Tipulidae","Tipulinae","Nephrotoma","Nephrotoma cornicina","7","7","7","43","0","0","99.27","448.14","0","448.71","36.2","0"
"AGAW2","08S","INS","BOLD:AAB0373","Arthropoda","Insecta","Neuroptera","Chrysopidae","Chrysopinae","Chrysoperla","Chrysoperla sp. isolate 82, Chrysoperla zastrowi, Chrysoperla plorabunda, Chrysoperla sp. isolate 71, Chrysoperla agilis, Italochrysa italica, Chrysoperla sp. isolate 88, Chrysoperla lucasina, Chrysoperla carnea, Chrysoperla mediterranea, Chrysoperla pallida, Chrysoperla nipponensis, Chrysoperla sillemi","7","7","7","43","0","0","99.45","432.57","0","435.29","31.23","0"
"AGAW2","08S","INS","BOLD:ABY5318","Arthropoda","Insecta","Diptera","Chloropidae","Oscinellinae","Incertella, Rhopalopterum","Rhopalopterum umbrosum, Rhopalopterum luteiceps, Incertella bispina, Rhopalopterum painteri","7","7","7","43","0","0","99.45","442","0","444.57","31.94","0"
"AGAW2","08S","INS","BOLD:ABZ3101","Arthropoda","Insecta","Hymenoptera","Braconidae","Aphidiinae","Aphidius","","7","7","7","43","0","0","98.23","420.57","0","431","25.8","0"
"AGAW2","08S","INS","BOLD:ADH9832","","","","","","","","6","6","6","47","0","0","98.36","446.17","0","450","32.93","0"
"AGAW2","08S","INS","BOLD:AAG2460","Arthropoda","Insecta","Diptera","Anthomyiidae","Anthomyiinae","Hydrophoria","Hydrophoria lancifer","5","5","5","48","0","0","98.54","436","0","441","34.03","0"
"AGAW2","08S","INS","BOLD:ABA1215","Arthropoda","Insecta","Diptera","Sciaridae","","Lycoriella","Lycoriella sativae","4","4","4","49","0","0","99.11","444","0","445.5","35.57","0"
"AGAW2","08S","INS","BOLD:AAN5220","Arthropoda","Insecta","Diptera","Cecidomyiidae","","","","4","4","4","49","0","0","98.14","436","0","436.75","27.01","0"
"AGAW2","08S","INS","BOLD:AAA3867","Arthropoda","Insecta","Lepidoptera","Noctuidae","Noctuinae","Apamea","Apamea amputatrix","4","4","4","49","0","0","99.26","442.25","0","444","33.55","0"
"AGAW2","08S","INS","BOLD:ACR3866","Arthropoda","Insecta","Psocodea","Caeciliusidae","Caeciliusinae","Valenzuela","","4","4","4","49","0","0","99.61","446.25","0","448","35.66","0"
"AGAW2","08S","INS","BOLD:AAQ0558","Arthropoda","Insecta","Thysanoptera","Thripidae","Thripinae","Anaphothrips","Anaphothrips obscurus","3","3","3","53","0","0","99.6","425.67","0","427","30.96","0"
"AGAW2","08S","INS","BOLD:AAN5242","Arthropoda","Insecta","Diptera","Cecidomyiidae","","","","3","3","3","53","0","0","98.3","387.67","0","390","28.28","0"
"AGAW2","08S","INS","BOLD:AAG7891","Arthropoda","Insecta","Hymenoptera","Platygastridae","Scelioninae","Telenomus","Telenomus podisi","3","3","3","53","0","0","99.15","435","0","442.33","27.06","0"
"AGAW2","08S","INS","BOLD:AAM6248","Arthropoda","Insecta","Diptera","Chironomidae","Chironominae","Microtendipes","Microtendipes pedellus grp.","3","3","3","53","0","0","99.24","402","0","404.33","30.26","0"
"AGAW2","08S","INS","BOLD:AAG2740","Arthropoda","Insecta","Diptera","Ephydridae","Parydrinae","Philygria","Philygria debilis","3","3","3","53","0","0","97.81","396.33","0","400.67","31.78","0"
"AGAW2","08S","INS","BOLD:AAA3181","Arthropoda","Insecta","Diptera","Muscidae","Muscinae","Stomoxys","Stomoxys calcitrans","2","2","2","58","0","0","98.45","430","0","432.5","32.81","0"
"AGAW2","08S","INS","BOLD:AAA6759","Arthropoda","Insecta","Lepidoptera","Hesperiidae","Hesperiinae","Thymelicus","Thymelicus lineola","2","2","2","58","0","0","98.77","448","0","450","30","0"
"AGAW2","08S","INS","BOLD:AAG1714","Arthropoda","Insecta","Diptera","Muscidae","Azeliinae","Muscina","Muscina pascuorum","2","2","2","58","0","0","99.88","431.5","0","450","34.22","0"
"AGAW2","08S","INS","BOLD:AAG1717","Arthropoda","Insecta","Diptera","Muscidae","Phaoniinae","Helina","Helina troene","2","2","2","58","0","0","99.55","449","0","450","30.89","0"
"AGAW2","08S","INS","BOLD:AAC9132","Arthropoda","Insecta","Diptera","Culicidae","Culicinae","Culiseta","Culiseta inornata","2","2","2","58","0","0","97.51","406","0","406.5","32.86","0"
"AGAW2","08S","INS","BOLD:AAU6592","Arthropoda","Insecta","Diptera","Ceratopogonidae","Dasyheleinae","Dasyhelea","Dasyhelea arenivaga","2","2","2","58","0","0","98.01","409.5","0","411","35.16","0"
"AGAW2","08S","INS","BOLD:AAA2632","Arthropoda","Insecta","Lepidoptera","Noctuidae","Noctuinae","Noctua","Noctua atlantica, Noctua pronuba","2","2","2","58","0","0","98.59","417","0","450","31.44","0"
"AGAW2","08S","INS","BOLD:ACE7444","Arthropoda","Insecta","Hemiptera","Miridae","Mirinae, Phylinae","Adelphocoris, Megalocoleus","Adelphocoris lineolatus, Megalocoleus molliculus","2","2","2","58","0","0","99","449.5","0","450","35.33","0"
"AGAW2","08S","INS","BOLD:ACC8511","Arthropoda","Insecta","Diptera","Cecidomyiidae","Cecidomyiinae","Sitodiplosis","Sitodiplosis mosellana","2","2","2","58","0","0","99.78","449","0","450","26.78","0"
"AGAW2","08S","INS","BOLD:AAB9140","Arthropoda","Insecta","Diptera","Calliphoridae","Chrysomyinae","Phormia","Phormia regina","2","2","2","58","0","0","99","448","0","450","32.89","0"
"AGAW2","08S","INS","BOLD:ACJ6984","Arthropoda","Insecta","Hymenoptera","Platygastridae","Scelioninae","Trissolcus","Trissolcus thyantae","2","2","2","58","0","0","99.66","438.5","0","450","27.22","0"
"AGAW2","08S","INS","BOLD:AAH2293","Arthropoda","Insecta","Diptera","Asilidae","Stenopogoninae","Dioctria, Eudioctria","Dioctria hyalipennis, Dioctria baumhaueri, Eudioctria propinqua","2","2","2","58","0","0","98.8","412","0","413.5","30.72","0"
"AGAW2","08S","INS","BOLD:ADG6553","Arthropoda","Insecta","Diptera","Chironomidae","","","","2","2","2","58","0","0","97.43","389","0","450","34.22","0"
"AGAW2","08S","INS","BOLD:AAA3654","Arthropoda","Insecta","Lepidoptera","Crambidae","Crambinae","Crambus","Crambus bigelovi, Crambus johnsoni, Crambus praefectellus","2","2","2","58","0","0","98.66","421","0","421.5","31.43","0"
"AGAW2","08S","INS","BOLD:AAN6433","Arthropoda","Insecta","Diptera","Sciaridae","","Corynoptera","Corynoptera deserta","2","2","2","58","0","0","100","447.5","0","450","30.56","0"
"AGAW2","08S","INS","BOLD:AAA5953","Arthropoda","Insecta","Lepidoptera","Gelechiidae","Gelechiinae","Coleotechnites","Coleotechnites sp., Coleotechnites piceaella, Coleotechnites laricis, Coleotechnites sp. 3, Coleotechnites carbonaria, Coleotechnites granti, Coleotechnites macleodi, Coleotechnites gibsonella, Coleotechnites coniferella, Coleotechnites martini","2","2","2","58","0","0","98.55","449","0","450","33.89","0"
"AGAW2","08S","INS","BOLD:AAA2590","Arthropoda","Insecta","Lepidoptera","Noctuidae","Noctuinae","Xestia","Xestia baja, Xestia smithii","1","1","1","74","0","0","99.78","449","0","450","32.89","0"
"AGAW2","08S","INS","BOLD:AAA6560","Arthropoda","Insecta","Lepidoptera","Crambidae","Crambinae","Crambus","Crambus multilinellus, Crambus saltuellus","1","1","1","74","0","0","100","450","0","450","31.11","0"
"AGAW2","08S","INS","BOLD:AAH7142","Arthropoda","Insecta","Diptera","Sarcophagidae","Sarcophaginae","Oxysarcodexia, Sarcophaginae_incertae_sedis","Oxysarcodexia cingarus, Sarcophaginae_incertae_sedis sp. 1 AGT, Oxysarcodexia ventricosa","1","1","1","74","0","0","99.33","449","0","450","34.22","0"
"AGAW2","08S","INS","BOLD:AAN5497","Arthropoda","Insecta","Diptera","Anthomyiidae","Pegomyinae","Pegomya","Pegomya solennis, Pegomya bicolor","1","1","1","74","0","0","98","450","0","450","29.78","0"
"AGAW2","08S","INS","BOLD:AAA8933","Arthropoda","Insecta","Coleoptera","Coccinellidae","Coccinellinae","Coccinella","Coccinella septempunctata","1","1","1","74","0","0","99.78","447","0","450","32.67","0"
"AGAW2","08S","INS","BOLD:AAG2123","Arthropoda","Insecta","Diptera","Tachinidae","Dexiinae","Dinera","Dinera grisescens","1","1","1","74","0","0","99.55","449","0","450","32.67","0"
"AGAW2","08S","INS","BOLD:AAH3664","Arthropoda","Insecta","Diptera","Cecidomyiidae","","","CecidInt3 sp. BOLD:AAH3664","1","1","1","74","0","0","97.33","449","0","450","27.56","0"
"AGAW2","08S","INS","BOLD:AAH3914","Arthropoda","Insecta","Diptera","Sciaridae","","Sciara","Sciara hamatilis, Sciara humeralis","1","1","1","74","0","0","99.33","450","0","450","30.89","0"
"AGAW2","08S","INS","BOLD:ACE7482","Arthropoda","Insecta","Hemiptera","Miridae","","","","1","1","1","74","0","0","97.47","434","0","436","35.55","0"
"AGAW2","08S","INS","BOLD:AAB7869","Arthropoda","Insecta","Lepidoptera","Tortricidae","Tortricinae","Phtheochroa","Phtheochroa baracana, Phtheochroa riscana, Phtheochroa birdana, Phtheochroa fulviplicana, Phtheochroa canariana","1","1","1","74","0","0","99.78","449","0","450","29.56","0"
"AGAW2","08S","INS","BOLD:AAA3889","Arthropoda","Insecta","Lepidoptera","Crambidae","Crambinae","Parapediasia","Parapediasia teterrellus","1","1","1","74","0","0","99.55","449","0","450","32.44","0"
"AGAW2","08S","INS","BOLD:AAA2326","Arthropoda","Insecta","Hymenoptera","Apidae","Apinae","Apis","Apis mellifera","1","1","1","74","0","0","98","450","0","450","26","0"
"AGAW2","08S","INS","BOLD:ABZ5071","Arthropoda","Insecta","Diptera","Chironomidae","Chironominae","Micropsectra","Micropsectra nigripila","1","1","1","74","0","0","97.67","386","0","389","36.5","0"
"AGAW2","08S","INS","BOLD:AAD1395","Arthropoda","Insecta","Diptera","Chironomidae","Chironominae","Polypedilum","Polypedilum convictum, Polypedilum flavum","1","1","1","74","0","0","98.89","449","0","450","33.56","0"
"AGAW2","08S","INS","BOLD:AAD2041","Arthropoda","Insecta","Diptera","Dolichopodidae","Sciapodinae","Condylostylus","Condylostylus inermis, Condylostylus nigrofemoratus, Condylostylus connectans, Condylostylus caudatus, Condylostylus flavipes","1","1","1","74","0","0","98.66","447","0","450","35.56","0"
"AGAW2","08S","INS","BOLD:AAA2815","Arthropoda","Insecta","Lepidoptera","Noctuidae","Noctuinae","Mythimna","Mythimna yukonensis, Mythimna oxygala","1","1","1","74","0","0","98.36","366","0","368","33.97","0"
"AGAW2","08S","INS","BOLD:AAP8989","Arthropoda","Insecta","Diptera","Micropezidae","Calobatinae","Compsobata","Compsobata univitta, Compsobata sp.","1","1","1","74","0","0","97.33","450","0","450","36","0"
"AGAW2","08S","INS","BOLD:ABV1297","Arthropoda","Insecta","Diptera","Sciaridae","","","","1","1","1","74","0","0","99.11","449","0","450","30.67","0"
"AGAW2","08S","INS","BOLD:AAB1098","Arthropoda","Insecta","Diptera","Culicidae","Culicinae","Ochlerotatus, Aedes","Aedes excrucians, Aedes cantans, Aedes stimulans, Aedes euedes, Aedes riparius, Aedes annulipes, Ochlerotatus cantans, Aedes fitchii, Aedes sp., Ochlerotatus annulipes, Ochlerotatus excrucians","1","1","1","74","0","0","98.42","444","0","445","35.28","0"
"AGAW2","08S","INS","BOLD:AAH0256","Arthropoda","Insecta","Coleoptera","Latridiidae","Corticariinae","Corticaria, Corticarina","","1","1","1","74","0","0","97.96","392","0","396","31.57","0"
"AGAW2","08S","INS","BOLD:ABX6298","Arthropoda","Insecta","Diptera","Tipulidae","Tipulinae","Nephrotoma","Nephrotoma ferruginea","1","1","1","74","0","0","99.18","367","0","370","33.78","0"
"AGAW2","08S","INS","BOLD:ABV9290","Arthropoda","Insecta","Diptera","Cecidomyiidae","","","","1","1","1","74","0","0","99.78","450","0","450","26","0"
"AGAW2","08S","INS","BOLD:ABV9293","Arthropoda","Insecta","Diptera","Chironomidae","Chironominae","Dicrotendipes","","1","1","1","74","0","0","97.46","354","0","450","37.33","0"
"AGAW2","08S","INS","BOLD:AAM6249","","","","","","","","1","1","1","74","0","0","99.78","449","0","450","32.89","0"
"AGAW2","08S","INS","BOLD:AAG1739","Arthropoda","Insecta","Diptera","Muscidae","Coenosiinae","Neodexiopsis","Neodexiopsis rufitibia","1","1","1","74","0","0","99.11","447","0","450","33.33","0"
"AGAW2","08S","INS","BOLD:AAN5148","Arthropoda","Insecta","Diptera","Ceratopogonidae","Forcipomyiinae","Forcipomyia","Forcipomyia bipunctata, Forcipomyia sp. CK-2012","1","1","1","74","0","0","97.89","427","0","428","35.28","0"
"AGAW2","08S","INS","BOLD:AAH4205","Arthropoda","Insecta","Diptera","Chloropidae","Chloropinae","Meromyza","Meromyza nigriseta, Meromyza sp.","1","1","1","74","0","0","97.99","448","0","450","32.89","0"
"AGAW2","08S","INS","BOLD:AAI4303","Arthropoda","Insecta","Diptera","Chironomidae","Chironominae","Chironomus","Chironomus melanescens","1","1","1","74","0","0","97.11","311","0","397","34.26","0"
"AGAW2","08S","INS","BOLD:ADF3937","","","","","","","","1","1","1","74","0","0","98.59","425","0","428","31.78","0"
"AGAW2","08S","INS","BOLD:AAD8086","Arthropoda","Insecta","Lepidoptera","Elachistidae","Elachistinae","Elachista","Elachista griseicornis, Elachista acenteta","1","1","1","74","0","0","98.88","447","0","450","30.67","0"
"AGAW2","08S","INS","BOLD:ABX0657","Arthropoda","Insecta","Coleoptera","Anthicidae","Notoxinae","Notoxus","Notoxus desertus","1","1","1","74","0","0","98.89","450","0","450","36.22","0"
"AGAW2","08S","INS","BOLD:AAM9255","Arthropoda","Insecta","Diptera","Sciaridae","","Pseudolycoriella","","1","1","1","74","0","0","97.56","450","0","450","32","0"
"AGAW2","08S","NAR","no match","no match","no match","no match","no match","no match","no match","no match","no match","no match","no match","no match","no match","no match","no match","no match","no match","no match","no match","no match"
"AGAW2","08S","NIN","BOLD:AAH6630","Arthropoda","Collembola","Entomobryomorpha","Entomobryidae","Entomobryinae","Entomobrya","Entomobrya unostrigata","1209","1015","1015","1","16.05","0","99.38","438.66","0","439.86","39.02","0"
"AGAW2","08S","NIN","BOLD:AAI4346","Arthropoda","Arachnida","Opiliones","Phalangiidae","","Paroligolophus, Phalangium","Paroligolophus agrestis, Phalangium opilio","61","58","58","2","4.92","0","99.01","412.28","0","420.59","35.35","0"
"AGAW2","08S","NIN","BOLD:ABW2639","Arthropoda","Arachnida","Trombidiformes","Anystidae","","","","46","42","42","3","8.7","0","99.56","445.87","0","444.28","31.42","0"
"AGAW2","08S","NIN","BOLD:AAC0175","Arthropoda","Arachnida","Araneae","Theridiidae","","Parasteatoda","Parasteatoda tepidariorum, Parasteatoda simulans","29","28","28","4","3.45","0","98.89","437.69","0","439.86","32.01","0"
"AGAW2","08S","NIN","BOLD:AAA5267","Arthropoda","Collembola","Entomobryomorpha","Isotomidae","","","","23","23","23","5","0","0","99.04","420.43","0","422.91","38.02","0"
"AGAW2","08S","NIN","BOLD:AAU6252","Arthropoda","Arachnida","Trombidiformes","Bdellidae","","Cyta","","17","16","16","6","5.88","0","99.4","414.18","0","415.47","31.12","0"
"AGAW2","08S","NIN","BOLD:AAM7961","Arthropoda","Arachnida","Trombidiformes","Anystidae","","","","11","11","11","7","0","0","99.37","438.36","0","434.36","31.34","0"
"AGAW2","08S","NIN","BOLD:AAB7914","Arthropoda","Collembola","Symphypleona","Bourletiellidae","","Bourletiella","","6","6","6","8","0","0","98.78","442.33","0","444.83","38.1","0"
"AGAW2","08S","NIN","BOLD:ACZ1295","Arthropoda","Arachnida","Trombidiformes","Anystidae","","","","5","5","5","9","0","0","98.8","379.8","0","396.2","31.09","0"
"AGAW2","08S","NIN","BOLD:ADD2059","Arthropoda","Arachnida","Mesostigmata","Phytoseiidae","","","","3","3","3","10","0","0","99.38","427.67","0","450","32.81","0"
"AGAW2","08S","NIN","BOLD:ACI9974","Arthropoda","Arachnida","Trombidiformes","Tydeidae","","","","1","1","1","11","0","0","99.24","394","0","393","40.2","0"
"AGAW2","08S","NIN","BOLD:AAA7242","Arthropoda","Collembola","Entomobryomorpha","Entomobryidae","Seirinae, Entomobryinae","Entomobrya, Lepidosira","Entomobrya multifasciata, Lepidosira sp.","1","1","1","11","0","0","99.1","445","0","450","39.56","0"
"AGAW2","08S","NIN","BOLD:AAB7915","Arthropoda","Collembola","Symphypleona","Bourletiellidae","","Deuterosminthurus","Deuterosminthurus sp1","1","1","1","11","0","0","99.78","448","0","450","35.11","0"
"AGAW2","08S","NIN","BOLD:AAA8924","Arthropoda","Collembola","Entomobryomorpha","Entomobryidae","Entomobryinae","Entomobrya","Entomobrya nivalis","1","1","1","11","0","0","98.78","409","0","411","38.69","0"
"AGAW2","08W","BAC","TAX:175768","Proteobacteria","Alphaproteobacteria","Rickettsiales","Rickettsiaceae",NA,"Wolbachia","Wolbachia pipientis","16","15","15","1","6.25","0","98.13","447.13","0","448.5","39.53","0"
"AGAW2","08W","BAC","TAX:91393","Proteobacteria","Alphaproteobacteria","Rickettsiales","Rickettsiaceae",NA,"Wolbachia","","15","15","15","2","0","0","99.12","391.87","0","419.8","39.26","0"
"AGAW2","08W","CHO","no match","no match","no match","no match","no match","no match","no match","no match","no match","no match","no match","no match","no match","no match","no match","no match","no match","no match","no match","no match"
"AGAW2","08W","INS","BOLD:AAG1711","Arthropoda","Insecta","Diptera","Muscidae","Phaoniinae","Helina","Helina depuncta","26220","19861","19861","1","24.25","0","99.44","433.36","0","435.66","31.34","0"
"AGAW2","08W","INS","BOLD:AAA2632","Arthropoda","Insecta","Lepidoptera","Noctuidae","Noctuinae","Noctua","Noctua atlantica, Noctua pronuba","16168","9436","9436","2","41.64","0","99.16","446.28","0","447.61","31.27","0"
"AGAW2","08W","INS","BOLD:AAA3453","Arthropoda","Insecta","Diptera","Anthomyiidae","Anthomyiinae","Delia","","8168","7047","7047","3","13.72","0","99.35","432.23","0","434.81","34.62","0"
"AGAW2","08W","INS","BOLD:AAG6745","Arthropoda","Insecta","Diptera","Calliphoridae","Calliphorinae","Pollenia","Pollenia pediculata","6470","5848","5848","4","9.61","0","99.32","407.71","0","409.47","33.44","0"
"AGAW2","08W","INS","BOLD:AAG2511","Arthropoda","Insecta","Diptera","Anthomyiidae","Anthomyiinae","Delia","Delia platura","3845","3303","3303","5","14.1","0","98.98","444.52","0","447.43","33.11","0"
"AGAW2","08W","INS","BOLD:AAB0373","Arthropoda","Insecta","Neuroptera","Chrysopidae","Chrysopinae","Chrysoperla","Chrysoperla sp. isolate 82, Chrysoperla zastrowi, Chrysoperla plorabunda, Chrysoperla sp. isolate 71, Chrysoperla agilis, Italochrysa italica, Chrysoperla sp. isolate 88, Chrysoperla lucasina, Chrysoperla carnea, Chrysoperla mediterranea, Chrysoperla pallida, Chrysoperla nipponensis, Chrysoperla sillemi","3299","2442","2442","6","25.98","0","99.48","437.43","0","438.38","31.32","0"
"AGAW2","08W","INS","BOLD:AAA7661","Arthropoda","Insecta","Diptera","Culicidae","Culicinae","Culiseta, Culex","Culiseta melanura, Culex articularis, Culex restuans, Culex acharistus","1004","838","838","7","16.53","0","99.03","437.33","0","439.23","33.27","0"
"AGAW2","08W","INS","BOLD:AAD0484","Arthropoda","Insecta","Diptera","Chironomidae","","","","627","565","565","8","9.89","0","98.85","444.16","0","445.93","34.62","0"
"AGAW2","08W","INS","BOLD:AAG5507","Arthropoda","Insecta","Diptera","Chironomidae","Chironominae","Chironomus","Chironomus atrella","555","541","541","9","2.52","0","98.91","429.79","0","434.1","35.89","0"
"AGAW2","08W","INS","BOLD:AAB2644","Arthropoda","Insecta","Diptera","Chironomidae","Orthocladiinae","Orthocladius","Orthocladius lapponicus, Orthocladius dorenus, Orthocladius mallochi","473","385","385","10","18.6","0","99.33","444.32","0","446.3","32.19","0"
"AGAW2","08W","INS","BOLD:AAN6154","Arthropoda","Insecta","Coleoptera","Latridiidae","Corticariinae","Melanophthalma","Melanophthalma inermis","424","361","361","11","14.86","0","98.84","439.7","0","441.27","31.31","0"
"AGAW2","08W","INS","BOLD:ABW1379","Arthropoda","Insecta","Diptera","Chloropidae","Oscinellinae","Malloewia","Malloewia abdominalis","330","251","251","12","23.94","0","98.31","443.9","0","447.06","31.55","0"
"AGAW2","08W","INS","BOLD:AAA4201","Arthropoda","Insecta","Lepidoptera","Noctuidae","Noctuinae","Apamea","Apamea sp., Apamea sordens, Apamea sp. nr. sordens, Apamea digitula","309","266","266","13","13.92","0","99.34","439.33","0","440.28","31.99","0"
"AGAW2","08W","INS","BOLD:AAD4214","Arthropoda","Insecta","Hymenoptera","Ichneumonidae","Diplazontinae","Diplazon","Diplazon laetatorius","295","240","240","14","18.64","0","99.55","445.07","0","447.24","25.78","0"
"AGAW2","08W","INS","BOLD:AAF9771","Arthropoda","Insecta","Diptera","Hybotidae","Tachydromiinae","Platypalpus","Platypalpus longiseta, Platypalpus trivialis, Platypalpus sp. 5, Platypalpus sp. 17, Platypalpus pallidiventris","294","229","229","15","22.11","0","99.52","445.9","0","446.7","35.14","0"
"AGAW2","08W","INS","BOLD:ACX5521","Arthropoda","Insecta","Diptera","Chironomidae","Orthocladiinae","","","263","250","250","16","4.94","0","99.11","416.62","0","418.65","34.58","0"
"AGAW2","08W","INS","BOLD:AAA5299","Arthropoda","Insecta","Diptera","Chironomidae","Orthocladiinae","Cricotopus","Cricotopus laetus, Cricotopus relucens, Cricotopus trifasciatus, Cricotopus sp., Cricotopus sp. 3ES, Cricotopus sylvestris, Cricotopus glacialis","191","177","177","17","7.33","0","99.19","432.74","0","435.28","34.55","0"
"AGAW2","08W","INS","BOLD:AAA4555","Arthropoda","Insecta","Orthoptera","Acrididae","Melanoplinae","Melanoplus","Melanoplus stupefactus, Melanoplus angustipennis, Melanoplus ponderosus, Melanoplus bruneri, Melanoplus frigidus, Melanoplus scapularis, Melanoplus cinereus, Melanoplus madeleineae, Melanoplus sanguinipes, Melanoplus nigrescens group, Melanoplus tunicae, Melanoplus walshi, Melanoplus femurrubrum, Melanoplus dawsoni, Melanoplus sp., Melanoplus fasciatus, Melanoplus bowditchi, Melanoplus stonei, Melanoplus bivittatus, Melanoplus huroni, Melanoplus thomasi, Melanoplus borealis, Melanoplus packardii","151","149","149","18","1.32","0","98.98","433.07","0","434.7","36.21","0"
"AGAW2","08W","INS","BOLD:AAN5199","Arthropoda","Insecta","Diptera","Cecidomyiidae","","","CecidInt20 sp. BOLD:AAN5199","142","107","107","19","24.65","0","99.62","432.29","0","446.49","26.84","0"
"AGAW2","08W","INS","BOLD:ADF5285","Arthropoda","Insecta","Hymenoptera","Braconidae","Opiinae","Opius","","100","74","74","20","26","0","98.58","388.53","0","447.21","25.62","0"
"AGAW2","08W","INS","BOLD:ACG3646","Arthropoda","Insecta","Coleoptera","Corylophidae","Corylophinae","Orthoperus","","90","68","68","21","24.44","0","98.37","446.14","0","447.28","34.15","0"
"AGAW2","08W","INS","BOLD:ACK4716","Arthropoda","Insecta","Diptera","Cecidomyiidae","","","","86","81","81","22","5.81","0","99.18","433","0","434.44","31.59","0"
"AGAW2","08W","INS","BOLD:ACK3431","Arthropoda","Insecta","Diptera","Cecidomyiidae","","","","79","77","77","23","2.53","0","98.98","421.35","0","423.04","32.72","0"
"AGAW2","08W","INS","BOLD:AAK8751","Arthropoda","Insecta","Diptera","Agromyzidae","Phytomyzinae","Liriomyza","Liriomyza sp. SB, Liriomyza trifoliearum","72","70","70","24","2.78","0","98.66","432.9","0","434.97","32.8","0"
"AGAW2","08W","INS","BOLD:AAG1741","Arthropoda","Insecta","Diptera","Muscidae","Coenosiinae","Coenosia","","64","59","59","25","7.81","0","99.24","447.16","0","448.42","33.62","0"
"AGAW2","08W","INS","BOLD:AAA9422","Arthropoda","Insecta","Hemiptera","Cicadellidae","Deltocephalinae","Macrosteles","Macrosteles quadrilineatus","57","57","57","26","0","0","98.3","393.04","0","400.12","35.42","0"
"AGAW2","08W","INS","BOLD:AAI4303","Arthropoda","Insecta","Diptera","Chironomidae","Chironominae","Chironomus","Chironomus melanescens","53","53","53","27","0","0","98.05","421.15","0","424.38","33.83","0"
"AGAW2","08W","INS","BOLD:AAH0135","Arthropoda","Insecta","Coleoptera","Phalacridae","Phalacrinae","Acylomus","Acylomus pugetanus, Acylomus ergoti","52","51","51","28","1.92","0","99.26","437.06","0","439.52","31.63","0"
"AGAW2","08W","INS","BOLD:AAH0134","Arthropoda","Insecta","Coleoptera","Phalacridae","Phalacrinae","Stilbus","Stilbus apicialis","51","49","49","29","3.92","0","99.4","426.25","0","427.25","31.97","0"
"AGAW2","08W","INS","BOLD:AAL4232","Arthropoda","Insecta","Diptera","Agromyzidae","Phytomyzinae","Phytoliriomyza","Phytoliriomyza arctica","42","33","33","30","21.43","0","99.63","441.48","0","443.02","32","0"
"AGAW2","08W","INS","BOLD:ACU7906","Arthropoda","Insecta","Diptera","Chironomidae","Orthocladiinae","","","36","36","36","31","0","0","98.8","413.36","0","415.5","35.62","0"
"AGAW2","08W","INS","BOLD:AAC4491","Arthropoda","Insecta","Diptera","Tipulidae","Tipulinae","Tipula","Tipula morrisoni, Tipula bicornis, Tipula johnsoniana, Tipula bicolor","33","33","33","32","0","0","99.08","437.73","0","440.03","33.95","0"
"AGAW2","08W","INS","BOLD:AAC8881","Arthropoda","Insecta","Lepidoptera","Gelechiidae","Gelechiinae","Syncopacma","Syncopacma palpilineella","32","27","27","33","15.63","0","99.56","410.94","0","446.34","33.2","0"
"AGAW2","08W","INS","BOLD:AAN5238","Arthropoda","Insecta","Diptera","Cecidomyiidae","","","","31","28","28","34","9.68","0","99.11","432.19","0","433.61","26.02","0"
"AGAW2","08W","INS","BOLD:AAB4894","Arthropoda","Insecta","Hemiptera","Aphididae","Aphidinae","Sitobion","Sitobion smilacifoliae, Sitobion avenae, Sitobion phyllanthi","31","31","31","34","0","0","97.81","439.48","0","440.42","25.44","0"
"AGAW2","08W","INS","BOLD:AAH8507","Arthropoda","Insecta","Hemiptera","Miridae","Phylinae","Phoenicocoris","Phoenicocoris strobicola","28","28","28","36","0","0","99.22","395.86","0","424.32","38.38","0"
"AGAW2","08W","INS","BOLD:ABY1179","Arthropoda","Insecta","Hymenoptera","Sierolomorphidae","","Sierolomorpha","","26","20","20","37","23.08","0","98.89","439.81","0","441.19","27.67","0"
"AGAW2","08W","INS","BOLD:AAG3154","Arthropoda","Insecta","Diptera","Psychodidae","Psychodinae","Psychoda","","26","24","24","37","7.69","0","98.41","417.77","0","436.5","32.57","0"
"AGAW2","08W","INS","BOLD:ABV1455","Arthropoda","Insecta","Diptera","Chironomidae","Chironominae","","","26","26","26","37","0","0","98.12","433.42","0","436.12","31.56","0"
"AGAW2","08W","INS","BOLD:ABA0863","Arthropoda","Insecta","Diptera","Cecidomyiidae","","","","25","24","24","40","4","0","99.37","438.52","0","439.96","29.75","0"
"AGAW2","08W","INS","BOLD:AAL7359","Arthropoda","Insecta","Diptera","Chironomidae","Chironominae","Tanytarsus","","24","24","24","41","0","0","98.82","436.5","0","437.92","36.05","0"
"AGAW2","08W","INS","BOLD:AAJ1250","Arthropoda","Insecta","Hymenoptera","Braconidae","Aphidiinae","Aphidius","Aphidius avenaphis, Aphidius uzbekistanicus","24","24","24","41","0","0","99.47","427","0","435.58","26.8","0"
"AGAW2","08W","INS","BOLD:ADF6661","Arthropoda","Insecta","Diptera","Cecidomyiidae","","","","22","18","18","43","18.18","0","99.19","413.23","0","438.59","29.51","0"
"AGAW2","08W","INS","BOLD:AAG7891","Arthropoda","Insecta","Hymenoptera","Platygastridae","Scelioninae","Telenomus","Telenomus podisi","21","21","21","44","0","0","99","426.76","0","435.81","27.02","0"
"AGAW2","08W","INS","BOLD:AAD7604","Arthropoda","Insecta","Coleoptera","Coccinellidae","Coccinellinae","Coleomegilla","Coleomegilla maculata","13","12","12","45","7.69","0","99.01","436.31","0","438.69","31.79","0"
"AGAW2","08W","INS","BOLD:AAV1136","Arthropoda","Insecta","Diptera","Scatopsidae","","","","12","12","12","46","0","0","98.67","414.42","0","416.5","36.63","0"
"AGAW2","08W","INS","BOLD:ABA7871","Arthropoda","Insecta","Diptera","Cecidomyiidae","","","","10","10","10","47","0","0","99.26","436.7","0","437.5","26.73","0"
"AGAW2","08W","INS","BOLD:AAN5255","Arthropoda","Insecta","Diptera","Cecidomyiidae","Cecidomyiinae","Dasineura","","9","9","9","48","0","0","99.12","417.44","0","447.56","26.88","0"
"AGAW2","08W","INS","BOLD:AAH3661","Arthropoda","Insecta","Diptera","Cecidomyiidae","","","","9","9","9","48","0","0","99.09","433.78","0","435.67","25.28","0"
"AGAW2","08W","INS","BOLD:AAH0256","Arthropoda","Insecta","Coleoptera","Latridiidae","Corticariinae","Corticaria, Corticarina","","7","7","7","50","0","0","98.66","430.43","0","436","31.68","0"
"AGAW2","08W","INS","BOLD:AAA6560","Arthropoda","Insecta","Lepidoptera","Crambidae","Crambinae","Crambus","Crambus multilinellus, Crambus saltuellus","6","6","6","51","0","0","99.68","441.83","0","441.83","30.84","0"
"AGAW2","08W","INS","BOLD:AAM7855","Arthropoda","Insecta","Thysanoptera","Thripidae","Thripinae","Frankliniella","Frankliniella tritici","5","5","5","52","0","0","99.82","448","0","450","31.51","0"
"AGAW2","08W","INS","BOLD:AAD2548","Arthropoda","Insecta","Hemiptera","Aphididae","Aphidinae","Acyrthosiphon","Acyrthosiphon caraganae, Acyrthosiphon sp. 1, Acyrthosiphon kondoi, Acyrthosiphon churchillense","5","5","5","52","0","0","99.22","445.2","0","450","25.91","0"
"AGAW2","08W","INS","BOLD:ACE4701","Arthropoda","Insecta","Lepidoptera","Crambidae","Crambinae","Crambus","Crambus multilinellus","5","5","5","52","0","0","98.96","426.6","0","427.8","31.59","0"
"AGAW2","08W","INS","BOLD:ACE7444","Arthropoda","Insecta","Hemiptera","Miridae","Mirinae, Phylinae","Adelphocoris, Megalocoleus","Adelphocoris lineolatus, Megalocoleus molliculus","4","4","4","55","0","0","99.78","444.5","0","445","35","0"
"AGAW2","08W","INS","BOLD:AAU8347","Arthropoda","Insecta","Hymenoptera","Platygastridae","Scelioninae","Telenomus","","4","4","4","55","0","0","100","403.5","0","450","25.83","0"
"AGAW2","08W","INS","BOLD:AAG2740","Arthropoda","Insecta","Diptera","Ephydridae","Parydrinae","Philygria","Philygria debilis","4","4","4","55","0","0","98.63","390.5","0","393.5","32.09","0"
"AGAW2","08W","INS","BOLD:AAP8739","Arthropoda","Insecta","Diptera","Platystomatidae","Platystomatinae","Rivellia","Rivellia steyskali","4","4","4","55","0","0","99.04","434.75","0","435.25","37.95","0"
"AGAW2","08W","INS","BOLD:AAA3181","Arthropoda","Insecta","Diptera","Muscidae","Muscinae","Stomoxys","Stomoxys calcitrans","3","3","3","59","0","0","97.91","435","0","437.33","32.92","0"
"AGAW2","08W","INS","BOLD:AAG1717","Arthropoda","Insecta","Diptera","Muscidae","Phaoniinae","Helina","Helina troene","3","3","3","59","0","0","99.39","438.33","0","439","30.89","0"
"AGAW2","08W","INS","BOLD:AAG0734","Arthropoda","Insecta","Thysanoptera","Thripidae","Thripinae","Frankliniella","Frankliniella tritici","3","3","3","59","0","0","98.29","448.33","0","450","30.22","0"
"AGAW2","08W","INS","BOLD:AAU6681","Arthropoda","Insecta","Diptera","Tipulidae","Tipulinae","Nephrotoma","Nephrotoma cornicina","3","3","3","59","0","0","99.04","449.33","0","450","36.15","0"
"AGAW2","08W","INS","BOLD:AAC0903","Arthropoda","Insecta","Diptera","Chironomidae","Chironominae","Chironomus","Chironomus acidophilus","3","3","3","59","0","0","99.69","428.33","0","429","36.28","0"
"AGAW2","08W","INS","BOLD:AAD2041","Arthropoda","Insecta","Diptera","Dolichopodidae","Sciapodinae","Condylostylus","Condylostylus inermis, Condylostylus nigrofemoratus, Condylostylus connectans, Condylostylus caudatus, Condylostylus flavipes","3","3","3","59","0","0","98.88","447.67","0","450","35.93","0"
"AGAW2","08W","INS","BOLD:AAA4277","Arthropoda","Insecta","Diptera","Syrphidae","Syrphinae","Toxomerus","Toxomerus marginatus, Toxomerus sp. CR-3, Toxomerus CR-3","3","3","3","59","0","0","99.18","449","0","450","30.81","0"
"AGAW2","08W","INS","BOLD:AAN5183","Arthropoda","Insecta","Diptera","Cecidomyiidae","","","","3","3","3","59","0","0","98.3","449.67","0","450","27.78","0"
"AGAW2","08W","INS","BOLD:AAG7276","Arthropoda","Insecta","Diptera","Sphaeroceridae","Limosininae","Leptocera","Leptocera erythrocera","2","2","2","67","0","0","97.33","396.5","0","397.5","33.46","0"
"AGAW2","08W","INS","BOLD:AAN5230","Arthropoda","Insecta","Diptera","Cecidomyiidae","","","","2","2","2","67","0","0","99","447","0","450","26.33","0"
"AGAW2","08W","INS","BOLD:AAY6412","Arthropoda","Insecta","Diptera","Sciaridae","","Bradysia","Bradysia urticae","2","2","2","67","0","0","100","448.5","0","450","32.33","0"
"AGAW2","08W","INS","BOLD:AAA3889","Arthropoda","Insecta","Lepidoptera","Crambidae","Crambinae","Parapediasia","Parapediasia teterrellus","2","2","2","67","0","0","99.16","431.5","0","432","31.68","0"
"AGAW2","08W","INS","BOLD:AAV5573","Arthropoda","Insecta","Diptera","Cecidomyiidae","","","","2","2","2","67","0","0","99.33","448.5","0","450","27.44","0"
"AGAW2","08W","INS","BOLD:AAA2815","Arthropoda","Insecta","Lepidoptera","Noctuidae","Noctuinae","Mythimna","Mythimna yukonensis, Mythimna oxygala","2","2","2","67","0","0","100","423","0","424.5","33.46","0"
"AGAW2","08W","INS","BOLD:AAF9187","Arthropoda","Insecta","Coleoptera","Curculionidae","Entiminae","Phyllobius","Phyllobius oblongus","2","2","2","67","0","0","99.88","434","0","434","38.81","0"
"AGAW2","08W","INS","BOLD:ACV2677","Arthropoda","Insecta","Diptera","Cecidomyiidae","","","","2","2","2","67","0","0","97.22","449","0","450","29.89","0"
"AGAW2","08W","INS","BOLD:AAC4197","Arthropoda","Insecta","Diptera","Chironomidae","Orthocladiinae","Paraphaenocladius","Paraphaenocladius impensus","2","2","2","67","0","0","99.78","449","0","450","35.33","0"
"AGAW2","08W","INS","BOLD:ADF3937","","","","","","","","2","2","2","67","0","0","98.01","428","0","450","32.11","0"
"AGAW2","08W","INS","BOLD:AAD8086","Arthropoda","Insecta","Lepidoptera","Elachistidae","Elachistinae","Elachista","Elachista griseicornis, Elachista acenteta","2","2","2","67","0","0","98.31","446","0","450","30.89","0"
"AGAW2","08W","INS","BOLD:AAG2460","Arthropoda","Insecta","Diptera","Anthomyiidae","Anthomyiinae","Hydrophoria","Hydrophoria lancifer","2","2","2","67","0","0","99.32","425","0","426.5","33.13","0"
"AGAW2","08W","INS","BOLD:ABW3687","Arthropoda","Insecta","Diptera","Lauxaniidae","Homoneurinae","Homoneura","Homoneura bispina","1","1","1","79","0","0","98.89","449","0","450","35.11","0"
"AGAW2","08W","INS","BOLD:AAA6292","Arthropoda","Insecta","Lepidoptera","Tortricidae","Tortricinae","Cnephasia","Cnephasia genitalana, Cnephasia stephensiana","1","1","1","79","0","0","99.78","450","0","450","36","0"
"AGAW2","08W","INS","BOLD:ABV1415","Arthropoda","Insecta","Diptera","Chironomidae","Chironominae","","","1","1","1","79","0","0","97.77","448","0","450","32.22","0"
"AGAW2","08W","INS","BOLD:AAA7067","Arthropoda","Insecta","Diptera","Culicidae","Culicinae","Aedes, Aedimorphus","Aedimorphus vexans, Aedes vexans","1","1","1","79","0","0","98.96","383","0","386","31.87","0"
"AGAW2","08W","INS","BOLD:AAH2306","Arthropoda","Insecta","Diptera","Asilidae","Asilinae","Efferia","Efferia aestuans","1","1","1","79","0","0","98.22","450","0","450","38","0"
"AGAW2","08W","INS","BOLD:AAM6780","Arthropoda","Insecta","Diptera","Dolichopodidae","Sympycninae","","","1","1","1","79","0","0","97.19","392","0","398","31.16","0"
"AGAW2","08W","INS","BOLD:AAH4135","Arthropoda","Insecta","Diptera","Chloropidae","Chloropinae","Thaumatomyia","Thaumatomyia glabra, Thaumatomyia grata, Thaumatomyia sp. 1 TAW","1","1","1","79","0","0","99","399","0","399","31.08","0"
"AGAW2","08W","INS","BOLD:AAB4657","Arthropoda","Insecta","Diptera","Chironomidae","Chironominae","Chironomus","Chironomus maturus","1","1","1","79","0","0","98.44","450","0","450","38","0"
"AGAW2","08W","INS","BOLD:ACL1796","Arthropoda","Insecta","Lepidoptera","Ypsolophidae","Ochsenheimeriinae","Ochsenheimeria","Ochsenheimeria vacculella","1","1","1","79","0","0","99.74","389","0","423","30.73","0"
"AGAW2","08W","INS","BOLD:AAE7186","Arthropoda","Insecta","Hymenoptera","Braconidae","Aphidiinae","Praon","Praon sp. 1 MM-2013, Praon volucre, Praon bicolor","1","1","1","79","0","0","100","450","0","450","25.78","0"
"AGAW2","08W","INS","BOLD:AAH3914","Arthropoda","Insecta","Diptera","Sciaridae","","Sciara","Sciara hamatilis, Sciara humeralis","1","1","1","79","0","0","98.87","444","0","450","30.89","0"
"AGAW2","08W","INS","BOLD:AAN5332","Arthropoda","Insecta","Diptera","Chironomidae","Chironominae","Polypedilum","","1","1","1","79","0","0","97.77","449","0","450","34.89","0"
"AGAW2","08W","INS","BOLD:AAH2293","Arthropoda","Insecta","Diptera","Asilidae","Stenopogoninae","Dioctria, Eudioctria","Dioctria hyalipennis, Dioctria baumhaueri, Eudioctria propinqua","1","1","1","79","0","0","100","448","0","450","30.44","0"
"AGAW2","08W","INS","BOLD:AAF9393","Arthropoda","Insecta","Coleoptera","Meloidae","Meloinae","Epicauta","Epicauta fabricii","1","1","1","79","0","0","97.18","390","0","391","40.66","0"
"AGAW2","08W","INS","BOLD:AAH3662","Arthropoda","Insecta","Diptera","Cecidomyiidae","","","CecidInt3 sp. BOLD:AAH3662","1","1","1","79","0","0","99.76","422","0","424","25.47","0"
"AGAW2","08W","INS","BOLD:AAB1098","Arthropoda","Insecta","Diptera","Culicidae","Culicinae","Ochlerotatus, Aedes","Aedes excrucians, Aedes cantans, Aedes stimulans, Aedes euedes, Aedes riparius, Aedes annulipes, Ochlerotatus cantans, Aedes fitchii, Aedes sp., Ochlerotatus annulipes, Ochlerotatus excrucians","1","1","1","79","0","0","99.11","449","0","450","35.78","0"
"AGAW2","08W","INS","BOLD:AAN5342","Arthropoda","Insecta","Diptera","Chironomidae","Orthocladiinae","Heleniella","","1","1","1","79","0","0","98.54","412","0","413","34.38","0"
"AGAW2","08W","INS","BOLD:AAM9013","Arthropoda","Insecta","Diptera","Keroplatidae","Macrocerinae","Macrocera","Macrocera pusilla","1","1","1","79","0","0","99.55","446","0","448","30.36","0"
"AGAW2","08W","INS","BOLD:ABV9290","Arthropoda","Insecta","Diptera","Cecidomyiidae","","","","1","1","1","79","0","0","99.33","448","0","450","25.78","0"
"AGAW2","08W","INS","BOLD:AAA3867","Arthropoda","Insecta","Lepidoptera","Noctuidae","Noctuinae","Apamea","Apamea amputatrix","1","1","1","79","0","0","99.1","444","0","450","33.11","0"
"AGAW2","08W","INS","BOLD:ABY5318","Arthropoda","Insecta","Diptera","Chloropidae","Oscinellinae","Incertella, Rhopalopterum","Rhopalopterum umbrosum, Rhopalopterum luteiceps, Incertella bispina, Rhopalopterum painteri","1","1","1","79","0","0","98.89","450","0","450","32.67","0"
"AGAW2","08W","INS","BOLD:AAE0845","Arthropoda","Insecta","Hemiptera","Pentatomidae","Pentatominae","Euschistus","Euschistus servus","1","1","1","79","0","0","99.1","446","0","450","37.78","0"
"AGAW2","08W","INS","BOLD:AAM6037","Arthropoda","Insecta","Diptera","Cecidomyiidae","Cecidomyiinae","Contarinia, Dasineura","","1","1","1","79","0","0","98.88","448","0","450","26.22","0"
"AGAW2","08W","INS","BOLD:AAN5267","Arthropoda","Insecta","Diptera","Cecidomyiidae","","","","1","1","1","79","0","0","100","450","0","450","30.67","0"
"AGAW2","08W","INS","BOLD:AAA3654","Arthropoda","Insecta","Lepidoptera","Crambidae","Crambinae","Crambus","Crambus bigelovi, Crambus johnsoni, Crambus praefectellus","1","1","1","79","0","0","98.89","450","0","450","31.33","0"
"AGAW2","08W","INS","BOLD:AAH7142","Arthropoda","Insecta","Diptera","Sarcophagidae","Sarcophaginae","Oxysarcodexia, Sarcophaginae_incertae_sedis","Oxysarcodexia cingarus, Sarcophaginae_incertae_sedis sp. 1 AGT, Oxysarcodexia ventricosa","1","1","1","79","0","0","99.33","450","0","450","34.22","0"
"AGAW2","08W","INS","BOLD:AAH4205","Arthropoda","Insecta","Diptera","Chloropidae","Chloropinae","Meromyza","Meromyza nigriseta, Meromyza sp.","1","1","1","79","0","0","98.22","450","0","450","33.56","0"
"AGAW2","08W","NAR","no match","no match","no match","no match","no match","no match","no match","no match","no match","no match","no match","no match","no match","no match","no match","no match","no match","no match","no match","no match"
"AGAW2","08W","NIN","BOLD:AAI4346","Arthropoda","Arachnida","Opiliones","Phalangiidae","","Paroligolophus, Phalangium","Paroligolophus agrestis, Phalangium opilio","1615","1424","1424","1","11.83","0","99.2","435.27","0","437.89","36.38","0"
"AGAW2","08W","NIN","BOLD:AAH6630","Arthropoda","Collembola","Entomobryomorpha","Entomobryidae","Entomobryinae","Entomobrya","Entomobrya unostrigata","1242","1038","1038","2","16.43","0","99.39","439.44","0","440.7","38.97","0"
"AGAW2","08W","NIN","BOLD:AAA5267","Arthropoda","Collembola","Entomobryomorpha","Isotomidae","","","","406","404","404","3","0.49","0","99","421.57","0","424.08","38.09","0"
"AGAW2","08W","NIN","BOLD:AAH0003","Arthropoda","Arachnida","Araneae","Linyphiidae","","Agyneta","Agyneta unimaculata","28","28","28","4","0","0","99.15","408.54","0","410.14","32.3","0"
"AGAW2","08W","NIN","BOLD:AAB7914","Arthropoda","Collembola","Symphypleona","Bourletiellidae","","Bourletiella","","28","26","26","4","7.14","0","99.64","448.04","0","450","38.18","0"
"AGAW2","08W","NIN","BOLD:ACC0359","Arthropoda","Collembola","Symphypleona","Bourletiellidae","","Bourletiella","","19","14","14","6","26.32","0","99.67","442.42","0","445.47","37.63","0"
"AGAW2","08W","NIN","BOLD:AAU6252","Arthropoda","Arachnida","Trombidiformes","Bdellidae","","Cyta","","16","15","15","7","6.25","0","99.17","423.31","0","424.06","31.3","0"
"AGAW2","08W","NIN","BOLD:ABX0027","Arthropoda","Collembola","Symphypleona","Bourletiellidae","","Bourletiella","","15","11","11","8","26.67","0","99.73","417.93","0","447.13","35.56","0"
"AGAW2","08W","NIN","BOLD:AAB7915","Arthropoda","Collembola","Symphypleona","Bourletiellidae","","Deuterosminthurus","Deuterosminthurus sp1","4","4","4","9","0","0","99.31","442","0","445.25","34.97","0"
"AGAW2","08W","NIN","BOLD:ACX8969","Arthropoda","Arachnida","Trombidiformes","Eupodidae","","","","4","4","4","9","0","0","98.53","401.25","0","405.5","27.21","0"
"AGAW2","09E","BAC","TAX:175768","Proteobacteria","Alphaproteobacteria","Rickettsiales","Rickettsiaceae",NA,"Wolbachia","Wolbachia pipientis","7","7","7","1","0","0","98.22","448.43","0","450","39.56","0"
"AGAW2","09E","BAC","TAX:91393","Proteobacteria","Alphaproteobacteria","Rickettsiales","Rickettsiaceae",NA,"Wolbachia","","6","6","6","2","0","0","99.19","392.17","0","412.33","38.84","0"
"AGAW2","09E","CHO","no match","no match","no match","no match","no match","no match","no match","no match","no match","no match","no match","no match","no match","no match","no match","no match","no match","no match","no match","no match"
"AGAW2","09E","INS","BOLD:AAA3453","Arthropoda","Insecta","Diptera","Anthomyiidae","Anthomyiinae","Delia","","19511","15951","15951","1","18.25","0","99.45","431.35","0","433.86","34.65","0"
"AGAW2","09E","INS","BOLD:AAG1711","Arthropoda","Insecta","Diptera","Muscidae","Phaoniinae","Helina","Helina depuncta","8707","6978","6978","2","19.86","0","99.43","432.83","0","435.12","31.37","0"
"AGAW2","09E","INS","BOLD:AAP8739","Arthropoda","Insecta","Diptera","Platystomatidae","Platystomatinae","Rivellia","Rivellia steyskali","6700","4449","4449","3","33.6","0","99.27","443.74","0","445.39","38.21","0"
"AGAW2","09E","INS","BOLD:AAA3867","Arthropoda","Insecta","Lepidoptera","Noctuidae","Noctuinae","Apamea","Apamea amputatrix","6427","4762","4762","4","25.91","0","99.32","440.76","0","442.53","33.72","0"
"AGAW2","09E","INS","BOLD:AAC9132","Arthropoda","Insecta","Diptera","Culicidae","Culicinae","Culiseta","Culiseta inornata","3804","3661","3661","5","3.76","0","98.54","401.3","0","404.1","32.41","0"
"AGAW2","09E","INS","BOLD:AAG6745","Arthropoda","Insecta","Diptera","Calliphoridae","Calliphorinae","Pollenia","Pollenia pediculata","3106","2848","2848","6","8.31","0","99.38","412.19","0","414.09","33.55","0"
"AGAW2","09E","INS","BOLD:AAN6154","Arthropoda","Insecta","Coleoptera","Latridiidae","Corticariinae","Melanophthalma","Melanophthalma inermis","2677","2098","2098","7","21.63","0","99.07","437.04","0","438.5","31.17","0"
"AGAW2","09E","INS","BOLD:AAD2041","Arthropoda","Insecta","Diptera","Dolichopodidae","Sciapodinae","Condylostylus","Condylostylus inermis, Condylostylus nigrofemoratus, Condylostylus connectans, Condylostylus caudatus, Condylostylus flavipes","2554","1781","1781","8","30.27","0","99.53","439.94","0","441.14","35.95","0"
"AGAW2","09E","INS","BOLD:AAB0373","Arthropoda","Insecta","Neuroptera","Chrysopidae","Chrysopinae","Chrysoperla","Chrysoperla sp. isolate 82, Chrysoperla zastrowi, Chrysoperla plorabunda, Chrysoperla sp. isolate 71, Chrysoperla agilis, Italochrysa italica, Chrysoperla sp. isolate 88, Chrysoperla lucasina, Chrysoperla carnea, Chrysoperla mediterranea, Chrysoperla pallida, Chrysoperla nipponensis, Chrysoperla sillemi","1786","1376","1376","9","22.96","0","99.52","435.96","0","436.84","31.33","0"
"AGAW2","09E","INS","BOLD:AAH4135","Arthropoda","Insecta","Diptera","Chloropidae","Chloropinae","Thaumatomyia","Thaumatomyia glabra, Thaumatomyia grata, Thaumatomyia sp. 1 TAW","1784","1669","1669","10","6.45","0","99.14","431.35","0","435.79","31.94","0"
"AGAW2","09E","INS","BOLD:AAH2306","Arthropoda","Insecta","Diptera","Asilidae","Asilinae","Efferia","Efferia aestuans","1277","1083","1083","11","15.19","0","99.24","435.52","0","436.88","37.96","0"
"AGAW2","09E","INS","BOLD:ABX9581","Arthropoda","Insecta","Hymenoptera","Chrysididae","Chrysidinae","Caenochrysis","","972","715","715","12","26.44","0","98.19","438.38","0","442.45","29.26","0"
"AGAW2","09E","INS","BOLD:ABY5318","Arthropoda","Insecta","Diptera","Chloropidae","Oscinellinae","Incertella, Rhopalopterum","Rhopalopterum umbrosum, Rhopalopterum luteiceps, Incertella bispina, Rhopalopterum painteri","749","601","601","13","19.76","0","99.03","440.53","0","441.54","32.28","0"
"AGAW2","09E","INS","BOLD:AAF9771","Arthropoda","Insecta","Diptera","Hybotidae","Tachydromiinae","Platypalpus","Platypalpus longiseta, Platypalpus trivialis, Platypalpus sp. 5, Platypalpus sp. 17, Platypalpus pallidiventris","675","505","505","14","25.19","0","99.27","442.54","0","443.35","34.9","0"
"AGAW2","09E","INS","BOLD:ACU7906","Arthropoda","Insecta","Diptera","Chironomidae","Orthocladiinae","","","579","575","575","15","0.69","0","98.9","402.61","0","411.04","36.23","0"
"AGAW2","09E","INS","BOLD:AAG5507","Arthropoda","Insecta","Diptera","Chironomidae","Chironominae","Chironomus","Chironomus atrella","572","564","564","16","1.4","0","98.83","420.09","0","427.84","36.02","0"
"AGAW2","09E","INS","BOLD:AAA5953","Arthropoda","Insecta","Lepidoptera","Gelechiidae","Gelechiinae","Coleotechnites","Coleotechnites sp., Coleotechnites piceaella, Coleotechnites laricis, Coleotechnites sp. 3, Coleotechnites carbonaria, Coleotechnites granti, Coleotechnites macleodi, Coleotechnites gibsonella, Coleotechnites coniferella, Coleotechnites martini","514","414","414","17","19.46","0","99.06","442.31","0","443.73","34.15","0"
"AGAW2","09E","INS","BOLD:AAA5299","Arthropoda","Insecta","Diptera","Chironomidae","Orthocladiinae","Cricotopus","Cricotopus laetus, Cricotopus relucens, Cricotopus trifasciatus, Cricotopus sp., Cricotopus sp. 3ES, Cricotopus sylvestris, Cricotopus glacialis","399","361","361","18","9.52","0","99.36","431.59","0","434.72","34.55","0"
"AGAW2","09E","INS","BOLD:AAG5457","Arthropoda","Insecta","Diptera","Chironomidae","Chironominae","","","350","342","342","19","2.29","0","98.91","431.72","0","433.98","34.44","0"
"AGAW2","09E","INS","BOLD:AAC4197","Arthropoda","Insecta","Diptera","Chironomidae","Orthocladiinae","Paraphaenocladius","Paraphaenocladius impensus","317","271","271","20","14.51","0","99.43","431.98","0","438.14","35.08","0"
"AGAW2","09E","INS","BOLD:AAB7030","Arthropoda","Insecta","Diptera","Chironomidae","Chironominae","Chironomus","Chironomus sp. BOLDAAP2996, Chironomus quinnitukqut, Chironomus cf. decorus 2g, Chironomus decorus group sp. 2 IP2013, Chironomus cf. decorus, Chironomus decorus gr.","304","301","301","21","0.99","0","98.54","427.04","0","433.15","34.52","0"
"AGAW2","09E","INS","BOLD:ACK1527","Arthropoda","Insecta","Diptera","Cecidomyiidae","","","","297","242","242","22","18.52","0","99.46","331.03","0","447.63","32.42","0"
"AGAW2","09E","INS","BOLD:AAN5199","Arthropoda","Insecta","Diptera","Cecidomyiidae","","","CecidInt20 sp. BOLD:AAN5199","234","181","181","23","22.65","0","99.56","439.93","0","446.72","26.66","0"
"AGAW2","09E","INS","BOLD:AAA7661","Arthropoda","Insecta","Diptera","Culicidae","Culicinae","Culiseta, Culex","Culiseta melanura, Culex articularis, Culex restuans, Culex acharistus","222","206","206","24","7.21","0","99.35","431.7","0","433.68","32.83","0"
"AGAW2","09E","INS","BOLD:ABA0822","Arthropoda","Insecta","Diptera","Cecidomyiidae","","","","213","197","197","25","7.51","0","99","422.34","0","423.47","25.67","0"
"AGAW2","09E","INS","BOLD:AAH3661","Arthropoda","Insecta","Diptera","Cecidomyiidae","","","","212","200","200","26","5.66","0","99.21","433.61","0","435.17","25.48","0"
"AGAW2","09E","INS","BOLD:ABW7897","Arthropoda","Insecta","Diptera","Cecidomyiidae","","","","204","154","154","27","24.51","0","99.67","440.71","0","442.14","29.65","0"
"AGAW2","09E","INS","BOLD:ACK3431","Arthropoda","Insecta","Diptera","Cecidomyiidae","","","","202","174","174","28","13.86","0","99.29","435.25","0","436.99","32.84","0"
"AGAW2","09E","INS","BOLD:AAL4232","Arthropoda","Insecta","Diptera","Agromyzidae","Phytomyzinae","Phytoliriomyza","Phytoliriomyza arctica","201","163","163","29","18.91","0","99.59","443.47","0","444.81","32.08","0"
"AGAW2","09E","INS","BOLD:AAN5332","Arthropoda","Insecta","Diptera","Chironomidae","Chironominae","Polypedilum","","193","166","166","30","13.99","0","98.11","440.48","0","441.72","34.43","0"
"AGAW2","09E","INS","BOLD:AAM9255","Arthropoda","Insecta","Diptera","Sciaridae","","Pseudolycoriella","","191","189","189","31","1.05","0","99.23","407.01","0","409.65","30.77","0"
"AGAW2","09E","INS","BOLD:AAH4171","Arthropoda","Insecta","Diptera","Chloropidae","Oscinellinae","Apallates","Apallates montanus, Apallates particeps","190","163","163","32","14.21","0","99.08","436.26","0","437.84","32.75","0"
"AGAW2","09E","INS","BOLD:ABV1297","Arthropoda","Insecta","Diptera","Sciaridae","","","","189","182","182","33","3.7","0","99.22","433.38","0","435.78","30.21","0"
"AGAW2","09E","INS","BOLD:AAN5342","Arthropoda","Insecta","Diptera","Chironomidae","Orthocladiinae","Heleniella","","178","165","165","34","7.3","0","98.89","425.72","0","431.76","35.17","0"
"AGAW2","09E","INS","BOLD:AAN6407","Arthropoda","Insecta","Diptera","Sphaeroceridae","Limosininae","Coproica","Coproica ferruginata","155","136","136","35","12.26","0","99.21","444.41","0","446.46","33.34","0"
"AGAW2","09E","INS","BOLD:ACX5521","Arthropoda","Insecta","Diptera","Chironomidae","Orthocladiinae","","","153","147","147","36","3.92","0","98.9","411.2","0","413.36","34.97","0"
"AGAW2","09E","INS","BOLD:ABZ6857","Arthropoda","Insecta","Hymenoptera","Braconidae","Microgastrinae","Microplitis","Microplitis jft46, Microplitis alaskensis","151","137","137","37","9.27","0","99.52","422.25","0","424.36","24.36","0"
"AGAW2","09E","INS","BOLD:AAH0135","Arthropoda","Insecta","Coleoptera","Phalacridae","Phalacrinae","Acylomus","Acylomus pugetanus, Acylomus ergoti","132","127","127","38","3.79","0","99.21","437.67","0","439.7","31.6","0"
"AGAW2","09E","INS","BOLD:AAL7359","Arthropoda","Insecta","Diptera","Chironomidae","Chironominae","Tanytarsus","","130","114","114","39","12.31","0","99.22","437","0","439.21","36.12","0"
"AGAW2","09E","INS","BOLD:ABZ4644","Arthropoda","Insecta","Diptera","Chloropidae","Oscinellinae","Malloewia","Malloewia nigripalpis","125","110","110","40","12","0","99.3","435.65","0","436.79","30.77","0"
"AGAW2","09E","INS","BOLD:AAD0484","Arthropoda","Insecta","Diptera","Chironomidae","","","","121","108","108","41","10.74","0","98.68","444.2","0","445.48","34.74","0"
"AGAW2","09E","INS","BOLD:ABU5525","Arthropoda","Insecta","Diptera","Chironomidae","Orthocladiinae","Limnophyes","Limnophyes sp. 14ES","117","116","116","42","0.85","0","98.88","404.8","0","408.09","37.24","0"
"AGAW2","09E","INS","BOLD:AAV5688","Arthropoda","Insecta","Diptera","Cecidomyiidae","","","","109","102","102","43","6.42","0","99.39","432.5","0","433.61","34.6","0"
"AGAW2","09E","INS","BOLD:AAN5358","Arthropoda","Insecta","Diptera","Chironomidae","Orthocladiinae","Smittia","Smittia stercoraria, Smittia sp. 22ES","89","78","78","44","12.36","0","99.57","442.33","0","444.12","37.47","0"
"AGAW2","09E","INS","BOLD:AAN5205","Arthropoda","Insecta","Diptera","Cecidomyiidae","","","CecidInt42 sp. BOLD:AAN5205","86","68","68","45","20.93","0","99.52","446.4","0","447.53","31.43","0"
"AGAW2","09E","INS","BOLD:AAH3664","Arthropoda","Insecta","Diptera","Cecidomyiidae","","","CecidInt3 sp. BOLD:AAH3664","83","82","82","46","1.2","0","99.04","428.37","0","431.54","26.94","0"
"AGAW2","09E","INS","BOLD:ABA6448","Arthropoda","Insecta","Diptera","Cecidomyiidae","","","","81","74","74","47","8.64","0","98.72","443.36","0","445.22","28.94","0"
"AGAW2","09E","INS","BOLD:AAN5183","Arthropoda","Insecta","Diptera","Cecidomyiidae","","","","76","66","66","48","13.16","0","98.33","440.66","0","441.75","27.74","0"
"AGAW2","09E","INS","BOLD:AAN6447","Arthropoda","Insecta","Diptera","Sciaridae","","Corynoptera","Corynoptera perpusilla","73","58","58","49","20.55","0","99.36","448.56","0","450","36.28","0"
"AGAW2","09E","INS","BOLD:AAN5238","Arthropoda","Insecta","Diptera","Cecidomyiidae","","","","69","55","55","50","20.29","0","99.35","433.3","0","434.94","25.67","0"
"AGAW2","09E","INS","BOLD:AAE3943","Arthropoda","Insecta","Hemiptera","Miridae","Phylinae","Plagiognathus","Plagiognathus chrysanthemi","66","64","64","51","3.03","0","99.52","390.98","0","448.29","36.31","0"
"AGAW2","09E","INS","BOLD:ABA0863","Arthropoda","Insecta","Diptera","Cecidomyiidae","","","","62","54","54","52","12.9","0","99.5","442.82","0","444.06","29.79","0"
"AGAW2","09E","INS","BOLD:AAU0577","Arthropoda","Insecta","Thysanoptera","Aeolothripidae","Aeolothripinae","Aeolothrips","Aeolothrips fasciatus","62","60","60","52","3.23","0","97.91","427.53","0","429.32","36.69","0"
"AGAW2","09E","INS","BOLD:ABV9277","","","","","","","","59","55","55","54","6.78","0","98.91","442.07","0","443.86","28.13","0"
"AGAW2","09E","INS","BOLD:ABV9319","Arthropoda","Insecta","Diptera","Chironomidae","Tanypodinae","Procladius","","50","46","46","55","8","0","99.23","437.34","0","439.58","34.79","0"
"AGAW2","09E","INS","BOLD:AAG7891","Arthropoda","Insecta","Hymenoptera","Platygastridae","Scelioninae","Telenomus","Telenomus podisi","50","40","40","55","20","0","98.85","426.76","0","435.08","27.17","0"
"AGAW2","09E","INS","BOLD:AAG6460","Arthropoda","Insecta","Diptera","Cecidomyiidae","","","","50","48","48","55","4","0","98.44","442.96","0","444.6","34.05","0"
"AGAW2","09E","INS","BOLD:AAN5267","Arthropoda","Insecta","Diptera","Cecidomyiidae","","","","49","39","39","58","20.41","0","99.34","447.76","0","448.51","30.65","0"
"AGAW2","09E","INS","BOLD:AAV6112","Arthropoda","Insecta","Diptera","Chloropidae","Oscinellinae","Oscinella","Oscinella sp. 2 AGT, Oscinella sp. TAW6","49","49","49","58","0","0","98.93","432.27","0","433.98","33.03","0"
"AGAW2","09E","INS","BOLD:ABA0858","Arthropoda","Insecta","Diptera","Cecidomyiidae","","","","44","40","40","60","9.09","0","99.45","432.16","0","432.95","26.06","0"
"AGAW2","09E","INS","BOLD:AAG6934","Arthropoda","Insecta","Diptera","Hybotidae","","","","44","40","40","60","9.09","0","99.45","432.52","0","434.59","38.78","0"
"AGAW2","09E","INS","BOLD:AAO8538","Arthropoda","Insecta","Diptera","Psychodidae","Psychodinae","Psychoda","","43","43","43","62","0","0","99.05","413.44","0","415.49","32.17","0"
"AGAW2","09E","INS","BOLD:ACK7314","Arthropoda","Insecta","Diptera","Chironomidae","Chironominae","Polypedilum","","41","41","41","63","0","0","98.96","389.12","0","395.05","35.21","0"
"AGAW2","09E","INS","BOLD:ADE0510","","","","","","","","38","38","38","64","0","0","97.52","385.37","0","413.55","35.92","0"
"AGAW2","09E","INS","BOLD:AAN5334","Arthropoda","Insecta","Diptera","Chironomidae","Chironominae","Polypedilum","","36","36","36","65","0","0","98.33","423.36","0","426.17","34.29","0"
"AGAW2","09E","INS","BOLD:AAD7604","Arthropoda","Insecta","Coleoptera","Coccinellidae","Coccinellinae","Coleomegilla","Coleomegilla maculata","35","35","35","66","0","0","98.99","437.77","0","440.17","31.79","0"
"AGAW2","09E","INS","BOLD:ABA7871","Arthropoda","Insecta","Diptera","Cecidomyiidae","","","","32","28","28","67","12.5","0","99.02","439.13","0","439.97","26.75","0"
"AGAW2","09E","INS","BOLD:AAM6037","Arthropoda","Insecta","Diptera","Cecidomyiidae","Cecidomyiinae","Contarinia, Dasineura","","32","26","26","67","18.75","0","98.69","444.56","0","445.47","26.39","0"
"AGAW2","09E","INS","BOLD:ACG3646","Arthropoda","Insecta","Coleoptera","Corylophidae","Corylophinae","Orthoperus","","31","26","26","69","16.13","0","99.11","443.13","0","444.48","33.45","0"
"AGAW2","09E","INS","BOLD:ACK1651","Arthropoda","Insecta","Diptera","Cecidomyiidae","","","","30","30","30","70","0","0","99.34","444","0","444.53","27.08","0"
"AGAW2","09E","INS","BOLD:AAD1485","Arthropoda","Insecta","Diptera","Chironomidae","Chironominae","Paratanytarsus","Paratanytarsus grimmii, Paratanytarsus sp.","30","24","24","70","20","0","99.49","442.83","0","444.07","33.78","0"
"AGAW2","09E","INS","BOLD:ACX6055","Arthropoda","Insecta","Diptera","Phoridae","Metopininae","Megaselia","Megaselia sp. AK-2010-2, Megaselia sp. BOLD:ACX6055","30","29","29","70","3.33","0","99.48","435.97","0","437.63","33.15","0"
"AGAW2","09E","INS","BOLD:AAV5594","Arthropoda","Insecta","Diptera","Cecidomyiidae","","","","29","26","26","73","10.34","0","99.34","443.93","0","444.93","26.43","0"
"AGAW2","09E","INS","BOLD:AAU8441","Arthropoda","Insecta","Hymenoptera","Ichneumonidae","Campopleginae","Meloboris","","27","24","24","74","11.11","0","99.77","445","0","446.04","25.45","0"
"AGAW2","09E","INS","BOLD:AAP6827","Arthropoda","Insecta","Diptera","Cecidomyiidae","","","","26","26","26","75","0","0","98.76","443.58","0","445.88","31.92","0"
"AGAW2","09E","INS","BOLD:AAH0256","Arthropoda","Insecta","Coleoptera","Latridiidae","Corticariinae","Corticaria, Corticarina","","26","25","25","75","3.85","0","98.62","416.38","0","426.04","31.75","0"
"AGAW2","09E","INS","BOLD:ACB9392","Arthropoda","Insecta","Diptera","Chironomidae","Chironominae","","","24","24","24","77","0","0","98.32","385.75","0","387.38","35.26","0"
"AGAW2","09E","INS","BOLD:AAH3662","Arthropoda","Insecta","Diptera","Cecidomyiidae","","","CecidInt3 sp. BOLD:AAH3662","20","20","20","78","0","0","98.4","431.1","0","432.2","25.98","0"
"AGAW2","09E","INS","BOLD:ACN7725","Arthropoda","Insecta","Hymenoptera","Platygastridae","","","","19","19","19","79","0","0","99.29","401.68","0","448.58","27.05","0"
"AGAW2","09E","INS","BOLD:AAP6790","Arthropoda","Insecta","Diptera","Chloropidae","Oscinellinae","Incertella","Incertella incerta","18","18","18","80","0","0","98.54","402.89","0","404.89","31.52","0"
"AGAW2","09E","INS","BOLD:AAG8091","Arthropoda","Insecta","Hymenoptera","Ichneumonidae","Campopleginae","Campoletis","Campoletis sp. AB.2","18","18","18","80","0","0","99.11","439.67","0","441.56","24.75","0"
"AGAW2","09E","INS","BOLD:AAH2876","Arthropoda","Insecta","Diptera","Cecidomyiidae","","","CecidInt51 sp. BOLD:AAH2876","18","16","16","80","11.11","0","99.23","439.39","0","441.17","26.26","0"
"AGAW2","09E","INS","BOLD:ACC8079","Arthropoda","Insecta","Diptera","Cecidomyiidae","","","","17","12","12","83","29.41","0","99.64","447.53","0","448.53","29.27","0"
"AGAW2","09E","INS","BOLD:ABV1375","Arthropoda","Insecta","Diptera","Cecidomyiidae","","","CecidInt3 sp. BOLD:ABV1375","16","16","16","84","0","0","98.97","439.19","0","440.13","26.44","0"
"AGAW2","09E","INS","BOLD:AAN5230","Arthropoda","Insecta","Diptera","Cecidomyiidae","","","","16","16","16","84","0","0","99.02","424.88","0","425.88","26.05","0"
"AGAW2","09E","INS","BOLD:AAP7026","Arthropoda","Insecta","Coleoptera","Latridiidae","Corticariinae","Melanophthalma","Melanophthalma villosa","15","12","12","86","20","0","97.74","405.33","0","438.33","31.34","0"
"AGAW2","09E","INS","BOLD:ADH9867","Arthropoda","Insecta","Diptera","","","","","15","15","15","86","0","0","98.69","403.27","0","406.13","37.23","0"
"AGAW2","09E","INS","BOLD:AAG2868","Arthropoda","Insecta","Hemiptera","Cicadellidae","Typhlocybinae","Empoasca","Empoasca sp., Empoasca fabae","14","12","12","88","14.29","0","99.56","429.57","0","443","32.28","0"
"AGAW2","09E","INS","BOLD:AAE7186","Arthropoda","Insecta","Hymenoptera","Braconidae","Aphidiinae","Praon","Praon sp. 1 MM-2013, Praon volucre, Praon bicolor","12","12","12","89","0","0","99.59","449.42","0","450","26.11","0"
"AGAW2","09E","INS","BOLD:AAG2511","Arthropoda","Insecta","Diptera","Anthomyiidae","Anthomyiinae","Delia","Delia platura","11","11","11","90","0","0","98.78","447.55","0","450","33.68","0"
"AGAW2","09E","INS","BOLD:AAC0903","Arthropoda","Insecta","Diptera","Chironomidae","Chironominae","Chironomus","Chironomus acidophilus","10","10","10","91","0","0","98.83","432.1","0","433","36.13","0"
"AGAW2","09E","INS","BOLD:ACX8294","Arthropoda","Insecta","Diptera","Cecidomyiidae","","","","9","9","9","92","0","0","99.72","445.33","0","446","32.33","0"
"AGAW2","09E","INS","BOLD:ADF6661","Arthropoda","Insecta","Diptera","Cecidomyiidae","","","","8","8","8","93","0","0","98.72","409.25","0","434.13","29.85","0"
"AGAW2","09E","INS","BOLD:AAN5220","Arthropoda","Insecta","Diptera","Cecidomyiidae","","","","7","7","7","94","0","0","97.85","444.29","0","447.29","26.67","0"
"AGAW2","09E","INS","BOLD:AAC0706","Arthropoda","Insecta","Diptera","Chironomidae","Chironominae","Dicrotendipes","Dicrotendipes tritomus","7","7","7","94","0","0","97.19","426.86","0","430","35.74","0"
"AGAW2","09E","INS","BOLD:AAN5242","Arthropoda","Insecta","Diptera","Cecidomyiidae","","","","6","6","6","96","0","0","99.42","405","0","406.83","28.25","0"
"AGAW2","09E","INS","BOLD:AAI4303","Arthropoda","Insecta","Diptera","Chironomidae","Chironominae","Chironomus","Chironomus melanescens","6","6","6","96","0","0","97.66","442","0","446.5","34.15","0"
"AGAW2","09E","INS","BOLD:AAN5255","Arthropoda","Insecta","Diptera","Cecidomyiidae","Cecidomyiinae","Dasineura","","6","6","6","96","0","0","98.79","424","0","449","26.91","0"
"AGAW2","09E","INS","BOLD:AAM7855","Arthropoda","Insecta","Thysanoptera","Thripidae","Thripinae","Frankliniella","Frankliniella tritici","6","5","5","96","16.67","0","99.42","443.83","0","445.83","31.13","0"
"AGAW2","09E","INS","BOLD:ABA0929","Arthropoda","Insecta","Diptera","Sciaridae","","Bradysia","Bradysia scabricornis","6","6","6","96","0","0","99.65","435.33","0","436.33","30.2","0"
"AGAW2","09E","INS","BOLD:ABV1292","Arthropoda","Insecta","Diptera","Cecidomyiidae","","","","6","5","5","96","16.67","0","99.7","438.33","0","439.5","28.29","0"
"AGAW2","09E","INS","BOLD:ABW1323","Arthropoda","Insecta","Diptera","Psychodidae","","","","5","5","5","102","0","0","98.91","361.2","0","366","29.4","0"
"AGAW2","09E","INS","BOLD:ACU5558","Arthropoda","Insecta","Hymenoptera","Bethylidae","","","","5","3","3","102","40","0","99.02","448.8","0","450","32.36","0"
"AGAW2","09E","INS","BOLD:AAV1136","Arthropoda","Insecta","Diptera","Scatopsidae","","","","5","5","5","102","0","0","99.22","415","0","416.8","36.33","0"
"AGAW2","09E","INS","BOLD:AAA4781","Arthropoda","Insecta","Hymenoptera","Braconidae","Microgastrinae","Glyptapanteles","Glyptapanteles carlossarmientoi","5","5","5","102","0","0","99.39","438","0","440.6","24.18","0"
"AGAW2","09E","INS","BOLD:AAA8933","Arthropoda","Insecta","Coleoptera","Coccinellidae","Coccinellinae","Coccinella","Coccinella septempunctata","4","4","4","106","0","0","98.77","446","0","450","32.72","0"
"AGAW2","09E","INS","BOLD:AAU6592","Arthropoda","Insecta","Diptera","Ceratopogonidae","Dasyheleinae","Dasyhelea","Dasyhelea arenivaga","4","4","4","106","0","0","99.67","389.5","0","390.75","34.56","0"
"AGAW2","09E","INS","BOLD:AAU6681","Arthropoda","Insecta","Diptera","Tipulidae","Tipulinae","Nephrotoma","Nephrotoma cornicina","4","4","4","106","0","0","99.66","441.25","0","442.25","36.18","0"
"AGAW2","09E","INS","BOLD:AAG2460","Arthropoda","Insecta","Diptera","Anthomyiidae","Anthomyiinae","Hydrophoria","Hydrophoria lancifer","4","4","4","106","0","0","98.57","430","0","431","33.21","0"
"AGAW2","09E","INS","BOLD:AAB0079","Arthropoda","Insecta","Diptera","Chironomidae","Orthocladiinae","Corynoneura","Corynoneura arctica","3","3","3","110","0","0","98.22","450.33","0","450","35.33","0"
"AGAW2","09E","INS","BOLD:AAY6412","Arthropoda","Insecta","Diptera","Sciaridae","","Bradysia","Bradysia urticae","3","3","3","110","0","0","99.4","437","0","440","31.65","0"
"AGAW2","09E","INS","BOLD:AAN5200","Arthropoda","Insecta","Diptera","Cecidomyiidae","","","","3","3","3","110","0","0","98.58","447.33","0","450","26.44","0"
"AGAW2","09E","INS","BOLD:AAA4188","Arthropoda","Insecta","Hymenoptera","Braconidae","Aphidiinae","Aphidius","Aphidius microlophii, Aphidius ervi","3","3","3","110","0","0","99.01","431","0","432.33","26.35","0"
"AGAW2","09E","INS","BOLD:AAV5796","Arthropoda","Insecta","Diptera","Cecidomyiidae","","","","3","3","3","110","0","0","99.25","430.33","0","432.33","29.1","0"
"AGAW2","09E","INS","BOLD:AAU6533","Arthropoda","Insecta","Diptera","Phoridae","Metopininae","Megaselia","","3","3","3","110","0","0","99.53","356","0","450","32.22","0"
"AGAW2","09E","INS","BOLD:AAG2740","Arthropoda","Insecta","Diptera","Ephydridae","Parydrinae","Philygria","Philygria debilis","3","3","3","110","0","0","98.28","405","0","408.33","32.6","0"
"AGAW2","09E","INS","BOLD:AAA6292","Arthropoda","Insecta","Lepidoptera","Tortricidae","Tortricinae","Cnephasia","Cnephasia genitalana, Cnephasia stephensiana","2","2","2","117","0","0","99.89","449","0","450","35.78","0"
"AGAW2","09E","INS","BOLD:AAG1714","Arthropoda","Insecta","Diptera","Muscidae","Azeliinae","Muscina","Muscina pascuorum","2","2","2","117","0","0","98.82","409.5","0","420","33.46","0"
"AGAW2","09E","INS","BOLD:ACA1917","Arthropoda","Insecta","Diptera","Cecidomyiidae","","","","2","2","2","117","0","0","99.18","425.5","0","427","25.14","0"
"AGAW2","09E","INS","BOLD:AAA6560","Arthropoda","Insecta","Lepidoptera","Crambidae","Crambinae","Crambus","Crambus multilinellus, Crambus saltuellus","2","2","2","117","0","0","99.32","441.5","0","442.5","31.41","0"
"AGAW2","09E","INS","BOLD:AAU8597","Arthropoda","Insecta","Hymenoptera","Platygastridae","","","","2","2","2","117","0","0","98.46","428","0","430.5","27.16","0"
"AGAW2","09E","INS","BOLD:AAG9626","Arthropoda","Insecta","Diptera","Dolichopodidae","","","","2","2","2","117","0","0","99.27","413.5","0","416","36.4","0"
"AGAW2","09E","INS","BOLD:AAF9306","Arthropoda","Insecta","Diptera","Psychodidae","Psychodinae","Psychoda","Psychoda alternata","2","2","2","117","0","0","98.11","448.5","0","450","31.11","0"
"AGAW2","09E","INS","BOLD:ABA5764","Arthropoda","Insecta","Hemiptera","Cicadellidae","Typhlocybinae","Empoasca","Empoasca coccinea","2","2","2","117","0","0","98.22","423","0","426","31.56","0"
"AGAW2","09E","INS","BOLD:ACV1014","Arthropoda","Insecta","Diptera","Cecidomyiidae","","","","2","2","2","117","0","0","98.13","376","0","380.5","25.37","0"
"AGAW2","09E","INS","BOLD:AAA4555","Arthropoda","Insecta","Orthoptera","Acrididae","Melanoplinae","Melanoplus","Melanoplus stupefactus, Melanoplus angustipennis, Melanoplus ponderosus, Melanoplus bruneri, Melanoplus frigidus, Melanoplus scapularis, Melanoplus cinereus, Melanoplus madeleineae, Melanoplus sanguinipes, Melanoplus nigrescens group, Melanoplus tunicae, Melanoplus walshi, Melanoplus femurrubrum, Melanoplus dawsoni, Melanoplus sp., Melanoplus fasciatus, Melanoplus bowditchi, Melanoplus stonei, Melanoplus bivittatus, Melanoplus huroni, Melanoplus thomasi, Melanoplus borealis, Melanoplus packardii","2","2","2","117","0","0","98.88","448.5","0","450","36.33","0"
"AGAW2","09E","INS","BOLD:AAE0845","Arthropoda","Insecta","Hemiptera","Pentatomidae","Pentatominae","Euschistus","Euschistus servus","2","2","2","117","0","0","98.74","413.5","0","412.5","38.22","0"
"AGAW2","09E","INS","BOLD:AAA3654","Arthropoda","Insecta","Lepidoptera","Crambidae","Crambinae","Crambus","Crambus bigelovi, Crambus johnsoni, Crambus praefectellus","2","2","2","117","0","0","98.89","449.5","0","450","32","0"
"AGAW2","09E","INS","BOLD:ADF5937","Arthropoda","Insecta","Diptera","Cecidomyiidae","","","","1","1","1","129","0","0","97.56","450","0","450","33.11","0"
"AGAW2","09E","INS","BOLD:AAN5162","Arthropoda","Insecta","Diptera","Ceratopogonidae","Dasyheleinae","Dasyhelea","","1","1","1","129","0","0","98","450","0","450","34.44","0"
"AGAW2","09E","INS","BOLD:AAA6759","Arthropoda","Insecta","Lepidoptera","Hesperiidae","Hesperiinae","Thymelicus","Thymelicus lineola","1","1","1","129","0","0","100","433","0","434","30.18","0"
"AGAW2","09E","INS","BOLD:AAP8651","Arthropoda","Insecta","Diptera","Tachinidae","Exoristinae","Oswaldia","Oswaldia albifacies","1","1","1","129","0","0","98.67","450","0","450","32.22","0"
"AGAW2","09E","INS","BOLD:AAQ0558","Arthropoda","Insecta","Thysanoptera","Thripidae","Thripinae","Anaphothrips","Anaphothrips obscurus","1","1","1","129","0","0","99.33","451","0","450","31.11","0"
"AGAW2","09E","INS","BOLD:ACJ6954","Arthropoda","Insecta","Diptera","Cecidomyiidae","","","","1","1","1","129","0","0","99.71","339","0","394","26.65","0"
"AGAW2","09E","INS","BOLD:ACE0699","Arthropoda","Insecta","Diptera","Chironomidae","","","","1","1","1","129","0","0","98.38","431","0","450","36.89","0"
"AGAW2","09E","INS","BOLD:AAG7276","Arthropoda","Insecta","Diptera","Sphaeroceridae","Limosininae","Leptocera","Leptocera erythrocera","1","1","1","129","0","0","99.78","450","0","450","32.89","0"
"AGAW2","09E","INS","BOLD:ABY9688","Arthropoda","Insecta","Diptera","Chloropidae","Chloropinae","Thaumatomyia","Thaumatomyia glabra","1","1","1","129","0","0","97.56","450","0","450","31.33","0"
"AGAW2","09E","INS","BOLD:AAA4213","Arthropoda","Insecta","Lepidoptera","Geometridae","Sterrhinae","Idaea","Idaea dimidiata","1","1","1","129","0","0","97.77","448","0","450","30.44","0"
"AGAW2","09E","INS","BOLD:AAC0635","Arthropoda","Insecta","Hemiptera","Miridae","Mirinae","Stenotus","Stenotus binotatus","1","1","1","129","0","0","98.88","447","0","450","35.11","0"
"AGAW2","09E","INS","BOLD:AAH0237","Arthropoda","Insecta","Coleoptera","Cleridae","Clerinae","Enoclerus","Enoclerus nigripes, Enoclerus rosmarus","1","1","1","129","0","0","97.09","447","0","450","32.44","0"
"AGAW2","09E","INS","BOLD:ACK3142","Arthropoda","Insecta","Diptera","Cecidomyiidae","","","","1","1","1","129","0","0","98.89","450","0","450","28.44","0"
"AGAW2","09E","INS","BOLD:AAG7036","Arthropoda","Insecta","Diptera","Agromyzidae","","","","1","1","1","129","0","0","99.22","386","0","386","33.68","0"
"AGAW2","09E","INS","BOLD:ADF9260","Arthropoda","Insecta","Coleoptera","Phalacridae","Phalacrinae","Acylomus","","1","1","1","129","0","0","98.66","449","0","450","31.33","0"
"AGAW2","09E","INS","BOLD:AAA2087","Arthropoda","Insecta","Lepidoptera","Erebidae","Herminiinae","Idia","Idia sp., Idia americalis","1","1","1","129","0","0","98.21","447","0","450","30.22","0"
"AGAW2","09E","INS","BOLD:AAF9187","Arthropoda","Insecta","Coleoptera","Curculionidae","Entiminae","Phyllobius","Phyllobius oblongus","1","1","1","129","0","0","97.56","450","0","450","39.33","0"
"AGAW2","09E","INS","BOLD:ABV1337","Arthropoda","Insecta","Diptera","Cecidomyiidae","","","","1","1","1","129","0","0","99.33","450","0","450","28.67","0"
"AGAW2","09E","INS","BOLD:ABX9225","Arthropoda","Insecta","Diptera","Limoniidae","Limoniinae","Rhipidia","","1","1","1","129","0","0","98.89","450","0","450","33.33","0"
"AGAW2","09E","INS","BOLD:AAM6229","Arthropoda","Insecta","Diptera","Chironomidae","Tanypodinae","","","1","1","1","129","0","0","98.44","450","0","450","33.33","0"
"AGAW2","09E","INS","BOLD:ABY5257","Arthropoda","Insecta","Lepidoptera","Noctuidae","Noctuinae","Apamea","Apamea devastator","1","1","1","129","0","0","97.98","445","0","450","31.33","0"
"AGAW2","09E","INS","BOLD:ACV2677","Arthropoda","Insecta","Diptera","Cecidomyiidae","","","","1","1","1","129","0","0","97.54","448","0","450","30","0"
"AGAW2","09E","INS","BOLD:AAD5425","Arthropoda","Insecta","Ephemeroptera","Baetidae","Baetinae","Callibaetis","Callibaetis fluctuans","1","1","1","129","0","0","99.11","450","0","450","40","0"
"AGAW2","09E","INS","BOLD:AAG1739","Arthropoda","Insecta","Diptera","Muscidae","Coenosiinae","Neodexiopsis","Neodexiopsis rufitibia","1","1","1","129","0","0","98.64","442","0","446","34.08","0"
"AGAW2","09E","INS","BOLD:ACD8563","Arthropoda","Insecta","Diptera","Cecidomyiidae","","","","1","1","1","129","0","0","97.11","450","0","450","25.78","0"
"AGAW2","09E","INS","BOLD:AAN8770","Arthropoda","Insecta","Diptera","Psychodidae","Psychodinae","Psychoda","Psychoda trinodulosa","1","1","1","129","0","0","100","448","0","450","34","0"
"AGAW2","09E","INS","BOLD:AAE9438","Arthropoda","Insecta","Hymenoptera","Ichneumonidae","Cryptinae","","","1","1","1","129","0","0","99.07","430","0","450","26.44","0"
"AGAW2","09E","INS","BOLD:ADF3937","","","","","","","","1","1","1","129","0","0","97.36","379","0","381","31.23","0"
"AGAW2","09E","INS","BOLD:AAG4944","Arthropoda","Insecta","Diptera","Mycetophilidae","Sciophilinae","Sciophila","Sciophila sp. CJB1","1","1","1","129","0","0","98.37","368","0","373","34.58","0"
"AGAW2","09E","INS","BOLD:AAC2637","Arthropoda","Insecta","Lepidoptera","Pterophoridae","Pterophorinae","Geina","Geina sheppardi","1","1","1","129","0","0","99.78","447","0","450","31.11","0"
"AGAW2","09E","INS","BOLD:AAV1295","Arthropoda","Insecta","Diptera","Sciaridae","","Bradysia","Bradysia difformis, Bradysia impatiens","1","1","1","129","0","0","99.53","423","0","450","31.56","0"
"AGAW2","09E","INS","BOLD:ACE4701","Arthropoda","Insecta","Lepidoptera","Crambidae","Crambinae","Crambus","Crambus multilinellus","1","1","1","129","0","0","98.19","443","0","450","31.33","0"
"AGAW2","09E","INS","BOLD:AAY6534","Arthropoda","Insecta","Coleoptera","Mordellidae","Mordellinae","Mordellina","Mordellina ancilla","1","1","1","129","0","0","99.11","450","0","450","39.11","0"
"AGAW2","09E","NAR","no match","no match","no match","no match","no match","no match","no match","no match","no match","no match","no match","no match","no match","no match","no match","no match","no match","no match","no match","no match"
"AGAW2","09E","NIN","BOLD:AAH6630","Arthropoda","Collembola","Entomobryomorpha","Entomobryidae","Entomobryinae","Entomobrya","Entomobrya unostrigata","834","725","725","1","13.07","0","99.25","438.44","0","439.59","39.2","0"
"AGAW2","09E","NIN","BOLD:ABW2639","Arthropoda","Arachnida","Trombidiformes","Anystidae","","","","421","326","326","2","22.57","0","99.54","447.15","0","446.31","31.47","0"
"AGAW2","09E","NIN","BOLD:AAA6381","Arthropoda","Arachnida","Araneae","Tetragnathidae","","Tetragnatha","Tetragnatha laboriosa","197","184","184","3","6.6","0","98.83","437.61","0","439.45","39.89","0"
"AGAW2","09E","NIN","BOLD:AAC6891","Arthropoda","Arachnida","Araneae","Salticidae","","Phidippus","Phidippus audax","113","88","88","4","22.12","0","99.73","402.02","0","444.27","32.04","0"
"AGAW2","09E","NIN","BOLD:AAA5267","Arthropoda","Collembola","Entomobryomorpha","Isotomidae","","","","70","70","70","5","0","0","98.98","414.99","0","417.84","38.08","0"
"AGAW2","09E","NIN","BOLD:ACZ1295","Arthropoda","Arachnida","Trombidiformes","Anystidae","","","","67","67","67","6","0","0","99.16","374.04","0","384.31","31.29","0"
"AGAW2","09E","NIN","BOLD:AAA8820","Arthropoda","Collembola","Entomobryomorpha","Entomobryidae","Lepidocyrtinae","Lepidocyrtus","Lepidocyrtus cyaneus","35","35","35","7","0","0","99.11","409.63","0","412.26","43.01","0"
"AGAW2","09E","NIN","BOLD:AAU6252","Arthropoda","Arachnida","Trombidiformes","Bdellidae","","Cyta","","26","24","24","8","7.69","0","99.49","435.73","0","436.46","31.65","0"
"AGAW2","09E","NIN","BOLD:AAA9476","Arthropoda","Arachnida","Araneae","Linyphiidae","","Grammonota","Grammonota gentilis","16","15","15","9","6.25","0","99.01","439.81","0","448.19","30.76","0"
"AGAW2","09E","NIN","BOLD:AAB7914","Arthropoda","Collembola","Symphypleona","Bourletiellidae","","Bourletiella","","5","4","4","10","20","0","99.58","443.2","0","443.6","38.09","0"
"AGAW2","09E","NIN","BOLD:AAI2833","Arthropoda","Arachnida","Araneae","Salticidae","","Phidippus","Phidippus princeps","3","3","3","11","0","0","97.25","448.67","0","450","32.59","0"
"AGAW2","09E","NIN","BOLD:AAB8452","Arthropoda","Collembola","Entomobryomorpha","Entomobryidae","Lepidocyrtinae","Lepidocyrtus","Lepidocyrtus paradoxus","3","3","3","11","0","0","99","419.67","0","419.67","42.35","0"
"AGAW2","09E","NIN","BOLD:AAI5320","Arthropoda","Arachnida","Araneae","Linyphiidae","","Islandiana","Islandiana flaveola","1","1","1","13","0","0","98.9","365","0","366","34.7","0"
"AGAW2","09E","NIN","BOLD:AAG5658","Arthropoda","Arachnida","Araneae","Mimetidae","","Mimetus","Mimetus epeiroides","1","1","1","13","0","0","99.78","449","0","450","35.11","0"
"AGAW2","09E","NIN","BOLD:ADD2059","Arthropoda","Arachnida","Mesostigmata","Phytoseiidae","","","","1","1","1","13","0","0","98.13","427","0","450","33.33","0"
"AGAW2","09N","BAC","TAX:175768","Proteobacteria","Alphaproteobacteria","Rickettsiales","Rickettsiaceae",NA,"Wolbachia","Wolbachia pipientis","46","37","37","1","19.57","0","97.94","448.17","0","449.57","39.49","0"
"AGAW2","09N","BAC","TAX:91393","Proteobacteria","Alphaproteobacteria","Rickettsiales","Rickettsiaceae",NA,"Wolbachia","","14","14","14","2","0","0","98.79","390.14","0","409.64","39.16","0"
"AGAW2","09N","BAC","TAX:91391","Proteobacteria","Alphaproteobacteria","Rickettsiales","",NA,"","","2","2","2","3","0","0","98.46","422","0","450","40","0"
"AGAW2","09N","BAC","TAX:703648","Proteobacteria","Alphaproteobacteria","Rickettsiales","Rickettsiaceae",NA,"Wolbachia","Wolbachia endosymbiont","1","1","1","4","0","0","97.22","395","0","450","39.56","0"
"AGAW2","09N","BAC","TAX:91389","Proteobacteria","","","",NA,"","","1","1","1","4","0","0","98.62","363","0","367","38.69","0"
"AGAW2","09N","CHO","no match","no match","no match","no match","no match","no match","no match","no match","no match","no match","no match","no match","no match","no match","no match","no match","no match","no match","no match","no match"
"AGAW2","09N","INS","BOLD:AAB9140","Arthropoda","Insecta","Diptera","Calliphoridae","Chrysomyinae","Phormia","Phormia regina","18288","10907","10907","1","40.36","0","99.6","443.51","0","444.87","33","0"
"AGAW2","09N","INS","BOLD:AAA3867","Arthropoda","Insecta","Lepidoptera","Noctuidae","Noctuinae","Apamea","Apamea amputatrix","15393","11914","11914","2","22.6","0","99.26","433.85","0","436.08","33.55","0"
"AGAW2","09N","INS","BOLD:AAC0903","Arthropoda","Insecta","Diptera","Chironomidae","Chironominae","Chironomus","Chironomus acidophilus","7749","5595","5595","3","27.8","0","99.5","438.37","0","440.91","36.3","0"
"AGAW2","09N","INS","BOLD:AAG6745","Arthropoda","Insecta","Diptera","Calliphoridae","Calliphorinae","Pollenia","Pollenia pediculata","5794","5136","5136","4","11.36","0","99.34","405.18","0","406.89","33.5","0"
"AGAW2","09N","INS","BOLD:AAG1711","Arthropoda","Insecta","Diptera","Muscidae","Phaoniinae","Helina","Helina depuncta","5612","4667","4667","5","16.84","0","99.32","432.32","0","434.62","31.36","0"
"AGAW2","09N","INS","BOLD:AAN6154","Arthropoda","Insecta","Coleoptera","Latridiidae","Corticariinae","Melanophthalma","Melanophthalma inermis","3970","3302","3302","6","16.83","0","98.77","438.24","0","439.63","31.32","0"
"AGAW2","09N","INS","BOLD:AAB0373","Arthropoda","Insecta","Neuroptera","Chrysopidae","Chrysopinae","Chrysoperla","Chrysoperla sp. isolate 82, Chrysoperla zastrowi, Chrysoperla plorabunda, Chrysoperla sp. isolate 71, Chrysoperla agilis, Italochrysa italica, Chrysoperla sp. isolate 88, Chrysoperla lucasina, Chrysoperla carnea, Chrysoperla mediterranea, Chrysoperla pallida, Chrysoperla nipponensis, Chrysoperla sillemi","3021","2306","2306","7","23.67","0","99.38","436.75","0","437.69","31.32","0"
"AGAW2","09N","INS","BOLD:AAA7374","Arthropoda","Insecta","Diptera","Syrphidae","Syrphinae","Sphaerophoria","Sphaerophoria bankowskae, Sphaerophoria infuscata, Sphaerophoria nr. philanthus, Sphaerophoria batava, Sphaerophoria sp., Sphaerophoria rueppellii, Sphaerophoria interrupta-group, Sphaerophoria sp. MA-1, Sphaerophoria cleoae, Sphaerophoria asymmetrica, Sphaerophoria fatarum, Sphaerophoria bengalensis, Sphaerophoria bifurcata, Sphaerophoria taeniata, Sphaerophoria interrupta, Sphaerophoria contigua, Sphaerophoria kaa, Sphaerophoria laurae, Sphaerophoria chongjini, Sphaerophoria boreoalpina, Sphaerophoria sp. BOLDAAA7374, Sphaerophoria macrogaster, Sphaerophoria sulphuripes, Sphaerophoria brevipilosa, Sphaerophoria philanthus, Sphaerophoria pyrrhina, Sphaerophoria abbreviata, Sphaerophoria scripta, Sphaerophoria potentillae, Sphaerophoria longipilosa, Sphaerophoria virgata","2934","2531","2531","8","13.74","0","99.29","428.68","0","432","30.72","0"
"AGAW2","09N","INS","BOLD:AAP8739","Arthropoda","Insecta","Diptera","Platystomatidae","Platystomatinae","Rivellia","Rivellia steyskali","2772","1959","1959","9","29.33","0","99.22","441.55","0","443.41","37.89","0"
"AGAW2","09N","INS","BOLD:ABY5257","Arthropoda","Insecta","Lepidoptera","Noctuidae","Noctuinae","Apamea","Apamea devastator","2625","1919","1919","10","26.9","0","99.04","443.83","0","445.37","30.69","0"
"AGAW2","09N","INS","BOLD:AAD5425","Arthropoda","Insecta","Ephemeroptera","Baetidae","Baetinae","Callibaetis","Callibaetis fluctuans","2488","2311","2311","11","7.11","0","98.71","398.11","0","400.32","39.58","0"
"AGAW2","09N","INS","BOLD:AAG2511","Arthropoda","Insecta","Diptera","Anthomyiidae","Anthomyiinae","Delia","Delia platura","1189","1065","1065","12","10.43","0","98.95","443.35","0","446.13","33.1","0"
"AGAW2","09N","INS","BOLD:AAB5609","Arthropoda","Insecta","Diptera","Muscidae","Coenosiinae","Coenosia","Coenosia sp.BOLD:AAB5609, Coenosia tigrina","1063","1014","1014","13","4.61","0","98.99","415.68","0","418.26","32.78","0"
"AGAW2","09N","INS","BOLD:AAA5299","Arthropoda","Insecta","Diptera","Chironomidae","Orthocladiinae","Cricotopus","Cricotopus laetus, Cricotopus relucens, Cricotopus trifasciatus, Cricotopus sp., Cricotopus sp. 3ES, Cricotopus sylvestris, Cricotopus glacialis","953","872","872","14","8.5","0","99.26","428.93","0","431.99","34.62","0"
"AGAW2","09N","INS","BOLD:AAA3453","Arthropoda","Insecta","Diptera","Anthomyiidae","Anthomyiinae","Delia","","917","855","855","15","6.76","0","99.29","430.86","0","433.63","34.68","0"
"AGAW2","09N","INS","BOLD:ABW7897","Arthropoda","Insecta","Diptera","Cecidomyiidae","","","","762","553","553","16","27.43","0","99.59","442.32","0","444","29.65","0"
"AGAW2","09N","INS","BOLD:AAA7067","Arthropoda","Insecta","Diptera","Culicidae","Culicinae","Aedes, Aedimorphus","Aedimorphus vexans, Aedes vexans","747","699","699","17","6.43","0","99.22","408.78","0","410.59","32.15","0"
"AGAW2","09N","INS","BOLD:AAB2384","Arthropoda","Insecta","Diptera","Syrphidae","Syrphinae","Eupeodes","Eupeodes duseki, Eupeodes volucris, Eupeodes nielseni, Eupeodes americanus, Eupeodes flukei, Eupeodes vockerothi, Eupeodes lucasi, Eupeodes montivagus, Eupeodes rufipunctatus, Eupeodes rojasi, Eupeodes goeldlini, Eupeodes luniger, Eupeodes punctifer, Eupeodes bucculatus, Eupeodes perplexus, Eupeodes abiskoensis, Eupeodes corollae, Eupeodes pomus, Eupeodes lundbecki, Eupeodes nitens, Eupeodes sp. BOLDAAB2384, Eupeodes curtus, Eupeodes latifasciatus","733","713","713","18","2.73","0","98.86","403.54","0","405.8","30.73","0"
"AGAW2","09N","INS","BOLD:AAF9949","Arthropoda","Insecta","Hemiptera","Miridae","Mirinae","Trigonotylus","Trigonotylus pulchellus, Trigonotylus americanus, Trigonotylus psammaecolor, Trigonotylus caelestialium, Trigonotylus viridis, Trigonotylus ruficornis","531","381","381","19","28.25","0","99.45","443","0","445.4","34.49","0"
"AGAW2","09N","INS","BOLD:AAD2041","Arthropoda","Insecta","Diptera","Dolichopodidae","Sciapodinae","Condylostylus","Condylostylus inermis, Condylostylus nigrofemoratus, Condylostylus connectans, Condylostylus caudatus, Condylostylus flavipes","512","387","387","20","24.41","0","99.44","440.85","0","442.08","35.97","0"
"AGAW2","09N","INS","BOLD:AAL4268","Arthropoda","Insecta","Diptera","Agromyzidae","Phytomyzinae","Phytomyza","Phytomyza flavofemorata, Phytomyza crassiseta","510","382","382","21","25.1","0","99.47","446.59","0","447.87","34.42","0"
"AGAW2","09N","INS","BOLD:AAH4135","Arthropoda","Insecta","Diptera","Chloropidae","Chloropinae","Thaumatomyia","Thaumatomyia glabra, Thaumatomyia grata, Thaumatomyia sp. 1 TAW","484","458","458","22","5.37","0","99.35","428.8","0","433.91","31.9","0"
"AGAW2","09N","INS","BOLD:ACX5521","Arthropoda","Insecta","Diptera","Chironomidae","Orthocladiinae","","","443","405","405","23","8.58","0","98.79","429.26","0","431.12","34.75","0"
"AGAW2","09N","INS","BOLD:AAF9771","Arthropoda","Insecta","Diptera","Hybotidae","Tachydromiinae","Platypalpus","Platypalpus longiseta, Platypalpus trivialis, Platypalpus sp. 5, Platypalpus sp. 17, Platypalpus pallidiventris","409","309","309","24","24.45","0","99.12","446.56","0","447.29","35.06","0"
"AGAW2","09N","INS","BOLD:AAG9668","Arthropoda","Insecta","Diptera","Dolichopodidae","Diaphorinae","Chrysotus","Chrysotus sp., Chrysotus sp. nr. obliquus","401","356","356","25","11.22","0","98.88","445.9","0","448.29","32.79","0"
"AGAW2","09N","INS","BOLD:AAG2868","Arthropoda","Insecta","Hemiptera","Cicadellidae","Typhlocybinae","Empoasca","Empoasca sp., Empoasca fabae","393","260","260","26","33.84","0","99.52","440.1","0","446.83","32.32","0"
"AGAW2","09N","INS","BOLD:ACK3431","Arthropoda","Insecta","Diptera","Cecidomyiidae","","","","368","352","352","27","4.35","0","99","424.82","0","426.51","32.65","0"
"AGAW2","09N","INS","BOLD:AAC4197","Arthropoda","Insecta","Diptera","Chironomidae","Orthocladiinae","Paraphaenocladius","Paraphaenocladius impensus","343","300","300","28","12.54","0","99.38","433.12","0","438.33","35.04","0"
"AGAW2","09N","INS","BOLD:AAF9028","Arthropoda","Insecta","Diptera","Limoniidae","","","","329","278","278","29","15.5","0","99.19","434.63","0","437.88","38.56","0"
"AGAW2","09N","INS","BOLD:AAG5453","Arthropoda","Insecta","Diptera","Chironomidae","Chironominae","Chironomus","Chironomus bifurcatus","315","261","261","30","17.14","0","98.31","440.24","0","444.28","34.89","0"
"AGAW2","09N","INS","BOLD:AAH0135","Arthropoda","Insecta","Coleoptera","Phalacridae","Phalacrinae","Acylomus","Acylomus pugetanus, Acylomus ergoti","305","276","276","31","9.51","0","99.29","434.97","0","436.9","31.61","0"
"AGAW2","09N","INS","BOLD:AAA7661","Arthropoda","Insecta","Diptera","Culicidae","Culicinae","Culiseta, Culex","Culiseta melanura, Culex articularis, Culex restuans, Culex acharistus","276","263","263","32","4.71","0","98.99","433.41","0","435.41","32.82","0"
"AGAW2","09N","INS","BOLD:AAD0484","Arthropoda","Insecta","Diptera","Chironomidae","","","","254","214","214","33","15.75","0","98.81","443.91","0","445.13","34.63","0"
"AGAW2","09N","INS","BOLD:ACU7906","Arthropoda","Insecta","Diptera","Chironomidae","Orthocladiinae","","","233","233","233","34","0","0","98.76","403.83","0","407.33","36.05","0"
"AGAW2","09N","INS","BOLD:AAN5161","Arthropoda","Insecta","Diptera","Ceratopogonidae","Dasyheleinae","Dasyhelea","","224","219","219","35","2.23","0","98.91","411.52","0","413.42","37.56","0"
"AGAW2","09N","INS","BOLD:AAB8013","Arthropoda","Insecta","Coleoptera","Coccinellidae","Coccinellinae","Hippodamia","Hippodamia expurgata, Hippodamia parenthesis, Hippodamia apicalis","200","172","172","36","14","0","99.41","445.56","0","445.5","34.89","0"
"AGAW2","09N","INS","BOLD:AAN5199","Arthropoda","Insecta","Diptera","Cecidomyiidae","","","CecidInt20 sp. BOLD:AAN5199","197","153","153","37","22.34","0","99.53","437.64","0","445.91","26.64","0"
"AGAW2","09N","INS","BOLD:AAG5507","Arthropoda","Insecta","Diptera","Chironomidae","Chironominae","Chironomus","Chironomus atrella","193","192","192","38","0.52","0","98.72","420.78","0","428.14","35.68","0"
"AGAW2","09N","INS","BOLD:AAN5342","Arthropoda","Insecta","Diptera","Chironomidae","Orthocladiinae","Heleniella","","164","161","161","39","1.83","0","98.92","417.01","0","425.01","35.07","0"
"AGAW2","09N","INS","BOLD:AAU6592","Arthropoda","Insecta","Diptera","Ceratopogonidae","Dasyheleinae","Dasyhelea","Dasyhelea arenivaga","152","148","148","40","2.63","0","99.06","402.63","0","404.7","34.69","0"
"AGAW2","09N","INS","BOLD:AAJ4275","Arthropoda","Insecta","Diptera","Chironomidae","Chironominae","Chironomus","Chironomus harpi","147","126","126","41","14.29","0","98.83","419.14","0","439.16","37.94","0"
"AGAW2","09N","INS","BOLD:AAD7604","Arthropoda","Insecta","Coleoptera","Coccinellidae","Coccinellinae","Coleomegilla","Coleomegilla maculata","142","132","132","42","7.04","0","98.86","429.85","0","432.43","31.71","0"
"AGAW2","09N","INS","BOLD:AAM9255","Arthropoda","Insecta","Diptera","Sciaridae","","Pseudolycoriella","","137","135","135","43","1.46","0","99.19","411.77","0","414.03","30.75","0"
"AGAW2","09N","INS","BOLD:AAA2409","Arthropoda","Insecta","Hymenoptera","Braconidae","Microgastrinae","Microplitis","Microplitis jft16","128","122","122","44","4.69","0","99.01","411.34","0","414.27","23.94","0"
"AGAW2","09N","INS","BOLD:AAN5267","Arthropoda","Insecta","Diptera","Cecidomyiidae","","","","119","90","90","45","24.37","0","99.58","448.39","0","449.05","30.6","0"
"AGAW2","09N","INS","BOLD:AAL4232","Arthropoda","Insecta","Diptera","Agromyzidae","Phytomyzinae","Phytoliriomyza","Phytoliriomyza arctica","116","95","95","46","18.1","0","99.35","440.38","0","441.86","32.23","0"
"AGAW2","09N","INS","BOLD:AAV5098","Arthropoda","Insecta","Diptera","Ceratopogonidae","Dasyheleinae","Dasyhelea","Dasyhelea turficola, Dasyhelea sp. 5ES","112","98","98","47","12.5","0","98.99","434.65","0","436.24","36.56","0"
"AGAW2","09N","INS","BOLD:AAV5157","","","","","","","","97","97","97","48","0","0","98.86","417.04","0","419.53","31.48","0"
"AGAW2","09N","INS","BOLD:ABW1379","Arthropoda","Insecta","Diptera","Chloropidae","Oscinellinae","Malloewia","Malloewia abdominalis","95","75","75","49","21.05","0","98.17","442.97","0","447.39","31.81","0"
"AGAW2","09N","INS","BOLD:AAG3154","Arthropoda","Insecta","Diptera","Psychodidae","Psychodinae","Psychoda","","83","65","65","50","21.69","0","98.72","443.71","0","445.93","31.82","0"
"AGAW2","09N","INS","BOLD:ABA0863","Arthropoda","Insecta","Diptera","Cecidomyiidae","","","","77","69","69","51","10.39","0","99.43","441.75","0","443.19","29.73","0"
"AGAW2","09N","INS","BOLD:ACK3589","Arthropoda","Insecta","Diptera","Cecidomyiidae","","","","77","65","65","51","15.58","0","98.85","448.78","0","449.78","32.69","0"
"AGAW2","09N","INS","BOLD:ABV9277","","","","","","","","73","67","67","53","8.22","0","99.2","437.01","0","438.44","28.11","0"
"AGAW2","09N","INS","BOLD:ACL8053","Arthropoda","Insecta","Diptera","Chironomidae","Chironominae","Tanytarsus","","69","68","68","54","1.45","0","99.25","444.07","0","445.59","33.53","0"
"AGAW2","09N","INS","BOLD:AAH3661","Arthropoda","Insecta","Diptera","Cecidomyiidae","","","","62","61","61","55","1.61","0","99.25","427.5","0","430.79","25.27","0"
"AGAW2","09N","INS","BOLD:ACK7314","Arthropoda","Insecta","Diptera","Chironomidae","Chironominae","Polypedilum","","57","57","57","56","0","0","98.8","395.81","0","398.91","35.24","0"
"AGAW2","09N","INS","BOLD:ABV1297","Arthropoda","Insecta","Diptera","Sciaridae","","","","57","57","57","56","0","0","99","433.6","0","435.91","30.23","0"
"AGAW2","09N","INS","BOLD:AAH0134","Arthropoda","Insecta","Coleoptera","Phalacridae","Phalacrinae","Stilbus","Stilbus apicialis","56","56","56","58","0","0","99.34","429.59","0","430.86","32.06","0"
"AGAW2","09N","INS","BOLD:AAG4905","Arthropoda","Insecta","Diptera","Mycetophilidae","Mycetophilinae","Mycetophila","","54","51","51","59","5.56","0","99.24","427.06","0","428.96","35.25","0"
"AGAW2","09N","INS","BOLD:ABU5525","Arthropoda","Insecta","Diptera","Chironomidae","Orthocladiinae","Limnophyes","Limnophyes sp. 14ES","54","54","54","59","0","0","99.07","397.54","0","400.43","37.2","0"
"AGAW2","09N","INS","BOLD:AAD4214","Arthropoda","Insecta","Hymenoptera","Ichneumonidae","Diplazontinae","Diplazon","Diplazon laetatorius","54","48","48","59","11.11","0","99.43","445.02","0","446.5","25.89","0"
"AGAW2","09N","INS","BOLD:ABZ6857","Arthropoda","Insecta","Hymenoptera","Braconidae","Microgastrinae","Microplitis","Microplitis jft46, Microplitis alaskensis","53","50","50","62","5.66","0","99.38","426.08","0","428.45","24.39","0"
"AGAW2","09N","INS","BOLD:AAH3664","Arthropoda","Insecta","Diptera","Cecidomyiidae","","","CecidInt3 sp. BOLD:AAH3664","52","52","52","63","0","0","99.12","434.71","0","437.04","27.14","0"
"AGAW2","09N","INS","BOLD:ACG3646","Arthropoda","Insecta","Coleoptera","Corylophidae","Corylophinae","Orthoperus","","51","45","45","64","11.76","0","98.99","440.59","0","442.43","33.77","0"
"AGAW2","09N","INS","BOLD:AAA4555","Arthropoda","Insecta","Orthoptera","Acrididae","Melanoplinae","Melanoplus","Melanoplus stupefactus, Melanoplus angustipennis, Melanoplus ponderosus, Melanoplus bruneri, Melanoplus frigidus, Melanoplus scapularis, Melanoplus cinereus, Melanoplus madeleineae, Melanoplus sanguinipes, Melanoplus nigrescens group, Melanoplus tunicae, Melanoplus walshi, Melanoplus femurrubrum, Melanoplus dawsoni, Melanoplus sp., Melanoplus fasciatus, Melanoplus bowditchi, Melanoplus stonei, Melanoplus bivittatus, Melanoplus huroni, Melanoplus thomasi, Melanoplus borealis, Melanoplus packardii","50","49","49","65","2","0","99.08","443","0","444.52","36.02","0"
"AGAW2","09N","INS","BOLD:AAN6447","Arthropoda","Insecta","Diptera","Sciaridae","","Corynoptera","Corynoptera perpusilla","44","38","38","66","13.64","0","99.45","448.34","0","449.43","36.28","0"
"AGAW2","09N","INS","BOLD:AAV5594","Arthropoda","Insecta","Diptera","Cecidomyiidae","","","","42","37","37","67","11.9","0","99.3","440.71","0","442.07","26.26","0"
"AGAW2","09N","INS","BOLD:AAP7026","Arthropoda","Insecta","Coleoptera","Latridiidae","Corticariinae","Melanophthalma","Melanophthalma villosa","39","34","34","68","12.82","0","98.48","390.36","0","442.54","31.08","0"
"AGAW2","09N","INS","BOLD:AAN5183","Arthropoda","Insecta","Diptera","Cecidomyiidae","","","","39","37","37","68","5.13","0","98.41","444.85","0","445.97","27.85","0"
"AGAW2","09N","INS","BOLD:AAV5688","Arthropoda","Insecta","Diptera","Cecidomyiidae","","","","35","34","34","70","2.86","0","99.36","435","0","436.4","34.33","0"
"AGAW2","09N","INS","BOLD:AAC4525","Arthropoda","Insecta","Diptera","Chironomidae","Chironominae","Tanytarsus","Tanytarsus guerlus","35","33","33","70","5.71","0","99.6","439.8","0","440.54","33.5","0"
"AGAW2","09N","INS","BOLD:AAG7891","Arthropoda","Insecta","Hymenoptera","Platygastridae","Scelioninae","Telenomus","Telenomus podisi","34","32","32","72","5.88","0","99.2","425.85","0","434.82","27.16","0"
"AGAW2","09N","INS","BOLD:ABV1194","Arthropoda","Insecta","Diptera","Chironomidae","Chironominae","Micropsectra","","34","25","25","72","26.47","0","99.42","438.03","0","439.12","36.16","0"
"AGAW2","09N","INS","BOLD:AAE3943","Arthropoda","Insecta","Hemiptera","Miridae","Phylinae","Plagiognathus","Plagiognathus chrysanthemi","34","34","34","72","0","0","99.67","390.94","0","447.53","36.53","0"
"AGAW2","09N","INS","BOLD:AAG2740","Arthropoda","Insecta","Diptera","Ephydridae","Parydrinae","Philygria","Philygria debilis","33","33","33","75","0","0","98.53","400.97","0","403.64","32.55","0"
"AGAW2","09N","INS","BOLD:AAN5220","Arthropoda","Insecta","Diptera","Cecidomyiidae","","","","31","31","31","76","0","0","98.13","408.58","0","425.81","26.38","0"
"AGAW2","09N","INS","BOLD:AAI1549","Arthropoda","Insecta","Hymenoptera","Braconidae","Euphorinae","Meteorus","","30","22","22","77","26.67","0","98.32","439.2","0","439.87","30.66","0"
"AGAW2","09N","INS","BOLD:AAD0300","Arthropoda","Insecta","Diptera","Chironomidae","Chironominae","Stempellinella","Stempellinella fimbriata","29","27","27","78","6.9","0","97.39","444.97","0","446.31","32.81","0"
"AGAW2","09N","INS","BOLD:ACC8079","Arthropoda","Insecta","Diptera","Cecidomyiidae","","","","28","23","23","79","17.86","0","99.55","445.46","0","447.07","29.21","0"
"AGAW2","09N","INS","BOLD:AAN5332","Arthropoda","Insecta","Diptera","Chironomidae","Chironominae","Polypedilum","","24","24","24","80","0","0","97.82","408.21","0","410.96","33.5","0"
"AGAW2","09N","INS","BOLD:AAZ5592","Arthropoda","Insecta","Diptera","Psychodidae","Psychodinae","","","21","20","20","81","4.76","0","98.96","436.76","0","438.86","32.52","0"
"AGAW2","09N","INS","BOLD:AAV5796","Arthropoda","Insecta","Diptera","Cecidomyiidae","","","","21","21","21","81","0","0","99.39","445.71","0","447.62","29.43","0"
"AGAW2","09N","INS","BOLD:ABX8236","Arthropoda","Insecta","Diptera","Cecidomyiidae","","","","21","21","21","81","0","0","98.79","415.29","0","420.48","28.85","0"
"AGAW2","09N","INS","BOLD:AAN5205","Arthropoda","Insecta","Diptera","Cecidomyiidae","","","CecidInt42 sp. BOLD:AAN5205","19","19","19","84","0","0","99.39","432.37","0","438.63","31.08","0"
"AGAW2","09N","INS","BOLD:AAG6934","Arthropoda","Insecta","Diptera","Hybotidae","","","","19","19","19","84","0","0","98.78","418.79","0","420.68","38.06","0"
"AGAW2","09N","INS","BOLD:ADF6661","Arthropoda","Insecta","Diptera","Cecidomyiidae","","","","19","18","18","84","5.26","0","99.39","408.37","0","430.42","29.16","0"
"AGAW2","09N","INS","BOLD:AAN8770","Arthropoda","Insecta","Diptera","Psychodidae","Psychodinae","Psychoda","Psychoda trinodulosa","15","14","14","87","6.67","0","99.06","441.6","0","442.73","33.95","0"
"AGAW2","09N","INS","BOLD:ABV0255","Arthropoda","Insecta","Diptera","Chironomidae","Orthocladiinae","Limnophyes","","14","14","14","88","0","0","99.18","426.93","0","428.43","34.26","0"
"AGAW2","09N","INS","BOLD:ABY1179","Arthropoda","Insecta","Hymenoptera","Sierolomorphidae","","Sierolomorpha","","14","13","13","88","7.14","0","99.6","439.79","0","440.93","28.14","0"
"AGAW2","09N","INS","BOLD:ACI7380","Arthropoda","Insecta","Diptera","Cecidomyiidae","","","","14","14","14","88","0","0","98.7","444","0","446.71","24.92","0"
"AGAW2","09N","INS","BOLD:AAN5260","Arthropoda","Insecta","Diptera","Cecidomyiidae","","","","12","12","12","91","0","0","98.98","408.75","0","409.67","25.89","0"
"AGAW2","09N","INS","BOLD:AAG5506","Arthropoda","Insecta","Diptera","Chironomidae","Chironominae","","","11","11","11","92","0","0","98.1","445","0","446","36.09","0"
"AGAW2","09N","INS","BOLD:ACK4901","Arthropoda","Insecta","Diptera","Chironomidae","Chironominae","","","11","11","11","92","0","0","99.01","423.91","0","425.82","36","0"
"AGAW2","09N","INS","BOLD:ACJ0111","Arthropoda","Insecta","Hymenoptera","Platygastridae","Scelioninae","Telenomus","","11","9","9","92","18.18","0","98.57","416.18","0","423.64","28.34","0"
"AGAW2","09N","INS","BOLD:ACW5117","Arthropoda","Insecta","Diptera","Chironomidae","Orthocladiinae","Smittia","Smittia leucopogon","11","11","11","92","0","0","99.24","430.18","0","431.55","31.44","0"
"AGAW2","09N","INS","BOLD:AAH3662","Arthropoda","Insecta","Diptera","Cecidomyiidae","","","CecidInt3 sp. BOLD:AAH3662","11","11","11","92","0","0","98.84","439","0","439.73","26.17","0"
"AGAW2","09N","INS","BOLD:AAL7819","Arthropoda","Insecta","Diptera","Psychodidae","Psychodinae","Psychoda","Psychoda sp., Psychoda sp. BOLD:AAL7819","10","10","10","97","0","0","99.63","426.2","0","427.1","32.88","0"
"AGAW2","09N","INS","BOLD:ADH9867","Arthropoda","Insecta","Diptera","","","","","10","10","10","97","0","0","98.93","396.2","0","399.6","37.09","0"
"AGAW2","09N","INS","BOLD:AAD5253","Arthropoda","Insecta","Diptera","Chironomidae","Orthocladiinae","Thienemanniella","Thienemanniella xena, Thienemanniella vittata","8","8","8","99","0","0","98.77","446.38","0","447.13","34.05","0"
"AGAW2","09N","INS","BOLD:AAF9306","Arthropoda","Insecta","Diptera","Psychodidae","Psychodinae","Psychoda","Psychoda alternata","8","8","8","99","0","0","99.61","446.25","0","446.75","31.53","0"
"AGAW2","09N","INS","BOLD:AAM7726","Arthropoda","Insecta","Coleoptera","Curculionidae","Curculioninae","Orchestes","Orchestes steppensis, Orchestes alni","8","8","8","99","0","0","99.2","429.63","0","431.75","33.24","0"
"AGAW2","09N","INS","BOLD:AAG6436","Arthropoda","Insecta","Diptera","Ceratopogonidae","Ceratopogoninae","Culicoides","Culicoides travisi","7","5","5","102","28.57","0","98.06","442.14","0","450","36.35","0"
"AGAW2","09N","INS","BOLD:ABV1292","Arthropoda","Insecta","Diptera","Cecidomyiidae","","","","7","7","7","102","0","0","99.33","448.86","0","450","28.19","0"
"AGAW2","09N","INS","BOLD:ADF3937","","","","","","","","7","7","7","102","0","0","99.58","399.71","0","405.29","31.3","0"
"AGAW2","09N","INS","BOLD:AAG6460","Arthropoda","Insecta","Diptera","Cecidomyiidae","","","","7","7","7","102","0","0","98.16","433.29","0","435","34.44","0"
"AGAW2","09N","INS","BOLD:AAA1513","Arthropoda","Insecta","Lepidoptera","Plutellidae","Plutellinae","Plutella, Rhigognostis","Plutella xylostella, Rhigognostis sp.","7","7","7","102","0","0","99.16","444.71","0","445.43","32.23","0"
"AGAW2","09N","INS","BOLD:AAE7186","Arthropoda","Insecta","Hymenoptera","Braconidae","Aphidiinae","Praon","Praon sp. 1 MM-2013, Praon volucre, Praon bicolor","6","6","6","107","0","0","98.97","438","0","446","26.07","0"
"AGAW2","09N","INS","BOLD:AAJ1250","Arthropoda","Insecta","Hymenoptera","Braconidae","Aphidiinae","Aphidius","Aphidius avenaphis, Aphidius uzbekistanicus","6","6","6","107","0","0","99.49","432.83","0","438","26.66","0"
"AGAW2","09N","INS","BOLD:ACM7977","Arthropoda","Insecta","Diptera","Cecidomyiidae","","","","6","6","6","107","0","0","99.44","449.33","0","450","24.93","0"
"AGAW2","09N","INS","BOLD:AAY6412","Arthropoda","Insecta","Diptera","Sciaridae","","Bradysia","Bradysia urticae","5","5","5","110","0","0","99.1","445.2","0","448.8","32.26","0"
"AGAW2","09N","INS","BOLD:AAM7855","Arthropoda","Insecta","Thysanoptera","Thripidae","Thripinae","Frankliniella","Frankliniella tritici","5","5","5","110","0","0","99.15","448","0","450","31.33","0"
"AGAW2","09N","INS","BOLD:AAG8091","Arthropoda","Insecta","Hymenoptera","Ichneumonidae","Campopleginae","Campoletis","Campoletis sp. AB.2","5","5","5","110","0","0","99.68","445","0","446.6","24.9","0"
"AGAW2","09N","INS","BOLD:AAM6037","Arthropoda","Insecta","Diptera","Cecidomyiidae","Cecidomyiinae","Contarinia, Dasineura","","5","5","5","110","0","0","98.17","448.6","0","450","26.53","0"
"AGAW2","09N","INS","BOLD:AAI4303","Arthropoda","Insecta","Diptera","Chironomidae","Chironominae","Chironomus","Chironomus melanescens","5","5","5","110","0","0","98.58","424.6","0","429","33.54","0"
"AGAW2","09N","INS","BOLD:AAA4781","Arthropoda","Insecta","Hymenoptera","Braconidae","Microgastrinae","Glyptapanteles","Glyptapanteles carlossarmientoi","5","5","5","110","0","0","98.59","426.8","0","430.2","23.89","0"
"AGAW2","09N","INS","BOLD:AAG2460","Arthropoda","Insecta","Diptera","Anthomyiidae","Anthomyiinae","Hydrophoria","Hydrophoria lancifer","5","5","5","110","0","0","98.62","448.6","0","449","33.94","0"
"AGAW2","09N","INS","BOLD:AAO8538","Arthropoda","Insecta","Diptera","Psychodidae","Psychodinae","Psychoda","","4","4","4","117","0","0","98.97","433.25","0","434.25","32.39","0"
"AGAW2","09N","INS","BOLD:ABW5534","Arthropoda","Insecta","Diptera","Cecidomyiidae","","","","4","4","4","117","0","0","98.54","426.25","0","428","25.71","0"
"AGAW2","09N","INS","BOLD:AAD1485","Arthropoda","Insecta","Diptera","Chironomidae","Chironominae","Paratanytarsus","Paratanytarsus grimmii, Paratanytarsus sp.","4","4","4","117","0","0","99.35","425.5","0","427","33.82","0"
"AGAW2","09N","INS","BOLD:AAV1136","Arthropoda","Insecta","Diptera","Scatopsidae","","","","4","4","4","117","0","0","98.38","437","0","439.75","35.86","0"
"AGAW2","09N","INS","BOLD:AAL9076","Arthropoda","Insecta","Diptera","Phoridae","Metopininae","Megaselia","","3","3","3","121","0","0","99.17","443.33","0","446","32.89","0"
"AGAW2","09N","INS","BOLD:ABA1215","Arthropoda","Insecta","Diptera","Sciaridae","","Lycoriella","Lycoriella sativae","3","3","3","121","0","0","99.62","445.67","0","446.67","35.82","0"
"AGAW2","09N","INS","BOLD:ACV5271","Arthropoda","Insecta","Diptera","Cecidomyiidae","","","","3","3","3","121","0","0","99.78","449.67","0","450","28.44","0"
"AGAW2","09N","INS","BOLD:AAH0256","Arthropoda","Insecta","Coleoptera","Latridiidae","Corticariinae","Corticaria, Corticarina","","3","3","3","121","0","0","98.22","448.67","0","450","31.19","0"
"AGAW2","09N","INS","BOLD:AAG1714","Arthropoda","Insecta","Diptera","Muscidae","Azeliinae","Muscina","Muscina pascuorum","2","2","2","125","0","0","99.88","430.5","0","450","34.33","0"
"AGAW2","09N","INS","BOLD:AAG1717","Arthropoda","Insecta","Diptera","Muscidae","Phaoniinae","Helina","Helina troene","2","2","2","125","0","0","99.44","449","0","450","30.89","0"
"AGAW2","09N","INS","BOLD:AAQ0558","Arthropoda","Insecta","Thysanoptera","Thripidae","Thripinae","Anaphothrips","Anaphothrips obscurus","2","2","2","125","0","0","99.18","425.5","0","427","31.5","0"
"AGAW2","09N","INS","BOLD:ABX4079","Arthropoda","Insecta","Diptera","Cecidomyiidae","","","","2","2","2","125","0","0","98.56","450","0","450","28.56","0"
"AGAW2","09N","INS","BOLD:AAN5242","Arthropoda","Insecta","Diptera","Cecidomyiidae","","","","2","2","2","125","0","0","99.39","431","0","431.5","28.59","0"
"AGAW2","09N","INS","BOLD:ABV1279","Arthropoda","Insecta","Diptera","Cecidomyiidae","","","","2","2","2","125","0","0","99.2","440","0","442","28.73","0"
"AGAW2","09N","INS","BOLD:AAN5238","Arthropoda","Insecta","Diptera","Cecidomyiidae","","","","2","2","2","125","0","0","97.31","446.5","0","450","27.11","0"
"AGAW2","09N","INS","BOLD:AAG0734","Arthropoda","Insecta","Thysanoptera","Thripidae","Thripinae","Frankliniella","Frankliniella tritici","2","2","2","125","0","0","98.27","398.5","0","402.5","29.44","0"
"AGAW2","09N","INS","BOLD:AAU6681","Arthropoda","Insecta","Diptera","Tipulidae","Tipulinae","Nephrotoma","Nephrotoma cornicina","2","2","2","125","0","0","99.89","450","0","450","36.67","0"
"AGAW2","09N","INS","BOLD:AAV5575","Arthropoda","Insecta","Diptera","Cecidomyiidae","","","CecidInt30 sp. BOLD:AAV5575","2","1","1","125","50","0","97.56","450","0","450","26.89","0"
"AGAW2","09N","INS","BOLD:AAD4936","Arthropoda","Insecta","Lepidoptera","Gracillariidae","Gracillariinae","Caloptilia","Caloptilia rhoifoliella, Caloptilia sassafrasella","2","2","2","125","0","0","99.74","374.5","0","375","28.93","0"
"AGAW2","09N","INS","BOLD:ACK3142","Arthropoda","Insecta","Diptera","Cecidomyiidae","","","","2","2","2","125","0","0","98.78","372","0","374","26.86","0"
"AGAW2","09N","INS","BOLD:AAA2815","Arthropoda","Insecta","Lepidoptera","Noctuidae","Noctuinae","Mythimna","Mythimna yukonensis, Mythimna oxygala","2","2","2","125","0","0","99.55","449","0","450","33.78","0"
"AGAW2","09N","INS","BOLD:AAA4277","Arthropoda","Insecta","Diptera","Syrphidae","Syrphinae","Toxomerus","Toxomerus marginatus, Toxomerus sp. CR-3, Toxomerus CR-3","2","2","2","125","0","0","99.44","449.5","0","450","30.56","0"
"AGAW2","09N","INS","BOLD:AAF9187","Arthropoda","Insecta","Coleoptera","Curculionidae","Entiminae","Phyllobius","Phyllobius oblongus","2","2","2","125","0","0","99.22","448.5","0","450","39","0"
"AGAW2","09N","INS","BOLD:AAG6743","Arthropoda","Insecta","Diptera","Sarcophagidae","Sarcophaginae","Sarcophaga","Sarcophaga sp. 1 MG, Sarcophaga subvicina","2","2","2","125","0","0","99.62","424","0","426.5","34.3","0"
"AGAW2","09N","INS","BOLD:ABY5318","Arthropoda","Insecta","Diptera","Chloropidae","Oscinellinae","Incertella, Rhopalopterum","Rhopalopterum umbrosum, Rhopalopterum luteiceps, Incertella bispina, Rhopalopterum painteri","2","2","2","125","0","0","99.22","450","0","450","31.89","0"
"AGAW2","09N","INS","BOLD:AAA3654","Arthropoda","Insecta","Lepidoptera","Crambidae","Crambinae","Crambus","Crambus bigelovi, Crambus johnsoni, Crambus praefectellus","2","2","2","125","0","0","98.38","400","0","401","31.09","0"
"AGAW2","09N","INS","BOLD:AAA5953","Arthropoda","Insecta","Lepidoptera","Gelechiidae","Gelechiinae","Coleotechnites","Coleotechnites sp., Coleotechnites piceaella, Coleotechnites laricis, Coleotechnites sp. 3, Coleotechnites carbonaria, Coleotechnites granti, Coleotechnites macleodi, Coleotechnites gibsonella, Coleotechnites coniferella, Coleotechnites martini","2","2","2","125","0","0","99","402.5","0","408.5","32.33","0"
"AGAW2","09N","INS","BOLD:ACE6002","Arthropoda","Insecta","Lepidoptera","Noctuidae","Noctuinae","Apamea","Apamea contradicta","2","2","2","125","0","0","97.09","447.5","0","450","31.11","0"
"AGAW2","09N","INS","BOLD:AAA4222","Arthropoda","Insecta","Lepidoptera","Erebidae","Hypeninae","Hypena","Hypena scabra","1","1","1","145","0","0","98.21","446","0","450","32","0"
"AGAW2","09N","INS","BOLD:AAA3181","Arthropoda","Insecta","Diptera","Muscidae","Muscinae","Stomoxys","Stomoxys calcitrans","1","1","1","145","0","0","98.66","448","0","450","33.56","0"
"AGAW2","09N","INS","BOLD:ABW3687","Arthropoda","Insecta","Diptera","Lauxaniidae","Homoneurinae","Homoneura","Homoneura bispina","1","1","1","145","0","0","98","449","0","450","36","0"
"AGAW2","09N","INS","BOLD:AAP7027","Arthropoda","Insecta","Coleoptera","Curculionidae","Curculioninae","Tychius","Tychius picirostris","1","1","1","145","0","0","100","450","0","450","34","0"
"AGAW2","09N","INS","BOLD:AAN5497","Arthropoda","Insecta","Diptera","Anthomyiidae","Pegomyinae","Pegomya","Pegomya solennis, Pegomya bicolor","1","1","1","145","0","0","99.33","450","0","450","29.56","0"
"AGAW2","09N","INS","BOLD:AAG8839","Arthropoda","Insecta","Hemiptera","Cicadellidae","Deltocephalinae","Errastunus, Latalus","Latalus ocellaris, Errastunus ocellaris","1","1","1","145","0","0","100","449","0","450","36","0"
"AGAW2","09N","INS","BOLD:AAR9127","Arthropoda","Insecta","Hemiptera","Tingidae","Tinginae","Corythucha","Corythucha sp. HP28, Corythucha marmorata","1","1","1","145","0","0","99.56","450","0","450","32.89","0"
"AGAW2","09N","INS","BOLD:ADF3809","Arthropoda","Insecta","Diptera","Dolichopodidae","","","","1","1","1","145","0","0","97.9","429","0","432","35.19","0"
"AGAW2","09N","INS","BOLD:AAN5230","Arthropoda","Insecta","Diptera","Cecidomyiidae","","","","1","1","1","145","0","0","97.78","360","0","361","24.38","0"
"AGAW2","09N","INS","BOLD:ACK4716","Arthropoda","Insecta","Diptera","Cecidomyiidae","","","","1","1","1","145","0","0","100","401","0","405","30.86","0"
"AGAW2","09N","INS","BOLD:AAA3889","Arthropoda","Insecta","Lepidoptera","Crambidae","Crambinae","Parapediasia","Parapediasia teterrellus","1","1","1","145","0","0","98.58","351","0","351","31.62","0"
"AGAW2","09N","INS","BOLD:AAG0996","Arthropoda","Insecta","Diptera","Chironomidae","Orthocladiinae","Cricotopus","Cricotopus sp. 18ES, Cricotopus bicinctus gr., Cricotopus bicinctus grp.","1","1","1","145","0","0","99.01","404","0","405","31.85","0"
"AGAW2","09N","INS","BOLD:AAA2632","Arthropoda","Insecta","Lepidoptera","Noctuidae","Noctuinae","Noctua","Noctua atlantica, Noctua pronuba","1","1","1","145","0","0","99.55","448","0","450","31.11","0"
"AGAW2","09N","INS","BOLD:ABV1455","Arthropoda","Insecta","Diptera","Chironomidae","Chironominae","","","1","1","1","145","0","0","98","450","0","450","32.44","0"
"AGAW2","09N","INS","BOLD:AAH2293","Arthropoda","Insecta","Diptera","Asilidae","Stenopogoninae","Dioctria, Eudioctria","Dioctria hyalipennis, Dioctria baumhaueri, Eudioctria propinqua","1","1","1","145","0","0","99.1","444","0","450","30.44","0"
"AGAW2","09N","INS","BOLD:AAM6295","Arthropoda","Insecta","Diptera","Chaoboridae","","","","1","1","1","145","0","0","98.44","449","0","450","38","0"
"AGAW2","09N","INS","BOLD:AAP5087","Arthropoda","Insecta","Diptera","Ceratopogonidae","Dasyheleinae","Dasyhelea","","1","1","1","145","0","0","98.44","450","0","450","35.33","0"
"AGAW2","09N","INS","BOLD:AAG6753","Arthropoda","Insecta","Diptera","Lauxaniidae","Lauxaniinae","Camptoprosopella","","1","1","1","145","0","0","99.27","413","0","450","35.56","0"
"AGAW2","09N","INS","BOLD:AAC1312","Arthropoda","Insecta","Diptera","Syrphidae","Syrphinae","Toxomerus","Toxomerus geminatus, Toxomerus 75-5, Toxomerus rombicus, Toxomerus sp. 75-5","1","1","1","145","0","0","100","447","0","450","30","0"
"AGAW2","09N","INS","BOLD:AAG9626","Arthropoda","Insecta","Diptera","Dolichopodidae","","","","1","1","1","145","0","0","98.22","450","0","450","37.56","0"
"AGAW2","09N","INS","BOLD:AAV1261","Arthropoda","Insecta","Diptera","Sciaridae","","Bradysia","Bradysia splendida","1","1","1","145","0","0","100","448","0","450","34","0"
"AGAW2","09N","INS","BOLD:AAG5499","Arthropoda","Insecta","Diptera","Chironomidae","Orthocladiinae","Psectrocladius","Psectrocladius obvius, Psectrocladius platypus","1","1","1","145","0","0","98.29","409","0","411","35.28","0"
"AGAW2","09N","INS","BOLD:AAP9022","Arthropoda","Insecta","Diptera","Cecidomyiidae","","","","1","1","1","145","0","0","99.11","447","0","450","27.33","0"
"AGAW2","09N","INS","BOLD:ACF7023","Arthropoda","Insecta","Diptera","Dolichopodidae","","","","1","1","1","145","0","0","97.32","224","0","411","33.09","0"
"AGAW2","09N","INS","BOLD:AAE0845","Arthropoda","Insecta","Hemiptera","Pentatomidae","Pentatominae","Euschistus","Euschistus servus","1","1","1","145","0","0","98.45","386","0","390","37.69","0"
"AGAW2","09N","INS","BOLD:AAB3656","Arthropoda","Insecta","Lepidoptera","Pterophoridae","Pterophorinae","Hellinsia","Hellinsia glenni, Hellinsia lacteodactylus, Hellinsia paleaceus","1","1","1","145","0","0","97.77","448","0","450","33.11","0"
"AGAW2","09N","INS","BOLD:AAU6656","Arthropoda","Insecta","Diptera","Limoniidae","Chioneinae","Neolimnophila","Neolimnophila placida","1","1","1","145","0","0","97.76","401","0","403","34","0"
"AGAW2","09N","INS","BOLD:ABX2489","Arthropoda","Insecta","Coleoptera","Phalacridae","Phalacrinae","Stilbus","","1","1","1","145","0","0","97.19","392","0","392","32.4","0"
"AGAW2","09N","INS","BOLD:AAZ9744","Arthropoda","Insecta","Hymenoptera","Braconidae","Brachistinae","Blacus","Blacus stelfoxi","1","1","1","145","0","0","98.23","451","0","450","32","0"
"AGAW2","09N","INS","BOLD:ABZ4994","Arthropoda","Insecta","Orthoptera","Acrididae","Melanoplinae","Melanoplus","Melanoplus gladstoni, Melanoplus sanguinipes","1","1","1","145","0","0","97.09","446","0","450","37.11","0"
"AGAW2","09N","INS","BOLD:AAY6450","Arthropoda","Insecta","Diptera","Cecidomyiidae","","","","1","1","1","145","0","0","100","449","0","450","27.78","0"
"AGAW2","09N","INS","BOLD:ABX6186","Arthropoda","Insecta","Diptera","Tipulidae","Tipulinae","Nephrotoma","Nephrotoma ferruginea","1","1","1","145","0","0","98","450","0","450","34.89","0"
"AGAW2","09N","INS","BOLD:AAH3644","Arthropoda","Insecta","Diptera","Cecidomyiidae","","","","1","1","1","145","0","0","99.27","410","0","412","26.21","0"
"AGAW2","09N","INS","BOLD:AAH7146","Arthropoda","Insecta","Diptera","Sarcophagidae","Sarcophaginae","Boettcheria","Boettcheria latisterna","1","1","1","145","0","0","99.51","405","0","407","34.4","0"
"AGAW2","09N","INS","BOLD:AAA7368","Arthropoda","Insecta","Lepidoptera","Cosmopterigidae","Cosmopteriginae","Limnaecia","Limnaecia phragmitella","1","1","1","145","0","0","99.55","448","0","450","30.89","0"
"AGAW2","09N","INS","BOLD:AAP8651","Arthropoda","Insecta","Diptera","Tachinidae","Exoristinae","Oswaldia","Oswaldia albifacies","1","1","1","145","0","0","98.23","451","0","450","32.67","0"
"AGAW2","09N","INS","BOLD:AAC2637","Arthropoda","Insecta","Lepidoptera","Pterophoridae","Pterophorinae","Geina","Geina sheppardi","1","1","1","145","0","0","99.78","449","0","450","30.89","0"
"AGAW2","09N","INS","BOLD:ACE4701","Arthropoda","Insecta","Lepidoptera","Crambidae","Crambinae","Crambus","Crambus multilinellus","1","1","1","145","0","0","97.54","448","0","450","30.89","0"
"AGAW2","09N","NAR","no match","no match","no match","no match","no match","no match","no match","no match","no match","no match","no match","no match","no match","no match","no match","no match","no match","no match","no match","no match"
"AGAW2","09N","NIN","BOLD:AAH6630","Arthropoda","Collembola","Entomobryomorpha","Entomobryidae","Entomobryinae","Entomobrya","Entomobrya unostrigata","2043","1694","1694","1","17.08","0","99.28","438.41","0","439.63","39.09","0"
"AGAW2","09N","NIN","BOLD:AAA5267","Arthropoda","Collembola","Entomobryomorpha","Isotomidae","","","","665","660","660","2","0.75","0","99","419.4","0","421.82","38.12","0"
"AGAW2","09N","NIN","BOLD:AAI4346","Arthropoda","Arachnida","Opiliones","Phalangiidae","","Paroligolophus, Phalangium","Paroligolophus agrestis, Phalangium opilio","647","618","618","3","4.48","0","98.93","412.09","0","420.7","35.42","0"
"AGAW2","09N","NIN","BOLD:AAA6383","","","","","","","","598","493","493","4","17.56","0","98.61","436.71","0","437.9","37.79","0"
"AGAW2","09N","NIN","BOLD:ACR6860","","","","","","","","179","167","167","5","6.7","0","99.12","384.6","0","437.54","40.29","0"
"AGAW2","09N","NIN","BOLD:ABW2639","Arthropoda","Arachnida","Trombidiformes","Anystidae","","","","63","51","51","6","19.05","0","99.62","447.38","0","446.38","31.47","0"
"AGAW2","09N","NIN","BOLD:AAU6252","Arthropoda","Arachnida","Trombidiformes","Bdellidae","","Cyta","","38","35","35","7","7.89","0","98.12","441.24","0","442.24","31.46","0"
"AGAW2","09N","NIN","BOLD:AAH8314","Arthropoda","Arachnida","Araneae","Linyphiidae","","Walckenaeria","Walckenaeria microspiralis","37","37","37","8","0","0","98.28","429.51","0","434.03","35.84","0"
"AGAW2","09N","NIN","BOLD:AAA8820","Arthropoda","Collembola","Entomobryomorpha","Entomobryidae","Lepidocyrtinae","Lepidocyrtus","Lepidocyrtus cyaneus","27","27","27","9","0","0","98.79","409.37","0","412.19","43.39","0"
"AGAW2","09N","NIN","BOLD:ACY2550","Arthropoda","Arachnida","Trombidiformes","Bdellidae","","Cyta","","17","17","17","10","0","0","99.47","385.71","0","416.12","31.15","0"
"AGAW2","09N","NIN","BOLD:ACZ1295","Arthropoda","Arachnida","Trombidiformes","Anystidae","","","","14","14","14","11","0","0","99.66","379.5","0","394.14","31.34","0"
"AGAW2","09N","NIN","BOLD:ACY9761","Arthropoda","Arachnida","Mesostigmata","Phytoseiidae","","","","13","13","13","12","0","0","99.57","431.69","0","433.85","31.73","0"
"AGAW2","09N","NIN","BOLD:AAB7914","Arthropoda","Collembola","Symphypleona","Bourletiellidae","","Bourletiella","","12","12","12","13","0","0","99.33","442.33","0","444.08","37.97","0"
"AGAW2","09N","NIN","BOLD:AAM7944","Arthropoda","Diplopoda","Julida","Julidae","","Brachyiulus","Brachyiulus pusillus","3","2","2","14","33.33","0","99.63","448.67","0","450","40.44","0"
"AGAW2","09N","NIN","BOLD:ABW5649","Arthropoda","Arachnida","Trombidiformes","Hydryphantidae","Euthyadinae","Parathyas","","2","2","2","15","0","0","99.39","426.5","0","427.5","31.56","0"
"AGAW2","09N","NIN","BOLD:AAB8452","Arthropoda","Collembola","Entomobryomorpha","Entomobryidae","Lepidocyrtinae","Lepidocyrtus","Lepidocyrtus paradoxus","2","2","2","15","0","0","98.54","445","0","450","41.22","0"
"AGAW2","09N","NIN","BOLD:ADD2059","Arthropoda","Arachnida","Mesostigmata","Phytoseiidae","","","","1","1","1","17","0","0","99.07","428","0","450","33.78","0"
"AGAW2","09N","NIN","BOLD:AAA6381","Arthropoda","Arachnida","Araneae","Tetragnathidae","","Tetragnatha","Tetragnatha laboriosa","1","1","1","17","0","0","100","448","0","450","40","0"
"AGAW2","09N","NIN","BOLD:ABY1984","Arthropoda","Arachnida","Mesostigmata","Phytoseiidae","","","","1","1","1","17","0","0","97.1","449","0","450","35.78","0"
"AGAW2","09N","NIN","BOLD:AAB3836","Arthropoda","Arachnida","Araneae","Philodromidae","","Philodromus","Philodromus cespitum","1","1","1","17","0","0","98.22","393","0","450","30.22","0"
"AGAW2","09S","BAC","TAX:91391","Proteobacteria","Alphaproteobacteria","Rickettsiales",NA,NA,NA,NA,"1","1","1","1","0","0","100","384","0","450","42.22","0"
"AGAW2","09S","CHO","no match","no match","no match","no match","no match","no match","no match","no match","no match","no match","no match","no match","no match","no match","no match","no match","no match","no match","no match","no match"
"AGAW2","09S","INS","BOLD:AAA3453","Arthropoda","Insecta","Diptera","Anthomyiidae","Anthomyiinae","Delia","","16974","14078","14078","1","17.06","0","99.39","431.82","0","434.42","34.63","0"
"AGAW2","09S","INS","BOLD:AAG1711","Arthropoda","Insecta","Diptera","Muscidae","Phaoniinae","Helina","Helina depuncta","12002","9506","9506","2","20.8","0","99.39","433.21","0","435.5","31.36","0"
"AGAW2","09S","INS","BOLD:AAG6745","Arthropoda","Insecta","Diptera","Calliphoridae","Calliphorinae","Pollenia","Pollenia pediculata","10032","8578","8578","3","14.49","0","99.28","430.14","0","432.16","33.46","0"
"AGAW2","09S","INS","BOLD:AAN5497","Arthropoda","Insecta","Diptera","Anthomyiidae","Pegomyinae","Pegomya","Pegomya solennis, Pegomya bicolor","7663","5047","5047","4","34.14","0","99.47","447.04","0","448.26","29.58","0"
"AGAW2","09S","INS","BOLD:AAA3867","Arthropoda","Insecta","Lepidoptera","Noctuidae","Noctuinae","Apamea","Apamea amputatrix","6771","5077","5077","5","25.02","0","99.3","441.99","0","443.6","33.67","0"
"AGAW2","09S","INS","BOLD:AAA4555","Arthropoda","Insecta","Orthoptera","Acrididae","Melanoplinae","Melanoplus","Melanoplus stupefactus, Melanoplus angustipennis, Melanoplus ponderosus, Melanoplus bruneri, Melanoplus frigidus, Melanoplus scapularis, Melanoplus cinereus, Melanoplus madeleineae, Melanoplus sanguinipes, Melanoplus nigrescens group, Melanoplus tunicae, Melanoplus walshi, Melanoplus femurrubrum, Melanoplus dawsoni, Melanoplus sp., Melanoplus fasciatus, Melanoplus bowditchi, Melanoplus stonei, Melanoplus bivittatus, Melanoplus huroni, Melanoplus thomasi, Melanoplus borealis, Melanoplus packardii","3627","3045","3045","6","16.05","0","99.22","442.8","0","444.21","36.18","0"
"AGAW2","09S","INS","BOLD:ACE7444","Arthropoda","Insecta","Hemiptera","Miridae","Mirinae, Phylinae","Adelphocoris, Megalocoleus","Adelphocoris lineolatus, Megalocoleus molliculus","3499","2120","2120","7","39.41","0","99.69","442.41","0","443.08","34.79","0"
"AGAW2","09S","INS","BOLD:AAA4277","Arthropoda","Insecta","Diptera","Syrphidae","Syrphinae","Toxomerus","Toxomerus marginatus, Toxomerus sp. CR-3, Toxomerus CR-3","1360","992","992","8","27.06","0","99.32","445.25","0","446.13","30.71","0"
"AGAW2","09S","INS","BOLD:AAC0903","Arthropoda","Insecta","Diptera","Chironomidae","Chironominae","Chironomus","Chironomus acidophilus","1358","1068","1068","9","21.35","0","99.48","438.52","0","441.67","36.26","0"
"AGAW2","09S","INS","BOLD:AAN6154","Arthropoda","Insecta","Coleoptera","Latridiidae","Corticariinae","Melanophthalma","Melanophthalma inermis","1342","1139","1139","10","15.13","0","98.86","438.73","0","440.12","31.33","0"
"AGAW2","09S","INS","BOLD:AAD0484","Arthropoda","Insecta","Diptera","Chironomidae","","","","490","414","414","11","15.51","0","98.79","444.98","0","446.34","34.55","0"
"AGAW2","09S","INS","BOLD:AAG2511","Arthropoda","Insecta","Diptera","Anthomyiidae","Anthomyiinae","Delia","Delia platura","482","447","447","12","7.26","0","98.82","444.29","0","446.99","33.14","0"
"AGAW2","09S","INS","BOLD:AAG9626","Arthropoda","Insecta","Diptera","Dolichopodidae","","","","480","389","389","13","18.96","0","98.66","437.02","0","438.99","36.16","0"
"AGAW2","09S","INS","BOLD:AAG5507","Arthropoda","Insecta","Diptera","Chironomidae","Chironominae","Chironomus","Chironomus atrella","337","335","335","14","0.59","0","99.01","422.82","0","428.54","35.85","0"
"AGAW2","09S","INS","BOLD:AAB0373","Arthropoda","Insecta","Neuroptera","Chrysopidae","Chrysopinae","Chrysoperla","Chrysoperla sp. isolate 82, Chrysoperla zastrowi, Chrysoperla plorabunda, Chrysoperla sp. isolate 71, Chrysoperla agilis, Italochrysa italica, Chrysoperla sp. isolate 88, Chrysoperla lucasina, Chrysoperla carnea, Chrysoperla mediterranea, Chrysoperla pallida, Chrysoperla nipponensis, Chrysoperla sillemi","320","272","272","15","15","0","99.43","434.38","0","435.33","31.34","0"
"AGAW2","09S","INS","BOLD:ACK3431","Arthropoda","Insecta","Diptera","Cecidomyiidae","","","","291","259","259","16","11","0","99.13","430.17","0","431.78","32.79","0"
"AGAW2","09S","INS","BOLD:ABZ4644","Arthropoda","Insecta","Diptera","Chloropidae","Oscinellinae","Malloewia","Malloewia nigripalpis","278","242","242","17","12.95","0","99.2","436.51","0","437.69","30.78","0"
"AGAW2","09S","INS","BOLD:AAF9771","Arthropoda","Insecta","Diptera","Hybotidae","Tachydromiinae","Platypalpus","Platypalpus longiseta, Platypalpus trivialis, Platypalpus sp. 5, Platypalpus sp. 17, Platypalpus pallidiventris","240","207","207","18","13.75","0","99.37","442.95","0","443.9","35.05","0"
"AGAW2","09S","INS","BOLD:ACK9434","Arthropoda","Insecta","Hemiptera","Cicadellidae","Macropsinae","Macropsis","Macropsis ocellata","227","181","181","19","20.26","0","99.27","446.99","0","447.76","33.98","0"
"AGAW2","09S","INS","BOLD:AAA8933","Arthropoda","Insecta","Coleoptera","Coccinellidae","Coccinellinae","Coccinella","Coccinella septempunctata","205","172","172","20","16.1","0","99.27","443.65","0","445.9","32.73","0"
"AGAW2","09S","INS","BOLD:AAA5299","Arthropoda","Insecta","Diptera","Chironomidae","Orthocladiinae","Cricotopus","Cricotopus laetus, Cricotopus relucens, Cricotopus trifasciatus, Cricotopus sp., Cricotopus sp. 3ES, Cricotopus sylvestris, Cricotopus glacialis","198","175","175","21","11.62","0","99.32","431.89","0","433.93","34.52","0"
"AGAW2","09S","INS","BOLD:AAG4781","Arthropoda","Insecta","Diptera","Agromyzidae","","","","171","135","135","22","21.05","0","98.57","439.96","0","442.29","31.95","0"
"AGAW2","09S","INS","BOLD:ABW7897","Arthropoda","Insecta","Diptera","Cecidomyiidae","","","","163","126","126","23","22.7","0","99.59","444.4","0","445.8","29.7","0"
"AGAW2","09S","INS","BOLD:AAI2688","Arthropoda","Insecta","Diptera","Chironomidae","Orthocladiinae","Parametriocnemus","","156","134","134","24","14.1","0","99.04","446.94","0","448.92","32.83","0"
"AGAW2","09S","INS","BOLD:ACX5521","Arthropoda","Insecta","Diptera","Chironomidae","Orthocladiinae","","","128","120","120","25","6.25","0","99.14","420.09","0","422.23","34.56","0"
"AGAW2","09S","INS","BOLD:AAN5199","Arthropoda","Insecta","Diptera","Cecidomyiidae","","","CecidInt20 sp. BOLD:AAN5199","102","82","82","26","19.61","0","99.56","434.85","0","447.56","26.81","0"
"AGAW2","09S","INS","BOLD:AAD7604","Arthropoda","Insecta","Coleoptera","Coccinellidae","Coccinellinae","Coleomegilla","Coleomegilla maculata","100","95","95","27","5","0","98.77","436.35","0","439.42","31.77","0"
"AGAW2","09S","INS","BOLD:AAP7026","Arthropoda","Insecta","Coleoptera","Latridiidae","Corticariinae","Melanophthalma","Melanophthalma villosa","97","77","77","28","20.62","0","98.29","408.14","0","440.42","30.11","0"
"AGAW2","09S","INS","BOLD:AAG2868","Arthropoda","Insecta","Hemiptera","Cicadellidae","Typhlocybinae","Empoasca","Empoasca sp., Empoasca fabae","84","56","56","29","33.33","0","99.5","446.24","0","449.54","32.23","0"
"AGAW2","09S","INS","BOLD:AAH4135","Arthropoda","Insecta","Diptera","Chloropidae","Chloropinae","Thaumatomyia","Thaumatomyia glabra, Thaumatomyia grata, Thaumatomyia sp. 1 TAW","82","81","81","30","1.22","0","99.39","424.38","0","431.28","31.84","0"
"AGAW2","09S","INS","BOLD:AAG9629","Arthropoda","Insecta","Diptera","Dolichopodidae","Dolichopodinae","Dolichopus","Dolichopus omnivagus, Dolichopus lobatus","74","72","72","31","2.7","0","98.93","441.42","0","442.91","39.1","0"
"AGAW2","09S","INS","BOLD:AAA3181","Arthropoda","Insecta","Diptera","Muscidae","Muscinae","Stomoxys","Stomoxys calcitrans","66","63","63","32","4.55","0","98.55","420.83","0","424.64","32.59","0"
"AGAW2","09S","INS","BOLD:AAL4232","Arthropoda","Insecta","Diptera","Agromyzidae","Phytomyzinae","Phytoliriomyza","Phytoliriomyza arctica","62","53","53","33","14.52","0","99.47","441.44","0","442.52","32.03","0"
"AGAW2","09S","INS","BOLD:AAN5205","Arthropoda","Insecta","Diptera","Cecidomyiidae","","","CecidInt42 sp. BOLD:AAN5205","61","54","54","34","11.48","0","99.44","446.38","0","447.93","31.33","0"
"AGAW2","09S","INS","BOLD:AAH3661","Arthropoda","Insecta","Diptera","Cecidomyiidae","","","","56","54","54","35","3.57","0","99.1","434.16","0","435.71","25.51","0"
"AGAW2","09S","INS","BOLD:ABU5525","Arthropoda","Insecta","Diptera","Chironomidae","Orthocladiinae","Limnophyes","Limnophyes sp. 14ES","51","50","50","36","1.96","0","98.83","408.65","0","411.35","37.22","0"
"AGAW2","09S","INS","BOLD:AAD4214","Arthropoda","Insecta","Hymenoptera","Ichneumonidae","Diplazontinae","Diplazon","Diplazon laetatorius","51","43","43","36","15.69","0","99.13","445.88","0","446.86","25.86","0"
"AGAW2","09S","INS","BOLD:ABY5318","Arthropoda","Insecta","Diptera","Chloropidae","Oscinellinae","Incertella, Rhopalopterum","Rhopalopterum umbrosum, Rhopalopterum luteiceps, Incertella bispina, Rhopalopterum painteri","50","44","44","38","12","0","98.57","443.7","0","444.04","32.34","0"
"AGAW2","09S","INS","BOLD:AAD2041","Arthropoda","Insecta","Diptera","Dolichopodidae","Sciapodinae","Condylostylus","Condylostylus inermis, Condylostylus nigrofemoratus, Condylostylus connectans, Condylostylus caudatus, Condylostylus flavipes","49","40","40","39","18.37","0","99.54","440.18","0","441.45","35.83","0"
"AGAW2","09S","INS","BOLD:AAH3664","Arthropoda","Insecta","Diptera","Cecidomyiidae","","","CecidInt3 sp. BOLD:AAH3664","49","49","49","39","0","0","99.04","426.69","0","431.43","26.79","0"
"AGAW2","09S","INS","BOLD:ABY1179","Arthropoda","Insecta","Hymenoptera","Sierolomorphidae","","Sierolomorpha","","43","41","41","41","4.65","0","99.59","441.12","0","442.95","28.16","0"
"AGAW2","09S","INS","BOLD:ABV0470","Arthropoda","Insecta","Diptera","Cecidomyiidae","","","","38","37","37","42","2.63","0","99.32","416.76","0","418.45","26.17","0"
"AGAW2","09S","INS","BOLD:ABV1194","Arthropoda","Insecta","Diptera","Chironomidae","Chironominae","Micropsectra","","36","34","34","43","5.56","0","99.46","432.81","0","435.03","35.93","0"
"AGAW2","09S","INS","BOLD:AAA7661","Arthropoda","Insecta","Diptera","Culicidae","Culicinae","Culiseta, Culex","Culiseta melanura, Culex articularis, Culex restuans, Culex acharistus","36","36","36","43","0","0","98.85","435.22","0","437.36","33.16","0"
"AGAW2","09S","INS","BOLD:ABY1165","Arthropoda","Insecta","Coleoptera","Anthicidae","","","","32","31","31","45","3.13","0","98.29","422.81","0","424.84","35.77","0"
"AGAW2","09S","INS","BOLD:ABV9277","","","","","","","","28","28","28","46","0","0","98.56","444.39","0","446.61","28.06","0"
"AGAW2","09S","INS","BOLD:AAC0706","Arthropoda","Insecta","Diptera","Chironomidae","Chironominae","Dicrotendipes","Dicrotendipes tritomus","27","25","25","47","7.41","0","98.43","432.41","0","434.48","35.94","0"
"AGAW2","09S","INS","BOLD:ACD4236","","","","","","","","26","26","26","48","0","0","98.78","400.08","0","408.81","33.98","0"
"AGAW2","09S","INS","BOLD:ACU7906","Arthropoda","Insecta","Diptera","Chironomidae","Orthocladiinae","","","21","21","21","49","0","0","98.64","398.86","0","402.86","35.87","0"
"AGAW2","09S","INS","BOLD:AAQ0558","Arthropoda","Insecta","Thysanoptera","Thripidae","Thripinae","Anaphothrips","Anaphothrips obscurus","18","18","18","50","0","0","99.26","438.44","0","440.39","31.59","0"
"AGAW2","09S","INS","BOLD:AAN5267","Arthropoda","Insecta","Diptera","Cecidomyiidae","","","","17","16","16","51","5.88","0","99.54","449.47","0","450","30.65","0"
"AGAW2","09S","INS","BOLD:AAV1136","Arthropoda","Insecta","Diptera","Scatopsidae","","","","16","16","16","52","0","0","98.76","421","0","422.38","36.19","0"
"AGAW2","09S","INS","BOLD:AAA4781","Arthropoda","Insecta","Hymenoptera","Braconidae","Microgastrinae","Glyptapanteles","Glyptapanteles carlossarmientoi","16","16","16","52","0","0","99.22","428.06","0","430.19","24.25","0"
"AGAW2","09S","INS","BOLD:AAN5183","Arthropoda","Insecta","Diptera","Cecidomyiidae","","","","15","15","15","54","0","0","98.37","442.07","0","443.07","27.74","0"
"AGAW2","09S","INS","BOLD:AAM9255","Arthropoda","Insecta","Diptera","Sciaridae","","Pseudolycoriella","","13","13","13","55","0","0","99.02","402.69","0","405.46","30.68","0"
"AGAW2","09S","INS","BOLD:ABX5069","Arthropoda","Insecta","Trichoptera","Hydroptilidae","Hydroptilinae","Hydroptila","Hydroptila consimilis","12","11","11","56","8.33","0","99.46","440.92","0","443","33.14","0"
"AGAW2","09S","INS","BOLD:AAL7819","Arthropoda","Insecta","Diptera","Psychodidae","Psychodinae","Psychoda","Psychoda sp., Psychoda sp. BOLD:AAL7819","11","11","11","57","0","0","99.78","399.91","0","402.18","32.76","0"
"AGAW2","09S","INS","BOLD:AAJ2720","Arthropoda","Insecta","Coleoptera","Staphylinidae","Aleocharinae","Aleochara","Aleochara verna","11","11","11","57","0","0","98.66","411.55","0","413.18","30.07","0"
"AGAW2","09S","INS","BOLD:AAH0134","Arthropoda","Insecta","Coleoptera","Phalacridae","Phalacrinae","Stilbus","Stilbus apicialis","9","9","9","59","0","0","99.37","430.44","0","431.89","32.01","0"
"AGAW2","09S","INS","BOLD:ABA0863","Arthropoda","Insecta","Diptera","Cecidomyiidae","","","","8","7","7","60","12.5","0","99.16","449","0","450","29.97","0"
"AGAW2","09S","INS","BOLD:AAG8091","Arthropoda","Insecta","Hymenoptera","Ichneumonidae","Campopleginae","Campoletis","Campoletis sp. AB.2","8","8","8","60","0","0","99.46","443.25","0","444.5","24.83","0"
"AGAW2","09S","INS","BOLD:ACI1304","Arthropoda","Insecta","Diptera","Muscidae","Muscinae","Stomoxys","Stomoxys indicus","7","7","7","62","0","0","97.51","416.14","0","421","32.33","0"
"AGAW2","09S","INS","BOLD:ABA6407","Arthropoda","Insecta","Diptera","Sciaridae","","Pseudolycoriella","","7","7","7","62","0","0","99.84","438.71","0","440.57","35.07","0"
"AGAW2","09S","INS","BOLD:AAZ0271","Arthropoda","Insecta","Diptera","Ceratopogonidae","Ceratopogoninae","Culicoides","","7","7","7","62","0","0","98.74","440","0","442.86","41.34","0"
"AGAW2","09S","INS","BOLD:ACE8878","Arthropoda","Insecta","Orthoptera","Acrididae","Melanoplinae","Melanoplus","Melanoplus scudderi group","6","6","6","65","0","0","97.44","449.83","0","450","35.48","0"
"AGAW2","09S","INS","BOLD:AAH0135","Arthropoda","Insecta","Coleoptera","Phalacridae","Phalacrinae","Acylomus","Acylomus pugetanus, Acylomus ergoti","6","6","6","65","0","0","99.52","449.17","0","450","32.04","0"
"AGAW2","09S","INS","BOLD:AAP8739","Arthropoda","Insecta","Diptera","Platystomatidae","Platystomatinae","Rivellia","Rivellia steyskali","6","6","6","65","0","0","98.7","439.83","0","441.33","37.89","0"
"AGAW2","09S","INS","BOLD:AAN5342","Arthropoda","Insecta","Diptera","Chironomidae","Orthocladiinae","Heleniella","","6","6","6","65","0","0","99.34","410.67","0","412","35.3","0"
"AGAW2","09S","INS","BOLD:ACF1257","","","","","","","","5","5","5","69","0","0","99.6","449.4","0","450","34.8","0"
"AGAW2","09S","INS","BOLD:ADF5759","Arthropoda","Insecta","Diptera","Cecidomyiidae","","","","5","5","5","69","0","0","99.5","437.8","0","438.8","27.51","0"
"AGAW2","09S","INS","BOLD:ACJ6954","Arthropoda","Insecta","Diptera","Cecidomyiidae","","","","3","3","3","71","0","0","99.61","338","0","418","26.06","0"
"AGAW2","09S","INS","BOLD:AAG7276","Arthropoda","Insecta","Diptera","Sphaeroceridae","Limosininae","Leptocera","Leptocera erythrocera","3","3","3","71","0","0","99","425","0","428.33","32.92","0"
"AGAW2","09S","INS","BOLD:ABA1215","Arthropoda","Insecta","Diptera","Sciaridae","","Lycoriella","Lycoriella sativae","3","3","3","71","0","0","99.21","405.67","0","408","35.17","0"
"AGAW2","09S","INS","BOLD:AAM7855","Arthropoda","Insecta","Thysanoptera","Thripidae","Thripinae","Frankliniella","Frankliniella tritici","3","3","3","71","0","0","99.76","430.33","0","432.33","31.04","0"
"AGAW2","09S","INS","BOLD:AAE7186","Arthropoda","Insecta","Hymenoptera","Braconidae","Aphidiinae","Praon","Praon sp. 1 MM-2013, Praon volucre, Praon bicolor","3","3","3","71","0","0","99.4","449","0","450","25.93","0"
"AGAW2","09S","INS","BOLD:AAU6681","Arthropoda","Insecta","Diptera","Tipulidae","Tipulinae","Nephrotoma","Nephrotoma cornicina","3","3","3","71","0","0","99.48","445.33","0","446.33","36.37","0"
"AGAW2","09S","INS","BOLD:AAU8713","Arthropoda","Insecta","Hymenoptera","Dryinidae","Anteoninae","Anteon","Anteon tripartitum, Anteon gaullei, Anteon cameroni, Anteon pubicorne","3","3","3","71","0","0","98.92","403","0","405.33","27.06","0"
"AGAW2","09S","INS","BOLD:AAH0256","Arthropoda","Insecta","Coleoptera","Latridiidae","Corticariinae","Corticaria, Corticarina","","3","3","3","71","0","0","98.67","421","0","431","31.99","0"
"AGAW2","09S","INS","BOLD:ADH9867","Arthropoda","Insecta","Diptera","","","","","3","3","3","71","0","0","98.74","407.67","0","411.33","37.34","0"
"AGAW2","09S","INS","BOLD:AAM6037","Arthropoda","Insecta","Diptera","Cecidomyiidae","Cecidomyiinae","Contarinia, Dasineura","","3","3","3","71","0","0","98.4","437.67","0","441","25.46","0"
"AGAW2","09S","INS","BOLD:ABZ4994","Arthropoda","Insecta","Orthoptera","Acrididae","Melanoplinae","Melanoplus","Melanoplus gladstoni, Melanoplus sanguinipes","3","3","3","71","0","0","97.67","431.33","0","425.33","35.95","0"
"AGAW2","09S","INS","BOLD:AAA3654","Arthropoda","Insecta","Lepidoptera","Crambidae","Crambinae","Crambus","Crambus bigelovi, Crambus johnsoni, Crambus praefectellus","3","2","2","71","33.33","0","98.89","450","0","450","31.85","0"
"AGAW2","09S","INS","BOLD:AAU7170","Arthropoda","Insecta","Coleoptera","Cryptophagidae","Atomariinae","Atomaria","Atomaria wollastoni","2","2","2","83","0","0","99.26","410.5","0","411","39.67","0"
"AGAW2","09S","INS","BOLD:AAN5230","Arthropoda","Insecta","Diptera","Cecidomyiidae","","","","2","2","2","83","0","0","99.44","448.5","0","450","26.33","0"
"AGAW2","09S","INS","BOLD:AAC0635","Arthropoda","Insecta","Hemiptera","Miridae","Mirinae","Stenotus","Stenotus binotatus","2","1","1","83","50","0","97.33","450","0","450","34","0"
"AGAW2","09S","INS","BOLD:AAG7891","Arthropoda","Insecta","Hymenoptera","Platygastridae","Scelioninae","Telenomus","Telenomus podisi","2","2","2","83","0","0","99.54","439","0","450","26.67","0"
"AGAW2","09S","INS","BOLD:ABZ5071","Arthropoda","Insecta","Diptera","Chironomidae","Chironominae","Micropsectra","Micropsectra nigripila","2","2","2","83","0","0","98.65","402.5","0","411.5","36.97","0"
"AGAW2","09S","INS","BOLD:AAA2815","Arthropoda","Insecta","Lepidoptera","Noctuidae","Noctuinae","Mythimna","Mythimna yukonensis, Mythimna oxygala","2","2","2","83","0","0","99.44","447.5","0","450","33.56","0"
"AGAW2","09S","INS","BOLD:AAF9187","Arthropoda","Insecta","Coleoptera","Curculionidae","Entiminae","Phyllobius","Phyllobius oblongus","2","2","2","83","0","0","99.22","448.5","0","450","38.78","0"
"AGAW2","09S","INS","BOLD:ACF4925","Arthropoda","Insecta","Hemiptera","Miridae","Mirinae","Adelphocoris","Adelphocoris detritus","2","1","1","83","50","0","97.33","449","0","450","36.44","0"
"AGAW2","09S","INS","BOLD:AAB7072","Arthropoda","Insecta","Lepidoptera","Coleophoridae","Coleophorinae","Coleophora","Coleophora coxi, Coleophora texanella","1","1","1","91","0","0","97.99","447","0","450","33.78","0"
"AGAW2","09S","INS","BOLD:ABW3687","Arthropoda","Insecta","Diptera","Lauxaniidae","Homoneurinae","Homoneura","Homoneura bispina","1","1","1","91","0","0","97.56","450","0","450","34","0"
"AGAW2","09S","INS","BOLD:AAA6292","Arthropoda","Insecta","Lepidoptera","Tortricidae","Tortricinae","Cnephasia","Cnephasia genitalana, Cnephasia stephensiana","1","1","1","91","0","0","100","449","0","450","35.56","0"
"AGAW2","09S","INS","BOLD:AAG1717","Arthropoda","Insecta","Diptera","Muscidae","Phaoniinae","Helina","Helina troene","1","1","1","91","0","0","98.45","453","0","450","30","0"
"AGAW2","09S","INS","BOLD:AAA2590","Arthropoda","Insecta","Lepidoptera","Noctuidae","Noctuinae","Xestia","Xestia baja, Xestia smithii","1","1","1","91","0","0","99.56","450","0","450","32.89","0"
"AGAW2","09S","INS","BOLD:ACF2568","Arthropoda","Insecta","Orthoptera","Acrididae","Melanoplinae","Melanoplus","Melanoplus scudderi group, Melanoplus scudderi, Melanoplus quercicola","1","1","1","91","0","0","97.1","449","0","450","36.67","0"
"AGAW2","09S","INS","BOLD:AAA6560","Arthropoda","Insecta","Lepidoptera","Crambidae","Crambinae","Crambus","Crambus multilinellus, Crambus saltuellus","1","1","1","91","0","0","100","450","0","450","31.11","0"
"AGAW2","09S","INS","BOLD:ABV9319","Arthropoda","Insecta","Diptera","Chironomidae","Tanypodinae","Procladius","","1","1","1","91","0","0","97.77","449","0","450","33.56","0"
"AGAW2","09S","INS","BOLD:AAN5220","Arthropoda","Insecta","Diptera","Cecidomyiidae","","","","1","1","1","91","0","0","97.77","448","0","450","27.11","0"
"AGAW2","09S","INS","BOLD:ABZ7164","Arthropoda","Insecta","Diptera","Cecidomyiidae","","","","1","1","1","91","0","0","97.45","432","0","433","25.87","0"
"AGAW2","09S","INS","BOLD:AAA7631","Arthropoda","Insecta","Hemiptera","Miridae","Mirinae","Adelphocoris","Adelphocoris lineolatus","1","1","1","91","0","0","97.56","450","0","450","35.11","0"
"AGAW2","09S","INS","BOLD:AAG0999","Arthropoda","Insecta","Diptera","Chironomidae","Orthocladiinae","","","1","1","1","91","0","0","97.51","402","0","403","34.49","0"
"AGAW2","09S","INS","BOLD:AAL7359","Arthropoda","Insecta","Diptera","Chironomidae","Chironominae","Tanytarsus","","1","1","1","91","0","0","100","449","0","450","36.44","0"
"AGAW2","09S","INS","BOLD:AAB9652","Arthropoda","Insecta","Trichoptera","Leptoceridae","Leptocerinae","Triaenodes","Triaenodes nox","1","1","1","91","0","0","98.19","386","0","390","30","0"
"AGAW2","09S","INS","BOLD:AAA8796","Arthropoda","Insecta","Lepidoptera","Tortricidae","Tortricinae","Acleris","Acleris forsskaleana","1","1","1","91","0","0","100","377","0","377","32.1","0"
"AGAW2","09S","INS","BOLD:ACE8875","Arthropoda","Insecta","Orthoptera","Acrididae","Melanoplinae","Melanoplus","Melanoplus bowditchi","1","1","1","91","0","0","97.77","448","0","450","36","0"
"AGAW2","09S","INS","BOLD:AAG9663","Arthropoda","Insecta","Diptera","Dolichopodidae","Dolichopodinae","Dolichopus","Dolichopus amphericus","1","1","1","91","0","0","97","434","0","449","36.53","0"
"AGAW2","09S","INS","BOLD:ACK3469","Arthropoda","Insecta","Coleoptera","Chrysomelidae","Eumolpinae","Brachypnoea","Brachypnoea puncticollis","1","1","1","91","0","0","99.53","425","0","433","35.33","0"
"AGAW2","09S","INS","BOLD:ACV5271","Arthropoda","Insecta","Diptera","Cecidomyiidae","","","","1","1","1","91","0","0","99.56","450","0","450","28.22","0"
"AGAW2","09S","INS","BOLD:AAA9246","Arthropoda","Insecta","Lepidoptera","Tortricidae","Tortricinae","Clepsis","Clepsis clemensiana","1","1","1","91","0","0","99.11","448","0","450","30","0"
"AGAW2","09S","INS","BOLD:ACM1176","Arthropoda","Insecta","Diptera","Anthomyiidae","Pegomyinae","Pegomya","Pegomya rugulosa","1","1","1","91","0","0","97.53","446","0","450","29.33","0"
"AGAW2","09S","INS","BOLD:AAG2898","Arthropoda","Insecta","Hemiptera","Cicadellidae","Deltocephalinae","Paraphlepsius","Paraphlepsius apertus, Paraphlepsius irroratus","1","1","1","91","0","0","99.33","449","0","450","32","0"
"AGAW2","09S","INS","BOLD:AAZ5967","Arthropoda","Insecta","Diptera","Limoniidae","Chioneinae","Molophilus","Molophilus forcipulus, Molophilus pubipennis","1","1","1","91","0","0","99.11","447","0","450","38","0"
"AGAW2","09S","INS","BOLD:ACY6524","Arthropoda","Insecta","Diptera","Chironomidae","Orthocladiinae","Psectrocladius","Psectrocladius cf. limbatellus","1","1","1","91","0","0","99.04","314","0","450","36","0"
"AGAW2","09S","INS","BOLD:AAA7374","Arthropoda","Insecta","Diptera","Syrphidae","Syrphinae","Sphaerophoria","Sphaerophoria bankowskae, Sphaerophoria infuscata, Sphaerophoria nr. philanthus, Sphaerophoria batava, Sphaerophoria sp., Sphaerophoria rueppellii, Sphaerophoria interrupta-group, Sphaerophoria sp. MA-1, Sphaerophoria cleoae, Sphaerophoria asymmetrica, Sphaerophoria fatarum, Sphaerophoria bengalensis, Sphaerophoria bifurcata, Sphaerophoria taeniata, Sphaerophoria interrupta, Sphaerophoria contigua, Sphaerophoria kaa, Sphaerophoria laurae, Sphaerophoria chongjini, Sphaerophoria boreoalpina, Sphaerophoria sp. BOLDAAA7374, Sphaerophoria macrogaster, Sphaerophoria sulphuripes, Sphaerophoria brevipilosa, Sphaerophoria philanthus, Sphaerophoria pyrrhina, Sphaerophoria abbreviata, Sphaerophoria scripta, Sphaerophoria potentillae, Sphaerophoria longipilosa, Sphaerophoria virgata","1","1","1","91","0","0","99.24","394","0","398","30.15","0"
"AGAW2","09S","INS","BOLD:ABX0893","Arthropoda","Insecta","Diptera","Cecidomyiidae","","","","1","1","1","91","0","0","97.33","449","0","450","26.89","0"
"AGAW2","09S","INS","BOLD:AAV0162","Arthropoda","Insecta","Hemiptera","Cicadellidae","Deltocephalinae","Eutettix","Eutettix variabilis","1","1","1","91","0","0","99.73","364","0","398","35.93","0"
"AGAW2","09S","INS","BOLD:ACK6937","Arthropoda","Insecta","Diptera","Cecidomyiidae","","","","1","1","1","91","0","0","100","407","0","408","25.74","0"
"AGAW2","09S","INS","BOLD:AAD8086","Arthropoda","Insecta","Lepidoptera","Elachistidae","Elachistinae","Elachista","Elachista griseicornis, Elachista acenteta","1","1","1","91","0","0","99.29","420","0","421","30.4","0"
"AGAW2","09S","INS","BOLD:AAP6465","Arthropoda","Insecta","Diptera","Sciaridae","","Corynoptera","Corynoptera cursor","1","1","1","91","0","0","98.44","450","0","450","32.22","0"
"AGAW2","09S","INS","BOLD:ACD3585","Arthropoda","Insecta","Hymenoptera","Eulophidae","Tetrastichinae","Tetrastichus","","1","1","1","91","0","0","100","395","0","450","28.89","0"
"AGAW2","09S","INS","BOLD:AAH3942","Arthropoda","Insecta","Diptera","Sciaridae","","","","1","1","1","91","0","0","99.04","415","0","416","31.49","0"
"AGAW2","09S","NAR","TAX:660402","Mollusca","Gastropoda","Stylommatophora","Agriolimacidae",NA,"Deroceras","Deroceras sp.","1","1","1","1","0","0","98.25","401","0","402","32.59","0"
"AGAW2","09S","NIN","BOLD:AAH6630","Arthropoda","Collembola","Entomobryomorpha","Entomobryidae","Entomobryinae","Entomobrya","Entomobrya unostrigata","2948","2399","2399","1","18.62","0","99.34","438.81","0","439.95","39.05","0"
"AGAW2","09S","NIN","BOLD:AAA5267","Arthropoda","Collembola","Entomobryomorpha","Isotomidae","","","","439","439","439","2","0","0","98.98","421.72","0","424.11","38.17","0"
"AGAW2","09S","NIN","BOLD:AAU6252","Arthropoda","Arachnida","Trombidiformes","Bdellidae","","Cyta","","120","105","105","3","12.5","0","98.85","436.16","0","437.34","31.49","0"
"AGAW2","09S","NIN","BOLD:ABW2639","Arthropoda","Arachnida","Trombidiformes","Anystidae","","","","55","51","51","4","7.27","0","99.69","446.15","0","442.75","31.42","0"
"AGAW2","09S","NIN","BOLD:AAA8820","Arthropoda","Collembola","Entomobryomorpha","Entomobryidae","Lepidocyrtinae","Lepidocyrtus","Lepidocyrtus cyaneus","30","30","30","5","0","0","99.04","420.57","0","423.33","43.33","0"
"AGAW2","09S","NIN","BOLD:ACY2550","Arthropoda","Arachnida","Trombidiformes","Bdellidae","","Cyta","","23","23","23","6","0","0","99.24","383.39","0","409.83","31.12","0"
"AGAW2","09S","NIN","BOLD:ACZ1295","Arthropoda","Arachnida","Trombidiformes","Anystidae","","","","9","9","9","7","0","0","99.48","385.44","0","419.67","31.31","0"
"AGAW2","09S","NIN","BOLD:AAA6381","Arthropoda","Arachnida","Araneae","Tetragnathidae","","Tetragnatha","Tetragnatha laboriosa","3","3","3","8","0","0","98.89","449.33","0","450","39.78","0"
"AGAW2","09S","NIN","BOLD:ADD2059","Arthropoda","Arachnida","Mesostigmata","Phytoseiidae","","","","2","2","2","9","0","0","99.65","427.5","0","450","33.33","0"
"AGAW2","09S","NIN","BOLD:ACC8538","Arthropoda","Arachnida","Mesostigmata","Blattisociidae","","","","2","2","2","9","0","0","99.74","417","0","417.5","32.9","0"
"AGAW2","09S","NIN","BOLD:ACM6751","Arthropoda","Arachnida","Trombidiformes","Anystidae","","","","1","1","1","11","0","0","100","402","0","450","29.56","0"
"AGAW2","09S","NIN","BOLD:ACI9974","Arthropoda","Arachnida","Trombidiformes","Tydeidae","","","","1","1","1","11","0","0","99.53","425","0","450","40.67","0"
"AGAW2","09S","NIN","BOLD:AAA7242","Arthropoda","Collembola","Entomobryomorpha","Entomobryidae","Seirinae, Entomobryinae","Entomobrya, Lepidosira","Entomobrya multifasciata, Lepidosira sp.","1","1","1","11","0","0","98.66","449","0","450","39.56","0"
"AGAW2","09S","NIN","BOLD:AAB7914","Arthropoda","Collembola","Symphypleona","Bourletiellidae","","Bourletiella","","1","1","1","11","0","0","97.78","450","0","450","38.67","0"
"AGAW2","09S","NIN","BOLD:ACF8020","Arthropoda","Arachnida","Mesostigmata","Ascidae","","","","1","1","1","11","0","0","99.78","449","0","450","37.56","0"
"AGAW2","09W","BAC","TAX:91393","Proteobacteria","Alphaproteobacteria","Rickettsiales","Rickettsiaceae",NA,"Wolbachia","","14","14","14","1","0","0","98.87","391.57","0","432.86","39.42","0"
"AGAW2","09W","BAC","TAX:175768","Proteobacteria","Alphaproteobacteria","Rickettsiales","Rickettsiaceae",NA,"Wolbachia","Wolbachia pipientis","9","9","9","2","0","0","98.19","447.56","0","448.44","39.69","0"
"AGAW2","09W","CHO","no match","no match","no match","no match","no match","no match","no match","no match","no match","no match","no match","no match","no match","no match","no match","no match","no match","no match","no match","no match"
"AGAW2","09W","INS","BOLD:AAA3453","Arthropoda","Insecta","Diptera","Anthomyiidae","Anthomyiinae","Delia","","35831","28288","28288","1","21.05","0","99.45","432.26","0","434.93","34.66","0"
"AGAW2","09W","INS","BOLD:AAG1711","Arthropoda","Insecta","Diptera","Muscidae","Phaoniinae","Helina","Helina depuncta","13523","10530","10530","2","22.13","0","99.4","433.62","0","435.87","31.37","0"
"AGAW2","09W","INS","BOLD:AAA4555","Arthropoda","Insecta","Orthoptera","Acrididae","Melanoplinae","Melanoplus","Melanoplus stupefactus, Melanoplus angustipennis, Melanoplus ponderosus, Melanoplus bruneri, Melanoplus frigidus, Melanoplus scapularis, Melanoplus cinereus, Melanoplus madeleineae, Melanoplus sanguinipes, Melanoplus nigrescens group, Melanoplus tunicae, Melanoplus walshi, Melanoplus femurrubrum, Melanoplus dawsoni, Melanoplus sp., Melanoplus fasciatus, Melanoplus bowditchi, Melanoplus stonei, Melanoplus bivittatus, Melanoplus huroni, Melanoplus thomasi, Melanoplus borealis, Melanoplus packardii","5458","4248","4248","3","22.17","0","99.32","447.06","0","448.43","36.18","0"
"AGAW2","09W","INS","BOLD:AAA3867","Arthropoda","Insecta","Lepidoptera","Noctuidae","Noctuinae","Apamea","Apamea amputatrix","2826","2191","2191","4","22.47","0","99.29","440.54","0","442.33","33.7","0"
"AGAW2","09W","INS","BOLD:AAN6154","Arthropoda","Insecta","Coleoptera","Latridiidae","Corticariinae","Melanophthalma","Melanophthalma inermis","2397","1984","1984","5","17.23","0","98.97","437.48","0","438.99","31.19","0"
"AGAW2","09W","INS","BOLD:AAC0635","Arthropoda","Insecta","Hemiptera","Miridae","Mirinae","Stenotus","Stenotus binotatus","2288","1588","1588","6","30.59","0","99.35","443.43","0","444.23","34.55","0"
"AGAW2","09W","INS","BOLD:AAB0373","Arthropoda","Insecta","Neuroptera","Chrysopidae","Chrysopinae","Chrysoperla","Chrysoperla sp. isolate 82, Chrysoperla zastrowi, Chrysoperla plorabunda, Chrysoperla sp. isolate 71, Chrysoperla agilis, Italochrysa italica, Chrysoperla sp. isolate 88, Chrysoperla lucasina, Chrysoperla carnea, Chrysoperla mediterranea, Chrysoperla pallida, Chrysoperla nipponensis, Chrysoperla sillemi","1515","1178","1178","7","22.24","0","99.23","437","0","437.93","31.35","0"
"AGAW2","09W","INS","BOLD:ACE7444","Arthropoda","Insecta","Hemiptera","Miridae","Mirinae, Phylinae","Adelphocoris, Megalocoleus","Adelphocoris lineolatus, Megalocoleus molliculus","967","663","663","8","31.44","0","99.68","442.88","0","443.49","34.82","0"
"AGAW2","09W","INS","BOLD:AAA8933","Arthropoda","Insecta","Coleoptera","Coccinellidae","Coccinellinae","Coccinella","Coccinella septempunctata","943","734","734","9","22.16","0","99.43","443.43","0","445.31","32.97","0"
"AGAW2","09W","INS","BOLD:AAD7604","Arthropoda","Insecta","Coleoptera","Coccinellidae","Coccinellinae","Coleomegilla","Coleomegilla maculata","833","742","742","10","10.92","0","99.02","434.31","0","436.79","31.72","0"
"AGAW2","09W","INS","BOLD:ACX5521","Arthropoda","Insecta","Diptera","Chironomidae","Orthocladiinae","","","722","643","643","11","10.94","0","99.27","423.64","0","425.63","34.58","0"
"AGAW2","09W","INS","BOLD:AAH3664","Arthropoda","Insecta","Diptera","Cecidomyiidae","","","CecidInt3 sp. BOLD:AAH3664","702","650","650","12","7.41","0","98.9","433.07","0","436.22","27.15","0"
"AGAW2","09W","INS","BOLD:AAE0707","Arthropoda","Insecta","Diptera","Chironomidae","Chironominae","Microtendipes","Microtendipes pedellus","632","582","582","13","7.91","0","97.77","420.71","0","423.43","31.51","0"
"AGAW2","09W","INS","BOLD:AAG9626","Arthropoda","Insecta","Diptera","Dolichopodidae","","","","565","483","483","14","14.51","0","98.44","430.5","0","432.13","36.77","0"
"AGAW2","09W","INS","BOLD:AAG0999","Arthropoda","Insecta","Diptera","Chironomidae","Orthocladiinae","","","476","331","331","15","30.46","0","98","446.07","0","447.02","35.75","0"
"AGAW2","09W","INS","BOLD:AAC4197","Arthropoda","Insecta","Diptera","Chironomidae","Orthocladiinae","Paraphaenocladius","Paraphaenocladius impensus","452","377","377","16","16.59","0","99.44","432.33","0","438.18","35.1","0"
"AGAW2","09W","INS","BOLD:ABV1430","Arthropoda","Insecta","Diptera","Dolichopodidae","Medeterinae","Medetera","","379","367","367","17","3.17","0","99.22","415.69","0","420.72","38.93","0"
"AGAW2","09W","INS","BOLD:AAN5199","Arthropoda","Insecta","Diptera","Cecidomyiidae","","","CecidInt20 sp. BOLD:AAN5199","362","276","276","18","23.76","0","99.55","437.35","0","446.91","26.73","0"
"AGAW2","09W","INS","BOLD:AAF9949","Arthropoda","Insecta","Hemiptera","Miridae","Mirinae","Trigonotylus","Trigonotylus pulchellus, Trigonotylus americanus, Trigonotylus psammaecolor, Trigonotylus caelestialium, Trigonotylus viridis, Trigonotylus ruficornis","317","250","250","19","21.14","0","99.45","443.91","0","446.08","34.55","0"
"AGAW2","09W","INS","BOLD:AAU6681","Arthropoda","Insecta","Diptera","Tipulidae","Tipulinae","Nephrotoma","Nephrotoma cornicina","310","273","273","20","11.94","0","99.53","441.4","0","442.67","36.32","0"
"AGAW2","09W","INS","BOLD:AAG7279","Arthropoda","Insecta","Diptera","Sphaeroceridae","Limosininae","Spelobia","Spelobia ochripes","293","238","238","21","18.77","0","98.78","437.67","0","441.53","35.06","0"
"AGAW2","09W","INS","BOLD:ABY1179","Arthropoda","Insecta","Hymenoptera","Sierolomorphidae","","Sierolomorpha","","266","205","205","22","22.93","0","99.63","441.64","0","443.27","28.18","0"
"AGAW2","09W","INS","BOLD:AAG2511","Arthropoda","Insecta","Diptera","Anthomyiidae","Anthomyiinae","Delia","Delia platura","231","221","221","23","4.33","0","98.79","443.68","0","446.55","33.24","0"
"AGAW2","09W","INS","BOLD:AAH0152","Arthropoda","Insecta","Coleoptera","Cantharidae","Cantharinae","Cantharis","Cantharis rufa","215","181","181","24","15.81","0","98.5","435.43","0","439.31","35.22","0"
"AGAW2","09W","INS","BOLD:AAM7657","","","","","","","","204","183","183","25","10.29","0","99.12","424.56","0","426.2","34.38","0"
"AGAW2","09W","INS","BOLD:ACK5443","Arthropoda","Insecta","Diptera","Cecidomyiidae","","","","181","160","160","26","11.6","0","98.78","436.9","0","438.02","25.49","0"
"AGAW2","09W","INS","BOLD:AAA5299","Arthropoda","Insecta","Diptera","Chironomidae","Orthocladiinae","Cricotopus","Cricotopus laetus, Cricotopus relucens, Cricotopus trifasciatus, Cricotopus sp., Cricotopus sp. 3ES, Cricotopus sylvestris, Cricotopus glacialis","162","151","151","27","6.79","0","99.13","430.36","0","432.53","34.68","0"
"AGAW2","09W","INS","BOLD:AAH3661","Arthropoda","Insecta","Diptera","Cecidomyiidae","","","","138","125","125","28","9.42","0","99.29","432.8","0","434.87","25.46","0"
"AGAW2","09W","INS","BOLD:ABA0863","Arthropoda","Insecta","Diptera","Cecidomyiidae","","","","129","108","108","29","16.28","0","99.41","441.05","0","442.26","29.73","0"
"AGAW2","09W","INS","BOLD:ABZ5071","Arthropoda","Insecta","Diptera","Chironomidae","Chironominae","Micropsectra","Micropsectra nigripila","126","120","120","30","4.76","0","98.1","427.92","0","444.11","37.01","0"
"AGAW2","09W","INS","BOLD:AAG5507","Arthropoda","Insecta","Diptera","Chironomidae","Chironominae","Chironomus","Chironomus atrella","125","124","124","31","0.8","0","98.74","424.52","0","428.42","35.73","0"
"AGAW2","09W","INS","BOLD:AAH0135","Arthropoda","Insecta","Coleoptera","Phalacridae","Phalacrinae","Acylomus","Acylomus pugetanus, Acylomus ergoti","123","116","116","32","5.69","0","99.25","436.61","0","438.39","31.65","0"
"AGAW2","09W","INS","BOLD:AAN5342","Arthropoda","Insecta","Diptera","Chironomidae","Orthocladiinae","Heleniella","","115","112","112","33","2.61","0","98.69","420.99","0","429.67","35.13","0"
"AGAW2","09W","INS","BOLD:ABV1297","Arthropoda","Insecta","Diptera","Sciaridae","","","","108","106","106","34","1.85","0","99.17","434.74","0","437.31","30.25","0"
"AGAW2","09W","INS","BOLD:AAL7819","Arthropoda","Insecta","Diptera","Psychodidae","Psychodinae","Psychoda","Psychoda sp., Psychoda sp. BOLD:AAL7819","96","92","92","35","4.17","0","99.19","415.97","0","417.88","32.66","0"
"AGAW2","09W","INS","BOLD:AAP6790","Arthropoda","Insecta","Diptera","Chloropidae","Oscinellinae","Incertella","Incertella incerta","82","66","66","36","19.51","0","99.02","441.04","0","442.79","31.44","0"
"AGAW2","09W","INS","BOLD:ACX6055","Arthropoda","Insecta","Diptera","Phoridae","Metopininae","Megaselia","Megaselia sp. AK-2010-2, Megaselia sp. BOLD:ACX6055","80","70","70","37","12.5","0","99.47","436.5","0","440.99","33.27","0"
"AGAW2","09W","INS","BOLD:ACB6899","Arthropoda","Insecta","Diptera","Cecidomyiidae","","","","69","63","63","38","8.7","0","99.46","414.16","0","441.36","26.33","0"
"AGAW2","09W","INS","BOLD:AAP8816","Arthropoda","Insecta","Diptera","Cecidomyiidae","Cecidomyiinae","Sitodiplosis","Sitodiplosis sp. BOLD:AAP8816","67","64","64","39","4.48","0","99.11","427.51","0","432.04","26.04","0"
"AGAW2","09W","INS","BOLD:AAG2868","Arthropoda","Insecta","Hemiptera","Cicadellidae","Typhlocybinae","Empoasca","Empoasca sp., Empoasca fabae","66","64","64","40","3.03","0","99.13","438.35","0","446.12","32.58","0"
"AGAW2","09W","INS","BOLD:AAN5260","Arthropoda","Insecta","Diptera","Cecidomyiidae","","","","65","65","65","41","0","0","99.18","407.03","0","408.66","25.98","0"
"AGAW2","09W","INS","BOLD:AAH0524","Arthropoda","Insecta","Hymenoptera","Ichneumonidae","Diplazontinae","Woldstedtius","Woldstedtius flavolineatus","63","63","63","42","0","0","99.12","407.7","0","410.59","26.94","0"
"AGAW2","09W","INS","BOLD:AAN5220","Arthropoda","Insecta","Diptera","Cecidomyiidae","","","","62","61","61","43","1.61","0","98.02","404.16","0","423.76","26.47","0"
"AGAW2","09W","INS","BOLD:AAG8864","Arthropoda","Insecta","Hemiptera","Miridae","Mirinae","Taedia","Taedia colon","60","60","60","44","0","0","98.81","411.07","0","412.6","35.09","0"
"AGAW2","09W","INS","BOLD:AAE3943","Arthropoda","Insecta","Hemiptera","Miridae","Phylinae","Plagiognathus","Plagiognathus chrysanthemi","60","43","43","44","28.33","0","99.12","392.85","0","448.85","35.93","0"
"AGAW2","09W","INS","BOLD:AAV1261","Arthropoda","Insecta","Diptera","Sciaridae","","Bradysia","Bradysia splendida","56","48","48","46","14.29","0","99.37","436.13","0","437.63","34.01","0"
"AGAW2","09W","INS","BOLD:AAH0134","Arthropoda","Insecta","Coleoptera","Phalacridae","Phalacrinae","Stilbus","Stilbus apicialis","52","50","50","47","3.85","0","99.44","424.62","0","425.85","31.87","0"
"AGAW2","09W","INS","BOLD:AAN5183","Arthropoda","Insecta","Diptera","Cecidomyiidae","","","","49","45","45","48","8.16","0","98.34","444.22","0","445.18","27.71","0"
"AGAW2","09W","INS","BOLD:ABY5318","Arthropoda","Insecta","Diptera","Chloropidae","Oscinellinae","Incertella, Rhopalopterum","Rhopalopterum umbrosum, Rhopalopterum luteiceps, Incertella bispina, Rhopalopterum painteri","46","41","41","49","10.87","0","99.26","443.7","0","444.98","31.94","0"
"AGAW2","09W","INS","BOLD:AAD0484","Arthropoda","Insecta","Diptera","Chironomidae","","","","44","42","42","50","4.55","0","98.89","447.18","0","448.14","34.63","0"
"AGAW2","09W","INS","BOLD:AAF9771","Arthropoda","Insecta","Diptera","Hybotidae","Tachydromiinae","Platypalpus","Platypalpus longiseta, Platypalpus trivialis, Platypalpus sp. 5, Platypalpus sp. 17, Platypalpus pallidiventris","37","33","33","51","10.81","0","99.19","443.92","0","444.78","34.77","0"
"AGAW2","09W","INS","BOLD:ACK3142","Arthropoda","Insecta","Diptera","Cecidomyiidae","","","","37","36","36","51","2.7","0","98.41","438.05","0","440.05","28.47","0"
"AGAW2","09W","INS","BOLD:ACV5892","Arthropoda","Insecta","Diptera","Cecidomyiidae","","","","34","31","31","53","8.82","0","97.8","447.18","0","448.82","25.44","0"
"AGAW2","09W","INS","BOLD:ABW1379","Arthropoda","Insecta","Diptera","Chloropidae","Oscinellinae","Malloewia","Malloewia abdominalis","33","30","30","54","9.09","0","99.27","434.33","0","441.88","31.45","0"
"AGAW2","09W","INS","BOLD:AAM6037","Arthropoda","Insecta","Diptera","Cecidomyiidae","Cecidomyiinae","Contarinia, Dasineura","","29","25","25","55","13.79","0","98.29","445.72","0","447.17","26.59","0"
"AGAW2","09W","INS","BOLD:ACL8053","Arthropoda","Insecta","Diptera","Chironomidae","Chironominae","Tanytarsus","","29","29","29","55","0","0","99.13","446.48","0","448.28","33.59","0"
"AGAW2","09W","INS","BOLD:ABA1215","Arthropoda","Insecta","Diptera","Sciaridae","","Lycoriella","Lycoriella sativae","27","25","25","57","7.41","0","99.26","437.07","0","438.63","35.7","0"
"AGAW2","09W","INS","BOLD:ACH3153","Arthropoda","Insecta","Hymenoptera","Ichneumonidae","Diplazontinae","Woldstedtius","Woldstedtius biguttatus","25","24","24","58","4","0","99.06","436.96","0","438.52","28.91","0"
"AGAW2","09W","INS","BOLD:ABA9946","Arthropoda","Insecta","Coleoptera","Cryptophagidae","Atomariinae","Atomaria","Atomaria fuscata","24","19","19","59","20.83","0","99.42","442.17","0","443.5","31.93","0"
"AGAW2","09W","INS","BOLD:AAF5980","Arthropoda","Insecta","Hemiptera","Cicadellidae","Typhlocybinae","Typhlocyba","Typhlocyba pomaria","22","20","20","60","9.09","0","97.92","429.95","0","430.82","37.14","0"
"AGAW2","09W","INS","BOLD:AAB8851","Arthropoda","Insecta","Diptera","Drosophilidae","Drosophilinae","Drosophila","Drosophila affinis, Drosophila algonquin","17","14","14","61","17.65","0","99.58","448.82","0","450","32.69","0"
"AGAW2","09W","INS","BOLD:AAM9255","Arthropoda","Insecta","Diptera","Sciaridae","","Pseudolycoriella","","17","17","17","61","0","0","99.05","413.94","0","416.53","30.7","0"
"AGAW2","09W","INS","BOLD:AAG8091","Arthropoda","Insecta","Hymenoptera","Ichneumonidae","Campopleginae","Campoletis","Campoletis sp. AB.2","15","15","15","63","0","0","99.15","430","0","431.8","24.53","0"
"AGAW2","09W","INS","BOLD:AAB4894","Arthropoda","Insecta","Hemiptera","Aphididae","Aphidinae","Sitobion","Sitobion smilacifoliae, Sitobion avenae, Sitobion phyllanthi","14","14","14","64","0","0","98.26","446.57","0","447","25.58","0"
"AGAW2","09W","INS","BOLD:AAV1136","Arthropoda","Insecta","Diptera","Scatopsidae","","","","13","13","13","65","0","0","99.2","429.15","0","431.08","36.43","0"
"AGAW2","09W","INS","BOLD:AAU0577","Arthropoda","Insecta","Thysanoptera","Aeolothripidae","Aeolothripinae","Aeolothrips","Aeolothrips fasciatus","11","11","11","66","0","0","98.04","428.91","0","439.73","37.02","0"
"AGAW2","09W","INS","BOLD:AAB9332","Arthropoda","Insecta","Hemiptera","Aphididae","Aphidinae","Nasonovia","Nasonovia ribisnigri","10","10","10","67","0","0","99.4","438.3","0","439.2","24.37","0"
"AGAW2","09W","INS","BOLD:AAN5267","Arthropoda","Insecta","Diptera","Cecidomyiidae","","","","10","9","9","67","10","0","99.53","449.6","0","450","30.58","0"
"AGAW2","09W","INS","BOLD:AAH0256","Arthropoda","Insecta","Coleoptera","Latridiidae","Corticariinae","Corticaria, Corticarina","","10","10","10","67","0","0","98.78","401.4","0","409.5","31.42","0"
"AGAW2","09W","INS","BOLD:AAN5205","Arthropoda","Insecta","Diptera","Cecidomyiidae","","","CecidInt42 sp. BOLD:AAN5205","9","9","9","70","0","0","99.5","447","0","448.11","31.61","0"
"AGAW2","09W","INS","BOLD:ACV5891","Arthropoda","Insecta","Diptera","Cecidomyiidae","","","","9","9","9","70","0","0","98.64","360.44","0","413.33","24.88","0"
"AGAW2","09W","INS","BOLD:AAV5594","Arthropoda","Insecta","Diptera","Cecidomyiidae","","","","7","7","7","72","0","0","99.56","422","0","422.86","26.33","0"
"AGAW2","09W","INS","BOLD:ABU5525","Arthropoda","Insecta","Diptera","Chironomidae","Orthocladiinae","Limnophyes","Limnophyes sp. 14ES","7","7","7","72","0","0","98.91","417","0","420","37.46","0"
"AGAW2","09W","INS","BOLD:ABU5545","Arthropoda","Insecta","Diptera","Mycetophilidae","Mycetophilinae","Cordyla","","7","7","7","72","0","0","98.85","410.14","0","449.57","37.46","0"
"AGAW2","09W","INS","BOLD:AAH3662","Arthropoda","Insecta","Diptera","Cecidomyiidae","","","CecidInt3 sp. BOLD:AAH3662","7","7","7","72","0","0","98.52","435.29","0","436.43","26.04","0"
"AGAW2","09W","INS","BOLD:ABZ4994","Arthropoda","Insecta","Orthoptera","Acrididae","Melanoplinae","Melanoplus","Melanoplus gladstoni, Melanoplus sanguinipes","7","7","7","72","0","0","97.2","449.14","0","450","36.44","0"
"AGAW2","09W","INS","BOLD:AAG6745","Arthropoda","Insecta","Diptera","Calliphoridae","Calliphorinae","Pollenia","Pollenia pediculata","7","7","7","72","0","0","99.19","439","0","440.14","33.66","0"
"AGAW2","09W","INS","BOLD:AAA7661","Arthropoda","Insecta","Diptera","Culicidae","Culicinae","Culiseta, Culex","Culiseta melanura, Culex articularis, Culex restuans, Culex acharistus","7","7","7","72","0","0","99.13","437.29","0","438.57","33.14","0"
"AGAW2","09W","INS","BOLD:AAP8739","Arthropoda","Insecta","Diptera","Platystomatidae","Platystomatinae","Rivellia","Rivellia steyskali","7","7","7","72","0","0","99.62","440","0","441.57","37.93","0"
"AGAW2","09W","INS","BOLD:AAN5225","Arthropoda","Insecta","Diptera","Cecidomyiidae","","","","6","5","5","80","16.67","0","99.52","449.5","0","450","28.37","0"
"AGAW2","09W","INS","BOLD:AAM7855","Arthropoda","Insecta","Thysanoptera","Thripidae","Thripinae","Frankliniella","Frankliniella tritici","6","6","6","80","0","0","99.37","447.33","0","450","31.48","0"
"AGAW2","09W","INS","BOLD:ACF1257","","","","","","","","6","4","4","80","33.33","0","99.96","448.17","0","448.17","34.7","0"
"AGAW2","09W","INS","BOLD:ACC8079","Arthropoda","Insecta","Diptera","Cecidomyiidae","","","","6","4","4","80","33.33","0","99.7","449.33","0","450","29.33","0"
"AGAW2","09W","INS","BOLD:AAU6966","Arthropoda","Insecta","Coleoptera","Throscidae","Throscinae","Trixagus","Trixagus chevrolati","6","6","6","80","0","0","99.64","412.33","0","450","39.56","0"
"AGAW2","09W","INS","BOLD:AAN5200","Arthropoda","Insecta","Diptera","Cecidomyiidae","","","","5","4","4","85","20","0","99.22","423.6","0","425.6","26.41","0"
"AGAW2","09W","INS","BOLD:ABV9277","","","","","","","","5","5","5","85","0","0","99.26","432.6","0","434.4","28.36","0"
"AGAW2","09W","INS","BOLD:AAB9140","Arthropoda","Insecta","Diptera","Calliphoridae","Chrysomyinae","Phormia","Phormia regina","5","5","5","85","0","0","98.93","444.4","0","444.6","32.97","0"
"AGAW2","09W","INS","BOLD:ABU5546","Arthropoda","Insecta","Diptera","Psychodidae","Psychodinae","Psychoda","","5","5","5","85","0","0","99.01","442.6","0","443.8","30.95","0"
"AGAW2","09W","INS","BOLD:AAI4303","Arthropoda","Insecta","Diptera","Chironomidae","Chironominae","Chironomus","Chironomus melanescens","5","5","5","85","0","0","98.02","432.8","0","434.2","34.31","0"
"AGAW2","09W","INS","BOLD:AAV1295","Arthropoda","Insecta","Diptera","Sciaridae","","Bradysia","Bradysia difformis, Bradysia impatiens","5","5","5","85","0","0","99.39","425","0","450","31.78","0"
"AGAW2","09W","INS","BOLD:AAA6560","Arthropoda","Insecta","Lepidoptera","Crambidae","Crambinae","Crambus","Crambus multilinellus, Crambus saltuellus","4","4","4","91","0","0","99.4","429.25","0","432.5","30.98","0"
"AGAW2","09W","INS","BOLD:AAG5453","Arthropoda","Insecta","Diptera","Chironomidae","Chironominae","Chironomus","Chironomus bifurcatus","4","4","4","91","0","0","98.82","446.75","0","450","34.61","0"
"AGAW2","09W","INS","BOLD:AAG9629","Arthropoda","Insecta","Diptera","Dolichopodidae","Dolichopodinae","Dolichopus","Dolichopus omnivagus, Dolichopus lobatus","4","4","4","91","0","0","98.52","410.75","0","413.25","38.34","0"
"AGAW2","09W","INS","BOLD:ACP4736","Arthropoda","Insecta","Diptera","Chironomidae","Orthocladiinae","Smittia","Smittia sp. 8ES","4","4","4","91","0","0","99.76","416.5","0","417.75","33.26","0"
"AGAW2","09W","INS","BOLD:AAV3714","Arthropoda","Insecta","Diptera","Hybotidae","Tachydromiinae","Micrempis","Micrempis bomboxynon","4","4","4","91","0","0","98.36","391.75","0","395.25","37.21","0"
"AGAW2","09W","INS","BOLD:AAC0022","Arthropoda","Insecta","Diptera","Chironomidae","Chironominae","Micropsectra","Micropsectra nigripila","3","3","3","96","0","0","97.31","446.67","0","450","36.89","0"
"AGAW2","09W","INS","BOLD:AAE7186","Arthropoda","Insecta","Hymenoptera","Braconidae","Aphidiinae","Praon","Praon sp. 1 MM-2013, Praon volucre, Praon bicolor","3","3","3","96","0","0","99.83","417.67","0","418","25.64","0"
"AGAW2","09W","INS","BOLD:AAA7631","Arthropoda","Insecta","Hemiptera","Miridae","Mirinae","Adelphocoris","Adelphocoris lineolatus","3","2","2","96","33.33","0","99.12","433.67","0","450","35.19","0"
"AGAW2","09W","INS","BOLD:AAO2130","Arthropoda","Insecta","Hymenoptera","Figitidae","Charipinae","Phaenoglyphis","Phaenoglyphis villosa","3","3","3","96","0","0","99.35","411","0","446.67","26.79","0"
"AGAW2","09W","INS","BOLD:AAA2815","Arthropoda","Insecta","Lepidoptera","Noctuidae","Noctuinae","Mythimna","Mythimna yukonensis, Mythimna oxygala","3","3","3","96","0","0","99.48","449.33","0","450","33.85","0"
"AGAW2","09W","INS","BOLD:ACE8878","Arthropoda","Insecta","Orthoptera","Acrididae","Melanoplinae","Melanoplus","Melanoplus scudderi group","3","3","3","96","0","0","97.7","448.33","0","450","35.93","0"
"AGAW2","09W","INS","BOLD:AAN6447","Arthropoda","Insecta","Diptera","Sciaridae","","Corynoptera","Corynoptera perpusilla","3","3","3","96","0","0","99.11","447","0","450","36.52","0"
"AGAW2","09W","INS","BOLD:ACW5121","Arthropoda","Insecta","Hymenoptera","Braconidae","Braconinae","Habrobracon","","3","3","3","96","0","0","98.59","449.67","0","450","25.41","0"
"AGAW2","09W","INS","BOLD:AAA3654","Arthropoda","Insecta","Lepidoptera","Crambidae","Crambinae","Crambus","Crambus bigelovi, Crambus johnsoni, Crambus praefectellus","3","3","3","96","0","0","98.29","449","0","450","31.85","0"
"AGAW2","09W","INS","BOLD:AAH3320","Arthropoda","Insecta","Coleoptera","Coccinellidae","Coccinellinae","Psyllobora","Psyllobora vigintimaculata","3","3","3","96","0","0","98.96","423.33","0","425.33","33.92","0"
"AGAW2","09W","INS","BOLD:AAQ0558","Arthropoda","Insecta","Thysanoptera","Thripidae","Thripinae","Anaphothrips","Anaphothrips obscurus","2","2","2","106","0","0","100","447","0","450","31.56","0"
"AGAW2","09W","INS","BOLD:ABV1415","Arthropoda","Insecta","Diptera","Chironomidae","Chironominae","","","2","2","2","106","0","0","99.65","415","0","416","32.92","0"
"AGAW2","09W","INS","BOLD:ACE4100","Arthropoda","Insecta","Orthoptera","Acrididae","Melanoplinae","Melanoplus","Melanoplus pinicola","2","2","2","106","0","0","97.1","448","0","450","35","0"
"AGAW2","09W","INS","BOLD:ACR4394","Arthropoda","Insecta","Diptera","Anthomyiidae","Anthomyiinae","Delia","Delia florilega","2","2","2","106","0","0","97.72","436","0","437.5","33.46","0"
"AGAW2","09W","INS","BOLD:AAG1717","Arthropoda","Insecta","Diptera","Muscidae","Phaoniinae","Helina","Helina troene","2","2","2","106","0","0","98.41","440.5","0","443.5","31.69","0"
"AGAW2","09W","INS","BOLD:ACU7906","Arthropoda","Insecta","Diptera","Chironomidae","Orthocladiinae","","","2","2","2","106","0","0","98.81","424.5","0","426","35.56","0"
"AGAW2","09W","INS","BOLD:AAA3889","Arthropoda","Insecta","Lepidoptera","Crambidae","Crambinae","Parapediasia","Parapediasia teterrellus","2","2","2","106","0","0","99.67","448.5","0","450","32.56","0"
"AGAW2","09W","INS","BOLD:ACJ0111","Arthropoda","Insecta","Hymenoptera","Platygastridae","Scelioninae","Telenomus","","2","2","2","106","0","0","98.55","404.5","0","410","27.96","0"
"AGAW2","09W","INS","BOLD:AAC0903","Arthropoda","Insecta","Diptera","Chironomidae","Chironominae","Chironomus","Chironomus acidophilus","2","2","2","106","0","0","99.55","448.5","0","450","36.33","0"
"AGAW2","09W","INS","BOLD:ABA6488","Arthropoda","Insecta","Diptera","Sciaridae","","Bradysia","Bradysia trivittata","2","2","2","106","0","0","99.38","400.5","0","404.5","32.76","0"
"AGAW2","09W","INS","BOLD:AAF9187","Arthropoda","Insecta","Coleoptera","Curculionidae","Entiminae","Phyllobius","Phyllobius oblongus","2","2","2","106","0","0","99.33","449.5","0","450","39.33","0"
"AGAW2","09W","INS","BOLD:AAH3644","Arthropoda","Insecta","Diptera","Cecidomyiidae","","","","2","2","2","106","0","0","100","449.5","0","450","27.22","0"
"AGAW2","09W","INS","BOLD:ACF4925","Arthropoda","Insecta","Hemiptera","Miridae","Mirinae","Adelphocoris","Adelphocoris detritus","2","2","2","106","0","0","97.33","449","0","450","36.56","0"
"AGAW2","09W","INS","BOLD:AAE7016","Arthropoda","Insecta","Lepidoptera","Gelechiidae","Dichomeridinae","Helcystogramma","Helcystogramma hystricella","2","2","2","106","0","0","98.75","421.5","0","421.5","31.3","0"
"AGAW2","09W","INS","BOLD:AAH0270","Arthropoda","Insecta","Coleoptera","Curculionidae","Entiminae","Trachyphloeus","Trachyphloeus bifoveolatus, Trachyphloeus angustisetulus","2","2","2","106","0","0","98.76","437.5","0","439","32.22","0"
"AGAW2","09W","INS","BOLD:AAG2740","Arthropoda","Insecta","Diptera","Ephydridae","Parydrinae","Philygria","Philygria debilis","2","2","2","106","0","0","98.16","352.5","0","352.5","32.48","0"
"AGAW2","09W","INS","BOLD:AAA5953","Arthropoda","Insecta","Lepidoptera","Gelechiidae","Gelechiinae","Coleotechnites","Coleotechnites sp., Coleotechnites piceaella, Coleotechnites laricis, Coleotechnites sp. 3, Coleotechnites carbonaria, Coleotechnites granti, Coleotechnites macleodi, Coleotechnites gibsonella, Coleotechnites coniferella, Coleotechnites martini","2","2","2","106","0","0","97.75","445.5","0","446.5","34.15","0"
"AGAW2","09W","INS","BOLD:ACE4701","Arthropoda","Insecta","Lepidoptera","Crambidae","Crambinae","Crambus","Crambus multilinellus","2","2","2","106","0","0","99.54","430.5","0","434","31.2","0"
"AGAW2","09W","INS","BOLD:AAA3181","Arthropoda","Insecta","Diptera","Muscidae","Muscinae","Stomoxys","Stomoxys calcitrans","1","1","1","124","0","0","97.97","443","0","450","32.89","0"
"AGAW2","09W","INS","BOLD:AAG9055","Arthropoda","Insecta","Hymenoptera","Vespidae","Vespinae","Vespula","Vespula germanica","1","1","1","124","0","0","97.68","431","0","431","31.32","0"
"AGAW2","09W","INS","BOLD:AAH4135","Arthropoda","Insecta","Diptera","Chloropidae","Chloropinae","Thaumatomyia","Thaumatomyia glabra, Thaumatomyia grata, Thaumatomyia sp. 1 TAW","1","1","1","124","0","0","98.44","450","0","450","32.22","0"
"AGAW2","09W","INS","BOLD:AAA6292","Arthropoda","Insecta","Lepidoptera","Tortricidae","Tortricinae","Cnephasia","Cnephasia genitalana, Cnephasia stephensiana","1","1","1","124","0","0","100","432","0","432","35.88","0"
"AGAW2","09W","INS","BOLD:AAA2590","Arthropoda","Insecta","Lepidoptera","Noctuidae","Noctuinae","Xestia","Xestia baja, Xestia smithii","1","1","1","124","0","0","99.78","450","0","450","33.11","0"
"AGAW2","09W","INS","BOLD:AAA3836","Arthropoda","Insecta","Lepidoptera","Noctuidae","Plusiinae","Autographa","Autographa precationis","1","1","1","124","0","0","99.78","448","0","450","32.22","0"
"AGAW2","09W","INS","BOLD:AAF9028","Arthropoda","Insecta","Diptera","Limoniidae","","","","1","1","1","124","0","0","97.23","433","0","450","38.67","0"
"AGAW2","09W","INS","BOLD:AAN5242","Arthropoda","Insecta","Diptera","Cecidomyiidae","","","","1","1","1","124","0","0","99.75","401","0","405","27.65","0"
"AGAW2","09W","INS","BOLD:AAA4751","Arthropoda","Insecta","Diptera","Culicidae","Culicinae","Culex","Culex pipiens sl, Culex nr. pipiens, Culex pipiens, Culex quinquefasciatus, Culex sp.","1","1","1","124","0","0","99.75","408","0","412","33.01","0"
"AGAW2","09W","INS","BOLD:AAB7869","Arthropoda","Insecta","Lepidoptera","Tortricidae","Tortricinae","Phtheochroa","Phtheochroa baracana, Phtheochroa riscana, Phtheochroa birdana, Phtheochroa fulviplicana, Phtheochroa canariana","1","1","1","124","0","0","99.78","450","0","450","29.56","0"
"AGAW2","09W","INS","BOLD:ACG3349","Arthropoda","Insecta","Diptera","Ceratopogonidae","Dasyheleinae","Dasyhelea","","1","1","1","124","0","0","97.33","450","0","450","39.11","0"
"AGAW2","09W","INS","BOLD:ACA9275","Arthropoda","Insecta","Diptera","Chironomidae","Chironominae","Chironomus","","1","1","1","124","0","0","97.66","384","0","389","35.22","0"
"AGAW2","09W","INS","BOLD:AAB4657","Arthropoda","Insecta","Diptera","Chironomidae","Chironominae","Chironomus","Chironomus maturus","1","1","1","124","0","0","98.76","402","0","450","38.67","0"
"AGAW2","09W","INS","BOLD:ACW5516","Arthropoda","Insecta","Hymenoptera","Braconidae","Braconinae","Habrobracon","","1","1","1","124","0","0","98","450","0","450","25.78","0"
"AGAW2","09W","INS","BOLD:AAB7030","Arthropoda","Insecta","Diptera","Chironomidae","Chironominae","Chironomus","Chironomus sp. BOLDAAP2996, Chironomus quinnitukqut, Chironomus cf. decorus 2g, Chironomus decorus group sp. 2 IP2013, Chironomus cf. decorus, Chironomus decorus gr.","1","1","1","124","0","0","98.93","375","0","379","33.77","0"
"AGAW2","09W","INS","BOLD:AAN5528","Arthropoda","Insecta","Diptera","Dolichopodidae","","","","1","1","1","124","0","0","98.34","421","0","425","38.82","0"
"AGAW2","09W","INS","BOLD:AAY6412","Arthropoda","Insecta","Diptera","Sciaridae","","Bradysia","Bradysia urticae","1","1","1","124","0","0","99.22","383","0","387","30.75","0"
"AGAW2","09W","INS","BOLD:AAH3914","Arthropoda","Insecta","Diptera","Sciaridae","","Sciara","Sciara hamatilis, Sciara humeralis","1","1","1","124","0","0","100","450","0","450","30.22","0"
"AGAW2","09W","INS","BOLD:AAG6435","Arthropoda","Insecta","Diptera","Ceratopogonidae","Forcipomyiinae","Forcipomyia","","1","1","1","124","0","0","98.5","399","0","445","32.81","0"
"AGAW2","09W","INS","BOLD:ACW5458","Arthropoda","Insecta","Hymenoptera","Braconidae","Braconinae","Habrobracon","","1","1","1","124","0","0","99.1","446","0","450","25.11","0"
"AGAW2","09W","INS","BOLD:AAP6421","Arthropoda","Insecta","Diptera","Phoridae","Metopininae","Pseudacteon","","1","1","1","124","0","0","98.86","351","0","352","34.09","0"
"AGAW2","09W","INS","BOLD:AAH2293","Arthropoda","Insecta","Diptera","Asilidae","Stenopogoninae","Dioctria, Eudioctria","Dioctria hyalipennis, Dioctria baumhaueri, Eudioctria propinqua","1","1","1","124","0","0","98.08","417","0","418","30.38","0"
"AGAW2","09W","INS","BOLD:ACO8630","","","","","","","","1","1","1","124","0","0","98.44","448","0","450","34","0"
"AGAW2","09W","INS","BOLD:ACK9434","Arthropoda","Insecta","Hemiptera","Cicadellidae","Macropsinae","Macropsis","Macropsis ocellata","1","1","1","124","0","0","97.77","449","0","450","34","0"
"AGAW2","09W","INS","BOLD:AAG6753","Arthropoda","Insecta","Diptera","Lauxaniidae","Lauxaniinae","Camptoprosopella","","1","1","1","124","0","0","98.79","412","0","450","35.33","0"
"AGAW2","09W","INS","BOLD:AAM6288","Arthropoda","Insecta","Diptera","Chironomidae","Chironominae","Chironomus","","1","1","1","124","0","0","100","450","0","450","35.33","0"
"AGAW2","09W","INS","BOLD:ABA8103","Arthropoda","Insecta","Hemiptera","Cicadellidae","Typhlocybinae","Empoasca","","1","1","1","124","0","0","98","449","0","450","33.11","0"
"AGAW2","09W","INS","BOLD:ACE6796","Arthropoda","Insecta","Hemiptera","Miridae","Mirinae","Taedia","Taedia scrupea","1","1","1","124","0","0","98.31","356","0","357","35.01","0"
"AGAW2","09W","INS","BOLD:AAB1098","Arthropoda","Insecta","Diptera","Culicidae","Culicinae","Ochlerotatus, Aedes","Aedes excrucians, Aedes cantans, Aedes stimulans, Aedes euedes, Aedes riparius, Aedes annulipes, Ochlerotatus cantans, Aedes fitchii, Aedes sp., Ochlerotatus annulipes, Ochlerotatus excrucians","1","1","1","124","0","0","98.5","400","0","399","34.59","0"
"AGAW2","09W","INS","BOLD:AAG2123","Arthropoda","Insecta","Diptera","Tachinidae","Dexiinae","Dinera","Dinera grisescens","1","1","1","124","0","0","98.96","384","0","386","31.35","0"
"AGAW2","09W","INS","BOLD:ABY5257","Arthropoda","Insecta","Lepidoptera","Noctuidae","Noctuinae","Apamea","Apamea devastator","1","1","1","124","0","0","97.33","450","0","450","31.33","0"
"AGAW2","09W","INS","BOLD:AAD4214","Arthropoda","Insecta","Hymenoptera","Ichneumonidae","Diplazontinae","Diplazon","Diplazon laetatorius","1","1","1","124","0","0","98.22","449","0","450","25.78","0"
"AGAW2","09W","INS","BOLD:AAB3656","Arthropoda","Insecta","Lepidoptera","Pterophoridae","Pterophorinae","Hellinsia","Hellinsia glenni, Hellinsia lacteodactylus, Hellinsia paleaceus","1","1","1","124","0","0","97.99","399","0","403","31.51","0"
"AGAW2","09W","INS","BOLD:ABX2489","Arthropoda","Insecta","Coleoptera","Phalacridae","Phalacrinae","Stilbus","","1","1","1","124","0","0","97.56","450","0","450","33.56","0"
"AGAW2","09W","INS","BOLD:ABX6186","Arthropoda","Insecta","Diptera","Tipulidae","Tipulinae","Nephrotoma","Nephrotoma ferruginea","1","1","1","124","0","0","98.43","447","0","450","34.44","0"
"AGAW2","09W","INS","BOLD:AAF4976","Arthropoda","Insecta","Coleoptera","Lampyridae","Lampyrinae","Ellychnia","Ellychnia aff. corrusca KSH611, Ellychnia sp., Ellychnia facula, Ellychnia sp. KSH1376, Ellychnia lacustris, Ellychnia corrusca, Ellychnia hatchi, Ellychnia californica","1","1","1","124","0","0","98.14","431","0","435","37.01","0"
"AGAW2","09W","INS","BOLD:AAC9062","Arthropoda","Insecta","Diptera","Culicidae","Culicinae","Ochlerotatus, Aedes","Aedes fitchii, Ochlerotatus fitchii, Aedes grossbecki, Aedes ventrovittis","1","1","1","124","0","0","99.33","447","0","450","33.11","0"
"AGAW2","09W","INS","BOLD:AAP8651","Arthropoda","Insecta","Diptera","Tachinidae","Exoristinae","Oswaldia","Oswaldia albifacies","1","1","1","124","0","0","97.77","449","0","450","32","0"
"AGAW2","09W","INS","BOLD:ABX6298","Arthropoda","Insecta","Diptera","Tipulidae","Tipulinae","Nephrotoma","Nephrotoma ferruginea","1","1","1","124","0","0","97.32","448","0","450","33.11","0"
"AGAW2","09W","INS","BOLD:ADF3937","","","","","","","","1","1","1","124","0","0","99.25","400","0","402","30.6","0"
"AGAW2","09W","INS","BOLD:ACK3431","Arthropoda","Insecta","Diptera","Cecidomyiidae","","","","1","1","1","124","0","0","99.33","450","0","450","32.89","0"
"AGAW2","09W","INS","BOLD:AAD8086","Arthropoda","Insecta","Lepidoptera","Elachistidae","Elachistinae","Elachista","Elachista griseicornis, Elachista acenteta","1","1","1","124","0","0","99.75","399","0","399","29.82","0"
"AGAW2","09W","INS","BOLD:AAC0706","Arthropoda","Insecta","Diptera","Chironomidae","Chironominae","Dicrotendipes","Dicrotendipes tritomus","1","1","1","124","0","0","98.43","383","0","386","35.75","0"
"AGAW2","09W","INS","BOLD:AAA5803","Arthropoda","Insecta","Hemiptera","Miridae","Mirinae","Lygus","Lygus lineolaris","1","1","1","124","0","0","98.21","448","0","450","34","0"
"AGAW2","09W","INS","BOLD:AAG2460","Arthropoda","Insecta","Diptera","Anthomyiidae","Anthomyiinae","Hydrophoria","Hydrophoria lancifer","1","1","1","124","0","0","97.56","450","0","450","34.67","0"
"AGAW2","09W","INS","BOLD:ACD3588","Arthropoda","Insecta","Diptera","Sciaridae","","Lycoriella","Lycoriella sativae","1","1","1","124","0","0","99.74","389","0","450","36","0"
"AGAW2","09W","NAR","no match","no match","no match","no match","no match","no match","no match","no match","no match","no match","no match","no match","no match","no match","no match","no match","no match","no match","no match","no match"
"AGAW2","09W","NIN","BOLD:AAH6630","Arthropoda","Collembola","Entomobryomorpha","Entomobryidae","Entomobryinae","Entomobrya","Entomobrya unostrigata","9905","8047","8047","1","18.76","0","99.27","437.23","0","438.53","39.09","0"
"AGAW2","09W","NIN","BOLD:AAA5267","Arthropoda","Collembola","Entomobryomorpha","Isotomidae","","","","4992","4834","4834","2","3.17","0","98.95","421.35","0","423.92","38.09","0"
"AGAW2","09W","NIN","BOLD:AAA7188","Arthropoda","Arachnida","Araneae","Philodromidae","","Tibellus","Tibellus maritimus, Tibellus oblongus","2780","2405","2405","3","13.49","0","98.21","408.79","0","411.27","31.87","0"
"AGAW2","09W","NIN","BOLD:AAI4346","Arthropoda","Arachnida","Opiliones","Phalangiidae","","Paroligolophus, Phalangium","Paroligolophus agrestis, Phalangium opilio","628","607","607","4","3.34","0","98.91","413.48","0","421.36","35.38","0"
"AGAW2","09W","NIN","BOLD:AAA8820","Arthropoda","Collembola","Entomobryomorpha","Entomobryidae","Lepidocyrtinae","Lepidocyrtus","Lepidocyrtus cyaneus","499","468","468","5","6.21","0","98.75","409.8","0","411.79","43.34","0"
"AGAW2","09W","NIN","BOLD:ADG1586","Arthropoda","Collembola","Entomobryomorpha","Entomobryidae","","","","158","150","150","6","5.06","0","99.42","398.35","0","439.58","39.98","0"
"AGAW2","09W","NIN","BOLD:AAA6381","Arthropoda","Arachnida","Araneae","Tetragnathidae","","Tetragnatha","Tetragnatha laboriosa","135","133","133","7","1.48","0","99","427.52","0","429.92","39.24","0"
"AGAW2","09W","NIN","BOLD:AAU6252","Arthropoda","Arachnida","Trombidiformes","Bdellidae","","Cyta","","66","63","63","8","4.55","0","99.41","422.47","0","423.73","31.37","0"
"AGAW2","09W","NIN","BOLD:AAB7914","Arthropoda","Collembola","Symphypleona","Bourletiellidae","","Bourletiella","","61","51","51","9","16.39","0","99.45","442.89","0","444.3","38.08","0"
"AGAW2","09W","NIN","BOLD:ABW2639","Arthropoda","Arachnida","Trombidiformes","Anystidae","","","","52","47","47","10","9.62","0","99.41","447.81","0","447.56","31.41","0"
"AGAW2","09W","NIN","BOLD:AAG1792","Arthropoda","Arachnida","Araneae","Tetragnathidae","","Glenognatha","Glenognatha foxi","21","21","21","11","0","0","99.12","444.24","0","446.05","31.62","0"
"AGAW2","09W","NIN","BOLD:AAM7961","Arthropoda","Arachnida","Trombidiformes","Anystidae","","","","15","12","12","12","20","0","99.64","451.33","0","445.47","31.17","0"
"AGAW2","09W","NIN","BOLD:ACC0359","Arthropoda","Collembola","Symphypleona","Bourletiellidae","","Bourletiella","","15","12","12","12","20","0","99.3","440.07","0","443.4","37.43","0"
"AGAW2","09W","NIN","BOLD:ACZ1295","Arthropoda","Arachnida","Trombidiformes","Anystidae","","","","8","8","8","14","0","0","99.31","374.63","0","392","31.26","0"
"AGAW2","09W","NIN","BOLD:ADF8421","Arthropoda","Collembola","Entomobryomorpha","","","","","7","7","7","15","0","0","98.1","376.43","0","415.43","44.51","0"
"AGAW2","09W","NIN","BOLD:ACC0536","Arthropoda","Arachnida","Mesostigmata","Blattisociidae","","","","6","6","6","16","0","0","99.39","423.83","0","424.83","36.08","0"
"AGAW2","09W","NIN","BOLD:AAH6568","Arthropoda","Arachnida","Sarcoptiformes","Tectocepheidae","","Tectocepheus","","3","3","3","17","0","0","98.16","419.67","0","431.33","38.77","0"
"AGAW2","09W","NIN","BOLD:AAA9476","Arthropoda","Arachnida","Araneae","Linyphiidae","","Grammonota","Grammonota gentilis","2","2","2","18","0","0","98.66","448","0","450","30.67","0"
"AGAW2","09W","NIN","BOLD:AAB0726","Arthropoda","Arachnida","Araneae","Lycosidae","","Trochosa","Trochosa ruricola","1","1","1","19","0","0","99.33","450","0","450","31.56","0"
"AGAW2","09W","NIN","BOLD:AAA7189","","","","","","","","1","1","1","19","0","0","97.49","359","0","362","31.49","0"
"AGAW2","09W","NIN","BOLD:AAH7470","Arthropoda","Diplopoda","Julida","Julidae","Julinae","Ophyiulus","Ophyiulus pilosus","1","1","1","19","0","0","99.78","449","0","450","35.78","0"
"AGAW2","09W","NIN","BOLD:AAA7242","Arthropoda","Collembola","Entomobryomorpha","Entomobryidae","Seirinae, Entomobryinae","Entomobrya, Lepidosira","Entomobrya multifasciata, Lepidosira sp.","1","1","1","19","0","0","100","449","0","450","39.56","0"
"AGAW2","10E","BAC","TAX:375853","Proteobacteria","Alphaproteobacteria","Rickettsiales","Rickettsiaceae",NA,"Wolbachia","Wolbachia sp. Hymenoptera_ASGLE646-10","152","88","88","1","42.11","0","97.88","449.26","0","449.8","40.66","0"
"AGAW2","10E","BAC","TAX:392873","Proteobacteria","Alphaproteobacteria","Rickettsiales","Rickettsiaceae",NA,"Wolbachia","Wolbachia endosymbiont of Scrapter algoensis","28","28","28","2","0","0","98.1","412.61","0","436","40.66","0"
"AGAW2","10E","BAC","TAX:91393","Proteobacteria","Alphaproteobacteria","Rickettsiales","Rickettsiaceae",NA,"Wolbachia","","17","17","17","3","0","0","99.42","387.82","0","408.41","39.41","0"
"AGAW2","10E","BAC","TAX:175768","Proteobacteria","Alphaproteobacteria","Rickettsiales","Rickettsiaceae",NA,"Wolbachia","Wolbachia pipientis","15","14","14","4","6.67","0","98.37","448.93","0","450","39.5","0"
"AGAW2","10E","BAC","TAX:376688","Proteobacteria","Alphaproteobacteria","Rickettsiales","Rickettsiaceae",NA,"Wolbachia","Wolbachia sp. elachBioLep01 BioLep373","10","10","10","5","0","0","98.18","446","0","447.5","40.67","0"
"AGAW2","10E","BAC","TAX:375807","Proteobacteria","Alphaproteobacteria","Rickettsiales","Rickettsiaceae",NA,"Wolbachia","Wolbachia sp. Cerapachys lividus_ASANV557-09","8","8","8","6","0","0","99.84","405.63","0","450","40.67","0"
"AGAW2","10E","BAC","TAX:375845","Proteobacteria","Alphaproteobacteria","Rickettsiales","Rickettsiaceae",NA,"Wolbachia","Wolbachia sp. Hymenoptera_ASGLE1440-10","7","7","7","7","0","0","98.31","438.71","0","442.29","40.79","0"
"AGAW2","10E","BAC","TAX:375772","Proteobacteria","Alphaproteobacteria","Rickettsiales","Rickettsiaceae",NA,"Wolbachia","Wolbachia sp. Leptogenys JCR25_ASANE483-10","5","5","5","8","0","0","97.49","446.6","0","450","40.49","0"
"AGAW2","10E","BAC","TAX:392874","Proteobacteria","Alphaproteobacteria","Rickettsiales","Rickettsiaceae",NA,"Wolbachia","Wolbachia endosymbiont of Hymenoptera","5","5","5","8","0","0","98.07","446","0","449.4","40.45","0"
"AGAW2","10E","BAC","TAX:375782","Proteobacteria","Alphaproteobacteria","Rickettsiales","Rickettsiaceae",NA,"Wolbachia","Wolbachia sp. Cerapachys L_MG11_ASANH426-10","3","3","3","10","0","0","98.07","448.33","0","450","40.74","0"
"AGAW2","10E","BAC","TAX:91391","Proteobacteria","Alphaproteobacteria","Rickettsiales","",NA,"","","3","3","3","10","0","0","97.6","429.67","0","437.67","40.29","0"
"AGAW2","10E","BAC","TAX:375847","Proteobacteria","Alphaproteobacteria","Rickettsiales","Rickettsiaceae",NA,"Wolbachia","Wolbachia sp. Hymenoptera_ASGLE1705-10","2","2","2","12","0","0","97.96","392","0","394.5","39.29","0"
"AGAW2","10E","BAC","TAX:392917","Proteobacteria","Alphaproteobacteria","Rickettsiales","Rickettsiaceae",NA,"Wolbachia","Wolbachia endosymbiont of Glyptapanteles Whitfield131","2","2","2","12","0","0","97.98","444.5","0","448","40.63","0"
"AGAW2","10E","BAC","TAX:392881","Proteobacteria","Alphaproteobacteria","Rickettsiales","Rickettsiaceae",NA,"Wolbachia","Wolbachia endosymbiont of Blastophaga sp. 1-ERE","2","2","2","12","0","0","98","425","0","430","40.1","0"
"AGAW2","10E","BAC","TAX:375841","Proteobacteria","Alphaproteobacteria","Rickettsiales","Rickettsiaceae",NA,"Wolbachia","Wolbachia sp. Pheidole fervens_ASPNA897-10","1","1","1","15","0","0","97.55","449","0","450","40.89","0"
"AGAW2","10E","BAC","TAX:392914","Proteobacteria","Alphaproteobacteria","Rickettsiales","Rickettsiaceae",NA,"Wolbachia","Wolbachia endosymbiont of chalJanzen01 Janzen01","1","1","1","15","0","0","98.21","446","0","450","41.11","0"
"AGAW2","10E","BAC","TAX:375810","Proteobacteria","Alphaproteobacteria","Rickettsiales","Rickettsiaceae",NA,"Wolbachia","Wolbachia sp. Cerapachys L_MG10_ASANV571-09","1","1","1","15","0","0","97.78","450","0","450","41.11","0"
"AGAW2","10E","BAC","TAX:392837","Proteobacteria","Alphaproteobacteria","Rickettsiales","Rickettsiaceae",NA,"Wolbachia","Wolbachia endosymbiont of Polyctor polyctor","1","1","1","15","0","0","97.58","414","0","424","39.62","0"
"AGAW2","10E","BAC","TAX:265077","Proteobacteria","Alphaproteobacteria","Rickettsiales","Rickettsiaceae",NA,"Wolbachia","Wolbachia sp. FormicidaeASPNA479-09","1","1","1","15","0","0","97.26","438","0","440","42.05","0"
"AGAW2","10E","BAC","TAX:453459","Proteobacteria","Alphaproteobacteria","Rickettsiales","Rickettsiaceae",NA,"Wolbachia","Wolbachia endosymbiont of Ichneumonidae","1","1","1","15","0","0","98.22","449","0","450","40.89","0"
"AGAW2","10E","BAC","TAX:453453","Proteobacteria","Alphaproteobacteria","Rickettsiales","Rickettsiaceae",NA,"Wolbachia","Wolbachia endosymbiont of Rogas luteus","1","1","1","15","0","0","97.54","447","0","450","40.89","0"
"AGAW2","10E","BAC","TAX:375763","Proteobacteria","Alphaproteobacteria","Rickettsiales","Rickettsiaceae",NA,"Wolbachia","Wolbachia sp. Pheidole nebulosa_ASLAM107-08","1","1","1","15","0","0","97.11","415","0","450","40.44","0"
"AGAW2","10E","CHO","no match","no match","no match","no match","no match","no match","no match","no match","no match","no match","no match","no match","no match","no match","no match","no match","no match","no match","no match","no match"
"AGAW2","10E","INS","BOLD:AAG1711","Arthropoda","Insecta","Diptera","Muscidae","Phaoniinae","Helina","Helina depuncta","17017","13376","13376","1","21.4","0","99.46","432.76","0","435.12","31.34","0"
"AGAW2","10E","INS","BOLD:AAA3453","Arthropoda","Insecta","Diptera","Anthomyiidae","Anthomyiinae","Delia","","7556","6667","6667","2","11.77","0","99.4","428.56","0","431.41","34.59","0"
"AGAW2","10E","INS","BOLD:AAG5499","Arthropoda","Insecta","Diptera","Chironomidae","Orthocladiinae","Psectrocladius","Psectrocladius obvius, Psectrocladius platypus","6034","4373","4373","3","27.53","0","99.39","440.62","0","442.64","35.52","0"
"AGAW2","10E","INS","BOLD:AAI4303","Arthropoda","Insecta","Diptera","Chironomidae","Chironominae","Chironomus","Chironomus melanescens","5671","4892","4892","4","13.74","0","98.8","429.84","0","434.13","34.2","0"
"AGAW2","10E","INS","BOLD:AAN6154","Arthropoda","Insecta","Coleoptera","Latridiidae","Corticariinae","Melanophthalma","Melanophthalma inermis","5006","4185","4185","5","16.4","0","99.04","435.49","0","437.12","31.26","0"
"AGAW2","10E","INS","BOLD:ACE7444","Arthropoda","Insecta","Hemiptera","Miridae","Mirinae, Phylinae","Adelphocoris, Megalocoleus","Adelphocoris lineolatus, Megalocoleus molliculus","2974","1968","1968","6","33.83","0","99.71","440.45","0","441.23","34.81","0"
"AGAW2","10E","INS","BOLD:ABY5318","Arthropoda","Insecta","Diptera","Chloropidae","Oscinellinae","Incertella, Rhopalopterum","Rhopalopterum umbrosum, Rhopalopterum luteiceps, Incertella bispina, Rhopalopterum painteri","2513","1877","1877","7","25.31","0","99.34","443.58","0","444.82","32.06","0"
"AGAW2","10E","INS","BOLD:AAU6681","Arthropoda","Insecta","Diptera","Tipulidae","Tipulinae","Nephrotoma","Nephrotoma cornicina","2111","1617","1617","8","23.4","0","99.64","442.5","0","443.73","36.3","0"
"AGAW2","10E","INS","BOLD:AAG2898","Arthropoda","Insecta","Hemiptera","Cicadellidae","Deltocephalinae","Paraphlepsius","Paraphlepsius apertus, Paraphlepsius irroratus","1841","1386","1386","9","24.71","0","99.63","440.5","0","441.47","31.86","0"
"AGAW2","10E","INS","BOLD:AAC0903","Arthropoda","Insecta","Diptera","Chironomidae","Chironominae","Chironomus","Chironomus acidophilus","1610","1269","1269","10","21.18","0","99.56","436.34","0","438.73","36.3","0"
"AGAW2","10E","INS","BOLD:AAA6759","Arthropoda","Insecta","Lepidoptera","Hesperiidae","Hesperiinae","Thymelicus","Thymelicus lineola","1525","1305","1305","11","14.43","0","99.51","437.37","0","439.24","29.93","0"
"AGAW2","10E","INS","BOLD:AAG2868","Arthropoda","Insecta","Hemiptera","Cicadellidae","Typhlocybinae","Empoasca","Empoasca sp., Empoasca fabae","1364","868","868","12","36.36","0","99.73","446.07","0","447.72","32.42","0"
"AGAW2","10E","INS","BOLD:AAA7661","Arthropoda","Insecta","Diptera","Culicidae","Culicinae","Culiseta, Culex","Culiseta melanura, Culex articularis, Culex restuans, Culex acharistus","1191","1086","1086","13","8.82","0","99.25","432.73","0","434.85","32.8","0"
"AGAW2","10E","INS","BOLD:AAG5507","Arthropoda","Insecta","Diptera","Chironomidae","Chironominae","Chironomus","Chironomus atrella","1098","1060","1060","14","3.46","0","98.81","425.2","0","428.74","35.79","0"
"AGAW2","10E","INS","BOLD:AAV5575","Arthropoda","Insecta","Diptera","Cecidomyiidae","","","CecidInt30 sp. BOLD:AAV5575","1029","741","741","15","27.99","0","99.56","442.95","0","444.15","25.9","0"
"AGAW2","10E","INS","BOLD:ACG3646","Arthropoda","Insecta","Coleoptera","Corylophidae","Corylophinae","Orthoperus","","941","722","722","16","23.27","0","99.12","442.34","0","444.29","33.6","0"
"AGAW2","10E","INS","BOLD:AAH3661","Arthropoda","Insecta","Diptera","Cecidomyiidae","","","","886","803","803","17","9.37","0","99.25","429.8","0","431.65","25.35","0"
"AGAW2","10E","INS","BOLD:AAK8751","Arthropoda","Insecta","Diptera","Agromyzidae","Phytomyzinae","Liriomyza","Liriomyza sp. SB, Liriomyza trifoliearum","809","697","697","18","13.84","0","98.97","435.17","0","436.47","32.61","0"
"AGAW2","10E","INS","BOLD:AAB4657","Arthropoda","Insecta","Diptera","Chironomidae","Chironominae","Chironomus","Chironomus maturus","776","662","662","19","14.69","0","99.52","435.62","0","438.2","37.6","0"
"AGAW2","10E","INS","BOLD:AAN5199","Arthropoda","Insecta","Diptera","Cecidomyiidae","","","CecidInt20 sp. BOLD:AAN5199","717","545","545","20","23.99","0","99.56","439.99","0","445.17","26.57","0"
"AGAW2","10E","INS","BOLD:ABA7198","Arthropoda","Insecta","Diptera","Xylomyidae","","Xylomya","Xylomya aterrima, Xylomya tenthredinoides","707","566","566","21","19.94","0","99.44","438.16","0","439.96","34.7","0"
"AGAW2","10E","INS","BOLD:AAK5928","Arthropoda","Insecta","Coleoptera","Chrysomelidae","Criocerinae","Oulema","Oulema melanopus","684","557","557","22","18.57","0","99.63","440.06","0","441.38","36.48","0"
"AGAW2","10E","INS","BOLD:ABW7897","Arthropoda","Insecta","Diptera","Cecidomyiidae","","","","681","502","502","23","26.28","0","99.65","440.68","0","443.27","29.62","0"
"AGAW2","10E","INS","BOLD:ABW1379","Arthropoda","Insecta","Diptera","Chloropidae","Oscinellinae","Malloewia","Malloewia abdominalis","525","428","428","24","18.48","0","98.36","445.22","0","446.94","31.54","0"
"AGAW2","10E","INS","BOLD:AAF9771","Arthropoda","Insecta","Diptera","Hybotidae","Tachydromiinae","Platypalpus","Platypalpus longiseta, Platypalpus trivialis, Platypalpus sp. 5, Platypalpus sp. 17, Platypalpus pallidiventris","484","368","368","25","23.97","0","99.55","440.09","0","440.89","35.05","0"
"AGAW2","10E","INS","BOLD:AAG3273","Arthropoda","Insecta","Diptera","Phoridae","Metopininae","Megaselia","Megaselia albicaudata","408","307","307","26","24.75","0","99.55","443.31","0","443.89","32.76","0"
"AGAW2","10E","INS","BOLD:AAJ4555","Arthropoda","Insecta","Diptera","Conopidae","Stylogastrinae","Stylogaster","Stylogaster neglecta","392","306","306","27","21.94","0","99.6","436.34","0","437.61","34.29","0"
"AGAW2","10E","INS","BOLD:ACC8079","Arthropoda","Insecta","Diptera","Cecidomyiidae","","","","385","276","276","28","28.31","0","99.53","446.14","0","448.01","29.2","0"
"AGAW2","10E","INS","BOLD:AAN5242","Arthropoda","Insecta","Diptera","Cecidomyiidae","","","","383","380","380","29","0.78","0","98.9","404.71","0","406.93","28.12","0"
"AGAW2","10E","INS","BOLD:AAA9422","Arthropoda","Insecta","Hemiptera","Cicadellidae","Deltocephalinae","Macrosteles","Macrosteles quadrilineatus","370","361","361","30","2.43","0","98.79","406.88","0","415.55","36.2","0"
"AGAW2","10E","INS","BOLD:AAH3664","Arthropoda","Insecta","Diptera","Cecidomyiidae","","","CecidInt3 sp. BOLD:AAH3664","365","350","350","31","4.11","0","99.09","425.96","0","429.35","26.94","0"
"AGAW2","10E","INS","BOLD:ABA0863","Arthropoda","Insecta","Diptera","Cecidomyiidae","","","","364","317","317","32","12.91","0","99.44","438.15","0","439.59","29.73","0"
"AGAW2","10E","INS","BOLD:AAN5205","Arthropoda","Insecta","Diptera","Cecidomyiidae","","","CecidInt42 sp. BOLD:AAN5205","302","268","268","33","11.26","0","99.07","437.82","0","439.33","31.44","0"
"AGAW2","10E","INS","BOLD:AAB9516","Arthropoda","Insecta","Diptera","Chironomidae","Orthocladiinae","Acricotopus","","302","273","273","33","9.6","0","98.47","421.96","0","427.78","36.48","0"
"AGAW2","10E","INS","BOLD:ACK3431","Arthropoda","Insecta","Diptera","Cecidomyiidae","","","","287","270","270","35","5.92","0","99.09","422","0","423.81","32.7","0"
"AGAW2","10E","INS","BOLD:AAG7891","Arthropoda","Insecta","Hymenoptera","Platygastridae","Scelioninae","Telenomus","Telenomus podisi","275","228","228","36","17.09","0","99.04","428.15","0","436.95","27.16","0"
"AGAW2","10E","INS","BOLD:ACU7906","Arthropoda","Insecta","Diptera","Chironomidae","Orthocladiinae","","","271","271","271","37","0","0","98.9","409.54","0","412.31","35.99","0"
"AGAW2","10E","INS","BOLD:AAN5183","Arthropoda","Insecta","Diptera","Cecidomyiidae","","","","246","209","209","38","15.04","0","98.2","441.83","0","442.88","27.8","0"
"AGAW2","10E","INS","BOLD:ACC5649","Arthropoda","Insecta","Diptera","Cecidomyiidae","","","CecidInt23 sp. BOLD:ACC5649","241","228","228","39","5.39","0","99.19","425.43","0","426.91","25.59","0"
"AGAW2","10E","INS","BOLD:AAD1485","Arthropoda","Insecta","Diptera","Chironomidae","Chironominae","Paratanytarsus","Paratanytarsus grimmii, Paratanytarsus sp.","237","198","198","40","16.46","0","99.59","443.95","0","445.26","33.82","0"
"AGAW2","10E","INS","BOLD:ABZ5071","Arthropoda","Insecta","Diptera","Chironomidae","Chironominae","Micropsectra","Micropsectra nigripila","233","228","228","41","2.15","0","98.23","423.12","0","437.97","37.05","0"
"AGAW2","10E","INS","BOLD:AAH0135","Arthropoda","Insecta","Coleoptera","Phalacridae","Phalacrinae","Acylomus","Acylomus pugetanus, Acylomus ergoti","229","210","210","42","8.3","0","99.37","437.54","0","439.58","31.66","0"
"AGAW2","10E","INS","BOLD:ABV1194","Arthropoda","Insecta","Diptera","Chironomidae","Chironominae","Micropsectra","","176","152","152","43","13.64","0","99.57","433.38","0","435.48","36.03","0"
"AGAW2","10E","INS","BOLD:ACX6055","Arthropoda","Insecta","Diptera","Phoridae","Metopininae","Megaselia","Megaselia sp. AK-2010-2, Megaselia sp. BOLD:ACX6055","167","144","144","44","13.77","0","99.47","437.42","0","439.32","33.13","0"
"AGAW2","10E","INS","BOLD:AAN5161","Arthropoda","Insecta","Diptera","Ceratopogonidae","Dasyheleinae","Dasyhelea","","165","162","162","45","1.82","0","99.06","399.12","0","401.1","37.36","0"
"AGAW2","10E","INS","BOLD:AAV5688","Arthropoda","Insecta","Diptera","Cecidomyiidae","","","","157","147","147","46","6.37","0","99.37","432.59","0","433.89","34.27","0"
"AGAW2","10E","INS","BOLD:AAM9013","Arthropoda","Insecta","Diptera","Keroplatidae","Macrocerinae","Macrocera","Macrocera pusilla","147","144","144","47","2.04","0","99.36","432.34","0","433.85","30.26","0"
"AGAW2","10E","INS","BOLD:AAN5332","Arthropoda","Insecta","Diptera","Chironomidae","Chironominae","Polypedilum","","142","142","142","48","0","0","97.77","407.54","0","411.29","33.06","0"
"AGAW2","10E","INS","BOLD:AAP6882","Arthropoda","Insecta","Diptera","Chironomidae","Chironominae","Dicrotendipes","","141","140","140","49","0.71","0","98","402.39","0","404.28","36.08","0"
"AGAW2","10E","INS","BOLD:AAO8538","Arthropoda","Insecta","Diptera","Psychodidae","Psychodinae","Psychoda","","120","118","118","50","1.67","0","98.81","424.64","0","426.95","32.76","0"
"AGAW2","10E","INS","BOLD:AAN8770","Arthropoda","Insecta","Diptera","Psychodidae","Psychodinae","Psychoda","Psychoda trinodulosa","110","100","100","51","9.09","0","99.29","437.43","0","438.68","33.64","0"
"AGAW2","10E","INS","BOLD:AAN5148","Arthropoda","Insecta","Diptera","Ceratopogonidae","Forcipomyiinae","Forcipomyia","Forcipomyia bipunctata, Forcipomyia sp. CK-2012","101","92","92","52","8.91","0","99.01","432.55","0","434.41","35.43","0"
"AGAW2","10E","INS","BOLD:ABU5525","Arthropoda","Insecta","Diptera","Chironomidae","Orthocladiinae","Limnophyes","Limnophyes sp. 14ES","90","90","90","53","0","0","99.02","397.72","0","402.02","37.12","0"
"AGAW2","10E","INS","BOLD:ABX0657","Arthropoda","Insecta","Coleoptera","Anthicidae","Notoxinae","Notoxus","Notoxus desertus","87","83","83","54","4.6","0","98.9","431.85","0","432.89","35.92","0"
"AGAW2","10E","INS","BOLD:AAZ5592","Arthropoda","Insecta","Diptera","Psychodidae","Psychodinae","","","82","80","80","55","2.44","0","99.05","428.95","0","431.8","32.41","0"
"AGAW2","10E","INS","BOLD:AAN5328","Arthropoda","Insecta","Diptera","Chironomidae","Chironominae","Micropsectra","","80","79","79","56","1.25","0","99.23","415.24","0","418.34","35.83","0"
"AGAW2","10E","INS","BOLD:ACE9668","Arthropoda","Insecta","Hymenoptera","Ichneumonidae","Ophioninae","Enicospilus","Enicospilus purgatusDHJ02","76","70","70","57","7.89","0","99.02","441.09","0","442.54","25.71","0"
"AGAW2","10E","INS","BOLD:AAG2740","Arthropoda","Insecta","Diptera","Ephydridae","Parydrinae","Philygria","Philygria debilis","72","72","72","58","0","0","97.89","399.72","0","401.76","32.18","0"
"AGAW2","10E","INS","BOLD:ACL1796","Arthropoda","Insecta","Lepidoptera","Ypsolophidae","Ochsenheimeriinae","Ochsenheimeria","Ochsenheimeria vacculella","59","59","59","59","0","0","98.96","407.75","0","414.37","32.11","0"
"AGAW2","10E","INS","BOLD:AAN5230","Arthropoda","Insecta","Diptera","Cecidomyiidae","","","","59","59","59","59","0","0","99.19","420.29","0","422.07","25.77","0"
"AGAW2","10E","INS","BOLD:ABV1297","Arthropoda","Insecta","Diptera","Sciaridae","","","","59","58","58","59","1.69","0","99.04","424.08","0","426.86","30.04","0"
"AGAW2","10E","INS","BOLD:ACM7774","Arthropoda","Insecta","Trichoptera","Hydroptilidae","Orthotrichiinae","Orthotrichia","Orthotrichia cristata","54","54","54","62","0","0","99.19","405.15","0","406.22","29.17","0"
"AGAW2","10E","INS","BOLD:AAV5609","Arthropoda","Insecta","Diptera","Cecidomyiidae","","","","53","42","42","63","20.75","0","99.13","446.91","0","447.53","27.81","0"
"AGAW2","10E","INS","BOLD:AAG8091","Arthropoda","Insecta","Hymenoptera","Ichneumonidae","Campopleginae","Campoletis","Campoletis sp. AB.2","52","52","52","64","0","0","99.17","435.6","0","437.94","24.58","0"
"AGAW2","10E","INS","BOLD:ABA1661","Arthropoda","Insecta","Coleoptera","Chrysomelidae","Galerucinae","Longitarsus","Longitarsus pellucidus","52","48","48","64","7.69","0","99.28","437.17","0","438.38","35.58","0"
"AGAW2","10E","INS","BOLD:ACK3142","Arthropoda","Insecta","Diptera","Cecidomyiidae","","","","42","42","42","66","0","0","98.25","435.76","0","437.02","27.98","0"
"AGAW2","10E","INS","BOLD:ACK1651","Arthropoda","Insecta","Diptera","Cecidomyiidae","","","","40","35","35","67","12.5","0","98.94","439.5","0","440.33","27.05","0"
"AGAW2","10E","INS","BOLD:ABV1337","Arthropoda","Insecta","Diptera","Cecidomyiidae","","","","38","34","34","68","10.53","0","98.91","432.92","0","437.63","28.43","0"
"AGAW2","10E","INS","BOLD:AAH3227","Arthropoda","Insecta","Psocodea","Lachesillidae","Lachesillinae","Lachesilla","","38","38","38","68","0","0","99.06","431","0","432.45","39.87","0"
"AGAW2","10E","INS","BOLD:AAN5249","Arthropoda","Insecta","Diptera","Cecidomyiidae","","","","36","33","33","70","8.33","0","99.13","441.58","0","443.31","33.06","0"
"AGAW2","10E","INS","BOLD:AAD2041","Arthropoda","Insecta","Diptera","Dolichopodidae","Sciapodinae","Condylostylus","Condylostylus inermis, Condylostylus nigrofemoratus, Condylostylus connectans, Condylostylus caudatus, Condylostylus flavipes","35","32","32","71","8.57","0","99.66","436.57","0","438.17","35.81","0"
"AGAW2","10E","INS","BOLD:AAG9626","Arthropoda","Insecta","Diptera","Dolichopodidae","","","","33","33","33","72","0","0","97.6","433.76","0","435.91","36.34","0"
"AGAW2","10E","INS","BOLD:ABV1475","Arthropoda","Insecta","Diptera","Cecidomyiidae","","","","33","30","30","72","9.09","0","99.27","445.67","0","447.03","26.28","0"
"AGAW2","10E","INS","BOLD:AAN5220","Arthropoda","Insecta","Diptera","Cecidomyiidae","","","","32","32","32","74","0","0","98.54","399.34","0","409.66","25.43","0"
"AGAW2","10E","INS","BOLD:AAM6037","Arthropoda","Insecta","Diptera","Cecidomyiidae","Cecidomyiinae","Contarinia, Dasineura","","31","30","30","75","3.23","0","98.33","440.68","0","441.71","26.54","0"
"AGAW2","10E","INS","BOLD:AAV5796","Arthropoda","Insecta","Diptera","Cecidomyiidae","","","","27","26","26","76","3.7","0","99.57","440.11","0","441.56","29.09","0"
"AGAW2","10E","INS","BOLD:AAP8739","Arthropoda","Insecta","Diptera","Platystomatidae","Platystomatinae","Rivellia","Rivellia steyskali","27","26","26","76","3.7","0","99.08","441.59","0","443.96","37.94","0"
"AGAW2","10E","INS","BOLD:AAD7604","Arthropoda","Insecta","Coleoptera","Coccinellidae","Coccinellinae","Coleomegilla","Coleomegilla maculata","26","25","25","78","3.85","0","99.02","437.69","0","440.12","31.83","0"
"AGAW2","10E","INS","BOLD:AAM9347","Arthropoda","Insecta","Diptera","Phoridae","Metopininae","Megaselia","","24","24","24","79","0","0","98.26","405.5","0","407.33","33.14","0"
"AGAW2","10E","INS","BOLD:ABA7871","Arthropoda","Insecta","Diptera","Cecidomyiidae","","","","21","21","21","80","0","0","99.02","441.9","0","442.95","26.36","0"
"AGAW2","10E","INS","BOLD:AAG6745","Arthropoda","Insecta","Diptera","Calliphoridae","Calliphorinae","Pollenia","Pollenia pediculata","19","19","19","81","0","0","99.23","422.84","0","424.89","33.71","0"
"AGAW2","10E","INS","BOLD:ACX5521","Arthropoda","Insecta","Diptera","Chironomidae","Orthocladiinae","","","18","18","18","82","0","0","99","403.56","0","405.78","35","0"
"AGAW2","10E","INS","BOLD:AAH0134","Arthropoda","Insecta","Coleoptera","Phalacridae","Phalacrinae","Stilbus","Stilbus apicialis","18","18","18","82","0","0","99.64","420.56","0","421.67","31.85","0"
"AGAW2","10E","INS","BOLD:AAM9255","Arthropoda","Insecta","Diptera","Sciaridae","","Pseudolycoriella","","18","18","18","82","0","0","99.01","409.5","0","411.83","31.18","0"
"AGAW2","10E","INS","BOLD:AAZ5599","Arthropoda","Insecta","Diptera","Chironomidae","Orthocladiinae","","","17","14","14","85","17.65","0","99.76","439.29","0","440.59","36.65","0"
"AGAW2","10E","INS","BOLD:AAG4944","Arthropoda","Insecta","Diptera","Mycetophilidae","Sciophilinae","Sciophila","Sciophila sp. CJB1","17","17","17","85","0","0","97.7","420.82","0","423.47","33.48","0"
"AGAW2","10E","INS","BOLD:AAN5238","Arthropoda","Insecta","Diptera","Cecidomyiidae","","","","17","17","17","85","0","0","99.37","418.35","0","420.41","25.11","0"
"AGAW2","10E","INS","BOLD:ACA8173","Arthropoda","Insecta","Diptera","Phoridae","Phorinae","Diplonevra","Diplonevra funebris","16","16","16","88","0","0","99.15","419.69","0","421.81","31.93","0"
"AGAW2","10E","INS","BOLD:AAN6447","Arthropoda","Insecta","Diptera","Sciaridae","","Corynoptera","Corynoptera perpusilla","15","14","14","89","6.67","0","99.35","448.6","0","450","36.16","0"
"AGAW2","10E","INS","BOLD:AAU8379","Arthropoda","Insecta","Hymenoptera","Braconidae","Alysiinae","Dinotrema","Dinotrema JFT01","15","13","13","89","13.33","0","99.7","442.87","0","444.2","24.22","0"
"AGAW2","10E","INS","BOLD:ABX9046","Arthropoda","Insecta","Diptera","Cecidomyiidae","","","","14","14","14","91","0","0","98.55","426.86","0","428.21","25.4","0"
"AGAW2","10E","INS","BOLD:AAN5260","Arthropoda","Insecta","Diptera","Cecidomyiidae","","","","14","14","14","91","0","0","99.26","410.64","0","412.86","26.03","0"
"AGAW2","10E","INS","BOLD:ABV0470","Arthropoda","Insecta","Diptera","Cecidomyiidae","","","","13","12","12","93","7.69","0","98.71","443.23","0","444.15","25.65","0"
"AGAW2","10E","INS","BOLD:ACK5742","Arthropoda","Insecta","Diptera","Cecidomyiidae","","","","11","10","10","94","9.09","0","99.41","443.55","0","443.91","29.17","0"
"AGAW2","10E","INS","BOLD:ABV1279","Arthropoda","Insecta","Diptera","Cecidomyiidae","","","","11","10","10","94","9.09","0","99.81","444.36","0","445.55","28.86","0"
"AGAW2","10E","INS","BOLD:AAN5215","Arthropoda","Insecta","Diptera","Cecidomyiidae","","","","11","11","11","94","0","0","99.19","420.18","0","422.09","29.08","0"
"AGAW2","10E","INS","BOLD:AAU8628","Arthropoda","Insecta","Hymenoptera","Bethylidae","","","","11","11","11","94","0","0","98.99","435.27","0","438.36","27.19","0"
"AGAW2","10E","INS","BOLD:AAH1932","Arthropoda","Insecta","Hymenoptera","Ichneumonidae","Cryptinae","Dichrogaster","Dichrogaster perlae, Dichrogaster modesta","10","10","10","98","0","0","98.96","431.9","0","434.6","26.12","0"
"AGAW2","10E","INS","BOLD:AAC0022","Arthropoda","Insecta","Diptera","Chironomidae","Chironominae","Micropsectra","Micropsectra nigripila","9","7","7","99","22.22","0","97.22","448.33","0","450","37.26","0"
"AGAW2","10E","INS","BOLD:AAZ0311","Arthropoda","Insecta","Diptera","Cecidomyiidae","","","","9","9","9","99","0","0","99.35","422.78","0","424.56","25.99","0"
"AGAW2","10E","INS","BOLD:ACE7472","Arthropoda","Insecta","Diptera","Cecidomyiidae","","","","9","9","9","99","0","0","98.34","449","0","450","27.56","0"
"AGAW2","10E","INS","BOLD:ACL4508","Arthropoda","Insecta","Coleoptera","Staphylinidae","","","","9","9","9","99","0","0","99.14","425.44","0","427.33","32.51","0"
"AGAW2","10E","INS","BOLD:AAG2511","Arthropoda","Insecta","Diptera","Anthomyiidae","Anthomyiinae","Delia","Delia platura","8","8","8","103","0","0","99.04","439.13","0","441.75","33.21","0"
"AGAW2","10E","INS","BOLD:ADH9867","Arthropoda","Insecta","Diptera","","","","","8","8","8","103","0","0","98.36","385.5","0","389.88","37.03","0"
"AGAW2","10E","INS","BOLD:AAB8143","Arthropoda","Insecta","Hemiptera","Nabidae","Nabinae","Nabis","Nabis inscriptus, Nabis americoferus, Nabis alternatus, Nabis sp.","7","7","7","105","0","0","98.95","448.14","0","449.86","37.09","0"
"AGAW2","10E","INS","BOLD:AAV5400","Arthropoda","Insecta","Diptera","Cecidomyiidae","","","","7","7","7","105","0","0","97.89","427.43","0","428","29.55","0"
"AGAW2","10E","INS","BOLD:AAN5267","Arthropoda","Insecta","Diptera","Cecidomyiidae","","","","7","7","7","105","0","0","99.43","449.57","0","450","30.67","0"
"AGAW2","10E","INS","BOLD:ACM7977","Arthropoda","Insecta","Diptera","Cecidomyiidae","","","","7","7","7","105","0","0","98.28","447.71","0","450","24.1","0"
"AGAW2","10E","INS","BOLD:AAA4222","Arthropoda","Insecta","Lepidoptera","Erebidae","Hypeninae","Hypena","Hypena scabra","7","7","7","105","0","0","99.28","404.71","0","410.14","32.49","0"
"AGAW2","10E","INS","BOLD:AAH2254","Arthropoda","Insecta","Diptera","Chironomidae","Chironominae, Tanypodinae","Tanytarsus","","6","6","6","110","0","0","97.91","420","0","423.33","34.84","0"
"AGAW2","10E","INS","BOLD:AAF9306","Arthropoda","Insecta","Diptera","Psychodidae","Psychodinae","Psychoda","Psychoda alternata","6","6","6","110","0","0","98.06","446.33","0","448","31.59","0"
"AGAW2","10E","INS","BOLD:AAN5501","Arthropoda","Insecta","Diptera","Hybotidae","","","","6","6","6","110","0","0","98.67","435.67","0","437.83","34.24","0"
"AGAW2","10E","INS","BOLD:AAL7442","Arthropoda","Insecta","Diptera","Cecidomyiidae","","","","5","5","5","113","0","0","99.62","421.4","0","422.8","26.43","0"
"AGAW2","10E","INS","BOLD:AAV1136","Arthropoda","Insecta","Diptera","Scatopsidae","","","","5","5","5","113","0","0","99.14","431","0","432.2","36.22","0"
"AGAW2","10E","INS","BOLD:ACB0705","Arthropoda","Insecta","Diptera","Cecidomyiidae","","","","5","5","5","113","0","0","99.55","439.8","0","440.4","25.69","0"
"AGAW2","10E","INS","BOLD:AAY6469","Arthropoda","Insecta","Diptera","Cecidomyiidae","","","","5","5","5","113","0","0","99.36","421.2","0","422.8","25.75","0"
"AGAW2","10E","INS","BOLD:ACM0918","Arthropoda","Insecta","Diptera","Cecidomyiidae","","","","4","3","3","117","25","0","97.72","449.75","0","450","27.44","0"
"AGAW2","10E","INS","BOLD:AAN5255","Arthropoda","Insecta","Diptera","Cecidomyiidae","Cecidomyiinae","Dasineura","","4","4","4","117","0","0","99.67","413","0","450","26.83","0"
"AGAW2","10E","INS","BOLD:ABA6848","Arthropoda","Insecta","Diptera","Mycetophilidae","Mycetophilinae","Exechia","","4","4","4","117","0","0","98.87","434.25","0","436","36.17","0"
"AGAW2","10E","INS","BOLD:AAY6412","Arthropoda","Insecta","Diptera","Sciaridae","","Bradysia","Bradysia urticae","4","4","4","117","0","0","98.6","414.25","0","415.5","32.28","0"
"AGAW2","10E","INS","BOLD:AAA2518","Arthropoda","Insecta","Lepidoptera","Crambidae","Crambinae","Chrysoteuchia","Chrysoteuchia topiaria, Chrysoteuchia topiarius","4","4","4","117","0","0","99.2","436","0","437","35.34","0"
"AGAW2","10E","INS","BOLD:ACK7314","Arthropoda","Insecta","Diptera","Chironomidae","Chironominae","Polypedilum","","4","4","4","117","0","0","99.68","407.75","0","410.5","35.53","0"
"AGAW2","10E","INS","BOLD:AAG2123","Arthropoda","Insecta","Diptera","Tachinidae","Dexiinae","Dinera","Dinera grisescens","4","4","4","117","0","0","99.44","415.5","0","416.5","31.86","0"
"AGAW2","10E","INS","BOLD:ACV5892","Arthropoda","Insecta","Diptera","Cecidomyiidae","","","","4","4","4","117","0","0","98.03","431.25","0","433","24.64","0"
"AGAW2","10E","INS","BOLD:ACB8843","Arthropoda","Insecta","Diptera","Cecidomyiidae","","","","3","3","3","125","0","0","97.23","445","0","447.67","27.03","0"
"AGAW2","10E","INS","BOLD:AAU8736","Arthropoda","Insecta","Hymenoptera","Diapriidae","Belytinae","Belyta","Belyta validicornis","3","2","2","125","33.33","0","99.29","439.67","0","450","24.67","0"
"AGAW2","10E","INS","BOLD:AAG8882","Arthropoda","Insecta","Hemiptera","Pentatomidae","Pentatominae","Thyanta","Thyanta pallidovirens, Thyanta custator, Thyanta accerra","3","3","3","125","0","0","99.26","450","0","450","36.52","0"
"AGAW2","10E","INS","BOLD:AAA2482","Arthropoda","Insecta","Lepidoptera","Noctuidae","Noctuinae","Mythimna","Mythimna unipuncta","3","3","3","125","0","0","98.74","447","0","450","30.89","0"
"AGAW2","10E","INS","BOLD:ACD7485","Arthropoda","Insecta","Diptera","Conopidae","Stylogastrinae","Stylogaster","Stylogaster neglecta, Stylogaster beresfordi","3","3","3","125","0","0","97.7","432.67","0","433","33.36","0"
"AGAW2","10E","INS","BOLD:AAH3662","Arthropoda","Insecta","Diptera","Cecidomyiidae","","","CecidInt3 sp. BOLD:AAH3662","3","3","3","125","0","0","99.48","449.33","0","450","26.15","0"
"AGAW2","10E","INS","BOLD:AAA2632","Arthropoda","Insecta","Lepidoptera","Noctuidae","Noctuinae","Noctua","Noctua atlantica, Noctua pronuba","3","3","3","125","0","0","98.51","447.67","0","450","30.96","0"
"AGAW2","10E","INS","BOLD:ABA6407","Arthropoda","Insecta","Diptera","Sciaridae","","Pseudolycoriella","","3","3","3","125","0","0","99.54","423.67","0","424.33","35.35","0"
"AGAW2","10E","INS","BOLD:AAV5356","Arthropoda","Insecta","Diptera","Cecidomyiidae","","","","3","3","3","125","0","0","97.99","448","0","450","31.78","0"
"AGAW2","10E","INS","BOLD:AAA4277","Arthropoda","Insecta","Diptera","Syrphidae","Syrphinae","Toxomerus","Toxomerus marginatus, Toxomerus sp. CR-3, Toxomerus CR-3","3","3","3","125","0","0","98.58","447.33","0","450","30.89","0"
"AGAW2","10E","INS","BOLD:AAA3181","Arthropoda","Insecta","Diptera","Muscidae","Muscinae","Stomoxys","Stomoxys calcitrans","2","2","2","135","0","0","98.52","443","0","446","33.41","0"
"AGAW2","10E","INS","BOLD:AAA3836","Arthropoda","Insecta","Lepidoptera","Noctuidae","Plusiinae","Autographa","Autographa precationis","2","2","2","135","0","0","99.44","449","0","450","32.33","0"
"AGAW2","10E","INS","BOLD:ACJ6954","Arthropoda","Insecta","Diptera","Cecidomyiidae","","","","2","2","2","135","0","0","98.96","337.5","0","450","26.89","0"
"AGAW2","10E","INS","BOLD:ACC8303","Arthropoda","Insecta","Diptera","Cecidomyiidae","","","","2","2","2","135","0","0","98.37","368","0","371","25.87","0"
"AGAW2","10E","INS","BOLD:AAI6205","Arthropoda","Insecta","Lepidoptera","Elachistidae","Elachistinae","Elachista","","2","2","2","135","0","0","98.89","449","0","450","33","0"
"AGAW2","10E","INS","BOLD:AAC9132","Arthropoda","Insecta","Diptera","Culicidae","Culicinae","Culiseta","Culiseta inornata","2","2","2","135","0","0","97.81","395.5","0","401","31.8","0"
"AGAW2","10E","INS","BOLD:AAG5453","Arthropoda","Insecta","Diptera","Chironomidae","Chironominae","Chironomus","Chironomus bifurcatus","2","2","2","135","0","0","98.33","448.5","0","450","35","0"
"AGAW2","10E","INS","BOLD:AAH3305","Arthropoda","Insecta","Coleoptera","Coccinellidae","Coccinellinae","Hippodamia","Hippodamia glacialis","2","2","2","135","0","0","99.36","407.5","0","410.5","33.63","0"
"AGAW2","10E","INS","BOLD:AAG3302","Arthropoda","Insecta","Diptera","Phoridae","Metopininae","Megaselia","","2","2","2","135","0","0","98.8","414","0","419.5","31.44","0"
"AGAW2","10E","INS","BOLD:ACL3515","Arthropoda","Insecta","Diptera","Cecidomyiidae","","","","2","2","2","135","0","0","99.62","422","0","425","26.08","0"
"AGAW2","10E","INS","BOLD:AAA2326","Arthropoda","Insecta","Hymenoptera","Apidae","Apinae","Apis","Apis mellifera","2","2","2","135","0","0","98.77","447.5","0","450","26.33","0"
"AGAW2","10E","INS","BOLD:AAZ5483","Arthropoda","Insecta","Diptera","Muscidae","Phaoniinae","Helina","Helina setiventris","2","2","2","135","0","0","98.32","447.5","0","450","29.78","0"
"AGAW2","10E","INS","BOLD:ACE0554","Arthropoda","Insecta","Diptera","Phoridae","Metopininae","Megaselia","","2","2","2","135","0","0","99.89","416.5","0","419","31.45","0"
"AGAW2","10E","INS","BOLD:ACU6062","Arthropoda","Insecta","Diptera","Cecidomyiidae","","","","2","2","2","135","0","0","98.88","447","0","450","26.33","0"
"AGAW2","10E","INS","BOLD:AAB0373","Arthropoda","Insecta","Neuroptera","Chrysopidae","Chrysopinae","Chrysoperla","Chrysoperla sp. isolate 82, Chrysoperla zastrowi, Chrysoperla plorabunda, Chrysoperla sp. isolate 71, Chrysoperla agilis, Italochrysa italica, Chrysoperla sp. isolate 88, Chrysoperla lucasina, Chrysoperla carnea, Chrysoperla mediterranea, Chrysoperla pallida, Chrysoperla nipponensis, Chrysoperla sillemi","2","2","2","135","0","0","99.74","415","0","416.5","31.43","0"
"AGAW2","10E","INS","BOLD:ACN0444","Arthropoda","Insecta","Hymenoptera","Megaspilidae","","","","2","2","2","135","0","0","98.23","395","0","392.5","32.36","0"
"AGAW2","10E","INS","BOLD:AAU6512","Arthropoda","Insecta","Diptera","Cecidomyiidae","","","CecidInt45 sp. BOLD:AAU6512","2","2","2","135","0","0","99.53","431","0","435","31.6","0"
"AGAW2","10E","INS","BOLD:AAH2295","Arthropoda","Insecta","Diptera","Asilidae","Leptogastrinae","Leptogaster","Leptogaster flavipes","2","2","2","135","0","0","98.43","413","0","450","35.33","0"
"AGAW2","10E","INS","BOLD:AAH2293","Arthropoda","Insecta","Diptera","Asilidae","Stenopogoninae","Dioctria, Eudioctria","Dioctria hyalipennis, Dioctria baumhaueri, Eudioctria propinqua","2","2","2","135","0","0","98.95","418.5","0","421.5","30.47","0"
"AGAW2","10E","INS","BOLD:AAD5253","Arthropoda","Insecta","Diptera","Chironomidae","Orthocladiinae","Thienemanniella","Thienemanniella xena, Thienemanniella vittata","2","2","2","135","0","0","98.12","426","0","426.5","34.08","0"
"AGAW2","10E","INS","BOLD:AAL2750","Arthropoda","Insecta","Diptera","Muscidae","Phaoniinae","Phaonia","Phaonia tuguriorum","2","2","2","135","0","0","98.74","435.5","0","439","32.34","0"
"AGAW2","10E","INS","BOLD:AAU6966","Arthropoda","Insecta","Coleoptera","Throscidae","Throscinae","Trixagus","Trixagus chevrolati","2","2","2","135","0","0","99.64","412","0","450","39.44","0"
"AGAW2","10E","INS","BOLD:AAX3741","Arthropoda","Insecta","Diptera","Agromyzidae","Phytomyzinae","Napomyza","Napomyza cichorii","2","2","2","135","0","0","99.21","382.5","0","440","30.91","0"
"AGAW2","10E","INS","BOLD:AAA4555","Arthropoda","Insecta","Orthoptera","Acrididae","Melanoplinae","Melanoplus","Melanoplus stupefactus, Melanoplus angustipennis, Melanoplus ponderosus, Melanoplus bruneri, Melanoplus frigidus, Melanoplus scapularis, Melanoplus cinereus, Melanoplus madeleineae, Melanoplus sanguinipes, Melanoplus nigrescens group, Melanoplus tunicae, Melanoplus walshi, Melanoplus femurrubrum, Melanoplus dawsoni, Melanoplus sp., Melanoplus fasciatus, Melanoplus bowditchi, Melanoplus stonei, Melanoplus bivittatus, Melanoplus huroni, Melanoplus thomasi, Melanoplus borealis, Melanoplus packardii","2","2","2","135","0","0","99.89","448.5","0","450","36.22","0"
"AGAW2","10E","INS","BOLD:AAN5162","Arthropoda","Insecta","Diptera","Ceratopogonidae","Dasyheleinae","Dasyhelea","","1","1","1","159","0","0","99.27","412","0","419","32.46","0"
"AGAW2","10E","INS","BOLD:AAG4462","Arthropoda","Insecta","Coleoptera","Chrysomelidae","Galerucinae","Crepidodera","Crepidodera solita, Crepidodera decora, Crepidodera bella, Crepidodera heikertingeri","1","1","1","159","0","0","99.23","390","0","391","35.29","0"
"AGAW2","10E","INS","BOLD:AAG9668","Arthropoda","Insecta","Diptera","Dolichopodidae","Diaphorinae","Chrysotus","Chrysotus sp., Chrysotus sp. nr. obliquus","1","1","1","159","0","0","98.66","448","0","450","33.11","0"
"AGAW2","10E","INS","BOLD:AAH2306","Arthropoda","Insecta","Diptera","Asilidae","Asilinae","Efferia","Efferia aestuans","1","1","1","159","0","0","99.33","450","0","450","38","0"
"AGAW2","10E","INS","BOLD:AAA8933","Arthropoda","Insecta","Coleoptera","Coccinellidae","Coccinellinae","Coccinella","Coccinella septempunctata","1","1","1","159","0","0","100","445","0","450","32.44","0"
"AGAW2","10E","INS","BOLD:ACM1467","Arthropoda","Insecta","Diptera","Cecidomyiidae","","","CecidInt34 sp. BOLD:ACM1467","1","1","1","159","0","0","97.1","379","0","380","27.11","0"
"AGAW2","10E","INS","BOLD:AAA3867","Arthropoda","Insecta","Lepidoptera","Noctuidae","Noctuinae","Apamea","Apamea amputatrix","1","1","1","159","0","0","99.77","440","0","442","33.71","0"
"AGAW2","10E","INS","BOLD:AAB8271","Arthropoda","Insecta","Lepidoptera","Autostichidae","Oegoconiinae","Oegoconia","Oegoconia quadripuncta, Oegoconia deauratella","1","1","1","159","0","0","99.74","384","0","386","34.46","0"
"AGAW2","10E","INS","BOLD:ABX5883","Arthropoda","Insecta","Lepidoptera","Tortricidae","Tortricinae","Choristoneura","Choristoneura retiniana, Choristoneura lambertiana, Choristoneura biennis, Choristoneura orae, Choristoneura fumiferana, Choristoneura occidentalis, Choristoneura freemani, Choristoneura pinus, Choristoneura occidentalis group, Choristoneura sp., Choristoneura carnana","1","1","1","159","0","0","98.2","445","0","450","31.11","0"
"AGAW2","10E","INS","BOLD:AAG0734","Arthropoda","Insecta","Thysanoptera","Thripidae","Thripinae","Frankliniella","Frankliniella tritici","1","1","1","159","0","0","98.66","447","0","450","30.22","0"
"AGAW2","10E","INS","BOLD:ACL8669","Arthropoda","Insecta","Lepidoptera","Psychidae","Psychinae","Psyche","Psyche casta","1","1","1","159","0","0","97.73","440","0","450","37.11","0"
"AGAW2","10E","INS","BOLD:AAG2017","Arthropoda","Insecta","Neuroptera","Chrysopidae","Chrysopinae","Chrysopa","Chrysopa oculata","1","1","1","159","0","0","98.24","397","0","399","31.33","0"
"AGAW2","10E","INS","BOLD:AAZ5506","Arthropoda","Insecta","Diptera","Dolichopodidae","Dolichopodinae","Dolichopus","Dolichopus sp. SEB39, Dolichopus gratus","1","1","1","159","0","0","99.33","451","0","450","41.33","0"
"AGAW2","10E","INS","BOLD:AAA5299","Arthropoda","Insecta","Diptera","Chironomidae","Orthocladiinae","Cricotopus","Cricotopus laetus, Cricotopus relucens, Cricotopus trifasciatus, Cricotopus sp., Cricotopus sp. 3ES, Cricotopus sylvestris, Cricotopus glacialis","1","1","1","159","0","0","97.77","449","0","450","36.22","0"
"AGAW2","10E","INS","BOLD:ADH9796","Arthropoda","Insecta","Diptera","","","","","1","1","1","159","0","0","97.32","447","0","450","34.67","0"
"AGAW2","10E","INS","BOLD:AAA3430","Arthropoda","Insecta","Lepidoptera","Geometridae","Larentiinae","Orthonama","Orthonama obstipata","1","1","1","159","0","0","99.55","445","0","450","32.67","0"
"AGAW2","10E","INS","BOLD:AAL7359","Arthropoda","Insecta","Diptera","Chironomidae","Chironominae","Tanytarsus","","1","1","1","159","0","0","100","448","0","450","36.22","0"
"AGAW2","10E","INS","BOLD:ACE6286","Arthropoda","Insecta","Diptera","Culicidae","Culicinae","Ochlerotatus, Aedes","Aedes dorsalis, Ochlerotatus dorsalis","1","1","1","159","0","0","98.44","450","0","450","31.78","0"
"AGAW2","10E","INS","BOLD:AAN5430","Arthropoda","Insecta","Diptera","Agromyzidae","","","","1","1","1","159","0","0","97.87","422","0","450","32.89","0"
"AGAW2","10E","INS","BOLD:AAM6040","Arthropoda","Insecta","Diptera","Cecidomyiidae","","","","1","1","1","159","0","0","99.56","450","0","450","27.11","0"
"AGAW2","10E","INS","BOLD:AAG7284","Arthropoda","Insecta","Diptera","Sphaeroceridae","Limosininae","Pullimosina","Pullimosina pullula","1","1","1","159","0","0","99.77","431","0","432","33.1","0"
"AGAW2","10E","INS","BOLD:AAU7126","Arthropoda","Insecta","Coleoptera","Cleridae","Hydnocerinae","Isohydnocera","Isohydnocera curtipennis","1","1","1","159","0","0","100","449","0","450","33.11","0"
"AGAW2","10E","INS","BOLD:ACI9753","Arthropoda","Insecta","Diptera","Dolichopodidae","Diaphorinae","Chrysotus","","1","1","1","159","0","0","97.51","362","0","367","31.61","0"
"AGAW2","10E","INS","BOLD:AAA5206","Arthropoda","Insecta","Lepidoptera","Erebidae","Herminiinae","Zanclognatha","Zanclognatha cruralis, Zanclognatha sp. 1, Zanclognatha jacchusalis, Zanclognatha minoralis, Zanclognatha minoralis sp. 1, Zanclognatha sp., Zanclognatha minoralis sp. 3, Zanclognatha gypsalis, Zanclognatha dentata, Zanclognatha theralis, Zanclognatha protumnusalis, Zanclognatha martha, Zanclognatha atrilineella, Zanclognatha obscuripennis, Zanclognatha minoralis sp. 2, Zanclognatha marcidilinea","1","1","1","159","0","0","99.11","448","0","450","31.56","0"
"AGAW2","10E","INS","BOLD:AAB1850","Arthropoda","Insecta","Hemiptera","Aphrophoridae","Aphrophorinae","Philaenus","Philaenus tesselatus, Philaenus spumarius, Philaenus maghresignus","1","1","1","159","0","0","99.33","449","0","450","33.33","0"
"AGAW2","10E","INS","BOLD:ACF0890","Arthropoda","Insecta","Hymenoptera","Ichneumonidae","Ophioninae","Enicospilus","Enicospilus purgatusDHJ02","1","1","1","159","0","0","97.55","449","0","450","26","0"
"AGAW2","10E","INS","BOLD:AAF9187","Arthropoda","Insecta","Coleoptera","Curculionidae","Entiminae","Phyllobius","Phyllobius oblongus","1","1","1","159","0","0","100","373","0","374","39.04","0"
"AGAW2","10E","INS","BOLD:AAJ7123","Arthropoda","Insecta","Diptera","Culicidae","Culicinae","Culiseta","Culiseta minnesotae","1","1","1","159","0","0","99.11","450","0","450","35.33","0"
"AGAW2","10E","INS","BOLD:ADH6991","Arthropoda","Insecta","Diptera","Cecidomyiidae","","","","1","1","1","159","0","0","97.91","382","0","385","26.49","0"
"AGAW2","10E","INS","BOLD:AAN5342","Arthropoda","Insecta","Diptera","Chironomidae","Orthocladiinae","Heleniella","","1","1","1","159","0","0","97.1","414","0","416","34.13","0"
"AGAW2","10E","INS","BOLD:AAB0651","Arthropoda","Insecta","Lepidoptera","Gelechiidae","Dichomeridinae","Dichomeris","Dichomeris leuconotella, Dichomeris gnoma","1","1","1","159","0","0","99.78","447","0","450","33.33","0"
"AGAW2","10E","INS","BOLD:AAE0845","Arthropoda","Insecta","Hemiptera","Pentatomidae","Pentatominae","Euschistus","Euschistus servus","1","1","1","159","0","0","98.8","415","0","416","37.02","0"
"AGAW2","10E","INS","BOLD:ABX2489","Arthropoda","Insecta","Coleoptera","Phalacridae","Phalacrinae","Stilbus","","1","1","1","159","0","0","97.3","407","0","407","32.43","0"
"AGAW2","10E","INS","BOLD:AAS1281","Arthropoda","Insecta","Diptera","Chironomidae","Chironominae","Chironomus","Chironomus nr. sollicitus","1","1","1","159","0","0","97.11","415","0","420","32.86","0"
"AGAW2","10E","INS","BOLD:AAA7374","Arthropoda","Insecta","Diptera","Syrphidae","Syrphinae","Sphaerophoria","Sphaerophoria bankowskae, Sphaerophoria infuscata, Sphaerophoria nr. philanthus, Sphaerophoria batava, Sphaerophoria sp., Sphaerophoria rueppellii, Sphaerophoria interrupta-group, Sphaerophoria sp. MA-1, Sphaerophoria cleoae, Sphaerophoria asymmetrica, Sphaerophoria fatarum, Sphaerophoria bengalensis, Sphaerophoria bifurcata, Sphaerophoria taeniata, Sphaerophoria interrupta, Sphaerophoria contigua, Sphaerophoria kaa, Sphaerophoria laurae, Sphaerophoria chongjini, Sphaerophoria boreoalpina, Sphaerophoria sp. BOLDAAA7374, Sphaerophoria macrogaster, Sphaerophoria sulphuripes, Sphaerophoria brevipilosa, Sphaerophoria philanthus, Sphaerophoria pyrrhina, Sphaerophoria abbreviata, Sphaerophoria scripta, Sphaerophoria potentillae, Sphaerophoria longipilosa, Sphaerophoria virgata","1","1","1","159","0","0","98.21","448","0","450","31.78","0"
"AGAW2","10E","INS","BOLD:ABV1237","Arthropoda","Insecta","Diptera","Chironomidae","Chironominae","Chironomus","","1","1","1","159","0","0","97.56","450","0","450","34.89","0"
"AGAW2","10E","INS","BOLD:ACF4925","Arthropoda","Insecta","Hemiptera","Miridae","Mirinae","Adelphocoris","Adelphocoris detritus","1","1","1","159","0","0","97.09","447","0","450","36.22","0"
"AGAW2","10E","INS","BOLD:AAB5640","Arthropoda","Insecta","Coleoptera","Coccinellidae","Coccinellinae","Harmonia","Harmonia axyridis","1","1","1","159","0","0","100","449","0","450","32.44","0"
"AGAW2","10E","INS","BOLD:ACE7482","Arthropoda","Insecta","Hemiptera","Miridae","","","","1","1","1","159","0","0","98.67","450","0","450","35.11","0"
"AGAW2","10E","INS","BOLD:ADF3937","","","","","","","","1","1","1","159","0","0","100","369","0","371","31.27","0"
"AGAW2","10E","INS","BOLD:AAB6095","Arthropoda","Insecta","Lepidoptera","Noctuidae","Noctuinae","Neoligia","Neoligia exhausta","1","1","1","159","0","0","99.19","372","0","376","32.98","0"
"AGAW2","10E","INS","BOLD:AAG2460","Arthropoda","Insecta","Diptera","Anthomyiidae","Anthomyiinae","Hydrophoria","Hydrophoria lancifer","1","1","1","159","0","0","97.89","426","0","427","34.66","0"
"AGAW2","10E","INS","BOLD:AAH3942","Arthropoda","Insecta","Diptera","Sciaridae","","","","1","1","1","159","0","0","99.76","425","0","427","32.32","0"
"AGAW2","10E","INS","BOLD:AAA6337","Arthropoda","Insecta","Lepidoptera","Noctuidae","Plusiinae","Anagrapha","Anagrapha falcifera","1","1","1","159","0","0","100","450","0","450","32.67","0"
"AGAW2","10E","INS","BOLD:AAA5803","Arthropoda","Insecta","Hemiptera","Miridae","Mirinae","Lygus","Lygus lineolaris","1","1","1","159","0","0","99.78","449","0","450","34.22","0"
"AGAW2","10E","NAR","BOLD:AAI9663","Mollusca","Gastropoda","Stylommatophora","Agriolimacidae",NA,"Deroceras","Deroceras reticulatum","7","6","6","1","14.29","0","99.26","406.14","0","444.29","35.74","0"
"AGAW2","10E","NAR","TAX:660402","Mollusca","Gastropoda","Stylommatophora","Agriolimacidae",NA,"Deroceras","Deroceras sp.","1","1","1","2","0","0","97.09","378","0","381","33.33","0"
"AGAW2","10E","NIN","BOLD:AAI4346","Arthropoda","Arachnida","Opiliones","Phalangiidae","","Paroligolophus, Phalangium","Paroligolophus agrestis, Phalangium opilio","1769","1553","1553","1","12.21","0","99.31","437.15","0","439.03","36.35","0"
"AGAW2","10E","NIN","BOLD:AAH6630","Arthropoda","Collembola","Entomobryomorpha","Entomobryidae","Entomobryinae","Entomobrya","Entomobrya unostrigata","692","603","603","2","12.86","0","99.33","435.06","0","436.26","38.99","0"
"AGAW2","10E","NIN","BOLD:AAA6381","Arthropoda","Arachnida","Araneae","Tetragnathidae","","Tetragnatha","Tetragnatha laboriosa","603","593","593","3","1.66","0","98.92","426","0","428.21","39.19","0"
"AGAW2","10E","NIN","BOLD:ABX0027","Arthropoda","Collembola","Symphypleona","Bourletiellidae","","Bourletiella","","101","81","81","4","19.8","0","99.64","416.86","0","444.8","35.63","0"
"AGAW2","10E","NIN","BOLD:ABW2639","Arthropoda","Arachnida","Trombidiformes","Anystidae","","","","45","44","44","5","2.22","0","99.51","447.8","0","445.69","31.48","0"
"AGAW2","10E","NIN","BOLD:AAV3042","Arthropoda","Arachnida","Araneae","Theridiidae","","Theridion","Theridion albidum","41","41","41","6","0","0","98.4","409.78","0","412.63","28.26","0"
"AGAW2","10E","NIN","BOLD:AAF9173","Arthropoda","Arachnida","Sarcoptiformes","Mochlozetidae","","Podoribates","Podoribates pratensis","39","34","34","7","12.82","0","99.67","431.03","0","431.72","42.8","0"
"AGAW2","10E","NIN","BOLD:AAU6252","Arthropoda","Arachnida","Trombidiformes","Bdellidae","","Cyta","","33","31","31","8","6.06","0","99.41","428.12","0","430.03","31.33","0"
"AGAW2","10E","NIN","BOLD:AAA9476","Arthropoda","Arachnida","Araneae","Linyphiidae","","Grammonota","Grammonota gentilis","27","24","24","9","11.11","0","98.8","431.93","0","433.19","30.82","0"
"AGAW2","10E","NIN","BOLD:AAA8820","Arthropoda","Collembola","Entomobryomorpha","Entomobryidae","Lepidocyrtinae","Lepidocyrtus","Lepidocyrtus cyaneus","26","26","26","10","0","0","98.94","404.5","0","406.38","43.61","0"
"AGAW2","10E","NIN","BOLD:AAM7961","Arthropoda","Arachnida","Trombidiformes","Anystidae","","","","17","14","14","11","17.65","0","99.67","438.06","0","444.65","31.31","0"
"AGAW2","10E","NIN","BOLD:ACX8969","Arthropoda","Arachnida","Trombidiformes","Eupodidae","","","","5","5","5","12","0","0","98.77","412","0","415.4","27.31","0"
"AGAW2","10E","NIN","BOLD:AAB8452","Arthropoda","Collembola","Entomobryomorpha","Entomobryidae","Lepidocyrtinae","Lepidocyrtus","Lepidocyrtus paradoxus","4","4","4","13","0","0","99.71","434.5","0","435","42.4","0"
"AGAW2","10E","NIN","BOLD:ACZ1295","Arthropoda","Arachnida","Trombidiformes","Anystidae","","","","3","3","3","14","0","0","99.72","367.67","0","370","31.16","0"
"AGAW2","10E","NIN","BOLD:AAB7601","Arthropoda","Arachnida","Araneae","Eutichuridae","","Cheiracanthium","Cheiracanthium mildei","3","3","3","14","0","0","97.67","405.67","0","408","28.61","0"
"AGAW2","10E","NIN","BOLD:AAA7242","Arthropoda","Collembola","Entomobryomorpha","Entomobryidae","Seirinae, Entomobryinae","Entomobrya, Lepidosira","Entomobrya multifasciata, Lepidosira sp.","1","1","1","16","0","0","98.06","360","0","363","40.22","0"
"AGAW2","10E","NIN","BOLD:AAB7914","Arthropoda","Collembola","Symphypleona","Bourletiellidae","","Bourletiella","","1","1","1","16","0","0","100","450","0","450","38.22","0"
"AGAW2","10E","NIN","BOLD:ACN5817","Arthropoda","Arachnida","Mesostigmata","Parasitidae","","","","1","1","1","16","0","0","100","365","0","450","41.56","0"
"AGAW2","10N","BAC","TAX:175768","Proteobacteria","Alphaproteobacteria","Rickettsiales","Rickettsiaceae",NA,"Wolbachia","Wolbachia pipientis","18","14","14","1","22.22","0","98.02","448.83","0","449.78","39.48","0"
"AGAW2","10N","BAC","TAX:91393","Proteobacteria","Alphaproteobacteria","Rickettsiales","Rickettsiaceae",NA,"Wolbachia","","12","12","12","2","0","0","98.91","390.67","0","418.75","39.1","0"
"AGAW2","10N","BAC","TAX:91391","Proteobacteria","Alphaproteobacteria","Rickettsiales","",NA,"","","9","9","9","3","0","0","99.38","398.22","0","430.78","40.81","0"
"AGAW2","10N","CHO","no match","no match","no match","no match","no match","no match","no match","no match","no match","no match","no match","no match","no match","no match","no match","no match","no match","no match","no match","no match"
"AGAW2","10N","INS","BOLD:AAG1711","Arthropoda","Insecta","Diptera","Muscidae","Phaoniinae","Helina","Helina depuncta","15391","12302","12302","1","20.07","0","99.43","431.41","0","433.79","31.34","0"
"AGAW2","10N","INS","BOLD:AAN6154","Arthropoda","Insecta","Coleoptera","Latridiidae","Corticariinae","Melanophthalma","Melanophthalma inermis","11321","9277","9277","2","18.05","0","98.9","434.69","0","436.35","31.31","0"
"AGAW2","10N","INS","BOLD:AAC9062","Arthropoda","Insecta","Diptera","Culicidae","Culicinae","Ochlerotatus, Aedes","Aedes fitchii, Ochlerotatus fitchii, Aedes grossbecki, Aedes ventrovittis","7166","5615","5615","3","21.64","0","99.52","429.51","0","431.28","33.09","0"
"AGAW2","10N","INS","BOLD:AAG6745","Arthropoda","Insecta","Diptera","Calliphoridae","Calliphorinae","Pollenia","Pollenia pediculata","7102","6227","6227","4","12.32","0","99.42","421.61","0","423.53","33.52","0"
"AGAW2","10N","INS","BOLD:AAH4135","Arthropoda","Insecta","Diptera","Chloropidae","Chloropinae","Thaumatomyia","Thaumatomyia glabra, Thaumatomyia grata, Thaumatomyia sp. 1 TAW","4575","3781","3781","5","17.36","0","99.16","434.17","0","437.48","32.07","0"
"AGAW2","10N","INS","BOLD:AAU6681","Arthropoda","Insecta","Diptera","Tipulidae","Tipulinae","Nephrotoma","Nephrotoma cornicina","4079","3095","3095","6","24.12","0","99.63","442.07","0","443.35","36.3","0"
"AGAW2","10N","INS","BOLD:AAG2123","Arthropoda","Insecta","Diptera","Tachinidae","Dexiinae","Dinera","Dinera grisescens","3767","2796","2796","7","25.78","0","99.37","432.39","0","434.04","31.83","0"
"AGAW2","10N","INS","BOLD:ACE7444","Arthropoda","Insecta","Hemiptera","Miridae","Mirinae, Phylinae","Adelphocoris, Megalocoleus","Adelphocoris lineolatus, Megalocoleus molliculus","3544","2298","2298","8","35.16","0","99.71","440.85","0","441.61","34.81","0"
"AGAW2","10N","INS","BOLD:ABW7897","Arthropoda","Insecta","Diptera","Cecidomyiidae","","","","3057","2109","2109","9","31.01","0","99.61","442.04","0","444.03","29.67","0"
"AGAW2","10N","INS","BOLD:AAH3661","Arthropoda","Insecta","Diptera","Cecidomyiidae","","","","1453","1296","1296","10","10.81","0","99.23","428.07","0","430.04","25.3","0"
"AGAW2","10N","INS","BOLD:AAG2868","Arthropoda","Insecta","Hemiptera","Cicadellidae","Typhlocybinae","Empoasca","Empoasca sp., Empoasca fabae","1447","935","935","11","35.38","0","99.59","446.05","0","447.52","32.29","0"
"AGAW2","10N","INS","BOLD:AAC0903","Arthropoda","Insecta","Diptera","Chironomidae","Chironominae","Chironomus","Chironomus acidophilus","1413","1106","1106","12","21.73","0","99.56","436.2","0","438.68","36.3","0"
"AGAW2","10N","INS","BOLD:AAA3453","Arthropoda","Insecta","Diptera","Anthomyiidae","Anthomyiinae","Delia","","1385","1277","1277","13","7.8","0","99.39","428.43","0","431.22","34.6","0"
"AGAW2","10N","INS","BOLD:AAP8739","Arthropoda","Insecta","Diptera","Platystomatidae","Platystomatinae","Rivellia","Rivellia steyskali","1173","885","885","14","24.55","0","99.31","440.33","0","442.17","37.99","0"
"AGAW2","10N","INS","BOLD:ACK3431","Arthropoda","Insecta","Diptera","Cecidomyiidae","","","","1071","950","950","15","11.3","0","99.2","425.72","0","427.6","32.3","0"
"AGAW2","10N","INS","BOLD:AAG3154","Arthropoda","Insecta","Diptera","Psychodidae","Psychodinae","Psychoda","","1031","910","910","16","11.74","0","98.36","436.06","0","438.48","31.92","0"
"AGAW2","10N","INS","BOLD:ABW1379","Arthropoda","Insecta","Diptera","Chloropidae","Oscinellinae","Malloewia","Malloewia abdominalis","809","623","623","17","22.99","0","98.24","444.51","0","446.97","31.62","0"
"AGAW2","10N","INS","BOLD:ABY5318","Arthropoda","Insecta","Diptera","Chloropidae","Oscinellinae","Incertella, Rhopalopterum","Rhopalopterum umbrosum, Rhopalopterum luteiceps, Incertella bispina, Rhopalopterum painteri","762","591","591","18","22.44","0","99.26","442.21","0","443.31","32.33","0"
"AGAW2","10N","INS","BOLD:ABA0863","Arthropoda","Insecta","Diptera","Cecidomyiidae","","","","668","572","572","19","14.37","0","99.4","439.14","0","440.54","29.74","0"
"AGAW2","10N","INS","BOLD:AAV5575","Arthropoda","Insecta","Diptera","Cecidomyiidae","","","CecidInt30 sp. BOLD:AAV5575","615","459","459","20","25.37","0","99.51","442.18","0","443.64","25.9","0"
"AGAW2","10N","INS","BOLD:ACU7906","Arthropoda","Insecta","Diptera","Chironomidae","Orthocladiinae","","","600","599","599","21","0.17","0","98.86","405.34","0","410.64","36.12","0"
"AGAW2","10N","INS","BOLD:AAH0135","Arthropoda","Insecta","Coleoptera","Phalacridae","Phalacrinae","Acylomus","Acylomus pugetanus, Acylomus ergoti","530","484","484","22","8.68","0","99.13","434.14","0","436.37","31.6","0"
"AGAW2","10N","INS","BOLD:AAN5225","Arthropoda","Insecta","Diptera","Cecidomyiidae","","","","507","387","387","23","23.67","0","99.2","447.16","0","448.24","28.37","0"
"AGAW2","10N","INS","BOLD:ABV1297","Arthropoda","Insecta","Diptera","Sciaridae","","","","477","458","458","24","3.98","0","99.09","430.44","0","432.93","30.19","0"
"AGAW2","10N","INS","BOLD:AAN5242","Arthropoda","Insecta","Diptera","Cecidomyiidae","","","","432","428","428","25","0.93","0","98.72","404.37","0","407.28","28.12","0"
"AGAW2","10N","INS","BOLD:AAD1485","Arthropoda","Insecta","Diptera","Chironomidae","Chironominae","Paratanytarsus","Paratanytarsus grimmii, Paratanytarsus sp.","400","323","323","26","19.25","0","99.52","442.41","0","443.93","33.78","0"
"AGAW2","10N","INS","BOLD:ACE0554","Arthropoda","Insecta","Diptera","Phoridae","Metopininae","Megaselia","","398","332","332","27","16.58","0","99.64","440.48","0","442.32","32.02","0"
"AGAW2","10N","INS","BOLD:AAA7661","Arthropoda","Insecta","Diptera","Culicidae","Culicinae","Culiseta, Culex","Culiseta melanura, Culex articularis, Culex restuans, Culex acharistus","396","372","372","28","6.06","0","99.24","431.54","0","433.68","32.67","0"
"AGAW2","10N","INS","BOLD:AAN5199","Arthropoda","Insecta","Diptera","Cecidomyiidae","","","CecidInt20 sp. BOLD:AAN5199","383","307","307","29","19.84","0","99.48","435.72","0","444.54","26.6","0"
"AGAW2","10N","INS","BOLD:AAD7604","Arthropoda","Insecta","Coleoptera","Coccinellidae","Coccinellinae","Coleomegilla","Coleomegilla maculata","379","343","343","30","9.5","0","99.05","432.14","0","434.73","31.73","0"
"AGAW2","10N","INS","BOLD:AAG5507","Arthropoda","Insecta","Diptera","Chironomidae","Chironominae","Chironomus","Chironomus atrella","359","355","355","31","1.11","0","98.74","425.46","0","428.63","35.79","0"
"AGAW2","10N","INS","BOLD:AAH3664","Arthropoda","Insecta","Diptera","Cecidomyiidae","","","CecidInt3 sp. BOLD:AAH3664","337","325","325","32","3.56","0","99.03","429.09","0","433.02","27.04","0"
"AGAW2","10N","INS","BOLD:ACP8377","Arthropoda","Insecta","Diptera","Psychodidae","Psychodinae","","","326","246","246","33","24.54","0","99.68","445.74","0","447.55","30.98","0"
"AGAW2","10N","INS","BOLD:AAG3273","Arthropoda","Insecta","Diptera","Phoridae","Metopininae","Megaselia","Megaselia albicaudata","325","232","232","34","28.62","0","99.54","444.37","0","444.91","32.7","0"
"AGAW2","10N","INS","BOLD:ABA0822","Arthropoda","Insecta","Diptera","Cecidomyiidae","","","","311","254","254","35","18.33","0","98.73","439.3","0","440.74","26.66","0"
"AGAW2","10N","INS","BOLD:AAC4197","Arthropoda","Insecta","Diptera","Chironomidae","Orthocladiinae","Paraphaenocladius","Paraphaenocladius impensus","304","266","266","36","12.5","0","99","433.2","0","435.66","34.64","0"
"AGAW2","10N","INS","BOLD:AAN5205","Arthropoda","Insecta","Diptera","Cecidomyiidae","","","CecidInt42 sp. BOLD:AAN5205","297","235","235","37","20.88","0","99.63","446.87","0","448.15","31.4","0"
"AGAW2","10N","INS","BOLD:AAO8538","Arthropoda","Insecta","Diptera","Psychodidae","Psychodinae","Psychoda","","249","235","235","38","5.62","0","99.03","427.94","0","429.92","32.55","0"
"AGAW2","10N","INS","BOLD:ABY1179","Arthropoda","Insecta","Hymenoptera","Sierolomorphidae","","Sierolomorpha","","240","205","205","39","14.58","0","99.55","437.85","0","439.46","28.15","0"
"AGAW2","10N","INS","BOLD:AAA4277","Arthropoda","Insecta","Diptera","Syrphidae","Syrphinae","Toxomerus","Toxomerus marginatus, Toxomerus sp. CR-3, Toxomerus CR-3","237","168","168","40","29.11","0","99.53","442.78","0","443.64","30.58","0"
"AGAW2","10N","INS","BOLD:AAA4555","Arthropoda","Insecta","Orthoptera","Acrididae","Melanoplinae","Melanoplus","Melanoplus stupefactus, Melanoplus angustipennis, Melanoplus ponderosus, Melanoplus bruneri, Melanoplus frigidus, Melanoplus scapularis, Melanoplus cinereus, Melanoplus madeleineae, Melanoplus sanguinipes, Melanoplus nigrescens group, Melanoplus tunicae, Melanoplus walshi, Melanoplus femurrubrum, Melanoplus dawsoni, Melanoplus sp., Melanoplus fasciatus, Melanoplus bowditchi, Melanoplus stonei, Melanoplus bivittatus, Melanoplus huroni, Melanoplus thomasi, Melanoplus borealis, Melanoplus packardii","222","201","201","41","9.46","0","99.19","445.72","0","447.33","36.54","0"
"AGAW2","10N","INS","BOLD:AAP7026","Arthropoda","Insecta","Coleoptera","Latridiidae","Corticariinae","Melanophthalma","Melanophthalma villosa","215","192","192","42","10.7","0","97.87","394.47","0","434.14","30.98","0"
"AGAW2","10N","INS","BOLD:AAN5260","Arthropoda","Insecta","Diptera","Cecidomyiidae","","","","203","197","197","43","2.96","0","99.14","405.06","0","406.97","25.87","0"
"AGAW2","10N","INS","BOLD:AAM9255","Arthropoda","Insecta","Diptera","Sciaridae","","Pseudolycoriella","","188","184","184","44","2.13","0","99.13","405.55","0","407.8","30.71","0"
"AGAW2","10N","INS","BOLD:AAH2870","Arthropoda","Insecta","Hemiptera","Aphididae","Calaphidinae","Euceraphis","Euceraphis papyrifericola","165","148","148","45","10.3","0","99.42","423.81","0","424.97","27.36","0"
"AGAW2","10N","INS","BOLD:ACX6055","Arthropoda","Insecta","Diptera","Phoridae","Metopininae","Megaselia","Megaselia sp. AK-2010-2, Megaselia sp. BOLD:ACX6055","158","140","140","46","11.39","0","99.45","435.6","0","437.56","33.05","0"
"AGAW2","10N","INS","BOLD:AAM9013","Arthropoda","Insecta","Diptera","Keroplatidae","Macrocerinae","Macrocera","Macrocera pusilla","152","145","145","47","4.61","0","99.38","434.48","0","436.19","30.32","0"
"AGAW2","10N","INS","BOLD:AAB0373","Arthropoda","Insecta","Neuroptera","Chrysopidae","Chrysopinae","Chrysoperla","Chrysoperla sp. isolate 82, Chrysoperla zastrowi, Chrysoperla plorabunda, Chrysoperla sp. isolate 71, Chrysoperla agilis, Italochrysa italica, Chrysoperla sp. isolate 88, Chrysoperla lucasina, Chrysoperla carnea, Chrysoperla mediterranea, Chrysoperla pallida, Chrysoperla nipponensis, Chrysoperla sillemi","143","124","124","48","13.29","0","99.56","431.11","0","432.28","31.3","0"
"AGAW2","10N","INS","BOLD:ABU5525","Arthropoda","Insecta","Diptera","Chironomidae","Orthocladiinae","Limnophyes","Limnophyes sp. 14ES","130","129","129","49","0.77","0","99.09","392.13","0","397.97","37.26","0"
"AGAW2","10N","INS","BOLD:ACG3646","Arthropoda","Insecta","Coleoptera","Corylophidae","Corylophinae","Orthoperus","","129","100","100","50","22.48","0","98.97","443.26","0","445","33.64","0"
"AGAW2","10N","INS","BOLD:AAG6431","Arthropoda","Insecta","Diptera","Ceratopogonidae","Forcipomyiinae","Atrichopogon","","123","120","120","51","2.44","0","98.14","421.66","0","427.41","36.47","0"
"AGAW2","10N","INS","BOLD:AAA5299","Arthropoda","Insecta","Diptera","Chironomidae","Orthocladiinae","Cricotopus","Cricotopus laetus, Cricotopus relucens, Cricotopus trifasciatus, Cricotopus sp., Cricotopus sp. 3ES, Cricotopus sylvestris, Cricotopus glacialis","122","114","114","52","6.56","0","99.37","436.25","0","438.09","34.59","0"
"AGAW2","10N","INS","BOLD:ACU7060","Arthropoda","Insecta","Diptera","Chironomidae","Chironominae","Microtendipes","","120","117","117","53","2.5","0","99.28","434.27","0","436.2","33.2","0"
"AGAW2","10N","INS","BOLD:AAN5249","Arthropoda","Insecta","Diptera","Cecidomyiidae","","","","113","111","111","54","1.77","0","98.75","430.58","0","432.01","33.4","0"
"AGAW2","10N","INS","BOLD:AAL7359","Arthropoda","Insecta","Diptera","Chironomidae","Chironominae","Tanytarsus","","111","105","105","55","5.41","0","99.31","438.49","0","440.24","36.16","0"
"AGAW2","10N","INS","BOLD:AAH2953","Arthropoda","Insecta","Diptera","Rhagionidae","Rhagioninae","Rhagio","Rhagio tringarius","106","92","92","56","13.21","0","99.22","438.1","0","440.6","32.23","0"
"AGAW2","10N","INS","BOLD:AAN5183","Arthropoda","Insecta","Diptera","Cecidomyiidae","","","","98","90","90","57","8.16","0","98.23","440.91","0","442.04","27.73","0"
"AGAW2","10N","INS","BOLD:AAZ5592","Arthropoda","Insecta","Diptera","Psychodidae","Psychodinae","","","93","86","86","58","7.53","0","99.12","431.53","0","433.68","32.49","0"
"AGAW2","10N","INS","BOLD:AAG7891","Arthropoda","Insecta","Hymenoptera","Platygastridae","Scelioninae","Telenomus","Telenomus podisi","89","81","81","59","8.99","0","99.05","431.72","0","440.66","27.12","0"
"AGAW2","10N","INS","BOLD:AAN5267","Arthropoda","Insecta","Diptera","Cecidomyiidae","","","","83","62","62","60","25.3","0","99.51","448.39","0","449.31","30.61","0"
"AGAW2","10N","INS","BOLD:AAL7819","Arthropoda","Insecta","Diptera","Psychodidae","Psychodinae","Psychoda","Psychoda sp., Psychoda sp. BOLD:AAL7819","82","77","77","61","6.1","0","99.28","414.21","0","416.16","32.68","0"
"AGAW2","10N","INS","BOLD:AAM9347","Arthropoda","Insecta","Diptera","Phoridae","Metopininae","Megaselia","","73","72","72","62","1.37","0","98.05","398.32","0","401","33.09","0"
"AGAW2","10N","INS","BOLD:AAF9771","Arthropoda","Insecta","Diptera","Hybotidae","Tachydromiinae","Platypalpus","Platypalpus longiseta, Platypalpus trivialis, Platypalpus sp. 5, Platypalpus sp. 17, Platypalpus pallidiventris","70","61","61","63","12.86","0","99.49","441.76","0","442.66","35.13","0"
"AGAW2","10N","INS","BOLD:AAN5220","Arthropoda","Insecta","Diptera","Cecidomyiidae","","","","69","69","69","64","0","0","97.97","419.59","0","424.9","25.78","0"
"AGAW2","10N","INS","BOLD:AAN5330","Arthropoda","Insecta","Diptera","Chironomidae","Orthocladiinae","Corynoneura","Corynoneura scutellata","63","63","63","65","0","0","98.85","411.32","0","418.22","34.01","0"
"AGAW2","10N","INS","BOLD:AAH3231","Arthropoda","Insecta","Psocodea","Psocidae","","","","59","59","59","66","0","0","98.49","413.15","0","415.17","38.62","0"
"AGAW2","10N","INS","BOLD:ABZ5071","Arthropoda","Insecta","Diptera","Chironomidae","Chironominae","Micropsectra","Micropsectra nigripila","57","57","57","67","0","0","98.18","416.54","0","429.12","36.83","0"
"AGAW2","10N","INS","BOLD:ABX5969","Arthropoda","Insecta","Hemiptera","Miridae","Mirinae","Neolygus","Neolygus inconspicuus, Neolygus communis","56","43","43","68","23.21","0","99.12","445.13","0","445.8","34.19","0"
"AGAW2","10N","INS","BOLD:ACC8079","Arthropoda","Insecta","Diptera","Cecidomyiidae","","","","56","40","40","68","28.57","0","99.57","447.7","0","449.07","29.28","0"
"AGAW2","10N","INS","BOLD:ABA6407","Arthropoda","Insecta","Diptera","Sciaridae","","Pseudolycoriella","","56","55","55","68","1.79","0","98.94","430.2","0","432.73","34.92","0"
"AGAW2","10N","INS","BOLD:ACK5742","Arthropoda","Insecta","Diptera","Cecidomyiidae","","","","53","48","48","71","9.43","0","99.41","439.17","0","440.25","29.05","0"
"AGAW2","10N","INS","BOLD:AAN5148","Arthropoda","Insecta","Diptera","Ceratopogonidae","Forcipomyiinae","Forcipomyia","Forcipomyia bipunctata, Forcipomyia sp. CK-2012","51","51","51","72","0","0","98.63","423.96","0","428.84","35.6","0"
"AGAW2","10N","INS","BOLD:AAP8816","Arthropoda","Insecta","Diptera","Cecidomyiidae","Cecidomyiinae","Sitodiplosis","Sitodiplosis sp. BOLD:AAP8816","50","45","45","73","10","0","99.51","440.8","0","442.14","26.34","0"
"AGAW2","10N","INS","BOLD:AAM8930","Arthropoda","Insecta","Psocodea","Caeciliusidae","Caeciliusinae","Valenzuela","","49","45","45","74","8.16","0","98.64","439.78","0","441.06","37.06","0"
"AGAW2","10N","INS","BOLD:AAH0134","Arthropoda","Insecta","Coleoptera","Phalacridae","Phalacrinae","Stilbus","Stilbus apicialis","48","44","44","75","8.33","0","99.24","426.35","0","428.02","31.9","0"
"AGAW2","10N","INS","BOLD:ABW8023","Arthropoda","Insecta","Diptera","Cecidomyiidae","","","","47","47","47","76","0","0","98.1","445.68","0","447.81","27.19","0"
"AGAW2","10N","INS","BOLD:AAN5230","Arthropoda","Insecta","Diptera","Cecidomyiidae","","","","45","45","45","77","0","0","99.3","421.04","0","422.93","25.85","0"
"AGAW2","10N","INS","BOLD:ACM1709","Arthropoda","Insecta","Diptera","Chironomidae","Tanypodinae","","","44","41","41","78","6.82","0","99.18","437.27","0","438.64","35.02","0"
"AGAW2","10N","INS","BOLD:ACF2457","Arthropoda","Insecta","Diptera","Chironomidae","Chironominae","Paratanytarsus","Paratanytarsus laccophilus","41","37","37","79","9.76","0","99.12","438.98","0","440.85","32.41","0"
"AGAW2","10N","INS","BOLD:ACV5892","Arthropoda","Insecta","Diptera","Cecidomyiidae","","","","38","37","37","80","2.63","0","97.98","441.95","0","443.53","25.2","0"
"AGAW2","10N","INS","BOLD:ABV3850","Arthropoda","Insecta","Coleoptera","Curculionidae","Entiminae","Sitona","Sitona lepidus","36","30","30","81","16.67","0","99.19","438.36","0","439.53","36.38","0"
"AGAW2","10N","INS","BOLD:ACA6930","Arthropoda","Insecta","Diptera","Cecidomyiidae","","","","35","32","32","82","8.57","0","98.47","427.2","0","444.09","27.81","0"
"AGAW2","10N","INS","BOLD:AAN5255","Arthropoda","Insecta","Diptera","Cecidomyiidae","Cecidomyiinae","Dasineura","","34","30","30","83","11.76","0","97.99","435.85","0","446.82","26.72","0"
"AGAW2","10N","INS","BOLD:AAF9306","Arthropoda","Insecta","Diptera","Psychodidae","Psychodinae","Psychoda","Psychoda alternata","33","33","33","84","0","0","98.32","436.36","0","437.3","31.04","0"
"AGAW2","10N","INS","BOLD:AAV5688","Arthropoda","Insecta","Diptera","Cecidomyiidae","","","","32","28","28","85","12.5","0","99.2","430.88","0","432.19","34.38","0"
"AGAW2","10N","INS","BOLD:AAN8770","Arthropoda","Insecta","Diptera","Psychodidae","Psychodinae","Psychoda","Psychoda trinodulosa","32","29","29","85","9.38","0","99.09","435.94","0","437.03","33.59","0"
"AGAW2","10N","INS","BOLD:AAN8446","Arthropoda","Insecta","Psocodea","Peripsocidae","","Peripsocus","Peripsocus madidus","31","31","31","87","0","0","99.09","421","0","422.19","41.52","0"
"AGAW2","10N","INS","BOLD:ACA7877","Arthropoda","Insecta","Diptera","Cecidomyiidae","","","","31","26","26","87","16.13","0","98.95","438.61","0","440.19","25.76","0"
"AGAW2","10N","INS","BOLD:AAG5453","Arthropoda","Insecta","Diptera","Chironomidae","Chironominae","Chironomus","Chironomus bifurcatus","28","28","28","89","0","0","98.16","447.93","0","450","34.91","0"
"AGAW2","10N","INS","BOLD:ACK3142","Arthropoda","Insecta","Diptera","Cecidomyiidae","","","","28","28","28","89","0","0","98.54","434.64","0","436.11","28.11","0"
"AGAW2","10N","INS","BOLD:AAD2041","Arthropoda","Insecta","Diptera","Dolichopodidae","Sciapodinae","Condylostylus","Condylostylus inermis, Condylostylus nigrofemoratus, Condylostylus connectans, Condylostylus caudatus, Condylostylus flavipes","26","18","18","91","30.77","0","99.71","441.23","0","442.46","36.02","0"
"AGAW2","10N","INS","BOLD:AAG6436","Arthropoda","Insecta","Diptera","Ceratopogonidae","Ceratopogoninae","Culicoides","Culicoides travisi","21","19","19","92","9.52","0","99.39","442.19","0","450","37.25","0"
"AGAW2","10N","INS","BOLD:AAY6730","Arthropoda","Insecta","Hemiptera","Derbidae","Cedusinae","Cedusa","Cedusa incisa","19","19","19","93","0","0","99.23","432.89","0","434.21","35.31","0"
"AGAW2","10N","INS","BOLD:AAN5221","Arthropoda","Insecta","Diptera","Cecidomyiidae","","","","19","19","19","93","0","0","99.07","411.58","0","413.68","25.41","0"
"AGAW2","10N","INS","BOLD:AAM6776","Arthropoda","Insecta","Diptera","Dolichopodidae","Medeterinae","Medetera","Medetera vittata","19","16","16","93","15.79","0","99.32","436.26","0","437.05","39.56","0"
"AGAW2","10N","INS","BOLD:ABA1215","Arthropoda","Insecta","Diptera","Sciaridae","","Lycoriella","Lycoriella sativae","18","16","16","96","11.11","0","99.44","434.94","0","436.61","35.8","0"
"AGAW2","10N","INS","BOLD:ADH9867","Arthropoda","Insecta","Diptera","","","","","18","18","18","96","0","0","98.81","425.22","0","429.11","37.45","0"
"AGAW2","10N","INS","BOLD:AAN5501","Arthropoda","Insecta","Diptera","Hybotidae","","","","17","17","17","98","0","0","98.8","447","0","448.29","35.51","0"
"AGAW2","10N","INS","BOLD:AAG3302","Arthropoda","Insecta","Diptera","Phoridae","Metopininae","Megaselia","","17","17","17","98","0","0","99.13","409.71","0","412.53","31.47","0"
"AGAW2","10N","INS","BOLD:ADG3489","Arthropoda","Insecta","Diptera","","","","","16","16","16","100","0","0","99.56","387.06","0","426.88","33.98","0"
"AGAW2","10N","INS","BOLD:ACX5521","Arthropoda","Insecta","Diptera","Chironomidae","Orthocladiinae","","","16","16","16","100","0","0","98.6","423.44","0","425.38","35.15","0"
"AGAW2","10N","INS","BOLD:AAK5928","Arthropoda","Insecta","Coleoptera","Chrysomelidae","Criocerinae","Oulema","Oulema melanopus","15","15","15","102","0","0","99.58","435.47","0","437.2","36.17","0"
"AGAW2","10N","INS","BOLD:AAI4303","Arthropoda","Insecta","Diptera","Chironomidae","Chironominae","Chironomus","Chironomus melanescens","14","14","14","103","0","0","98.21","428","0","431.5","34.05","0"
"AGAW2","10N","INS","BOLD:AAV5400","Arthropoda","Insecta","Diptera","Cecidomyiidae","","","","14","14","14","103","0","0","97.81","427.57","0","429.57","29.04","0"
"AGAW2","10N","INS","BOLD:AAA6560","Arthropoda","Insecta","Lepidoptera","Crambidae","Crambinae","Crambus","Crambus multilinellus, Crambus saltuellus","12","12","12","105","0","0","99.43","437.33","0","438.75","30.87","0"
"AGAW2","10N","INS","BOLD:AAA4222","Arthropoda","Insecta","Lepidoptera","Erebidae","Hypeninae","Hypena","Hypena scabra","11","11","11","106","0","0","99.42","388.55","0","390.82","32.28","0"
"AGAW2","10N","INS","BOLD:AAU8379","Arthropoda","Insecta","Hymenoptera","Braconidae","Alysiinae","Dinotrema","Dinotrema JFT01","11","11","11","106","0","0","99.41","447.09","0","448.55","24.49","0"
"AGAW2","10N","INS","BOLD:AAG2511","Arthropoda","Insecta","Diptera","Anthomyiidae","Anthomyiinae","Delia","Delia platura","9","9","9","108","0","0","99.41","431.11","0","433.89","33.26","0"
"AGAW2","10N","INS","BOLD:ACV5891","Arthropoda","Insecta","Diptera","Cecidomyiidae","","","","8","8","8","109","0","0","99.17","361.38","0","430","24.93","0"
"AGAW2","10N","INS","BOLD:ACF1257","","","","","","","","8","8","8","109","0","0","98.91","433.75","0","435.13","34.86","0"
"AGAW2","10N","INS","BOLD:AAV5561","Arthropoda","Insecta","Diptera","Cecidomyiidae","","","","7","7","7","111","0","0","98.5","431","0","433","27.33","0"
"AGAW2","10N","INS","BOLD:ACK1456","Arthropoda","Insecta","Diptera","Cecidomyiidae","","","","7","7","7","111","0","0","99.47","431","0","448.43","25.93","0"
"AGAW2","10N","INS","BOLD:AAE7309","Arthropoda","Insecta","Hemiptera","Aphididae","Aphidinae","Aphis","Aphis sambuci, Aphis horii","7","6","6","111","14.29","0","99.87","443.43","0","445.14","27.65","0"
"AGAW2","10N","INS","BOLD:AAU8628","Arthropoda","Insecta","Hymenoptera","Bethylidae","","","","7","7","7","111","0","0","98.7","400.29","0","402.71","26.57","0"
"AGAW2","10N","INS","BOLD:AAA2482","Arthropoda","Insecta","Lepidoptera","Noctuidae","Noctuinae","Mythimna","Mythimna unipuncta","6","6","6","115","0","0","99.22","413.5","0","416.33","30.4","0"
"AGAW2","10N","INS","BOLD:AAM6037","Arthropoda","Insecta","Diptera","Cecidomyiidae","Cecidomyiinae","Contarinia, Dasineura","","6","5","5","115","16.67","0","97.81","448.5","0","450","26.96","0"
"AGAW2","10N","INS","BOLD:ACF5830","Arthropoda","Insecta","Diptera","Chloropidae","Chloropinae","Thaumatomyia","","6","6","6","115","0","0","97.53","409.33","0","410.5","31.56","0"
"AGAW2","10N","INS","BOLD:AAC1374","Arthropoda","Insecta","Hemiptera","Aphididae","Aphidinae","Aphis","Aphis nasturtii, Aphis davletshinae, Aphis sp. rostella like, Aphis cf. rostella, Aphis althaeae, Aphis umbrella","5","5","5","118","0","0","99.55","439.8","0","440.6","26.13","0"
"AGAW2","10N","INS","BOLD:AAG2017","Arthropoda","Insecta","Neuroptera","Chrysopidae","Chrysopinae","Chrysopa","Chrysopa oculata","5","5","5","118","0","0","99","412.6","0","413.6","31.41","0"
"AGAW2","10N","INS","BOLD:AAZ0271","Arthropoda","Insecta","Diptera","Ceratopogonidae","Ceratopogoninae","Culicoides","","5","5","5","118","0","0","98.3","435.2","0","438.8","41.42","0"
"AGAW2","10N","INS","BOLD:AAA3181","Arthropoda","Insecta","Diptera","Muscidae","Muscinae","Stomoxys","Stomoxys calcitrans","4","4","4","121","0","0","98.85","435.25","0","439.5","32.8","0"
"AGAW2","10N","INS","BOLD:ABA0856","Arthropoda","Insecta","Diptera","Cecidomyiidae","","","","4","4","4","121","0","0","98.29","424.5","0","425.75","27.98","0"
"AGAW2","10N","INS","BOLD:ADG3385","Arthropoda","Insecta","Diptera","Cecidomyiidae","","","","4","4","4","121","0","0","99.04","391.5","0","443.75","25.23","0"
"AGAW2","10N","INS","BOLD:AAY6412","Arthropoda","Insecta","Diptera","Sciaridae","","Bradysia","Bradysia urticae","4","4","4","121","0","0","98.97","433.75","0","438.25","31.58","0"
"AGAW2","10N","INS","BOLD:AAA6822","Arthropoda","Insecta","Orthoptera","Acrididae","Gomphocerinae","Chorthippus","Chorthippus curtipennis","4","4","4","121","0","0","99.55","449.25","0","450","35.22","0"
"AGAW2","10N","INS","BOLD:AAA5206","Arthropoda","Insecta","Lepidoptera","Erebidae","Herminiinae","Zanclognatha","Zanclognatha cruralis, Zanclognatha sp. 1, Zanclognatha jacchusalis, Zanclognatha minoralis, Zanclognatha minoralis sp. 1, Zanclognatha sp., Zanclognatha minoralis sp. 3, Zanclognatha gypsalis, Zanclognatha dentata, Zanclognatha theralis, Zanclognatha protumnusalis, Zanclognatha martha, Zanclognatha atrilineella, Zanclognatha obscuripennis, Zanclognatha minoralis sp. 2, Zanclognatha marcidilinea","4","4","4","121","0","0","98.83","448.75","0","450","31.17","0"
"AGAW2","10N","INS","BOLD:ADF6661","Arthropoda","Insecta","Diptera","Cecidomyiidae","","","","4","3","3","121","25","0","99.46","418.5","0","450","29.89","0"
"AGAW2","10N","INS","BOLD:AAB0651","Arthropoda","Insecta","Lepidoptera","Gelechiidae","Dichomeridinae","Dichomeris","Dichomeris leuconotella, Dichomeris gnoma","4","3","3","121","25","0","99.83","447.5","0","450","33.39","0"
"AGAW2","10N","INS","BOLD:ABY5257","Arthropoda","Insecta","Lepidoptera","Noctuidae","Noctuinae","Apamea","Apamea devastator","4","4","4","121","0","0","99.49","443","0","443.75","30.65","0"
"AGAW2","10N","INS","BOLD:AAG5499","Arthropoda","Insecta","Diptera","Chironomidae","Orthocladiinae","Psectrocladius","Psectrocladius obvius, Psectrocladius platypus","4","4","4","121","0","0","99.16","448.25","0","450","35.72","0"
"AGAW2","10N","INS","BOLD:ABV1237","Arthropoda","Insecta","Diptera","Chironomidae","Chironominae","Chironomus","","4","4","4","121","0","0","99.43","436.5","0","436.75","34.9","0"
"AGAW2","10N","INS","BOLD:ABW6101","Arthropoda","Insecta","Diptera","Cecidomyiidae","","","","4","4","4","121","0","0","99.32","428","0","428.25","29.39","0"
"AGAW2","10N","INS","BOLD:AAA2518","Arthropoda","Insecta","Lepidoptera","Crambidae","Crambinae","Chrysoteuchia","Chrysoteuchia topiaria, Chrysoteuchia topiarius","3","3","3","133","0","0","97.84","447.67","0","450","35.11","0"
"AGAW2","10N","INS","BOLD:ABA0858","Arthropoda","Insecta","Diptera","Cecidomyiidae","","","","3","3","3","133","0","0","99.75","416.67","0","417.67","25.51","0"
"AGAW2","10N","INS","BOLD:AAC9132","Arthropoda","Insecta","Diptera","Culicidae","Culicinae","Culiseta","Culiseta inornata","3","3","3","133","0","0","99.12","372","0","374","32.72","0"
"AGAW2","10N","INS","BOLD:AAE7186","Arthropoda","Insecta","Hymenoptera","Braconidae","Aphidiinae","Praon","Praon sp. 1 MM-2013, Praon volucre, Praon bicolor","3","3","3","133","0","0","99.25","444","0","447","25.35","0"
"AGAW2","10N","INS","BOLD:AAH3225","Arthropoda","Insecta","Psocodea","Psocidae","Psocinae","Metylophorus","Metylophorus novaescotiae","3","3","3","133","0","0","98.57","443.67","0","445","34.91","0"
"AGAW2","10N","INS","BOLD:AAD8027","Arthropoda","Insecta","Diptera","Culicidae","Culicinae","Aedes","Aedes fitchii","3","3","3","133","0","0","99.39","433","0","433.67","34.42","0"
"AGAW2","10N","INS","BOLD:AAV5796","Arthropoda","Insecta","Diptera","Cecidomyiidae","","","","3","3","3","133","0","0","98.96","446.67","0","450","29.11","0"
"AGAW2","10N","INS","BOLD:AAH2868","Arthropoda","Insecta","Hemiptera","Aphididae","Drepanosiphinae","Drepanaphis","Drepanaphis acerifoliae","3","3","3","133","0","0","99.42","426.67","0","428","29.46","0"
"AGAW2","10N","INS","BOLD:AAZ8146","","","","","","","","3","3","3","133","0","0","99.23","397.33","0","419","24.55","0"
"AGAW2","10N","INS","BOLD:AAA7374","Arthropoda","Insecta","Diptera","Syrphidae","Syrphinae","Sphaerophoria","Sphaerophoria bankowskae, Sphaerophoria infuscata, Sphaerophoria nr. philanthus, Sphaerophoria batava, Sphaerophoria sp., Sphaerophoria rueppellii, Sphaerophoria interrupta-group, Sphaerophoria sp. MA-1, Sphaerophoria cleoae, Sphaerophoria asymmetrica, Sphaerophoria fatarum, Sphaerophoria bengalensis, Sphaerophoria bifurcata, Sphaerophoria taeniata, Sphaerophoria interrupta, Sphaerophoria contigua, Sphaerophoria kaa, Sphaerophoria laurae, Sphaerophoria chongjini, Sphaerophoria boreoalpina, Sphaerophoria sp. BOLDAAA7374, Sphaerophoria macrogaster, Sphaerophoria sulphuripes, Sphaerophoria brevipilosa, Sphaerophoria philanthus, Sphaerophoria pyrrhina, Sphaerophoria abbreviata, Sphaerophoria scripta, Sphaerophoria potentillae, Sphaerophoria longipilosa, Sphaerophoria virgata","3","3","3","133","0","0","98.58","445.67","0","450","30.81","0"
"AGAW2","10N","INS","BOLD:ACF4925","Arthropoda","Insecta","Hemiptera","Miridae","Mirinae","Adelphocoris","Adelphocoris detritus","3","3","3","133","0","0","97.4","421.67","0","423.67","36.01","0"
"AGAW2","10N","INS","BOLD:ABW3687","Arthropoda","Insecta","Diptera","Lauxaniidae","Homoneurinae","Homoneura","Homoneura bispina","2","2","2","144","0","0","98.51","408","0","409","34.13","0"
"AGAW2","10N","INS","BOLD:AAQ0558","Arthropoda","Insecta","Thysanoptera","Thripidae","Thripinae","Anaphothrips","Anaphothrips obscurus","2","2","2","144","0","0","99.11","450.5","0","450","31.44","0"
"AGAW2","10N","INS","BOLD:AAG7276","Arthropoda","Insecta","Diptera","Sphaeroceridae","Limosininae","Leptocera","Leptocera erythrocera","2","2","2","144","0","0","99.27","403.5","0","410","31.6","0"
"AGAW2","10N","INS","BOLD:AAH2306","Arthropoda","Insecta","Diptera","Asilidae","Asilinae","Efferia","Efferia aestuans","2","2","2","144","0","0","98.05","424","0","424.5","37.53","0"
"AGAW2","10N","INS","BOLD:AAA4751","Arthropoda","Insecta","Diptera","Culicidae","Culicinae","Culex","Culex pipiens sl, Culex nr. pipiens, Culex pipiens, Culex quinquefasciatus, Culex sp.","2","2","2","144","0","0","97.58","436","0","437","32.25","0"
"AGAW2","10N","INS","BOLD:AAM6781","Arthropoda","Insecta","Diptera","Dolichopodidae","Neurigoninae","Neurigona","Neurigona tenuis","2","2","2","144","0","0","99.33","449","0","450","36","0"
"AGAW2","10N","INS","BOLD:ACL2677","Arthropoda","Insecta","Psocodea","Mesopsocidae","","Mesopsocus","Mesopsocus laticeps","2","2","2","144","0","0","99.78","449","0","450","40.78","0"
"AGAW2","10N","INS","BOLD:ACL1796","Arthropoda","Insecta","Lepidoptera","Ypsolophidae","Ochsenheimeriinae","Ochsenheimeria","Ochsenheimeria vacculella","2","2","2","144","0","0","98.66","447","0","450","33","0"
"AGAW2","10N","INS","BOLD:AAO7783","Arthropoda","Insecta","Diptera","Dolichopodidae","Diaphorinae","","","2","2","2","144","0","0","98.21","448","0","450","33.56","0"
"AGAW2","10N","INS","BOLD:AAG8882","Arthropoda","Insecta","Hemiptera","Pentatomidae","Pentatominae","Thyanta","Thyanta pallidovirens, Thyanta custator, Thyanta accerra","2","2","2","144","0","0","99.11","449.5","0","450","36.22","0"
"AGAW2","10N","INS","BOLD:ABA5202","Arthropoda","Insecta","Diptera","Limoniidae","Limnophilinae","Pseudolimnophila","Pseudolimnophila luteipennis","2","2","2","144","0","0","98.11","449","0","450","33.78","0"
"AGAW2","10N","INS","BOLD:AAV1136","Arthropoda","Insecta","Diptera","Scatopsidae","","","","2","2","2","144","0","0","99.14","414.5","0","414.5","36.06","0"
"AGAW2","10N","INS","BOLD:ACE6286","Arthropoda","Insecta","Diptera","Culicidae","Culicinae","Ochlerotatus, Aedes","Aedes dorsalis, Ochlerotatus dorsalis","2","2","2","144","0","0","98.43","445.5","0","450","31.56","0"
"AGAW2","10N","INS","BOLD:AAH0237","Arthropoda","Insecta","Coleoptera","Cleridae","Clerinae","Enoclerus","Enoclerus nigripes, Enoclerus rosmarus","2","1","1","144","50","0","98.67","450","0","450","32.67","0"
"AGAW2","10N","INS","BOLD:AAZ5483","Arthropoda","Insecta","Diptera","Muscidae","Phaoniinae","Helina","Helina setiventris","2","2","2","144","0","0","98.78","424","0","427.5","28.85","0"
"AGAW2","10N","INS","BOLD:ACU6062","Arthropoda","Insecta","Diptera","Cecidomyiidae","","","","2","2","2","144","0","0","98.82","423.5","0","425","25.85","0"
"AGAW2","10N","INS","BOLD:AAA2632","Arthropoda","Insecta","Lepidoptera","Noctuidae","Noctuinae","Noctua","Noctua atlantica, Noctua pronuba","2","2","2","144","0","0","98.77","447","0","450","30.89","0"
"AGAW2","10N","INS","BOLD:ADI0607","","","","","","","","2","2","2","144","0","0","98.33","449.5","0","450","37.22","0"
"AGAW2","10N","INS","BOLD:AAA6337","Arthropoda","Insecta","Lepidoptera","Noctuidae","Plusiinae","Anagrapha","Anagrapha falcifera","2","2","2","144","0","0","99.67","448","0","450","32.44","0"
"AGAW2","10N","INS","BOLD:AAH2293","Arthropoda","Insecta","Diptera","Asilidae","Stenopogoninae","Dioctria, Eudioctria","Dioctria hyalipennis, Dioctria baumhaueri, Eudioctria propinqua","2","2","2","144","0","0","99.19","432","0","435","30.68","0"
"AGAW2","10N","INS","BOLD:AAG6929","Arthropoda","Insecta","Diptera","Dolichopodidae","Medeterinae","Medetera","Medetera vittata","2","2","2","144","0","0","99.33","449","0","450","41.11","0"
"AGAW2","10N","INS","BOLD:AAH3662","Arthropoda","Insecta","Diptera","Cecidomyiidae","","","CecidInt3 sp. BOLD:AAH3662","2","2","2","144","0","0","99","448.5","0","450","26.11","0"
"AGAW2","10N","INS","BOLD:AAL2750","Arthropoda","Insecta","Diptera","Muscidae","Phaoniinae","Phaonia","Phaonia tuguriorum","2","2","2","144","0","0","98.78","449","0","450","32.67","0"
"AGAW2","10N","INS","BOLD:AAF9316","Arthropoda","Insecta","Diptera","Psychodidae","Psychodinae","Psychoda","","2","2","2","144","0","0","99.78","448.5","0","450","31.67","0"
"AGAW2","10N","INS","BOLD:AAJ7123","Arthropoda","Insecta","Diptera","Culicidae","Culicinae","Culiseta","Culiseta minnesotae","2","2","2","144","0","0","98.73","432.5","0","434","34.2","0"
"AGAW2","10N","INS","BOLD:AAG6743","Arthropoda","Insecta","Diptera","Sarcophagidae","Sarcophaginae","Sarcophaga","Sarcophaga sp. 1 MG, Sarcophaga subvicina","2","2","2","144","0","0","100","443","0","444","35.36","0"
"AGAW2","10N","INS","BOLD:AAI3064","","","","","","","","2","2","2","144","0","0","99.11","447.5","0","450","33.44","0"
"AGAW2","10N","INS","BOLD:ACE9865","Arthropoda","Insecta","Lepidoptera","Noctuidae","Noctuinae","Feltia","Feltia subgothica, Feltia herilis, Feltia tricosa","2","2","2","144","0","0","99.55","447.5","0","450","31.11","0"
"AGAW2","10N","INS","BOLD:ACV4278","Arthropoda","Insecta","Diptera","Limoniidae","Chioneinae","Erioptera","Erioptera chlorophylla","2","2","2","144","0","0","98.43","446","0","449","33.3","0"
"AGAW2","10N","INS","BOLD:ABZ4994","Arthropoda","Insecta","Orthoptera","Acrididae","Melanoplinae","Melanoplus","Melanoplus gladstoni, Melanoplus sanguinipes","2","2","2","144","0","0","97.1","430.5","0","432","35.04","0"
"AGAW2","10N","INS","BOLD:ACX7686","Arthropoda","Insecta","Diptera","Cecidomyiidae","","","","2","2","2","144","0","0","99.87","395","0","450","28.33","0"
"AGAW2","10N","INS","BOLD:ADB5456","Arthropoda","Insecta","Hymenoptera","Eulophidae","Entedoninae","Neochrysocharis","","2","2","2","144","0","0","99.15","413","0","450","25.11","0"
"AGAW2","10N","INS","BOLD:ABA7198","Arthropoda","Insecta","Diptera","Xylomyidae","","Xylomya","Xylomya aterrima, Xylomya tenthredinoides","2","2","2","144","0","0","99.62","424.5","0","426.5","34.19","0"
"AGAW2","10N","INS","BOLD:AAP3767","Arthropoda","Insecta","Diptera","Cecidomyiidae","","","","2","2","2","144","0","0","98.22","449","0","450","29","0"
"AGAW2","10N","INS","BOLD:ADH9310","Arthropoda","Insecta","Diptera","Anthomyiidae","Anthomyiinae","Anthomyia","Anthomyia procellaris, Anthomyia pluvialis","2","2","2","144","0","0","99.08","409","0","409","32.53","0"
"AGAW2","10N","INS","BOLD:ACF5831","Arthropoda","Insecta","Diptera","Chloropidae","","","","2","2","2","144","0","0","97.75","445.5","0","450","32.44","0"
"AGAW2","10N","INS","BOLD:AAN5162","Arthropoda","Insecta","Diptera","Ceratopogonidae","Dasyheleinae","Dasyhelea","","1","1","1","180","0","0","99.56","450","0","450","34.22","0"
"AGAW2","10N","INS","BOLD:AAN5161","Arthropoda","Insecta","Diptera","Ceratopogonidae","Dasyheleinae","Dasyhelea","","1","1","1","180","0","0","98","449","0","450","36.89","0"
"AGAW2","10N","INS","BOLD:AAA9899","Arthropoda","Insecta","Hemiptera","Aphididae","Aphidinae","Rhopalosiphum","Rhopalosiphum padi","1","1","1","180","0","0","99.23","388","0","390","24.62","0"
"AGAW2","10N","INS","BOLD:ACL1784","Arthropoda","Insecta","Psocodea","Lachesillidae","Lachesillinae","Lachesilla","Lachesilla forcepeta","1","1","1","180","0","0","99.32","442","0","449","37.42","0"
"AGAW2","10N","INS","BOLD:AAM9105","Arthropoda","Insecta","Diptera","Muscidae","Coenosiinae","Coenosia","","1","1","1","180","0","0","99.11","450","0","450","37.11","0"
"AGAW2","10N","INS","BOLD:AAA8933","Arthropoda","Insecta","Coleoptera","Coccinellidae","Coccinellinae","Coccinella","Coccinella septempunctata","1","1","1","180","0","0","99.11","447","0","450","32.89","0"
"AGAW2","10N","INS","BOLD:ABY9688","Arthropoda","Insecta","Diptera","Chloropidae","Chloropinae","Thaumatomyia","Thaumatomyia glabra","1","1","1","180","0","0","98.25","400","0","405","30.37","0"
"AGAW2","10N","INS","BOLD:ACA9905","Arthropoda","Insecta","Diptera","Limoniidae","Chioneinae","Erioptera","Erioptera chlorophylla","1","1","1","180","0","0","99.78","450","0","450","33.56","0"
"AGAW2","10N","INS","BOLD:AAB4658","Arthropoda","Insecta","Diptera","Chironomidae","Chironominae","Chironomus","Chironomus dilutus","1","1","1","180","0","0","97.33","449","0","450","38.89","0"
"AGAW2","10N","INS","BOLD:AAC1042","Arthropoda","Insecta","Lepidoptera","Crambidae","Spilomelinae","Lygropia","Lygropia rivulalis","1","1","1","180","0","0","98","450","0","450","28.67","0"
"AGAW2","10N","INS","BOLD:AAL1297","Arthropoda","Insecta","Coleoptera","Cantharidae","Cantharinae","Rhagonycha","Rhagonycha fulva","1","1","1","180","0","0","99.75","400","0","404","32.67","0"
"AGAW2","10N","INS","BOLD:ADF2969","Arthropoda","Insecta","Diptera","Sarcophagidae","Sarcophaginae","Blaesoxipha","Blaesoxipha falciformis","1","1","1","180","0","0","99.5","404","0","405","37.28","0"
"AGAW2","10N","INS","BOLD:AAB8271","Arthropoda","Insecta","Lepidoptera","Autostichidae","Oegoconiinae","Oegoconia","Oegoconia quadripuncta, Oegoconia deauratella","1","1","1","180","0","0","97.89","380","0","379","35.36","0"
"AGAW2","10N","INS","BOLD:AAV5594","Arthropoda","Insecta","Diptera","Cecidomyiidae","","","","1","1","1","180","0","0","97.76","447","0","450","27.33","0"
"AGAW2","10N","INS","BOLD:AAB4657","Arthropoda","Insecta","Diptera","Chironomidae","Chironominae","Chironomus","Chironomus maturus","1","1","1","180","0","0","98.97","390","0","392","37.24","0"
"AGAW2","10N","INS","BOLD:AAN5233","Arthropoda","Insecta","Diptera","Cecidomyiidae","","","","1","1","1","180","0","0","98.89","451","0","450","26.89","0"
"AGAW2","10N","INS","BOLD:AAU6592","Arthropoda","Insecta","Diptera","Ceratopogonidae","Dasyheleinae","Dasyhelea","Dasyhelea arenivaga","1","1","1","180","0","0","100","397","0","397","34.51","0"
"AGAW2","10N","INS","BOLD:ADG6150","Arthropoda","Insecta","Diptera","Ceratopogonidae","Dasyheleinae","Dasyhelea","","1","1","1","180","0","0","100","386","0","450","38.89","0"
"AGAW2","10N","INS","BOLD:ABV1422","Arthropoda","Insecta","Diptera","Cecidomyiidae","","","","1","1","1","180","0","0","99.55","449","0","450","27.11","0"
"AGAW2","10N","INS","BOLD:ADC6100","Arthropoda","Insecta","Hemiptera","Miridae","Mirinae","Adelphocoris","Adelphocoris fasciaticollis","1","1","1","180","0","0","98.64","368","0","371","34.5","0"
"AGAW2","10N","INS","BOLD:ABY2946","Arthropoda","Insecta","Diptera","Tabanidae","Tabaninae","Hybomitra","Hybomitra trepida","1","1","1","180","0","0","98.88","448","0","450","33.11","0"
"AGAW2","10N","INS","BOLD:AAH3914","Arthropoda","Insecta","Diptera","Sciaridae","","Sciara","Sciara hamatilis, Sciara humeralis","1","1","1","180","0","0","100","450","0","450","30.22","0"
"AGAW2","10N","INS","BOLD:AAG2717","Arthropoda","Insecta","Orthoptera","Tettigoniidae","Phaneropterinae","Scudderia","Scudderia curvicauda, Scudderia pistillata","1","1","1","180","0","0","99.11","449","0","450","39.56","0"
"AGAW2","10N","INS","BOLD:ABW1321","Arthropoda","Insecta","Diptera","Cecidomyiidae","","","","1","1","1","180","0","0","99.33","448","0","450","32","0"
"AGAW2","10N","INS","BOLD:AAB7869","Arthropoda","Insecta","Lepidoptera","Tortricidae","Tortricinae","Phtheochroa","Phtheochroa baracana, Phtheochroa riscana, Phtheochroa birdana, Phtheochroa fulviplicana, Phtheochroa canariana","1","1","1","180","0","0","98.89","449","0","450","30.67","0"
"AGAW2","10N","INS","BOLD:ACI9753","Arthropoda","Insecta","Diptera","Dolichopodidae","Diaphorinae","Chrysotus","","1","1","1","180","0","0","97.55","408","0","429","33.1","0"
"AGAW2","10N","INS","BOLD:AAB5609","Arthropoda","Insecta","Diptera","Muscidae","Coenosiinae","Coenosia","Coenosia sp.BOLD:AAB5609, Coenosia tigrina","1","1","1","180","0","0","99.75","402","0","402","32.09","0"
"AGAW2","10N","INS","BOLD:AAB6702","Arthropoda","Insecta","Coleoptera","Cerambycidae","Cerambycinae","Clytus","Clytus ruricola","1","1","1","180","0","0","99.3","429","0","450","38.22","0"
"AGAW2","10N","INS","BOLD:ACE7482","Arthropoda","Insecta","Hemiptera","Miridae","","","","1","1","1","180","0","0","97.05","441","0","442","36.43","0"
"AGAW2","10N","INS","BOLD:AAM6295","Arthropoda","Insecta","Diptera","Chaoboridae","","","","1","1","1","180","0","0","100","450","0","450","37.11","0"
"AGAW2","10N","INS","BOLD:ACX5247","Arthropoda","Insecta","Diptera","Muscidae","Coenosiinae","Pentacricia","Pentacricia aldrichii","1","1","1","180","0","0","98.89","449","0","450","29.11","0"
"AGAW2","10N","INS","BOLD:AAA9251","Arthropoda","Insecta","Lepidoptera","Gelechiidae","Gelechiinae","Scrobipalpa","Scrobipalpa atriplicella, Scrobipalpa nr. atriplicella sp. 1","1","1","1","180","0","0","99.54","432","0","434","35.25","0"
"AGAW2","10N","INS","BOLD:AAV3890","Arthropoda","Insecta","Diptera","Dolichopodidae","","","","1","1","1","180","0","0","99.06","425","0","426","38.26","0"
"AGAW2","10N","INS","BOLD:AAB0269","Arthropoda","Insecta","Lepidoptera","Tortricidae","Tortricinae","Acleris","Acleris albicomana","1","1","1","180","0","0","98.22","450","0","450","33.33","0"
"AGAW2","10N","INS","BOLD:AAW5869","Arthropoda","Insecta","Coleoptera","Dermestidae","Megatominae","Anthrenus","Anthrenus sp. SLB-2003","1","1","1","180","0","0","100","443","0","450","49.33","0"
"AGAW2","10N","INS","BOLD:AAN5342","Arthropoda","Insecta","Diptera","Chironomidae","Orthocladiinae","Heleniella","","1","1","1","180","0","0","100","439","0","441","35.6","0"
"AGAW2","10N","INS","BOLD:AAE3943","Arthropoda","Insecta","Hemiptera","Miridae","Phylinae","Plagiognathus","Plagiognathus chrysanthemi","1","1","1","180","0","0","99.74","392","0","450","36.89","0"
"AGAW2","10N","INS","BOLD:AAA3867","Arthropoda","Insecta","Lepidoptera","Noctuidae","Noctuinae","Apamea","Apamea amputatrix","1","1","1","180","0","0","98.43","447","0","450","33.11","0"
"AGAW2","10N","INS","BOLD:AAB1352","Arthropoda","Insecta","Lepidoptera","Erebidae","Hypeninae","Hypena","Hypena sp.","1","1","1","180","0","0","99.56","450","0","450","32.44","0"
"AGAW2","10N","INS","BOLD:AAB3533","Arthropoda","Insecta","Lepidoptera","Geometridae","Sterrhinae","Haematopis","Haematopis grataria","1","1","1","180","0","0","100","403","0","404","30.69","0"
"AGAW2","10N","INS","BOLD:AAH0152","Arthropoda","Insecta","Coleoptera","Cantharidae","Cantharinae","Cantharis","Cantharis rufa","1","1","1","180","0","0","98.99","398","0","399","34.84","0"
"AGAW2","10N","INS","BOLD:AAG2898","Arthropoda","Insecta","Hemiptera","Cicadellidae","Deltocephalinae","Paraphlepsius","Paraphlepsius apertus, Paraphlepsius irroratus","1","1","1","180","0","0","97.78","450","0","450","32.22","0"
"AGAW2","10N","INS","BOLD:AAD1169","Arthropoda","Insecta","Lepidoptera","Cosmopterigidae","Cosmopteriginae","Cosmopterix","Cosmopterix scirpicola, Cosmopterix fernaldella","1","1","1","180","0","0","100","450","0","450","32.89","0"
"AGAW2","10N","INS","BOLD:AAE0845","Arthropoda","Insecta","Hemiptera","Pentatomidae","Pentatominae","Euschistus","Euschistus servus","1","1","1","180","0","0","97.72","438","0","438","38.58","0"
"AGAW2","10N","INS","BOLD:AAI8935","Arthropoda","Insecta","Coleoptera","Latridiidae","Corticariinae","Cortinicara","Cortinicara gibbosa","1","1","1","180","0","0","99.11","449","0","450","33.78","0"
"AGAW2","10N","INS","BOLD:AAY6753","Arthropoda","Insecta","Hemiptera","Miridae","Orthotylinae","Paraproba","Paraproba capitata","1","1","1","180","0","0","98.88","445","0","450","36.44","0"
"AGAW2","10N","INS","BOLD:AAQ0271","Arthropoda","Insecta","Diptera","Dolichopodidae","","","","1","1","1","180","0","0","99.33","450","0","450","37.11","0"
"AGAW2","10N","INS","BOLD:AAZ2075","Arthropoda","Insecta","Hemiptera","Anthocoridae","Anthocorinae","Orius","Orius insidiosus","1","1","1","180","0","0","99.78","450","0","450","35.11","0"
"AGAW2","10N","INS","BOLD:AAG6471","Arthropoda","Insecta","Diptera","Ceratopogonidae","Dasyheleinae","Dasyhelea","","1","1","1","180","0","0","99.25","400","0","404","32.92","0"
"AGAW2","10N","INS","BOLD:AAG8670","Arthropoda","Insecta","Hemiptera","Miridae","Mirinae","Phytocoris","","1","1","1","180","0","0","98.89","449","0","450","34.22","0"
"AGAW2","10N","INS","BOLD:ACB0984","Arthropoda","Insecta","Psocodea","Stenopsocidae","","Graphopsocus","Graphopsocus cruciatus","1","1","1","180","0","0","99.28","414","0","450","37.78","0"
"AGAW2","10N","INS","BOLD:AAC0706","Arthropoda","Insecta","Diptera","Chironomidae","Chironominae","Dicrotendipes","Dicrotendipes tritomus","1","1","1","180","0","0","98.66","447","0","448","35.71","0"
"AGAW2","10N","INS","BOLD:AAB9516","Arthropoda","Insecta","Diptera","Chironomidae","Orthocladiinae","Acricotopus","","1","1","1","180","0","0","97.33","449","0","450","36.22","0"
"AGAW2","10N","INS","BOLD:AAY6534","Arthropoda","Insecta","Coleoptera","Mordellidae","Mordellinae","Mordellina","Mordellina ancilla","1","1","1","180","0","0","98.89","450","0","450","38.44","0"
"AGAW2","10N","NAR","BOLD:AAL1512","Mollusca","Gastropoda","Stylommatophora","Agriolimacidae",NA,"Deroceras","Deroceras laeve","260","257","257","1","1.15","0","97.7","393.98","0","397.54","33.43","0"
"AGAW2","10N","NAR","TAX:660402","Mollusca","Gastropoda","Stylommatophora","Agriolimacidae",NA,"Deroceras","Deroceras sp.","2","2","2","2","0","0","97.62","418.5","0","422","33.74","0"
"AGAW2","10N","NAR","BOLD:AAI9663","Mollusca","Gastropoda","Stylommatophora","Agriolimacidae",NA,"Deroceras","Deroceras reticulatum","1","1","1","3","0","0","97.44","391","0","391","35.55","0"
"AGAW2","10N","NIN","BOLD:AAH6630","Arthropoda","Collembola","Entomobryomorpha","Entomobryidae","Entomobryinae","Entomobrya","Entomobrya unostrigata","2583","2114","2114","1","18.16","0","99.39","436.94","0","438.11","39.05","0"
"AGAW2","10N","NIN","BOLD:ABW2639","Arthropoda","Arachnida","Trombidiformes","Anystidae","","","","518","419","419","2","19.11","0","99.52","446.47","0","445.39","31.46","0"
"AGAW2","10N","NIN","BOLD:AAA6381","Arthropoda","Arachnida","Araneae","Tetragnathidae","","Tetragnatha","Tetragnatha laboriosa","385","369","369","3","4.16","0","98.91","430.41","0","432.75","39.54","0"
"AGAW2","10N","NIN","BOLD:AAA8820","Arthropoda","Collembola","Entomobryomorpha","Entomobryidae","Lepidocyrtinae","Lepidocyrtus","Lepidocyrtus cyaneus","246","229","229","4","6.91","0","98.86","410.08","0","412.23","43.29","0"
"AGAW2","10N","NIN","BOLD:AAA6383","","","","","","","","104","90","90","5","13.46","0","98.07","434.8","0","436.41","37.51","0"
"AGAW2","10N","NIN","BOLD:AAB7914","Arthropoda","Collembola","Symphypleona","Bourletiellidae","","Bourletiella","","86","64","64","6","25.58","0","99.71","444.65","0","445.83","38.08","0"
"AGAW2","10N","NIN","BOLD:ACZ1295","Arthropoda","Arachnida","Trombidiformes","Anystidae","","","","64","64","64","7","0","0","99.21","374.33","0","388.84","31.34","0"
"AGAW2","10N","NIN","BOLD:AAC0175","Arthropoda","Arachnida","Araneae","Theridiidae","","Parasteatoda","Parasteatoda tepidariorum, Parasteatoda simulans","40","37","37","8","7.5","0","99.35","435.2","0","437.5","31.9","0"
"AGAW2","10N","NIN","BOLD:AAB8452","Arthropoda","Collembola","Entomobryomorpha","Entomobryidae","Lepidocyrtinae","Lepidocyrtus","Lepidocyrtus paradoxus","8","7","7","9","12.5","0","99.91","443.25","0","445.75","42.05","0"
"AGAW2","10N","NIN","BOLD:ACX8969","Arthropoda","Arachnida","Trombidiformes","Eupodidae","","","","8","8","8","9","0","0","99.11","411.13","0","414.25","27.3","0"
"AGAW2","10N","NIN","BOLD:AAU6252","Arthropoda","Arachnida","Trombidiformes","Bdellidae","","Cyta","","4","4","4","11","0","0","99.83","446.5","0","448.25","31.62","0"
"AGAW2","10N","NIN","BOLD:ACI9974","Arthropoda","Arachnida","Trombidiformes","Tydeidae","","","","3","3","3","12","0","0","99.53","423.67","0","450","40.74","0"
"AGAW2","10N","NIN","BOLD:AAA7246","Arthropoda","Collembola","Entomobryomorpha","Entomobryidae","Entomobryinae","Entomobrya","Entomobrya nicoleti, Entomobrya multifasciata, Entomobrya intermedia","1","1","1","13","0","0","99.11","449","0","450","44","0"
"AGAW2","10N","NIN","BOLD:ACM6751","Arthropoda","Arachnida","Trombidiformes","Anystidae","","","","1","1","1","13","0","0","98.76","403","0","450","29.56","0"
"AGAW2","10N","NIN","BOLD:ACX8940","Arthropoda","Arachnida","Trombidiformes","Tarsonemidae","","","","1","1","1","13","0","0","97.33","450","0","450","25.33","0"
"AGAW2","10N","NIN","BOLD:AAO8147","Arthropoda","Arachnida","Trombidiformes","Eupodidae","","Eupodes","","1","1","1","13","0","0","99.55","448","0","450","28.67","0"
"AGAW2","10N","NIN","BOLD:ACY2550","Arthropoda","Arachnida","Trombidiformes","Bdellidae","","Cyta","","1","1","1","13","0","0","99.72","358","0","360","31.11","0"
"AGAW2","10N","NIN","BOLD:ACP9010","Arthropoda","Collembola","Symphypleona","Bourletiellidae","","","","1","1","1","13","0","0","98.89","450","0","450","38.89","0"
"AGAW2","10N","NIN","BOLD:AAN6627","Arthropoda","Arachnida","Mesostigmata","Parasitidae","","","","1","1","1","13","0","0","99.32","441","0","450","39.11","0"
"AGAW2","10S","BAC","TAX:703648","Proteobacteria","Alphaproteobacteria","Rickettsiales","Rickettsiaceae",NA,"Wolbachia","Wolbachia endosymbiont","1","1","1","1","0","0","97.72","395","0","413","39.47","0"
"AGAW2","10S","BAC","TAX:91391","Proteobacteria","Alphaproteobacteria","Rickettsiales","",NA,"","","1","1","1","1","0","0","98.11","423","0","450","40.22","0"
"AGAW2","10S","CHO","no match","no match","no match","no match","no match","no match","no match","no match","no match","no match","no match","no match","no match","no match","no match","no match","no match","no match","no match","no match"
"AGAW2","10S","INS","BOLD:AAG1711","Arthropoda","Insecta","Diptera","Muscidae","Phaoniinae","Helina","Helina depuncta","29325","22233","22233","1","24.18","0","99.45","433.02","0","435.37","31.34","0"
"AGAW2","10S","INS","BOLD:AAA3453","Arthropoda","Insecta","Diptera","Anthomyiidae","Anthomyiinae","Delia","","11399","9731","9731","2","14.63","0","99.4","429.07","0","431.74","34.61","0"
"AGAW2","10S","INS","BOLD:AAA4555","Arthropoda","Insecta","Orthoptera","Acrididae","Melanoplinae","Melanoplus","Melanoplus stupefactus, Melanoplus angustipennis, Melanoplus ponderosus, Melanoplus bruneri, Melanoplus frigidus, Melanoplus scapularis, Melanoplus cinereus, Melanoplus madeleineae, Melanoplus sanguinipes, Melanoplus nigrescens group, Melanoplus tunicae, Melanoplus walshi, Melanoplus femurrubrum, Melanoplus dawsoni, Melanoplus sp., Melanoplus fasciatus, Melanoplus bowditchi, Melanoplus stonei, Melanoplus bivittatus, Melanoplus huroni, Melanoplus thomasi, Melanoplus borealis, Melanoplus packardii","9430","7028","7028","3","25.47","0","99.38","446.9","0","448.09","36.21","0"
"AGAW2","10S","INS","BOLD:ACE7444","Arthropoda","Insecta","Hemiptera","Miridae","Mirinae, Phylinae","Adelphocoris, Megalocoleus","Adelphocoris lineolatus, Megalocoleus molliculus","6076","3786","3786","4","37.69","0","99.7","439.81","0","440.6","34.79","0"
"AGAW2","10S","INS","BOLD:AAG6745","Arthropoda","Insecta","Diptera","Calliphoridae","Calliphorinae","Pollenia","Pollenia pediculata","5064","4253","4253","5","16.02","0","99.36","433.72","0","435.75","33.46","0"
"AGAW2","10S","INS","BOLD:AAN6154","Arthropoda","Insecta","Coleoptera","Latridiidae","Corticariinae","Melanophthalma","Melanophthalma inermis","3735","3117","3117","6","16.55","0","98.92","435.84","0","437.41","31.23","0"
"AGAW2","10S","INS","BOLD:AAI4303","Arthropoda","Insecta","Diptera","Chironomidae","Chironominae","Chironomus","Chironomus melanescens","1773","1608","1608","7","9.31","0","98.18","426.93","0","431.36","33.98","0"
"AGAW2","10S","INS","BOLD:AAH3225","Arthropoda","Insecta","Psocodea","Psocidae","Psocinae","Metylophorus","Metylophorus novaescotiae","1266","1025","1025","8","19.04","0","98.21","438.49","0","439.61","34.71","0"
"AGAW2","10S","INS","BOLD:AAU6681","Arthropoda","Insecta","Diptera","Tipulidae","Tipulinae","Nephrotoma","Nephrotoma cornicina","949","750","750","9","20.97","0","99.65","442.64","0","443.79","36.29","0"
"AGAW2","10S","INS","BOLD:AAB8271","Arthropoda","Insecta","Lepidoptera","Autostichidae","Oegoconiinae","Oegoconia","Oegoconia quadripuncta, Oegoconia deauratella","818","807","807","10","1.34","0","99","401.74","0","403.73","34.38","0"
"AGAW2","10S","INS","BOLD:AAG2868","Arthropoda","Insecta","Hemiptera","Cicadellidae","Typhlocybinae","Empoasca","Empoasca sp., Empoasca fabae","507","333","333","11","34.32","0","99.64","443.71","0","446.75","32.28","0"
"AGAW2","10S","INS","BOLD:AAH4276","Arthropoda","Insecta","Lepidoptera","Gelechiidae","Gelechiinae","Bryotropha","Bryotropha hodgesi, Bryotropha altitudophila","505","414","414","12","18.02","0","99.44","434.15","0","436","33.53","0"
"AGAW2","10S","INS","BOLD:AAG5507","Arthropoda","Insecta","Diptera","Chironomidae","Chironominae","Chironomus","Chironomus atrella","392","383","383","13","2.3","0","98.79","427.34","0","430.6","35.73","0"
"AGAW2","10S","INS","BOLD:AAD2041","Arthropoda","Insecta","Diptera","Dolichopodidae","Sciapodinae","Condylostylus","Condylostylus inermis, Condylostylus nigrofemoratus, Condylostylus connectans, Condylostylus caudatus, Condylostylus flavipes","372","281","281","14","24.46","0","99.53","441.23","0","442.53","35.93","0"
"AGAW2","10S","INS","BOLD:AAH3661","Arthropoda","Insecta","Diptera","Cecidomyiidae","","","","365","329","329","15","9.86","0","99.25","428.62","0","430.95","25.36","0"
"AGAW2","10S","INS","BOLD:ABY5318","Arthropoda","Insecta","Diptera","Chloropidae","Oscinellinae","Incertella, Rhopalopterum","Rhopalopterum umbrosum, Rhopalopterum luteiceps, Incertella bispina, Rhopalopterum painteri","336","256","256","16","23.81","0","99.08","442.15","0","443.52","32.21","0"
"AGAW2","10S","INS","BOLD:ABA6506","Arthropoda","Insecta","Diptera","Chironomidae","Orthocladiinae","Orthocladius","Orthocladius smolandicus","328","289","289","17","11.89","0","98.75","432.03","0","433.72","35.24","0"
"AGAW2","10S","INS","BOLD:AAV5575","Arthropoda","Insecta","Diptera","Cecidomyiidae","","","CecidInt30 sp. BOLD:AAV5575","274","216","216","18","21.17","0","99.54","441.74","0","442.97","25.86","0"
"AGAW2","10S","INS","BOLD:AAG2511","Arthropoda","Insecta","Diptera","Anthomyiidae","Anthomyiinae","Delia","Delia platura","264","247","247","19","6.44","0","98.92","443.88","0","446.78","33.12","0"
"AGAW2","10S","INS","BOLD:AAN5205","Arthropoda","Insecta","Diptera","Cecidomyiidae","","","CecidInt42 sp. BOLD:AAN5205","261","203","203","20","22.22","0","99.61","447.39","0","448.83","31.4","0"
"AGAW2","10S","INS","BOLD:AAG2898","Arthropoda","Insecta","Hemiptera","Cicadellidae","Deltocephalinae","Paraphlepsius","Paraphlepsius apertus, Paraphlepsius irroratus","251","184","184","21","26.69","0","99.6","440.24","0","441.18","31.85","0"
"AGAW2","10S","INS","BOLD:ACE5778","Arthropoda","Insecta","Diptera","Dolichopodidae","","","","232","220","220","22","5.17","0","99.29","429.8","0","431.48","38.02","0"
"AGAW2","10S","INS","BOLD:AAH0135","Arthropoda","Insecta","Coleoptera","Phalacridae","Phalacrinae","Acylomus","Acylomus pugetanus, Acylomus ergoti","215","197","197","23","8.37","0","99.23","435.53","0","437.77","31.63","0"
"AGAW2","10S","INS","BOLD:AAD1485","Arthropoda","Insecta","Diptera","Chironomidae","Chironominae","Paratanytarsus","Paratanytarsus grimmii, Paratanytarsus sp.","214","178","178","24","16.82","0","99.5","443.05","0","444.57","33.79","0"
"AGAW2","10S","INS","BOLD:AAN5242","Arthropoda","Insecta","Diptera","Cecidomyiidae","","","","201","200","200","25","0.5","0","98.56","402.8","0","404.92","28","0"
"AGAW2","10S","INS","BOLD:ACM0749","Arthropoda","Insecta","Coleoptera","Anthicidae","Notoxinae","Notoxus","Notoxus anchora","174","171","171","26","1.72","0","99.19","414.16","0","415.93","35.33","0"
"AGAW2","10S","INS","BOLD:AAG7276","Arthropoda","Insecta","Diptera","Sphaeroceridae","Limosininae","Leptocera","Leptocera erythrocera","173","158","158","27","8.67","0","99.19","436.67","0","438.75","32.8","0"
"AGAW2","10S","INS","BOLD:AAH3664","Arthropoda","Insecta","Diptera","Cecidomyiidae","","","CecidInt3 sp. BOLD:AAH3664","135","134","134","28","0.74","0","98.98","432.76","0","434.87","27.15","0"
"AGAW2","10S","INS","BOLD:ABW7897","Arthropoda","Insecta","Diptera","Cecidomyiidae","","","","102","75","75","29","26.47","0","99.71","444.23","0","446.39","29.71","0"
"AGAW2","10S","INS","BOLD:ACC8079","Arthropoda","Insecta","Diptera","Cecidomyiidae","","","","98","76","76","30","22.45","0","99.33","447.3","0","448.57","29.24","0"
"AGAW2","10S","INS","BOLD:ADB4698","Arthropoda","Insecta","Diptera","Cecidomyiidae","","","","93","84","84","31","9.68","0","99.39","425.14","0","426.35","31.29","0"
"AGAW2","10S","INS","BOLD:AAN5183","Arthropoda","Insecta","Diptera","Cecidomyiidae","","","","91","85","85","32","6.59","0","98.36","436.43","0","437.62","27.73","0"
"AGAW2","10S","INS","BOLD:AAF9771","Arthropoda","Insecta","Diptera","Hybotidae","Tachydromiinae","Platypalpus","Platypalpus longiseta, Platypalpus trivialis, Platypalpus sp. 5, Platypalpus sp. 17, Platypalpus pallidiventris","84","70","70","33","16.67","0","99.02","446.68","0","447.48","35.05","0"
"AGAW2","10S","INS","BOLD:AAD5253","Arthropoda","Insecta","Diptera","Chironomidae","Orthocladiinae","Thienemanniella","Thienemanniella xena, Thienemanniella vittata","74","66","66","34","10.81","0","98.61","436.07","0","439.15","34.4","0"
"AGAW2","10S","INS","BOLD:AAN5260","Arthropoda","Insecta","Diptera","Cecidomyiidae","","","","68","66","66","35","2.94","0","99.17","405.19","0","406.84","25.88","0"
"AGAW2","10S","INS","BOLD:ACC7073","Arthropoda","Insecta","Coleoptera","Mordellidae","Mordellinae","Pseudotolida","Pseudotolida arida","65","59","59","36","9.23","0","99.58","441.66","0","442.83","38.12","0"
"AGAW2","10S","INS","BOLD:AAY6730","Arthropoda","Insecta","Hemiptera","Derbidae","Cedusinae","Cedusa","Cedusa incisa","60","60","60","37","0","0","99.03","419.58","0","421.05","35.3","0"
"AGAW2","10S","INS","BOLD:ADF6661","Arthropoda","Insecta","Diptera","Cecidomyiidae","","","","57","43","43","38","24.56","0","99.24","418.25","0","448.12","29.96","0"
"AGAW2","10S","INS","BOLD:AAA5299","Arthropoda","Insecta","Diptera","Chironomidae","Orthocladiinae","Cricotopus","Cricotopus laetus, Cricotopus relucens, Cricotopus trifasciatus, Cricotopus sp., Cricotopus sp. 3ES, Cricotopus sylvestris, Cricotopus glacialis","55","51","51","39","7.27","0","99.25","427.87","0","429.27","34.27","0"
"AGAW2","10S","INS","BOLD:AAG7891","Arthropoda","Insecta","Hymenoptera","Platygastridae","Scelioninae","Telenomus","Telenomus podisi","54","51","51","40","5.56","0","98.84","426.26","0","434.07","26.95","0"
"AGAW2","10S","INS","BOLD:ACX6055","Arthropoda","Insecta","Diptera","Phoridae","Metopininae","Megaselia","Megaselia sp. AK-2010-2, Megaselia sp. BOLD:ACX6055","53","53","53","41","0","0","99.45","433.55","0","436.3","33","0"
"AGAW2","10S","INS","BOLD:AAA7661","Arthropoda","Insecta","Diptera","Culicidae","Culicinae","Culiseta, Culex","Culiseta melanura, Culex articularis, Culex restuans, Culex acharistus","49","48","48","42","2.04","0","99.34","430.8","0","432.88","32.7","0"
"AGAW2","10S","INS","BOLD:ABW1379","Arthropoda","Insecta","Diptera","Chloropidae","Oscinellinae","Malloewia","Malloewia abdominalis","48","45","45","43","6.25","0","98.32","442.77","0","444.67","31.5","0"
"AGAW2","10S","INS","BOLD:ACK3431","Arthropoda","Insecta","Diptera","Cecidomyiidae","","","","47","46","46","44","2.13","0","99.34","422.3","0","423.85","32.67","0"
"AGAW2","10S","INS","BOLD:ABA0863","Arthropoda","Insecta","Diptera","Cecidomyiidae","","","","46","34","34","45","26.09","0","99.59","434.61","0","435.74","29.66","0"
"AGAW2","10S","INS","BOLD:AAJ2720","Arthropoda","Insecta","Coleoptera","Staphylinidae","Aleocharinae","Aleochara","Aleochara verna","44","44","44","46","0","0","98.64","408.93","0","410.27","30.03","0"
"AGAW2","10S","INS","BOLD:ABZ5071","Arthropoda","Insecta","Diptera","Chironomidae","Chironominae","Micropsectra","Micropsectra nigripila","41","41","41","47","0","0","98.2","428.27","0","443.44","37.03","0"
"AGAW2","10S","INS","BOLD:ACB6899","Arthropoda","Insecta","Diptera","Cecidomyiidae","","","","41","36","36","47","12.2","0","99.63","415.93","0","442.8","26.22","0"
"AGAW2","10S","INS","BOLD:AAN5199","Arthropoda","Insecta","Diptera","Cecidomyiidae","","","CecidInt20 sp. BOLD:AAN5199","41","35","35","47","14.63","0","99.53","439.76","0","442.54","26.51","0"
"AGAW2","10S","INS","BOLD:AAG3236","Arthropoda","Insecta","Diptera","Phoridae","Phorinae","Conicera, Diplonevra","Diplonevra nitidula","36","35","35","50","2.78","0","99.57","448.22","0","450","31.96","0"
"AGAW2","10S","INS","BOLD:ABV1194","Arthropoda","Insecta","Diptera","Chironomidae","Chironominae","Micropsectra","","35","31","31","51","11.43","0","99.55","432.23","0","434.11","35.88","0"
"AGAW2","10S","INS","BOLD:ACK5742","Arthropoda","Insecta","Diptera","Cecidomyiidae","","","","32","28","28","52","12.5","0","99.56","440.16","0","441.25","28.87","0"
"AGAW2","10S","INS","BOLD:AAN8770","Arthropoda","Insecta","Diptera","Psychodidae","Psychodinae","Psychoda","Psychoda trinodulosa","32","31","31","52","3.13","0","99.43","441.03","0","441.56","33.72","0"
"AGAW2","10S","INS","BOLD:AAA4277","Arthropoda","Insecta","Diptera","Syrphidae","Syrphinae","Toxomerus","Toxomerus marginatus, Toxomerus sp. CR-3, Toxomerus CR-3","31","28","28","54","9.68","0","99.35","446.61","0","447.84","30.62","0"
"AGAW2","10S","INS","BOLD:ABU5525","Arthropoda","Insecta","Diptera","Chironomidae","Orthocladiinae","Limnophyes","Limnophyes sp. 14ES","31","31","31","54","0","0","98.99","405.81","0","408.84","36.92","0"
"AGAW2","10S","INS","BOLD:AAP8739","Arthropoda","Insecta","Diptera","Platystomatidae","Platystomatinae","Rivellia","Rivellia steyskali","30","29","29","56","3.33","0","99.27","442.13","0","443.23","38.06","0"
"AGAW2","10S","INS","BOLD:AAN5148","Arthropoda","Insecta","Diptera","Ceratopogonidae","Forcipomyiinae","Forcipomyia","Forcipomyia bipunctata, Forcipomyia sp. CK-2012","29","24","24","57","17.24","0","98.8","434.31","0","435.48","35.69","0"
"AGAW2","10S","INS","BOLD:AAC0903","Arthropoda","Insecta","Diptera","Chironomidae","Chironominae","Chironomus","Chironomus acidophilus","28","28","28","58","0","0","99.36","436.18","0","438.11","36.24","0"
"AGAW2","10S","INS","BOLD:ACJ6954","Arthropoda","Insecta","Diptera","Cecidomyiidae","","","","27","27","27","59","0","0","99.48","337.89","0","428.19","26.61","0"
"AGAW2","10S","INS","BOLD:ABA0752","Arthropoda","Insecta","Diptera","Agromyzidae","Phytomyzinae","Amauromyza","Amauromyza karli","25","25","25","60","0","0","99.13","426","0","427.56","34.01","0"
"AGAW2","10S","INS","BOLD:ACS4012","Arthropoda","Insecta","Diptera","Cecidomyiidae","","","","24","19","19","61","20.83","0","98.76","415.88","0","440.58","27.1","0"
"AGAW2","10S","INS","BOLD:AAZ5592","Arthropoda","Insecta","Diptera","Psychodidae","Psychodinae","","","24","24","24","61","0","0","99.32","428.42","0","431.63","32.47","0"
"AGAW2","10S","INS","BOLD:AAS1281","Arthropoda","Insecta","Diptera","Chironomidae","Chironominae","Chironomus","Chironomus nr. sollicitus","21","15","15","63","28.57","0","97.11","447.62","0","450","33.61","0"
"AGAW2","10S","INS","BOLD:ACG3646","Arthropoda","Insecta","Coleoptera","Corylophidae","Corylophinae","Orthoperus","","21","19","19","63","9.52","0","98.34","445.52","0","446.76","34.18","0"
"AGAW2","10S","INS","BOLD:AAV1261","Arthropoda","Insecta","Diptera","Sciaridae","","Bradysia","Bradysia splendida","21","20","20","63","4.76","0","99.17","431.67","0","432.9","34.07","0"
"AGAW2","10S","INS","BOLD:ABV1297","Arthropoda","Insecta","Diptera","Sciaridae","","","","19","19","19","66","0","0","99.3","432.11","0","434.42","30.33","0"
"AGAW2","10S","INS","BOLD:ACF2457","Arthropoda","Insecta","Diptera","Chironomidae","Chironominae","Paratanytarsus","Paratanytarsus laccophilus","19","19","19","66","0","0","99.32","444.32","0","445.79","32.57","0"
"AGAW2","10S","INS","BOLD:AAV5689","Arthropoda","Insecta","Diptera","Cecidomyiidae","","","","17","17","17","68","0","0","98.59","440.18","0","441.65","26.29","0"
"AGAW2","10S","INS","BOLD:ACX5521","Arthropoda","Insecta","Diptera","Chironomidae","Orthocladiinae","","","17","17","17","68","0","0","99.09","404.24","0","406.47","34.99","0"
"AGAW2","10S","INS","BOLD:AAA4222","Arthropoda","Insecta","Lepidoptera","Erebidae","Hypeninae","Hypena","Hypena scabra","15","15","15","70","0","0","99.31","402.07","0","404","32.51","0"
"AGAW2","10S","INS","BOLD:AAN6447","Arthropoda","Insecta","Diptera","Sciaridae","","Corynoptera","Corynoptera perpusilla","12","11","11","71","8.33","0","99.37","448.25","0","449.67","36.43","0"
"AGAW2","10S","INS","BOLD:AAA4781","Arthropoda","Insecta","Hymenoptera","Braconidae","Microgastrinae","Glyptapanteles","Glyptapanteles carlossarmientoi","12","12","12","71","0","0","99.27","419.75","0","424.67","23.78","0"
"AGAW2","10S","INS","BOLD:ACG5862","Arthropoda","Insecta","Diptera","Agromyzidae","","","","11","11","11","73","0","0","98.36","425.36","0","428","33.73","0"
"AGAW2","10S","INS","BOLD:ACF1257","","","","","","","","11","11","11","73","0","0","99.64","448.82","0","450","34.57","0"
"AGAW2","10S","INS","BOLD:AAH0134","Arthropoda","Insecta","Coleoptera","Phalacridae","Phalacrinae","Stilbus","Stilbus apicialis","10","10","10","75","0","0","99.61","437.3","0","438.8","32.11","0"
"AGAW2","10S","INS","BOLD:AAL7819","Arthropoda","Insecta","Diptera","Psychodidae","Psychodinae","Psychoda","Psychoda sp., Psychoda sp. BOLD:AAL7819","9","9","9","76","0","0","99.35","415.11","0","416.78","32.76","0"
"AGAW2","10S","INS","BOLD:AAK5928","Arthropoda","Insecta","Coleoptera","Chrysomelidae","Criocerinae","Oulema","Oulema melanopus","9","9","9","76","0","0","99.33","435","0","435.78","36.38","0"
"AGAW2","10S","INS","BOLD:ACE8878","Arthropoda","Insecta","Orthoptera","Acrididae","Melanoplinae","Melanoplus","Melanoplus scudderi group","8","8","8","78","0","0","97.45","450.25","0","450","35.81","0"
"AGAW2","10S","INS","BOLD:AAO3132","Arthropoda","Insecta","Coleoptera","Chrysomelidae","Galerucinae","Neocrepidodera","Neocrepidodera ferruginea","8","8","8","78","0","0","98.79","370.13","0","373.63","34.77","0"
"AGAW2","10S","INS","BOLD:ABZ4994","Arthropoda","Insecta","Orthoptera","Acrididae","Melanoplinae","Melanoplus","Melanoplus gladstoni, Melanoplus sanguinipes","8","8","8","78","0","0","97.42","449.75","0","450","36.42","0"
"AGAW2","10S","INS","BOLD:ACC5649","Arthropoda","Insecta","Diptera","Cecidomyiidae","","","CecidInt23 sp. BOLD:ACC5649","7","7","7","81","0","0","99.3","422.43","0","424.43","25.15","0"
"AGAW2","10S","INS","BOLD:AAN5238","Arthropoda","Insecta","Diptera","Cecidomyiidae","","","","7","6","6","81","14.29","0","99.52","433.43","0","434.29","25.49","0"
"AGAW2","10S","INS","BOLD:AAC1374","Arthropoda","Insecta","Hemiptera","Aphididae","Aphidinae","Aphis","Aphis nasturtii, Aphis davletshinae, Aphis sp. rostella like, Aphis cf. rostella, Aphis althaeae, Aphis umbrella","7","7","7","81","0","0","99.4","442.43","0","443.29","26.01","0"
"AGAW2","10S","INS","BOLD:AAV6213","Arthropoda","Insecta","Diptera","Chironomidae","Chironominae","Paratanytarsus","","7","7","7","81","0","0","99.32","421.86","0","424.14","34.39","0"
"AGAW2","10S","INS","BOLD:AAQ0558","Arthropoda","Insecta","Thysanoptera","Thripidae","Thripinae","Anaphothrips","Anaphothrips obscurus","6","6","6","85","0","0","98.96","441.17","0","442.83","31.42","0"
"AGAW2","10S","INS","BOLD:AAO8538","Arthropoda","Insecta","Diptera","Psychodidae","Psychodinae","Psychoda","","6","6","6","85","0","0","98.84","422.83","0","424.5","32.88","0"
"AGAW2","10S","INS","BOLD:ABA7871","Arthropoda","Insecta","Diptera","Cecidomyiidae","","","","6","5","5","85","16.67","0","99.44","444.83","0","445.33","26.82","0"
"AGAW2","10S","INS","BOLD:ACP6694","Arthropoda","Insecta","Diptera","Chironomidae","Orthocladiinae","","","6","6","6","85","0","0","99.04","438.33","0","440.67","33.62","0"
"AGAW2","10S","INS","BOLD:AAG2123","Arthropoda","Insecta","Diptera","Tachinidae","Dexiinae","Dinera","Dinera grisescens","6","6","6","85","0","0","99.05","437.83","0","439","31.87","0"
"AGAW2","10S","INS","BOLD:AAA8933","Arthropoda","Insecta","Coleoptera","Coccinellidae","Coccinellinae","Coccinella","Coccinella septempunctata","5","5","5","90","0","0","99.78","448.4","0","448.8","33.2","0"
"AGAW2","10S","INS","BOLD:AAG5453","Arthropoda","Insecta","Diptera","Chironomidae","Chironominae","Chironomus","Chironomus bifurcatus","5","5","5","90","0","0","98.35","448.8","0","450","35.07","0"
"AGAW2","10S","INS","BOLD:AAA3181","Arthropoda","Insecta","Diptera","Muscidae","Muscinae","Stomoxys","Stomoxys calcitrans","4","4","4","92","0","0","99.2","405.75","0","409.5","33.08","0"
"AGAW2","10S","INS","BOLD:AAP7026","Arthropoda","Insecta","Coleoptera","Latridiidae","Corticariinae","Melanophthalma","Melanophthalma villosa","4","4","4","92","0","0","98.98","400","0","446.75","31.78","0"
"AGAW2","10S","INS","BOLD:ACU2412","Arthropoda","Insecta","Diptera","Cecidomyiidae","Cecidomyiinae","Contarinia","","4","4","4","92","0","0","98.57","397","0","397.25","26.28","0"
"AGAW2","10S","INS","BOLD:ACF4925","Arthropoda","Insecta","Hemiptera","Miridae","Mirinae","Adelphocoris","Adelphocoris detritus","4","4","4","92","0","0","97.61","449","0","450","36.28","0"
"AGAW2","10S","INS","BOLD:AAN5220","Arthropoda","Insecta","Diptera","Cecidomyiidae","","","","4","4","4","92","0","0","98.4","402.5","0","415","25.81","0"
"AGAW2","10S","INS","BOLD:ACV5271","Arthropoda","Insecta","Diptera","Cecidomyiidae","","","","4","4","4","92","0","0","99.38","430","0","432.25","28.24","0"
"AGAW2","10S","INS","BOLD:AAF9306","Arthropoda","Insecta","Diptera","Psychodidae","Psychodinae","Psychoda","Psychoda alternata","4","4","4","92","0","0","98.54","445.75","0","448","31.42","0"
"AGAW2","10S","INS","BOLD:ABY5257","Arthropoda","Insecta","Lepidoptera","Noctuidae","Noctuinae","Apamea","Apamea devastator","4","4","4","92","0","0","99.21","447.5","0","447.75","30.94","0"
"AGAW2","10S","INS","BOLD:AAC0022","Arthropoda","Insecta","Diptera","Chironomidae","Chironominae","Micropsectra","Micropsectra nigripila","3","3","3","100","0","0","97.4","448.67","0","450","37.11","0"
"AGAW2","10S","INS","BOLD:AAB4658","Arthropoda","Insecta","Diptera","Chironomidae","Chironominae","Chironomus","Chironomus dilutus","3","3","3","100","0","0","98.19","441.67","0","443.67","37.79","0"
"AGAW2","10S","INS","BOLD:AAL1297","Arthropoda","Insecta","Coleoptera","Cantharidae","Cantharinae","Rhagonycha","Rhagonycha fulva","3","3","3","100","0","0","98.3","423.67","0","429","33.18","0"
"AGAW2","10S","INS","BOLD:AAN5529","Arthropoda","Insecta","Diptera","Dolichopodidae","","","","3","3","3","100","0","0","98.15","416.67","0","418.33","38.21","0"
"AGAW2","10S","INS","BOLD:AAN5230","Arthropoda","Insecta","Diptera","Cecidomyiidae","","","","3","3","3","100","0","0","99.52","419.67","0","421.33","25.75","0"
"AGAW2","10S","INS","BOLD:ABW3189","Arthropoda","Insecta","Hymenoptera","Platygastridae","Scelioninae","Telenomus","","3","3","3","100","0","0","98.9","408.67","0","416.33","24.79","0"
"AGAW2","10S","INS","BOLD:ACU6062","Arthropoda","Insecta","Diptera","Cecidomyiidae","","","","3","3","3","100","0","0","99.14","377.33","0","380.33","25.16","0"
"AGAW2","10S","INS","BOLD:AAB4894","Arthropoda","Insecta","Hemiptera","Aphididae","Aphidinae","Sitobion","Sitobion smilacifoliae, Sitobion avenae, Sitobion phyllanthi","3","3","3","100","0","0","97.48","437.67","0","438","25","0"
"AGAW2","10S","INS","BOLD:AAH2293","Arthropoda","Insecta","Diptera","Asilidae","Stenopogoninae","Dioctria, Eudioctria","Dioctria hyalipennis, Dioctria baumhaueri, Eudioctria propinqua","3","3","3","100","0","0","99.01","423","0","425.33","30.23","0"
"AGAW2","10S","INS","BOLD:AAA9251","Arthropoda","Insecta","Lepidoptera","Gelechiidae","Gelechiinae","Scrobipalpa","Scrobipalpa atriplicella, Scrobipalpa nr. atriplicella sp. 1","3","3","3","100","0","0","99.15","398.33","0","399","34.47","0"
"AGAW2","10S","INS","BOLD:AAN5342","Arthropoda","Insecta","Diptera","Chironomidae","Orthocladiinae","Heleniella","","3","3","3","100","0","0","98.54","411.33","0","412.67","35.11","0"
"AGAW2","10S","INS","BOLD:ADH9867","Arthropoda","Insecta","Diptera","","","","","3","3","3","100","0","0","98.81","447","0","450","37.33","0"
"AGAW2","10S","INS","BOLD:AAD1169","Arthropoda","Insecta","Lepidoptera","Cosmopterigidae","Cosmopteriginae","Cosmopterix","Cosmopterix scirpicola, Cosmopterix fernaldella","3","3","3","100","0","0","99.03","447","0","450","32.59","0"
"AGAW2","10S","INS","BOLD:AAN5267","Arthropoda","Insecta","Diptera","Cecidomyiidae","","","","3","3","3","100","0","0","99.85","449.33","0","450","30.59","0"
"AGAW2","10S","INS","BOLD:AAH4701","Arthropoda","Insecta","Lepidoptera","Pterophoridae","Pterophorinae","Sphenarches","Sphenarches ontario","2","2","2","114","0","0","99.22","447.5","0","450","30.89","0"
"AGAW2","10S","INS","BOLD:AAD0952","Arthropoda","Insecta","Lepidoptera","Pyralidae","Chrysauginae","Galasa","Galasa nigrinodis, Galasa nigripunctalis","2","2","2","114","0","0","98.9","391.5","0","406","31.83","0"
"AGAW2","10S","INS","BOLD:AAA3933","Arthropoda","Insecta","Lepidoptera","Erebidae","Herminiinae","Palthis","Palthis angulalis","2","2","2","114","0","0","98.22","410.5","0","413","31.96","0"
"AGAW2","10S","INS","BOLD:ACE4100","Arthropoda","Insecta","Orthoptera","Acrididae","Melanoplinae","Melanoplus","Melanoplus pinicola","2","2","2","114","0","0","97.39","401","0","402.5","34.85","0"
"AGAW2","10S","INS","BOLD:AAF3338","Arthropoda","Insecta","Lepidoptera","Noctuidae","Noctuinae","Striacosta","Striacosta albicosta","2","2","2","114","0","0","98.08","415.5","0","419","33.35","0"
"AGAW2","10S","INS","BOLD:AAA2518","Arthropoda","Insecta","Lepidoptera","Crambidae","Crambinae","Chrysoteuchia","Chrysoteuchia topiaria, Chrysoteuchia topiarius","2","2","2","114","0","0","98.88","395.5","0","395.5","35.19","0"
"AGAW2","10S","INS","BOLD:AAP8816","Arthropoda","Insecta","Diptera","Cecidomyiidae","Cecidomyiinae","Sitodiplosis","Sitodiplosis sp. BOLD:AAP8816","2","2","2","114","0","0","99.66","433.5","0","437","25.85","0"
"AGAW2","10S","INS","BOLD:AAY6412","Arthropoda","Insecta","Diptera","Sciaridae","","Bradysia","Bradysia urticae","2","2","2","114","0","0","99.01","402.5","0","406.5","31","0"
"AGAW2","10S","INS","BOLD:ACU7906","Arthropoda","Insecta","Diptera","Chironomidae","Orthocladiinae","","","2","2","2","114","0","0","98.92","373","0","381","37.27","0"
"AGAW2","10S","INS","BOLD:AAG8882","Arthropoda","Insecta","Hemiptera","Pentatomidae","Pentatominae","Thyanta","Thyanta pallidovirens, Thyanta custator, Thyanta accerra","2","2","2","114","0","0","99.44","450","0","450","36.22","0"
"AGAW2","10S","INS","BOLD:AAB5609","Arthropoda","Insecta","Diptera","Muscidae","Coenosiinae","Coenosia","Coenosia sp.BOLD:AAB5609, Coenosia tigrina","2","2","2","114","0","0","100","447","0","450","33.33","0"
"AGAW2","10S","INS","BOLD:AAA2482","Arthropoda","Insecta","Lepidoptera","Noctuidae","Noctuinae","Mythimna","Mythimna unipuncta","2","2","2","114","0","0","99.75","400","0","406","30.11","0"
"AGAW2","10S","INS","BOLD:ACK3142","Arthropoda","Insecta","Diptera","Cecidomyiidae","","","","2","2","2","114","0","0","98.62","434.5","0","438","27.84","0"
"AGAW2","10S","INS","BOLD:AAG6743","Arthropoda","Insecta","Diptera","Sarcophagidae","Sarcophaginae","Sarcophaga","Sarcophaga sp. 1 MG, Sarcophaga subvicina","2","2","2","114","0","0","99.78","450","0","450","35.33","0"
"AGAW2","10S","INS","BOLD:AAO7694","Arthropoda","Insecta","Diptera","Cecidomyiidae","","","","2","2","2","114","0","0","98.77","449","0","450","27","0"
"AGAW2","10S","INS","BOLD:ACE9865","Arthropoda","Insecta","Lepidoptera","Noctuidae","Noctuinae","Feltia","Feltia subgothica, Feltia herilis, Feltia tricosa","2","2","2","114","0","0","99.66","447.5","0","450","31","0"
"AGAW2","10S","INS","BOLD:AAN5161","Arthropoda","Insecta","Diptera","Ceratopogonidae","Dasyheleinae","Dasyhelea","","1","1","1","130","0","0","98.1","369","0","369","36.86","0"
"AGAW2","10S","INS","BOLD:ABY1165","Arthropoda","Insecta","Coleoptera","Anthicidae","","","","1","1","1","130","0","0","98.08","417","0","417","35.01","0"
"AGAW2","10S","INS","BOLD:AAA3836","Arthropoda","Insecta","Lepidoptera","Noctuidae","Plusiinae","Autographa","Autographa precationis","1","1","1","130","0","0","98.22","449","0","450","32.44","0"
"AGAW2","10S","INS","BOLD:ACE0699","Arthropoda","Insecta","Diptera","Chironomidae","","","","1","1","1","130","0","0","98.38","431","0","450","36.44","0"
"AGAW2","10S","INS","BOLD:AAA6560","Arthropoda","Insecta","Lepidoptera","Crambidae","Crambinae","Crambus","Crambus multilinellus, Crambus saltuellus","1","1","1","130","0","0","98.63","438","0","436","31.19","0"
"AGAW2","10S","INS","BOLD:AAV5796","Arthropoda","Insecta","Diptera","Cecidomyiidae","","","","1","1","1","130","0","0","99.78","448","0","450","29.33","0"
"AGAW2","10S","INS","BOLD:AAB4903","Arthropoda","Insecta","Lepidoptera","Glyphipterigidae","Acrolepiinae","Acrolepiopsis","Acrolepiopsis assectella","1","1","1","130","0","0","100","426","0","450","31.11","0"
"AGAW2","10S","INS","BOLD:AAH2306","Arthropoda","Insecta","Diptera","Asilidae","Asilinae","Efferia","Efferia aestuans","1","1","1","130","0","0","100","449","0","450","38.22","0"
"AGAW2","10S","INS","BOLD:AAH2307","Arthropoda","Insecta","Diptera","Asilidae","","","","1","1","1","130","0","0","98.14","431","0","450","45.56","0"
"AGAW2","10S","INS","BOLD:AAA4751","Arthropoda","Insecta","Diptera","Culicidae","Culicinae","Culex","Culex pipiens sl, Culex nr. pipiens, Culex pipiens, Culex quinquefasciatus, Culex sp.","1","1","1","130","0","0","99.78","447","0","450","32.89","0"
"AGAW2","10S","INS","BOLD:AAM6780","Arthropoda","Insecta","Diptera","Dolichopodidae","Sympycninae","","","1","1","1","130","0","0","98.43","445","0","450","31.78","0"
"AGAW2","10S","INS","BOLD:AAB2076","Arthropoda","Insecta","Lepidoptera","Tortricidae","Olethreutinae","Ancylis","Ancylis metamelana, Ancylis sp.","1","1","1","130","0","0","100","446","0","450","30.89","0"
"AGAW2","10S","INS","BOLD:ACR4394","Arthropoda","Insecta","Diptera","Anthomyiidae","Anthomyiinae","Delia","Delia florilega","1","1","1","130","0","0","99.78","450","0","450","33.33","0"
"AGAW2","10S","INS","BOLD:AAB0483","Arthropoda","Insecta","Lepidoptera","Tortricidae","Olethreutinae","Zeiraphera","Zeiraphera vancouverana","1","1","1","130","0","0","99.1","446","0","450","33.11","0"
"AGAW2","10S","INS","BOLD:AAC1042","Arthropoda","Insecta","Lepidoptera","Crambidae","Spilomelinae","Lygropia","Lygropia rivulalis","1","1","1","130","0","0","99.78","448","0","450","28.44","0"
"AGAW2","10S","INS","BOLD:AAF9949","Arthropoda","Insecta","Hemiptera","Miridae","Mirinae","Trigonotylus","Trigonotylus pulchellus, Trigonotylus americanus, Trigonotylus psammaecolor, Trigonotylus caelestialium, Trigonotylus viridis, Trigonotylus ruficornis","1","1","1","130","0","0","100","450","0","450","34.44","0"
"AGAW2","10S","INS","BOLD:AAM7398","Arthropoda","Insecta","Hymenoptera","Ichneumonidae","Campopleginae","Hyposoter","","1","1","1","130","0","0","98.86","350","0","354","24.29","0"
"AGAW2","10S","INS","BOLD:ADF2969","Arthropoda","Insecta","Diptera","Sarcophagidae","Sarcophaginae","Blaesoxipha","Blaesoxipha falciformis","1","1","1","130","0","0","99.15","353","0","358","37.43","0"
"AGAW2","10S","INS","BOLD:AAC4403","Arthropoda","Insecta","Diptera","Chironomidae","Chironominae","Micropsectra","Micropsectra polita","1","1","1","130","0","0","99.11","448","0","450","34.89","0"
"AGAW2","10S","INS","BOLD:AAA7636","Arthropoda","Insecta","Hymenoptera","Braconidae","Alysiinae","Dinotrema","Dinotrema jft02, Dinotrema jft01","1","1","1","130","0","0","97.78","450","0","450","25.11","0"
"AGAW2","10S","INS","BOLD:AAU6592","Arthropoda","Insecta","Diptera","Ceratopogonidae","Dasyheleinae","Dasyhelea","Dasyhelea arenivaga","1","1","1","130","0","0","98.33","419","0","422","33.65","0"
"AGAW2","10S","INS","BOLD:ACV2677","Arthropoda","Insecta","Diptera","Cecidomyiidae","","","","1","1","1","130","0","0","97.32","448","0","450","30","0"
"AGAW2","10S","INS","BOLD:ABX5883","Arthropoda","Insecta","Lepidoptera","Tortricidae","Tortricinae","Choristoneura","Choristoneura retiniana, Choristoneura lambertiana, Choristoneura biennis, Choristoneura orae, Choristoneura fumiferana, Choristoneura occidentalis, Choristoneura freemani, Choristoneura pinus, Choristoneura occidentalis group, Choristoneura sp., Choristoneura carnana","1","1","1","130","0","0","99.55","449","0","450","31.33","0"
"AGAW2","10S","INS","BOLD:AAA7631","Arthropoda","Insecta","Hemiptera","Miridae","Mirinae","Adelphocoris","Adelphocoris lineolatus","1","1","1","130","0","0","99.11","449","0","450","35.33","0"
"AGAW2","10S","INS","BOLD:AAC9132","Arthropoda","Insecta","Diptera","Culicidae","Culicinae","Culiseta","Culiseta inornata","1","1","1","130","0","0","98.86","351","0","353","32.29","0"
"AGAW2","10S","INS","BOLD:AAM9105","Arthropoda","Insecta","Diptera","Muscidae","Coenosiinae","Coenosia","","1","1","1","130","0","0","99.15","352","0","353","37.96","0"
"AGAW2","10S","INS","BOLD:ACE7482","Arthropoda","Insecta","Hemiptera","Miridae","","","","1","1","1","130","0","0","97.65","426","0","428","35.75","0"
"AGAW2","10S","INS","BOLD:AAM9347","Arthropoda","Insecta","Diptera","Phoridae","Metopininae","Megaselia","","1","1","1","130","0","0","98.1","368","0","366","33.61","0"
"AGAW2","10S","INS","BOLD:AAV1136","Arthropoda","Insecta","Diptera","Scatopsidae","","","","1","1","1","130","0","0","97.77","449","0","450","36","0"
"AGAW2","10S","INS","BOLD:AAH0237","Arthropoda","Insecta","Coleoptera","Cleridae","Clerinae","Enoclerus","Enoclerus nigripes, Enoclerus rosmarus","1","1","1","130","0","0","97.99","448","0","450","32.22","0"
"AGAW2","10S","INS","BOLD:AAZ5483","Arthropoda","Insecta","Diptera","Muscidae","Phaoniinae","Helina","Helina setiventris","1","1","1","130","0","0","99.33","448","0","450","29.11","0"
"AGAW2","10S","INS","BOLD:ABA0822","Arthropoda","Insecta","Diptera","Cecidomyiidae","","","","1","1","1","130","0","0","98.41","440","0","441","26.3","0"
"AGAW2","10S","INS","BOLD:AAB0373","Arthropoda","Insecta","Neuroptera","Chrysopidae","Chrysopinae","Chrysoperla","Chrysoperla sp. isolate 82, Chrysoperla zastrowi, Chrysoperla plorabunda, Chrysoperla sp. isolate 71, Chrysoperla agilis, Italochrysa italica, Chrysoperla sp. isolate 88, Chrysoperla lucasina, Chrysoperla carnea, Chrysoperla mediterranea, Chrysoperla pallida, Chrysoperla nipponensis, Chrysoperla sillemi","1","1","1","130","0","0","100","353","0","355","31.27","0"
"AGAW2","10S","INS","BOLD:AAA2632","Arthropoda","Insecta","Lepidoptera","Noctuidae","Noctuinae","Noctua","Noctua atlantica, Noctua pronuba","1","1","1","130","0","0","99.56","451","0","450","31.56","0"
"AGAW2","10S","INS","BOLD:AAN5255","Arthropoda","Insecta","Diptera","Cecidomyiidae","Cecidomyiinae","Dasineura","","1","1","1","130","0","0","98.22","450","0","450","26.67","0"
"AGAW2","10S","INS","BOLD:ACV5891","Arthropoda","Insecta","Diptera","Cecidomyiidae","","","","1","1","1","130","0","0","100","361","0","450","25.56","0"
"AGAW2","10S","INS","BOLD:AAA7039","Arthropoda","Insecta","Lepidoptera","Tortricidae","Tortricinae","Sparganothis","Sparganothis bistriata, Sparganothis robinsonana, Sparganothis sulfureana, Sparganothis lycopodiana, Sparganothis tristriata","1","1","1","130","0","0","97.74","443","0","450","31.56","0"
"AGAW2","10S","INS","BOLD:ABX5095","Arthropoda","Insecta","Hemiptera","Miridae","Mirinae","Neolygus","Neolygus tiliae","1","1","1","130","0","0","99.55","449","0","450","33.56","0"
"AGAW2","10S","INS","BOLD:AAA5206","Arthropoda","Insecta","Lepidoptera","Erebidae","Herminiinae","Zanclognatha","Zanclognatha cruralis, Zanclognatha sp. 1, Zanclognatha jacchusalis, Zanclognatha minoralis, Zanclognatha minoralis sp. 1, Zanclognatha sp., Zanclognatha minoralis sp. 3, Zanclognatha gypsalis, Zanclognatha dentata, Zanclognatha theralis, Zanclognatha protumnusalis, Zanclognatha martha, Zanclognatha atrilineella, Zanclognatha obscuripennis, Zanclognatha minoralis sp. 2, Zanclognatha marcidilinea","1","1","1","130","0","0","97.44","390","0","394","30.96","0"
"AGAW2","10S","INS","BOLD:AAJ7105","Arthropoda","Insecta","Diptera","Agromyzidae","Agromyzinae","Agromyza","Agromyza frontella","1","1","1","130","0","0","99","401","0","450","36.44","0"
"AGAW2","10S","INS","BOLD:AAF9393","Arthropoda","Insecta","Coleoptera","Meloidae","Meloinae","Epicauta","Epicauta fabricii","1","1","1","130","0","0","99.5","404","0","408","40.2","0"
"AGAW2","10S","INS","BOLD:ACE8875","Arthropoda","Insecta","Orthoptera","Acrididae","Melanoplinae","Melanoplus","Melanoplus bowditchi","1","1","1","130","0","0","97.1","449","0","450","36","0"
"AGAW2","10S","INS","BOLD:AAL2750","Arthropoda","Insecta","Diptera","Muscidae","Phaoniinae","Phaonia","Phaonia tuguriorum","1","1","1","130","0","0","98","449","0","450","33.11","0"
"AGAW2","10S","INS","BOLD:ACB0412","Arthropoda","Insecta","Diptera","Agromyzidae","Phytomyzinae","Liriomyza","","1","1","1","130","0","0","98.22","449","0","450","34.44","0"
"AGAW2","10S","INS","BOLD:AAF1051","Arthropoda","Insecta","Diptera","Agromyzidae","Phytomyzinae","Cerodontha","Cerodontha muscina, Cerodontha calamagrostis","1","1","1","130","0","0","99.78","446","0","450","36.44","0"
"AGAW2","10S","INS","BOLD:AAH7161","Arthropoda","Insecta","Diptera","Sarcophagidae","Sarcophaginae","Ravinia","Ravinia derelicta","1","1","1","130","0","0","97.57","412","0","417","32.61","0"
"AGAW2","10S","INS","BOLD:AAV6112","Arthropoda","Insecta","Diptera","Chloropidae","Oscinellinae","Oscinella","Oscinella sp. 2 AGT, Oscinella sp. TAW6","1","1","1","130","0","0","100","412","0","417","31.65","0"
"AGAW2","10S","INS","BOLD:AAA3867","Arthropoda","Insecta","Lepidoptera","Noctuidae","Noctuinae","Apamea","Apamea amputatrix","1","1","1","130","0","0","98.22","449","0","450","33.78","0"
"AGAW2","10S","INS","BOLD:AAB4321","Arthropoda","Insecta","Lepidoptera","Gelechiidae","Gelechiinae","Metzneria","Metzneria lappella","1","1","1","130","0","0","99.29","425","0","450","32","0"
"AGAW2","10S","INS","BOLD:ACE4897","Arthropoda","Insecta","Diptera","Tachinidae","Tachininae","Microphthalma","","1","1","1","130","0","0","99.48","381","0","383","28.46","0"
"AGAW2","10S","INS","BOLD:AAB0651","Arthropoda","Insecta","Lepidoptera","Gelechiidae","Dichomeridinae","Dichomeris","Dichomeris leuconotella, Dichomeris gnoma","1","1","1","130","0","0","99.47","379","0","380","32.37","0"
"AGAW2","10S","INS","BOLD:AAG2355","Arthropoda","Insecta","Diptera","Tachinidae","Tachininae","Strongygaster","Strongygaster triangulifera, Strongygaster sp.","1","1","1","130","0","0","98.67","377","0","383","30.29","0"
"AGAW2","10S","INS","BOLD:AAP9022","Arthropoda","Insecta","Diptera","Cecidomyiidae","","","","1","1","1","130","0","0","100","449","0","450","27.56","0"
"AGAW2","10S","INS","BOLD:AAE0845","Arthropoda","Insecta","Hemiptera","Pentatomidae","Pentatominae","Euschistus","Euschistus servus","1","1","1","130","0","0","98.44","450","0","450","37.78","0"
"AGAW2","10S","INS","BOLD:ABA5934","Arthropoda","Insecta","Hymenoptera","Ichneumonidae","Cryptinae","","","1","1","1","130","0","0","98.44","450","0","450","30.22","0"
"AGAW2","10S","INS","BOLD:AAM6230","Arthropoda","Insecta","Diptera","Chironomidae","Chironominae","Chironomus","","1","1","1","130","0","0","97.98","445","0","450","33.78","0"
"AGAW2","10S","INS","BOLD:AAA3797","Arthropoda","Insecta","Hymenoptera","Megachilidae","Megachilinae","Megachile","Megachile sp. aff rotundata, Megachile cf. rotundata, Megachile rotundata","1","1","1","130","0","0","98.96","384","0","441","25.4","0"
"AGAW2","10S","INS","BOLD:AAC9062","Arthropoda","Insecta","Diptera","Culicidae","Culicinae","Ochlerotatus, Aedes","Aedes fitchii, Ochlerotatus fitchii, Aedes grossbecki, Aedes ventrovittis","1","1","1","130","0","0","98.89","450","0","450","32.67","0"
"AGAW2","10S","INS","BOLD:AAG6475","Arthropoda","Insecta","Diptera","Ceratopogonidae","Dasyheleinae","Dasyhelea","","1","1","1","130","0","0","99.01","406","0","409","29.58","0"
"AGAW2","10S","INS","BOLD:AAA7368","Arthropoda","Insecta","Lepidoptera","Cosmopterigidae","Cosmopteriginae","Limnaecia","Limnaecia phragmitella","1","1","1","130","0","0","100","405","0","412","30.1","0"
"AGAW2","10S","INS","BOLD:ADF3937","","","","","","","","1","1","1","130","0","0","97.14","384","0","386","31.35","0"
"AGAW2","10S","INS","BOLD:ACB0984","Arthropoda","Insecta","Psocodea","Stenopsocidae","","Graphopsocus","Graphopsocus cruciatus","1","1","1","130","0","0","99.76","413","0","450","37.56","0"
"AGAW2","10S","INS","BOLD:ABV1422","Arthropoda","Insecta","Diptera","Cecidomyiidae","","","","1","1","1","130","0","0","97.86","421","0","426","26.76","0"
"AGAW2","10S","INS","BOLD:AAY6469","Arthropoda","Insecta","Diptera","Cecidomyiidae","","","","1","1","1","130","0","0","99.78","449","0","450","26.22","0"
"AGAW2","10S","INS","BOLD:AAG2460","Arthropoda","Insecta","Diptera","Anthomyiidae","Anthomyiinae","Hydrophoria","Hydrophoria lancifer","1","1","1","130","0","0","98.44","449","0","450","34","0"
"AGAW2","10S","INS","BOLD:AAA5803","Arthropoda","Insecta","Hemiptera","Miridae","Mirinae","Lygus","Lygus lineolaris","1","1","1","130","0","0","98.88","447","0","450","34.22","0"
"AGAW2","10S","NAR","BOLD:AAI9663","Mollusca","Gastropoda","Stylommatophora","Agriolimacidae",NA,"Deroceras","Deroceras reticulatum","2","2","2","1","0","0","99.14","408","0","450","35.78","0"
"AGAW2","10S","NIN","BOLD:AAH6630","Arthropoda","Collembola","Entomobryomorpha","Entomobryidae","Entomobryinae","Entomobrya","Entomobrya unostrigata","1964","1657","1657","1","15.63","0","99.23","435.51","0","436.76","39.15","0"
"AGAW2","10S","NIN","BOLD:ABW2639","Arthropoda","Arachnida","Trombidiformes","Anystidae","","","","246","218","218","2","11.38","0","99.51","446.3","0","445.94","31.47","0"
"AGAW2","10S","NIN","BOLD:AAI4346","Arthropoda","Arachnida","Opiliones","Phalangiidae","","Paroligolophus, Phalangium","Paroligolophus agrestis, Phalangium opilio","134","129","129","3","3.73","0","99.32","436.01","0","437.62","36.3","0"
"AGAW2","10S","NIN","BOLD:AAA6381","Arthropoda","Arachnida","Araneae","Tetragnathidae","","Tetragnatha","Tetragnatha laboriosa","107","107","107","4","0","0","99.08","426.14","0","428.5","39.36","0"
"AGAW2","10S","NIN","BOLD:AAU6252","Arthropoda","Arachnida","Trombidiformes","Bdellidae","","Cyta","","71","64","64","5","9.86","0","99.28","428.68","0","429.9","31.46","0"
"AGAW2","10S","NIN","BOLD:ADG1586","Arthropoda","Collembola","Entomobryomorpha","Entomobryidae","","","","60","59","59","6","1.67","0","99.52","397.62","0","435.82","40.04","0"
"AGAW2","10S","NIN","BOLD:AAA8820","Arthropoda","Collembola","Entomobryomorpha","Entomobryidae","Lepidocyrtinae","Lepidocyrtus","Lepidocyrtus cyaneus","47","46","46","7","2.13","0","98.97","400.34","0","403.45","43.29","0"
"AGAW2","10S","NIN","BOLD:ACZ1295","Arthropoda","Arachnida","Trombidiformes","Anystidae","","","","31","31","31","8","0","0","99.39","373.58","0","382.81","31.3","0"
"AGAW2","10S","NIN","BOLD:AAA9476","Arthropoda","Arachnida","Araneae","Linyphiidae","","Grammonota","Grammonota gentilis","20","20","20","9","0","0","98.84","426.85","0","432.7","30.78","0"
"AGAW2","10S","NIN","BOLD:AAB7915","Arthropoda","Collembola","Symphypleona","Bourletiellidae","","Deuterosminthurus","Deuterosminthurus sp1","8","8","8","10","0","0","98.52","390.38","0","392.38","34.24","0"
"AGAW2","10S","NIN","BOLD:AAM7961","Arthropoda","Arachnida","Trombidiformes","Anystidae","","","","7","7","7","11","0","0","99.94","447.71","0","450","31.11","0"
"AGAW2","10S","NIN","BOLD:ACJ7454","Arthropoda","Collembola","Symphypleona","Bourletiellidae","","","","2","2","2","12","0","0","99.63","402","0","440","38.98","0"
"AGAW2","10S","NIN","BOLD:ACL9406","Arthropoda","Arachnida","Trombidiformes","Cunaxidae","","","","2","2","2","12","0","0","99.76","415.5","0","437","25.16","0"
"AGAW2","10S","NIN","BOLD:ACX8969","Arthropoda","Arachnida","Trombidiformes","Eupodidae","","","","2","2","2","12","0","0","99.18","423.5","0","425","27.75","0"
"AGAW2","10S","NIN","BOLD:AAF9285","Arthropoda","Arachnida","Mesostigmata","Parasitidae","","","","1","1","1","15","0","0","97.99","448","0","450","34.67","0"
"AGAW2","10S","NIN","BOLD:AAD1564","Arthropoda","Arachnida","Araneae","Clubionidae","","Clubiona","Clubiona abboti","1","1","1","15","0","0","98.56","417","0","420","30.95","0"
"AGAW2","10S","NIN","BOLD:AAB8452","Arthropoda","Collembola","Entomobryomorpha","Entomobryidae","Lepidocyrtinae","Lepidocyrtus","Lepidocyrtus paradoxus","1","1","1","15","0","0","100","449","0","450","42.22","0"
"AGAW2","10S","NIN","BOLD:AAB7914","Arthropoda","Collembola","Symphypleona","Bourletiellidae","","Bourletiella","","1","1","1","15","0","0","100","447","0","450","37.78","0"
"AGAW2","10S","NIN","BOLD:ACY2550","Arthropoda","Arachnida","Trombidiformes","Bdellidae","","Cyta","","1","1","1","15","0","0","99.2","376","0","378","30.95","0"
"AGAW2","10W","BAC","TAX:91391","Proteobacteria","Alphaproteobacteria","Rickettsiales","",NA,"","","41","32","32","1","21.95","0","99.01","405.71","0","446.8","39.78","0"
"AGAW2","10W","BAC","TAX:175768","Proteobacteria","Alphaproteobacteria","Rickettsiales","Rickettsiaceae",NA,"Wolbachia","Wolbachia pipientis","6","6","6","2","0","0","98.14","448.17","0","449.17","39.85","0"
"AGAW2","10W","BAC","TAX:91393","Proteobacteria","Alphaproteobacteria","Rickettsiales","Rickettsiaceae",NA,"Wolbachia","","1","1","1","3","0","0","99.23","392","0","427","39.11","0"
"AGAW2","10W","CHO","no match","no match","no match","no match","no match","no match","no match","no match","no match","no match","no match","no match","no match","no match","no match","no match","no match","no match","no match","no match"
"AGAW2","10W","INS","BOLD:AAA3453","Arthropoda","Insecta","Diptera","Anthomyiidae","Anthomyiinae","Delia","","26417","22362","22362","1","15.35","0","99.39","427.65","0","430.38","34.59","0"
"AGAW2","10W","INS","BOLD:AAA4555","Arthropoda","Insecta","Orthoptera","Acrididae","Melanoplinae","Melanoplus","Melanoplus stupefactus, Melanoplus angustipennis, Melanoplus ponderosus, Melanoplus bruneri, Melanoplus frigidus, Melanoplus scapularis, Melanoplus cinereus, Melanoplus madeleineae, Melanoplus sanguinipes, Melanoplus nigrescens group, Melanoplus tunicae, Melanoplus walshi, Melanoplus femurrubrum, Melanoplus dawsoni, Melanoplus sp., Melanoplus fasciatus, Melanoplus bowditchi, Melanoplus stonei, Melanoplus bivittatus, Melanoplus huroni, Melanoplus thomasi, Melanoplus borealis, Melanoplus packardii","9433","6911","6911","2","26.74","0","99.42","447.11","0","448.21","36.23","0"
"AGAW2","10W","INS","BOLD:ACE7444","Arthropoda","Insecta","Hemiptera","Miridae","Mirinae, Phylinae","Adelphocoris, Megalocoleus","Adelphocoris lineolatus, Megalocoleus molliculus","7644","4785","4785","3","37.4","0","99.7","439.55","0","440.31","34.78","0"
"AGAW2","10W","INS","BOLD:AAN6154","Arthropoda","Insecta","Coleoptera","Latridiidae","Corticariinae","Melanophthalma","Melanophthalma inermis","5672","4638","4638","4","18.23","0","98.87","434.87","0","436.3","31.22","0"
"AGAW2","10W","INS","BOLD:ABY2946","Arthropoda","Insecta","Diptera","Tabanidae","Tabaninae","Hybomitra","Hybomitra trepida","5419","3638","3638","5","32.87","0","98.6","437.33","0","438.82","33.19","0"
"AGAW2","10W","INS","BOLD:AAA8933","Arthropoda","Insecta","Coleoptera","Coccinellidae","Coccinellinae","Coccinella","Coccinella septempunctata","3386","2620","2620","6","22.62","0","99.51","443.77","0","445.66","32.93","0"
"AGAW2","10W","INS","BOLD:AAG2868","Arthropoda","Insecta","Hemiptera","Cicadellidae","Typhlocybinae","Empoasca","Empoasca sp., Empoasca fabae","2595","1658","1658","7","36.11","0","99.56","445.09","0","447.39","32.29","0"
"AGAW2","10W","INS","BOLD:AAC0903","Arthropoda","Insecta","Diptera","Chironomidae","Chironominae","Chironomus","Chironomus acidophilus","2360","1869","1869","8","20.81","0","99.49","436.69","0","438.96","36.29","0"
"AGAW2","10W","INS","BOLD:AAU6681","Arthropoda","Insecta","Diptera","Tipulidae","Tipulinae","Nephrotoma","Nephrotoma cornicina","2172","1668","1668","9","23.2","0","99.62","442.43","0","443.66","36.28","0"
"AGAW2","10W","INS","BOLD:AAA3181","Arthropoda","Insecta","Diptera","Muscidae","Muscinae","Stomoxys","Stomoxys calcitrans","1681","1431","1431","10","14.87","0","98.74","429.88","0","433.58","32.94","0"
"AGAW2","10W","INS","BOLD:AAH4135","Arthropoda","Insecta","Diptera","Chloropidae","Chloropinae","Thaumatomyia","Thaumatomyia glabra, Thaumatomyia grata, Thaumatomyia sp. 1 TAW","1476","1341","1341","11","9.15","0","99.5","431.07","0","435.14","31.88","0"
"AGAW2","10W","INS","BOLD:AAG2511","Arthropoda","Insecta","Diptera","Anthomyiidae","Anthomyiinae","Delia","Delia platura","1261","1166","1166","12","7.53","0","98.92","443.14","0","446.13","33.16","0"
"AGAW2","10W","INS","BOLD:AAH3661","Arthropoda","Insecta","Diptera","Cecidomyiidae","","","","1002","889","889","13","11.28","0","99.26","429.61","0","431.61","25.33","0"
"AGAW2","10W","INS","BOLD:ACF1257","","","","","","","","875","564","564","14","35.54","0","99.7","445.39","0","446.02","34.62","0"
"AGAW2","10W","INS","BOLD:ABA0863","Arthropoda","Insecta","Diptera","Cecidomyiidae","","","","811","666","666","15","17.88","0","99.34","438.26","0","439.61","29.78","0"
"AGAW2","10W","INS","BOLD:AAD1169","Arthropoda","Insecta","Lepidoptera","Cosmopterigidae","Cosmopteriginae","Cosmopterix","Cosmopterix scirpicola, Cosmopterix fernaldella","803","680","680","16","15.32","0","99.4","438.94","0","440.82","32.67","0"
"AGAW2","10W","INS","BOLD:AAJ4275","Arthropoda","Insecta","Diptera","Chironomidae","Chironominae","Chironomus","Chironomus harpi","794","710","710","17","10.58","0","98.26","415.71","0","431.57","38.23","0"
"AGAW2","10W","INS","BOLD:AAF9771","Arthropoda","Insecta","Diptera","Hybotidae","Tachydromiinae","Platypalpus","Platypalpus longiseta, Platypalpus trivialis, Platypalpus sp. 5, Platypalpus sp. 17, Platypalpus pallidiventris","733","514","514","18","29.88","0","98.96","445.59","0","446.21","34.99","0"
"AGAW2","10W","INS","BOLD:AAG2123","Arthropoda","Insecta","Diptera","Tachinidae","Dexiinae","Dinera","Dinera grisescens","679","539","539","19","20.62","0","99.38","430.32","0","432.18","31.8","0"
"AGAW2","10W","INS","BOLD:AAK5928","Arthropoda","Insecta","Coleoptera","Chrysomelidae","Criocerinae","Oulema","Oulema melanopus","674","543","543","20","19.44","0","99.4","441.17","0","442.35","36.25","0"
"AGAW2","10W","INS","BOLD:AAK8751","Arthropoda","Insecta","Diptera","Agromyzidae","Phytomyzinae","Liriomyza","Liriomyza sp. SB, Liriomyza trifoliearum","665","537","537","21","19.25","0","99.47","437.09","0","438.22","33.12","0"
"AGAW2","10W","INS","BOLD:AAH0134","Arthropoda","Insecta","Coleoptera","Phalacridae","Phalacrinae","Stilbus","Stilbus apicialis","637","552","552","22","13.34","0","99.39","422.83","0","424.07","31.88","0"
"AGAW2","10W","INS","BOLD:AAG9668","Arthropoda","Insecta","Diptera","Dolichopodidae","Diaphorinae","Chrysotus","Chrysotus sp., Chrysotus sp. nr. obliquus","635","582","582","23","8.35","0","98.88","444.1","0","446.78","32.79","0"
"AGAW2","10W","INS","BOLD:AAG5453","Arthropoda","Insecta","Diptera","Chironomidae","Chironominae","Chironomus","Chironomus bifurcatus","604","493","493","24","18.38","0","98.34","442.31","0","445.71","34.95","0"
"AGAW2","10W","INS","BOLD:AAA5803","Arthropoda","Insecta","Hemiptera","Miridae","Mirinae","Lygus","Lygus lineolaris","544","386","386","25","29.04","0","99.32","446.17","0","447.85","34.09","0"
"AGAW2","10W","INS","BOLD:AAG1711","Arthropoda","Insecta","Diptera","Muscidae","Phaoniinae","Helina","Helina depuncta","491","434","434","26","11.61","0","99.37","432.92","0","435.27","31.33","0"
"AGAW2","10W","INS","BOLD:ABY5318","Arthropoda","Insecta","Diptera","Chloropidae","Oscinellinae","Incertella, Rhopalopterum","Rhopalopterum umbrosum, Rhopalopterum luteiceps, Incertella bispina, Rhopalopterum painteri","412","330","330","27","19.9","0","98.95","444.42","0","445.58","31.99","0"
"AGAW2","10W","INS","BOLD:ABW7897","Arthropoda","Insecta","Diptera","Cecidomyiidae","","","","397","306","306","28","22.92","0","99.56","442.11","0","443.89","29.62","0"
"AGAW2","10W","INS","BOLD:AAN5242","Arthropoda","Insecta","Diptera","Cecidomyiidae","","","","394","392","392","29","0.51","0","98.75","402.84","0","405.13","28.15","0"
"AGAW2","10W","INS","BOLD:AAV5575","Arthropoda","Insecta","Diptera","Cecidomyiidae","","","CecidInt30 sp. BOLD:AAV5575","367","272","272","30","25.89","0","99.54","442.49","0","443.37","25.91","0"
"AGAW2","10W","INS","BOLD:AAG4781","Arthropoda","Insecta","Diptera","Agromyzidae","","","","367","279","279","30","23.98","0","99.03","443.38","0","445.16","32.45","0"
"AGAW2","10W","INS","BOLD:AAH3664","Arthropoda","Insecta","Diptera","Cecidomyiidae","","","CecidInt3 sp. BOLD:AAH3664","341","327","327","32","4.11","0","99.19","427.35","0","430.41","26.88","0"
"AGAW2","10W","INS","BOLD:ACU7906","Arthropoda","Insecta","Diptera","Chironomidae","Orthocladiinae","","","318","316","316","33","0.63","0","98.97","399.58","0","409.12","36.35","0"
"AGAW2","10W","INS","BOLD:AAV5594","Arthropoda","Insecta","Diptera","Cecidomyiidae","","","","313","266","266","34","15.02","0","99.37","438.03","0","439.69","26.37","0"
"AGAW2","10W","INS","BOLD:ACE0699","Arthropoda","Insecta","Diptera","Chironomidae","","","","290","218","218","35","24.83","0","98.39","426.04","0","442.92","36.37","0"
"AGAW2","10W","INS","TAX:487439","Arthropoda","Insecta","Coleoptera","Chrysomelidae","Eumolpinae","Paria","Paria thoracica","272","259","259","36","4.78","0","99.41","401.97","0","403.69","38.81","0"
"AGAW2","10W","INS","BOLD:AAH0135","Arthropoda","Insecta","Coleoptera","Phalacridae","Phalacrinae","Acylomus","Acylomus pugetanus, Acylomus ergoti","217","205","205","37","5.53","0","99.08","431.93","0","433.92","31.48","0"
"AGAW2","10W","INS","BOLD:AAA7661","Arthropoda","Insecta","Diptera","Culicidae","Culicinae","Culiseta, Culex","Culiseta melanura, Culex articularis, Culex restuans, Culex acharistus","208","198","198","38","4.81","0","99.4","430.92","0","432.96","32.79","0"
"AGAW2","10W","INS","BOLD:AAN5430","Arthropoda","Insecta","Diptera","Agromyzidae","","","","205","189","189","39","7.8","0","98.76","415.56","0","434.92","32.05","0"
"AGAW2","10W","INS","BOLD:ADF6661","Arthropoda","Insecta","Diptera","Cecidomyiidae","","","","156","139","139","40","10.9","0","99.21","415.3","0","441.47","29.7","0"
"AGAW2","10W","INS","BOLD:AAD5253","Arthropoda","Insecta","Diptera","Chironomidae","Orthocladiinae","Thienemanniella","Thienemanniella xena, Thienemanniella vittata","149","128","128","41","14.09","0","98.45","437.7","0","440.33","34.44","0"
"AGAW2","10W","INS","BOLD:ACU8099","Arthropoda","Insecta","Diptera","Chironomidae","","","","127","125","125","42","1.57","0","98.95","428.24","0","430.39","33.44","0"
"AGAW2","10W","INS","BOLD:ABV1297","Arthropoda","Insecta","Diptera","Sciaridae","","","","126","124","124","43","1.59","0","99.04","428.4","0","430.88","30.21","0"
"AGAW2","10W","INS","BOLD:AAM9255","Arthropoda","Insecta","Diptera","Sciaridae","","Pseudolycoriella","","107","107","107","44","0","0","99.19","410.74","0","413.1","30.76","0"
"AGAW2","10W","INS","BOLD:ABX3618","Arthropoda","Insecta","Coleoptera","Staphylinidae","Paederinae","Lobrathium","","101","71","71","45","29.7","0","99.49","436.18","0","439.12","31.11","0"
"AGAW2","10W","INS","BOLD:AAP5169","Arthropoda","Insecta","Diptera","Chloropidae","Oscinellinae","Elachiptera","Elachiptera nigriceps","92","81","81","46","11.96","0","99.28","428.74","0","429.91","31.6","0"
"AGAW2","10W","INS","BOLD:AAK4963","Arthropoda","Insecta","Diptera","Chironomidae","Chironominae","Paracladopelma","Paracladopelma winnelli","84","79","79","47","5.95","0","99.12","433.99","0","435.8","34.65","0"
"AGAW2","10W","INS","BOLD:ABV1455","Arthropoda","Insecta","Diptera","Chironomidae","Chironominae","","","82","79","79","48","3.66","0","98.13","435.11","0","437.06","31.6","0"
"AGAW2","10W","INS","BOLD:AAN5199","Arthropoda","Insecta","Diptera","Cecidomyiidae","","","CecidInt20 sp. BOLD:AAN5199","77","65","65","49","15.58","0","99.5","439.18","0","448.35","26.7","0"
"AGAW2","10W","INS","BOLD:ACK3431","Arthropoda","Insecta","Diptera","Cecidomyiidae","","","","75","72","72","50","4","0","99.22","428.16","0","430.77","32.27","0"
"AGAW2","10W","INS","BOLD:AAI1549","Arthropoda","Insecta","Hymenoptera","Braconidae","Euphorinae","Meteorus","","75","63","63","50","16","0","98.92","441.8","0","442.99","31.17","0"
"AGAW2","10W","INS","BOLD:AAN5260","Arthropoda","Insecta","Diptera","Cecidomyiidae","","","","73","72","72","52","1.37","0","99.04","402.63","0","404.47","25.92","0"
"AGAW2","10W","INS","BOLD:AAN5501","Arthropoda","Insecta","Diptera","Hybotidae","","","","72","71","71","53","1.39","0","99.13","410.49","0","412.49","34.06","0"
"AGAW2","10W","INS","BOLD:AAG6745","Arthropoda","Insecta","Diptera","Calliphoridae","Calliphorinae","Pollenia","Pollenia pediculata","70","70","70","54","0","0","99.36","418.77","0","420.8","33.51","0"
"AGAW2","10W","INS","BOLD:AAF9789","Arthropoda","Insecta","Diptera","Hybotidae","","","","70","70","70","54","0","0","99.33","422","0","423.13","32.94","0"
"AGAW2","10W","INS","BOLD:AAP8739","Arthropoda","Insecta","Diptera","Platystomatidae","Platystomatinae","Rivellia","Rivellia steyskali","69","62","62","56","10.14","0","99.21","439.59","0","441","38","0"
"AGAW2","10W","INS","BOLD:AAG8091","Arthropoda","Insecta","Hymenoptera","Ichneumonidae","Campopleginae","Campoletis","Campoletis sp. AB.2","62","62","62","57","0","0","99.11","431.24","0","433.44","24.62","0"
"AGAW2","10W","INS","BOLD:ADG1318","","","","","","","","53","49","49","58","7.55","0","99.46","438.7","0","441","25.13","0"
"AGAW2","10W","INS","BOLD:AAG7891","Arthropoda","Insecta","Hymenoptera","Platygastridae","Scelioninae","Telenomus","Telenomus podisi","52","48","48","59","7.69","0","98.84","431.42","0","439.62","26.79","0"
"AGAW2","10W","INS","BOLD:AAN5230","Arthropoda","Insecta","Diptera","Cecidomyiidae","","","","50","49","49","60","2","0","99.3","426.3","0","427.84","25.99","0"
"AGAW2","10W","INS","BOLD:AAM9013","Arthropoda","Insecta","Diptera","Keroplatidae","Macrocerinae","Macrocera","Macrocera pusilla","49","48","48","61","2.04","0","99.24","424.86","0","426.84","29.99","0"
"AGAW2","10W","INS","BOLD:ABW1379","Arthropoda","Insecta","Diptera","Chloropidae","Oscinellinae","Malloewia","Malloewia abdominalis","49","42","42","61","14.29","0","98.48","440.37","0","445.14","31.49","0"
"AGAW2","10W","INS","BOLD:AAA4222","Arthropoda","Insecta","Lepidoptera","Erebidae","Hypeninae","Hypena","Hypena scabra","48","48","48","63","0","0","98.93","395.33","0","397.54","32.38","0"
"AGAW2","10W","INS","BOLD:AAV5688","Arthropoda","Insecta","Diptera","Cecidomyiidae","","","","47","47","47","64","0","0","99.47","419.64","0","420.85","34.16","0"
"AGAW2","10W","INS","BOLD:AAD7604","Arthropoda","Insecta","Coleoptera","Coccinellidae","Coccinellinae","Coleomegilla","Coleomegilla maculata","46","42","42","65","8.7","0","98.95","435.83","0","438.7","31.75","0"
"AGAW2","10W","INS","BOLD:ABA0858","Arthropoda","Insecta","Diptera","Cecidomyiidae","","","","44","43","43","66","2.27","0","99.24","425.52","0","426.43","25.99","0"
"AGAW2","10W","INS","BOLD:AAM6037","Arthropoda","Insecta","Diptera","Cecidomyiidae","Cecidomyiinae","Contarinia, Dasineura","","44","41","41","66","6.82","0","98.98","439.32","0","441.39","26.51","0"
"AGAW2","10W","INS","BOLD:AAF0738","Arthropoda","Insecta","Trichoptera","Hydroptilidae","Orthotrichiinae","Orthotrichia","Orthotrichia aegerfasciella","42","37","37","68","11.9","0","98.69","445.12","0","446.57","31.07","0"
"AGAW2","10W","INS","BOLD:ACC8079","Arthropoda","Insecta","Diptera","Cecidomyiidae","","","","41","38","38","69","7.32","0","99.37","441.46","0","443.24","29","0"
"AGAW2","10W","INS","BOLD:AAN5205","Arthropoda","Insecta","Diptera","Cecidomyiidae","","","CecidInt42 sp. BOLD:AAN5205","40","40","40","70","0","0","98.71","429.33","0","430.88","31.18","0"
"AGAW2","10W","INS","BOLD:ACX6055","Arthropoda","Insecta","Diptera","Phoridae","Metopininae","Megaselia","Megaselia sp. AK-2010-2, Megaselia sp. BOLD:ACX6055","40","38","38","70","5","0","99.42","431.08","0","432.48","32.97","0"
"AGAW2","10W","INS","BOLD:AAN5183","Arthropoda","Insecta","Diptera","Cecidomyiidae","","","","36","36","36","72","0","0","98.25","436.06","0","437.33","27.72","0"
"AGAW2","10W","INS","BOLD:ACH2915","Arthropoda","Insecta","Diptera","Cecidomyiidae","","","","35","35","35","73","0","0","98","383.23","0","418.69","32.48","0"
"AGAW2","10W","INS","BOLD:ABU5525","Arthropoda","Insecta","Diptera","Chironomidae","Orthocladiinae","Limnophyes","Limnophyes sp. 14ES","34","34","34","74","0","0","99.04","389.74","0","393.74","37.42","0"
"AGAW2","10W","INS","BOLD:AAD1485","Arthropoda","Insecta","Diptera","Chironomidae","Chironominae","Paratanytarsus","Paratanytarsus grimmii, Paratanytarsus sp.","33","31","31","75","6.06","0","99.64","442.61","0","444.15","33.75","0"
"AGAW2","10W","INS","BOLD:AAO8538","Arthropoda","Insecta","Diptera","Psychodidae","Psychodinae","Psychoda","","29","27","27","76","6.9","0","98.87","429.97","0","432.14","32.91","0"
"AGAW2","10W","INS","BOLD:ACV5892","Arthropoda","Insecta","Diptera","Cecidomyiidae","","","","29","28","28","76","3.45","0","97.86","446","0","447.1","25.41","0"
"AGAW2","10W","INS","BOLD:ACJ1941","Arthropoda","Insecta","Diptera","Mycetophilidae","Mycetophilinae","Zygomyia","","29","25","25","76","13.79","0","99.58","444.41","0","446.03","32.2","0"
"AGAW2","10W","INS","BOLD:ACE4233","Arthropoda","Insecta","Hymenoptera","Ichneumonidae","Orthocentrinae","Megastylus","Megastylus jft01","21","21","21","79","0","0","98.93","427.9","0","429.38","28.13","0"
"AGAW2","10W","INS","BOLD:AAN5255","Arthropoda","Insecta","Diptera","Cecidomyiidae","Cecidomyiinae","Dasineura","","15","15","15","80","0","0","99.01","426","0","446.07","26.95","0"
"AGAW2","10W","INS","BOLD:ACV5891","Arthropoda","Insecta","Diptera","Cecidomyiidae","","","","14","14","14","81","0","0","98.69","361.07","0","409.36","24.66","0"
"AGAW2","10W","INS","BOLD:AAN5220","Arthropoda","Insecta","Diptera","Cecidomyiidae","","","","14","14","14","81","0","0","98.67","401.64","0","420.93","26.12","0"
"AGAW2","10W","INS","BOLD:AAG5541","Arthropoda","Insecta","Diptera","Chironomidae","Chironominae","Polypedilum","","14","14","14","81","0","0","97.18","414.57","0","416.93","33.67","0"
"AGAW2","10W","INS","BOLD:AAB0373","Arthropoda","Insecta","Neuroptera","Chrysopidae","Chrysopinae","Chrysoperla","Chrysoperla sp. isolate 82, Chrysoperla zastrowi, Chrysoperla plorabunda, Chrysoperla sp. isolate 71, Chrysoperla agilis, Italochrysa italica, Chrysoperla sp. isolate 88, Chrysoperla lucasina, Chrysoperla carnea, Chrysoperla mediterranea, Chrysoperla pallida, Chrysoperla nipponensis, Chrysoperla sillemi","13","12","12","84","7.69","0","99.38","427.23","0","428.46","31.14","0"
"AGAW2","10W","INS","BOLD:AAI4303","Arthropoda","Insecta","Diptera","Chironomidae","Chironominae","Chironomus","Chironomus melanescens","13","12","12","84","7.69","0","98.89","429.08","0","432.23","34.42","0"
"AGAW2","10W","INS","BOLD:ACK5742","Arthropoda","Insecta","Diptera","Cecidomyiidae","","","","12","12","12","86","0","0","99.25","433.08","0","433.67","28.74","0"
"AGAW2","10W","INS","BOLD:ACD3276","Arthropoda","Insecta","Diptera","Cecidomyiidae","","","","12","12","12","86","0","0","99.18","446.92","0","447.67","28.22","0"
"AGAW2","10W","INS","BOLD:ABV1279","Arthropoda","Insecta","Diptera","Cecidomyiidae","","","","11","11","11","88","0","0","99.42","438.27","0","440.18","28.67","0"
"AGAW2","10W","INS","BOLD:AAN5148","Arthropoda","Insecta","Diptera","Ceratopogonidae","Forcipomyiinae","Forcipomyia","Forcipomyia bipunctata, Forcipomyia sp. CK-2012","11","11","11","88","0","0","99.02","416.45","0","419.18","35.75","0"
"AGAW2","10W","INS","BOLD:AAA3933","Arthropoda","Insecta","Lepidoptera","Erebidae","Herminiinae","Palthis","Palthis angulalis","10","10","10","90","0","0","99.04","417.6","0","419.9","31.85","0"
"AGAW2","10W","INS","BOLD:AAV1295","Arthropoda","Insecta","Diptera","Sciaridae","","Bradysia","Bradysia difformis, Bradysia impatiens","10","9","9","90","10","0","99.25","417","0","443.5","31.6","0"
"AGAW2","10W","INS","BOLD:ABZ4994","Arthropoda","Insecta","Orthoptera","Acrididae","Melanoplinae","Melanoplus","Melanoplus gladstoni, Melanoplus sanguinipes","10","10","10","90","0","0","97.22","449.1","0","450","36.38","0"
"AGAW2","10W","INS","BOLD:AAB4657","Arthropoda","Insecta","Diptera","Chironomidae","Chironominae","Chironomus","Chironomus maturus","9","9","9","93","0","0","99.23","447.67","0","448.67","37.54","0"
"AGAW2","10W","INS","BOLD:AAA3867","Arthropoda","Insecta","Lepidoptera","Noctuidae","Noctuinae","Apamea","Apamea amputatrix","9","9","9","93","0","0","99.19","433.33","0","435.56","33.46","0"
"AGAW2","10W","INS","BOLD:AAN5334","Arthropoda","Insecta","Diptera","Chironomidae","Chironominae","Polypedilum","","9","9","9","93","0","0","98.19","423.78","0","427.22","34.17","0"
"AGAW2","10W","INS","BOLD:ACV2677","Arthropoda","Insecta","Diptera","Cecidomyiidae","","","","8","8","8","96","0","0","97.6","448.88","0","450","29.86","0"
"AGAW2","10W","INS","BOLD:AAL7819","Arthropoda","Insecta","Diptera","Psychodidae","Psychodinae","Psychoda","Psychoda sp., Psychoda sp. BOLD:AAL7819","8","8","8","96","0","0","98.94","434.75","0","436.25","32.87","0"
"AGAW2","10W","INS","BOLD:AAG6743","Arthropoda","Insecta","Diptera","Sarcophagidae","Sarcophaginae","Sarcophaga","Sarcophaga sp. 1 MG, Sarcophaga subvicina","8","8","8","96","0","0","99.29","435.38","0","436.25","34.74","0"
"AGAW2","10W","INS","BOLD:AAG7012","Arthropoda","Insecta","Diptera","Limoniidae","Limoniinae","Geranomyia","","8","8","8","96","0","0","98.75","438.88","0","440.38","33.41","0"
"AGAW2","10W","INS","BOLD:AAA6560","Arthropoda","Insecta","Lepidoptera","Crambidae","Crambinae","Crambus","Crambus multilinellus, Crambus saltuellus","7","7","7","100","0","0","99.52","437.29","0","437.71","30.77","0"
"AGAW2","10W","INS","BOLD:AAB1352","Arthropoda","Insecta","Lepidoptera","Erebidae","Hypeninae","Hypena","Hypena sp.","7","7","7","100","0","0","99.14","449.14","0","450","32.6","0"
"AGAW2","10W","INS","BOLD:AAN5342","Arthropoda","Insecta","Diptera","Chironomidae","Orthocladiinae","Heleniella","","7","7","7","100","0","0","98.5","426.57","0","432.71","34.97","0"
"AGAW2","10W","INS","BOLD:AAI1526","Arthropoda","Insecta","Hemiptera","Aphididae","Aphidinae","Macrosiphoniella","Macrosiphoniella tapuskae","7","4","4","100","42.86","0","99.52","449.57","0","450","25.56","0"
"AGAW2","10W","INS","BOLD:AAA4277","Arthropoda","Insecta","Diptera","Syrphidae","Syrphinae","Toxomerus","Toxomerus marginatus, Toxomerus sp. CR-3, Toxomerus CR-3","7","7","7","100","0","0","98.71","423.43","0","425.29","30.72","0"
"AGAW2","10W","INS","BOLD:ACE4100","Arthropoda","Insecta","Orthoptera","Acrididae","Melanoplinae","Melanoplus","Melanoplus pinicola","6","6","6","105","0","0","97.58","406","0","408.33","35","0"
"AGAW2","10W","INS","BOLD:AAM6097","Arthropoda","Insecta","Diptera","Cecidomyiidae","","","","6","6","6","105","0","0","99.07","445","0","446.5","34.38","0"
"AGAW2","10W","INS","BOLD:AAA3836","Arthropoda","Insecta","Lepidoptera","Noctuidae","Plusiinae","Autographa","Autographa precationis","6","6","6","105","0","0","99.48","447.83","0","450","32.3","0"
"AGAW2","10W","INS","BOLD:ACI1304","Arthropoda","Insecta","Diptera","Muscidae","Muscinae","Stomoxys","Stomoxys indicus","6","6","6","105","0","0","97.47","421.5","0","426","32.77","0"
"AGAW2","10W","INS","BOLD:ACE8878","Arthropoda","Insecta","Orthoptera","Acrididae","Melanoplinae","Melanoplus","Melanoplus scudderi group","6","6","6","105","0","0","97.66","448.67","0","450","35.41","0"
"AGAW2","10W","INS","BOLD:ABX2489","Arthropoda","Insecta","Coleoptera","Phalacridae","Phalacrinae","Stilbus","","6","6","6","105","0","0","97.61","413.67","0","415.33","32.39","0"
"AGAW2","10W","INS","BOLD:AAC9062","Arthropoda","Insecta","Diptera","Culicidae","Culicinae","Ochlerotatus, Aedes","Aedes fitchii, Ochlerotatus fitchii, Aedes grossbecki, Aedes ventrovittis","6","6","6","105","0","0","99.13","439","0","440.17","32.78","0"
"AGAW2","10W","INS","BOLD:AAG5507","Arthropoda","Insecta","Diptera","Chironomidae","Chironominae","Chironomus","Chironomus atrella","5","5","5","112","0","0","99.17","431.2","0","432.2","36.21","0"
"AGAW2","10W","INS","BOLD:AAC5210","Arthropoda","Insecta","Diptera","Culicidae","Culicinae","Aedes, Aedimorphus","Aedes japonicus, Aedimorphus caecus","5","4","4","112","20","0","98.98","449.4","0","450","34.8","0"
"AGAW2","10W","INS","BOLD:AAG8882","Arthropoda","Insecta","Hemiptera","Pentatomidae","Pentatominae","Thyanta","Thyanta pallidovirens, Thyanta custator, Thyanta accerra","5","5","5","112","0","0","99.6","448.6","0","450","36.13","0"
"AGAW2","10W","INS","BOLD:AAA2326","Arthropoda","Insecta","Hymenoptera","Apidae","Apinae","Apis","Apis mellifera","5","5","5","112","0","0","99.82","448.6","0","450","26.36","0"
"AGAW2","10W","INS","BOLD:ACG3646","Arthropoda","Insecta","Coleoptera","Corylophidae","Corylophinae","Orthoperus","","5","5","5","112","0","0","97.75","443.8","0","445","33.89","0"
"AGAW2","10W","INS","BOLD:AAA7374","Arthropoda","Insecta","Diptera","Syrphidae","Syrphinae","Sphaerophoria","Sphaerophoria bankowskae, Sphaerophoria infuscata, Sphaerophoria nr. philanthus, Sphaerophoria batava, Sphaerophoria sp., Sphaerophoria rueppellii, Sphaerophoria interrupta-group, Sphaerophoria sp. MA-1, Sphaerophoria cleoae, Sphaerophoria asymmetrica, Sphaerophoria fatarum, Sphaerophoria bengalensis, Sphaerophoria bifurcata, Sphaerophoria taeniata, Sphaerophoria interrupta, Sphaerophoria contigua, Sphaerophoria kaa, Sphaerophoria laurae, Sphaerophoria chongjini, Sphaerophoria boreoalpina, Sphaerophoria sp. BOLDAAA7374, Sphaerophoria macrogaster, Sphaerophoria sulphuripes, Sphaerophoria brevipilosa, Sphaerophoria philanthus, Sphaerophoria pyrrhina, Sphaerophoria abbreviata, Sphaerophoria scripta, Sphaerophoria potentillae, Sphaerophoria longipilosa, Sphaerophoria virgata","5","5","5","112","0","0","98.7","439","0","441","30.92","0"
"AGAW2","10W","INS","BOLD:ACF2568","Arthropoda","Insecta","Orthoptera","Acrididae","Melanoplinae","Melanoplus","Melanoplus scudderi group, Melanoplus scudderi, Melanoplus quercicola","4","4","4","118","0","0","97.23","397.75","0","400.5","36.38","0"
"AGAW2","10W","INS","BOLD:AAF3338","Arthropoda","Insecta","Lepidoptera","Noctuidae","Noctuinae","Striacosta","Striacosta albicosta","4","4","4","118","0","0","98.85","418.25","0","420.25","33.92","0"
"AGAW2","10W","INS","BOLD:AAL1297","Arthropoda","Insecta","Coleoptera","Cantharidae","Cantharinae","Rhagonycha","Rhagonycha fulva","4","4","4","118","0","0","98.73","414.5","0","419","33.19","0"
"AGAW2","10W","INS","BOLD:AAZ5483","Arthropoda","Insecta","Diptera","Muscidae","Phaoniinae","Helina","Helina setiventris","4","4","4","118","0","0","99.37","443.5","0","445.75","29.32","0"
"AGAW2","10W","INS","BOLD:AAA2482","Arthropoda","Insecta","Lepidoptera","Noctuidae","Noctuinae","Mythimna","Mythimna unipuncta","4","4","4","118","0","0","99.48","400.75","0","403.75","30.08","0"
"AGAW2","10W","INS","BOLD:AAA2632","Arthropoda","Insecta","Lepidoptera","Noctuidae","Noctuinae","Noctua","Noctua atlantica, Noctua pronuba","4","4","4","118","0","0","97.94","449.75","0","450","31.17","0"
"AGAW2","10W","INS","BOLD:AAH2295","Arthropoda","Insecta","Diptera","Asilidae","Leptogastrinae","Leptogaster","Leptogaster flavipes","4","4","4","118","0","0","98.88","397.5","0","425","34.83","0"
"AGAW2","10W","INS","BOLD:ACE0554","Arthropoda","Insecta","Diptera","Phoridae","Metopininae","Megaselia","","4","4","4","118","0","0","99.78","447.75","0","450","32.17","0"
"AGAW2","10W","INS","BOLD:AAG6929","Arthropoda","Insecta","Diptera","Dolichopodidae","Medeterinae","Medetera","Medetera vittata","4","3","3","118","25","0","98.94","449.5","0","450","40.89","0"
"AGAW2","10W","INS","BOLD:AAN5267","Arthropoda","Insecta","Diptera","Cecidomyiidae","","","","4","4","4","118","0","0","99.83","448.5","0","450","30.67","0"
"AGAW2","10W","INS","BOLD:AAC4197","Arthropoda","Insecta","Diptera","Chironomidae","Orthocladiinae","Paraphaenocladius","Paraphaenocladius impensus","4","4","4","118","0","0","98.64","440.25","0","444","34.69","0"
"AGAW2","10W","INS","BOLD:AAL4232","Arthropoda","Insecta","Diptera","Agromyzidae","Phytomyzinae","Phytoliriomyza","Phytoliriomyza arctica","3","3","3","129","0","0","99.93","449.67","0","450","32.22","0"
"AGAW2","10W","INS","BOLD:AAA4213","Arthropoda","Insecta","Lepidoptera","Geometridae","Sterrhinae","Idaea","Idaea dimidiata","3","3","3","129","0","0","97.96","420","0","423","29.85","0"
"AGAW2","10W","INS","BOLD:ABA5202","Arthropoda","Insecta","Diptera","Limoniidae","Limnophilinae","Pseudolimnophila","Pseudolimnophila luteipennis","3","3","3","129","0","0","98.81","448","0","450","33.78","0"
"AGAW2","10W","INS","BOLD:AAA5299","Arthropoda","Insecta","Diptera","Chironomidae","Orthocladiinae","Cricotopus","Cricotopus laetus, Cricotopus relucens, Cricotopus trifasciatus, Cricotopus sp., Cricotopus sp. 3ES, Cricotopus sylvestris, Cricotopus glacialis","3","3","3","129","0","0","98.36","406.67","0","408","34.14","0"
"AGAW2","10W","INS","BOLD:ACR4394","Arthropoda","Insecta","Diptera","Anthomyiidae","Anthomyiinae","Delia","Delia florilega","3","3","3","129","0","0","99.55","444","0","445","33.19","0"
"AGAW2","10W","INS","BOLD:ACJ6984","Arthropoda","Insecta","Hymenoptera","Platygastridae","Scelioninae","Trissolcus","Trissolcus thyantae","3","3","3","129","0","0","99.7","439","0","450","27.04","0"
"AGAW2","10W","INS","BOLD:AAL2750","Arthropoda","Insecta","Diptera","Muscidae","Phaoniinae","Phaonia","Phaonia tuguriorum","3","3","3","129","0","0","99.84","409","0","412.33","31.21","0"
"AGAW2","10W","INS","BOLD:AAC2657","Arthropoda","Insecta","Lepidoptera","Coleophoridae","Coleophorinae","Coleophora","Coleophora salinoidella, Coleophora JFL205, Coleophora littorella, Coleophora lineapulvella","3","3","3","129","0","0","99.11","449.33","0","450","32.22","0"
"AGAW2","10W","INS","BOLD:ABV1237","Arthropoda","Insecta","Diptera","Chironomidae","Chironominae","Chironomus","","3","3","3","129","0","0","99.63","445.67","0","445","35.36","0"
"AGAW2","10W","INS","BOLD:AAN5162","Arthropoda","Insecta","Diptera","Ceratopogonidae","Dasyheleinae","Dasyhelea","","2","2","2","138","0","0","99.44","449.5","0","450","34.11","0"
"AGAW2","10W","INS","BOLD:AAH4701","Arthropoda","Insecta","Lepidoptera","Pterophoridae","Pterophorinae","Sphenarches","Sphenarches ontario","2","2","2","138","0","0","98.66","446.5","0","450","30.44","0"
"AGAW2","10W","INS","BOLD:AAN5332","Arthropoda","Insecta","Diptera","Chironomidae","Chironominae","Polypedilum","","2","2","2","138","0","0","97.9","426.5","0","428","34.07","0"
"AGAW2","10W","INS","BOLD:AAD0952","Arthropoda","Insecta","Lepidoptera","Pyralidae","Chrysauginae","Galasa","Galasa nigrinodis, Galasa nigripunctalis","2","2","2","138","0","0","99.49","395.5","0","398.5","32.25","0"
"AGAW2","10W","INS","BOLD:AAA5206","Arthropoda","Insecta","Lepidoptera","Erebidae","Herminiinae","Zanclognatha","Zanclognatha cruralis, Zanclognatha sp. 1, Zanclognatha jacchusalis, Zanclognatha minoralis, Zanclognatha minoralis sp. 1, Zanclognatha sp., Zanclognatha minoralis sp. 3, Zanclognatha gypsalis, Zanclognatha dentata, Zanclognatha theralis, Zanclognatha protumnusalis, Zanclognatha martha, Zanclognatha atrilineella, Zanclognatha obscuripennis, Zanclognatha minoralis sp. 2, Zanclognatha marcidilinea","2","2","2","138","0","0","97.46","411.5","0","413.5","30.63","0"
"AGAW2","10W","INS","BOLD:AAH3920","Arthropoda","Insecta","Diptera","Sciaridae","","Dichopygina, Scatopsciara","","2","2","2","138","0","0","99.25","398","0","450","36.22","0"
"AGAW2","10W","INS","BOLD:AAB5962","Arthropoda","Insecta","Lepidoptera","Tortricidae","Tortricinae","Phalonidia","Phalonidia lepidana","2","2","2","138","0","0","100","449","0","450","33.22","0"
"AGAW2","10W","INS","BOLD:ABX5883","Arthropoda","Insecta","Lepidoptera","Tortricidae","Tortricinae","Choristoneura","Choristoneura retiniana, Choristoneura lambertiana, Choristoneura biennis, Choristoneura orae, Choristoneura fumiferana, Choristoneura occidentalis, Choristoneura freemani, Choristoneura pinus, Choristoneura occidentalis group, Choristoneura sp., Choristoneura carnana","2","2","2","138","0","0","98.77","447","0","450","31.11","0"
"AGAW2","10W","INS","BOLD:ACE5981","","","","","","","","2","2","2","138","0","0","98.02","376.5","0","377.5","35.65","0"
"AGAW2","10W","INS","BOLD:ACB1322","Arthropoda","Insecta","Coleoptera","Mordellidae","Mordellinae","Mordellistena","Mordellistena trifasciata","2","2","2","138","0","0","99.75","404","0","443.5","41.94","0"
"AGAW2","10W","INS","BOLD:AAV1136","Arthropoda","Insecta","Diptera","Scatopsidae","","","","2","2","2","138","0","0","97.99","448.5","0","450","37","0"
"AGAW2","10W","INS","BOLD:ABV1188","Arthropoda","Insecta","Diptera","Chironomidae","Chironominae","Tanytarsus","","2","2","2","138","0","0","97.55","326","0","379.5","34.66","0"
"AGAW2","10W","INS","BOLD:AAG1743","Arthropoda","Insecta","Diptera","Muscidae","Phaoniinae","Phaonia","Phaonia fuscana","2","2","2","138","0","0","97.31","408.5","0","410","32.56","0"
"AGAW2","10W","INS","BOLD:ACG5862","Arthropoda","Insecta","Diptera","Agromyzidae","","","","2","2","2","138","0","0","98.89","450","0","450","34","0"
"AGAW2","10W","INS","BOLD:AAH2293","Arthropoda","Insecta","Diptera","Asilidae","Stenopogoninae","Dioctria, Eudioctria","Dioctria hyalipennis, Dioctria baumhaueri, Eudioctria propinqua","2","2","2","138","0","0","98.78","406.5","0","408","30.61","0"
"AGAW2","10W","INS","BOLD:AAH3662","Arthropoda","Insecta","Diptera","Cecidomyiidae","","","CecidInt3 sp. BOLD:AAH3662","2","2","2","138","0","0","99.45","364","0","368.5","25.38","0"
"AGAW2","10W","INS","BOLD:AAA9251","Arthropoda","Insecta","Lepidoptera","Gelechiidae","Gelechiinae","Scrobipalpa","Scrobipalpa atriplicella, Scrobipalpa nr. atriplicella sp. 1","2","2","2","138","0","0","98.95","434","0","435.5","33.98","0"
"AGAW2","10W","INS","BOLD:AAN6447","Arthropoda","Insecta","Diptera","Sciaridae","","Corynoptera","Corynoptera perpusilla","2","2","2","138","0","0","99.65","437.5","0","438.5","36.61","0"
"AGAW2","10W","INS","BOLD:AAG5499","Arthropoda","Insecta","Diptera","Chironomidae","Orthocladiinae","Psectrocladius","Psectrocladius obvius, Psectrocladius platypus","2","2","2","138","0","0","99.78","448.5","0","450","35.67","0"
"AGAW2","10W","INS","BOLD:ACJ0111","Arthropoda","Insecta","Hymenoptera","Platygastridae","Scelioninae","Telenomus","","2","2","2","138","0","0","97.91","398.5","0","400.5","28.64","0"
"AGAW2","10W","INS","BOLD:ACA2933","Arthropoda","Insecta","Psocodea","Stenopsocidae","","Graphopsocus","Graphopsocus cruciatus","2","2","2","138","0","0","99.54","435.5","0","438","38.35","0"
"AGAW2","10W","INS","BOLD:ADH9867","Arthropoda","Insecta","Diptera","","","","","2","2","2","138","0","0","98.7","404.5","0","406.5","37.37","0"
"AGAW2","10W","INS","BOLD:AAF3508","Arthropoda","Insecta","Hymenoptera","Ichneumonidae","Orthocentrinae","Megastylus","Megastylus jft01","2","2","2","138","0","0","98.44","449","0","450","27.89","0"
"AGAW2","10W","INS","BOLD:AAG8842","Arthropoda","Insecta","Hemiptera","Cicadellidae","Deltocephalinae","Scaphytopius","Scaphytopius acutus, Scaphytopius latus, Scaphytopius oregonensis","2","2","2","138","0","0","99.76","435","0","435.5","35.24","0"
"AGAW2","10W","INS","BOLD:AAM6295","Arthropoda","Insecta","Diptera","Chaoboridae","","","","2","2","2","138","0","0","99.67","450","0","450","37.22","0"
"AGAW2","10W","INS","BOLD:AAN5221","Arthropoda","Insecta","Diptera","Cecidomyiidae","","","","2","2","2","138","0","0","98.71","401","0","401.5","25.21","0"
"AGAW2","10W","INS","BOLD:AAD8027","Arthropoda","Insecta","Diptera","Culicidae","Culicinae","Aedes","Aedes fitchii","2","2","2","138","0","0","100","450","0","450","34.67","0"
"AGAW2","10W","INS","BOLD:ACF2457","Arthropoda","Insecta","Diptera","Chironomidae","Chironominae","Paratanytarsus","Paratanytarsus laccophilus","2","2","2","138","0","0","99.56","450","0","450","32.78","0"
"AGAW2","10W","INS","BOLD:AAC0635","Arthropoda","Insecta","Hemiptera","Miridae","Mirinae","Stenotus","Stenotus binotatus","2","1","1","138","50","0","100","450","0","450","34.67","0"
"AGAW2","10W","INS","BOLD:AAI1345","Arthropoda","Insecta","Diptera","Limoniidae","Limoniinae","Dicranomyia","Dicranomyia haeretica, Dicranomyia (Idiopyga) halterella, Dicranomyia (Dicranomyia) haeretica, Dicranomyia (Dicranomyia) longipennis, Dicranomyia moniliformis","1","1","1","167","0","0","98.44","449","0","450","34.22","0"
"AGAW2","10W","INS","BOLD:AAZ5592","Arthropoda","Insecta","Diptera","Psychodidae","Psychodinae","","","1","1","1","167","0","0","99.33","450","0","450","33.33","0"
"AGAW2","10W","INS","BOLD:AAE1681","Arthropoda","Insecta","Diptera","Tabanidae","Tabaninae","Hybomitra","Hybomitra hearlei, Hybomitra frosti, Hybomitra sexfasciata, Hybomitra itasca, Hybomitra minuscula","1","1","1","167","0","0","97.33","450","0","450","32.44","0"
"AGAW2","10W","INS","BOLD:ACH4728","Arthropoda","Insecta","Psocodea","Elipsocidae","Elipsocinae","Cuneopalpus","Cuneopalpus cyanops","1","1","1","167","0","0","99.78","448","0","450","38.67","0"
"AGAW2","10W","INS","BOLD:ACF4734","Arthropoda","Insecta","Diptera","Syrphidae","Syrphinae","Platycheirus","Platycheirus hyperboreus","1","1","1","167","0","0","99.33","447","0","450","31.56","0"
"AGAW2","10W","INS","BOLD:ACE6318","Arthropoda","Insecta","Psocodea","Psocidae","Psocinae","Trichadenotecnum","Trichadenotecnum alexanderae complex","1","1","1","167","0","0","98.85","436","0","440","36.82","0"
"AGAW2","10W","INS","BOLD:AAO7783","Arthropoda","Insecta","Diptera","Dolichopodidae","Diaphorinae","","","1","1","1","167","0","0","98.22","450","0","450","32.89","0"
"AGAW2","10W","INS","BOLD:ABU5546","Arthropoda","Insecta","Diptera","Psychodidae","Psychodinae","Psychoda","","1","1","1","167","0","0","98.2","445","0","450","32","0"
"AGAW2","10W","INS","BOLD:AAA2518","Arthropoda","Insecta","Lepidoptera","Crambidae","Crambinae","Chrysoteuchia","Chrysoteuchia topiaria, Chrysoteuchia topiarius","1","1","1","167","0","0","99.55","449","0","450","35.78","0"
"AGAW2","10W","INS","BOLD:AAH2306","Arthropoda","Insecta","Diptera","Asilidae","Asilinae","Efferia","Efferia aestuans","1","1","1","167","0","0","99.56","450","0","450","38.44","0"
"AGAW2","10W","INS","BOLD:AAE3675","Arthropoda","Insecta","Diptera","Chironomidae","Chironominae","Paratanytarsus","Paratanytarsus sp. TE03","1","1","1","167","0","0","100","449","0","450","36.22","0"
"AGAW2","10W","INS","BOLD:ABA6163","Arthropoda","Insecta","Psocodea","Caeciliusidae","Caeciliusinae","Valenzuela","Valenzuela piceus","1","1","1","167","0","0","99.11","450","0","450","37.11","0"
"AGAW2","10W","INS","BOLD:AAP7026","Arthropoda","Insecta","Coleoptera","Latridiidae","Corticariinae","Melanophthalma","Melanophthalma villosa","1","1","1","167","0","0","98.2","388","0","450","31.33","0"
"AGAW2","10W","INS","BOLD:AAN5309","Arthropoda","Insecta","Diptera","Chironomidae","Chironominae","","","1","1","1","167","0","0","97.34","376","0","373","37.8","0"
"AGAW2","10W","INS","BOLD:AAM6781","Arthropoda","Insecta","Diptera","Dolichopodidae","Neurigoninae","Neurigona","Neurigona tenuis","1","1","1","167","0","0","99.33","448","0","450","35.56","0"
"AGAW2","10W","INS","BOLD:AAA8640","Arthropoda","Insecta","Lepidoptera","Tortricidae","Olethreutinae","Eucosma, Sonia, Pelochrista","Pelochrista suadana, Sonia new sp. 1, Sonia new sp. 2, Sonia new sp. 3, Sonia paraplesiana, Sonia canadana","1","1","1","167","0","0","99.11","449","0","450","32.89","0"
"AGAW2","10W","INS","BOLD:AAW0101","","","","","","","","1","1","1","167","0","0","100","338","0","367","31.88","0"
"AGAW2","10W","INS","BOLD:AAP2602","Arthropoda","Insecta","Psocodea","Psocidae","","","","1","1","1","167","0","0","99.55","447","0","450","37.33","0"
"AGAW2","10W","INS","BOLD:AAB4658","Arthropoda","Insecta","Diptera","Chironomidae","Chironominae","Chironomus","Chironomus dilutus","1","1","1","167","0","0","98.66","448","0","450","38","0"
"AGAW2","10W","INS","BOLD:ACA9275","Arthropoda","Insecta","Diptera","Chironomidae","Chironominae","Chironomus","","1","1","1","167","0","0","99.78","449","0","450","35.56","0"
"AGAW2","10W","INS","BOLD:AAC1042","Arthropoda","Insecta","Lepidoptera","Crambidae","Spilomelinae","Lygropia","Lygropia rivulalis","1","1","1","167","0","0","100","448","0","450","28.89","0"
"AGAW2","10W","INS","BOLD:ACL1796","Arthropoda","Insecta","Lepidoptera","Ypsolophidae","Ochsenheimeriinae","Ochsenheimeria","Ochsenheimeria vacculella","1","1","1","167","0","0","98.48","395","0","450","32.67","0"
"AGAW2","10W","INS","BOLD:ADF2969","Arthropoda","Insecta","Diptera","Sarcophagidae","Sarcophaginae","Blaesoxipha","Blaesoxipha falciformis","1","1","1","167","0","0","97.78","450","0","450","37.56","0"
"AGAW2","10W","INS","BOLD:ACS9320","Arthropoda","Insecta","Diptera","Cecidomyiidae","","","","1","1","1","167","0","0","97.76","446","0","450","30","0"
"AGAW2","10W","INS","BOLD:ACE6862","Arthropoda","Insecta","Diptera","Chironomidae","Chironominae","Polypedilum","","1","1","1","167","0","0","97.56","450","0","450","33.33","0"
"AGAW2","10W","INS","BOLD:AAN5238","Arthropoda","Insecta","Diptera","Cecidomyiidae","","","","1","1","1","167","0","0","98.22","450","0","450","26.67","0"
"AGAW2","10W","INS","BOLD:AAJ7123","Arthropoda","Insecta","Diptera","Culicidae","Culicinae","Culiseta","Culiseta minnesotae","1","1","1","167","0","0","99.78","449","0","450","34.89","0"
"AGAW2","10W","INS","BOLD:AAU6592","Arthropoda","Insecta","Diptera","Ceratopogonidae","Dasyheleinae","Dasyhelea","Dasyhelea arenivaga","1","1","1","167","0","0","98.67","375","0","379","34.3","0"
"AGAW2","10W","INS","BOLD:AAA7631","Arthropoda","Insecta","Hemiptera","Miridae","Mirinae","Adelphocoris","Adelphocoris lineolatus","1","1","1","167","0","0","97.55","449","0","450","36.22","0"
"AGAW2","10W","INS","BOLD:ACL8669","Arthropoda","Insecta","Lepidoptera","Psychidae","Psychinae","Psyche","Psyche casta","1","1","1","167","0","0","100","377","0","378","37.3","0"
"AGAW2","10W","INS","BOLD:AAG6433","Arthropoda","Insecta","Diptera","Ceratopogonidae","Forcipomyiinae","Forcipomyia","Forcipomyia sp. BOLD:AAG6433, Forcipomyia sp. 17","1","1","1","167","0","0","99.11","449","0","450","32","0"
"AGAW2","10W","INS","BOLD:AAG2017","Arthropoda","Insecta","Neuroptera","Chrysopidae","Chrysopinae","Chrysopa","Chrysopa oculata","1","1","1","167","0","0","99.33","449","0","450","31.56","0"
"AGAW2","10W","INS","BOLD:AAG3154","Arthropoda","Insecta","Diptera","Psychodidae","Psychodinae","Psychoda","","1","1","1","167","0","0","98.22","450","0","450","32","0"
"AGAW2","10W","INS","BOLD:AAM9105","Arthropoda","Insecta","Diptera","Muscidae","Coenosiinae","Coenosia","","1","1","1","167","0","0","99.26","406","0","408","36.27","0"
"AGAW2","10W","INS","BOLD:AAG9897","Arthropoda","Insecta","Dermaptera","Forficulidae","Forficulinae","Forficula","Forficula auricularia-A","1","1","1","167","0","0","99.77","428","0","450","46.44","0"
"AGAW2","10W","INS","BOLD:AAB5609","Arthropoda","Insecta","Diptera","Muscidae","Coenosiinae","Coenosia","Coenosia sp.BOLD:AAB5609, Coenosia tigrina","1","1","1","167","0","0","97.32","448","0","450","32.89","0"
"AGAW2","10W","INS","BOLD:AAG2334","Arthropoda","Insecta","Diptera","Tachinidae","Phasiinae","Gymnoclytia","Gymnoclytia occidua, Gymnoclytia sp. 1, Gymnoclytia sp. 2","1","1","1","167","0","0","98.87","443","0","450","30.44","0"
"AGAW2","10W","INS","BOLD:AAD2041","Arthropoda","Insecta","Diptera","Dolichopodidae","Sciapodinae","Condylostylus","Condylostylus inermis, Condylostylus nigrofemoratus, Condylostylus connectans, Condylostylus caudatus, Condylostylus flavipes","1","1","1","167","0","0","98","449","0","450","35.11","0"
"AGAW2","10W","INS","BOLD:AAA7039","Arthropoda","Insecta","Lepidoptera","Tortricidae","Tortricinae","Sparganothis","Sparganothis bistriata, Sparganothis robinsonana, Sparganothis sulfureana, Sparganothis lycopodiana, Sparganothis tristriata","1","1","1","167","0","0","98.89","450","0","450","32","0"
"AGAW2","10W","INS","BOLD:AAU7126","Arthropoda","Insecta","Coleoptera","Cleridae","Hydnocerinae","Isohydnocera","Isohydnocera curtipennis","1","1","1","167","0","0","99.78","449","0","450","33.56","0"
"AGAW2","10W","INS","BOLD:AAF5980","Arthropoda","Insecta","Hemiptera","Cicadellidae","Typhlocybinae","Typhlocyba","Typhlocyba pomaria","1","1","1","167","0","0","97.55","449","0","450","37.11","0"
"AGAW2","10W","INS","BOLD:AAB9332","Arthropoda","Insecta","Hemiptera","Aphididae","Aphidinae","Nasonovia","Nasonovia ribisnigri","1","1","1","167","0","0","99.78","449","0","450","25.33","0"
"AGAW2","10W","INS","BOLD:ACE8875","Arthropoda","Insecta","Orthoptera","Acrididae","Melanoplinae","Melanoplus","Melanoplus bowditchi","1","1","1","167","0","0","97.1","449","0","450","36.67","0"
"AGAW2","10W","INS","BOLD:ACL6109","Arthropoda","Insecta","Psocodea","Psocidae","","","","1","1","1","167","0","0","97.24","434","0","435","37.24","0"
"AGAW2","10W","INS","BOLD:AAB7533","Arthropoda","Insecta","Lepidoptera","Coleophoridae","Coleophorinae","Coleophora","Coleophora detractella","1","1","1","167","0","0","98","401","0","402","32.09","0"
"AGAW2","10W","INS","BOLD:AAF9187","Arthropoda","Insecta","Coleoptera","Curculionidae","Entiminae","Phyllobius","Phyllobius oblongus","1","1","1","167","0","0","99.55","449","0","450","39.33","0"
"AGAW2","10W","INS","BOLD:AAV4870","Arthropoda","Insecta","Diptera","Agromyzidae","","","","1","1","1","167","0","0","98.66","447","0","450","29.78","0"
"AGAW2","10W","INS","BOLD:AAB5969","Arthropoda","Insecta","Lepidoptera","Noctuidae","Noctuinae","Loscopia","Loscopia velata","1","1","1","167","0","0","99.77","431","0","434","31.11","0"
"AGAW2","10W","INS","BOLD:AAH7158","Arthropoda","Insecta","Diptera","Sarcophagidae","Sarcophaginae","Ravinia","Ravinia stimulans","1","1","1","167","0","0","98.88","445","0","450","33.56","0"
"AGAW2","10W","INS","BOLD:AAG1741","Arthropoda","Insecta","Diptera","Muscidae","Coenosiinae","Coenosia","","1","1","1","167","0","0","98.88","448","0","450","33.78","0"
"AGAW2","10W","INS","BOLD:ACU7270","Arthropoda","Insecta","Diptera","Cecidomyiidae","","","","1","1","1","167","0","0","99.27","411","0","415","23.61","0"
"AGAW2","10W","INS","BOLD:AAH0255","Arthropoda","Insecta","Coleoptera","Chrysomelidae","Galerucinae","Phyllotreta","Phyllotreta cruciferae, Phyllotreta albionica","1","1","1","167","0","0","99.78","447","0","450","32.67","0"
"AGAW2","10W","INS","BOLD:AAC9070","Arthropoda","Insecta","Hemiptera","Miridae","Mirinae","Poecilocapsus","Poecilocapsus lineatus","1","1","1","167","0","0","98.89","451","0","450","39.56","0"
"AGAW2","10W","INS","BOLD:AAN5201","Arthropoda","Insecta","Diptera","Cecidomyiidae","","","","1","1","1","167","0","0","100","450","0","450","28","0"
"AGAW2","10W","INS","BOLD:AAU6533","Arthropoda","Insecta","Diptera","Phoridae","Metopininae","Megaselia","","1","1","1","167","0","0","100","356","0","450","32.22","0"
"AGAW2","10W","INS","BOLD:AAG2355","Arthropoda","Insecta","Diptera","Tachinidae","Tachininae","Strongygaster","Strongygaster triangulifera, Strongygaster sp.","1","1","1","167","0","0","100","443","0","444","31.53","0"
"AGAW2","10W","INS","BOLD:AAP9022","Arthropoda","Insecta","Diptera","Cecidomyiidae","","","","1","1","1","167","0","0","99.74","391","0","393","27.48","0"
"AGAW2","10W","INS","BOLD:AAA4178","Arthropoda","Insecta","Lepidoptera","Geometridae","Larentiinae","Eupithecia","Eupithecia miserulata, Eupithecia chlorofasciata","1","1","1","167","0","0","100","445","0","450","30.89","0"
"AGAW2","10W","INS","BOLD:ACE9865","Arthropoda","Insecta","Lepidoptera","Noctuidae","Noctuinae","Feltia","Feltia subgothica, Feltia herilis, Feltia tricosa","1","1","1","167","0","0","99.56","450","0","450","31.11","0"
"AGAW2","10W","INS","BOLD:ACF8174","Arthropoda","Insecta","Diptera","Chironomidae","Chironominae","","","1","1","1","167","0","0","97.94","436","0","442","33.48","0"
"AGAW2","10W","INS","BOLD:ABV1430","Arthropoda","Insecta","Diptera","Dolichopodidae","Medeterinae","Medetera","","1","1","1","167","0","0","99.22","387","0","397","37.78","0"
"AGAW2","10W","INS","BOLD:AAN5661","Arthropoda","Insecta","Diptera","Chloropidae","Oscinellinae","Olcella","Olcella parva","1","1","1","167","0","0","97.68","431","0","435","31.03","0"
"AGAW2","10W","INS","BOLD:AAL4268","Arthropoda","Insecta","Diptera","Agromyzidae","Phytomyzinae","Phytomyza","Phytomyza flavofemorata, Phytomyza crassiseta","1","1","1","167","0","0","99.78","450","0","450","34.67","0"
"AGAW2","10W","INS","BOLD:AAP6471","Arthropoda","Insecta","Diptera","Sciaridae","","Pseudolycoriella","","1","1","1","167","0","0","98.89","450","0","450","36.67","0"
"AGAW2","10W","INS","BOLD:AAB6095","Arthropoda","Insecta","Lepidoptera","Noctuidae","Noctuinae","Neoligia","Neoligia exhausta","1","1","1","167","0","0","99.04","417","0","417","31.41","0"
"AGAW2","10W","INS","BOLD:AAB9516","Arthropoda","Insecta","Diptera","Chironomidae","Orthocladiinae","Acricotopus","","1","1","1","167","0","0","98.89","450","0","450","36.89","0"
"AGAW2","10W","INS","BOLD:ADH9310","Arthropoda","Insecta","Diptera","Anthomyiidae","Anthomyiinae","Anthomyia","Anthomyia procellaris, Anthomyia pluvialis","1","1","1","167","0","0","99.78","446","0","450","31.78","0"
"AGAW2","10W","INS","BOLD:AAH3914","Arthropoda","Insecta","Diptera","Sciaridae","","Sciara","Sciara hamatilis, Sciara humeralis","1","1","1","167","0","0","99.09","438","0","439","30.75","0"
"AGAW2","10W","NAR","BOLD:AAL1512","Mollusca","Gastropoda","Stylommatophora","Agriolimacidae",NA,"Deroceras","Deroceras laeve","2897","2827","2827","1","2.42","0","97.86","382.86","0","386.18","33.03","0"
"AGAW2","10W","NAR","TAX:660402","Mollusca","Gastropoda","Stylommatophora","Agriolimacidae",NA,"Deroceras","Deroceras sp.","607","605","605","2","0.33","0","97.75","392.85","0","395.5","32.77","0"
"AGAW2","10W","NAR","TAX:106138","","","","",NA,"","","8","8","8","3","0","0","97.71","389.13","0","392.13","32.65","0"
"AGAW2","10W","NAR","BOLD:AAG1548","Mollusca","Gastropoda","Stylommatophora","Agriolimacidae",NA,"Deroceras","Deroceras laeve","5","4","4","4","20","0","97.13","445.4","0","447","32.48","0"
"AGAW2","10W","NAR","BOLD:AAI9663","Mollusca","Gastropoda","Stylommatophora","Agriolimacidae",NA,"Deroceras","Deroceras reticulatum","3","3","3","5","0","0","99.75","405.67","0","450","35.78","0"
"AGAW2","10W","NIN","BOLD:AAH6630","Arthropoda","Collembola","Entomobryomorpha","Entomobryidae","Entomobryinae","Entomobrya","Entomobrya unostrigata","5000","4078","4078","1","18.44","0","99.33","436.85","0","438.09","39.08","0"
"AGAW2","10W","NIN","BOLD:AAA6381","Arthropoda","Arachnida","Araneae","Tetragnathidae","","Tetragnatha","Tetragnatha laboriosa","402","391","391","2","2.74","0","98.89","426.68","0","428.58","39.15","0"
"AGAW2","10W","NIN","BOLD:ACK1257","Arthropoda","Arachnida","Trombidiformes","Erythraeidae","","Balaustium","","190","147","147","3","22.63","0","99.65","443.89","0","444.76","32.71","0"
"AGAW2","10W","NIN","BOLD:AAI5320","Arthropoda","Arachnida","Araneae","Linyphiidae","","Islandiana","Islandiana flaveola","75","72","72","4","4","0","98.88","410.71","0","426.49","34.05","0"
"AGAW2","10W","NIN","BOLD:ABW2639","Arthropoda","Arachnida","Trombidiformes","Anystidae","","","","39","33","33","5","15.38","0","99.67","449.15","0","448.82","31.48","0"
"AGAW2","10W","NIN","BOLD:AAA8820","Arthropoda","Collembola","Entomobryomorpha","Entomobryidae","Lepidocyrtinae","Lepidocyrtus","Lepidocyrtus cyaneus","14","14","14","6","0","0","98.9","410.79","0","413.43","43.11","0"
"AGAW2","10W","NIN","BOLD:ACI9974","Arthropoda","Arachnida","Trombidiformes","Tydeidae","","","","12","11","11","7","8.33","0","99","424.67","0","450","40.91","0"
"AGAW2","10W","NIN","BOLD:ACC0359","Arthropoda","Collembola","Symphypleona","Bourletiellidae","","Bourletiella","","11","11","11","8","0","0","98.88","437.45","0","441.18","37.72","0"
"AGAW2","10W","NIN","BOLD:ACV5610","","","","","","","","9","9","9","9","0","0","99.72","436.67","0","439.11","37.94","0"
"AGAW2","10W","NIN","BOLD:AAB7914","Arthropoda","Collembola","Symphypleona","Bourletiellidae","","Bourletiella","","7","7","7","10","0","0","99.74","448.14","0","450","38.32","0"
"AGAW2","10W","NIN","BOLD:AAU6252","Arthropoda","Arachnida","Trombidiformes","Bdellidae","","Cyta","","5","4","4","11","20","0","99.47","449.6","0","450","31.78","0"
"AGAW2","10W","NIN","BOLD:ACZ1295","Arthropoda","Arachnida","Trombidiformes","Anystidae","","","","4","4","4","12","0","0","98.65","367.25","0","369.25","30.94","0"
"AGAW2","10W","NIN","BOLD:ACX8969","Arthropoda","Arachnida","Trombidiformes","Eupodidae","","","","3","3","3","13","0","0","97.88","426.33","0","429.67","27.86","0"
"AGAW2","10W","NIN","BOLD:ACX9836","Arthropoda","Arachnida","Trombidiformes","Tarsonemidae","","","","1","1","1","14","0","0","100","395","0","402","25.12","0"
"AGAW2","10W","NIN","BOLD:AAA7242","Arthropoda","Collembola","Entomobryomorpha","Entomobryidae","Seirinae, Entomobryinae","Entomobrya, Lepidosira","Entomobrya multifasciata, Lepidosira sp.","1","1","1","14","0","0","99.11","447","0","450","39.33","0"
"AGAW2","10W","NIN","BOLD:ACR6863","","","","","","","","1","1","1","14","0","0","97","434","0","437","38.9","0"
"AGAW2","10W","NIN","BOLD:AAO8147","Arthropoda","Arachnida","Trombidiformes","Eupodidae","","Eupodes","","1","1","1","14","0","0","99.15","354","0","356","26.97","0"
"AGAW2","10W","NIN","BOLD:AAB0726","Arthropoda","Arachnida","Araneae","Lycosidae","","Trochosa","Trochosa ruricola","1","1","1","14","0","0","98.44","449","0","450","32","0"
"AGAW2","10W","NIN","BOLD:AAI4346","Arthropoda","Arachnida","Opiliones","Phalangiidae","","Paroligolophus, Phalangium","Paroligolophus agrestis, Phalangium opilio","1","1","1","14","0","0","99.56","450","0","450","36.67","0"
"AGAW2","11E","BAC","TAX:175768","Proteobacteria","Alphaproteobacteria","Rickettsiales","Rickettsiaceae",NA,"Wolbachia","Wolbachia pipientis","6","6","6","1","0","0","98.06","439.67","0","441.33","39.46","0"
"AGAW2","11E","BAC","TAX:375853","Proteobacteria","Alphaproteobacteria","Rickettsiales","Rickettsiaceae",NA,"Wolbachia","Wolbachia sp. Hymenoptera_ASGLE646-10","4","4","4","2","0","0","97.83","449.5","0","450","40.39","0"
"AGAW2","11E","BAC","TAX:91393","Proteobacteria","Alphaproteobacteria","Rickettsiales","Rickettsiaceae",NA,"Wolbachia","","2","2","2","3","0","0","99.49","392","0","409.5","39.07","0"
"AGAW2","11E","BAC","TAX:392874","Proteobacteria","Alphaproteobacteria","Rickettsiales","Rickettsiaceae",NA,"Wolbachia","Wolbachia endosymbiont of Hymenoptera","1","1","1","4","0","0","98.43","445","0","450","40","0"
"AGAW2","11E","BAC","TAX:376688","Proteobacteria","Alphaproteobacteria","Rickettsiales","Rickettsiaceae",NA,"Wolbachia","Wolbachia sp. elachBioLep01 BioLep373","1","1","1","4","0","0","99.33","448","0","450","40","0"
"AGAW2","11E","CHO","no match","no match","no match","no match","no match","no match","no match","no match","no match","no match","no match","no match","no match","no match","no match","no match","no match","no match","no match","no match"
"AGAW2","11E","INS","BOLD:AAH2306","Arthropoda","Insecta","Diptera","Asilidae","Asilinae","Efferia","Efferia aestuans","7055","5784","5784","1","18.02","0","99.3","434.07","0","435.48","37.98","0"
"AGAW2","11E","INS","BOLD:AAA2482","Arthropoda","Insecta","Lepidoptera","Noctuidae","Noctuinae","Mythimna","Mythimna unipuncta","4236","3841","3841","2","9.32","0","99.21","414.5","0","417.57","30.13","0"
"AGAW2","11E","INS","BOLD:AAG1711","Arthropoda","Insecta","Diptera","Muscidae","Phaoniinae","Helina","Helina depuncta","2453","2002","2002","3","18.39","0","99.49","435.36","0","437.62","31.35","0"
"AGAW2","11E","INS","BOLD:ABY5318","Arthropoda","Insecta","Diptera","Chloropidae","Oscinellinae","Incertella, Rhopalopterum","Rhopalopterum umbrosum, Rhopalopterum luteiceps, Incertella bispina, Rhopalopterum painteri","1999","1459","1459","4","27.01","0","99.39","443.52","0","444.65","32.2","0"
"AGAW2","11E","INS","BOLD:AAA4555","Arthropoda","Insecta","Orthoptera","Acrididae","Melanoplinae","Melanoplus","Melanoplus stupefactus, Melanoplus angustipennis, Melanoplus ponderosus, Melanoplus bruneri, Melanoplus frigidus, Melanoplus scapularis, Melanoplus cinereus, Melanoplus madeleineae, Melanoplus sanguinipes, Melanoplus nigrescens group, Melanoplus tunicae, Melanoplus walshi, Melanoplus femurrubrum, Melanoplus dawsoni, Melanoplus sp., Melanoplus fasciatus, Melanoplus bowditchi, Melanoplus stonei, Melanoplus bivittatus, Melanoplus huroni, Melanoplus thomasi, Melanoplus borealis, Melanoplus packardii","1766","1383","1383","5","21.69","0","99.48","447.48","0","448.52","36.21","0"
"AGAW2","11E","INS","BOLD:AAD2041","Arthropoda","Insecta","Diptera","Dolichopodidae","Sciapodinae","Condylostylus","Condylostylus inermis, Condylostylus nigrofemoratus, Condylostylus connectans, Condylostylus caudatus, Condylostylus flavipes","1577","1141","1141","6","27.65","0","99.62","437.79","0","439.02","35.95","0"
"AGAW2","11E","INS","BOLD:AAK5928","Arthropoda","Insecta","Coleoptera","Chrysomelidae","Criocerinae","Oulema","Oulema melanopus","1575","1246","1246","7","20.89","0","99.58","440.49","0","441.69","36.41","0"
"AGAW2","11E","INS","BOLD:AAA3453","Arthropoda","Insecta","Diptera","Anthomyiidae","Anthomyiinae","Delia","","1556","1435","1435","8","7.78","0","99.43","428.71","0","431.3","34.6","0"
"AGAW2","11E","INS","BOLD:ACE7444","Arthropoda","Insecta","Hemiptera","Miridae","Mirinae, Phylinae","Adelphocoris, Megalocoleus","Adelphocoris lineolatus, Megalocoleus molliculus","1500","991","991","9","33.93","0","99.73","440.92","0","441.6","34.8","0"
"AGAW2","11E","INS","BOLD:AAN6154","Arthropoda","Insecta","Coleoptera","Latridiidae","Corticariinae","Melanophthalma","Melanophthalma inermis","1487","1252","1252","10","15.8","0","98.83","433.96","0","436.09","31.49","0"
"AGAW2","11E","INS","BOLD:AAV5575","Arthropoda","Insecta","Diptera","Cecidomyiidae","","","CecidInt30 sp. BOLD:AAV5575","1437","1009","1009","11","29.78","0","99.66","443.34","0","444.54","25.9","0"
"AGAW2","11E","INS","BOLD:ACE0554","Arthropoda","Insecta","Diptera","Phoridae","Metopininae","Megaselia","","1391","1132","1132","12","18.62","0","99.53","442.3","0","444.2","32.07","0"
"AGAW2","11E","INS","BOLD:AAU6681","Arthropoda","Insecta","Diptera","Tipulidae","Tipulinae","Nephrotoma","Nephrotoma cornicina","1210","948","948","13","21.65","0","99.67","442.97","0","444.06","36.32","0"
"AGAW2","11E","INS","BOLD:ACU7906","Arthropoda","Insecta","Diptera","Chironomidae","Orthocladiinae","","","1128","1122","1122","14","0.53","0","98.88","405.57","0","410.47","36.13","0"
"AGAW2","11E","INS","BOLD:AAH3661","Arthropoda","Insecta","Diptera","Cecidomyiidae","","","","806","714","714","15","11.41","0","99.33","430.72","0","432.37","25.36","0"
"AGAW2","11E","INS","BOLD:ABA0822","Arthropoda","Insecta","Diptera","Cecidomyiidae","","","","772","689","689","16","10.75","0","98.96","425.86","0","427.07","25.99","0"
"AGAW2","11E","INS","BOLD:AAN5199","Arthropoda","Insecta","Diptera","Cecidomyiidae","","","CecidInt20 sp. BOLD:AAN5199","699","520","520","17","25.61","0","99.64","442.37","0","444.79","26.52","0"
"AGAW2","11E","INS","BOLD:AAN5242","Arthropoda","Insecta","Diptera","Cecidomyiidae","","","","573","563","563","18","1.75","0","98.82","404.64","0","407.12","28.18","0"
"AGAW2","11E","INS","BOLD:AAU7126","Arthropoda","Insecta","Coleoptera","Cleridae","Hydnocerinae","Isohydnocera","Isohydnocera curtipennis","442","347","347","19","21.49","0","99.53","439.46","0","440.78","33","0"
"AGAW2","11E","INS","BOLD:ABW7897","Arthropoda","Insecta","Diptera","Cecidomyiidae","","","","385","277","277","20","28.05","0","99.71","444.2","0","445.48","29.7","0"
"AGAW2","11E","INS","BOLD:ACH4728","Arthropoda","Insecta","Psocodea","Elipsocidae","Elipsocinae","Cuneopalpus","Cuneopalpus cyanops","358","313","313","21","12.57","0","99.43","422.73","0","425.54","38.84","0"
"AGAW2","11E","INS","BOLD:ADF6661","Arthropoda","Insecta","Diptera","Cecidomyiidae","","","","333","268","268","22","19.52","0","99.24","415.78","0","441.98","29.78","0"
"AGAW2","11E","INS","BOLD:ACX5521","Arthropoda","Insecta","Diptera","Chironomidae","Orthocladiinae","","","304","296","296","23","2.63","0","98.94","407.09","0","409.02","35.03","0"
"AGAW2","11E","INS","BOLD:ACX6055","Arthropoda","Insecta","Diptera","Phoridae","Metopininae","Megaselia","Megaselia sp. AK-2010-2, Megaselia sp. BOLD:ACX6055","230","196","196","24","14.78","0","99.57","438.37","0","439.94","33.17","0"
"AGAW2","11E","INS","BOLD:AAB8143","Arthropoda","Insecta","Hemiptera","Nabidae","Nabinae","Nabis","Nabis inscriptus, Nabis americoferus, Nabis alternatus, Nabis sp.","228","195","195","25","14.47","0","99.19","433.66","0","435.25","36.91","0"
"AGAW2","11E","INS","BOLD:ACG5862","Arthropoda","Insecta","Diptera","Agromyzidae","","","","226","219","219","26","3.1","0","98.71","438.67","0","440.89","34.03","0"
"AGAW2","11E","INS","BOLD:AAD2144","Arthropoda","Insecta","Diptera","Chironomidae","Chironominae","Tanytarsus","Tanytarsus wirthi","219","201","201","27","8.22","0","99.35","434.8","0","437","36.19","0"
"AGAW2","11E","INS","BOLD:AAF9771","Arthropoda","Insecta","Diptera","Hybotidae","Tachydromiinae","Platypalpus","Platypalpus longiseta, Platypalpus trivialis, Platypalpus sp. 5, Platypalpus sp. 17, Platypalpus pallidiventris","209","154","154","28","26.32","0","99.04","446.74","0","447.41","35.06","0"
"AGAW2","11E","INS","BOLD:AAG7891","Arthropoda","Insecta","Hymenoptera","Platygastridae","Scelioninae","Telenomus","Telenomus podisi","167","153","153","29","8.38","0","99.13","430.8","0","439.32","27.01","0"
"AGAW2","11E","INS","BOLD:AAV5853","Arthropoda","Insecta","Diptera","Chironomidae","Tanypodinae","","","158","156","156","30","1.27","0","99.35","391.78","0","393.78","35.58","0"
"AGAW2","11E","INS","BOLD:AAL4232","Arthropoda","Insecta","Diptera","Agromyzidae","Phytomyzinae","Phytoliriomyza","Phytoliriomyza arctica","157","124","124","31","21.02","0","99.62","443.12","0","444.35","32.11","0"
"AGAW2","11E","INS","BOLD:ABA1661","Arthropoda","Insecta","Coleoptera","Chrysomelidae","Galerucinae","Longitarsus","Longitarsus pellucidus","154","132","132","32","14.29","0","99.46","425.54","0","426.7","35.63","0"
"AGAW2","11E","INS","BOLD:AAM9013","Arthropoda","Insecta","Diptera","Keroplatidae","Macrocerinae","Macrocera","Macrocera pusilla","153","141","141","33","7.84","0","99.38","431.67","0","433.1","30.31","0"
"AGAW2","11E","INS","BOLD:AAN5183","Arthropoda","Insecta","Diptera","Cecidomyiidae","","","","149","137","137","34","8.05","0","98.43","440.15","0","441.4","27.8","0"
"AGAW2","11E","INS","BOLD:AAH1693","Arthropoda","Insecta","Hymenoptera","Ichneumonidae","Cryptinae","Cryptus","Cryptus albitarsis","124","121","121","35","2.42","0","98.52","415.3","0","416.85","28.15","0"
"AGAW2","11E","INS","BOLD:AAH0256","Arthropoda","Insecta","Coleoptera","Latridiidae","Corticariinae","Corticaria, Corticarina","","121","121","121","36","0","0","98.35","413.85","0","421.19","31.33","0"
"AGAW2","11E","INS","BOLD:AAA4781","Arthropoda","Insecta","Hymenoptera","Braconidae","Microgastrinae","Glyptapanteles","Glyptapanteles carlossarmientoi","116","108","108","37","6.9","0","99.16","431.53","0","434.48","24.19","0"
"AGAW2","11E","INS","BOLD:ABA0863","Arthropoda","Insecta","Diptera","Cecidomyiidae","","","","114","97","97","38","14.91","0","99.46","438.72","0","440","29.66","0"
"AGAW2","11E","INS","BOLD:AAG2740","Arthropoda","Insecta","Diptera","Ephydridae","Parydrinae","Philygria","Philygria debilis","107","105","105","39","1.87","0","97.91","407.73","0","410.39","33.36","0"
"AGAW2","11E","INS","BOLD:AAG6471","Arthropoda","Insecta","Diptera","Ceratopogonidae","Dasyheleinae","Dasyhelea","","104","97","97","40","6.73","0","99.08","416.63","0","418.94","34.08","0"
"AGAW2","11E","INS","BOLD:ACK3431","Arthropoda","Insecta","Diptera","Cecidomyiidae","","","","100","93","93","41","7","0","99.2","430.38","0","431.88","32.74","0"
"AGAW2","11E","INS","BOLD:ABW8023","Arthropoda","Insecta","Diptera","Cecidomyiidae","","","","87","71","71","42","18.39","0","98.17","439.54","0","440.47","27.24","0"
"AGAW2","11E","INS","BOLD:ACC8079","Arthropoda","Insecta","Diptera","Cecidomyiidae","","","","85","66","66","43","22.35","0","99.53","448.6","0","449.99","29.21","0"
"AGAW2","11E","INS","BOLD:AAA5953","Arthropoda","Insecta","Lepidoptera","Gelechiidae","Gelechiinae","Coleotechnites","Coleotechnites sp., Coleotechnites piceaella, Coleotechnites laricis, Coleotechnites sp. 3, Coleotechnites carbonaria, Coleotechnites granti, Coleotechnites macleodi, Coleotechnites gibsonella, Coleotechnites coniferella, Coleotechnites martini","81","67","67","44","17.28","0","99.5","443.38","0","444.83","34.48","0"
"AGAW2","11E","INS","BOLD:AAH4171","Arthropoda","Insecta","Diptera","Chloropidae","Oscinellinae","Apallates","Apallates montanus, Apallates particeps","80","74","74","45","7.5","0","99.27","433.55","0","435.41","32.65","0"
"AGAW2","11E","INS","BOLD:AAN5205","Arthropoda","Insecta","Diptera","Cecidomyiidae","","","CecidInt42 sp. BOLD:AAN5205","79","69","69","46","12.66","0","99.53","440.81","0","442.2","31.33","0"
"AGAW2","11E","INS","BOLD:ABA0782","Arthropoda","Insecta","Diptera","Chloropidae","Oscinellinae","Gaurax","","79","68","68","46","13.92","0","99.46","427.65","0","428.63","33.3","0"
"AGAW2","11E","INS","BOLD:ACE9668","Arthropoda","Insecta","Hymenoptera","Ichneumonidae","Ophioninae","Enicospilus","Enicospilus purgatusDHJ02","76","73","73","48","3.95","0","98.97","440.75","0","441.8","25.8","0"
"AGAW2","11E","INS","BOLD:AAG8091","Arthropoda","Insecta","Hymenoptera","Ichneumonidae","Campopleginae","Campoletis","Campoletis sp. AB.2","61","60","60","49","1.64","0","99.04","434.43","0","436.49","24.63","0"
"AGAW2","11E","INS","BOLD:AAC4403","Arthropoda","Insecta","Diptera","Chironomidae","Chironominae","Micropsectra","Micropsectra polita","59","59","59","50","0","0","98.95","410.98","0","413.15","34.6","0"
"AGAW2","11E","INS","BOLD:ABW1379","Arthropoda","Insecta","Diptera","Chloropidae","Oscinellinae","Malloewia","Malloewia abdominalis","59","52","52","50","11.86","0","98.36","446.03","0","447.86","31.53","0"
"AGAW2","11E","INS","BOLD:ACM5699","Arthropoda","Insecta","Diptera","Cecidomyiidae","","","","57","48","48","52","15.79","0","97.76","445.28","0","446.21","26.09","0"
"AGAW2","11E","INS","BOLD:ADF7493","Arthropoda","Insecta","Diptera","Phoridae","Metopininae","Megaselia","","54","53","53","53","1.85","0","99.5","388.54","0","420.67","31.94","0"
"AGAW2","11E","INS","BOLD:AAG2868","Arthropoda","Insecta","Hemiptera","Cicadellidae","Typhlocybinae","Empoasca","Empoasca sp., Empoasca fabae","48","38","38","54","20.83","0","99.71","437.56","0","441.98","32.42","0"
"AGAW2","11E","INS","BOLD:ACK5742","Arthropoda","Insecta","Diptera","Cecidomyiidae","","","","46","41","41","55","10.87","0","99.6","442.13","0","442.98","29.02","0"
"AGAW2","11E","INS","BOLD:AAN6447","Arthropoda","Insecta","Diptera","Sciaridae","","Corynoptera","Corynoptera perpusilla","41","35","35","56","14.63","0","99.39","447.54","0","448.59","36.41","0"
"AGAW2","11E","INS","BOLD:AAH0134","Arthropoda","Insecta","Coleoptera","Phalacridae","Phalacrinae","Stilbus","Stilbus apicialis","40","36","36","57","10","0","99.37","423.2","0","424.23","31.9","0"
"AGAW2","11E","INS","BOLD:ABZ5071","Arthropoda","Insecta","Diptera","Chironomidae","Chironominae","Micropsectra","Micropsectra nigripila","39","39","39","58","0","0","98.27","425.46","0","441","36.84","0"
"AGAW2","11E","INS","BOLD:ABU5525","Arthropoda","Insecta","Diptera","Chironomidae","Orthocladiinae","Limnophyes","Limnophyes sp. 14ES","38","38","38","59","0","0","99.04","399.24","0","402.71","37.27","0"
"AGAW2","11E","INS","BOLD:AAA7374","Arthropoda","Insecta","Diptera","Syrphidae","Syrphinae","Sphaerophoria","Sphaerophoria bankowskae, Sphaerophoria infuscata, Sphaerophoria nr. philanthus, Sphaerophoria batava, Sphaerophoria sp., Sphaerophoria rueppellii, Sphaerophoria interrupta-group, Sphaerophoria sp. MA-1, Sphaerophoria cleoae, Sphaerophoria asymmetrica, Sphaerophoria fatarum, Sphaerophoria bengalensis, Sphaerophoria bifurcata, Sphaerophoria taeniata, Sphaerophoria interrupta, Sphaerophoria contigua, Sphaerophoria kaa, Sphaerophoria laurae, Sphaerophoria chongjini, Sphaerophoria boreoalpina, Sphaerophoria sp. BOLDAAA7374, Sphaerophoria macrogaster, Sphaerophoria sulphuripes, Sphaerophoria brevipilosa, Sphaerophoria philanthus, Sphaerophoria pyrrhina, Sphaerophoria abbreviata, Sphaerophoria scripta, Sphaerophoria potentillae, Sphaerophoria longipilosa, Sphaerophoria virgata","37","35","35","60","5.41","0","99.13","430.97","0","432.78","30.95","0"
"AGAW2","11E","INS","BOLD:AAG3275","Arthropoda","Insecta","Diptera","Phoridae","Metopininae","Megaselia","Megaselia minuta, Megaselia nubilipennis","36","34","34","61","5.56","0","99.14","428.58","0","430.83","34.76","0"
"AGAW2","11E","INS","BOLD:ABA7871","Arthropoda","Insecta","Diptera","Cecidomyiidae","","","","35","33","33","62","5.71","0","98.88","442.89","0","444.03","26.89","0"
"AGAW2","11E","INS","BOLD:AAN5238","Arthropoda","Insecta","Diptera","Cecidomyiidae","","","","33","28","28","63","15.15","0","99.13","436.3","0","441.27","25.92","0"
"AGAW2","11E","INS","BOLD:AAN8770","Arthropoda","Insecta","Diptera","Psychodidae","Psychodinae","Psychoda","Psychoda trinodulosa","31","31","31","64","0","0","99.7","442.29","0","443.42","33.88","0"
"AGAW2","11E","INS","BOLD:AAD1485","Arthropoda","Insecta","Diptera","Chironomidae","Chironominae","Paratanytarsus","Paratanytarsus grimmii, Paratanytarsus sp.","30","28","28","65","6.67","0","99.23","438.4","0","439.57","33.82","0"
"AGAW2","11E","INS","BOLD:AAN5220","Arthropoda","Insecta","Diptera","Cecidomyiidae","","","","21","21","21","66","0","0","98.09","403.24","0","420.43","26.33","0"
"AGAW2","11E","INS","BOLD:AAV5594","Arthropoda","Insecta","Diptera","Cecidomyiidae","","","","20","16","16","67","20","0","99.33","444.05","0","445.4","26.17","0"
"AGAW2","11E","INS","BOLD:ACF4786","Arthropoda","Insecta","Hymenoptera","Braconidae","Aphidiinae","Aphidius","Aphidius urticae, Aphidius matricariae","17","17","17","68","0","0","99.2","393.82","0","396.76","25.85","0"
"AGAW2","11E","INS","BOLD:AAH0135","Arthropoda","Insecta","Coleoptera","Phalacridae","Phalacrinae","Acylomus","Acylomus pugetanus, Acylomus ergoti","17","16","16","68","5.88","0","99.36","429.65","0","431.29","31.59","0"
"AGAW2","11E","INS","BOLD:AAP7026","Arthropoda","Insecta","Coleoptera","Latridiidae","Corticariinae","Melanophthalma","Melanophthalma villosa","16","14","14","70","12.5","0","97.94","406.38","0","442.69","30.91","0"
"AGAW2","11E","INS","BOLD:AAU8379","Arthropoda","Insecta","Hymenoptera","Braconidae","Alysiinae","Dinotrema","Dinotrema JFT01","16","16","16","70","0","0","99.53","439.31","0","442","24.23","0"
"AGAW2","11E","INS","BOLD:AAU6595","Arthropoda","Insecta","Diptera","Sciaridae","","Corynoptera","Corynoptera perpusilla","15","15","15","72","0","0","98.96","398.47","0","400.93","35.32","0"
"AGAW2","11E","INS","BOLD:AAN5255","Arthropoda","Insecta","Diptera","Cecidomyiidae","Cecidomyiinae","Dasineura","","15","15","15","72","0","0","98.85","433.4","0","447.87","26.89","0"
"AGAW2","11E","INS","BOLD:AAJ1250","Arthropoda","Insecta","Hymenoptera","Braconidae","Aphidiinae","Aphidius","Aphidius avenaphis, Aphidius uzbekistanicus","13","13","13","74","0","0","99.61","429.08","0","434.69","26.69","0"
"AGAW2","11E","INS","BOLD:ABV1178","Arthropoda","Insecta","Diptera","Sphaeroceridae","Limosininae","Leptocera, Rachispoda","Rachispoda limosa, Leptocera limosa","13","13","13","74","0","0","99.48","439.38","0","442.31","31.38","0"
"AGAW2","11E","INS","BOLD:ACE4233","Arthropoda","Insecta","Hymenoptera","Ichneumonidae","Orthocentrinae","Megastylus","Megastylus jft01","13","13","13","74","0","0","98.95","429.85","0","431","27.73","0"
"AGAW2","11E","INS","BOLD:ACO4990","Arthropoda","Insecta","Diptera","Cecidomyiidae","","","","12","12","12","77","0","0","98.74","362.5","0","388.75","25.26","0"
"AGAW2","11E","INS","BOLD:ABV1194","Arthropoda","Insecta","Diptera","Chironomidae","Chironominae","Micropsectra","","12","12","12","77","0","0","99.47","427.92","0","429.67","36.09","0"
"AGAW2","11E","INS","BOLD:ACV1014","Arthropoda","Insecta","Diptera","Cecidomyiidae","","","","11","11","11","79","0","0","97.87","369.09","0","371.09","25.27","0"
"AGAW2","11E","INS","BOLD:AAM6257","Arthropoda","Insecta","Diptera","Chironomidae","Orthocladiinae","","","11","11","11","79","0","0","98.7","435.27","0","436.73","35.47","0"
"AGAW2","11E","INS","BOLD:ADF3937","","","","","","","","11","11","11","79","0","0","98.28","403.91","0","408.55","31.32","0"
"AGAW2","11E","INS","BOLD:ACO5153","Arthropoda","Insecta","Diptera","Cecidomyiidae","","","","11","11","11","79","0","0","97.88","431.82","0","442.55","26.24","0"
"AGAW2","11E","INS","BOLD:AAV5688","Arthropoda","Insecta","Diptera","Cecidomyiidae","","","","10","10","10","83","0","0","99.47","437.6","0","439.1","34.72","0"
"AGAW2","11E","INS","BOLD:ACV2677","Arthropoda","Insecta","Diptera","Cecidomyiidae","","","","10","10","10","83","0","0","97.44","449.2","0","450","30.11","0"
"AGAW2","11E","INS","BOLD:AAA8469","Arthropoda","Insecta","Hymenoptera","Braconidae","Euphorinae","Peristenus","Peristenus sp.","9","9","9","85","0","0","99.82","421.22","0","447","28.73","0"
"AGAW2","11E","INS","BOLD:AAG6745","Arthropoda","Insecta","Diptera","Calliphoridae","Calliphorinae","Pollenia","Pollenia pediculata","9","8","8","85","11.11","0","99.57","440.33","0","442.33","33.6","0"
"AGAW2","11E","INS","BOLD:AAH3664","Arthropoda","Insecta","Diptera","Cecidomyiidae","","","CecidInt3 sp. BOLD:AAH3664","9","9","9","85","0","0","98.83","430","0","431.89","26.96","0"
"AGAW2","11E","INS","BOLD:AAA2305","Arthropoda","Insecta","Trichoptera","Philopotamidae","Philopotaminae","Dolophilodes","Dolophilodes sp. BOLDAAA2305","9","9","9","85","0","0","98.18","437.44","0","437.78","36.15","0"
"AGAW2","11E","INS","BOLD:ABX8236","Arthropoda","Insecta","Diptera","Cecidomyiidae","","","","9","9","9","85","0","0","99.27","418.11","0","420","28.91","0"
"AGAW2","11E","INS","BOLD:AAO8538","Arthropoda","Insecta","Diptera","Psychodidae","Psychodinae","Psychoda","","8","8","8","90","0","0","99.17","434.63","0","435","32.54","0"
"AGAW2","11E","INS","BOLD:ABV1337","Arthropoda","Insecta","Diptera","Cecidomyiidae","","","","7","7","7","91","0","0","98.95","449.14","0","450","28.25","0"
"AGAW2","11E","INS","BOLD:AAM6248","Arthropoda","Insecta","Diptera","Chironomidae","Chironominae","Microtendipes","Microtendipes pedellus grp.","7","5","5","91","28.57","0","99.38","424.71","0","426.29","31.33","0"
"AGAW2","11E","INS","BOLD:AAA4222","Arthropoda","Insecta","Lepidoptera","Erebidae","Hypeninae","Hypena","Hypena scabra","6","6","6","93","0","0","98.73","381","0","383.33","32.4","0"
"AGAW2","11E","INS","BOLD:ACK3142","Arthropoda","Insecta","Diptera","Cecidomyiidae","","","","6","6","6","93","0","0","98.52","403.5","0","405","27.54","0"
"AGAW2","11E","INS","BOLD:ABA0929","Arthropoda","Insecta","Diptera","Sciaridae","","Bradysia","Bradysia scabricornis","6","6","6","93","0","0","99.03","426.33","0","429.33","29.94","0"
"AGAW2","11E","INS","BOLD:AAG2123","Arthropoda","Insecta","Diptera","Tachinidae","Dexiinae","Dinera","Dinera grisescens","5","5","5","96","0","0","99.46","435.8","0","436.8","31.93","0"
"AGAW2","11E","INS","BOLD:AAN5249","Arthropoda","Insecta","Diptera","Cecidomyiidae","","","","4","4","4","97","0","0","98.5","425.25","0","428","33.12","0"
"AGAW2","11E","INS","BOLD:AAG3302","Arthropoda","Insecta","Diptera","Phoridae","Metopininae","Megaselia","","4","4","4","97","0","0","98.92","406.5","0","409.75","31.27","0"
"AGAW2","11E","INS","BOLD:ACU6062","Arthropoda","Insecta","Diptera","Cecidomyiidae","","","","4","4","4","97","0","0","98.71","446.25","0","450","26.61","0"
"AGAW2","11E","INS","BOLD:AAC0903","Arthropoda","Insecta","Diptera","Chironomidae","Chironominae","Chironomus","Chironomus acidophilus","4","4","4","97","0","0","99.52","432.25","0","435","35.73","0"
"AGAW2","11E","INS","BOLD:AAG2511","Arthropoda","Insecta","Diptera","Anthomyiidae","Anthomyiinae","Delia","Delia platura","4","4","4","97","0","0","99.22","446.75","0","450","33","0"
"AGAW2","11E","INS","BOLD:AAP8739","Arthropoda","Insecta","Diptera","Platystomatidae","Platystomatinae","Rivellia","Rivellia steyskali","4","4","4","97","0","0","99.55","446.75","0","449","38.36","0"
"AGAW2","11E","INS","BOLD:AAB7361","Arthropoda","Insecta","Diptera","Chironomidae","Orthocladiinae","Limnophyes","Limnophyes natalensis","3","3","3","103","0","0","98.25","351.33","0","378.67","38.13","0"
"AGAW2","11E","INS","BOLD:AAA4188","Arthropoda","Insecta","Hymenoptera","Braconidae","Aphidiinae","Aphidius","Aphidius microlophii, Aphidius ervi","3","3","3","103","0","0","99.38","431.67","0","435.67","25.78","0"
"AGAW2","11E","INS","BOLD:ACK1456","Arthropoda","Insecta","Diptera","Cecidomyiidae","","","","3","3","3","103","0","0","99.13","423.67","0","431.33","25.17","0"
"AGAW2","11E","INS","BOLD:AAF9306","Arthropoda","Insecta","Diptera","Psychodidae","Psychodinae","Psychoda","Psychoda alternata","3","3","3","103","0","0","97.88","430.67","0","431.33","31.04","0"
"AGAW2","11E","INS","BOLD:ACD8731","Arthropoda","Insecta","Diptera","Phoridae","Phorinae","Borophaga","Borophaga subsultans","3","3","3","103","0","0","99.76","427","0","428","33.16","0"
"AGAW2","11E","INS","BOLD:ABX2489","Arthropoda","Insecta","Coleoptera","Phalacridae","Phalacrinae","Stilbus","","3","3","3","103","0","0","97.97","429.33","0","431.67","33.03","0"
"AGAW2","11E","INS","BOLD:ACF4925","Arthropoda","Insecta","Hemiptera","Miridae","Mirinae","Adelphocoris","Adelphocoris detritus","3","3","3","103","0","0","97.92","449.33","0","450","35.56","0"
"AGAW2","11E","INS","BOLD:AAA3933","Arthropoda","Insecta","Lepidoptera","Erebidae","Herminiinae","Palthis","Palthis angulalis","2","2","2","110","0","0","99.21","381.5","0","383.5","31.81","0"
"AGAW2","11E","INS","BOLD:ACJ6954","Arthropoda","Insecta","Diptera","Cecidomyiidae","","","","2","2","2","110","0","0","100","339","0","442.5","26.9","0"
"AGAW2","11E","INS","BOLD:AAG5507","Arthropoda","Insecta","Diptera","Chironomidae","Chironominae","Chironomus","Chironomus atrella","2","2","2","110","0","0","98.43","446","0","450","35.67","0"
"AGAW2","11E","INS","BOLD:AAM6781","Arthropoda","Insecta","Diptera","Dolichopodidae","Neurigoninae","Neurigona","Neurigona tenuis","2","2","2","110","0","0","98.27","402","0","403","35.6","0"
"AGAW2","11E","INS","BOLD:AAB4657","Arthropoda","Insecta","Diptera","Chironomidae","Chironominae","Chironomus","Chironomus maturus","2","2","2","110","0","0","98.24","422","0","435","37.57","0"
"AGAW2","11E","INS","BOLD:AAN5230","Arthropoda","Insecta","Diptera","Cecidomyiidae","","","","2","2","2","110","0","0","99.19","439","0","440.5","26.34","0"
"AGAW2","11E","INS","BOLD:AAN5201","Arthropoda","Insecta","Diptera","Cecidomyiidae","","","","2","2","2","110","0","0","99.78","449","0","450","27.89","0"
"AGAW2","11E","INS","BOLD:AAH2293","Arthropoda","Insecta","Diptera","Asilidae","Stenopogoninae","Dioctria, Eudioctria","Dioctria hyalipennis, Dioctria baumhaueri, Eudioctria propinqua","2","2","2","110","0","0","98.11","397","0","399","30.57","0"
"AGAW2","11E","INS","BOLD:ACE8878","Arthropoda","Insecta","Orthoptera","Acrididae","Melanoplinae","Melanoplus","Melanoplus scudderi group","2","2","2","110","0","0","97.67","450","0","450","35.67","0"
"AGAW2","11E","INS","BOLD:ABV1444","Arthropoda","Insecta","Diptera","Cecidomyiidae","","","","2","2","2","110","0","0","99.67","449","0","450","30.22","0"
"AGAW2","11E","INS","BOLD:AAV6112","Arthropoda","Insecta","Diptera","Chloropidae","Oscinellinae","Oscinella","Oscinella sp. 2 AGT, Oscinella sp. TAW6","2","2","2","110","0","0","98.4","438.5","0","439.5","34.13","0"
"AGAW2","11E","INS","BOLD:AAB0651","Arthropoda","Insecta","Lepidoptera","Gelechiidae","Dichomeridinae","Dichomeris","Dichomeris leuconotella, Dichomeris gnoma","2","2","2","110","0","0","98.88","448","0","450","33.33","0"
"AGAW2","11E","INS","BOLD:ABY5257","Arthropoda","Insecta","Lepidoptera","Noctuidae","Noctuinae","Apamea","Apamea devastator","2","2","2","110","0","0","99.22","448.5","0","450","30.33","0"
"AGAW2","11E","INS","BOLD:AAN5260","Arthropoda","Insecta","Diptera","Cecidomyiidae","","","","2","2","2","110","0","0","98.16","381.5","0","382","26.57","0"
"AGAW2","11E","INS","BOLD:ACX6289","Arthropoda","Insecta","Diptera","Phoridae","Metopininae","Megaselia","","2","2","2","110","0","0","99.6","373.5","0","374.5","37.92","0"
"AGAW2","11E","INS","BOLD:AAF1729","Arthropoda","Insecta","Psocodea","Lachesillidae","Lachesillinae","Lachesilla","Lachesilla pedicularia","2","2","2","110","0","0","99.44","450","0","450","37.44","0"
"AGAW2","11E","INS","BOLD:ABW3288","","","","","","","","2","2","2","110","0","0","99.02","403","0","403.5","29.57","0"
"AGAW2","11E","INS","BOLD:AAU8628","Arthropoda","Insecta","Hymenoptera","Bethylidae","","","","2","2","2","110","0","0","99.66","446.5","0","450","27.11","0"
"AGAW2","11E","INS","BOLD:ACE7482","Arthropoda","Insecta","Hemiptera","Miridae","","","","2","2","2","110","0","0","98.17","382.5","0","383","33.81","0"
"AGAW2","11E","INS","BOLD:ABV1422","Arthropoda","Insecta","Diptera","Cecidomyiidae","","","","2","2","2","110","0","0","99.29","419.5","0","420","26.77","0"
"AGAW2","11E","INS","BOLD:AAA3181","Arthropoda","Insecta","Diptera","Muscidae","Muscinae","Stomoxys","Stomoxys calcitrans","1","1","1","130","0","0","99.09","441","0","444","33.33","0"
"AGAW2","11E","INS","BOLD:ADB1634","Arthropoda","Insecta","Hymenoptera","Ceraphronidae","","","","1","1","1","130","0","0","100","338","0","450","30.22","0"
"AGAW2","11E","INS","BOLD:AAN5161","Arthropoda","Insecta","Diptera","Ceratopogonidae","Dasyheleinae","Dasyhelea","","1","1","1","130","0","0","97.87","376","0","377","37.4","0"
"AGAW2","11E","INS","BOLD:AAA3836","Arthropoda","Insecta","Lepidoptera","Noctuidae","Plusiinae","Autographa","Autographa precationis","1","1","1","130","0","0","98.44","448","0","450","32.22","0"
"AGAW2","11E","INS","BOLD:AAF9028","Arthropoda","Insecta","Diptera","Limoniidae","","","","1","1","1","130","0","0","100","449","0","450","38.67","0"
"AGAW2","11E","INS","BOLD:AAF3338","Arthropoda","Insecta","Lepidoptera","Noctuidae","Noctuinae","Striacosta","Striacosta albicosta","1","1","1","130","0","0","98.66","447","0","450","34.22","0"
"AGAW2","11E","INS","BOLD:AAA6560","Arthropoda","Insecta","Lepidoptera","Crambidae","Crambinae","Crambus","Crambus multilinellus, Crambus saltuellus","1","1","1","130","0","0","98.62","434","0","435","31.49","0"
"AGAW2","11E","INS","BOLD:AAY6730","Arthropoda","Insecta","Hemiptera","Derbidae","Cedusinae","Cedusa","Cedusa incisa","1","1","1","130","0","0","97.33","450","0","450","35.33","0"
"AGAW2","11E","INS","BOLD:AAA2518","Arthropoda","Insecta","Lepidoptera","Crambidae","Crambinae","Chrysoteuchia","Chrysoteuchia topiaria, Chrysoteuchia topiarius","1","1","1","130","0","0","98","450","0","450","35.11","0"
"AGAW2","11E","INS","BOLD:ABY8248","Arthropoda","Insecta","Diptera","Cecidomyiidae","","","","1","1","1","130","0","0","97.91","430","0","450","26","0"
"AGAW2","11E","INS","BOLD:ABA0858","Arthropoda","Insecta","Diptera","Cecidomyiidae","","","","1","1","1","130","0","0","100","416","0","416","25","0"
"AGAW2","11E","INS","BOLD:AAB2076","Arthropoda","Insecta","Lepidoptera","Tortricidae","Olethreutinae","Ancylis","Ancylis metamelana, Ancylis sp.","1","1","1","130","0","0","99.78","450","0","450","31.33","0"
"AGAW2","11E","INS","BOLD:AAH4135","Arthropoda","Insecta","Diptera","Chloropidae","Chloropinae","Thaumatomyia","Thaumatomyia glabra, Thaumatomyia grata, Thaumatomyia sp. 1 TAW","1","1","1","130","0","0","99.1","442","0","450","32.89","0"
"AGAW2","11E","INS","BOLD:ACZ0513","Arthropoda","Insecta","Coleoptera","Chrysomelidae","Galerucinae","Longitarsus","Longitarsus pellucidus","1","1","1","130","0","0","97.32","447","0","450","34","0"
"AGAW2","11E","INS","BOLD:AAV5098","Arthropoda","Insecta","Diptera","Ceratopogonidae","Dasyheleinae","Dasyhelea","Dasyhelea turficola, Dasyhelea sp. 5ES","1","1","1","130","0","0","99.33","449","0","450","36.89","0"
"AGAW2","11E","INS","BOLD:AAC0022","Arthropoda","Insecta","Diptera","Chironomidae","Chironominae","Micropsectra","Micropsectra nigripila","1","1","1","130","0","0","97.77","448","0","450","36.67","0"
"AGAW2","11E","INS","BOLD:AAE7186","Arthropoda","Insecta","Hymenoptera","Braconidae","Aphidiinae","Praon","Praon sp. 1 MM-2013, Praon volucre, Praon bicolor","1","1","1","130","0","0","100","449","0","450","26","0"
"AGAW2","11E","INS","BOLD:AAG5453","Arthropoda","Insecta","Diptera","Chironomidae","Chironominae","Chironomus","Chironomus bifurcatus","1","1","1","130","0","0","97.75","445","0","450","34.67","0"
"AGAW2","11E","INS","BOLD:ACT3332","Arthropoda","Insecta","Diptera","Cecidomyiidae","","","","1","1","1","130","0","0","97.77","449","0","450","26.22","0"
"AGAW2","11E","INS","BOLD:AAH3225","Arthropoda","Insecta","Psocodea","Psocidae","Psocinae","Metylophorus","Metylophorus novaescotiae","1","1","1","130","0","0","99.51","412","0","415","32.77","0"
"AGAW2","11E","INS","BOLD:ACD3276","Arthropoda","Insecta","Diptera","Cecidomyiidae","","","","1","1","1","130","0","0","99.78","448","0","450","28","0"
"AGAW2","11E","INS","BOLD:AAZ5483","Arthropoda","Insecta","Diptera","Muscidae","Phaoniinae","Helina","Helina setiventris","1","1","1","130","0","0","97.77","448","0","450","30.22","0"
"AGAW2","11E","INS","BOLD:AAG1743","Arthropoda","Insecta","Diptera","Muscidae","Phaoniinae","Phaonia","Phaonia fuscana","1","1","1","130","0","0","98.06","360","0","361","33.52","0"
"AGAW2","11E","INS","BOLD:ACJ0111","Arthropoda","Insecta","Hymenoptera","Platygastridae","Scelioninae","Telenomus","","1","1","1","130","0","0","98.86","440","0","450","29.11","0"
"AGAW2","11E","INS","BOLD:AAN5215","Arthropoda","Insecta","Diptera","Cecidomyiidae","","","","1","1","1","130","0","0","99.78","446","0","450","29.11","0"
"AGAW2","11E","INS","BOLD:AAH2295","Arthropoda","Insecta","Diptera","Asilidae","Leptogastrinae","Leptogaster","Leptogaster flavipes","1","1","1","130","0","0","99.02","410","0","450","34.44","0"
"AGAW2","11E","INS","BOLD:AAA5206","Arthropoda","Insecta","Lepidoptera","Erebidae","Herminiinae","Zanclognatha","Zanclognatha cruralis, Zanclognatha sp. 1, Zanclognatha jacchusalis, Zanclognatha minoralis, Zanclognatha minoralis sp. 1, Zanclognatha sp., Zanclognatha minoralis sp. 3, Zanclognatha gypsalis, Zanclognatha dentata, Zanclognatha theralis, Zanclognatha protumnusalis, Zanclognatha martha, Zanclognatha atrilineella, Zanclognatha obscuripennis, Zanclognatha minoralis sp. 2, Zanclognatha marcidilinea","1","1","1","130","0","0","98.43","445","0","450","31.11","0"
"AGAW2","11E","INS","BOLD:AAL2750","Arthropoda","Insecta","Diptera","Muscidae","Phaoniinae","Phaonia","Phaonia tuguriorum","1","1","1","130","0","0","100","450","0","450","32.89","0"
"AGAW2","11E","INS","BOLD:ACF1257","","","","","","","","1","1","1","130","0","0","98.66","449","0","450","35.11","0"
"AGAW2","11E","INS","BOLD:AAN5341","Arthropoda","Insecta","Diptera","Chironomidae","Orthocladiinae","Camptocladius","Camptocladius stercorarius","1","1","1","130","0","0","97.95","391","0","393","35.37","0"
"AGAW2","11E","INS","BOLD:AAB1352","Arthropoda","Insecta","Lepidoptera","Erebidae","Hypeninae","Hypena","Hypena sp.","1","1","1","130","0","0","99.1","446","0","450","32.89","0"
"AGAW2","11E","INS","BOLD:AAI3260","Arthropoda","Insecta","Diptera","Mycetophilidae","Mycetophilinae","Mycetophila","Mycetophila caudata","1","1","1","130","0","0","99.33","450","0","450","33.11","0"
"AGAW2","11E","INS","BOLD:AAA3867","Arthropoda","Insecta","Lepidoptera","Noctuidae","Noctuinae","Apamea","Apamea amputatrix","1","1","1","130","0","0","99.78","449","0","450","33.78","0"
"AGAW2","11E","INS","BOLD:AAI4303","Arthropoda","Insecta","Diptera","Chironomidae","Chironominae","Chironomus","Chironomus melanescens","1","1","1","130","0","0","98.21","446","0","450","34.89","0"
"AGAW2","11E","INS","BOLD:ABA6407","Arthropoda","Insecta","Diptera","Sciaridae","","Pseudolycoriella","","1","1","1","130","0","0","99.33","446","0","450","34.67","0"
"AGAW2","11E","INS","BOLD:ABZ3705","Arthropoda","Insecta","Lepidoptera","Crambidae","Pyraustinae","Sitochroa","Sitochroa chortalis","1","1","1","130","0","0","100","450","0","450","30","0"
"AGAW2","11E","INS","BOLD:ADH9867","Arthropoda","Insecta","Diptera","","","","","1","1","1","130","0","0","98.88","448","0","450","38.22","0"
"AGAW2","11E","INS","BOLD:ACF7023","Arthropoda","Insecta","Diptera","Dolichopodidae","","","","1","1","1","130","0","0","97.8","227","0","450","34.44","0"
"AGAW2","11E","INS","BOLD:AAE0845","Arthropoda","Insecta","Hemiptera","Pentatomidae","Pentatominae","Euschistus","Euschistus servus","1","1","1","130","0","0","97.99","448","0","450","38.44","0"
"AGAW2","11E","INS","BOLD:ABZ4994","Arthropoda","Insecta","Orthoptera","Acrididae","Melanoplinae","Melanoplus","Melanoplus gladstoni, Melanoplus sanguinipes","1","1","1","130","0","0","97.11","450","0","450","36.67","0"
"AGAW2","11E","INS","BOLD:AAC4197","Arthropoda","Insecta","Diptera","Chironomidae","Orthocladiinae","Paraphaenocladius","Paraphaenocladius impensus","1","1","1","130","0","0","99.78","450","0","450","35.11","0"
"AGAW2","11E","INS","BOLD:AAA1513","Arthropoda","Insecta","Lepidoptera","Plutellidae","Plutellinae","Plutella, Rhigognostis","Plutella xylostella, Rhigognostis sp.","1","1","1","130","0","0","98.99","397","0","402","30.6","0"
"AGAW2","11E","INS","BOLD:AAA5803","Arthropoda","Insecta","Hemiptera","Miridae","Mirinae","Lygus","Lygus lineolaris","1","1","1","130","0","0","98","449","0","450","33.56","0"
"AGAW2","11E","NAR","BOLD:AAL1512","Mollusca","Gastropoda","Stylommatophora","Agriolimacidae",NA,"Deroceras","Deroceras laeve","10","10","10","1","0","0","97.73","388.9","0","392.3","32.77","0"
"AGAW2","11E","NAR","TAX:660402","Mollusca","Gastropoda","Stylommatophora","Agriolimacidae",NA,"Deroceras","Deroceras sp.","2","2","2","2","0","0","97.81","366.5","0","371","32.74","0"
"AGAW2","11E","NAR","BOLD:AAI9663","Mollusca","Gastropoda","Stylommatophora","Agriolimacidae",NA,"Deroceras","Deroceras reticulatum","1","1","1","3","0","0","100","407","0","450","36.44","0"
"AGAW2","11E","NIN","BOLD:AAH6630","Arthropoda","Collembola","Entomobryomorpha","Entomobryidae","Entomobryinae","Entomobrya","Entomobrya unostrigata","230","216","216","1","6.09","0","99.29","433.73","0","435.2","39.15","0"
"AGAW2","11E","NIN","BOLD:ACR6860","","","","","","","","165","152","152","2","7.88","0","99.09","385.37","0","439.75","40.39","0"
"AGAW2","11E","NIN","BOLD:ABW2639","Arthropoda","Arachnida","Trombidiformes","Anystidae","","","","100","91","91","3","9","0","99.5","445.54","0","445.42","31.37","0"
"AGAW2","11E","NIN","BOLD:AAA6381","Arthropoda","Arachnida","Araneae","Tetragnathidae","","Tetragnatha","Tetragnatha laboriosa","50","50","50","4","0","0","98.88","407.68","0","409.36","39.17","0"
"AGAW2","11E","NIN","BOLD:AAA8820","Arthropoda","Collembola","Entomobryomorpha","Entomobryidae","Lepidocyrtinae","Lepidocyrtus","Lepidocyrtus cyaneus","31","31","31","5","0","0","98.74","402.23","0","403.81","43.42","0"
"AGAW2","11E","NIN","BOLD:AAB7914","Arthropoda","Collembola","Symphypleona","Bourletiellidae","","Bourletiella","","20","17","17","6","15","0","99.77","449.4","0","450","38.2","0"
"AGAW2","11E","NIN","BOLD:ACZ1295","Arthropoda","Arachnida","Trombidiformes","Anystidae","","","","11","11","11","7","0","0","99.03","372.55","0","375.36","31.19","0"
"AGAW2","11E","NIN","BOLD:AAU6252","Arthropoda","Arachnida","Trombidiformes","Bdellidae","","Cyta","","6","6","6","8","0","0","98.33","431.83","0","433.83","31.45","0"
"AGAW2","11E","NIN","BOLD:ACI9974","Arthropoda","Arachnida","Trombidiformes","Tydeidae","","","","5","5","5","9","0","0","99.86","425.4","0","450","41.11","0"
"AGAW2","11E","NIN","BOLD:ACV5610","","","","","","","","4","3","3","10","25","0","99.28","448.25","0","450","38.22","0"
"AGAW2","11E","NIN","BOLD:AAI4346","Arthropoda","Arachnida","Opiliones","Phalangiidae","","Paroligolophus, Phalangium","Paroligolophus agrestis, Phalangium opilio","1","1","1","11","0","0","100","450","0","450","36.89","0"
"AGAW2","11E","NIN","BOLD:AAO8147","Arthropoda","Arachnida","Trombidiformes","Eupodidae","","Eupodes","","1","1","1","11","0","0","99.33","446","0","450","28.67","0"
"AGAW2","11E","NIN","BOLD:ACI9748","Arthropoda","Arachnida","Araneae","Salticidae","","Dendryphantes","Dendryphantes sp. 1GAB","1","1","1","11","0","0","99.47","377","0","427","31.15","0"
"AGAW2","11N","BAC","TAX:91391","Proteobacteria","Alphaproteobacteria","Rickettsiales","",NA,"","","4","4","4","1","0","0","99.31","398.75","0","414.25","39.89","0"
"AGAW2","11N","BAC","TAX:392874","Proteobacteria","Alphaproteobacteria","Rickettsiales","Rickettsiaceae",NA,"Wolbachia","Wolbachia endosymbiont of Hymenoptera","3","3","3","2","0","0","98.81","450","0","450","40.44","0"
"AGAW2","11N","BAC","TAX:91393","Proteobacteria","Alphaproteobacteria","Rickettsiales","Rickettsiaceae",NA,"Wolbachia","","2","2","2","3","0","0","99.62","392.5","0","422","39.55","0"
"AGAW2","11N","BAC","TAX:375845","Proteobacteria","Alphaproteobacteria","Rickettsiales","Rickettsiaceae",NA,"Wolbachia","Wolbachia sp. Hymenoptera_ASGLE1440-10","1","1","1","4","0","0","99.33","446","0","450","40.22","0"
"AGAW2","11N","BAC","TAX:376688","Proteobacteria","Alphaproteobacteria","Rickettsiales","Rickettsiaceae",NA,"Wolbachia","Wolbachia sp. elachBioLep01 BioLep373","1","1","1","4","0","0","99.11","449","0","450","40.44","0"
"AGAW2","11N","BAC","TAX:392873","Proteobacteria","Alphaproteobacteria","Rickettsiales","Rickettsiaceae",NA,"Wolbachia","Wolbachia endosymbiont of Scrapter algoensis","1","1","1","4","0","0","98.3","412","0","415","39.28","0"
"AGAW2","11N","BAC","TAX:175768","Proteobacteria","Alphaproteobacteria","Rickettsiales","Rickettsiaceae",NA,"Wolbachia","Wolbachia pipientis","1","1","1","4","0","0","98.22","449","0","450","39.56","0"
"AGAW2","11N","CHO","no match","no match","no match","no match","no match","no match","no match","no match","no match","no match","no match","no match","no match","no match","no match","no match","no match","no match","no match","no match"
"AGAW2","11N","INS","BOLD:AAG1711","Arthropoda","Insecta","Diptera","Muscidae","Phaoniinae","Helina","Helina depuncta","13894","10923","10923","1","21.38","0","99.46","433.59","0","435.93","31.34","0"
"AGAW2","11N","INS","BOLD:AAG8882","Arthropoda","Insecta","Hemiptera","Pentatomidae","Pentatominae","Thyanta","Thyanta pallidovirens, Thyanta custator, Thyanta accerra","11020","6005","6005","2","45.51","0","99.48","444.2","0","445.73","36.2","0"
"AGAW2","11N","INS","BOLD:AAG2123","Arthropoda","Insecta","Diptera","Tachinidae","Dexiinae","Dinera","Dinera grisescens","8021","5655","5655","3","29.5","0","99.41","432.05","0","433.82","31.83","0"
"AGAW2","11N","INS","BOLD:AAP8739","Arthropoda","Insecta","Diptera","Platystomatidae","Platystomatinae","Rivellia","Rivellia steyskali","4494","3000","3000","4","33.24","0","99.33","440.6","0","442.2","37.98","0"
"AGAW2","11N","INS","BOLD:AAU6681","Arthropoda","Insecta","Diptera","Tipulidae","Tipulinae","Nephrotoma","Nephrotoma cornicina","3902","2942","2942","5","24.6","0","99.65","442.13","0","443.36","36.31","0"
"AGAW2","11N","INS","BOLD:AAK5928","Arthropoda","Insecta","Coleoptera","Chrysomelidae","Criocerinae","Oulema","Oulema melanopus","3253","2536","2536","6","22.04","0","99.53","440.52","0","441.86","36.39","0"
"AGAW2","11N","INS","BOLD:AAA3453","Arthropoda","Insecta","Diptera","Anthomyiidae","Anthomyiinae","Delia","","3191","2846","2846","7","10.81","0","99.41","428.75","0","431.46","34.62","0"
"AGAW2","11N","INS","BOLD:AAA4555","Arthropoda","Insecta","Orthoptera","Acrididae","Melanoplinae","Melanoplus","Melanoplus stupefactus, Melanoplus angustipennis, Melanoplus ponderosus, Melanoplus bruneri, Melanoplus frigidus, Melanoplus scapularis, Melanoplus cinereus, Melanoplus madeleineae, Melanoplus sanguinipes, Melanoplus nigrescens group, Melanoplus tunicae, Melanoplus walshi, Melanoplus femurrubrum, Melanoplus dawsoni, Melanoplus sp., Melanoplus fasciatus, Melanoplus bowditchi, Melanoplus stonei, Melanoplus bivittatus, Melanoplus huroni, Melanoplus thomasi, Melanoplus borealis, Melanoplus packardii","2871","2222","2222","8","22.61","0","99.41","445.95","0","447.2","36.19","0"
"AGAW2","11N","INS","BOLD:AAF3338","Arthropoda","Insecta","Lepidoptera","Noctuidae","Noctuinae","Striacosta","Striacosta albicosta","1928","1562","1562","9","18.98","0","98.94","439","0","440.52","34.44","0"
"AGAW2","11N","INS","BOLD:AAN6154","Arthropoda","Insecta","Coleoptera","Latridiidae","Corticariinae","Melanophthalma","Melanophthalma inermis","1928","1595","1595","9","17.27","0","99.14","435.75","0","437.28","31.17","0"
"AGAW2","11N","INS","BOLD:AAG6745","Arthropoda","Insecta","Diptera","Calliphoridae","Calliphorinae","Pollenia","Pollenia pediculata","1738","1480","1480","11","14.84","0","99.43","434.07","0","436.33","33.48","0"
"AGAW2","11N","INS","BOLD:ACU7906","Arthropoda","Insecta","Diptera","Chironomidae","Orthocladiinae","","","1625","1614","1614","12","0.68","0","98.9","406.04","0","410.28","36.03","0"
"AGAW2","11N","INS","BOLD:ACG5862","Arthropoda","Insecta","Diptera","Agromyzidae","","","","1332","1179","1179","13","11.49","0","98.89","438.64","0","440.85","33.88","0"
"AGAW2","11N","INS","BOLD:AAH3225","Arthropoda","Insecta","Psocodea","Psocidae","Psocinae","Metylophorus","Metylophorus novaescotiae","1310","1044","1044","14","20.31","0","98.25","439.28","0","440.41","34.74","0"
"AGAW2","11N","INS","BOLD:AAA3181","Arthropoda","Insecta","Diptera","Muscidae","Muscinae","Stomoxys","Stomoxys calcitrans","1169","1006","1006","15","13.94","0","98.74","428.3","0","432.06","32.91","0"
"AGAW2","11N","INS","BOLD:AAH2307","Arthropoda","Insecta","Diptera","Asilidae","","","","1107","732","732","16","33.88","0","99.68","428.6","0","446.07","46.09","0"
"AGAW2","11N","INS","BOLD:AAD2041","Arthropoda","Insecta","Diptera","Dolichopodidae","Sciapodinae","Condylostylus","Condylostylus inermis, Condylostylus nigrofemoratus, Condylostylus connectans, Condylostylus caudatus, Condylostylus flavipes","1002","747","747","17","25.45","0","99.59","437.26","0","438.47","36.11","0"
"AGAW2","11N","INS","BOLD:AAH3661","Arthropoda","Insecta","Diptera","Cecidomyiidae","","","","875","771","771","18","11.89","0","99.27","429.36","0","431.44","25.33","0"
"AGAW2","11N","INS","BOLD:ACE7444","Arthropoda","Insecta","Hemiptera","Miridae","Mirinae, Phylinae","Adelphocoris, Megalocoleus","Adelphocoris lineolatus, Megalocoleus molliculus","849","583","583","19","31.33","0","99.69","440.72","0","441.49","34.8","0"
"AGAW2","11N","INS","BOLD:ABY5318","Arthropoda","Insecta","Diptera","Chloropidae","Oscinellinae","Incertella, Rhopalopterum","Rhopalopterum umbrosum, Rhopalopterum luteiceps, Incertella bispina, Rhopalopterum painteri","794","599","599","20","24.56","0","99.46","443.33","0","444.45","32.17","0"
"AGAW2","11N","INS","BOLD:AAN5242","Arthropoda","Insecta","Diptera","Cecidomyiidae","","","","715","698","698","21","2.38","0","98.79","404.24","0","406.47","28.12","0"
"AGAW2","11N","INS","BOLD:ABW7897","Arthropoda","Insecta","Diptera","Cecidomyiidae","","","","701","512","512","22","26.96","0","99.62","442.49","0","444.18","29.64","0"
"AGAW2","11N","INS","BOLD:AAV5575","Arthropoda","Insecta","Diptera","Cecidomyiidae","","","CecidInt30 sp. BOLD:AAV5575","651","483","483","23","25.81","0","99.6","444.01","0","445.31","25.92","0"
"AGAW2","11N","INS","BOLD:AAG5507","Arthropoda","Insecta","Diptera","Chironomidae","Chironominae","Chironomus","Chironomus atrella","630","618","618","24","1.9","0","99.04","425","0","429.87","36.09","0"
"AGAW2","11N","INS","BOLD:AAB5609","Arthropoda","Insecta","Diptera","Muscidae","Coenosiinae","Coenosia","Coenosia sp.BOLD:AAB5609, Coenosia tigrina","487","477","477","25","2.05","0","99.08","412.54","0","415.93","32.82","0"
"AGAW2","11N","INS","BOLD:AAG2898","Arthropoda","Insecta","Hemiptera","Cicadellidae","Deltocephalinae","Paraphlepsius","Paraphlepsius apertus, Paraphlepsius irroratus","314","260","260","26","17.2","0","99.34","437.92","0","439.17","32.01","0"
"AGAW2","11N","INS","BOLD:ABA0822","Arthropoda","Insecta","Diptera","Cecidomyiidae","","","","300","277","277","27","7.67","0","98.77","431.67","0","433.03","26.12","0"
"AGAW2","11N","INS","BOLD:ABW1379","Arthropoda","Insecta","Diptera","Chloropidae","Oscinellinae","Malloewia","Malloewia abdominalis","287","234","234","28","18.47","0","98.37","445.23","0","447.62","31.54","0"
"AGAW2","11N","INS","BOLD:AAD1485","Arthropoda","Insecta","Diptera","Chironomidae","Chironominae","Paratanytarsus","Paratanytarsus grimmii, Paratanytarsus sp.","263","235","235","29","10.65","0","99.51","442.36","0","444","33.78","0"
"AGAW2","11N","INS","BOLD:AAN5199","Arthropoda","Insecta","Diptera","Cecidomyiidae","","","CecidInt20 sp. BOLD:AAN5199","255","210","210","30","17.65","0","99.58","441.03","0","444.49","26.5","0"
"AGAW2","11N","INS","BOLD:AAN5183","Arthropoda","Insecta","Diptera","Cecidomyiidae","","","","242","206","206","31","14.88","0","98.52","440.33","0","441.52","27.74","0"
"AGAW2","11N","INS","BOLD:ACX5521","Arthropoda","Insecta","Diptera","Chironomidae","Orthocladiinae","","","242","232","232","31","4.13","0","98.94","415.64","0","417.65","34.93","0"
"AGAW2","11N","INS","BOLD:AAN5162","Arthropoda","Insecta","Diptera","Ceratopogonidae","Dasyheleinae","Dasyhelea","","238","233","233","33","2.1","0","99.11","429.21","0","431.59","33.49","0"
"AGAW2","11N","INS","BOLD:AAN5161","Arthropoda","Insecta","Diptera","Ceratopogonidae","Dasyheleinae","Dasyhelea","","230","228","228","34","0.87","0","99.01","400.9","0","403.19","37.38","0"
"AGAW2","11N","INS","BOLD:ACC8079","Arthropoda","Insecta","Diptera","Cecidomyiidae","","","","194","142","142","35","26.8","0","99.45","447.77","0","449.24","29.29","0"
"AGAW2","11N","INS","BOLD:AAG6929","Arthropoda","Insecta","Diptera","Dolichopodidae","Medeterinae","Medetera","Medetera vittata","177","168","168","36","5.08","0","98.87","434.01","0","435.04","40.6","0"
"AGAW2","11N","INS","BOLD:AAA4781","Arthropoda","Insecta","Hymenoptera","Braconidae","Microgastrinae","Glyptapanteles","Glyptapanteles carlossarmientoi","158","154","154","37","2.53","0","99.15","426.03","0","428.75","24.06","0"
"AGAW2","11N","INS","BOLD:ABA6931","Arthropoda","Insecta","Coleoptera","Latridiidae","Corticariinae","Corticarina","Corticarina longipennis","144","139","139","38","3.47","0","97.9","428.67","0","430.44","33.26","0"
"AGAW2","11N","INS","BOLD:ABA0863","Arthropoda","Insecta","Diptera","Cecidomyiidae","","","","139","121","121","39","12.95","0","99.48","438.55","0","439.99","29.74","0"
"AGAW2","11N","INS","BOLD:AAD7604","Arthropoda","Insecta","Coleoptera","Coccinellidae","Coccinellinae","Coleomegilla","Coleomegilla maculata","125","116","116","40","7.2","0","99.05","433.89","0","436.27","31.69","0"
"AGAW2","11N","INS","BOLD:AAH0135","Arthropoda","Insecta","Coleoptera","Phalacridae","Phalacrinae","Acylomus","Acylomus pugetanus, Acylomus ergoti","120","117","117","41","2.5","0","99.26","433.1","0","435.33","31.5","0"
"AGAW2","11N","INS","BOLD:AAH0256","Arthropoda","Insecta","Coleoptera","Latridiidae","Corticariinae","Corticaria, Corticarina","","116","99","99","42","14.66","0","98.82","430.25","0","435.91","31.46","0"
"AGAW2","11N","INS","BOLD:ADF6661","Arthropoda","Insecta","Diptera","Cecidomyiidae","","","","103","85","85","43","17.48","0","99.25","416.52","0","444.09","29.95","0"
"AGAW2","11N","INS","BOLD:ABU5525","Arthropoda","Insecta","Diptera","Chironomidae","Orthocladiinae","Limnophyes","Limnophyes sp. 14ES","101","101","101","44","0","0","98.97","391.9","0","394.66","37.23","0"
"AGAW2","11N","INS","BOLD:AAG5541","Arthropoda","Insecta","Diptera","Chironomidae","Chironominae","Polypedilum","","95","95","95","45","0","0","97.58","392.95","0","395.24","32.06","0"
"AGAW2","11N","INS","BOLD:ACK5742","Arthropoda","Insecta","Diptera","Cecidomyiidae","","","","92","80","80","46","13.04","0","99.45","441.48","0","442.42","29.03","0"
"AGAW2","11N","INS","BOLD:AAA2482","Arthropoda","Insecta","Lepidoptera","Noctuidae","Noctuinae","Mythimna","Mythimna unipuncta","91","87","87","47","4.4","0","99.27","415.69","0","418.93","30.17","0"
"AGAW2","11N","INS","BOLD:ACE0554","Arthropoda","Insecta","Diptera","Phoridae","Metopininae","Megaselia","","84","74","74","48","11.9","0","99.71","441.29","0","443.5","32.02","0"
"AGAW2","11N","INS","BOLD:AAG2740","Arthropoda","Insecta","Diptera","Ephydridae","Parydrinae","Philygria","Philygria debilis","84","81","81","48","3.57","0","98.2","425.51","0","428.35","33.6","0"
"AGAW2","11N","INS","BOLD:ABV1297","Arthropoda","Insecta","Diptera","Sciaridae","","","","80","80","80","50","0","0","99.17","429.94","0","432.99","30.14","0"
"AGAW2","11N","INS","BOLD:AAN6447","Arthropoda","Insecta","Diptera","Sciaridae","","Corynoptera","Corynoptera perpusilla","77","67","67","51","12.99","0","99.44","447.22","0","448.73","36.25","0"
"AGAW2","11N","INS","BOLD:AAB0651","Arthropoda","Insecta","Lepidoptera","Gelechiidae","Dichomeridinae","Dichomeris","Dichomeris leuconotella, Dichomeris gnoma","76","61","61","52","19.74","0","99.72","446.5","0","447.74","33.22","0"
"AGAW2","11N","INS","BOLD:AAF9771","Arthropoda","Insecta","Diptera","Hybotidae","Tachydromiinae","Platypalpus","Platypalpus longiseta, Platypalpus trivialis, Platypalpus sp. 5, Platypalpus sp. 17, Platypalpus pallidiventris","68","49","49","53","27.94","0","98.99","445.59","0","446.31","35.01","0"
"AGAW2","11N","INS","BOLD:AAA7661","Arthropoda","Insecta","Diptera","Culicidae","Culicinae","Culiseta, Culex","Culiseta melanura, Culex articularis, Culex restuans, Culex acharistus","65","62","62","54","4.62","0","99.33","437.4","0","439.17","32.84","0"
"AGAW2","11N","INS","BOLD:AAM6295","Arthropoda","Insecta","Diptera","Chaoboridae","","","","61","53","53","55","13.11","0","99.67","445.9","0","447.02","37.12","0"
"AGAW2","11N","INS","BOLD:ACE6318","Arthropoda","Insecta","Psocodea","Psocidae","Psocinae","Trichadenotecnum","Trichadenotecnum alexanderae complex","56","53","53","56","5.36","0","99.3","430.66","0","432.86","36.35","0"
"AGAW2","11N","INS","BOLD:AAL7534","Arthropoda","Insecta","Diptera","Anisopodidae","Anisopodinae","Sylvicola","Sylvicola stackelbergi, Sylvicola stackelbergii 1, Sylvicola punctatus","56","51","51","56","8.93","0","99.22","410.39","0","412.63","33","0"
"AGAW2","11N","INS","BOLD:ADI1144","","","","","","","","56","48","48","56","14.29","0","99.39","414.84","0","432.75","30.55","0"
"AGAW2","11N","INS","BOLD:ACL1796","Arthropoda","Insecta","Lepidoptera","Ypsolophidae","Ochsenheimeriinae","Ochsenheimeria","Ochsenheimeria vacculella","51","45","45","59","11.76","0","99.09","432.22","0","439.63","32.61","0"
"AGAW2","11N","INS","BOLD:AAA4277","Arthropoda","Insecta","Diptera","Syrphidae","Syrphinae","Toxomerus","Toxomerus marginatus, Toxomerus sp. CR-3, Toxomerus CR-3","51","44","44","59","13.73","0","99.64","441.55","0","442.76","30.51","0"
"AGAW2","11N","INS","BOLD:AAG8091","Arthropoda","Insecta","Hymenoptera","Ichneumonidae","Campopleginae","Campoletis","Campoletis sp. AB.2","51","51","51","59","0","0","99.32","433.25","0","435.63","24.65","0"
"AGAW2","11N","INS","BOLD:AAG2868","Arthropoda","Insecta","Hemiptera","Cicadellidae","Typhlocybinae","Empoasca","Empoasca sp., Empoasca fabae","50","36","36","62","28","0","99.69","441.82","0","446.32","32.43","0"
"AGAW2","11N","INS","BOLD:ACK3431","Arthropoda","Insecta","Diptera","Cecidomyiidae","","","","44","41","41","63","6.82","0","99.13","423.64","0","425.45","32.26","0"
"AGAW2","11N","INS","BOLD:AAF9306","Arthropoda","Insecta","Diptera","Psychodidae","Psychodinae","Psychoda","Psychoda alternata","44","44","44","63","0","0","98.16","428.52","0","429.86","31.1","0"
"AGAW2","11N","INS","BOLD:AAG7891","Arthropoda","Insecta","Hymenoptera","Platygastridae","Scelioninae","Telenomus","Telenomus podisi","43","40","40","65","6.98","0","99.31","425.79","0","434.56","27","0"
"AGAW2","11N","INS","BOLD:AAV5098","Arthropoda","Insecta","Diptera","Ceratopogonidae","Dasyheleinae","Dasyhelea","Dasyhelea turficola, Dasyhelea sp. 5ES","42","37","37","66","11.9","0","98.96","427.33","0","429.52","36.36","0"
"AGAW2","11N","INS","BOLD:AAN5260","Arthropoda","Insecta","Diptera","Cecidomyiidae","","","","42","41","41","66","2.38","0","99.04","410.52","0","412.14","25.83","0"
"AGAW2","11N","INS","BOLD:AAM9013","Arthropoda","Insecta","Diptera","Keroplatidae","Macrocerinae","Macrocera","Macrocera pusilla","42","41","41","66","2.38","0","99.27","431.43","0","433.05","30.36","0"
"AGAW2","11N","INS","BOLD:ACG3646","Arthropoda","Insecta","Coleoptera","Corylophidae","Corylophinae","Orthoperus","","39","38","38","69","2.56","0","98.83","439.46","0","441.92","33.88","0"
"AGAW2","11N","INS","BOLD:AAV1261","Arthropoda","Insecta","Diptera","Sciaridae","","Bradysia","Bradysia splendida","38","37","37","70","2.63","0","99.5","435.87","0","437.29","34.03","0"
"AGAW2","11N","INS","BOLD:ABV1279","Arthropoda","Insecta","Diptera","Cecidomyiidae","","","","35","32","32","71","8.57","0","99.54","438.23","0","440.34","28.7","0"
"AGAW2","11N","INS","BOLD:AAN5205","Arthropoda","Insecta","Diptera","Cecidomyiidae","","","CecidInt42 sp. BOLD:AAN5205","33","31","31","72","6.06","0","99.02","438.79","0","444.36","31.19","0"
"AGAW2","11N","INS","BOLD:AAN5255","Arthropoda","Insecta","Diptera","Cecidomyiidae","Cecidomyiinae","Dasineura","","30","28","28","73","6.67","0","98.85","423.8","0","446.7","26.72","0"
"AGAW2","11N","INS","BOLD:AAV1136","Arthropoda","Insecta","Diptera","Scatopsidae","","","","30","30","30","73","0","0","98.9","418.27","0","420.13","36.72","0"
"AGAW2","11N","INS","BOLD:ACI1304","Arthropoda","Insecta","Diptera","Muscidae","Muscinae","Stomoxys","Stomoxys indicus","28","28","28","75","0","0","97.69","429.29","0","432.64","32.6","0"
"AGAW2","11N","INS","BOLD:AAI3014","Arthropoda","Insecta","Lepidoptera","Gracillariidae","Phyllocnistinae","Phyllocnistis","Phyllocnistis vitegenella, Phyllocnistis vitifoliella","28","20","20","75","28.57","0","99.4","446.75","0","449.75","29.65","0"
"AGAW2","11N","INS","BOLD:ACV5349","Arthropoda","Insecta","Diptera","Chironomidae","Tanypodinae","","","26","23","23","77","11.54","0","98.12","387.73","0","437.04","36.62","0"
"AGAW2","11N","INS","BOLD:AAN5166","Arthropoda","Insecta","Diptera","Ceratopogonidae","Ceratopogoninae","Culicoides","","26","26","26","77","0","0","99.08","431.46","0","433.58","34.8","0"
"AGAW2","11N","INS","BOLD:AAH0134","Arthropoda","Insecta","Coleoptera","Phalacridae","Phalacrinae","Stilbus","Stilbus apicialis","24","24","24","79","0","0","99.41","427.17","0","428.21","31.97","0"
"AGAW2","11N","INS","BOLD:AAM6040","Arthropoda","Insecta","Diptera","Cecidomyiidae","","","","23","19","19","80","17.39","0","99.41","441.65","0","442.74","27.06","0"
"AGAW2","11N","INS","BOLD:AAG1000","Arthropoda","Insecta","Diptera","Chironomidae","Orthocladiinae","Orthocladius","Orthocladius carlatus","23","23","23","80","0","0","97.94","403.13","0","405.43","33.5","0"
"AGAW2","11N","INS","BOLD:ADF3937","","","","","","","","23","23","23","80","0","0","98.72","406.09","0","414.3","31.03","0"
"AGAW2","11N","INS","BOLD:AAU6592","Arthropoda","Insecta","Diptera","Ceratopogonidae","Dasyheleinae","Dasyhelea","Dasyhelea arenivaga","22","22","22","83","0","0","98.88","411.91","0","413.91","35.03","0"
"AGAW2","11N","INS","BOLD:ABV0266","Arthropoda","Insecta","Diptera","Chloropidae","Oscinellinae","Liohippelates","Liohippelates bishoppi","20","20","20","84","0","0","98.75","441.25","0","445.05","30.55","0"
"AGAW2","11N","INS","BOLD:ABZ5071","Arthropoda","Insecta","Diptera","Chironomidae","Chironominae","Micropsectra","Micropsectra nigripila","20","20","20","84","0","0","98.3","422.6","0","436.5","36.76","0"
"AGAW2","11N","INS","BOLD:ACL9726","Arthropoda","Insecta","Hemiptera","Delphacidae","Delphacinae","Pissonotus","Pissonotus dorsalis","20","20","20","84","0","0","97.69","407.25","0","410.8","35.44","0"
"AGAW2","11N","INS","BOLD:AAN5220","Arthropoda","Insecta","Diptera","Cecidomyiidae","","","","20","20","20","84","0","0","98.62","413.6","0","422.45","25.87","0"
"AGAW2","11N","INS","BOLD:ACX6055","Arthropoda","Insecta","Diptera","Phoridae","Metopininae","Megaselia","Megaselia sp. AK-2010-2, Megaselia sp. BOLD:ACX6055","19","18","18","88","5.26","0","99.64","444.16","0","445.84","33.2","0"
"AGAW2","11N","INS","BOLD:ABA0845","Arthropoda","Insecta","Diptera","Cecidomyiidae","","","","17","13","13","89","23.53","0","99.25","446.59","0","447.24","31.04","0"
"AGAW2","11N","INS","BOLD:ABA1661","Arthropoda","Insecta","Coleoptera","Chrysomelidae","Galerucinae","Longitarsus","Longitarsus pellucidus","17","17","17","89","0","0","99.51","426.65","0","428.24","35.56","0"
"AGAW2","11N","INS","BOLD:ABA0811","Arthropoda","Insecta","Diptera","Cecidomyiidae","","","","17","15","15","89","11.76","0","99.24","442.82","0","444.59","26.34","0"
"AGAW2","11N","INS","BOLD:AAG9668","Arthropoda","Insecta","Diptera","Dolichopodidae","Diaphorinae","Chrysotus","Chrysotus sp., Chrysotus sp. nr. obliquus","15","14","14","92","6.67","0","99.03","446.33","0","449.93","32.82","0"
"AGAW2","11N","INS","BOLD:ACK1651","Arthropoda","Insecta","Diptera","Cecidomyiidae","","","","15","14","14","92","6.67","0","99.59","436.8","0","438.33","26.69","0"
"AGAW2","11N","INS","BOLD:AAC4197","Arthropoda","Insecta","Diptera","Chironomidae","Orthocladiinae","Paraphaenocladius","Paraphaenocladius impensus","15","12","12","92","20","0","99.62","430.93","0","432.6","35.01","0"
"AGAW2","11N","INS","BOLD:AAA3867","Arthropoda","Insecta","Lepidoptera","Noctuidae","Noctuinae","Apamea","Apamea amputatrix","14","14","14","95","0","0","99.27","438.86","0","440.79","33.73","0"
"AGAW2","11N","INS","BOLD:ACP6694","Arthropoda","Insecta","Diptera","Chironomidae","Orthocladiinae","","","13","12","12","96","7.69","0","99.45","428.92","0","431.08","34.06","0"
"AGAW2","11N","INS","BOLD:AAF9949","Arthropoda","Insecta","Hemiptera","Miridae","Mirinae","Trigonotylus","Trigonotylus pulchellus, Trigonotylus americanus, Trigonotylus psammaecolor, Trigonotylus caelestialium, Trigonotylus viridis, Trigonotylus ruficornis","11","9","9","97","18.18","0","99.84","449","0","450","34.14","0"
"AGAW2","11N","INS","BOLD:AAG3302","Arthropoda","Insecta","Diptera","Phoridae","Metopininae","Megaselia","","11","11","11","97","0","0","99.37","412.55","0","415.91","31.32","0"
"AGAW2","11N","INS","BOLD:AAG3275","Arthropoda","Insecta","Diptera","Phoridae","Metopininae","Megaselia","Megaselia minuta, Megaselia nubilipennis","10","10","10","99","0","0","99.08","434.8","0","436.3","34.49","0"
"AGAW2","11N","INS","BOLD:AAG3154","Arthropoda","Insecta","Diptera","Psychodidae","Psychodinae","Psychoda","","10","10","10","99","0","0","98.42","416","0","417.7","32.1","0"
"AGAW2","11N","INS","BOLD:AAC0903","Arthropoda","Insecta","Diptera","Chironomidae","Chironominae","Chironomus","Chironomus acidophilus","10","10","10","99","0","0","99.34","434.8","0","435.5","36.38","0"
"AGAW2","11N","INS","BOLD:AAI1549","Arthropoda","Insecta","Hymenoptera","Braconidae","Euphorinae","Meteorus","","10","8","8","99","20","0","99.09","448.2","0","450","31.56","0"
"AGAW2","11N","INS","BOLD:ABA0856","Arthropoda","Insecta","Diptera","Cecidomyiidae","","","","9","8","8","103","11.11","0","99.63","449.78","0","450","29.41","0"
"AGAW2","11N","INS","BOLD:ACB1322","Arthropoda","Insecta","Coleoptera","Mordellidae","Mordellinae","Mordellistena","Mordellistena trifasciata","9","9","9","103","0","0","99.61","403.78","0","442.11","41.57","0"
"AGAW2","11N","INS","BOLD:AAL7819","Arthropoda","Insecta","Diptera","Psychodidae","Psychodinae","Psychoda","Psychoda sp., Psychoda sp. BOLD:AAL7819","9","9","9","103","0","0","99.58","409.22","0","410.78","32.79","0"
"AGAW2","11N","INS","BOLD:ACV5892","Arthropoda","Insecta","Diptera","Cecidomyiidae","","","","9","9","9","103","0","0","97.76","445.56","0","446.67","25.34","0"
"AGAW2","11N","INS","BOLD:ACV1014","Arthropoda","Insecta","Diptera","Cecidomyiidae","","","","8","8","8","107","0","0","98.06","368.13","0","370.75","25.28","0"
"AGAW2","11N","INS","BOLD:ADH8410","","","","","","","","7","7","7","108","0","0","99.03","420.14","0","423","33.31","0"
"AGAW2","11N","INS","BOLD:AAY6534","Arthropoda","Insecta","Coleoptera","Mordellidae","Mordellinae","Mordellina","Mordellina ancilla","7","5","5","108","28.57","0","99.97","449.71","0","450","38.89","0"
"AGAW2","11N","INS","BOLD:ACJ6954","Arthropoda","Insecta","Diptera","Cecidomyiidae","","","","6","6","6","110","0","0","99.9","337.67","0","440.83","26.83","0"
"AGAW2","11N","INS","BOLD:AAA4222","Arthropoda","Insecta","Lepidoptera","Erebidae","Hypeninae","Hypena","Hypena scabra","6","6","6","110","0","0","99.23","375.5","0","377.17","32.26","0"
"AGAW2","11N","INS","BOLD:ABV1422","Arthropoda","Insecta","Diptera","Cecidomyiidae","","","","6","6","6","110","0","0","98.81","417.67","0","420.5","26.59","0"
"AGAW2","11N","INS","BOLD:ABA6407","Arthropoda","Insecta","Diptera","Sciaridae","","Pseudolycoriella","","6","6","6","110","0","0","99.37","448.17","0","450","35.19","0"
"AGAW2","11N","INS","BOLD:ADH9867","Arthropoda","Insecta","Diptera","","","","","6","6","6","110","0","0","98.93","405.83","0","409.33","37.69","0"
"AGAW2","11N","INS","BOLD:ACF4925","Arthropoda","Insecta","Hemiptera","Miridae","Mirinae","Adelphocoris","Adelphocoris detritus","6","3","3","110","50","0","97.32","448.17","0","449.33","36.46","0"
"AGAW2","11N","INS","BOLD:AAN8770","Arthropoda","Insecta","Diptera","Psychodidae","Psychodinae","Psychoda","Psychoda trinodulosa","6","5","5","110","16.67","0","98.79","423.17","0","424","33.32","0"
"AGAW2","11N","INS","BOLD:AAP2601","Arthropoda","Insecta","Psocodea","Psocidae","Psocinae","Trichadenotecnum","Trichadenotecnum alexanderae complex","5","5","5","117","0","0","98.13","436.4","0","440.2","36.16","0"
"AGAW2","11N","INS","BOLD:ACV2677","Arthropoda","Insecta","Diptera","Cecidomyiidae","","","","5","4","4","117","20","0","97.87","450","0","450","29.78","0"
"AGAW2","11N","INS","BOLD:ACU7686","Arthropoda","Insecta","Diptera","Cecidomyiidae","","","","5","5","5","117","0","0","97.3","429.4","0","430.2","25.72","0"
"AGAW2","11N","INS","BOLD:ABZ4994","Arthropoda","Insecta","Orthoptera","Acrididae","Melanoplinae","Melanoplus","Melanoplus gladstoni, Melanoplus sanguinipes","5","4","4","117","20","0","97.22","436.4","0","437.6","35.98","0"
"AGAW2","11N","INS","BOLD:AAA1259","Arthropoda","Insecta","Diptera","Cecidomyiidae","","","","4","4","4","121","0","0","97.5","418.75","0","420.75","31.16","0"
"AGAW2","11N","INS","BOLD:ACE8878","Arthropoda","Insecta","Orthoptera","Acrididae","Melanoplinae","Melanoplus","Melanoplus scudderi group","4","4","4","121","0","0","97.66","449.75","0","450","35.61","0"
"AGAW2","11N","INS","BOLD:ACL4792","Arthropoda","Insecta","Diptera","Cecidomyiidae","","","","4","4","4","121","0","0","97.91","431.25","0","432.5","27.49","0"
"AGAW2","11N","INS","BOLD:ACN0443","Arthropoda","Insecta","Diptera","Chironomidae","Chironominae","Cladotanytarsus","Cladotanytarsus sp. TE04","4","4","4","121","0","0","99.04","414.75","0","416","39.07","0"
"AGAW2","11N","INS","BOLD:ABA0860","Arthropoda","Insecta","Diptera","Cecidomyiidae","","","","3","3","3","125","0","0","98.77","400","0","401","25.25","0"
"AGAW2","11N","INS","BOLD:AAB9578","Arthropoda","Insecta","Coleoptera","Curculionidae","Scolytinae","Xyleborinus","Xyleborinus saxeseni","3","2","2","125","33.33","0","98.89","449.33","0","450","38.67","0"
"AGAW2","11N","INS","BOLD:ACD8681","Arthropoda","Insecta","Diptera","Cecidomyiidae","Cecidomyiinae","Ozirhincus, Asphondylia","Asphondylia sp., Ozirhincus hungaricus, Ozirhincus tanaceti","3","3","3","125","0","0","99.66","426.33","0","427.33","25.49","0"
"AGAW2","11N","INS","BOLD:AAR3352","Arthropoda","Insecta","Coleoptera","Staphylinidae","Oxytelinae","Anotylus","Anotylus insecatus","3","3","3","125","0","0","99.33","420.33","0","424.33","35.68","0"
"AGAW2","11N","INS","BOLD:AAZ5483","Arthropoda","Insecta","Diptera","Muscidae","Phaoniinae","Helina","Helina setiventris","3","3","3","125","0","0","97.55","413.33","0","417","29.6","0"
"AGAW2","11N","INS","BOLD:AAB0373","Arthropoda","Insecta","Neuroptera","Chrysopidae","Chrysopinae","Chrysoperla","Chrysoperla sp. isolate 82, Chrysoperla zastrowi, Chrysoperla plorabunda, Chrysoperla sp. isolate 71, Chrysoperla agilis, Italochrysa italica, Chrysoperla sp. isolate 88, Chrysoperla lucasina, Chrysoperla carnea, Chrysoperla mediterranea, Chrysoperla pallida, Chrysoperla nipponensis, Chrysoperla sillemi","3","3","3","125","0","0","99.16","410","0","411","31.04","0"
"AGAW2","11N","INS","BOLD:AAN5337","Arthropoda","Insecta","Diptera","Chironomidae","Tanypodinae","","","3","3","3","125","0","0","98.59","448.33","0","450","34.96","0"
"AGAW2","11N","INS","BOLD:AAN5334","Arthropoda","Insecta","Diptera","Chironomidae","Chironominae","Polypedilum","","3","3","3","125","0","0","97.6","416","0","418.67","34.54","0"
"AGAW2","11N","INS","BOLD:AAH3664","Arthropoda","Insecta","Diptera","Cecidomyiidae","","","CecidInt3 sp. BOLD:AAH3664","3","3","3","125","0","0","99.53","419.33","0","427","26.82","0"
"AGAW2","11N","INS","BOLD:ABX7480","Arthropoda","Insecta","Diptera","Cecidomyiidae","","","","3","3","3","125","0","0","97.66","403.67","0","425.67","26.99","0"
"AGAW2","11N","INS","BOLD:ACO7137","Arthropoda","Insecta","Diptera","Cecidomyiidae","","","","3","3","3","125","0","0","98.39","419.33","0","417","27.95","0"
"AGAW2","11N","INS","BOLD:ACF1257","","","","","","","","3","3","3","125","0","0","99.7","428","0","429.33","34.23","0"
"AGAW2","11N","INS","BOLD:ADH7839","Arthropoda","Insecta","Diptera","","","","","3","3","3","125","0","0","99.07","438.67","0","441.33","25.89","0"
"AGAW2","11N","INS","BOLD:AAU8628","Arthropoda","Insecta","Hymenoptera","Bethylidae","","","","3","3","3","125","0","0","99.85","447.67","0","450","27.26","0"
"AGAW2","11N","INS","BOLD:AAY6450","Arthropoda","Insecta","Diptera","Cecidomyiidae","","","","2","2","2","139","0","0","98.65","446","0","450","27.67","0"
"AGAW2","11N","INS","BOLD:ABY1165","Arthropoda","Insecta","Coleoptera","Anthicidae","","","","2","2","2","139","0","0","97.8","412.5","0","414","35.99","0"
"AGAW2","11N","INS","BOLD:AAC0022","Arthropoda","Insecta","Diptera","Chironomidae","Chironominae","Micropsectra","Micropsectra nigripila","2","2","2","139","0","0","97.65","447.5","0","450","37","0"
"AGAW2","11N","INS","BOLD:ACE4100","Arthropoda","Insecta","Orthoptera","Acrididae","Melanoplinae","Melanoplus","Melanoplus pinicola","2","2","2","139","0","0","97.33","450","0","450","35.11","0"
"AGAW2","11N","INS","BOLD:AAY6730","Arthropoda","Insecta","Hemiptera","Derbidae","Cedusinae","Cedusa","Cedusa incisa","2","2","2","139","0","0","99.4","415.5","0","417.5","35.42","0"
"AGAW2","11N","INS","BOLD:ACK5443","Arthropoda","Insecta","Diptera","Cecidomyiidae","","","","2","2","2","139","0","0","99.22","449","0","450","26","0"
"AGAW2","11N","INS","BOLD:AAB4657","Arthropoda","Insecta","Diptera","Chironomidae","Chironominae","Chironomus","Chironomus maturus","2","2","2","139","0","0","99.43","440","0","441","38.09","0"
"AGAW2","11N","INS","BOLD:AAG5453","Arthropoda","Insecta","Diptera","Chironomidae","Chironominae","Chironomus","Chironomus bifurcatus","2","2","2","139","0","0","98.21","448","0","450","35","0"
"AGAW2","11N","INS","BOLD:AAM9347","Arthropoda","Insecta","Diptera","Phoridae","Metopininae","Megaselia","","2","2","2","139","0","0","97.92","398.5","0","401","33.26","0"
"AGAW2","11N","INS","BOLD:ACE2759","Arthropoda","Insecta","Thysanoptera","Aeolothripidae","Aeolothripinae","Aeolothrips","Aeolothrips albicinctus","2","2","2","139","0","0","98.77","446.5","0","450","38.44","0"
"AGAW2","11N","INS","BOLD:AAV5400","Arthropoda","Insecta","Diptera","Cecidomyiidae","","","","2","2","2","139","0","0","98.14","401","0","403.5","28.5","0"
"AGAW2","11N","INS","BOLD:ADH8687","Arthropoda","Insecta","Coleoptera","","","","","2","2","2","139","0","0","100","438","0","450","34.89","0"
"AGAW2","11N","INS","BOLD:ABY5257","Arthropoda","Insecta","Lepidoptera","Noctuidae","Noctuinae","Apamea","Apamea devastator","2","2","2","139","0","0","98.78","449.5","0","450","30.67","0"
"AGAW2","11N","INS","BOLD:ACN6329","Arthropoda","Insecta","Diptera","Cecidomyiidae","","","","2","2","2","139","0","0","99.08","435.5","0","436","26.37","0"
"AGAW2","11N","INS","BOLD:AAN8091","Arthropoda","Insecta","Hymenoptera","Trichogrammatidae","","Trichogramma","","2","2","2","139","0","0","99.26","407.5","0","422","24.85","0"
"AGAW2","11N","INS","BOLD:ABX0893","Arthropoda","Insecta","Diptera","Cecidomyiidae","","","","2","2","2","139","0","0","97.33","448.5","0","450","26.89","0"
"AGAW2","11N","INS","BOLD:ACJ6984","Arthropoda","Insecta","Hymenoptera","Platygastridae","Scelioninae","Trissolcus","Trissolcus thyantae","2","2","2","139","0","0","99.51","413.5","0","420","26.65","0"
"AGAW2","11N","INS","BOLD:AAD0484","Arthropoda","Insecta","Diptera","Chironomidae","","","","2","2","2","139","0","0","99.33","448.5","0","450","34.67","0"
"AGAW2","11N","INS","BOLD:AAH4701","Arthropoda","Insecta","Lepidoptera","Pterophoridae","Pterophorinae","Sphenarches","Sphenarches ontario","1","1","1","157","0","0","97.53","446","0","450","31.33","0"
"AGAW2","11N","INS","BOLD:AAH4135","Arthropoda","Insecta","Diptera","Chloropidae","Chloropinae","Thaumatomyia","Thaumatomyia glabra, Thaumatomyia grata, Thaumatomyia sp. 1 TAW","1","1","1","157","0","0","97.21","394","0","397","31.74","0"
"AGAW2","11N","INS","BOLD:AAD0952","Arthropoda","Insecta","Lepidoptera","Pyralidae","Chrysauginae","Galasa","Galasa nigrinodis, Galasa nigripunctalis","1","1","1","157","0","0","98.35","364","0","364","32.14","0"
"AGAW2","11N","INS","BOLD:AAN8447","Arthropoda","Insecta","Psocodea","Caeciliusidae","Caeciliusinae","Valenzuela","Valenzuela flavidus","1","1","1","157","0","0","98.28","349","0","351","34.76","0"
"AGAW2","11N","INS","BOLD:ADI0836","","","","","","","","1","1","1","157","0","0","97.95","391","0","395","32.15","0"
"AGAW2","11N","INS","BOLD:AAB4903","Arthropoda","Insecta","Lepidoptera","Glyphipterigidae","Acrolepiinae","Acrolepiopsis","Acrolepiopsis assectella","1","1","1","157","0","0","100","424","0","450","30.89","0"
"AGAW2","11N","INS","BOLD:AAH2306","Arthropoda","Insecta","Diptera","Asilidae","Asilinae","Efferia","Efferia aestuans","1","1","1","157","0","0","99.75","402","0","408","37.01","0"
"AGAW2","11N","INS","BOLD:AAP4633","Arthropoda","Insecta","Psocodea","Psocidae","Psocinae","Loensia","","1","1","1","157","0","0","99.56","450","0","450","37.11","0"
"AGAW2","11N","INS","BOLD:AAD9052","Arthropoda","Insecta","Lepidoptera","Elachistidae","Elachistinae","Elachista","Elachista herbigrada, Elachista rianella, Elachista illectella, Elachista sp., Elachista tuorella","1","1","1","157","0","0","97.42","349","0","352","30.68","0"
"AGAW2","11N","INS","BOLD:AAA8933","Arthropoda","Insecta","Coleoptera","Coccinellidae","Coccinellinae","Coccinella","Coccinella septempunctata","1","1","1","157","0","0","99.33","448","0","450","32.67","0"
"AGAW2","11N","INS","BOLD:ADG8758","Arthropoda","Insecta","Hymenoptera","Braconidae","Brachistinae","Blacus","Blacus stelfoxi","1","1","1","157","0","0","99.55","449","0","450","32.67","0"
"AGAW2","11N","INS","BOLD:AAO8538","Arthropoda","Insecta","Diptera","Psychodidae","Psychodinae","Psychoda","","1","1","1","157","0","0","99.21","381","0","382","31.94","0"
"AGAW2","11N","INS","BOLD:AAB4658","Arthropoda","Insecta","Diptera","Chironomidae","Chironominae","Chironomus","Chironomus dilutus","1","1","1","157","0","0","99.28","418","0","421","37.05","0"
"AGAW2","11N","INS","BOLD:ACA9275","Arthropoda","Insecta","Diptera","Chironomidae","Chironominae","Chironomus","","1","1","1","157","0","0","98.43","445","0","450","35.56","0"
"AGAW2","11N","INS","BOLD:ABZ8007","Arthropoda","Insecta","Lepidoptera","Noctuidae","Cuculliinae","Cucullia","Cucullia similaris, Cucullia postera, Cucullia sp., Cucullia florea","1","1","1","157","0","0","100","449","0","450","31.33","0"
"AGAW2","11N","INS","BOLD:AAG3273","Arthropoda","Insecta","Diptera","Phoridae","Metopininae","Megaselia","Megaselia albicaudata","1","1","1","157","0","0","99.33","449","0","450","32.67","0"
"AGAW2","11N","INS","BOLD:AAY6728","Arthropoda","Insecta","Hemiptera","Cicadellidae","Aphrodinae","Anoscopus","Anoscopus serratulae","1","1","1","157","0","0","100","450","0","450","32.67","0"
"AGAW2","11N","INS","BOLD:AAU6537","Arthropoda","Insecta","Diptera","Sciaridae","","Corynoptera","Corynoptera cuniculata","1","1","1","157","0","0","97.12","417","0","422","33.18","0"
"AGAW2","11N","INS","BOLD:AAN5238","Arthropoda","Insecta","Diptera","Cecidomyiidae","","","","1","1","1","157","0","0","99.32","439","0","440","26.36","0"
"AGAW2","11N","INS","BOLD:AAE7186","Arthropoda","Insecta","Hymenoptera","Braconidae","Aphidiinae","Praon","Praon sp. 1 MM-2013, Praon volucre, Praon bicolor","1","1","1","157","0","0","97.55","449","0","450","25.78","0"
"AGAW2","11N","INS","BOLD:AAN5230","Arthropoda","Insecta","Diptera","Cecidomyiidae","","","","1","1","1","157","0","0","98.22","449","0","450","26","0"
"AGAW2","11N","INS","BOLD:AAE7796","Arthropoda","Insecta","Hemiptera","Aphididae","Aphidinae","Metopolophium","Metopolophium dirhodum, Metopolophium albidum","1","1","1","157","0","0","98.89","449","0","450","26.22","0"
"AGAW2","11N","INS","BOLD:AAM9105","Arthropoda","Insecta","Diptera","Muscidae","Coenosiinae","Coenosia","","1","1","1","157","0","0","98.89","450","0","450","37.33","0"
"AGAW2","11N","INS","BOLD:AAR9498","Arthropoda","Insecta","Diptera","Chironomidae","Chironominae","Cladotanytarsus","Cladotanytarsus sp. TE04","1","1","1","157","0","0","97.48","437","0","441","39","0"
"AGAW2","11N","INS","BOLD:ACE6286","Arthropoda","Insecta","Diptera","Culicidae","Culicinae","Ochlerotatus, Aedes","Aedes dorsalis, Ochlerotatus dorsalis","1","1","1","157","0","0","97.55","449","0","450","32.89","0"
"AGAW2","11N","INS","BOLD:AAA2632","Arthropoda","Insecta","Lepidoptera","Noctuidae","Noctuinae","Noctua","Noctua atlantica, Noctua pronuba","1","1","1","157","0","0","98.66","447","0","450","30.44","0"
"AGAW2","11N","INS","BOLD:AAH3720","Arthropoda","Insecta","Diptera","Cecidomyiidae","","","CecidInt34 sp. BOLD:AAH3720","1","1","1","157","0","0","99.27","413","0","416","27.16","0"
"AGAW2","11N","INS","BOLD:ABA8948","Arthropoda","Insecta","Coleoptera","Chrysomelidae","","","","1","1","1","157","0","0","97.16","388","0","393","35.88","0"
"AGAW2","11N","INS","BOLD:AAA7067","Arthropoda","Insecta","Diptera","Culicidae","Culicinae","Aedes, Aedimorphus","Aedimorphus vexans, Aedes vexans","1","1","1","157","0","0","99.48","387","0","389","31.62","0"
"AGAW2","11N","INS","BOLD:AAH2295","Arthropoda","Insecta","Diptera","Asilidae","Leptogastrinae","Leptogaster","Leptogaster flavipes","1","1","1","157","0","0","97.09","413","0","450","34.67","0"
"AGAW2","11N","INS","BOLD:AAH2293","Arthropoda","Insecta","Diptera","Asilidae","Stenopogoninae","Dioctria, Eudioctria","Dioctria hyalipennis, Dioctria baumhaueri, Eudioctria propinqua","1","1","1","157","0","0","98.46","390","0","397","29.47","0"
"AGAW2","11N","INS","BOLD:ACI9753","Arthropoda","Insecta","Diptera","Dolichopodidae","Diaphorinae","Chrysotus","","1","1","1","157","0","0","98.54","411","0","450","32","0"
"AGAW2","11N","INS","BOLD:AAC8842","Arthropoda","Insecta","Diptera","Chironomidae","Chironominae","Paratanytarsus","Paratanytarsus laccophilus","1","1","1","157","0","0","98.42","443","0","448","33.48","0"
"AGAW2","11N","INS","BOLD:AAB0269","Arthropoda","Insecta","Lepidoptera","Tortricidae","Tortricinae","Acleris","Acleris albicomana","1","1","1","157","0","0","99.31","433","0","435","33.1","0"
"AGAW2","11N","INS","BOLD:ABW1460","Arthropoda","Insecta","Coleoptera","Chrysomelidae","Criocerinae","Oulema","Oulema erichsonii","1","1","1","157","0","0","99.55","449","0","450","35.56","0"
"AGAW2","11N","INS","BOLD:AAC5848","Arthropoda","Insecta","Lepidoptera","Psychidae","Psychinae","Psyche","Psyche casta","1","1","1","157","0","0","98.25","399","0","402","37.81","0"
"AGAW2","11N","INS","BOLD:AAP9022","Arthropoda","Insecta","Diptera","Cecidomyiidae","","","","1","1","1","157","0","0","99.76","424","0","425","27.06","0"
"AGAW2","11N","INS","BOLD:AAE3015","Arthropoda","Insecta","Hemiptera","Aphididae","Aphidinae","Capitophorus","Capitophorus elaeagni","1","1","1","157","0","0","99.78","449","0","450","25.56","0"
"AGAW2","11N","INS","BOLD:AAN5501","Arthropoda","Insecta","Diptera","Hybotidae","","","","1","1","1","157","0","0","99.55","447","0","450","35.11","0"
"AGAW2","11N","INS","BOLD:ABX2489","Arthropoda","Insecta","Coleoptera","Phalacridae","Phalacrinae","Stilbus","","1","1","1","157","0","0","97.52","403","0","405","32.1","0"
"AGAW2","11N","INS","BOLD:ADH7513","","","","","","","","1","1","1","157","0","0","98.35","423","0","424","33.02","0"
"AGAW2","11N","INS","BOLD:AAB0483","Arthropoda","Insecta","Lepidoptera","Tortricidae","Olethreutinae","Zeiraphera","Zeiraphera vancouverana","1","1","1","157","0","0","97.09","447","0","450","32.89","0"
"AGAW2","11N","INS","BOLD:AAA1962","Arthropoda","Insecta","Diptera","Sarcophagidae","Sarcophaginae","Helicobia, sarcJanzen01, Sarcophaga","sarcJanzen01 Janzen14, Helicobia rapax, Sarcophaga rapax","1","1","1","157","0","0","98.21","446","0","450","35.11","0"
"AGAW2","11N","INS","BOLD:ACV5891","Arthropoda","Insecta","Diptera","Cecidomyiidae","","","","1","1","1","157","0","0","97.79","362","0","450","27.33","0"
"AGAW2","11N","INS","BOLD:ACD8563","Arthropoda","Insecta","Diptera","Cecidomyiidae","","","","1","1","1","157","0","0","97.33","450","0","450","25.56","0"
"AGAW2","11N","INS","BOLD:ACB0984","Arthropoda","Insecta","Psocodea","Stenopsocidae","","Graphopsocus","Graphopsocus cruciatus","1","1","1","157","0","0","99.27","413","0","450","37.56","0"
"AGAW2","11N","INS","BOLD:AAF9014","Arthropoda","Insecta","Diptera","Limoniidae","Chioneinae","Symplecta","Symplecta cana, Symplecta sp. BOLDAAF9014","1","1","1","157","0","0","98.96","384","0","384","33.33","0"
"AGAW2","11N","INS","BOLD:AAA7374","Arthropoda","Insecta","Diptera","Syrphidae","Syrphinae","Sphaerophoria","Sphaerophoria bankowskae, Sphaerophoria infuscata, Sphaerophoria nr. philanthus, Sphaerophoria batava, Sphaerophoria sp., Sphaerophoria rueppellii, Sphaerophoria interrupta-group, Sphaerophoria sp. MA-1, Sphaerophoria cleoae, Sphaerophoria asymmetrica, Sphaerophoria fatarum, Sphaerophoria bengalensis, Sphaerophoria bifurcata, Sphaerophoria taeniata, Sphaerophoria interrupta, Sphaerophoria contigua, Sphaerophoria kaa, Sphaerophoria laurae, Sphaerophoria chongjini, Sphaerophoria boreoalpina, Sphaerophoria sp. BOLDAAA7374, Sphaerophoria macrogaster, Sphaerophoria sulphuripes, Sphaerophoria brevipilosa, Sphaerophoria philanthus, Sphaerophoria pyrrhina, Sphaerophoria abbreviata, Sphaerophoria scripta, Sphaerophoria potentillae, Sphaerophoria longipilosa, Sphaerophoria virgata","1","1","1","157","0","0","97.36","416","0","422","30.33","0"
"AGAW2","11N","INS","BOLD:AAA5803","Arthropoda","Insecta","Hemiptera","Miridae","Mirinae","Lygus","Lygus lineolaris","1","1","1","157","0","0","99.55","449","0","450","34.44","0"
"AGAW2","11N","NAR","BOLD:AAI9663","Mollusca","Gastropoda","Stylommatophora","Agriolimacidae",NA,"Deroceras","Deroceras reticulatum","1","1","1","1","0","0","99.75","407","0","450","36","0"
"AGAW2","11N","NIN","BOLD:AAH6630","Arthropoda","Collembola","Entomobryomorpha","Entomobryidae","Entomobryinae","Entomobrya","Entomobrya unostrigata","467","414","414","1","11.35","0","99.33","434.1","0","435.48","39.03","0"
"AGAW2","11N","NIN","BOLD:AAF9173","Arthropoda","Arachnida","Sarcoptiformes","Mochlozetidae","","Podoribates","Podoribates pratensis","219","166","166","2","24.2","0","99.71","434.3","0","435.15","42.89","0"
"AGAW2","11N","NIN","BOLD:AAI4346","Arthropoda","Arachnida","Opiliones","Phalangiidae","","Paroligolophus, Phalangium","Paroligolophus agrestis, Phalangium opilio","203","197","197","3","2.96","0","98.72","407.24","0","414.48","35.34","0"
"AGAW2","11N","NIN","BOLD:ABW2639","Arthropoda","Arachnida","Trombidiformes","Anystidae","","","","84","69","69","4","17.86","0","99.6","445.85","0","445.1","31.41","0"
"AGAW2","11N","NIN","BOLD:AAA6381","Arthropoda","Arachnida","Araneae","Tetragnathidae","","Tetragnatha","Tetragnatha laboriosa","77","77","77","5","0","0","99.2","428.21","0","430.51","39.57","0"
"AGAW2","11N","NIN","BOLD:AAB7914","Arthropoda","Collembola","Symphypleona","Bourletiellidae","","Bourletiella","","57","47","47","6","17.54","0","99.63","445.68","0","446.6","38.16","0"
"AGAW2","11N","NIN","BOLD:ABX0027","Arthropoda","Collembola","Symphypleona","Bourletiellidae","","Bourletiella","","37","30","30","7","18.92","0","99.62","418.32","0","447.57","35.76","0"
"AGAW2","11N","NIN","BOLD:AAU6252","Arthropoda","Arachnida","Trombidiformes","Bdellidae","","Cyta","","30","27","27","8","10","0","99.43","420.6","0","422.27","31.26","0"
"AGAW2","11N","NIN","BOLD:AAV3042","Arthropoda","Arachnida","Araneae","Theridiidae","","Theridion","Theridion albidum","26","26","26","9","0","0","98.88","418.12","0","419.58","27.88","0"
"AGAW2","11N","NIN","BOLD:ACZ1295","Arthropoda","Arachnida","Trombidiformes","Anystidae","","","","13","13","13","10","0","0","99.24","377.38","0","387.62","31.12","0"
"AGAW2","11N","NIN","BOLD:ACL3993","Arthropoda","Arachnida","Mesostigmata","Ascidae","","","","12","12","12","11","0","0","99.52","426.5","0","437.58","40.84","0"
"AGAW2","11N","NIN","BOLD:ACI9974","Arthropoda","Arachnida","Trombidiformes","Tydeidae","","","","11","10","10","12","9.09","0","99.26","415.18","0","440.91","40.88","0"
"AGAW2","11N","NIN","BOLD:AAA8820","Arthropoda","Collembola","Entomobryomorpha","Entomobryidae","Lepidocyrtinae","Lepidocyrtus","Lepidocyrtus cyaneus","8","8","8","13","0","0","98.58","408.63","0","411","43.29","0"
"AGAW2","11N","NIN","BOLD:ACL2796","Arthropoda","Arachnida","Trombidiformes","Microtrombidiidae","","","","6","6","6","14","0","0","99.7","449.33","0","450","34.78","0"
"AGAW2","11N","NIN","BOLD:AAO8147","Arthropoda","Arachnida","Trombidiformes","Eupodidae","","Eupodes","","4","4","4","15","0","0","99.25","410","0","411","27.66","0"
"AGAW2","11N","NIN","BOLD:ACX8969","Arthropoda","Arachnida","Trombidiformes","Eupodidae","","","","2","2","2","16","0","0","98.6","426.5","0","427.5","28.19","0"
"AGAW2","11N","NIN","BOLD:AAZ2181","Arthropoda","Collembola","Symphypleona","Bourletiellidae","","","","1","1","1","17","0","0","97.53","365","0","365","39.18","0"
"AGAW2","11N","NIN","BOLD:ACJ7454","Arthropoda","Collembola","Symphypleona","Bourletiellidae","","","","1","1","1","17","0","0","99.5","404","0","450","38.89","0"
"AGAW2","11N","NIN","BOLD:AAB7915","Arthropoda","Collembola","Symphypleona","Bourletiellidae","","Deuterosminthurus","Deuterosminthurus sp1","1","1","1","17","0","0","99.33","450","0","450","35.11","0"
"AGAW2","11N","NIN","BOLD:ACI0834","Arthropoda","Arachnida","Trombidiformes","Eriophyidae","","Aceria","Aceria tosichella","1","1","1","17","0","0","98.44","448","0","450","34.89","0"
"AGAW2","11S","BAC","TAX:392874","Proteobacteria","Alphaproteobacteria","Rickettsiales","Rickettsiaceae",NA,"Wolbachia","Wolbachia endosymbiont of Hymenoptera","6","6","6","1","0","0","98.85","449.5","0","450","40.26","0"
"AGAW2","11S","BAC","TAX:376688","Proteobacteria","Alphaproteobacteria","Rickettsiales","Rickettsiaceae",NA,"Wolbachia","Wolbachia sp. elachBioLep01 BioLep373","2","2","2","2","0","0","99.33","450","0","450","40.11","0"
"AGAW2","11S","BAC","TAX:453459","Proteobacteria","Alphaproteobacteria","Rickettsiales","Rickettsiaceae",NA,"Wolbachia","Wolbachia endosymbiont of Ichneumonidae","1","1","1","3","0","0","98.66","449","0","450","40.44","0"
"AGAW2","11S","BAC","TAX:375853","Proteobacteria","Alphaproteobacteria","Rickettsiales","Rickettsiaceae",NA,"Wolbachia","Wolbachia sp. Hymenoptera_ASGLE646-10","1","1","1","3","0","0","97.55","449","0","450","40.44","0"
"AGAW2","11S","BAC","TAX:375847","Proteobacteria","Alphaproteobacteria","Rickettsiales","Rickettsiaceae",NA,"Wolbachia","Wolbachia sp. Hymenoptera_ASGLE1705-10","1","1","1","3","0","0","100","368","0","369","39.84","0"
"AGAW2","11S","BAC","TAX:91391","Proteobacteria","Alphaproteobacteria","Rickettsiales","",NA,"","","1","1","1","3","0","0","98.58","422","0","450","40","0"
"AGAW2","11S","BAC","TAX:375762","Proteobacteria","Alphaproteobacteria","Rickettsiales","Rickettsiaceae",NA,"Wolbachia","Wolbachia sp. Myrmicinae_ACGAN556-09","1","1","1","3","0","0","99.33","450","0","450","40.44","0"
"AGAW2","11S","CHO","no match","no match","no match","no match","no match","no match","no match","no match","no match","no match","no match","no match","no match","no match","no match","no match","no match","no match","no match","no match"
"AGAW2","11S","INS","BOLD:AAK5928","Arthropoda","Insecta","Coleoptera","Chrysomelidae","Criocerinae","Oulema","Oulema melanopus","8339","6157","6157","1","26.17","0","99.58","439.52","0","440.88","36.47","0"
"AGAW2","11S","INS","BOLD:AAA2482","Arthropoda","Insecta","Lepidoptera","Noctuidae","Noctuinae","Mythimna","Mythimna unipuncta","5701","5092","5092","2","10.68","0","99.24","414.86","0","418.19","30.13","0"
"AGAW2","11S","INS","BOLD:AAA4555","Arthropoda","Insecta","Orthoptera","Acrididae","Melanoplinae","Melanoplus","Melanoplus stupefactus, Melanoplus angustipennis, Melanoplus ponderosus, Melanoplus bruneri, Melanoplus frigidus, Melanoplus scapularis, Melanoplus cinereus, Melanoplus madeleineae, Melanoplus sanguinipes, Melanoplus nigrescens group, Melanoplus tunicae, Melanoplus walshi, Melanoplus femurrubrum, Melanoplus dawsoni, Melanoplus sp., Melanoplus fasciatus, Melanoplus bowditchi, Melanoplus stonei, Melanoplus bivittatus, Melanoplus huroni, Melanoplus thomasi, Melanoplus borealis, Melanoplus packardii","5518","4203","4203","3","23.83","0","99.41","446.45","0","447.65","36.19","0"
"AGAW2","11S","INS","BOLD:AAA3453","Arthropoda","Insecta","Diptera","Anthomyiidae","Anthomyiinae","Delia","","5281","4609","4609","4","12.72","0","99.45","429.93","0","432.62","34.62","0"
"AGAW2","11S","INS","BOLD:AAB0651","Arthropoda","Insecta","Lepidoptera","Gelechiidae","Dichomeridinae","Dichomeris","Dichomeris leuconotella, Dichomeris gnoma","4603","2954","2954","5","35.82","0","99.69","446.48","0","447.62","33.19","0"
"AGAW2","11S","INS","BOLD:AAA4277","Arthropoda","Insecta","Diptera","Syrphidae","Syrphinae","Toxomerus","Toxomerus marginatus, Toxomerus sp. CR-3, Toxomerus CR-3","3331","2150","2150","6","35.45","0","99.61","443.71","0","444.69","30.59","0"
"AGAW2","11S","INS","BOLD:AAF3338","Arthropoda","Insecta","Lepidoptera","Noctuidae","Noctuinae","Striacosta","Striacosta albicosta","1629","1487","1487","7","8.72","0","98.44","422.76","0","424.95","33.86","0"
"AGAW2","11S","INS","BOLD:AAN6154","Arthropoda","Insecta","Coleoptera","Latridiidae","Corticariinae","Melanophthalma","Melanophthalma inermis","1506","1264","1264","8","16.07","0","98.95","435.38","0","437","31.46","0"
"AGAW2","11S","INS","BOLD:AAA3867","Arthropoda","Insecta","Lepidoptera","Noctuidae","Noctuinae","Apamea","Apamea amputatrix","992","814","814","9","17.94","0","99.34","439.6","0","441.28","33.72","0"
"AGAW2","11S","INS","BOLD:ACB1322","Arthropoda","Insecta","Coleoptera","Mordellidae","Mordellinae","Mordellistena","Mordellistena trifasciata","786","703","703","10","10.56","0","99.43","393.74","0","421.65","41.25","0"
"AGAW2","11S","INS","BOLD:AAU6681","Arthropoda","Insecta","Diptera","Tipulidae","Tipulinae","Nephrotoma","Nephrotoma cornicina","655","543","543","11","17.1","0","99.66","442.54","0","443.7","36.29","0"
"AGAW2","11S","INS","BOLD:ACU7906","Arthropoda","Insecta","Diptera","Chironomidae","Orthocladiinae","","","599","599","599","12","0","0","98.93","400.3","0","411.54","36.46","0"
"AGAW2","11S","INS","BOLD:AAV5575","Arthropoda","Insecta","Diptera","Cecidomyiidae","","","CecidInt30 sp. BOLD:AAV5575","595","448","448","13","24.71","0","99.65","442.15","0","443.45","25.88","0"
"AGAW2","11S","INS","BOLD:AAG2868","Arthropoda","Insecta","Hemiptera","Cicadellidae","Typhlocybinae","Empoasca","Empoasca sp., Empoasca fabae","447","301","301","14","32.66","0","99.74","446.03","0","448.06","32.4","0"
"AGAW2","11S","INS","BOLD:AAN5242","Arthropoda","Insecta","Diptera","Cecidomyiidae","","","","382","379","379","15","0.79","0","98.91","403.03","0","405.42","28.13","0"
"AGAW2","11S","INS","BOLD:ABY5318","Arthropoda","Insecta","Diptera","Chloropidae","Oscinellinae","Incertella, Rhopalopterum","Rhopalopterum umbrosum, Rhopalopterum luteiceps, Incertella bispina, Rhopalopterum painteri","355","263","263","16","25.92","0","99.65","439.97","0","440.99","32.19","0"
"AGAW2","11S","INS","BOLD:AAA3181","Arthropoda","Insecta","Diptera","Muscidae","Muscinae","Stomoxys","Stomoxys calcitrans","315","281","281","17","10.79","0","98.71","423.91","0","427.57","32.83","0"
"AGAW2","11S","INS","BOLD:AAN5337","Arthropoda","Insecta","Diptera","Chironomidae","Tanypodinae","","","304","266","266","18","12.5","0","98.26","428.6","0","430.78","34.72","0"
"AGAW2","11S","INS","BOLD:AAH3661","Arthropoda","Insecta","Diptera","Cecidomyiidae","","","","263","244","244","19","7.22","0","99.31","429.26","0","431.03","25.3","0"
"AGAW2","11S","INS","BOLD:ACX5521","Arthropoda","Insecta","Diptera","Chironomidae","Orthocladiinae","","","226","213","213","20","5.75","0","99.26","416.79","0","418.67","34.65","0"
"AGAW2","11S","INS","BOLD:AAD1485","Arthropoda","Insecta","Diptera","Chironomidae","Chironominae","Paratanytarsus","Paratanytarsus grimmii, Paratanytarsus sp.","153","124","124","21","18.95","0","99.6","443.46","0","444.96","33.84","0"
"AGAW2","11S","INS","BOLD:AAN5199","Arthropoda","Insecta","Diptera","Cecidomyiidae","","","CecidInt20 sp. BOLD:AAN5199","151","120","120","22","20.53","0","99.56","439.15","0","444.43","26.6","0"
"AGAW2","11S","INS","BOLD:ADF6661","Arthropoda","Insecta","Diptera","Cecidomyiidae","","","","142","120","120","23","15.49","0","99.27","416.52","0","443.18","29.75","0"
"AGAW2","11S","INS","BOLD:AAM9347","Arthropoda","Insecta","Diptera","Phoridae","Metopininae","Megaselia","","140","140","140","24","0","0","98.08","405.74","0","408.12","33.08","0"
"AGAW2","11S","INS","BOLD:AAG6929","Arthropoda","Insecta","Diptera","Dolichopodidae","Medeterinae","Medetera","Medetera vittata","131","99","99","25","24.43","0","98.88","442.85","0","445.34","40.93","0"
"AGAW2","11S","INS","BOLD:ACC8079","Arthropoda","Insecta","Diptera","Cecidomyiidae","","","","123","87","87","26","29.27","0","99.51","446.72","0","449.25","29.26","0"
"AGAW2","11S","INS","BOLD:ACE0554","Arthropoda","Insecta","Diptera","Phoridae","Metopininae","Megaselia","","103","91","91","27","11.65","0","99.58","444.04","0","445.97","32.04","0"
"AGAW2","11S","INS","BOLD:AAN5260","Arthropoda","Insecta","Diptera","Cecidomyiidae","","","","103","102","102","27","0.97","0","99.11","405.29","0","406.91","25.92","0"
"AGAW2","11S","INS","BOLD:ACN6329","Arthropoda","Insecta","Diptera","Cecidomyiidae","","","","93","90","90","29","3.23","0","98.56","438.65","0","440.19","26.62","0"
"AGAW2","11S","INS","BOLD:AAM9013","Arthropoda","Insecta","Diptera","Keroplatidae","Macrocerinae","Macrocera","Macrocera pusilla","72","66","66","30","8.33","0","99.52","431.79","0","433.51","30.27","0"
"AGAW2","11S","INS","BOLD:ABA0822","Arthropoda","Insecta","Diptera","Cecidomyiidae","","","","71","69","69","31","2.82","0","98.91","428.3","0","429.61","26.02","0"
"AGAW2","11S","INS","BOLD:ABZ5071","Arthropoda","Insecta","Diptera","Chironomidae","Chironominae","Micropsectra","Micropsectra nigripila","69","69","69","32","0","0","98.2","415.88","0","428.58","36.9","0"
"AGAW2","11S","INS","BOLD:AAN5328","Arthropoda","Insecta","Diptera","Chironomidae","Chironominae","Micropsectra","","67","67","67","33","0","0","99.13","410.72","0","413.78","35.78","0"
"AGAW2","11S","INS","BOLD:AAA7661","Arthropoda","Insecta","Diptera","Culicidae","Culicinae","Culiseta, Culex","Culiseta melanura, Culex articularis, Culex restuans, Culex acharistus","67","64","64","33","4.48","0","99.22","430.31","0","432.67","32.96","0"
"AGAW2","11S","INS","BOLD:AAH0135","Arthropoda","Insecta","Coleoptera","Phalacridae","Phalacrinae","Acylomus","Acylomus pugetanus, Acylomus ergoti","63","61","61","35","3.17","0","99.31","433.03","0","435.19","31.56","0"
"AGAW2","11S","INS","BOLD:AAG7891","Arthropoda","Insecta","Hymenoptera","Platygastridae","Scelioninae","Telenomus","Telenomus podisi","61","54","54","36","11.48","0","99.33","428.39","0","436.56","27.24","0"
"AGAW2","11S","INS","BOLD:ACP6694","Arthropoda","Insecta","Diptera","Chironomidae","Orthocladiinae","","","59","56","56","37","5.08","0","98.98","415.86","0","418.05","33.93","0"
"AGAW2","11S","INS","BOLD:AAH0256","Arthropoda","Insecta","Coleoptera","Latridiidae","Corticariinae","Corticaria, Corticarina","","53","45","45","38","15.09","0","99.11","437.51","0","442.32","31.51","0"
"AGAW2","11S","INS","BOLD:AAN5334","Arthropoda","Insecta","Diptera","Chironomidae","Chironominae","Polypedilum","","50","50","50","39","0","0","98.19","424.8","0","427.94","35.08","0"
"AGAW2","11S","INS","BOLD:AAG3236","Arthropoda","Insecta","Diptera","Phoridae","Phorinae","Conicera, Diplonevra","Diplonevra nitidula","46","41","41","40","10.87","0","99.7","448.11","0","450","31.92","0"
"AGAW2","11S","INS","BOLD:AAN5183","Arthropoda","Insecta","Diptera","Cecidomyiidae","","","","41","40","40","41","2.44","0","98.15","442.93","0","444.54","28","0"
"AGAW2","11S","INS","BOLD:AAP7030","Arthropoda","Insecta","Coleoptera","Cryptophagidae","Atomariinae","Atomaria","Atomaria ephippiata","37","36","36","42","2.7","0","99.49","433.22","0","434.73","35.94","0"
"AGAW2","11S","INS","BOLD:AAF9771","Arthropoda","Insecta","Diptera","Hybotidae","Tachydromiinae","Platypalpus","Platypalpus longiseta, Platypalpus trivialis, Platypalpus sp. 5, Platypalpus sp. 17, Platypalpus pallidiventris","36","31","31","43","13.89","0","99.47","427.42","0","428.64","34.44","0"
"AGAW2","11S","INS","BOLD:ABU5525","Arthropoda","Insecta","Diptera","Chironomidae","Orthocladiinae","Limnophyes","Limnophyes sp. 14ES","33","33","33","44","0","0","98.68","410.52","0","412.39","37.11","0"
"AGAW2","11S","INS","BOLD:ABW1379","Arthropoda","Insecta","Diptera","Chloropidae","Oscinellinae","Malloewia","Malloewia abdominalis","32","25","25","45","21.88","0","98.43","445.78","0","447.16","31.42","0"
"AGAW2","11S","INS","BOLD:ACK3431","Arthropoda","Insecta","Diptera","Cecidomyiidae","","","","29","29","29","46","0","0","99.11","428.21","0","429.72","32.3","0"
"AGAW2","11S","INS","BOLD:ABA0863","Arthropoda","Insecta","Diptera","Cecidomyiidae","","","","24","24","24","47","0","0","99.56","440.79","0","442.17","29.75","0"
"AGAW2","11S","INS","BOLD:AAL7359","Arthropoda","Insecta","Diptera","Chironomidae","Chironominae","Tanytarsus","","23","21","21","48","8.7","0","99.5","443.96","0","445.65","36.27","0"
"AGAW2","11S","INS","BOLD:ABV1194","Arthropoda","Insecta","Diptera","Chironomidae","Chironominae","Micropsectra","","21","20","20","49","4.76","0","99.62","436.43","0","437.67","36","0"
"AGAW2","11S","INS","BOLD:ACK5742","Arthropoda","Insecta","Diptera","Cecidomyiidae","","","","20","19","19","50","5","0","99.6","433.35","0","433.45","28.73","0"
"AGAW2","11S","INS","BOLD:ACI1304","Arthropoda","Insecta","Diptera","Muscidae","Muscinae","Stomoxys","Stomoxys indicus","17","17","17","51","0","0","97.51","418.76","0","422.47","32.49","0"
"AGAW2","11S","INS","BOLD:ACR3866","Arthropoda","Insecta","Psocodea","Caeciliusidae","Caeciliusinae","Valenzuela","","17","15","15","51","11.76","0","99.47","438.12","0","439.76","35.39","0"
"AGAW2","11S","INS","BOLD:AAH0134","Arthropoda","Insecta","Coleoptera","Phalacridae","Phalacrinae","Stilbus","Stilbus apicialis","16","15","15","53","6.25","0","99.1","424.94","0","426.19","32.43","0"
"AGAW2","11S","INS","BOLD:ADF3937","","","","","","","","15","15","15","54","0","0","99.24","406.4","0","418.53","31.3","0"
"AGAW2","11S","INS","BOLD:ACL1796","Arthropoda","Insecta","Lepidoptera","Ypsolophidae","Ochsenheimeriinae","Ochsenheimeria","Ochsenheimeria vacculella","14","14","14","55","0","0","99.06","407.86","0","420","32.2","0"
"AGAW2","11S","INS","BOLD:ACJ6954","Arthropoda","Insecta","Diptera","Cecidomyiidae","","","","13","13","13","56","0","0","99.68","338.23","0","422.85","26.37","0"
"AGAW2","11S","INS","BOLD:ACM0208","Arthropoda","Insecta","Diptera","Cecidomyiidae","","","","12","9","9","57","25","0","97.83","449.17","0","450","27.07","0"
"AGAW2","11S","INS","BOLD:AAG6745","Arthropoda","Insecta","Diptera","Calliphoridae","Calliphorinae","Pollenia","Pollenia pediculata","12","12","12","57","0","0","99.03","425.58","0","427.5","33.75","0"
"AGAW2","11S","INS","BOLD:AAC4403","Arthropoda","Insecta","Diptera","Chironomidae","Chironominae","Micropsectra","Micropsectra polita","12","12","12","57","0","0","97.1","428.67","0","430.25","34.98","0"
"AGAW2","11S","INS","BOLD:ACA6930","Arthropoda","Insecta","Diptera","Cecidomyiidae","","","","12","12","12","57","0","0","98.4","413","0","427.83","27.58","0"
"AGAW2","11S","INS","BOLD:AAG1711","Arthropoda","Insecta","Diptera","Muscidae","Phaoniinae","Helina","Helina depuncta","11","11","11","61","0","0","99.33","439.82","0","442.18","31.37","0"
"AGAW2","11S","INS","BOLD:AAU0577","Arthropoda","Insecta","Thysanoptera","Aeolothripidae","Aeolothripinae","Aeolothrips","Aeolothrips fasciatus","11","11","11","61","0","0","98.1","431.09","0","433.36","36.66","0"
"AGAW2","11S","INS","BOLD:AAA4222","Arthropoda","Insecta","Lepidoptera","Erebidae","Hypeninae","Hypena","Hypena scabra","9","9","9","63","0","0","99.13","390.89","0","392.78","32.17","0"
"AGAW2","11S","INS","BOLD:AAC0903","Arthropoda","Insecta","Diptera","Chironomidae","Chironominae","Chironomus","Chironomus acidophilus","9","9","9","63","0","0","99.39","435.89","0","437.33","36.38","0"
"AGAW2","11S","INS","BOLD:AAN5249","Arthropoda","Insecta","Diptera","Cecidomyiidae","","","","8","8","8","65","0","0","98.64","436.38","0","438.5","33.31","0"
"AGAW2","11S","INS","BOLD:AAN5205","Arthropoda","Insecta","Diptera","Cecidomyiidae","","","CecidInt42 sp. BOLD:AAN5205","8","7","7","65","12.5","0","99.55","448.13","0","450","31.33","0"
"AGAW2","11S","INS","BOLD:ACK3142","Arthropoda","Insecta","Diptera","Cecidomyiidae","","","","8","8","8","65","0","0","98.49","435.63","0","437.25","28.38","0"
"AGAW2","11S","INS","BOLD:AAF9014","Arthropoda","Insecta","Diptera","Limoniidae","Chioneinae","Symplecta","Symplecta cana, Symplecta sp. BOLDAAF9014","8","8","8","65","0","0","98.69","423.75","0","428.88","33.64","0"
"AGAW2","11S","INS","BOLD:AAH3664","Arthropoda","Insecta","Diptera","Cecidomyiidae","","","CecidInt3 sp. BOLD:AAH3664","7","6","6","69","14.29","0","99.62","448.43","0","449.43","27.4","0"
"AGAW2","11S","INS","BOLD:AAO0694","Arthropoda","Insecta","Coleoptera","Chrysomelidae","Criocerinae","Oulema","Oulema duftschmidi, Oulema cfr. melanopus","7","7","7","69","0","0","98.03","395.29","0","397.86","36.88","0"
"AGAW2","11S","INS","BOLD:AAG2740","Arthropoda","Insecta","Diptera","Ephydridae","Parydrinae","Philygria","Philygria debilis","7","7","7","69","0","0","98.83","439","0","442.71","32.99","0"
"AGAW2","11S","INS","BOLD:AAP8739","Arthropoda","Insecta","Diptera","Platystomatidae","Platystomatinae","Rivellia","Rivellia steyskali","7","7","7","69","0","0","99.1","439.57","0","440.57","38.16","0"
"AGAW2","11S","INS","BOLD:AAO8538","Arthropoda","Insecta","Diptera","Psychodidae","Psychodinae","Psychoda","","6","6","6","73","0","0","98.56","441.17","0","444.17","32.94","0"
"AGAW2","11S","INS","BOLD:ACE8878","Arthropoda","Insecta","Orthoptera","Acrididae","Melanoplinae","Melanoplus","Melanoplus scudderi group","6","6","6","73","0","0","97.67","443.83","0","444.67","35.37","0"
"AGAW2","11S","INS","BOLD:ACK5443","Arthropoda","Insecta","Diptera","Cecidomyiidae","","","","6","6","6","73","0","0","97.92","448.83","0","450","25.59","0"
"AGAW2","11S","INS","BOLD:ADH9867","Arthropoda","Insecta","Diptera","","","","","6","6","6","73","0","0","98.4","422.67","0","425","37.64","0"
"AGAW2","11S","INS","BOLD:ABW3206","Arthropoda","Insecta","Hymenoptera","Pteromalidae","Pteromalinae","Cecidostiba","","5","5","5","77","0","0","98.54","424","0","436.4","28.83","0"
"AGAW2","11S","INS","BOLD:AAN5220","Arthropoda","Insecta","Diptera","Cecidomyiidae","","","","5","5","5","77","0","0","98.26","410.8","0","431","25.58","0"
"AGAW2","11S","INS","BOLD:ABV9018","Arthropoda","Insecta","Diptera","Mycetophilidae","Sciophilinae","Sciophila","","5","5","5","77","0","0","99.28","445.6","0","447.6","32.44","0"
"AGAW2","11S","INS","BOLD:ABW3288","","","","","","","","5","5","5","77","0","0","99.36","427.6","0","429.6","30.23","0"
"AGAW2","11S","INS","BOLD:AAV5796","Arthropoda","Insecta","Diptera","Cecidomyiidae","","","","5","5","5","77","0","0","99.44","421","0","424.6","28.76","0"
"AGAW2","11S","INS","BOLD:AAH3720","Arthropoda","Insecta","Diptera","Cecidomyiidae","","","CecidInt34 sp. BOLD:AAH3720","5","5","5","77","0","0","98.15","404.4","0","406.2","27.75","0"
"AGAW2","11S","INS","BOLD:ABV1297","Arthropoda","Insecta","Diptera","Sciaridae","","","","5","5","5","77","0","0","98.92","421.4","0","424.4","29.81","0"
"AGAW2","11S","INS","BOLD:AAG2123","Arthropoda","Insecta","Diptera","Tachinidae","Dexiinae","Dinera","Dinera grisescens","4","4","4","84","0","0","99.08","431.5","0","433.25","31.7","0"
"AGAW2","11S","INS","BOLD:AAN6447","Arthropoda","Insecta","Diptera","Sciaridae","","Corynoptera","Corynoptera perpusilla","4","4","4","84","0","0","99.39","448.25","0","450","36.44","0"
"AGAW2","11S","INS","BOLD:ACV2677","Arthropoda","Insecta","Diptera","Cecidomyiidae","","","","4","4","4","84","0","0","97.38","449","0","450","30.11","0"
"AGAW2","11S","INS","BOLD:ABZ4994","Arthropoda","Insecta","Orthoptera","Acrididae","Melanoplinae","Melanoplus","Melanoplus gladstoni, Melanoplus sanguinipes","4","4","4","84","0","0","97.22","449.5","0","450","36.28","0"
"AGAW2","11S","INS","BOLD:ACV5892","Arthropoda","Insecta","Diptera","Cecidomyiidae","","","","4","4","4","84","0","0","98","449.25","0","450","25.44","0"
"AGAW2","11S","INS","BOLD:AAP7026","Arthropoda","Insecta","Coleoptera","Latridiidae","Corticariinae","Melanophthalma","Melanophthalma villosa","3","3","3","89","0","0","97.77","404","0","419.33","30.1","0"
"AGAW2","11S","INS","BOLD:ACW5117","Arthropoda","Insecta","Diptera","Chironomidae","Orthocladiinae","Smittia","Smittia leucopogon","3","3","3","89","0","0","99.1","435","0","436","31.18","0"
"AGAW2","11S","INS","BOLD:AAA5206","Arthropoda","Insecta","Lepidoptera","Erebidae","Herminiinae","Zanclognatha","Zanclognatha cruralis, Zanclognatha sp. 1, Zanclognatha jacchusalis, Zanclognatha minoralis, Zanclognatha minoralis sp. 1, Zanclognatha sp., Zanclognatha minoralis sp. 3, Zanclognatha gypsalis, Zanclognatha dentata, Zanclognatha theralis, Zanclognatha protumnusalis, Zanclognatha martha, Zanclognatha atrilineella, Zanclognatha obscuripennis, Zanclognatha minoralis sp. 2, Zanclognatha marcidilinea","3","3","3","89","0","0","98.68","429.33","0","430.67","31.42","0"
"AGAW2","11S","INS","BOLD:AAI1549","Arthropoda","Insecta","Hymenoptera","Braconidae","Euphorinae","Meteorus","","3","3","3","89","0","0","99.26","450.67","0","450","31.19","0"
"AGAW2","11S","INS","BOLD:AAB9332","Arthropoda","Insecta","Hemiptera","Aphididae","Aphidinae","Nasonovia","Nasonovia ribisnigri","3","3","3","89","0","0","99.85","430","0","431.67","24.6","0"
"AGAW2","11S","INS","BOLD:ABW1460","Arthropoda","Insecta","Coleoptera","Chrysomelidae","Criocerinae","Oulema","Oulema erichsonii","3","3","3","89","0","0","97.8","379","0","380.33","34.44","0"
"AGAW2","11S","INS","BOLD:AAG2511","Arthropoda","Insecta","Diptera","Anthomyiidae","Anthomyiinae","Delia","Delia platura","3","3","3","89","0","0","98.35","440","0","442","32.43","0"
"AGAW2","11S","INS","BOLD:AAP2527","Arthropoda","Insecta","Diptera","Mycetophilidae","Mycetophilinae","Brevicornu","Brevicornu fissicauda","3","3","3","89","0","0","99.33","449","0","450","36.67","0"
"AGAW2","11S","INS","BOLD:AAG6743","Arthropoda","Insecta","Diptera","Sarcophagidae","Sarcophaginae","Sarcophaga","Sarcophaga sp. 1 MG, Sarcophaga subvicina","3","3","3","89","0","0","99.7","448","0","449","35.04","0"
"AGAW2","11S","INS","BOLD:AAN9916","Arthropoda","Insecta","Coleoptera","Staphylinidae","Scydmaeninae","Stenichnus","Stenichnus scutellaris","3","3","3","89","0","0","99.76","433.33","0","435.33","27.68","0"
"AGAW2","11S","INS","BOLD:ADG4648","Arthropoda","Insecta","Diptera","Cecidomyiidae","","","","2","2","2","99","0","0","99.49","388.5","0","450","27.22","0"
"AGAW2","11S","INS","BOLD:ABX4079","Arthropoda","Insecta","Diptera","Cecidomyiidae","","","","2","2","2","99","0","0","98.47","429","0","433","27.69","0"
"AGAW2","11S","INS","BOLD:ACE4100","Arthropoda","Insecta","Orthoptera","Acrididae","Melanoplinae","Melanoplus","Melanoplus pinicola","2","2","2","99","0","0","97.99","446","0","450","35.44","0"
"AGAW2","11S","INS","BOLD:ACF4786","Arthropoda","Insecta","Hymenoptera","Braconidae","Aphidiinae","Aphidius","Aphidius urticae, Aphidius matricariae","2","2","2","99","0","0","99.62","395","0","398.5","25.6","0"
"AGAW2","11S","INS","BOLD:AAA8933","Arthropoda","Insecta","Coleoptera","Coccinellidae","Coccinellinae","Coccinella","Coccinella septempunctata","2","2","2","99","0","0","99.56","450.5","0","450","32.89","0"
"AGAW2","11S","INS","BOLD:ABA5313","Arthropoda","Insecta","Coleoptera","Staphylinidae","Aleocharinae","Amischa","Amischa analis, Amischa bifoveolata, Amischa nigrofusca","2","2","2","99","0","0","97.97","442","0","443.5","33.82","0"
"AGAW2","11S","INS","BOLD:AAY6412","Arthropoda","Insecta","Diptera","Sciaridae","","Bradysia","Bradysia urticae","2","2","2","99","0","0","99.89","448.5","0","450","32.33","0"
"AGAW2","11S","INS","BOLD:AAL7442","Arthropoda","Insecta","Diptera","Cecidomyiidae","","","","2","2","2","99","0","0","99.67","448.5","0","450","26.67","0"
"AGAW2","11S","INS","BOLD:AAB0373","Arthropoda","Insecta","Neuroptera","Chrysopidae","Chrysopinae","Chrysoperla","Chrysoperla sp. isolate 82, Chrysoperla zastrowi, Chrysoperla plorabunda, Chrysoperla sp. isolate 71, Chrysoperla agilis, Italochrysa italica, Chrysoperla sp. isolate 88, Chrysoperla lucasina, Chrysoperla carnea, Chrysoperla mediterranea, Chrysoperla pallida, Chrysoperla nipponensis, Chrysoperla sillemi","2","2","2","99","0","0","99.89","448.5","0","450","31.33","0"
"AGAW2","11S","INS","BOLD:AAD2041","Arthropoda","Insecta","Diptera","Dolichopodidae","Sciapodinae","Condylostylus","Condylostylus inermis, Condylostylus nigrofemoratus, Condylostylus connectans, Condylostylus caudatus, Condylostylus flavipes","2","2","2","99","0","0","99.61","380.5","0","382.5","35.43","0"
"AGAW2","11S","INS","BOLD:AAY6469","Arthropoda","Insecta","Diptera","Cecidomyiidae","","","","2","2","2","99","0","0","99","447.5","0","450","26.44","0"
"AGAW2","11S","INS","BOLD:ACE8875","Arthropoda","Insecta","Orthoptera","Acrididae","Melanoplinae","Melanoplus","Melanoplus bowditchi","2","2","2","99","0","0","97.33","449","0","450","36.11","0"
"AGAW2","11S","INS","BOLD:AAY6112","Arthropoda","Insecta","Diptera","Cecidomyiidae","","","","2","2","2","99","0","0","100","447.5","0","450","26.67","0"
"AGAW2","11S","INS","BOLD:AAV1136","Arthropoda","Insecta","Diptera","Scatopsidae","","","","2","2","2","99","0","0","98.4","398","0","398.5","37.03","0"
"AGAW2","11S","INS","BOLD:AAA3797","Arthropoda","Insecta","Hymenoptera","Megachilidae","Megachilinae","Megachile","Megachile sp. aff rotundata, Megachile cf. rotundata, Megachile rotundata","2","2","2","99","0","0","100","449","0","450","24.11","0"
"AGAW2","11S","INS","BOLD:ABW3687","Arthropoda","Insecta","Diptera","Lauxaniidae","Homoneurinae","Homoneura","Homoneura bispina","1","1","1","114","0","0","97.54","447","0","450","36","0"
"AGAW2","11S","INS","BOLD:AAC0022","Arthropoda","Insecta","Diptera","Chironomidae","Chironominae","Micropsectra","Micropsectra nigripila","1","1","1","114","0","0","97.32","448","0","450","36.89","0"
"AGAW2","11S","INS","BOLD:AAA3933","Arthropoda","Insecta","Lepidoptera","Erebidae","Herminiinae","Palthis","Palthis angulalis","1","1","1","114","0","0","99.55","447","0","450","32","0"
"AGAW2","11S","INS","BOLD:AAV5688","Arthropoda","Insecta","Diptera","Cecidomyiidae","","","","1","1","1","114","0","0","97.55","449","0","450","34.89","0"
"AGAW2","11S","INS","BOLD:AAG5507","Arthropoda","Insecta","Diptera","Chironomidae","Chironominae","Chironomus","Chironomus atrella","1","1","1","114","0","0","98.67","450","0","450","36.67","0"
"AGAW2","11S","INS","BOLD:AAA6560","Arthropoda","Insecta","Lepidoptera","Crambidae","Crambinae","Crambus","Crambus multilinellus, Crambus saltuellus","1","1","1","114","0","0","98.66","448","0","450","31.33","0"
"AGAW2","11S","INS","BOLD:ABA0856","Arthropoda","Insecta","Diptera","Cecidomyiidae","","","","1","1","1","114","0","0","97.77","448","0","450","29.78","0"
"AGAW2","11S","INS","BOLD:ABA6163","Arthropoda","Insecta","Psocodea","Caeciliusidae","Caeciliusinae","Valenzuela","Valenzuela piceus","1","1","1","114","0","0","99.33","450","0","450","36.89","0"
"AGAW2","11S","INS","BOLD:AAM6781","Arthropoda","Insecta","Diptera","Dolichopodidae","Neurigoninae","Neurigona","Neurigona tenuis","1","1","1","114","0","0","97.1","449","0","450","35.33","0"
"AGAW2","11S","INS","BOLD:AAD0952","Arthropoda","Insecta","Lepidoptera","Pyralidae","Chrysauginae","Galasa","Galasa nigrinodis, Galasa nigripunctalis","1","1","1","114","0","0","97.42","427","0","450","32.44","0"
"AGAW2","11S","INS","BOLD:AAB4321","Arthropoda","Insecta","Lepidoptera","Gelechiidae","Gelechiinae","Metzneria","Metzneria lappella","1","1","1","114","0","0","98.36","366","0","450","31.33","0"
"AGAW2","11S","INS","BOLD:AAG5453","Arthropoda","Insecta","Diptera","Chironomidae","Chironominae","Chironomus","Chironomus bifurcatus","1","1","1","114","0","0","98.44","448","0","450","35.11","0"
"AGAW2","11S","INS","BOLD:AAL1297","Arthropoda","Insecta","Coleoptera","Cantharidae","Cantharinae","Rhagonycha","Rhagonycha fulva","1","1","1","114","0","0","97.06","442","0","450","33.78","0"
"AGAW2","11S","INS","BOLD:AAG3273","Arthropoda","Insecta","Diptera","Phoridae","Metopininae","Megaselia","Megaselia albicaudata","1","1","1","114","0","0","98.66","448","0","450","32.67","0"
"AGAW2","11S","INS","BOLD:AAC5210","Arthropoda","Insecta","Diptera","Culicidae","Culicinae","Aedes, Aedimorphus","Aedes japonicus, Aedimorphus caecus","1","1","1","114","0","0","100","433","0","436","34.17","0"
"AGAW2","11S","INS","BOLD:AAN5255","Arthropoda","Insecta","Diptera","Cecidomyiidae","Cecidomyiinae","Dasineura","","1","1","1","114","0","0","98.22","450","0","450","26.89","0"
"AGAW2","11S","INS","BOLD:AAN5238","Arthropoda","Insecta","Diptera","Cecidomyiidae","","","","1","1","1","114","0","0","100","447","0","450","25.78","0"
"AGAW2","11S","INS","BOLD:ABW5534","Arthropoda","Insecta","Diptera","Cecidomyiidae","","","","1","1","1","114","0","0","100","449","0","450","26","0"
"AGAW2","11S","INS","BOLD:AAC3517","Arthropoda","Insecta","Hemiptera","Delphacidae","Delphacinae","Kosswigianella, Muirodelphax, Delphacodes, Nothodelphax, Elachodelphax","Muirodelphax arvensis, Delphacodes nitens, Delphacodes lutulenta, Muirodelphax nigrinota, Kosswigianella analis, Delphacodes campestris, Muirodelphax anufrievi, Delphacodes indistincta, Muirodelphax unda, Muirodelphax nr. arvensis, Delphacodes stramineosa, Nothodelphax consimilis, Muirodelphax atrolabis, Muirodelphax atralabis, Kosswigianella lutulenta","1","1","1","114","0","0","99.78","449","0","450","37.11","0"
"AGAW2","11S","INS","BOLD:ABY2946","Arthropoda","Insecta","Diptera","Tabanidae","Tabaninae","Hybomitra","Hybomitra trepida","1","1","1","114","0","0","98.89","450","0","450","33.33","0"
"AGAW2","11S","INS","BOLD:AAG2017","Arthropoda","Insecta","Neuroptera","Chrysopidae","Chrysopinae","Chrysopa","Chrysopa oculata","1","1","1","114","0","0","99.54","438","0","440","32.27","0"
"AGAW2","11S","INS","BOLD:AAA8640","Arthropoda","Insecta","Lepidoptera","Tortricidae","Olethreutinae","Eucosma, Sonia, Pelochrista","Pelochrista suadana, Sonia new sp. 1, Sonia new sp. 2, Sonia new sp. 3, Sonia paraplesiana, Sonia canadana","1","1","1","114","0","0","99.55","447","0","450","33.11","0"
"AGAW2","11S","INS","BOLD:AAH1932","Arthropoda","Insecta","Hymenoptera","Ichneumonidae","Cryptinae","Dichrogaster","Dichrogaster perlae, Dichrogaster modesta","1","1","1","114","0","0","99.78","449","0","450","26.67","0"
"AGAW2","11S","INS","BOLD:AAA2632","Arthropoda","Insecta","Lepidoptera","Noctuidae","Noctuinae","Noctua","Noctua atlantica, Noctua pronuba","1","1","1","114","0","0","98","450","0","450","31.11","0"
"AGAW2","11S","INS","BOLD:ACE7444","Arthropoda","Insecta","Hemiptera","Miridae","Mirinae, Phylinae","Adelphocoris, Megalocoleus","Adelphocoris lineolatus, Megalocoleus molliculus","1","1","1","114","0","0","99.56","450","0","450","35.11","0"
"AGAW2","11S","INS","BOLD:ACV5891","Arthropoda","Insecta","Diptera","Cecidomyiidae","","","","1","1","1","114","0","0","99.17","362","0","435","26.9","0"
"AGAW2","11S","INS","BOLD:ABA8948","Arthropoda","Insecta","Coleoptera","Chrysomelidae","","","","1","1","1","114","0","0","97.25","400","0","401","36.41","0"
"AGAW2","11S","INS","BOLD:AAU7126","Arthropoda","Insecta","Coleoptera","Cleridae","Hydnocerinae","Isohydnocera","Isohydnocera curtipennis","1","1","1","114","0","0","99.75","396","0","401","31.92","0"
"AGAW2","11S","INS","BOLD:ACI9753","Arthropoda","Insecta","Diptera","Dolichopodidae","Diaphorinae","Chrysotus","","1","1","1","114","0","0","99.02","409","0","450","32.44","0"
"AGAW2","11S","INS","BOLD:ACK9434","Arthropoda","Insecta","Hemiptera","Cicadellidae","Macropsinae","Macropsis","Macropsis ocellata","1","1","1","114","0","0","97.33","450","0","450","34","0"
"AGAW2","11S","INS","BOLD:ABA1661","Arthropoda","Insecta","Coleoptera","Chrysomelidae","Galerucinae","Longitarsus","Longitarsus pellucidus","1","1","1","114","0","0","99.55","449","0","450","35.33","0"
"AGAW2","11S","INS","BOLD:ACL7238","Arthropoda","Insecta","Diptera","Xylomyidae","","Solva","","1","1","1","114","0","0","98.22","450","0","450","35.11","0"
"AGAW2","11S","INS","BOLD:ABV9293","Arthropoda","Insecta","Diptera","Chironomidae","Chironominae","Dicrotendipes","","1","1","1","114","0","0","97.75","355","0","450","34.89","0"
"AGAW2","11S","INS","BOLD:ACE4897","Arthropoda","Insecta","Diptera","Tachinidae","Tachininae","Microphthalma","","1","1","1","114","0","0","98.42","444","0","450","28.67","0"
"AGAW2","11S","INS","BOLD:ABY5257","Arthropoda","Insecta","Lepidoptera","Noctuidae","Noctuinae","Apamea","Apamea devastator","1","1","1","114","0","0","99.03","413","0","415","29.4","0"
"AGAW2","11S","INS","BOLD:AAG5499","Arthropoda","Insecta","Diptera","Chironomidae","Orthocladiinae","Psectrocladius","Psectrocladius obvius, Psectrocladius platypus","1","1","1","114","0","0","99.78","450","0","450","36","0"
"AGAW2","11S","INS","BOLD:ACK6578","Arthropoda","Insecta","Hymenoptera","Trichogrammatidae","","","","1","1","1","114","0","0","98.1","421","0","450","25.11","0"
"AGAW2","11S","INS","BOLD:ACV3832","","","","","","","","1","1","1","114","0","0","97.05","407","0","450","36","0"
"AGAW2","11S","INS","BOLD:AAV1261","Arthropoda","Insecta","Diptera","Sciaridae","","Bradysia","Bradysia splendida","1","1","1","114","0","0","99.11","447","0","450","33.56","0"
"AGAW2","11S","INS","BOLD:ACX5531","Arthropoda","Insecta","Diptera","Chloropidae","Oscinellinae","Conioscinella","Conioscinella triorbiculata","1","1","1","114","0","0","99.22","387","0","389","30.33","0"
"AGAW2","11S","INS","BOLD:ABX2489","Arthropoda","Insecta","Coleoptera","Phalacridae","Phalacrinae","Stilbus","","1","1","1","114","0","0","97.78","450","0","450","33.33","0"
"AGAW2","11S","INS","BOLD:AAU6592","Arthropoda","Insecta","Diptera","Ceratopogonidae","Dasyheleinae","Dasyhelea","Dasyhelea arenivaga","1","1","1","114","0","0","98.44","448","0","450","35.78","0"
"AGAW2","11S","INS","BOLD:ADG7735","Arthropoda","Insecta","Diptera","Cecidomyiidae","","","","1","1","1","114","0","0","97.7","392","0","427","26.46","0"
"AGAW2","11S","INS","BOLD:AAC9062","Arthropoda","Insecta","Diptera","Culicidae","Culicinae","Ochlerotatus, Aedes","Aedes fitchii, Ochlerotatus fitchii, Aedes grossbecki, Aedes ventrovittis","1","1","1","114","0","0","98.66","447","0","450","33.56","0"
"AGAW2","11S","INS","BOLD:AAN6435","Arthropoda","Insecta","Diptera","Sciaridae","","Bradysia","Bradysia angustipennis","1","1","1","114","0","0","99.75","397","0","440","35","0"
"AGAW2","11S","INS","BOLD:ADB5456","Arthropoda","Insecta","Hymenoptera","Eulophidae","Entedoninae","Neochrysocharis","","1","1","1","114","0","0","99.51","412","0","450","24.67","0"
"AGAW2","11S","INS","BOLD:ABZ8007","Arthropoda","Insecta","Lepidoptera","Noctuidae","Cuculliinae","Cucullia","Cucullia similaris, Cucullia postera, Cucullia sp., Cucullia florea","1","1","1","114","0","0","97.1","448","0","450","30.89","0"
"AGAW2","11S","INS","BOLD:AAH3225","Arthropoda","Insecta","Psocodea","Psocidae","Psocinae","Metylophorus","Metylophorus novaescotiae","1","1","1","114","0","0","97.78","450","0","450","34.44","0"
"AGAW2","11S","INS","BOLD:ABV1422","Arthropoda","Insecta","Diptera","Cecidomyiidae","","","","1","1","1","114","0","0","98.44","448","0","450","27.11","0"
"AGAW2","11S","INS","BOLD:ADH8687","Arthropoda","Insecta","Coleoptera","","","","","1","1","1","114","0","0","99.77","439","0","450","35.11","0"
"AGAW2","11S","INS","BOLD:ABW7897","Arthropoda","Insecta","Diptera","Cecidomyiidae","","","","1","1","1","114","0","0","97.33","450","0","450","29.78","0"
"AGAW2","11S","INS","BOLD:ACC7241","","","","","","","","1","1","1","114","0","0","97.11","346","0","352","28.69","0"
"AGAW2","11S","NAR","BOLD:AAI9663","Mollusca","Gastropoda","Stylommatophora","Agriolimacidae",NA,"Deroceras","Deroceras reticulatum","1","1","1","1","0","0","99.51","411","0","450","36.44","0"
"AGAW2","11S","NIN","BOLD:AAH6630","Arthropoda","Collembola","Entomobryomorpha","Entomobryidae","Entomobryinae","Entomobrya","Entomobrya unostrigata","811","707","707","1","12.82","0","99.34","436.05","0","437.27","39.08","0"
"AGAW2","11S","NIN","BOLD:ABW2639","Arthropoda","Arachnida","Trombidiformes","Anystidae","","","","264","207","207","2","21.59","0","99.57","445.91","0","444.35","31.42","0"
"AGAW2","11S","NIN","BOLD:AAU6252","Arthropoda","Arachnida","Trombidiformes","Bdellidae","","Cyta","","154","143","143","3","7.14","0","99.24","423.92","0","425.35","31.32","0"
"AGAW2","11S","NIN","BOLD:ACZ1295","Arthropoda","Arachnida","Trombidiformes","Anystidae","","","","46","46","46","4","0","0","99.37","374.91","0","383.02","31.32","0"
"AGAW2","11S","NIN","BOLD:AAA8820","Arthropoda","Collembola","Entomobryomorpha","Entomobryidae","Lepidocyrtinae","Lepidocyrtus","Lepidocyrtus cyaneus","41","40","40","5","2.44","0","98.87","436.46","0","438.32","43.27","0"
"AGAW2","11S","NIN","BOLD:AAB7914","Arthropoda","Collembola","Symphypleona","Bourletiellidae","","Bourletiella","","31","26","26","6","16.13","0","99.41","438.52","0","440.23","37.99","0"
"AGAW2","11S","NIN","BOLD:AAM7961","Arthropoda","Arachnida","Trombidiformes","Anystidae","","","","26","24","24","7","7.69","0","99.44","441.35","0","439.08","31.32","0"
"AGAW2","11S","NIN","BOLD:AAZ2181","Arthropoda","Collembola","Symphypleona","Bourletiellidae","","","","14","11","11","8","21.43","0","97.41","449.29","0","450","39.65","0"
"AGAW2","11S","NIN","BOLD:ACY2550","Arthropoda","Arachnida","Trombidiformes","Bdellidae","","Cyta","","10","10","10","9","0","0","99.74","388.7","0","408.6","30.71","0"
"AGAW2","11S","NIN","BOLD:ACC0359","Arthropoda","Collembola","Symphypleona","Bourletiellidae","","Bourletiella","","5","5","5","10","0","0","99.3","437.6","0","441.6","37.58","0"
"AGAW2","11S","NIN","BOLD:ACI9974","Arthropoda","Arachnida","Trombidiformes","Tydeidae","","","","3","3","3","11","0","0","98.82","424.33","0","450","40.89","0"
"AGAW2","11S","NIN","BOLD:AAO8147","Arthropoda","Arachnida","Trombidiformes","Eupodidae","","Eupodes","","3","3","3","11","0","0","98.88","448.33","0","450","28.81","0"
"AGAW2","11S","NIN","BOLD:AAH6568","Arthropoda","Arachnida","Sarcoptiformes","Tectocepheidae","","Tectocepheus","","2","2","2","13","0","0","98.46","421","0","431.5","37.39","0"
"AGAW2","11S","NIN","BOLD:ACX8969","Arthropoda","Arachnida","Trombidiformes","Eupodidae","","","","2","2","2","13","0","0","98.12","432.5","0","433.5","28.24","0"
"AGAW2","11S","NIN","BOLD:AAA7242","Arthropoda","Collembola","Entomobryomorpha","Entomobryidae","Seirinae, Entomobryinae","Entomobrya, Lepidosira","Entomobrya multifasciata, Lepidosira sp.","1","1","1","15","0","0","98.44","449","0","450","39.78","0"
"AGAW2","11S","NIN","BOLD:AAI6251","Arthropoda","Arachnida","Araneae","Dictynidae","","Emblyna","Emblyna hentzi","1","1","1","15","0","0","98.66","447","0","450","32","0"
"AGAW2","11W","BAC","TAX:175768","Proteobacteria","Alphaproteobacteria","Rickettsiales","Rickettsiaceae",NA,"Wolbachia","Wolbachia pipientis","12","10","10","1","16.67","0","99.67","449","0","450","39.8","0"
"AGAW2","11W","BAC","TAX:392874","Proteobacteria","Alphaproteobacteria","Rickettsiales","Rickettsiaceae",NA,"Wolbachia","Wolbachia endosymbiont of Hymenoptera","7","6","6","2","14.29","0","98.89","449","0","450","40.29","0"
"AGAW2","11W","BAC","TAX:376688","Proteobacteria","Alphaproteobacteria","Rickettsiales","Rickettsiaceae",NA,"Wolbachia","Wolbachia sp. elachBioLep01 BioLep373","6","5","5","3","16.67","0","99","449.33","0","450","40.48","0"
"AGAW2","11W","BAC","TAX:375847","Proteobacteria","Alphaproteobacteria","Rickettsiales","Rickettsiaceae",NA,"Wolbachia","Wolbachia sp. Hymenoptera_ASGLE1705-10","5","5","5","4","0","0","99.09","412.6","0","413.4","39.85","0"
"AGAW2","11W","BAC","TAX:91391","Proteobacteria","Alphaproteobacteria","Rickettsiales","",NA,"","","3","3","3","5","0","0","98.71","415","0","430.67","39.69","0"
"AGAW2","11W","BAC","TAX:392873","Proteobacteria","Alphaproteobacteria","Rickettsiales","Rickettsiaceae",NA,"Wolbachia","Wolbachia endosymbiont of Scrapter algoensis","2","2","2","6","0","0","98.27","405","0","408","39.1","0"
"AGAW2","11W","BAC","TAX:375845","Proteobacteria","Alphaproteobacteria","Rickettsiales","Rickettsiaceae",NA,"Wolbachia","Wolbachia sp. Hymenoptera_ASGLE1440-10","1","1","1","7","0","0","99.51","411","0","418","38.52","0"
"AGAW2","11W","BAC","TAX:392822","Proteobacteria","Alphaproteobacteria","Rickettsiales","Rickettsiaceae",NA,"Wolbachia","Wolbachia endosymbiont of Caligo telamonius","1","1","1","7","0","0","99.33","448","0","450","40.44","0"
"AGAW2","11W","BAC","TAX:375772","Proteobacteria","Alphaproteobacteria","Rickettsiales","Rickettsiaceae",NA,"Wolbachia","Wolbachia sp. Leptogenys JCR25_ASANE483-10","1","1","1","7","0","0","97.99","448","0","450","40.22","0"
"AGAW2","11W","BAC","TAX:392914","Proteobacteria","Alphaproteobacteria","Rickettsiales","Rickettsiaceae",NA,"Wolbachia","Wolbachia endosymbiont of chalJanzen01 Janzen01","1","1","1","7","0","0","99.11","447","0","450","40","0"
"AGAW2","11W","BAC","TAX:375762","Proteobacteria","Alphaproteobacteria","Rickettsiales","Rickettsiaceae",NA,"Wolbachia","Wolbachia sp. Myrmicinae_ACGAN556-09","1","1","1","7","0","0","99.55","443","0","445","40.45","0"
"AGAW2","11W","CHO","no match","no match","no match","no match","no match","no match","no match","no match","no match","no match","no match","no match","no match","no match","no match","no match","no match","no match","no match","no match"
"AGAW2","11W","INS","BOLD:AAK5928","Arthropoda","Insecta","Coleoptera","Chrysomelidae","Criocerinae","Oulema","Oulema melanopus","11300","8500","8500","1","24.78","0","99.57","439.57","0","440.91","36.47","0"
"AGAW2","11W","INS","BOLD:AAA3181","Arthropoda","Insecta","Diptera","Muscidae","Muscinae","Stomoxys","Stomoxys calcitrans","7418","6081","6081","2","18.02","0","98.92","428.36","0","432.12","32.97","0"
"AGAW2","11W","INS","BOLD:AAG1711","Arthropoda","Insecta","Diptera","Muscidae","Phaoniinae","Helina","Helina depuncta","5471","4449","4449","3","18.68","0","99.48","434.17","0","436.45","31.35","0"
"AGAW2","11W","INS","BOLD:AAL1297","Arthropoda","Insecta","Coleoptera","Cantharidae","Cantharinae","Rhagonycha","Rhagonycha fulva","3528","3204","3204","4","9.18","0","99.1","412.24","0","416.41","33.44","0"
"AGAW2","11W","INS","BOLD:AAA4555","Arthropoda","Insecta","Orthoptera","Acrididae","Melanoplinae","Melanoplus","Melanoplus stupefactus, Melanoplus angustipennis, Melanoplus ponderosus, Melanoplus bruneri, Melanoplus frigidus, Melanoplus scapularis, Melanoplus cinereus, Melanoplus madeleineae, Melanoplus sanguinipes, Melanoplus nigrescens group, Melanoplus tunicae, Melanoplus walshi, Melanoplus femurrubrum, Melanoplus dawsoni, Melanoplus sp., Melanoplus fasciatus, Melanoplus bowditchi, Melanoplus stonei, Melanoplus bivittatus, Melanoplus huroni, Melanoplus thomasi, Melanoplus borealis, Melanoplus packardii","3351","2566","2566","5","23.43","0","99.46","447.24","0","448.43","36.18","0"
"AGAW2","11W","INS","BOLD:AAC0903","Arthropoda","Insecta","Diptera","Chironomidae","Chironominae","Chironomus","Chironomus acidophilus","2728","2096","2096","6","23.17","0","99.55","437.59","0","439.57","36.33","0"
"AGAW2","11W","INS","BOLD:AAG5453","Arthropoda","Insecta","Diptera","Chironomidae","Chironominae","Chironomus","Chironomus bifurcatus","2047","1520","1520","7","25.74","0","98.02","441.78","0","445.16","35.03","0"
"AGAW2","11W","INS","BOLD:AAA2518","Arthropoda","Insecta","Lepidoptera","Crambidae","Crambinae","Chrysoteuchia","Chrysoteuchia topiaria, Chrysoteuchia topiarius","1785","1520","1520","8","14.85","0","99.28","429.68","0","431.59","35.27","0"
"AGAW2","11W","INS","BOLD:AAV6112","Arthropoda","Insecta","Diptera","Chloropidae","Oscinellinae","Oscinella","Oscinella sp. 2 AGT, Oscinella sp. TAW6","1365","1056","1056","9","22.64","0","99.62","441.82","0","442.89","33.22","0"
"AGAW2","11W","INS","BOLD:AAU6681","Arthropoda","Insecta","Diptera","Tipulidae","Tipulinae","Nephrotoma","Nephrotoma cornicina","1017","820","820","10","19.37","0","99.67","442.61","0","443.83","36.32","0"
"AGAW2","11W","INS","BOLD:ADG1318","","","","","","","","929","783","783","11","15.72","0","99.56","437.61","0","439.96","25.03","0"
"AGAW2","11W","INS","BOLD:ABY7666","Arthropoda","Insecta","Diptera","Culicidae","Culicinae","Culex","Culex territans","690","527","527","12","23.62","0","98.68","445.6","0","447.14","33.57","0"
"AGAW2","11W","INS","BOLD:AAN6154","Arthropoda","Insecta","Coleoptera","Latridiidae","Corticariinae","Melanophthalma","Melanophthalma inermis","651","540","540","13","17.05","0","99.1","437.13","0","438.59","31.26","0"
"AGAW2","11W","INS","BOLD:AAH3661","Arthropoda","Insecta","Diptera","Cecidomyiidae","","","","561","503","503","14","10.34","0","99.33","428.7","0","430.66","25.25","0"
"AGAW2","11W","INS","BOLD:AAA4178","Arthropoda","Insecta","Lepidoptera","Geometridae","Larentiinae","Eupithecia","Eupithecia miserulata, Eupithecia chlorofasciata","467","381","381","15","18.42","0","99.58","444.76","0","446.51","31.39","0"
"AGAW2","11W","INS","BOLD:ABW1379","Arthropoda","Insecta","Diptera","Chloropidae","Oscinellinae","Malloewia","Malloewia abdominalis","453","345","345","16","23.84","0","98.53","445.06","0","446.65","31.48","0"
"AGAW2","11W","INS","BOLD:AAG8493","Arthropoda","Insecta","Diptera","Drosophilidae","Drosophilinae","Scaptomyza","Scaptomyza pallida","417","413","413","17","0.96","0","99.14","395.14","0","397.01","33.03","0"
"AGAW2","11W","INS","BOLD:AAN5242","Arthropoda","Insecta","Diptera","Cecidomyiidae","","","","415","415","415","18","0","0","98.57","409.19","0","411.24","28.23","0"
"AGAW2","11W","INS","BOLD:ACU7906","Arthropoda","Insecta","Diptera","Chironomidae","Orthocladiinae","","","356","356","356","19","0","0","98.98","407.49","0","410.2","35.99","0"
"AGAW2","11W","INS","BOLD:AAN5162","Arthropoda","Insecta","Diptera","Ceratopogonidae","Dasyheleinae","Dasyhelea","","353","315","315","20","10.76","0","99.5","431.2","0","433.37","33.75","0"
"AGAW2","11W","INS","BOLD:AAN5661","Arthropoda","Insecta","Diptera","Chloropidae","Oscinellinae","Olcella","Olcella parva","345","281","281","21","18.55","0","98.91","426.59","0","427.79","31.18","0"
"AGAW2","11W","INS","BOLD:ACY6524","Arthropoda","Insecta","Diptera","Chironomidae","Orthocladiinae","Psectrocladius","Psectrocladius cf. limbatellus","339","334","334","22","1.47","0","98.57","419.31","0","425.94","36.03","0"
"AGAW2","11W","INS","BOLD:AAV5575","Arthropoda","Insecta","Diptera","Cecidomyiidae","","","CecidInt30 sp. BOLD:AAV5575","307","230","230","23","25.08","0","99.66","443.43","0","444.26","25.88","0"
"AGAW2","11W","INS","BOLD:AAA1513","Arthropoda","Insecta","Lepidoptera","Plutellidae","Plutellinae","Plutella, Rhigognostis","Plutella xylostella, Rhigognostis sp.","292","280","280","24","4.11","0","99.12","421.99","0","423.89","31.98","0"
"AGAW2","11W","INS","BOLD:ABW7897","Arthropoda","Insecta","Diptera","Cecidomyiidae","","","","270","202","202","25","25.19","0","99.71","443.56","0","444.89","29.66","0"
"AGAW2","11W","INS","BOLD:AAA7067","Arthropoda","Insecta","Diptera","Culicidae","Culicinae","Aedes, Aedimorphus","Aedimorphus vexans, Aedes vexans","261","247","247","26","5.36","0","99.1","411.17","0","412.98","32.2","0"
"AGAW2","11W","INS","BOLD:AAA6560","Arthropoda","Insecta","Lepidoptera","Crambidae","Crambinae","Crambus","Crambus multilinellus, Crambus saltuellus","244","204","204","27","16.39","0","99.36","436.64","0","438.09","30.64","0"
"AGAW2","11W","INS","BOLD:AAN5183","Arthropoda","Insecta","Diptera","Cecidomyiidae","","","","194","174","174","28","10.31","0","98.37","442.34","0","443.55","27.94","0"
"AGAW2","11W","INS","BOLD:ABY5318","Arthropoda","Insecta","Diptera","Chloropidae","Oscinellinae","Incertella, Rhopalopterum","Rhopalopterum umbrosum, Rhopalopterum luteiceps, Incertella bispina, Rhopalopterum painteri","183","150","150","29","18.03","0","99.44","444.47","0","445.52","32.14","0"
"AGAW2","11W","INS","BOLD:AAN5309","Arthropoda","Insecta","Diptera","Chironomidae","Chironominae","","","170","160","160","30","5.88","0","97.44","425.96","0","427.91","36.61","0"
"AGAW2","11W","INS","BOLD:AAH0135","Arthropoda","Insecta","Coleoptera","Phalacridae","Phalacrinae","Acylomus","Acylomus pugetanus, Acylomus ergoti","151","137","137","31","9.27","0","99.19","430.92","0","433.25","31.55","0"
"AGAW2","11W","INS","BOLD:ACR3866","Arthropoda","Insecta","Psocodea","Caeciliusidae","Caeciliusinae","Valenzuela","","142","126","126","32","11.27","0","99.55","435.35","0","436.71","35.42","0"
"AGAW2","11W","INS","BOLD:ACC8079","Arthropoda","Insecta","Diptera","Cecidomyiidae","","","","140","98","98","33","30","0","99.54","447.06","0","448.59","29.3","0"
"AGAW2","11W","INS","BOLD:AAA7661","Arthropoda","Insecta","Diptera","Culicidae","Culicinae","Culiseta, Culex","Culiseta melanura, Culex articularis, Culex restuans, Culex acharistus","136","126","126","34","7.35","0","99.37","435.67","0","437.79","32.79","0"
"AGAW2","11W","INS","BOLD:ACA2933","Arthropoda","Insecta","Psocodea","Stenopsocidae","","Graphopsocus","Graphopsocus cruciatus","129","97","97","35","24.81","0","99.3","440.88","0","441.81","38.6","0"
"AGAW2","11W","INS","BOLD:AAG9626","Arthropoda","Insecta","Diptera","Dolichopodidae","","","","111","107","107","36","3.6","0","98.09","417.07","0","419.45","36.43","0"
"AGAW2","11W","INS","BOLD:ABA1661","Arthropoda","Insecta","Coleoptera","Chrysomelidae","Galerucinae","Longitarsus","Longitarsus pellucidus","90","82","82","37","8.89","0","99.52","431.56","0","432.93","35.62","0"
"AGAW2","11W","INS","BOLD:ACI1304","Arthropoda","Insecta","Diptera","Muscidae","Muscinae","Stomoxys","Stomoxys indicus","89","83","83","38","6.74","0","97.71","423","0","425.93","32.65","0"
"AGAW2","11W","INS","BOLD:AAM9013","Arthropoda","Insecta","Diptera","Keroplatidae","Macrocerinae","Macrocera","Macrocera pusilla","82","77","77","39","6.1","0","99.43","428.62","0","430.22","30.21","0"
"AGAW2","11W","INS","BOLD:ADF6661","Arthropoda","Insecta","Diptera","Cecidomyiidae","","","","80","61","61","40","23.75","0","99.04","416.18","0","443.36","29.96","0"
"AGAW2","11W","INS","BOLD:AAP8739","Arthropoda","Insecta","Diptera","Platystomatidae","Platystomatinae","Rivellia","Rivellia steyskali","78","59","59","41","24.36","0","99.33","437.85","0","440.06","37.93","0"
"AGAW2","11W","INS","BOLD:AAG2740","Arthropoda","Insecta","Diptera","Ephydridae","Parydrinae","Philygria","Philygria debilis","74","71","71","42","4.05","0","98.36","428.65","0","431.42","32.88","0"
"AGAW2","11W","INS","BOLD:AAN5199","Arthropoda","Insecta","Diptera","Cecidomyiidae","","","CecidInt20 sp. BOLD:AAN5199","69","56","56","43","18.84","0","99.74","445.43","0","447.36","26.51","0"
"AGAW2","11W","INS","BOLD:ACX6055","Arthropoda","Insecta","Diptera","Phoridae","Metopininae","Megaselia","Megaselia sp. AK-2010-2, Megaselia sp. BOLD:ACX6055","69","57","57","43","17.39","0","99.55","435.23","0","437.13","33.13","0"
"AGAW2","11W","INS","BOLD:AAM9347","Arthropoda","Insecta","Diptera","Phoridae","Metopininae","Megaselia","","67","66","66","45","1.49","0","97.9","398.75","0","400.49","32.82","0"
"AGAW2","11W","INS","BOLD:AAG7891","Arthropoda","Insecta","Hymenoptera","Platygastridae","Scelioninae","Telenomus","Telenomus podisi","67","63","63","45","5.97","0","99.11","421.69","0","429.82","26.86","0"
"AGAW2","11W","INS","BOLD:AAU6592","Arthropoda","Insecta","Diptera","Ceratopogonidae","Dasyheleinae","Dasyhelea","Dasyhelea arenivaga","65","65","65","47","0","0","99.16","396","0","398.14","34.59","0"
"AGAW2","11W","INS","BOLD:AAN5238","Arthropoda","Insecta","Diptera","Cecidomyiidae","","","","57","49","49","48","14.04","0","99.55","423.12","0","424.84","25.44","0"
"AGAW2","11W","INS","BOLD:AAG2868","Arthropoda","Insecta","Hemiptera","Cicadellidae","Typhlocybinae","Empoasca","Empoasca sp., Empoasca fabae","55","39","39","49","29.09","0","99.73","445.84","0","446.62","32.26","0"
"AGAW2","11W","INS","BOLD:AAH0256","Arthropoda","Insecta","Coleoptera","Latridiidae","Corticariinae","Corticaria, Corticarina","","52","50","50","50","3.85","0","98.74","420.9","0","426.79","31.47","0"
"AGAW2","11W","INS","BOLD:AAN5260","Arthropoda","Insecta","Diptera","Cecidomyiidae","","","","50","50","50","51","0","0","99.15","401.56","0","403.28","25.9","0"
"AGAW2","11W","INS","BOLD:ABA0822","Arthropoda","Insecta","Diptera","Cecidomyiidae","","","","43","38","38","52","11.63","0","98.92","429.91","0","431","25.99","0"
"AGAW2","11W","INS","BOLD:AAA4781","Arthropoda","Insecta","Hymenoptera","Braconidae","Microgastrinae","Glyptapanteles","Glyptapanteles carlossarmientoi","33","32","32","53","3.03","0","99.29","427.67","0","430.24","24.11","0"
"AGAW2","11W","INS","BOLD:AAN6447","Arthropoda","Insecta","Diptera","Sciaridae","","Corynoptera","Corynoptera perpusilla","27","23","23","54","14.81","0","99.32","447.22","0","448.56","36.42","0"
"AGAW2","11W","INS","BOLD:AAV5573","Arthropoda","Insecta","Diptera","Cecidomyiidae","","","","26","22","22","55","15.38","0","99.16","446.27","0","447.12","27.42","0"
"AGAW2","11W","INS","BOLD:ABA0863","Arthropoda","Insecta","Diptera","Cecidomyiidae","","","","24","23","23","56","4.17","0","99.55","439.54","0","440.83","29.81","0"
"AGAW2","11W","INS","BOLD:AAD1485","Arthropoda","Insecta","Diptera","Chironomidae","Chironominae","Paratanytarsus","Paratanytarsus grimmii, Paratanytarsus sp.","17","16","16","57","5.88","0","99.62","440.24","0","441.18","33.75","0"
"AGAW2","11W","INS","BOLD:AAF9771","Arthropoda","Insecta","Diptera","Hybotidae","Tachydromiinae","Platypalpus","Platypalpus longiseta, Platypalpus trivialis, Platypalpus sp. 5, Platypalpus sp. 17, Platypalpus pallidiventris","15","10","10","58","33.33","0","99.78","449.27","0","450","35.26","0"
"AGAW2","11W","INS","BOLD:AAC4197","Arthropoda","Insecta","Diptera","Chironomidae","Orthocladiinae","Paraphaenocladius","Paraphaenocladius impensus","15","13","13","58","13.33","0","99.46","443.33","0","444.4","34.97","0"
"AGAW2","11W","INS","BOLD:AAN5330","Arthropoda","Insecta","Diptera","Chironomidae","Orthocladiinae","Corynoneura","Corynoneura scutellata","14","14","14","60","0","0","98.8","415.5","0","420.29","33.84","0"
"AGAW2","11W","INS","BOLD:ADH8410","","","","","","","","12","12","12","61","0","0","98.34","414.83","0","416.92","33.64","0"
"AGAW2","11W","INS","BOLD:AAY6730","Arthropoda","Insecta","Hemiptera","Derbidae","Cedusinae","Cedusa","Cedusa incisa","12","12","12","61","0","0","99.21","433.92","0","435.67","35.55","0"
"AGAW2","11W","INS","BOLD:AAD7604","Arthropoda","Insecta","Coleoptera","Coccinellidae","Coccinellinae","Coleomegilla","Coleomegilla maculata","12","12","12","61","0","0","99.19","428","0","431.17","31.78","0"
"AGAW2","11W","INS","BOLD:ADI0836","","","","","","","","12","12","12","61","0","0","98.22","370.42","0","372.33","33.06","0"
"AGAW2","11W","INS","BOLD:AAG6745","Arthropoda","Insecta","Diptera","Calliphoridae","Calliphorinae","Pollenia","Pollenia pediculata","11","11","11","65","0","0","99.13","432.18","0","434.27","33.66","0"
"AGAW2","11W","INS","BOLD:AAN5255","Arthropoda","Insecta","Diptera","Cecidomyiidae","Cecidomyiinae","Dasineura","","11","10","10","65","9.09","0","98.77","422.45","0","441.82","26.72","0"
"AGAW2","11W","INS","BOLD:ADF3937","","","","","","","","11","11","11","65","0","0","98.37","393.82","0","398.27","31.18","0"
"AGAW2","11W","INS","BOLD:AAB8143","Arthropoda","Insecta","Hemiptera","Nabidae","Nabinae","Nabis","Nabis inscriptus, Nabis americoferus, Nabis alternatus, Nabis sp.","10","10","10","68","0","0","99.26","441.9","0","443.8","37.17","0"
"AGAW2","11W","INS","BOLD:AAO0694","Arthropoda","Insecta","Coleoptera","Chrysomelidae","Criocerinae","Oulema","Oulema duftschmidi, Oulema cfr. melanopus","10","10","10","68","0","0","97.88","419.8","0","421.7","37.02","0"
"AGAW2","11W","INS","BOLD:AAP6882","Arthropoda","Insecta","Diptera","Chironomidae","Chironominae","Dicrotendipes","","10","10","10","68","0","0","98.01","418","0","427.6","35.95","0"
"AGAW2","11W","INS","BOLD:AAO4092","Arthropoda","Insecta","Psocodea","Caeciliusidae","Caeciliusinae","Valenzuela","Valenzuela burmeisteri","8","8","8","71","0","0","99.3","421.5","0","423.38","34.98","0"
"AGAW2","11W","INS","BOLD:ACV2677","Arthropoda","Insecta","Diptera","Cecidomyiidae","","","","7","7","7","72","0","0","97.8","448.43","0","450","30.13","0"
"AGAW2","11W","INS","BOLD:AAO7847","Arthropoda","Insecta","Hymenoptera","Braconidae","Microgastrinae","Choeras","","6","6","6","73","0","0","98.04","438","0","439.83","24.73","0"
"AGAW2","11W","INS","BOLD:ABZ4994","Arthropoda","Insecta","Orthoptera","Acrididae","Melanoplinae","Melanoplus","Melanoplus gladstoni, Melanoplus sanguinipes","6","6","6","73","0","0","97.18","449.17","0","450","36.26","0"
"AGAW2","11W","INS","BOLD:ADH8687","Arthropoda","Insecta","Coleoptera","","","","","6","5","5","73","16.67","0","99.85","437.5","0","446.83","35.09","0"
"AGAW2","11W","INS","BOLD:AAA4222","Arthropoda","Insecta","Lepidoptera","Erebidae","Hypeninae","Hypena","Hypena scabra","5","5","5","76","0","0","99.15","387.8","0","390.2","31.9","0"
"AGAW2","11W","INS","BOLD:ABY1179","Arthropoda","Insecta","Hymenoptera","Sierolomorphidae","","Sierolomorpha","","5","5","5","76","0","0","99.91","449","0","450","28.27","0"
"AGAW2","11W","INS","BOLD:AAC3517","Arthropoda","Insecta","Hemiptera","Delphacidae","Delphacinae","Kosswigianella, Muirodelphax, Delphacodes, Nothodelphax, Elachodelphax","Muirodelphax arvensis, Delphacodes nitens, Delphacodes lutulenta, Muirodelphax nigrinota, Kosswigianella analis, Delphacodes campestris, Muirodelphax anufrievi, Delphacodes indistincta, Muirodelphax unda, Muirodelphax nr. arvensis, Delphacodes stramineosa, Nothodelphax consimilis, Muirodelphax atrolabis, Muirodelphax atralabis, Kosswigianella lutulenta","4","4","4","78","0","0","99.72","449.25","0","450","37.17","0"
"AGAW2","11W","INS","BOLD:AAE7186","Arthropoda","Insecta","Hymenoptera","Braconidae","Aphidiinae","Praon","Praon sp. 1 MM-2013, Praon volucre, Praon bicolor","4","3","3","78","25","0","99.88","440","0","450","25.89","0"
"AGAW2","11W","INS","BOLD:ACD3276","Arthropoda","Insecta","Diptera","Cecidomyiidae","","","","4","4","4","78","0","0","99.41","421","0","422.5","27.96","0"
"AGAW2","11W","INS","BOLD:AAG4905","Arthropoda","Insecta","Diptera","Mycetophilidae","Mycetophilinae","Mycetophila","","4","4","4","78","0","0","99.68","420.75","0","422","35.3","0"
"AGAW2","11W","INS","BOLD:ABV1203","Arthropoda","Insecta","Diptera","Chironomidae","Orthocladiinae","","","4","4","4","78","0","0","99.77","447","0","447.75","34.34","0"
"AGAW2","11W","INS","BOLD:AAU6966","Arthropoda","Insecta","Coleoptera","Throscidae","Throscinae","Trixagus","Trixagus chevrolati","4","4","4","78","0","0","99.76","412.5","0","450","39.78","0"
"AGAW2","11W","INS","BOLD:ABV1337","Arthropoda","Insecta","Diptera","Cecidomyiidae","","","","4","3","3","78","25","0","98.96","438.5","0","438.5","28.2","0"
"AGAW2","11W","INS","BOLD:AAA3453","Arthropoda","Insecta","Diptera","Anthomyiidae","Anthomyiinae","Delia","","3","3","3","85","0","0","98.98","426.33","0","430","34.57","0"
"AGAW2","11W","INS","BOLD:AAN5205","Arthropoda","Insecta","Diptera","Cecidomyiidae","","","CecidInt42 sp. BOLD:AAN5205","3","3","3","85","0","0","98.88","448","0","450","31.56","0"
"AGAW2","11W","INS","BOLD:AAH0134","Arthropoda","Insecta","Coleoptera","Phalacridae","Phalacrinae","Stilbus","Stilbus apicialis","3","3","3","85","0","0","99.93","424.67","0","426.33","32.09","0"
"AGAW2","11W","INS","BOLD:ABA6032","Arthropoda","Insecta","Hymenoptera","Platygastridae","Scelioninae","Scelio","","3","3","3","85","0","0","99.28","416.67","0","417","26.68","0"
"AGAW2","11W","INS","BOLD:AAG2123","Arthropoda","Insecta","Diptera","Tachinidae","Dexiinae","Dinera","Dinera grisescens","3","3","3","85","0","0","99.55","445","0","446","32.29","0"
"AGAW2","11W","INS","BOLD:AAH4135","Arthropoda","Insecta","Diptera","Chloropidae","Chloropinae","Thaumatomyia","Thaumatomyia glabra, Thaumatomyia grata, Thaumatomyia sp. 1 TAW","2","2","2","90","0","0","99.06","429.5","0","433","31.73","0"
"AGAW2","11W","INS","BOLD:AAA8933","Arthropoda","Insecta","Coleoptera","Coccinellidae","Coccinellinae","Coccinella","Coccinella septempunctata","2","2","2","90","0","0","99.22","448","0","450","33.11","0"
"AGAW2","11W","INS","BOLD:AAB4657","Arthropoda","Insecta","Diptera","Chironomidae","Chironominae","Chironomus","Chironomus maturus","2","2","2","90","0","0","99.22","450","0","450","37.67","0"
"AGAW2","11W","INS","BOLD:AAF9949","Arthropoda","Insecta","Hemiptera","Miridae","Mirinae","Trigonotylus","Trigonotylus pulchellus, Trigonotylus americanus, Trigonotylus psammaecolor, Trigonotylus caelestialium, Trigonotylus viridis, Trigonotylus ruficornis","2","2","2","90","0","0","100","419","0","421.5","34.38","0"
"AGAW2","11W","INS","BOLD:ACL4165","Arthropoda","Insecta","Diptera","Cecidomyiidae","","","","2","2","2","90","0","0","98","409.5","0","411","28.72","0"
"AGAW2","11W","INS","BOLD:ACE7444","Arthropoda","Insecta","Hemiptera","Miridae","Mirinae, Phylinae","Adelphocoris, Megalocoleus","Adelphocoris lineolatus, Megalocoleus molliculus","2","2","2","90","0","0","100","449.5","0","450","34.67","0"
"AGAW2","11W","INS","BOLD:AAU9449","Arthropoda","Insecta","Hymenoptera","Braconidae","Alysiinae","","","2","2","2","90","0","0","97.22","449","0","450","26","0"
"AGAW2","11W","INS","BOLD:AAL2750","Arthropoda","Insecta","Diptera","Muscidae","Phaoniinae","Phaonia","Phaonia tuguriorum","2","2","2","90","0","0","99.53","429","0","432.5","31.86","0"
"AGAW2","11W","INS","BOLD:AAA3867","Arthropoda","Insecta","Lepidoptera","Noctuidae","Noctuinae","Apamea","Apamea amputatrix","2","2","2","90","0","0","98.71","415","0","416","33.38","0"
"AGAW2","11W","INS","BOLD:ABA0858","Arthropoda","Insecta","Diptera","Cecidomyiidae","","","","2","2","2","90","0","0","100","367","0","370","25.13","0"
"AGAW2","11W","INS","BOLD:AAH1772","Arthropoda","Insecta","Hymenoptera","Ichneumonidae","Orthocentrinae","Helictes","Helictes borealis","2","2","2","90","0","0","97.31","427.5","0","450","27.67","0"
"AGAW2","11W","INS","BOLD:ACK3142","Arthropoda","Insecta","Diptera","Cecidomyiidae","","","","2","2","2","90","0","0","99.16","423.5","0","424","28.14","0"
"AGAW2","11W","INS","BOLD:ABZ3101","Arthropoda","Insecta","Hymenoptera","Braconidae","Aphidiinae","Aphidius","","2","2","2","90","0","0","99.2","439","0","450","25.67","0"
"AGAW2","11W","INS","BOLD:AAU0577","Arthropoda","Insecta","Thysanoptera","Aeolothripidae","Aeolothripinae","Aeolothrips","Aeolothrips fasciatus","1","1","1","103","0","0","98.21","448","0","450","36.89","0"
"AGAW2","11W","INS","BOLD:AAN5161","Arthropoda","Insecta","Diptera","Ceratopogonidae","Dasyheleinae","Dasyhelea","","1","1","1","103","0","0","98.79","412","0","415","36.14","0"
"AGAW2","11W","INS","BOLD:AAP7026","Arthropoda","Insecta","Coleoptera","Latridiidae","Corticariinae","Melanophthalma","Melanophthalma villosa","1","1","1","103","0","0","97.82","412","0","450","31.56","0"
"AGAW2","11W","INS","BOLD:ACE4100","Arthropoda","Insecta","Orthoptera","Acrididae","Melanoplinae","Melanoplus","Melanoplus pinicola","1","1","1","103","0","0","98.85","347","0","350","34.86","0"
"AGAW2","11W","INS","BOLD:ACK5742","Arthropoda","Insecta","Diptera","Cecidomyiidae","","","","1","1","1","103","0","0","99.76","424","0","424","28.54","0"
"AGAW2","11W","INS","BOLD:AAG5507","Arthropoda","Insecta","Diptera","Chironomidae","Chironominae","Chironomus","Chironomus atrella","1","1","1","103","0","0","97.98","445","0","450","36.67","0"
"AGAW2","11W","INS","BOLD:AAF3338","Arthropoda","Insecta","Lepidoptera","Noctuidae","Noctuinae","Striacosta","Striacosta albicosta","1","1","1","103","0","0","98.9","363","0","367","32.7","0"
"AGAW2","11W","INS","BOLD:AAG9668","Arthropoda","Insecta","Diptera","Dolichopodidae","Diaphorinae","Chrysotus","Chrysotus sp., Chrysotus sp. nr. obliquus","1","1","1","103","0","0","98.87","444","0","450","32.44","0"
"AGAW2","11W","INS","BOLD:AAP8816","Arthropoda","Insecta","Diptera","Cecidomyiidae","Cecidomyiinae","Sitodiplosis","Sitodiplosis sp. BOLD:AAP8816","1","1","1","103","0","0","99.74","389","0","415","25.3","0"
"AGAW2","11W","INS","BOLD:AAI4303","Arthropoda","Insecta","Diptera","Chironomidae","Chironominae","Chironomus","Chironomus melanescens","1","1","1","103","0","0","97.98","446","0","450","33.33","0"
"AGAW2","11W","INS","BOLD:AAA8640","Arthropoda","Insecta","Lepidoptera","Tortricidae","Olethreutinae","Eucosma, Sonia, Pelochrista","Pelochrista suadana, Sonia new sp. 1, Sonia new sp. 2, Sonia new sp. 3, Sonia paraplesiana, Sonia canadana","1","1","1","103","0","0","97.76","446","0","450","32.44","0"
"AGAW2","11W","INS","BOLD:AAA2955","Arthropoda","Insecta","Lepidoptera","Tortricidae","Tortricinae","Argyrotaenia","Argyrotaenia occultana, Argyrotaenia ljungiana, Argyrotaenia pinatubana, Argyrotaenia floridana, Argyrotaenia velutinana, Argyrotaenia tabulana, Argyrotaenia gogana","1","1","1","103","0","0","97.78","450","0","450","32.67","0"
"AGAW2","11W","INS","BOLD:AAH1054","Arthropoda","Insecta","Hymenoptera","Braconidae","Microgastrinae","Choeras","","1","1","1","103","0","0","99.76","420","0","450","25.33","0"
"AGAW2","11W","INS","BOLD:AAC1042","Arthropoda","Insecta","Lepidoptera","Crambidae","Spilomelinae","Lygropia","Lygropia rivulalis","1","1","1","103","0","0","99.78","447","0","450","28.89","0"
"AGAW2","11W","INS","BOLD:AAG3273","Arthropoda","Insecta","Diptera","Phoridae","Metopininae","Megaselia","Megaselia albicaudata","1","1","1","103","0","0","99.56","451","0","450","32.89","0"
"AGAW2","11W","INS","BOLD:ADF2969","Arthropoda","Insecta","Diptera","Sarcophagidae","Sarcophaginae","Blaesoxipha","Blaesoxipha falciformis","1","1","1","103","0","0","98.74","396","0","398","36.93","0"
"AGAW2","11W","INS","BOLD:AAH3914","Arthropoda","Insecta","Diptera","Sciaridae","","Sciara","Sciara hamatilis, Sciara humeralis","1","1","1","103","0","0","97.44","352","0","354","29.38","0"
"AGAW2","11W","INS","BOLD:AAV1136","Arthropoda","Insecta","Diptera","Scatopsidae","","","","1","1","1","103","0","0","98.41","441","0","442","37.1","0"
"AGAW2","11W","INS","BOLD:AAA5299","Arthropoda","Insecta","Diptera","Chironomidae","Orthocladiinae","Cricotopus","Cricotopus laetus, Cricotopus relucens, Cricotopus trifasciatus, Cricotopus sp., Cricotopus sp. 3ES, Cricotopus sylvestris, Cricotopus glacialis","1","1","1","103","0","0","97.53","446","0","450","34","0"
"AGAW2","11W","INS","BOLD:ABZ5071","Arthropoda","Insecta","Diptera","Chironomidae","Chironominae","Micropsectra","Micropsectra nigripila","1","1","1","103","0","0","98.06","360","0","360","36.67","0"
"AGAW2","11W","INS","BOLD:AAZ5483","Arthropoda","Insecta","Diptera","Muscidae","Phaoniinae","Helina","Helina setiventris","1","1","1","103","0","0","99.55","445","0","450","28.89","0"
"AGAW2","11W","INS","BOLD:ACE0554","Arthropoda","Insecta","Diptera","Phoridae","Metopininae","Megaselia","","1","1","1","103","0","0","99.11","449","0","450","32.44","0"
"AGAW2","11W","INS","BOLD:AAB5609","Arthropoda","Insecta","Diptera","Muscidae","Coenosiinae","Coenosia","Coenosia sp.BOLD:AAB5609, Coenosia tigrina","1","1","1","103","0","0","97.85","418","0","420","31.67","0"
"AGAW2","11W","INS","BOLD:AAB0373","Arthropoda","Insecta","Neuroptera","Chrysopidae","Chrysopinae","Chrysoperla","Chrysoperla sp. isolate 82, Chrysoperla zastrowi, Chrysoperla plorabunda, Chrysoperla sp. isolate 71, Chrysoperla agilis, Italochrysa italica, Chrysoperla sp. isolate 88, Chrysoperla lucasina, Chrysoperla carnea, Chrysoperla mediterranea, Chrysoperla pallida, Chrysoperla nipponensis, Chrysoperla sillemi","1","1","1","103","0","0","99.78","450","0","450","31.56","0"
"AGAW2","11W","INS","BOLD:ADH4889","Arthropoda","Insecta","Diptera","","","","","1","1","1","103","0","0","99.11","450","0","450","35.78","0"
"AGAW2","11W","INS","BOLD:AAG2334","Arthropoda","Insecta","Diptera","Tachinidae","Phasiinae","Gymnoclytia","Gymnoclytia occidua, Gymnoclytia sp. 1, Gymnoclytia sp. 2","1","1","1","103","0","0","98.42","443","0","450","30.44","0"
"AGAW2","11W","INS","BOLD:AAD2041","Arthropoda","Insecta","Diptera","Dolichopodidae","Sciapodinae","Condylostylus","Condylostylus inermis, Condylostylus nigrofemoratus, Condylostylus connectans, Condylostylus caudatus, Condylostylus flavipes","1","1","1","103","0","0","98.99","397","0","400","35.25","0"
"AGAW2","11W","INS","BOLD:AAU7126","Arthropoda","Insecta","Coleoptera","Cleridae","Hydnocerinae","Isohydnocera","Isohydnocera curtipennis","1","1","1","103","0","0","99.55","449","0","450","33.33","0"
"AGAW2","11W","INS","BOLD:AAH2293","Arthropoda","Insecta","Diptera","Asilidae","Stenopogoninae","Dioctria, Eudioctria","Dioctria hyalipennis, Dioctria baumhaueri, Eudioctria propinqua","1","1","1","103","0","0","97.94","389","0","391","29.92","0"
"AGAW2","11W","INS","BOLD:ACE8878","Arthropoda","Insecta","Orthoptera","Acrididae","Melanoplinae","Melanoplus","Melanoplus scudderi group","1","1","1","103","0","0","97.56","450","0","450","35.56","0"
"AGAW2","11W","INS","BOLD:AAB1352","Arthropoda","Insecta","Lepidoptera","Erebidae","Hypeninae","Hypena","Hypena sp.","1","1","1","103","0","0","99.78","450","0","450","32.89","0"
"AGAW2","11W","INS","BOLD:ABW1460","Arthropoda","Insecta","Coleoptera","Chrysomelidae","Criocerinae","Oulema","Oulema erichsonii","1","1","1","103","0","0","97.78","360","0","361","33.8","0"
"AGAW2","11W","INS","BOLD:ACL7238","Arthropoda","Insecta","Diptera","Xylomyidae","","Solva","","1","1","1","103","0","0","99.56","450","0","450","35.56","0"
"AGAW2","11W","INS","BOLD:ACE9865","Arthropoda","Insecta","Lepidoptera","Noctuidae","Noctuinae","Feltia","Feltia subgothica, Feltia herilis, Feltia tricosa","1","1","1","103","0","0","100","450","0","450","30.89","0"
"AGAW2","11W","INS","BOLD:AAE3943","Arthropoda","Insecta","Hemiptera","Miridae","Phylinae","Plagiognathus","Plagiognathus chrysanthemi","1","1","1","103","0","0","98.98","392","0","450","36","0"
"AGAW2","11W","INS","BOLD:AAA7374","Arthropoda","Insecta","Diptera","Syrphidae","Syrphinae","Sphaerophoria","Sphaerophoria bankowskae, Sphaerophoria infuscata, Sphaerophoria nr. philanthus, Sphaerophoria batava, Sphaerophoria sp., Sphaerophoria rueppellii, Sphaerophoria interrupta-group, Sphaerophoria sp. MA-1, Sphaerophoria cleoae, Sphaerophoria asymmetrica, Sphaerophoria fatarum, Sphaerophoria bengalensis, Sphaerophoria bifurcata, Sphaerophoria taeniata, Sphaerophoria interrupta, Sphaerophoria contigua, Sphaerophoria kaa, Sphaerophoria laurae, Sphaerophoria chongjini, Sphaerophoria boreoalpina, Sphaerophoria sp. BOLDAAA7374, Sphaerophoria macrogaster, Sphaerophoria sulphuripes, Sphaerophoria brevipilosa, Sphaerophoria philanthus, Sphaerophoria pyrrhina, Sphaerophoria abbreviata, Sphaerophoria scripta, Sphaerophoria potentillae, Sphaerophoria longipilosa, Sphaerophoria virgata","1","1","1","103","0","0","99.52","417","0","419","30.31","0"
"AGAW2","11W","INS","BOLD:ADB5456","Arthropoda","Insecta","Hymenoptera","Eulophidae","Entedoninae","Neochrysocharis","","1","1","1","103","0","0","98.06","413","0","450","25.56","0"
"AGAW2","11W","INS","BOLD:AAC1374","Arthropoda","Insecta","Hemiptera","Aphididae","Aphidinae","Aphis","Aphis nasturtii, Aphis davletshinae, Aphis sp. rostella like, Aphis cf. rostella, Aphis althaeae, Aphis umbrella","1","1","1","103","0","0","99.78","449","0","450","26.22","0"
"AGAW2","11W","INS","BOLD:AAO7890","Arthropoda","Insecta","Diptera","Agromyzidae","","","","1","1","1","103","0","0","97.99","398","0","445","35.28","0"
"AGAW2","11W","INS","BOLD:AAH3225","Arthropoda","Insecta","Psocodea","Psocidae","Psocinae","Metylophorus","Metylophorus novaescotiae","1","1","1","103","0","0","99.33","450","0","450","34.89","0"
"AGAW2","11W","INS","BOLD:ABV1422","Arthropoda","Insecta","Diptera","Cecidomyiidae","","","","1","1","1","103","0","0","97.49","398","0","401","26.93","0"
"AGAW2","11W","INS","BOLD:AAM9255","Arthropoda","Insecta","Diptera","Sciaridae","","Pseudolycoriella","","1","1","1","103","0","0","98.2","445","0","450","30.67","0"
"AGAW2","11W","INS","BOLD:ACZ7632","Arthropoda","Insecta","Thysanoptera","Thripidae","Thripinae","Aptinothrips","Aptinothrips rufus","1","1","1","103","0","0","98.4","375","0","450","31.78","0"
"AGAW2","11W","NAR","BOLD:AAL1512","Mollusca","Gastropoda","Stylommatophora","Agriolimacidae",NA,"Deroceras","Deroceras laeve","1243","1232","1232","1","0.88","0","97.76","388.78","0","392.05","33.29","0"
"AGAW2","11W","NAR","BOLD:AAI9663","Mollusca","Gastropoda","Stylommatophora","Agriolimacidae",NA,"Deroceras","Deroceras reticulatum","207","182","182","2","12.08","0","99.59","400.57","0","436.26","35.67","0"
"AGAW2","11W","NAR","TAX:660402","Mollusca","Gastropoda","Stylommatophora","Agriolimacidae",NA,"Deroceras","Deroceras sp.","122","122","122","3","0","0","97.8","394.7","0","397.85","32.76","0"
"AGAW2","11W","NAR","TAX:106138","","","","",NA,"","","1","1","1","4","0","0","97.97","395","0","401","32.92","0"
"AGAW2","11W","NIN","BOLD:AAH6630","Arthropoda","Collembola","Entomobryomorpha","Entomobryidae","Entomobryinae","Entomobrya","Entomobrya unostrigata","1046","898","898","1","14.15","0","99.42","436.63","0","437.84","38.98","0"
"AGAW2","11W","NIN","BOLD:AAI4346","Arthropoda","Arachnida","Opiliones","Phalangiidae","","Paroligolophus, Phalangium","Paroligolophus agrestis, Phalangium opilio","687","633","633","2","7.86","0","99.28","433.08","0","435.74","36.21","0"
"AGAW2","11W","NIN","BOLD:ABW2639","Arthropoda","Arachnida","Trombidiformes","Anystidae","","","","622","509","509","3","18.17","0","99.51","447.77","0","446.58","31.45","0"
"AGAW2","11W","NIN","BOLD:AAA6381","Arthropoda","Arachnida","Araneae","Tetragnathidae","","Tetragnatha","Tetragnatha laboriosa","246","235","235","4","4.47","0","99.23","424.81","0","426.88","39.37","0"
"AGAW2","11W","NIN","BOLD:ACZ1295","Arthropoda","Arachnida","Trombidiformes","Anystidae","","","","87","87","87","5","0","0","99.39","372.72","0","380.2","31.31","0"
"AGAW2","11W","NIN","BOLD:AAB7914","Arthropoda","Collembola","Symphypleona","Bourletiellidae","","Bourletiella","","40","37","37","6","7.5","0","99.58","446.4","0","447.63","38.17","0"
"AGAW2","11W","NIN","BOLD:AAU6252","Arthropoda","Arachnida","Trombidiformes","Bdellidae","","Cyta","","32","29","29","7","9.38","0","99.07","424.97","0","426.66","31.29","0"
"AGAW2","11W","NIN","BOLD:AAB7601","Arthropoda","Arachnida","Araneae","Eutichuridae","","Cheiracanthium","Cheiracanthium mildei","29","26","26","8","10.34","0","99.54","429.03","0","430.24","28.93","0"
"AGAW2","11W","NIN","BOLD:AAM7961","Arthropoda","Arachnida","Trombidiformes","Anystidae","","","","21","20","20","9","4.76","0","99.53","449.86","0","446.48","31.29","0"
"AGAW2","11W","NIN","BOLD:AAA5267","Arthropoda","Collembola","Entomobryomorpha","Isotomidae","","","","14","14","14","10","0","0","99.2","420.36","0","422.71","37.96","0"
"AGAW2","11W","NIN","BOLD:AAF9093","Arthropoda","Arachnida","Mesostigmata","Parasitidae","","","","8","8","8","11","0","0","99.27","426.63","0","427.63","40.39","0"
"AGAW2","11W","NIN","BOLD:AAA9476","Arthropoda","Arachnida","Araneae","Linyphiidae","","Grammonota","Grammonota gentilis","6","6","6","12","0","0","98.8","437.33","0","438.5","30.78","0"
"AGAW2","11W","NIN","BOLD:ACX8969","Arthropoda","Arachnida","Trombidiformes","Eupodidae","","","","4","4","4","13","0","0","99.01","429.25","0","433.25","27.39","0"
"AGAW2","11W","NIN","BOLD:ACY2550","Arthropoda","Arachnida","Trombidiformes","Bdellidae","","Cyta","","3","3","3","14","0","0","99.04","382.67","0","389.67","31.31","0"
"AGAW2","11W","NIN","BOLD:ACV5609","","","","","","","","2","2","2","15","0","0","99.22","448.5","0","450","38.89","0"
"AGAW2","11W","NIN","BOLD:ACC0359","Arthropoda","Collembola","Symphypleona","Bourletiellidae","","Bourletiella","","1","1","1","16","0","0","99.09","441","0","450","37.78","0"
"AGAW2","11W","NIN","BOLD:AAF9285","Arthropoda","Arachnida","Mesostigmata","Parasitidae","","","","1","1","1","16","0","0","97.46","394","0","399","34.59","0"
"AGAW2","11W","NIN","BOLD:AAB0726","Arthropoda","Arachnida","Araneae","Lycosidae","","Trochosa","Trochosa ruricola","1","1","1","16","0","0","98.66","448","0","450","31.11","0"
"AGAW3","02E","BAC","TAX:91391","Proteobacteria","Alphaproteobacteria","Rickettsiales",NA,NA,NA,NA,"5","5","5","1","0","0","98.19","418.2","0","440.6","39.98","0"
"AGAW3","02E","CHO","no match","no match","no match","no match","no match","no match","no match","no match","no match","no match","no match","no match","no match","no match","no match","no match","no match","no match","no match","no match"
"AGAW3","02E","INS","BOLD:AAG2511","Arthropoda","Insecta","Diptera","Anthomyiidae","Anthomyiinae","Delia","Delia platura","94163","61013","61013","1","35.2","0","99.43","443.27","0","445.57","33.18","0"
"AGAW3","02E","INS","BOLD:AAA3453","Arthropoda","Insecta","Diptera","Anthomyiidae","Anthomyiinae","Delia","","10470","7758","7758","2","25.9","0","99.22","444.47","0","446.31","34.66","0"
"AGAW3","02E","INS","BOLD:AAD0853","Arthropoda","Insecta","Diptera","Scathophagidae","Scathophaginae","Scathophaga","Scathophaga lutaria, Scathophaga hiemalis, Scathophaga furcata, Scathophaga stercoraria, Scathophaga inquinata, Scathophaga suilla","5830","4021","4021","3","31.03","0","98.83","445.09","0","446.4","32.19","0"
"AGAW3","02E","INS","BOLD:AAB2384","Arthropoda","Insecta","Diptera","Syrphidae","Syrphinae","Eupeodes","Eupeodes duseki, Eupeodes volucris, Eupeodes nielseni, Eupeodes americanus, Eupeodes flukei, Eupeodes vockerothi, Eupeodes lucasi, Eupeodes montivagus, Eupeodes rufipunctatus, Eupeodes rojasi, Eupeodes goeldlini, Eupeodes luniger, Eupeodes punctifer, Eupeodes bucculatus, Eupeodes perplexus, Eupeodes abiskoensis, Eupeodes corollae, Eupeodes pomus, Eupeodes lundbecki, Eupeodes nitens, Eupeodes sp. BOLDAAB2384, Eupeodes curtus, Eupeodes latifasciatus","4858","4134","4134","4","14.9","0","99.02","428.51","0","430.54","30.78","0"
"AGAW3","02E","INS","BOLD:AAL7534","Arthropoda","Insecta","Diptera","Anisopodidae","Anisopodinae","Sylvicola","Sylvicola stackelbergi, Sylvicola stackelbergii 1, Sylvicola punctatus","2204","1624","1624","5","26.32","0","99.26","438.13","0","439.98","33.72","0"
"AGAW3","02E","INS","BOLD:AAD2085","Arthropoda","Insecta","Lepidoptera","Bucculatricidae","","Bucculatrix","Bucculatrix angustata","1247","910","910","6","27.02","0","99.18","442.93","0","448.92","32.91","0"
"AGAW3","02E","INS","BOLD:AAB2644","Arthropoda","Insecta","Diptera","Chironomidae","Orthocladiinae","Orthocladius","Orthocladius lapponicus, Orthocladius dorenus, Orthocladius mallochi","814","711","711","7","12.65","0","98.89","441.06","0","442.98","32.67","0"
"AGAW3","02E","INS","BOLD:AAN5358","Arthropoda","Insecta","Diptera","Chironomidae","Orthocladiinae","Smittia","Smittia stercoraria, Smittia sp. 22ES","574","509","509","8","11.32","0","99.22","442.61","0","444.67","37.25","0"
"AGAW3","02E","INS","BOLD:ABZ3875","Arthropoda","Insecta","Diptera","Chloropidae","Oscinellinae","Oscinella","Oscinella sp. BOLD:ABZ3875, Oscinella sp. 4, Oscinella sp. 2 AGT, Oscinella nigerrima","403","344","344","9","14.64","0","99.26","437.13","0","440.37","32.75","0"
"AGAW3","02E","INS","BOLD:AAF9014","Arthropoda","Insecta","Diptera","Limoniidae","Chioneinae","Symplecta","Symplecta cana, Symplecta sp. BOLDAAF9014","360","319","319","10","11.39","0","98.73","431.1","0","436.94","33.58","0"
"AGAW3","02E","INS","BOLD:AAC4197","Arthropoda","Insecta","Diptera","Chironomidae","Orthocladiinae","Paraphaenocladius","Paraphaenocladius impensus","282","256","256","11","9.22","0","99.38","438.45","0","440.48","35.07","0"
"AGAW3","02E","INS","BOLD:AAP4697","Arthropoda","Insecta","Diptera","Phoridae","Phorinae","Triphleba","","214","181","181","12","15.42","0","99.29","444.94","0","445.91","32.67","0"
"AGAW3","02E","INS","BOLD:ACB1079","Arthropoda","Insecta","Diptera","Cecidomyiidae","Micromyinae","Campylomyza","Campylomyza dilatata, CecidInt44 sp. BOLD:ACB1079","141","105","105","13","25.53","0","98.3","443.85","0","446.72","28.3","0"
"AGAW3","02E","INS","BOLD:AAP6827","Arthropoda","Insecta","Diptera","Cecidomyiidae","","","","112","105","105","14","6.25","0","98.99","444.13","0","446.02","31.83","0"
"AGAW3","02E","INS","BOLD:AAN5267","Arthropoda","Insecta","Diptera","Cecidomyiidae","","","","74","60","60","15","18.92","0","99.54","449.03","0","450","30.67","0"
"AGAW3","02E","INS","BOLD:AAH0134","Arthropoda","Insecta","Coleoptera","Phalacridae","Phalacrinae","Stilbus","Stilbus apicialis","73","73","73","16","0","0","99.27","426.64","0","427.78","31.94","0"
"AGAW3","02E","INS","BOLD:ACR4394","Arthropoda","Insecta","Diptera","Anthomyiidae","Anthomyiinae","Delia","Delia florilega","70","70","70","17","0","0","98.86","432.63","0","434.43","32.97","0"
"AGAW3","02E","INS","BOLD:ACR4350","Arthropoda","Insecta","Diptera","Sciaridae","","Bradysia","Bradysia placida","64","61","61","18","4.69","0","99.53","419.42","0","441.36","35.29","0"
"AGAW3","02E","INS","BOLD:AAP6876","Arthropoda","Insecta","Diptera","Chironomidae","Chironominae","Tanytarsus","","59","48","48","19","18.64","0","99.3","448.14","0","448.75","35.39","0"
"AGAW3","02E","INS","BOLD:AAB1098","Arthropoda","Insecta","Diptera","Culicidae","Culicinae","Ochlerotatus, Aedes","Aedes excrucians, Aedes cantans, Aedes stimulans, Aedes euedes, Aedes riparius, Aedes annulipes, Ochlerotatus cantans, Aedes fitchii, Aedes sp., Ochlerotatus annulipes, Ochlerotatus excrucians","56","56","56","20","0","0","98.6","429.21","0","430.84","35.24","0"
"AGAW3","02E","INS","BOLD:AAB9140","Arthropoda","Insecta","Diptera","Calliphoridae","Chrysomyinae","Phormia","Phormia regina","47","46","46","21","2.13","0","99.09","446.47","0","447.45","33.09","0"
"AGAW3","02E","INS","BOLD:AAN6433","Arthropoda","Insecta","Diptera","Sciaridae","","Corynoptera","Corynoptera deserta","29","28","28","22","3.45","0","99.46","435.48","0","437.55","30.33","0"
"AGAW3","02E","INS","BOLD:AAZ5599","Arthropoda","Insecta","Diptera","Chironomidae","Orthocladiinae","","","28","28","28","23","0","0","99.32","414.18","0","415.57","36.6","0"
"AGAW3","02E","INS","BOLD:ABA6465","Arthropoda","Insecta","Diptera","Chironomidae","Orthocladiinae","","","20","19","19","24","5","0","97.95","446.15","0","447.4","33.72","0"
"AGAW3","02E","INS","BOLD:ABA1215","Arthropoda","Insecta","Diptera","Sciaridae","","Lycoriella","Lycoriella sativae","15","13","13","25","13.33","0","99.39","437","0","437.93","35.6","0"
"AGAW3","02E","INS","BOLD:AAN6447","Arthropoda","Insecta","Diptera","Sciaridae","","Corynoptera","Corynoptera perpusilla","15","13","13","25","13.33","0","99.52","449.13","0","450","36.37","0"
"AGAW3","02E","INS","BOLD:AAB0868","Arthropoda","Insecta","Diptera","Calliphoridae","Calliphorinae","Cynomya","Cynomya cadaverina, Cynomya mortuorum","15","14","14","25","6.67","0","99.1","445.27","0","446.2","34.8","0"
"AGAW3","02E","INS","BOLD:AAM9254","Arthropoda","Insecta","Diptera","Sciaridae","","Bradysia","Bradysia pallipes","14","14","14","28","0","0","98.43","440.5","0","441.07","33.39","0"
"AGAW3","02E","INS","BOLD:AAQ0602","Arthropoda","Insecta","Diptera","Chironomidae","Chironominae","Micropsectra","","9","9","9","29","0","0","98.56","448.78","0","450","36.25","0"
"AGAW3","02E","INS","BOLD:AAH3920","Arthropoda","Insecta","Diptera","Sciaridae","","Dichopygina, Scatopsciara","","8","8","8","30","0","0","98.51","417.5","0","442.13","35.3","0"
"AGAW3","02E","INS","BOLD:AAG2462","Arthropoda","Insecta","Diptera","Anthomyiidae","Anthomyiinae","Botanophila","Botanophila fugax","8","8","8","30","0","0","98.86","445.38","0","447.13","31.87","0"
"AGAW3","02E","INS","BOLD:AAM9242","Arthropoda","Insecta","Diptera","Sciaridae","","Corynoptera","Corynoptera melanochaeta","7","7","7","32","0","0","98.65","444.57","0","445.71","34.41","0"
"AGAW3","02E","INS","BOLD:ABA1260","Arthropoda","Insecta","Diptera","Phoridae","Metopininae","Metopina, Megaselia","","7","7","7","32","0","0","99.76","409.71","0","449.29","35.77","0"
"AGAW3","02E","INS","BOLD:ACC7611","Arthropoda","Insecta","Diptera","Cecidomyiidae","","","","6","6","6","34","0","0","97.65","447.67","0","450","28.37","0"
"AGAW3","02E","INS","BOLD:AAD4355","Arthropoda","Insecta","Diptera","Culicidae","Culicinae","Aedes","Aedes provocans","6","6","6","34","0","0","99.55","439.33","0","442.33","31.68","0"
"AGAW3","02E","INS","BOLD:AAN5355","Arthropoda","Insecta","Diptera","Chironomidae","Orthocladiinae","Smittia","Smittia cf. stercoraria, Smittia stercoraria","6","6","6","34","0","0","98.6","429.33","0","431.67","36.25","0"
"AGAW3","02E","INS","BOLD:ABX2489","Arthropoda","Insecta","Coleoptera","Phalacridae","Phalacrinae","Stilbus","","5","5","5","37","0","0","97.73","429.6","0","431.2","32.9","0"
"AGAW3","02E","INS","BOLD:AAN6151","Arthropoda","Insecta","Coleoptera","Chrysomelidae","Galerucinae","Longitarsus","Longitarsus pratensis, Longitarsus reichei, Longitarsus scutellaris","5","5","5","37","0","0","99.59","440.4","0","442.6","33.64","0"
"AGAW3","02E","INS","BOLD:AAN8770","Arthropoda","Insecta","Diptera","Psychodidae","Psychodinae","Psychoda","Psychoda trinodulosa","4","4","4","39","0","0","98.82","441.5","0","441.75","33.89","0"
"AGAW3","02E","INS","BOLD:AAV0234","Arthropoda","Insecta","Hemiptera","Triozidae","Triozinae","Trioza","","3","3","3","40","0","0","98.21","446.33","0","449","37.49","0"
"AGAW3","02E","INS","BOLD:AAA4222","Arthropoda","Insecta","Lepidoptera","Erebidae","Hypeninae","Hypena","Hypena scabra","3","3","3","40","0","0","99.03","367.67","0","370.67","32.19","0"
"AGAW3","02E","INS","BOLD:AAA9251","Arthropoda","Insecta","Lepidoptera","Gelechiidae","Gelechiinae","Scrobipalpa","Scrobipalpa atriplicella, Scrobipalpa nr. atriplicella sp. 1","3","3","3","40","0","0","98.93","437.67","0","440.33","33.75","0"
"AGAW3","02E","INS","BOLD:AAN5356","Arthropoda","Insecta","Diptera","Chironomidae","Orthocladiinae","Smittia","","2","2","2","43","0","0","97.99","447.5","0","450","34.56","0"
"AGAW3","02E","INS","BOLD:AAG1742","Arthropoda","Insecta","Diptera","Muscidae","Phaoniinae","Helina","Helina rufitibia","2","2","2","43","0","0","98.78","449","0","450","30.33","0"
"AGAW3","02E","INS","BOLD:AAB1024","Arthropoda","Insecta","Lepidoptera","Noctuidae","Noctuinae","Pseudohermonassa","Pseudohermonassa bicarnea","2","2","2","43","0","0","99.78","449","0","450","33.78","0"
"AGAW3","02E","INS","BOLD:AAG6743","Arthropoda","Insecta","Diptera","Sarcophagidae","Sarcophaginae","Sarcophaga","Sarcophaga sp. 1 MG, Sarcophaga subvicina","2","2","2","43","0","0","98.65","441.5","0","442.5","35.37","0"
"AGAW3","02E","INS","BOLD:AAQ0582","Arthropoda","Insecta","Diptera","Anthomyiidae","Anthomyiinae","Lasiomma","","2","2","2","43","0","0","99.07","438","0","438","31.61","0"
"AGAW3","02E","INS","BOLD:ACL0119","Arthropoda","Insecta","Diptera","Chironomidae","Orthocladiinae","Paraphaenocladius","","1","1","1","48","0","0","99.55","449","0","450","33.33","0"
"AGAW3","02E","INS","BOLD:ACV3368","","","","","","","","1","1","1","48","0","0","98.89","450","0","450","33.56","0"
"AGAW3","02E","INS","BOLD:AAZ9744","Arthropoda","Insecta","Hymenoptera","Braconidae","Brachistinae","Blacus","Blacus stelfoxi","1","1","1","48","0","0","98.22","449","0","450","32.22","0"
"AGAW3","02E","INS","BOLD:AAY6412","Arthropoda","Insecta","Diptera","Sciaridae","","Bradysia","Bradysia urticae","1","1","1","48","0","0","100","449","0","450","32.22","0"
"AGAW3","02E","INS","BOLD:AAA7374","Arthropoda","Insecta","Diptera","Syrphidae","Syrphinae","Sphaerophoria","Sphaerophoria bankowskae, Sphaerophoria infuscata, Sphaerophoria nr. philanthus, Sphaerophoria batava, Sphaerophoria sp., Sphaerophoria rueppellii, Sphaerophoria interrupta-group, Sphaerophoria sp. MA-1, Sphaerophoria cleoae, Sphaerophoria asymmetrica, Sphaerophoria fatarum, Sphaerophoria bengalensis, Sphaerophoria bifurcata, Sphaerophoria taeniata, Sphaerophoria interrupta, Sphaerophoria contigua, Sphaerophoria kaa, Sphaerophoria laurae, Sphaerophoria chongjini, Sphaerophoria boreoalpina, Sphaerophoria sp. BOLDAAA7374, Sphaerophoria macrogaster, Sphaerophoria sulphuripes, Sphaerophoria brevipilosa, Sphaerophoria philanthus, Sphaerophoria pyrrhina, Sphaerophoria abbreviata, Sphaerophoria scripta, Sphaerophoria potentillae, Sphaerophoria longipilosa, Sphaerophoria virgata","1","1","1","48","0","0","99.11","449","0","450","31.78","0"
"AGAW3","02E","INS","BOLD:AAK5928","Arthropoda","Insecta","Coleoptera","Chrysomelidae","Criocerinae","Oulema","Oulema melanopus","1","1","1","48","0","0","98.44","449","0","450","36","0"
"AGAW3","02E","INS","BOLD:AAC9062","Arthropoda","Insecta","Diptera","Culicidae","Culicinae","Ochlerotatus, Aedes","Aedes fitchii, Ochlerotatus fitchii, Aedes grossbecki, Aedes ventrovittis","1","1","1","48","0","0","98.44","449","0","450","32.89","0"
"AGAW3","02E","INS","BOLD:AAH3914","Arthropoda","Insecta","Diptera","Sciaridae","","Sciara","Sciara hamatilis, Sciara humeralis","1","1","1","48","0","0","100","447","0","450","30","0"
"AGAW3","02E","INS","BOLD:AAG9055","Arthropoda","Insecta","Hymenoptera","Vespidae","Vespinae","Vespula","Vespula germanica","1","1","1","48","0","0","97.11","450","0","450","30.67","0"
"AGAW3","02E","INS","BOLD:ABU9891","Arthropoda","Insecta","Diptera","Muscidae","Phaoniinae","Phaonia","","1","1","1","48","0","0","99.55","447","0","450","33.33","0"
"AGAW3","02E","INS","BOLD:AAH3661","Arthropoda","Insecta","Diptera","Cecidomyiidae","","","","1","1","1","48","0","0","99.56","450","0","450","25.78","0"
"AGAW3","02E","INS","BOLD:AAN5348","Arthropoda","Insecta","Diptera","Chironomidae","Orthocladiinae","Parametriocnemus","","1","1","1","48","0","0","98.86","352","0","355","34.08","0"
"AGAW3","02E","INS","BOLD:ACP4736","Arthropoda","Insecta","Diptera","Chironomidae","Orthocladiinae","Smittia","Smittia sp. 8ES","1","1","1","48","0","0","99.11","448","0","450","33.11","0"
"AGAW3","02E","INS","BOLD:AAB7105","Arthropoda","Insecta","Hymenoptera","Vespidae","Polistinae","Polistes","Polistes dominula","1","1","1","48","0","0","99.78","449","0","450","27.11","0"
"AGAW3","02E","INS","BOLD:AAB4093","Arthropoda","Insecta","Lepidoptera","Noctuidae","Noctuinae","Elaphria","Elaphria grata","1","1","1","48","0","0","98.07","415","0","416","29.33","0"
"AGAW3","02E","INS","BOLD:AAB2076","Arthropoda","Insecta","Lepidoptera","Tortricidae","Olethreutinae","Ancylis","Ancylis metamelana, Ancylis sp.","1","1","1","48","0","0","100","447","0","450","31.33","0"
"AGAW3","02E","INS","BOLD:AAA4121","Arthropoda","Insecta","Diptera","Simuliidae","Simuliinae","Simulium","Simulium vittatum complex, Simulium vittatum, Simulium sp. BOLDAAA4121, Simulium argus, Simulium vittatum s.str.","1","1","1","48","0","0","98.96","385","0","387","32.56","0"
"AGAW3","02E","INS","BOLD:AAA2482","Arthropoda","Insecta","Lepidoptera","Noctuidae","Noctuinae","Mythimna","Mythimna unipuncta","1","1","1","48","0","0","99.11","448","0","450","30.44","0"
"AGAW3","02E","INS","BOLD:AAF6798","Arthropoda","Insecta","Diptera","Chironomidae","Tanypodinae","Procladius","Procladius signatus","1","1","1","48","0","0","97.33","449","0","450","33.33","0"
"AGAW3","02E","INS","BOLD:AAG2123","Arthropoda","Insecta","Diptera","Tachinidae","Dexiinae","Dinera","Dinera grisescens","1","1","1","48","0","0","98.78","409","0","411","30.41","0"
"AGAW3","02E","INS","BOLD:AAG7306","Arthropoda","Insecta","Diptera","Sphaeroceridae","Limosininae","Minilimosina","Minilimosina vitripennis","1","1","1","48","0","0","99.32","444","0","450","33.78","0"
"AGAW3","02E","INS","BOLD:ACJ0205","Arthropoda","Insecta","Diptera","Chironomidae","Orthocladiinae","Thienemanniella","","1","1","1","48","0","0","99.23","389","0","431","33.64","0"
"AGAW3","02E","INS","BOLD:AAA3080","Arthropoda","Insecta","Trichoptera","Limnephilidae","Limnephilinae","Limnephilus","Limnephilus submonilifer","1","1","1","48","0","0","99.76","411","0","415","33.49","0"
"AGAW3","02E","INS","BOLD:AAG6460","Arthropoda","Insecta","Diptera","Cecidomyiidae","","","","1","1","1","48","0","0","97.55","449","0","450","34","0"
"AGAW3","02E","NAR","BOLD:AAL1512","Mollusca","Gastropoda","Stylommatophora","Agriolimacidae",NA,"Deroceras","Deroceras laeve","2","2","2","1","0","0","97.9","357.5","0","359","33.99","0"
"AGAW3","02E","NIN","BOLD:ABW0506","Arthropoda","Arachnida","Opiliones","Phalangiidae","Phalangiinae","Rilaena","Rilaena triangularis","264","182","182","1","31.06","0","99.67","446.15","0","446.83","39.76","0"
"AGAW3","02E","NIN","BOLD:ADH9176","","","","","","","","8","8","8","2","0","0","98.31","428.38","0","450","38.31","0"
"AGAW3","02E","NIN","BOLD:ADI0704","","","","","","","","8","8","8","2","0","0","98.63","389.63","0","426.38","37.71","0"
"AGAW3","02E","NIN","BOLD:ADI0775","Arthropoda","Arachnida","Opiliones","","","","","5","5","5","4","0","0","98.04","448.4","0","450","39.91","0"
"AGAW3","02E","NIN","BOLD:AAV0589","Arthropoda","Arachnida","Mesostigmata","Blattisociidae","Platyseiinae","Cheiroseius","","4","4","4","5","0","0","99.28","448.75","0","450","27.94","0"
"AGAW3","02E","NIN","BOLD:ACL9406","Arthropoda","Arachnida","Trombidiformes","Cunaxidae","","","","2","2","2","6","0","0","98.62","399.5","0","428","24.51","0"
"AGAW3","02E","NIN","BOLD:AAC6891","Arthropoda","Arachnida","Araneae","Salticidae","","Phidippus","Phidippus audax","1","1","1","7","0","0","99.01","404","0","450","31.33","0"
"AGAW3","02N","BAC","TAX:91391","Proteobacteria","Alphaproteobacteria","Rickettsiales","",NA,"","","207","143","143","1","30.92","0","99.52","386.95","0","445.65","39.95","0"
"AGAW3","02N","BAC","TAX:175768","Proteobacteria","Alphaproteobacteria","Rickettsiales","Rickettsiaceae",NA,"Wolbachia","Wolbachia pipientis","40","28","28","2","30","0","98.72","447.1","0","448.1","39.6","0"
"AGAW3","02N","BAC","TAX:91393","Proteobacteria","Alphaproteobacteria","Rickettsiales","Rickettsiaceae",NA,"Wolbachia","","11","11","11","3","0","0","99.63","391.73","0","426.09","39.24","0"
"AGAW3","02N","BAC","TAX:392851","Proteobacteria","Alphaproteobacteria","Rickettsiales","Rickettsiaceae",NA,"Wolbachia","Wolbachia endosymbiont of Formicidae","3","3","3","4","0","0","98","431.67","0","450","42","0"
"AGAW3","02N","BAC","TAX:91389","Proteobacteria","","","",NA,"","","1","1","1","5","0","0","99.45","362","0","417","37.65","0"
"AGAW3","02N","CHO","no match","no match","no match","no match","no match","no match","no match","no match","no match","no match","no match","no match","no match","no match","no match","no match","no match","no match","no match","no match"
"AGAW3","02N","INS","BOLD:AAG2511","Arthropoda","Insecta","Diptera","Anthomyiidae","Anthomyiinae","Delia","Delia platura","54428","36314","36314","1","33.28","0","99.46","443.09","0","445.48","33.14","0"
"AGAW3","02N","INS","BOLD:AAN5355","Arthropoda","Insecta","Diptera","Chironomidae","Orthocladiinae","Smittia","Smittia cf. stercoraria, Smittia stercoraria","12325","10332","10332","2","16.17","0","99.23","420.36","0","445.2","36.29","0"
"AGAW3","02N","INS","BOLD:AAP9022","Arthropoda","Insecta","Diptera","Cecidomyiidae","","","","8342","5328","5328","3","36.13","0","99.7","446.56","0","447.62","27.66","0"
"AGAW3","02N","INS","BOLD:AAD0853","Arthropoda","Insecta","Diptera","Scathophagidae","Scathophaginae","Scathophaga","Scathophaga lutaria, Scathophaga hiemalis, Scathophaga furcata, Scathophaga stercoraria, Scathophaga inquinata, Scathophaga suilla","5166","3698","3698","4","28.42","0","99.4","443.94","0","445.69","31.94","0"
"AGAW3","02N","INS","BOLD:AAN5358","Arthropoda","Insecta","Diptera","Chironomidae","Orthocladiinae","Smittia","Smittia stercoraria, Smittia sp. 22ES","4551","3770","3770","5","17.16","0","99.22","442.77","0","444.72","37.33","0"
"AGAW3","02N","INS","BOLD:AAB1098","Arthropoda","Insecta","Diptera","Culicidae","Culicinae","Ochlerotatus, Aedes","Aedes excrucians, Aedes cantans, Aedes stimulans, Aedes euedes, Aedes riparius, Aedes annulipes, Ochlerotatus cantans, Aedes fitchii, Aedes sp., Ochlerotatus annulipes, Ochlerotatus excrucians","2973","2651","2651","6","10.83","0","98.6","433.5","0","434.93","35.37","0"
"AGAW3","02N","INS","BOLD:AAB2076","Arthropoda","Insecta","Lepidoptera","Tortricidae","Olethreutinae","Ancylis","Ancylis metamelana, Ancylis sp.","2479","1827","1827","7","26.3","0","99.53","440.93","0","442.65","31.03","0"
"AGAW3","02N","INS","BOLD:AAN5267","Arthropoda","Insecta","Diptera","Cecidomyiidae","","","","2355","1561","1561","8","33.72","0","99.45","447.73","0","448.83","30.63","0"
"AGAW3","02N","INS","BOLD:AAP2970","Arthropoda","Insecta","Diptera","Anthomyiidae","Anthomyiinae","Anthomyia","Anthomyia procellaris, Anthomyia pluvialis","2132","1652","1652","9","22.51","0","99.3","442.39","0","444.28","33.2","0"
"AGAW3","02N","INS","BOLD:AAB2644","Arthropoda","Insecta","Diptera","Chironomidae","Orthocladiinae","Orthocladius","Orthocladius lapponicus, Orthocladius dorenus, Orthocladius mallochi","1912","1475","1475","10","22.86","0","99.42","446.61","0","448.35","32.3","0"
"AGAW3","02N","INS","BOLD:AAN5348","Arthropoda","Insecta","Diptera","Chironomidae","Orthocladiinae","Parametriocnemus","","1896","1668","1668","11","12.03","0","98.76","436.22","0","437.98","34","0"
"AGAW3","02N","INS","BOLD:ACN8090","Arthropoda","Insecta","Diptera","Chironomidae","Orthocladiinae","","","1123","830","830","12","26.09","0","98.42","440.1","0","441.72","33.54","0"
"AGAW3","02N","INS","BOLD:AAN1738","Arthropoda","Insecta","Coleoptera","Staphylinidae","Oxytelinae","Anotylus","Anotylus tetracarinatus","1078","710","710","13","34.14","0","99.6","445.34","0","446.97","33.76","0"
"AGAW3","02N","INS","BOLD:AAE4298","Arthropoda","Insecta","Diptera","Chironomidae","Orthocladiinae","Cricotopus","Cricotopus tremulus","885","821","821","14","7.23","0","98.4","421.02","0","430.03","32.69","0"
"AGAW3","02N","INS","BOLD:AAA8109","Arthropoda","Insecta","Lepidoptera","Gelechiidae","Dichomeridinae","Dichomeris","Dichomeris ligulella","803","532","532","15","33.75","0","98.77","446.32","0","447.26","31.93","0"
"AGAW3","02N","INS","BOLD:ACF2821","Arthropoda","Insecta","Diptera","Mycetophilidae","Mycetophilinae","Mycetophila","Mycetophila fungorum","774","636","636","16","17.83","0","99.27","439.1","0","441.48","35.29","0"
"AGAW3","02N","INS","BOLD:AAN9511","Arthropoda","Insecta","Coleoptera","Staphylinidae","Tachyporinae","Tachyporus","Tachyporus dispar, Tachyporus rulomoides, Tachyporus chrysomelinus","482","429","429","17","11","0","99.15","433.8","0","436.14","32.47","0"
"AGAW3","02N","INS","BOLD:AAA9251","Arthropoda","Insecta","Lepidoptera","Gelechiidae","Gelechiinae","Scrobipalpa","Scrobipalpa atriplicella, Scrobipalpa nr. atriplicella sp. 1","459","394","394","18","14.16","0","99.46","440.71","0","442.6","34.45","0"
"AGAW3","02N","INS","BOLD:AAH2753","Arthropoda","Insecta","Coleoptera","Anthicidae","Anthicinae","Anthicus","Anthicus cervinus","455","396","396","19","12.97","0","98.17","418.17","0","434.32","33.57","0"
"AGAW3","02N","INS","BOLD:ACP4736","Arthropoda","Insecta","Diptera","Chironomidae","Orthocladiinae","Smittia","Smittia sp. 8ES","447","389","389","20","12.98","0","98.94","434.35","0","436.3","32.95","0"
"AGAW3","02N","INS","BOLD:AAI1527","Arthropoda","Insecta","Diptera","Chironomidae","Chironominae","Micropsectra","","242","232","232","21","4.13","0","99.02","423.03","0","425.29","33.89","0"
"AGAW3","02N","INS","BOLD:AAC1644","Arthropoda","Insecta","Lepidoptera","Gelechiidae","Gelechiinae","Scrobipalpa","Scrobipalpa acuminatella","239","212","212","22","11.3","0","98.61","443.31","0","444.55","34.08","0"
"AGAW3","02N","INS","BOLD:AAY6412","Arthropoda","Insecta","Diptera","Sciaridae","","Bradysia","Bradysia urticae","201","167","167","23","16.92","0","99.38","439.51","0","441.85","32.15","0"
"AGAW3","02N","INS","BOLD:ABV0255","Arthropoda","Insecta","Diptera","Chironomidae","Orthocladiinae","Limnophyes","","186","157","157","24","15.59","0","99.31","440.95","0","442.25","34.77","0"
"AGAW3","02N","INS","BOLD:AAM9254","Arthropoda","Insecta","Diptera","Sciaridae","","Bradysia","Bradysia pallipes","154","153","153","25","0.65","0","98.82","428.47","0","429","33.34","0"
"AGAW3","02N","INS","BOLD:AAQ0638","Arthropoda","Insecta","Diptera","Cecidomyiidae","","","","151","112","112","26","25.83","0","97.52","437.14","0","438.57","29.19","0"
"AGAW3","02N","INS","BOLD:AAG6460","Arthropoda","Insecta","Diptera","Cecidomyiidae","","","","149","132","132","27","11.41","0","98.58","440.72","0","442.15","34.13","0"
"AGAW3","02N","INS","BOLD:ACK1651","Arthropoda","Insecta","Diptera","Cecidomyiidae","","","","147","126","126","28","14.29","0","98.72","437.48","0","439.06","27.08","0"
"AGAW3","02N","INS","BOLD:ABA5313","Arthropoda","Insecta","Coleoptera","Staphylinidae","Aleocharinae","Amischa","Amischa analis, Amischa bifoveolata, Amischa nigrofusca","147","121","121","28","17.69","0","99.24","446.39","0","448.03","33.31","0"
"AGAW3","02N","INS","BOLD:ABU5525","Arthropoda","Insecta","Diptera","Chironomidae","Orthocladiinae","Limnophyes","Limnophyes sp. 14ES","117","113","113","30","3.42","0","99.21","417.91","0","420.87","37.43","0"
"AGAW3","02N","INS","BOLD:AAC4197","Arthropoda","Insecta","Diptera","Chironomidae","Orthocladiinae","Paraphaenocladius","Paraphaenocladius impensus","111","107","107","31","3.6","0","99.16","434.79","0","441.18","34.79","0"
"AGAW3","02N","INS","BOLD:ACP6694","Arthropoda","Insecta","Diptera","Chironomidae","Orthocladiinae","","","76","72","72","32","5.26","0","99.22","427.04","0","428.7","33.98","0"
"AGAW3","02N","INS","BOLD:AAP7026","Arthropoda","Insecta","Coleoptera","Latridiidae","Corticariinae","Melanophthalma","Melanophthalma villosa","68","57","57","33","16.18","0","97.73","402.75","0","449.32","31.34","0"
"AGAW3","02N","INS","BOLD:ABA1215","Arthropoda","Insecta","Diptera","Sciaridae","","Lycoriella","Lycoriella sativae","68","59","59","33","13.24","0","99.39","443.24","0","444.79","35.78","0"
"AGAW3","02N","INS","BOLD:AAN8770","Arthropoda","Insecta","Diptera","Psychodidae","Psychodinae","Psychoda","Psychoda trinodulosa","63","56","56","35","11.11","0","99.4","447","0","447.46","33.98","0"
"AGAW3","02N","INS","BOLD:AAG1000","Arthropoda","Insecta","Diptera","Chironomidae","Orthocladiinae","Orthocladius","Orthocladius carlatus","51","51","51","36","0","0","97.95","433.88","0","435.94","34.03","0"
"AGAW3","02N","INS","BOLD:AAL0908","Arthropoda","Insecta","Coleoptera","Chrysomelidae","Galerucinae","Dibolia, Psylliodes","Dibolia borealis","42","41","41","37","2.38","0","99","441.93","0","444.57","34.07","0"
"AGAW3","02N","INS","BOLD:AAA3453","Arthropoda","Insecta","Diptera","Anthomyiidae","Anthomyiinae","Delia","","42","41","41","37","2.38","0","98.84","444.05","0","445.83","34.48","0"
"AGAW3","02N","INS","BOLD:ABV9274","Arthropoda","Insecta","Diptera","Cecidomyiidae","","","","42","42","42","37","0","0","99.42","427.07","0","428.55","33.29","0"
"AGAW3","02N","INS","BOLD:AAB9140","Arthropoda","Insecta","Diptera","Calliphoridae","Chrysomyinae","Phormia","Phormia regina","41","40","40","40","2.44","0","99.13","442.34","0","443.85","33.04","0"
"AGAW3","02N","INS","BOLD:ACW5117","Arthropoda","Insecta","Diptera","Chironomidae","Orthocladiinae","Smittia","Smittia leucopogon","33","33","33","41","0","0","99.36","438.52","0","439.82","31.22","0"
"AGAW3","02N","INS","BOLD:ACL7849","","","","","","","","29","25","25","42","13.79","0","99.58","376.1","0","445","33.63","0"
"AGAW3","02N","INS","BOLD:ACI4909","","","","","","","","29","29","29","42","0","0","99.38","447.17","0","449.03","36.58","0"
"AGAW3","02N","INS","BOLD:AAM9252","Arthropoda","Insecta","Diptera","Sciaridae","","Bradysia, Leptosciarella","Bradysia vagans, Leptosciarella zanti","28","26","26","44","7.14","0","99.53","447.11","0","448.04","39","0"
"AGAW3","02N","INS","BOLD:ADH9867","Arthropoda","Insecta","Diptera","","","","","26","22","22","45","15.38","0","99.19","441.46","0","444.5","37.65","0"
"AGAW3","02N","INS","BOLD:ACF4289","Arthropoda","Insecta","Coleoptera","Staphylinidae","","","","23","22","22","46","4.35","0","99.1","440.17","0","446.78","33.58","0"
"AGAW3","02N","INS","BOLD:ADH4889","Arthropoda","Insecta","Diptera","","","","","22","22","22","47","0","0","98.87","443.55","0","444.77","35.45","0"
"AGAW3","02N","INS","BOLD:AAN5356","Arthropoda","Insecta","Diptera","Chironomidae","Orthocladiinae","Smittia","","22","22","22","47","0","0","98.46","448.95","0","450","34.94","0"
"AGAW3","02N","INS","BOLD:AAF9314","Arthropoda","Insecta","Diptera","Psychodidae","Psychodinae","Psychoda","Psychoda sp., Psychoda minuta","20","17","17","49","15","0","98.31","448.95","0","450","35.26","0"
"AGAW3","02N","INS","BOLD:AAU6603","Arthropoda","Insecta","Diptera","Chironomidae","Orthocladiinae","","","15","15","15","50","0","0","98.42","438.73","0","441.13","35.28","0"
"AGAW3","02N","INS","BOLD:ABU5520","Arthropoda","Insecta","Diptera","Sciaridae","","Corynoptera","","14","13","13","51","7.14","0","99.03","422.5","0","423.79","35.3","0"
"AGAW3","02N","INS","BOLD:AAA0178","Arthropoda","Insecta","Lepidoptera","Gelechiidae","Dichomeridinae","Dichomeris","Dichomeris santarosensis","14","12","12","51","14.29","0","97.26","328.43","0","449.93","31.94","0"
"AGAW3","02N","INS","BOLD:AAH3920","Arthropoda","Insecta","Diptera","Sciaridae","","Dichopygina, Scatopsciara","","13","13","13","53","0","0","99.14","395.92","0","440.69","36.3","0"
"AGAW3","02N","INS","BOLD:AAB0868","Arthropoda","Insecta","Diptera","Calliphoridae","Calliphorinae","Cynomya","Cynomya cadaverina, Cynomya mortuorum","11","10","10","54","9.09","0","99.14","437.36","0","438.18","34.74","0"
"AGAW3","02N","INS","BOLD:AAG4944","Arthropoda","Insecta","Diptera","Mycetophilidae","Sciophilinae","Sciophila","Sciophila sp. CJB1","11","11","11","54","0","0","98.03","425.91","0","429.18","33.73","0"
"AGAW3","02N","INS","BOLD:AAN5178","Arthropoda","Insecta","Diptera","Cecidomyiidae","Cecidomyiinae","Rhopalomyia","Rhopalomyia sp. BOLD:AAN5178","8","8","8","56","0","0","99.38","439.63","0","444.13","26.9","0"
"AGAW3","02N","INS","BOLD:AAZ5626","Arthropoda","Insecta","Diptera","Sciaridae","","Bradysia","Bradysia cellarum","7","7","7","57","0","0","99.39","427.14","0","429.43","31.86","0"
"AGAW3","02N","INS","BOLD:ABZ8841","Arthropoda","Insecta","Coleoptera","Staphylinidae","Tachyporinae","Tachyporus","Tachyporus hypnorum","6","6","6","58","0","0","97.88","448.5","0","449.17","33.02","0"
"AGAW3","02N","INS","BOLD:ABY0357","Arthropoda","Insecta","Hymenoptera","Megaspilidae","","","","6","6","6","58","0","0","98.63","448.5","0","450","30.19","0"
"AGAW3","02N","INS","BOLD:AAM6047","Arthropoda","Insecta","Diptera","Cecidomyiidae","","","","5","5","5","60","0","0","99.23","434","0","436.4","32.12","0"
"AGAW3","02N","INS","BOLD:ABU9891","Arthropoda","Insecta","Diptera","Muscidae","Phaoniinae","Phaonia","","4","4","4","61","0","0","98.62","434.5","0","437.5","32.73","0"
"AGAW3","02N","INS","BOLD:ABW2820","Arthropoda","Insecta","Coleoptera","Staphylinidae","Aleocharinae","Philhygra","","4","4","4","61","0","0","99.33","449.25","0","450","36.94","0"
"AGAW3","02N","INS","BOLD:AAG1742","Arthropoda","Insecta","Diptera","Muscidae","Phaoniinae","Helina","Helina rufitibia","4","4","4","61","0","0","98.26","445.25","0","450","30","0"
"AGAW3","02N","INS","BOLD:AAB2384","Arthropoda","Insecta","Diptera","Syrphidae","Syrphinae","Eupeodes","Eupeodes duseki, Eupeodes volucris, Eupeodes nielseni, Eupeodes americanus, Eupeodes flukei, Eupeodes vockerothi, Eupeodes lucasi, Eupeodes montivagus, Eupeodes rufipunctatus, Eupeodes rojasi, Eupeodes goeldlini, Eupeodes luniger, Eupeodes punctifer, Eupeodes bucculatus, Eupeodes perplexus, Eupeodes abiskoensis, Eupeodes corollae, Eupeodes pomus, Eupeodes lundbecki, Eupeodes nitens, Eupeodes sp. BOLDAAB2384, Eupeodes curtus, Eupeodes latifasciatus","4","4","4","61","0","0","98.4","423.75","0","427","31.32","0"
"AGAW3","02N","INS","BOLD:AAH1890","Arthropoda","Insecta","Hymenoptera","Ichneumonidae","Cryptinae","Stibeutes","Stibeutes brevicornis, Stibeutes heinemanni","3","3","3","65","0","0","99.52","431.67","0","433","29.61","0"
"AGAW3","02N","INS","BOLD:ACR4394","Arthropoda","Insecta","Diptera","Anthomyiidae","Anthomyiinae","Delia","Delia florilega","3","3","3","65","0","0","98.85","423.67","0","426","33.07","0"
"AGAW3","02N","INS","BOLD:AAD0245","Arthropoda","Insecta","Hymenoptera","Ichneumonidae","Orthocentrinae","Stenomacrus","Stenomacrus jft17, Stenomacrus JFT17","3","3","3","65","0","0","98.57","397","0","398.67","26.64","0"
"AGAW3","02N","INS","BOLD:AAN5501","Arthropoda","Insecta","Diptera","Hybotidae","","","","3","3","3","65","0","0","98.87","418","0","420","33.93","0"
"AGAW3","02N","INS","BOLD:AAA3470","Arthropoda","Insecta","Lepidoptera","Gelechiidae","","gelJanzen01","gelJanzen01 Janzen390, gelJanzen01 Janzen391","3","2","2","65","33.33","0","97.11","450","0","450","31.93","0"
"AGAW3","02N","INS","BOLD:AAG2462","Arthropoda","Insecta","Diptera","Anthomyiidae","Anthomyiinae","Botanophila","Botanophila fugax","3","3","3","65","0","0","98.44","448.33","0","450","32.22","0"
"AGAW3","02N","INS","BOLD:AAA4222","Arthropoda","Insecta","Lepidoptera","Erebidae","Hypeninae","Hypena","Hypena scabra","2","2","2","71","0","0","98.89","449","0","450","32.67","0"
"AGAW3","02N","INS","BOLD:AAB7361","Arthropoda","Insecta","Diptera","Chironomidae","Orthocladiinae","Limnophyes","Limnophyes natalensis","2","2","2","71","0","0","99.23","358.5","0","386","38.21","0"
"AGAW3","02N","INS","BOLD:AAQ0582","Arthropoda","Insecta","Diptera","Anthomyiidae","Anthomyiinae","Lasiomma","","2","2","2","71","0","0","99.11","449.5","0","450","32","0"
"AGAW3","02N","INS","BOLD:ADH9310","Arthropoda","Insecta","Diptera","Anthomyiidae","Anthomyiinae","Anthomyia","Anthomyia procellaris, Anthomyia pluvialis","2","2","2","71","0","0","97.66","428.5","0","431.5","32.43","0"
"AGAW3","02N","INS","BOLD:ACX5521","Arthropoda","Insecta","Diptera","Chironomidae","Orthocladiinae","","","2","2","2","71","0","0","98.35","418","0","420","34.64","0"
"AGAW3","02N","INS","BOLD:AAD2085","Arthropoda","Insecta","Lepidoptera","Bucculatricidae","","Bucculatrix","Bucculatrix angustata","2","2","2","71","0","0","99.22","450","0","450","33.22","0"
"AGAW3","02N","INS","BOLD:AAA3181","Arthropoda","Insecta","Diptera","Muscidae","Muscinae","Stomoxys","Stomoxys calcitrans","1","1","1","77","0","0","99.55","448","0","450","33.11","0"
"AGAW3","02N","INS","BOLD:ACU5687","Arthropoda","Insecta","Coleoptera","Chrysomelidae","Galerucinae","Dibolia","Dibolia chelones","1","1","1","77","0","0","98.43","445","0","450","34.44","0"
"AGAW3","02N","INS","BOLD:ACF5890","Arthropoda","Insecta","Diptera","Chironomidae","Chironominae","","","1","1","1","77","0","0","99.33","447","0","448","31.25","0"
"AGAW3","02N","INS","BOLD:AAK8831","Arthropoda","Insecta","Diptera","Limoniidae","Limoniinae","Metalimnobia, Limonia","","1","1","1","77","0","0","99.78","448","0","450","33.11","0"
"AGAW3","02N","INS","BOLD:AAG1711","Arthropoda","Insecta","Diptera","Muscidae","Phaoniinae","Helina","Helina depuncta","1","1","1","77","0","0","100","448","0","450","31.56","0"
"AGAW3","02N","INS","BOLD:AAG7276","Arthropoda","Insecta","Diptera","Sphaeroceridae","Limosininae","Leptocera","Leptocera erythrocera","1","1","1","77","0","0","99.78","450","0","450","33.11","0"
"AGAW3","02N","INS","BOLD:AAA3750","Arthropoda","Insecta","Diptera","Culicidae","Culicinae","Ochlerotatus, Aedes","Aedes cataphylla, Aedes implicatus, Aedes nigripes, Aedes impiger, Aedes leucomelas, Aedes sp., Ochlerotatus nigripes","1","1","1","77","0","0","98.88","356","0","357","34.17","0"
"AGAW3","02N","INS","BOLD:AAA4121","Arthropoda","Insecta","Diptera","Simuliidae","Simuliinae","Simulium","Simulium vittatum complex, Simulium vittatum, Simulium sp. BOLDAAA4121, Simulium argus, Simulium vittatum s.str.","1","1","1","77","0","0","97.02","403","0","404","32.67","0"
"AGAW3","02N","INS","BOLD:AAL7534","Arthropoda","Insecta","Diptera","Anisopodidae","Anisopodinae","Sylvicola","Sylvicola stackelbergi, Sylvicola stackelbergii 1, Sylvicola punctatus","1","1","1","77","0","0","100","449","0","450","34","0"
"AGAW3","02N","INS","BOLD:ABW5541","Arthropoda","Insecta","Diptera","Sciaridae","","Lycoriella","Lycoriella agraria","1","1","1","77","0","0","97.53","446","0","450","37.11","0"
"AGAW3","02N","INS","BOLD:AAA3882","Arthropoda","Insecta","Lepidoptera","Pterophoridae","Pterophorinae","Emmelina","Emmelina monodactyla","1","1","1","77","0","0","99.06","424","0","438","32.65","0"
"AGAW3","02N","INS","BOLD:AAB5609","Arthropoda","Insecta","Diptera","Muscidae","Coenosiinae","Coenosia","Coenosia sp.BOLD:AAB5609, Coenosia tigrina","1","1","1","77","0","0","97.94","388","0","391","32.23","0"
"AGAW3","02N","INS","BOLD:AAA2482","Arthropoda","Insecta","Lepidoptera","Noctuidae","Noctuinae","Mythimna","Mythimna unipuncta","1","1","1","77","0","0","99.78","447","0","450","30.89","0"
"AGAW3","02N","INS","BOLD:AAB7872","Arthropoda","Insecta","Diptera","Chironomidae","Orthocladiinae","Orthocladius","Orthocladius oliveri","1","1","1","77","0","0","100","449","0","450","33.11","0"
"AGAW3","02N","INS","BOLD:AAM7064","Arthropoda","Insecta","Diptera","Chironomidae","Orthocladiinae","Smittia","Smittia sp. 14ES","1","1","1","77","0","0","98.46","389","0","400","31.5","0"
"AGAW3","02N","INS","BOLD:ABW1509","Arthropoda","Insecta","Diptera","Chironomidae","Orthocladiinae","","","1","1","1","77","0","0","100","449","0","450","35.33","0"
"AGAW3","02N","INS","BOLD:AAK5928","Arthropoda","Insecta","Coleoptera","Chrysomelidae","Criocerinae","Oulema","Oulema melanopus","1","1","1","77","0","0","98.44","450","0","450","35.33","0"
"AGAW3","02N","INS","BOLD:ACP4925","Arthropoda","Insecta","Diptera","Sciaridae","","","","1","1","1","77","0","0","97.94","389","0","450","36.22","0"
"AGAW3","02N","INS","BOLD:AAB1024","Arthropoda","Insecta","Lepidoptera","Noctuidae","Noctuinae","Pseudohermonassa","Pseudohermonassa bicarnea","1","1","1","77","0","0","99.77","442","0","446","34.08","0"
"AGAW3","02N","INS","BOLD:ACM1547","Arthropoda","Insecta","Hymenoptera","Diapriidae","","","","1","1","1","77","0","0","99.52","419","0","450","22.67","0"
"AGAW3","02N","INS","BOLD:ABW2407","Arthropoda","Insecta","Diptera","Anthomyiidae","Anthomyiinae","Hylemya","","1","1","1","77","0","0","97.98","446","0","450","32.67","0"
"AGAW3","02N","INS","BOLD:AAH0134","Arthropoda","Insecta","Coleoptera","Phalacridae","Phalacrinae","Stilbus","Stilbus apicialis","1","1","1","77","0","0","99.33","449","0","450","31.78","0"
"AGAW3","02N","INS","BOLD:AAN6435","Arthropoda","Insecta","Diptera","Sciaridae","","Bradysia","Bradysia angustipennis","1","1","1","77","0","0","99.75","398","0","450","34.44","0"
"AGAW3","02N","INS","BOLD:AAB5640","Arthropoda","Insecta","Coleoptera","Coccinellidae","Coccinellinae","Harmonia","Harmonia axyridis","1","1","1","77","0","0","99.78","450","0","450","32.22","0"
"AGAW3","02N","INS","BOLD:ACI2754","Arthropoda","Insecta","Coleoptera","Chrysomelidae","Galerucinae","Dibolia","Dibolia melampyri","1","1","1","77","0","0","98.19","386","0","408","32.6","0"
"AGAW3","02N","INS","BOLD:AAA5803","Arthropoda","Insecta","Hemiptera","Miridae","Mirinae","Lygus","Lygus lineolaris","1","1","1","77","0","0","97.55","449","0","450","33.56","0"
"AGAW3","02N","NAR","no match","no match","no match","no match","no match","no match","no match","no match","no match","no match","no match","no match","no match","no match","no match","no match","no match","no match","no match","no match"
"AGAW3","02N","NIN","BOLD:AAB0726","Arthropoda","Arachnida","Araneae","Lycosidae",NA,"Trochosa","Trochosa ruricola","894","833","833","1","6.82","0","98.91","439.84","0","442.16","31.14","0"
"AGAW3","02N","NIN","BOLD:ABX0027","Arthropoda","Collembola","Symphypleona","Bourletiellidae",NA,"Bourletiella","","48","42","42","2","12.5","0","99.56","416.79","0","446.31","35.66","0"
"AGAW3","02N","NIN","BOLD:AAN6707","Arthropoda","Arachnida","Mesostigmata","Ascidae",NA,"Arctoseius","","38","37","37","3","2.63","0","99.55","443.42","0","446.39","42.39","0"
"AGAW3","02N","NIN","BOLD:ADG3289","Arthropoda","Arachnida","Mesostigmata","Ascidae",NA,"","","23","18","18","4","21.74","0","99.61","391.48","0","448.43","36.04","0"
"AGAW3","02N","NIN","BOLD:ACF8020","Arthropoda","Arachnida","Mesostigmata","Ascidae",NA,"","","13","12","12","5","7.69","0","99.43","448.77","0","450","37.01","0"
"AGAW3","02N","NIN","BOLD:ABW2693","Arthropoda","Arachnida","Mesostigmata","Ascidae",NA,"","","12","11","11","6","8.33","0","99.29","443.92","0","445.67","35.82","0"
"AGAW3","02N","NIN","BOLD:ACL9406","Arthropoda","Arachnida","Trombidiformes","Cunaxidae",NA,"","","2","2","2","7","0","0","100","415","0","439.5","25.26","0"
"AGAW3","02N","NIN","BOLD:AAI4087","Arthropoda","Arachnida","Araneae","Clubionidae",NA,"Clubiona","Clubiona pallidula","1","1","1","8","0","0","99.33","449","0","450","31.78","0"
"AGAW3","02N","NIN","BOLD:ACF8021","Arthropoda","Arachnida","Mesostigmata","Ascidae",NA,"Arctoseius","Arctoseius cetratus","1","1","1","8","0","0","99.04","417","0","419","37.71","0"
"AGAW3","02S","BAC","TAX:175768","Proteobacteria","Alphaproteobacteria","Rickettsiales","Rickettsiaceae",NA,"Wolbachia","Wolbachia pipientis","21","17","17","1","19.05","0","98.74","445.95","0","446.76","39.51","0"
"AGAW3","02S","BAC","TAX:91393","Proteobacteria","Alphaproteobacteria","Rickettsiales","Rickettsiaceae",NA,"Wolbachia","","6","6","6","2","0","0","99.53","391.67","0","438.33","39.2","0"
"AGAW3","02S","BAC","TAX:91391","Proteobacteria","Alphaproteobacteria","Rickettsiales","",NA,"","","2","2","2","3","0","0","99.05","419","0","433","38.9","0"
"AGAW3","02S","BAC","TAX:91389","Proteobacteria","","","",NA,"","","1","1","1","4","0","0","99.45","362","0","409","37.65","0"
"AGAW3","02S","CHO","no match","no match","no match","no match","no match","no match","no match","no match","no match","no match","no match","no match","no match","no match","no match","no match","no match","no match","no match","no match"
"AGAW3","02S","INS","BOLD:AAG2511","Arthropoda","Insecta","Diptera","Anthomyiidae","Anthomyiinae","Delia","Delia platura","91974","57057","57057","1","37.96","0","99.48","443.31","0","445.61","33.16","0"
"AGAW3","02S","INS","BOLD:AAB2384","Arthropoda","Insecta","Diptera","Syrphidae","Syrphinae","Eupeodes","Eupeodes duseki, Eupeodes volucris, Eupeodes nielseni, Eupeodes americanus, Eupeodes flukei, Eupeodes vockerothi, Eupeodes lucasi, Eupeodes montivagus, Eupeodes rufipunctatus, Eupeodes rojasi, Eupeodes goeldlini, Eupeodes luniger, Eupeodes punctifer, Eupeodes bucculatus, Eupeodes perplexus, Eupeodes abiskoensis, Eupeodes corollae, Eupeodes pomus, Eupeodes lundbecki, Eupeodes nitens, Eupeodes sp. BOLDAAB2384, Eupeodes curtus, Eupeodes latifasciatus","17424","14592","14592","2","16.25","0","98.66","428.08","0","430.64","31.05","0"
"AGAW3","02S","INS","BOLD:AAA3453","Arthropoda","Insecta","Diptera","Anthomyiidae","Anthomyiinae","Delia","","5049","3836","3836","3","24.02","0","99.16","444.66","0","446.39","34.63","0"
"AGAW3","02S","INS","BOLD:AAH0127","Arthropoda","Insecta","Coleoptera","Pyrochroidae","Pedilinae","Pedilus","Pedilus lugubris","1057","811","811","4","23.27","0","99.4","397.18","0","442.94","32.32","0"
"AGAW3","02S","INS","BOLD:AAN5355","Arthropoda","Insecta","Diptera","Chironomidae","Orthocladiinae","Smittia","Smittia cf. stercoraria, Smittia stercoraria","585","493","493","5","15.73","0","99.29","407.6","0","447.22","36.21","0"
"AGAW3","02S","INS","BOLD:ACV5074","","","","","","","","483","375","375","6","22.36","0","99.33","435.76","0","436.91","31.3","0"
"AGAW3","02S","INS","BOLD:AAD0853","Arthropoda","Insecta","Diptera","Scathophagidae","Scathophaginae","Scathophaga","Scathophaga lutaria, Scathophaga hiemalis, Scathophaga furcata, Scathophaga stercoraria, Scathophaga inquinata, Scathophaga suilla","398","326","326","7","18.09","0","99.03","441.28","0","443.63","32.1","0"
"AGAW3","02S","INS","BOLD:AAB9140","Arthropoda","Insecta","Diptera","Calliphoridae","Chrysomyinae","Phormia","Phormia regina","319","237","237","8","25.71","0","99.29","443.9","0","444.94","33.06","0"
"AGAW3","02S","INS","BOLD:ACJ8761","Arthropoda","Insecta","Diptera","Chironomidae","Chironominae","Chironomus","","271","214","214","9","21.03","0","99.34","445.73","0","447","39.31","0"
"AGAW3","02S","INS","BOLD:AAH3301","Arthropoda","Insecta","Coleoptera","Coccinellidae","Chilocorinae","Chilocorus","Chilocorus stigma","179","139","139","10","22.35","0","98.62","446.12","0","447.25","30.43","0"
"AGAW3","02S","INS","BOLD:AAK5928","Arthropoda","Insecta","Coleoptera","Chrysomelidae","Criocerinae","Oulema","Oulema melanopus","78","62","62","11","20.51","0","99.5","448.94","0","449.71","36.29","0"
"AGAW3","02S","INS","BOLD:ACA4263","Arthropoda","Insecta","Diptera","Chironomidae","Orthocladiinae","Bryophaenocladius","","75","69","69","12","8","0","98.84","394.95","0","436.57","34.48","0"
"AGAW3","02S","INS","BOLD:ABU9891","Arthropoda","Insecta","Diptera","Muscidae","Phaoniinae","Phaonia","","48","48","48","13","0","0","99.08","432.56","0","436.88","32.45","0"
"AGAW3","02S","INS","BOLD:AAL7370","Arthropoda","Insecta","Diptera","Chironomidae","Tanypodinae","Procladius","","40","38","38","14","5","0","98.9","437.48","0","439.45","32.35","0"
"AGAW3","02S","INS","BOLD:AAL9076","Arthropoda","Insecta","Diptera","Phoridae","Metopininae","Megaselia","","26","26","26","15","0","0","98.52","420.19","0","429.27","32.59","0"
"AGAW3","02S","INS","BOLD:AAC4197","Arthropoda","Insecta","Diptera","Chironomidae","Orthocladiinae","Paraphaenocladius","Paraphaenocladius impensus","26","23","23","15","11.54","0","98.95","441.92","0","443.5","34.65","0"
"AGAW3","02S","INS","BOLD:AAB2644","Arthropoda","Insecta","Diptera","Chironomidae","Orthocladiinae","Orthocladius","Orthocladius lapponicus, Orthocladius dorenus, Orthocladius mallochi","25","25","25","17","0","0","98.79","438.08","0","440.36","32.68","0"
"AGAW3","02S","INS","BOLD:AAP6827","Arthropoda","Insecta","Diptera","Cecidomyiidae","","","","25","23","23","17","8","0","99.07","447.16","0","448.08","31.49","0"
"AGAW3","02S","INS","BOLD:ACF4289","Arthropoda","Insecta","Coleoptera","Staphylinidae","","","","20","20","20","19","0","0","98.56","438.45","0","439.75","34.07","0"
"AGAW3","02S","INS","BOLD:AAO8538","Arthropoda","Insecta","Diptera","Psychodidae","Psychodinae","Psychoda","","17","17","17","20","0","0","99.29","428.18","0","430.18","32.63","0"
"AGAW3","02S","INS","BOLD:AAM9117","Arthropoda","Insecta","Hymenoptera","Ichneumonidae","Cryptinae","","","16","15","15","21","6.25","0","99.3","416.69","0","430.75","28.27","0"
"AGAW3","02S","INS","BOLD:AAG1010","Arthropoda","Insecta","Diptera","Chironomidae","Orthocladiinae","Paraphaenocladius","","16","16","16","21","0","0","98.66","439.88","0","441.38","33.13","0"
"AGAW3","02S","INS","BOLD:AAZ5592","Arthropoda","Insecta","Diptera","Psychodidae","Psychodinae","","","15","13","13","23","13.33","0","99.02","439.33","0","440.33","32.71","0"
"AGAW3","02S","INS","BOLD:ACW5117","Arthropoda","Insecta","Diptera","Chironomidae","Orthocladiinae","Smittia","Smittia leucopogon","14","13","13","24","7.14","0","99.19","435.71","0","436.71","31.09","0"
"AGAW3","02S","INS","BOLD:AAL7534","Arthropoda","Insecta","Diptera","Anisopodidae","Anisopodinae","Sylvicola","Sylvicola stackelbergi, Sylvicola stackelbergii 1, Sylvicola punctatus","14","14","14","24","0","0","99.05","437.64","0","439","33.74","0"
"AGAW3","02S","INS","BOLD:AAN5358","Arthropoda","Insecta","Diptera","Chironomidae","Orthocladiinae","Smittia","Smittia stercoraria, Smittia sp. 22ES","14","14","14","24","0","0","99.36","445.43","0","446.71","37.47","0"
"AGAW3","02S","INS","BOLD:AAG2462","Arthropoda","Insecta","Diptera","Anthomyiidae","Anthomyiinae","Botanophila","Botanophila fugax","13","12","12","27","7.69","0","99.42","447.77","0","448.85","31.71","0"
"AGAW3","02S","INS","BOLD:AAB0868","Arthropoda","Insecta","Diptera","Calliphoridae","Calliphorinae","Cynomya","Cynomya cadaverina, Cynomya mortuorum","11","11","11","28","0","0","99.21","440.27","0","441.27","34.99","0"
"AGAW3","02S","INS","BOLD:AAH2753","Arthropoda","Insecta","Coleoptera","Anthicidae","Anthicinae","Anthicus","Anthicus cervinus","11","11","11","28","0","0","98.4","416.09","0","431.64","34.71","0"
"AGAW3","02S","INS","BOLD:AAG6745","Arthropoda","Insecta","Diptera","Calliphoridae","Calliphorinae","Pollenia","Pollenia pediculata","11","11","11","28","0","0","99.21","419.36","0","420.91","33.42","0"
"AGAW3","02S","INS","BOLD:AAV5688","Arthropoda","Insecta","Diptera","Cecidomyiidae","","","","10","10","10","31","0","0","99.45","429.1","0","430.8","34.08","0"
"AGAW3","02S","INS","BOLD:AAV0234","Arthropoda","Insecta","Hemiptera","Triozidae","Triozinae","Trioza","","7","7","7","32","0","0","97.96","431.29","0","432.71","37.76","0"
"AGAW3","02S","INS","BOLD:AAB1098","Arthropoda","Insecta","Diptera","Culicidae","Culicinae","Ochlerotatus, Aedes","Aedes excrucians, Aedes cantans, Aedes stimulans, Aedes euedes, Aedes riparius, Aedes annulipes, Ochlerotatus cantans, Aedes fitchii, Aedes sp., Ochlerotatus annulipes, Ochlerotatus excrucians","7","7","7","32","0","0","98.38","443.14","0","444","35.23","0"
"AGAW3","02S","INS","BOLD:ADH4889","Arthropoda","Insecta","Diptera","","","","","6","6","6","34","0","0","99.22","446.5","0","447.83","35.84","0"
"AGAW3","02S","INS","BOLD:AAN5267","Arthropoda","Insecta","Diptera","Cecidomyiidae","","","","6","6","6","34","0","0","99.59","448.67","0","450","30.63","0"
"AGAW3","02S","INS","BOLD:AAD4355","Arthropoda","Insecta","Diptera","Culicidae","Culicinae","Aedes","Aedes provocans","6","6","6","34","0","0","99.51","446","0","447.17","32.24","0"
"AGAW3","02S","INS","BOLD:AAH0134","Arthropoda","Insecta","Coleoptera","Phalacridae","Phalacrinae","Stilbus","Stilbus apicialis","5","5","5","37","0","0","99.48","417.6","0","418.8","32.08","0"
"AGAW3","02S","INS","BOLD:ACJ0208","Arthropoda","Insecta","Diptera","Chironomidae","Orthocladiinae","","","5","5","5","37","0","0","98.44","437","0","450","34.84","0"
"AGAW3","02S","INS","BOLD:AAA4121","Arthropoda","Insecta","Diptera","Simuliidae","Simuliinae","Simulium","Simulium vittatum complex, Simulium vittatum, Simulium sp. BOLDAAA4121, Simulium argus, Simulium vittatum s.str.","4","4","4","39","0","0","99.08","434.25","0","435.5","33.12","0"
"AGAW3","02S","INS","BOLD:AAU6967","Arthropoda","Insecta","Coleoptera","Chrysomelidae","Galerucinae","Psylliodes","Psylliodes affinis","4","4","4","39","0","0","99.58","428.75","0","434.75","34.88","0"
"AGAW3","02S","INS","BOLD:ACB1079","Arthropoda","Insecta","Diptera","Cecidomyiidae","Micromyinae","Campylomyza","Campylomyza dilatata, CecidInt44 sp. BOLD:ACB1079","4","4","4","39","0","0","98.44","449","0","449.75","28.46","0"
"AGAW3","02S","INS","BOLD:AAA4222","Arthropoda","Insecta","Lepidoptera","Erebidae","Hypeninae","Hypena","Hypena scabra","4","4","4","39","0","0","98.73","416.75","0","419.5","32.34","0"
"AGAW3","02S","INS","BOLD:AAG9055","Arthropoda","Insecta","Hymenoptera","Vespidae","Vespinae","Vespula","Vespula germanica","3","3","3","43","0","0","99.2","428.33","0","431","30.8","0"
"AGAW3","02S","INS","BOLD:AAN6433","Arthropoda","Insecta","Diptera","Sciaridae","","Corynoptera","Corynoptera deserta","3","3","3","43","0","0","99.04","450","0","450","30.89","0"
"AGAW3","02S","INS","BOLD:AAA8401","Arthropoda","Insecta","Hymenoptera","Braconidae","Microgastrinae","Dolichogenidea","Dolichogenidea absona","3","3","3","43","0","0","99.39","433.67","0","450","25.93","0"
"AGAW3","02S","INS","BOLD:AAH3920","Arthropoda","Insecta","Diptera","Sciaridae","","Dichopygina, Scatopsciara","","3","3","3","43","0","0","98.74","398.33","0","437","35.2","0"
"AGAW3","02S","INS","BOLD:AAD7604","Arthropoda","Insecta","Coleoptera","Coccinellidae","Coccinellinae","Coleomegilla","Coleomegilla maculata","3","3","3","43","0","0","98.73","446.67","0","450","32.3","0"
"AGAW3","02S","INS","BOLD:AAG6743","Arthropoda","Insecta","Diptera","Sarcophagidae","Sarcophaginae","Sarcophaga","Sarcophaga sp. 1 MG, Sarcophaga subvicina","3","3","3","43","0","0","99.85","449","0","450","35.19","0"
"AGAW3","02S","INS","BOLD:AAF9014","Arthropoda","Insecta","Diptera","Limoniidae","Chioneinae","Symplecta","Symplecta cana, Symplecta sp. BOLDAAF9014","3","3","3","43","0","0","97.81","442.33","0","450","34.74","0"
"AGAW3","02S","INS","BOLD:ACW1780","Arthropoda","Insecta","Diptera","Cecidomyiidae","","","","2","2","2","50","0","0","99.36","388","0","450","28.78","0"
"AGAW3","02S","INS","BOLD:ACR4394","Arthropoda","Insecta","Diptera","Anthomyiidae","Anthomyiinae","Delia","Delia florilega","2","2","2","50","0","0","97.78","402","0","407","32.66","0"
"AGAW3","02S","INS","BOLD:AAG0991","Arthropoda","Insecta","Diptera","Chironomidae","Orthocladiinae","Orthocladius","","2","2","2","50","0","0","98.13","413","0","415.5","31.99","0"
"AGAW3","02S","INS","BOLD:AAA4264","Arthropoda","Insecta","Diptera","Simuliidae","Simuliinae","Simulium","Simulium venustum/truncatum, Simulium venustum s.str., Simulium verecundum/verecundum complex, Simulium venustum complex, Simulium venustum, Simulium tormentor, Simulium truncatum, Simulium sp. BOLDAAA4264, Simulium venustum complex sp. JMR-2009","2","2","2","50","0","0","98.55","448.5","0","450","36.78","0"
"AGAW3","02S","INS","BOLD:AAN5341","Arthropoda","Insecta","Diptera","Chironomidae","Orthocladiinae","Camptocladius","Camptocladius stercorarius","2","2","2","50","0","0","97.7","353.5","0","417.5","35.59","0"
"AGAW3","02S","INS","BOLD:ACM1547","Arthropoda","Insecta","Hymenoptera","Diapriidae","","","","2","2","2","50","0","0","99.53","424.5","0","450","22.44","0"
"AGAW3","02S","INS","BOLD:AAN6447","Arthropoda","Insecta","Diptera","Sciaridae","","Corynoptera","Corynoptera perpusilla","2","2","2","50","0","0","98.54","448","0","450","36.22","0"
"AGAW3","02S","INS","BOLD:AAG1000","Arthropoda","Insecta","Diptera","Chironomidae","Orthocladiinae","Orthocladius","Orthocladius carlatus","2","2","2","50","0","0","98.6","434.5","0","437","33.38","0"
"AGAW3","02S","INS","BOLD:AAM7726","Arthropoda","Insecta","Coleoptera","Curculionidae","Curculioninae","Orchestes","Orchestes steppensis, Orchestes alni","2","2","2","50","0","0","98.89","450","0","450","33.56","0"
"AGAW3","02S","INS","BOLD:AAC9062","Arthropoda","Insecta","Diptera","Culicidae","Culicinae","Ochlerotatus, Aedes","Aedes fitchii, Ochlerotatus fitchii, Aedes grossbecki, Aedes ventrovittis","2","2","2","50","0","0","99.55","449","0","450","33.56","0"
"AGAW3","02S","INS","BOLD:AAK8831","Arthropoda","Insecta","Diptera","Limoniidae","Limoniinae","Metalimnobia, Limonia","","1","1","1","60","0","0","99.78","450","0","450","33.11","0"
"AGAW3","02S","INS","BOLD:AAG1711","Arthropoda","Insecta","Diptera","Muscidae","Phaoniinae","Helina","Helina depuncta","1","1","1","60","0","0","98.89","450","0","450","31.33","0"
"AGAW3","02S","INS","BOLD:AAB2641","Arthropoda","Insecta","Diptera","Chironomidae","Orthocladiinae","Orthocladius","","1","1","1","60","0","0","98.89","450","0","450","32.67","0"
"AGAW3","02S","INS","BOLD:AAG7276","Arthropoda","Insecta","Diptera","Sphaeroceridae","Limosininae","Leptocera","Leptocera erythrocera","1","1","1","60","0","0","99.11","450","0","450","32.89","0"
"AGAW3","02S","INS","BOLD:AAA8933","Arthropoda","Insecta","Coleoptera","Coccinellidae","Coccinellinae","Coccinella","Coccinella septempunctata","1","1","1","60","0","0","100","450","0","450","33.11","0"
"AGAW3","02S","INS","BOLD:AAA2955","Arthropoda","Insecta","Lepidoptera","Tortricidae","Tortricinae","Argyrotaenia","Argyrotaenia occultana, Argyrotaenia ljungiana, Argyrotaenia pinatubana, Argyrotaenia floridana, Argyrotaenia velutinana, Argyrotaenia tabulana, Argyrotaenia gogana","1","1","1","60","0","0","99.11","448","0","450","32","0"
"AGAW3","02S","INS","BOLD:AAI2688","Arthropoda","Insecta","Diptera","Chironomidae","Orthocladiinae","Parametriocnemus","","1","1","1","60","0","0","99.11","450","0","450","32.89","0"
"AGAW3","02S","INS","BOLD:AAV1366","Arthropoda","Insecta","Diptera","Sciaridae","","Bradysia","Bradysia fenestralis","1","1","1","60","0","0","99.33","447","0","450","33.33","0"
"AGAW3","02S","INS","BOLD:AAY6412","Arthropoda","Insecta","Diptera","Sciaridae","","Bradysia","Bradysia urticae","1","1","1","60","0","0","97.1","449","0","450","32.67","0"
"AGAW3","02S","INS","BOLD:AAH3914","Arthropoda","Insecta","Diptera","Sciaridae","","Sciara","Sciara hamatilis, Sciara humeralis","1","1","1","60","0","0","99.55","448","0","450","30.67","0"
"AGAW3","02S","INS","BOLD:AAG7312","Arthropoda","Insecta","Diptera","Sphaeroceridae","Limosininae","Spelobia","Spelobia clunipes, Spelobia sp. B TAW","1","1","1","60","0","0","97.32","448","0","450","32.67","0"
"AGAW3","02S","INS","BOLD:ABW7747","Arthropoda","Insecta","Coleoptera","Staphylinidae","Aleocharinae","Nehemitropia","Nehemitropia lividipennis","1","1","1","60","0","0","99.11","450","0","450","36.67","0"
"AGAW3","02S","INS","BOLD:AAG1742","Arthropoda","Insecta","Diptera","Muscidae","Phaoniinae","Helina","Helina rufitibia","1","1","1","60","0","0","98.66","448","0","450","30.44","0"
"AGAW3","02S","INS","BOLD:ABU5520","Arthropoda","Insecta","Diptera","Sciaridae","","Corynoptera","","1","1","1","60","0","0","99.56","450","0","450","35.56","0"
"AGAW3","02S","INS","BOLD:AAN5351","Arthropoda","Insecta","Diptera","Chironomidae","Tanypodinae","Conchapelopia","Conchapelopia telema","1","1","1","60","0","0","98","450","0","450","34.44","0"
"AGAW3","02S","INS","BOLD:AAI1269","Arthropoda","Insecta","Coleoptera","Carabidae","Harpalinae","Lebia","Lebia bivittata","1","1","1","60","0","0","100","448","0","450","33.78","0"
"AGAW3","02S","INS","BOLD:AAN1738","Arthropoda","Insecta","Coleoptera","Staphylinidae","Oxytelinae","Anotylus","Anotylus tetracarinatus","1","1","1","60","0","0","99.78","449","0","450","33.78","0"
"AGAW3","02S","INS","BOLD:AAD5253","Arthropoda","Insecta","Diptera","Chironomidae","Orthocladiinae","Thienemanniella","Thienemanniella xena, Thienemanniella vittata","1","1","1","60","0","0","99.33","450","0","450","34.44","0"
"AGAW3","02S","INS","BOLD:AAA2482","Arthropoda","Insecta","Lepidoptera","Noctuidae","Noctuinae","Mythimna","Mythimna unipuncta","1","1","1","60","0","0","99.49","396","0","397","29.72","0"
"AGAW3","02S","INS","BOLD:AAB1024","Arthropoda","Insecta","Lepidoptera","Noctuidae","Noctuinae","Pseudohermonassa","Pseudohermonassa bicarnea","1","1","1","60","0","0","97.77","449","0","450","33.56","0"
"AGAW3","02S","INS","BOLD:AAG2123","Arthropoda","Insecta","Diptera","Tachinidae","Dexiinae","Dinera","Dinera grisescens","1","1","1","60","0","0","99.78","450","0","450","32.22","0"
"AGAW3","02S","INS","BOLD:AAG3316","Arthropoda","Insecta","Diptera","Phoridae","Metopininae","Megaselia","","1","1","1","60","0","0","99.33","449","0","450","33.56","0"
"AGAW3","02S","INS","BOLD:ACF6547","Arthropoda","Insecta","Diptera","Chironomidae","Orthocladiinae","Orthocladius","","1","1","1","60","0","0","99.72","351","0","450","32.22","0"
"AGAW3","02S","INS","BOLD:ABV1430","Arthropoda","Insecta","Diptera","Dolichopodidae","Medeterinae","Medetera","","1","1","1","60","0","0","99.78","446","0","450","39.33","0"
"AGAW3","02S","INS","BOLD:ACP4736","Arthropoda","Insecta","Diptera","Chironomidae","Orthocladiinae","Smittia","Smittia sp. 8ES","1","1","1","60","0","0","98.89","450","0","450","33.56","0"
"AGAW3","02S","INS","BOLD:AAB5640","Arthropoda","Insecta","Coleoptera","Coccinellidae","Coccinellinae","Harmonia","Harmonia axyridis","1","1","1","60","0","0","100","448","0","450","32.22","0"
"AGAW3","02S","INS","BOLD:AAE4299","Arthropoda","Insecta","Diptera","Chironomidae","Orthocladiinae","Cricotopus","","1","1","1","60","0","0","99.33","449","0","450","34","0"
"AGAW3","02S","NAR","no match","no match","no match","no match","no match","no match","no match","no match","no match","no match","no match","no match","no match","no match","no match","no match","no match","no match","no match","no match"
"AGAW3","02S","NIN","BOLD:AAB0726","Arthropoda","Arachnida","Araneae","Lycosidae","","Trochosa","Trochosa ruricola","198","187","187","1","5.56","0","98.87","441.36","0","443.27","31.13","0"
"AGAW3","02S","NIN","BOLD:AAI2838","Arthropoda","Arachnida","Araneae","Philodromidae","","Philodromus","Philodromus marxi, Philodromus imbecillus","158","134","134","2","15.19","0","98.71","441.93","0","442.96","32.76","0"
"AGAW3","02S","NIN","BOLD:AAB7667","Arthropoda","Arachnida","Araneae","Lycosidae","","Pardosa","Pardosa saxatilis","18","18","18","3","0","0","98.35","421.61","0","426.22","31.46","0"
"AGAW3","02S","NIN","BOLD:AAH8314","Arthropoda","Arachnida","Araneae","Linyphiidae","","Walckenaeria","Walckenaeria microspiralis","9","9","9","4","0","0","98.28","394.44","0","396.67","35.51","0"
"AGAW3","02S","NIN","BOLD:AAA4664","Arthropoda","Collembola","Entomobryomorpha","Entomobryidae","Lepidocyrtinae","Willowsia","Willowsia buski","1","1","1","5","0","0","98.88","448","0","450","41.78","0"
"AGAW3","02S","NIN","BOLD:AAN4906","Arthropoda","Collembola","Symphypleona","Bourletiellidae","","","","1","1","1","5","0","0","99.33","450","0","450","36.44","0"
"AGAW3","02S","NIN","BOLD:AAI4087","Arthropoda","Arachnida","Araneae","Clubionidae","","Clubiona","Clubiona pallidula","1","1","1","5","0","0","100","450","0","450","32","0"
"AGAW3","02S","NIN","BOLD:ACF1714","Arthropoda","Collembola","Entomobryomorpha","Entomobryidae","Entomobryinae","Entomobrya","Entomobrya assuta","1","1","1","5","0","0","98.21","448","0","450","39.78","0"
"AGAW3","02W","BAC","TAX:175768","Proteobacteria","Alphaproteobacteria","Rickettsiales","Rickettsiaceae",NA,"Wolbachia","Wolbachia pipientis","5","4","4","1","20","0","98.89","448.8","0","450","39.47","0"
"AGAW3","02W","BAC","TAX:375853","Proteobacteria","Alphaproteobacteria","Rickettsiales","Rickettsiaceae",NA,"Wolbachia","Wolbachia sp. Hymenoptera_ASGLE646-10","1","1","1","2","0","0","97.78","450","0","450","40.89","0"
"AGAW3","02W","BAC","TAX:91393","Proteobacteria","Alphaproteobacteria","Rickettsiales","Rickettsiaceae",NA,"Wolbachia","","1","1","1","2","0","0","99.23","388","0","450","40.67","0"
"AGAW3","02W","CHO","no match","no match","no match","no match","no match","no match","no match","no match","no match","no match","no match","no match","no match","no match","no match","no match","no match","no match","no match","no match"
"AGAW3","02W","INS","BOLD:AAB0868","Arthropoda","Insecta","Diptera","Calliphoridae","Calliphorinae","Cynomya","Cynomya cadaverina, Cynomya mortuorum","40566","24966","24966","1","38.46","0","99.53","437.82","0","439.57","34.98","0"
"AGAW3","02W","INS","BOLD:AAG2511","Arthropoda","Insecta","Diptera","Anthomyiidae","Anthomyiinae","Delia","Delia platura","37483","25853","25853","2","31.03","0","99.36","443.86","0","446.19","33.22","0"
"AGAW3","02W","INS","BOLD:AAA3453","Arthropoda","Insecta","Diptera","Anthomyiidae","Anthomyiinae","Delia","","12490","9101","9101","3","27.13","0","99.34","444.19","0","446.11","34.77","0"
"AGAW3","02W","INS","BOLD:AAB2076","Arthropoda","Insecta","Lepidoptera","Tortricidae","Olethreutinae","Ancylis","Ancylis metamelana, Ancylis sp.","6793","4325","4325","4","36.33","0","99.62","442.54","0","444.05","31.04","0"
"AGAW3","02W","INS","BOLD:AAA2482","Arthropoda","Insecta","Lepidoptera","Noctuidae","Noctuinae","Mythimna","Mythimna unipuncta","2694","2339","2339","5","13.18","0","99.25","431.49","0","434.06","30.46","0"
"AGAW3","02W","INS","BOLD:AAB1098","Arthropoda","Insecta","Diptera","Culicidae","Culicinae","Ochlerotatus, Aedes","Aedes excrucians, Aedes cantans, Aedes stimulans, Aedes euedes, Aedes riparius, Aedes annulipes, Ochlerotatus cantans, Aedes fitchii, Aedes sp., Ochlerotatus annulipes, Ochlerotatus excrucians","964","893","893","6","7.37","0","98.6","427.15","0","428.69","34.81","0"
"AGAW3","02W","INS","BOLD:AAG6454","Arthropoda","Insecta","Diptera","Ceratopogonidae","Dasyheleinae","Dasyhelea","","914","782","782","7","14.44","0","98.8","434.89","0","436.15","36.94","0"
"AGAW3","02W","INS","BOLD:AAA8109","Arthropoda","Insecta","Lepidoptera","Gelechiidae","Dichomeridinae","Dichomeris","Dichomeris ligulella","347","255","255","8","26.51","0","98.58","445.89","0","446.72","32.12","0"
"AGAW3","02W","INS","BOLD:AAD8505","Arthropoda","Insecta","Lepidoptera","Gelechiidae","Gelechiinae","Chrysoesthia","Chrysoesthia sexguttella","215","209","209","9","2.79","0","98.9","427.18","0","429.81","29.66","0"
"AGAW3","02W","INS","BOLD:AAP4697","Arthropoda","Insecta","Diptera","Phoridae","Phorinae","Triphleba","","197","171","171","10","13.2","0","99.3","445.11","0","446.53","32.82","0"
"AGAW3","02W","INS","BOLD:AAN5358","Arthropoda","Insecta","Diptera","Chironomidae","Orthocladiinae","Smittia","Smittia stercoraria, Smittia sp. 22ES","175","155","155","11","11.43","0","99.43","443.57","0","445.58","37.43","0"
"AGAW3","02W","INS","BOLD:ABV1200","Arthropoda","Insecta","Diptera","Chironomidae","Orthocladiinae","Metriocnemus","","164","136","136","12","17.07","0","99.65","441.66","0","443.18","40.87","0"
"AGAW3","02W","INS","BOLD:AAM7726","Arthropoda","Insecta","Coleoptera","Curculionidae","Curculioninae","Orchestes","Orchestes steppensis, Orchestes alni","74","68","68","13","8.11","0","99.38","441.53","0","443.14","32.95","0"
"AGAW3","02W","INS","BOLD:AAX0836","Arthropoda","Insecta","Diptera","Chironomidae","Orthocladiinae","Heterotrissocladius","Heterotrissocladius changi","65","61","61","14","6.15","0","98.37","426.88","0","435.38","37.43","0"
"AGAW3","02W","INS","BOLD:AAC4197","Arthropoda","Insecta","Diptera","Chironomidae","Orthocladiinae","Paraphaenocladius","Paraphaenocladius impensus","62","58","58","15","6.45","0","99.08","432.27","0","445.48","34.98","0"
"AGAW3","02W","INS","BOLD:AAA7661","Arthropoda","Insecta","Diptera","Culicidae","Culicinae","Culiseta, Culex","Culiseta melanura, Culex articularis, Culex restuans, Culex acharistus","60","57","57","16","5","0","99.43","433.85","0","435.38","32.88","0"
"AGAW3","02W","INS","BOLD:AAN5267","Arthropoda","Insecta","Diptera","Cecidomyiidae","","","","47","36","36","17","23.4","0","99.53","447.85","0","448.94","30.61","0"
"AGAW3","02W","INS","BOLD:AAB9140","Arthropoda","Insecta","Diptera","Calliphoridae","Chrysomyinae","Phormia","Phormia regina","46","46","46","18","0","0","99.13","442.59","0","443.67","33.23","0"
"AGAW3","02W","INS","BOLD:AAY1310","Arthropoda","Insecta","Diptera","Agromyzidae","Phytomyzinae","Phytomyza","Phytomyza agromyzina","45","38","38","19","15.56","0","99.15","442.84","0","444.69","32.47","0"
"AGAW3","02W","INS","BOLD:AAA3080","Arthropoda","Insecta","Trichoptera","Limnephilidae","Limnephilinae","Limnephilus","Limnephilus submonilifer","43","43","43","20","0","0","98.81","432.49","0","434.79","34.12","0"
"AGAW3","02W","INS","BOLD:AAN6447","Arthropoda","Insecta","Diptera","Sciaridae","","Corynoptera","Corynoptera perpusilla","41","36","36","21","12.2","0","99.45","446.98","0","448.93","36.1","0"
"AGAW3","02W","INS","BOLD:AAG4944","Arthropoda","Insecta","Diptera","Mycetophilidae","Sciophilinae","Sciophila","Sciophila sp. CJB1","39","39","39","22","0","0","97.88","408.79","0","411.77","34.07","0"
"AGAW3","02W","INS","BOLD:AAN6435","Arthropoda","Insecta","Diptera","Sciaridae","","Bradysia","Bradysia angustipennis","30","28","28","23","6.67","0","99.76","396.33","0","440.53","34.13","0"
"AGAW3","02W","INS","BOLD:AAH3920","Arthropoda","Insecta","Diptera","Sciaridae","","Dichopygina, Scatopsciara","","28","28","28","24","0","0","98.36","406.71","0","431.79","35.35","0"
"AGAW3","02W","INS","BOLD:AAM9252","Arthropoda","Insecta","Diptera","Sciaridae","","Bradysia, Leptosciarella","Bradysia vagans, Leptosciarella zanti","27","27","27","25","0","0","99.49","442.78","0","444.41","38.97","0"
"AGAW3","02W","INS","BOLD:ABX0657","Arthropoda","Insecta","Coleoptera","Anthicidae","Notoxinae","Notoxus","Notoxus desertus","23","22","22","26","4.35","0","99.26","447.3","0","448.17","35.94","0"
"AGAW3","02W","INS","BOLD:AAY6412","Arthropoda","Insecta","Diptera","Sciaridae","","Bradysia","Bradysia urticae","22","20","20","27","9.09","0","99.6","448.09","0","449.45","32.47","0"
"AGAW3","02W","INS","BOLD:AAF9314","Arthropoda","Insecta","Diptera","Psychodidae","Psychodinae","Psychoda","Psychoda sp., Psychoda minuta","15","15","15","28","0","0","98.51","440.6","0","450","34.49","0"
"AGAW3","02W","INS","BOLD:AAN5356","Arthropoda","Insecta","Diptera","Chironomidae","Orthocladiinae","Smittia","","14","13","13","29","7.14","0","98.45","445.43","0","446.5","35.3","0"
"AGAW3","02W","INS","BOLD:AAM7855","Arthropoda","Insecta","Thysanoptera","Thripidae","Thripinae","Frankliniella","Frankliniella tritici","14","14","14","29","0","0","98.89","426.64","0","429.43","31.35","0"
"AGAW3","02W","INS","BOLD:AAB2644","Arthropoda","Insecta","Diptera","Chironomidae","Orthocladiinae","Orthocladius","Orthocladius lapponicus, Orthocladius dorenus, Orthocladius mallochi","13","13","13","31","0","0","98.97","445.31","0","447.31","32.81","0"
"AGAW3","02W","INS","BOLD:AAN5355","Arthropoda","Insecta","Diptera","Chironomidae","Orthocladiinae","Smittia","Smittia cf. stercoraria, Smittia stercoraria","12","12","12","32","0","0","99.09","434.08","0","446.83","36.22","0"
"AGAW3","02W","INS","BOLD:AAL7819","Arthropoda","Insecta","Diptera","Psychodidae","Psychodinae","Psychoda","Psychoda sp., Psychoda sp. BOLD:AAL7819","8","7","7","33","12.5","0","99.71","410.5","0","412.25","32.58","0"
"AGAW3","02W","INS","BOLD:AAD0853","Arthropoda","Insecta","Diptera","Scathophagidae","Scathophaginae","Scathophaga","Scathophaga lutaria, Scathophaga hiemalis, Scathophaga furcata, Scathophaga stercoraria, Scathophaga inquinata, Scathophaga suilla","8","8","8","33","0","0","98.83","446.75","0","449","32.13","0"
"AGAW3","02W","INS","BOLD:AAA4222","Arthropoda","Insecta","Lepidoptera","Erebidae","Hypeninae","Hypena","Hypena scabra","7","7","7","35","0","0","98.74","418.29","0","421.29","32.47","0"
"AGAW3","02W","INS","BOLD:AAB2384","Arthropoda","Insecta","Diptera","Syrphidae","Syrphinae","Eupeodes","Eupeodes duseki, Eupeodes volucris, Eupeodes nielseni, Eupeodes americanus, Eupeodes flukei, Eupeodes vockerothi, Eupeodes lucasi, Eupeodes montivagus, Eupeodes rufipunctatus, Eupeodes rojasi, Eupeodes goeldlini, Eupeodes luniger, Eupeodes punctifer, Eupeodes bucculatus, Eupeodes perplexus, Eupeodes abiskoensis, Eupeodes corollae, Eupeodes pomus, Eupeodes lundbecki, Eupeodes nitens, Eupeodes sp. BOLDAAB2384, Eupeodes curtus, Eupeodes latifasciatus","7","7","7","35","0","0","98.89","442.43","0","444","31.07","0"
"AGAW3","02W","INS","BOLD:ABA5798","Arthropoda","Insecta","Hemiptera","Cicadellidae","Typhlocybinae","Erythroneura","Erythroneura elegans","7","7","7","35","0","0","98.37","444.71","0","447.29","36.09","0"
"AGAW3","02W","INS","BOLD:ACJ5126","Arthropoda","Insecta","Diptera","Chironomidae","Orthocladiinae","Metriocnemus","","6","6","6","38","0","0","99.18","430","0","433","34.64","0"
"AGAW3","02W","INS","BOLD:AAK8831","Arthropoda","Insecta","Diptera","Limoniidae","Limoniinae","Metalimnobia, Limonia","","5","5","5","39","0","0","99.15","447.8","0","450","33.29","0"
"AGAW3","02W","INS","BOLD:AAG1742","Arthropoda","Insecta","Diptera","Muscidae","Phaoniinae","Helina","Helina rufitibia","5","5","5","39","0","0","99.42","448.4","0","448.2","30.39","0"
"AGAW3","02W","INS","BOLD:ACR4394","Arthropoda","Insecta","Diptera","Anthomyiidae","Anthomyiinae","Delia","Delia florilega","5","5","5","39","0","0","98.35","448.4","0","450","33.42","0"
"AGAW3","02W","INS","BOLD:AAG6743","Arthropoda","Insecta","Diptera","Sarcophagidae","Sarcophaginae","Sarcophaga","Sarcophaga sp. 1 MG, Sarcophaga subvicina","4","4","4","42","0","0","99.05","448","0","449.75","34.96","0"
"AGAW3","02W","INS","BOLD:AAQ0582","Arthropoda","Insecta","Diptera","Anthomyiidae","Anthomyiinae","Lasiomma","","4","4","4","42","0","0","99.33","449","0","450","31.67","0"
"AGAW3","02W","INS","BOLD:AAB4093","Arthropoda","Insecta","Lepidoptera","Noctuidae","Noctuinae","Elaphria","Elaphria grata","3","3","3","44","0","0","98.78","407.33","0","410","29.51","0"
"AGAW3","02W","INS","BOLD:ABA4842","Arthropoda","Insecta","Coleoptera","Elateridae","Agrypninae","Aeolus","Aeolus mellillus","3","3","3","44","0","0","99.55","447.67","0","450","39.04","0"
"AGAW3","02W","INS","BOLD:ACC7611","Arthropoda","Insecta","Diptera","Cecidomyiidae","","","","2","2","2","46","0","0","97.1","449","0","450","28.56","0"
"AGAW3","02W","INS","BOLD:ABU5546","Arthropoda","Insecta","Diptera","Psychodidae","Psychodinae","Psychoda","","2","2","2","46","0","0","97.7","419","0","422","31.15","0"
"AGAW3","02W","INS","BOLD:AAN8770","Arthropoda","Insecta","Diptera","Psychodidae","Psychodinae","Psychoda","Psychoda trinodulosa","2","2","2","46","0","0","99.67","450","0","450","33.56","0"
"AGAW3","02W","INS","BOLD:ABU9891","Arthropoda","Insecta","Diptera","Muscidae","Phaoniinae","Phaonia","","2","2","2","46","0","0","99.42","412","0","416","32.41","0"
"AGAW3","02W","INS","BOLD:AAA4121","Arthropoda","Insecta","Diptera","Simuliidae","Simuliinae","Simulium","Simulium vittatum complex, Simulium vittatum, Simulium sp. BOLDAAA4121, Simulium argus, Simulium vittatum s.str.","2","2","2","46","0","0","99.89","449.5","0","450","33.89","0"
"AGAW3","02W","INS","BOLD:ACJ0208","Arthropoda","Insecta","Diptera","Chironomidae","Orthocladiinae","","","2","2","2","46","0","0","97.71","436.5","0","448","34.04","0"
"AGAW3","02W","INS","BOLD:AAB5640","Arthropoda","Insecta","Coleoptera","Coccinellidae","Coccinellinae","Harmonia","Harmonia axyridis","2","2","2","46","0","0","98","449.5","0","450","32.22","0"
"AGAW3","02W","INS","BOLD:AAA8933","Arthropoda","Insecta","Coleoptera","Coccinellidae","Coccinellinae","Coccinella","Coccinella septempunctata","2","2","2","46","0","0","100","449","0","450","33.11","0"
"AGAW3","02W","INS","BOLD:AAB1024","Arthropoda","Insecta","Lepidoptera","Noctuidae","Noctuinae","Pseudohermonassa","Pseudohermonassa bicarnea","2","2","2","46","0","0","99.78","449.5","0","450","33.78","0"
"AGAW3","02W","INS","BOLD:AAA0178","Arthropoda","Insecta","Lepidoptera","Gelechiidae","Dichomeridinae","Dichomeris","Dichomeris santarosensis","2","2","2","46","0","0","97.87","329","0","450","32","0"
"AGAW3","02W","INS","BOLD:AAP9022","Arthropoda","Insecta","Diptera","Cecidomyiidae","","","","2","2","2","46","0","0","99.56","450","0","450","28","0"
"AGAW3","02W","INS","BOLD:ACB1079","Arthropoda","Insecta","Diptera","Cecidomyiidae","Micromyinae","Campylomyza","Campylomyza dilatata, CecidInt44 sp. BOLD:ACB1079","2","2","2","46","0","0","98","449","0","450","28.33","0"
"AGAW3","02W","INS","BOLD:AAG1711","Arthropoda","Insecta","Diptera","Muscidae","Phaoniinae","Helina","Helina depuncta","1","1","1","58","0","0","99.78","448","0","450","31.78","0"
"AGAW3","02W","INS","BOLD:AAN8583","Arthropoda","Insecta","Diptera","Mycetophilidae","Sciophilinae","Sciophila","Sciophila pseudoflexuosa, Sciophila lutea","1","1","1","58","0","0","97.46","393","0","396","33.84","0"
"AGAW3","02W","INS","BOLD:AAK5928","Arthropoda","Insecta","Coleoptera","Chrysomelidae","Criocerinae","Oulema","Oulema melanopus","1","1","1","58","0","0","99.78","449","0","450","36.67","0"
"AGAW3","02W","INS","BOLD:AAD4355","Arthropoda","Insecta","Diptera","Culicidae","Culicinae","Aedes","Aedes provocans","1","1","1","58","0","0","97.41","348","0","351","31.91","0"
"AGAW3","02W","INS","BOLD:AAA3836","Arthropoda","Insecta","Lepidoptera","Noctuidae","Plusiinae","Autographa","Autographa precationis","1","1","1","58","0","0","99.78","448","0","450","32.22","0"
"AGAW3","02W","INS","BOLD:AAL2750","Arthropoda","Insecta","Diptera","Muscidae","Phaoniinae","Phaonia","Phaonia tuguriorum","1","1","1","58","0","0","99.78","450","0","450","33.33","0"
"AGAW3","02W","INS","BOLD:AAH0127","Arthropoda","Insecta","Coleoptera","Pyrochroidae","Pedilinae","Pedilus","Pedilus lugubris","1","1","1","58","0","0","97.66","384","0","385","31.17","0"
"AGAW3","02W","INS","BOLD:AAH3914","Arthropoda","Insecta","Diptera","Sciaridae","","Sciara","Sciara hamatilis, Sciara humeralis","1","1","1","58","0","0","100","450","0","450","30.44","0"
"AGAW3","02W","INS","BOLD:AAH3661","Arthropoda","Insecta","Diptera","Cecidomyiidae","","","","1","1","1","58","0","0","99.56","450","0","450","25.78","0"
"AGAW3","02W","INS","BOLD:AAA9251","Arthropoda","Insecta","Lepidoptera","Gelechiidae","Gelechiinae","Scrobipalpa","Scrobipalpa atriplicella, Scrobipalpa nr. atriplicella sp. 1","1","1","1","58","0","0","100","449","0","450","34.44","0"
"AGAW3","02W","INS","BOLD:AAA3470","Arthropoda","Insecta","Lepidoptera","Gelechiidae","","gelJanzen01","gelJanzen01 Janzen390, gelJanzen01 Janzen391","1","1","1","58","0","0","97.03","437","0","438","32.88","0"
"AGAW3","02W","INS","BOLD:AAA3750","Arthropoda","Insecta","Diptera","Culicidae","Culicinae","Ochlerotatus, Aedes","Aedes cataphylla, Aedes implicatus, Aedes nigripes, Aedes impiger, Aedes leucomelas, Aedes sp., Ochlerotatus nigripes","1","1","1","58","0","0","99.78","449","0","450","34","0"
"AGAW3","02W","INS","BOLD:ACE2618","Arthropoda","Insecta","Coleoptera","Ptiliidae","Acrotrichinae","Acrotrichis","","1","1","1","58","0","0","99.31","434","0","450","35.11","0"
"AGAW3","02W","INS","BOLD:ABV9277","","","","","","","","1","1","1","58","0","0","99.33","450","0","450","28.22","0"
"AGAW3","02W","INS","BOLD:AAO8538","Arthropoda","Insecta","Diptera","Psychodidae","Psychodinae","Psychoda","","1","1","1","58","0","0","99.33","450","0","450","33.11","0"
"AGAW3","02W","INS","BOLD:AAG6745","Arthropoda","Insecta","Diptera","Calliphoridae","Calliphorinae","Pollenia","Pollenia pediculata","1","1","1","58","0","0","100","448","0","450","34","0"
"AGAW3","02W","INS","BOLD:AAK8678","Arthropoda","Insecta","Lepidoptera","Noctuidae","Noctuinae","Mythimna","Mythimna pAH01Cl","1","1","1","58","0","0","97.1","448","0","450","30","0"
"AGAW3","02W","INS","BOLD:AAG8493","Arthropoda","Insecta","Diptera","Drosophilidae","Drosophilinae","Scaptomyza","Scaptomyza pallida","1","1","1","58","0","0","98.44","448","0","450","34.67","0"
"AGAW3","02W","INS","BOLD:AAU8485","Arthropoda","Insecta","Hymenoptera","Ichneumonidae","Cryptinae","","","1","1","1","58","0","0","98.89","450","0","450","30.67","0"
"AGAW3","02W","INS","BOLD:AAY7999","Arthropoda","Insecta","Diptera","Culicidae","Culicinae","Aedes","Aedes aloponotum","1","1","1","58","0","0","98","449","0","450","34.89","0"
"AGAW3","02W","INS","BOLD:AAG2462","Arthropoda","Insecta","Diptera","Anthomyiidae","Anthomyiinae","Botanophila","Botanophila fugax","1","1","1","58","0","0","99.78","447","0","450","31.78","0"
"AGAW3","02W","NAR","no match","no match","no match","no match","no match","no match","no match","no match","no match","no match","no match","no match","no match","no match","no match","no match","no match","no match","no match","no match"
"AGAW3","02W","NIN","BOLD:AAB0726","Arthropoda","Arachnida","Araneae","Lycosidae","","Trochosa","Trochosa ruricola","391","370","370","1","5.37","0","98.88","440.83","0","442.91","31.14","0"
"AGAW3","02W","NIN","BOLD:AAD1564","Arthropoda","Arachnida","Araneae","Clubionidae","","Clubiona","Clubiona abboti","244","224","224","2","8.2","0","98.18","441.07","0","443.35","31.65","0"
"AGAW3","02W","NIN","BOLD:ACA2358","Arthropoda","Arachnida","Araneae","Linyphiidae","","Glyphesis","Glyphesis scopulifer","14","14","14","3","0","0","98.68","433.93","0","440.64","38.36","0"
"AGAW3","02W","NIN","BOLD:ACL9406","Arthropoda","Arachnida","Trombidiformes","Cunaxidae","","","","5","5","5","4","0","0","98.84","411","0","436.8","25.04","0"
"AGAW3","02W","NIN","BOLD:ACE5102","Arthropoda","Collembola","Entomobryomorpha","Entomobryidae","Entomobryinae","Entomobrya","Entomobrya atrocincta","4","4","4","5","0","0","98.65","399.5","0","431.5","38.14","0"
"AGAW3","02W","NIN","BOLD:ADH9851","Arthropoda","Collembola","Entomobryomorpha","","","","","1","1","1","6","0","0","97.54","448","0","450","39.11","0"
"AGAW3","02W","NIN","BOLD:AAB5726","Arthropoda","Arachnida","Araneae","Agelenidae","","Agelenopsis","Agelenopsis pennsylvanica, Agelenopsis emertoni, Agelenopsis potteri, Agelenopsis actuosa, Agelenopsis kastoni","1","1","1","6","0","0","98.86","440","0","441","33.56","0"
"AGAW3","05E","BAC","TAX:392874","Proteobacteria","Alphaproteobacteria","Rickettsiales","Rickettsiaceae",NA,"Wolbachia","Wolbachia endosymbiont of Hymenoptera","1","1","1","1","0","0","98.89","450","0","450","40.44","0"
"AGAW3","05E","BAC","TAX:375847","Proteobacteria","Alphaproteobacteria","Rickettsiales","Rickettsiaceae",NA,"Wolbachia","Wolbachia sp. Hymenoptera_ASGLE1705-10","1","1","1","1","0","0","98.39","436","0","437","40.96","0"
"AGAW3","05E","CHO","no match","no match","no match","no match","no match","no match","no match","no match","no match","no match","no match","no match","no match","no match","no match","no match","no match","no match","no match","no match"
"AGAW3","05E","INS","BOLD:AAH7158","Arthropoda","Insecta","Diptera","Sarcophagidae","Sarcophaginae","Ravinia","Ravinia stimulans","20062","16648","16648","1","17.02","0","98.95","418.84","0","422.36","33.37","0"
"AGAW3","05E","INS","BOLD:ACF5939","Arthropoda","Insecta","Diptera","Tachinidae","Exoristinae","Winthemia","Winthemia sinuata, Winthemia illinoensis","17717","15225","15225","2","14.07","0","99.18","428.18","0","429.93","29.56","0"
"AGAW3","05E","INS","BOLD:AAA3453","Arthropoda","Insecta","Diptera","Anthomyiidae","Anthomyiinae","Delia","","4263","3835","3835","3","10.04","0","99.29","430.83","0","433.21","34.68","0"
"AGAW3","05E","INS","BOLD:AAF9187","Arthropoda","Insecta","Coleoptera","Curculionidae","Entiminae","Phyllobius","Phyllobius oblongus","3352","2622","2622","4","21.78","0","99.44","436.39","0","438.97","38.86","0"
"AGAW3","05E","INS","BOLD:AAG2511","Arthropoda","Insecta","Diptera","Anthomyiidae","Anthomyiinae","Delia","Delia platura","3200","2805","2805","5","12.34","0","98.89","444.74","0","447.16","33.17","0"
"AGAW3","05E","INS","BOLD:AAB2384","Arthropoda","Insecta","Diptera","Syrphidae","Syrphinae","Eupeodes","Eupeodes duseki, Eupeodes volucris, Eupeodes nielseni, Eupeodes americanus, Eupeodes flukei, Eupeodes vockerothi, Eupeodes lucasi, Eupeodes montivagus, Eupeodes rufipunctatus, Eupeodes rojasi, Eupeodes goeldlini, Eupeodes luniger, Eupeodes punctifer, Eupeodes bucculatus, Eupeodes perplexus, Eupeodes abiskoensis, Eupeodes corollae, Eupeodes pomus, Eupeodes lundbecki, Eupeodes nitens, Eupeodes sp. BOLDAAB2384, Eupeodes curtus, Eupeodes latifasciatus","3095","2943","2943","6","4.91","0","98.84","400.13","0","402.56","30.72","0"
"AGAW3","05E","INS","BOLD:AAG9055","Arthropoda","Insecta","Hymenoptera","Vespidae","Vespinae","Vespula","Vespula germanica","2857","2327","2327","7","18.55","0","99.49","434.6","0","435.82","30.63","0"
"AGAW3","05E","INS","BOLD:AAO3939","Arthropoda","Insecta","Diptera","Limoniidae","Limoniinae","Dicranomyia","","2857","2341","2341","7","18.06","0","99.15","437.6","0","439.19","36.24","0"
"AGAW3","05E","INS","BOLD:AAC9132","Arthropoda","Insecta","Diptera","Culicidae","Culicinae","Culiseta","Culiseta inornata","789","781","781","9","1.01","0","98.13","404.57","0","407.04","32.93","0"
"AGAW3","05E","INS","BOLD:ACG5862","Arthropoda","Insecta","Diptera","Agromyzidae","","","","698","632","632","10","9.46","0","98.64","439.33","0","441.13","34.08","0"
"AGAW3","05E","INS","BOLD:AAG6588","Arthropoda","Insecta","Coleoptera","Chrysomelidae","Cassidinae","Deloyala","Deloyala guttata","678","639","639","11","5.75","0","98.72","425.96","0","427.17","31.15","0"
"AGAW3","05E","INS","BOLD:AAC1312","Arthropoda","Insecta","Diptera","Syrphidae","Syrphinae","Toxomerus","Toxomerus geminatus, Toxomerus 75-5, Toxomerus rombicus, Toxomerus sp. 75-5","519","437","437","12","15.8","0","98.91","435.58","0","436.98","29.92","0"
"AGAW3","05E","INS","BOLD:ADD1920","Arthropoda","Insecta","Hymenoptera","Ichneumonidae","Diplazontinae","Enizemum","Enizemum cf. ornatum","375","267","267","13","28.8","0","99.6","446.75","0","447.42","26.91","0"
"AGAW3","05E","INS","BOLD:AAU8613","Arthropoda","Insecta","Hymenoptera","Ichneumonidae","Diplazontinae","Sussaba","Sussaba dorsalis","342","272","272","14","20.47","0","99.54","446.63","0","447.44","30.25","0"
"AGAW3","05E","INS","BOLD:ACF0198","Arthropoda","Insecta","Diptera","Tachinidae","Exoristinae","Winthemia","Winthemia quadripustulata, Winthemia venusta, Winthemia sinuata, Winthemia occidentis, Winthemia erythura, Winthemia illinoensis","339","339","339","15","0","0","98.71","400.56","0","406.51","29.35","0"
"AGAW3","05E","INS","BOLD:AAB7030","Arthropoda","Insecta","Diptera","Chironomidae","Chironominae","Chironomus","Chironomus sp. BOLDAAP2996, Chironomus quinnitukqut, Chironomus cf. decorus 2g, Chironomus decorus group sp. 2 IP2013, Chironomus cf. decorus, Chironomus decorus gr.","295","294","294","16","0.34","0","98.3","387.19","0","395.24","33.84","0"
"AGAW3","05E","INS","BOLD:AAM9013","Arthropoda","Insecta","Diptera","Keroplatidae","Macrocerinae","Macrocera","Macrocera pusilla","285","268","268","17","5.96","0","99.18","432.33","0","433.49","30.25","0"
"AGAW3","05E","INS","BOLD:AAF9771","Arthropoda","Insecta","Diptera","Hybotidae","Tachydromiinae","Platypalpus","Platypalpus longiseta, Platypalpus trivialis, Platypalpus sp. 5, Platypalpus sp. 17, Platypalpus pallidiventris","279","225","225","18","19.35","0","98.91","444.06","0","444.48","35.03","0"
"AGAW3","05E","INS","BOLD:ACL8669","Arthropoda","Insecta","Lepidoptera","Psychidae","Psychinae","Psyche","Psyche casta","209","209","209","19","0","0","99.01","399.33","0","402.71","37.27","0"
"AGAW3","05E","INS","BOLD:AAA5565","Arthropoda","Insecta","Hemiptera","Aphididae","Aphidinae","Aphis","Aphis fabae, Aphis newtoni, Aphis solanella, Aphis ilicis, Aphis hederae, Aphis viburni, Aphis lambersi, Aphis fukii","199","174","174","20","12.56","0","99.44","431.68","0","434.43","26.79","0"
"AGAW3","05E","INS","BOLD:AAN5355","Arthropoda","Insecta","Diptera","Chironomidae","Orthocladiinae","Smittia","Smittia cf. stercoraria, Smittia stercoraria","189","169","169","21","10.58","0","99.34","397.59","0","441.82","36.25","0"
"AGAW3","05E","INS","BOLD:AAQ0265","Arthropoda","Insecta","Diptera","Hybotidae","Tachydromiinae","Platypalpus","Platypalpus pulicarius","189","142","142","21","24.87","0","98.39","446.99","0","447.72","35.45","0"
"AGAW3","05E","INS","BOLD:ACK7296","Arthropoda","Insecta","Diptera","Cecidomyiidae","","","","184","164","164","23","10.87","0","99.55","426.86","0","428.21","24.19","0"
"AGAW3","05E","INS","BOLD:AAF9014","Arthropoda","Insecta","Diptera","Limoniidae","Chioneinae","Symplecta","Symplecta cana, Symplecta sp. BOLDAAF9014","172","165","165","24","4.07","0","98.56","429.49","0","435.65","33.99","0"
"AGAW3","05E","INS","BOLD:AAU6967","Arthropoda","Insecta","Coleoptera","Chrysomelidae","Galerucinae","Psylliodes","Psylliodes affinis","166","134","134","25","19.28","0","99.55","434.61","0","444.47","34.95","0"
"AGAW3","05E","INS","BOLD:AAC4403","Arthropoda","Insecta","Diptera","Chironomidae","Chironominae","Micropsectra","Micropsectra polita","148","146","146","26","1.35","0","98.81","415.73","0","417.44","34.65","0"
"AGAW3","05E","INS","BOLD:AAD8086","Arthropoda","Insecta","Lepidoptera","Elachistidae","Elachistinae","Elachista","Elachista griseicornis, Elachista acenteta","129","122","122","27","5.43","0","99.28","430.67","0","433.41","30.79","0"
"AGAW3","05E","INS","BOLD:AAP2528","Arthropoda","Insecta","Diptera","Keroplatidae","Keroplatinae","Orfelia","Orfelia nemoralis","97","95","95","28","2.06","0","99.18","413.72","0","415.43","33.21","0"
"AGAW3","05E","INS","BOLD:ACF6291","Arthropoda","Insecta","Coleoptera","Curculionidae","","","","94","92","92","29","2.13","0","98.92","434.13","0","435.77","33","0"
"AGAW3","05E","INS","BOLD:AAG6934","Arthropoda","Insecta","Diptera","Hybotidae","","","","88","85","85","30","3.41","0","99.28","426.57","0","428.17","38.63","0"
"AGAW3","05E","INS","BOLD:AAN5249","Arthropoda","Insecta","Diptera","Cecidomyiidae","","","","88","87","87","30","1.14","0","98.68","424.13","0","425.33","33.28","0"
"AGAW3","05E","INS","BOLD:AAC2312","Arthropoda","Insecta","Hymenoptera","Apidae","Xylocopinae","Xylocopa","Xylocopa virginica","88","74","74","30","15.91","0","99.05","438.61","0","439.49","22.59","0"
"AGAW3","05E","INS","BOLD:AAH3664","Arthropoda","Insecta","Diptera","Cecidomyiidae","","","CecidInt3 sp. BOLD:AAH3664","83","78","78","33","6.02","0","99.3","423.61","0","427.27","26.93","0"
"AGAW3","05E","INS","BOLD:AAN5342","Arthropoda","Insecta","Diptera","Chironomidae","Orthocladiinae","Heleniella","","83","81","81","33","2.41","0","98.7","427.39","0","434.23","35.18","0"
"AGAW3","05E","INS","BOLD:ACK3431","Arthropoda","Insecta","Diptera","Cecidomyiidae","","","","80","76","76","35","5","0","99.19","422.55","0","424.45","32.11","0"
"AGAW3","05E","INS","BOLD:AAU8501","Arthropoda","Insecta","Hymenoptera","Ichneumonidae","Ctenopelmatinae","Perilissus","Perilissus albitarsis, Perilissus rufoniger","75","73","73","36","2.67","0","99.16","445.08","0","446.92","33.01","0"
"AGAW3","05E","INS","BOLD:AAG6756","Arthropoda","Insecta","Diptera","Sarcophagidae","Sarcophaginae","Boettcheria","Boettcheria cimbicis","70","62","62","37","11.43","0","99.3","437.67","0","439.23","34.69","0"
"AGAW3","05E","INS","BOLD:AAB1098","Arthropoda","Insecta","Diptera","Culicidae","Culicinae","Ochlerotatus, Aedes","Aedes excrucians, Aedes cantans, Aedes stimulans, Aedes euedes, Aedes riparius, Aedes annulipes, Ochlerotatus cantans, Aedes fitchii, Aedes sp., Ochlerotatus annulipes, Ochlerotatus excrucians","51","50","50","38","1.96","0","98.73","432.29","0","433.55","35.36","0"
"AGAW3","05E","INS","BOLD:AAH3968","Arthropoda","Insecta","Diptera","Sciaridae","","Cratyna","Cratyna cryptospina","45","43","43","39","4.44","0","99.24","443.29","0","445.33","27.45","0"
"AGAW3","05E","INS","BOLD:AAG5192","Arthropoda","Insecta","Coleoptera","Curculionidae","Entiminae","Barypeithes","Barypeithes pellucidus","43","41","41","40","4.65","0","98.99","436.3","0","437.28","39.22","0"
"AGAW3","05E","INS","BOLD:ADH8378","Arthropoda","Insecta","Hymenoptera","Ichneumonidae","","","","37","32","32","41","13.51","0","99.52","386.08","0","432.59","26.65","0"
"AGAW3","05E","INS","BOLD:AAR9709","Arthropoda","Insecta","Lepidoptera","Gelechiidae","Gelechiinae","Chrysoesthia","Chrysoesthia lingulacella, Chrysoesthia drurella","33","33","33","42","0","0","99.09","433.21","0","434.88","31.79","0"
"AGAW3","05E","INS","BOLD:AAL7819","Arthropoda","Insecta","Diptera","Psychodidae","Psychodinae","Psychoda","Psychoda sp., Psychoda sp. BOLD:AAL7819","27","27","27","43","0","0","99.26","414.48","0","415.7","32.86","0"
"AGAW3","05E","INS","BOLD:AAN5330","Arthropoda","Insecta","Diptera","Chironomidae","Orthocladiinae","Corynoneura","Corynoneura scutellata","20","20","20","44","0","0","98.89","408.3","0","415.95","33.87","0"
"AGAW3","05E","INS","BOLD:AAP2527","Arthropoda","Insecta","Diptera","Mycetophilidae","Mycetophilinae","Brevicornu","Brevicornu fissicauda","20","18","18","44","10","0","99.46","447.5","0","448.7","36.83","0"
"AGAW3","05E","INS","BOLD:AAN6433","Arthropoda","Insecta","Diptera","Sciaridae","","Corynoptera","Corynoptera deserta","16","16","16","46","0","0","98.87","438.44","0","439.63","30.5","0"
"AGAW3","05E","INS","BOLD:AAE3911","Arthropoda","Insecta","Trichoptera","Hydroptilidae","Hydroptilinae","Oxyethira","Oxyethira rivicola","16","15","15","46","6.25","0","98.42","443.81","0","445.06","36.11","0"
"AGAW3","05E","INS","BOLD:AAA4222","Arthropoda","Insecta","Lepidoptera","Erebidae","Hypeninae","Hypena","Hypena scabra","15","15","15","48","0","0","99.24","403.93","0","405.2","32.56","0"
"AGAW3","05E","INS","BOLD:AAB1850","Arthropoda","Insecta","Hemiptera","Aphrophoridae","Aphrophorinae","Philaenus","Philaenus tesselatus, Philaenus spumarius, Philaenus maghresignus","12","12","12","49","0","0","99.24","442.58","0","443.83","33.05","0"
"AGAW3","05E","INS","BOLD:AAN5205","Arthropoda","Insecta","Diptera","Cecidomyiidae","","","CecidInt42 sp. BOLD:AAN5205","11","11","11","50","0","0","99.51","449.64","0","450","31.66","0"
"AGAW3","05E","INS","BOLD:ACK3142","Arthropoda","Insecta","Diptera","Cecidomyiidae","","","","11","11","11","50","0","0","98.51","425.18","0","426.82","28.08","0"
"AGAW3","05E","INS","BOLD:AAI8935","Arthropoda","Insecta","Coleoptera","Latridiidae","Corticariinae","Cortinicara","Cortinicara gibbosa","11","11","11","50","0","0","98.87","424","0","424.73","33.71","0"
"AGAW3","05E","INS","BOLD:ABV1225","Arthropoda","Insecta","Diptera","Chironomidae","Chironominae","Tanytarsus","","11","11","11","50","0","0","98.2","443.73","0","445","37.57","0"
"AGAW3","05E","INS","BOLD:AAN5267","Arthropoda","Insecta","Diptera","Cecidomyiidae","","","","10","9","9","54","10","0","99.42","449.6","0","450","30.71","0"
"AGAW3","05E","INS","BOLD:AAA7374","Arthropoda","Insecta","Diptera","Syrphidae","Syrphinae","Sphaerophoria","Sphaerophoria bankowskae, Sphaerophoria infuscata, Sphaerophoria nr. philanthus, Sphaerophoria batava, Sphaerophoria sp., Sphaerophoria rueppellii, Sphaerophoria interrupta-group, Sphaerophoria sp. MA-1, Sphaerophoria cleoae, Sphaerophoria asymmetrica, Sphaerophoria fatarum, Sphaerophoria bengalensis, Sphaerophoria bifurcata, Sphaerophoria taeniata, Sphaerophoria interrupta, Sphaerophoria contigua, Sphaerophoria kaa, Sphaerophoria laurae, Sphaerophoria chongjini, Sphaerophoria boreoalpina, Sphaerophoria sp. BOLDAAA7374, Sphaerophoria macrogaster, Sphaerophoria sulphuripes, Sphaerophoria brevipilosa, Sphaerophoria philanthus, Sphaerophoria pyrrhina, Sphaerophoria abbreviata, Sphaerophoria scripta, Sphaerophoria potentillae, Sphaerophoria longipilosa, Sphaerophoria virgata","10","10","10","54","0","0","99.16","421.4","0","423.3","31.1","0"
"AGAW3","05E","INS","BOLD:ACW5117","Arthropoda","Insecta","Diptera","Chironomidae","Orthocladiinae","Smittia","Smittia leucopogon","9","9","9","56","0","0","99.43","416.44","0","417.89","31.56","0"
"AGAW3","05E","INS","BOLD:AAB2644","Arthropoda","Insecta","Diptera","Chironomidae","Orthocladiinae","Orthocladius","Orthocladius lapponicus, Orthocladius dorenus, Orthocladius mallochi","7","7","7","57","0","0","98.49","426.71","0","428.86","32.54","0"
"AGAW3","05E","INS","BOLD:AAQ2892","Arthropoda","Insecta","Hymenoptera","Braconidae","Rhyssalinae","Proacrisis","","7","5","5","57","28.57","0","99.78","450.29","0","450","28","0"
"AGAW3","05E","INS","BOLD:AAN5358","Arthropoda","Insecta","Diptera","Chironomidae","Orthocladiinae","Smittia","Smittia stercoraria, Smittia sp. 22ES","7","7","7","57","0","0","99.59","449","0","450","37.65","0"
"AGAW3","05E","INS","BOLD:AAM9252","Arthropoda","Insecta","Diptera","Sciaridae","","Bradysia, Leptosciarella","Bradysia vagans, Leptosciarella zanti","7","7","7","57","0","0","99.53","427.71","0","429","38.77","0"
"AGAW3","05E","INS","BOLD:AAE3675","Arthropoda","Insecta","Diptera","Chironomidae","Chironominae","Paratanytarsus","Paratanytarsus sp. TE03","6","6","6","61","0","0","99.18","449.5","0","450","36.41","0"
"AGAW3","05E","INS","BOLD:ABV9089","Arthropoda","Insecta","Diptera","Cecidomyiidae","","","","6","6","6","61","0","0","98.66","448.33","0","450","27.15","0"
"AGAW3","05E","INS","BOLD:AAG2868","Arthropoda","Insecta","Hemiptera","Cicadellidae","Typhlocybinae","Empoasca","Empoasca sp., Empoasca fabae","6","6","6","61","0","0","99.74","448.83","0","450","32.15","0"
"AGAW3","05E","INS","BOLD:ADH4889","Arthropoda","Insecta","Diptera","","","","","6","5","5","61","16.67","0","99.08","436.5","0","437.5","35.61","0"
"AGAW3","05E","INS","BOLD:AAG6743","Arthropoda","Insecta","Diptera","Sarcophagidae","Sarcophaginae","Sarcophaga","Sarcophaga sp. 1 MG, Sarcophaga subvicina","6","6","6","61","0","0","99.16","422.83","0","424.33","34.87","0"
"AGAW3","05E","INS","BOLD:AAN5271","Arthropoda","Insecta","Diptera","Cecidomyiidae","","","CecidInt9 sp. BOLD:AAN5271","6","6","6","61","0","0","99.15","449.5","0","450","26.19","0"
"AGAW3","05E","INS","BOLD:AAG6441","Arthropoda","Insecta","Diptera","Ceratopogonidae","Ceratopogoninae","Culicoides","Culicoides stilobezzioides","5","5","5","67","0","0","98.4","448.8","0","450","34.84","0"
"AGAW3","05E","INS","BOLD:AAN5356","Arthropoda","Insecta","Diptera","Chironomidae","Orthocladiinae","Smittia","","5","5","5","67","0","0","99.06","447.6","0","450","35.33","0"
"AGAW3","05E","INS","BOLD:AAN8770","Arthropoda","Insecta","Diptera","Psychodidae","Psychodinae","Psychoda","Psychoda trinodulosa","5","5","5","67","0","0","99.2","429","0","430.8","33.58","0"
"AGAW3","05E","INS","BOLD:AAA7661","Arthropoda","Insecta","Diptera","Culicidae","Culicinae","Culiseta, Culex","Culiseta melanura, Culex articularis, Culex restuans, Culex acharistus","5","5","5","67","0","0","98.47","412","0","415.8","32.68","0"
"AGAW3","05E","INS","BOLD:ABU5520","Arthropoda","Insecta","Diptera","Sciaridae","","Corynoptera","","4","4","4","71","0","0","98.83","411","0","412.5","34.9","0"
"AGAW3","05E","INS","BOLD:AAG5242","Arthropoda","Insecta","Coleoptera","Curculionidae","Entiminae","Barypeithes","Barypeithes pellucidus","4","4","4","71","0","0","98.01","441.5","0","442","39.74","0"
"AGAW3","05E","INS","BOLD:AAG4653","Arthropoda","Insecta","Diptera","Syrphidae","Syrphinae","Heringia","Heringia latitarsis, Heringia pubescens, Heringia jakutorum, Heringia rita, Heringia fulvimanus, Heringia sp. BOLDAAG4653, Heringia nr. pubescens, Heringia coxalis, Heringia larusi, Heringia vitripennis","4","4","4","71","0","0","99.61","447.5","0","450","31.56","0"
"AGAW3","05E","INS","BOLD:AAB4657","Arthropoda","Insecta","Diptera","Chironomidae","Chironominae","Chironomus","Chironomus maturus","3","3","3","74","0","0","98.6","426.33","0","427.33","37.34","0"
"AGAW3","05E","INS","BOLD:ABV9276","Arthropoda","Insecta","Diptera","Chironomidae","Chironominae","Tanytarsus","","3","3","3","74","0","0","97.97","418.67","0","420","36.89","0"
"AGAW3","05E","INS","BOLD:AAD1485","Arthropoda","Insecta","Diptera","Chironomidae","Chironominae","Paratanytarsus","Paratanytarsus grimmii, Paratanytarsus sp.","3","3","3","74","0","0","99.47","444","0","446","33.78","0"
"AGAW3","05E","INS","BOLD:AAA2815","Arthropoda","Insecta","Lepidoptera","Noctuidae","Noctuinae","Mythimna","Mythimna yukonensis, Mythimna oxygala","3","3","3","74","0","0","98.95","403","0","403.67","33.46","0"
"AGAW3","05E","INS","BOLD:AAN6437","Arthropoda","Insecta","Diptera","Sciaridae","","Corynoptera","Corynoptera saccata","3","3","3","74","0","0","98.85","431.33","0","432","39.39","0"
"AGAW3","05E","INS","BOLD:AAH3914","Arthropoda","Insecta","Diptera","Sciaridae","","Sciara","Sciara hamatilis, Sciara humeralis","3","3","3","74","0","0","99.26","449.67","0","450","30.67","0"
"AGAW3","05E","INS","BOLD:AAA3181","Arthropoda","Insecta","Diptera","Muscidae","Muscinae","Stomoxys","Stomoxys calcitrans","2","2","2","80","0","0","98.34","390.5","0","394.5","32.45","0"
"AGAW3","05E","INS","BOLD:AAA3933","Arthropoda","Insecta","Lepidoptera","Erebidae","Herminiinae","Palthis","Palthis angulalis","2","2","2","80","0","0","99.33","448","0","450","32.11","0"
"AGAW3","05E","INS","BOLD:ACT0257","","","","","","","","2","2","2","80","0","0","100","449.5","0","450","37.89","0"
"AGAW3","05E","INS","BOLD:AAA4121","Arthropoda","Insecta","Diptera","Simuliidae","Simuliinae","Simulium","Simulium vittatum complex, Simulium vittatum, Simulium sp. BOLDAAA4121, Simulium argus, Simulium vittatum s.str.","2","2","2","80","0","0","98.15","381.5","0","383.5","33.25","0"
"AGAW3","05E","INS","BOLD:AAG5453","Arthropoda","Insecta","Diptera","Chironomidae","Chironominae","Chironomus","Chironomus bifurcatus","2","2","2","80","0","0","98.37","406.5","0","408.5","34.88","0"
"AGAW3","05E","INS","BOLD:AAA2482","Arthropoda","Insecta","Lepidoptera","Noctuidae","Noctuinae","Mythimna","Mythimna unipuncta","2","2","2","80","0","0","99.78","411","0","416","30.48","0"
"AGAW3","05E","INS","BOLD:ABU5525","Arthropoda","Insecta","Diptera","Chironomidae","Orthocladiinae","Limnophyes","Limnophyes sp. 14ES","2","2","2","80","0","0","97.54","366","0","367","36.92","0"
"AGAW3","05E","INS","BOLD:ACF2924","","","","","","","","2","2","2","80","0","0","98.21","448","0","450","25.89","0"
"AGAW3","05E","INS","BOLD:AAC2498","Arthropoda","Insecta","Diptera","Muscidae","Phaoniinae","Helina","Helina evecta, Helina aldrichi","2","2","2","80","0","0","97.98","394.5","0","401","29.69","0"
"AGAW3","05E","INS","BOLD:AAP4829","Arthropoda","Insecta","Diptera","Tachinidae","Exoristinae","Chrysoexorista","Chrysoexorista sp. 1","2","2","2","80","0","0","98.72","425","0","425","30.46","0"
"AGAW3","05E","INS","BOLD:AAN5268","Arthropoda","Insecta","Diptera","Cecidomyiidae","","","","2","2","2","80","0","0","98.67","436","0","436","29.93","0"
"AGAW3","05E","INS","BOLD:AAC5848","Arthropoda","Insecta","Lepidoptera","Psychidae","Psychinae","Psyche","Psyche casta","2","2","2","80","0","0","98.16","382","0","383","37.99","0"
"AGAW3","05E","INS","BOLD:AAD2548","Arthropoda","Insecta","Hemiptera","Aphididae","Aphidinae","Acyrthosiphon","Acyrthosiphon caraganae, Acyrthosiphon sp. 1, Acyrthosiphon kondoi, Acyrthosiphon churchillense","2","2","2","80","0","0","99.78","450","0","450","25.89","0"
"AGAW3","05E","INS","BOLD:AAF9306","Arthropoda","Insecta","Diptera","Psychodidae","Psychodinae","Psychoda","Psychoda alternata","2","2","2","80","0","0","97.86","404.5","0","408","30.53","0"
"AGAW3","05E","INS","BOLD:AAN5501","Arthropoda","Insecta","Diptera","Hybotidae","","","","2","2","2","80","0","0","99.33","449.5","0","450","34.78","0"
"AGAW3","05E","INS","BOLD:AAH1867","Arthropoda","Insecta","Hymenoptera","Ichneumonidae","Ctenopelmatinae","","","2","2","2","80","0","0","98.01","378.5","0","379.5","24.51","0"
"AGAW3","05E","INS","BOLD:AAC9062","Arthropoda","Insecta","Diptera","Culicidae","Culicinae","Ochlerotatus, Aedes","Aedes fitchii, Ochlerotatus fitchii, Aedes grossbecki, Aedes ventrovittis","2","2","2","80","0","0","98.13","426.5","0","427","32.77","0"
"AGAW3","05E","INS","BOLD:AAB5577","Arthropoda","Insecta","Diptera","Syrphidae","Syrphinae","Syrphus","Syrphus sexmaculata, Syrphus sexmaculatus, Syrphus opinator, Syrphus rectus, Syrphus vitripennis","2","2","2","80","0","0","98.63","387","0","390","30.94","0"
"AGAW3","05E","INS","BOLD:ACV4333","Arthropoda","Insecta","Diptera","Chironomidae","Chironominae","Tanytarsus","","2","2","2","80","0","0","97.17","370.5","0","372.5","35.7","0"
"AGAW3","05E","INS","BOLD:AAA4782","Arthropoda","Insecta","Hymenoptera","Braconidae","Microgastrinae","Protapanteles, Glyptapanteles","","2","2","2","80","0","0","98.94","400","0","400.5","24.79","0"
"AGAW3","05E","INS","BOLD:ADH9310","Arthropoda","Insecta","Diptera","Anthomyiidae","Anthomyiinae","Anthomyia","Anthomyia procellaris, Anthomyia pluvialis","2","2","2","80","0","0","99.61","407","0","407","32.71","0"
"AGAW3","05E","INS","BOLD:ABV1243","Arthropoda","Insecta","Diptera","Sarcophagidae","Paramacronychiinae, Miltogramminae","Miltogramminae_incertae_sedis, Brachicoma","Miltogramminae_incertae_sedis sp. 1 MG, Brachicoma sarcophagina","1","1","1","101","0","0","99.33","450","0","450","34.89","0"
"AGAW3","05E","INS","BOLD:AAH4135","Arthropoda","Insecta","Diptera","Chloropidae","Chloropinae","Thaumatomyia","Thaumatomyia glabra, Thaumatomyia grata, Thaumatomyia sp. 1 TAW","1","1","1","101","0","0","99.33","449","0","450","31.78","0"
"AGAW3","05E","INS","BOLD:AAG1717","Arthropoda","Insecta","Diptera","Muscidae","Phaoniinae","Helina","Helina troene","1","1","1","101","0","0","100","423","0","424","30.66","0"
"AGAW3","05E","INS","BOLD:ACK2748","Arthropoda","Insecta","Diptera","Cecidomyiidae","","","","1","1","1","101","0","0","97.11","450","0","450","27.78","0"
"AGAW3","05E","INS","BOLD:AAG5507","Arthropoda","Insecta","Diptera","Chironomidae","Chironominae","Chironomus","Chironomus atrella","1","1","1","101","0","0","99.26","408","0","409","35.94","0"
"AGAW3","05E","INS","BOLD:AAG9668","Arthropoda","Insecta","Diptera","Dolichopodidae","Diaphorinae","Chrysotus","Chrysotus sp., Chrysotus sp. nr. obliquus","1","1","1","101","0","0","99.55","448","0","450","32.22","0"
"AGAW3","05E","INS","BOLD:AAG9713","Arthropoda","Insecta","Diptera","Dolichopodidae","Sympycninae","Sympycnus","Sympycnus lineatus","1","1","1","101","0","0","99.56","450","0","450","35.33","0"
"AGAW3","05E","INS","BOLD:AAI4303","Arthropoda","Insecta","Diptera","Chironomidae","Chironominae","Chironomus","Chironomus melanescens","1","1","1","101","0","0","97.75","445","0","450","34.44","0"
"AGAW3","05E","INS","BOLD:AAP8998","","","","","","","","1","1","1","101","0","0","97.54","448","0","450","33.56","0"
"AGAW3","05E","INS","BOLD:AAK1128","Arthropoda","Insecta","Hymenoptera","Ichneumonidae","Diplazontinae","Sussaba","Sussaba cultriformis, Sussaba dorsalis","1","1","1","101","0","0","97.51","361","0","363","28.1","0"
"AGAW3","05E","INS","BOLD:ACE8783","Arthropoda","Insecta","Diptera","Chloropidae","Oscinellinae","Siphonella","Siphonella oscinina","1","1","1","101","0","0","97.25","436","0","437","34.78","0"
"AGAW3","05E","INS","BOLD:AAA3817","Arthropoda","Insecta","Lepidoptera","Geometridae","Larentiinae","Xanthorhoe","Xanthorhoe ferrugata","1","1","1","101","0","0","97.42","426","0","426","33.1","0"
"AGAW3","05E","INS","BOLD:AAG2462","Arthropoda","Insecta","Diptera","Anthomyiidae","Anthomyiinae","Botanophila","Botanophila fugax","1","1","1","101","0","0","98.78","411","0","413","31.23","0"
"AGAW3","05E","INS","BOLD:AAG6433","Arthropoda","Insecta","Diptera","Ceratopogonidae","Forcipomyiinae","Forcipomyia","Forcipomyia sp. BOLD:AAG6433, Forcipomyia sp. 17","1","1","1","101","0","0","99.31","433","0","433","32.79","0"
"AGAW3","05E","INS","BOLD:AAA5448","Arthropoda","Insecta","Lepidoptera","Geometridae","Ennominae","Eutrapela","Eutrapela clemataria","1","1","1","101","0","0","99.56","450","0","450","30","0"
"AGAW3","05E","INS","BOLD:AAH3661","Arthropoda","Insecta","Diptera","Cecidomyiidae","","","","1","1","1","101","0","0","99.78","450","0","450","26.22","0"
"AGAW3","05E","INS","BOLD:AAG7312","Arthropoda","Insecta","Diptera","Sphaeroceridae","Limosininae","Spelobia","Spelobia clunipes, Spelobia sp. B TAW","1","1","1","101","0","0","99.78","449","0","450","33.78","0"
"AGAW3","05E","INS","BOLD:AAG5457","Arthropoda","Insecta","Diptera","Chironomidae","Chironominae","","","1","1","1","101","0","0","100","450","0","450","34.22","0"
"AGAW3","05E","INS","BOLD:ACK4716","Arthropoda","Insecta","Diptera","Cecidomyiidae","","","","1","1","1","101","0","0","99.55","449","0","450","32","0"
"AGAW3","05E","INS","BOLD:AAK7242","Arthropoda","Insecta","Coleoptera","Melandryidae","Melandryinae","Melandrya","Melandrya striata","1","1","1","101","0","0","97.88","378","0","377","32.1","0"
"AGAW3","05E","INS","BOLD:AAG8493","Arthropoda","Insecta","Diptera","Drosophilidae","Drosophilinae","Scaptomyza","Scaptomyza pallida","1","1","1","101","0","0","97.41","424","0","426","33.8","0"
"AGAW3","05E","INS","BOLD:AAC0903","Arthropoda","Insecta","Diptera","Chironomidae","Chironominae","Chironomus","Chironomus acidophilus","1","1","1","101","0","0","99.11","450","0","450","36.44","0"
"AGAW3","05E","INS","BOLD:AAB9140","Arthropoda","Insecta","Diptera","Calliphoridae","Chrysomyinae","Phormia","Phormia regina","1","1","1","101","0","0","98.66","447","0","450","32.89","0"
"AGAW3","05E","INS","BOLD:AAA9101","Arthropoda","Insecta","Lepidoptera","Pyralidae","Phycitinae","Moodna, Caudellia","Moodna ostrinella, Moodna bisinuella, Caudellia nigrella","1","1","1","101","0","0","98.21","447","0","450","33.56","0"
"AGAW3","05E","INS","BOLD:ABU5546","Arthropoda","Insecta","Diptera","Psychodidae","Psychodinae","Psychoda","","1","1","1","101","0","0","99.26","406","0","409","29.58","0"
"AGAW3","05E","INS","BOLD:ABZ5047","Arthropoda","Insecta","Diptera","Tachinidae","Exoristinae","Winthemia","Winthemia sminthurae","1","1","1","101","0","0","97.14","385","0","392","29.34","0"
"AGAW3","05E","INS","BOLD:AAJ7105","Arthropoda","Insecta","Diptera","Agromyzidae","Agromyzinae","Agromyza","Agromyza frontella","1","1","1","101","0","0","100","399","0","450","36.89","0"
"AGAW3","05E","INS","BOLD:AAA9251","Arthropoda","Insecta","Lepidoptera","Gelechiidae","Gelechiinae","Scrobipalpa","Scrobipalpa atriplicella, Scrobipalpa nr. atriplicella sp. 1","1","1","1","101","0","0","97.48","357","0","359","33.7","0"
"AGAW3","05E","INS","BOLD:AAL2750","Arthropoda","Insecta","Diptera","Muscidae","Phaoniinae","Phaonia","Phaonia tuguriorum","1","1","1","101","0","0","99.55","447","0","450","33.11","0"
"AGAW3","05E","INS","BOLD:ABA3960","Arthropoda","Insecta","Coleoptera","Chrysomelidae","Eumolpinae","Paria","Paria fragariae","1","1","1","101","0","0","97.1","448","0","450","36.67","0"
"AGAW3","05E","INS","BOLD:ACF2821","Arthropoda","Insecta","Diptera","Mycetophilidae","Mycetophilinae","Mycetophila","Mycetophila fungorum","1","1","1","101","0","0","99.55","449","0","450","35.78","0"
"AGAW3","05E","INS","BOLD:AAH0256","Arthropoda","Insecta","Coleoptera","Latridiidae","Corticariinae","Corticaria, Corticarina","","1","1","1","101","0","0","97.12","347","0","351","31.05","0"
"AGAW3","05E","INS","BOLD:ABW2407","Arthropoda","Insecta","Diptera","Anthomyiidae","Anthomyiinae","Hylemya","","1","1","1","101","0","0","97.14","419","0","434","32.49","0"
"AGAW3","05E","INS","BOLD:AAB1787","Arthropoda","Insecta","Hemiptera","Aphididae","Aphidinae","Rhopalosiphum, Acyrthosiphon","Acyrthosiphon pisum, Acyrthosiphon pisum 5A, Rhopalosiphum maidis","1","1","1","101","0","0","97.77","449","0","450","26","0"
"AGAW3","05E","INS","BOLD:AAE0844","Arthropoda","Insecta","Hemiptera","Pentatomidae","Pentatominae","Euschistus","Euschistus variolarius","1","1","1","101","0","0","98.72","390","0","390","35.64","0"
"AGAW3","05E","INS","BOLD:AAP9022","Arthropoda","Insecta","Diptera","Cecidomyiidae","","","","1","1","1","101","0","0","100","450","0","450","27.56","0"
"AGAW3","05E","INS","BOLD:AAH1888","Arthropoda","Insecta","Hymenoptera","Ichneumonidae","Ctenopelmatinae","","","1","1","1","101","0","0","98.44","450","0","450","29.33","0"
"AGAW3","05E","INS","BOLD:AAA4555","Arthropoda","Insecta","Orthoptera","Acrididae","Melanoplinae","Melanoplus","Melanoplus stupefactus, Melanoplus angustipennis, Melanoplus ponderosus, Melanoplus bruneri, Melanoplus frigidus, Melanoplus scapularis, Melanoplus cinereus, Melanoplus madeleineae, Melanoplus sanguinipes, Melanoplus nigrescens group, Melanoplus tunicae, Melanoplus walshi, Melanoplus femurrubrum, Melanoplus dawsoni, Melanoplus sp., Melanoplus fasciatus, Melanoplus bowditchi, Melanoplus stonei, Melanoplus bivittatus, Melanoplus huroni, Melanoplus thomasi, Melanoplus borealis, Melanoplus packardii","1","1","1","101","0","0","99.33","447","0","450","36.67","0"
"AGAW3","05E","INS","BOLD:ADH9867","Arthropoda","Insecta","Diptera","","","","","1","1","1","101","0","0","99.44","355","0","357","36.97","0"
"AGAW3","05E","INS","BOLD:AAF9041","Arthropoda","Insecta","Diptera","Tipulidae","Tipulinae","Tipula","Tipula oleracea","1","1","1","101","0","0","98.76","404","0","405","34.81","0"
"AGAW3","05E","INS","BOLD:AAG1004","Arthropoda","Insecta","Diptera","Chironomidae","Orthocladiinae","Cricotopus","Cricotopus trifascia, Cricotopus trifascia grp.","1","1","1","101","0","0","98.21","390","0","394","36.29","0"
"AGAW3","05E","INS","BOLD:AAM6037","Arthropoda","Insecta","Diptera","Cecidomyiidae","Cecidomyiinae","Contarinia, Dasineura","","1","1","1","101","0","0","97.53","446","0","450","26.67","0"
"AGAW3","05E","INS","BOLD:ACW7612","Arthropoda","Insecta","Diptera","Chironomidae","","","","1","1","1","101","0","0","99.77","434","0","450","36.44","0"
"AGAW3","05E","INS","BOLD:AAU6630","Arthropoda","Insecta","Diptera","Fanniidae","Fanniinae","Fannia","Fannia armata","1","1","1","101","0","0","99.78","449","0","450","33.56","0"
"AGAW3","05E","INS","BOLD:ACP4736","Arthropoda","Insecta","Diptera","Chironomidae","Orthocladiinae","Smittia","Smittia sp. 8ES","1","1","1","101","0","0","100","443","0","443","33.86","0"
"AGAW3","05E","INS","BOLD:AAA1962","Arthropoda","Insecta","Diptera","Sarcophagidae","Sarcophaginae","Helicobia, sarcJanzen01, Sarcophaga","sarcJanzen01 Janzen14, Helicobia rapax, Sarcophaga rapax","1","1","1","101","0","0","99.78","449","0","450","34.89","0"
"AGAW3","05E","INS","BOLD:ACF7232","Arthropoda","Insecta","Diptera","Chironomidae","Orthocladiinae","Parakiefferiella","","1","1","1","101","0","0","99.22","383","0","450","37.56","0"
"AGAW3","05E","INS","BOLD:ACX5521","Arthropoda","Insecta","Diptera","Chironomidae","Orthocladiinae","","","1","1","1","101","0","0","99.56","450","0","450","34.89","0"
"AGAW3","05E","INS","BOLD:AAE4298","Arthropoda","Insecta","Diptera","Chironomidae","Orthocladiinae","Cricotopus","Cricotopus tremulus","1","1","1","101","0","0","97.67","387","0","388","31.7","0"
"AGAW3","05E","INS","BOLD:AAG1701","Arthropoda","Insecta","Diptera","Muscidae","Coenosiinae","Macrorchis","Macrorchis ausoba","1","1","1","101","0","0","98.44","449","0","450","33.33","0"
"AGAW3","05E","INS","BOLD:AAE7016","Arthropoda","Insecta","Lepidoptera","Gelechiidae","Dichomeridinae","Helcystogramma","Helcystogramma hystricella","1","1","1","101","0","0","98.21","447","0","450","32.22","0"
"AGAW3","05E","INS","BOLD:ABX3225","Arthropoda","Insecta","Coleoptera","Chrysomelidae","Galerucinae","Psylliodes","Psylliodes cucullatus","1","1","1","101","0","0","97.03","438","0","440","35.23","0"
"AGAW3","05E","INS","BOLD:AAA1513","Arthropoda","Insecta","Lepidoptera","Plutellidae","Plutellinae","Plutella, Rhigognostis","Plutella xylostella, Rhigognostis sp.","1","1","1","101","0","0","99.78","450","0","450","32.44","0"
"AGAW3","05E","INS","BOLD:AAA5953","Arthropoda","Insecta","Lepidoptera","Gelechiidae","Gelechiinae","Coleotechnites","Coleotechnites sp., Coleotechnites piceaella, Coleotechnites laricis, Coleotechnites sp. 3, Coleotechnites carbonaria, Coleotechnites granti, Coleotechnites macleodi, Coleotechnites gibsonella, Coleotechnites coniferella, Coleotechnites martini","1","1","1","101","0","0","97.77","449","0","450","34.22","0"
"AGAW3","05E","INS","BOLD:AAD0484","Arthropoda","Insecta","Diptera","Chironomidae","","","","1","1","1","101","0","0","99.78","448","0","450","35.11","0"
"AGAW3","05E","INS","BOLD:AAA8451","Arthropoda","Insecta","Lepidoptera","Hepialidae","","Korscheltellus","Korscheltellus lupulina","1","1","1","101","0","0","98.22","450","0","450","28.22","0"
"AGAW3","05E","NAR","no match","no match","no match","no match","no match","no match","no match","no match","no match","no match","no match","no match","no match","no match","no match","no match","no match","no match","no match","no match"
"AGAW3","05E","NIN","BOLD:AAB0726","Arthropoda","Arachnida","Araneae","Lycosidae","","Trochosa","Trochosa ruricola","939","886","886","1","5.64","0","98.92","434.91","0","437.21","31.16","0"
"AGAW3","05E","NIN","BOLD:AAI4346","Arthropoda","Arachnida","Opiliones","Phalangiidae","","Paroligolophus, Phalangium","Paroligolophus agrestis, Phalangium opilio","406","393","393","2","3.2","0","98.81","424.44","0","431.51","35.85","0"
"AGAW3","05E","NIN","BOLD:AAB7914","Arthropoda","Collembola","Symphypleona","Bourletiellidae","","Bourletiella","","64","58","58","3","9.38","0","99.37","440.7","0","441.98","38.02","0"
"AGAW3","05E","NIN","BOLD:ABW2639","Arthropoda","Arachnida","Trombidiformes","Anystidae","","","","63","59","59","4","6.35","0","99.34","446.6","0","442.19","31.36","0"
"AGAW3","05E","NIN","BOLD:AAD1564","Arthropoda","Arachnida","Araneae","Clubionidae","","Clubiona","Clubiona abboti","37","37","37","5","0","0","98.57","417.7","0","419.3","31.99","0"
"AGAW3","05E","NIN","BOLD:ACC0359","Arthropoda","Collembola","Symphypleona","Bourletiellidae","","Bourletiella","","22","22","22","6","0","0","99.24","442.36","0","445.86","37.56","0"
"AGAW3","05E","NIN","BOLD:AAV0589","Arthropoda","Arachnida","Mesostigmata","Blattisociidae","Platyseiinae","Cheiroseius","","21","18","18","7","14.29","0","99.65","445","0","446.19","27.46","0"
"AGAW3","05E","NIN","BOLD:ACZ1295","Arthropoda","Arachnida","Trombidiformes","Anystidae","","","","15","15","15","8","0","0","98.67","372.93","0","375.53","31.02","0"
"AGAW3","05E","NIN","BOLD:AAA7246","Arthropoda","Collembola","Entomobryomorpha","Entomobryidae","Entomobryinae","Entomobrya","Entomobrya nicoleti, Entomobrya multifasciata, Entomobrya intermedia","14","14","14","9","0","0","98.71","438.43","0","439.5","43.45","0"
"AGAW3","05E","NIN","BOLD:AAH6630","Arthropoda","Collembola","Entomobryomorpha","Entomobryidae","Entomobryinae","Entomobrya","Entomobrya unostrigata","7","7","7","10","0","0","98.52","445.14","0","445.86","39.76","0"
"AGAW3","05E","NIN","BOLD:AAM7961","Arthropoda","Arachnida","Trombidiformes","Anystidae","","","","6","6","6","11","0","0","99.56","454.83","0","450","31.33","0"
"AGAW3","05E","NIN","BOLD:AAN6642","Arthropoda","Arachnida","Mesostigmata","Blattisociidae","Platyseiinae","Cheiroseius","","3","3","3","12","0","0","99.93","424","0","425.33","26.13","0"
"AGAW3","05E","NIN","BOLD:ACK4848","Arthropoda","Arachnida","Trombidiformes","Eupodidae","","","","1","1","1","13","0","0","98.92","371","0","374","33.96","0"
"AGAW3","05N","BAC","no match","no match","no match","no match","no match","no match","no match","no match","no match","no match","no match","no match","no match","no match","no match","no match","no match","no match","no match","no match"
"AGAW3","05N","CHO","no match","no match","no match","no match","no match","no match","no match","no match","no match","no match","no match","no match","no match","no match","no match","no match","no match","no match","no match","no match"
"AGAW3","05N","INS","BOLD:AAA4222","Arthropoda","Insecta","Lepidoptera","Erebidae","Hypeninae","Hypena","Hypena scabra","26597","22725","22725","1","14.56","0","99.24","395.73","0","397.4","32.43","0"
"AGAW3","05N","INS","BOLD:AAF9187","Arthropoda","Insecta","Coleoptera","Curculionidae","Entiminae","Phyllobius","Phyllobius oblongus","23559","16701","16701","2","29.11","0","99.48","436.98","0","439.34","38.89","0"
"AGAW3","05N","INS","BOLD:AAA3933","Arthropoda","Insecta","Lepidoptera","Erebidae","Herminiinae","Palthis","Palthis angulalis","12957","10710","10710","3","17.34","0","98.99","425.21","0","427.36","31.91","0"
"AGAW3","05N","INS","BOLD:ABW7995","Arthropoda","Insecta","Diptera","Dolichopodidae","Dolichopodinae","Gymnopternus","Gymnopternus nigribarbus","3022","2698","2698","4","10.72","0","99.33","407.86","0","420.98","35.92","0"
"AGAW3","05N","INS","BOLD:AAA3654","Arthropoda","Insecta","Lepidoptera","Crambidae","Crambinae","Crambus","Crambus bigelovi, Crambus johnsoni, Crambus praefectellus","2570","2168","2168","5","15.64","0","98.97","435.72","0","437.74","31.58","0"
"AGAW3","05N","INS","BOLD:AAA7661","Arthropoda","Insecta","Diptera","Culicidae","Culicinae","Culiseta, Culex","Culiseta melanura, Culex articularis, Culex restuans, Culex acharistus","1143","1054","1054","6","7.79","0","99.25","433.07","0","434.9","32.84","0"
"AGAW3","05N","INS","BOLD:AAF9014","Arthropoda","Insecta","Diptera","Limoniidae","Chioneinae","Symplecta","Symplecta cana, Symplecta sp. BOLDAAF9014","973","889","889","7","8.63","0","98.58","426.94","0","432.82","34.25","0"
"AGAW3","05N","INS","BOLD:ACA7493","Arthropoda","Insecta","Diptera","Chironomidae","Chironominae","","","945","932","932","8","1.38","0","97.43","379.67","0","389.74","39.48","0"
"AGAW3","05N","INS","BOLD:AAJ7105","Arthropoda","Insecta","Diptera","Agromyzidae","Agromyzinae","Agromyza","Agromyza frontella","832","653","653","9","21.51","0","99.55","397.89","0","444.01","36.5","0"
"AGAW3","05N","INS","BOLD:AAD0484","Arthropoda","Insecta","Diptera","Chironomidae","","","","773","667","667","10","13.71","0","99.11","445.26","0","446.51","34.72","0"
"AGAW3","05N","INS","BOLD:AAG2511","Arthropoda","Insecta","Diptera","Anthomyiidae","Anthomyiinae","Delia","Delia platura","683","640","640","11","6.3","0","99.01","443.93","0","446.47","33.16","0"
"AGAW3","05N","INS","BOLD:ABY4763","Arthropoda","Insecta","Hemiptera","Aphididae","Aphidinae","Uroleucon","Uroleucon taraxaci","641","455","455","12","29.02","0","99.61","442.94","0","444.08","24.98","0"
"AGAW3","05N","INS","BOLD:AAE4298","Arthropoda","Insecta","Diptera","Chironomidae","Orthocladiinae","Cricotopus","Cricotopus tremulus","610","558","558","13","8.52","0","98.96","428.86","0","431.01","31.99","0"
"AGAW3","05N","INS","BOLD:AAH1890","Arthropoda","Insecta","Hymenoptera","Ichneumonidae","Cryptinae","Stibeutes","Stibeutes brevicornis, Stibeutes heinemanni","554","426","426","14","23.1","0","99.6","442.48","0","444.65","29.6","0"
"AGAW3","05N","INS","BOLD:AAN5355","Arthropoda","Insecta","Diptera","Chironomidae","Orthocladiinae","Smittia","Smittia cf. stercoraria, Smittia stercoraria","536","535","535","15","0.19","0","98.95","430.31","0","433.74","36.16","0"
"AGAW3","05N","INS","BOLD:ACK4716","Arthropoda","Insecta","Diptera","Cecidomyiidae","","","","479","452","452","16","5.64","0","99.01","427.75","0","428.76","31.71","0"
"AGAW3","05N","INS","BOLD:AAH3664","Arthropoda","Insecta","Diptera","Cecidomyiidae","","","CecidInt3 sp. BOLD:AAH3664","352","340","340","17","3.41","0","99.01","425.13","0","427.52","27.1","0"
"AGAW3","05N","INS","BOLD:ACB9392","Arthropoda","Insecta","Diptera","Chironomidae","Chironominae","","","268","267","267","18","0.37","0","98.82","419.9","0","422.2","34.78","0"
"AGAW3","05N","INS","BOLD:AAG5192","Arthropoda","Insecta","Coleoptera","Curculionidae","Entiminae","Barypeithes","Barypeithes pellucidus","261","239","239","19","8.43","0","98.87","429.08","0","430.57","39.03","0"
"AGAW3","05N","INS","BOLD:ACA8176","Arthropoda","Insecta","Diptera","Chironomidae","","","","175","161","161","20","8","0","99.26","427.98","0","446.82","37.43","0"
"AGAW3","05N","INS","BOLD:ABU5525","Arthropoda","Insecta","Diptera","Chironomidae","Orthocladiinae","Limnophyes","Limnophyes sp. 14ES","168","168","168","21","0","0","98.92","395.67","0","398.38","37.4","0"
"AGAW3","05N","INS","BOLD:AAH3968","Arthropoda","Insecta","Diptera","Sciaridae","","Cratyna","Cratyna cryptospina","117","113","113","22","3.42","0","99.42","440.48","0","443.15","27.37","0"
"AGAW3","05N","INS","BOLD:AAN8770","Arthropoda","Insecta","Diptera","Psychodidae","Psychodinae","Psychoda","Psychoda trinodulosa","110","97","97","23","11.82","0","99.02","439.65","0","440.29","33.36","0"
"AGAW3","05N","INS","BOLD:AAD5253","Arthropoda","Insecta","Diptera","Chironomidae","Orthocladiinae","Thienemanniella","Thienemanniella xena, Thienemanniella vittata","108","81","81","24","25","0","98.62","447.21","0","447.89","33.99","0"
"AGAW3","05N","INS","BOLD:AAN5205","Arthropoda","Insecta","Diptera","Cecidomyiidae","","","CecidInt42 sp. BOLD:AAN5205","107","89","89","25","16.82","0","99.55","446.92","0","447.86","31.51","0"
"AGAW3","05N","INS","BOLD:AAI3449","Arthropoda","Insecta","Hemiptera","Aphididae","Aphidinae","Ovatus","Ovatus crataegarius","105","90","90","26","14.29","0","99.57","447.02","0","447.8","24.94","0"
"AGAW3","05N","INS","BOLD:AAU6967","Arthropoda","Insecta","Coleoptera","Chrysomelidae","Galerucinae","Psylliodes","Psylliodes affinis","101","74","74","27","26.73","0","99.48","435.23","0","445.19","34.98","0"
"AGAW3","05N","INS","BOLD:AAN6447","Arthropoda","Insecta","Diptera","Sciaridae","","Corynoptera","Corynoptera perpusilla","90","65","65","28","27.78","0","99.37","448.26","0","449.22","36.39","0"
"AGAW3","05N","INS","BOLD:ACM0208","Arthropoda","Insecta","Diptera","Cecidomyiidae","","","","71","57","57","29","19.72","0","97.94","448.55","0","449.17","27.19","0"
"AGAW3","05N","INS","BOLD:AAN5332","Arthropoda","Insecta","Diptera","Chironomidae","Chironominae","Polypedilum","","66","66","66","30","0","0","97.62","406.32","0","409.61","32.8","0"
"AGAW3","05N","INS","BOLD:AAK4963","Arthropoda","Insecta","Diptera","Chironomidae","Chironominae","Paracladopelma","Paracladopelma winnelli","54","54","54","31","0","0","98.71","428.74","0","432.48","35.04","0"
"AGAW3","05N","INS","BOLD:AAN5267","Arthropoda","Insecta","Diptera","Cecidomyiidae","","","","54","45","45","31","16.67","0","99.44","446.07","0","448.19","30.68","0"
"AGAW3","05N","INS","BOLD:AAN5249","Arthropoda","Insecta","Diptera","Cecidomyiidae","","","","53","52","52","33","1.89","0","98.7","423.26","0","424.3","33.4","0"
"AGAW3","05N","INS","BOLD:ADH6290","","","","","","","","46","45","45","34","2.17","0","99.41","433.59","0","434.7","38.91","0"
"AGAW3","05N","INS","BOLD:AAL7819","Arthropoda","Insecta","Diptera","Psychodidae","Psychodinae","Psychoda","Psychoda sp., Psychoda sp. BOLD:AAL7819","42","42","42","35","0","0","99.34","415.64","0","416.12","32.71","0"
"AGAW3","05N","INS","BOLD:AAH0134","Arthropoda","Insecta","Coleoptera","Phalacridae","Phalacrinae","Stilbus","Stilbus apicialis","39","39","39","36","0","0","99.01","427.85","0","429.28","31.72","0"
"AGAW3","05N","INS","BOLD:AAN5356","Arthropoda","Insecta","Diptera","Chironomidae","Orthocladiinae","Smittia","","39","35","35","36","10.26","0","99.09","443.69","0","444.67","35.23","0"
"AGAW3","05N","INS","BOLD:AAU9508","Arthropoda","Insecta","Hymenoptera","Dryinidae","Anteoninae","Anteon","Anteon tenuicorne, Anteon cf. pubicorne, Anteon pubicorne","37","37","37","38","0","0","99.14","437.38","0","438.81","29.25","0"
"AGAW3","05N","INS","BOLD:AAN5358","Arthropoda","Insecta","Diptera","Chironomidae","Orthocladiinae","Smittia","Smittia stercoraria, Smittia sp. 22ES","29","29","29","39","0","0","99.11","441.72","0","443.14","37.29","0"
"AGAW3","05N","INS","BOLD:AAF1729","Arthropoda","Insecta","Psocodea","Lachesillidae","Lachesillinae","Lachesilla","Lachesilla pedicularia","23","22","22","40","4.35","0","99.16","446.7","0","448.43","37.3","0"
"AGAW3","05N","INS","BOLD:AAA3453","Arthropoda","Insecta","Diptera","Anthomyiidae","Anthomyiinae","Delia","","21","21","21","41","0","0","99.24","432.86","0","435.62","34.68","0"
"AGAW3","05N","INS","BOLD:ABU5520","Arthropoda","Insecta","Diptera","Sciaridae","","Corynoptera","","20","20","20","42","0","0","99.31","402.6","0","403.85","35.99","0"
"AGAW3","05N","INS","BOLD:AAN5260","Arthropoda","Insecta","Diptera","Cecidomyiidae","","","","20","20","20","42","0","0","98.9","409.1","0","410.65","26.17","0"
"AGAW3","05N","INS","BOLD:AAG2868","Arthropoda","Insecta","Hemiptera","Cicadellidae","Typhlocybinae","Empoasca","Empoasca sp., Empoasca fabae","19","17","17","44","10.53","0","99.47","426.37","0","446.21","32.43","0"
"AGAW3","05N","INS","BOLD:AAG6934","Arthropoda","Insecta","Diptera","Hybotidae","","","","18","18","18","45","0","0","98.85","417.5","0","419.61","38.07","0"
"AGAW3","05N","INS","BOLD:AAN6154","Arthropoda","Insecta","Coleoptera","Latridiidae","Corticariinae","Melanophthalma","Melanophthalma inermis","17","17","17","46","0","0","98.18","428.06","0","429.47","30.71","0"
"AGAW3","05N","INS","BOLD:AAA9899","Arthropoda","Insecta","Hemiptera","Aphididae","Aphidinae","Rhopalosiphum","Rhopalosiphum padi","12","11","11","47","8.33","0","99.72","436.83","0","437.25","25.58","0"
"AGAW3","05N","INS","BOLD:ADG4648","Arthropoda","Insecta","Diptera","Cecidomyiidae","","","","12","12","12","47","0","0","99.03","385.5","0","430.33","26.84","0"
"AGAW3","05N","INS","BOLD:AAN6437","Arthropoda","Insecta","Diptera","Sciaridae","","Corynoptera","Corynoptera saccata","11","11","11","49","0","0","99.4","415.64","0","416.36","39.6","0"
"AGAW3","05N","INS","BOLD:ABA6407","Arthropoda","Insecta","Diptera","Sciaridae","","Pseudolycoriella","","11","11","11","49","0","0","99.24","432.64","0","435.64","34.96","0"
"AGAW3","05N","INS","BOLD:AAA7374","Arthropoda","Insecta","Diptera","Syrphidae","Syrphinae","Sphaerophoria","Sphaerophoria bankowskae, Sphaerophoria infuscata, Sphaerophoria nr. philanthus, Sphaerophoria batava, Sphaerophoria sp., Sphaerophoria rueppellii, Sphaerophoria interrupta-group, Sphaerophoria sp. MA-1, Sphaerophoria cleoae, Sphaerophoria asymmetrica, Sphaerophoria fatarum, Sphaerophoria bengalensis, Sphaerophoria bifurcata, Sphaerophoria taeniata, Sphaerophoria interrupta, Sphaerophoria contigua, Sphaerophoria kaa, Sphaerophoria laurae, Sphaerophoria chongjini, Sphaerophoria boreoalpina, Sphaerophoria sp. BOLDAAA7374, Sphaerophoria macrogaster, Sphaerophoria sulphuripes, Sphaerophoria brevipilosa, Sphaerophoria philanthus, Sphaerophoria pyrrhina, Sphaerophoria abbreviata, Sphaerophoria scripta, Sphaerophoria potentillae, Sphaerophoria longipilosa, Sphaerophoria virgata","11","11","11","49","0","0","98.84","435.64","0","437.36","31.37","0"
"AGAW3","05N","INS","BOLD:ABU5546","Arthropoda","Insecta","Diptera","Psychodidae","Psychodinae","Psychoda","","10","10","10","52","0","0","98.63","435","0","436.3","31.43","0"
"AGAW3","05N","INS","BOLD:AAB1098","Arthropoda","Insecta","Diptera","Culicidae","Culicinae","Ochlerotatus, Aedes","Aedes excrucians, Aedes cantans, Aedes stimulans, Aedes euedes, Aedes riparius, Aedes annulipes, Ochlerotatus cantans, Aedes fitchii, Aedes sp., Ochlerotatus annulipes, Ochlerotatus excrucians","10","10","10","52","0","0","98.47","421.9","0","424","35.2","0"
"AGAW3","05N","INS","BOLD:AAG6743","Arthropoda","Insecta","Diptera","Sarcophagidae","Sarcophaginae","Sarcophaga","Sarcophaga sp. 1 MG, Sarcophaga subvicina","10","10","10","52","0","0","99.33","423.9","0","425.6","34.49","0"
"AGAW3","05N","INS","BOLD:ADH9867","Arthropoda","Insecta","Diptera","","","","","10","10","10","52","0","0","98.27","413.6","0","416.6","37.37","0"
"AGAW3","05N","INS","BOLD:AAA3080","Arthropoda","Insecta","Trichoptera","Limnephilidae","Limnephilinae","Limnephilus","Limnephilus submonilifer","10","10","10","52","0","0","99.47","389.4","0","391.8","34.36","0"
"AGAW3","05N","INS","BOLD:AAV1261","Arthropoda","Insecta","Diptera","Sciaridae","","Bradysia","Bradysia splendida","9","9","9","57","0","0","99.2","435","0","436","33.95","0"
"AGAW3","05N","INS","BOLD:ACV4945","Arthropoda","Insecta","Diptera","Cecidomyiidae","","","","7","7","7","58","0","0","99.26","448","0","450","28.86","0"
"AGAW3","05N","INS","BOLD:ADH4889","Arthropoda","Insecta","Diptera","","","","","7","7","7","58","0","0","98.18","446.71","0","448.57","35.7","0"
"AGAW3","05N","INS","BOLD:AAB9140","Arthropoda","Insecta","Diptera","Calliphoridae","Chrysomyinae","Phormia","Phormia regina","7","7","7","58","0","0","99.34","426","0","427.86","33.06","0"
"AGAW3","05N","INS","BOLD:AAQ2943","Arthropoda","Insecta","Hymenoptera","Braconidae","Rhyssalinae","Acrisis","","6","6","6","61","0","0","98.27","415","0","442.83","26.78","0"
"AGAW3","05N","INS","BOLD:AAO3939","Arthropoda","Insecta","Diptera","Limoniidae","Limoniinae","Dicranomyia","","6","6","6","61","0","0","98.91","446.17","0","447.17","36.52","0"
"AGAW3","05N","INS","BOLD:AAN6433","Arthropoda","Insecta","Diptera","Sciaridae","","Corynoptera","Corynoptera deserta","5","5","5","63","0","0","98.6","428.8","0","430","30.22","0"
"AGAW3","05N","INS","BOLD:AAB2644","Arthropoda","Insecta","Diptera","Chironomidae","Orthocladiinae","Orthocladius","Orthocladius lapponicus, Orthocladius dorenus, Orthocladius mallochi","4","4","4","64","0","0","98.37","427.25","0","428.5","32.02","0"
"AGAW3","05N","INS","BOLD:AAN9807","Arthropoda","Insecta","Coleoptera","Curculionidae","Ceutorhynchinae","Ceutorhynchus","Ceutorhynchus floralis, Ceutorhynchus cakilis, Ceutorhynchus typhae","4","4","4","64","0","0","98.89","449.5","0","450","34.61","0"
"AGAW3","05N","INS","BOLD:AAA9251","Arthropoda","Insecta","Lepidoptera","Gelechiidae","Gelechiinae","Scrobipalpa","Scrobipalpa atriplicella, Scrobipalpa nr. atriplicella sp. 1","4","4","4","64","0","0","98.43","419","0","421.5","34.1","0"
"AGAW3","05N","INS","BOLD:AAN5342","Arthropoda","Insecta","Diptera","Chironomidae","Orthocladiinae","Heleniella","","4","4","4","64","0","0","99.32","439.75","0","440","35.15","0"
"AGAW3","05N","INS","BOLD:AAC0299","Arthropoda","Insecta","Diptera","Chironomidae","Chironominae","Tanytarsus","Tanytarsus sp. 1","4","3","3","64","25","0","98.61","450","0","450","36","0"
"AGAW3","05N","INS","BOLD:AAB7490","Arthropoda","Insecta","Diptera","Chironomidae","Orthocladiinae","Tvetenia","Tvetenia paucunca","3","3","3","69","0","0","98.84","435","0","435.33","34.05","0"
"AGAW3","05N","INS","BOLD:AAB4657","Arthropoda","Insecta","Diptera","Chironomidae","Chironominae","Chironomus","Chironomus maturus","3","3","3","69","0","0","98.67","449.67","0","450","37.85","0"
"AGAW3","05N","INS","BOLD:AAB7030","Arthropoda","Insecta","Diptera","Chironomidae","Chironominae","Chironomus","Chironomus sp. BOLDAAP2996, Chironomus quinnitukqut, Chironomus cf. decorus 2g, Chironomus decorus group sp. 2 IP2013, Chironomus cf. decorus, Chironomus decorus gr.","3","3","3","69","0","0","98.26","434.33","0","435.67","35.01","0"
"AGAW3","05N","INS","BOLD:AAV5325","Arthropoda","Insecta","Diptera","Cecidomyiidae","","","CecidInt12 sp. BOLD:AAV5325","3","3","3","69","0","0","98.67","450.67","0","450","28.52","0"
"AGAW3","05N","INS","BOLD:AAD0853","Arthropoda","Insecta","Diptera","Scathophagidae","Scathophaginae","Scathophaga","Scathophaga lutaria, Scathophaga hiemalis, Scathophaga furcata, Scathophaga stercoraria, Scathophaga inquinata, Scathophaga suilla","3","3","3","69","0","0","98.7","429.67","0","431","31.68","0"
"AGAW3","05N","INS","BOLD:AAP9022","Arthropoda","Insecta","Diptera","Cecidomyiidae","","","","3","3","3","69","0","0","99.03","448","0","450","27.7","0"
"AGAW3","05N","INS","BOLD:ADH9310","Arthropoda","Insecta","Diptera","Anthomyiidae","Anthomyiinae","Anthomyia","Anthomyia procellaris, Anthomyia pluvialis","3","3","3","69","0","0","99.37","387","0","387.67","32.3","0"
"AGAW3","05N","INS","BOLD:AAM6037","Arthropoda","Insecta","Diptera","Cecidomyiidae","Cecidomyiinae","Contarinia, Dasineura","","3","3","3","69","0","0","97.81","411","0","412","26.02","0"
"AGAW3","05N","INS","BOLD:AAG5242","Arthropoda","Insecta","Coleoptera","Curculionidae","Entiminae","Barypeithes","Barypeithes pellucidus","3","3","3","69","0","0","98.52","400.33","0","401.33","39.17","0"
"AGAW3","05N","INS","BOLD:AAO8538","Arthropoda","Insecta","Diptera","Psychodidae","Psychodinae","Psychoda","","2","2","2","78","0","0","98.84","393.5","0","398","31.28","0"
"AGAW3","05N","INS","BOLD:AAP7035","Arthropoda","Insecta","Diptera","Chironomidae","Chironominae","Tanytarsus","","2","2","2","78","0","0","97.43","410","0","405.5","38.7","0"
"AGAW3","05N","INS","BOLD:AAA4121","Arthropoda","Insecta","Diptera","Simuliidae","Simuliinae","Simulium","Simulium vittatum complex, Simulium vittatum, Simulium sp. BOLDAAA4121, Simulium argus, Simulium vittatum s.str.","2","2","2","78","0","0","98.56","412.5","0","414.5","33.62","0"
"AGAW3","05N","INS","BOLD:AAF9041","Arthropoda","Insecta","Diptera","Tipulidae","Tipulinae","Tipula","Tipula oleracea","2","2","2","78","0","0","99.47","372.5","0","373","35.93","0"
"AGAW3","05N","INS","BOLD:ADG0064","Arthropoda","Insecta","Hymenoptera","Braconidae","Aphidiinae","Aphidius","","2","2","2","78","0","0","98.58","386.5","0","425.5","25.7","0"
"AGAW3","05N","INS","BOLD:AAM9347","Arthropoda","Insecta","Diptera","Phoridae","Metopininae","Megaselia","","2","2","2","78","0","0","97.92","361","0","366","33.2","0"
"AGAW3","05N","INS","BOLD:AAG0995","Arthropoda","Insecta","Diptera","Chironomidae","Orthocladiinae","Parakiefferiella","","2","2","2","78","0","0","98.61","361","0","364.5","39.37","0"
"AGAW3","05N","INS","BOLD:ACL4768","Arthropoda","Insecta","Coleoptera","Chrysomelidae","Galerucinae","Chaetocnema","Chaetocnema confinis","2","2","2","78","0","0","99.28","347","0","392","32.54","0"
"AGAW3","05N","INS","BOLD:AAG5457","Arthropoda","Insecta","Diptera","Chironomidae","Chironominae","","","2","2","2","78","0","0","98.24","399.5","0","401","33.84","0"
"AGAW3","05N","INS","BOLD:AAA2815","Arthropoda","Insecta","Lepidoptera","Noctuidae","Noctuinae","Mythimna","Mythimna yukonensis, Mythimna oxygala","2","2","2","78","0","0","99.55","448","0","450","33.33","0"
"AGAW3","05N","INS","BOLD:AAA9101","Arthropoda","Insecta","Lepidoptera","Pyralidae","Phycitinae","Moodna, Caudellia","Moodna ostrinella, Moodna bisinuella, Caudellia nigrella","2","2","2","78","0","0","100","416.5","0","417","34.17","0"
"AGAW3","05N","INS","BOLD:ACF2821","Arthropoda","Insecta","Diptera","Mycetophilidae","Mycetophilinae","Mycetophila","Mycetophila fungorum","2","2","2","78","0","0","98.55","408","0","412.5","34.9","0"
"AGAW3","05N","INS","BOLD:AAU6966","Arthropoda","Insecta","Coleoptera","Throscidae","Throscinae","Trixagus","Trixagus chevrolati","2","2","2","78","0","0","100","413","0","450","39.78","0"
"AGAW3","05N","INS","BOLD:AAB2384","Arthropoda","Insecta","Diptera","Syrphidae","Syrphinae","Eupeodes","Eupeodes duseki, Eupeodes volucris, Eupeodes nielseni, Eupeodes americanus, Eupeodes flukei, Eupeodes vockerothi, Eupeodes lucasi, Eupeodes montivagus, Eupeodes rufipunctatus, Eupeodes rojasi, Eupeodes goeldlini, Eupeodes luniger, Eupeodes punctifer, Eupeodes bucculatus, Eupeodes perplexus, Eupeodes abiskoensis, Eupeodes corollae, Eupeodes pomus, Eupeodes lundbecki, Eupeodes nitens, Eupeodes sp. BOLDAAB2384, Eupeodes curtus, Eupeodes latifasciatus","2","2","2","78","0","0","98","399","0","401","30.81","0"
"AGAW3","05N","INS","BOLD:AAG1701","Arthropoda","Insecta","Diptera","Muscidae","Coenosiinae","Macrorchis","Macrorchis ausoba","2","2","2","78","0","0","99.31","437.5","0","441.5","32.84","0"
"AGAW3","05N","INS","BOLD:AAG4653","Arthropoda","Insecta","Diptera","Syrphidae","Syrphinae","Heringia","Heringia latitarsis, Heringia pubescens, Heringia jakutorum, Heringia rita, Heringia fulvimanus, Heringia sp. BOLDAAG4653, Heringia nr. pubescens, Heringia coxalis, Heringia larusi, Heringia vitripennis","2","2","2","78","0","0","99.06","426.5","0","426.5","30.69","0"
"AGAW3","05N","INS","BOLD:AAC9062","Arthropoda","Insecta","Diptera","Culicidae","Culicinae","Ochlerotatus, Aedes","Aedes fitchii, Ochlerotatus fitchii, Aedes grossbecki, Aedes ventrovittis","2","2","2","78","0","0","100","426.5","0","427.5","32.84","0"
"AGAW3","05N","INS","BOLD:AAG2462","Arthropoda","Insecta","Diptera","Anthomyiidae","Anthomyiinae","Botanophila","Botanophila fugax","2","2","2","78","0","0","97.78","450","0","450","31.56","0"
"AGAW3","05N","INS","BOLD:AAA3181","Arthropoda","Insecta","Diptera","Muscidae","Muscinae","Stomoxys","Stomoxys calcitrans","1","1","1","96","0","0","99.11","448","0","450","33.33","0"
"AGAW3","05N","INS","BOLD:ABV1243","Arthropoda","Insecta","Diptera","Sarcophagidae","Paramacronychiinae, Miltogramminae","Miltogramminae_incertae_sedis, Brachicoma","Miltogramminae_incertae_sedis sp. 1 MG, Brachicoma sarcophagina","1","1","1","96","0","0","99.78","450","0","450","34.22","0"
"AGAW3","05N","INS","BOLD:ABA6514","Arthropoda","Insecta","Diptera","Chironomidae","Chironominae","Rheotanytarsus","","1","1","1","96","0","0","97.04","372","0","374","36.63","0"
"AGAW3","05N","INS","BOLD:ACW5117","Arthropoda","Insecta","Diptera","Chironomidae","Orthocladiinae","Smittia","Smittia leucopogon","1","1","1","96","0","0","100","449","0","450","31.56","0"
"AGAW3","05N","INS","BOLD:AAN1738","Arthropoda","Insecta","Coleoptera","Staphylinidae","Oxytelinae","Anotylus","Anotylus tetracarinatus","1","1","1","96","0","0","100","450","0","450","33.78","0"
"AGAW3","05N","INS","BOLD:AAN5227","Arthropoda","Insecta","Diptera","Cecidomyiidae","","","","1","1","1","96","0","0","98","450","0","450","27.56","0"
"AGAW3","05N","INS","BOLD:AAI4303","Arthropoda","Insecta","Diptera","Chironomidae","Chironominae","Chironomus","Chironomus melanescens","1","1","1","96","0","0","97.31","446","0","450","34.22","0"
"AGAW3","05N","INS","BOLD:ACA3626","Arthropoda","Insecta","Diptera","Ceratopogonidae","Forcipomyiinae","Atrichopogon","Atrichopogon sp. 2, Atrichopogon sp. BOLD:ACA3626","1","1","1","96","0","0","99.75","393","0","395","33.92","0"
"AGAW3","05N","INS","BOLD:ACD8765","Arthropoda","Insecta","Diptera","Dolichopodidae","Dolichopodinae","Gymnopternus","Gymnopternus spectabilis","1","1","1","96","0","0","99.78","450","0","450","34.89","0"
"AGAW3","05N","INS","BOLD:AAC4403","Arthropoda","Insecta","Diptera","Chironomidae","Chironominae","Micropsectra","Micropsectra polita","1","1","1","96","0","0","99.55","448","0","450","34.67","0"
"AGAW3","05N","INS","BOLD:ACL3790","Arthropoda","Insecta","Diptera","Chironomidae","Chironominae","Tanytarsus","","1","1","1","96","0","0","97.32","448","0","450","35.78","0"
"AGAW3","05N","INS","BOLD:ACC8521","Arthropoda","Insecta","Hemiptera","Cicadellidae","Typhlocybinae","Empoasca","","1","1","1","96","0","0","99.11","450","0","450","33.56","0"
"AGAW3","05N","INS","BOLD:AAG5443","Arthropoda","Insecta","Diptera","Chironomidae","Orthocladiinae","","","1","1","1","96","0","0","97.74","354","0","356","33.99","0"
"AGAW3","05N","INS","BOLD:ACU7906","Arthropoda","Insecta","Diptera","Chironomidae","Orthocladiinae","","","1","1","1","96","0","0","98.66","449","0","450","36.22","0"
"AGAW3","05N","INS","BOLD:AAY3269","Arthropoda","Insecta","Diptera","Chironomidae","Diamesinae","Potthastia","Potthastia gaedii","1","1","1","96","0","0","98.22","450","0","450","34.89","0"
"AGAW3","05N","INS","BOLD:ABZ5071","Arthropoda","Insecta","Diptera","Chironomidae","Chironominae","Micropsectra","Micropsectra nigripila","1","1","1","96","0","0","98.61","433","0","450","36.67","0"
"AGAW3","05N","INS","BOLD:ACL3456","Arthropoda","Insecta","Hemiptera","Aphididae","Aphidinae","Uroleucon","Uroleucon solidaginis","1","1","1","96","0","0","97.33","450","0","450","24.89","0"
"AGAW3","05N","INS","BOLD:AAB5609","Arthropoda","Insecta","Diptera","Muscidae","Coenosiinae","Coenosia","Coenosia sp.BOLD:AAB5609, Coenosia tigrina","1","1","1","96","0","0","97.36","416","0","418","32.06","0"
"AGAW3","05N","INS","BOLD:AAG1743","Arthropoda","Insecta","Diptera","Muscidae","Phaoniinae","Phaonia","Phaonia fuscana","1","1","1","96","0","0","97.82","367","0","368","33.97","0"
"AGAW3","05N","INS","BOLD:ACW9475","Arthropoda","Insecta","Coleoptera","Chrysomelidae","Galerucinae","Chaetocnema","Chaetocnema confinis","1","1","1","96","0","0","97.09","412","0","414","32.13","0"
"AGAW3","05N","INS","BOLD:ADA3072","Arthropoda","Insecta","Diptera","Phoridae","Metopininae","Megaselia","Megaselia arcticae","1","1","1","96","0","0","98.22","449","0","450","32","0"
"AGAW3","05N","INS","BOLD:AAN5219","Arthropoda","Insecta","Diptera","Cecidomyiidae","","","","1","1","1","96","0","0","99.33","448","0","450","28.89","0"
"AGAW3","05N","INS","BOLD:AAI2681","Arthropoda","Insecta","Diptera","Chironomidae","Orthocladiinae","Parakiefferiella","","1","1","1","96","0","0","99.78","450","0","450","34.22","0"
"AGAW3","05N","INS","BOLD:AAL2750","Arthropoda","Insecta","Diptera","Muscidae","Phaoniinae","Phaonia","Phaonia tuguriorum","1","1","1","96","0","0","97.56","450","0","450","33.11","0"
"AGAW3","05N","INS","BOLD:AAG5423","Arthropoda","Insecta","Diptera","Chironomidae","Chironominae","Dicrotendipes","","1","1","1","96","0","0","97.33","450","0","450","36.89","0"
"AGAW3","05N","INS","BOLD:ACP6694","Arthropoda","Insecta","Diptera","Chironomidae","Orthocladiinae","","","1","1","1","96","0","0","99.72","354","0","356","33.71","0"
"AGAW3","05N","INS","BOLD:AAA4188","Arthropoda","Insecta","Hymenoptera","Braconidae","Aphidiinae","Aphidius","Aphidius microlophii, Aphidius ervi","1","1","1","96","0","0","99.55","447","0","450","26.44","0"
"AGAW3","05N","INS","BOLD:ABA3960","Arthropoda","Insecta","Coleoptera","Chrysomelidae","Eumolpinae","Paria","Paria fragariae","1","1","1","96","0","0","98.22","450","0","450","36.89","0"
"AGAW3","05N","INS","BOLD:AAD8855","Arthropoda","Insecta","Diptera","Chironomidae","Chironominae","Tanytarsus","Tanytarsus glabrescens","1","1","1","96","0","0","97.85","372","0","374","35.83","0"
"AGAW3","05N","INS","BOLD:AAG6745","Arthropoda","Insecta","Diptera","Calliphoridae","Calliphorinae","Pollenia","Pollenia pediculata","1","1","1","96","0","0","99.11","448","0","450","33.78","0"
"AGAW3","05N","INS","BOLD:AAG2128","Arthropoda","Insecta","Diptera","Tachinidae","Exoristinae","Cryptomeigenia","Cryptomeigenia sp. 3, Cryptomeigenia triangularis, Cryptomeigenia muscoides, Cryptomeigenia CR 82, Cryptomeigenia sp. GER5, Cryptomeigenia theutis, Cryptomeigenia sp. GER3","1","1","1","96","0","0","100","446","0","450","31.11","0"
"AGAW3","05N","INS","BOLD:ABW2407","Arthropoda","Insecta","Diptera","Anthomyiidae","Anthomyiinae","Hylemya","","1","1","1","96","0","0","98.76","404","0","406","31.77","0"
"AGAW3","05N","INS","BOLD:ACG5862","Arthropoda","Insecta","Diptera","Agromyzidae","","","","1","1","1","96","0","0","98.86","439","0","440","34.09","0"
"AGAW3","05N","INS","BOLD:AAB1787","Arthropoda","Insecta","Hemiptera","Aphididae","Aphidinae","Rhopalosiphum, Acyrthosiphon","Acyrthosiphon pisum, Acyrthosiphon pisum 5A, Rhopalosiphum maidis","1","1","1","96","0","0","99.78","448","0","450","26","0"
"AGAW3","05N","INS","BOLD:AAH7158","Arthropoda","Insecta","Diptera","Sarcophagidae","Sarcophaginae","Ravinia","Ravinia stimulans","1","1","1","96","0","0","99.5","399","0","401","33.42","0"
"AGAW3","05N","INS","BOLD:AAE0844","Arthropoda","Insecta","Hemiptera","Pentatomidae","Pentatominae","Euschistus","Euschistus variolarius","1","1","1","96","0","0","98.88","448","0","450","36.22","0"
"AGAW3","05N","INS","BOLD:AAU6656","Arthropoda","Insecta","Diptera","Limoniidae","Chioneinae","Neolimnophila","Neolimnophila placida","1","1","1","96","0","0","97.33","449","0","450","35.11","0"
"AGAW3","05N","INS","BOLD:ABX6186","Arthropoda","Insecta","Diptera","Tipulidae","Tipulinae","Nephrotoma","Nephrotoma ferruginea","1","1","1","96","0","0","98.44","450","0","450","34.22","0"
"AGAW3","05N","INS","BOLD:ACE7414","","","","","","","","1","1","1","96","0","0","99.78","450","0","450","33.33","0"
"AGAW3","05N","INS","BOLD:AAI3059","Arthropoda","Insecta","Lepidoptera","Tineidae","Nemapogoninae","Nemapogon","Nemapogon variatella","1","1","1","96","0","0","100","450","0","450","30.44","0"
"AGAW3","05N","INS","BOLD:ACF7232","Arthropoda","Insecta","Diptera","Chironomidae","Orthocladiinae","Parakiefferiella","","1","1","1","96","0","0","99.48","382","0","450","38.22","0"
"AGAW3","05N","INS","BOLD:ACX5521","Arthropoda","Insecta","Diptera","Chironomidae","Orthocladiinae","","","1","1","1","96","0","0","98.67","376","0","377","35.28","0"
"AGAW3","05N","INS","BOLD:ACB0984","Arthropoda","Insecta","Psocodea","Stenopsocidae","","Graphopsocus","Graphopsocus cruciatus","1","1","1","96","0","0","97.32","411","0","450","36.67","0"
"AGAW3","05N","INS","BOLD:AAG0161","Arthropoda","Insecta","Diptera","Milichiidae","Madizinae","Leptometopa","Leptometopa latipes","1","1","1","96","0","0","99.33","450","0","450","32.67","0"
"AGAW3","05N","INS","BOLD:AAG2460","Arthropoda","Insecta","Diptera","Anthomyiidae","Anthomyiinae","Hydrophoria","Hydrophoria lancifer","1","1","1","96","0","0","99.56","450","0","450","34.22","0"
"AGAW3","05N","INS","BOLD:AAY6534","Arthropoda","Insecta","Coleoptera","Mordellidae","Mordellinae","Mordellina","Mordellina ancilla","1","1","1","96","0","0","98","450","0","450","39.33","0"
"AGAW3","05N","NAR","no match","no match","no match","no match","no match","no match","no match","no match","no match","no match","no match","no match","no match","no match","no match","no match","no match","no match","no match","no match"
"AGAW3","05N","NIN","BOLD:ABW2639","Arthropoda","Arachnida","Trombidiformes","Anystidae","","","","285","274","274","1","3.86","0","99.31","447.45","0","442.98","31.32","0"
"AGAW3","05N","NIN","BOLD:AAI4346","Arthropoda","Arachnida","Opiliones","Phalangiidae","","Paroligolophus, Phalangium","Paroligolophus agrestis, Phalangium opilio","175","173","173","2","1.14","0","99.12","430.22","0","434.83","36.18","0"
"AGAW3","05N","NIN","BOLD:AAA7246","Arthropoda","Collembola","Entomobryomorpha","Entomobryidae","Entomobryinae","Entomobrya","Entomobrya nicoleti, Entomobrya multifasciata, Entomobrya intermedia","144","137","137","3","4.86","0","98.95","443.1","0","444.28","43.54","0"
"AGAW3","05N","NIN","BOLD:AAH6630","Arthropoda","Collembola","Entomobryomorpha","Entomobryidae","Entomobryinae","Entomobrya","Entomobrya unostrigata","138","133","133","4","3.62","0","99.16","437.14","0","438.41","39.17","0"
"AGAW3","05N","NIN","BOLD:ACV5610","","","","","","","","98","73","73","5","25.51","0","99.64","444.97","0","446.19","38.11","0"
"AGAW3","05N","NIN","BOLD:AAA5267","Arthropoda","Collembola","Entomobryomorpha","Isotomidae","","","","68","68","68","6","0","0","98.95","417.51","0","419.34","38.08","0"
"AGAW3","05N","NIN","BOLD:ACZ1295","Arthropoda","Arachnida","Trombidiformes","Anystidae","","","","60","60","60","7","0","0","99.1","373.97","0","383.27","31.16","0"
"AGAW3","05N","NIN","BOLD:AAM7961","Arthropoda","Arachnida","Trombidiformes","Anystidae","","","","35","31","31","8","11.43","0","99.5","443.71","0","440.06","31.3","0"
"AGAW3","05N","NIN","BOLD:AAB7914","Arthropoda","Collembola","Symphypleona","Bourletiellidae","","Bourletiella","","28","28","28","9","0","0","99.49","442.14","0","443.57","38.13","0"
"AGAW3","05N","NIN","BOLD:AAV0589","Arthropoda","Arachnida","Mesostigmata","Blattisociidae","Platyseiinae","Cheiroseius","","19","19","19","10","0","0","99.25","448.79","0","450","27.6","0"
"AGAW3","05N","NIN","BOLD:AAN6642","Arthropoda","Arachnida","Mesostigmata","Blattisociidae","Platyseiinae","Cheiroseius","","17","15","15","11","11.76","0","99.46","427.82","0","435.71","26.42","0"
"AGAW3","05N","NIN","BOLD:AAC7413","Arthropoda","Arachnida","Araneae","Thomisidae","","Ozyptila","Ozyptila praticola","14","14","14","12","0","0","98.43","406","0","448.71","30.84","0"
"AGAW3","05N","NIN","BOLD:ACJ7454","Arthropoda","Collembola","Symphypleona","Bourletiellidae","","","","11","4","4","13","63.64","0","99.88","401.64","0","444.18","39","0"
"AGAW3","05N","NIN","BOLD:ABW2693","Arthropoda","Arachnida","Mesostigmata","Ascidae","","","","2","2","2","14","0","0","99.22","448.5","0","450","35.33","0"
"AGAW3","05N","NIN","BOLD:AAN6711","Arthropoda","Arachnida","Mesostigmata","Parasitidae","","","","1","1","1","15","0","0","97.32","448","0","450","39.56","0"
"AGAW3","05N","NIN","BOLD:ABW0506","Arthropoda","Arachnida","Opiliones","Phalangiidae","Phalangiinae","Rilaena","Rilaena triangularis","1","1","1","15","0","0","97.09","447","0","450","39.56","0"
"AGAW3","05N","NIN","BOLD:AAD2346","Arthropoda","Arachnida","Araneae","Thomisidae","","Xysticus","Xysticus punctatus","1","1","1","15","0","0","98.51","404","0","408","30.88","0"
"AGAW3","05S","BAC","TAX:375853","Proteobacteria","Alphaproteobacteria","Rickettsiales","Rickettsiaceae",NA,"Wolbachia","Wolbachia sp. Hymenoptera_ASGLE646-10","33","19","19","1","42.42","0","97.67","449.03","0","449.58","40.87","0"
"AGAW3","05S","BAC","TAX:91393","Proteobacteria","Alphaproteobacteria","Rickettsiales","Rickettsiaceae",NA,"Wolbachia","","6","6","6","2","0","0","99.07","383.5","0","412.17","39.78","0"
"AGAW3","05S","BAC","TAX:375845","Proteobacteria","Alphaproteobacteria","Rickettsiales","Rickettsiaceae",NA,"Wolbachia","Wolbachia sp. Hymenoptera_ASGLE1440-10","3","3","3","3","0","0","97.95","438.33","0","441.33","41.09","0"
"AGAW3","05S","BAC","TAX:392873","Proteobacteria","Alphaproteobacteria","Rickettsiales","Rickettsiaceae",NA,"Wolbachia","Wolbachia endosymbiont of Scrapter algoensis","3","3","3","3","0","0","97.9","413","0","442.67","40.59","0"
"AGAW3","05S","BAC","TAX:376688","Proteobacteria","Alphaproteobacteria","Rickettsiales","Rickettsiaceae",NA,"Wolbachia","Wolbachia sp. elachBioLep01 BioLep373","2","2","2","5","0","0","98.16","436","0","438.5","41.05","0"
"AGAW3","05S","BAC","TAX:453459","Proteobacteria","Alphaproteobacteria","Rickettsiales","Rickettsiaceae",NA,"Wolbachia","Wolbachia endosymbiont of Ichneumonidae","1","1","1","6","0","0","97.76","446","0","450","40.44","0"
"AGAW3","05S","BAC","TAX:376694","Proteobacteria","Alphaproteobacteria","Rickettsiales","Rickettsiaceae",NA,"Wolbachia","Wolbachia sp. Anacrusis nephrodes","1","1","1","6","0","0","98.43","447","0","450","41.11","0"
"AGAW3","05S","BAC","TAX:375772","Proteobacteria","Alphaproteobacteria","Rickettsiales","Rickettsiaceae",NA,"Wolbachia","Wolbachia sp. Leptogenys JCR25_ASANE483-10","1","1","1","6","0","0","97.53","446","0","450","40.89","0"
"AGAW3","05S","BAC","TAX:375782","Proteobacteria","Alphaproteobacteria","Rickettsiales","Rickettsiaceae",NA,"Wolbachia","Wolbachia sp. Cerapachys L_MG11_ASANH426-10","1","1","1","6","0","0","98","449","0","450","40.89","0"
"AGAW3","05S","BAC","TAX:392874","Proteobacteria","Alphaproteobacteria","Rickettsiales","Rickettsiaceae",NA,"Wolbachia","Wolbachia endosymbiont of Hymenoptera","1","1","1","6","0","0","97.31","446","0","450","40.67","0"
"AGAW3","05S","BAC","TAX:265077","Proteobacteria","Alphaproteobacteria","Rickettsiales","Rickettsiaceae",NA,"Wolbachia","Wolbachia sp. FormicidaeASPNA479-09","1","1","1","6","0","0","97.78","451","0","450","41.11","0"
"AGAW3","05S","BAC","TAX:91391","Proteobacteria","Alphaproteobacteria","Rickettsiales","",NA,"","","1","1","1","6","0","0","97.39","422","0","450","40.22","0"
"AGAW3","05S","CHO","no match","no match","no match","no match","no match","no match","no match","no match","no match","no match","no match","no match","no match","no match","no match","no match","no match","no match","no match","no match"
"AGAW3","05S","INS","BOLD:AAG2511","Arthropoda","Insecta","Diptera","Anthomyiidae","Anthomyiinae","Delia","Delia platura","14839","11883","11883","1","19.92","0","99.12","444.49","0","447.1","33.14","0"
"AGAW3","05S","INS","BOLD:AAB2384","Arthropoda","Insecta","Diptera","Syrphidae","Syrphinae","Eupeodes","Eupeodes duseki, Eupeodes volucris, Eupeodes nielseni, Eupeodes americanus, Eupeodes flukei, Eupeodes vockerothi, Eupeodes lucasi, Eupeodes montivagus, Eupeodes rufipunctatus, Eupeodes rojasi, Eupeodes goeldlini, Eupeodes luniger, Eupeodes punctifer, Eupeodes bucculatus, Eupeodes perplexus, Eupeodes abiskoensis, Eupeodes corollae, Eupeodes pomus, Eupeodes lundbecki, Eupeodes nitens, Eupeodes sp. BOLDAAB2384, Eupeodes curtus, Eupeodes latifasciatus","9907","9119","9119","2","7.95","0","98.89","401.94","0","404.52","30.72","0"
"AGAW3","05S","INS","BOLD:AAG6768","Arthropoda","Insecta","Diptera","Scathophagidae","Scathophaginae","Cordilura","Cordilura pleuritica","4917","4020","4020","3","18.24","0","98.66","434.35","0","435.84","33.97","0"
"AGAW3","05S","INS","BOLD:AAA7374","Arthropoda","Insecta","Diptera","Syrphidae","Syrphinae","Sphaerophoria","Sphaerophoria bankowskae, Sphaerophoria infuscata, Sphaerophoria nr. philanthus, Sphaerophoria batava, Sphaerophoria sp., Sphaerophoria rueppellii, Sphaerophoria interrupta-group, Sphaerophoria sp. MA-1, Sphaerophoria cleoae, Sphaerophoria asymmetrica, Sphaerophoria fatarum, Sphaerophoria bengalensis, Sphaerophoria bifurcata, Sphaerophoria taeniata, Sphaerophoria interrupta, Sphaerophoria contigua, Sphaerophoria kaa, Sphaerophoria laurae, Sphaerophoria chongjini, Sphaerophoria boreoalpina, Sphaerophoria sp. BOLDAAA7374, Sphaerophoria macrogaster, Sphaerophoria sulphuripes, Sphaerophoria brevipilosa, Sphaerophoria philanthus, Sphaerophoria pyrrhina, Sphaerophoria abbreviata, Sphaerophoria scripta, Sphaerophoria potentillae, Sphaerophoria longipilosa, Sphaerophoria virgata","4659","4156","4156","4","10.8","0","99.11","426.37","0","429.82","30.73","0"
"AGAW3","05S","INS","BOLD:AAE0844","Arthropoda","Insecta","Hemiptera","Pentatomidae","Pentatominae","Euschistus","Euschistus variolarius","3802","3248","3248","5","14.57","0","99.08","423.17","0","424.17","36.18","0"
"AGAW3","05S","INS","BOLD:AAF9187","Arthropoda","Insecta","Coleoptera","Curculionidae","Entiminae","Phyllobius","Phyllobius oblongus","2478","2005","2005","6","19.09","0","99.43","437.56","0","439.86","38.87","0"
"AGAW3","05S","INS","BOLD:AAA2482","Arthropoda","Insecta","Lepidoptera","Noctuidae","Noctuinae","Mythimna","Mythimna unipuncta","1154","1097","1097","7","4.94","0","99.08","415.85","0","419.25","30.12","0"
"AGAW3","05S","INS","BOLD:ABA4842","Arthropoda","Insecta","Coleoptera","Elateridae","Agrypninae","Aeolus","Aeolus mellillus","578","419","419","8","27.51","0","99.59","439.79","0","441.15","38.95","0"
"AGAW3","05S","INS","BOLD:AAF1049","Arthropoda","Insecta","Diptera","Agromyzidae","Phytomyzinae","Cerodontha","Cerodontha fasciata","571","464","464","9","18.74","0","99.64","395.73","0","435.22","31.22","0"
"AGAW3","05S","INS","BOLD:AAP8998","","","","","","","","539","501","501","10","7.05","0","98.16","427.65","0","429.47","32.76","0"
"AGAW3","05S","INS","BOLD:AAC9132","Arthropoda","Insecta","Diptera","Culicidae","Culicinae","Culiseta","Culiseta inornata","433","433","433","11","0","0","98.77","404.24","0","406.89","32.36","0"
"AGAW3","05S","INS","BOLD:AAN5342","Arthropoda","Insecta","Diptera","Chironomidae","Orthocladiinae","Heleniella","","274","259","259","12","5.47","0","98.8","424.62","0","430.72","35.14","0"
"AGAW3","05S","INS","BOLD:ACK7296","Arthropoda","Insecta","Diptera","Cecidomyiidae","","","","260","225","225","13","13.46","0","99.41","430.63","0","431.98","24.24","0"
"AGAW3","05S","INS","BOLD:ABV1350","Arthropoda","Insecta","Diptera","Agromyzidae","","","","258","257","257","14","0.39","0","98.79","396.6","0","400.32","36.36","0"
"AGAW3","05S","INS","BOLD:AAG5468","Arthropoda","Insecta","Diptera","Chironomidae","Tanypodinae","","","239","206","206","15","13.81","0","99.13","431.67","0","440.16","34.69","0"
"AGAW3","05S","INS","BOLD:AAO3939","Arthropoda","Insecta","Diptera","Limoniidae","Limoniinae","Dicranomyia","","207","185","185","16","10.63","0","99","437.62","0","439.02","36.24","0"
"AGAW3","05S","INS","BOLD:AAF9771","Arthropoda","Insecta","Diptera","Hybotidae","Tachydromiinae","Platypalpus","Platypalpus longiseta, Platypalpus trivialis, Platypalpus sp. 5, Platypalpus sp. 17, Platypalpus pallidiventris","187","155","155","17","17.11","0","99.42","442.75","0","443.43","35.18","0"
"AGAW3","05S","INS","BOLD:AAM7726","Arthropoda","Insecta","Coleoptera","Curculionidae","Curculioninae","Orchestes","Orchestes steppensis, Orchestes alni","176","147","147","18","16.48","0","99","437.21","0","440.86","33.27","0"
"AGAW3","05S","INS","BOLD:ACR8658","Arthropoda","Insecta","Diptera","Chironomidae","Chironominae","Tanytarsus","","167","159","159","19","4.79","0","99.3","380.25","0","418.23","35.62","0"
"AGAW3","05S","INS","BOLD:ACJ7008","Arthropoda","Insecta","Diptera","Sciaridae","","Leptosciarella","Leptosciarella belingi","137","133","133","20","2.92","0","99.72","349.73","0","417.76","32.67","0"
"AGAW3","05S","INS","BOLD:AAA5565","Arthropoda","Insecta","Hemiptera","Aphididae","Aphidinae","Aphis","Aphis fabae, Aphis newtoni, Aphis solanella, Aphis ilicis, Aphis hederae, Aphis viburni, Aphis lambersi, Aphis fukii","128","115","115","21","10.16","0","99.65","434.02","0","436.86","26.97","0"
"AGAW3","05S","INS","BOLD:AAG5467","Arthropoda","Insecta","Diptera","Chironomidae","Chironominae","Tanytarsus","","115","114","114","22","0.87","0","98.89","405.72","0","409.17","31.46","0"
"AGAW3","05S","INS","BOLD:AAH3664","Arthropoda","Insecta","Diptera","Cecidomyiidae","","","CecidInt3 sp. BOLD:AAH3664","114","110","110","23","3.51","0","99.19","428.09","0","430.62","26.87","0"
"AGAW3","05S","INS","BOLD:AAA9251","Arthropoda","Insecta","Lepidoptera","Gelechiidae","Gelechiinae","Scrobipalpa","Scrobipalpa atriplicella, Scrobipalpa nr. atriplicella sp. 1","110","108","108","24","1.82","0","99.36","414.62","0","416.98","34.22","0"
"AGAW3","05S","INS","BOLD:AAV1055","Arthropoda","Insecta","Diptera","Stratiomyidae","Beridinae","Allognosta","Allognosta fuscitarsis","97","84","84","25","13.4","0","98.37","440.86","0","441.86","35.63","0"
"AGAW3","05S","INS","BOLD:AAG7036","Arthropoda","Insecta","Diptera","Agromyzidae","","","","93","91","91","26","2.15","0","98.91","392.88","0","398.41","33.34","0"
"AGAW3","05S","INS","BOLD:AAH4208","Arthropoda","Insecta","Diptera","Chloropidae","Oscinellinae","Elachiptera","Elachiptera erythropleura, Elachiptera costata","76","68","68","27","10.53","0","98.95","441.72","0","443.2","31.43","0"
"AGAW3","05S","INS","BOLD:AAD7604","Arthropoda","Insecta","Coleoptera","Coccinellidae","Coccinellinae","Coleomegilla","Coleomegilla maculata","73","72","72","28","1.37","0","98.78","433.01","0","435.49","31.7","0"
"AGAW3","05S","INS","BOLD:AAF9014","Arthropoda","Insecta","Diptera","Limoniidae","Chioneinae","Symplecta","Symplecta cana, Symplecta sp. BOLDAAF9014","72","69","69","29","4.17","0","98.42","421.69","0","427.57","34.01","0"
"AGAW3","05S","INS","BOLD:AAN5343","Arthropoda","Insecta","Diptera","Chironomidae","Chironominae","Lauterborniella","Lauterborniella agrayloides","59","57","57","30","3.39","0","98.16","431.46","0","433.69","33.62","0"
"AGAW3","05S","INS","BOLD:AAG5192","Arthropoda","Insecta","Coleoptera","Curculionidae","Entiminae","Barypeithes","Barypeithes pellucidus","51","43","43","31","15.69","0","99.67","438.86","0","440","39.48","0"
"AGAW3","05S","INS","BOLD:ABU5525","Arthropoda","Insecta","Diptera","Chironomidae","Orthocladiinae","Limnophyes","Limnophyes sp. 14ES","46","46","46","32","0","0","98.91","399.35","0","402.65","37.52","0"
"AGAW3","05S","INS","BOLD:AAA3453","Arthropoda","Insecta","Diptera","Anthomyiidae","Anthomyiinae","Delia","","45","45","45","33","0","0","98.79","425.33","0","428.04","34.36","0"
"AGAW3","05S","INS","BOLD:AAM9252","Arthropoda","Insecta","Diptera","Sciaridae","","Bradysia, Leptosciarella","Bradysia vagans, Leptosciarella zanti","35","35","35","34","0","0","99.48","435.97","0","437.51","38.86","0"
"AGAW3","05S","INS","BOLD:AAU6533","Arthropoda","Insecta","Diptera","Phoridae","Metopininae","Megaselia","","34","30","30","35","11.76","0","99.82","355.71","0","440.5","32.22","0"
"AGAW3","05S","INS","BOLD:ADH9832","","","","","","","","33","32","32","36","3.03","0","98.87","448.36","0","449.67","33.32","0"
"AGAW3","05S","INS","BOLD:AAM9347","Arthropoda","Insecta","Diptera","Phoridae","Metopininae","Megaselia","","29","29","29","37","0","0","97.98","392.72","0","395.28","33.19","0"
"AGAW3","05S","INS","BOLD:AAN6149","Arthropoda","Insecta","Coleoptera","Coccinellidae","Coccinellinae","Stethorus","Stethorus punctillum","26","26","26","38","0","0","97.85","393.23","0","410.08","36.54","0"
"AGAW3","05S","INS","BOLD:AAN8694","Arthropoda","Insecta","Diptera","Phoridae","Metopininae","Megaselia","","24","24","24","39","0","0","97.99","382.67","0","384.79","32.3","0"
"AGAW3","05S","INS","BOLD:ACK4716","Arthropoda","Insecta","Diptera","Cecidomyiidae","","","","21","21","21","40","0","0","99.09","434.52","0","435.57","31.62","0"
"AGAW3","05S","INS","BOLD:AAA4222","Arthropoda","Insecta","Lepidoptera","Erebidae","Hypeninae","Hypena","Hypena scabra","14","14","14","41","0","0","98.7","381","0","382.86","32.37","0"
"AGAW3","05S","INS","BOLD:AAG0734","Arthropoda","Insecta","Thysanoptera","Thripidae","Thripinae","Frankliniella","Frankliniella tritici","11","11","11","42","0","0","99.1","405","0","426.91","29.43","0"
"AGAW3","05S","INS","BOLD:ABA0863","Arthropoda","Insecta","Diptera","Cecidomyiidae","","","","9","9","9","43","0","0","99.43","449","0","450","30.32","0"
"AGAW3","05S","INS","BOLD:AAH0256","Arthropoda","Insecta","Coleoptera","Latridiidae","Corticariinae","Corticaria, Corticarina","","9","9","9","43","0","0","98.04","414.78","0","420.11","31.62","0"
"AGAW3","05S","INS","BOLD:AAG6743","Arthropoda","Insecta","Diptera","Sarcophagidae","Sarcophaginae","Sarcophaga","Sarcophaga sp. 1 MG, Sarcophaga subvicina","8","8","8","45","0","0","99.09","424.63","0","426.63","34.84","0"
"AGAW3","05S","INS","BOLD:AAY6412","Arthropoda","Insecta","Diptera","Sciaridae","","Bradysia","Bradysia urticae","7","7","7","46","0","0","99.13","416.71","0","419.57","31.65","0"
"AGAW3","05S","INS","BOLD:AAN8770","Arthropoda","Insecta","Diptera","Psychodidae","Psychodinae","Psychoda","Psychoda trinodulosa","7","7","7","46","0","0","99.29","444.57","0","445.71","33.94","0"
"AGAW3","05S","INS","BOLD:AAN5267","Arthropoda","Insecta","Diptera","Cecidomyiidae","","","","7","5","5","46","28.57","0","99.82","439.57","0","449.71","30.53","0"
"AGAW3","05S","INS","BOLD:AAG3236","Arthropoda","Insecta","Diptera","Phoridae","Phorinae","Conicera, Diplonevra","Diplonevra nitidula","7","7","7","46","0","0","99.27","449","0","450","32.03","0"
"AGAW3","05S","INS","BOLD:ADH9867","Arthropoda","Insecta","Diptera","","","","","7","7","7","46","0","0","98.62","436.71","0","440.86","37.41","0"
"AGAW3","05S","INS","BOLD:AAP6465","Arthropoda","Insecta","Diptera","Sciaridae","","Corynoptera","Corynoptera cursor","7","6","6","46","14.29","0","98.7","439.71","0","441.29","32.26","0"
"AGAW3","05S","INS","BOLD:AAP6369","Arthropoda","Insecta","Coleoptera","Elateridae","Agrypninae","Aeolus","Aeolus mellillus","7","7","7","46","0","0","99.21","435.43","0","435.86","39.11","0"
"AGAW3","05S","INS","BOLD:AAN5356","Arthropoda","Insecta","Diptera","Chironomidae","Orthocladiinae","Smittia","","6","6","6","53","0","0","98.25","448.83","0","450","35.07","0"
"AGAW3","05S","INS","BOLD:AAG2462","Arthropoda","Insecta","Diptera","Anthomyiidae","Anthomyiinae","Botanophila","Botanophila fugax","5","5","5","54","0","0","98.67","442.2","0","442.8","31.78","0"
"AGAW3","05S","INS","BOLD:AAN8172","Arthropoda","Insecta","Hymenoptera","Ichneumonidae","Ophioninae","Ophion","Ophion idoneus","4","4","4","55","0","0","98.92","415.25","0","450","27.89","0"
"AGAW3","05S","INS","BOLD:ACF5939","Arthropoda","Insecta","Diptera","Tachinidae","Exoristinae","Winthemia","Winthemia sinuata, Winthemia illinoensis","4","4","4","55","0","0","98.74","419.5","0","421.75","29.21","0"
"AGAW3","05S","INS","BOLD:ADH9310","Arthropoda","Insecta","Diptera","Anthomyiidae","Anthomyiinae","Anthomyia","Anthomyia procellaris, Anthomyia pluvialis","4","4","4","55","0","0","99.82","394.75","0","396.25","32.38","0"
"AGAW3","05S","INS","BOLD:AAN5355","Arthropoda","Insecta","Diptera","Chironomidae","Orthocladiinae","Smittia","Smittia cf. stercoraria, Smittia stercoraria","3","3","3","58","0","0","98.82","430","0","432.67","36.18","0"
"AGAW3","05S","INS","BOLD:AAB9140","Arthropoda","Insecta","Diptera","Calliphoridae","Chrysomyinae","Phormia","Phormia regina","3","3","3","58","0","0","99.63","448.33","0","450","33.11","0"
"AGAW3","05S","INS","BOLD:AAG1701","Arthropoda","Insecta","Diptera","Muscidae","Coenosiinae","Macrorchis","Macrorchis ausoba","3","3","3","58","0","0","98.62","405","0","406.67","32.87","0"
"AGAW3","05S","INS","BOLD:AAB2644","Arthropoda","Insecta","Diptera","Chironomidae","Orthocladiinae","Orthocladius","Orthocladius lapponicus, Orthocladius dorenus, Orthocladius mallochi","2","2","2","61","0","0","98.76","388.5","0","402","32.07","0"
"AGAW3","05S","INS","BOLD:AAA3836","Arthropoda","Insecta","Lepidoptera","Noctuidae","Plusiinae","Autographa","Autographa precationis","2","2","2","61","0","0","99.44","449","0","450","32.22","0"
"AGAW3","05S","INS","BOLD:AAB4657","Arthropoda","Insecta","Diptera","Chironomidae","Chironominae","Chironomus","Chironomus maturus","2","2","2","61","0","0","99.67","450","0","450","37.67","0"
"AGAW3","05S","INS","BOLD:AAH3914","Arthropoda","Insecta","Diptera","Sciaridae","","Sciara","Sciara hamatilis, Sciara humeralis","2","2","2","61","0","0","99.56","450","0","450","30.67","0"
"AGAW3","05S","INS","BOLD:AAA5299","Arthropoda","Insecta","Diptera","Chironomidae","Orthocladiinae","Cricotopus","Cricotopus laetus, Cricotopus relucens, Cricotopus trifasciatus, Cricotopus sp., Cricotopus sp. 3ES, Cricotopus sylvestris, Cricotopus glacialis","2","2","2","61","0","0","98.07","444","0","445.5","35.13","0"
"AGAW3","05S","INS","BOLD:AAG8493","Arthropoda","Insecta","Diptera","Drosophilidae","Drosophilinae","Scaptomyza","Scaptomyza pallida","2","2","2","61","0","0","98.14","417","0","419","33.61","0"
"AGAW3","05S","INS","BOLD:AAD0484","Arthropoda","Insecta","Diptera","Chironomidae","","","","2","2","2","61","0","0","97.76","445.5","0","450","33.89","0"
"AGAW3","05S","INS","BOLD:AAC2498","Arthropoda","Insecta","Diptera","Muscidae","Phaoniinae","Helina","Helina evecta, Helina aldrichi","2","2","2","61","0","0","98.1","448.5","0","450","30.67","0"
"AGAW3","05S","INS","BOLD:AAH1888","Arthropoda","Insecta","Hymenoptera","Ichneumonidae","Ctenopelmatinae","","","2","2","2","61","0","0","99.1","404","0","405","28.35","0"
"AGAW3","05S","INS","BOLD:ACX5521","Arthropoda","Insecta","Diptera","Chironomidae","Orthocladiinae","","","2","2","2","61","0","0","97.4","426","0","428.5","34.04","0"
"AGAW3","05S","INS","BOLD:AAG4653","Arthropoda","Insecta","Diptera","Syrphidae","Syrphinae","Heringia","Heringia latitarsis, Heringia pubescens, Heringia jakutorum, Heringia rita, Heringia fulvimanus, Heringia sp. BOLDAAG4653, Heringia nr. pubescens, Heringia coxalis, Heringia larusi, Heringia vitripennis","2","2","2","61","0","0","99.22","449","0","450","31.78","0"
"AGAW3","05S","INS","BOLD:AAA3181","Arthropoda","Insecta","Diptera","Muscidae","Muscinae","Stomoxys","Stomoxys calcitrans","1","1","1","72","0","0","99.33","446","0","450","33.56","0"
"AGAW3","05S","INS","BOLD:ABV1243","Arthropoda","Insecta","Diptera","Sarcophagidae","Paramacronychiinae, Miltogramminae","Miltogramminae_incertae_sedis, Brachicoma","Miltogramminae_incertae_sedis sp. 1 MG, Brachicoma sarcophagina","1","1","1","72","0","0","99.78","450","0","450","34.22","0"
"AGAW3","05S","INS","BOLD:ADE4394","Arthropoda","Insecta","Diptera","Chironomidae","Chironominae","Tanytarsus","","1","1","1","72","0","0","98.11","423","0","450","35.78","0"
"AGAW3","05S","INS","BOLD:AAH4135","Arthropoda","Insecta","Diptera","Chloropidae","Chloropinae","Thaumatomyia","Thaumatomyia glabra, Thaumatomyia grata, Thaumatomyia sp. 1 TAW","1","1","1","72","0","0","97.75","445","0","448","31.92","0"
"AGAW3","05S","INS","BOLD:AAO1339","Arthropoda","Insecta","Coleoptera","Anthribidae","Anthribinae","Anthribus","Anthribus nebulosus","1","1","1","72","0","0","99.73","369","0","371","36.39","0"
"AGAW3","05S","INS","BOLD:AAG9668","Arthropoda","Insecta","Diptera","Dolichopodidae","Diaphorinae","Chrysotus","Chrysotus sp., Chrysotus sp. nr. obliquus","1","1","1","72","0","0","97.54","448","0","450","32.44","0"
"AGAW3","05S","INS","BOLD:ABV0271","Arthropoda","Insecta","Diptera","Chloropidae","Oscinellinae","","","1","1","1","72","0","0","99.78","450","0","450","33.33","0"
"AGAW3","05S","INS","BOLD:AAI4303","Arthropoda","Insecta","Diptera","Chironomidae","Chironominae","Chironomus","Chironomus melanescens","1","1","1","72","0","0","98.43","447","0","450","34.22","0"
"AGAW3","05S","INS","BOLD:AAB7490","Arthropoda","Insecta","Diptera","Chironomidae","Orthocladiinae","Tvetenia","Tvetenia paucunca","1","1","1","72","0","0","97.09","447","0","450","33.56","0"
"AGAW3","05S","INS","BOLD:ACD1218","Arthropoda","Insecta","Diptera","Sciaridae","","Leptosciarella","Leptosciarella scutellata","1","1","1","72","0","0","99.75","398","0","398","33.92","0"
"AGAW3","05S","INS","BOLD:ADI0355","Arthropoda","Insecta","Diptera","","","","","1","1","1","72","0","0","98.37","430","0","434","37.1","0"
"AGAW3","05S","INS","BOLD:AAJ7105","Arthropoda","Insecta","Diptera","Agromyzidae","Agromyzinae","Agromyza","Agromyza frontella","1","1","1","72","0","0","99.5","400","0","450","36.44","0"
"AGAW3","05S","INS","BOLD:AAV6211","Arthropoda","Insecta","Diptera","Chironomidae","Chironominae","Paratanytarsus","Paratanytarsus sp. 2ES","1","1","1","72","0","0","97.31","372","0","380","31.84","0"
"AGAW3","05S","INS","BOLD:AAH0127","Arthropoda","Insecta","Coleoptera","Pyrochroidae","Pedilinae","Pedilus","Pedilus lugubris","1","1","1","72","0","0","99.75","398","0","450","33.11","0"
"AGAW3","05S","INS","BOLD:AAB8817","Arthropoda","Insecta","Diptera","Muscidae","Azeliinae","Muscina","Muscina levida","1","1","1","72","0","0","99.56","450","0","450","33.56","0"
"AGAW3","05S","INS","BOLD:AAA5448","Arthropoda","Insecta","Lepidoptera","Geometridae","Ennominae","Eutrapela","Eutrapela clemataria","1","1","1","72","0","0","97.72","395","0","396","28.79","0"
"AGAW3","05S","INS","BOLD:ABV9274","Arthropoda","Insecta","Diptera","Cecidomyiidae","","","","1","1","1","72","0","0","97.05","440","0","440","33.41","0"
"AGAW3","05S","INS","BOLD:AAE0707","Arthropoda","Insecta","Diptera","Chironomidae","Chironominae","Microtendipes","Microtendipes pedellus","1","1","1","72","0","0","99.14","347","0","352","30.11","0"
"AGAW3","05S","INS","BOLD:AAG2868","Arthropoda","Insecta","Hemiptera","Cicadellidae","Typhlocybinae","Empoasca","Empoasca sp., Empoasca fabae","1","1","1","72","0","0","99.78","450","0","450","32.67","0"
"AGAW3","05S","INS","BOLD:AAH0134","Arthropoda","Insecta","Coleoptera","Phalacridae","Phalacrinae","Stilbus","Stilbus apicialis","1","1","1","72","0","0","100","404","0","404","30.94","0"
"AGAW3","05S","INS","BOLD:ACF6291","Arthropoda","Insecta","Coleoptera","Curculionidae","","","","1","1","1","72","0","0","98.65","446","0","450","32.44","0"
"AGAW3","05S","INS","BOLD:ABU5520","Arthropoda","Insecta","Diptera","Sciaridae","","Corynoptera","","1","1","1","72","0","0","99.52","414","0","414","35.02","0"
"AGAW3","05S","INS","BOLD:AAD1485","Arthropoda","Insecta","Diptera","Chironomidae","Chironominae","Paratanytarsus","Paratanytarsus grimmii, Paratanytarsus sp.","1","1","1","72","0","0","98","450","0","450","33.33","0"
"AGAW3","05S","INS","BOLD:AAG5457","Arthropoda","Insecta","Diptera","Chironomidae","Chironominae","","","1","1","1","72","0","0","99.76","413","0","415","33.25","0"
"AGAW3","05S","INS","BOLD:AAU6630","Arthropoda","Insecta","Diptera","Fanniidae","Fanniinae","Fannia","Fannia armata","1","1","1","72","0","0","99.78","448","0","450","33.78","0"
"AGAW3","05S","INS","BOLD:AAN5219","Arthropoda","Insecta","Diptera","Cecidomyiidae","","","","1","1","1","72","0","0","99.78","450","0","450","29.11","0"
"AGAW3","05S","INS","BOLD:AAA2815","Arthropoda","Insecta","Lepidoptera","Noctuidae","Noctuinae","Mythimna","Mythimna yukonensis, Mythimna oxygala","1","1","1","72","0","0","98.89","450","0","450","33.56","0"
"AGAW3","05S","INS","BOLD:AAU6970","Arthropoda","Insecta","Coleoptera","Cleridae","Clerinae","Enoclerus","Enoclerus nigripes","1","1","1","72","0","0","98.51","403","0","403","32.51","0"
"AGAW3","05S","INS","BOLD:AAD5253","Arthropoda","Insecta","Diptera","Chironomidae","Orthocladiinae","Thienemanniella","Thienemanniella xena, Thienemanniella vittata","1","1","1","72","0","0","97.35","453","0","450","32.44","0"
"AGAW3","05S","INS","BOLD:AAA9101","Arthropoda","Insecta","Lepidoptera","Pyralidae","Phycitinae","Moodna, Caudellia","Moodna ostrinella, Moodna bisinuella, Caudellia nigrella","1","1","1","72","0","0","98.67","450","0","450","34","0"
"AGAW3","05S","INS","BOLD:AAF0889","Arthropoda","Insecta","Diptera","Tabanidae","Tabaninae","Hybomitra","Hybomitra lasiophthalma","1","1","1","72","0","0","98.05","411","0","413","31.23","0"
"AGAW3","05S","INS","BOLD:AAH7158","Arthropoda","Insecta","Diptera","Sarcophagidae","Sarcophaginae","Ravinia","Ravinia stimulans","1","1","1","72","0","0","97.79","408","0","413","32.93","0"
"AGAW3","05S","INS","BOLD:ABA9949","Arthropoda","Insecta","Coleoptera","Chrysomelidae","Galerucinae","Chaetocnema","Chaetocnema concinna","1","1","1","72","0","0","98.44","450","0","450","36.22","0"
"AGAW3","05S","INS","BOLD:AAD0853","Arthropoda","Insecta","Diptera","Scathophagidae","Scathophaginae","Scathophaga","Scathophaga lutaria, Scathophaga hiemalis, Scathophaga furcata, Scathophaga stercoraria, Scathophaga inquinata, Scathophaga suilla","1","1","1","72","0","0","98","449","0","450","32.22","0"
"AGAW3","05S","INS","BOLD:AAH0145","Arthropoda","Insecta","Coleoptera","Carabidae","Harpalinae","Anisodactylus","Anisodactylus pitychrous, Anisodactylus sanctaecrucis, Anisodactylus alternans, Anisodactylus discoideus","1","1","1","72","0","0","99.11","450","0","450","32.44","0"
"AGAW3","05S","INS","BOLD:AAD2548","Arthropoda","Insecta","Hemiptera","Aphididae","Aphidinae","Acyrthosiphon","Acyrthosiphon caraganae, Acyrthosiphon sp. 1, Acyrthosiphon kondoi, Acyrthosiphon churchillense","1","1","1","72","0","0","97.33","450","0","450","26.89","0"
"AGAW3","05S","INS","BOLD:AAG2128","Arthropoda","Insecta","Diptera","Tachinidae","Exoristinae","Cryptomeigenia","Cryptomeigenia sp. 3, Cryptomeigenia triangularis, Cryptomeigenia muscoides, Cryptomeigenia CR 82, Cryptomeigenia sp. GER5, Cryptomeigenia theutis, Cryptomeigenia sp. GER3","1","1","1","72","0","0","100","449","0","450","31.11","0"
"AGAW3","05S","INS","BOLD:AAG0314","Arthropoda","Insecta","Diptera","Chironomidae","Tanypodinae","Psectrotanypus","Psectrotanypus sp. ES01","1","1","1","72","0","0","97.67","430","0","435","33.1","0"
"AGAW3","05S","INS","BOLD:AAM6184","Arthropoda","Insecta","Diptera","Ceratopogonidae","Ceratopogoninae","Culicoides","","1","1","1","72","0","0","98.06","361","0","365","37.53","0"
"AGAW3","05S","INS","BOLD:AAM6227","Arthropoda","Insecta","Diptera","Chironomidae","Tanypodinae","Procladius","Procladius denticulatus, Procladius sp.","1","1","1","72","0","0","99.11","450","0","450","31.33","0"
"AGAW3","05S","INS","BOLD:AAU8501","Arthropoda","Insecta","Hymenoptera","Ichneumonidae","Ctenopelmatinae","Perilissus","Perilissus albitarsis, Perilissus rufoniger","1","1","1","72","0","0","99.33","450","0","450","33.11","0"
"AGAW3","05S","INS","BOLD:AAQ0582","Arthropoda","Insecta","Diptera","Anthomyiidae","Anthomyiinae","Lasiomma","","1","1","1","72","0","0","98.67","450","0","450","32.44","0"
"AGAW3","05S","INS","BOLD:AAB1787","Arthropoda","Insecta","Hemiptera","Aphididae","Aphidinae","Rhopalosiphum, Acyrthosiphon","Acyrthosiphon pisum, Acyrthosiphon pisum 5A, Rhopalosiphum maidis","1","1","1","72","0","0","99.11","450","0","450","26","0"
"AGAW3","05S","INS","BOLD:ACA2933","Arthropoda","Insecta","Psocodea","Stenopsocidae","","Graphopsocus","Graphopsocus cruciatus","1","1","1","72","0","0","98.76","403","0","406","37.44","0"
"AGAW3","05S","INS","BOLD:AAG1004","Arthropoda","Insecta","Diptera","Chironomidae","Orthocladiinae","Cricotopus","Cricotopus trifascia, Cricotopus trifascia grp.","1","1","1","72","0","0","98.22","449","0","450","37.11","0"
"AGAW3","05S","INS","BOLD:AAC9062","Arthropoda","Insecta","Diptera","Culicidae","Culicinae","Ochlerotatus, Aedes","Aedes fitchii, Ochlerotatus fitchii, Aedes grossbecki, Aedes ventrovittis","1","1","1","72","0","0","98.43","445","0","450","32.67","0"
"AGAW3","05S","INS","BOLD:AAA3654","Arthropoda","Insecta","Lepidoptera","Crambidae","Crambinae","Crambus","Crambus bigelovi, Crambus johnsoni, Crambus praefectellus","1","1","1","72","0","0","97.47","396","0","396","30.81","0"
"AGAW3","05S","INS","BOLD:ABX6186","Arthropoda","Insecta","Diptera","Tipulidae","Tipulinae","Nephrotoma","Nephrotoma ferruginea","1","1","1","72","0","0","100","450","0","450","34.44","0"
"AGAW3","05S","INS","BOLD:ACB1116","Arthropoda","Insecta","Diptera","Phoridae","Metopininae","Megaselia","","1","1","1","72","0","0","97.1","448","0","450","32.22","0"
"AGAW3","05S","INS","BOLD:AAL4232","Arthropoda","Insecta","Diptera","Agromyzidae","Phytomyzinae","Phytoliriomyza","Phytoliriomyza arctica","1","1","1","72","0","0","100","416","0","418","31.1","0"
"AGAW3","05S","INS","BOLD:AAA7661","Arthropoda","Insecta","Diptera","Culicidae","Culicinae","Culiseta, Culex","Culiseta melanura, Culex articularis, Culex restuans, Culex acharistus","1","1","1","72","0","0","99.33","450","0","450","32.44","0"
"AGAW3","05S","INS","BOLD:ABW7897","Arthropoda","Insecta","Diptera","Cecidomyiidae","","","","1","1","1","72","0","0","98.65","446","0","450","29.78","0"
"AGAW3","05S","INS","BOLD:ABV1229","Arthropoda","Insecta","Diptera","Chironomidae","Orthocladiinae","Eukiefferiella","","1","1","1","72","0","0","97.1","448","0","450","36.44","0"
"AGAW3","05S","INS","BOLD:ABW1310","Arthropoda","Insecta","Diptera","Anthomyiidae","","","","1","1","1","72","0","0","99.75","403","0","406","34.98","0"
"AGAW3","05S","NAR","no match","no match","no match","no match","no match","no match","no match","no match","no match","no match","no match","no match","no match","no match","no match","no match","no match","no match","no match","no match"
"AGAW3","05S","NIN","BOLD:AAA7188","Arthropoda","Arachnida","Araneae","Philodromidae",NA,"Tibellus","Tibellus maritimus, Tibellus oblongus","866","813","813","1","6.12","0","98.37","403.09","0","405.25","32.23","0"
"AGAW3","05S","NIN","BOLD:AAB0726","Arthropoda","Arachnida","Araneae","Lycosidae",NA,"Trochosa","Trochosa ruricola","540","517","517","2","4.26","0","98.91","435.01","0","437.28","31.18","0"
"AGAW3","05S","NIN","BOLD:AAC7802","Arthropoda","Arachnida","Araneae","Lycosidae",NA,"Pardosa","Pardosa distincta","314","263","263","3","16.24","0","98.35","436.66","0","438.21","29.86","0"
"AGAW3","05S","NIN","BOLD:ABW2639","Arthropoda","Arachnida","Trombidiformes","Anystidae",NA,"","","130","119","119","4","8.46","0","99.44","448.89","0","445.04","31.4","0"
"AGAW3","05S","NIN","BOLD:ACV5609","","","","",NA,"","","65","56","56","5","13.85","0","98.91","441.45","0","442.66","38.73","0"
"AGAW3","05S","NIN","BOLD:AAB7914","Arthropoda","Collembola","Symphypleona","Bourletiellidae",NA,"Bourletiella","","45","41","41","6","8.89","0","99.49","445.53","0","446.44","38.09","0"
"AGAW3","05S","NIN","BOLD:ACV5610","","","","",NA,"","","22","22","22","7","0","0","99.47","432","0","433.32","38.07","0"
"AGAW3","05S","NIN","BOLD:ACC0359","Arthropoda","Collembola","Symphypleona","Bourletiellidae",NA,"Bourletiella","","19","17","17","8","10.53","0","99.2","444.26","0","447.21","38.04","0"
"AGAW3","05S","NIN","BOLD:ACZ1295","Arthropoda","Arachnida","Trombidiformes","Anystidae",NA,"","","18","18","18","9","0","0","99.31","370.56","0","374.11","31.18","0"
"AGAW3","05S","NIN","BOLD:ACC8538","Arthropoda","Arachnida","Mesostigmata","Blattisociidae",NA,"","","16","16","16","10","0","0","99.44","398.19","0","399.5","32.69","0"
"AGAW3","05S","NIN","BOLD:AAD1564","Arthropoda","Arachnida","Araneae","Clubionidae",NA,"Clubiona","Clubiona abboti","8","8","8","11","0","0","98.16","409","0","411.25","31.55","0"
"AGAW3","05S","NIN","BOLD:AAM7961","Arthropoda","Arachnida","Trombidiformes","Anystidae",NA,"","","5","5","5","12","0","0","99.56","424.4","0","444","31.16","0"
"AGAW3","05S","NIN","BOLD:AAA6381","Arthropoda","Arachnida","Araneae","Tetragnathidae",NA,"Tetragnatha","Tetragnatha laboriosa","1","1","1","13","0","0","99.75","394","0","395","39.75","0"
"AGAW3","05S","NIN","BOLD:AAI4346","Arthropoda","Arachnida","Opiliones","Phalangiidae",NA,"Paroligolophus, Phalangium","Paroligolophus agrestis, Phalangium opilio","1","1","1","13","0","0","98.76","402","0","407","33.42","0"
"AGAW3","05W","BAC","TAX:376688","Proteobacteria","Alphaproteobacteria","Rickettsiales","Rickettsiaceae",NA,"Wolbachia","Wolbachia sp. elachBioLep01 BioLep373","2","2","2","1","0","0","98.22","449.5","0","450","40.56","0"
"AGAW3","05W","BAC","TAX:175768","Proteobacteria","Alphaproteobacteria","Rickettsiales","Rickettsiaceae",NA,"Wolbachia","Wolbachia pipientis","2","2","2","1","0","0","97.77","448","0","450","39.44","0"
"AGAW3","05W","BAC","TAX:453453","Proteobacteria","Alphaproteobacteria","Rickettsiales","Rickettsiaceae",NA,"Wolbachia","Wolbachia endosymbiont of Rogas luteus","1","1","1","3","0","0","97.56","410","0","410","40.49","0"
"AGAW3","05W","BAC","TAX:91393","Proteobacteria","Alphaproteobacteria","Rickettsiales","Rickettsiaceae",NA,"Wolbachia","","1","1","1","3","0","0","99.49","392","0","450","39.78","0"
"AGAW3","05W","BAC","TAX:392917","Proteobacteria","Alphaproteobacteria","Rickettsiales","Rickettsiaceae",NA,"Wolbachia","Wolbachia endosymbiont of Glyptapanteles Whitfield131","1","1","1","3","0","0","97.93","435","0","435","41.15","0"
"AGAW3","05W","CHO","no match","no match","no match","no match","no match","no match","no match","no match","no match","no match","no match","no match","no match","no match","no match","no match","no match","no match","no match","no match"
"AGAW3","05W","INS","BOLD:AAA2482","Arthropoda","Insecta","Lepidoptera","Noctuidae","Noctuinae","Mythimna","Mythimna unipuncta","14234","12460","12460","1","12.46","0","98.94","414.6","0","417.82","30.36","0"
"AGAW3","05W","INS","BOLD:AAG2511","Arthropoda","Insecta","Diptera","Anthomyiidae","Anthomyiinae","Delia","Delia platura","5450","4738","4738","2","13.06","0","99.03","444.26","0","446.73","33.17","0"
"AGAW3","05W","INS","BOLD:ABX6186","Arthropoda","Insecta","Diptera","Tipulidae","Tipulinae","Nephrotoma","Nephrotoma ferruginea","4706","3867","3867","3","17.83","0","99.33","430.45","0","432.17","34.31","0"
"AGAW3","05W","INS","BOLD:AAP9022","Arthropoda","Insecta","Diptera","Cecidomyiidae","","","","4152","3281","3281","4","20.98","0","99.47","438.98","0","440.23","27.52","0"
"AGAW3","05W","INS","BOLD:ABV0261","Arthropoda","Insecta","Diptera","Chironomidae","Orthocladiinae","","","3587","3247","3247","5","9.48","0","99.21","422.31","0","423.78","33.17","0"
"AGAW3","05W","INS","BOLD:AAG5507","Arthropoda","Insecta","Diptera","Chironomidae","Chironominae","Chironomus","Chironomus atrella","2070","2014","2014","6","2.71","0","98.96","419.54","0","423.37","35.65","0"
"AGAW3","05W","INS","BOLD:AAG6588","Arthropoda","Insecta","Coleoptera","Chrysomelidae","Cassidinae","Deloyala","Deloyala guttata","1122","1027","1027","7","8.47","0","98.87","427.13","0","428.74","31.31","0"
"AGAW3","05W","INS","BOLD:AAG2868","Arthropoda","Insecta","Hemiptera","Cicadellidae","Typhlocybinae","Empoasca","Empoasca sp., Empoasca fabae","843","579","579","8","31.32","0","99.65","440.76","0","446.3","32.44","0"
"AGAW3","05W","INS","BOLD:AAB5609","Arthropoda","Insecta","Diptera","Muscidae","Coenosiinae","Coenosia","Coenosia sp.BOLD:AAB5609, Coenosia tigrina","829","821","821","9","0.97","0","99","410.07","0","412.58","32.76","0"
"AGAW3","05W","INS","BOLD:AAG5192","Arthropoda","Insecta","Coleoptera","Curculionidae","Entiminae","Barypeithes","Barypeithes pellucidus","810","674","674","10","16.79","0","99.39","439.99","0","441.16","39.41","0"
"AGAW3","05W","INS","BOLD:AAH0349","Arthropoda","Insecta","Coleoptera","Chrysomelidae","Cassidinae","Charidotella","Charidotella sexpunctata, Charidotella purpurata","615","575","575","11","6.5","0","99.27","423.97","0","425.49","32.81","0"
"AGAW3","05W","INS","BOLD:AAN5342","Arthropoda","Insecta","Diptera","Chironomidae","Orthocladiinae","Heleniella","","594","575","575","12","3.2","0","99.07","401.48","0","412.2","34.85","0"
"AGAW3","05W","INS","BOLD:AAA5299","Arthropoda","Insecta","Diptera","Chironomidae","Orthocladiinae","Cricotopus","Cricotopus laetus, Cricotopus relucens, Cricotopus trifasciatus, Cricotopus sp., Cricotopus sp. 3ES, Cricotopus sylvestris, Cricotopus glacialis","544","500","500","13","8.09","0","99.32","428.98","0","430.7","34.56","0"
"AGAW3","05W","INS","BOLD:AAO3939","Arthropoda","Insecta","Diptera","Limoniidae","Limoniinae","Dicranomyia","","469","443","443","14","5.54","0","98.97","433.76","0","436.17","36.44","0"
"AGAW3","05W","INS","BOLD:AAN5219","Arthropoda","Insecta","Diptera","Cecidomyiidae","","","","417","327","327","15","21.58","0","99.36","434.98","0","436","28.74","0"
"AGAW3","05W","INS","BOLD:AAF9014","Arthropoda","Insecta","Diptera","Limoniidae","Chioneinae","Symplecta","Symplecta cana, Symplecta sp. BOLDAAF9014","356","356","356","16","0","0","97.91","418.88","0","422.84","33.78","0"
"AGAW3","05W","INS","BOLD:AAE3015","Arthropoda","Insecta","Hemiptera","Aphididae","Aphidinae","Capitophorus","Capitophorus elaeagni","214","206","206","17","3.74","0","99.32","432.39","0","434.21","25.06","0"
"AGAW3","05W","INS","BOLD:ABZ0908","Arthropoda","Insecta","Diptera","Tipulidae","Tipulinae","Nephrotoma","","139","135","135","18","2.88","0","97.48","366.33","0","369.61","34.11","0"
"AGAW3","05W","INS","BOLD:AAU8501","Arthropoda","Insecta","Hymenoptera","Ichneumonidae","Ctenopelmatinae","Perilissus","Perilissus albitarsis, Perilissus rufoniger","107","106","106","19","0.93","0","99","447.09","0","449.45","32.98","0"
"AGAW3","05W","INS","BOLD:ABW7897","Arthropoda","Insecta","Diptera","Cecidomyiidae","","","","88","78","78","20","11.36","0","99.57","443.61","0","444.86","29.76","0"
"AGAW3","05W","INS","BOLD:ABV9277","","","","","","","","68","63","63","21","7.35","0","98.42","430.41","0","432.01","27.95","0"
"AGAW3","05W","INS","BOLD:AAE9431","Arthropoda","Insecta","Coleoptera","Cerambycidae","Lamiinae","Tetrops","Tetrops praeusta, Tetrops starkii","62","62","62","22","0","0","98.82","421.95","0","424.58","35","0"
"AGAW3","05W","INS","BOLD:AAG3628","Arthropoda","Insecta","Diptera","Mycetophilidae","Mycetophilinae","Mycetophila","Mycetophila luctuosa","61","53","53","23","13.11","0","99.48","444.57","0","446.56","32.76","0"
"AGAW3","05W","INS","BOLD:ACJ2981","Arthropoda","Insecta","Lepidoptera","Elachistidae","Elachistinae","Elachista","","53","47","47","24","11.32","0","99.46","443.25","0","444.4","32.46","0"
"AGAW3","05W","INS","BOLD:AAA7661","Arthropoda","Insecta","Diptera","Culicidae","Culicinae","Culiseta, Culex","Culiseta melanura, Culex articularis, Culex restuans, Culex acharistus","46","44","44","25","4.35","0","99.06","438.74","0","440.52","33.02","0"
"AGAW3","05W","INS","BOLD:ACL8125","","","","","","","","40","39","39","26","2.5","0","99.35","440.4","0","441.68","26.89","0"
"AGAW3","05W","INS","BOLD:AAH0134","Arthropoda","Insecta","Coleoptera","Phalacridae","Phalacrinae","Stilbus","Stilbus apicialis","35","35","35","27","0","0","99.13","431.54","0","432.97","32.1","0"
"AGAW3","05W","INS","BOLD:AAH3664","Arthropoda","Insecta","Diptera","Cecidomyiidae","","","CecidInt3 sp. BOLD:AAH3664","28","28","28","28","0","0","99.14","423.21","0","425.68","26.95","0"
"AGAW3","05W","INS","BOLD:ABU5525","Arthropoda","Insecta","Diptera","Chironomidae","Orthocladiinae","Limnophyes","Limnophyes sp. 14ES","26","26","26","29","0","0","98.92","395.73","0","402.08","37.58","0"
"AGAW3","05W","INS","BOLD:AAM9242","Arthropoda","Insecta","Diptera","Sciaridae","","Corynoptera","Corynoptera melanochaeta","24","24","24","30","0","0","99.6","440.46","0","441.79","34.57","0"
"AGAW3","05W","INS","BOLD:AAA3453","Arthropoda","Insecta","Diptera","Anthomyiidae","Anthomyiinae","Delia","","20","20","20","31","0","0","99","434.6","0","438.55","34.58","0"
"AGAW3","05W","INS","BOLD:AAA4222","Arthropoda","Insecta","Lepidoptera","Erebidae","Hypeninae","Hypena","Hypena scabra","18","18","18","32","0","0","99.03","398.56","0","400.11","32.5","0"
"AGAW3","05W","INS","BOLD:AAY6450","Arthropoda","Insecta","Diptera","Cecidomyiidae","","","","11","11","11","33","0","0","99.36","438.55","0","440.45","27.38","0"
"AGAW3","05W","INS","BOLD:AAA7374","Arthropoda","Insecta","Diptera","Syrphidae","Syrphinae","Sphaerophoria","Sphaerophoria bankowskae, Sphaerophoria infuscata, Sphaerophoria nr. philanthus, Sphaerophoria batava, Sphaerophoria sp., Sphaerophoria rueppellii, Sphaerophoria interrupta-group, Sphaerophoria sp. MA-1, Sphaerophoria cleoae, Sphaerophoria asymmetrica, Sphaerophoria fatarum, Sphaerophoria bengalensis, Sphaerophoria bifurcata, Sphaerophoria taeniata, Sphaerophoria interrupta, Sphaerophoria contigua, Sphaerophoria kaa, Sphaerophoria laurae, Sphaerophoria chongjini, Sphaerophoria boreoalpina, Sphaerophoria sp. BOLDAAA7374, Sphaerophoria macrogaster, Sphaerophoria sulphuripes, Sphaerophoria brevipilosa, Sphaerophoria philanthus, Sphaerophoria pyrrhina, Sphaerophoria abbreviata, Sphaerophoria scripta, Sphaerophoria potentillae, Sphaerophoria longipilosa, Sphaerophoria virgata","10","10","10","34","0","0","98.9","409.2","0","411.3","30.95","0"
"AGAW3","05W","INS","BOLD:AAG5242","Arthropoda","Insecta","Coleoptera","Curculionidae","Entiminae","Barypeithes","Barypeithes pellucidus","9","9","9","35","0","0","98.35","415.56","0","416.89","39.39","0"
"AGAW3","05W","INS","BOLD:ACL4768","Arthropoda","Insecta","Coleoptera","Chrysomelidae","Galerucinae","Chaetocnema","Chaetocnema confinis","8","8","8","36","0","0","99.32","348.88","0","415","32.45","0"
"AGAW3","05W","INS","BOLD:AAM7487","Arthropoda","Insecta","Hymenoptera","Diapriidae","Diapriinae","Trichopria","Trichopria nixoni","6","6","6","37","0","0","99.68","416.67","0","418.17","21.47","0"
"AGAW3","05W","INS","BOLD:AAL7819","Arthropoda","Insecta","Diptera","Psychodidae","Psychodinae","Psychoda","Psychoda sp., Psychoda sp. BOLD:AAL7819","6","6","6","37","0","0","99.39","412.67","0","415","32.69","0"
"AGAW3","05W","INS","BOLD:ADH4973","Arthropoda","Insecta","Diptera","Tipulidae","Tipulinae","Nephrotoma","","6","6","6","37","0","0","97.58","385.83","0","389.83","33.63","0"
"AGAW3","05W","INS","BOLD:AAF9187","Arthropoda","Insecta","Coleoptera","Curculionidae","Entiminae","Phyllobius","Phyllobius oblongus","6","6","6","37","0","0","99.22","433.67","0","434.17","38.93","0"
"AGAW3","05W","INS","BOLD:AAN5267","Arthropoda","Insecta","Diptera","Cecidomyiidae","","","","6","5","5","37","16.67","0","99.81","448.67","0","450","30.63","0"
"AGAW3","05W","INS","BOLD:ABW3140","Arthropoda","Insecta","Hymenoptera","Platygastridae","","","","6","5","5","37","16.67","0","99.25","422.83","0","427","26.41","0"
"AGAW3","05W","INS","BOLD:AAN6437","Arthropoda","Insecta","Diptera","Sciaridae","","Corynoptera","Corynoptera saccata","6","6","6","37","0","0","98.81","403.17","0","404.67","39.72","0"
"AGAW3","05W","INS","BOLD:AAG2462","Arthropoda","Insecta","Diptera","Anthomyiidae","Anthomyiinae","Botanophila","Botanophila fugax","5","5","5","44","0","0","98.35","442.2","0","443.4","31.66","0"
"AGAW3","05W","INS","BOLD:ACF5939","Arthropoda","Insecta","Diptera","Tachinidae","Exoristinae","Winthemia","Winthemia sinuata, Winthemia illinoensis","4","4","4","45","0","0","98.83","430.75","0","433","29.36","0"
"AGAW3","05W","INS","BOLD:AAC9062","Arthropoda","Insecta","Diptera","Culicidae","Culicinae","Ochlerotatus, Aedes","Aedes fitchii, Ochlerotatus fitchii, Aedes grossbecki, Aedes ventrovittis","4","4","4","45","0","0","99.6","443.5","0","444.5","33.24","0"
"AGAW3","05W","INS","BOLD:ACB2589","Arthropoda","Insecta","Hymenoptera","Platygastridae","Scelioninae","Telenomus","Telenomus podisi","3","3","3","47","0","0","98.09","435.67","0","444","26.13","0"
"AGAW3","05W","INS","BOLD:ACW9475","Arthropoda","Insecta","Coleoptera","Chrysomelidae","Galerucinae","Chaetocnema","Chaetocnema confinis","3","3","3","47","0","0","97.57","412","0","450","32.44","0"
"AGAW3","05W","INS","BOLD:AAD0484","Arthropoda","Insecta","Diptera","Chironomidae","","","","3","3","3","47","0","0","98.89","449","0","450","34.96","0"
"AGAW3","05W","INS","BOLD:AAN5355","Arthropoda","Insecta","Diptera","Chironomidae","Orthocladiinae","Smittia","Smittia cf. stercoraria, Smittia stercoraria","3","3","3","47","0","0","99.3","405.67","0","413.67","35.94","0"
"AGAW3","05W","INS","BOLD:AAC2498","Arthropoda","Insecta","Diptera","Muscidae","Phaoniinae","Helina","Helina evecta, Helina aldrichi","3","3","3","47","0","0","97.77","435.67","0","438.67","30.23","0"
"AGAW3","05W","INS","BOLD:AAD7145","Arthropoda","Insecta","Diptera","Muscidae","Mydaeinae","Myospila","Myospila meditabunda","2","2","2","52","0","0","97.89","450","0","450","32","0"
"AGAW3","05W","INS","BOLD:AAU6630","Arthropoda","Insecta","Diptera","Fanniidae","Fanniinae","Fannia","Fannia armata","2","2","2","52","0","0","99.3","420","0","422.5","33.25","0"
"AGAW3","05W","INS","BOLD:AAG5453","Arthropoda","Insecta","Diptera","Chironomidae","Chironominae","Chironomus","Chironomus bifurcatus","2","2","2","52","0","0","97.77","447.5","0","450","35.22","0"
"AGAW3","05W","INS","BOLD:AAF9041","Arthropoda","Insecta","Diptera","Tipulidae","Tipulinae","Tipula","Tipula oleracea","2","2","2","52","0","0","98.1","391","0","392.5","35.18","0"
"AGAW3","05W","INS","BOLD:AAV6213","Arthropoda","Insecta","Diptera","Chironomidae","Chironominae","Paratanytarsus","","2","2","2","52","0","0","98.39","325","0","437.5","35.31","0"
"AGAW3","05W","INS","BOLD:AAH3914","Arthropoda","Insecta","Diptera","Sciaridae","","Sciara","Sciara hamatilis, Sciara humeralis","2","2","2","52","0","0","99.55","448","0","450","30.67","0"
"AGAW3","05W","INS","BOLD:AAE0707","Arthropoda","Insecta","Diptera","Chironomidae","Chironominae","Microtendipes","Microtendipes pedellus","2","2","2","52","0","0","97.72","411","0","412.5","31.49","0"
"AGAW3","05W","INS","BOLD:AAV5573","Arthropoda","Insecta","Diptera","Cecidomyiidae","","","","2","2","2","52","0","0","97.77","448","0","450","27.56","0"
"AGAW3","05W","INS","BOLD:AAL2750","Arthropoda","Insecta","Diptera","Muscidae","Phaoniinae","Phaonia","Phaonia tuguriorum","2","2","2","52","0","0","99.89","449","0","450","33.22","0"
"AGAW3","05W","INS","BOLD:AAG6745","Arthropoda","Insecta","Diptera","Calliphoridae","Calliphorinae","Pollenia","Pollenia pediculata","2","2","2","52","0","0","98.9","426.5","0","429.5","33.39","0"
"AGAW3","05W","INS","BOLD:AAG2128","Arthropoda","Insecta","Diptera","Tachinidae","Exoristinae","Cryptomeigenia","Cryptomeigenia sp. 3, Cryptomeigenia triangularis, Cryptomeigenia muscoides, Cryptomeigenia CR 82, Cryptomeigenia sp. GER5, Cryptomeigenia theutis, Cryptomeigenia sp. GER3","2","2","2","52","0","0","98.89","449.5","0","450","31.33","0"
"AGAW3","05W","INS","BOLD:AAH3548","Arthropoda","Insecta","Diptera","Sphaeroceridae","Copromyzinae","Lotophila","Lotophila sp., Lotophila atra","1","1","1","63","0","0","99.11","449","0","450","32.44","0"
"AGAW3","05W","INS","BOLD:AAG9055","Arthropoda","Insecta","Hymenoptera","Vespidae","Vespinae","Vespula","Vespula germanica","1","1","1","63","0","0","98.89","449","0","450","30.67","0"
"AGAW3","05W","INS","BOLD:AAH4135","Arthropoda","Insecta","Diptera","Chloropidae","Chloropinae","Thaumatomyia","Thaumatomyia glabra, Thaumatomyia grata, Thaumatomyia sp. 1 TAW","1","1","1","63","0","0","98.15","378","0","380","31.32","0"
"AGAW3","05W","INS","BOLD:AAB2644","Arthropoda","Insecta","Diptera","Chironomidae","Orthocladiinae","Orthocladius","Orthocladius lapponicus, Orthocladius dorenus, Orthocladius mallochi","1","1","1","63","0","0","100","449","0","450","32.22","0"
"AGAW3","05W","INS","BOLD:AAA3933","Arthropoda","Insecta","Lepidoptera","Erebidae","Herminiinae","Palthis","Palthis angulalis","1","1","1","63","0","0","99.33","446","0","450","31.78","0"
"AGAW3","05W","INS","BOLD:ACT0257","","","","","","","","1","1","1","63","0","0","98.89","361","0","363","37.47","0"
"AGAW3","05W","INS","BOLD:AAA3836","Arthropoda","Insecta","Lepidoptera","Noctuidae","Plusiinae","Autographa","Autographa precationis","1","1","1","63","0","0","100","447","0","450","31.78","0"
"AGAW3","05W","INS","BOLD:ABX6298","Arthropoda","Insecta","Diptera","Tipulidae","Tipulinae","Nephrotoma","Nephrotoma ferruginea","1","1","1","63","0","0","98.37","368","0","369","33.6","0"
"AGAW3","05W","INS","BOLD:AAP7035","Arthropoda","Insecta","Diptera","Chironomidae","Chironominae","Tanytarsus","","1","1","1","63","0","0","99.55","448","0","450","37.56","0"
"AGAW3","05W","INS","BOLD:AAI4303","Arthropoda","Insecta","Diptera","Chironomidae","Chironominae","Chironomus","Chironomus melanescens","1","1","1","63","0","0","97.76","446","0","450","33.78","0"
"AGAW3","05W","INS","BOLD:AAM7657","","","","","","","","1","1","1","63","0","0","99.11","449","0","450","34.44","0"
"AGAW3","05W","INS","BOLD:AAH1888","Arthropoda","Insecta","Hymenoptera","Ichneumonidae","Ctenopelmatinae","","","1","1","1","63","0","0","98.44","450","0","450","29.33","0"
"AGAW3","05W","INS","BOLD:AAA4121","Arthropoda","Insecta","Diptera","Simuliidae","Simuliinae","Simulium","Simulium vittatum complex, Simulium vittatum, Simulium sp. BOLDAAA4121, Simulium argus, Simulium vittatum s.str.","1","1","1","63","0","0","98.42","444","0","450","33.78","0"
"AGAW3","05W","INS","BOLD:ABW2407","Arthropoda","Insecta","Diptera","Anthomyiidae","Anthomyiinae","Hylemya","","1","1","1","63","0","0","99.55","449","0","450","32.44","0"
"AGAW3","05W","INS","BOLD:ABA0755","Arthropoda","Insecta","Diptera","Drosophilidae","","","","1","1","1","63","0","0","98.89","450","0","450","31.56","0"
"AGAW3","05W","INS","BOLD:AAN6406","Arthropoda","Insecta","Diptera","Sphaeroceridae","Limosininae","Opalimosina","Opalimosina mirabilis","1","1","1","63","0","0","98.12","373","0","377","31.56","0"
"AGAW3","05W","INS","BOLD:AAD8749","Arthropoda","Insecta","Lepidoptera","Gelechiidae","Pexicopiinae","Platyedra","Platyedra subcinerea","1","1","1","63","0","0","98.22","449","0","450","31.56","0"
"AGAW3","05W","INS","BOLD:AAH3920","Arthropoda","Insecta","Diptera","Sciaridae","","Dichopygina, Scatopsciara","","1","1","1","63","0","0","97.24","398","0","450","36.22","0"
"AGAW3","05W","INS","BOLD:AAU6592","Arthropoda","Insecta","Diptera","Ceratopogonidae","Dasyheleinae","Dasyhelea","Dasyhelea arenivaga","1","1","1","63","0","0","99.75","398","0","399","34.09","0"
"AGAW3","05W","INS","BOLD:AAF9771","Arthropoda","Insecta","Diptera","Hybotidae","Tachydromiinae","Platypalpus","Platypalpus longiseta, Platypalpus trivialis, Platypalpus sp. 5, Platypalpus sp. 17, Platypalpus pallidiventris","1","1","1","63","0","0","98.44","449","0","450","35.33","0"
"AGAW3","05W","INS","BOLD:AAB9096","Arthropoda","Insecta","Lepidoptera","Gracillariidae","Gracillariinae","Caloptilia","Caloptilia bimaculatella","1","1","1","63","0","0","100","449","0","450","30.44","0"
"AGAW3","05W","INS","BOLD:ACQ8463","Arthropoda","Insecta","Diptera","Limoniidae","Limoniinae","Antocha","","1","1","1","63","0","0","97.32","448","0","450","37.56","0"
"AGAW3","05W","INS","BOLD:ACK4716","Arthropoda","Insecta","Diptera","Cecidomyiidae","","","","1","1","1","63","0","0","99.33","449","0","450","31.56","0"
"AGAW3","05W","INS","BOLD:ABX2597","Arthropoda","Insecta","Hymenoptera","Proctotrupidae","","","","1","1","1","63","0","0","98.89","449","0","450","26.44","0"
"AGAW3","05W","INS","BOLD:AAG1503","Arthropoda","Insecta","Diptera","Chloropidae","Oscinellinae","Tricimba","Tricimba melancholica","1","1","1","63","0","0","97.77","448","0","450","30.89","0"
"AGAW3","05W","INS","BOLD:AAB9140","Arthropoda","Insecta","Diptera","Calliphoridae","Chrysomyinae","Phormia","Phormia regina","1","1","1","63","0","0","97.11","450","0","450","33.11","0"
"AGAW3","05W","INS","BOLD:AAG6768","Arthropoda","Insecta","Diptera","Scathophagidae","Scathophaginae","Cordilura","Cordilura pleuritica","1","1","1","63","0","0","97.78","450","0","450","34","0"
"AGAW3","05W","INS","BOLD:AAN5356","Arthropoda","Insecta","Diptera","Chironomidae","Orthocladiinae","Smittia","","1","1","1","63","0","0","97.32","448","0","450","34.89","0"
"AGAW3","05W","INS","BOLD:AAJ7105","Arthropoda","Insecta","Diptera","Agromyzidae","Agromyzinae","Agromyza","Agromyza frontella","1","1","1","63","0","0","97.49","399","0","424","34.67","0"
"AGAW3","05W","INS","BOLD:AAH0145","Arthropoda","Insecta","Coleoptera","Carabidae","Harpalinae","Anisodactylus","Anisodactylus pitychrous, Anisodactylus sanctaecrucis, Anisodactylus alternans, Anisodactylus discoideus","1","1","1","63","0","0","98.89","451","0","450","32.22","0"
"AGAW3","05W","INS","BOLD:AAB1098","Arthropoda","Insecta","Diptera","Culicidae","Culicinae","Ochlerotatus, Aedes","Aedes excrucians, Aedes cantans, Aedes stimulans, Aedes euedes, Aedes riparius, Aedes annulipes, Ochlerotatus cantans, Aedes fitchii, Aedes sp., Ochlerotatus annulipes, Ochlerotatus excrucians","1","1","1","63","0","0","98.44","450","0","450","35.78","0"
"AGAW3","05W","INS","BOLD:AAN5249","Arthropoda","Insecta","Diptera","Cecidomyiidae","","","","1","1","1","63","0","0","98.01","403","0","402","32.59","0"
"AGAW3","05W","INS","BOLD:AAN6447","Arthropoda","Insecta","Diptera","Sciaridae","","Corynoptera","Corynoptera perpusilla","1","1","1","63","0","0","97.54","448","0","450","36.44","0"
"AGAW3","05W","INS","BOLD:AAG3236","Arthropoda","Insecta","Diptera","Phoridae","Phorinae","Conicera, Diplonevra","Diplonevra nitidula","1","1","1","63","0","0","100","449","0","450","31.78","0"
"AGAW3","05W","INS","BOLD:AAB1787","Arthropoda","Insecta","Hemiptera","Aphididae","Aphidinae","Rhopalosiphum, Acyrthosiphon","Acyrthosiphon pisum, Acyrthosiphon pisum 5A, Rhopalosiphum maidis","1","1","1","63","0","0","99.33","449","0","450","26.44","0"
"AGAW3","05W","INS","BOLD:AAH7158","Arthropoda","Insecta","Diptera","Sarcophagidae","Sarcophaginae","Ravinia","Ravinia stimulans","1","1","1","63","0","0","99.55","447","0","450","33.56","0"
"AGAW3","05W","INS","BOLD:ADH9867","Arthropoda","Insecta","Diptera","","","","","1","1","1","63","0","0","98.88","445","0","450","37.56","0"
"AGAW3","05W","INS","BOLD:AAP6465","Arthropoda","Insecta","Diptera","Sciaridae","","Corynoptera","Corynoptera cursor","1","1","1","63","0","0","98.67","450","0","450","32.22","0"
"AGAW3","05W","INS","BOLD:AAM6037","Arthropoda","Insecta","Diptera","Cecidomyiidae","Cecidomyiinae","Contarinia, Dasineura","","1","1","1","63","0","0","98","449","0","450","26.67","0"
"AGAW3","05W","INS","BOLD:ACJ1150","Arthropoda","Insecta","Hymenoptera","Ichneumonidae","Campopleginae","","","1","1","1","63","0","0","97.1","449","0","450","26.67","0"
"AGAW3","05W","INS","BOLD:AAA3654","Arthropoda","Insecta","Lepidoptera","Crambidae","Crambinae","Crambus","Crambus bigelovi, Crambus johnsoni, Crambus praefectellus","1","1","1","63","0","0","99.46","370","0","373","31.37","0"
"AGAW3","05W","INS","BOLD:ACP4736","Arthropoda","Insecta","Diptera","Chironomidae","Orthocladiinae","Smittia","Smittia sp. 8ES","1","1","1","63","0","0","97.39","383","0","384","32.55","0"
"AGAW3","05W","INS","BOLD:ACB0984","Arthropoda","Insecta","Psocodea","Stenopsocidae","","Graphopsocus","Graphopsocus cruciatus","1","1","1","63","0","0","97.09","413","0","450","37.56","0"
"AGAW3","05W","INS","BOLD:AAE4298","Arthropoda","Insecta","Diptera","Chironomidae","Orthocladiinae","Cricotopus","Cricotopus tremulus","1","1","1","63","0","0","99.55","448","0","450","32","0"
"AGAW3","05W","INS","BOLD:AAE7016","Arthropoda","Insecta","Lepidoptera","Gelechiidae","Dichomeridinae","Helcystogramma","Helcystogramma hystricella","1","1","1","63","0","0","97.56","450","0","450","30.89","0"
"AGAW3","05W","INS","BOLD:AAG2740","Arthropoda","Insecta","Diptera","Ephydridae","Parydrinae","Philygria","Philygria debilis","1","1","1","63","0","0","97.77","449","0","450","34","0"
"AGAW3","05W","INS","BOLD:ABV1229","Arthropoda","Insecta","Diptera","Chironomidae","Orthocladiinae","Eukiefferiella","","1","1","1","63","0","0","97.77","448","0","450","36.22","0"
"AGAW3","05W","INS","BOLD:ADH9310","Arthropoda","Insecta","Diptera","Anthomyiidae","Anthomyiinae","Anthomyia","Anthomyia procellaris, Anthomyia pluvialis","1","1","1","63","0","0","99.17","360","0","363","31.96","0"
"AGAW3","05W","INS","BOLD:AAG2465","Arthropoda","Insecta","Diptera","Anthomyiidae","Anthomyiinae","Delia","Delia radicum","1","1","1","63","0","0","99.5","403","0","404","33.66","0"
"AGAW3","05W","NAR","no match","no match","no match","no match","no match","no match","no match","no match","no match","no match","no match","no match","no match","no match","no match","no match","no match","no match","no match","no match"
"AGAW3","05W","NIN","BOLD:AAC1568","Arthropoda","Arachnida","Araneae","Thomisidae","","Xysticus","Xysticus elegans","2839","2333","2333","1","17.82","0","97.83","402.76","0","437.53","31.71","0"
"AGAW3","05W","NIN","BOLD:AAI4346","Arthropoda","Arachnida","Opiliones","Phalangiidae","","Paroligolophus, Phalangium","Paroligolophus agrestis, Phalangium opilio","924","898","898","2","2.81","0","98.89","415.51","0","425.74","35.53","0"
"AGAW3","05W","NIN","BOLD:ABW2639","Arthropoda","Arachnida","Trombidiformes","Anystidae","","","","385","362","362","3","5.97","0","99.28","446.72","0","443.32","31.29","0"
"AGAW3","05W","NIN","BOLD:ACI9908","Arthropoda","Collembola","Symphypleona","Bourletiellidae","","","","95","86","86","4","9.47","0","99.35","434.2","0","436","34.91","0"
"AGAW3","05W","NIN","BOLD:AAD1564","Arthropoda","Arachnida","Araneae","Clubionidae","","Clubiona","Clubiona abboti","68","68","68","5","0","0","98.32","406.12","0","408.63","31.95","0"
"AGAW3","05W","NIN","BOLD:ACZ1295","Arthropoda","Arachnida","Trombidiformes","Anystidae","","","","61","61","61","6","0","0","98.95","373","0","383.87","31.07","0"
"AGAW3","05W","NIN","BOLD:AAB7914","Arthropoda","Collembola","Symphypleona","Bourletiellidae","","Bourletiella","","43","38","38","7","11.63","0","99.47","443.91","0","445.26","38.05","0"
"AGAW3","05W","NIN","BOLD:AAV0589","Arthropoda","Arachnida","Mesostigmata","Blattisociidae","Platyseiinae","Cheiroseius","","30","30","30","8","0","0","98.87","445.13","0","446.5","27.56","0"
"AGAW3","05W","NIN","BOLD:ACC0359","Arthropoda","Collembola","Symphypleona","Bourletiellidae","","Bourletiella","","25","20","20","9","20","0","99.71","445.88","0","449.32","37.48","0"
"AGAW3","05W","NIN","BOLD:ACA2345","Arthropoda","Arachnida","Araneae","Linyphiidae","","Agyneta","Agyneta sheffordiana","24","24","24","10","0","0","99.2","437.5","0","438.54","35.5","0"
"AGAW3","05W","NIN","BOLD:AAN6642","Arthropoda","Arachnida","Mesostigmata","Blattisociidae","Platyseiinae","Cheiroseius","","3","3","3","11","0","0","100","449.33","0","450","26.44","0"
"AGAW3","05W","NIN","BOLD:AAM7961","Arthropoda","Arachnida","Trombidiformes","Anystidae","","","","2","2","2","12","0","0","99.57","467.5","0","450","31.44","0"
"AGAW3","05W","NIN","BOLD:AAH6630","Arthropoda","Collembola","Entomobryomorpha","Entomobryidae","Entomobryinae","Entomobrya","Entomobrya unostrigata","1","1","1","13","0","0","99.55","448","0","450","38.67","0"
"AGAW3","05W","NIN","BOLD:AAN4906","Arthropoda","Collembola","Symphypleona","Bourletiellidae","","","","1","1","1","13","0","0","98.35","364","0","367","35.97","0"
"AGAW3","05W","NIN","BOLD:AAB7601","Arthropoda","Arachnida","Araneae","Eutichuridae","","Cheiracanthium","Cheiracanthium mildei","1","1","1","13","0","0","98.44","448","0","450","28.67","0"
"AGAW3","05W","NIN","BOLD:AAA7246","Arthropoda","Collembola","Entomobryomorpha","Entomobryidae","Entomobryinae","Entomobrya","Entomobrya nicoleti, Entomobrya multifasciata, Entomobrya intermedia","1","1","1","13","0","0","97.1","448","0","450","43.78","0"
"AGAW3","08E","BAC","TAX:175768","Proteobacteria","Alphaproteobacteria","Rickettsiales","Rickettsiaceae",NA,"Wolbachia","Wolbachia pipientis","183","148","148","1","19.13","0","99.27","442.77","0","443.94","39.62","0"
"AGAW3","08E","BAC","TAX:91391","Proteobacteria","Alphaproteobacteria","Rickettsiales","",NA,"","","24","19","19","2","20.83","0","99.16","402.21","0","447.46","39.13","0"
"AGAW3","08E","BAC","TAX:91393","Proteobacteria","Alphaproteobacteria","Rickettsiales","Rickettsiaceae",NA,"Wolbachia","","23","23","23","3","0","0","98.91","390.48","0","419.57","39.12","0"
"AGAW3","08E","BAC","TAX:376688","Proteobacteria","Alphaproteobacteria","Rickettsiales","Rickettsiaceae",NA,"Wolbachia","Wolbachia sp. elachBioLep01 BioLep373","2","2","2","4","0","0","99.78","449","0","450","40.44","0"
"AGAW3","08E","CHO","no match","no match","no match","no match","no match","no match","no match","no match","no match","no match","no match","no match","no match","no match","no match","no match","no match","no match","no match","no match"
"AGAW3","08E","INS","BOLD:AAG1711","Arthropoda","Insecta","Diptera","Muscidae","Phaoniinae","Helina","Helina depuncta","16295","12497","12497","1","23.31","0","99.42","434.2","0","436.51","31.36","0"
"AGAW3","08E","INS","BOLD:AAB0373","Arthropoda","Insecta","Neuroptera","Chrysopidae","Chrysopinae","Chrysoperla","Chrysoperla sp. isolate 82, Chrysoperla zastrowi, Chrysoperla plorabunda, Chrysoperla sp. isolate 71, Chrysoperla agilis, Italochrysa italica, Chrysoperla sp. isolate 88, Chrysoperla lucasina, Chrysoperla carnea, Chrysoperla mediterranea, Chrysoperla pallida, Chrysoperla nipponensis, Chrysoperla sillemi","4576","3307","3307","2","27.73","0","99.27","437.5","0","438.35","31.58","0"
"AGAW3","08E","INS","BOLD:AAI4303","Arthropoda","Insecta","Diptera","Chironomidae","Chironominae","Chironomus","Chironomus melanescens","3798","3396","3396","3","10.58","0","98.58","430.23","0","435.07","34.09","0"
"AGAW3","08E","INS","BOLD:AAP8651","Arthropoda","Insecta","Diptera","Tachinidae","Exoristinae","Oswaldia","Oswaldia albifacies","3592","2851","2851","4","20.63","0","98.37","440.94","0","442.67","31.88","0"
"AGAW3","08E","INS","BOLD:AAA3181","Arthropoda","Insecta","Diptera","Muscidae","Muscinae","Stomoxys","Stomoxys calcitrans","3505","2939","2939","5","16.15","0","98.87","428.09","0","431.79","32.8","0"
"AGAW3","08E","INS","BOLD:AAC0903","Arthropoda","Insecta","Diptera","Chironomidae","Chironominae","Chironomus","Chironomus acidophilus","3354","2527","2527","6","24.66","0","99.46","437.63","0","440.93","36.27","0"
"AGAW3","08E","INS","BOLD:AAA4222","Arthropoda","Insecta","Lepidoptera","Erebidae","Hypeninae","Hypena","Hypena scabra","3312","3114","3114","7","5.98","0","99.17","402.01","0","404","32.38","0"
"AGAW3","08E","INS","BOLD:AAC0635","Arthropoda","Insecta","Hemiptera","Miridae","Mirinae","Stenotus","Stenotus binotatus","2708","1904","1904","8","29.69","0","99.37","442.03","0","442.98","34.59","0"
"AGAW3","08E","INS","BOLD:AAZ5483","Arthropoda","Insecta","Diptera","Muscidae","Phaoniinae","Helina","Helina setiventris","2410","2020","2020","9","16.18","0","99.13","441.46","0","444.01","29.3","0"
"AGAW3","08E","INS","BOLD:AAA3075","Arthropoda","Insecta","Lepidoptera","Geometridae","Larentiinae","Eupithecia, Pasiphila","Pasiphila rectangulata","2153","1693","1693","10","21.37","0","99.35","430.02","0","431.43","30.89","0"
"AGAW3","08E","INS","BOLD:AAG1739","Arthropoda","Insecta","Diptera","Muscidae","Coenosiinae","Neodexiopsis","Neodexiopsis rufitibia","1731","1350","1350","11","22.01","0","99.39","441.9","0","443.95","33.91","0"
"AGAW3","08E","INS","BOLD:AAH4285","Arthropoda","Insecta","Lepidoptera","Cosmopterigidae","Cosmopteriginae","Cosmopterix","Cosmopterix montisella","1634","1249","1249","12","23.56","0","99.44","446.88","0","448","32.58","0"
"AGAW3","08E","INS","BOLD:AAA4277","Arthropoda","Insecta","Diptera","Syrphidae","Syrphinae","Toxomerus","Toxomerus marginatus, Toxomerus sp. CR-3, Toxomerus CR-3","1521","1063","1063","13","30.11","0","99.26","444.58","0","445.45","30.49","0"
"AGAW3","08E","INS","BOLD:AAA8279","Arthropoda","Insecta","Lepidoptera","Noctuidae","Noctuinae","Neoligia","Neoligia subjuncta","1488","1200","1200","14","19.35","0","99.4","438.46","0","439.81","30.73","0"
"AGAW3","08E","INS","BOLD:AAG5507","Arthropoda","Insecta","Diptera","Chironomidae","Chironominae","Chironomus","Chironomus atrella","1344","1288","1288","15","4.17","0","98.79","425.52","0","429.94","35.79","0"
"AGAW3","08E","INS","BOLD:ADG1318","","","","","","","","1307","1073","1073","16","17.9","0","99.48","438.36","0","440.64","25.04","0"
"AGAW3","08E","INS","BOLD:AAA7661","Arthropoda","Insecta","Diptera","Culicidae","Culicinae","Culiseta, Culex","Culiseta melanura, Culex articularis, Culex restuans, Culex acharistus","1294","1166","1166","17","9.89","0","99.12","434.69","0","436.74","33.01","0"
"AGAW3","08E","INS","BOLD:ABY5318","Arthropoda","Insecta","Diptera","Chloropidae","Oscinellinae","Incertella, Rhopalopterum","Rhopalopterum umbrosum, Rhopalopterum luteiceps, Incertella bispina, Rhopalopterum painteri","1248","974","974","18","21.96","0","99.1","442.94","0","444.1","32.13","0"
"AGAW3","08E","INS","BOLD:AAH2293","Arthropoda","Insecta","Diptera","Asilidae","Stenopogoninae","Dioctria, Eudioctria","Dioctria hyalipennis, Dioctria baumhaueri, Eudioctria propinqua","1246","1199","1199","19","3.77","0","98.9","423.5","0","425.68","30.54","0"
"AGAW3","08E","INS","BOLD:AAB1098","Arthropoda","Insecta","Diptera","Culicidae","Culicinae","Ochlerotatus, Aedes","Aedes excrucians, Aedes cantans, Aedes stimulans, Aedes euedes, Aedes riparius, Aedes annulipes, Ochlerotatus cantans, Aedes fitchii, Aedes sp., Ochlerotatus annulipes, Ochlerotatus excrucians","1214","1048","1048","20","13.67","0","99.25","432.98","0","434.44","35.04","0"
"AGAW3","08E","INS","BOLD:AAA4201","Arthropoda","Insecta","Lepidoptera","Noctuidae","Noctuinae","Apamea","Apamea sp., Apamea sordens, Apamea sp. nr. sordens, Apamea digitula","1210","1003","1003","21","17.11","0","99.27","445.33","0","446.39","31.92","0"
"AGAW3","08E","INS","BOLD:AAF9771","Arthropoda","Insecta","Diptera","Hybotidae","Tachydromiinae","Platypalpus","Platypalpus longiseta, Platypalpus trivialis, Platypalpus sp. 5, Platypalpus sp. 17, Platypalpus pallidiventris","1171","873","873","22","25.45","0","99.13","444.4","0","445.09","34.99","0"
"AGAW3","08E","INS","BOLD:AAA5953","Arthropoda","Insecta","Lepidoptera","Gelechiidae","Gelechiinae","Coleotechnites","Coleotechnites sp., Coleotechnites piceaella, Coleotechnites laricis, Coleotechnites sp. 3, Coleotechnites carbonaria, Coleotechnites granti, Coleotechnites macleodi, Coleotechnites gibsonella, Coleotechnites coniferella, Coleotechnites martini","1146","867","867","23","24.35","0","99.38","443.18","0","444.51","34.17","0"
"AGAW3","08E","INS","BOLD:AAG2868","Arthropoda","Insecta","Hemiptera","Cicadellidae","Typhlocybinae","Empoasca","Empoasca sp., Empoasca fabae","975","657","657","24","32.62","0","99.49","441.93","0","447.46","32.33","0"
"AGAW3","08E","INS","BOLD:AAL7359","Arthropoda","Insecta","Diptera","Chironomidae","Chironominae","Tanytarsus","","867","774","774","25","10.73","0","99.23","436.88","0","438.91","36.12","0"
"AGAW3","08E","INS","BOLD:AAE7016","Arthropoda","Insecta","Lepidoptera","Gelechiidae","Dichomeridinae","Helcystogramma","Helcystogramma hystricella","825","773","773","26","6.3","0","99.29","420.12","0","422.11","30.9","0"
"AGAW3","08E","INS","BOLD:ACL8669","Arthropoda","Insecta","Lepidoptera","Psychidae","Psychinae","Psyche","Psyche casta","796","789","789","27","0.88","0","98.97","403.43","0","407.35","37.23","0"
"AGAW3","08E","INS","BOLD:AAD2041","Arthropoda","Insecta","Diptera","Dolichopodidae","Sciapodinae","Condylostylus","Condylostylus inermis, Condylostylus nigrofemoratus, Condylostylus connectans, Condylostylus caudatus, Condylostylus flavipes","683","523","523","28","23.43","0","99.47","440.18","0","441.43","35.93","0"
"AGAW3","08E","INS","BOLD:AAN5342","Arthropoda","Insecta","Diptera","Chironomidae","Orthocladiinae","Heleniella","","656","593","593","29","9.6","0","98.74","425.29","0","431.29","35.19","0"
"AGAW3","08E","INS","BOLD:AAD0484","Arthropoda","Insecta","Diptera","Chironomidae","","","","646","551","551","30","14.71","0","99.05","445.31","0","446.73","34.75","0"
"AGAW3","08E","INS","BOLD:AAH3661","Arthropoda","Insecta","Diptera","Cecidomyiidae","","","","627","549","549","31","12.44","0","99.3","434.36","0","436.03","25.48","0"
"AGAW3","08E","INS","BOLD:ABA6163","Arthropoda","Insecta","Psocodea","Caeciliusidae","Caeciliusinae","Valenzuela","Valenzuela piceus","556","447","447","32","19.6","0","99.51","427.31","0","429.1","36.64","0"
"AGAW3","08E","INS","BOLD:AAC0706","Arthropoda","Insecta","Diptera","Chironomidae","Chironominae","Dicrotendipes","Dicrotendipes tritomus","506","481","481","33","4.94","0","98.59","420.28","0","424.14","35.78","0"
"AGAW3","08E","INS","BOLD:AAH4135","Arthropoda","Insecta","Diptera","Chloropidae","Chloropinae","Thaumatomyia","Thaumatomyia glabra, Thaumatomyia grata, Thaumatomyia sp. 1 TAW","435","412","412","34","5.29","0","98.98","428.87","0","434.06","32.28","0"
"AGAW3","08E","INS","BOLD:AAP8739","Arthropoda","Insecta","Diptera","Platystomatidae","Platystomatinae","Rivellia","Rivellia steyskali","428","327","327","35","23.6","0","99.08","440.92","0","442.75","38.15","0"
"AGAW3","08E","INS","BOLD:AAN5332","Arthropoda","Insecta","Diptera","Chironomidae","Chironominae","Polypedilum","","417","385","385","36","7.67","0","97.99","426.4","0","429.09","33.91","0"
"AGAW3","08E","INS","BOLD:AAD1395","Arthropoda","Insecta","Diptera","Chironomidae","Chironominae","Polypedilum","Polypedilum convictum, Polypedilum flavum","378","321","321","37","15.08","0","98.38","444.06","0","446.17","33.2","0"
"AGAW3","08E","INS","BOLD:AAN6154","Arthropoda","Insecta","Coleoptera","Latridiidae","Corticariinae","Melanophthalma","Melanophthalma inermis","370","325","325","38","12.16","0","98.94","438.6","0","440","31.1","0"
"AGAW3","08E","INS","BOLD:ACF1538","Arthropoda","Insecta","Lepidoptera","Blastobasidae","Blastobasinae","Pigritia","","337","284","284","39","15.73","0","99.41","443.04","0","444.48","31.02","0"
"AGAW3","08E","INS","BOLD:ACE0554","Arthropoda","Insecta","Diptera","Phoridae","Metopininae","Megaselia","","307","269","269","40","12.38","0","99.44","442.17","0","444.26","31.98","0"
"AGAW3","08E","INS","BOLD:AAZ4462","Arthropoda","Insecta","Hymenoptera","Braconidae","Homolobinae","Homolobus","Homolobus truncator","287","228","228","41","20.56","0","99.35","444.75","0","446.32","25.51","0"
"AGAW3","08E","INS","BOLD:AAB5081","Arthropoda","Insecta","Hemiptera","Miridae","Mirinae","Leptopterna","Leptopterna dolabrata","282","274","274","42","2.84","0","99.1","407","0","408.39","39.39","0"
"AGAW3","08E","INS","BOLD:AAG7276","Arthropoda","Insecta","Diptera","Sphaeroceridae","Limosininae","Leptocera","Leptocera erythrocera","258","236","236","43","8.53","0","99.18","434.99","0","437.21","32.78","0"
"AGAW3","08E","INS","BOLD:AAL7819","Arthropoda","Insecta","Diptera","Psychodidae","Psychodinae","Psychoda","Psychoda sp., Psychoda sp. BOLD:AAL7819","257","236","236","44","8.17","0","99.32","415.85","0","417.55","32.6","0"
"AGAW3","08E","INS","BOLD:ABA0772","Arthropoda","Insecta","Diptera","Chironomidae","Chironominae","Polypedilum","","256","242","242","45","5.47","0","98.64","419.11","0","423.03","33.13","0"
"AGAW3","08E","INS","BOLD:AAH3644","Arthropoda","Insecta","Diptera","Cecidomyiidae","","","","235","202","202","46","14.04","0","99.2","439.73","0","441.33","27.18","0"
"AGAW3","08E","INS","BOLD:ACV3832","","","","","","","","234","202","202","47","13.68","0","98.63","396.29","0","427.53","35.56","0"
"AGAW3","08E","INS","BOLD:AAH2372","Arthropoda","Insecta","Coleoptera","Elateridae","Elaterinae","Dalopius","Dalopius pallidus","219","178","178","48","18.72","0","99.43","440.68","0","441.79","36.6","0"
"AGAW3","08E","INS","BOLD:AAB2384","Arthropoda","Insecta","Diptera","Syrphidae","Syrphinae","Eupeodes","Eupeodes duseki, Eupeodes volucris, Eupeodes nielseni, Eupeodes americanus, Eupeodes flukei, Eupeodes vockerothi, Eupeodes lucasi, Eupeodes montivagus, Eupeodes rufipunctatus, Eupeodes rojasi, Eupeodes goeldlini, Eupeodes luniger, Eupeodes punctifer, Eupeodes bucculatus, Eupeodes perplexus, Eupeodes abiskoensis, Eupeodes corollae, Eupeodes pomus, Eupeodes lundbecki, Eupeodes nitens, Eupeodes sp. BOLDAAB2384, Eupeodes curtus, Eupeodes latifasciatus","216","209","209","49","3.24","0","98.84","402.98","0","405.61","30.71","0"
"AGAW3","08E","INS","BOLD:AAE0707","Arthropoda","Insecta","Diptera","Chironomidae","Chironominae","Microtendipes","Microtendipes pedellus","204","191","191","50","6.37","0","97.76","437.53","0","440","31.43","0"
"AGAW3","08E","INS","BOLD:AAN5160","Arthropoda","Insecta","Diptera","Ceratopogonidae","Forcipomyiinae","Atrichopogon","","200","195","195","51","2.5","0","99.11","429.13","0","432.13","37.32","0"
"AGAW3","08E","INS","BOLD:AAA1525","Arthropoda","Insecta","Trichoptera","Leptoceridae","Leptocerinae","Oecetis","Oecetis inconspicua","198","161","161","52","18.69","0","98.66","448.04","0","448.98","32.2","0"
"AGAW3","08E","INS","BOLD:ACG6561","Arthropoda","Insecta","Diptera","Chironomidae","Orthocladiinae","Heterotrissocladius","","182","168","168","53","7.69","0","99.34","441.09","0","443.3","37.82","0"
"AGAW3","08E","INS","BOLD:ACP4736","Arthropoda","Insecta","Diptera","Chironomidae","Orthocladiinae","Smittia","Smittia sp. 8ES","176","165","165","54","6.25","0","99.07","425.61","0","427.77","32.98","0"
"AGAW3","08E","INS","BOLD:ACE6783","","","","","","","","173","144","144","55","16.76","0","99.53","419.23","0","420.94","25.91","0"
"AGAW3","08E","INS","BOLD:ACX5521","Arthropoda","Insecta","Diptera","Chironomidae","Orthocladiinae","","","168","167","167","56","0.6","0","98.79","410.92","0","413.14","34.97","0"
"AGAW3","08E","INS","BOLD:AAG4827","Arthropoda","Insecta","Diptera","Anthomyzidae","Anthomyzinae","Mumetopia","Mumetopia occipitalis","164","155","155","57","5.49","0","98.91","441.16","0","442.81","36.46","0"
"AGAW3","08E","INS","BOLD:AAN6406","Arthropoda","Insecta","Diptera","Sphaeroceridae","Limosininae","Opalimosina","Opalimosina mirabilis","163","151","151","58","7.36","0","99.04","427.64","0","431.5","32.14","0"
"AGAW3","08E","INS","BOLD:AAC9132","Arthropoda","Insecta","Diptera","Culicidae","Culicinae","Culiseta","Culiseta inornata","138","138","138","59","0","0","98.84","403.86","0","407.8","32.37","0"
"AGAW3","08E","INS","BOLD:AAH3653","Arthropoda","Insecta","Diptera","Cecidomyiidae","","","","122","108","108","60","11.48","0","99.18","437.54","0","438.85","26.03","0"
"AGAW3","08E","INS","BOLD:ABA7871","Arthropoda","Insecta","Diptera","Cecidomyiidae","","","","115","106","106","61","7.83","0","99.1","439.89","0","440.72","26.72","0"
"AGAW3","08E","INS","BOLD:AAV5575","Arthropoda","Insecta","Diptera","Cecidomyiidae","","","CecidInt30 sp. BOLD:AAV5575","100","80","80","62","20","0","99.44","444.06","0","444.99","25.97","0"
"AGAW3","08E","INS","BOLD:ACI1304","Arthropoda","Insecta","Diptera","Muscidae","Muscinae","Stomoxys","Stomoxys indicus","85","85","85","63","0","0","97.62","421.68","0","424.51","32.57","0"
"AGAW3","08E","INS","BOLD:ABY9689","Arthropoda","Insecta","Diptera","Chloropidae","Chloropinae","Thaumatomyia","","80","76","76","64","5","0","99.09","433.03","0","434.71","32.41","0"
"AGAW3","08E","INS","BOLD:AAU6533","Arthropoda","Insecta","Diptera","Phoridae","Metopininae","Megaselia","","79","73","73","65","7.59","0","99.77","355.82","0","440.38","32.17","0"
"AGAW3","08E","INS","BOLD:AAU7126","Arthropoda","Insecta","Coleoptera","Cleridae","Hydnocerinae","Isohydnocera","Isohydnocera curtipennis","78","64","64","66","17.95","0","99.29","439.62","0","441.19","32.74","0"
"AGAW3","08E","INS","BOLD:AAV1055","Arthropoda","Insecta","Diptera","Stratiomyidae","Beridinae","Allognosta","Allognosta fuscitarsis","78","64","64","66","17.95","0","98.35","442.23","0","443.47","35.76","0"
"AGAW3","08E","INS","BOLD:ACU7906","Arthropoda","Insecta","Diptera","Chironomidae","Orthocladiinae","","","77","77","77","68","0","0","98.97","407.4","0","410.34","36.06","0"
"AGAW3","08E","INS","BOLD:AAN5199","Arthropoda","Insecta","Diptera","Cecidomyiidae","","","CecidInt20 sp. BOLD:AAN5199","76","63","63","69","17.11","0","99.43","438.95","0","447.28","26.61","0"
"AGAW3","08E","INS","BOLD:AAF7638","Arthropoda","Insecta","Diptera","Chironomidae","Tanypodinae","Rheopelopia","Rheopelopia ornata","74","74","74","70","0","0","98.66","423.3","0","425.77","35.34","0"
"AGAW3","08E","INS","BOLD:AAG6934","Arthropoda","Insecta","Diptera","Hybotidae","","","","72","64","64","71","11.11","0","99.12","432.18","0","434.17","38.66","0"
"AGAW3","08E","INS","BOLD:ABY8233","Arthropoda","Insecta","Diptera","Tachinidae","Exoristinae","Oswaldia","Oswaldia conica","70","70","70","72","0","0","97.9","422.61","0","425.24","31.63","0"
"AGAW3","08E","INS","BOLD:ACG8783","Arthropoda","Insecta","Diptera","Chironomidae","Chironominae","Polypedilum","","67","47","47","73","29.85","0","99","447.54","0","448.49","33.52","0"
"AGAW3","08E","INS","BOLD:ABV3226","Arthropoda","Insecta","Diptera","Sphaeroceridae","Limosininae","Coproica","Coproica hirticula","67","63","63","73","5.97","0","98.98","446.39","0","448.04","35","0"
"AGAW3","08E","INS","BOLD:AAC4197","Arthropoda","Insecta","Diptera","Chironomidae","Orthocladiinae","Paraphaenocladius","Paraphaenocladius impensus","61","59","59","75","3.28","0","98.89","426.82","0","437.38","34.66","0"
"AGAW3","08E","INS","BOLD:AAP8999","Arthropoda","Insecta","Diptera","Chironomidae","Tanypodinae","","","60","59","59","76","1.67","0","98.75","432.17","0","433.65","35.98","0"
"AGAW3","08E","INS","BOLD:AAG5463","Arthropoda","Insecta","Diptera","Chironomidae","Chironominae","Tanytarsus","","59","58","58","77","1.69","0","97.95","428.19","0","433.58","35.49","0"
"AGAW3","08E","INS","BOLD:AAN5205","Arthropoda","Insecta","Diptera","Cecidomyiidae","","","CecidInt42 sp. BOLD:AAN5205","55","48","48","78","12.73","0","99.26","445.91","0","446.87","31.47","0"
"AGAW3","08E","INS","BOLD:AAV0162","Arthropoda","Insecta","Hemiptera","Cicadellidae","Deltocephalinae","Eutettix","Eutettix variabilis","53","46","46","79","13.21","0","99.76","364.45","0","434.83","36.6","0"
"AGAW3","08E","INS","BOLD:AAH0524","Arthropoda","Insecta","Hymenoptera","Ichneumonidae","Diplazontinae","Woldstedtius","Woldstedtius flavolineatus","52","52","52","80","0","0","99","413.52","0","415.88","27.17","0"
"AGAW3","08E","INS","BOLD:ACW5117","Arthropoda","Insecta","Diptera","Chironomidae","Orthocladiinae","Smittia","Smittia leucopogon","46","42","42","81","8.7","0","99.43","421.78","0","423.41","31.35","0"
"AGAW3","08E","INS","BOLD:AAN5230","Arthropoda","Insecta","Diptera","Cecidomyiidae","","","","46","46","46","81","0","0","99.14","426.78","0","428.37","26.02","0"
"AGAW3","08E","INS","BOLD:AAN8770","Arthropoda","Insecta","Diptera","Psychodidae","Psychodinae","Psychoda","Psychoda trinodulosa","46","45","45","81","2.17","0","99.32","442.07","0","442.91","33.57","0"
"AGAW3","08E","INS","BOLD:AAH0134","Arthropoda","Insecta","Coleoptera","Phalacridae","Phalacrinae","Stilbus","Stilbus apicialis","46","43","43","81","6.52","0","99.3","424.02","0","425.39","31.87","0"
"AGAW3","08E","INS","BOLD:AAE8368","Arthropoda","Insecta","Hymenoptera","Braconidae","Macrocentrinae","Macrocentrus","Macrocentrus sp.","45","43","43","85","4.44","0","98.97","428.22","0","432.69","24.01","0"
"AGAW3","08E","INS","BOLD:AAD4214","Arthropoda","Insecta","Hymenoptera","Ichneumonidae","Diplazontinae","Diplazon","Diplazon laetatorius","44","42","42","86","4.55","0","99.28","445.2","0","446.48","25.75","0"
"AGAW3","08E","INS","BOLD:AAP3003","Arthropoda","Insecta","Diptera","Chironomidae","Chironominae","","","44","43","43","86","2.27","0","97.43","415.25","0","416.95","33.8","0"
"AGAW3","08E","INS","BOLD:AAN5225","Arthropoda","Insecta","Diptera","Cecidomyiidae","","","","43","35","35","88","18.6","0","99.08","447.93","0","448.65","28.39","0"
"AGAW3","08E","INS","BOLD:AAD9052","Arthropoda","Insecta","Lepidoptera","Elachistidae","Elachistinae","Elachista","Elachista herbigrada, Elachista rianella, Elachista illectella, Elachista sp., Elachista tuorella","43","35","35","88","18.6","0","99.43","446.14","0","446.91","31.72","0"
"AGAW3","08E","INS","BOLD:AAN5183","Arthropoda","Insecta","Diptera","Cecidomyiidae","","","","38","35","35","90","7.89","0","98.34","447.5","0","448.34","27.77","0"
"AGAW3","08E","INS","BOLD:AAG8839","Arthropoda","Insecta","Hemiptera","Cicadellidae","Deltocephalinae","Errastunus, Latalus","Latalus ocellaris, Errastunus ocellaris","36","29","29","91","19.44","0","99.64","447.44","0","448.67","35.89","0"
"AGAW3","08E","INS","BOLD:ACE5127","Arthropoda","Insecta","Diptera","Tachinidae","Exoristinae","Oswaldia","Oswaldia anorbitalis, Oswaldia albifacies","35","34","34","92","2.86","0","97.59","422.57","0","424.6","31.82","0"
"AGAW3","08E","INS","BOLD:AAU6967","Arthropoda","Insecta","Coleoptera","Chrysomelidae","Galerucinae","Psylliodes","Psylliodes affinis","35","29","29","92","17.14","0","99.5","436.17","0","445.97","34.94","0"
"AGAW3","08E","INS","BOLD:AAG5192","Arthropoda","Insecta","Coleoptera","Curculionidae","Entiminae","Barypeithes","Barypeithes pellucidus","35","33","33","92","5.71","0","98.59","429.91","0","431.49","39.19","0"
"AGAW3","08E","INS","BOLD:AAN5309","Arthropoda","Insecta","Diptera","Chironomidae","Chironominae","","","33","30","30","95","9.09","0","97.39","433.42","0","434.79","36.12","0"
"AGAW3","08E","INS","BOLD:ACX2014","Arthropoda","Insecta","Diptera","Chironomidae","Chironominae","Tanytarsus","Tanytarsus mendax","33","33","33","95","0","0","97.48","385.82","0","402.15","35.89","0"
"AGAW3","08E","INS","BOLD:AAA4781","Arthropoda","Insecta","Hymenoptera","Braconidae","Microgastrinae","Glyptapanteles","Glyptapanteles carlossarmientoi","32","32","32","97","0","0","99.27","428.59","0","431.09","24.05","0"
"AGAW3","08E","INS","BOLD:AAG6432","Arthropoda","Insecta","Diptera","Ceratopogonidae","Forcipomyiinae","Atrichopogon","","31","27","27","98","12.9","0","98.33","437.94","0","439.23","34.16","0"
"AGAW3","08E","INS","BOLD:ACF3458","Arthropoda","Insecta","Hymenoptera","Ichneumonidae","Cryptinae","Cryptus","Cryptus rufovinctus","31","30","30","98","3.23","0","99.41","445.48","0","446.71","29.05","0"
"AGAW3","08E","INS","BOLD:AAG3275","Arthropoda","Insecta","Diptera","Phoridae","Metopininae","Megaselia","Megaselia minuta, Megaselia nubilipennis","31","30","30","98","3.23","0","99.21","430.23","0","431.55","34.42","0"
"AGAW3","08E","INS","BOLD:AAB1281","Arthropoda","Insecta","Hymenoptera","Ichneumonidae","Pimplinae","Itoplectis","Itoplectis sp., Itoplectis conquisitor","31","27","27","98","12.9","0","98.71","439.71","0","441.06","28.87","0"
"AGAW3","08E","INS","BOLD:AAM9254","Arthropoda","Insecta","Diptera","Sciaridae","","Bradysia","Bradysia pallipes","31","30","30","98","3.23","0","98.96","437","0","437.71","33.1","0"
"AGAW3","08E","INS","BOLD:AAG5457","Arthropoda","Insecta","Diptera","Chironomidae","Chironominae","","","30","28","28","103","6.67","0","98.63","437.47","0","440.8","34.35","0"
"AGAW3","08E","INS","BOLD:ACC8079","Arthropoda","Insecta","Diptera","Cecidomyiidae","","","","30","25","25","103","16.67","0","99.35","445.47","0","446.57","29.21","0"
"AGAW3","08E","INS","BOLD:AAH0152","Arthropoda","Insecta","Coleoptera","Cantharidae","Cantharinae","Cantharis","Cantharis rufa","30","29","29","103","3.33","0","98.6","433.07","0","436.93","35.2","0"
"AGAW3","08E","INS","BOLD:AAN5383","Arthropoda","Insecta","Diptera","Chironomidae","Chironominae","Dicrotendipes","","29","29","29","106","0","0","97.7","406.17","0","407.76","36.86","0"
"AGAW3","08E","INS","BOLD:AAM9347","Arthropoda","Insecta","Diptera","Phoridae","Metopininae","Megaselia","","29","28","28","106","3.45","0","97.91","397.62","0","400.07","32.99","0"
"AGAW3","08E","INS","BOLD:AAV5688","Arthropoda","Insecta","Diptera","Cecidomyiidae","","","","28","27","27","108","3.57","0","99.2","427.96","0","428.96","34.35","0"
"AGAW3","08E","INS","BOLD:AAG4905","Arthropoda","Insecta","Diptera","Mycetophilidae","Mycetophilinae","Mycetophila","","28","28","28","108","0","0","99.05","423.71","0","425.86","35.27","0"
"AGAW3","08E","INS","BOLD:AAN5242","Arthropoda","Insecta","Diptera","Cecidomyiidae","","","","26","26","26","110","0","0","99.17","409.42","0","411.08","28.29","0"
"AGAW3","08E","INS","BOLD:AAG9897","Arthropoda","Insecta","Dermaptera","Forficulidae","Forficulinae","Forficula","Forficula auricularia-A","26","26","26","110","0","0","98.99","394.46","0","404.88","45.73","0"
"AGAW3","08E","INS","BOLD:ACE9668","Arthropoda","Insecta","Hymenoptera","Ichneumonidae","Ophioninae","Enicospilus","Enicospilus purgatusDHJ02","24","24","24","112","0","0","99.08","442.67","0","443.67","25.85","0"
"AGAW3","08E","INS","BOLD:ABU5546","Arthropoda","Insecta","Diptera","Psychodidae","Psychodinae","Psychoda","","23","23","23","113","0","0","98.72","433.48","0","434.87","31.37","0"
"AGAW3","08E","INS","BOLD:AAV5573","Arthropoda","Insecta","Diptera","Cecidomyiidae","","","","23","22","22","113","4.35","0","98.99","440.96","0","442.39","27.55","0"
"AGAW3","08E","INS","BOLD:AAA3453","Arthropoda","Insecta","Diptera","Anthomyiidae","Anthomyiinae","Delia","","22","22","22","115","0","0","98.96","423.86","0","427.77","34.75","0"
"AGAW3","08E","INS","BOLD:AAL7815","Arthropoda","Insecta","Diptera","Psychodidae","Psychodinae","Psychoda","","22","22","22","115","0","0","97.94","422.41","0","424.5","34.48","0"
"AGAW3","08E","INS","BOLD:AAM6037","Arthropoda","Insecta","Diptera","Cecidomyiidae","Cecidomyiinae","Contarinia, Dasineura","","22","20","20","115","9.09","0","98.36","445.55","0","447.18","26.26","0"
"AGAW3","08E","INS","BOLD:AAU8556","Arthropoda","Insecta","Hymenoptera","Braconidae","Rogadinae","Aleiodes","","22","21","21","115","4.55","0","99.19","395.32","0","396.77","24.78","0"
"AGAW3","08E","INS","BOLD:ABA6514","Arthropoda","Insecta","Diptera","Chironomidae","Chironominae","Rheotanytarsus","","20","20","20","119","0","0","99.03","433.8","0","435.85","36.08","0"
"AGAW3","08E","INS","BOLD:AAD1485","Arthropoda","Insecta","Diptera","Chironomidae","Chironominae","Paratanytarsus","Paratanytarsus grimmii, Paratanytarsus sp.","20","17","17","119","15","0","99.63","443.75","0","444.85","33.79","0"
"AGAW3","08E","INS","BOLD:ADH4889","Arthropoda","Insecta","Diptera","","","","","20","18","18","119","10","0","99.04","441.05","0","442.4","35.29","0"
"AGAW3","08E","INS","BOLD:ACM7774","Arthropoda","Insecta","Trichoptera","Hydroptilidae","Orthotrichiinae","Orthotrichia","Orthotrichia cristata","20","20","20","119","0","0","98.96","407.85","0","410","29.15","0"
"AGAW3","08E","INS","BOLD:AAM9255","Arthropoda","Insecta","Diptera","Sciaridae","","Pseudolycoriella","","20","20","20","119","0","0","99.39","399.45","0","402.1","30.55","0"
"AGAW3","08E","INS","BOLD:ACX6055","Arthropoda","Insecta","Diptera","Phoridae","Metopininae","Megaselia","Megaselia sp. AK-2010-2, Megaselia sp. BOLD:ACX6055","19","19","19","124","0","0","99.51","436.37","0","438.53","33.19","0"
"AGAW3","08E","INS","BOLD:AAQ2557","Arthropoda","Insecta","Diptera","Sciaridae","","Peyerimhoffia, Cratyna","Cratyna vagabunda, Peyerimhoffia vagabunda","19","19","19","124","0","0","99.22","427.74","0","430.11","35.53","0"
"AGAW3","08E","INS","BOLD:AAN5238","Arthropoda","Insecta","Diptera","Cecidomyiidae","","","","19","18","18","124","5.26","0","99.39","432.68","0","434","25.55","0"
"AGAW3","08E","INS","BOLD:AAG7891","Arthropoda","Insecta","Hymenoptera","Platygastridae","Scelioninae","Telenomus","Telenomus podisi","19","18","18","124","5.26","0","98.87","431","0","440.84","27.25","0"
"AGAW3","08E","INS","BOLD:ABV1188","Arthropoda","Insecta","Diptera","Chironomidae","Chironominae","Tanytarsus","","19","19","19","124","0","0","98.15","418.26","0","423.05","33.74","0"
"AGAW3","08E","INS","BOLD:ABA0929","Arthropoda","Insecta","Diptera","Sciaridae","","Bradysia","Bradysia scabricornis","19","17","17","124","10.53","0","99.35","425.16","0","426.58","30.17","0"
"AGAW3","08E","INS","BOLD:AAP9021","Arthropoda","Insecta","Diptera","Cecidomyiidae","","","","19","19","19","124","0","0","99.46","408.21","0","410.84","29.98","0"
"AGAW3","08E","INS","BOLD:ABA0856","Arthropoda","Insecta","Diptera","Cecidomyiidae","","","","18","15","15","131","16.67","0","98.88","449.61","0","450","28.91","0"
"AGAW3","08E","INS","BOLD:AAG6952","Arthropoda","Insecta","Diptera","Chloropidae","Oscinellinae","Incertella","","18","16","16","131","11.11","0","99.06","448.28","0","449","31.5","0"
"AGAW3","08E","INS","BOLD:AAV5594","Arthropoda","Insecta","Diptera","Cecidomyiidae","","","","18","15","15","131","16.67","0","99.14","439.61","0","441.06","26.13","0"
"AGAW3","08E","INS","BOLD:AAF9891","Arthropoda","Insecta","Orthoptera","Gryllidae","Oecanthinae","Oecanthus","Oecanthus pini, Oecanthus nigricornis, Oecanthus celerinictus, Oecanthus quadripunctatus, Oecanthus argentinus","15","11","11","134","26.67","0","99.63","449.47","0","450","33.63","0"
"AGAW3","08E","INS","BOLD:ACJ0111","Arthropoda","Insecta","Hymenoptera","Platygastridae","Scelioninae","Telenomus","","15","13","13","134","13.33","0","98.78","430.47","0","443.8","28.83","0"
"AGAW3","08E","INS","BOLD:AAN5267","Arthropoda","Insecta","Diptera","Cecidomyiidae","","","","15","11","11","134","26.67","0","99.33","449.13","0","450","30.67","0"
"AGAW3","08E","INS","BOLD:ABA1215","Arthropoda","Insecta","Diptera","Sciaridae","","Lycoriella","Lycoriella sativae","14","14","14","137","0","0","99.17","447.64","0","449.14","35.92","0"
"AGAW3","08E","INS","BOLD:AAG3154","Arthropoda","Insecta","Diptera","Psychodidae","Psychodinae","Psychoda","","14","13","13","137","7.14","0","98.27","441.64","0","443.5","32.62","0"
"AGAW3","08E","INS","BOLD:AAH0135","Arthropoda","Insecta","Coleoptera","Phalacridae","Phalacrinae","Acylomus","Acylomus pugetanus, Acylomus ergoti","13","13","13","139","0","0","99.02","422.31","0","425.15","31.05","0"
"AGAW3","08E","INS","BOLD:AAM6248","Arthropoda","Insecta","Diptera","Chironomidae","Chironominae","Microtendipes","Microtendipes pedellus grp.","12","12","12","140","0","0","99.16","414.33","0","415.92","31.14","0"
"AGAW3","08E","INS","BOLD:AAN5255","Arthropoda","Insecta","Diptera","Cecidomyiidae","Cecidomyiinae","Dasineura","","11","10","10","141","9.09","0","98.56","422.64","0","446","26.81","0"
"AGAW3","08E","INS","BOLD:AAP7834","Arthropoda","Insecta","Coleoptera","Phalacridae","Phalacrinae","Stilbus","","10","10","10","142","0","0","99.07","434.7","0","435.4","32.98","0"
"AGAW3","08E","INS","BOLD:AAJ2399","Arthropoda","Insecta","Hemiptera","Miridae","Phylinae","Amblytylus","Amblytylus nasutus","10","9","9","142","10","0","99.91","449.1","0","450","39.82","0"
"AGAW3","08E","INS","BOLD:ABV2532","Arthropoda","Insecta","Hemiptera","Aleyrodidae","","","","10","9","9","142","10","0","99.71","430.8","0","433.2","37.17","0"
"AGAW3","08E","INS","BOLD:AAG2511","Arthropoda","Insecta","Diptera","Anthomyiidae","Anthomyiinae","Delia","Delia platura","10","10","10","142","0","0","99.06","447","0","449.2","33.33","0"
"AGAW3","08E","INS","BOLD:AAH1890","Arthropoda","Insecta","Hymenoptera","Ichneumonidae","Cryptinae","Stibeutes","Stibeutes brevicornis, Stibeutes heinemanni","10","10","10","142","0","0","99.47","443.5","0","444.5","29.6","0"
"AGAW3","08E","INS","BOLD:AAM7855","Arthropoda","Insecta","Thysanoptera","Thripidae","Thripinae","Frankliniella","Frankliniella tritici","9","8","8","147","11.11","0","99.45","447.89","0","450","31.41","0"
"AGAW3","08E","INS","BOLD:ABW6103","Arthropoda","Insecta","Diptera","Cecidomyiidae","","","","9","8","8","147","11.11","0","99.22","433.78","0","434.78","25.66","0"
"AGAW3","08E","INS","BOLD:AAV1261","Arthropoda","Insecta","Diptera","Sciaridae","","Bradysia","Bradysia splendida","9","9","9","147","0","0","99.12","428.22","0","430","34.08","0"
"AGAW3","08E","INS","BOLD:ACG0715","Arthropoda","Insecta","Diptera","Sphaeroceridae","Limosininae","Pullimosina","Pullimosina heteroneura","9","9","9","147","0","0","99.07","441.78","0","443.44","33.99","0"
"AGAW3","08E","INS","BOLD:ACH3153","Arthropoda","Insecta","Hymenoptera","Ichneumonidae","Diplazontinae","Woldstedtius","Woldstedtius biguttatus","8","8","8","151","0","0","99.05","404.25","0","406.25","28.37","0"
"AGAW3","08E","INS","BOLD:AAH3920","Arthropoda","Insecta","Diptera","Sciaridae","","Dichopygina, Scatopsciara","","8","8","8","151","0","0","98.67","413.38","0","445.63","35.5","0"
"AGAW3","08E","INS","BOLD:AAG6745","Arthropoda","Insecta","Diptera","Calliphoridae","Calliphorinae","Pollenia","Pollenia pediculata","8","8","8","151","0","0","99.08","409.25","0","411","33.87","0"
"AGAW3","08E","INS","BOLD:ACC8511","Arthropoda","Insecta","Diptera","Cecidomyiidae","Cecidomyiinae","Sitodiplosis","Sitodiplosis mosellana","7","7","7","154","0","0","98.69","418","0","438.71","27.06","0"
"AGAW3","08E","INS","BOLD:ACJ6954","Arthropoda","Insecta","Diptera","Cecidomyiidae","","","","7","7","7","154","0","0","99.49","338.14","0","422.71","26.42","0"
"AGAW3","08E","INS","BOLD:AAO7847","Arthropoda","Insecta","Hymenoptera","Braconidae","Microgastrinae","Choeras","","7","7","7","154","0","0","98.41","448.71","0","450","25.59","0"
"AGAW3","08E","INS","BOLD:ABU5520","Arthropoda","Insecta","Diptera","Sciaridae","","Corynoptera","","7","7","7","154","0","0","99.19","426","0","428.14","35.52","0"
"AGAW3","08E","INS","BOLD:ABU5525","Arthropoda","Insecta","Diptera","Chironomidae","Orthocladiinae","Limnophyes","Limnophyes sp. 14ES","7","7","7","154","0","0","99","402.14","0","404.14","37.29","0"
"AGAW3","08E","INS","BOLD:AAG3266","Arthropoda","Insecta","Diptera","Phoridae","Metopininae","Megaselia","","6","6","6","159","0","0","99.15","407","0","407","31.59","0"
"AGAW3","08E","INS","BOLD:ACK5742","Arthropoda","Insecta","Diptera","Cecidomyiidae","","","","5","4","4","160","20","0","99.69","445.8","0","445.8","29.29","0"
"AGAW3","08E","INS","BOLD:AAY6397","Arthropoda","Insecta","Diptera","Cecidomyiidae","","","","5","4","4","160","20","0","98.51","429.8","0","430.6","28.12","0"
"AGAW3","08E","INS","BOLD:ACX1262","Arthropoda","Insecta","Diptera","Tachinidae","Exoristinae","Vibrissina, Eribella","Eribella sp. 1, Vibrissina leibyi","5","5","5","160","0","0","97.51","408.6","0","410.6","32.05","0"
"AGAW3","08E","INS","BOLD:ACF1485","Arthropoda","Insecta","Hymenoptera","Platygastridae","Scelioninae","","","5","5","5","160","0","0","98.65","414","0","416.8","23.93","0"
"AGAW3","08E","INS","BOLD:ABA8048","Arthropoda","Insecta","Hymenoptera","Ichneumonidae","Cryptinae","","","5","5","5","160","0","0","99.38","438.4","0","439","29.37","0"
"AGAW3","08E","INS","BOLD:ACE3123","Arthropoda","Insecta","Diptera","Sciaridae","","Corynoptera","","5","5","5","160","0","0","97.84","443.8","0","445.4","34.89","0"
"AGAW3","08E","INS","BOLD:ABW7969","Arthropoda","Insecta","Diptera","Sciaridae","","Bradysia","Bradysia ocellaris","5","5","5","160","0","0","99.76","415.4","0","415.8","34.31","0"
"AGAW3","08E","INS","BOLD:AAU8237","Arthropoda","Insecta","Hymenoptera","Ichneumonidae","Ctenopelmatinae","","","4","4","4","167","0","0","98.98","422.75","0","424.5","28.07","0"
"AGAW3","08E","INS","BOLD:AAA3654","Arthropoda","Insecta","Lepidoptera","Crambidae","Crambinae","Crambus","Crambus bigelovi, Crambus johnsoni, Crambus praefectellus","4","4","4","167","0","0","98.88","444.75","0","446","31.56","0"
"AGAW3","08E","INS","BOLD:AAA6292","Arthropoda","Insecta","Lepidoptera","Tortricidae","Tortricinae","Cnephasia","Cnephasia genitalana, Cnephasia stephensiana","3","3","3","169","0","0","99.33","449.33","0","450","35.85","0"
"AGAW3","08E","INS","BOLD:AAA2632","Arthropoda","Insecta","Lepidoptera","Noctuidae","Noctuinae","Noctua","Noctua atlantica, Noctua pronuba","3","3","3","169","0","0","98.51","449","0","450","31.78","0"
"AGAW3","08E","INS","BOLD:ABA6271","Arthropoda","Insecta","Hymenoptera","Braconidae","Alysiinae","Dinotrema","","3","3","3","169","0","0","99.46","417.33","0","418","24.82","0"
"AGAW3","08E","INS","BOLD:AAA2815","Arthropoda","Insecta","Lepidoptera","Noctuidae","Noctuinae","Mythimna","Mythimna yukonensis, Mythimna oxygala","3","3","3","169","0","0","99.07","432.33","0","434","33.16","0"
"AGAW3","08E","INS","BOLD:AAN5260","Arthropoda","Insecta","Diptera","Cecidomyiidae","","","","3","3","3","169","0","0","98.1","404.67","0","406","25.7","0"
"AGAW3","08E","INS","BOLD:ABX2489","Arthropoda","Insecta","Coleoptera","Phalacridae","Phalacrinae","Stilbus","","3","3","3","169","0","0","97.69","392.67","0","393","32.64","0"
"AGAW3","08E","INS","BOLD:AAA6560","Arthropoda","Insecta","Lepidoptera","Crambidae","Crambinae","Crambus","Crambus multilinellus, Crambus saltuellus","2","2","2","175","0","0","99.89","449","0","450","31","0"
"AGAW3","08E","INS","BOLD:AAU6681","Arthropoda","Insecta","Diptera","Tipulidae","Tipulinae","Nephrotoma","Nephrotoma cornicina","2","2","2","175","0","0","99.44","450","0","450","36.44","0"
"AGAW3","08E","INS","BOLD:AAH1054","Arthropoda","Insecta","Hymenoptera","Braconidae","Microgastrinae","Choeras","","2","2","2","175","0","0","99.88","418","0","450","25.89","0"
"AGAW3","08E","INS","BOLD:ACD1218","Arthropoda","Insecta","Diptera","Sciaridae","","Leptosciarella","Leptosciarella scutellata","2","2","2","175","0","0","99.75","403.5","0","405","33.7","0"
"AGAW3","08E","INS","BOLD:AAI5897","Arthropoda","Insecta","Lepidoptera","Gelechiidae","Gelechiinae","Coleotechnites","","2","2","2","175","0","0","97.34","394.5","0","395","33.04","0"
"AGAW3","08E","INS","BOLD:ADF6661","Arthropoda","Insecta","Diptera","Cecidomyiidae","","","","2","2","2","175","0","0","98.81","418.5","0","448","29.8","0"
"AGAW3","08E","INS","BOLD:ACF2404","Arthropoda","Insecta","Diptera","Tachinidae","Exoristinae","Oswaldia","Oswaldia aurifrons","2","2","2","175","0","0","97.86","443","0","447","31.88","0"
"AGAW3","08E","INS","BOLD:ABV1297","Arthropoda","Insecta","Diptera","Sciaridae","","","","2","2","2","175","0","0","100","389","0","390.5","29.85","0"
"AGAW3","08E","INS","BOLD:AAA9246","Arthropoda","Insecta","Lepidoptera","Tortricidae","Tortricinae","Clepsis","Clepsis clemensiana","2","2","2","175","0","0","97.98","386","0","388.5","29.62","0"
"AGAW3","08E","INS","BOLD:ABW7897","Arthropoda","Insecta","Diptera","Cecidomyiidae","","","","2","2","2","175","0","0","98.33","449","0","450","30.22","0"
"AGAW3","08E","INS","BOLD:ADH7839","Arthropoda","Insecta","Diptera","","","","","2","2","2","175","0","0","98.49","430","0","432.5","25.89","0"
"AGAW3","08E","INS","BOLD:ABV2748","Arthropoda","Insecta","Hymenoptera","Braconidae","Alysiinae","","","2","2","2","175","0","0","99.53","407.5","0","410","28.44","0"
"AGAW3","08E","INS","BOLD:AAG5242","Arthropoda","Insecta","Coleoptera","Curculionidae","Entiminae","Barypeithes","Barypeithes pellucidus","2","2","2","175","0","0","97.95","437","0","441.5","39.86","0"
"AGAW3","08E","INS","BOLD:AAA3774","Arthropoda","Insecta","Diptera","Chironomidae","Chironominae","Tanytarsus","Tanytarsus mendax","2","2","2","175","0","0","97.25","363","0","365.5","35.26","0"
"AGAW3","08E","INS","BOLD:ABX0657","Arthropoda","Insecta","Coleoptera","Anthicidae","Notoxinae","Notoxus","Notoxus desertus","2","2","2","175","0","0","99.4","415","0","418","36.35","0"
"AGAW3","08E","INS","BOLD:ADH8884","","","","","","","","2","2","2","175","0","0","98.42","376","0","379","33.11","0"
"AGAW3","08E","INS","BOLD:AAN5355","Arthropoda","Insecta","Diptera","Chironomidae","Orthocladiinae","Smittia","Smittia cf. stercoraria, Smittia stercoraria","2","2","2","175","0","0","99.87","389","0","450","36.44","0"
"AGAW3","08E","INS","BOLD:ABW3687","Arthropoda","Insecta","Diptera","Lauxaniidae","Homoneurinae","Homoneura","Homoneura bispina","1","1","1","192","0","0","98.22","450","0","450","34.44","0"
"AGAW3","08E","INS","BOLD:AAH1693","Arthropoda","Insecta","Hymenoptera","Ichneumonidae","Cryptinae","Cryptus","Cryptus albitarsis","1","1","1","192","0","0","97.07","376","0","378","28.84","0"
"AGAW3","08E","INS","BOLD:AAH2868","Arthropoda","Insecta","Hemiptera","Aphididae","Drepanosiphinae","Drepanaphis","Drepanaphis acerifoliae","1","1","1","192","0","0","99.78","450","0","450","30","0"
"AGAW3","08E","INS","BOLD:AAY6730","Arthropoda","Insecta","Hemiptera","Derbidae","Cedusinae","Cedusa","Cedusa incisa","1","1","1","192","0","0","99.48","387","0","389","34.7","0"
"AGAW3","08E","INS","BOLD:AAC8881","Arthropoda","Insecta","Lepidoptera","Gelechiidae","Gelechiinae","Syncopacma","Syncopacma palpilineella","1","1","1","192","0","0","100","403","0","406","32.02","0"
"AGAW3","08E","INS","BOLD:AAG1748","Arthropoda","Insecta","Diptera","Muscidae","Coenosiinae","Coenosia","","1","1","1","192","0","0","99.11","450","0","450","34.44","0"
"AGAW3","08E","INS","BOLD:AAA4751","Arthropoda","Insecta","Diptera","Culicidae","Culicinae","Culex","Culex pipiens sl, Culex nr. pipiens, Culex pipiens, Culex quinquefasciatus, Culex sp.","1","1","1","192","0","0","99.54","435","0","441","32.88","0"
"AGAW3","08E","INS","BOLD:AAM6781","Arthropoda","Insecta","Diptera","Dolichopodidae","Neurigoninae","Neurigona","Neurigona tenuis","1","1","1","192","0","0","98.54","412","0","414","35.27","0"
"AGAW3","08E","INS","BOLD:AAO8538","Arthropoda","Insecta","Diptera","Psychodidae","Psychodinae","Psychoda","","1","1","1","192","0","0","98.02","404","0","405","30.86","0"
"AGAW3","08E","INS","BOLD:AAM7886","Arthropoda","Insecta","Diptera","Tachinidae","Exoristinae","Oswaldia","","1","1","1","192","0","0","97.12","416","0","421","30.88","0"
"AGAW3","08E","INS","BOLD:AAB4657","Arthropoda","Insecta","Diptera","Chironomidae","Chironominae","Chironomus","Chironomus maturus","1","1","1","192","0","0","99.33","450","0","450","37.56","0"
"AGAW3","08E","INS","BOLD:ABX9225","Arthropoda","Insecta","Diptera","Limoniidae","Limoniinae","Rhipidia","","1","1","1","192","0","0","99.33","450","0","450","33.56","0"
"AGAW3","08E","INS","BOLD:AAH3914","Arthropoda","Insecta","Diptera","Sciaridae","","Sciara","Sciara hamatilis, Sciara humeralis","1","1","1","192","0","0","99.56","450","0","450","30.67","0"
"AGAW3","08E","INS","BOLD:ACL7849","","","","","","","","1","1","1","192","0","0","100","377","0","450","33.78","0"
"AGAW3","08E","INS","BOLD:AAB7869","Arthropoda","Insecta","Lepidoptera","Tortricidae","Tortricinae","Phtheochroa","Phtheochroa baracana, Phtheochroa riscana, Phtheochroa birdana, Phtheochroa fulviplicana, Phtheochroa canariana","1","1","1","192","0","0","97.38","382","0","383","28.72","0"
"AGAW3","08E","INS","BOLD:AAA5299","Arthropoda","Insecta","Diptera","Chironomidae","Orthocladiinae","Cricotopus","Cricotopus laetus, Cricotopus relucens, Cricotopus trifasciatus, Cricotopus sp., Cricotopus sp. 3ES, Cricotopus sylvestris, Cricotopus glacialis","1","1","1","192","0","0","99.06","426","0","428","35.28","0"
"AGAW3","08E","INS","BOLD:AAD5253","Arthropoda","Insecta","Diptera","Chironomidae","Orthocladiinae","Thienemanniella","Thienemanniella xena, Thienemanniella vittata","1","1","1","192","0","0","98.88","445","0","447","34.23","0"
"AGAW3","08E","INS","BOLD:AAB8013","Arthropoda","Insecta","Coleoptera","Coccinellidae","Coccinellinae","Hippodamia","Hippodamia expurgata, Hippodamia parenthesis, Hippodamia apicalis","1","1","1","192","0","0","99.75","404","0","405","33.83","0"
"AGAW3","08E","INS","BOLD:AAA2482","Arthropoda","Insecta","Lepidoptera","Noctuidae","Noctuinae","Mythimna","Mythimna unipuncta","1","1","1","192","0","0","99.11","450","0","450","30.89","0"
"AGAW3","08E","INS","BOLD:AAB9140","Arthropoda","Insecta","Diptera","Calliphoridae","Chrysomyinae","Phormia","Phormia regina","1","1","1","192","0","0","100","450","0","450","33.11","0"
"AGAW3","08E","INS","BOLD:ACV5892","Arthropoda","Insecta","Diptera","Cecidomyiidae","","","","1","1","1","192","0","0","99.78","447","0","450","24.89","0"
"AGAW3","08E","INS","BOLD:AAP3813","Arthropoda","Insecta","Diptera","Chloropidae","Oscinellinae","Rhopalopterum","Rhopalopterum carbonarium","1","1","1","192","0","0","97.33","450","0","450","32","0"
"AGAW3","08E","INS","BOLD:ACU6062","Arthropoda","Insecta","Diptera","Cecidomyiidae","","","","1","1","1","192","0","0","99.33","447","0","450","26.44","0"
"AGAW3","08E","INS","BOLD:AAG5499","Arthropoda","Insecta","Diptera","Chironomidae","Orthocladiinae","Psectrocladius","Psectrocladius obvius, Psectrocladius platypus","1","1","1","192","0","0","97.76","446","0","450","36.22","0"
"AGAW3","08E","INS","BOLD:AAH0256","Arthropoda","Insecta","Coleoptera","Latridiidae","Corticariinae","Corticaria, Corticarina","","1","1","1","192","0","0","98.62","434","0","435","32.64","0"
"AGAW3","08E","INS","BOLD:AAC5848","Arthropoda","Insecta","Lepidoptera","Psychidae","Psychinae","Psyche","Psyche casta","1","1","1","192","0","0","98.73","394","0","397","38.29","0"
"AGAW3","08E","INS","BOLD:ABV9290","Arthropoda","Insecta","Diptera","Cecidomyiidae","","","","1","1","1","192","0","0","99.11","450","0","450","26","0"
"AGAW3","08E","INS","BOLD:AAG9626","Arthropoda","Insecta","Diptera","Dolichopodidae","","","","1","1","1","192","0","0","97.33","449","0","450","35.56","0"
"AGAW3","08E","INS","BOLD:ABW1379","Arthropoda","Insecta","Diptera","Chloropidae","Oscinellinae","Malloewia","Malloewia abdominalis","1","1","1","192","0","0","97.54","448","0","450","31.78","0"
"AGAW3","08E","INS","BOLD:AAA3867","Arthropoda","Insecta","Lepidoptera","Noctuidae","Noctuinae","Apamea","Apamea amputatrix","1","1","1","192","0","0","99.55","449","0","450","34","0"
"AGAW3","08E","INS","BOLD:ACZ5284","Arthropoda","Insecta","Diptera","Sciomyzidae","Sciomyzinae","Dictya","Dictya stricta, Dictya expansa, Dictya floridensis, Dictya borealis, Dictya gaigei, Dictya sp. TAW1, Dictya umbroides, Dictya texensis, Dictya steyskali, Dictya pictipes","1","1","1","192","0","0","97.09","447","0","450","35.11","0"
"AGAW3","08E","INS","BOLD:AAA4555","Arthropoda","Insecta","Orthoptera","Acrididae","Melanoplinae","Melanoplus","Melanoplus stupefactus, Melanoplus angustipennis, Melanoplus ponderosus, Melanoplus bruneri, Melanoplus frigidus, Melanoplus scapularis, Melanoplus cinereus, Melanoplus madeleineae, Melanoplus sanguinipes, Melanoplus nigrescens group, Melanoplus tunicae, Melanoplus walshi, Melanoplus femurrubrum, Melanoplus dawsoni, Melanoplus sp., Melanoplus fasciatus, Melanoplus bowditchi, Melanoplus stonei, Melanoplus bivittatus, Melanoplus huroni, Melanoplus thomasi, Melanoplus borealis, Melanoplus packardii","1","1","1","192","0","0","98.2","389","0","388","35.82","0"
"AGAW3","08E","INS","BOLD:AAF8971","Arthropoda","Insecta","Diptera","Limoniidae","","","","1","1","1","192","0","0","99.33","448","0","450","36","0"
"AGAW3","08E","INS","BOLD:AAE0845","Arthropoda","Insecta","Hemiptera","Pentatomidae","Pentatominae","Euschistus","Euschistus servus","1","1","1","192","0","0","98.66","449","0","450","37.78","0"
"AGAW3","08E","INS","BOLD:AAV5352","Arthropoda","Insecta","Diptera","Cecidomyiidae","","","","1","1","1","192","0","0","99.78","450","0","450","27.11","0"
"AGAW3","08E","INS","BOLD:AAU6656","Arthropoda","Insecta","Diptera","Limoniidae","Chioneinae","Neolimnophila","Neolimnophila placida","1","1","1","192","0","0","98","450","0","450","34.67","0"
"AGAW3","08E","INS","BOLD:ABX6186","Arthropoda","Insecta","Diptera","Tipulidae","Tipulinae","Nephrotoma","Nephrotoma ferruginea","1","1","1","192","0","0","98","449","0","450","34.44","0"
"AGAW3","08E","INS","BOLD:ADH4973","Arthropoda","Insecta","Diptera","Tipulidae","Tipulinae","Nephrotoma","","1","1","1","192","0","0","98.44","450","0","450","33.78","0"
"AGAW3","08E","INS","BOLD:ABZ5558","","","","","","","","1","1","1","192","0","0","97.04","405","0","409","27.63","0"
"AGAW3","08E","INS","BOLD:ABV0488","Arthropoda","Insecta","Diptera","Cecidomyiidae","","","","1","1","1","192","0","0","99.03","414","0","415","26.75","0"
"AGAW3","08E","INS","BOLD:ABX0893","Arthropoda","Insecta","Diptera","Cecidomyiidae","","","","1","1","1","192","0","0","97.33","449","0","450","26.89","0"
"AGAW3","08E","INS","BOLD:AAL4232","Arthropoda","Insecta","Diptera","Agromyzidae","Phytomyzinae","Phytoliriomyza","Phytoliriomyza arctica","1","1","1","192","0","0","99.11","448","0","450","32","0"
"AGAW3","08E","INS","BOLD:ACB0984","Arthropoda","Insecta","Psocodea","Stenopsocidae","","Graphopsocus","Graphopsocus cruciatus","1","1","1","192","0","0","99.27","413","0","450","37.78","0"
"AGAW3","08E","INS","BOLD:AAK8756","Arthropoda","Insecta","Diptera","Agromyzidae","Phytomyzinae","Liriomyza","Liriomyza sp. 2 AGT, Liriomyza fricki","1","1","1","192","0","0","99.48","382","0","384","31.25","0"
"AGAW3","08E","INS","BOLD:ACP1329","Arthropoda","Insecta","Hemiptera","Cicadellidae","Macropsinae","Macropsis","","1","1","1","192","0","0","99.33","449","0","450","32.44","0"
"AGAW3","08E","INS","BOLD:ACE1806","Arthropoda","Insecta","Diptera","Camillidae","","Camilla","Camilla atrimana","1","1","1","192","0","0","97.77","449","0","450","36.44","0"
"AGAW3","08E","INS","BOLD:ADH7661","Arthropoda","Insecta","Lepidoptera","","","","","1","1","1","192","0","0","97.3","445","0","450","34.22","0"
"AGAW3","08E","NAR","no match","no match","no match","no match","no match","no match","no match","no match","no match","no match","no match","no match","no match","no match","no match","no match","no match","no match","no match","no match"
"AGAW3","08E","NIN","BOLD:AAD1564","Arthropoda","Arachnida","Araneae","Clubionidae","","Clubiona","Clubiona abboti","940","859","859","1","8.62","0","98.17","426.42","0","429.24","31.55","0"
"AGAW3","08E","NIN","BOLD:ABW2639","Arthropoda","Arachnida","Trombidiformes","Anystidae","","","","803","667","667","2","16.94","0","99.34","446.04","0","445.73","31.29","0"
"AGAW3","08E","NIN","BOLD:AAA6381","Arthropoda","Arachnida","Araneae","Tetragnathidae","","Tetragnatha","Tetragnatha laboriosa","718","680","680","3","5.29","0","99.23","428.46","0","430.71","39.41","0"
"AGAW3","08E","NIN","BOLD:AAB7914","Arthropoda","Collembola","Symphypleona","Bourletiellidae","","Bourletiella","","687","537","537","4","21.83","0","99.56","444.38","0","445.78","38.13","0"
"AGAW3","08E","NIN","BOLD:AAA7246","Arthropoda","Collembola","Entomobryomorpha","Entomobryidae","Entomobryinae","Entomobrya","Entomobrya nicoleti, Entomobrya multifasciata, Entomobrya intermedia","402","310","310","5","22.89","0","99.17","442.31","0","443.42","43.42","0"
"AGAW3","08E","NIN","BOLD:AAB0726","Arthropoda","Arachnida","Araneae","Lycosidae","","Trochosa","Trochosa ruricola","179","176","176","6","1.68","0","98.76","437.68","0","440.71","31.12","0"
"AGAW3","08E","NIN","BOLD:AAH6630","Arthropoda","Collembola","Entomobryomorpha","Entomobryidae","Entomobryinae","Entomobrya","Entomobrya unostrigata","152","130","130","7","14.47","0","99.41","438.9","0","440.02","39.01","0"
"AGAW3","08E","NIN","BOLD:AAA5267","Arthropoda","Collembola","Entomobryomorpha","Isotomidae","","","","141","141","141","8","0","0","99.09","419.07","0","421.82","38.12","0"
"AGAW3","08E","NIN","BOLD:ACZ1295","Arthropoda","Arachnida","Trombidiformes","Anystidae","","","","117","116","116","9","0.85","0","99.14","374.85","0","383.91","31.1","0"
"AGAW3","08E","NIN","BOLD:ACC0359","Arthropoda","Collembola","Symphypleona","Bourletiellidae","","Bourletiella","","80","65","65","10","18.75","0","99.48","442.18","0","445.46","37.67","0"
"AGAW3","08E","NIN","BOLD:ACI9908","Arthropoda","Collembola","Symphypleona","Bourletiellidae","","","","52","41","41","11","21.15","0","99.51","438.5","0","440.12","35.04","0"
"AGAW3","08E","NIN","BOLD:ACV5609","","","","","","","","49","45","45","12","8.16","0","98.94","446.29","0","447.57","38.79","0"
"AGAW3","08E","NIN","BOLD:ACJ7454","Arthropoda","Collembola","Symphypleona","Bourletiellidae","","","","31","27","27","13","12.9","0","99.67","402.48","0","446.42","38.97","0"
"AGAW3","08E","NIN","BOLD:AAM7961","Arthropoda","Arachnida","Trombidiformes","Anystidae","","","","29","28","28","14","3.45","0","99.23","442.45","0","442.17","31.41","0"
"AGAW3","08E","NIN","BOLD:AAB8452","Arthropoda","Collembola","Entomobryomorpha","Entomobryidae","Lepidocyrtinae","Lepidocyrtus","Lepidocyrtus paradoxus","15","13","13","15","13.33","0","99.51","447.53","0","443.73","42.09","0"
"AGAW3","08E","NIN","BOLD:AAN4465","","","","","","","","14","14","14","16","0","0","99.35","437.07","0","438.21","46.15","0"
"AGAW3","08E","NIN","BOLD:ACV5610","","","","","","","","14","10","10","16","28.57","0","99.69","447.43","0","448.71","38.09","0"
"AGAW3","08E","NIN","BOLD:AAZ2181","Arthropoda","Collembola","Symphypleona","Bourletiellidae","","","","13","12","12","18","7.69","0","97.38","448.46","0","450","39.42","0"
"AGAW3","08E","NIN","BOLD:AAC9042","Arthropoda","Arachnida","Araneae","Salticidae","","Salticus","Salticus scenicus","8","8","8","19","0","0","99.3","446","0","447.5","33.24","0"
"AGAW3","08E","NIN","BOLD:ACR6863","","","","","","","","3","3","3","20","0","0","98.31","433.67","0","433.67","38.88","0"
"AGAW3","08E","NIN","BOLD:AAF9285","Arthropoda","Arachnida","Mesostigmata","Parasitidae","","","","2","2","2","21","0","0","99.18","423.5","0","426","34.25","0"
"AGAW3","08E","NIN","BOLD:ACP9010","Arthropoda","Collembola","Symphypleona","Bourletiellidae","","","","2","2","2","21","0","0","97.99","448.5","0","450","38.78","0"
"AGAW3","08E","NIN","BOLD:ACR6862","","","","","","","","1","1","1","23","0","0","99.11","450","0","450","39.33","0"
"AGAW3","08N","BAC","TAX:175768","Proteobacteria","Alphaproteobacteria","Rickettsiales","Rickettsiaceae",NA,"Wolbachia","Wolbachia pipientis","127","109","109","1","14.17","0","98.87","443.8","0","445.28","39.52","0"
"AGAW3","08N","BAC","TAX:91393","Proteobacteria","Alphaproteobacteria","Rickettsiales","Rickettsiaceae",NA,"Wolbachia","","96","96","96","2","0","0","99.48","390.25","0","420.56","39.12","0"
"AGAW3","08N","BAC","TAX:91391","Proteobacteria","Alphaproteobacteria","Rickettsiales","",NA,"","","2","2","2","3","0","0","98.34","421.5","0","450","40","0"
"AGAW3","08N","BAC","TAX:392851","Proteobacteria","Alphaproteobacteria","Rickettsiales","Rickettsiaceae",NA,"Wolbachia","Wolbachia endosymbiont of Formicidae","1","1","1","4","0","0","98.65","443","0","450","42","0"
"AGAW3","08N","BAC","TAX:392909","Proteobacteria","Alphaproteobacteria","Rickettsiales","Rickettsiaceae",NA,"Wolbachia","Wolbachia endosymbiont of Microgastrinae","1","1","1","4","0","0","99.55","446","0","450","41.11","0"
"AGAW3","08N","BAC","TAX:91389","Proteobacteria","","","",NA,"","","1","1","1","4","0","0","97.78","361","0","366","37.7","0"
"AGAW3","08N","CHO","no match","no match","no match","no match","no match","no match","no match","no match","no match","no match","no match","no match","no match","no match","no match","no match","no match","no match","no match","no match"
"AGAW3","08N","INS","BOLD:AAA9246","Arthropoda","Insecta","Lepidoptera","Tortricidae","Tortricinae","Clepsis","Clepsis clemensiana","19452","16145","16145","1","17","0","99.36","427.55","0","429.32","30.2","0"
"AGAW3","08N","INS","BOLD:AAA6292","Arthropoda","Insecta","Lepidoptera","Tortricidae","Tortricinae","Cnephasia","Cnephasia genitalana, Cnephasia stephensiana","10223","6974","6974","2","31.78","0","99.53","438.78","0","439.94","35.41","0"
"AGAW3","08N","INS","BOLD:AAC0903","Arthropoda","Insecta","Diptera","Chironomidae","Chironominae","Chironomus","Chironomus acidophilus","6766","5131","5131","3","24.16","0","99.38","439.05","0","441.96","36.32","0"
"AGAW3","08N","INS","BOLD:AAG1711","Arthropoda","Insecta","Diptera","Muscidae","Phaoniinae","Helina","Helina depuncta","6693","5367","5367","4","19.81","0","99.36","433.22","0","435.49","31.37","0"
"AGAW3","08N","INS","BOLD:AAA3867","Arthropoda","Insecta","Lepidoptera","Noctuidae","Noctuinae","Apamea","Apamea amputatrix","5092","3919","3919","5","23.04","0","99.23","440.27","0","442.12","33.72","0"
"AGAW3","08N","INS","BOLD:AAI4303","Arthropoda","Insecta","Diptera","Chironomidae","Chironominae","Chironomus","Chironomus melanescens","4305","3916","3916","6","9.04","0","98.21","429.23","0","434.69","33.74","0"
"AGAW3","08N","INS","BOLD:AAA4201","Arthropoda","Insecta","Lepidoptera","Noctuidae","Noctuinae","Apamea","Apamea sp., Apamea sordens, Apamea sp. nr. sordens, Apamea digitula","3092","2383","2383","7","22.93","0","99.24","446.14","0","447.15","31.94","0"
"AGAW3","08N","INS","BOLD:AAG6745","Arthropoda","Insecta","Diptera","Calliphoridae","Calliphorinae","Pollenia","Pollenia pediculata","2853","2589","2589","8","9.25","0","99.3","418.49","0","420.46","33.5","0"
"AGAW3","08N","INS","BOLD:AAG5507","Arthropoda","Insecta","Diptera","Chironomidae","Chironominae","Chironomus","Chironomus atrella","2833","2725","2725","9","3.81","0","99.07","420.66","0","429.63","35.91","0"
"AGAW3","08N","INS","BOLD:ABW3687","Arthropoda","Insecta","Diptera","Lauxaniidae","Homoneurinae","Homoneura","Homoneura bispina","2379","1658","1658","10","30.31","0","98.58","441.07","0","442.15","34.86","0"
"AGAW3","08N","INS","BOLD:ACL8669","Arthropoda","Insecta","Lepidoptera","Psychidae","Psychinae","Psyche","Psyche casta","1910","1887","1887","11","1.2","0","98.99","404.38","0","408.55","37.19","0"
"AGAW3","08N","INS","BOLD:AAA3889","Arthropoda","Insecta","Lepidoptera","Crambidae","Crambinae","Parapediasia","Parapediasia teterrellus","1340","1210","1210","12","9.7","0","99.23","429.3","0","430.93","32.15","0"
"AGAW3","08N","INS","BOLD:ABV9293","Arthropoda","Insecta","Diptera","Chironomidae","Chironominae","Dicrotendipes","","1225","966","966","13","21.14","0","99.67","355.79","0","436.68","34.8","0"
"AGAW3","08N","INS","BOLD:AAC0635","Arthropoda","Insecta","Hemiptera","Miridae","Mirinae","Stenotus","Stenotus binotatus","1167","870","870","14","25.45","0","99.15","442.53","0","443.47","34.71","0"
"AGAW3","08N","INS","BOLD:ABY5318","Arthropoda","Insecta","Diptera","Chloropidae","Oscinellinae","Incertella, Rhopalopterum","Rhopalopterum umbrosum, Rhopalopterum luteiceps, Incertella bispina, Rhopalopterum painteri","1013","771","771","15","23.89","0","98.56","442.9","0","443.91","32.24","0"
"AGAW3","08N","INS","BOLD:AAB5081","Arthropoda","Insecta","Hemiptera","Miridae","Mirinae","Leptopterna","Leptopterna dolabrata","923","883","883","16","4.33","0","99.12","405.01","0","406.31","39.38","0"
"AGAW3","08N","INS","BOLD:ABA6163","Arthropoda","Insecta","Psocodea","Caeciliusidae","Caeciliusinae","Valenzuela","Valenzuela piceus","812","692","692","17","14.78","0","99.45","425.15","0","427.09","36.61","0"
"AGAW3","08N","INS","BOLD:ACI9753","Arthropoda","Insecta","Diptera","Dolichopodidae","Diaphorinae","Chrysotus","","777","709","709","18","8.75","0","99.35","405.09","0","433.31","32.19","0"
"AGAW3","08N","INS","BOLD:AAA7661","Arthropoda","Insecta","Diptera","Culicidae","Culicinae","Culiseta, Culex","Culiseta melanura, Culex articularis, Culex restuans, Culex acharistus","774","711","711","19","8.14","0","99.21","433.46","0","435.51","32.91","0"
"AGAW3","08N","INS","BOLD:AAC1312","Arthropoda","Insecta","Diptera","Syrphidae","Syrphinae","Toxomerus","Toxomerus geminatus, Toxomerus 75-5, Toxomerus rombicus, Toxomerus sp. 75-5","744","593","593","20","20.3","0","99.42","437.23","0","439.3","30.05","0"
"AGAW3","08N","INS","BOLD:AAN5341","Arthropoda","Insecta","Diptera","Chironomidae","Orthocladiinae","Camptocladius","Camptocladius stercorarius","683","647","647","21","5.27","0","98.58","407.29","0","410.52","35.54","0"
"AGAW3","08N","INS","BOLD:AAA2636","Arthropoda","Insecta","Lepidoptera","Noctuidae","Noctuinae","Lacinipolia","Lacinipolia renigera","619","538","538","22","13.09","0","99.32","438.33","0","439.57","34.19","0"
"AGAW3","08N","INS","BOLD:AAU5981","Arthropoda","Insecta","Diptera","Pipunculidae","Pipunculinae","Pipunculus","Pipunculus sp. ON15","596","576","576","23","3.36","0","98.63","415.79","0","419.34","33.46","0"
"AGAW3","08N","INS","BOLD:ACK7314","Arthropoda","Insecta","Diptera","Chironomidae","Chironominae","Polypedilum","","445","439","439","24","1.35","0","98.49","386.65","0","392.04","35.13","0"
"AGAW3","08N","INS","BOLD:AAA2482","Arthropoda","Insecta","Lepidoptera","Noctuidae","Noctuinae","Mythimna","Mythimna unipuncta","440","408","408","25","7.27","0","99.09","416.8","0","420","30.16","0"
"AGAW3","08N","INS","BOLD:AAG2868","Arthropoda","Insecta","Hemiptera","Cicadellidae","Typhlocybinae","Empoasca","Empoasca sp., Empoasca fabae","436","293","293","26","32.8","0","99.56","440.07","0","447.66","32.44","0"
"AGAW3","08N","INS","BOLD:ADF3809","Arthropoda","Insecta","Diptera","Dolichopodidae","","","","389","354","354","27","9","0","98","432.15","0","433.41","35.35","0"
"AGAW3","08N","INS","BOLD:AAV1055","Arthropoda","Insecta","Diptera","Stratiomyidae","Beridinae","Allognosta","Allognosta fuscitarsis","371","334","334","28","9.97","0","99.09","432.77","0","434.3","35.92","0"
"AGAW3","08N","INS","BOLD:AAN5332","Arthropoda","Insecta","Diptera","Chironomidae","Chironominae","Polypedilum","","357","351","351","29","1.68","0","97.81","415.16","0","418.41","33.39","0"
"AGAW3","08N","INS","BOLD:AAG7276","Arthropoda","Insecta","Diptera","Sphaeroceridae","Limosininae","Leptocera","Leptocera erythrocera","355","328","328","30","7.61","0","99.18","434.51","0","437.03","32.84","0"
"AGAW3","08N","INS","BOLD:AAG0999","Arthropoda","Insecta","Diptera","Chironomidae","Orthocladiinae","","","322","250","250","31","22.36","0","97.51","438.84","0","439.94","35.76","0"
"AGAW3","08N","INS","BOLD:AAF9771","Arthropoda","Insecta","Diptera","Hybotidae","Tachydromiinae","Platypalpus","Platypalpus longiseta, Platypalpus trivialis, Platypalpus sp. 5, Platypalpus sp. 17, Platypalpus pallidiventris","294","231","231","32","21.43","0","99.07","445.3","0","445.93","35","0"
"AGAW3","08N","INS","BOLD:AAL7359","Arthropoda","Insecta","Diptera","Chironomidae","Chironominae","Tanytarsus","","257","242","242","33","5.84","0","99.11","438.44","0","440.3","36.09","0"
"AGAW3","08N","INS","BOLD:AAC4197","Arthropoda","Insecta","Diptera","Chironomidae","Orthocladiinae","Paraphaenocladius","Paraphaenocladius impensus","249","222","222","34","10.84","0","99.22","430.96","0","436.28","35.01","0"
"AGAW3","08N","INS","BOLD:AAH3661","Arthropoda","Insecta","Diptera","Cecidomyiidae","","","","240","217","217","35","9.58","0","99.18","431.58","0","433.19","25.38","0"
"AGAW3","08N","INS","BOLD:ACE0554","Arthropoda","Insecta","Diptera","Phoridae","Metopininae","Megaselia","","221","188","188","36","14.93","0","99.45","443.66","0","445.29","32.02","0"
"AGAW3","08N","INS","BOLD:AAD0484","Arthropoda","Insecta","Diptera","Chironomidae","","","","212","184","184","37","13.21","0","99.03","443.55","0","444.93","34.63","0"
"AGAW3","08N","INS","BOLD:AAG5010","Arthropoda","Insecta","Hymenoptera","Braconidae","Rogadinae","Aleiodes","","190","147","147","38","22.63","0","99.63","444.22","0","445.69","27.37","0"
"AGAW3","08N","INS","BOLD:AAD1485","Arthropoda","Insecta","Diptera","Chironomidae","Chironominae","Paratanytarsus","Paratanytarsus grimmii, Paratanytarsus sp.","181","162","162","39","10.5","0","99.35","441.09","0","442.56","33.84","0"
"AGAW3","08N","INS","BOLD:AAB1098","Arthropoda","Insecta","Diptera","Culicidae","Culicinae","Ochlerotatus, Aedes","Aedes excrucians, Aedes cantans, Aedes stimulans, Aedes euedes, Aedes riparius, Aedes annulipes, Ochlerotatus cantans, Aedes fitchii, Aedes sp., Ochlerotatus annulipes, Ochlerotatus excrucians","179","165","165","40","7.82","0","98.65","430.02","0","431.32","35.27","0"
"AGAW3","08N","INS","BOLD:AAB2384","Arthropoda","Insecta","Diptera","Syrphidae","Syrphinae","Eupeodes","Eupeodes duseki, Eupeodes volucris, Eupeodes nielseni, Eupeodes americanus, Eupeodes flukei, Eupeodes vockerothi, Eupeodes lucasi, Eupeodes montivagus, Eupeodes rufipunctatus, Eupeodes rojasi, Eupeodes goeldlini, Eupeodes luniger, Eupeodes punctifer, Eupeodes bucculatus, Eupeodes perplexus, Eupeodes abiskoensis, Eupeodes corollae, Eupeodes pomus, Eupeodes lundbecki, Eupeodes nitens, Eupeodes sp. BOLDAAB2384, Eupeodes curtus, Eupeodes latifasciatus","178","175","175","41","1.69","0","98.81","409.11","0","411.6","30.75","0"
"AGAW3","08N","INS","BOLD:AAU6967","Arthropoda","Insecta","Coleoptera","Chrysomelidae","Galerucinae","Psylliodes","Psylliodes affinis","174","134","134","42","22.99","0","99.48","435.8","0","445.67","34.95","0"
"AGAW3","08N","INS","BOLD:AAN5342","Arthropoda","Insecta","Diptera","Chironomidae","Orthocladiinae","Heleniella","","172","162","162","43","5.81","0","98.86","420.78","0","428.99","35.19","0"
"AGAW3","08N","INS","BOLD:AAG6432","Arthropoda","Insecta","Diptera","Ceratopogonidae","Forcipomyiinae","Atrichopogon","","171","149","149","44","12.87","0","98.57","439.26","0","440.37","34.2","0"
"AGAW3","08N","INS","BOLD:AAA3882","Arthropoda","Insecta","Lepidoptera","Pterophoridae","Pterophorinae","Emmelina","Emmelina monodactyla","139","139","139","45","0","0","98.73","402.15","0","409.82","31.42","0"
"AGAW3","08N","INS","BOLD:AAM6288","Arthropoda","Insecta","Diptera","Chironomidae","Chironominae","Chironomus","","129","101","101","46","21.71","0","99.48","445.87","0","447.28","35.14","0"
"AGAW3","08N","INS","BOLD:AAN6154","Arthropoda","Insecta","Coleoptera","Latridiidae","Corticariinae","Melanophthalma","Melanophthalma inermis","121","109","109","47","9.92","0","99.07","435.97","0","437.17","31.14","0"
"AGAW3","08N","INS","BOLD:AAP6790","Arthropoda","Insecta","Diptera","Chloropidae","Oscinellinae","Incertella","Incertella incerta","118","116","116","48","1.69","0","98.97","420.15","0","421.82","31.47","0"
"AGAW3","08N","INS","BOLD:ACK4716","Arthropoda","Insecta","Diptera","Cecidomyiidae","","","","112","107","107","49","4.46","0","99.14","425.85","0","427.02","31.61","0"
"AGAW3","08N","INS","BOLD:AAN5160","Arthropoda","Insecta","Diptera","Ceratopogonidae","Forcipomyiinae","Atrichopogon","","107","103","103","50","3.74","0","99.03","431.34","0","434.21","37.44","0"
"AGAW3","08N","INS","BOLD:AAL7819","Arthropoda","Insecta","Diptera","Psychodidae","Psychodinae","Psychoda","Psychoda sp., Psychoda sp. BOLD:AAL7819","98","89","89","51","9.18","0","99.44","416.56","0","418.27","32.61","0"
"AGAW3","08N","INS","BOLD:ACJ7008","Arthropoda","Insecta","Diptera","Sciaridae","","Leptosciarella","Leptosciarella belingi","90","85","85","52","5.56","0","99.73","349.67","0","419.48","32.62","0"
"AGAW3","08N","INS","BOLD:ACL6355","Arthropoda","Insecta","Diptera","Chironomidae","Chironominae","Polypedilum","","84","83","83","53","1.19","0","98.67","412.49","0","415.1","33.95","0"
"AGAW3","08N","INS","BOLD:AAA5299","Arthropoda","Insecta","Diptera","Chironomidae","Orthocladiinae","Cricotopus","Cricotopus laetus, Cricotopus relucens, Cricotopus trifasciatus, Cricotopus sp., Cricotopus sp. 3ES, Cricotopus sylvestris, Cricotopus glacialis","79","74","74","54","6.33","0","99.44","425.23","0","429.94","34.5","0"
"AGAW3","08N","INS","BOLD:AAN8770","Arthropoda","Insecta","Diptera","Psychodidae","Psychodinae","Psychoda","Psychoda trinodulosa","77","68","68","55","11.69","0","99.28","444.53","0","446.95","33.94","0"
"AGAW3","08N","INS","BOLD:AAE2703","Arthropoda","Insecta","Diptera","Drosophilidae","Drosophilinae","Chymomyza","Chymomyza amoena","76","76","76","56","0","0","98.78","394.32","0","397.54","31.86","0"
"AGAW3","08N","INS","BOLD:ABW1511","Arthropoda","Insecta","Diptera","Ceratopogonidae","","","","68","56","56","57","17.65","0","97.12","418","0","419.85","36.18","0"
"AGAW3","08N","INS","BOLD:AAF9393","Arthropoda","Insecta","Coleoptera","Meloidae","Meloinae","Epicauta","Epicauta fabricii","63","58","58","58","7.94","0","98.38","437.41","0","439.06","40.78","0"
"AGAW3","08N","INS","BOLD:ADH4889","Arthropoda","Insecta","Diptera","","","","","60","54","54","59","10","0","98.93","446.8","0","448.15","35.41","0"
"AGAW3","08N","INS","BOLD:AAG5457","Arthropoda","Insecta","Diptera","Chironomidae","Chironominae","","","57","57","57","60","0","0","98.66","432.28","0","434.84","34.42","0"
"AGAW3","08N","INS","BOLD:AAP9021","Arthropoda","Insecta","Diptera","Cecidomyiidae","","","","57","57","57","60","0","0","99.06","418.02","0","419.84","30.03","0"
"AGAW3","08N","INS","BOLD:AAM9347","Arthropoda","Insecta","Diptera","Phoridae","Metopininae","Megaselia","","52","52","52","62","0","0","98.17","397.35","0","399.54","32.8","0"
"AGAW3","08N","INS","BOLD:ABA0856","Arthropoda","Insecta","Diptera","Cecidomyiidae","","","","50","42","42","63","16","0","98.96","446.6","0","448.08","29.01","0"
"AGAW3","08N","INS","BOLD:AAA3080","Arthropoda","Insecta","Trichoptera","Limnephilidae","Limnephilinae","Limnephilus","Limnephilus submonilifer","47","47","47","64","0","0","99.09","399.89","0","401.49","33.94","0"
"AGAW3","08N","INS","BOLD:AAL7751","Arthropoda","Insecta","Diptera","Sphaeroceridae","Limosininae","Coproica","Coproica acutangula","46","37","37","65","19.57","0","98.93","442.74","0","444.11","33.46","0"
"AGAW3","08N","INS","BOLD:ABU5525","Arthropoda","Insecta","Diptera","Chironomidae","Orthocladiinae","Limnophyes","Limnophyes sp. 14ES","46","46","46","65","0","0","99.03","403.04","0","409","37.2","0"
"AGAW3","08N","INS","BOLD:AAG8493","Arthropoda","Insecta","Diptera","Drosophilidae","Drosophilinae","Scaptomyza","Scaptomyza pallida","44","44","44","67","0","0","98.97","400.41","0","402.45","33.15","0"
"AGAW3","08N","INS","BOLD:AAV5575","Arthropoda","Insecta","Diptera","Cecidomyiidae","","","CecidInt30 sp. BOLD:AAV5575","43","33","33","68","23.26","0","99.62","445.6","0","446.51","25.93","0"
"AGAW3","08N","INS","BOLD:ACF2457","Arthropoda","Insecta","Diptera","Chironomidae","Chironominae","Paratanytarsus","Paratanytarsus laccophilus","43","43","43","68","0","0","99.23","441.05","0","443.02","32.6","0"
"AGAW3","08N","INS","BOLD:AAH1890","Arthropoda","Insecta","Hymenoptera","Ichneumonidae","Cryptinae","Stibeutes","Stibeutes brevicornis, Stibeutes heinemanni","42","34","34","70","19.05","0","99.45","446.86","0","448.6","29.76","0"
"AGAW3","08N","INS","BOLD:AAG5192","Arthropoda","Insecta","Coleoptera","Curculionidae","Entiminae","Barypeithes","Barypeithes pellucidus","37","32","32","71","13.51","0","99.47","439.68","0","440.7","39.48","0"
"AGAW3","08N","INS","BOLD:AAG8839","Arthropoda","Insecta","Hemiptera","Cicadellidae","Deltocephalinae","Errastunus, Latalus","Latalus ocellaris, Errastunus ocellaris","36","29","29","72","19.44","0","99.62","447.72","0","449.06","35.87","0"
"AGAW3","08N","INS","BOLD:AAP7834","Arthropoda","Insecta","Coleoptera","Phalacridae","Phalacrinae","Stilbus","","30","29","29","73","3.33","0","98.68","429.63","0","431.57","32.58","0"
"AGAW3","08N","INS","BOLD:AAC4523","Arthropoda","Insecta","Diptera","Chironomidae","Chironominae","Tanytarsus","Tanytarsus guerlus","30","28","28","73","6.67","0","98.86","426.67","0","429.33","32.42","0"
"AGAW3","08N","INS","BOLD:AAA3453","Arthropoda","Insecta","Diptera","Anthomyiidae","Anthomyiinae","Delia","","28","28","28","75","0","0","99.06","429.43","0","431.96","34.61","0"
"AGAW3","08N","INS","BOLD:ABA0929","Arthropoda","Insecta","Diptera","Sciaridae","","Bradysia","Bradysia scabricornis","28","28","28","75","0","0","99.09","422.75","0","424.64","30.08","0"
"AGAW3","08N","INS","BOLD:ABW7897","Arthropoda","Insecta","Diptera","Cecidomyiidae","","","","28","23","23","75","17.86","0","99.36","445.18","0","447","29.78","0"
"AGAW3","08N","INS","BOLD:AAH3920","Arthropoda","Insecta","Diptera","Sciaridae","","Dichopygina, Scatopsciara","","27","27","27","78","0","0","98.44","387.7","0","402.59","35.47","0"
"AGAW3","08N","INS","BOLD:ACF6513","Arthropoda","Insecta","Diptera","Psychodidae","Psychodinae","Psychoda","Psychoda cinerea","27","26","26","78","3.7","0","99.33","433.26","0","434.67","30.79","0"
"AGAW3","08N","INS","BOLD:ACK3602","","","","","","","","27","27","27","78","0","0","98.99","411.59","0","414.41","26.27","0"
"AGAW3","08N","INS","BOLD:AAF5705","Arthropoda","Insecta","Hemiptera","Miridae","Orthotylinae","Blepharidopterus","Blepharidopterus chlorionis","27","24","24","78","11.11","0","99.36","360.85","0","447.07","42.5","0"
"AGAW3","08N","INS","BOLD:AAG6934","Arthropoda","Insecta","Diptera","Hybotidae","","","","25","24","24","82","4","0","99.22","434.4","0","436.12","38.72","0"
"AGAW3","08N","INS","BOLD:AAH3653","Arthropoda","Insecta","Diptera","Cecidomyiidae","","","","24","23","23","83","4.17","0","99.05","435.5","0","437.04","25.59","0"
"AGAW3","08N","INS","BOLD:AAH0134","Arthropoda","Insecta","Coleoptera","Phalacridae","Phalacrinae","Stilbus","Stilbus apicialis","23","23","23","84","0","0","99.3","421.35","0","422.7","31.85","0"
"AGAW3","08N","INS","BOLD:AAN5183","Arthropoda","Insecta","Diptera","Cecidomyiidae","","","","23","22","22","84","4.35","0","98.36","439.65","0","440.74","27.79","0"
"AGAW3","08N","INS","BOLD:AAN5242","Arthropoda","Insecta","Diptera","Cecidomyiidae","","","","21","21","21","86","0","0","98.82","412.57","0","413.86","28.15","0"
"AGAW3","08N","INS","BOLD:AAG5242","Arthropoda","Insecta","Coleoptera","Curculionidae","Entiminae","Barypeithes","Barypeithes pellucidus","20","20","20","87","0","0","98.91","427","0","431.25","39.89","0"
"AGAW3","08N","INS","BOLD:ABU5520","Arthropoda","Insecta","Diptera","Sciaridae","","Corynoptera","","20","20","20","87","0","0","98.75","414","0","416.1","35.11","0"
"AGAW3","08N","INS","BOLD:ABV1397","Arthropoda","Insecta","Diptera","Cecidomyiidae","","","","19","18","18","89","5.26","0","98.87","434.21","0","435.21","29.33","0"
"AGAW3","08N","INS","BOLD:AAV5688","Arthropoda","Insecta","Diptera","Cecidomyiidae","","","","18","18","18","90","0","0","99.24","436.39","0","437.44","34.4","0"
"AGAW3","08N","INS","BOLD:ABV9319","Arthropoda","Insecta","Diptera","Chironomidae","Tanypodinae","Procladius","","18","18","18","90","0","0","99.11","427.83","0","429.61","34.18","0"
"AGAW3","08N","INS","BOLD:ACE3123","Arthropoda","Insecta","Diptera","Sciaridae","","Corynoptera","","17","16","16","92","5.88","0","99.4","439.18","0","440","35.07","0"
"AGAW3","08N","INS","BOLD:ACF8174","Arthropoda","Insecta","Diptera","Chironomidae","Chironominae","","","17","17","17","92","0","0","98.89","430.12","0","432.71","32.46","0"
"AGAW3","08N","INS","BOLD:ACJ6954","Arthropoda","Insecta","Diptera","Cecidomyiidae","","","","16","16","16","94","0","0","99.58","337.81","0","422.13","26.81","0"
"AGAW3","08N","INS","BOLD:AAV5594","Arthropoda","Insecta","Diptera","Cecidomyiidae","","","","14","13","13","95","7.14","0","99.18","443.29","0","444","26.36","0"
"AGAW3","08N","INS","BOLD:AAM7855","Arthropoda","Insecta","Thysanoptera","Thripidae","Thripinae","Frankliniella","Frankliniella tritici","14","14","14","95","0","0","99.17","434.5","0","437.14","31.31","0"
"AGAW3","08N","INS","BOLD:AAG2511","Arthropoda","Insecta","Diptera","Anthomyiidae","Anthomyiinae","Delia","Delia platura","14","14","14","95","0","0","98.8","446.93","0","449","33.55","0"
"AGAW3","08N","INS","BOLD:AAE3943","Arthropoda","Insecta","Hemiptera","Miridae","Phylinae","Plagiognathus","Plagiognathus chrysanthemi","13","13","13","98","0","0","98.74","392.08","0","447.54","35.64","0"
"AGAW3","08N","INS","BOLD:AAD0245","Arthropoda","Insecta","Hymenoptera","Ichneumonidae","Orthocentrinae","Stenomacrus","Stenomacrus jft17, Stenomacrus JFT17","13","13","13","98","0","0","98.61","422.08","0","425.15","26.49","0"
"AGAW3","08N","INS","BOLD:AAG6468","Arthropoda","Insecta","Diptera","Ceratopogonidae","Ceratopogoninae","Culicoides","Culicoides biguttatus","12","12","12","100","0","0","98.01","431","0","431.83","37.3","0"
"AGAW3","08N","INS","BOLD:AAA4781","Arthropoda","Insecta","Hymenoptera","Braconidae","Microgastrinae","Glyptapanteles","Glyptapanteles carlossarmientoi","12","12","12","100","0","0","99.26","422.33","0","424.92","24.05","0"
"AGAW3","08N","INS","BOLD:AAP8739","Arthropoda","Insecta","Diptera","Platystomatidae","Platystomatinae","Rivellia","Rivellia steyskali","12","12","12","100","0","0","98.76","441.17","0","442.5","38.28","0"
"AGAW3","08N","INS","BOLD:AAG7891","Arthropoda","Insecta","Hymenoptera","Platygastridae","Scelioninae","Telenomus","Telenomus podisi","11","11","11","103","0","0","98.61","416.91","0","426.09","26.61","0"
"AGAW3","08N","INS","BOLD:AAJ7123","Arthropoda","Insecta","Diptera","Culicidae","Culicinae","Culiseta","Culiseta minnesotae","11","11","11","103","0","0","97.73","438.64","0","439.55","34.88","0"
"AGAW3","08N","INS","BOLD:AAF5258","Arthropoda","Insecta","Hymenoptera","Braconidae","Microgastrinae","Microplitis","Microplitis jft114, Microplitis jft58","11","9","9","103","18.18","0","99.28","440","0","441.64","25.62","0"
"AGAW3","08N","INS","BOLD:AAP2527","Arthropoda","Insecta","Diptera","Mycetophilidae","Mycetophilinae","Brevicornu","Brevicornu fissicauda","10","9","9","106","10","0","99.53","449.2","0","450","37.04","0"
"AGAW3","08N","INS","BOLD:AAQ0598","Arthropoda","Insecta","Diptera","Chironomidae","Orthocladiinae","Limnophyes","Limnophyes minimus","10","10","10","106","0","0","99.46","443.2","0","444.3","37.89","0"
"AGAW3","08N","INS","BOLD:ACP8563","Arthropoda","Insecta","Hymenoptera","Braconidae","Euphorinae","Centistes","","10","10","10","106","0","0","99.27","425.3","0","444.3","32.98","0"
"AGAW3","08N","INS","BOLD:AAP6827","Arthropoda","Insecta","Diptera","Cecidomyiidae","","","","9","9","9","109","0","0","98.41","443.89","0","446.22","31.19","0"
"AGAW3","08N","INS","BOLD:AAB0373","Arthropoda","Insecta","Neuroptera","Chrysopidae","Chrysopinae","Chrysoperla","Chrysoperla sp. isolate 82, Chrysoperla zastrowi, Chrysoperla plorabunda, Chrysoperla sp. isolate 71, Chrysoperla agilis, Italochrysa italica, Chrysoperla sp. isolate 88, Chrysoperla lucasina, Chrysoperla carnea, Chrysoperla mediterranea, Chrysoperla pallida, Chrysoperla nipponensis, Chrysoperla sillemi","9","9","9","109","0","0","99.15","436.11","0","437.11","31.39","0"
"AGAW3","08N","INS","BOLD:AAH0135","Arthropoda","Insecta","Coleoptera","Phalacridae","Phalacrinae","Acylomus","Acylomus pugetanus, Acylomus ergoti","9","9","9","109","0","0","98.96","423.67","0","426.11","31.35","0"
"AGAW3","08N","INS","BOLD:AAM9228","Arthropoda","Insecta","Diptera","Sciaridae","","","","9","9","9","109","0","0","99.43","449.11","0","449.67","36","0"
"AGAW3","08N","INS","BOLD:AAH1912","","","","","","","","8","8","8","113","0","0","98.63","410.88","0","412.25","27.9","0"
"AGAW3","08N","INS","BOLD:ADF3808","Arthropoda","Insecta","Diptera","Dolichopodidae","","","","8","8","8","113","0","0","97.68","382","0","393.5","34.66","0"
"AGAW3","08N","INS","BOLD:AAF9311","Arthropoda","Insecta","Diptera","Psychodidae","Psychodinae","Psychoda","Psychoda cf. satchelli, Psychoda satchelli","7","7","7","115","0","0","99.74","435.57","0","436.43","34.97","0"
"AGAW3","08N","INS","BOLD:ACX6055","Arthropoda","Insecta","Diptera","Phoridae","Metopininae","Megaselia","Megaselia sp. AK-2010-2, Megaselia sp. BOLD:ACX6055","7","7","7","115","0","0","98.39","432.86","0","434","32.62","0"
"AGAW3","08N","INS","BOLD:AAN5225","Arthropoda","Insecta","Diptera","Cecidomyiidae","","","","6","6","6","117","0","0","99.59","448.83","0","450","28.33","0"
"AGAW3","08N","INS","BOLD:AAV5573","Arthropoda","Insecta","Diptera","Cecidomyiidae","","","","6","4","4","117","33.33","0","99.81","449","0","450","27.63","0"
"AGAW3","08N","INS","BOLD:AAM6037","Arthropoda","Insecta","Diptera","Cecidomyiidae","Cecidomyiinae","Contarinia, Dasineura","","6","6","6","117","0","0","98.47","436.5","0","438","26.49","0"
"AGAW3","08N","INS","BOLD:AAN5267","Arthropoda","Insecta","Diptera","Cecidomyiidae","","","","6","5","5","117","16.67","0","99.18","448.67","0","450","30.56","0"
"AGAW3","08N","INS","BOLD:AAN6435","Arthropoda","Insecta","Diptera","Sciaridae","","Bradysia","Bradysia angustipennis","6","5","5","117","16.67","0","99.87","397.67","0","440.83","33.98","0"
"AGAW3","08N","INS","BOLD:ABA0814","Arthropoda","Insecta","Diptera","Cecidomyiidae","","","","6","6","6","117","0","0","98.4","416.5","0","419.67","29.71","0"
"AGAW3","08N","INS","BOLD:ABA0863","Arthropoda","Insecta","Diptera","Cecidomyiidae","","","","5","4","4","123","20","0","99.87","448.6","0","450","29.96","0"
"AGAW3","08N","INS","BOLD:AAU9508","Arthropoda","Insecta","Hymenoptera","Dryinidae","Anteoninae","Anteon","Anteon tenuicorne, Anteon cf. pubicorne, Anteon pubicorne","5","5","5","123","0","0","99.24","445.2","0","445.8","29.79","0"
"AGAW3","08N","INS","BOLD:AAN5199","Arthropoda","Insecta","Diptera","Cecidomyiidae","","","CecidInt20 sp. BOLD:AAN5199","5","5","5","123","0","0","98.84","448.8","0","450","26.76","0"
"AGAW3","08N","INS","BOLD:ABV2748","Arthropoda","Insecta","Hymenoptera","Braconidae","Alysiinae","","","5","5","5","123","0","0","99.28","437","0","438.2","28.42","0"
"AGAW3","08N","INS","BOLD:ADH9867","Arthropoda","Insecta","Diptera","","","","","5","5","5","123","0","0","98.63","413.6","0","417.4","37.42","0"
"AGAW3","08N","INS","BOLD:ACD0207","Arthropoda","Insecta","Hymenoptera","Braconidae","Alysiinae","Dinotrema","","5","5","5","123","0","0","99.68","420.6","0","422.8","23.84","0"
"AGAW3","08N","INS","BOLD:ACU4215","Arthropoda","Insecta","Diptera","Dolichopodidae","","","","4","4","4","129","0","0","97.36","444.25","0","445","34.89","0"
"AGAW3","08N","INS","BOLD:AAA4277","Arthropoda","Insecta","Diptera","Syrphidae","Syrphinae","Toxomerus","Toxomerus marginatus, Toxomerus sp. CR-3, Toxomerus CR-3","4","4","4","129","0","0","99.36","440","0","441.5","30.69","0"
"AGAW3","08N","INS","BOLD:ABA7871","Arthropoda","Insecta","Diptera","Cecidomyiidae","","","","4","4","4","129","0","0","98.76","444","0","445","27.25","0"
"AGAW3","08N","INS","BOLD:AAC5848","Arthropoda","Insecta","Lepidoptera","Psychidae","Psychinae","Psyche","Psyche casta","4","4","4","129","0","0","99.32","410.75","0","412.75","37.47","0"
"AGAW3","08N","INS","BOLD:AAN6447","Arthropoda","Insecta","Diptera","Sciaridae","","Corynoptera","Corynoptera perpusilla","4","4","4","129","0","0","99.44","447","0","450","36.33","0"
"AGAW3","08N","INS","BOLD:AAV5352","Arthropoda","Insecta","Diptera","Cecidomyiidae","","","","4","4","4","129","0","0","98.71","447.25","0","450","26.33","0"
"AGAW3","08N","INS","BOLD:AAA3654","Arthropoda","Insecta","Lepidoptera","Crambidae","Crambinae","Crambus","Crambus bigelovi, Crambus johnsoni, Crambus praefectellus","4","4","4","129","0","0","99.21","437.75","0","438.5","31.22","0"
"AGAW3","08N","INS","BOLD:AAM9255","Arthropoda","Insecta","Diptera","Sciaridae","","Pseudolycoriella","","4","4","4","129","0","0","99.11","391.75","0","396.75","30.56","0"
"AGAW3","08N","INS","BOLD:AAM6097","Arthropoda","Insecta","Diptera","Cecidomyiidae","","","","3","3","3","137","0","0","98.95","395.67","0","397.67","34.1","0"
"AGAW3","08N","INS","BOLD:AAU6681","Arthropoda","Insecta","Diptera","Tipulidae","Tipulinae","Nephrotoma","Nephrotoma cornicina","3","3","3","137","0","0","99.55","445","0","447","36.39","0"
"AGAW3","08N","INS","BOLD:AAN5255","Arthropoda","Insecta","Diptera","Cecidomyiidae","Cecidomyiinae","Dasineura","","3","3","3","137","0","0","97.92","449.33","0","450","26.81","0"
"AGAW3","08N","INS","BOLD:AAG0996","Arthropoda","Insecta","Diptera","Chironomidae","Orthocladiinae","Cricotopus","Cricotopus sp. 18ES, Cricotopus bicinctus gr., Cricotopus bicinctus grp.","3","3","3","137","0","0","98.43","446.67","0","448","33.41","0"
"AGAW3","08N","INS","BOLD:ABA6514","Arthropoda","Insecta","Diptera","Chironomidae","Chironominae","Rheotanytarsus","","3","3","3","137","0","0","99.05","399","0","401","36.4","0"
"AGAW3","08N","INS","BOLD:ACY2979","","","","","","","","3","3","3","137","0","0","98.77","428.33","0","430.33","34.7","0"
"AGAW3","08N","INS","BOLD:ACD9543","Arthropoda","Insecta","Diptera","Sciaridae","","Bradysia","Bradysia sp. BOLD:ACD9543","3","3","3","137","0","0","99.16","442.33","0","444","34.83","0"
"AGAW3","08N","INS","BOLD:AAC0706","Arthropoda","Insecta","Diptera","Chironomidae","Chironominae","Dicrotendipes","Dicrotendipes tritomus","3","3","3","137","0","0","97.87","437.67","0","448.67","36.92","0"
"AGAW3","08N","INS","BOLD:ACM7774","Arthropoda","Insecta","Trichoptera","Hydroptilidae","Orthotrichiinae","Orthotrichia","Orthotrichia cristata","3","3","3","137","0","0","98.59","386.33","0","386.33","28.83","0"
"AGAW3","08N","INS","BOLD:AAD8086","Arthropoda","Insecta","Lepidoptera","Elachistidae","Elachistinae","Elachista","Elachista griseicornis, Elachista acenteta","3","3","3","137","0","0","99.1","407.67","0","409.33","30.37","0"
"AGAW3","08N","INS","BOLD:AAG1717","Arthropoda","Insecta","Diptera","Muscidae","Phaoniinae","Helina","Helina troene","2","2","2","147","0","0","99.67","444","0","444","31.42","0"
"AGAW3","08N","INS","BOLD:ACU7906","Arthropoda","Insecta","Diptera","Chironomidae","Orthocladiinae","","","2","2","2","147","0","0","99.28","411.5","0","415","35.42","0"
"AGAW3","08N","INS","BOLD:AAA6560","Arthropoda","Insecta","Lepidoptera","Crambidae","Crambinae","Crambus","Crambus multilinellus, Crambus saltuellus","2","2","2","147","0","0","99.33","450","0","450","31.11","0"
"AGAW3","08N","INS","BOLD:ABA1215","Arthropoda","Insecta","Diptera","Sciaridae","","Lycoriella","Lycoriella sativae","2","2","2","147","0","0","99.89","450","0","450","35.89","0"
"AGAW3","08N","INS","BOLD:ACF4786","Arthropoda","Insecta","Hymenoptera","Braconidae","Aphidiinae","Aphidius","Aphidius urticae, Aphidius matricariae","2","2","2","147","0","0","99.08","414","0","419","26.11","0"
"AGAW3","08N","INS","BOLD:AAC9132","Arthropoda","Insecta","Diptera","Culicidae","Culicinae","Culiseta","Culiseta inornata","2","2","2","147","0","0","98.33","400","0","402","32.8","0"
"AGAW3","08N","INS","BOLD:ACR8658","Arthropoda","Insecta","Diptera","Chironomidae","Chironominae","Tanytarsus","","2","2","2","147","0","0","98.85","392.5","0","426.5","36.11","0"
"AGAW3","08N","INS","BOLD:ACM7639","Arthropoda","Insecta","Diptera","Cecidomyiidae","","","","2","2","2","147","0","0","97.72","417","0","417.5","28","0"
"AGAW3","08N","INS","BOLD:ACL4768","Arthropoda","Insecta","Coleoptera","Chrysomelidae","Galerucinae","Chaetocnema","Chaetocnema confinis","2","2","2","147","0","0","99.28","349","0","414","32.37","0"
"AGAW3","08N","INS","BOLD:AAG0906","Arthropoda","Insecta","Neuroptera","Hemerobiidae","Microminae","Micromus","Micromus posticus","2","2","2","147","0","0","99.09","441.5","0","444.5","32.96","0"
"AGAW3","08N","INS","BOLD:AAN5383","Arthropoda","Insecta","Diptera","Chironomidae","Chironominae","Dicrotendipes","","2","2","2","147","0","0","97.41","410.5","0","414.5","36.45","0"
"AGAW3","08N","INS","BOLD:AAB9404","Arthropoda","Insecta","Lepidoptera","Tortricidae","Tortricinae","Archips","Archips rosana, Archips grisea","2","2","2","147","0","0","98.42","444","0","446","31.84","0"
"AGAW3","08N","INS","BOLD:ACE7444","Arthropoda","Insecta","Hemiptera","Miridae","Mirinae, Phylinae","Adelphocoris, Megalocoleus","Adelphocoris lineolatus, Megalocoleus molliculus","2","2","2","147","0","0","99.33","447.5","0","450","34.89","0"
"AGAW3","08N","INS","BOLD:AAA2815","Arthropoda","Insecta","Lepidoptera","Noctuidae","Noctuinae","Mythimna","Mythimna yukonensis, Mythimna oxygala","2","2","2","147","0","0","99.1","443.5","0","445","33.6","0"
"AGAW3","08N","INS","BOLD:AAD5601","Arthropoda","Insecta","Lepidoptera","Noctuidae","Noctuinae","Apamea","Apamea impulsa","2","2","2","147","0","0","97.66","448","0","450","31.22","0"
"AGAW3","08N","INS","BOLD:AAF9187","Arthropoda","Insecta","Coleoptera","Curculionidae","Entiminae","Phyllobius","Phyllobius oblongus","2","2","2","147","0","0","99.22","449","0","450","39","0"
"AGAW3","08N","INS","BOLD:AAD2169","Arthropoda","Insecta","Diptera","Chironomidae","Chironominae","Tanytarsus","Tanytarsus volgensis","2","2","2","147","0","0","98.89","427","0","431.5","39.49","0"
"AGAW3","08N","INS","BOLD:ABX0130","Arthropoda","Insecta","Diptera","Cecidomyiidae","","","CecidInt43 sp. BOLD:ABX0130","2","2","2","147","0","0","98.88","433.5","0","434.5","30.01","0"
"AGAW3","08N","INS","BOLD:AAA7374","Arthropoda","Insecta","Diptera","Syrphidae","Syrphinae","Sphaerophoria","Sphaerophoria bankowskae, Sphaerophoria infuscata, Sphaerophoria nr. philanthus, Sphaerophoria batava, Sphaerophoria sp., Sphaerophoria rueppellii, Sphaerophoria interrupta-group, Sphaerophoria sp. MA-1, Sphaerophoria cleoae, Sphaerophoria asymmetrica, Sphaerophoria fatarum, Sphaerophoria bengalensis, Sphaerophoria bifurcata, Sphaerophoria taeniata, Sphaerophoria interrupta, Sphaerophoria contigua, Sphaerophoria kaa, Sphaerophoria laurae, Sphaerophoria chongjini, Sphaerophoria boreoalpina, Sphaerophoria sp. BOLDAAA7374, Sphaerophoria macrogaster, Sphaerophoria sulphuripes, Sphaerophoria brevipilosa, Sphaerophoria philanthus, Sphaerophoria pyrrhina, Sphaerophoria abbreviata, Sphaerophoria scripta, Sphaerophoria potentillae, Sphaerophoria longipilosa, Sphaerophoria virgata","2","2","2","147","0","0","99.55","447.5","0","450","31.11","0"
"AGAW3","08N","INS","BOLD:AAA3181","Arthropoda","Insecta","Diptera","Muscidae","Muscinae","Stomoxys","Stomoxys calcitrans","1","1","1","166","0","0","98.27","404","0","408","31.86","0"
"AGAW3","08N","INS","BOLD:ADE4394","Arthropoda","Insecta","Diptera","Chironomidae","Chironominae","Tanytarsus","","1","1","1","166","0","0","97.18","426","0","450","35.33","0"
"AGAW3","08N","INS","BOLD:AAG3920","Arthropoda","Insecta","Diptera","Chironomidae","Tanypodinae","Procladius","Procladius sp. 1ES","1","1","1","166","0","0","99.33","447","0","450","33.33","0"
"AGAW3","08N","INS","BOLD:AAA6759","Arthropoda","Insecta","Lepidoptera","Hesperiidae","Hesperiinae","Thymelicus","Thymelicus lineola","1","1","1","166","0","0","97.51","441","0","444","30.63","0"
"AGAW3","08N","INS","BOLD:ACD9062","Arthropoda","Insecta","Hemiptera","Miridae","Mirinae","Capsus","Capsus ater, Capsus cinctus","1","1","1","166","0","0","98.61","432","0","450","36.22","0"
"AGAW3","08N","INS","BOLD:AAE4612","Arthropoda","Insecta","Hemiptera","Miridae","Phylinae","Plagiognathus","Plagiognathus suffuscipennis, Plagiognathus tumidifrons","1","1","1","166","0","0","100","448","0","449","38.31","0"
"AGAW3","08N","INS","BOLD:AAZ5973","Arthropoda","Insecta","Diptera","Sepsidae","Sepsinae","Themira","Themira annulipes","1","1","1","166","0","0","98.79","413","0","416","34.86","0"
"AGAW3","08N","INS","BOLD:ACK3431","Arthropoda","Insecta","Diptera","Cecidomyiidae","","","","1","1","1","166","0","0","97.77","449","0","450","33.11","0"
"AGAW3","08N","INS","BOLD:AAH4135","Arthropoda","Insecta","Diptera","Chloropidae","Chloropinae","Thaumatomyia","Thaumatomyia glabra, Thaumatomyia grata, Thaumatomyia sp. 1 TAW","1","1","1","166","0","0","100","446","0","447","32.44","0"
"AGAW3","08N","INS","BOLD:AAB4657","Arthropoda","Insecta","Diptera","Chironomidae","Chironominae","Chironomus","Chironomus maturus","1","1","1","166","0","0","98.22","449","0","450","36.67","0"
"AGAW3","08N","INS","BOLD:AAG5453","Arthropoda","Insecta","Diptera","Chironomidae","Chironominae","Chironomus","Chironomus bifurcatus","1","1","1","166","0","0","98.08","364","0","367","34.88","0"
"AGAW3","08N","INS","BOLD:AAN5238","Arthropoda","Insecta","Diptera","Cecidomyiidae","","","","1","1","1","166","0","0","98.44","450","0","450","26.89","0"
"AGAW3","08N","INS","BOLD:AAA4213","Arthropoda","Insecta","Lepidoptera","Geometridae","Sterrhinae","Idaea","Idaea dimidiata","1","1","1","166","0","0","98.65","445","0","447","30.2","0"
"AGAW3","08N","INS","BOLD:AAN5230","Arthropoda","Insecta","Diptera","Cecidomyiidae","","","","1","1","1","166","0","0","97.35","378","0","384","24.74","0"
"AGAW3","08N","INS","BOLD:AAS1281","Arthropoda","Insecta","Diptera","Chironomidae","Chironominae","Chironomus","Chironomus nr. sollicitus","1","1","1","166","0","0","97.11","450","0","450","33.33","0"
"AGAW3","08N","INS","BOLD:AAY6412","Arthropoda","Insecta","Diptera","Sciaridae","","Bradysia","Bradysia urticae","1","1","1","166","0","0","99.55","445","0","450","31.78","0"
"AGAW3","08N","INS","BOLD:AAH3914","Arthropoda","Insecta","Diptera","Sciaridae","","Sciara","Sciara hamatilis, Sciara humeralis","1","1","1","166","0","0","99.33","449","0","450","30.67","0"
"AGAW3","08N","INS","BOLD:AAZ5504","Arthropoda","Insecta","Diptera","Dolichopodidae","","","","1","1","1","166","0","0","97.07","444","0","450","34.22","0"
"AGAW3","08N","INS","BOLD:AAB7869","Arthropoda","Insecta","Lepidoptera","Tortricidae","Tortricinae","Phtheochroa","Phtheochroa baracana, Phtheochroa riscana, Phtheochroa birdana, Phtheochroa fulviplicana, Phtheochroa canariana","1","1","1","166","0","0","98.43","447","0","450","29.78","0"
"AGAW3","08N","INS","BOLD:AAA2087","Arthropoda","Insecta","Lepidoptera","Erebidae","Herminiinae","Idia","Idia sp., Idia americalis","1","1","1","166","0","0","99.55","449","0","450","30.22","0"
"AGAW3","08N","INS","BOLD:ABA0822","Arthropoda","Insecta","Diptera","Cecidomyiidae","","","","1","1","1","166","0","0","97.97","444","0","450","27.56","0"
"AGAW3","08N","INS","BOLD:AAA2632","Arthropoda","Insecta","Lepidoptera","Noctuidae","Noctuinae","Noctua","Noctua atlantica, Noctua pronuba","1","1","1","166","0","0","99.56","450","0","450","31.33","0"
"AGAW3","08N","INS","BOLD:AAA8796","Arthropoda","Insecta","Lepidoptera","Tortricidae","Tortricinae","Acleris","Acleris forsskaleana","1","1","1","166","0","0","98.57","420","0","423","31.44","0"
"AGAW3","08N","INS","BOLD:ACV5892","Arthropoda","Insecta","Diptera","Cecidomyiidae","","","","1","1","1","166","0","0","99.26","406","0","409","23.96","0"
"AGAW3","08N","INS","BOLD:AAF9317","Arthropoda","Insecta","Diptera","Psychodidae","Psychodinae","Psychoda","Psychoda cf. phalaenoides, Psychoda phalaenoides","1","1","1","166","0","0","97.1","449","0","450","32.89","0"
"AGAW3","08N","INS","BOLD:AAU6970","Arthropoda","Insecta","Coleoptera","Cleridae","Clerinae","Enoclerus","Enoclerus nigripes","1","1","1","166","0","0","97.54","447","0","450","32.44","0"
"AGAW3","08N","INS","BOLD:ADF6661","Arthropoda","Insecta","Diptera","Cecidomyiidae","","","","1","1","1","166","0","0","97.85","418","0","450","30.22","0"
"AGAW3","08N","INS","BOLD:AAL2750","Arthropoda","Insecta","Diptera","Muscidae","Phaoniinae","Phaonia","Phaonia tuguriorum","1","1","1","166","0","0","98.16","380","0","383","31.59","0"
"AGAW3","08N","INS","BOLD:ACP4925","Arthropoda","Insecta","Diptera","Sciaridae","","","","1","1","1","166","0","0","97.69","389","0","450","36.22","0"
"AGAW3","08N","INS","BOLD:ACE4862","","","","","","","","1","1","1","166","0","0","97.11","450","0","450","31.78","0"
"AGAW3","08N","INS","BOLD:AAD4214","Arthropoda","Insecta","Hymenoptera","Ichneumonidae","Diplazontinae","Diplazon","Diplazon laetatorius","1","1","1","166","0","0","99.78","447","0","450","25.78","0"
"AGAW3","08N","INS","BOLD:AAM9013","Arthropoda","Insecta","Diptera","Keroplatidae","Macrocerinae","Macrocera","Macrocera pusilla","1","1","1","166","0","0","99.78","449","0","450","30.44","0"
"AGAW3","08N","INS","BOLD:AAJ1250","Arthropoda","Insecta","Hymenoptera","Braconidae","Aphidiinae","Aphidius","Aphidius avenaphis, Aphidius uzbekistanicus","1","1","1","166","0","0","98.76","404","0","405","26.67","0"
"AGAW3","08N","INS","BOLD:AAG6743","Arthropoda","Insecta","Diptera","Sarcophagidae","Sarcophaginae","Sarcophaga","Sarcophaga sp. 1 MG, Sarcophaga subvicina","1","1","1","166","0","0","99.55","448","0","450","35.78","0"
"AGAW3","08N","INS","BOLD:AAG2123","Arthropoda","Insecta","Diptera","Tachinidae","Dexiinae","Dinera","Dinera grisescens","1","1","1","166","0","0","99.78","449","0","450","32.22","0"
"AGAW3","08N","INS","BOLD:ACZ5284","Arthropoda","Insecta","Diptera","Sciomyzidae","Sciomyzinae","Dictya","Dictya stricta, Dictya expansa, Dictya floridensis, Dictya borealis, Dictya gaigei, Dictya sp. TAW1, Dictya umbroides, Dictya texensis, Dictya steyskali, Dictya pictipes","1","1","1","166","0","0","99.55","449","0","450","36.22","0"
"AGAW3","08N","INS","BOLD:AAA4555","Arthropoda","Insecta","Orthoptera","Acrididae","Melanoplinae","Melanoplus","Melanoplus stupefactus, Melanoplus angustipennis, Melanoplus ponderosus, Melanoplus bruneri, Melanoplus frigidus, Melanoplus scapularis, Melanoplus cinereus, Melanoplus madeleineae, Melanoplus sanguinipes, Melanoplus nigrescens group, Melanoplus tunicae, Melanoplus walshi, Melanoplus femurrubrum, Melanoplus dawsoni, Melanoplus sp., Melanoplus fasciatus, Melanoplus bowditchi, Melanoplus stonei, Melanoplus bivittatus, Melanoplus huroni, Melanoplus thomasi, Melanoplus borealis, Melanoplus packardii","1","1","1","166","0","0","97.46","354","0","357","37.25","0"
"AGAW3","08N","INS","BOLD:AAG1739","Arthropoda","Insecta","Diptera","Muscidae","Coenosiinae","Neodexiopsis","Neodexiopsis rufitibia","1","1","1","166","0","0","100","449","0","450","34.22","0"
"AGAW3","08N","INS","BOLD:ACO2010","Arthropoda","Insecta","Hymenoptera","Braconidae","Aphidiinae","Aphidius","","1","1","1","166","0","0","98.9","365","0","450","27.56","0"
"AGAW3","08N","INS","BOLD:AAM6230","Arthropoda","Insecta","Diptera","Chironomidae","Chironominae","Chironomus","","1","1","1","166","0","0","98.42","443","0","450","34.67","0"
"AGAW3","08N","INS","BOLD:AAD2041","Arthropoda","Insecta","Diptera","Dolichopodidae","Sciapodinae","Condylostylus","Condylostylus inermis, Condylostylus nigrofemoratus, Condylostylus connectans, Condylostylus caudatus, Condylostylus flavipes","1","1","1","166","0","0","99.47","374","0","375","35.47","0"
"AGAW3","08N","INS","BOLD:AAH4205","Arthropoda","Insecta","Diptera","Chloropidae","Chloropinae","Meromyza","Meromyza nigriseta, Meromyza sp.","1","1","1","166","0","0","100","450","0","450","32.89","0"
"AGAW3","08N","INS","BOLD:AAN5221","Arthropoda","Insecta","Diptera","Cecidomyiidae","","","","1","1","1","166","0","0","99.01","403","0","404","24.5","0"
"AGAW3","08N","INS","BOLD:ACE4701","Arthropoda","Insecta","Lepidoptera","Crambidae","Crambinae","Crambus","Crambus multilinellus","1","1","1","166","0","0","99.55","449","0","450","31.78","0"
"AGAW3","08N","INS","BOLD:AAM9253","Arthropoda","Insecta","Diptera","Sciaridae","","Sciara","Sciara hebes","1","1","1","166","0","0","99.11","447","0","450","29.33","0"
"AGAW3","08N","INS","BOLD:AAG2460","Arthropoda","Insecta","Diptera","Anthomyiidae","Anthomyiinae","Hydrophoria","Hydrophoria lancifer","1","1","1","166","0","0","99.78","449","0","450","34","0"
"AGAW3","08N","NAR","no match","no match","no match","no match","no match","no match","no match","no match","no match","no match","no match","no match","no match","no match","no match","no match","no match","no match","no match","no match"
"AGAW3","08N","NIN","BOLD:AAN6394","Arthropoda","Arachnida","Araneae","Anyphaenidae","","Hibana","Hibana gracilis","1093","1057","1057","1","3.29","0","98.86","427.39","0","429.94","30.87","0"
"AGAW3","08N","NIN","BOLD:AAB7914","Arthropoda","Collembola","Symphypleona","Bourletiellidae","","Bourletiella","","1000","765","765","2","23.5","0","99.52","444.89","0","446.32","38.13","0"
"AGAW3","08N","NIN","BOLD:AAC9044","Arthropoda","Arachnida","Araneae","Salticidae","","Salticus","Salticus scenicus","791","736","736","3","6.95","0","99.32","410.01","0","422.03","33.37","0"
"AGAW3","08N","NIN","BOLD:ABW2639","Arthropoda","Arachnida","Trombidiformes","Anystidae","","","","685","599","599","4","12.55","0","99.38","445.81","0","445.12","31.29","0"
"AGAW3","08N","NIN","BOLD:AAH6630","Arthropoda","Collembola","Entomobryomorpha","Entomobryidae","Entomobryinae","Entomobrya","Entomobrya unostrigata","553","481","481","5","13.02","0","99.45","438.11","0","439.16","38.93","0"
"AGAW3","08N","NIN","BOLD:AAI4346","Arthropoda","Arachnida","Opiliones","Phalangiidae","","Paroligolophus, Phalangium","Paroligolophus agrestis, Phalangium opilio","445","405","405","6","8.99","0","99.25","439.91","0","441.6","36.39","0"
"AGAW3","08N","NIN","BOLD:ACC0359","Arthropoda","Collembola","Symphypleona","Bourletiellidae","","Bourletiella","","216","185","185","7","14.35","0","99.25","436.92","0","440.78","37.61","0"
"AGAW3","08N","NIN","BOLD:ACI9908","Arthropoda","Collembola","Symphypleona","Bourletiellidae","","","","207","173","173","8","16.43","0","99.44","435.39","0","437.87","34.85","0"
"AGAW3","08N","NIN","BOLD:AAA7246","Arthropoda","Collembola","Entomobryomorpha","Entomobryidae","Entomobryinae","Entomobrya","Entomobrya nicoleti, Entomobrya multifasciata, Entomobrya intermedia","156","132","132","9","15.38","0","99.1","444.19","0","445.22","43.43","0"
"AGAW3","08N","NIN","BOLD:ACZ1295","Arthropoda","Arachnida","Trombidiformes","Anystidae","","","","125","125","125","10","0","0","99.14","376.3","0","390.41","31.16","0"
"AGAW3","08N","NIN","BOLD:AAD1564","Arthropoda","Arachnida","Araneae","Clubionidae","","Clubiona","Clubiona abboti","96","89","89","11","7.29","0","98.2","423.75","0","426.39","31.99","0"
"AGAW3","08N","NIN","BOLD:ACV5609","","","","","","","","86","77","77","12","10.47","0","98.95","445.26","0","446.29","38.74","0"
"AGAW3","08N","NIN","BOLD:AAM7961","Arthropoda","Arachnida","Trombidiformes","Anystidae","","","","64","57","57","13","10.94","0","99.42","446.09","0","442.95","31.3","0"
"AGAW3","08N","NIN","BOLD:ACJ7454","Arthropoda","Collembola","Symphypleona","Bourletiellidae","","","","64","46","46","13","28.13","0","99.63","402.91","0","446.97","38.99","0"
"AGAW3","08N","NIN","BOLD:AAA8820","Arthropoda","Collembola","Entomobryomorpha","Entomobryidae","Lepidocyrtinae","Lepidocyrtus","Lepidocyrtus cyaneus","40","40","40","15","0","0","98.67","410.53","0","411.9","43.33","0"
"AGAW3","08N","NIN","BOLD:AAZ2181","Arthropoda","Collembola","Symphypleona","Bourletiellidae","","","","37","28","28","16","24.32","0","97.44","441.08","0","442.32","39.45","0"
"AGAW3","08N","NIN","BOLD:ACL2932","Arthropoda","Collembola","Symphypleona","Bourletiellidae","","Bourletiella","","35","35","35","17","0","0","99.11","426.11","0","427.77","38.36","0"
"AGAW3","08N","NIN","BOLD:ACV5610","","","","","","","","28","21","21","18","25","0","99.56","445.29","0","446.07","38.18","0"
"AGAW3","08N","NIN","BOLD:ACC8538","Arthropoda","Arachnida","Mesostigmata","Blattisociidae","","","","17","17","17","19","0","0","99.27","395.88","0","397.65","32.66","0"
"AGAW3","08N","NIN","BOLD:AAN6706","Arthropoda","Arachnida","Mesostigmata","Ascidae","","","","8","8","8","20","0","0","99.54","416","0","417.75","33.97","0"
"AGAW3","08N","NIN","BOLD:ACP9010","Arthropoda","Collembola","Symphypleona","Bourletiellidae","","","","5","5","5","21","0","0","98.88","445.6","0","447.2","38.64","0"
"AGAW3","08N","NIN","BOLD:ACM6751","Arthropoda","Arachnida","Trombidiformes","Anystidae","","","","5","5","5","21","0","0","99.56","394","0","431.8","29.17","0"
"AGAW3","08N","NIN","BOLD:AAB8452","Arthropoda","Collembola","Entomobryomorpha","Entomobryidae","Lepidocyrtinae","Lepidocyrtus","Lepidocyrtus paradoxus","5","5","5","21","0","0","99.73","449","0","450","42.13","0"
"AGAW3","08N","NIN","BOLD:AAA8824","Arthropoda","Collembola","Entomobryomorpha","Entomobryidae","Lepidocyrtinae","Lepidocyrtus","Lepidocyrtus violaceus","4","4","4","24","0","0","99.03","396.5","0","399.5","44.34","0"
"AGAW3","08N","NIN","BOLD:AAF9285","Arthropoda","Arachnida","Mesostigmata","Parasitidae","","","","4","4","4","24","0","0","98.47","393.75","0","395.25","35.17","0"
"AGAW3","08N","NIN","BOLD:ADD2059","Arthropoda","Arachnida","Mesostigmata","Phytoseiidae","","","","3","3","3","26","0","0","99.22","427","0","450","33.33","0"
"AGAW3","08N","NIN","BOLD:AAA9476","Arthropoda","Arachnida","Araneae","Linyphiidae","","Grammonota","Grammonota gentilis","2","2","2","27","0","0","98.91","417.5","0","418","30.23","0"
"AGAW3","08N","NIN","BOLD:ADE0357","Arthropoda","Arachnida","Mesostigmata","Phytoseiidae","","","","2","2","2","27","0","0","97.59","420","0","420.5","32.8","0"
"AGAW3","08N","NIN","BOLD:ADE0382","Arthropoda","Arachnida","Mesostigmata","Phytoseiidae","","","","2","2","2","27","0","0","98.33","449","0","450","33.11","0"
"AGAW3","08N","NIN","BOLD:AAA5267","Arthropoda","Collembola","Entomobryomorpha","Isotomidae","","","","1","1","1","30","0","0","99.55","449","0","448","38.62","0"
"AGAW3","08N","NIN","BOLD:AAN4465","","","","","","","","1","1","1","30","0","0","100","450","0","450","45.78","0"
"AGAW3","08N","NIN","BOLD:AAG5658","Arthropoda","Arachnida","Araneae","Mimetidae","","Mimetus","Mimetus epeiroides","1","1","1","30","0","0","97.11","450","0","450","34","0"
"AGAW3","08N","NIN","BOLD:AAO8147","Arthropoda","Arachnida","Trombidiformes","Eupodidae","","Eupodes","","1","1","1","30","0","0","99.2","377","0","378","27.51","0"
"AGAW3","08S","BAC","TAX:175768","Proteobacteria","Alphaproteobacteria","Rickettsiales","Rickettsiaceae",NA,"Wolbachia","Wolbachia pipientis","151","129","129","1","14.57","0","99.2","442.08","0","443.05","39.66","0"
"AGAW3","08S","BAC","TAX:91393","Proteobacteria","Alphaproteobacteria","Rickettsiales","Rickettsiaceae",NA,"Wolbachia","","68","68","68","2","0","0","99.47","390.59","0","424.28","39.15","0"
"AGAW3","08S","BAC","TAX:91391","Proteobacteria","Alphaproteobacteria","Rickettsiales","",NA,"","","27","26","26","3","3.7","0","98.96","416.67","0","446.19","39.75","0"
"AGAW3","08S","BAC","TAX:91389","Proteobacteria","","","",NA,"","","4","4","4","4","0","0","98.96","361.5","0","366","38.46","0"
"AGAW3","08S","CHO","no match","no match","no match","no match","no match","no match","no match","no match","no match","no match","no match","no match","no match","no match","no match","no match","no match","no match","no match","no match"
"AGAW3","08S","INS","BOLD:AAG1711","Arthropoda","Insecta","Diptera","Muscidae","Phaoniinae","Helina","Helina depuncta","31304","22878","22878","1","26.92","0","99.46","431.83","0","434.08","31.37","0"
"AGAW3","08S","INS","BOLD:AAA3453","Arthropoda","Insecta","Diptera","Anthomyiidae","Anthomyiinae","Delia","","4887","4336","4336","2","11.27","0","99.31","430.6","0","433.44","34.64","0"
"AGAW3","08S","INS","BOLD:AAB0373","Arthropoda","Insecta","Neuroptera","Chrysopidae","Chrysopinae","Chrysoperla","Chrysoperla sp. isolate 82, Chrysoperla zastrowi, Chrysoperla plorabunda, Chrysoperla sp. isolate 71, Chrysoperla agilis, Italochrysa italica, Chrysoperla sp. isolate 88, Chrysoperla lucasina, Chrysoperla carnea, Chrysoperla mediterranea, Chrysoperla pallida, Chrysoperla nipponensis, Chrysoperla sillemi","4872","3502","3502","3","28.12","0","99.43","436.81","0","437.72","31.37","0"
"AGAW3","08S","INS","BOLD:AAC0903","Arthropoda","Insecta","Diptera","Chironomidae","Chironominae","Chironomus","Chironomus acidophilus","4282","3302","3302","4","22.89","0","99.31","436.6","0","440.12","36.31","0"
"AGAW3","08S","INS","BOLD:AAC0635","Arthropoda","Insecta","Hemiptera","Miridae","Mirinae","Stenotus","Stenotus binotatus","3480","2516","2516","5","27.7","0","99.2","441.87","0","442.86","34.67","0"
"AGAW3","08S","INS","BOLD:ACZ5284","Arthropoda","Insecta","Diptera","Sciomyzidae","Sciomyzinae","Dictya","Dictya stricta, Dictya expansa, Dictya floridensis, Dictya borealis, Dictya gaigei, Dictya sp. TAW1, Dictya umbroides, Dictya texensis, Dictya steyskali, Dictya pictipes","2781","1988","1988","6","28.51","0","99.12","443.65","0","444.67","35.82","0"
"AGAW3","08S","INS","BOLD:AAP8739","Arthropoda","Insecta","Diptera","Platystomatidae","Platystomatinae","Rivellia","Rivellia steyskali","2714","1801","1801","7","33.64","0","99.25","441.61","0","443.68","37.97","0"
"AGAW3","08S","INS","BOLD:AAE0844","Arthropoda","Insecta","Hemiptera","Pentatomidae","Pentatominae","Euschistus","Euschistus variolarius","2706","2273","2273","8","16","0","98.98","425.22","0","426.43","36.65","0"
"AGAW3","08S","INS","BOLD:AAD2041","Arthropoda","Insecta","Diptera","Dolichopodidae","Sciapodinae","Condylostylus","Condylostylus inermis, Condylostylus nigrofemoratus, Condylostylus connectans, Condylostylus caudatus, Condylostylus flavipes","1983","1517","1517","9","23.5","0","99.38","437.54","0","438.83","35.98","0"
"AGAW3","08S","INS","BOLD:AAA4277","Arthropoda","Insecta","Diptera","Syrphidae","Syrphinae","Toxomerus","Toxomerus marginatus, Toxomerus sp. CR-3, Toxomerus CR-3","1583","1061","1061","10","32.98","0","99.4","444.95","0","445.81","30.64","0"
"AGAW3","08S","INS","BOLD:AAF9771","Arthropoda","Insecta","Diptera","Hybotidae","Tachydromiinae","Platypalpus","Platypalpus longiseta, Platypalpus trivialis, Platypalpus sp. 5, Platypalpus sp. 17, Platypalpus pallidiventris","1420","1017","1017","11","28.38","0","99.36","445.41","0","446.17","35.1","0"
"AGAW3","08S","INS","BOLD:AAG5507","Arthropoda","Insecta","Diptera","Chironomidae","Chironominae","Chironomus","Chironomus atrella","1401","1351","1351","12","3.57","0","98.82","425.15","0","429.29","36.05","0"
"AGAW3","08S","INS","BOLD:AAD0484","Arthropoda","Insecta","Diptera","Chironomidae","","","","1287","1130","1130","13","12.2","0","98.86","445.33","0","446.66","34.77","0"
"AGAW3","08S","INS","BOLD:ABW3687","Arthropoda","Insecta","Diptera","Lauxaniidae","Homoneurinae","Homoneura","Homoneura bispina","1270","921","921","14","27.48","0","98.53","442","0","443.09","34.85","0"
"AGAW3","08S","INS","BOLD:AAB7612","Arthropoda","Insecta","Lepidoptera","Noctuidae","Noctuinae","Apamea","Apamea alia","1024","959","959","15","6.35","0","99.07","420.86","0","422.74","30.75","0"
"AGAW3","08S","INS","BOLD:AAG7036","Arthropoda","Insecta","Diptera","Agromyzidae","","","","938","833","833","16","11.19","0","98.84","388.38","0","392.54","33.31","0"
"AGAW3","08S","INS","BOLD:AAL7359","Arthropoda","Insecta","Diptera","Chironomidae","Chironominae","Tanytarsus","","907","765","765","17","15.66","0","99.17","437.99","0","439.96","36.11","0"
"AGAW3","08S","INS","BOLD:ABV9293","Arthropoda","Insecta","Diptera","Chironomidae","Chironominae","Dicrotendipes","","886","733","733","18","17.27","0","98.88","355.64","0","422.09","35.54","0"
"AGAW3","08S","INS","BOLD:AAC9132","Arthropoda","Insecta","Diptera","Culicidae","Culicinae","Culiseta","Culiseta inornata","847","832","832","19","1.77","0","98.75","401.07","0","403.71","32.16","0"
"AGAW3","08S","INS","BOLD:AAG2868","Arthropoda","Insecta","Hemiptera","Cicadellidae","Typhlocybinae","Empoasca","Empoasca sp., Empoasca fabae","641","410","410","20","36.04","0","99.66","442.13","0","448.41","32.43","0"
"AGAW3","08S","INS","BOLD:AAN5332","Arthropoda","Insecta","Diptera","Chironomidae","Chironominae","Polypedilum","","430","390","390","21","9.3","0","98.07","426.3","0","427.93","33.95","0"
"AGAW3","08S","INS","BOLD:ABV2554","Arthropoda","Insecta","Hemiptera","Cicadellidae","Deltocephalinae","Sorhoanus, Stenometopiellus","Stenometopiellus sigillatus, Sorhoanus uhleri, Sorhoanus pascuellus","425","329","329","22","22.59","0","99.39","444.4","0","445.57","39.4","0"
"AGAW3","08S","INS","BOLD:AAA4201","Arthropoda","Insecta","Lepidoptera","Noctuidae","Noctuinae","Apamea","Apamea sp., Apamea sordens, Apamea sp. nr. sordens, Apamea digitula","398","336","336","23","15.58","0","99.25","446.87","0","447.75","31.92","0"
"AGAW3","08S","INS","BOLD:AAG2511","Arthropoda","Insecta","Diptera","Anthomyiidae","Anthomyiinae","Delia","Delia platura","383","353","353","24","7.83","0","98.84","444.15","0","446.79","33.18","0"
"AGAW3","08S","INS","BOLD:AAA2482","Arthropoda","Insecta","Lepidoptera","Noctuidae","Noctuinae","Mythimna","Mythimna unipuncta","374","345","345","25","7.75","0","99.05","417.43","0","420.55","30.19","0"
"AGAW3","08S","INS","BOLD:AAA1513","Arthropoda","Insecta","Lepidoptera","Plutellidae","Plutellinae","Plutella, Rhigognostis","Plutella xylostella, Rhigognostis sp.","348","271","271","26","22.13","0","99.25","447.09","0","448.01","32.24","0"
"AGAW3","08S","INS","BOLD:AAH4205","Arthropoda","Insecta","Diptera","Chloropidae","Chloropinae","Meromyza","Meromyza nigriseta, Meromyza sp.","345","234","234","27","32.17","0","99.52","446.62","0","447.57","32.87","0"
"AGAW3","08S","INS","BOLD:AAN5342","Arthropoda","Insecta","Diptera","Chironomidae","Orthocladiinae","Heleniella","","334","313","313","28","6.29","0","98.75","418.24","0","427.12","35.09","0"
"AGAW3","08S","INS","BOLD:ADH9832","","","","","","","","313","249","249","29","20.45","0","98.94","448.15","0","449.13","33.42","0"
"AGAW3","08S","INS","BOLD:AAH3661","Arthropoda","Insecta","Diptera","Cecidomyiidae","","","","290","272","272","30","6.21","0","99.13","431.44","0","433.1","25.36","0"
"AGAW3","08S","INS","BOLD:AAU6533","Arthropoda","Insecta","Diptera","Phoridae","Metopininae","Megaselia","","278","232","232","31","16.55","0","99.69","355.88","0","444.01","32.21","0"
"AGAW3","08S","INS","BOLD:AAA7661","Arthropoda","Insecta","Diptera","Culicidae","Culicinae","Culiseta, Culex","Culiseta melanura, Culex articularis, Culex restuans, Culex acharistus","244","235","235","32","3.69","0","99.18","434.91","0","437.16","32.83","0"
"AGAW3","08S","INS","BOLD:AAJ2399","Arthropoda","Insecta","Hemiptera","Miridae","Phylinae","Amblytylus","Amblytylus nasutus","226","130","130","33","42.48","0","99.76","447.86","0","448.38","39.99","0"
"AGAW3","08S","INS","BOLD:AAN5341","Arthropoda","Insecta","Diptera","Chironomidae","Orthocladiinae","Camptocladius","Camptocladius stercorarius","209","205","205","34","1.91","0","99.04","411.37","0","416.34","35.53","0"
"AGAW3","08S","INS","BOLD:AAV5688","Arthropoda","Insecta","Diptera","Cecidomyiidae","","","","162","146","146","35","9.88","0","99.47","433.04","0","434.15","34.33","0"
"AGAW3","08S","INS","BOLD:AAG6432","Arthropoda","Insecta","Diptera","Ceratopogonidae","Forcipomyiinae","Atrichopogon","","134","118","118","36","11.94","0","98.6","440.51","0","441.9","33.97","0"
"AGAW3","08S","INS","BOLD:AAH3644","Arthropoda","Insecta","Diptera","Cecidomyiidae","","","","133","119","119","37","10.53","0","99.35","439.89","0","441.28","27.18","0"
"AGAW3","08S","INS","BOLD:ABA4245","Arthropoda","Insecta","Hemiptera","Miridae","Phylinae","Atractotomus","Atractotomus mali","131","120","120","38","8.4","0","99.2","426.7","0","427.79","41.9","0"
"AGAW3","08S","INS","BOLD:AAB1098","Arthropoda","Insecta","Diptera","Culicidae","Culicinae","Ochlerotatus, Aedes","Aedes excrucians, Aedes cantans, Aedes stimulans, Aedes euedes, Aedes riparius, Aedes annulipes, Ochlerotatus cantans, Aedes fitchii, Aedes sp., Ochlerotatus annulipes, Ochlerotatus excrucians","129","120","120","39","6.98","0","98.58","430.27","0","431.49","34.84","0"
"AGAW3","08S","INS","BOLD:ABY5318","Arthropoda","Insecta","Diptera","Chloropidae","Oscinellinae","Incertella, Rhopalopterum","Rhopalopterum umbrosum, Rhopalopterum luteiceps, Incertella bispina, Rhopalopterum painteri","120","83","83","40","30.83","0","99.31","445.85","0","446.86","31.89","0"
"AGAW3","08S","INS","BOLD:ABW1511","Arthropoda","Insecta","Diptera","Ceratopogonidae","","","","102","56","56","41","45.1","0","97.11","433.56","0","434.7","36.55","0"
"AGAW3","08S","INS","BOLD:AAH0134","Arthropoda","Insecta","Coleoptera","Phalacridae","Phalacrinae","Stilbus","Stilbus apicialis","88","77","77","42","12.5","0","99.16","429.05","0","429.94","31.84","0"
"AGAW3","08S","INS","BOLD:AAG4827","Arthropoda","Insecta","Diptera","Anthomyzidae","Anthomyzinae","Mumetopia","Mumetopia occipitalis","86","83","83","43","3.49","0","98.58","438.09","0","439.48","36.41","0"
"AGAW3","08S","INS","BOLD:AAG5192","Arthropoda","Insecta","Coleoptera","Curculionidae","Entiminae","Barypeithes","Barypeithes pellucidus","81","68","68","44","16.05","0","99.47","435.36","0","436.59","39.29","0"
"AGAW3","08S","INS","BOLD:AAV5575","Arthropoda","Insecta","Diptera","Cecidomyiidae","","","CecidInt30 sp. BOLD:AAV5575","76","55","55","45","27.63","0","99.65","444.99","0","446.22","25.92","0"
"AGAW3","08S","INS","BOLD:AAG6934","Arthropoda","Insecta","Diptera","Hybotidae","","","","75","74","74","46","1.33","0","99.12","420.71","0","422.8","38.33","0"
"AGAW3","08S","INS","BOLD:AAH2372","Arthropoda","Insecta","Coleoptera","Elateridae","Elaterinae","Dalopius","Dalopius pallidus","72","65","65","47","9.72","0","99.44","438.58","0","439.76","36.68","0"
"AGAW3","08S","INS","BOLD:ABV2748","Arthropoda","Insecta","Hymenoptera","Braconidae","Alysiinae","","","71","63","63","48","11.27","0","99.35","438.99","0","439.89","28.15","0"
"AGAW3","08S","INS","BOLD:AAN5183","Arthropoda","Insecta","Diptera","Cecidomyiidae","","","","66","58","58","49","12.12","0","98.45","445.97","0","446.58","27.87","0"
"AGAW3","08S","INS","BOLD:AAV6213","Arthropoda","Insecta","Diptera","Chironomidae","Chironominae","Paratanytarsus","","64","63","63","50","1.56","0","99.08","406.19","0","420.42","34.25","0"
"AGAW3","08S","INS","BOLD:ACE0554","Arthropoda","Insecta","Diptera","Phoridae","Metopininae","Megaselia","","64","53","53","50","17.19","0","99.48","445.41","0","446.77","32.06","0"
"AGAW3","08S","INS","BOLD:AAU7040","Arthropoda","Insecta","Coleoptera","Corylophidae","Corylophinae","Orthoperus","Orthoperus scutellaris","61","52","52","52","14.75","0","98.23","422.62","0","429.57","33.1","0"
"AGAW3","08S","INS","BOLD:AAH3300","Arthropoda","Insecta","Coleoptera","Coccinellidae","Coccinellinae","Cycloneda","Cycloneda munda","61","61","61","52","0","0","98.65","418.85","0","426.16","35.53","0"
"AGAW3","08S","INS","BOLD:AAG0464","Arthropoda","Insecta","Diptera","Heleomyzidae","Heleomyzinae","","","60","60","60","54","0","0","97.8","409.95","0","411.92","36.26","0"
"AGAW3","08S","INS","BOLD:AAG1741","Arthropoda","Insecta","Diptera","Muscidae","Coenosiinae","Coenosia","","55","47","47","55","14.55","0","98.93","444.38","0","445.27","33.83","0"
"AGAW3","08S","INS","BOLD:ABV1455","Arthropoda","Insecta","Diptera","Chironomidae","Chironominae","","","53","52","52","56","1.89","0","98.15","430.64","0","432.94","31.6","0"
"AGAW3","08S","INS","BOLD:AAH3653","Arthropoda","Insecta","Diptera","Cecidomyiidae","","","","51","49","49","57","3.92","0","98.97","433.41","0","435.12","25.77","0"
"AGAW3","08S","INS","BOLD:AAN5148","Arthropoda","Insecta","Diptera","Ceratopogonidae","Forcipomyiinae","Forcipomyia","Forcipomyia bipunctata, Forcipomyia sp. CK-2012","50","50","50","58","0","0","98.88","421.02","0","423.78","35.55","0"
"AGAW3","08S","INS","BOLD:AAB8911","Arthropoda","Insecta","Coleoptera","Curculionidae","Entiminae","Polydrusus","Polydrusus formosus","48","33","33","59","31.25","0","99.48","442.9","0","443.52","34.52","0"
"AGAW3","08S","INS","BOLD:AAN5199","Arthropoda","Insecta","Diptera","Cecidomyiidae","","","CecidInt20 sp. BOLD:AAN5199","43","37","37","60","13.95","0","99.19","428.05","0","444.74","26.81","0"
"AGAW3","08S","INS","BOLD:ABA0822","Arthropoda","Insecta","Diptera","Cecidomyiidae","","","","42","39","39","61","7.14","0","98.7","434.45","0","435.86","26.57","0"
"AGAW3","08S","INS","BOLD:AAG8493","Arthropoda","Insecta","Diptera","Drosophilidae","Drosophilinae","Scaptomyza","Scaptomyza pallida","37","37","37","62","0","0","98.7","399.81","0","401.68","32.96","0"
"AGAW3","08S","INS","BOLD:AAH0135","Arthropoda","Insecta","Coleoptera","Phalacridae","Phalacrinae","Acylomus","Acylomus pugetanus, Acylomus ergoti","29","29","29","63","0","0","99.11","427.17","0","429.52","31.49","0"
"AGAW3","08S","INS","BOLD:AAB6773","Arthropoda","Insecta","Hymenoptera","Braconidae","Euphorinae","Microctonus, Perilitus","Microctonus aethiopoides, Microctonus sp., Perilitus aethiopoides","29","22","22","63","24.14","0","99.26","447.86","0","448.31","27.82","0"
"AGAW3","08S","INS","BOLD:ABA7871","Arthropoda","Insecta","Diptera","Cecidomyiidae","","","","28","27","27","65","3.57","0","99.23","439.32","0","439.89","26.42","0"
"AGAW3","08S","INS","BOLD:AAN5160","Arthropoda","Insecta","Diptera","Ceratopogonidae","Forcipomyiinae","Atrichopogon","","27","26","26","66","3.7","0","99.23","434.22","0","437.3","37.03","0"
"AGAW3","08S","INS","BOLD:AAG3236","Arthropoda","Insecta","Diptera","Phoridae","Phorinae","Conicera, Diplonevra","Diplonevra nitidula","27","27","27","66","0","0","99.54","447.33","0","449.44","31.96","0"
"AGAW3","08S","INS","BOLD:AAL7819","Arthropoda","Insecta","Diptera","Psychodidae","Psychodinae","Psychoda","Psychoda sp., Psychoda sp. BOLD:AAL7819","26","26","26","68","0","0","99.46","405.08","0","406.69","32.57","0"
"AGAW3","08S","INS","BOLD:AAN6447","Arthropoda","Insecta","Diptera","Sciaridae","","Corynoptera","Corynoptera perpusilla","26","24","24","68","7.69","0","99.2","447.85","0","449.46","36.32","0"
"AGAW3","08S","INS","BOLD:AAN5242","Arthropoda","Insecta","Diptera","Cecidomyiidae","","","","24","24","24","70","0","0","98.61","420.17","0","421.5","28.44","0"
"AGAW3","08S","INS","BOLD:AAN6154","Arthropoda","Insecta","Coleoptera","Latridiidae","Corticariinae","Melanophthalma","Melanophthalma inermis","24","21","21","70","12.5","0","98.95","430.63","0","431.75","31.18","0"
"AGAW3","08S","INS","BOLD:ACC8511","Arthropoda","Insecta","Diptera","Cecidomyiidae","Cecidomyiinae","Sitodiplosis","Sitodiplosis mosellana","23","22","22","72","4.35","0","99.41","428.61","0","430.3","26.54","0"
"AGAW3","08S","INS","BOLD:AAA4781","Arthropoda","Insecta","Hymenoptera","Braconidae","Microgastrinae","Glyptapanteles","Glyptapanteles carlossarmientoi","23","23","23","72","0","0","99.11","436.65","0","439.39","24.3","0"
"AGAW3","08S","INS","BOLD:AAU9508","Arthropoda","Insecta","Hymenoptera","Dryinidae","Anteoninae","Anteon","Anteon tenuicorne, Anteon cf. pubicorne, Anteon pubicorne","22","22","22","74","0","0","99.27","431.18","0","433.09","28.94","0"
"AGAW3","08S","INS","BOLD:AAG2754","Arthropoda","Insecta","Diptera","Ephydridae","Ilytheinae","Nostima","Nostima picta, Nostima approximata","19","19","19","75","0","0","98.79","438.32","0","440.05","32.82","0"
"AGAW3","08S","INS","BOLD:AAQ0558","Arthropoda","Insecta","Thysanoptera","Thripidae","Thripinae","Anaphothrips","Anaphothrips obscurus","17","17","17","76","0","0","99.46","434.59","0","437.18","31.37","0"
"AGAW3","08S","INS","BOLD:ACK3431","Arthropoda","Insecta","Diptera","Cecidomyiidae","","","","17","17","17","76","0","0","99.04","430.53","0","431.94","32.9","0"
"AGAW3","08S","INS","BOLD:ADH4889","Arthropoda","Insecta","Diptera","","","","","16","15","15","78","6.25","0","98.96","448.88","0","450","35.56","0"
"AGAW3","08S","INS","BOLD:ADF6661","Arthropoda","Insecta","Diptera","Cecidomyiidae","","","","16","16","16","78","0","0","99.24","418.44","0","448.81","29.84","0"
"AGAW3","08S","INS","BOLD:ACC9085","Arthropoda","Insecta","Diptera","Limoniidae","Limoniinae","Dicranomyia","","14","14","14","80","0","0","99.56","441","0","442.5","31.84","0"
"AGAW3","08S","INS","BOLD:AAG7891","Arthropoda","Insecta","Hymenoptera","Platygastridae","Scelioninae","Telenomus","Telenomus podisi","14","13","13","80","7.14","0","98.99","438.64","0","450","27.33","0"
"AGAW3","08S","INS","BOLD:AAN9511","Arthropoda","Insecta","Coleoptera","Staphylinidae","Tachyporinae","Tachyporus","Tachyporus dispar, Tachyporus rulomoides, Tachyporus chrysomelinus","12","12","12","82","0","0","99.09","416","0","418.92","32.13","0"
"AGAW3","08S","INS","BOLD:ACE5981","","","","","","","","12","11","11","82","8.33","0","98.85","431.58","0","433.33","36.37","0"
"AGAW3","08S","INS","BOLD:AAN5238","Arthropoda","Insecta","Diptera","Cecidomyiidae","","","","11","10","10","84","9.09","0","99.54","430.18","0","431.18","26.04","0"
"AGAW3","08S","INS","BOLD:ACC8079","Arthropoda","Insecta","Diptera","Cecidomyiidae","","","","11","11","11","84","0","0","99.13","448","0","450","29.11","0"
"AGAW3","08S","INS","BOLD:AAV1055","Arthropoda","Insecta","Diptera","Stratiomyidae","Beridinae","Allognosta","Allognosta fuscitarsis","11","11","11","84","0","0","99.16","422","0","423.36","35.87","0"
"AGAW3","08S","INS","BOLD:AAN5267","Arthropoda","Insecta","Diptera","Cecidomyiidae","","","","11","9","9","84","18.18","0","99.63","440","0","441.45","30.47","0"
"AGAW3","08S","INS","BOLD:AAN5260","Arthropoda","Insecta","Diptera","Cecidomyiidae","","","","10","10","10","88","0","0","99.21","403.4","0","404.9","25.94","0"
"AGAW3","08S","INS","BOLD:AAM9255","Arthropoda","Insecta","Diptera","Sciaridae","","Pseudolycoriella","","10","10","10","88","0","0","98.88","428.2","0","430.8","31.21","0"
"AGAW3","08S","INS","BOLD:AAD1485","Arthropoda","Insecta","Diptera","Chironomidae","Chironominae","Paratanytarsus","Paratanytarsus grimmii, Paratanytarsus sp.","9","9","9","90","0","0","99.53","447.67","0","448.22","33.89","0"
"AGAW3","08S","INS","BOLD:ABU5525","Arthropoda","Insecta","Diptera","Chironomidae","Orthocladiinae","Limnophyes","Limnophyes sp. 14ES","9","9","9","90","0","0","98.95","398.22","0","401.33","37.2","0"
"AGAW3","08S","INS","BOLD:AAV1295","Arthropoda","Insecta","Diptera","Sciaridae","","Bradysia","Bradysia difformis, Bradysia impatiens","9","8","8","90","11.11","0","99.51","414.33","0","438.11","31.38","0"
"AGAW3","08S","INS","BOLD:AAH0152","Arthropoda","Insecta","Coleoptera","Cantharidae","Cantharinae","Cantharis","Cantharis rufa","8","8","8","93","0","0","98.56","432.63","0","436.5","35.27","0"
"AGAW3","08S","INS","BOLD:AAU6967","Arthropoda","Insecta","Coleoptera","Chrysomelidae","Galerucinae","Psylliodes","Psylliodes affinis","7","6","6","94","14.29","0","99.51","438","0","447.43","35.06","0"
"AGAW3","08S","INS","BOLD:AAG6745","Arthropoda","Insecta","Diptera","Calliphoridae","Calliphorinae","Pollenia","Pollenia pediculata","7","7","7","94","0","0","99.12","426.57","0","427.86","33.44","0"
"AGAW3","08S","INS","BOLD:ABA0863","Arthropoda","Insecta","Diptera","Cecidomyiidae","","","","6","6","6","96","0","0","99.34","433","0","433.67","29.1","0"
"AGAW3","08S","INS","BOLD:ABA0856","Arthropoda","Insecta","Diptera","Cecidomyiidae","","","","6","4","4","96","33.33","0","99.22","446.33","0","446.33","28.98","0"
"AGAW3","08S","INS","BOLD:ABU5520","Arthropoda","Insecta","Diptera","Sciaridae","","Corynoptera","","6","6","6","96","0","0","98.21","417.67","0","420.33","35.1","0"
"AGAW3","08S","INS","BOLD:AAL7815","Arthropoda","Insecta","Diptera","Psychodidae","Psychodinae","Psychoda","","6","5","5","96","16.67","0","99.07","449.33","0","450","34.41","0"
"AGAW3","08S","INS","BOLD:AAP2527","Arthropoda","Insecta","Diptera","Mycetophilidae","Mycetophilinae","Brevicornu","Brevicornu fissicauda","6","3","3","96","50","0","99.59","450","0","450","36.96","0"
"AGAW3","08S","INS","BOLD:AAA7636","Arthropoda","Insecta","Hymenoptera","Braconidae","Alysiinae","Dinotrema","Dinotrema jft02, Dinotrema jft01","5","5","5","101","0","0","98.31","413.2","0","415.6","23.63","0"
"AGAW3","08S","INS","BOLD:AAM7855","Arthropoda","Insecta","Thysanoptera","Thripidae","Thripinae","Frankliniella","Frankliniella tritici","5","5","5","101","0","0","98.68","443.6","0","446","31.34","0"
"AGAW3","08S","INS","BOLD:ABU5546","Arthropoda","Insecta","Diptera","Psychodidae","Psychodinae","Psychoda","","5","5","5","101","0","0","98.78","443.2","0","444.6","31.17","0"
"AGAW3","08S","INS","BOLD:AAF9187","Arthropoda","Insecta","Coleoptera","Curculionidae","Entiminae","Phyllobius","Phyllobius oblongus","5","4","4","101","20","0","99.78","449.6","0","450","38.98","0"
"AGAW3","08S","INS","BOLD:AAM9254","Arthropoda","Insecta","Diptera","Sciaridae","","Bradysia","Bradysia pallipes","5","5","5","101","0","0","98.67","414.4","0","415.6","33.04","0"
"AGAW3","08S","INS","BOLD:AAY6397","Arthropoda","Insecta","Diptera","Cecidomyiidae","","","","4","4","4","106","0","0","97.82","415.75","0","416.75","27.76","0"
"AGAW3","08S","INS","BOLD:AAA5299","Arthropoda","Insecta","Diptera","Chironomidae","Orthocladiinae","Cricotopus","Cricotopus laetus, Cricotopus relucens, Cricotopus trifasciatus, Cricotopus sp., Cricotopus sp. 3ES, Cricotopus sylvestris, Cricotopus glacialis","4","4","4","106","0","0","98.88","441","0","443","34.59","0"
"AGAW3","08S","INS","BOLD:AAG9897","Arthropoda","Insecta","Dermaptera","Forficulidae","Forficulinae","Forficula","Forficula auricularia-A","4","4","4","106","0","0","99.12","428.5","0","450","45.78","0"
"AGAW3","08S","INS","BOLD:AAA2632","Arthropoda","Insecta","Lepidoptera","Noctuidae","Noctuinae","Noctua","Noctua atlantica, Noctua pronuba","4","4","4","106","0","0","98.44","448.75","0","450","31.61","0"
"AGAW3","08S","INS","BOLD:AAG8821","Arthropoda","Insecta","Hemiptera","Cicadellidae","Deltocephalinae","Doratura","Doratura stylata","4","4","4","106","0","0","99.15","415.5","0","418","34.73","0"
"AGAW3","08S","INS","BOLD:AAH3662","Arthropoda","Insecta","Diptera","Cecidomyiidae","","","CecidInt3 sp. BOLD:AAH3662","4","4","4","106","0","0","99.22","449","0","450","26.06","0"
"AGAW3","08S","INS","BOLD:AAN5220","Arthropoda","Insecta","Diptera","Cecidomyiidae","","","","4","4","4","106","0","0","98.13","443","0","446","26.57","0"
"AGAW3","08S","INS","BOLD:AAA4555","Arthropoda","Insecta","Orthoptera","Acrididae","Melanoplinae","Melanoplus","Melanoplus stupefactus, Melanoplus angustipennis, Melanoplus ponderosus, Melanoplus bruneri, Melanoplus frigidus, Melanoplus scapularis, Melanoplus cinereus, Melanoplus madeleineae, Melanoplus sanguinipes, Melanoplus nigrescens group, Melanoplus tunicae, Melanoplus walshi, Melanoplus femurrubrum, Melanoplus dawsoni, Melanoplus sp., Melanoplus fasciatus, Melanoplus bowditchi, Melanoplus stonei, Melanoplus bivittatus, Melanoplus huroni, Melanoplus thomasi, Melanoplus borealis, Melanoplus packardii","4","4","4","106","0","0","99.11","449.25","0","450","36.11","0"
"AGAW3","08S","INS","BOLD:AAU8556","Arthropoda","Insecta","Hymenoptera","Braconidae","Rogadinae","Aleiodes","","4","4","4","106","0","0","99.02","409.5","0","411.75","24.89","0"
"AGAW3","08S","INS","BOLD:AAN5225","Arthropoda","Insecta","Diptera","Cecidomyiidae","","","","3","3","3","115","0","0","98.29","449.67","0","450","28.52","0"
"AGAW3","08S","INS","BOLD:AAN5230","Arthropoda","Insecta","Diptera","Cecidomyiidae","","","","3","3","3","115","0","0","99.69","429.67","0","431.67","26.08","0"
"AGAW3","08S","INS","BOLD:ACE7444","Arthropoda","Insecta","Hemiptera","Miridae","Mirinae, Phylinae","Adelphocoris, Megalocoleus","Adelphocoris lineolatus, Megalocoleus molliculus","3","3","3","115","0","0","99.18","449.33","0","450","34.67","0"
"AGAW3","08S","INS","BOLD:AAI1549","Arthropoda","Insecta","Hymenoptera","Braconidae","Euphorinae","Meteorus","","3","3","3","115","0","0","98.62","431","0","433","31.22","0"
"AGAW3","08S","INS","BOLD:AAG9626","Arthropoda","Insecta","Diptera","Dolichopodidae","","","","3","3","3","115","0","0","98.43","403","0","406","36.19","0"
"AGAW3","08S","INS","BOLD:AAH1867","Arthropoda","Insecta","Hymenoptera","Ichneumonidae","Ctenopelmatinae","","","3","3","3","115","0","0","99.04","401.33","0","402.67","25.06","0"
"AGAW3","08S","INS","BOLD:AAI4303","Arthropoda","Insecta","Diptera","Chironomidae","Chironominae","Chironomus","Chironomus melanescens","3","3","3","115","0","0","97.98","445.67","0","449.33","34.64","0"
"AGAW3","08S","INS","BOLD:AAA5953","Arthropoda","Insecta","Lepidoptera","Gelechiidae","Gelechiinae","Coleotechnites","Coleotechnites sp., Coleotechnites piceaella, Coleotechnites laricis, Coleotechnites sp. 3, Coleotechnites carbonaria, Coleotechnites granti, Coleotechnites macleodi, Coleotechnites gibsonella, Coleotechnites coniferella, Coleotechnites martini","3","3","3","115","0","0","99.85","449.33","0","450","34.37","0"
"AGAW3","08S","INS","BOLD:ABW0395","Arthropoda","Insecta","Diptera","Tachinidae","Phasiinae","Phasia","Phasia fenestrata","2","2","2","123","0","0","99.22","448","0","450","30.78","0"
"AGAW3","08S","INS","BOLD:ACJ6954","Arthropoda","Insecta","Diptera","Cecidomyiidae","","","","2","2","2","123","0","0","98.82","338.5","0","450","26.78","0"
"AGAW3","08S","INS","BOLD:AAG7276","Arthropoda","Insecta","Diptera","Sphaeroceridae","Limosininae","Leptocera","Leptocera erythrocera","2","2","2","123","0","0","97.28","441.5","0","442.5","33.33","0"
"AGAW3","08S","INS","BOLD:ABA1215","Arthropoda","Insecta","Diptera","Sciaridae","","Lycoriella","Lycoriella sativae","2","2","2","123","0","0","98.35","435.5","0","439","35.86","0"
"AGAW3","08S","INS","BOLD:AAG9668","Arthropoda","Insecta","Diptera","Dolichopodidae","Diaphorinae","Chrysotus","Chrysotus sp., Chrysotus sp. nr. obliquus","2","2","2","123","0","0","99.22","449.5","0","450","33.11","0"
"AGAW3","08S","INS","BOLD:AAN5497","Arthropoda","Insecta","Diptera","Anthomyiidae","Pegomyinae","Pegomya","Pegomya solennis, Pegomya bicolor","2","2","2","123","0","0","98.66","448.5","0","450","29.33","0"
"AGAW3","08S","INS","BOLD:AAA3867","Arthropoda","Insecta","Lepidoptera","Noctuidae","Noctuinae","Apamea","Apamea amputatrix","2","2","2","123","0","0","99.44","449.5","0","450","33.78","0"
"AGAW3","08S","INS","BOLD:AAG1717","Arthropoda","Insecta","Diptera","Muscidae","Phaoniinae","Helina","Helina troene","2","2","2","123","0","0","99.89","441","0","443.5","31.23","0"
"AGAW3","08S","INS","BOLD:AAA2815","Arthropoda","Insecta","Lepidoptera","Noctuidae","Noctuinae","Mythimna","Mythimna yukonensis, Mythimna oxygala","2","2","2","123","0","0","99.21","445","0","448.5","33.11","0"
"AGAW3","08S","INS","BOLD:AAN5198","Arthropoda","Insecta","Diptera","Cecidomyiidae","","","","2","2","2","123","0","0","98.88","436.5","0","437","26.19","0"
"AGAW3","08S","INS","BOLD:AAC4197","Arthropoda","Insecta","Diptera","Chironomidae","Orthocladiinae","Paraphaenocladius","Paraphaenocladius impensus","2","2","2","123","0","0","99.44","449.5","0","450","35.33","0"
"AGAW3","08S","INS","BOLD:AAH0256","Arthropoda","Insecta","Coleoptera","Latridiidae","Corticariinae","Corticaria, Corticarina","","2","2","2","123","0","0","99","449.5","0","450","31.56","0"
"AGAW3","08S","INS","BOLD:ACF7023","Arthropoda","Insecta","Diptera","Dolichopodidae","","","","2","2","2","123","0","0","97.79","226","0","393.5","35.62","0"
"AGAW3","08S","INS","BOLD:ABV9322","Arthropoda","Insecta","Diptera","Cecidomyiidae","","","","2","2","2","123","0","0","97.89","450","0","450","27.89","0"
"AGAW3","08S","INS","BOLD:AAA3654","Arthropoda","Insecta","Lepidoptera","Crambidae","Crambinae","Crambus","Crambus bigelovi, Crambus johnsoni, Crambus praefectellus","2","2","2","123","0","0","99.47","401.5","0","404","31.4","0"
"AGAW3","08S","INS","BOLD:AAP6471","Arthropoda","Insecta","Diptera","Sciaridae","","Pseudolycoriella","","2","2","2","123","0","0","98.77","447.5","0","450","36.33","0"
"AGAW3","08S","INS","BOLD:ACE4701","Arthropoda","Insecta","Lepidoptera","Crambidae","Crambinae","Crambus","Crambus multilinellus","2","2","2","123","0","0","99.89","449.5","0","450","32.11","0"
"AGAW3","08S","INS","BOLD:AAA3181","Arthropoda","Insecta","Diptera","Muscidae","Muscinae","Stomoxys","Stomoxys calcitrans","1","1","1","140","0","0","98.96","385","0","389","32.65","0"
"AGAW3","08S","INS","BOLD:AAH4135","Arthropoda","Insecta","Diptera","Chloropidae","Chloropinae","Thaumatomyia","Thaumatomyia glabra, Thaumatomyia grata, Thaumatomyia sp. 1 TAW","1","1","1","140","0","0","99.78","449","0","450","31.78","0"
"AGAW3","08S","INS","BOLD:AAA6292","Arthropoda","Insecta","Lepidoptera","Tortricidae","Tortricinae","Cnephasia","Cnephasia genitalana, Cnephasia stephensiana","1","1","1","140","0","0","100","449","0","450","35.78","0"
"AGAW3","08S","INS","BOLD:AAG1714","Arthropoda","Insecta","Diptera","Muscidae","Azeliinae","Muscina","Muscina pascuorum","1","1","1","140","0","0","99.77","430","0","450","34.44","0"
"AGAW3","08S","INS","BOLD:AAB9404","Arthropoda","Insecta","Lepidoptera","Tortricidae","Tortricinae","Archips","Archips rosana, Archips grisea","1","1","1","140","0","0","99.78","449","0","450","31.56","0"
"AGAW3","08S","INS","BOLD:ACL6253","Arthropoda","Insecta","Coleoptera","Carabidae","Harpalinae","Apenes","Apenes lucidula","1","1","1","140","0","0","100","418","0","418","30.62","0"
"AGAW3","08S","INS","BOLD:AAL4232","Arthropoda","Insecta","Diptera","Agromyzidae","Phytomyzinae","Phytoliriomyza","Phytoliriomyza arctica","1","1","1","140","0","0","98.1","368","0","368","32.07","0"
"AGAW3","08S","INS","BOLD:AAA4751","Arthropoda","Insecta","Diptera","Culicidae","Culicinae","Culex","Culex pipiens sl, Culex nr. pipiens, Culex pipiens, Culex quinquefasciatus, Culex sp.","1","1","1","140","0","0","99.14","350","0","350","33.71","0"
"AGAW3","08S","INS","BOLD:AAM9228","Arthropoda","Insecta","Diptera","Sciaridae","","","","1","1","1","140","0","0","97.87","423","0","432","34.72","0"
"AGAW3","08S","INS","BOLD:AAA8933","Arthropoda","Insecta","Coleoptera","Coccinellidae","Coccinellinae","Coccinella","Coccinella septempunctata","1","1","1","140","0","0","99.78","449","0","450","32.67","0"
"AGAW3","08S","INS","BOLD:AAG8839","Arthropoda","Insecta","Hemiptera","Cicadellidae","Deltocephalinae","Errastunus, Latalus","Latalus ocellaris, Errastunus ocellaris","1","1","1","140","0","0","99.78","450","0","450","35.78","0"
"AGAW3","08S","INS","BOLD:ACA9275","Arthropoda","Insecta","Diptera","Chironomidae","Chironominae","Chironomus","","1","1","1","140","0","0","100","448","0","450","35.56","0"
"AGAW3","08S","INS","BOLD:AAU6592","Arthropoda","Insecta","Diptera","Ceratopogonidae","Dasyheleinae","Dasyhelea","Dasyhelea arenivaga","1","1","1","140","0","0","99.73","369","0","369","35.23","0"
"AGAW3","08S","INS","BOLD:ACV2677","Arthropoda","Insecta","Diptera","Cecidomyiidae","","","","1","1","1","140","0","0","97.33","450","0","450","29.78","0"
"AGAW3","08S","INS","BOLD:ACU7906","Arthropoda","Insecta","Diptera","Chironomidae","Orthocladiinae","","","1","1","1","140","0","0","97.58","414","0","414","35.51","0"
"AGAW3","08S","INS","BOLD:AAU6681","Arthropoda","Insecta","Diptera","Tipulidae","Tipulinae","Nephrotoma","Nephrotoma cornicina","1","1","1","140","0","0","100","447","0","450","36.22","0"
"AGAW3","08S","INS","BOLD:AAA3889","Arthropoda","Insecta","Lepidoptera","Crambidae","Crambinae","Parapediasia","Parapediasia teterrellus","1","1","1","140","0","0","100","449","0","450","32.44","0"
"AGAW3","08S","INS","BOLD:AAZ5483","Arthropoda","Insecta","Diptera","Muscidae","Phaoniinae","Helina","Helina setiventris","1","1","1","140","0","0","99.78","449","0","450","29.11","0"
"AGAW3","08S","INS","BOLD:ACF8552","Arthropoda","Insecta","Coleoptera","Anthicidae","Anthicinae","Malporus","Malporus formicarius","1","1","1","140","0","0","99.73","364","0","394","34.77","0"
"AGAW3","08S","INS","BOLD:AAN8770","Arthropoda","Insecta","Diptera","Psychodidae","Psychodinae","Psychoda","Psychoda trinodulosa","1","1","1","140","0","0","99.11","448","0","450","33.11","0"
"AGAW3","08S","INS","BOLD:AAL2750","Arthropoda","Insecta","Diptera","Muscidae","Phaoniinae","Phaonia","Phaonia tuguriorum","1","1","1","140","0","0","99.76","422","0","425","32","0"
"AGAW3","08S","INS","BOLD:AAZ0285","Arthropoda","Insecta","Diptera","Cecidomyiidae","","","","1","1","1","140","0","0","97.56","450","0","450","25.56","0"
"AGAW3","08S","INS","BOLD:AAG2123","Arthropoda","Insecta","Diptera","Tachinidae","Dexiinae","Dinera","Dinera grisescens","1","1","1","140","0","0","99.08","434","0","434","32.26","0"
"AGAW3","08S","INS","BOLD:AAF1049","Arthropoda","Insecta","Diptera","Agromyzidae","Phytomyzinae","Cerodontha","Cerodontha fasciata","1","1","1","140","0","0","99.25","402","0","450","32","0"
"AGAW3","08S","INS","BOLD:ABX6186","Arthropoda","Insecta","Diptera","Tipulidae","Tipulinae","Nephrotoma","Nephrotoma ferruginea","1","1","1","140","0","0","98.4","375","0","378","35.19","0"
"AGAW3","08S","INS","BOLD:AAG5242","Arthropoda","Insecta","Coleoptera","Curculionidae","Entiminae","Barypeithes","Barypeithes pellucidus","1","1","1","140","0","0","98.67","450","0","450","39.78","0"
"AGAW3","08S","INS","BOLD:AAH1890","Arthropoda","Insecta","Hymenoptera","Ichneumonidae","Cryptinae","Stibeutes","Stibeutes brevicornis, Stibeutes heinemanni","1","1","1","140","0","0","100","450","0","450","29.56","0"
"AGAW3","08S","INS","BOLD:ACB1116","Arthropoda","Insecta","Diptera","Phoridae","Metopininae","Megaselia","","1","1","1","140","0","0","97.55","449","0","450","32.67","0"
"AGAW3","08S","INS","BOLD:ADF3937","","","","","","","","1","1","1","140","0","0","98.67","377","0","381","31.76","0"
"AGAW3","08S","INS","BOLD:ACV5892","Arthropoda","Insecta","Diptera","Cecidomyiidae","","","","1","1","1","140","0","0","97.78","450","0","450","25.56","0"
"AGAW3","08S","INS","BOLD:ACD9117","Arthropoda","Insecta","Hemiptera","Cicadellidae","Eurymelinae","Idiocerus","Idiocerus levis","1","1","1","140","0","0","99.71","349","0","413","31.23","0"
"AGAW3","08S","INS","BOLD:AAG2460","Arthropoda","Insecta","Diptera","Anthomyiidae","Anthomyiinae","Hydrophoria","Hydrophoria lancifer","1","1","1","140","0","0","97.33","449","0","450","33.78","0"
"AGAW3","08S","NAR","BOLD:AAI9663","Mollusca","Gastropoda","Stylommatophora","Agriolimacidae",NA,"Deroceras","Deroceras reticulatum","258","237","237","1","8.14","0","99.55","400.11","0","434.04","35.62","0"
"AGAW3","08S","NIN","BOLD:AAB7914","Arthropoda","Collembola","Symphypleona","Bourletiellidae","","Bourletiella","","955","749","749","1","21.57","0","99.45","443.91","0","445.51","38.12","0"
"AGAW3","08S","NIN","BOLD:ABW2639","Arthropoda","Arachnida","Trombidiformes","Anystidae","","","","703","595","595","2","15.36","0","99.37","446.27","0","445.41","31.32","0"
"AGAW3","08S","NIN","BOLD:ACI9908","Arthropoda","Collembola","Symphypleona","Bourletiellidae","","","","415","346","346","3","16.63","0","99.38","435.52","0","437.71","34.88","0"
"AGAW3","08S","NIN","BOLD:AAA7246","Arthropoda","Collembola","Entomobryomorpha","Entomobryidae","Entomobryinae","Entomobrya","Entomobrya nicoleti, Entomobrya multifasciata, Entomobrya intermedia","253","211","211","4","16.6","0","99.09","442.77","0","444.01","43.42","0"
"AGAW3","08S","NIN","BOLD:ACC0359","Arthropoda","Collembola","Symphypleona","Bourletiellidae","","Bourletiella","","218","178","178","5","18.35","0","99.08","435.17","0","438.98","37.77","0"
"AGAW3","08S","NIN","BOLD:AAH6630","Arthropoda","Collembola","Entomobryomorpha","Entomobryidae","Entomobryinae","Entomobrya","Entomobrya unostrigata","217","194","194","6","10.6","0","99.46","442.04","0","443.26","38.82","0"
"AGAW3","08S","NIN","BOLD:ACZ1295","Arthropoda","Arachnida","Trombidiformes","Anystidae","","","","124","123","123","7","0.81","0","99.1","375.48","0","388.65","31.13","0"
"AGAW3","08S","NIN","BOLD:AAA4123","Arthropoda","Arachnida","Araneae","Araneidae","","Neoscona","Neoscona arabesca","110","110","110","8","0","0","98.48","378.31","0","425.62","30.53","0"
"AGAW3","08S","NIN","BOLD:AAC6891","Arthropoda","Arachnida","Araneae","Salticidae","","Phidippus","Phidippus audax","107","93","93","9","13.08","0","99.82","402.99","0","445.22","32.07","0"
"AGAW3","08S","NIN","BOLD:ACV5610","","","","","","","","79","66","66","10","16.46","0","99.5","441.06","0","441.95","38.09","0"
"AGAW3","08S","NIN","BOLD:AAM7961","Arthropoda","Arachnida","Trombidiformes","Anystidae","","","","62","56","56","11","9.68","0","99.34","438.4","0","437.11","31.26","0"
"AGAW3","08S","NIN","BOLD:AAB7667","Arthropoda","Arachnida","Araneae","Lycosidae","","Pardosa","Pardosa saxatilis","60","60","60","12","0","0","99.36","408.03","0","412.67","31.18","0"
"AGAW3","08S","NIN","BOLD:ACV5609","","","","","","","","41","41","41","13","0","0","98.68","439.73","0","442.1","38.6","0"
"AGAW3","08S","NIN","BOLD:AAZ2181","Arthropoda","Collembola","Symphypleona","Bourletiellidae","","","","40","31","31","14","22.5","0","97.46","439.18","0","440.55","39.42","0"
"AGAW3","08S","NIN","BOLD:ACJ7454","Arthropoda","Collembola","Symphypleona","Bourletiellidae","","","","39","33","33","15","15.38","0","99.37","403.87","0","448.31","38.98","0"
"AGAW3","08S","NIN","BOLD:ACL2932","Arthropoda","Collembola","Symphypleona","Bourletiellidae","","Bourletiella","","23","23","23","16","0","0","99.19","442.91","0","444.78","38.67","0"
"AGAW3","08S","NIN","BOLD:AAA8820","Arthropoda","Collembola","Entomobryomorpha","Entomobryidae","Lepidocyrtinae","Lepidocyrtus","Lepidocyrtus cyaneus","18","18","18","17","0","0","98.87","421.89","0","423.17","43.34","0"
"AGAW3","08S","NIN","BOLD:AAB8452","Arthropoda","Collembola","Entomobryomorpha","Entomobryidae","Lepidocyrtinae","Lepidocyrtus","Lepidocyrtus paradoxus","6","6","6","18","0","0","99.08","455.5","0","450","41.89","0"
"AGAW3","08S","NIN","BOLD:AAO8147","Arthropoda","Arachnida","Trombidiformes","Eupodidae","","Eupodes","","3","3","3","19","0","0","98.63","431.67","0","432.33","28.89","0"
"AGAW3","08S","NIN","BOLD:AAA7242","Arthropoda","Collembola","Entomobryomorpha","Entomobryidae","Seirinae, Entomobryinae","Entomobrya, Lepidosira","Entomobrya multifasciata, Lepidosira sp.","2","2","2","20","0","0","99","449","0","450","39.44","0"
"AGAW3","08S","NIN","BOLD:AAI2833","Arthropoda","Arachnida","Araneae","Salticidae","","Phidippus","Phidippus princeps","1","1","1","21","0","0","97.11","450","0","450","32.67","0"
"AGAW3","08S","NIN","BOLD:AAI4346","Arthropoda","Arachnida","Opiliones","Phalangiidae","","Paroligolophus, Phalangium","Paroligolophus agrestis, Phalangium opilio","1","1","1","21","0","0","99.07","432","0","450","36.67","0"
"AGAW3","08S","NIN","BOLD:AAB3836","Arthropoda","Arachnida","Araneae","Philodromidae","","Philodromus","Philodromus cespitum","1","1","1","21","0","0","97.73","397","0","409","29.58","0"
"AGAW3","08W","BAC","TAX:175768","Proteobacteria","Alphaproteobacteria","Rickettsiales","Rickettsiaceae",NA,"Wolbachia","Wolbachia pipientis","24","23","23","1","4.17","0","98.53","443.67","0","445","39.4","0"
"AGAW3","08W","BAC","TAX:91393","Proteobacteria","Alphaproteobacteria","Rickettsiales","Rickettsiaceae",NA,"Wolbachia","","14","14","14","2","0","0","99.42","389.79","0","411.14","39.02","0"
"AGAW3","08W","BAC","TAX:91391","Proteobacteria","Alphaproteobacteria","Rickettsiales","",NA,"","","10","9","9","3","10","0","99.25","396.8","0","449.1","39.84","0"
"AGAW3","08W","BAC","TAX:375847","Proteobacteria","Alphaproteobacteria","Rickettsiales","Rickettsiaceae",NA,"Wolbachia","Wolbachia sp. Hymenoptera_ASGLE1705-10","3","2","2","4","33.33","0","98.52","449.33","0","450","40.07","0"
"AGAW3","08W","CHO","no match","no match","no match","no match","no match","no match","no match","no match","no match","no match","no match","no match","no match","no match","no match","no match","no match","no match","no match","no match"
"AGAW3","08W","INS","BOLD:AAG1711","Arthropoda","Insecta","Diptera","Muscidae","Phaoniinae","Helina","Helina depuncta","35978","25728","25728","1","28.49","0","99.49","433.95","0","436.2","31.37","0"
"AGAW3","08W","INS","BOLD:AAP8739","Arthropoda","Insecta","Diptera","Platystomatidae","Platystomatinae","Rivellia","Rivellia steyskali","14707","8719","8719","2","40.72","0","99.31","442.18","0","443.85","38","0"
"AGAW3","08W","INS","BOLD:AAA3453","Arthropoda","Insecta","Diptera","Anthomyiidae","Anthomyiinae","Delia","","3612","3200","3200","3","11.41","0","99.33","431.29","0","434.24","34.65","0"
"AGAW3","08W","INS","BOLD:AAD0484","Arthropoda","Insecta","Diptera","Chironomidae","","","","1938","1599","1599","4","17.49","0","98.91","445.05","0","446.44","34.74","0"
"AGAW3","08W","INS","BOLD:AAA4555","Arthropoda","Insecta","Orthoptera","Acrididae","Melanoplinae","Melanoplus","Melanoplus stupefactus, Melanoplus angustipennis, Melanoplus ponderosus, Melanoplus bruneri, Melanoplus frigidus, Melanoplus scapularis, Melanoplus cinereus, Melanoplus madeleineae, Melanoplus sanguinipes, Melanoplus nigrescens group, Melanoplus tunicae, Melanoplus walshi, Melanoplus femurrubrum, Melanoplus dawsoni, Melanoplus sp., Melanoplus fasciatus, Melanoplus bowditchi, Melanoplus stonei, Melanoplus bivittatus, Melanoplus huroni, Melanoplus thomasi, Melanoplus borealis, Melanoplus packardii","1908","1600","1600","5","16.14","0","99.32","446.99","0","447.86","36.04","0"
"AGAW3","08W","INS","BOLD:AAG3920","Arthropoda","Insecta","Diptera","Chironomidae","Tanypodinae","Procladius","Procladius sp. 1ES","1683","1040","1040","6","38.21","0","98.76","446.93","0","447.65","33.76","0"
"AGAW3","08W","INS","BOLD:AAC0635","Arthropoda","Insecta","Hemiptera","Miridae","Mirinae","Stenotus","Stenotus binotatus","1518","1114","1114","7","26.61","0","99.29","442.46","0","443.37","34.6","0"
"AGAW3","08W","INS","BOLD:ABY5318","Arthropoda","Insecta","Diptera","Chloropidae","Oscinellinae","Incertella, Rhopalopterum","Rhopalopterum umbrosum, Rhopalopterum luteiceps, Incertella bispina, Rhopalopterum painteri","1471","1047","1047","8","28.82","0","99.04","444.2","0","445.35","32.2","0"
"AGAW3","08W","INS","BOLD:AAD9195","Arthropoda","Insecta","Lepidoptera","Argyresthiidae","","Argyresthia","Argyresthia affinis, Argyresthia thuiella, Argyresthia austerella","1381","1095","1095","9","20.71","0","99.06","440.79","0","442.34","33.01","0"
"AGAW3","08W","INS","BOLD:AAG5507","Arthropoda","Insecta","Diptera","Chironomidae","Chironominae","Chironomus","Chironomus atrella","1167","1015","1015","10","13.02","0","98.76","430.57","0","434.8","36.27","0"
"AGAW3","08W","INS","BOLD:AAG6588","Arthropoda","Insecta","Coleoptera","Chrysomelidae","Cassidinae","Deloyala","Deloyala guttata","835","762","762","11","8.74","0","98.91","426.67","0","428.41","31.26","0"
"AGAW3","08W","INS","BOLD:AAF9771","Arthropoda","Insecta","Diptera","Hybotidae","Tachydromiinae","Platypalpus","Platypalpus longiseta, Platypalpus trivialis, Platypalpus sp. 5, Platypalpus sp. 17, Platypalpus pallidiventris","679","528","528","12","22.24","0","99.36","440.39","0","441.19","34.96","0"
"AGAW3","08W","INS","BOLD:AAA4781","Arthropoda","Insecta","Hymenoptera","Braconidae","Microgastrinae","Glyptapanteles","Glyptapanteles carlossarmientoi","553","498","498","13","9.95","0","99.21","429.51","0","432.42","24.16","0"
"AGAW3","08W","INS","BOLD:AAG5192","Arthropoda","Insecta","Coleoptera","Curculionidae","Entiminae","Barypeithes","Barypeithes pellucidus","358","315","315","14","12.01","0","98.95","433.04","0","434.85","39.04","0"
"AGAW3","08W","INS","BOLD:AAF4976","Arthropoda","Insecta","Coleoptera","Lampyridae","Lampyrinae","Ellychnia","Ellychnia aff. corrusca KSH611, Ellychnia sp., Ellychnia facula, Ellychnia sp. KSH1376, Ellychnia lacustris, Ellychnia corrusca, Ellychnia hatchi, Ellychnia californica","351","323","323","15","7.98","0","98.71","422.52","0","424.6","36.51","0"
"AGAW3","08W","INS","BOLD:AAH3644","Arthropoda","Insecta","Diptera","Cecidomyiidae","","","","328","274","274","16","16.46","0","99.36","438.96","0","440.28","27.18","0"
"AGAW3","08W","INS","BOLD:AAC0903","Arthropoda","Insecta","Diptera","Chironomidae","Chironominae","Chironomus","Chironomus acidophilus","324","275","275","17","15.12","0","99.45","434.1","0","438.93","36.29","0"
"AGAW3","08W","INS","BOLD:AAH3661","Arthropoda","Insecta","Diptera","Cecidomyiidae","","","","312","281","281","18","9.94","0","99.27","431.73","0","433.54","25.42","0"
"AGAW3","08W","INS","BOLD:AAN6154","Arthropoda","Insecta","Coleoptera","Latridiidae","Corticariinae","Melanophthalma","Melanophthalma inermis","312","239","239","18","23.4","0","98.85","440.38","0","441.55","30.96","0"
"AGAW3","08W","INS","BOLD:AAN5342","Arthropoda","Insecta","Diptera","Chironomidae","Orthocladiinae","Heleniella","","292","288","288","20","1.37","0","99.12","419.39","0","424","35.19","0"
"AGAW3","08W","INS","BOLD:AAM7855","Arthropoda","Insecta","Thysanoptera","Thripidae","Thripinae","Frankliniella","Frankliniella tritici","160","141","141","21","11.88","0","99.34","442.4","0","444.75","31.23","0"
"AGAW3","08W","INS","BOLD:AAM9013","Arthropoda","Insecta","Diptera","Keroplatidae","Macrocerinae","Macrocera","Macrocera pusilla","142","137","137","22","3.52","0","99.29","435.54","0","437.08","30.35","0"
"AGAW3","08W","INS","BOLD:AAN5183","Arthropoda","Insecta","Diptera","Cecidomyiidae","","","","121","99","99","23","18.18","0","98.5","444.53","0","445.33","27.87","0"
"AGAW3","08W","INS","BOLD:AAH1890","Arthropoda","Insecta","Hymenoptera","Ichneumonidae","Cryptinae","Stibeutes","Stibeutes brevicornis, Stibeutes heinemanni","107","81","81","24","24.3","0","99.6","443.58","0","445.57","29.64","0"
"AGAW3","08W","INS","BOLD:AAF6935","Arthropoda","Insecta","Coleoptera","Coccinellidae","Coccinellinae","Propylaea","Propylaea quatuordecimpunctata","102","80","80","25","21.57","0","99.6","444.34","0","445.11","32.07","0"
"AGAW3","08W","INS","BOLD:ABV9319","Arthropoda","Insecta","Diptera","Chironomidae","Tanypodinae","Procladius","","97","93","93","26","4.12","0","99.14","427.31","0","430.15","34.42","0"
"AGAW3","08W","INS","BOLD:AAF9789","Arthropoda","Insecta","Diptera","Hybotidae","","","","93","90","90","27","3.23","0","99.2","423.25","0","424.39","33.1","0"
"AGAW3","08W","INS","BOLD:AAG7891","Arthropoda","Insecta","Hymenoptera","Platygastridae","Scelioninae","Telenomus","Telenomus podisi","71","57","57","28","19.72","0","99.26","433.58","0","442.93","27.33","0"
"AGAW3","08W","INS","BOLD:AAM9254","Arthropoda","Insecta","Diptera","Sciaridae","","Bradysia","Bradysia pallipes","71","69","69","28","2.82","0","98.87","425.54","0","426.31","33.16","0"
"AGAW3","08W","INS","BOLD:ACD1218","Arthropoda","Insecta","Diptera","Sciaridae","","Leptosciarella","Leptosciarella scutellata","68","64","64","30","5.88","0","99.13","426.29","0","427.69","35.14","0"
"AGAW3","08W","INS","BOLD:AAV5688","Arthropoda","Insecta","Diptera","Cecidomyiidae","","","","64","64","64","31","0","0","99.34","432.19","0","433.2","34.3","0"
"AGAW3","08W","INS","BOLD:AAA4183","Arthropoda","Insecta","Hemiptera","Aphididae","Aphidinae","Aphis","Aphis spiraecola","55","39","39","32","29.09","0","99.52","447.4","0","447.84","26.43","0"
"AGAW3","08W","INS","BOLD:AAA5299","Arthropoda","Insecta","Diptera","Chironomidae","Orthocladiinae","Cricotopus","Cricotopus laetus, Cricotopus relucens, Cricotopus trifasciatus, Cricotopus sp., Cricotopus sp. 3ES, Cricotopus sylvestris, Cricotopus glacialis","54","53","53","33","1.85","0","99.29","431.74","0","436.37","34.47","0"
"AGAW3","08W","INS","BOLD:ACE1806","Arthropoda","Insecta","Diptera","Camillidae","","Camilla","Camilla atrimana","50","50","50","34","0","0","98.44","427","0","429.56","35.24","0"
"AGAW3","08W","INS","BOLD:AAG9897","Arthropoda","Insecta","Dermaptera","Forficulidae","Forficulinae","Forficula","Forficula auricularia-A","49","49","49","35","0","0","99.05","381.1","0","388.04","45.54","0"
"AGAW3","08W","INS","BOLD:AAG2868","Arthropoda","Insecta","Hemiptera","Cicadellidae","Typhlocybinae","Empoasca","Empoasca sp., Empoasca fabae","47","39","39","36","17.02","0","99.28","436.72","0","447.79","32.36","0"
"AGAW3","08W","INS","BOLD:ADH8663","Arthropoda","Insecta","Hymenoptera","","","","","44","44","44","37","0","0","99","390.55","0","393","27.06","0"
"AGAW3","08W","INS","BOLD:ACW5117","Arthropoda","Insecta","Diptera","Chironomidae","Orthocladiinae","Smittia","Smittia leucopogon","41","40","40","38","2.44","0","99.22","423.49","0","424.73","31.42","0"
"AGAW3","08W","INS","BOLD:AAO7890","Arthropoda","Insecta","Diptera","Agromyzidae","","","","37","35","35","39","5.41","0","98.8","393.05","0","428.73","34.15","0"
"AGAW3","08W","INS","BOLD:AAD1485","Arthropoda","Insecta","Diptera","Chironomidae","Chironominae","Paratanytarsus","Paratanytarsus grimmii, Paratanytarsus sp.","36","35","35","40","2.78","0","99.37","439.97","0","441.58","33.7","0"
"AGAW3","08W","INS","BOLD:AAH0135","Arthropoda","Insecta","Coleoptera","Phalacridae","Phalacrinae","Acylomus","Acylomus pugetanus, Acylomus ergoti","35","32","32","41","8.57","0","98.85","436.17","0","438.06","31.67","0"
"AGAW3","08W","INS","BOLD:AAN8770","Arthropoda","Insecta","Diptera","Psychodidae","Psychodinae","Psychoda","Psychoda trinodulosa","30","29","29","42","3.33","0","98.81","432.1","0","433.43","33.01","0"
"AGAW3","08W","INS","BOLD:ABU5546","Arthropoda","Insecta","Diptera","Psychodidae","Psychodinae","Psychoda","","28","28","28","43","0","0","98.67","435.93","0","437.14","31.3","0"
"AGAW3","08W","INS","BOLD:AAG8839","Arthropoda","Insecta","Hemiptera","Cicadellidae","Deltocephalinae","Errastunus, Latalus","Latalus ocellaris, Errastunus ocellaris","25","18","18","44","28","0","99.65","449.36","0","449.88","35.96","0"
"AGAW3","08W","INS","BOLD:ACY8145","Arthropoda","Insecta","Hymenoptera","Ichneumonidae","Ctenopelmatinae","","","25","25","25","44","0","0","99.26","414.88","0","423.52","26.89","0"
"AGAW3","08W","INS","BOLD:ABA7871","Arthropoda","Insecta","Diptera","Cecidomyiidae","","","","24","24","24","46","0","0","99.09","439.13","0","440.13","26.77","0"
"AGAW3","08W","INS","BOLD:AAA2528","Arthropoda","Insecta","Trichoptera","Hydropsychidae","Hydropsychinae","Hydropsyche","Hydropsyche sparna","23","23","23","47","0","0","98.21","439.61","0","442.7","29.39","0"
"AGAW3","08W","INS","BOLD:AAN6447","Arthropoda","Insecta","Diptera","Sciaridae","","Corynoptera","Corynoptera perpusilla","23","20","20","47","13.04","0","99.28","445.04","0","446.48","36.24","0"
"AGAW3","08W","INS","BOLD:ACJ0111","Arthropoda","Insecta","Hymenoptera","Platygastridae","Scelioninae","Telenomus","","22","22","22","49","0","0","98.58","420.27","0","429.91","28.79","0"
"AGAW3","08W","INS","BOLD:ABW1511","Arthropoda","Insecta","Diptera","Ceratopogonidae","","","","21","15","15","50","28.57","0","97.12","428.24","0","429.57","36.46","0"
"AGAW3","08W","INS","BOLD:ABV2748","Arthropoda","Insecta","Hymenoptera","Braconidae","Alysiinae","","","21","18","18","50","14.29","0","99.61","432.57","0","433.62","28.4","0"
"AGAW3","08W","INS","BOLD:AAN5225","Arthropoda","Insecta","Diptera","Cecidomyiidae","","","","20","17","17","52","15","0","99.38","448.1","0","449.15","28.37","0"
"AGAW3","08W","INS","BOLD:AAV5573","Arthropoda","Insecta","Diptera","Cecidomyiidae","","","","19","19","19","53","0","0","99.24","449.32","0","450","27.94","0"
"AGAW3","08W","INS","BOLD:ABU5525","Arthropoda","Insecta","Diptera","Chironomidae","Orthocladiinae","Limnophyes","Limnophyes sp. 14ES","18","18","18","54","0","0","99.19","388.72","0","391.33","37.36","0"
"AGAW3","08W","INS","BOLD:AAN5242","Arthropoda","Insecta","Diptera","Cecidomyiidae","","","","17","17","17","55","0","0","98.6","413.88","0","416","28.28","0"
"AGAW3","08W","INS","BOLD:AAP9021","Arthropoda","Insecta","Diptera","Cecidomyiidae","","","","17","17","17","55","0","0","98.93","414.24","0","416.47","29.79","0"
"AGAW3","08W","INS","BOLD:AAH3664","Arthropoda","Insecta","Diptera","Cecidomyiidae","","","CecidInt3 sp. BOLD:AAH3664","16","16","16","57","0","0","99.22","434.06","0","439.5","26.97","0"
"AGAW3","08W","INS","BOLD:AAH0134","Arthropoda","Insecta","Coleoptera","Phalacridae","Phalacrinae","Stilbus","Stilbus apicialis","15","14","14","58","6.67","0","99.38","407.93","0","408.87","31.69","0"
"AGAW3","08W","INS","BOLD:ACC7516","Arthropoda","Insecta","Diptera","Cecidomyiidae","","","","13","13","13","59","0","0","99.2","429.69","0","431.77","26.71","0"
"AGAW3","08W","INS","BOLD:AAG5242","Arthropoda","Insecta","Coleoptera","Curculionidae","Entiminae","Barypeithes","Barypeithes pellucidus","13","12","12","59","7.69","0","97.98","436","0","437.15","39.86","0"
"AGAW3","08W","INS","BOLD:AAL7359","Arthropoda","Insecta","Diptera","Chironomidae","Chironominae","Tanytarsus","","12","12","12","61","0","0","98.96","438","0","439.42","36.16","0"
"AGAW3","08W","INS","BOLD:ACJ6954","Arthropoda","Insecta","Diptera","Cecidomyiidae","","","","11","11","11","62","0","0","99.36","338","0","426.73","26.42","0"
"AGAW3","08W","INS","BOLD:AAG2511","Arthropoda","Insecta","Diptera","Anthomyiidae","Anthomyiinae","Delia","Delia platura","10","10","10","63","0","0","98.76","442.5","0","444.9","33.35","0"
"AGAW3","08W","INS","BOLD:AAF5705","Arthropoda","Insecta","Hemiptera","Miridae","Orthotylinae","Blepharidopterus","Blepharidopterus chlorionis","10","8","8","63","20","0","99.56","360.9","0","450","42.49","0"
"AGAW3","08W","INS","BOLD:AAG9196","Arthropoda","Insecta","Hymenoptera","Ichneumonidae","Cryptinae","","","9","9","9","65","0","0","98.42","448.56","0","449.44","29.86","0"
"AGAW3","08W","INS","BOLD:AAH3920","Arthropoda","Insecta","Diptera","Sciaridae","","Dichopygina, Scatopsciara","","9","9","9","65","0","0","98.42","406","0","426","35.1","0"
"AGAW3","08W","INS","BOLD:AAN5267","Arthropoda","Insecta","Diptera","Cecidomyiidae","","","","9","8","8","65","11.11","0","99.68","444.44","0","444.89","30.48","0"
"AGAW3","08W","INS","BOLD:AAB0373","Arthropoda","Insecta","Neuroptera","Chrysopidae","Chrysopinae","Chrysoperla","Chrysoperla sp. isolate 82, Chrysoperla zastrowi, Chrysoperla plorabunda, Chrysoperla sp. isolate 71, Chrysoperla agilis, Italochrysa italica, Chrysoperla sp. isolate 88, Chrysoperla lucasina, Chrysoperla carnea, Chrysoperla mediterranea, Chrysoperla pallida, Chrysoperla nipponensis, Chrysoperla sillemi","7","7","7","68","0","0","99.52","449.86","0","450","31.52","0"
"AGAW3","08W","INS","BOLD:ABA0822","Arthropoda","Insecta","Diptera","Cecidomyiidae","","","","7","6","6","68","14.29","0","98.95","449.71","0","450","26.92","0"
"AGAW3","08W","INS","BOLD:AAL7819","Arthropoda","Insecta","Diptera","Psychodidae","Psychodinae","Psychoda","Psychoda sp., Psychoda sp. BOLD:AAL7819","6","6","6","70","0","0","99.3","429.17","0","430.33","32.74","0"
"AGAW3","08W","INS","BOLD:ADF6661","Arthropoda","Insecta","Diptera","Cecidomyiidae","","","","6","6","6","70","0","0","99","418.67","0","450","30.26","0"
"AGAW3","08W","INS","BOLD:AAF9187","Arthropoda","Insecta","Coleoptera","Curculionidae","Entiminae","Phyllobius","Phyllobius oblongus","6","6","6","70","0","0","99.59","449.33","0","450","39.04","0"
"AGAW3","08W","INS","BOLD:ACA4421","Arthropoda","Insecta","Coleoptera","Chrysomelidae","Galerucinae","Crepidodera","Crepidodera sculpturata","5","5","5","73","0","0","98.97","448.6","0","449.6","36.65","0"
"AGAW3","08W","INS","BOLD:ADH4889","Arthropoda","Insecta","Diptera","","","","","5","5","5","73","0","0","98.75","437.6","0","438.6","35.19","0"
"AGAW3","08W","INS","BOLD:AAG6745","Arthropoda","Insecta","Diptera","Calliphoridae","Calliphorinae","Pollenia","Pollenia pediculata","5","5","5","73","0","0","98.6","393.2","0","394.8","34.02","0"
"AGAW3","08W","INS","BOLD:AAA3867","Arthropoda","Insecta","Lepidoptera","Noctuidae","Noctuinae","Apamea","Apamea amputatrix","5","5","5","73","0","0","98.96","423.8","0","427.2","33.57","0"
"AGAW3","08W","INS","BOLD:AAA5953","Arthropoda","Insecta","Lepidoptera","Gelechiidae","Gelechiinae","Coleotechnites","Coleotechnites sp., Coleotechnites piceaella, Coleotechnites laricis, Coleotechnites sp. 3, Coleotechnites carbonaria, Coleotechnites granti, Coleotechnites macleodi, Coleotechnites gibsonella, Coleotechnites coniferella, Coleotechnites martini","5","5","5","73","0","0","99.02","449.2","0","450","34.67","0"
"AGAW3","08W","INS","BOLD:AAP7834","Arthropoda","Insecta","Coleoptera","Phalacridae","Phalacrinae","Stilbus","","4","4","4","78","0","0","98.93","418","0","419.25","32.73","0"
"AGAW3","08W","INS","BOLD:AAD2041","Arthropoda","Insecta","Diptera","Dolichopodidae","Sciapodinae","Condylostylus","Condylostylus inermis, Condylostylus nigrofemoratus, Condylostylus connectans, Condylostylus caudatus, Condylostylus flavipes","4","4","4","78","0","0","99.16","448.75","0","450","36.06","0"
"AGAW3","08W","INS","BOLD:AAU8736","Arthropoda","Insecta","Hymenoptera","Diapriidae","Belytinae","Belyta","Belyta validicornis","3","3","3","80","0","0","99.93","448","0","450","24.74","0"
"AGAW3","08W","INS","BOLD:AAM9355","Arthropoda","Insecta","Diptera","Phoridae","Metopininae","Megaselia","Megaselia sp. BOLD:AAM9355","3","3","3","80","0","0","99.11","447.33","0","450","35.04","0"
"AGAW3","08W","INS","BOLD:AAA2815","Arthropoda","Insecta","Lepidoptera","Noctuidae","Noctuinae","Mythimna","Mythimna yukonensis, Mythimna oxygala","3","3","3","80","0","0","99.03","448.33","0","450","33.63","0"
"AGAW3","08W","INS","BOLD:AAA4277","Arthropoda","Insecta","Diptera","Syrphidae","Syrphinae","Toxomerus","Toxomerus marginatus, Toxomerus sp. CR-3, Toxomerus CR-3","3","3","3","80","0","0","98.14","448.67","0","450","31.41","0"
"AGAW3","08W","INS","BOLD:AAH0152","Arthropoda","Insecta","Coleoptera","Cantharidae","Cantharinae","Cantharis","Cantharis rufa","3","2","2","80","33.33","0","98.51","446","0","450","35.41","0"
"AGAW3","08W","INS","BOLD:AAH1888","Arthropoda","Insecta","Hymenoptera","Ichneumonidae","Ctenopelmatinae","","","3","3","3","80","0","0","99.5","424.67","0","428.33","28.91","0"
"AGAW3","08W","INS","BOLD:AAA3181","Arthropoda","Insecta","Diptera","Muscidae","Muscinae","Stomoxys","Stomoxys calcitrans","2","2","2","86","0","0","98.44","447.5","0","450","33.11","0"
"AGAW3","08W","INS","BOLD:AAG1717","Arthropoda","Insecta","Diptera","Muscidae","Phaoniinae","Helina","Helina troene","2","2","2","86","0","0","98.31","442.5","0","443.5","30.67","0"
"AGAW3","08W","INS","BOLD:AAI4303","Arthropoda","Insecta","Diptera","Chironomidae","Chironominae","Chironomus","Chironomus melanescens","2","2","2","86","0","0","98.21","415.5","0","421","34.46","0"
"AGAW3","08W","INS","BOLD:AAU6681","Arthropoda","Insecta","Diptera","Tipulidae","Tipulinae","Nephrotoma","Nephrotoma cornicina","2","2","2","86","0","0","99.77","441","0","442.5","36.5","0"
"AGAW3","08W","INS","BOLD:AAN5255","Arthropoda","Insecta","Diptera","Cecidomyiidae","Cecidomyiinae","Dasineura","","2","2","2","86","0","0","99.1","426","0","450","26.78","0"
"AGAW3","08W","INS","BOLD:AAP7843","Arthropoda","Insecta","Coleoptera","Cantharidae","Malthininae","Malthodes","Malthodes pumilus","2","2","2","86","0","0","99.44","447.5","0","450","38","0"
"AGAW3","08W","INS","BOLD:AAB7869","Arthropoda","Insecta","Lepidoptera","Tortricidae","Tortricinae","Phtheochroa","Phtheochroa baracana, Phtheochroa riscana, Phtheochroa birdana, Phtheochroa fulviplicana, Phtheochroa canariana","2","2","2","86","0","0","99.56","450","0","450","29.56","0"
"AGAW3","08W","INS","BOLD:ACF4901","Arthropoda","Insecta","Diptera","Cecidomyiidae","","","","2","2","2","86","0","0","98.87","444","0","447","28.3","0"
"AGAW3","08W","INS","BOLD:AAA2632","Arthropoda","Insecta","Lepidoptera","Noctuidae","Noctuinae","Noctua","Noctua atlantica, Noctua pronuba","2","2","2","86","0","0","98.32","445.5","0","447.5","31.84","0"
"AGAW3","08W","INS","BOLD:ACD9543","Arthropoda","Insecta","Diptera","Sciaridae","","Bradysia","Bradysia sp. BOLD:ACD9543","2","2","2","86","0","0","99.42","407.5","0","409","34.18","0"
"AGAW3","08W","INS","BOLD:AAG9626","Arthropoda","Insecta","Diptera","Dolichopodidae","","","","2","2","2","86","0","0","97.89","450","0","450","37.22","0"
"AGAW3","08W","INS","BOLD:AAF9306","Arthropoda","Insecta","Diptera","Psychodidae","Psychodinae","Psychoda","Psychoda alternata","2","2","2","86","0","0","97.55","449.5","0","450","31.67","0"
"AGAW3","08W","INS","BOLD:ACE0699","Arthropoda","Insecta","Diptera","Chironomidae","","","","2","2","2","86","0","0","97.45","431","0","450","36.44","0"
"AGAW3","08W","INS","BOLD:ADH9867","Arthropoda","Insecta","Diptera","","","","","2","2","2","86","0","0","99.66","443.5","0","448","37.39","0"
"AGAW3","08W","INS","BOLD:AAA3654","Arthropoda","Insecta","Lepidoptera","Crambidae","Crambinae","Crambus","Crambus bigelovi, Crambus johnsoni, Crambus praefectellus","2","2","2","86","0","0","98.87","436","0","438","31.39","0"
"AGAW3","08W","INS","BOLD:AAP7026","Arthropoda","Insecta","Coleoptera","Latridiidae","Corticariinae","Melanophthalma","Melanophthalma villosa","2","2","2","86","0","0","98.59","389","0","450","31","0"
"AGAW3","08W","INS","BOLD:AAA7374","Arthropoda","Insecta","Diptera","Syrphidae","Syrphinae","Sphaerophoria","Sphaerophoria bankowskae, Sphaerophoria infuscata, Sphaerophoria nr. philanthus, Sphaerophoria batava, Sphaerophoria sp., Sphaerophoria rueppellii, Sphaerophoria interrupta-group, Sphaerophoria sp. MA-1, Sphaerophoria cleoae, Sphaerophoria asymmetrica, Sphaerophoria fatarum, Sphaerophoria bengalensis, Sphaerophoria bifurcata, Sphaerophoria taeniata, Sphaerophoria interrupta, Sphaerophoria contigua, Sphaerophoria kaa, Sphaerophoria laurae, Sphaerophoria chongjini, Sphaerophoria boreoalpina, Sphaerophoria sp. BOLDAAA7374, Sphaerophoria macrogaster, Sphaerophoria sulphuripes, Sphaerophoria brevipilosa, Sphaerophoria philanthus, Sphaerophoria pyrrhina, Sphaerophoria abbreviata, Sphaerophoria scripta, Sphaerophoria potentillae, Sphaerophoria longipilosa, Sphaerophoria virgata","2","2","2","86","0","0","99.1","424","0","424.5","31.16","0"
"AGAW3","08W","INS","BOLD:AAG2460","Arthropoda","Insecta","Diptera","Anthomyiidae","Anthomyiinae","Hydrophoria","Hydrophoria lancifer","2","2","2","86","0","0","97.65","446.5","0","450","34.56","0"
"AGAW3","08W","INS","BOLD:AAZ5592","Arthropoda","Insecta","Diptera","Psychodidae","Psychodinae","","","1","1","1","104","0","0","98.92","371","0","373","31.9","0"
"AGAW3","08W","INS","BOLD:AAH4135","Arthropoda","Insecta","Diptera","Chloropidae","Chloropinae","Thaumatomyia","Thaumatomyia glabra, Thaumatomyia grata, Thaumatomyia sp. 1 TAW","1","1","1","104","0","0","99.33","449","0","450","32.44","0"
"AGAW3","08W","INS","BOLD:ACE4100","Arthropoda","Insecta","Orthoptera","Acrididae","Melanoplinae","Melanoplus","Melanoplus pinicola","1","1","1","104","0","0","97.11","450","0","450","36.44","0"
"AGAW3","08W","INS","BOLD:AAG7276","Arthropoda","Insecta","Diptera","Sphaeroceridae","Limosininae","Leptocera","Leptocera erythrocera","1","1","1","104","0","0","97.36","379","0","382","33.25","0"
"AGAW3","08W","INS","BOLD:ABA6163","Arthropoda","Insecta","Psocodea","Caeciliusidae","Caeciliusinae","Valenzuela","Valenzuela piceus","1","1","1","104","0","0","98.98","393","0","397","36.52","0"
"AGAW3","08W","INS","BOLD:AAO8538","Arthropoda","Insecta","Diptera","Psychodidae","Psychodinae","Psychoda","","1","1","1","104","0","0","99.29","421","0","425","32","0"
"AGAW3","08W","INS","BOLD:AAB4657","Arthropoda","Insecta","Diptera","Chironomidae","Chironominae","Chironomus","Chironomus maturus","1","1","1","104","0","0","99.56","450","0","450","38","0"
"AGAW3","08W","INS","BOLD:ADH5210","Arthropoda","Insecta","Diptera","Cecidomyiidae","","","","1","1","1","104","0","0","98.89","450","0","450","26.22","0"
"AGAW3","08W","INS","BOLD:AAA9811","Arthropoda","Insecta","Lepidoptera","Geometridae","Ennominae","Somatolophia, Nemeris, Eusarca","Somatolophia incana, Eusarca confusaria, Somatolophia haydenata, Nemeris speciosa, Somatolophia pallescens","1","1","1","104","0","0","99.78","448","0","450","31.11","0"
"AGAW3","08W","INS","BOLD:AAL8278","Arthropoda","Insecta","Hymenoptera","Ichneumonidae","","","","1","1","1","104","0","0","97.31","372","0","375","26.13","0"
"AGAW3","08W","INS","BOLD:ABA6849","Arthropoda","Insecta","Diptera","Mycetophilidae","Mycetophilinae","Exechia","","1","1","1","104","0","0","99.77","442","0","445","35.96","0"
"AGAW3","08W","INS","BOLD:AAN5230","Arthropoda","Insecta","Diptera","Cecidomyiidae","","","","1","1","1","104","0","0","99.56","450","0","450","26.22","0"
"AGAW3","08W","INS","BOLD:ACL7849","","","","","","","","1","1","1","104","0","0","99.73","373","0","415","32.29","0"
"AGAW3","08W","INS","BOLD:AAA3889","Arthropoda","Insecta","Lepidoptera","Crambidae","Crambinae","Parapediasia","Parapediasia teterrellus","1","1","1","104","0","0","98.15","433","0","436","32.57","0"
"AGAW3","08W","INS","BOLD:AAG9705","Arthropoda","Insecta","Diptera","Dolichopodidae","","","","1","1","1","104","0","0","99.11","449","0","450","32.22","0"
"AGAW3","08W","INS","BOLD:AAZ5483","Arthropoda","Insecta","Diptera","Muscidae","Phaoniinae","Helina","Helina setiventris","1","1","1","104","0","0","99.11","448","0","450","30.22","0"
"AGAW3","08W","INS","BOLD:ADH4973","Arthropoda","Insecta","Diptera","Tipulidae","Tipulinae","Nephrotoma","","1","1","1","104","0","0","98.89","449","0","450","34.67","0"
"AGAW3","08W","INS","BOLD:ACK7314","Arthropoda","Insecta","Diptera","Chironomidae","Chironominae","Polypedilum","","1","1","1","104","0","0","97.52","443","0","450","35.56","0"
"AGAW3","08W","INS","BOLD:ACE7444","Arthropoda","Insecta","Hemiptera","Miridae","Mirinae, Phylinae","Adelphocoris, Megalocoleus","Adelphocoris lineolatus, Megalocoleus molliculus","1","1","1","104","0","0","100","449","0","450","35.11","0"
"AGAW3","08W","INS","BOLD:ABU5520","Arthropoda","Insecta","Diptera","Sciaridae","","Corynoptera","","1","1","1","104","0","0","99.25","401","0","402","34.83","0"
"AGAW3","08W","INS","BOLD:AAH4171","Arthropoda","Insecta","Diptera","Chloropidae","Oscinellinae","Apallates","Apallates montanus, Apallates particeps","1","1","1","104","0","0","100","450","0","450","32.89","0"
"AGAW3","08W","INS","BOLD:AAH2293","Arthropoda","Insecta","Diptera","Asilidae","Stenopogoninae","Dioctria, Eudioctria","Dioctria hyalipennis, Dioctria baumhaueri, Eudioctria propinqua","1","1","1","104","0","0","97.99","447","0","450","30.67","0"
"AGAW3","08W","INS","BOLD:AAH2753","Arthropoda","Insecta","Coleoptera","Anthicidae","Anthicinae","Anthicus","Anthicus cervinus","1","1","1","104","0","0","98.37","430","0","450","34.22","0"
"AGAW3","08W","INS","BOLD:ACI9753","Arthropoda","Insecta","Diptera","Dolichopodidae","Diaphorinae","Chrysotus","","1","1","1","104","0","0","99.02","410","0","450","32.44","0"
"AGAW3","08W","INS","BOLD:AAA2087","Arthropoda","Insecta","Lepidoptera","Erebidae","Herminiinae","Idia","Idia sp., Idia americalis","1","1","1","104","0","0","99.55","449","0","450","30.44","0"
"AGAW3","08W","INS","BOLD:AAP5087","Arthropoda","Insecta","Diptera","Ceratopogonidae","Dasyheleinae","Dasyhelea","","1","1","1","104","0","0","99.11","448","0","450","34.89","0"
"AGAW3","08W","INS","BOLD:AAG6753","Arthropoda","Insecta","Diptera","Lauxaniidae","Lauxaniinae","Camptoprosopella","","1","1","1","104","0","0","97.1","449","0","450","36.44","0"
"AGAW3","08W","INS","BOLD:AAP5169","Arthropoda","Insecta","Diptera","Chloropidae","Oscinellinae","Elachiptera","Elachiptera nigriceps","1","1","1","104","0","0","99.78","450","0","450","31.78","0"
"AGAW3","08W","INS","BOLD:ABV1356","Arthropoda","Insecta","Diptera","Ceratopogonidae","Dasyheleinae","Dasyhelea","","1","1","1","104","0","0","98","449","0","450","38.22","0"
"AGAW3","08W","INS","BOLD:AAK5607","Arthropoda","Insecta","Diptera","Agromyzidae","Agromyzinae","Ophiomyia","Ophiomyia nasuta","1","1","1","104","0","0","97.09","447","0","450","34.22","0"
"AGAW3","08W","INS","BOLD:AAD0591","Arthropoda","Insecta","Hemiptera","Rhopalidae","Rhopalinae","Harmostes","Harmostes reflexulus","1","1","1","104","0","0","98.22","449","0","450","33.33","0"
"AGAW3","08W","INS","BOLD:AAN5199","Arthropoda","Insecta","Diptera","Cecidomyiidae","","","CecidInt20 sp. BOLD:AAN5199","1","1","1","104","0","0","98.67","450","0","450","26.67","0"
"AGAW3","08W","INS","BOLD:ABY5257","Arthropoda","Insecta","Lepidoptera","Noctuidae","Noctuinae","Apamea","Apamea devastator","1","1","1","104","0","0","99.11","450","0","450","30.67","0"
"AGAW3","08W","INS","BOLD:AAF1049","Arthropoda","Insecta","Diptera","Agromyzidae","Phytomyzinae","Cerodontha","Cerodontha fasciata","1","1","1","104","0","0","99.5","401","0","450","31.78","0"
"AGAW3","08W","INS","BOLD:ABV1237","Arthropoda","Insecta","Diptera","Chironomidae","Chironominae","Chironomus","","1","1","1","104","0","0","98.88","448","0","450","34.44","0"
"AGAW3","08W","INS","BOLD:AAC2637","Arthropoda","Insecta","Lepidoptera","Pterophoridae","Pterophorinae","Geina","Geina sheppardi","1","1","1","104","0","0","98.88","448","0","450","31.56","0"
"AGAW3","08W","INS","BOLD:AAA7661","Arthropoda","Insecta","Diptera","Culicidae","Culicinae","Culiseta, Culex","Culiseta melanura, Culex articularis, Culex restuans, Culex acharistus","1","1","1","104","0","0","99.56","450","0","450","33.11","0"
"AGAW3","08W","INS","BOLD:ACE4701","Arthropoda","Insecta","Lepidoptera","Crambidae","Crambinae","Crambus","Crambus multilinellus","1","1","1","104","0","0","100","448","0","450","31.56","0"
"AGAW3","08W","NAR","no match","no match","no match","no match","no match","no match","no match","no match","no match","no match","no match","no match","no match","no match","no match","no match","no match","no match","no match","no match"
"AGAW3","08W","NIN","BOLD:AAB7914","Arthropoda","Collembola","Symphypleona","Bourletiellidae","","Bourletiella","","2273","1724","1724","1","24.15","0","99.54","444.66","0","446.16","38.13","0"
"AGAW3","08W","NIN","BOLD:AAD1564","Arthropoda","Arachnida","Araneae","Clubionidae","","Clubiona","Clubiona abboti","2000","1833","1833","2","8.35","0","98.15","428.58","0","430.99","31.67","0"
"AGAW3","08W","NIN","BOLD:AAG5658","Arthropoda","Arachnida","Araneae","Mimetidae","","Mimetus","Mimetus epeiroides","1651","1393","1393","3","15.63","0","99.15","433.69","0","435.44","34.63","0"
"AGAW3","08W","NIN","BOLD:ACI9908","Arthropoda","Collembola","Symphypleona","Bourletiellidae","","","","1606","1321","1321","4","17.75","0","99.43","432.92","0","435.26","34.84","0"
"AGAW3","08W","NIN","BOLD:AAB3836","Arthropoda","Arachnida","Araneae","Philodromidae","","Philodromus","Philodromus cespitum","1585","1336","1336","5","15.71","0","99.68","391.28","0","429.32","30.89","0"
"AGAW3","08W","NIN","BOLD:ABW2639","Arthropoda","Arachnida","Trombidiformes","Anystidae","","","","1440","1125","1125","6","21.88","0","99.45","447.52","0","446.12","31.38","0"
"AGAW3","08W","NIN","BOLD:AAH6630","Arthropoda","Collembola","Entomobryomorpha","Entomobryidae","Entomobryinae","Entomobrya","Entomobrya unostrigata","1120","952","952","7","15","0","99.28","439.27","0","440.47","39.02","0"
"AGAW3","08W","NIN","BOLD:ACC0359","Arthropoda","Collembola","Symphypleona","Bourletiellidae","","Bourletiella","","737","568","568","8","22.93","0","99.26","439.99","0","443.38","37.74","0"
"AGAW3","08W","NIN","BOLD:AAN4465","","","","","","","","459","387","387","9","15.69","0","98.49","434.09","0","435.32","46.31","0"
"AGAW3","08W","NIN","BOLD:AAI4346","Arthropoda","Arachnida","Opiliones","Phalangiidae","","Paroligolophus, Phalangium","Paroligolophus agrestis, Phalangium opilio","379","367","367","10","3.17","0","99.08","420.09","0","426.17","35.68","0"
"AGAW3","08W","NIN","BOLD:ACV5610","","","","","","","","366","251","251","11","31.42","0","99.5","446.24","0","447.16","38.18","0"
"AGAW3","08W","NIN","BOLD:ACZ1295","Arthropoda","Arachnida","Trombidiformes","Anystidae","","","","222","221","221","12","0.45","0","99.12","376.29","0","387.77","31.17","0"
"AGAW3","08W","NIN","BOLD:AAA5267","Arthropoda","Collembola","Entomobryomorpha","Isotomidae","","","","220","218","218","13","0.91","0","98.97","420.53","0","423.03","38.15","0"
"AGAW3","08W","NIN","BOLD:AAA7246","Arthropoda","Collembola","Entomobryomorpha","Entomobryidae","Entomobryinae","Entomobrya","Entomobrya nicoleti, Entomobrya multifasciata, Entomobrya intermedia","197","168","168","14","14.72","0","99.12","441.82","0","443.13","43.43","0"
"AGAW3","08W","NIN","BOLD:ACV5609","","","","","","","","173","146","146","15","15.61","0","98.79","443.89","0","445.13","38.68","0"
"AGAW3","08W","NIN","BOLD:ACJ7454","Arthropoda","Collembola","Symphypleona","Bourletiellidae","","","","134","97","97","16","27.61","0","99.58","403.39","0","447.55","38.94","0"
"AGAW3","08W","NIN","BOLD:AAF8190","Arthropoda","Arachnida","Araneae","Thomisidae","","Xysticus","Xysticus luctans","121","116","116","17","4.13","0","98.02","402.88","0","441.5","29.6","0"
"AGAW3","08W","NIN","BOLD:ACK1257","Arthropoda","Arachnida","Trombidiformes","Erythraeidae","","Balaustium","","113","91","91","18","19.47","0","99.61","445.1","0","446.2","32.7","0"
"AGAW3","08W","NIN","BOLD:AAE7853","Arthropoda","Arachnida","Araneae","Theridiidae","","Thymoites","Thymoites unimaculatus","95","91","91","19","4.21","0","98.28","430.27","0","432.38","29.36","0"
"AGAW3","08W","NIN","BOLD:ADG1586","Arthropoda","Collembola","Entomobryomorpha","Entomobryidae","","","","83","77","77","20","7.23","0","99.48","395.42","0","433.05","39.98","0"
"AGAW3","08W","NIN","BOLD:AAA8824","Arthropoda","Collembola","Entomobryomorpha","Entomobryidae","Lepidocyrtinae","Lepidocyrtus","Lepidocyrtus violaceus","63","62","62","21","1.59","0","99.14","423.95","0","425.94","44.27","0"
"AGAW3","08W","NIN","BOLD:AAA6910","Arthropoda","Arachnida","Araneae","Theridiidae","","Enoplognatha","Parasteatoda lunata, Enoplognatha ovata, Enoplognatha nr. latimana, Enoplognatha nr. ovata, Enoplognatha latimana","58","54","54","22","6.9","0","98.62","424","0","437.6","31.55","0"
"AGAW3","08W","NIN","BOLD:AAM7961","Arthropoda","Arachnida","Trombidiformes","Anystidae","","","","28","28","28","23","0","0","99.28","443.39","0","440.39","31.39","0"
"AGAW3","08W","NIN","BOLD:AAZ2181","Arthropoda","Collembola","Symphypleona","Bourletiellidae","","","","22","17","17","24","22.73","0","97.37","445.73","0","446.73","39.6","0"
"AGAW3","08W","NIN","BOLD:ACX4635","Arthropoda","Collembola","Symphypleona","Bourletiellidae","","","","12","9","9","25","25","0","99.41","382.92","0","445","39.02","0"
"AGAW3","08W","NIN","BOLD:AAB8452","Arthropoda","Collembola","Entomobryomorpha","Entomobryidae","Lepidocyrtinae","Lepidocyrtus","Lepidocyrtus paradoxus","7","7","7","26","0","0","99.9","449","0","450","42.25","0"
"AGAW3","08W","NIN","BOLD:AAN6555","Arthropoda","Collembola","Entomobryomorpha","Entomobryidae","Lepidocyrtinae","Lepidocyrtus","Lepidocyrtus cyaneus","1","1","1","27","0","0","100","401","0","401","43.64","0"
"AGAW3","08W","NIN","BOLD:ACF7013","Arthropoda","Collembola","Entomobryomorpha","Entomobryidae","Lepidocyrtinae","Lepidocyrtus","","1","1","1","27","0","0","99.78","450","0","450","43.56","0"
"AGAW3","08W","NIN","BOLD:AAA7242","Arthropoda","Collembola","Entomobryomorpha","Entomobryidae","Seirinae, Entomobryinae","Entomobrya, Lepidosira","Entomobrya multifasciata, Lepidosira sp.","1","1","1","27","0","0","99.75","403","0","406","39.16","0"
"AGAW3","09E","BAC","TAX:175768","Proteobacteria","Alphaproteobacteria","Rickettsiales","Rickettsiaceae",NA,"Wolbachia","Wolbachia pipientis","149","119","119","1","20.13","0","99.47","442.8","0","443.97","39.68","0"
"AGAW3","09E","BAC","TAX:91391","Proteobacteria","Alphaproteobacteria","Rickettsiales","",NA,"","","70","51","51","2","27.14","0","99.56","421.26","0","447.51","39.52","0"
"AGAW3","09E","BAC","TAX:91393","Proteobacteria","Alphaproteobacteria","Rickettsiales","Rickettsiaceae",NA,"Wolbachia","","5","5","5","3","0","0","99.01","382.6","0","420.6","38.87","0"
"AGAW3","09E","BAC","TAX:392874","Proteobacteria","Alphaproteobacteria","Rickettsiales","Rickettsiaceae",NA,"Wolbachia","Wolbachia endosymbiont of Hymenoptera","3","2","2","4","33.33","0","97.11","450","0","450","40.44","0"
"AGAW3","09E","BAC","TAX:375847","Proteobacteria","Alphaproteobacteria","Rickettsiales","Rickettsiaceae",NA,"Wolbachia","Wolbachia sp. Hymenoptera_ASGLE1705-10","2","2","2","5","0","0","97.3","407","0","407.5","39.51","0"
"AGAW3","09E","BAC","TAX:91389","Proteobacteria","","","",NA,"","","1","1","1","6","0","0","99.17","361","0","363","38.84","0"
"AGAW3","09E","BAC","TAX:376099","Proteobacteria","Alphaproteobacteria","Rickettsiales","Rickettsiaceae",NA,"Wolbachia","Wolbachia sp. Hymenoptera","1","1","1","6","0","0","97.16","388","0","391","39.39","0"
"AGAW3","09E","CHO","no match","no match","no match","no match","no match","no match","no match","no match","no match","no match","no match","no match","no match","no match","no match","no match","no match","no match","no match","no match"
"AGAW3","09E","INS","BOLD:AAG1711","Arthropoda","Insecta","Diptera","Muscidae","Phaoniinae","Helina","Helina depuncta","9331","7421","7421","1","20.47","0","99.41","434.6","0","436.87","31.37","0"
"AGAW3","09E","INS","BOLD:AAP8739","Arthropoda","Insecta","Diptera","Platystomatidae","Platystomatinae","Rivellia","Rivellia steyskali","8291","5459","5459","2","34.16","0","99.13","442.89","0","444.56","37.99","0"
"AGAW3","09E","INS","BOLD:AAI4303","Arthropoda","Insecta","Diptera","Chironomidae","Chironominae","Chironomus","Chironomus melanescens","5162","4581","4581","3","11.26","0","98.3","428.57","0","433.86","34.3","0"
"AGAW3","09E","INS","BOLD:ABV1237","Arthropoda","Insecta","Diptera","Chironomidae","Chironominae","Chironomus","","4804","3762","3762","4","21.69","0","99.18","441.96","0","442.69","34.89","0"
"AGAW3","09E","INS","BOLD:AAD2041","Arthropoda","Insecta","Diptera","Dolichopodidae","Sciapodinae","Condylostylus","Condylostylus inermis, Condylostylus nigrofemoratus, Condylostylus connectans, Condylostylus caudatus, Condylostylus flavipes","3181","2248","2248","5","29.33","0","99.5","438.71","0","439.9","35.97","0"
"AGAW3","09E","INS","BOLD:AAC0903","Arthropoda","Insecta","Diptera","Chironomidae","Chironominae","Chironomus","Chironomus acidophilus","2756","2136","2136","6","22.5","0","99.47","438.33","0","441.22","36.29","0"
"AGAW3","09E","INS","BOLD:ABY5318","Arthropoda","Insecta","Diptera","Chloropidae","Oscinellinae","Incertella, Rhopalopterum","Rhopalopterum umbrosum, Rhopalopterum luteiceps, Incertella bispina, Rhopalopterum painteri","2637","2087","2087","7","20.86","0","99.25","441.97","0","443.09","32.15","0"
"AGAW3","09E","INS","BOLD:AAN5342","Arthropoda","Insecta","Diptera","Chironomidae","Orthocladiinae","Heleniella","","2481","2228","2228","8","10.2","0","98.92","423.03","0","429.11","35.14","0"
"AGAW3","09E","INS","BOLD:AAA6759","Arthropoda","Insecta","Lepidoptera","Hesperiidae","Hesperiinae","Thymelicus","Thymelicus lineola","1975","1638","1638","9","17.06","0","99.22","437.9","0","439.75","29.69","0"
"AGAW3","09E","INS","BOLD:AAB0373","Arthropoda","Insecta","Neuroptera","Chrysopidae","Chrysopinae","Chrysoperla","Chrysoperla sp. isolate 82, Chrysoperla zastrowi, Chrysoperla plorabunda, Chrysoperla sp. isolate 71, Chrysoperla agilis, Italochrysa italica, Chrysoperla sp. isolate 88, Chrysoperla lucasina, Chrysoperla carnea, Chrysoperla mediterranea, Chrysoperla pallida, Chrysoperla nipponensis, Chrysoperla sillemi","1697","1317","1317","10","22.39","0","99.41","436.39","0","437.32","31.39","0"
"AGAW3","09E","INS","BOLD:AAA2815","Arthropoda","Insecta","Lepidoptera","Noctuidae","Noctuinae","Mythimna","Mythimna yukonensis, Mythimna oxygala","1353","1279","1279","11","5.47","0","98.97","422.81","0","424.97","33.65","0"
"AGAW3","09E","INS","BOLD:AAA4277","Arthropoda","Insecta","Diptera","Syrphidae","Syrphinae","Toxomerus","Toxomerus marginatus, Toxomerus sp. CR-3, Toxomerus CR-3","1352","969","969","12","28.33","0","99.3","444.4","0","445.34","30.45","0"
"AGAW3","09E","INS","BOLD:AAC9062","Arthropoda","Insecta","Diptera","Culicidae","Culicinae","Ochlerotatus, Aedes","Aedes fitchii, Ochlerotatus fitchii, Aedes grossbecki, Aedes ventrovittis","1332","1137","1137","13","14.64","0","99.3","431.99","0","433.82","33.03","0"
"AGAW3","09E","INS","BOLD:AAB7030","Arthropoda","Insecta","Diptera","Chironomidae","Chironominae","Chironomus","Chironomus sp. BOLDAAP2996, Chironomus quinnitukqut, Chironomus cf. decorus 2g, Chironomus decorus group sp. 2 IP2013, Chironomus cf. decorus, Chironomus decorus gr.","1176","1153","1153","14","1.96","0","98.38","420.62","0","425.36","34.48","0"
"AGAW3","09E","INS","BOLD:AAA3867","Arthropoda","Insecta","Lepidoptera","Noctuidae","Noctuinae","Apamea","Apamea amputatrix","1124","918","918","15","18.33","0","99.24","439.75","0","441.68","33.72","0"
"AGAW3","09E","INS","BOLD:AAB9652","Arthropoda","Insecta","Trichoptera","Leptoceridae","Leptocerinae","Triaenodes","Triaenodes nox","1104","876","876","16","20.65","0","97.71","436.52","0","438.04","31.71","0"
"AGAW3","09E","INS","BOLD:AAL7359","Arthropoda","Insecta","Diptera","Chironomidae","Chironominae","Tanytarsus","","1028","931","931","17","9.44","0","99.24","436.29","0","438.35","36.12","0"
"AGAW3","09E","INS","BOLD:ACO8630","","","","","","","","1019","714","714","18","29.93","0","98.23","444.48","0","445.54","34.1","0"
"AGAW3","09E","INS","BOLD:AAG2868","Arthropoda","Insecta","Hemiptera","Cicadellidae","Typhlocybinae","Empoasca","Empoasca sp., Empoasca fabae","975","659","659","19","32.41","0","99.51","442.19","0","447.39","32.32","0"
"AGAW3","09E","INS","BOLD:AAN6154","Arthropoda","Insecta","Coleoptera","Latridiidae","Corticariinae","Melanophthalma","Melanophthalma inermis","946","807","807","20","14.69","0","98.92","435.57","0","437.46","31.35","0"
"AGAW3","09E","INS","BOLD:AAY6730","Arthropoda","Insecta","Hemiptera","Derbidae","Cedusinae","Cedusa","Cedusa incisa","829","707","707","21","14.72","0","99.13","424.33","0","425.97","35.38","0"
"AGAW3","09E","INS","BOLD:AAG8839","Arthropoda","Insecta","Hemiptera","Cicadellidae","Deltocephalinae","Errastunus, Latalus","Latalus ocellaris, Errastunus ocellaris","822","507","507","22","38.32","0","99.72","447.58","0","448.67","35.98","0"
"AGAW3","09E","INS","BOLD:ACB9399","Arthropoda","Insecta","Diptera","Chironomidae","Chironominae","Parachironomus","","817","719","719","23","12","0","99.03","430.98","0","433.79","36.85","0"
"AGAW3","09E","INS","BOLD:AAA4222","Arthropoda","Insecta","Lepidoptera","Erebidae","Hypeninae","Hypena","Hypena scabra","787","761","761","24","3.3","0","99.16","401.78","0","403.61","32.39","0"
"AGAW3","09E","INS","BOLD:ACL8669","Arthropoda","Insecta","Lepidoptera","Psychidae","Psychinae","Psyche","Psyche casta","775","773","773","25","0.26","0","99.01","402.93","0","407.08","37.16","0"
"AGAW3","09E","INS","BOLD:AAA3453","Arthropoda","Insecta","Diptera","Anthomyiidae","Anthomyiinae","Delia","","693","641","641","26","7.5","0","99.37","431.88","0","434.61","34.66","0"
"AGAW3","09E","INS","BOLD:AAG2898","Arthropoda","Insecta","Hemiptera","Cicadellidae","Deltocephalinae","Paraphlepsius","Paraphlepsius apertus, Paraphlepsius irroratus","663","510","510","27","23.08","0","99.59","442.64","0","443.48","31.9","0"
"AGAW3","09E","INS","BOLD:AAN5160","Arthropoda","Insecta","Diptera","Ceratopogonidae","Forcipomyiinae","Atrichopogon","","656","616","616","28","6.1","0","99.2","428.11","0","431.07","37.27","0"
"AGAW3","09E","INS","BOLD:ACA9275","Arthropoda","Insecta","Diptera","Chironomidae","Chironominae","Chironomus","","632","576","576","29","8.86","0","99.38","430.07","0","432.28","35.4","0"
"AGAW3","09E","INS","BOLD:AAZ5973","Arthropoda","Insecta","Diptera","Sepsidae","Sepsinae","Themira","Themira annulipes","627","516","516","30","17.7","0","99.2","437.44","0","438.79","35.67","0"
"AGAW3","09E","INS","BOLD:AAA2314","Arthropoda","Insecta","Lepidoptera","Crambidae","Crambinae","Pediasia","Pediasia trisecta, Pediasia laciniellus, Pediasia sp.","622","519","519","31","16.56","0","98.49","436.78","0","441.1","33.19","0"
"AGAW3","09E","INS","BOLD:AAG5457","Arthropoda","Insecta","Diptera","Chironomidae","Chironominae","","","573","539","539","32","5.93","0","98.89","433.45","0","435.67","34.47","0"
"AGAW3","09E","INS","BOLD:AAM9255","Arthropoda","Insecta","Diptera","Sciaridae","","Pseudolycoriella","","504","490","490","33","2.78","0","99.1","413.2","0","415.89","30.81","0"
"AGAW3","09E","INS","BOLD:ACE8375","Arthropoda","Insecta","Lepidoptera","Crambidae","Spilomelinae","Desmia","Desmia maculalis","498","454","454","34","8.84","0","99.06","423.21","0","425.31","30.45","0"
"AGAW3","09E","INS","BOLD:AAH3661","Arthropoda","Insecta","Diptera","Cecidomyiidae","","","","478","431","431","35","9.83","0","99.3","431.96","0","433.67","25.41","0"
"AGAW3","09E","INS","BOLD:AAG1741","Arthropoda","Insecta","Diptera","Muscidae","Coenosiinae","Coenosia","","469","379","379","36","19.19","0","98.95","440.86","0","441.88","33.34","0"
"AGAW3","09E","INS","BOLD:AAC9132","Arthropoda","Insecta","Diptera","Culicidae","Culicinae","Culiseta","Culiseta inornata","461","459","459","37","0.43","0","98.51","397.11","0","399.96","32.34","0"
"AGAW3","09E","INS","BOLD:AAF9771","Arthropoda","Insecta","Diptera","Hybotidae","Tachydromiinae","Platypalpus","Platypalpus longiseta, Platypalpus trivialis, Platypalpus sp. 5, Platypalpus sp. 17, Platypalpus pallidiventris","446","340","340","38","23.77","0","99.24","445.62","0","446.27","35.04","0"
"AGAW3","09E","INS","BOLD:AAG5507","Arthropoda","Insecta","Diptera","Chironomidae","Chironominae","Chironomus","Chironomus atrella","443","442","442","39","0.23","0","98.71","414.5","0","419.7","35.86","0"
"AGAW3","09E","INS","BOLD:AAH3914","Arthropoda","Insecta","Diptera","Sciaridae","","Sciara","Sciara hamatilis, Sciara humeralis","433","355","355","40","18.01","0","99.35","443.98","0","445.29","30.39","0"
"AGAW3","09E","INS","BOLD:AAP5169","Arthropoda","Insecta","Diptera","Chloropidae","Oscinellinae","Elachiptera","Elachiptera nigriceps","429","337","337","41","21.45","0","99.24","430.84","0","431.9","31.2","0"
"AGAW3","09E","INS","BOLD:AAG7276","Arthropoda","Insecta","Diptera","Sphaeroceridae","Limosininae","Leptocera","Leptocera erythrocera","400","382","382","42","4.5","0","99.07","420.15","0","423.72","32.68","0"
"AGAW3","09E","INS","BOLD:AAE0844","Arthropoda","Insecta","Hemiptera","Pentatomidae","Pentatominae","Euschistus","Euschistus variolarius","390","333","333","43","14.62","0","98.83","422.57","0","423.76","36.64","0"
"AGAW3","09E","INS","BOLD:AAG4781","Arthropoda","Insecta","Diptera","Agromyzidae","","","","370","293","293","44","20.81","0","98.92","443.24","0","445.61","32.23","0"
"AGAW3","09E","INS","BOLD:AAG9668","Arthropoda","Insecta","Diptera","Dolichopodidae","Diaphorinae","Chrysotus","Chrysotus sp., Chrysotus sp. nr. obliquus","362","324","324","45","10.5","0","98.93","445.34","0","447.91","32.78","0"
"AGAW3","09E","INS","BOLD:ABW5431","Arthropoda","Insecta","Diptera","Agromyzidae","","","","340","286","286","46","15.88","0","99.18","435.27","0","436.37","34.11","0"
"AGAW3","09E","INS","BOLD:AAA7374","Arthropoda","Insecta","Diptera","Syrphidae","Syrphinae","Sphaerophoria","Sphaerophoria bankowskae, Sphaerophoria infuscata, Sphaerophoria nr. philanthus, Sphaerophoria batava, Sphaerophoria sp., Sphaerophoria rueppellii, Sphaerophoria interrupta-group, Sphaerophoria sp. MA-1, Sphaerophoria cleoae, Sphaerophoria asymmetrica, Sphaerophoria fatarum, Sphaerophoria bengalensis, Sphaerophoria bifurcata, Sphaerophoria taeniata, Sphaerophoria interrupta, Sphaerophoria contigua, Sphaerophoria kaa, Sphaerophoria laurae, Sphaerophoria chongjini, Sphaerophoria boreoalpina, Sphaerophoria sp. BOLDAAA7374, Sphaerophoria macrogaster, Sphaerophoria sulphuripes, Sphaerophoria brevipilosa, Sphaerophoria philanthus, Sphaerophoria pyrrhina, Sphaerophoria abbreviata, Sphaerophoria scripta, Sphaerophoria potentillae, Sphaerophoria longipilosa, Sphaerophoria virgata","324","263","263","47","18.83","0","98.83","441.55","0","443.16","31.08","0"
"AGAW3","09E","INS","BOLD:ACF2821","Arthropoda","Insecta","Diptera","Mycetophilidae","Mycetophilinae","Mycetophila","Mycetophila fungorum","308","282","282","48","8.44","0","99.07","430.66","0","433.69","35.16","0"
"AGAW3","09E","INS","BOLD:AAN8770","Arthropoda","Insecta","Diptera","Psychodidae","Psychodinae","Psychoda","Psychoda trinodulosa","305","261","261","49","14.43","0","99.23","439.59","0","440.98","33.66","0"
"AGAW3","09E","INS","BOLD:ACF4734","Arthropoda","Insecta","Diptera","Syrphidae","Syrphinae","Platycheirus","Platycheirus hyperboreus","301","255","255","50","15.28","0","99.13","439.11","0","440.77","31.4","0"
"AGAW3","09E","INS","BOLD:ADG1318","","","","","","","","298","271","271","51","9.06","0","99.4","437.34","0","439.8","25","0"
"AGAW3","09E","INS","BOLD:AAA6337","Arthropoda","Insecta","Lepidoptera","Noctuidae","Plusiinae","Anagrapha","Anagrapha falcifera","289","246","246","52","14.88","0","99.27","440.72","0","443.01","32.24","0"
"AGAW3","09E","INS","BOLD:ACB9147","Arthropoda","Insecta","Psocodea","Psocidae","","","","282","246","246","53","12.77","0","97.96","429.83","0","431.58","36.83","0"
"AGAW3","09E","INS","BOLD:AAA2482","Arthropoda","Insecta","Lepidoptera","Noctuidae","Noctuinae","Mythimna","Mythimna unipuncta","281","274","274","54","2.49","0","99.04","413.59","0","416.9","30.12","0"
"AGAW3","09E","INS","BOLD:AAF9393","Arthropoda","Insecta","Coleoptera","Meloidae","Meloinae","Epicauta","Epicauta fabricii","280","254","254","55","9.29","0","99","430.84","0","432.51","41.15","0"
"AGAW3","09E","INS","BOLD:AAE0707","Arthropoda","Insecta","Diptera","Chironomidae","Chironominae","Microtendipes","Microtendipes pedellus","268","259","259","56","3.36","0","99.09","435.33","0","437.68","30.75","0"
"AGAW3","09E","INS","BOLD:AAP2870","Arthropoda","Insecta","Diptera","Dolichopodidae","","","","267","229","229","57","14.23","0","99.15","439.37","0","440.62","38.36","0"
"AGAW3","09E","INS","BOLD:AAA4555","Arthropoda","Insecta","Orthoptera","Acrididae","Melanoplinae","Melanoplus","Melanoplus stupefactus, Melanoplus angustipennis, Melanoplus ponderosus, Melanoplus bruneri, Melanoplus frigidus, Melanoplus scapularis, Melanoplus cinereus, Melanoplus madeleineae, Melanoplus sanguinipes, Melanoplus nigrescens group, Melanoplus tunicae, Melanoplus walshi, Melanoplus femurrubrum, Melanoplus dawsoni, Melanoplus sp., Melanoplus fasciatus, Melanoplus bowditchi, Melanoplus stonei, Melanoplus bivittatus, Melanoplus huroni, Melanoplus thomasi, Melanoplus borealis, Melanoplus packardii","254","232","232","58","8.66","0","99.33","444.4","0","446.1","36.2","0"
"AGAW3","09E","INS","BOLD:AAB2384","Arthropoda","Insecta","Diptera","Syrphidae","Syrphinae","Eupeodes","Eupeodes duseki, Eupeodes volucris, Eupeodes nielseni, Eupeodes americanus, Eupeodes flukei, Eupeodes vockerothi, Eupeodes lucasi, Eupeodes montivagus, Eupeodes rufipunctatus, Eupeodes rojasi, Eupeodes goeldlini, Eupeodes luniger, Eupeodes punctifer, Eupeodes bucculatus, Eupeodes perplexus, Eupeodes abiskoensis, Eupeodes corollae, Eupeodes pomus, Eupeodes lundbecki, Eupeodes nitens, Eupeodes sp. BOLDAAB2384, Eupeodes curtus, Eupeodes latifasciatus","247","231","231","59","6.48","0","98.17","417.02","0","418.81","31.37","0"
"AGAW3","09E","INS","BOLD:AAA7661","Arthropoda","Insecta","Diptera","Culicidae","Culicinae","Culiseta, Culex","Culiseta melanura, Culex articularis, Culex restuans, Culex acharistus","246","234","234","60","4.88","0","99.26","433.76","0","435.93","32.85","0"
"AGAW3","09E","INS","BOLD:ABV1297","Arthropoda","Insecta","Diptera","Sciaridae","","","","241","238","238","61","1.24","0","98.97","432.12","0","434.6","30.29","0"
"AGAW3","09E","INS","BOLD:AAF9028","Arthropoda","Insecta","Diptera","Limoniidae","","","","229","228","228","62","0.44","0","98.84","401.73","0","405.22","38.21","0"
"AGAW3","09E","INS","BOLD:ADH7798","Arthropoda","Insecta","Hemiptera","Cicadellidae","Deltocephalinae","Colladonus","Colladonus clitellarius","221","196","196","63","11.31","0","98.88","381.96","0","436.13","35.54","0"
"AGAW3","09E","INS","BOLD:AAA5299","Arthropoda","Insecta","Diptera","Chironomidae","Orthocladiinae","Cricotopus","Cricotopus laetus, Cricotopus relucens, Cricotopus trifasciatus, Cricotopus sp., Cricotopus sp. 3ES, Cricotopus sylvestris, Cricotopus glacialis","211","195","195","64","7.58","0","99","436.78","0","438.15","34.86","0"
"AGAW3","09E","INS","BOLD:AAG4905","Arthropoda","Insecta","Diptera","Mycetophilidae","Mycetophilinae","Mycetophila","","199","186","186","65","6.53","0","99.21","421.42","0","423.06","35.23","0"
"AGAW3","09E","INS","BOLD:AAB6702","Arthropoda","Insecta","Coleoptera","Cerambycidae","Cerambycinae","Clytus","Clytus ruricola","190","153","153","66","19.47","0","99.15","422.09","0","438.33","38.58","0"
"AGAW3","09E","INS","BOLD:AAH0237","Arthropoda","Insecta","Coleoptera","Cleridae","Clerinae","Enoclerus","Enoclerus nigripes, Enoclerus rosmarus","189","185","185","67","2.12","0","98.33","420.82","0","423.25","32.14","0"
"AGAW3","09E","INS","BOLD:AAB5081","Arthropoda","Insecta","Hemiptera","Miridae","Mirinae","Leptopterna","Leptopterna dolabrata","186","183","183","68","1.61","0","99.18","399.91","0","401.34","39.38","0"
"AGAW3","09E","INS","BOLD:AAC4403","Arthropoda","Insecta","Diptera","Chironomidae","Chironominae","Micropsectra","Micropsectra polita","185","184","184","69","0.54","0","98.88","418.42","0","420.58","34.54","0"
"AGAW3","09E","INS","BOLD:AAC0635","Arthropoda","Insecta","Hemiptera","Miridae","Mirinae","Stenotus","Stenotus binotatus","185","155","155","69","16.22","0","99.22","444.09","0","444.82","34.73","0"
"AGAW3","09E","INS","BOLD:AAA4781","Arthropoda","Insecta","Hymenoptera","Braconidae","Microgastrinae","Glyptapanteles","Glyptapanteles carlossarmientoi","185","173","173","69","6.49","0","99.21","431.17","0","434.01","24.24","0"
"AGAW3","09E","INS","BOLD:AAG5423","Arthropoda","Insecta","Diptera","Chironomidae","Chironominae","Dicrotendipes","","177","171","171","72","3.39","0","98.51","414.59","0","417.12","37.21","0"
"AGAW3","09E","INS","BOLD:AAC4197","Arthropoda","Insecta","Diptera","Chironomidae","Orthocladiinae","Paraphaenocladius","Paraphaenocladius impensus","172","145","145","73","15.7","0","99.32","433.51","0","435.96","34.86","0"
"AGAW3","09E","INS","BOLD:AAV1055","Arthropoda","Insecta","Diptera","Stratiomyidae","Beridinae","Allognosta","Allognosta fuscitarsis","162","150","150","74","7.41","0","99.06","431.97","0","433.49","35.71","0"
"AGAW3","09E","INS","BOLD:AAG9626","Arthropoda","Insecta","Diptera","Dolichopodidae","","","","161","134","134","75","16.77","0","98.54","441.4","0","443.07","36.13","0"
"AGAW3","09E","INS","BOLD:AAG2511","Arthropoda","Insecta","Diptera","Anthomyiidae","Anthomyiinae","Delia","Delia platura","159","155","155","76","2.52","0","98.98","442.18","0","444.97","33.1","0"
"AGAW3","09E","INS","BOLD:AAN5341","Arthropoda","Insecta","Diptera","Chironomidae","Orthocladiinae","Camptocladius","Camptocladius stercorarius","149","148","148","77","0.67","0","98.74","400.8","0","403.81","35.61","0"
"AGAW3","09E","INS","BOLD:AAH4135","Arthropoda","Insecta","Diptera","Chloropidae","Chloropinae","Thaumatomyia","Thaumatomyia glabra, Thaumatomyia grata, Thaumatomyia sp. 1 TAW","147","136","136","78","7.48","0","98.64","424.26","0","427.38","31.42","0"
"AGAW3","09E","INS","BOLD:AAG6588","Arthropoda","Insecta","Coleoptera","Chrysomelidae","Cassidinae","Deloyala","Deloyala guttata","147","144","144","78","2.04","0","98.99","420.08","0","422.1","31.29","0"
"AGAW3","09E","INS","BOLD:AAB8911","Arthropoda","Insecta","Coleoptera","Curculionidae","Entiminae","Polydrusus","Polydrusus formosus","143","111","111","80","22.38","0","99.4","445.02","0","445.99","34.55","0"
"AGAW3","09E","INS","BOLD:AAD1485","Arthropoda","Insecta","Diptera","Chironomidae","Chironominae","Paratanytarsus","Paratanytarsus grimmii, Paratanytarsus sp.","137","118","118","81","13.87","0","99.43","445.17","0","446.43","33.85","0"
"AGAW3","09E","INS","BOLD:AAG6463","Arthropoda","Insecta","Diptera","Ceratopogonidae","","","","136","129","129","82","5.15","0","99.41","430.22","0","432.49","31.1","0"
"AGAW3","09E","INS","BOLD:AAG8842","Arthropoda","Insecta","Hemiptera","Cicadellidae","Deltocephalinae","Scaphytopius","Scaphytopius acutus, Scaphytopius latus, Scaphytopius oregonensis","135","99","99","83","26.67","0","99.47","443.06","0","444.13","35.62","0"
"AGAW3","09E","INS","BOLD:AAG6432","Arthropoda","Insecta","Diptera","Ceratopogonidae","Forcipomyiinae","Atrichopogon","","133","117","117","84","12.03","0","98.28","436.47","0","437.99","34","0"
"AGAW3","09E","INS","BOLD:ACU6949","Arthropoda","Insecta","Diptera","Dolichopodidae","","","","133","129","129","84","3.01","0","98.56","409.49","0","411.36","34.66","0"
"AGAW3","09E","INS","BOLD:AAH0349","Arthropoda","Insecta","Coleoptera","Chrysomelidae","Cassidinae","Charidotella","Charidotella sexpunctata, Charidotella purpurata","131","124","124","86","5.34","0","99.24","423.83","0","425.75","32.72","0"
"AGAW3","09E","INS","BOLD:AAN8212","Arthropoda","Insecta","Hymenoptera","Braconidae","Euphorinae","Meteorus","","131","125","125","86","4.58","0","98.49","428.02","0","430.2","25.88","0"
"AGAW3","09E","INS","BOLD:AAE3183","Arthropoda","Insecta","Lepidoptera","Noctuidae","Noctuinae","Apamea","Apamea cariosa","130","116","116","88","10.77","0","99.06","438.01","0","439.46","32.73","0"
"AGAW3","09E","INS","BOLD:AAA8279","Arthropoda","Insecta","Lepidoptera","Noctuidae","Noctuinae","Neoligia","Neoligia subjuncta","130","116","116","88","10.77","0","99.29","435.64","0","436.92","30.67","0"
"AGAW3","09E","INS","BOLD:AAG9736","Arthropoda","Insecta","Diptera","Dolichopodidae","Dolichopodinae","Dolichopus","Dolichopus burnesi, Dolichopus albiciliatus","128","127","127","90","0.78","0","98.9","391.3","0","393.91","35.76","0"
"AGAW3","09E","INS","BOLD:AAH0134","Arthropoda","Insecta","Coleoptera","Phalacridae","Phalacrinae","Stilbus","Stilbus apicialis","119","105","105","91","11.76","0","99.33","430.35","0","431.49","31.96","0"
"AGAW3","09E","INS","BOLD:ACK3589","Arthropoda","Insecta","Diptera","Cecidomyiidae","","","","117","95","95","92","18.8","0","98.95","448.63","0","449.41","32.37","0"
"AGAW3","09E","INS","BOLD:AAB1098","Arthropoda","Insecta","Diptera","Culicidae","Culicinae","Ochlerotatus, Aedes","Aedes excrucians, Aedes cantans, Aedes stimulans, Aedes euedes, Aedes riparius, Aedes annulipes, Ochlerotatus cantans, Aedes fitchii, Aedes sp., Ochlerotatus annulipes, Ochlerotatus excrucians","109","104","104","93","4.59","0","98.78","435.45","0","436.77","35.33","0"
"AGAW3","09E","INS","BOLD:AAN6406","Arthropoda","Insecta","Diptera","Sphaeroceridae","Limosininae","Opalimosina","Opalimosina mirabilis","105","101","101","94","3.81","0","99.21","429.72","0","434.66","32.38","0"
"AGAW3","09E","INS","BOLD:ABY1165","Arthropoda","Insecta","Coleoptera","Anthicidae","","","","104","96","96","95","7.69","0","98.42","425.71","0","427.02","35.76","0"
"AGAW3","09E","INS","BOLD:AAL4232","Arthropoda","Insecta","Diptera","Agromyzidae","Phytomyzinae","Phytoliriomyza","Phytoliriomyza arctica","104","92","92","95","11.54","0","99.21","428.02","0","429.77","31.96","0"
"AGAW3","09E","INS","BOLD:AAN5225","Arthropoda","Insecta","Diptera","Cecidomyiidae","","","","102","79","79","97","22.55","0","99.32","445.04","0","446.05","28.33","0"
"AGAW3","09E","INS","BOLD:AAC0706","Arthropoda","Insecta","Diptera","Chironomidae","Chironominae","Dicrotendipes","Dicrotendipes tritomus","87","87","87","98","0","0","97.91","405.62","0","412.31","36.12","0"
"AGAW3","09E","INS","BOLD:AAU6681","Arthropoda","Insecta","Diptera","Tipulidae","Tipulinae","Nephrotoma","Nephrotoma cornicina","83","81","81","99","2.41","0","99.66","442.01","0","443.18","36.28","0"
"AGAW3","09E","INS","BOLD:ACE0554","Arthropoda","Insecta","Diptera","Phoridae","Metopininae","Megaselia","","83","81","81","99","2.41","0","99.24","439.11","0","441.31","32.2","0"
"AGAW3","09E","INS","BOLD:ABV1455","Arthropoda","Insecta","Diptera","Chironomidae","Chironominae","","","71","65","65","101","8.45","0","98.66","426.62","0","428.62","31.9","0"
"AGAW3","09E","INS","BOLD:AAH0256","Arthropoda","Insecta","Coleoptera","Latridiidae","Corticariinae","Corticaria, Corticarina","","70","66","66","102","5.71","0","98.53","427.53","0","434.6","31.9","0"
"AGAW3","09E","INS","BOLD:AAG7279","Arthropoda","Insecta","Diptera","Sphaeroceridae","Limosininae","Spelobia","Spelobia ochripes","64","46","46","103","28.13","0","97.78","441.81","0","445.58","34.4","0"
"AGAW3","09E","INS","BOLD:ADF9260","Arthropoda","Insecta","Coleoptera","Phalacridae","Phalacrinae","Acylomus","","64","57","57","103","10.94","0","98.77","445.11","0","446.27","31.73","0"
"AGAW3","09E","INS","BOLD:AAA4178","Arthropoda","Insecta","Lepidoptera","Geometridae","Larentiinae","Eupithecia","Eupithecia miserulata, Eupithecia chlorofasciata","62","59","59","105","4.84","0","99.5","445.08","0","446.77","31.39","0"
"AGAW3","09E","INS","BOLD:AAH3300","Arthropoda","Insecta","Coleoptera","Coccinellidae","Coccinellinae","Cycloneda","Cycloneda munda","58","57","57","106","1.72","0","98.92","417.38","0","421.05","35.48","0"
"AGAW3","09E","INS","BOLD:AAU6966","Arthropoda","Insecta","Coleoptera","Throscidae","Throscinae","Trixagus","Trixagus chevrolati","56","44","44","107","21.43","0","99.51","398.52","0","425.54","39.99","0"
"AGAW3","09E","INS","BOLD:AAA4271","Arthropoda","Insecta","Lepidoptera","Geometridae","Larentiinae","Costaconvexa","Costaconvexa centrostrigaria","53","50","50","108","5.66","0","99.15","444.6","0","446.58","31.62","0"
"AGAW3","09E","INS","BOLD:ACD3384","Arthropoda","Insecta","Hemiptera","Miridae","Orthotylinae","Orthotylus","Orthotylus robiniae","53","45","45","108","15.09","0","99.71","436.89","0","437.83","41.55","0"
"AGAW3","09E","INS","BOLD:AAA7822","Arthropoda","Insecta","Lepidoptera","Coleophoridae","Coleophorinae","Coleophora","Coleophora pruniella, Coleophora prunifoliae, Coleophora leucochrysella, Coleophora cf. pruniella","51","47","47","110","7.84","0","99.66","444.84","0","446.37","32.31","0"
"AGAW3","09E","INS","BOLD:AAN5161","Arthropoda","Insecta","Diptera","Ceratopogonidae","Dasyheleinae","Dasyhelea","","50","50","50","111","0","0","98.98","398.72","0","400.9","37.37","0"
"AGAW3","09E","INS","BOLD:AAN6408","Arthropoda","Insecta","Diptera","Sphaeroceridae","Limosininae","Spelobia, Bifronsina","Bifronsina bifrons, Spelobia luteilabris, Spelobia bifrons","50","45","45","111","10","0","99.16","437.96","0","440.26","32.59","0"
"AGAW3","09E","INS","BOLD:AAG3275","Arthropoda","Insecta","Diptera","Phoridae","Metopininae","Megaselia","Megaselia minuta, Megaselia nubilipennis","50","48","48","111","4","0","98.49","433.12","0","434.88","35.06","0"
"AGAW3","09E","INS","BOLD:AAG8491","Arthropoda","Insecta","Diptera","Drosophilidae","Drosophilinae","Scaptomyza","Scaptomyza trochanterata, Scaptomyza adusta","50","50","50","111","0","0","98.94","410.64","0","414.74","33.71","0"
"AGAW3","09E","INS","BOLD:AAM6655","Arthropoda","Insecta","Diptera","Hybotidae","Tachydromiinae","Platypalpus","Platypalpus stabilis","48","40","40","115","16.67","0","99.51","447.77","0","448.52","35.15","0"
"AGAW3","09E","INS","BOLD:AAV5688","Arthropoda","Insecta","Diptera","Cecidomyiidae","","","","42","40","40","116","4.76","0","99.41","433.05","0","434.33","34.37","0"
"AGAW3","09E","INS","BOLD:AAQ0271","Arthropoda","Insecta","Diptera","Dolichopodidae","","","","41","37","37","117","9.76","0","98.13","438.9","0","440.51","37.09","0"
"AGAW3","09E","INS","BOLD:AAH4208","Arthropoda","Insecta","Diptera","Chloropidae","Oscinellinae","Elachiptera","Elachiptera erythropleura, Elachiptera costata","40","37","37","118","7.5","0","99.11","434.48","0","437.98","31.33","0"
"AGAW3","09E","INS","BOLD:AAA7067","Arthropoda","Insecta","Diptera","Culicidae","Culicinae","Aedes, Aedimorphus","Aedimorphus vexans, Aedes vexans","39","38","38","119","2.56","0","98.02","424.59","0","426.18","33.04","0"
"AGAW3","09E","INS","BOLD:AAL7819","Arthropoda","Insecta","Diptera","Psychodidae","Psychodinae","Psychoda","Psychoda sp., Psychoda sp. BOLD:AAL7819","39","36","36","119","7.69","0","99.38","424.08","0","426.56","32.71","0"
