## Supplementary figures and images for "CommDivMap: Modelling and mapping species richness at different spatial scales"

### Suppl File 2 - animated GIF linear model

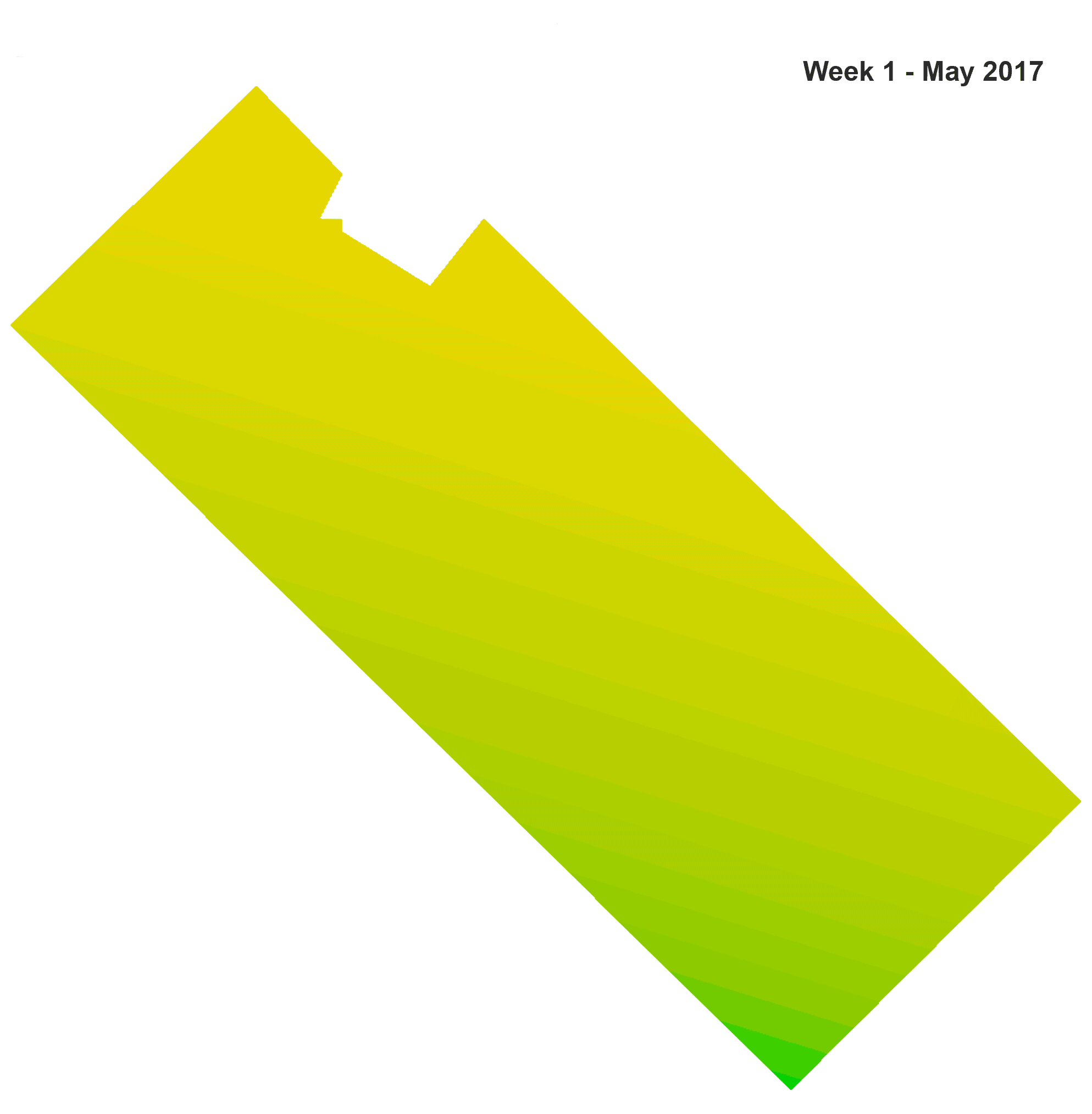

### Suppl File 3 - animated GIF binomial model

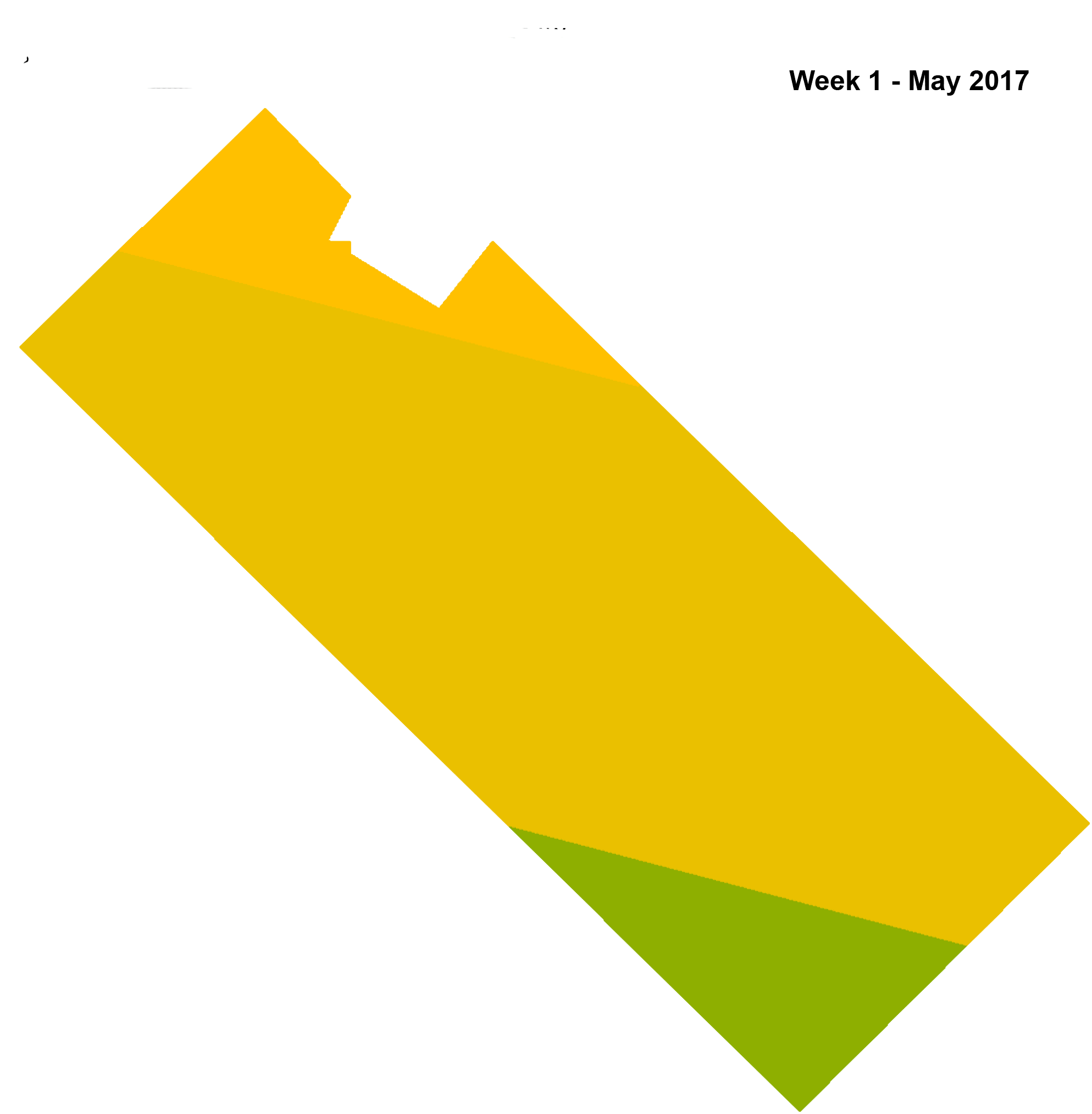
